# Impaired autophagic flux in skeletal muscle of plectin-related epidermolysis bullosa simplex with muscular dystrophy

**DOI:** 10.1101/2025.02.26.640276

**Authors:** Michaela M. Zrelski, Margret Eckhard, Petra Fichtinger, Sabrina Hösele, Andy Sombke, Leonid Mill, Monika Kustermann, Wolfgang M. Schmidt, Fiona Norwood, Ursula Schlötzer-Schrehardt, Gerhard Wiche, Rolf Schröder, Lilli Winter

## Abstract

**Background:** Plectin, a multi-functional cytolinker and intermediate filament (IF) stabilizing protein, is essential for muscle fiber integrity and function. Mutations in the human plectin gene (*PLEC*) cause autosomal recessive epidermolysis bullosa simplex with muscular dystrophy (EBS-MD). The disorganization and aggregation of desmin IFs in conjunction with degenerative changes of the myofibrillar apparatus are key features in the skeletal muscle pathology of EBS-MD. We performed a comprehensive analysis addressing protein homeostasis in this rare protein aggregation disease by using human EBS-MD tissue, plectin knock-out mice, and plectin-deficient cells.

**Methods:** Protein degradation pathways were analyzed in muscles from EBS-MD patients, muscle-specific conditional plectin knockout (MCK-Cre/cKO) mice, as well as immortalized plectin-deficient (*Plec^-/-^*) myoblasts by electron and immunofluorescence microscopy. To obtain a comprehensive picture of autophagic processes, we evaluated the transcriptional regulation and expression levels of autophagic markers in plectin-deficient muscles and myoblasts (RNA-Seq, qRT-PCR, immunoblotting). Autophagic turnover was dynamically assessed by measuring baseline autophagy as well as specific inhibition and activation in mCherry-EGFP-LC3B-expressing *Plec^+/+^*and *Plec^-/-^* myoblasts, and by monitoring primary wild-type (WT) and plectin-deficient (P0) myoblasts using organelle-specific dyes. Analyses of chloroquine (CQ)-treated MCK-Cre/cKO mice corroborated that loss of plectin coincides with impaired autophagic clearance.

**Results:** Our study identified massive accumulation of degradative vacuoles as well as LC3 and SQSTM1-positive patches in EBS-MD patient and MCK-Cre/cKO mouse muscles and *Plec^-/-^* myoblasts. While the transcriptional regulation of autophagy-related proteins remained largely unaltered, protein levels of downstream targets of the autophagosomal degradation machinery were elevated in MCK-Cre/cKO muscle lysates (e.g. LAMP2, BAG3, and SQSTM1 to ∼160, ∼150, and ∼140% of control samples, respectively; *P*<0.05). Autophagosome turnover was compromised in mCherry-EGFP-LC3B-expressing *Plec^-/-^*myoblasts compared to *Plec^+/+^* cells (∼40% reduction in median red:green ratio, reduced puncta number, smaller puncta; *P*<0.01). By labelling autophagic compartments with CYTO-ID dye or lysosomes with LYSO-ID, we found reduced signal intensities in P0 cells (*P*<0.001). Treatment of primary myoblasts with CQ led to drastic swelling of autophagic vacuoles in WT myoblasts, while the swelling in P0 cells was moderate, establishing a defect in their autophagic clearance. Finally, CQ-treatment of MCK-Cre/cKO mice reassured in vivo the concept that autophagic flux is impaired in plectin-deficient muscles.

**Conclusions:** Our work demonstrates that the characteristic protein aggregation pathology in EBS-MD is linked to an impaired autophagic flux. The obtained results open a new perspective on the understanding of the protein aggregation pathology in plectin-related disorders and provides a basis for further pharmacological intervention.

## Introduction

Skeletal muscles are elaborately assembled machines possessing a precisely organized myofibrillar apparatus, designated for contraction and force generation. The principal components of the extrasarcomeric cytoskeleton are desmin intermediate filaments (IFs) forming a three-dimensional scaffold around the myofibrillar Z-disk and throughout the whole myofiber [1]. Plectin, an extraordinarily large (>500 kDa) multi-modular cytolinker protein, can be considered a central connector of the various cytoskeletal filament systems. It harbors a functional actin-binding domain, binding sites for microtubule-associated proteins, and most importantly a binding domain for all types of IF proteins [2]. Distinct plectin isoforms in skeletal muscle bestow a central role in the interconnection, but also in the targeting and anchorage of desmin IFs to sites of strategic importance, i.e. the Z-disks (plectin 1d), costameres (plectin 1f), mitochondria (plectin 1b), and the nuclear/ER membrane system (plectin 1) [3–6]. Thus, plectin orchestrates the structural and functional organization of filamentous cytoskeletal networks, IFs in particular, and thereby substantially contributes to the fundamental biomechanical properties of stress-bearing tissues such as muscle.

As a consequence, the ablation or expression of mutant plectin in human and mouse leads to a faulty organization of the desmin IF system in striated muscle, thereby inflicting a reduced mechanical stress tolerance and a progressive myopathic process. Accordingly, pathogenic sequence alterations of the human plectin gene (*PLEC*), cause a multitude of clinical entities subsumed under the term “plectinopathies”. Epidermolysis bullosa simplex with muscular dystrophy (EBS-MD, MIM #226670), a rare autosomal-recessive disease with congenital skin blistering and late-onset progressive muscle weakness, is the most prevalent plectinopathy [7, 8]. Due to its universal expression, diseases caused by *PLEC* mutations frequently display multisystemic manifestations including more and more additional symptoms [7]. The morphological hallmark of the skeletal muscle pathology in most plectinopathies is the presence of desmin-positive protein aggregates, degenerative changes of the myofibrillar apparatus, and mitochondrial abnormalities [5, 7, 9–11].

While loss of IF network function in conjunction with increased mechanical vulnerability and protein aggregation unambiguously promotes the progressive muscle damage in EBS-MD, imbalanced protein homeostasis is highly anticipated to play a role in the pathogenesis of the disease. To this end, we examined protein quality control mechanisms, such as autophagy, in skeletal muscle specimens from EBS-MD patients, muscle-specific conditional plectin knockout mice (MCK-Cre/cKO), and plectin-deficient myoblasts.

## Materials and methods

### Human skeletal muscle biopsy material

Tissue samples of previously reported EBS-MD patients [9, 11, 12] were obtained from the Institute of Neuropathology, University Hospital Erlangen, and the Department of Neurology, King’s College Hospital London and used for immunofluorescence and electron microscopy as described in the Supplementary Methods. Human plectin reference sequence: NM_000445.5, *PLEC* mutations are listed in the Supplementary Methods. The study was approved by the local ethics committees of participating institutions and conducted according to the principles of the Declaration of Helsinki (2013). Written informed consent was obtained from all participants.

### Animals

Wild-type mice homozygous for the floxed plectin allele (*Plec^fl/fl^*) [13] or MCK-Cre/cKO [5], both in a C57BL/6 background, were housed under specific and pathogen-free conditions in a standard environment with free access to water and food and handled in accordance with the Austrian Federal Government laws and regulations. Animals were sacrificed through cervical dislocation and muscles were subsequently processed for analyses as described in the Supplementary Methods. To determine in vivo the autophagic flux, 20-week-old wild-type and MCK-Cre/cKO animals were injected intraperitoneally with 0.9% saline (control; Sigma-Aldrich, S8776) or with CQ (10 mg/kg in 0.9% saline; Sigma-Adrich, C6628) 4 h ante mortem [14]. All experiments involving animals were performed according to Austrian Federal Government laws and regulations and were approved by the Austrian Federal Ministry of Education, Science and Research (BMBWF-66.009/0226-V/3b/2019 and 2022-0.660.178).

### Cell culture

Immortalized skeletal myoblasts were derived from plectin-expressing (*Plec^+/+^*) or plectin-deficient (*Plec^-/-^*) littermates, both crossed into a p53-deficient (*p53^-/-^*) background, as described [10]. To generate myoblast cell lines stably expressing pBabe puro mCherry-EGFP-LC3B (from J. Debnath [Addgene, 22418] [15]), retroviral supernatants were generated by transfecting phoenix eco cells as previously described [16], with the modification that viral particles were released into F-10-based growth medium. Primary myoblasts were isolated from de-skinned front and hind limbs of neonatal WT or P0 mice as described [10]. Primary human dermal fibroblasts from EBS-MD patients [17, 18] were obtained from the Department of Dermatology, Medical Center University of Freiburg and cultivated as described [19]. *PLEC* mutations are listed in the Supplementary Methods. The study was approved by the ethics committee of the University of Freiburg (ethics number 293-14) and conducted according to the principles of the Declaration of Helsinki (2013). Written informed consent was obtained from all participants. Myoblast and fibroblast cell cultures were used for analyses as described in the Supplementary Methods

## Results

### Increased autophagic build-up in skeletal muscles from EBS-MD patients

Ultrastructural evaluation of a muscle biopsy from a EBS-MD patient [11] showed numerous autophagic vacuoles in the sarcoplasma of myofibers, either filled with cargo remnants appearing as electron-dense deposits or membrane fragments, and frequently located in close proximity to damaged mitochondria (Figure 1A). In addition, subsarcolemmal vacuoles containing myelinated bodies were present. Immunostaining of cryopreserved muscle sections from this as well as from two additional EBS-MD patients [9, 11, 12] showed a marked disruption of the desmin IF networks in conjunction with desmin-positive protein aggregates. Microscopic analysis of LC3, marking autophagosomal structures, revealed a diffuse sarcoplasmic labeling in a subset of EBS-MD muscle fibers as well as a perinuclear labeling, which was also observed in control samples (Figure 1B). In addition, focal deposits of LC3 were frequently noted in EBS-MD muscles, while no LC3 accumulation was seen in control samples and diffuse sarcoplasmic staining was consistently absent. Staining intensities of SQSTM1, a protein that connects polyubiquitinated proteins with LC3, were elevated in all EBS-MD patient muscles, with some fibers displaying massive SQSTM1-labeled subsarcolemmal as well as focal dense sarcoplasmic staining patterns (Figure 1C). While a significant proportion of the LC3-positive patches or SQSTM1-positive deposits clearly co-localized with desmin-positive aggregates (Figure 1B,C, magnifications I-VI, arrows), the majority appeared independent of the collapsed IF network (Figure 1B,C, magnifications I-VI, arrowheads). Taken together, increased LC3 and SQSTM1 immunoreactivity and ultrastructural evidence of abundant autophagic vacuoles indicated augmented autophagy in skeletal muscles of EBS-MD patients.

**Figure 1.**
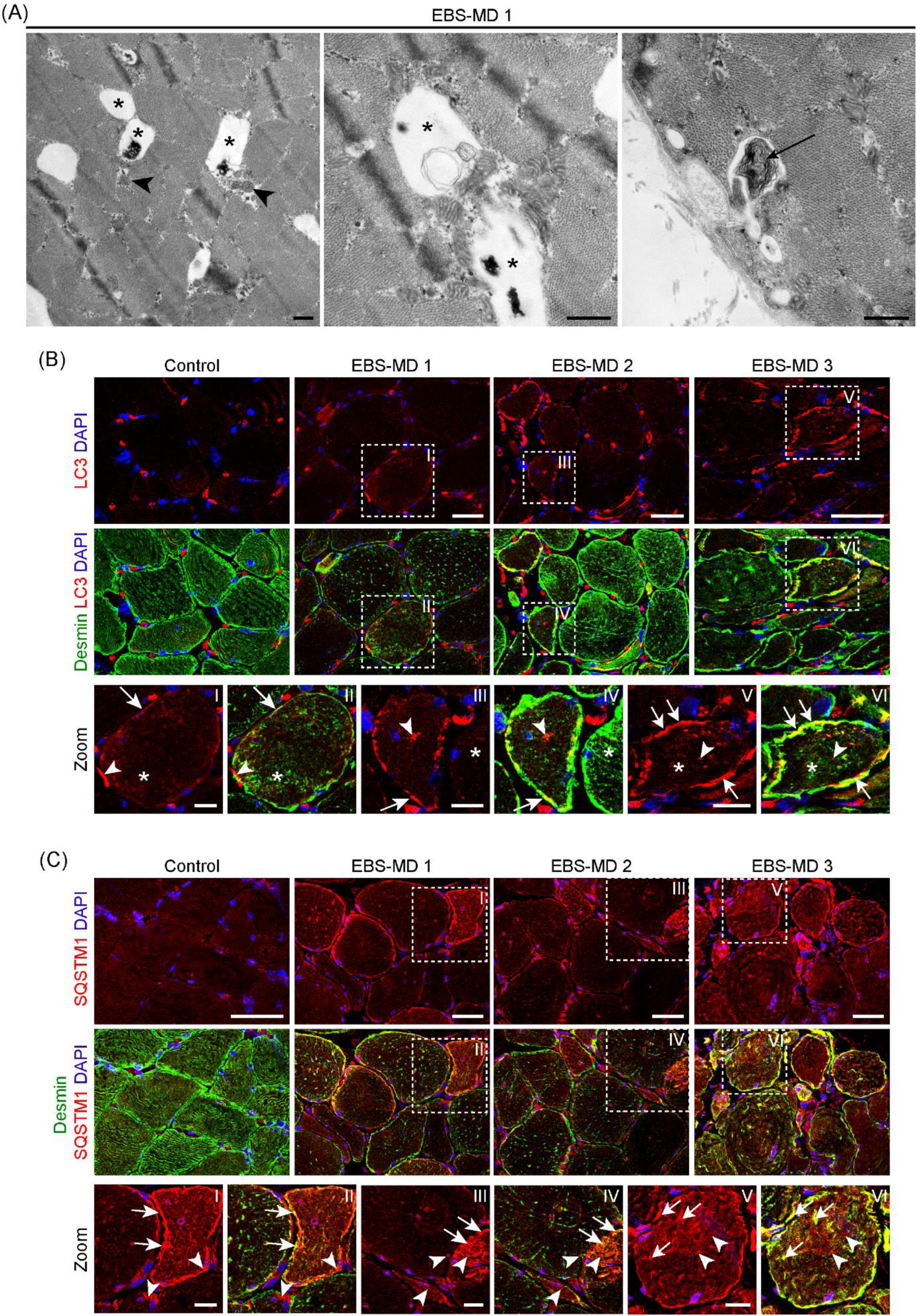
Identification of degradative vacuoles and accumulation of autophagy marker proteins in EBS-MD patient muscle. (A) Representative electron micrographs of muscle sections derived from EBS-MD patient 1 (EBS-MD 1) displaying various degradative vacuoles (asterisks), partially filled with cytoplasmic material and/or membrane remnants and often associated with damaged mitochondria (arrowheads), as well as the occurrence of myelinated bodies (arrow). Scale bars: 500 nm. (B) Double immunostaining of frozen muscle sections derived from a healthy control (Control) and EBS-MD patients (EBS-MD 1, EBS-MD 2, EBS-MD 3) using antibodies to LC3 (red) and desmin (green). Nuclei were visualized using DAPI. Boxed areas indicate magnifications of individual muscle fibers (as denoted by the boxes I-VI). Note the accumulation of LC3-positive patches within EBS-MD patient myofibers, either co-localizing with desmin aggregates (arrows) or without desmin association (arrowheads). Also, note the occurrence of desmin-positive protein aggregates not associating with LC3 protein signals (asterisks). Scale bars: 50 µm; magnifications I-VI 20 µm. (C) Double immunostaining of frozen muscle sections derived from a healthy control and EBS-MD patients using antibodies to SQSTM1 (red) and desmin (green). Nuclei were visualized using DAPI. Boxed areas indicate magnifications of individual muscle fibers (as denoted by the boxes I-VI). Note the occurrence of individual EBS-MD myofibers displaying massive SQSTM1-positive areas, while others harbor small SQSTM1-positive aggregates, either co-localizing with desmin aggregates (arrows) or without desmin association (arrowheads). Scale bars: 50 µm; magnifications I-VI 20 µm.

### Plectin-deficient mouse muscles mirror the autophagy-related pathology in human EBS-MD muscles

In cross and longitudinal muscle sections from 13-week-old MCK-Cre/cKO mice, an established model for the human EBS-MD muscle pathology [5], a focal accumulation of LC3-positive patches was noted (Figure 2A). While some LC3-stained deposits co-localized with desmin-labeled aggregates, the majority appeared independent of the collapsed desmin IF network. Perinuclear LC3 stains were detected in wild-type and diseased muscle samples. Likewise, microscopic analyses of SQSTM1 revealed plectin-deficient muscle fibers with markedly accumulated protein signals, only partially co-localizing with desmin aggregates (Figure 2B). The LC3 and SQSTM1 patterns in plectin-deficient muscles thus strongly mirror the results obtained in EBS-MD muscle. We then quantified the SQSTM1 signal intensities in whole muscle sections with MIRA Vision (https://www.mira.vision/), an artificial intelligence (AI)-based system, and analyzed the frequency distribution of SQSTM1-positive immunosignals from individual myofibers by binning and plotting the relative signal intensities as histograms (Figure 2C). Compared to wild-type, the MCK-Cre/cKO histogram displayed an expanded distribution and a clear shift towards higher binned intensities, highlighting a higher percentage of intensely-labeled myofibers. In addition, transmission electron microscopy analyses of soleus muscles from aged MCK-Cre/cKO mice showed an increased autophagic build-up (Figure 2D). Here, autophagic vacuoles were partially filled with glycogen and/or membrane remnants and their cargo was engulfed by a double-membrane. Again, morphologically altered mitochondria, sometimes containing inclusions, were frequently observed in close vicinity to autophagic vacuoles.

**Figure 2.**
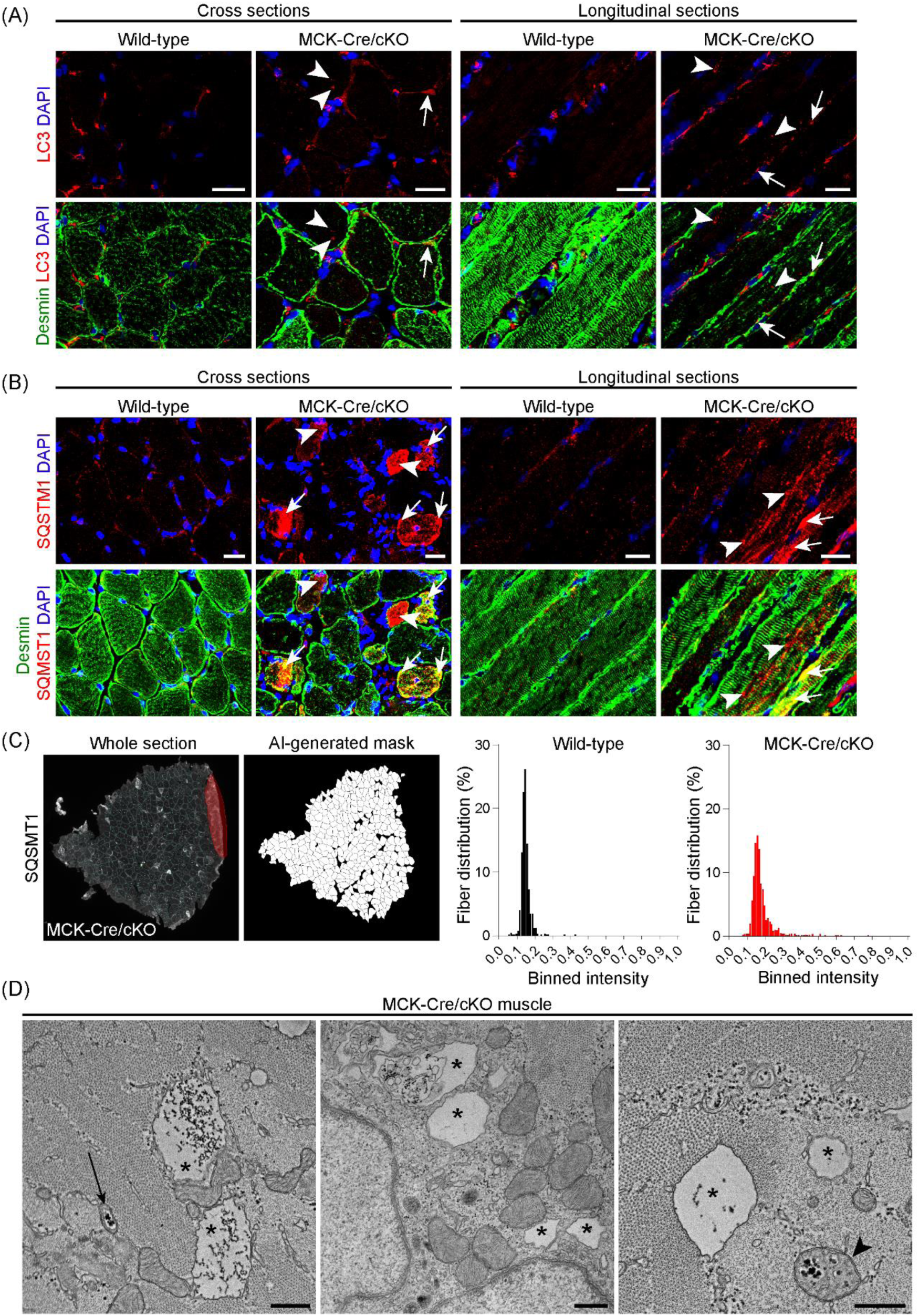
Accumulation of autophagy marker proteins and ultrastructural visualization of degradative vacuoles in plectin-deficient mouse muscle. (A) Double immunostaining of cross and longitudinal soleus muscle sections from wild-type and plectin-deficient (MCK-Cre/cKO) mice using antibodies to LC3 (red) and desmin (green). Nuclei were visualized with DAPI. Note the accumulation of LC3-positive patches in plectin-deficient muscles, either co-localizing with desmin aggregates (arrows) or without desmin association (arrowheads). Scale bars: 20 µm. (B) Double immunostaining of cross and longitudinal soleus muscle sections from wild-type and MCK-Cre/cKO animals using antibodies to SQSTM1 (red) and desmin (green). Nuclei were visualized with DAPI. Note the occurrence of individual plectin-deficient myofibers displaying massive SQSTM1-positive areas, while others harbor small SQSTM1-positive aggregates, either co-localizing with desmin aggregates (arrows) or without desmin association (arrowheads). Scale bars: 20 µm. (C) Artificial intelligence (AI)-based evaluation of SQSTM1 signal intensities within individual myofibers using MIRA Vision. Whole soleus muscle sections were scanned, myofibers automatically identified in an AI-generated mask, and signal intensities obtained and binned for each genotype (bin size = 0.1). Histograms represent the frequency distribution of binned intensities obtained from two animals each (wild-type, n = 673 fibers; MCK-Cre/cKO, n = 1128 fibers). Note the expanded distribution of the histogram and a shift towards the right (i.e. higher intensities) for plectin-deficient muscles. (D) Representative electron micrographs of soleus muscle cross sections obtained from 40-week-old MCK-Cre/cKO mice. Note the occurrence of various degradative vacuoles, including vacuoles with their cargo engulfed in a double-membrane (arrow), and pathologically enlarged vacuoles that are partially filled with glycogen and/or membrane remnants (asterisks). In addition, pathologically altered mitochondria with inclusions can be observed (arrowhead). Scale bars: 500 nm.

### No major alterations in the transcriptional regulation of autophagy in MCK-Cre/cKO muscles

To explore if autophagy-related genes (ATGs) or pathways were altered in plectinopathies, we employed RNA-sequencing (RNA-seq)-based transcriptomic analyses of wild-type and MCK-Cre/cKO muscles. Differential expression analysis using *edgeR* revealed 2169 genes up- and 2215 genes downregulated in MCK-Cre/cKO samples (Figure 3A). Upon functional annotation by using the Kyoto Encyclopedia of Genes and Genomes (KEGG) “mmu04140 Autophagy – animal” pathway analysis, only one third (n = 47) out of 142 genes were differentially expressed. Transcription factor EB (TFEB), a key molecule regulating the autophagy-lysosomal pathway at the transcriptional level [20], accumulated in subsarcolemmal regions of MCK-Cre/cKO muscles; in addition small TFEB-positive deposits within myofibers were found, both partially co-localizing with desmin-positive aggregates (Figure 3B). However, the number of TFEB-positive nuclei was in the same range in both wild-type and MCK-Cre/cKO muscles. Even though the phosphorylated, non-active confirmation of TFEB (pTFEB) appeared increased to ∼115% in plectin-deficient muscles, the protein levels of unphosphorylated, active TFEB were similar in both genotypes (Figure S1A). Together, these data imply that the skeletal muscle pathology in MCK-Cre/cKO muscles is not associated with marked transcriptional changes in autophagy-relevant genes.

**Figure 3.**
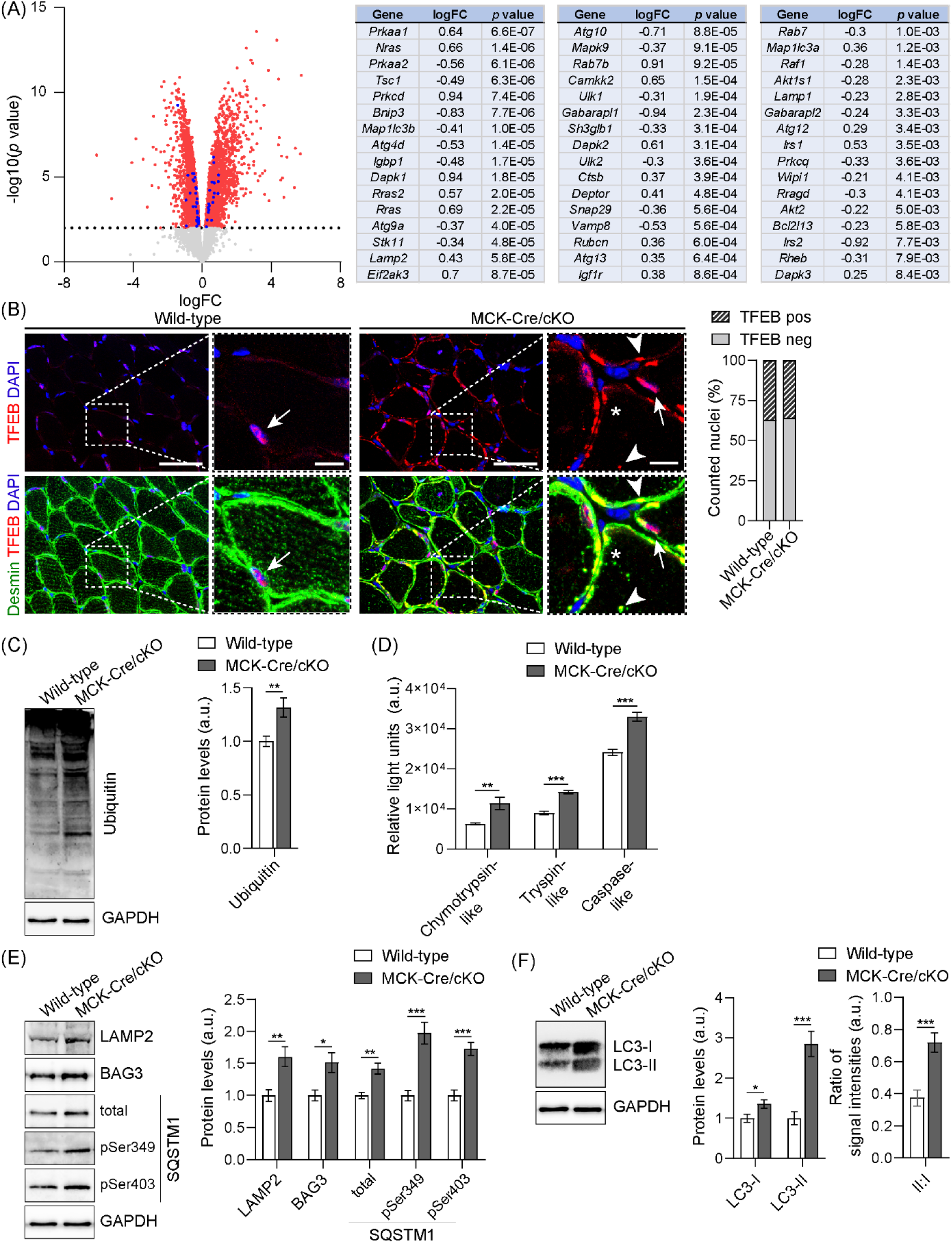
Increased protein levels of autophagic substrates in MCK-Cre/cKO muscles. (A) RNA-Seq analysis of mouse soleus muscle. Volcano plot illustrates differentially expressed genes in MCK-Cre/cKO compared to wild-type samples: significantly up- and downregulated genes are highlighted in red; the dotted line represents the cut-off with *P =* 0.01. Genes from the Kyoto Encyclopedia of Genes and Genomes (KEGG) pathway “mmu04140 Autophagy - animal” are highlighted in blue and listed on the right. logFC, log fold change; n = 5 animals per genotype. (B) Double immunostaining of soleus muscle sections from wild-type and MCK-Cre/cKO mice using antibodies to TFEB (red) and desmin (green). Nuclei were visualized with DAPI. Panels on the right are magnifications of boxed areas indicated in the panels on the left. Note that while a nuclear localization of TFEB was preserved in both genotypes (arrows), concentrated TFEB-positive subsarcolemmal and intermyofibrillar areas occurred only in MCK-Cre/cKO fibers. Also note that some desmin-positive aggregates co-localized with TFEB-positive signals (arrowheads), while others did not. Scale bars: 50 µm, magnifications 10 µm. Quantification of nuclei with or without TFEB localization (wild-type, n=1082 nuclei; MCK-Cre/cKO, n=1466 nuclei; two animals each). (C) Immunoblotting of wild-type and MCK-Cre/cKO muscle lysates using antibodies to ubiquitin and GAPDH. Signal intensities of immunoblots were densitometrically measured and normalized to the total protein content as analyzed by Coomassie staining (not shown). Mean ± SEM; n = 8. (D) Chymotrypsin-trypsin-, and caspase-like proteasomal activities were measured in wild-type and MCK-Cre/cKO muscle lysates derived from 13-week-old mice. Mean ± SEM; samples were measured as triplicates, n = 3 animals each. (E) Immunoblotting of wild-type and MCK-Cre/cKO muscle lysates using antibodies to LAMP2, BAG3, total and two phosphorylated forms of SQSTM1, and GAPDH. Signal intensities of protein bands were densitometrically measured and normalized to the total protein content as analyzed by Coomassie staining (not shown). Mean ± SEM; n = 7-8. (F) Immunoblotting of wild-type and MCK-Cre/cKO muscle lysates using antibodies to LC3 and GAPDH. Signal intensities of upper (non-lipidated, LC3-I) and lower (lipidated, LC3-II) protein bands were densitometrically measured and normalized to the total protein content (as analyzed by Coomassie staining, not shown). From these values, the LC3-II to LC3-I ratios were calculated. Mean ± SEM; n = 10. For (C-F): \**P* < 0.05, \*\**P* < 0.01, \*\*\**P* < 0.001 (two-tailed, unpaired *t*-test with Welch’s correction); ns, not significant.

### Accumulation of autophagy marker proteins and increased proteasomal function in MCK-Cre/cKO muscles

When the expression levels of PQC-related proteins required for the induction of autophagy or the build-up of the phagophore, such as mTOR, ULK1, ATG7, Beclin-1, ATG5 and ATG3 [21] were quantitatively assessed, all evaluated proteins revealed no obvious differences between the two genotypes, indicating unaltered autophagy initiation (Figure S1B). In contrast, the amounts of ubiquitinated proteins were ∼1.3-fold increased in MCK-Cre/cKO compared to wild-type muscle lysates (Figure 3C). In addition, while mRNA levels of proteasomal components were largely unaltered (Figure S1C), chymotrypsin-, trypsin-, and caspase-like proteasomal activities were significantly elevated to ∼180%, ∼155%, and ∼140% in MCK-Cre/cKO compared to wild-type muscles, respectively (Figure 3D). When normalized to the total levels of proteasomes (increased to ∼130% in MCK-Cre/cKO; Figure S1D), the proteasomal activities were basically unchanged in MCK-Cre/cKO compared to wild-type muscles (Figure S1E), indicating that plectin deficiency leads to increased levels of proteasomes, with normal proteasomal activity per proteasome. Notably, several downstream targets of the autophagosomal degradation machinery, such as lysosome-associated membrane protein 2 (LAMP2), BAG3, and SQSTM1, were significantly elevated in MCK-Cre/cKO muscle lysates to ∼160, ∼150, and ∼140%, respectively (Figure 3E). Especially the protein levels of SQSTM1 marked for degradation, denoted by phosphorylation at serine (Ser) 349 or Ser403, were drastically increased in plectin-deficient muscles to ∼200% and ∼ 170% of wild-type levels (Figure 3E). Quantitative evaluation of lipidated, membrane-bound LC3-II (Figure 3F, lower band) versus cytoplasmic LC3-I form (upper band) [22], revealed that both versions were significantly elevated, to ∼140% (LC3-I) and ∼290% (LC3-II) in MCK-Cre/cKO muscle lysates, respectively. The ratio of LC3-II to LC3-I appeared more than twice as high in plectin-deficient compared to wild-type muscles, indicating massively increased levels of membrane-associated LC3 in MCK-Cre/cKO samples.

Because of the progressive nature of plectinopathy-associated muscle degeneration, we assessed whether the observed autophagy-associated pathological alterations amplify in an age-dependent manner. In skeletal muscle lysates from 40-week-old MCK-Cre/cKO mice mTOR was decreased to ∼70%, while ULK1 and Beclin-1 were increased to ∼150% and ∼200% of wild-type levels, respectively, while protein levels of ATG3, ATG5, and ATG7 remained comparable between both genotypes (Figure S2A). Levels of ubiquitinated proteins (Figure S2B) as well as chymotrypsin- and trypsin-like proteasomal activities were significantly elevated in conjunction with increased protein levels of proteasomal subunits, while caspase-like proteasomal activities were reduced in aged MCK-Cre/cKO muscles compared to wild-type samples (Figure S2C,D). Increased amounts of LAMP2, BAG3, SQSTM1, LC3-I, and LC3-II in aged MCK-Cre/cKO muscle lysates (Figure S2E,F) appeared reminiscent of the changes observed in 13-week-old animals. Contrary to young mice, the ratio of LC3-II to LC3-I was lower in MCK-Cre/cKO muscles from aged animals compared to their wild-type counterparts. These analyses suggested that altered autophagic degradation, manifesting as prominent accumulation of autophagy effector proteins while leaving the upstream machineries largely intact, already manifested at a younger age.

### Impaired autophagosome turnover in plectin deficient myoblasts: evaluation of basal conditions, inhibition, and activation of autophagy

Accumulation of autophagic compartments and downstream target proteins, as observed in plectinopathic muscles, could either result from increased autophagy or impaired later degradation [22]. However, as autophagy is an utterly active process, unravelling autophagic flux in plectin-deficiency requires more dynamic examination approaches and the need of corresponding cell culture models. Immortalized *Plec^-/-^* myoblasts, when differentiated into multinucleated myotubes, reproduced critical pathological alterations of plectinopathic muscle fibers, including Z-disk aberrations and accumulation of desmin-positive protein aggregates [10]. On a single cell level, *Plec^-/-^* myoblasts recapitulated the autophagy-associated alterations observed in plectinopathy patients and mice, including enrichment of SQSTM1-positive bodies in the cytoplasm (Figure S3A) and numerous autophagic vacuoles, as observed in electron microscopy (Figure S3B). While mRNA expression levels of *Ulk1, Becn1*, *Lamp2*, *Bag3*, *Sqstm1*, *Map1lc3a* and *Map1lc3b* were not altered (Figure S3C), the protein levels of Beclin-1 were significantly increased (to ∼180%) compared to *Plec^+/+^* cells (Figure S3D). No differences were observed for mTOR or ULK1 protein levels (Figure S3D) or the total amount of polyubiquitinated proteins (Figure S3E). However, while the chymotrypsin-like proteasomal activity was slightly reduced (Figure S4A,B), the downstream targets of the autophagosomal degradation machinery, e.g. BAG3 and SQSTM1, and especially the Ser349 and Ser403 phosphorylated SQSTM1 versions, were substantially enriched in *Plec^-/-^* myoblasts (Figure S4C). While the overall LC3 protein levels were reduced, the LC3-II to LC3-I ratio displayed a significant increase (to ∼140%), indicating that a higher proportion of LC3 was membrane-associated in *Plec^-/-^* cells (Figure S4D). Altogether, our experiments confirmed that plectin deficiency evokes striking alterations in the autophagic pathway in human, mice, and myoblasts, and demonstrated the applicability of the cell system for further analyses.

Next, we generated *Plec^+/+^* and *Plec^-/-^* myoblast cell lines stably expressing a mCherry-EGFP-LC3B reporter [15] by retroviral transduction. In these cells, autophagic flux can be determined due to the quenching of EGFP in the acidic lysosome, whereas the more stable mCherry remains preserved, thereby enabling to follow the transitions of autophagic compartments from autophagosomes (yellow) to autolysosomes (red); accordingly, the red:green signal ratio is considered as a measure of autophagic flux [23]. At basal conditions, mCherry-EGFP-LC3B-expressing *Plec^+/+^* myoblasts displayed distinct puncta formation in the reddish spectrum, while plectin-deficient cells harbored primarily green puncta as well as diffuse cytoplasmic EGFP-LC3B signals. *Plec^-/-^* myoblasts displayed a significantly reduced mean red:green ratio of ∼0.4 (*Plec^+/+^*: ∼0.7), had only ∼69 puncta per cell (*Plec^+/+^*: ∼86), and the puncta were smaller, i.e. a reduced median puncta volume of ∼0.39 µm^3^ in *Plec^-/-^* myoblasts (diameter [d] = 0.90 µm; *Plec^+/+^*: ∼0.54 µm^3^, corresponding to d*a = 1.00 µm, assumed as a perfect sphere; Figure 4A). Together, these data pointed towards diminished formation and volume of autophagic compartments in mCherry-EGFP-LC3B-expressing *Plec^-/-^* myoblasts at basal conditions.

**Figure 4.**
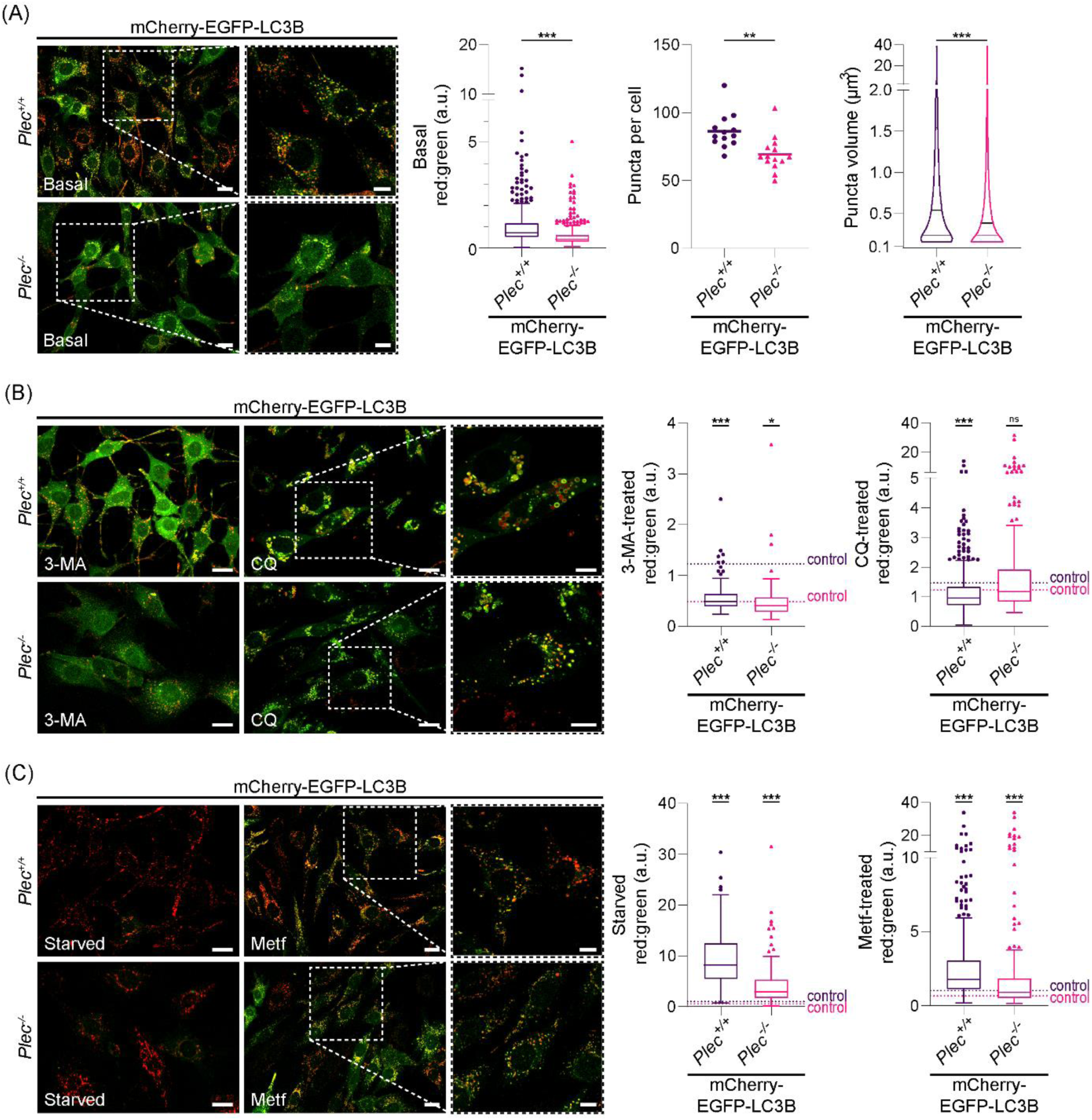
Impaired autophagic flux in mCherry-EGFP-LC3B-expressing plectin-deficient myoblast cell lines. (A) Immortalized (*p53^-/-^*) plectin-expressing (*Plec^+/+^*) and plectin-deficient (*Plec^-/-^*) myoblasts stably expressing mCherry-EGFP-LC3B at basal conditions. Panels on the right are single plane magnifications of the boxed areas indicated in the confocal images in the panels on the left. Note increased cytoplasmic LC3B signals *Plec^-/-^* myoblasts. Scale bars: 20 µm, magnifications 10 µm. Red:green signal ratios of mCherry-EGFP-LC3B-expressing *Plec^+/+^* and *Plec^-/-^*myoblasts at basal conditions: box plots show the median and Tukey whiskers (*Plec^+/+^*, n = 468 cells; *Plec^-/-^*, n= 455 cells); \*\*\**P* < 0.001 (two-tailed Mann Whitney test). Number of mCherry-positive puncta per cell was determined by using the 3D Objects Counter; each dot represents a single field-of-view, the line represents the mean (*Plec^+/+^*, n = 13 field-of-views; *Plec^-/-^*, n = 14 field-of-views); \*\**P* < 0.01 (two-tailed, unpaired t-test with Welch’s correction). Volume of mCherry-positive puncta was determined by using the 3D Objects Counter; violin plots show median, the 25^th^ and 75^th^ percentile (*Plec^+/+^*, n = 38926 puncta [468 cells]; *Plec^-/-^*, n = 29302 puncta [455 cells]); \*\*\**P* < 0.001 (two-tailed Mann Whitney test). (B) mCherry-EGFP-LC3B-expressing *Plec^+/+^* and *Plec^-/-^*myoblasts were treated with 9 mM 3-methyladenine (3-MA) or 50 µM chloroquine (CQ) for 3 h. Panels on the right are single plane magnifications of the boxed areas indicated in the confocal images in the center panels. Note the increased cytoplasmic LC3B signals upon 3-MA treatment, and the massive swelling of vesicles upon CQ-treatment in *Plec^+/+^*myoblasts, while *Plec^-/-^* myoblasts remained largely unaltered. Also note the occurrence of partially acidified autolysosomes in CQ-treated *Plec^+/+^* myoblasts, as denoted by the formation of green ring-like structures. Scale bars: 20 µm, magnifications 10 µm. Red:green signal ratios of 3-MA- and CQ-treated mCherry-EGFP-LC3B-expressing *Plec^+/+^* and *Plec^-/-^* myoblasts: dotted lines represent the median values of the respective cells at control conditions, box plots show the median and Tukey whiskers (*Plec^+/+^*, n = 103/113 [control/3-MA] and 202/260 [control/CQ] cells; *Plec^-/-^*, n = 89/81 [control/3-MA] and 142/206 [control/CQ] cells); \**P* < 0.05, \*\*\**P* < 0.001 (two-way ANOVA of the ranked dataset with Tukey’s post-hoc correction for multiple comparisons); ns, not significant. (C) mCherry-EGFP-LC3B-expressing *Plec^+/+^* and *Plec^-/-^* myoblasts were starved for 24 h or treated with 100 mM metformin (Metf) for 48 h. Panels on the right are single plane magnifications of the boxed areas indicated in the confocal images in the center panels. Note the increased red signals in both cell lines compared to control conditions. Scale bars: 20 µm, magnifications 10 µm. Red:green signal ratios of starved or Metf-treated mCherry-EGFP-LC3B-expressing *Plec^+/+^* and *Plec^-/-^* myoblasts: dotted lines represent the median values of the respective cells at control conditions, box plots show the median and Tukey whiskers (*Plec^+/+^*, n = 185/177 [control/starved] and 500/460 [control/Metf] cells; *Plec^-/-^*, n = 176/177 [control/ starved] and 630/285 [control/ Metf] cells). \**P* < 0.05, \*\*\**P* < 0.001 (two-way ANOVA of the ranked dataset with Tukey’s post-hoc correction for multiple comparisons); ns, not significant.

To explore the effects of impaired autophagic flux, cells were next treated with commonly used inhibitors of autophagy (Figure 4B). Application of 3-methyladenine (3-MA), an inhibitor of autophagic sequestration, for 3 h, led to enhanced cytoplasmic signals in mCherry-EGFP-LC3B-expressing *Plec^+/+^* myoblasts and a significant drop in the signal ratios to ∼40% of untreated controls. In contrast, the subcellular distribution of mCherry-EGFP-LC3B in 3-MA-treated *Plec^-/-^* myoblasts remained largely unaltered and the signal ratios were slightly reduced to ∼80% of untreated controls. Inhibition of late-stage autophagy by application of CQ for 3 h caused swollen and ballooned autophagic vacuoles in *Plec^+/+^* cells; defective degradation was emphasized by the occurrence of partially acidified vesicles (Figure 4B, magnifications), accompanied by significantly reduced red:green ratios (*Plec^+/+^*: ∼1.5 at control conditions vs. ∼1.0 after CQ-treatment). Notably, CQ-treated *Plec^-/-^* cells appeared similar to untreated *Plec^-/-^* controls and the red:green ratios remained unaffected (*Plec^-/-^*: ∼1.2 at control conditions vs. ∼1.2 after CQ-treatment). Treatment with bafilomycin A1 (Baf A1), inhibiting the autophagosome/lysosome fusion, had similar effects as the CQ-treatment (Figure S5A). Long-term 24 h application of CQ or Baf A1 led to almost exclusive bright yellow signals as well as comparable red:green signal ratios in mCherry-EGFP-LC3B-expressing *Plec^+/+^* and *Plec^-/-^*cells (Figure S5B), indicating an indistinguishable accumulation of autophagic vesicles after elongated inhibition. 24 h treatment with 3-MA, on the other hand, caused a return to baseline levels (Figure S5C).

To actively trigger autophagy, cells were either starved or treated with metformin (Metf) (Figure 4C). While 3 h starvation slightly activated autophagy (Figure S5D), 24 h starvation led to bright red signals in both genotypes and a ∼8 and ∼6-fold increase in red:green ratios for *Plec^+/+^* and *Plec^-/-^* cells, respectively. 48 h Metf-treatment, though less invasive, also triggered autophagic turnover, as illustrated by distinct puncta formation (Figure 4C, magnifications). Notably, the median red:green ratio for Metf-treated *Plec^-/-^* myoblasts (∼0.9) was comparable to that for untreated *Plec^+/+^* cells (∼1.0; *P* = 0.34). Collectively, our experiments indicated significantly compromised autophagic flux in mCherry-EGFP-LC3B-expressing *Plec^-/-^* myoblasts.

In addition, we used the cationic amphiphilic tracer dye CYTO-ID, comprehensively labeling autophagic compartments [24], and quantified autophagosomal staining in *Plec^+/+^* and *Plec^-/-^* myoblasts cell lines using flow cytometry (Figure 5A). As expected, the average median fluorescence intensity (MFI) of *Plec^-/-^*myoblasts was significantly reduced to ∼3.3×10^5^ (*Plec^+/+^*: ∼4.8×10^5^). Upon CQ-treatment, the MFI of *Plec^+/+^* myoblasts increased to ∼150%, while the MFI of CQ-treated *Plec^-/-^* cells only increased to ∼137% compared to the respective controls, indicating a lower response. To address the question whether the downstream clearance was also affected in plectin-deficient cells, acidic compartments such as endosomes, lysosomes, and autolysosomes were investigated by flow cytometry of LYSO-ID-stained myoblasts (Figure 5B). Here, basal conditions revealed a trend towards a ∼20% reduced labeling in *Plec^-/-^* myoblasts. In CQ-treated *Plec^+/+^* myoblasts the MFI increased to ∼940%, while *Plec^-/-^* cells reached only a MFI-increase of ∼570% upon CQ-treatment, pointing towards explicit differences between the two genotypes. Finally, lysosomal degradation capacities were assessed by measuring cathepsin B activity in *Plec^+/+^* and *Plec^-/-^*myoblasts (Figure 5C). Notably, the Magic Red signal intensities were significantly decreased in *Plec^-/-^* myoblasts, proving reduced levels of cathepsin B-mediated proteolysis, i.e. diminished functional lysosomes in plectin-deficient cells.

**Figure 5.**
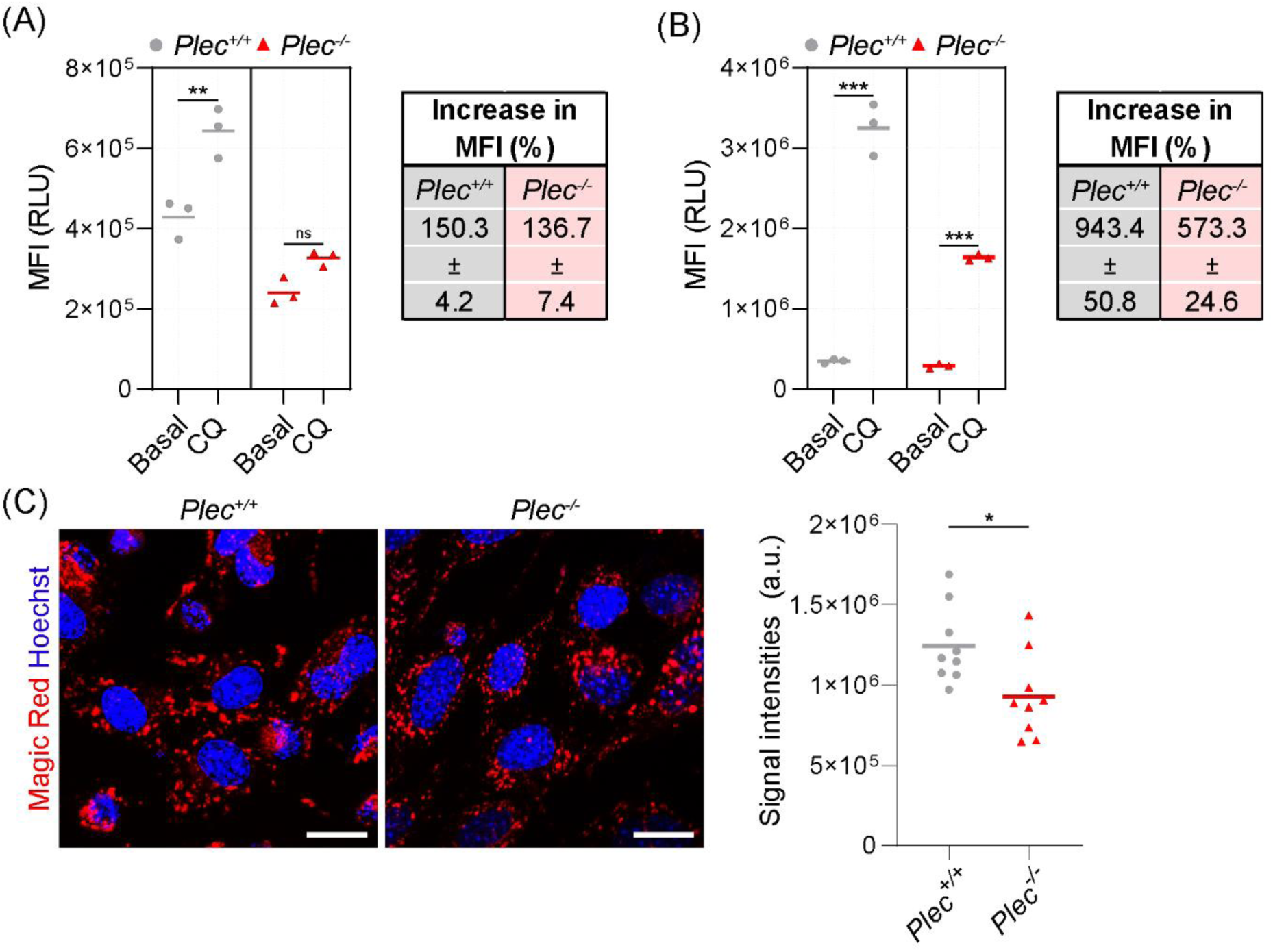
Impaired autophagic flux, reduced lysosomal capacities, and reduced intracellular cathepsin B protease activity in *Plec^-/-^* myoblasts. (A) Flow cytometry of CYTO-ID-stained *Plec^+/+^* and *Plec^-/-^* myoblasts at control conditions or treated with 50 µM CQ for 3 h and calculation of the increase in median fluorescence intensity (MFI) upon CQ-treatment (CQ:control). Line shows the mean, n = 3 experiments (7.5×10^4^ - 9.5×10^4^ cells/experiment); RLU, relative light units. (B) Flow cytometry of LYSO-ID-stained *Plec^+/+^*and *Plec^-/-^* myoblasts at control conditions or treated with 50 µM CQ for 3 h calculation of the increase MFI upon CQ-treatment (CQ:control). Line shows the mean; n = 3 experiments (4.1×10^4^ - 9.4×10^4^ cells/experiment). (C) *Plec^+/+^* and *Plec^-/-^* myoblasts were stained with the Magic Red Cathepsin B Assay Kit. Nuclei were visualized with Hoechst. Scale bars: 20 µm. Magic Red signal intensities were calculated by normalizing the RawIntDens of a field-of-view to the number of cells. Each dot represents a single field-of-view, the line represents the mean (*Plec^+/+^*, n = 9 [723 cells]; *Plec^-/-^*, n = 9 [748 cells] field-of-views). For (A-C): \**P* < 0.05, \*\**P* < 0.01, \*\*\**P* < 0.001 (two-tailed, unpaired *t*-test with Welch’s correction); ns, not significant.

### Impaired autophagic flux and reduced lysosomal capacities in primary plectin-deficient myoblasts and fibroblasts

As the immortalized cell lines were derived from *p53^-/-^* mice, our plectinopathy cell models represent double knockout systems, i.e. they lack p53 in addition to plectin [10]. Thus, phenotypes observed on the molecular or cellular level, attributed to the lack of plectin, could also (at least in part) be due to p53 deficiency. To establish the specificity of the phenotypic traits, key observations were confirmed in primary myoblast cultures derived from normal (not *p53^-/-^*) neonatal wild-type (WT) and plectin-deficient (P0) mice [25]. When primary P0 myoblasts were analyzed using the CYTO-ID dye by life-cell imaging (Figure 6A), their autophagic vesicles appeared smaller than in WT cells and the median CYTO-ID signal intensities were significantly decreased to ∼1.6×10^3^ (WT: ∼5.0×10^3^). Notably, while both genotypes harbored a similar number of puncta per cell (WT: ∼34, P0: ∼32), the median puncta volume was significantly reduced to ∼0.42 µm^3^ (d = 0.93 µm) in P0 myoblasts (WT: ∼0.54 µm^3^, d = 1.00 µm), suggesting that smaller puncta were the cause of the reduced CYTO-ID signal intensities. CQ-treatment led to a drastic swelling of autophagic vacuoles in WT myoblasts, while the swelling in P0 cells was moderate (Figure 6B). This was also reflected by the ∼1.3-fold higher CYTO-ID signal intensities in WT vs. P0 myoblasts. LYSO-ID staining of P0 myoblasts revealed a significant drop in the basal fluorescence intensities (∼60% of WT levels), and a reduction to only ∼19 puncta per P0 myoblast (WT: ∼30 puncta). Because the median puncta volume remained similar in both genotypes, reduced signal intensities correlated to a reduced number of acidic compartments. Consistently, application of CQ resulted in less vesicle swelling in P0 myoblasts, coinciding with a reduced median signal intensity (P0: ∼1.8×10^3^, WT: ∼5.0×10^3^; Figure 6D). Together, our dynamic monitoring of immortalized and primary plectin-deficient myoblasts unequivocally established a defect in their autophagic flux and lysosomal degradation.

**Figure 6.**
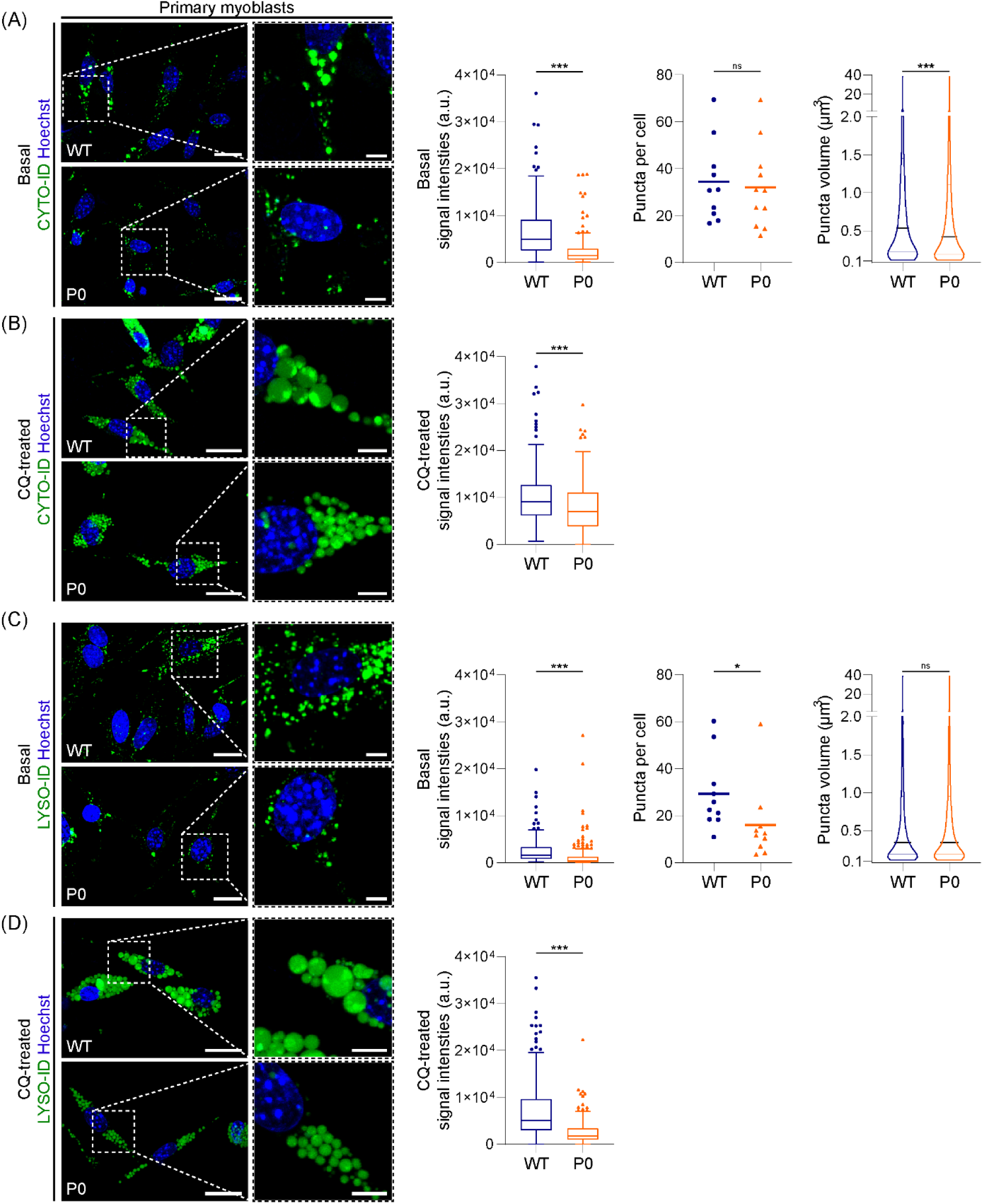
Impaired autophagic flux and reduced capacities of acidic compartments in primary plectin-deficient myoblasts. (A) Primary murine plectin-expressing myoblasts (WT) or plectin-deficient (P0) myoblasts were stained with CYTO-ID at control conditions. Nuclei were visualized with Hoechst. Panels on the right are magnifications of the boxed areas indicated in the panels on the left. Scale bars: 20 µm, magnifications 5 µm. CYTO-ID signal intensities in WT and P0 myoblasts at control conditions were calculated by normalizing the RawIntDens to cell areas. Box plots show the median and Tukey whiskers (WT, n = 177 cells; P0, n = 145 cells). Number of CYTO-ID-positive puncta per cell was determined by using the 3D Objects Counter. Each dot represents a single field-of-view, the line represents the mean (WT, n = 10 field-of-views; P0, n = 14 field-of-views). Volume of CYTO-ID-positive puncta was determined by using the 3D Objects Counter. Violin plots show median, the 25^th^ and 75^th^ percentile (WT, n = 7005 puncta [177 cells]; P0, n = 4474 puncta [145 cells]). (B) WT and P0 myoblasts were treated with 50 µM CQ for 3 h and stained with CYTO-ID. Nuclei were visualized with Hoechst. Panels on the right are magnifications of the boxed areas indicated in the panels on the left. Note the massively swollen vesicles in CQ-treated WT myoblasts, while CQ-treatment of P0 myoblasts caused marginal vesicle swelling. Scale bars: 20 µm, 5 µm magnifications. CYTO-ID signal intensities in CQ-treated WT and P0 myoblasts were calculated by normalizing the RawIntDens to cell areas. Box plots show the median and Tukey whiskers (WT, n = 187 cells; P0, n = 172 cells). (C) WT and P0 myoblasts were stained with LYSO-ID at control conditions. Nuclei were visualized with Hoechst. Panels on the right are magnifications of the boxed areas indicated in the panels on the left. Scale bars: 20 µm, magnifications 5 µm. LYSO-ID signal intensities in WT and P0 myoblasts at control conditions were calculated by normalizing the RawIntDens to cell areas. Box plots show the median and Tukey whiskers (WT, n = 166 cells; P0, n = 155 cells). Number of LYSO-ID-positive puncta per cell was determined by using the 3D Objects Counter. Each dot represents a single field-of-view, the line represents the mean (WT, n = 10 field-of-views; P0, n = 10 field-of-views). Volume of LYSO-ID-positive puncta was determined by using the 3D Objects Counter. Violin plots show median, the 25^th^ and 75^th^ percentile (WT, n = 4574 puncta [166 cells]; P0, n = 2452 puncta [155 cells]). (D) WT and P0 myoblasts were treated with 50 µM CQ for 3 h and stained with LYSO-ID. Nuclei were visualized with Hoechst. Panels on the right are magnifications of the boxed areas indicated in the panels on the left. Note the massively swollen vesicles in CQ-treated WT myoblasts, while CQ-treatment of P0 myoblasts caused marginal vesicle swelling. Scale bars: 20 µm, magnifications 5 µm. LYSO-ID signal intensities in CQ-treated WT and P0 were calculated by normalizing the RawIntDens to cell areas. Box plots show the median and Tukey whiskers (WT, n = 246; P0, n = 186 cells). For (A-D): \**P* < 0.05, \*\*\**P* < 0.001 (two-tailed Mann Whitney test)); ns, not significant.

To recapitulate the apparent block in autophagy in patient-derived cells, primary human dermal fibroblasts from two EBS-MD patients [17–19]) as well as from two healthy controls were immunolabeled for LC3 or SQSTM1 (Figure 7A). Both autophagy marker proteins were clearly enhanced in EBS-MD fibroblasts, with relative SQSTM1signal intensities significantly elevated in EBS-MD fibroblasts 1 and 2 to ∼140% and ∼190%, respectively, of control levels (Figure 7B). Flow cytometric analyses of CYTO-ID-labeled human fibroblasts at basal conditions revealed markedly increased MFIs of EBS-MD 1 and 2 to ∼6.8×10^5^ and ∼1.2×10^6^, respectively (control 1: ∼4.3×10^5^, control 2: ∼4.5×10^5^; Figure 7C). Incubation with CQ for 3 h increased MFIs to comparable levels in EBS-MD and control fibroblasts, which is a significantly lower proportional change in MFIs in EBS-MD fibroblasts compared to control cells, confirming that plectin deficiency coincides with impaired autophagic turnover, not only in myoblasts, but also in other cell types.

**Figure 7.**
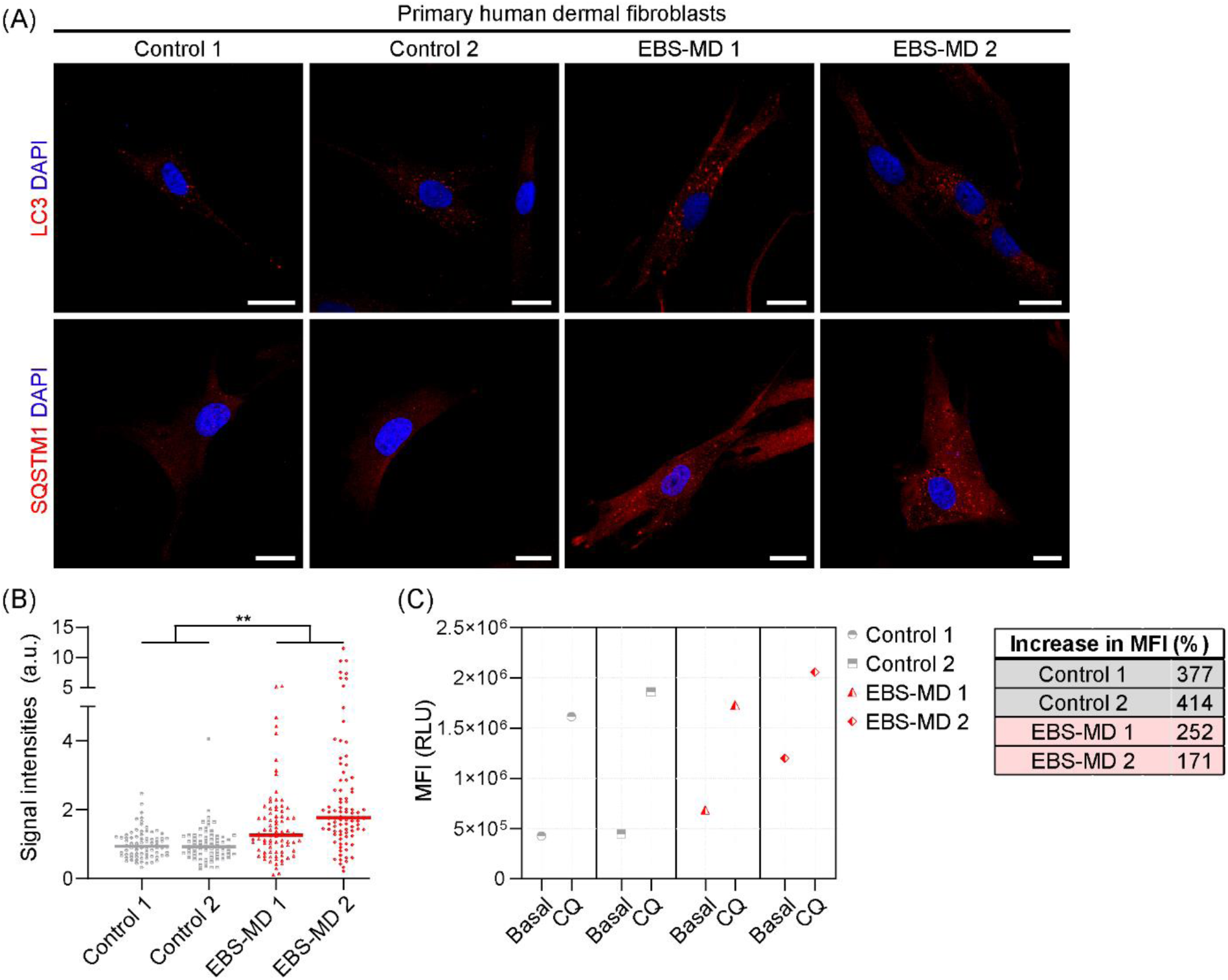
Reduced autophagic capacities in EBS-MD patient fibroblasts. (A) Immunostaining of primary human dermal fibroblasts obtained from healthy controls or EBS-MD patients using antibodies to LC3 or SQSMT1. Nuclei were visualized with DAPI. Scale bars: 20 µm. (B) SQSTM1 signal intensities in control and EBS-MD fibroblasts were calculated by normalizing the RawIntDens to cell areas. Each dot represents a single cell, the line represents the median (control 2, n = 78 cells; control 3, n = 73 cells; EBS-MD 4, n = 82 cells; EBS-MD 5, n = 88 cells); \*\**P* < 0.01 (Kruskal-Wallis test with Dunn’s correction for multiple comparisons). (C) Flow cytometry of CYTO-ID-stained control and EBS-MD fibroblasts at control conditions or treated with 50 µM CQ for 3 h and calculation of the increase in MFI upon treatment with CQ (CQ:control). Each dot represents the MFI (control 2, n = 8.3×10^4^/8.5×10^4^ [control/CQ treated] cells; control 3, n = 7.8×10^4^/6.8×10^4^ [control/CQ treated] cells; EBS-MD 1, n = 8.2×10^4^/7.2×10^4^ [control/CQ treated] cells; EBS-MD 2, n = 7.5×10^4^/4.4×10^4^ [control/CQ treated] cells).

**Figure 8.**
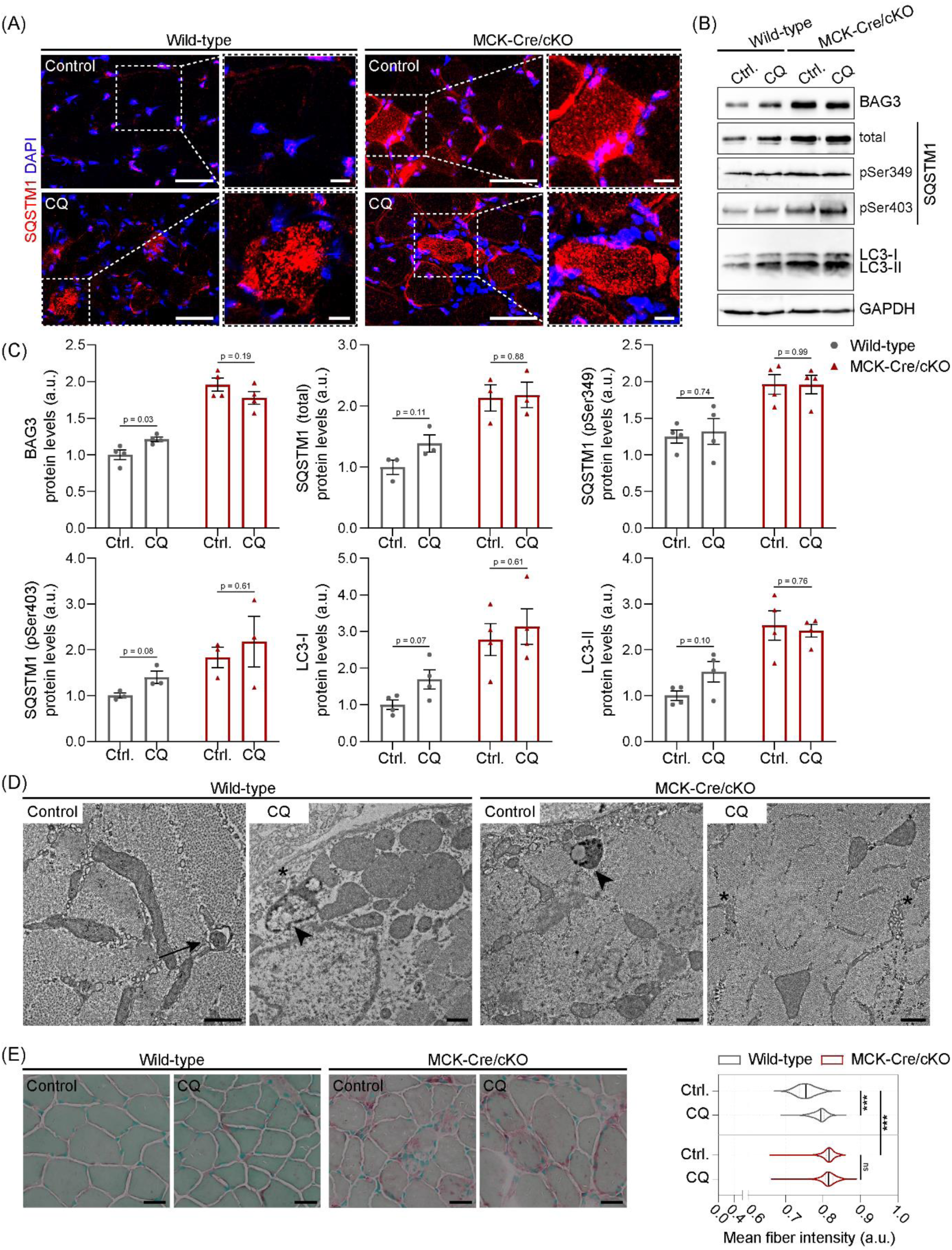
Chloroquine treatment of wild-type and MCK-Cre/cKO mice. (A) Immunostaining using antibodies to SQSTM1 of frozen muscle sections prepared from wild-type and MCK-Cre/cKO mice treated for 4 h ante mortem with either 0.9% saline-(control) or 10 mg/ml CQ-treated. Nuclei were visualized with DAPI. Panels on the right are magnifications of the boxed areas indicated in the panels on the left. Note enhanced SQSTM1 signals in wild-type muscles upon CQ-treatment, whereas the SQSTM1 signals in MCK-Cre/cKO muscles remain similar to saline-treated samples. Scale bars: 20 µm, magnifications 10 µm. (B) Immunoblotting of saline-(Ctrl.) or CQ-treated wild-type and MCK-Cre/cKO muscle lysates using antibodies to BAG3, total and phosphorylated forms of SQSTM1, LC3, and GAPDH. (C) Signal intensities of protein bands as shown in (B) were densitometrically measured and normalized to the total protein content as analyzed by Coomassie staining (not shown). Mean ± SEM; n = 3-4; all *P* values are indicated (unpaired, two-tailed *t*-test with Welch’s correction). (D) Representative electron micrographs of soleus muscle cross sections obtained from saline- or CQ-treated wild-type and MCK-Cre/cKO mice. Note the regular engulfment of cargo (arrow) in saline-treated wild-type muscle as well as the formation of membrane whirls (asterisks) and swollen, degradative vacuoles (arrowhead) in CQ-treated wild-type muscles. Also note that both saline- and CQ-treated plectin-deficient muscles of MCK-Cre/cKO mice display a comparable occurrence of swollen, degradative vacuoles (arrowhead) and membrane whirls (asterisks); no aggravation of autophagolytic changes was observed upon CQ-treatment. Scale bars: 500 nm. (E) Soleus muscle sections from saline- or CQ-treated wild-type and MCK-Cre/cKO mice were stained with acid phosphatase (AP). Note the increased AP staining (red) within CQ-treated wild-type, as well as saline- and CQ-treated MCK-Cre/cKO myofibers. Scale bars: 25 µm. AI-based evaluation of AP signal intensities within individual myofibers using MIRA Vision (wild-type, n = 1129/1359 fibers [control/CQ]; MCK-Cre/cKO, n = 852/1611 fibers [control/CQ]; two animals each); \*\*\**P* < 0.001 (two-way ANOVA of the ranked dataset with Tukey’s post-hoc correction for multiple comparisons); ns, not significant.

### Compromised autophagic flux in plectin-deficient skeletal muscle

Having identified impaired autophagic flux in plectin-deficient myoblasts, we aimed at consolidating this observation in mice. Accordingly, autophagy was inhibited in wild-type and MCK-Cre/cKO mice by intraperitoneal application of CQ for 4 h [14], and frozen muscles were subsequently evaluated. In CQ-treated wild-type muscles we identified multiple fibers with massively enhanced SQSTM1 signals compared to saline-treated controls, indicating accumulation of downstream substrates upon inhibition of autophagy (Figure 7A). Contrary, even though increased SQSTM1 staining pattern was already evident in saline-treated MCK-Cre/cKO muscles, CQ-treatment did not aggravate the pathology. In keeping with this finding, evaluation of BAG3, SQSTM1, Ser403 phosphorylated SQSTM1, LC3-I, and LC3-II revealed an overall trend towards increased autophagy marker proteins in CQ-treated vs. saline-treated wild-type muscles, while CQ-treated MCK-Cre/cKO muscles remained comparable to saline-treated controls (Figure 7B;C). Ultrastructural changes in CQ-treated wild-type samples included numerous swollen vacuoles and membrane whirls, while CQ-treated MCK-Cre/cKO samples showed similar results as saline-treated controls (Figure 7D). In addition, when accumulation of lysosomes was assessed by acid phosphatase enzymatic reactions on muscle sections and AI-based evaluation, relative mean intensities significantly increased in CQ-treated vs. saline-treated wild-type muscles (saline-treated: ∼0.75, CQ-treated: ∼0.79; Figure 7E), while CQ treatment did not shift signals in MCK-Cre/cKO myofibers (saline-treated: ∼0.82, CQ-treated: ∼0.82). Taken together, autophagy inhibition by CQ-treatment had only tenuous effects on the autophagy-associated pathology in MCK-Cre/cKO, reassuring in vivo the concept that autophagic flux is impaired in plectin-deficient muscle.

## Discussion

The present work demonstrated that the characteristic desmin protein aggregation pathology in plectin-deficient skeletal muscles and cells is associated with major alterations in protein quality control pathways. As alterations in virtually every step of the autophagic cascade can contribute to disease processes in myopathies [26], we elaborately explored autophagy-relevant pathways. Our experiments revealed that plectin-deficent muscles significantly accumulate downstream autophagic effector proteins, while leaving the upstream transcriptional regulation as well as the composition of the autophagic machinery largely intact. Since the observed alterations were already present in muscles from young MCK-Cre/cKO mice, these changes are likely contributing to the early pathology of this progressive and devastating muscle disease. At least in part, our results are in line with studies on muscle-specific ATG5 or ATG7 knockout mice showing that muscles that are incapable of performing autophagy develop functional and morphological alterations, including muscle loss, protein aggregates, misalignment of Z-lines and accumulation of abnormal mitochondria [27, 28]. In addition, loss of autophagy in both studies triggered a compensatory increase of the ubiquitin-proteasome degradation pathway [27, 28], similar to the increased levels of ubiquitinated proteins and proteasomal activities we observed in MCK-Cre/cKO muscles.

Since autophagy is a highly dynamic process, analyses on the tissue level only allow a steady assessment of the pathway intermediates without measuring their rate of turnover and degradation, i.e. increased numbers of autophagosomes do not inevitably correspond to increased autophagic activity, but might rather stem from suppressed steps in the autophagic pathway downstream of autophagosome formation [22]. As the exposure to inhibiting or activating conditions is an essential prerequisite for conclusively monitoring autophagic flux [23, 29], we measured baseline, impaired (3-MA, CQ, or Baf A1 treatments) or enhanced autophagy (starvation or Metf treatment) in plectin-deficient myoblasts. Notably, mCherry-EGFP-LC3B-expressing *Plec^-/-^* myoblasts exhibited explicit signs of reduced autophagic flux, i.e. they did rarely respond to the different inhibitors as red:green signal ratios barely dropped. However, long-term impairment of autophagy revealed bright yellow signals, indicating that *Plec^-/-^* myoblasts were in principle capable of performing mCherry-EGFP-LC3B turnover, albeit at a significantly lower rate than *Plec^+/+^* cells, and activation of autophagic flux by starvation or Metf-treatment was more efficient in *Plec^+/+^* than in *Plec^-/-^* myoblasts. Dynamic evaluation of CYTO-ID-, LYSO-ID-, and/or Magic Red-stained primary and immortalized plectin-deficient myoblasts and EBS-MD patient-derived fibroblasts further substantiated that loss of plectin unequivocally provokes a dysfunction in the autophagosomal-lysosomal turnover on the cellular level. Ultimately, a pharmacological blockade of autophagic flux in vivo indicated that only wild-type muscles accumulated downstream substrates, while in MCK-Cre/cKO muscles, albeit already displaying hallmarks of increased autophagy, application of CQ did not aggravate the observed pathology. In conclusion, our study convincingly demonstrated that loss of plectin leads to markedly reduced autophagic clearance capacities in skeletal muscle.

Plectin’s most prominent role is obviously the mechanical stabilization, interconnection, and recruitment of various types of IFs [2], as highlighted by several studies demonstrating increased bundling or even complete collapse of keratin, vimentin, or desmin IF networks in murine *Plec^-/-^*keratinocytes, fibroblasts, or myoblasts, respectively [10, 30, 31]. Plectin-mediated connection between keratin 8 (KRT8) and actin or mitochondria was implicated for correct autophagy and mitophagy, respectively, under oxidative stress in retinal pigment epithelial cells [32, 33]. Intact vimentin networks were prerequisite for autophagosome and lysosome positioning in HEK cells [34], and probably regulate mTORC1 signaling by facilitating its localization to lysosomes [35]. While a direct role for desmin IFs in autophagic transport has not yet been established, desmin deficiency as well as desminopathy-causing mutations lead to imbalanced protein homeostasis in murine muscles [36, 37]. Furthermore, the mere expression of various desmin mutants impact autophagic flux in C2C12 myoblasts [38] and desminopathy patients with autophagic vacuolar myopathies have been reported [39]. In the case of plectinopathies, our analyses suggest that the characteristic accumulation and aggregation of desmin in *PLEC* mutant skeletal muscle cells and tissues results from an impaired autophagic turnover. In conclusion, our study opens a new perspective on the current understanding of the protein aggregation pathology in plectin-related disorders and provides a basis for further pharmacological intervention studies addressing protein homeostasis.

## Supporting information

Supplemental Information_RNA Seq

## Acknowledgements

We thank Dimitra Kiritsi (Department of Dermatology, Medical Center University of Freiburg, Faculty of Medicine, University of Freiburg, Germany) for providing control and EBS-MD fibroblasts, and Renate Kain (Department of Pathology, Medical University of Vienna, Austria) for providing 3-MA. We especially thank Virginie Hubert and Renate Kain (Department of Pathology, Medical University of Vienna, Austria) for helpful discussion and technical advice. The authors of this manuscript certify that they comply with the ethical guidelines for authorship and publishing in the *Journal of Cachexia, Sarcopenia and Muscle* [40].

## References

1. Agnetti G, Herrmann H, Cohen S. New roles for desmin in the maintenance of muscle homeostasis. FEBS J. 2022;289:2755–70. doi:10.1111/febs.15864

2. Wiche G. Plectin-Mediated Intermediate Filament Functions: Why Isoforms Matter. Cells. 2021;10:doi:10.3390/cells10082154

3. Rezniczek GA, Abrahamsberg C, Fuchs P, Spazierer D, Wiche G. Plectin 5’-transcript diversity: short alternative sequences determine stability of gene products, initiation of translation and subcellular localization of isoforms. Hum Mol Genet. 2003;12:3181–94. doi:10.1093/hmg/ddg345

4. Rezniczek GA, Konieczny P, Nikolic B, Reipert S, Schneller D, Abrahamsberg C, et al. Plectin 1f scaffolding at the sarcolemma of dystrophic (mdx) muscle fibers through multiple interactions with beta-dystroglycan. J Cell Biol. 2007;176:965–77. doi:10.1083/jcb.200604179

5. Konieczny P, Fuchs P, Reipert S, Kunz WS, Zeold A, Fischer I, et al. Myofiber integrity depends on desmin network targeting to Z-disks and costameres via distinct plectin isoforms. J Cell Biol. 2008;181:667–81. doi:10.1083/jcb.200711058

6. Winter L, Staszewska-Daca I, Zittrich S, Elhamine F, Zrelski MM, Schmidt K, et al. Z-Disk-Associated Plectin (Isoform 1d): Spatial Arrangement, Interaction Partners, and Role in Filamin C Homeostasis. Cells. 2023;12:doi:10.3390/cells12091259

7. Zrelski MM, Kustermann M, Winter L. Muscle-Related Plectinopathies. Cells. 2021;10:doi:10.3390/cells10092480

8. Vahidnezhad H, Youssefian L, Harvey N, Tavasoli AR, Saeidian AH, Sotoudeh S, et al. Mutation update: The spectra of PLEC sequence variants and related plectinopathies. Hum Mutat. 2022;43:1706–31. doi:10.1002/humu.24434

9. Schröder R, Kunz WS, Rouan F, Pfendner E, Tolksdorf K, Kappes-Horn K, et al. Disorganization of the desmin cytoskeleton and mitochondrial dysfunction in plectin-related epidermolysis bullosa simplex with muscular dystrophy. J Neuropathol Exp Neurol. 2002;61:520–30. doi:10.1093/jnen/61.6.520

10. Winter L, Staszewska I, Mihailovska E, Fischer I, Goldmann WH, Schroder R, et al. Chemical chaperone ameliorates pathological protein aggregation in plectin-deficient muscle. J Clin Invest. 2014;124:1144–57. doi:10.1172/JCI71919

11. Winter L, Türk M, Harter PN, Mittelbronn M, Kornblum C, Norwood F, et al. Downstream effects of plectin mutations in epidermolysis bullosa simplex with muscular dystrophy. Acta Neuropathol Commun. 2016;4:44. doi:10.1186/s40478-016-0314-7

12. Mellerio JE, Smith FJ, McMillan JR, McLean WH, McGrath JA, Morrison GA, et al. Recessive epidermolysis bullosa simplex associated with plectin mutations: infantile respiratory complications in two unrelated cases. Br J Dermatol. 1997;137:898–906.

13. Ackerl R, Walko G, Fuchs P, Fischer I, Schmuth M, Wiche G. Conditional targeting of plectin in prenatal and adult mouse stratified epithelia causes keratinocyte fragility and lesional epidermal barrier defects. Journal of Cell Science. 2007;120:2435–43. doi:10.1242/jcs.004481

14. Moulis M, Vindis C. Methods for Measuring Autophagy in Mice. Cells. 2017;6:doi:10.3390/cells6020014

15. N’Diaye EN, Kajihara KK, Hsieh I, Morisaki H, Debnath J, Brown EJ. PLIC proteins or ubiquilins regulate autophagy-dependent cell survival during nutrient starvation. EMBO Rep. 2009;10:173–9. doi:10.1038/embor.2008.238

16. Swift S, Lorens J, Achacoso P, Nolan GP. Rapid production of retroviruses for efficient gene delivery to mammalian cells using 293T cell-based systems. Curr Protoc Immunol. 2001;Chapter 10:Unit 10 7C. doi:10.1002/0471142735.im1017cs31

17. Kunz M, Rouan F, Pulkkinen L, Hamm H, Jeschke R, Bruckner-Tuderman L, et al. Mutation reports: epidermolysis bullosa simplex associated with severe mucous membrane involvement and novel mutations in the plectin gene. J Invest Dermatol. 2000;114:376–80. doi:10.1046/j.1523-1747.2000.00856.x

18. Natsuga K, Nishie W, Akiyama M, Nakamura H, Shinkuma S, McMillan JR, et al. Plectin expression patterns determine two distinct subtypes of epidermolysis bullosa simplex. Hum Mutat. 2010;31:308–16. doi:10.1002/humu.21189

19. Zrelski MM, Hösele S, Kustermann M, Fichtinger P, Kah D, Athanasiou I, et al. Plectin Deficiency in Fibroblasts Deranges Intermediate Filament and Organelle Morphology, Migration, and Adhesion. J Invest Dermatol. 2023;doi:10.1016/j.jid.2023.08.020

20. Napolitano G, Ballabio A. TFEB at a glance. J Cell Sci. 2016;129:2475–81. doi:10.1242/jcs.146365

21. Zhao YG, Codogno P, Zhang H. Machinery, regulation and pathophysiological implications of autophagosome maturation. Nat Rev Mol Cell Biol. 2021;22:733–50. doi:10.1038/s41580-021-00392-4

22. Mizushima N, Yoshimori T, Levine B. Methods in mammalian autophagy research. Cell. 2010;140:313–26. doi:10.1016/j.cell.2010.01.028

23. Klionsky DJ, Abdel-Aziz AK, Abdelfatah S, Abdellatif M, Abdoli A, Abel S, et al. Guidelines for the use and interpretation of assays for monitoring autophagy (4th edition)(1). Autophagy. 2021;17:1–382. doi:10.1080/15548627.2020.1797280

24. Guo S, Liang Y, Murphy SF, Huang A, Shen H, Kelly DF, et al. A rapid and high content assay that measures cyto-ID-stained autophagic compartments and estimates autophagy flux with potential clinical applications. Autophagy. 2015;11:560–72. doi:10.1080/15548627.2015.1017181

25. Andrä K, Lassmann H, Bittner R, Shorny S, Fässler R, Propst F, et al. Targeted inactivation of plectin reveals essential function in maintaining the integrity of skin, muscle, and heart cytoarchitecture. Genes Dev. 1997;11:3143–56. doi:10.1101/gad.11.23.3143

26. Margeta M. Autophagy Defects in Skeletal Myopathies. Annu Rev Pathol. 2020;15:261–85. doi:10.1146/annurev-pathmechdis-012419-032618

27. Raben N, Hill V, Shea L, Takikita S, Baum R, Mizushima N, et al. Suppression of autophagy in skeletal muscle uncovers the accumulation of ubiquitinated proteins and their potential role in muscle damage in Pompe disease. Hum Mol Genet. 2008;17:3897–908. doi:10.1093/hmg/ddn292

28. Masiero E, Agatea L, Mammucari C, Blaauw B, Loro E, Komatsu M, et al. Autophagy is required to maintain muscle mass. Cell Metab. 2009;10:507–15. doi:10.1016/j.cmet.2009.10.008

29. Mizushima N, Murphy LO. Autophagy Assays for Biological Discovery and Therapeutic Development. Trends Biochem Sci. 2020;45:1080–93. doi:10.1016/j.tibs.2020.07.006

30. Osmanagic-Myers S, Gregor M, Walko G, Burgstaller G, Reipert S, Wiche G. Plectin-controlled keratin cytoarchitecture affects MAP kinases involved in cellular stress response and migration. J Cell Biol. 2006;174:557–68. doi:10.1083/jcb.200605172

31. Burgstaller G, Gregor M, Winter L, Wiche G. Keeping the vimentin network under control: cell-matrix adhesion-associated plectin 1f affects cell shape and polarity of fibroblasts. Mol Biol Cell. 2010;21:3362–75. doi:10.1091/mbc.E10-02-0094

32. Baek A, Son S, Baek YM, Kim DE. KRT8 (keratin 8) attenuates necrotic cell death by facilitating mitochondrial fission-mediated mitophagy through interaction with PLEC (plectin). Autophagy. 2021;17:3939–56. doi:10.1080/15548627.2021.1897962

33. Son S, Baek A, Lee JH, Kim DE. Autophagosome-lysosome fusion is facilitated by plectin-stabilized actin and keratin 8 during macroautophagic process. Cell Mol Life Sci. 2022;79:95. doi:10.1007/s00018-022-04144-1

34. Biskou O, Casanova V, Hooper KM, Kemp S, Wright GP, Satsangi J, et al. The type III intermediate filament vimentin regulates organelle distribution and modulates autophagy. PLoS One. 2019;14:e0209665. doi:10.1371/journal.pone.0209665

35. Mohanasundaram P, Coelho-Rato LS, Modi MK, Urbanska M, Lautenschläger F, Cheng F, et al. Cytoskeletal vimentin regulates cell size and autophagy through mTORC1 signaling. PLoS Biol. 2022;20:e3001737. doi:10.1371/journal.pbio.3001737

36. Winter L, Unger A, Berwanger C, Sporrer M, Turk M, Chevessier F, et al. Imbalances in protein homeostasis caused by mutant desmin. Neuropathol Appl Neurobiol. 2019;45:476–94. doi:10.1111/nan.12516

37. Joanne P, Hovhannisyan Y, Bencze M, Daher MT, Parlakian A, Toutirais G, et al. Absence of Desmin Results in Impaired Adaptive Response to Mechanical Overloading of Skeletal Muscle. Front Cell Dev Biol. 2021;9:662133. doi:10.3389/fcell.2021.662133

38. Sukhareva KS, Smolina NA, Churkina AI, Kalugina KK, Zhuk SV, Khudiakov AA, et al. Desmin mutations impact the autophagy flux in C2C12 cell in mutation-specific manner. Cell Tissue Res. 2023;393:357–75. doi:10.1007/s00441-023-03790-6

39. Weihl CC, Iyadurai S, Baloh RH, Pittman SK, Schmidt RE, Lopate G, et al. Autophagic vacuolar pathology in desminopathies. Neuromuscul Disord. 2015;25:199–206. doi:10.1016/j.nmd.2014.12.002

40. von Haehling S, Coats AJS, Anker SD. Ethical guidelines for publishing in the Journal of Cachexia, Sarcopenia and Muscle: update 2021. J Cachexia Sarcopenia Muscle. 2021;12:2259–61. doi:10.1002/jcsm.12899

