## Supplemental Information for "Impaired autophagic flux in skeletal muscle of plectin-related epidermolysis bullosa simplex with muscular dystrophy"

*Supplementary information*

Michaela M. Zrelski[a], Margret Eckhard[a], Petra Fichtinger[a], Sabrina Hösele[a], Andy Sombke[a], Leonid Mill[b], Monika Kustermann[c], Wolfgang M. Schmidt[c], Fiona Norwood[d], Ursula Schlötzer-Schrehardt[e], Gerhard Wiche[f], Rolf Schröder[g], and Lilli Winter[a]

[a]Division of Cell and Developmental Biology, Center for Anatomy and Cell Biology, Medical University of Vienna, Vienna, Austria
[b]MIRA Vision Microscopy GmbH, Wangen, Germany
[c]Neuromuscular Research Group, Division of Cell and Developmental Biology, Center for Anatomy and Cell Biology, Medical University of Vienna, Vienna, Austria
[d] Department of Neurology, Ruskin Wing, King's College Hospital, London, UK
[e]Department of Ophthalmology, University Hospital Erlangen, Friedrich-Alexander University Erlangen-Nürnberg, Erlangen, Germany
[f]Department of Biochemistry and Cell Biology, Max Perutz Laboratories, University of Vienna, Vienna, Austria
[g]Institute of Neuropathology, University Hospital Erlangen, Friedrich-Alexander University Erlangen-Nürnberg, Erlangen, Germany

### Supplemental Methods

#### EBS-MD patient-derived material used in this study

| Material | # | Mutation 1 DNA | Protein | Mutation 2 DNA | Protein | Ref. |
|---|---|---|---|---|---|---|
| Muscle | 1 | 2264_2266delTCT | Phe755del | 3119_3120delAA | Lys1040Argfs*139 | [1] |
| Muscle | 2 | 13459_13474dup | Glu4492Glyfs*48 | 13459_13474dup | Glu4492Glyfs*48 | [3] |
| Muscle | 3 | 5018_5036del | Leu1673Argfs*64 | 5018_5036del | Leu1673Argfs*64 | [2] |
| Fibrobl. | 1 | 4643_4667dup | Lys1558Glyfs*89 | 7120C>T | Gln2374* | [5,6] |
| Fibrobl. | 2 | 5137C>T | Gln1713* | 7051C>T | Arg2351* | [4-6] |

Fibrobl., fibroblasts; Ref., references

#### Transmission electron microscopy

Human biopsy material from the left M. triceps brachii from EBS-MD patient 1 was fixed in freshly prepared 2.5% glutaraldehyde in 0.1 M Sørensen's phosphate buffer, pH 7.2, post-fixed in 2% buffered osmium tetroxide, dehydrated in graded ethanol concentrations, and embedded in epoxy resin. 1 µm semithin sections for orientation were stained with toluidine blue. Ultrathin sections were stained with uranyl acetate and lead citrate and examined with a LEO 906E transmission electron microscope (Carl Zeiss Microscopy GmbH, Oberkochen, Germany).

Freshly isolated murine soleus muscles were pinned to agarose plates, fixed with 2% paraformaldehyde (PFA) and 2.5% glutaraldehyde (Sigma-Aldrich, G5882) in 0.1 M sodium cacodylate buffer (CB, pH 7.4; PanReac AppliChem, A2140,0250) overnight at 4°C. Samples were subsequently washed in CB [7]. Postfixation was performed in a solution of 1.5% potassium hexacyanoferrate (III) (Sigma-Aldrich, 60299) and 1% osmium tetroxide (Electron Microscopy Sciences, 19110) in CB for 1h. Immortalized myoblasts were washed in PBS and fixed for 10 min in 4% PFA and 2.5% glutaraldehyde in PBS. Cells were carefully scraped off the dishes and pelleted by centrifugation (5 min, 200 x g, 10°C) and fixed for another hour in the same fixative. For postfixation, samples were incubated in 1% osmium tetroxide in water for 1 h. Subsequently, all samples were washed in either CB (M. soleus) or PBS (myoblasts), dehydrated in an ascending ethanol series (30% - 100%), and embedded in epoxy resin (Serva, Epon 812). Ultrathin sections (40-50 nm) were prepared with a UC7 ultramicrotome (Leica Biosystems, Germany), mounted on copper mesh grids and contrasted with either 2% (soleus) or 1% (myoblasts) uranyl acetate (Merck, 8473) and 3% lead citrate (Sigma-Aldrich, 15326). TEM analysis was carried out at a Tecnai G2 20 transmission electron microscope (FEI Company, USA) operating at 80kV, equipped with a FEI Eagle 4K CCD-camera. Subsequent image enhancement was conducted using Fiji [8], utilizing global contrast and brightness adjustments as well as the CLAHE filter for local contrast improvement.

**Immunofluorescence microscopy**

Muscles were snap-frozen in isopentane cooled with dry ice. Thin sections (5-10 µm) from human skeletal muscles were fixed with pre-chilled acetone, blocked with 4% bovine serum albumin (BSA; Pan Biotech, P06-1391050) in PBS, and immunostained as previously described [1]. Thin sections of murine M. soleus were fixed with pre-chilled acetone and immunostained using the M.O.M Basic Kit (Vector Laboratories, BMK-2202) [9]. Immortalized myoblasts were grown on Geltrex-coated (Gibco, A1413202) glass coverslips (#1.5), human dermal fibroblasts on glass coverslips, washed with PBS, and fixed with 3.7% PFA for 10 min at room temperature. Following a 10 min permeabilization with 0.05% Triton X-100 (Sigma-Aldrich, T8787), cells were immunostained as previously described [10]. Nuclei were visualized with DAPI (Sigma Aldrich, 10236276001; 2 µg/ml). Microscopy was performed using an Olympus FLUOVIEW FV3000 confocal microscope equipped with PlanApo N 60x 1.4 NA and UPLAN FLN 40x 1.3 NA objective lenses (Olympus, Japan). Z-stacks were recorded using the Olympus FluoView software and processed with ImageJ software to generate maximum intensity projections unless stated otherwise. Cell contours were measured, raw integrated density (RawIntDens) and cell area were analyzed, and the relative signal intensities were calculated by normalizing the RawIntDens to the cell area.

**Antibodies**

The following primary antibodies were used for immunofluorescence micsroscopy and immunoblotting: rabbit monoclonal antibodies (mAbs) to LC3A/B (Cell Signaling Technology, D3U4C), mouse mAbs to desmin (Dako, D33), rabbit mAbs to p62/SQSTM1 (Sigma-Aldrich, P0067), rabbit polyclonal antibodies (pAbs) to TFEB (Bethyl Laboratories, A303-673A), rabbit pAbs to GAPDH (Sigma-Aldrich, G9545]), rabbit mAbs to mTOR (Cell Signaling Technology, 7C10), rabbit mAbs to ULK1 (Cell Signaling Technology, D8H5), rabbit mAbs to Beclin-1 (Cell Signaling Technology, D40C5), rabbit mAbs to ATG7 (Cell Signaling Technology, D12B11), rabbit mAbs to ATG5 (Cell Signaling Technology, D5F5U), rabbit pAbs to ATG3 (Cell Signaling Technology, 3414), mouse mAbs to ubiquitin (Enzo Life Sciences, P4D1), rabbit pAbs to LAMP2 (Invitrogen, PA1-655), rabbit pAbs to BAG3 (Proteintech Group, 10599), rabbit mAbs to phospho-SQSTM1 (Ser349; Cell Signaling Technology, E7MIA), rabbit mAbs to phospho-SQSTM1 (Ser403; Cell Signaling Technology, D8D6T), mouse mAbs to α-actinin (Sigma-Aldrich, EA-53), and mouse mAbs to proteasome 20S (α1, 2, 3, 5, 6 & 7 subunits; Enzo Life Sciences, MCP231). For immunofluorescence microscopy primary antibodies were used in combination with donkey anti-mouse IgG Alexa fluor 488$^+$, donkey anti-mouse IgG Alexa fluor 555$^+$, donkey anti-rabbit IgG Alexa fluor 488$^+$, and donkey anti-rabbit IgG Alexa fluor 555$^+$ (all from Invitrogen, A32766, A32773, A32790, A32794, respectively). Biotinylated anti-mouse antibodies from the M.O.M. Basic Kit were detected using streptavidin-conjugated Alexa fluor 488 (Invitrogen, S11223). For immunoblotting analyses, HRP-conjugated secondary antibodies were used (Jackson ImmunoResearch Laboratories, AB_10015289 [mouse], AB_2313567 [rabbit]).

**AI-based evaluation of whole muscle sections**

Whole soleus muscle sections, either immunolabeled using antibodies to SQSTM1 or stained with acid phosphatase enzymatic reactions, were scanned with an Olympus VS-BX slide scanner equipped with an UPLSAPO 2 40x NA0.95 objective lens (Olympus). To quantify signal intensities an AI-algorithm modified from the MIRA Vision platform (MIRA Vision Microscopy, Germany; https://www.mira.vision/), originally designed for recognizing hematoxylin-eosin-stained fibers in whole muscle sections, was used. Individual myofibers were automatically identified in an AI-generated mask; signal intensities were obtained for each fiber and normalized to the color depth of the image. For SQSTM1 analysis, intensities were binned within each genotype (bin size = 0.1) and presented as histograms displaying the frequency distribution of binned intensities obtained from two animals per genotype.

**RNA-sequencing (RNA-Seq)**

Cryosections of M. soleus from 13-week-old male mice snap-frozen in isopentane cooled with dry ice were dissociated by gentle trituration in RNAeasy Mini Kit (Qiagen, 74104) isolation buffer supplemented with 2-mercaptoethanol (Sigma-Aldrich, 63689). Total RNA was isolated using the RNAeasy Mini Kit following the manufacturer's protocol and stored at -80°C. RNA aliquots were shipped on dry ice to CeGaT GmbH (Tübingen, Germany), where RNA quality control, library preparation and next-generation sequencing was performed. RNA-Seq was carried out from libraries prepared employing the SMARTer Stranded Total RNA-Seq Kit - Pico Input Mammalian (Takara Bio), starting from 3.36 ng RNA input (quantified by a Qubit fluorometer, Thermo Fisher Scientific; the average RNA quality, measured by Bioanalyzer RNA, Agilent, was RIN ~9 [minimum 8.3]). 100 bp paired-end sequencing (with an average output of ~61 million read pairs yielding ~12 Gbp) was performed on a NovaSeq 6000

Sequencing System (Illumina, USA). Raw sequencing reads were demultiplexed with *bcl2fastq*, adapter trimming was performed with *Skewer*, and residual 3 bp corresponding to the SMART adapter were removed from mate reads by *cutadapt*. Data analysis starting from *.fastq.gz files was then performed, starting by filtering for full-length read pairs and mapping to the mouse reference genome sequence (build mm10/GRCm38.p6) using the algorithm *Hisat2* [11] and *samtools* for conversion to sorted *.bam files. Visualization of alignments was done in Integrative Genomics Viewer (IGV) [12]. Transcript expression quantification was conducted by using the algorithm toolset *Salmon v1.1* [13] with an index prepared from the GENCODE release M23 (GRCm38.p6) annotation set (optional parameters for the *salmon quant* function were as follows: --libType ISR, --validateMappings, --seqBias). Salmon quantification results were then analyzed in *R* (3.6.1) and *BioConductor* (3.9) using the *edgeR* (3.26.8) and *tximeta* (1.1.18) packages [14, 15]. Normalized gene expression levels (counts per million, cpm) were calculated, log-transformed and then finally used for downstream differential gene expression analyses, which was finally focused on alterations in the KEGG "04140 – mmu Autophagy – animal" and "mmu03050 Proteasome" pathway networks. Full data of differentially expressed genes are available in the Supplemental Table 1.

**Preparation of muscle and cell lysates, SDS-PAGE, and immunoblotting**

For quantitative immunoblotting, snap-frozen 3-6 M. triceps surae from age-matched wild-type or MCK-Cre/cKO mice were pooled and processed as previously described [9], heated to 60°C for 10 min, and stored at -80°C. Lysates were thawed on ice, supplemented with 6x SDS sample buffer consisting of 500 mM Tris-HCl pH 6.8, 600 mM DDT (Sigma-Aldrich, 11583786001), 10% SDS, 0.1% bromphenol blue, and 30% glycerol (Sigma-Aldrich, G5516), and heated to 60°C for 10 min (for LC3) or 95°C for 5 min. Myoblasts were washed with PBS and directly scraped off in 6x SDS sample buffer, DNA sheared by pressing the samples through a 27-gauge needle, heated to 60°C for 10 min (for LC3) or 95°C for 5 min, and stored at -20°C [9]. SDS-PAGE was performed according to [16]. Protein levels in lysates were determined by Coomassie staining of gels (Coomassie Brilliant Blue R 250 [Sigma-Aldrich, B0149], 50% methanol, 10% acetic acid), and quantification and normalization using ImageJ software (NIH, USA). For immunoblotting, proteins were transferred to either polyvinylidene difluoride (PVDF; Hybond 0.2 µm; Amersham, GE10600021) for LC3 blots or nitrocellulose (Protran 0.2 µm; Amersham, GE10600001) membranes using a Mini PROTEAN Tetra Cell blot apparatus (Bio-Rad Laboratories). Membranes were scanned with Fusion FX (Vilber Lourmat, Germany), and the amounts of protein contained in individual bands were quantified using ImageJ software.

#### Analysis of proteasomal activities

Chymotrypsin-, trypsin- and caspase-like proteasomal activities of M. soleus from 13-week-old or 30-week-old animals, or of $Plec^{+/+}$ and $Plec^{-/-}$ myoblasts, were measured using the Proteasome-Glo Assay (Promega, G8531) as described in [17].

#### Real-time quantitative PCR (RT-qPCR)

Total RNA from immortalized myoblasts was isolated using the RNeasy Mini Kit according to the manufacturer’s instructions, and then reverse transcribed into cDNA using superscript IV reverse transcriptase (Invitrogen, 18090010). RT-qPCR was performed using the SensiMix HI-ROX Kit (Meridian Bioscience, QT605-05) and a CFX96 Touch System (Bio-Rad Laboratories, USA). Relative gene expression levels were determined according to a modified $2^{-\Delta\Delta CT}$ equation, and normalization was performed against a common calibrator calculated from $Plec^{+/+}$ myoblasts [18, 19]. *Tbp* and *Hprt* were used as internal reference genes.

#### List of oligonucleotide primers

| Gene | Accession number | Forward | Reverse | Reference |
|---|---|---|---|---|
| *Ulk1* | NM_001347394_1 | CACTGCGTGGCTCACCTAAG | AGCCAACAGGGTCAGCAAAT | [20] |
| *Becn1* | NM_019584_4 | CAGGAACTCACAGCTCCATTAC | CCATCCTGGCGAGTTTCAATA | [21] |
| *Lamp2* | NM_001017959_2 | TAGGAGCCGTTCAGTCCAAT | GTGTGTCGCCTTGTCAGGTA | [22] |
| *Bag3* | NM_013863_5 | CTGGGAGATCAAAATCGACCC | GCTGAAGATGCAGTGTCCTTAG | [23] |
| *Sqstm1* | NM_011018_3 | TGCTCTTCGGAAGTCAGCAA | CCCGACTCCATCTGTTCCTC | [22] |
| *Map1cl3a* | NM_025735_3 | CTTCGCCGACCGCTGTAA | CGCCGGATGATCTTGACC | [24] |
| *Map1cl3b* | NM_026160_5 | CGATACAAGGGGGAGAAGCA | ACTTCGGAGATGGGAGTGGA | [22] |
| *Hprt* | NM_013556.2 | TGACACTGGCAAAACAATGCA | GGTCCTTTTCACCAGCAAGCT | - |
| *Tbp* | NM_013684.3 | CCTTGTACCCTTCACCAATGAC | ACAGCCAAGATTCACGGTAGA | - |

#### Generation of mCherry-EGFP-LC3B-expressing myoblasts

Immortalized skeletal myoblasts were derived from $Plec^{+/+}$ or $Plec^{-/-}$ littermates, both crossed into a p53-deficient (*p53*$^{-/-}$) background, as previously described [9] and used at passages numbers 25-35. Myoblasts were cultivated in F-10-based growth medium consisting of Ham's F-10 (Gibco, 31550031) supplemented with 20% fetal calf serum (FCS; Sigma Aldrich, F7524), 50 U/ml penicillin, and 50 µg/ml streptomycin (Gibco, 10378016), 25 µg/ml amphotericin B (Gibco, 15290026) and human basic fibroblast growth factor (rhFGF; Promega, G5071) on collagen-coated (0.01% PureCol Bovine Collagen type I [Cellsystems, 5005-B] in phosphate-buffered saline [PBS; Gibco, 10010056]) Nunc$^{TM}$ cell culture dishes (Thermo Scientific) at 37°C and 5% $CO_2$. To generate myoblasts stably expressing pBabe puro mCherry-EGFP-LC3B (from J. Debnath [Addgene, 22418] [25]), retroviral supernatants were generated by transfecting phoenix eco cells as previously described [26]. Phoenix eco cells (ATCC, CRL-3214) were routinely cultured in DMEM supplemented with 10% FCS, 2 mM L-glutamine, 50 U/ml penicillin, and 50µg/ml streptomycin on Nunc$^{TM}$ cell culture dishes at 37°C and 5% $CO_2$, however, viral particles were released into F-10-based growth medium. $1.2x10^5$ *Plec*$^{+/+}$ and *Plec*$^{-/-}$ myoblasts were seeded in 6 cm dishes, transduced using the retroviral supernatant supplemented with 5 µg/ml polybrene (Sigma-Aldrich, TR-1003-G) for 48 h and, after a recovery phase of 24 h in F-10-based growth medium, selected through the addition of 5 µg/ml puromycin (Sigma-Aldrich, P8833). Transduced myoblasts were cultivated in F-10-based growth medium supplemented with 2.5 µg/ml puromycin.

**Life cell analyses of vesicle dynamics**

To evaluate dynamics of autophagic vesicles at basal conditions, $2x10^4$ mCherry-EGFP-LC3B-expressing myoblasts were seeded in collagen-coated µ-Slides (Ibidi, 80806) and imaged after 24 h. For measuring autophagic flux, cells were treated with inhibitors (50 µM CQ [Enzo Life Sciences; in F-10 medium supplemented with 20% FCS], 200 nM Baf A1 [Sigma Aldrich, B1793; in F-10 medium supplemented with 20% FCS] or 9 mM 3-MA [Sigma-Aldrich, M9281; in DMEM medium supplemented with 10% FCS]) for 24 h, or with an activator (100 mM Metf [Sigma Aldrich, PHR1084; in F-10 medium supplemented with 20% FCS]) for 48 h. To starve cells, myoblasts were cultivated in DMEM or DMEM supplemented with 10 mM 3-MA for 24 h. Live cell imaging was performed using an Olympus FLUOVIEW FV3000 confocal microscope (40X objective) equipped with a cellVivo environmental chamber (Olympus) at 37°C. At least 3 randomly chosen field-of-views were acquired per condition and experiment. To calculate red:green ratios, cell contours were measured and RawIntDens of red and green signals of individual cells were analyzed using ImageJ software.

To investigate vesicle dynamics in primary myoblasts, WT and P0 cells were seeded in collagen-coated µ-Slides 24 h prior staining with either CYTO-ID® Autophagy detection kit 2.0

(CYDO-ID; Enzo Life Sciences, ENZ-KIT175; 1:500) or LYSO-ID® Green detection kit (LYSO-ID; Enzo Life Sciences, ENZ-51034; 1:1000) according to the manufacturer's instructions. After washing, cells in assay buffer supplemented with 5% FCS were analyzed using an Olympus FLUOVIEW FV3000 confocal microscope (60X objective) equipped with a cellVivo environmental chamber at 37°C. For measuring vesicle turnover, cells were treated with 50 µM CQ for 3 h. Nuclei were visualized with Hoechst 33342 (Enzo Life Sciences; 1:1000). At least 4 randomly chosen field-of-views were acquired per condition and experiment.

**Analysis of puncta number and volumes**

Maximum intensity projections were generated from Z-stacks, cell contours measured, and either the red (mCherry-EGFP-LC3B myoblasts) or green channel (primary myoblasts) from the original Z-Stacks selected for further analysis. Unevenly illuminated background was corrected by using the Rolling Ball Background Subtraction plugin (ImageJ, 50 pixel), pictures were thresholded, watershed, and subjected to the 3D Objects Counter algorithm (ImageJ) [27]. The lower cut-off was set to 2 (mCherry-EGFP-LC3B myoblasts) or 3 voxels (primary myoblasts), corresponding to a minimum size of 0.154/0.115 $\mu m^3$ (r = 0.333/0.302 µm), to ensure to measure only puncta above the physical resolution limit. Due to limitations of the algorithm distinguishing puncta in close proximity to each other, volumes of puncta larger than 38.35 $\mu m^3$ (mCherry-EGFP-LC3B myoblasts) or 38.57 $\mu m^3$ (primary myoblasts) were set to 38.35 and 38.57 $\mu m^3$, respectively. The number of puncta per cell was calculated by dividing the number of counted objects by the number of cells in each frame.

**Flow cytometry**

Cells were seeded 48 h prior to the experiment, trypsinized, counted and stained according to the manufacturer's instructions using $5x10^5$ cells per 500 µl staining solution (CYTO-ID, 1:1000; LYSO-ID, 1:500). Cells were washed, resuspended in assay buffer supplemented with 5% FCS, kept at 4°C, and measured with a CytoFlex (Beckman Coulter, USA) flow cytometer. To determine vesicle turnover, cells were treated with 50 µM CQ for 3 h.

**Analysis of cathepsin B activity**

$2x10^4$ immortalized myoblasts were seeded in collagen-coated µ-Slides 48 h prior to staining with Magic Red Cathepsin B Assay Kit (ImmunoChemistry Technologies, 6133; 1:260) for 45 min according to the manufacturers' instruction. Nuclei were stained with Hoechst 33342

(ImmunoChemistry Technologies, 639; 1 µg/ml). After washing, cells were kept in PBS and imaged with a FLUOVIEW FV3000 confocal microscope (40X objective) equipped with a cellVivo environmental chamber (Olympus) set to 37°C. At least 5 randomly chosen field-of-views were acquired per condition and experiment.

**Histology**

Muscles were snap-frozen in isopentane cooled with dry ice. Thin sections (7 µm) were air-dried, stained with acid phosphatase enzymatic reactions, and counterstained as described previously [28]. After dehydration by an ascending ethanol series to xylene, the specimens were mounted with DPX (Sigma-Aldrich, 1.00579).

**Statistical analysis**

Data analyses and statistical evaluations were performed using MS Excel or GraphPad Prism. The number of experiments is indicated in the figure legends. Comparisons between two groups were performed using either parametric (two-tailed, unpaired *t*-test with Welch's correction) or nonparametric (two-tailed, Mann-Whitney test) methods, depending on the normal distribution as determined by the D'Agostino-Pearson normality test. Comparisons among values of multiple groups with nonparametric normal distribution were performed using a Kruskal-Wallis test with Dunn's correction for multiple comparisons (one variable). In cases of two dependent variables, data were converted into ranks and analyzed using a two-way ANOVA with Tukey's post-hoc correction for multiple comparisons. *P*-values are *<0.05, **<0.01, and ***<0.001; a P-value <0.05 was considered statistically significant, except for the RNA-Seq analyses where a *P*-value < 0.01 was considered statistically significant. In general, three independent experiments were evaluated.

**References**


1. Winter L, Türk M, Harter PN, Mittelbronn M, Kornblum C, Norwood F, et al. Downstream effects of plectin mutations in epidermolysis bullosa simplex with muscular dystrophy. Acta Neuropathol Commun. 2016;4:44. doi:10.1186/s40478-016-0314-7
2. Mellerio JE, Smith FJ, McMillan JR, McLean WH, McGrath JA, Morrison GA, et al. Recessive epidermolysis bullosa simplex associated with plectin mutations: infantile respiratory complications in two unrelated cases. Br J Dermatol. 1997;137:898-906.
3. Schröder R, Kunz WS, Rouan F, Pfendner E, Tolksdorf K, Kappes-Horn K, et al. Disorganization of the desmin cytoskeleton and mitochondrial dysfunction in plectin-related epidermolysis bullosa simplex with muscular dystrophy. J Neuropathol Exp Neurol. 2002;61:520-30. doi:10.1093/jnen/61.6.520

4. Kunz M, Rouan F, Pulkkinen L, Hamm H, Jeschke R, Bruckner-Tuderman L, et al. Mutation reports: epidermolysis bullosa simplex associated with severe mucous membrane involvement and novel mutations in the plectin gene. J Invest Dermatol. 2000;114:376-80. doi:10.1046/j.1523-1747.2000.00856.x
5. Natsuga K, Nishie W, Akiyama M, Nakamura H, Shinkuma S, McMillan JR, et al. Plectin expression patterns determine two distinct subtypes of epidermolysis bullosa simplex. Hum Mutat. 2010;31:308-16. doi:10.1002/humu.21189
6. Zrelski MM, Hösele S, Kustermann M, Fichtinger P, Kah D, Athanasiou I, et al. Plectin Deficiency in Fibroblasts Deranges Intermediate Filament and Organelle Morphology, Migration, and Adhesion. J Invest Dermatol. 2023;doi:10.1016/j.jid.2023.08.020
7. Winter L, Staszewska-Daca I, Zittrich S, Elhamine F, Zrelski MM, Schmidt K, et al. Z-Disk-Associated Plectin (Isoform 1d): Spatial Arrangement, Interaction Partners, and Role in Filamin C Homeostasis. Cells. 2023;12:doi:10.3390/cells12091259
8. Schindelin J, Arganda-Carreras I, Frise E, Kaynig V, Longair M, Pietzsch T, et al. Fiji: an open-source platform for biological-image analysis. Nat Methods. 2012;9:676-82. doi:10.1038/nmeth.2019
9. Winter L, Staszewska I, Mihailovska E, Fischer I, Goldmann WH, Schroder R, et al. Chemical chaperone ameliorates pathological protein aggregation in plectin-deficient muscle. J Clin Invest. 2014;124:1144-57. doi:10.1172/JCI71919
10. Winter L, Abrahamsberg C, Wiche G. Plectin isoform 1b mediates mitochondrion-intermediate filament network linkage and controls organelle shape. J Cell Biol. 2008;181:903-11. doi:10.1083/jcb.200710151
11. Kim D, Paggi JM, Park C, Bennett C, Salzberg SL. Graph-based genome alignment and genotyping with HISAT2 and HISAT-genotype. Nat Biotechnol. 2019;37:907-15. doi:10.1038/s41587-019-0201-4
12. Robinson JT, Thorvaldsdottir H, Winckler W, Guttman M, Lander ES, Getz G, et al. Integrative genomics viewer. Nat Biotechnol. 2011;29:24-6. doi:10.1038/nbt.1754
13. Patro R, Duggal G, Love MI, Irizarry RA, Kingsford C. Salmon provides fast and bias-aware quantification of transcript expression. Nat Methods. 2017;14:417-9. doi:10.1038/nmeth.4197
14. Robinson MD, McCarthy DJ, Smyth GK. edgeR: a Bioconductor package for differential expression analysis of digital gene expression data. Bioinformatics. 2010;26:139-40. doi:10.1093/bioinformatics/btp616
15. Pertea M, Love MI, Soneson C, Hickey PF, Johnson LK, Pierce NT, et al. Tximeta: Reference sequence checksums for provenance identification in RNA-seq. PLOS Computational Biology. 2020;16:doi:10.1371/journal.pcbi.1007664
16. Laemmli UK. Cleavage of structural proteins during the assembly of the head of bacteriophage T4. Nature. 1970;227:680-5. doi:10.1038/227680a0
17. Strucksberg KH, Tangavelou K, Schröder R, Clemen CS. Proteasomal activity in skeletal muscle: a matter of assay design, muscle type, and age. Anal Biochem. 2010;399:225-9. doi:10.1016/j.ab.2009.12.026
18. Vandesompele J, De Preter K, Pattyn F, Poppe B, Van Roy N, De Paepe A, et al. Accurate normalization of real-time quantitative RT-PCR data by geometric averaging of multiple

internal control genes. Genome Biol. 2002;3:RESEARCH0034. doi:10.1186/gb-2002-3-7-research0034
19. Hellemans J, Mortier G, De Paepe A, Speleman F, Vandesompele J. qBase relative quantification framework and software for management and automated analysis of real-time quantitative PCR data. Genome Biol. 2007;8:R19. doi:10.1186/gb-2007-8-2-r19
20. Goldberg AA, Nkengfac B, Sanchez AMJ, Moroz N, Qureshi ST, Koromilas AE, et al. Regulation of ULK1 Expression and Autophagy by STAT1. J Biol Chem. 2017;292:1899-909. doi:10.1074/jbc.M116.771584
21. Ren J, Xu X, Wang Q, Ren SY, Dong M, Zhang Y. Permissive role of AMPK and autophagy in adiponectin deficiency-accentuated myocardial injury and inflammation in endotoxemia. J Mol Cell Cardiol. 2016;93:18-31. doi:10.1016/j.yjmcc.2016.02.002
22. Yamamoto J, Kamata S, Miura A, Nagata T, Kainuma R, Ishii I. Differential adaptive responses to 1- or 2-day fasting in various mouse tissues revealed by quantitative PCR analysis. FEBS Open Bio. 2015;5:357-68. doi:10.1016/j.fob.2015.04.012
23. Tang M, Ji C, Pallo S, Rahman I, Johnson GVW. Nrf2 mediates the expression of BAG3 and autophagy cargo adaptor proteins and tau clearance in an age-dependent manner. Neurobiol Aging. 2018;63:128-39. doi:10.1016/j.neurobiolaging.2017.12.001
24. Shi Y, Jia M, Xu L, Fang Z, Wu W, Zhang Q, et al. miR-96 and autophagy are involved in the beneficial effect of grape seed proanthocyanidins against high-fat-diet-induced dyslipidemia in mice. Phytother Res. 2019;33:1222-32. doi:10.1002/ptr.6318
25. N'Diaye EN, Kajihara KK, Hsieh I, Morisaki H, Debnath J, Brown EJ. PLIC proteins or ubiquilins regulate autophagy-dependent cell survival during nutrient starvation. EMBO Rep. 2009;10:173-9. doi:10.1038/embor.2008.238
26. Swift S, Lorens J, Achacoso P, Nolan GP. Rapid production of retroviruses for efficient gene delivery to mammalian cells using 293T cell-based systems. Curr Protoc Immunol. 2001;Chapter 10:Unit 10 7C. doi:10.1002/0471142735.im1017cs31
27. Bolte S, Cordelieres FP. A guided tour into subcellular colocalization analysis in light microscopy. J Microsc. 2006;224:213-32. doi:10.1111/j.1365-2818.2006.01706.x
28. Hildyard JCW, Foster EMA, Wells DJ, Piercy RJ. Rapid histological quantification of muscle fibrosis and lysosomal activity using the HSB colour space. bioRxiv. 2022;2022.08.02.502489. doi:10.1101/2022.08.02.502489

### Supplemental Information

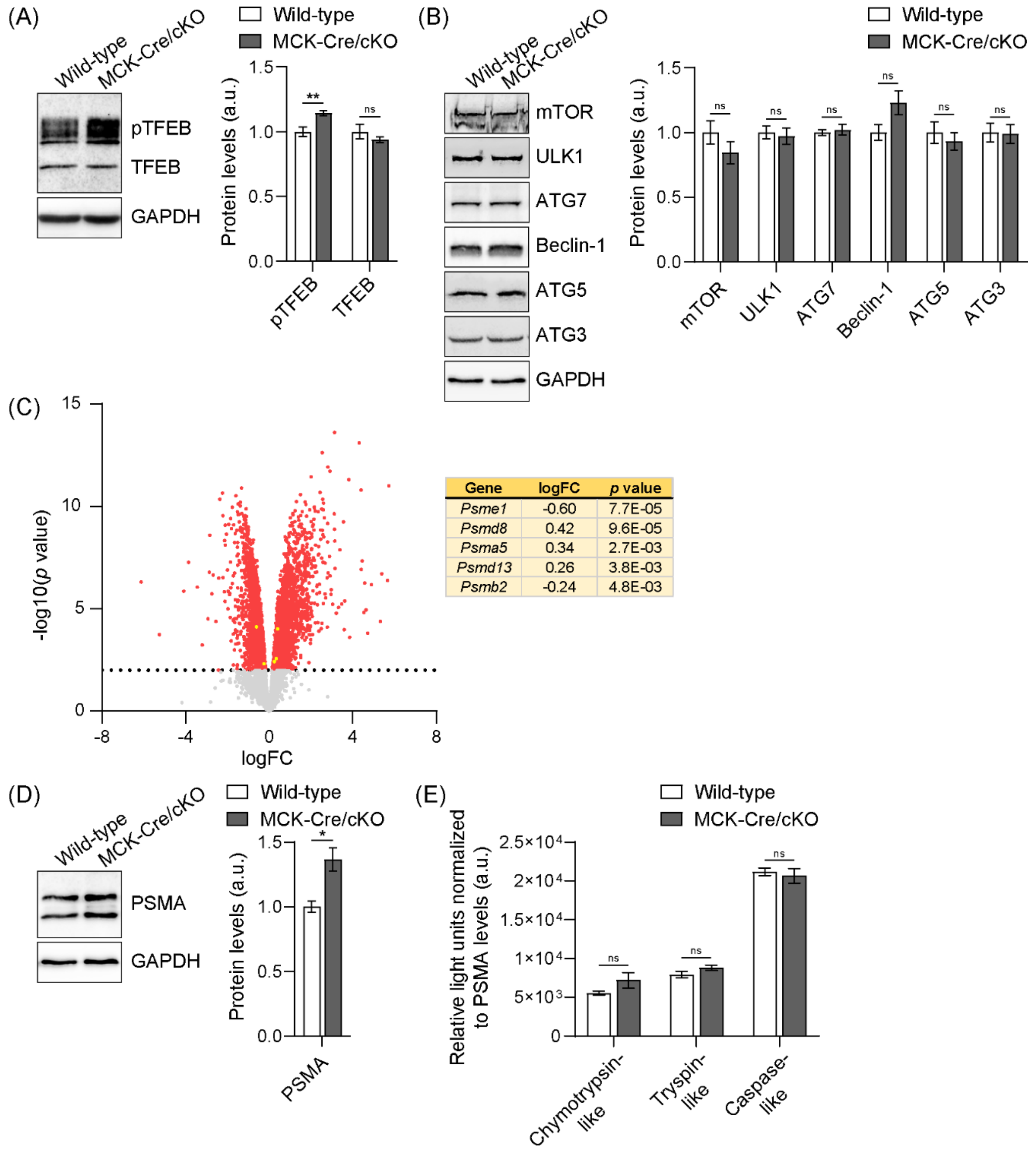


| Gene | logFC | *p* value |
|---|---|---|
| *Psme1* | -0.60 | 7.7E-05 |
| *Psmd8* | 0.42 | 9.6E-05 |
| *Psma5* | 0.34 | 2.7E-03 |
| *Psmd13* | 0.26 | 3.8E-03 |
| *Psmb2* | -0.24 | 4.8E-03 |

**Figure S1:** Expression of TFEB, proteins required for the induction of autophagy and proteasomal subunits, and proteasomal activities in 13-week-old plectin-deficient muscles. (A) Immunoblotting of muscle lysates from 13-week-old wild-type and MCK-Cre/cKO mice using antibodies to TFEB and GAPDH. Signal intensities of upper (corresponding to the phosphorylated versions, pTFEB) and lower (non-phosphorylated, TFEB) protein bands were densitometrically measured and normalized to the total protein content as analyzed by

Coomassie staining (not shown). Mean ± SEM; n = 8. (B) Immunoblotting of wild-type and MCK-Cre/cKO muscle lysates using antibodies mTOR, ULK1, ATG7, Beclin-1, ATG5, ATG3, and GAPDH. Signal intensities of protein bands were densitometrically measured and normalized to the total protein content as analyzed by Coomassie staining (not shown). Mean ± SEM; n = 8-10. (C) RNA-Seq analysis of mouse soleus muscle (as also shown in Figure 3A). Volcano plot illustrates differentially expressed genes in MCK-Cre/cKO compared to wild-type samples. All significantly up- and downregulated genes are highlighted in red; the dotted line represents the cut-off with $P = 0.01$. Significantly up- and downregulated genes from the KEGG pathway "mmu03050 Proteasome" are highlighted in yellow and listed on the right. logFC, log fold change; n = 5 animals per genotype. (D) Immunoblotting of wild-type and MCK-Cre/cKO muscle lysates, derived from 13-week-old animals, using antibodies to 20S α1, 2, 3, 5, 6, and 7 proteasomal subunits (PSMA), and GAPDH. Signal intensities of protein bands were densitometrically measured and normalized to the total protein content as analyzed by Coomassie staining (not shown). Mean ± SEM; n = 8. (E) Chymotrypsin-, trypsin-, and caspase-like proteasomal activities as assessed in Figure 3D were normalized to the proteasomal protein content as analyzed by immunoblotting (not shown). Mean ± SEM; samples were measured as triplicates, n = 3 animals each.

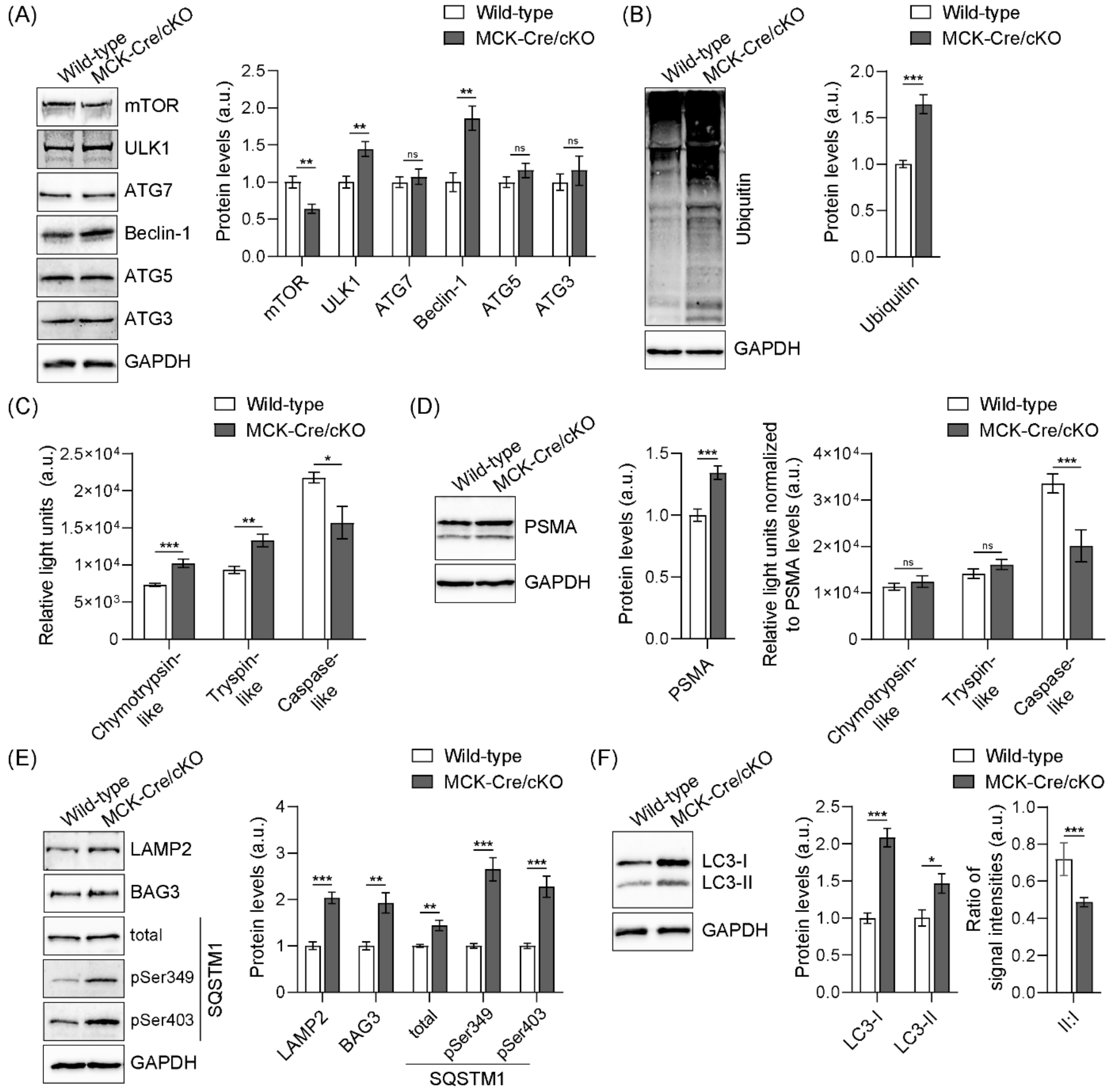


**Figure S2:** Evaluation of protein quality control mechanisms in muscles from aged mice. (A) Immunoblotting of muscle lysates from 40-week-old wild-type and MCK-Cre/cKO animals using antibodies mTOR, ULK1, ATG7, Beclin-1, ATG5, ATG3, and GAPDH. Signal intensities of protein bands were densitometrically measured and normalized to the total protein content as analyzed by Coomassie staining (not shown). Mean ± SEM; n = 6-8. (B) Immunoblotting of muscle lysates from 40-week-old wild-type and MCK-Cre/cKO animals using antibodies to ubiquitin and GAPDH. Signal intensities of immunoblots were densitometrically measured and normalized to the total protein content as analyzed by Coomassie staining (not shown). Mean ± SEM; n = 8. (C) Chymotrypsin-, trypsin-, and caspase-like proteasomal activities were measured in wild-type and MCK-Cre/cKO muscle lysates derived from 40-week-old mice. Mean ± SEM; samples were measured as triplicates, n = 3 animals each. (D) Immunoblotting of wild-type and MCK-Cre/cKO muscle lysates, derived from 40-week-old animals, using antibodies to 20S α1, 2, 3, 5, 6, and 7 proteasomal subunits (PSMA), and GAPDH. Signal intensities of protein bands were densitometrically measured and normalized to the total protein content as analyzed by Coomassie staining (not shown). Mean ± SEM; n = 8. Chymotrypsin-, trypsin-, and caspase-like proteasomal activities as assessed in (C) were normalized to the proteasomal protein content as analyzed by immunoblotting (not shown). Mean ± SEM; samples were measured as triplicates, n = 3 animals each. (D) Immunoblotting of muscle lysates obtained from 40-week-old wild-type and MCK-Cre/cKO animals using antibodies to LAMP2, BAG3, total and phosphorylated forms of SQSTM1, and GAPDH. Signal intensities of protein bands were densitometrically measured and normalized to the total protein content as analyzed by Coomassie staining (not shown). Mean ± SEM; n = 7-8. (E) Immunoblotting of muscle lysates obtained from 40-week-old wild-type and MCK-Cre/cKO animals using antibodies to LC3 and GAPDH. Signal intensities of upper (non-lipidated, LC3-I) and lower (lipidated, LC3-II) protein bands were densitometrically measured and normalized to the total protein content (as analyzed by Coomassie staining, not shown). From these values, the LC3-II to LC3-I ratios were calculated. Mean ± SEM; n = 8. For (A-F): *$P < 0.05$, **$P < 0.01$, ***$P < 0.001$ (two-tailed, unpaired *t*-test with Welch's correction); ns, not significant.

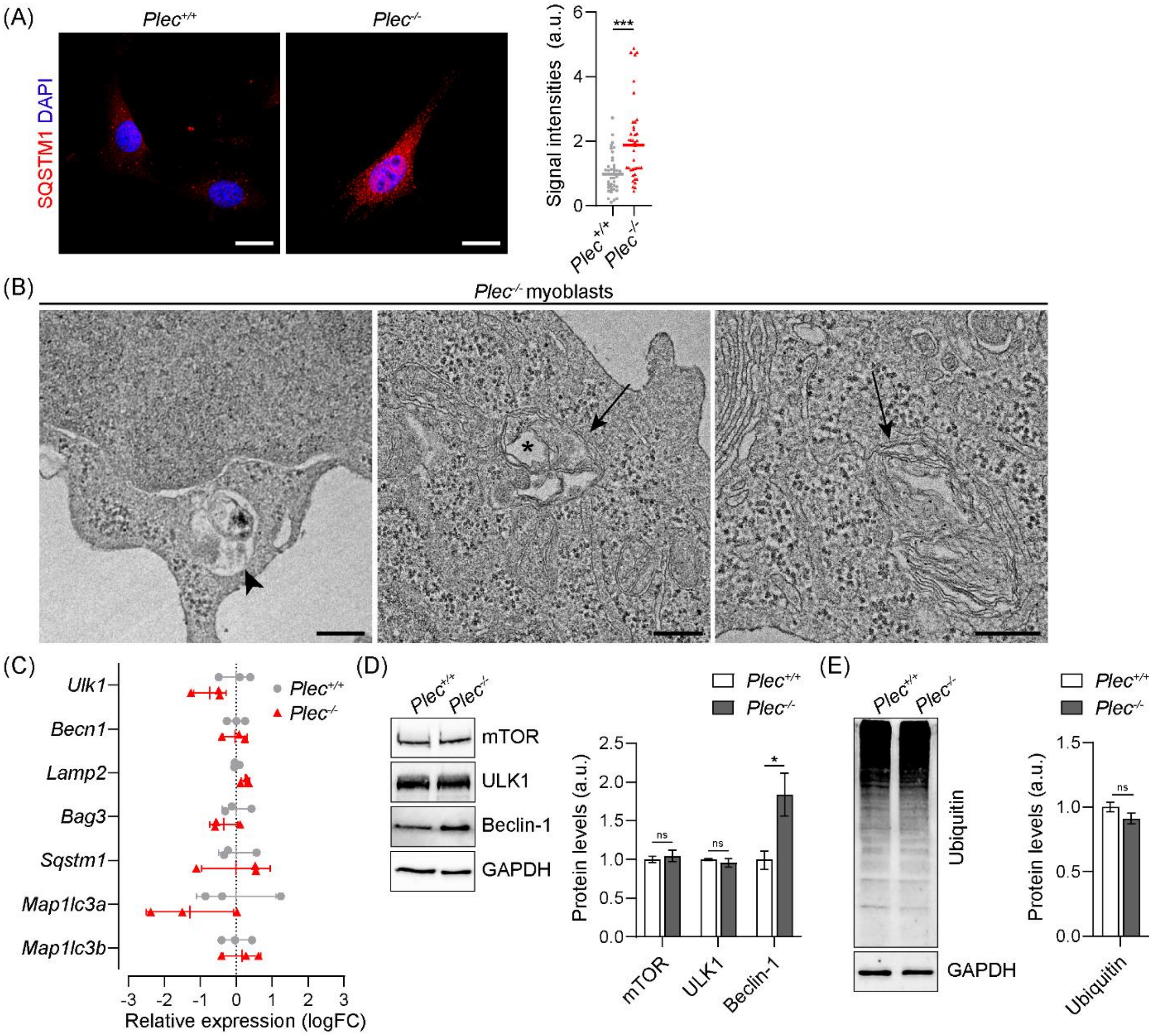


**Figure S3:** Accumulation of SQSTM1 and degradative vacuoles in plectin-deficient myoblasts, but unaltered autophagy regulation. (A) Immunostaining of immortalized (*p53*$^{-/-}$) murine plectin-positive (*Plec*$^{+/+}$) and plectin-deficient (*Plec*$^{-/-}$) myoblasts using antibodies SQSMT1. Nuclei were visualized with DAPI. Scale bars: 20 µm. SQSTM1 signal intensities in *Plec*$^{+/+}$ and *Plec*$^{-/-}$ myoblasts were calculated by normalizing the raw integrated densities (RawIntDens) to cell areas. Each dot represents a single cell, the line represents the median (*Plec*$^{+/+}$, n = 41 cells; *Plec*$^{-/-}$, n= 40 cells); ***$P < 0.001$ (two-tailed Mann-Whitney test). (B) Representative electron micrographs of *Plec*$^{-/-}$ myoblasts. Note the occurrence of degradative vacuoles (arrowhead) and membrane whirls (arrows), partially presenting with large vacuoles (asterisk). Scale bars: 250 nm. (C) Real-time quantitative PCR (RT-qPCR) analyses of ULK1 (*Ulk1)*, Beclin-1 (*Becn1*), LAMP2 (*Lamp2*), BAG3 (*Bag3*), SQSTM1 (*Sqstm1*), LC3A (*Map1lc3a*), and LC3B (*Map1lc3b*) mRNA expression. Relative gene expression values are depicted as logFC and were normalized to *Tbp* and *Hprt.* Samples were measured as triplicates, n = 3 experiments. (D) Immunoblotting of *Plec*$^{+/+}$ and *Plec*$^{-/-}$ myoblast cell lysates using antibodies to mTOR, ULK1, Beclin-1, and GAPDH. Signal intensities of protein bands were densitometrically measured and normalized to the total protein content as analyzed by Coomassie staining (not shown). Mean ± SEM; n = 8. (E) Immunoblotting of *Plec*$^{+/+}$ and *Plec*$^{-/-}$ myoblast cell lysates using antibodies to ubiquitin and GAPDH. Signal intensities of protein bandswere densitometrically measured and normalized to the total protein content as analyzed by Coomassie staining (not shown). Mean ± SEM; n = 8. For (D and E): *$P < 0.05$ (two-tailed, unpaired *t*-test with Welch's correction); ns, not significant.

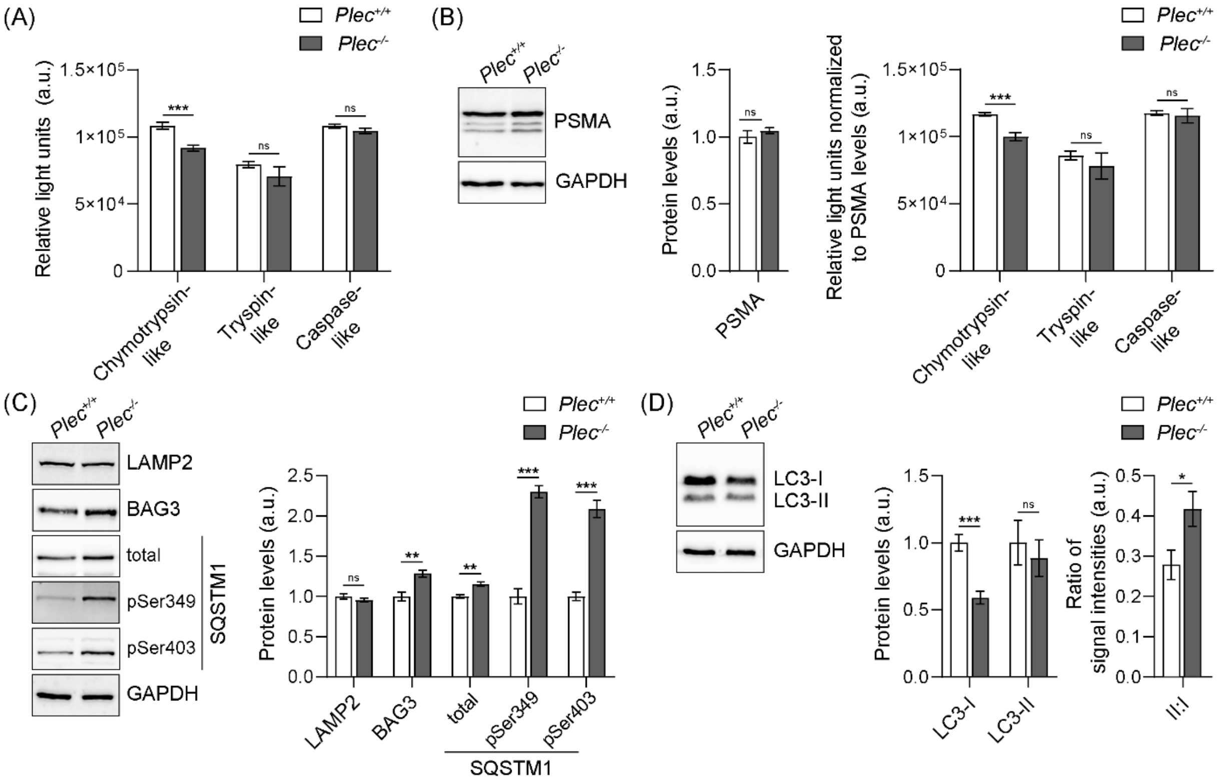


**Figure S4:** Evaluation of proteasomal activities and autophagic marker proteins in plectin-deficient myoblasts. (A) Chymotrypsin-, trypsin-, and caspase-like proteasomal activities were measured in *Plec*$^{+/+}$ and *Plec*$^{-/-}$ myoblast cell lysates. Mean ± SEM; samples were measured as triplicates, n = 2 experiments. (B) Immunoblotting of *Plec*$^{+/+}$ and *Plec*$^{-/-}$ myoblast cell lysates using antibodies to 20S α1, 2, 3, 5, 6, and 7 proteasomal subunits (PSMA), and GAPDH. Signal intensities of protein bands were densitometrically measured and normalized to the total protein content as analyzed by Coomassie staining (not shown). Mean ± SEM; n = 8. Chymotrypsin-, trypsin-, and caspase-like proteasomal activities as assessed in (A) were normalized to the proteasomal protein content as analyzed by immunoblotting (not shown). Mean ± SEM; samples were measured as triplicates, n = 3 animals each. (C) Immunoblotting of *Plec*$^{+/+}$ and *Plec*$^{-/-}$ myoblast cell lysates using antibodies to LAMP2, BAG3, total and phosphorylated forms of SQSTM1, and GAPDH. Signal intensities of protein bands were densitometrically measured and normalized to the total protein content as analyzed by Coomassie staining (not shown). Mean ± SEM; n = 7-8. (D) Immunoblotting of *Plec*$^{+/+}$ and *Plec*$^{-/-}$ myoblast cell lysates using antibodies to LC3 and GAPDH. Signal intensities of upper (non-lipidated, LC3-I) and lower (lipidated, LC3-II) protein bands were densitometrically measured and normalized to the total protein content (as analyzed by Coomassie staining, not shown). From these values, the LC3-II to LC3-I ratios were calculated. Mean ± SEM; n = 8. For (A-D): $*P < 0.05$, $**P < 0.01$, $***P < 0.001$ (two-tailed, unpaired *t*-test with Welch's correction); ns, not significant.

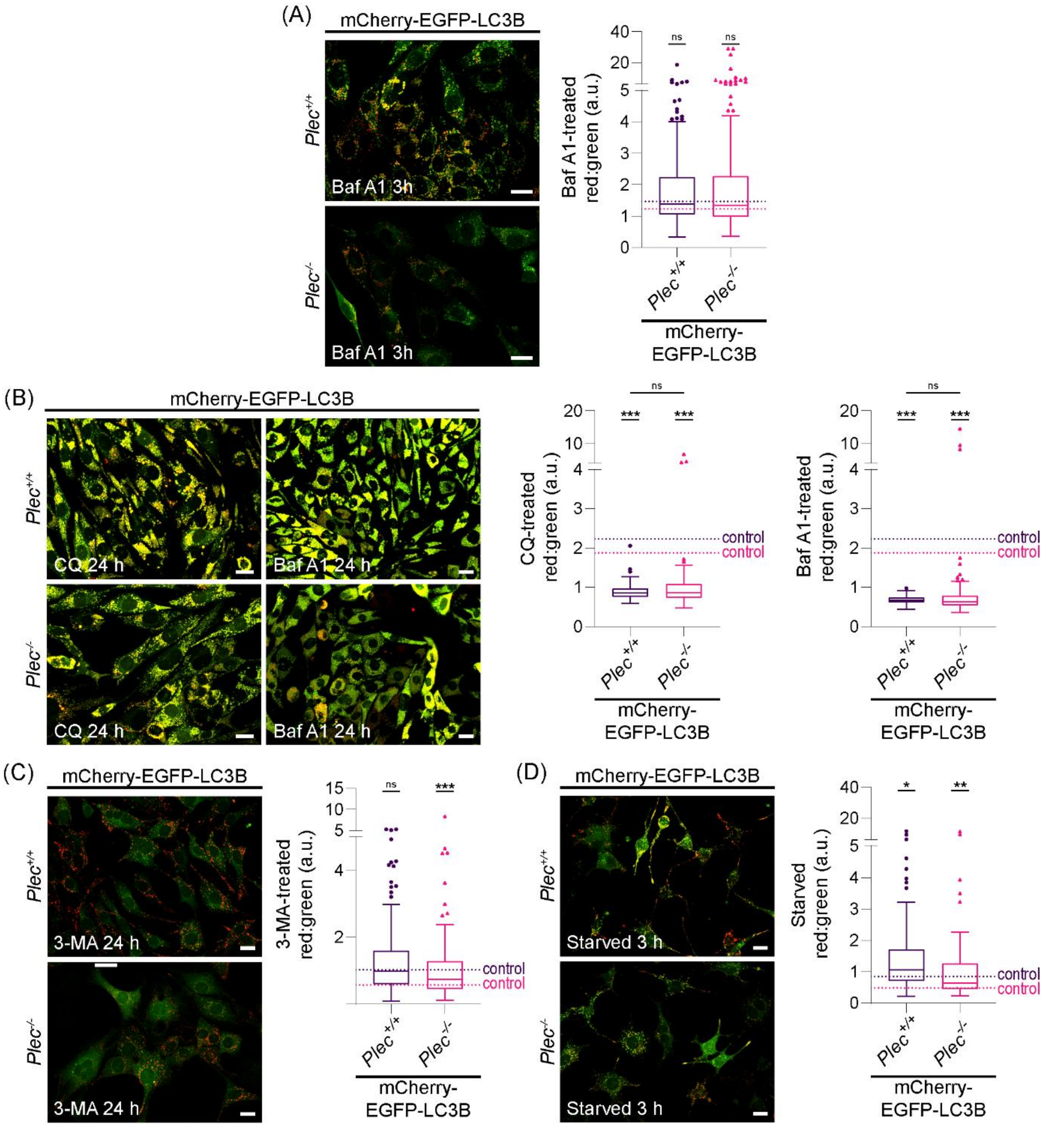


**Figure S5:** Impaired autophagic flux in *Plec*$^{-/-}$ myoblast cell lines. (A) mCherry-EGFP-LC3B-expressing *Plec*$^{+/+}$ and *Plec*$^{-/-}$ myoblasts were treated with 200 nM bafilomycin A1 (Baf A1) for 3 h. Scale bars: 20 µm. Red:green signal ratios of Baf A1-treated mCherry-EGFP-LC3B-expressing *Plec*$^{+/+}$ and *Plec*$^{-/-}$ myoblasts: dotted lines represent the median values of the respective cells at control conditions. Box plots show the median and Tukey whiskers (*Plec*$^{+/+}$, n = 202/263 [control/Baf A1]; *Plec*$^{-/-}$, n = 142/202 [control/Baf A1] cells). (B) mCherry-EGFP-LC3B-expressing *Plec*$^{+/+}$ and *Plec*$^{-/-}$ myoblasts were treated with 50 µM chloroquine (CQ) or 200 nM Baf A1 for 24 h. Note the massive swelling of vesicles in both *Plec*$^{+/+}$ and *Plec*$^{-/-}$ cells as well as the increased bright yellow signals compared to 3 h CQ-treated cells shown in Figure

5B. Scale bars: 20 μm. Red:green signal ratios of 24 h CQ and Baf A1-treated mCherry-EGFP-LC3B-expressing *Plec*$^{+/+}$ and *Plec*$^{-/-}$ myoblasts: dotted lines represent the median values of the respective cells at control conditions. Box plots show the median and Tukey whiskers (*Plec*$^{+/+}$, n = 113/149/177 [control/CQ/Baf A1]; *Plec*$^{-/-}$, n = 112/131/129 [control/CQ/Baf A1] cells). (C) mCherry-EGFP-LC3B-expressing *Plec*$^{+/+}$ and *Plec*$^{-/-}$ myoblasts were treated with 9 mM 3-methyladenine (3-MA) for 24 h. Scale bars: 20 μm. Red:green signal ratios of 24 h 3-MA-treated mCherry-EGFP-LC3B-expressing *Plec*$^{+/+}$ and *Plec*$^{-/-}$ myoblasts: dotted lines represent the median values of the respective cells at control conditions. Box plots show the median and Tukey whiskers (*Plec*$^{+/+}$, n = 185/226 [control/3-MA]; *Plec*$^{-/-}$, n = 176/196 [control/3-MA] cells). (D) mCherry-EGFP-LC3B-expressing *Plec*$^{+/+}$ and *Plec*$^{-/-}$ myoblasts were starved for 3 h. Scale bars: 20 μm. Red:green signal ratios of 3h-starved mCherry-EGFP-LC3B-expressing *Plec*$^{+/+}$ and *Plec*$^{-/-}$ myoblasts: dotted lines represent the median values of the respective cells at control conditions. Box plots show the median and Tukey whiskers (*Plec*$^{+/+}$, n = 103/99 [control/starved]; *Plec*$^{-/-}$, n = 89/92 [control/starved] cells). For (A-D): **P* < 0.05, ***P* < 0.01, ****P* < 0.001 (two-way ANOVA of the ranked dataset with Tukey's post-hoc correction for multiple comparisons); ns, not significant.
