## Supplemental Information_RNA Seq for "Impaired autophagic flux in skeletal muscle of plectin-related epidermolysis bullosa simplex with muscular dystrophy"

| Symbol | Genes | Entrez | logFC | AveExpr | t | P.Value | adj.P.Val |
|---|---|---|---|---|---|---|---|
| Myl4 | ENSMUSG00000061086 | 17896 | 3.14380928526113 | 3.85558507394462 | 33.602856013269 | 2.4700241786113 | 2.69998342964004e-10 |
| Myh3 | ENSMUSG00000020908 | 17883 | 4.31970427667252 | 6.96301557692128 | 30.7606214385414 | 7.9620202567198 | 4.35164217131024e-10 |
| Eda2r | ENSMUSG00000034457 | 245527 | 2.55016703112115 | 4.10786430930429 | 28.3014107996155 | 2.3945015688827 | 8.72476554981903e-10 |
| Npnt | ENSMUSG00000040998 | 114249 | 2.82227744290143 | 4.44086357549252 | 25.0083651282272 | 1.2212595426959 | 3.33739701530247e-09 |
| Runx1 | ENSMUSG00000022952 | 12394 | 2.91700619337499 | 4.17562472365354 | 24.1849384492368 | 1.8956771891689 | 4.14432947096111e-09 |
| Tubb6 | ENSMUSG00000001473 | 67951 | 2.02031957488405 | 4.74302770519814 | 23.0101478414847 | 3.6414202551981 | 6.63406080159519e-09 |
| Serpinb1a | ENSMUSG00000044734 | 66222 | 3.82600004449594 | 3.03889820556911 | 22.4734031081641 | 4.9594083686966 | 7.74447041117474e-09 |
| Mybph | ENSMUSG00000042451 | 53311 | 5.74256092565916 | 3.3758333256557 | 21.3363795981219 | 9.7721701862672 | 1.33524490382609e-08 |
| Hsdl2 | ENSMUSG00000028383 | 72479 | -1.31671319473881 | 5.69390307688924 | -20.9104345065445 | 1.2711086898068 | 1.54383212091984e-08 |
| Myh8 | ENSMUSG00000055775 | 17885 | 4.41250513566256 | 4.32724624998562 | 20.5517717902199 | 1.5924052058651 | 1.74065813053123e-08 |
| Peg3 | ENSMUSG00000002265 | 18616 | 2.45507714074309 | 5.4196247155893 | 20.129453399437 | 2.0863141008434 | 2.07322722148361e-08 |
| Ephx2 | ENSMUSG00000022040 | 13850 | -2.22388229713764 | 4.87500124169922 | -19.9912405317237 | 2.2818155322252 | 2.07854379856289e-08 |
| Actc1 | ENSMUSG00000068614 | 11464 | 1.50301955406381 | 5.0327876580786 | 19.726075681948 | 2.7140956925417 | 2.28213692424416e-08 |
| Grb14 | ENSMUSG00000026888 | 50915 | -1.57580236681012 | 4.4611865478433 | -19.5447586533335 | 3.0598060980887 | 2.38905288987199e-08 |
| Col8a1 | ENSMUSG00000068196 | 12837 | 2.57246702549679 | 5.17127191471176 | 18.9713623638916 | 4.5017745716378 | 3.13879368752184e-08 |
| Nos1 | ENSMUSG00000029361 | 18125 | -2.34949157721296 | 4.17905472512117 | -18.941578430881 | 4.5943371146600 | 3.13879368752184e-08 |
| Eef1a1 | ENSMUSG00000037742 | 13627 | 1.31737317099794 | 8.68735884646565 | 18.6025788084983 | 5.8043363978400 | 3.65352507427976e-08 |
| Maged2 | ENSMUSG00000025268 | 80884 | 1.724291949755 | 4.2726621531795 | 18.5071367367442 | 6.2036608843384 | 3.65352507427976e-08 |
| Adk | ENSMUSG00000039197 | 11534 | -1.37628618039337 | 4.7467218783011 | -18.4736938858902 | 6.3504689791707 | 3.65352507427976e-08 |
| Ahnak2 | ENSMUSG00000072812 | 100041194 | 2.61607729813688 | 6.233394536756 | 18.3513097600051 | 6.9202908014082 | 3.78228493750965e-08 |
| Eci1 | ENSMUSG00000024132 | 13177 | -1.48127996943267 | 5.63163032587351 | -18.2192950265086 | 7.5969544429622 | 3.95439566742951e-08 |
| Pde10a | ENSMUSG00000023868 | 23984 | 1.76765419774403 | 3.42049543781369 | 18.1326389635555 | 8.0794628367162 | 4.0143912849157e-08 |
| Gpcpd1 | ENSMUSG00000027346 | 74182 | -1.37552947882263 | 5.79983712788566 | -17.8871950169918 | 9.6333377165331 | 4.57834845997494e-08 |
| Itgam | ENSMUSG00000030786 | 16409 | 2.17618657864017 | 2.76350942913272 | 17.7303941768997 | 1.0791710403836 | 4.82086101826672e-08 |
| Lpl | ENSMUSG00000015568 | 16956 | -1.75100612704071 | 9.56607422023671 | -17.7009202740614 | 1.1025663293081 | 4.82086101826672e-08 |
| Lyz2 | ENSMUSG00000069516 | 17105 | 2.17248850671042 | 5.77875356779725 | 17.5886606887139 | 1.1967763650594 | 5.0315240178711e-08 |
| Ptp4a3 | ENSMUSG00000059895 | 19245 | -1.09481799534287 | 7.4621415771973 | -17.4546772656718 | 1.3206491127748 | 5.2426994602709e-08 |
| Myl6b | ENSMUSG00000039824 | 216459 | 2.52075637383031 | 6.07082145619645 | 17.4320125615857 | 1.3429291454357 | 5.2426994602709e-08 |
| Igf2 | ENSMUSG00000048583 | 16002 | 1.94632332338385 | 5.00449836153958 | 17.3210474244912 | 1.4579794105819 | 5.39499195385771e-08 |
| Aif1l | ENSMUSG00000001864 | 108897 | 2.59993354960718 | 2.68059887642164 | 17.3002912586207 | 1.4806491502674 | 5.39499195385771e-08 |
| Pam | ENSMUSG00000026335 | 18484 | 1.25250052056752 | 6.60531381259457 | 17.1406834042777 | 1.6681197436800 | 5.72635088700503e-08 |
| Ccr2 | ENSMUSG00000049103 | 12772 | 2.21799466483452 | 2.95335919790121 | 17.1236024655092 | 1.6896405055230 | 5.72635088700503e-08 |
| Tnnt2 | ENSMUSG00000026414 | 21956 | 2.35451662667307 | 4.03276915125394 | 17.034059449837 | 1.8074331086196 | 5.72635088700503e-08 |
| Myo5a | ENSMUSG00000034593 | 17918 | 1.59890526309707 | 3.74429854262315 | 17.0329834263771 | 1.8089010009260 | 5.72635088700503e-08 |
| Rhou | ENSMUSG00000039960 | 69581 | -1.50478214108305 | 4.19656651947893 | -17.0150734755779 | 1.8335219197253 | 5.72635088700503e-08 |
| Perm1 | ENSMUSG00000078486 | 74183 | -1.3023448495415 | 7.96545070490964 | -16.7519284508293 | 2.2398079654793 | 6.65974934941977e-08 |
| Camk2d | ENSMUSG00000053819 | 108058 | 1.13663741728913 | 5.77688051556802 | 16.7407171043923 | 2.2591336952824 | 6.65974934941977e-08 |
| Tnc | ENSMUSG00000028364 | 21923 | 3.12718159509059 | 4.43937098498253 | 16.6936807978575 | 2.3421818949381 | 6.65974934941977e-08 |
| Acadm | ENSMUSG00000062908 | 11364 | -1.40026349340796 | 7.73994806873533 | -16.6749884260679 | 2.3760884148510 | 6.65974934941977e-08 |
| Retsat | ENSMUSG00000056666 | 67442 | -1.46542429312298 | 3.994941972249 | -16.5745132903558 | 2.5675825128188 | 6.91645467580883e-08 |
| Abhd2 | ENSMUSG00000039202 | 54608 | 1.15139762801799 | 4.72934441326779 | 16.5611738001624 | 2.5942241488259 | 6.91645467580883e-08 |
| Plekhh2 | ENSMUSG00000040852 | 213556 | 1.52579697654766 | 3.73453076685972 | 16.4653067169758 | 2.7946245859939 | 7.1287980634988e-08 |
| Padi2 | ENSMUSG00000028927 | 18600 | -2.21248538329301 | 5.15936373311557 | -16.4500652788191 | 2.8279846194219 | 7.1287980634988e-08 |
| Cd36 | ENSMUSG00000002944 | 12491 | -1.14468790772742 | 7.97946805287774 | -16.431356144495 | 2.8695189350832 | 7.1287980634988e-08 |
| Lpin1 | ENSMUSG00000020593 | 14245 | -1.2110338505657 | 6.74605912925526 | -16.3761122907659 | 2.9960195780981 | 7.14562981800884e-08 |
| Pcdh7 | ENSMUSG00000029108 | 54216 | 1.79938952624937 | 3.55491096968945 | 16.3714206041023 | 3.0070347784137 | 7.14562981800884e-08 |
| Rilpl1 | ENSMUSG00000029392 | 75695 | -1.08373268092938 | 6.18807056409412 | -16.3419736949439 | 3.0771687793927 | 7.15670892075374e-08 |
| Plekho1 | ENSMUSG00000015745 | 67220 | 1.40970409759378 | 4.46586833738978 | 16.25928191825 | 3.2836526116308 | 7.47783472869527e-08 |
| S100a4 | ENSMUSG00000001020 | 20198 | 2.64923149566636 | 2.05220877504383 | 16.1517500968934 | 3.5746359475683 | 7.95070621649665e-08 |
| Vwa5a | ENSMUSG00000023186 | 67776 | 1.03253092366179 | 4.85041587081838 | 16.1300047419168 | 3.6367698364727 | 7.95070621649665e-08 |
| Nqo1 | ENSMUSG00000003849 | 18104 | -1.83874881994114 | 2.9168923619885 | -16.0462026850671 | 3.8872597567111 | 8.33169341188414e-08 |
| Col3a1 | ENSMUSG00000026043 | 12825 | 2.15917087567223 | 9.45520221865379 | 15.9337893930917 | 4.2526901718587 | 8.91593074547111e-08 |
| Adssl1 | ENSMUSG00000011148 | 11565 | -1.17965990803321 | 7.75992312058011 | -15.9133576900926 | 4.3229743802942 | 8.91593074547111e-08 |
| Pptc7 | ENSMUSG00000038582 | 320717 | -1.11125433647406 | 6.34080871741296 | -15.8458366237792 | 4.5642323794228 | 9.17372172790252e-08 |
| Cox7a1 | ENSMUSG00000074218 | 12865 | -1.14010989192584 | 6.92053740369767 | -15.8318965106597 | 4.6158146101421 | 9.17372172790252e-08 |
| Ankrd10 | ENSMUSG00000031508 | 102334 | 1.42375525480253 | 4.19309923465921 | 15.7690955313318 | 4.8560608696253 | 9.4788573867633e-08 |
| Cul7 | ENSMUSG00000038545 | 66515 | 1.23686908233542 | 3.84351080234009 | 15.6168160724424 | 5.4960112254792 | 1.04306719689529e-07 |
| Apoe | ENSMUSG00000002985 | 11816 | 2.02185169137151 | 6.3353067652626 | 15.6079497529085 | 5.5359544247293 | 1.04306719689529e-07 |
| Plec | ENSMUSG00000022565 | 18810 | -1.42000109219697 | 10.5776817497818 | -15.5798438511557 | 5.6646407852937 | 1.04306719689529e-07 |
| Lgals3 | ENSMUSG00000050335 | 16854 | 3.56903560499648 | 2.47663844703039 | 15.56681651856 | 5.7253711292395 | 1.04306719689529e-07 |
| Anxa4 | ENSMUSG00000029994 | 11746 | 1.10157878754531 | 4.4706794101676 | 15.4693350523955 | 6.2025469559395 | 1.09844556238309e-07 |
| Ppfibp1 | ENSMUSG00000016487 | 67533 | 1.14584364957775 | 6.70304279325261 | 15.4639109734814 | 6.2303197207713 | 1.09844556238309e-07 |
| Etfdh | ENSMUSG00000027809 | 66841 | -1.09764920565449 | 6.48342477207109 | -15.4345596400711 | 6.3829344904599 | 1.10748979230504e-07 |
| Cilp | ENSMUSG00000042254 | 214425 | 2.81327936592838 | 7.61259331315795 | 15.3658439067405 | 6.7561068976962 | 1.15392194529246e-07 |
| Chrnb1 | ENSMUSG00000041189 | 11443 | 1.51136097251443 | 4.03276851120342 | 15.3031530315388 | 7.1169818406176 | 1.19685736153526e-07 |
| Uck2 | ENSMUSG00000026558 | 80914 | 1.56038366004355 | 4.47556530432855 | 15.2372103508999 | 7.5189541946561 | 1.23265073385038e-07 |
| Adhfe1 | ENSMUSG00000025911 | 76187 | -1.92315693465454 | 2.98448870518224 | -15.2314260076628 | 7.5553562499291 | 1.23265073385038e-07 |
| Klf5 | ENSMUSG00000005148 | 12224 | 2.11613483118708 | 3.28486271300096 | 15.1923394851109 | 7.8063353928893 | 1.25486841440696e-07 |
| Tigar | ENSMUSG00000038028 | 319801 | -1.21212784471693 | 4.26560266420311 | -15.1726876397003 | 7.9358810670468 | 1.25720457889694e-07 |
| Alpl | ENSMUSG00000028766 | 11647 | -1.69142900053761 | 3.068896568258 | -15.1487302463276 | 8.0969301556208 | 1.2643934790156e-07 |
| Magix | ENSMUSG00000031147 | 54634 | -2.09977199396153 | 3.07357570826986 | -15.1253959787124 | 8.2571581414617 | 1.27125345977914e-07 |
| Cyb5r3 | ENSMUSG00000018042 | 109754 | 1.00998135152429 | 5.76119578950028 | 15.0598500336631 | 8.7256724172978 | 1.32472673879838e-07 |
| Lrrc14b | ENSMUSG00000021579 | 432779 | -1.08707414645654 | 5.42297596251391 | -15.0106273240635 | 9.0961956052383 | 1.35069851927445e-07 |
| Cdh13 | ENSMUSG00000031841 | 12554 | 1.12264678765469 | 6.99352664356931 | 15.0044496761027 | 9.1438743414426 | 1.35069851927445e-07 |
| Aspn | ENSMUSG00000021388 | 66695 | 2.08507198392991 | 5.53044778275512 | 14.9569822211633 | 9.5192741056596 | 1.36982455264224e-07 |
| Tmsb10 | ENSMUSG00000079523 | 19240 | 1.43103104748107 | 4.17907631282214 | 14.9563995163312 | 9.5239837161110 | 1.36982455264224e-07 |
| Wnk2 | ENSMUSG00000037989 | 75607 | -0.991146701078056 | 5.88612447920891 | -14.8869144245275 | 1.0103872455556 | 1.43435623132055e-07 |
| Ppif | ENSMUSG00000021868 | 105675 | -1.35749946734347 | 3.83121207952593 | -14.8304539863851 | 1.0602895137783 | 1.48590059937318e-07 |
| Csf1r | ENSMUSG00000024621 | 12978 | 1.55519320676651 | 4.41508962861272 | 14.7879871841133 | 1.0995599705779 | 1.52142911878326e-07 |
| Sar1b | ENSMUSG00000020386 | 66397 | -1.11525516303409 | 4.97305877701639 | -14.6755829307676 | 1.2112141849501 | 1.65497278196129e-07 |
| Plin5 | ENSMUSG00000011305 | 66968 | -1.7936084012139 | 3.94156289751921 | -14.6510863528338 | 1.2371222094257 | 1.66950405817695e-07 |
| Acat1 | ENSMUSG00000032047 | 110446 | -1.24114252231677 | 7.19421599467252 | -14.6212241536551 | 1.2695098772567 | 1.69231859369433e-07 |
| Vash2 | ENSMUSG00000037568 | 226841 | 3.14222062016296 | 1.590615955793 | 14.5856051074737 | 1.3093324806368 | 1.7243751019086e-07 |
| Anpep | ENSMUSG00000039062 | 16790 | 1.70952837363024 | 3.98586105722263 | 14.5583835729195 | 1.3406673077498 | 1.72659978767157e-07 |
| Stab1 | ENSMUSG00000042286 | 192187 | 1.18338415095873 | 4.36333588667524 | 14.5567161560159 | 1.3426125876139 | 1.72659978767157e-07 |
| Fdft1 | ENSMUSG00000021273 | 14137 | -1.39422417306219 | 4.90954689992855 | -14.4980916906381 | 1.4129634730747 | 1.77942699600801e-07 |
| Sord | ENSMUSG00000027227 | 20322 | -0.981607331364449 | 5.18896017026297 | -14.473487502599 | 1.4436553656844 | 1.77942699600801e-07 |
| Dclk1 | ENSMUSG00000027797 | 13175 | 2.47079236461522 | 3.28299188978607 | 14.4719605588868 | 1.4455834826092 | 1.77942699600801e-07 |
| Col5a2 | ENSMUSG00000026042 | 12832 | 2.16079205111469 | 6.53343222082161 | 14.4694132572475 | 1.4488061718480 | 1.77942699600801e-07 |
| Myog | ENSMUSG00000026459 | 17928 | 2.5525521451002 | 3.00965966445878 | 14.4438437312563 | 1.4815844498542 | 1.78262849570929e-07 |
| Rxrg | ENSMUSG00000015843 | 20183 | -1.54493450152309 | 2.5174734683606 | -14.441961231299 | 1.4840288455726 | 1.78262849570929e-07 |
| Oxct1 | ENSMUSG00000022186 | 67041 | -1.03638449657289 | 8.35141461396818 | -14.3432059241025 | 1.6185138587706 | 1.90925149987813e-07 |
| Obsl1 | ENSMUSG00000026211 | 98733 | 1.2210840513524 | 4.8728104703374 | 14.3391040929197 | 1.6243746179550 | 1.90925149987813e-07 |
| Tbc1d4 | ENSMUSG00000033083 | 210789 | -1.34021842956365 | 5.14942252688682 | -14.321842586996 | 1.6492881604655 | 1.91791158319669e-07 |
| Hadh | ENSMUSG00000027984 | 15107 | -1.0173984477537 | 6.90036779389766 | -14.2571670544039 | 1.7463332797475 | 2.00938621904424e-07 |
| Ech1 | ENSMUSG00000053898 | 51798 | -1.28757696622558 | 6.51866313409946 | -14.2196301094039 | 1.8054450215981 | 2.0299905691191e-07 |
| Ncam1 | ENSMUSG00000039542 | 17967 | 1.59117623651249 | 4.7803242774506 | 14.2165088023011 | 1.8104558591701 | 2.0299905691191e-07 |
| Map1b | ENSMUSG00000052727 | 17755 | 1.23908437993315 | 6.58413618532667 | 14.1981183630045 | 1.8402829827816 | 2.0299905691191e-07 |
| Pi16 | ENSMUSG00000024011 | 74116 | 1.7408412313307 | 5.36496433266066 | 14.1863931891809 | 1.8595740145992 | 2.0299905691191e-07 |
| Decr1 | ENSMUSG00000028223 | 67460 | -1.00194272913653 | 6.05596547812056 | -14.1830251465219 | 1.8651552614579 | 2.0299905691191e-07 |
| Col23a1 | ENSMUSG00000063564 | 237759 | 1.93563590262924 | 2.87212702543865 | 14.1767115611354 | 1.8756659727475 | 2.0299905691191e-07 |
| Atp13a3 | ENSMUSG00000022533 | 224088 | 1.0299322858415 | 5.96893154450826 | 14.1400794943936 | 1.9379142755626 | 2.07679813197796e-07 |
| Fst | ENSMUSG00000021765 | 14313 | 1.83887534851636 | 3.29324301168351 | 14.1184146641832 | 1.9757662936283 | 2.09680595685937e-07 |
| Egfr | ENSMUSG00000020122 | 13649 | 0.956899830459991 | 4.68624584010618 | 14.0710218325705 | 2.0613555574659 | 2.16660361525583e-07 |
| Ccdc85c | ENSMUSG00000084883 | 668158 | -1.61975585222442 | 3.35653196520992 | -14.0148770028064 | 2.1679151209718 | 2.2569028749851e-07 |
| Gadd45a | ENSMUSG00000036390 | 13197 | 2.04178065343339 | 2.96193160698227 | 13.9972786332434 | 2.2025172245438 | 2.26807242586389e-07 |
| Igfbp4 | ENSMUSG00000017493 | 16010 | 1.37316291101819 | 5.75815959230457 | 13.9728989035194 | 2.2514325640887 | 2.26807242586389e-07 |
| Slc4a3 | ENSMUSG00000006576 | 20536 | 1.42153445715266 | 3.71379690896978 | 13.9716921335987 | 2.2538838156705 | 2.26807242586389e-07 |
| Cfh | ENSMUSG00000026365 | 12628 | 1.64685119908717 | 5.04518718387413 | 13.9678828182134 | 2.2616402380309 | 2.26807242586389e-07 |
| Ppp1r1a | ENSMUSG00000022490 | 58200 | -2.03090953683478 | 4.78747491661355 | -13.9473454649469 | 2.3039527973296 | 2.27264676930482e-07 |
| Hadhb | ENSMUSG00000059447 | 231086 | -1.16630971972136 | 7.52631203897225 | -13.9378199016927 | 2.3238648480432 | 2.27264676930482e-07 |
| Tuba1b | ENSMUSG00000023004 | 22143 | 1.15406263732447 | 4.88493016217671 | 13.928835548332 | 2.3428143514586 | 2.27264676930482e-07 |
| Aldh6a1 | ENSMUSG00000021238 | 104776 | -1.00148424454421 | 6.01442630064647 | -13.925747834306 | 2.3493649705557 | 2.27264676930482e-07 |
| Acaa2 | ENSMUSG00000036880 | 52538 | -1.16820864706696 | 7.02325174343032 | -13.9054050139296 | 2.3930153167492 | 2.29251535855811e-07 |
| Slc25a20 | ENSMUSG00000032602 | 57279 | -1.12155711822206 | 5.08791349549571 | -13.8967498187115 | 2.4118494761154 | 2.29251535855811e-07 |

| | | | | | | | |
|---|---|---|---|---|---|---|---|
| Ppfia4 | ENSMUSG00000026458 | 68507 | 2.58567013084752 | 1.5747803897641 | 13.873024528502 | 2.4642947882882 | 2.32217295954991e-07 |
| Mybpc2 | ENSMUSG00000038670 | 233199 | -2.82838050904653 | 5.58722272074515 | -13.8518824563335 | 2.5120584682744 | 2.34214779364964e-07 |
| Amd1 | ENSMUSG00000075232 | 11702 | -1.13654751919747 | 5.16873315890084 | -13.8447713826824 | 2.5283454363796 | 2.34214779364964e-07 |
| Cotl1 | ENSMUSG00000031827 | 72042 | 1.54786412153446 | 3.19666449813209 | 13.7860546019634 | 2.6672258815934 | 2.43660852631185e-07 |
| Phyh | ENSMUSG00000026664 | 16922 | -1.12249117977913 | 6.36060715594953 | -13.7829071508487 | 2.6748972935451 | 2.43660852631185e-07 |
| Tmem65 | ENSMUSG00000062373 | 74868 | -1.04860367850003 | 6.59338588430762 | -13.7223742147134 | 2.8271226380649 | 2.55398988071796e-07 |
| Rhoc | ENSMUSG00000002233 | 11853 | 0.951792497016932 | 4.33103102973901 | 13.662824781479 | 2.9859495182912 | 2.67536181839682e-07 |
| Ccdc80 | ENSMUSG00000022665 | 67896 | 1.29441588874967 | 6.01642115745245 | 13.6487416239306 | 3.0248899645488 | 2.68821725223442e-07 |
| Mpeg1 | ENSMUSG00000046805 | 17476 | 2.62787009169099 | 3.17044985178672 | 13.6268114302747 | 3.0866132384877 | 2.72094913789595e-07 |
| Igf2r | ENSMUSG00000023830 | 16004 | 0.772710022586526 | 7.37351714409454 | 13.6075300985458 | 3.1419946560653 | 2.73889434714147e-07 |
| H2-Aa | ENSMUSG00000036594 | 14960 | 2.0437881605287 | 4.34575955957088 | 13.6023401598071 | 3.1570824969337 | 2.73889434714147e-07 |
| Slc27a1 | ENSMUSG00000031808 | 26457 | -1.31254209853322 | 5.4799069228079 | -13.5753866725286 | 3.2366961672699 | 2.77483440851662e-07 |
| Tspo | ENSMUSG00000041736 | 12257 | 1.75858967721255 | 2.03924811652894 | 13.5711913253933 | 3.2492800685218 | 2.77483440851662e-07 |
| Oplah | ENSMUSG00000022562 | 75475 | -1.22342357866527 | 4.66632593916396 | -13.547951036531 | 3.3199435558006 | 2.81242765407081e-07 |
| Dbt | ENSMUSG00000000340 | 13171 | -1.0411941467071 | 4.22324655146925 | -13.5399148931391 | 3.3447588969829 | 2.81242765407081e-07 |
| Sulf2 | ENSMUSG00000006800 | 72043 | 1.45514453364742 | 4.58007523337592 | 13.4673715258305 | 3.5779617692675 | 2.97240848085455e-07 |
| Adamts2 | ENSMUSG00000036545 | 216725 | 1.4498827393895 | 4.57041313706694 | 13.4563655370934 | 3.6148357075973 | 2.97240848085455e-07 |
| Col5a1 | ENSMUSG00000026837 | 12831 | 1.836870510349 | 6.7317847907452 | 13.4482170675249 | 3.6423977163360 | 2.97240848085455e-07 |
| Fzd10 | ENSMUSG00000081683 | 93897 | 1.97593565317392 | 2.94507553722517 | 13.4474061789267 | 3.6451527988389 | 2.97240848085455e-07 |
| Hadha | ENSMUSG00000025745 | 97212 | -1.03443569379479 | 8.42212473728879 | -13.4398354625945 | 3.6709829376577 | 2.97240848085455e-07 |
| Lmna | ENSMUSG00000028063 | 16905 | 0.847999423950013 | 7.40299694488809 | 13.3863119153415 | 3.8592763181932 | 3.10189334074781e-07 |
| Tmem176b | ENSMUSG00000029810 | 65963 | 1.37233494140846 | 3.18163475653792 | 13.3552286274339 | 3.9733454182999 | 3.17026560346252e-07 |
| Armh4 | ENSMUSG00000036242 | 67419 | 2.47789013777267 | 3.07286599836744 | 13.3427694153114 | 4.0200757040516 | 3.184307791376e-07 |
| Asb2 | ENSMUSG00000021200 | 65256 | -1.06213477752208 | 7.53773801600988 | -13.3007696575647 | 4.1819822869960 | 3.28872290497512e-07 |
| Optn | ENSMUSG00000026672 | 71648 | -0.915550723083882 | 6.48952882829509 | -13.2856856824312 | 4.2418202935657 | 3.2909756255344e-07 |
| C1qa | ENSMUSG00000036887 | 12259 | 1.75181923796993 | 3.33012874269254 | 13.2848753960213 | 4.2450604995000 | 3.2909756255344e-07 |
| Ndufa3 | ENSMUSG00000035674 | 66091 | -1.01346623635987 | 5.76494234022876 | -13.2671657274051 | 4.3165437803548 | 3.30884199342556e-07 |
| Dnajc27 | ENSMUSG00000020657 | 217378 | -1.0727396479381 | 5.26029772575774 | -13.2641982705214 | 4.3286470136296 | 3.30884199342556e-07 |
| Tex2 | ENSMUSG00000040548 | 21763 | -0.752707488310888 | 6.04143666975439 | -13.252487271465 | 4.3767675337965 | 3.3223920772174e-07 |
| Nin | ENSMUSG00000021068 | 18080 | 0.807209554264495 | 6.59491393501585 | 13.2017509625456 | 4.5919495765507 | 3.46081949162864e-07 |
| Ccdc141 | ENSMUSG00000044033 | 545428 | 1.36872625881965 | 3.25647647824851 | 13.19476653575 | 4.6224466725622 | 3.46081949162864e-07 |
| Slc25a34 | ENSMUSG00000040740 | 384071 | -2.11597765039967 | 3.23864532435632 | -13.1708689248467 | 4.7284432785740 | 3.51609615497232e-07 |
| Postn | ENSMUSG00000027750 | 50706 | 3.1500748941232 | 6.93188754736433 | 13.1271861088004 | 4.9289726585338 | 3.64044595475898e-07 |
| Cacnb1 | ENSMUSG00000020882 | 12295 | 0.863167020577521 | 6.71871625264922 | 13.0744716125609 | 5.1831460453543 | 3.78989553752527e-07 |
| Rbm38 | ENSMUSG00000027510 | 56190 | -0.964955565986957 | 6.48445294194546 | -13.0703245308252 | 5.2037282438414 | 3.78989553752527e-07 |
| Vegfb | ENSMUSG00000024962 | 22340 | -0.966676118261233 | 6.47400114368904 | -13.0639905912648 | 5.2353327798583 | 3.78989553752527e-07 |
| Sort1 | ENSMUSG00000068747 | 20661 | 1.20396641289295 | 3.8269458592714 | 13.0416977241418 | 5.3482102754766 | 3.84613727113393e-07 |
| Thbs4 | ENSMUSG00000021702 | 21828 | 2.80682481958651 | 8.44444289860216 | 13.0138926179845 | 5.4926579175425 | 3.92419893442205e-07 |
| Fabp3 | ENSMUSG00000028773 | 14077 | -1.4989820073137 | 9.46606715704761 | -12.9475064190135 | 5.8546757090570 | 4.15567923218851e-07 |
| Efcab6 | ENSMUSG00000022441 | 77627 | -1.64670262220387 | 3.11237416349254 | -12.93811565913 | 5.9079110971524 | 4.16641136793379e-07 |
| Cpt2 | ENSMUSG00000028607 | 12896 | -0.904189903403824 | 5.53553433955861 | -12.9177116457878 | 6.0253717549369 | 4.21518372377424e-07 |
| Dennd4b | ENSMUSG00000042404 | 229541 | -0.766480341771933 | 5.78982104993999 | -12.9127700693487 | 6.0541930713800 | 4.21518372377424e-07 |
| Asb14 | ENSMUSG00000021898 | 142687 | -1.2473154831491 | 4.40307886586411 | -12.9056408249449 | 6.0960337639756 | 4.21745221987457e-07 |
| Lsp1 | ENSMUSG00000018819 | 16985 | 1.3621566636003 | 4.89309753398278 | 12.8810642397743 | 6.2426569010930 | 4.29172846451873e-07 |
| Bace1 | ENSMUSG00000032086 | 23821 | 0.943106149528808 | 4.21306107678717 | 12.8469933747232 | 6.4521910215810 | 4.40805625355638e-07 |
| Sh3pxd2b | ENSMUSG00000040711 | 268396 | 1.29158020257632 | 4.06943353329238 | 12.7928985874644 | 6.8004676033446 | 4.59482230176857e-07 |
| Ldhb | ENSMUSG00000030246 | 16832 | -1.12221624821068 | 8.76427276607179 | -12.791514970978 | 6.8096351009652 | 4.59482230176857e-07 |
| Cpt1b | ENSMUSG00000078937 | 12895 | -0.872527511693107 | 7.57268180929666 | -12.7807410112967 | 6.8814742295373 | 4.61480949712101e-07 |
| Coq10a | ENSMUSG00000039914 | 210582 | -0.875093584723107 | 7.0198122760996 | -12.7609093954393 | 7.0158351553303 | 4.65243291007721e-07 |
| Cpne2 | ENSMUSG00000034361 | 234577 | 1.82835799192057 | 2.96326667427577 | 12.7599069494385 | 7.0227008522801 | 4.65243291007721e-07 |
| Cox8b | ENSMUSG00000025488 | 12869 | -1.05880675764056 | 7.5022601834633 | -12.685503373168 | 7.5528934283425 | 4.97353482320557e-07 |
| Maob | ENSMUSG00000040147 | 109731 | -2.10031948205149 | 2.32687592366377 | -12.6570960452164 | 7.7664703812774 | 5.08355016393678e-07 |
| Ccl6 | ENSMUSG00000018927 | 20305 | 2.22637768632433 | 3.22530144135015 | 12.6508015584663 | 7.8146628145169 | 5.08464757294552e-07 |
| Asb10 | ENSMUSG00000038204 | 117590 | -1.19067797270145 | 4.4597568237991 | -12.6156373998117 | 8.0898368092988 | 5.23254474333997e-07 |
| Sdhb | ENSMUSG00000009863 | 67680 | -1.0378476158391 | 7.6627607053648 | -12.603590886795 | 8.1864724513526 | 5.26390178621978e-07 |
| Tceal7 | ENSMUSG00000079428 | 100040972 | 1.61667036327167 | 3.81687186091423 | 12.5822485623048 | 8.3607201824145 | 5.34450481368264e-07 |
| Tmem43 | ENSMUSG00000030095 | 74122 | 0.960298600366936 | 4.31999152635899 | 12.4705338163371 | 9.3398447527425 | 5.92808436632187e-07 |
| Lox | ENSMUSG00000024529 | 16948 | 2.38202735689772 | 4.55799390491328 | 12.4617971181975 | 9.4214228717793 | 5.92808436632187e-07 |
| Nadk2 | ENSMUSG00000022253 | 68646 | -1.17691412808957 | 4.01890746929093 | -12.4602079522861 | 9.4363432416064 | 5.92808436632187e-07 |
| Acadvl | ENSMUSG00000018574 | 11370 | -1.00555910249231 | 7.55865504321474 | -12.4397478532912 | 9.6307109044347 | 6.01561719407866e-07 |
| Il13ra1 | ENSMUSG00000017057 | 16164 | 1.80243986475434 | 2.82388819950314 | 12.4165595981153 | 9.8561804208100 | 6.11080354907052e-07 |
| Osbpl6 | ENSMUSG00000042359 | 99031 | -1.50933465633605 | 3.1874028024486 | -12.4126339351746 | 9.8949064878371 | 6.11080354907052e-07 |
| Lgmn | ENSMUSG00000021190 | 19141 | 0.932950652078911 | 5.41260836814995 | 12.335044968953 | 1.0694637062782 | 6.53654697853212e-07 |
| Sod2 | ENSMUSG00000006818 | 20656 | -1.06962172231743 | 7.23826043352572 | -12.3341842173024 | 1.0703887193827 | 6.53654697853212e-07 |
| Prr33 | ENSMUSG00000043795 | 677289 | -1.20543257398851 | 5.264806631407 | -12.3096301666637 | 1.0971392383906 | 6.64978525870766e-07 |
| Kctd12 | ENSMUSG00000098557 | 239217 | 1.03673475447541 | 4.4487742289484 | 12.3019164183308 | 1.1056897747605 | 6.64978525870766e-07 |
| Pde4d | ENSMUSG00000021699 | 238871 | -0.915050469046161 | 6.44295454969334 | -12.3005765382339 | 1.1071822496430 | 6.64978525870766e-07 |
| Tead4 | ENSMUSG00000030353 | 21679 | 0.965196747752417 | 3.84034413054852 | 12.2849708443857 | 1.1247246483196 | 6.71823231190298e-07 |
| Anxa1 | ENSMUSG00000024659 | 16952 | 1.59841552521645 | 4.60040326890291 | 12.2659503054701 | 1.1465082180928 | 6.81113115868067e-07 |
| Panx1 | ENSMUSG00000031934 | 55991 | 1.26392302224175 | 2.96131020899568 | 12.2520127332474 | 1.1627564492712 | 6.87031932269433e-07 |
| Ctsz | ENSMUSG00000016256 | 64138 | 1.27160810400604 | 4.1173681492706 | 12.2449224162212 | 1.1711165049252 | 6.88251318028893e-07 |
| C4b | ENSMUSG00000073418 | 12268 | 2.04910604750887 | 3.97833692708567 | 12.2349278549531 | 1.1830101431078 | 6.91523201834861e-07 |
| Wwp1 | ENSMUSG00000041058 | 107568 | -0.877791447389045 | 7.06384433246954 | -12.2178315217411 | 1.2036551878371 | 6.99848662672785e-07 |
| Gm2a | ENSMUSG00000000594 | 14667 | 1.10268183752432 | 3.43552559890781 | 12.1923313624694 | 1.2351669035248 | 7.14370868911646e-07 |
| Mmut | ENSMUSG00000023921 | 17850 | -0.815161315451553 | 5.09460045128455 | -12.1837027727845 | 1.2460282882386 | 7.16859748354537e-07 |
| Ptprj | ENSMUSG00000025314 | 19271 | 1.09635763411455 | 3.75408818253349 | 12.1717808226179 | 1.2612034562877 | 7.21791360245097e-07 |
| Fras1 | ENSMUSG00000034687 | 231470 | 1.66694313028751 | 3.52994716427385 | 12.1422356359407 | 1.2996683654657 | 7.39930984526379e-07 |
| Gamt | ENSMUSG00000020150 | 14431 | 2.0379511408055 | 1.69120592519129 | 12.1316640612028 | 1.3137343603944 | 7.44063745775715e-07 |
| Nrcam | ENSMUSG00000020598 | 319504 | 3.39437385015217 | 1.11155463682936 | 12.1254430341392 | 1.3220876815912 | 7.4493507461208e-07 |
| Tmtc1 | ENSMUSG00000030306 | 387314 | -0.841022849702513 | 6.10997693261993 | -12.0906551333804 | 1.3698573080998 | 7.67892833009949e-07 |
| Plp2 | ENSMUSG00000031146 | 18824 | 0.973552097897282 | 4.00003313198852 | 12.0441799317761 | 1.4365730562208 | 8.01182657017872e-07 |
| Col14a1 | ENSMUSG00000022371 | 12818 | 1.60310462120991 | 5.03655224291602 | 12.0108942868164 | 1.4864842438005 | 8.24810115176835e-07 |
| Mtfp1 | ENSMUSG00000004748 | 67900 | -1.19957815858871 | 4.09664234270586 | -11.9821066419026 | 1.5311454748345 | 8.45300564920024e-07 |
| Crat | ENSMUSG00000026853 | 12908 | -0.822021860855715 | 7.81700802077255 | -11.9674361563722 | 1.5544547272697 | 8.53856513758076e-07 |
| Phka1 | ENSMUSG00000034055 | 18679 | -0.890419469743939 | 6.20839702453749 | -11.9559373817188 | 1.5729895173351 | 8.59717420699511e-07 |
| C77080 | ENSMUSG00000050390 | 97130 | 1.47693427326908 | 2.17227870734635 | 11.9441011958392 | 1.5923149696189 | 8.65949996661924e-07 |
| Etfb | ENSMUSG00000004610 | 110826 | -1.06225976232343 | 6.7797582517065 | -11.9081340347015 | 1.6526124960366 | 8.94292435355291e-07 |
| Ndufa12 | ENSMUSG00000020022 | 66414 | -0.897295539908276 | 6.10979493023729 | -11.8900587181802 | 1.6838317213002 | 9.06697760863704e-07 |
| Vcan | ENSMUSG00000021614 | 13003 | 1.61623346817207 | 5.00303101354948 | 11.8847583145622 | 1.6931054378958 | 9.07222330472544e-07 |
| Acadl | ENSMUSG00000026003 | 11363 | -0.835640449967697 | 8.13901797587744 | -11.8793886032793 | 1.7025560664698 | 9.07836115247886e-07 |
| Acot9 | ENSMUSG00000025287 | 56360 | 0.969186291058369 | 3.97445807613885 | 11.8662118741452 | 1.7259865485569 | 9.15862085547394e-07 |
| Phkg1 | ENSMUSG00000025537 | 18682 | -1.21135058094383 | 4.96904201507953 | -11.8528890534671 | 1.7500273000181 | 9.24132773743905e-07 |
| C1qb | ENSMUSG00000036905 | 12260 | 1.84194050911796 | 3.454431589156 | 11.8455679447509 | 1.7633901358021 | 9.25278487382842e-07 |
| Isoc2a | ENSMUSG00000086784 | 664994 | -1.1154978571964 | 2.95108007344267 | -11.8424433852464 | 1.7691263732779 | 9.25278487382842e-07 |
| Etfa | ENSMUSG00000032314 | 110842 | -0.93284266058797 | 6.70559185913477 | -11.8326167837562 | 1.7872968554068 | 9.2556309908562e-07 |
| Mrc1 | ENSMUSG00000026712 | 17533 | 1.50558879547196 | 4.62971314066691 | 11.8322925612453 | 1.7878997654104 | 9.2556309908562e-07 |
| Plxnb2 | ENSMUSG00000036606 | 140570 | 1.14131021656647 | 4.97598544359573 | 11.8268986760444 | 1.7979619289960 | 9.2556309908562e-07 |
| Mfn1 | ENSMUSG00000027668 | 67414 | -1.01257724257478 | 7.32955932626239 | -11.8239224733865 | 1.8035398417824 | 9.2556309908562e-07 |
| Ckmt2 | ENSMUSG00000021622 | 76722 | -1.08507821747028 | 9.80630156722225 | -11.809769686243 | 1.8303186641606 | 9.34916510184135e-07 |
| Unc93b1 | ENSMUSG00000036908 | 54445 | 1.36779818062832 | 3.5878341900213 | 11.7786695742805 | 1.8906673481217 | 9.60573654774847e-07 |
| Scn4a | ENSMUSG00000001027 | 110880 | -0.821772090052399 | 7.11014255587832 | -11.7749007361113 | 1.8981237712136 | 9.60573654774847e-07 |
| Acadsb | ENSMUSG00000030861 | 66885 | -0.84220650317957 | 5.86212168376951 | -11.7122231576969 | 2.0268422970184 | 1.01811159643623e-06 |
| Col6a3 | ENSMUSG00000048126 | 12835 | 1.68469605093097 | 7.0574203854666 | 11.7093309773609 | 2.0330025134224 | 1.01811159643623e-06 |
| Esyt1 | ENSMUSG00000025366 | 23943 | 0.84973116739004 | 5.19479601911173 | 11.7061679559159 | 2.0397625068112 | 1.01811159643623e-06 |
| Ptpn3 | ENSMUSG00000038764 | 545622 | -1.013081025905 | 6.6653690111254 | -11.6857778396325 | 2.0839203793344 | 1.0354242575684e-06 |
| Tln1 | ENSMUSG00000028465 | 21894 | 0.727129432150895 | 7.53333858854895 | 11.6794143203944 | 2.0979096860836 | 1.0376584062706e-06 |
| Rin1 | ENSMUSG00000024883 | 225870 | 2.38583672961314 | 2.28037967466757 | 11.6646859151247 | 2.1306741418240 | 1.04911707406662e-06 |
| Phyhd1 | ENSMUSG00000079484 | 227696 | -1.48314549296419 | 3.46134193238288 | -11.6587354005805 | 2.1440661110353 | 1.05097698025681e-06 |
| Lgals1 | ENSMUSG00000068220 | 16852 | 0.885449068429824 | 5.96369576546014 | 11.6340941353337 | 2.2004870246225 | 1.06575494361145e-06 |
| Acacb | ENSMUSG00000042010 | 100705 | -0.795870528891703 | 8.09974440141346 | -11.6339818799329 | 2.2007476534735 | 1.06575494361145e-06 |
| Dlgap4 | ENSMUSG00000061689 | 228836 | 0.956679495800699 | 5.34700087422502 | 11.6328128953302 | 2.2034637019137 | 1.06575494361145e-06 |
| Cd74 | ENSMUSG00000024610 | 16149 | 1.92624174422317 | 6.07250064519924 | 11.6258257398183 | 2.2197726495581 | 1.06891342873658e-06 |
| Erbb2 | ENSMUSG00000062312 | 13866 | 1.18388771950223 | 2.32929095577355 | 11.6067688879037 | 2.2649125595132 | 1.08586663105435e-06 |
| Klhl34 | ENSMUSG00000047485 | 245683 | -0.806775360412908 | 5.38397903394678 | -11.5986016845249 | 2.2845573377348 | 1.09050202003403e-06 |
| Myof | ENSMUSG00000048612 | 226101 | 1.22317748816927 | 4.08723342534252 | 11.5764427585034 | 2.3387789107239 | 1.10958897628778e-06 |
| Cyp2u1 | ENSMUSG00000027983 | 71519 | -1.10106952242483 | 3.19884879907819 | -11.573997715962 | 2.3448454260587 | 1.10958897628778e-06 |

| | | | | | | | |
|---|---|---|---|---|---|---|---|
| Svep1 | ENSMUSG00000028369 | 64817 | 1.43001854542279 | 5.0069063006176 | 11.5676135329939 | 2.3607648855098 | 1.11062388426996e-06 |
| Antxr2 | ENSMUSG00000029338 | 71914 | 0.847676868325009 | 4.2676608914188 | 11.5649849051164 | 2.3673530787201 | 1.11062388426996e-06 |
| Fgd3 | ENSMUSG00000037946 | 30938 | 2.34804782645173 | 2.72022127109034 | 11.5409833756998 | 2.4284246571634 | 1.13092446064394e-06 |
| Nckap1l | ENSMUSG00000022488 | 105855 | 1.75360447096364 | 2.25188994017218 | 11.539862746135 | 2.4313168809013 | 1.13092446064394e-06 |
| Osbpl3 | ENSMUSG00000029822 | 71720 | 1.29056662796838 | 3.27086602163615 | 11.5281390919172 | 2.4617958238812 | 1.14024958266296e-06 |
| Ncs1 | ENSMUSG00000062661 | 14299 | 1.1666950785523 | 2.90352351323362 | 11.5194534678654 | 2.4846395233344 | 1.14597445694383e-06 |
| Acss2 | ENSMUSG00000027605 | 60525 | -0.915319182081584 | 4.87607988541883 | -11.5047487576681 | 2.5238312683689 | 1.15915964682944e-06 |
| Pkdcc | ENSMUSG00000024247 | 106522 | -1.52304287769189 | 4.50537132398792 | -11.4610816112078 | 2.6441555861586 | 1.2093416197615e-06 |
| Cd44 | ENSMUSG00000005087 | 12505 | 1.7749586690274 | 4.22615472061731 | 11.4413791986248 | 2.7004372382325 | 1.22993664379667e-06 |
| Tbc1d16 | ENSMUSG00000039976 | 207592 | -1.05288670300299 | 3.8195365200238 | -11.4296563780033 | 2.7345302578460 | 1.24029669080975e-06 |
| Ndrg4 | ENSMUSG00000036564 | 234593 | 2.90308449939112 | 3.73191119085047 | 11.4171069094386 | 2.7715372435234 | 1.2474546424469e-06 |
| Echs1 | ENSMUSG00000025465 | 93747 | -0.944333506122983 | 5.65557289703477 | -11.4165688472367 | 2.7731358349153 | 1.2474546424469e-06 |
| Slc25a24 | ENSMUSG00000040322 | 229731 | 1.39429113681197 | 3.01607756137697 | 11.3980365813996 | 2.8288007278742 | 1.26727953919647e-06 |
| Suclg2 | ENSMUSG00000061838 | 20917 | -0.799657839230743 | 5.22185191309399 | -11.3740317642551 | 2.9026829255960 | 1.29507049223228e-06 |
| Srgap3 | ENSMUSG00000030257 | 259302 | 1.27289800532346 | 2.96752267262369 | 11.363968438516 | 2.9342662769813 | 1.30384002738547e-06 |
| Sync | ENSMUSG00000001333 | 68828 | 0.813966400825413 | 5.3865429660682 | 11.3568728590447 | 2.9567558862201 | 1.30851411304745e-06 |
| Tnnt3 | ENSMUSG00000061723 | 21957 | -1.23979819698705 | 11.7936482220882 | -11.3392046358895 | 3.0135591329865 | 1.3282747936563e-06 |
| Plekhh1 | ENSMUSG00000060716 | 211945 | 1.7468587061635 | 1.87478413766827 | 11.3269597102726 | 3.0536092864726 | 1.33638155010821e-06 |
| Stbd1 | ENSMUSG00000047963 | 52331 | 1.24896982586788 | 3.209561127838 | 11.3253953447611 | 3.0587667310169 | 1.33638155010821e-06 |
| Pde4a | ENSMUSG00000032177 | 18577 | -1.14233035683921 | 4.33087687188201 | -11.3224119126983 | 3.0686283878617 | 1.33638155010821e-06 |
| Hmcn1 | ENSMUSG00000066842 | 545370 | 1.83009927682146 | 2.85060207912091 | 11.3070845262198 | 3.1198305546705 | 1.35328840448825e-06 |
| Gatd3a | ENSMUSG00000053329 | 28295 | -0.813516445682407 | 6.12973670250939 | -11.2830689179297 | 3.2018987727955 | 1.38339745001691e-06 |
| Pdk2 | ENSMUSG00000038967 | 18604 | -0.830657811372522 | 7.93036830080331 | -11.2640699215852 | 3.2684558684885 | 1.4065941377342e-06 |
| Trp63 | ENSMUSG00000022510 | 22061 | 2.13547211216438 | 2.75905712183849 | 11.2465235162062 | 3.3312369748323 | 1.42799024987814e-06 |
| Ctss | ENSMUSG00000038642 | 13040 | 2.52114312424144 | 3.953739733993 | 11.2383512014806 | 3.3609157666013 | 1.43185488015657e-06 |
| Pirb | ENSMUSG00000058818 | 18733 | 1.93530675443713 | 2.49136332385112 | 11.2359891631099 | 3.3695462855696 | 1.43185488015657e-06 |
| Tanc2 | ENSMUSG00000053580 | 77097 | 0.889964986706312 | 4.34566153563861 | 11.2332594975745 | 3.3795495295983 | 1.43185488015657e-06 |
| Ppm1l | ENSMUSG00000027784 | 242083 | -0.880530194087119 | 5.39264591145971 | -11.2288687709999 | 3.3957065208389 | 1.43285826850072e-06 |
| Suclg1 | ENSMUSG00000052738 | 56451 | -0.896022252791287 | 6.82035055999507 | -11.2255066434786 | 3.4081342037342 | 1.43285826850072e-06 |
| Dhrs4 | ENSMUSG00000022210 | 28200 | -0.964872672023513 | 4.34889439973602 | -11.2014121043444 | 3.4986297157196 | 1.46507288242026e-06 |
| Ivd | ENSMUSG00000027332 | 56357 | -0.769421957669386 | 6.43890690006653 | -11.1974162087154 | 3.5138840192736 | 1.46507288242026e-06 |
| Map1a | ENSMUSG00000027254 | 17754 | 1.34915451103829 | 6.43791042382129 | 11.1945245339272 | 3.5249672315115 | 1.46507288242026e-06 |
| Carnmt1 | ENSMUSG00000024726 | 67383 | -0.960548088437272 | 6.69796813771774 | -11.1806126536065 | 3.5788127063192 | 1.48181824593847e-06 |
| Itgb2 | ENSMUSG00000000290 | 16414 | 1.88910223613166 | 2.58770045736766 | 11.1632875788985 | 3.6470991432045 | 1.50439398997617e-06 |
| Mcrip2 | ENSMUSG00000025732 | 68241 | -1.05299024031326 | 3.78507554008552 | -11.1372666495219 | 3.7522855763699 | 1.54196367050001e-06 |
| Pank1 | ENSMUSG00000033610 | 75735 | -1.23781008437655 | 4.26792753503067 | -11.126412490572 | 3.7971149237012 | 1.55454169404412e-06 |
| Plaat1 | ENSMUSG00000022525 | 27281 | -1.12607392497303 | 4.06945919106608 | -11.1086828447958 | 3.8715749178746 | 1.57911139654058e-06 |
| Uchl1 | ENSMUSG00000029223 | 22223 | 1.7618518452388 | 3.78611240606905 | 11.079271105952 | 3.9985545450023 | 1.62484013871454e-06 |
| Laptm5 | ENSMUSG00000028581 | 16792 | 1.72811474250143 | 3.26672879298264 | 11.0736586677848 | 4.0232862354695 | 1.62883488295992e-06 |
| Atpaf1 | ENSMUSG00000028710 | 230649 | -0.892630102289155 | 4.61028762293608 | -11.0663014584427 | 4.0559539349185 | 1.63600119788174e-06 |
| Igsf1 | ENSMUSG00000031111 | 209268 | 1.89360558520681 | 2.05153956016618 | 11.04882721369 | 4.1346835181658 | 1.6616259388629e-06 |
| Bcat2 | ENSMUSG00000030826 | 12036 | -0.946184042994437 | 4.04732738061302 | -11.0393262053188 | 4.1781729496063 | 1.67295269275263e-06 |
| H2-Ab1 | ENSMUSG00000073421 | 14961 | 2.09768838671995 | 4.79476108442803 | 11.0215144019859 | 4.2610245891646 | 1.69989999212259e-06 |
| Eppk1 | ENSMUSG00000115388 | 223650 | 2.16601231957675 | 1.7283383422689 | 11.0144497197021 | 4.2943699872423 | 1.70592406790665e-06 |
| Cybb | ENSMUSG00000015340 | 13058 | 1.83666670395252 | 2.86436333186598 | 11.0065138568751 | 4.3321597292244 | 1.70592406790665e-06 |
| Hsd17b4 | ENSMUSG00000024507 | 15488 | -0.68158015669843 | 5.4262068615388 | -11.0041168038971 | 4.3436438983404 | 1.70592406790665e-06 |
| Vdac1 | ENSMUSG00000020402 | 22333 | -0.815828947409836 | 8.60457479275787 | -11.0025125432846 | 4.3513479785269 | 1.70592406790665e-06 |
| Trim47 | ENSMUSG00000020773 | 217333 | 1.20036674270418 | 3.89267751037864 | 11.0019285249501 | 4.3541562066229 | 1.70592406790665e-06 |
| Stk40 | ENSMUSG00000042608 | 74178 | -0.576585649003905 | 6.06180880821498 | -10.9748073881018 | 4.4867210554753 | 1.75158385205005e-06 |
| Acot2 | ENSMUSG00000021226 | 171210 | -1.43028560077747 | 4.23448789774025 | -10.9704272081317 | 4.5085317281524 | 1.75236548315554e-06 |
| Dock8 | ENSMUSG00000052085 | 76088 | 0.90289687359682 | 2.92703200238307 | 10.9679762177494 | 4.5207855296849 | 1.75236548315554e-06 |
| Gstm1 | ENSMUSG00000058135 | 14862 | -0.782651138178941 | 5.17346379801785 | -10.9516735389314 | 4.6032009607341 | 1.77800670324328e-06 |
| Casr | ENSMUSG00000051980 | 12374 | 4.46794847962245 | 1.00992576918694 | 10.947336726451 | 4.6253938960663 | 1.78028805203876e-06 |
| Chpt1 | ENSMUSG00000060002 | 212862 | -0.921609306309421 | 6.42065445474731 | -10.939202578281 | 4.6673275311322 | 1.79012481553706e-06 |
| Sirpa | ENSMUSG00000037902 | 19261 | 1.23372327124266 | 3.75144684622643 | 10.9292912766537 | 4.7189718173588 | 1.80155988915386e-06 |
| Pkp2 | ENSMUSG00000041957 | 67451 | 1.93216971784735 | 2.25150555749153 | 10.9270118052853 | 4.7309353583000 | 1.80155988915386e-06 |
| Prkg1 | ENSMUSG00000052920 | 19091 | -0.780121652682758 | 5.77133887703351 | -10.9240391977382 | 4.7465853817245 | 1.80155988915386e-06 |
| Uqcc1 | ENSMUSG00000005882 | 56046 | -0.886251762937091 | 5.1794965910681 | -10.9197604618611 | 4.7692088842794 | 1.80388312505395e-06 |
| Hspd1 | ENSMUSG00000025980 | 15510 | -0.700996477888718 | 6.49666493525322 | -10.9092226247489 | 4.8254189481725 | 1.81885015594737e-06 |
| Tmem143 | ENSMUSG00000002781 | 70209 | -0.871042678629437 | 4.11801180736309 | -10.9003729478221 | 4.8731702414280 | 1.83053690409107e-06 |
| Bnc2 | ENSMUSG00000028487 | 242509 | 1.26503275721346 | 2.69778432521373 | 10.8881384169285 | 4.9400175973212 | 1.84126601050907e-06 |
| Arpp21 | ENSMUSG00000032503 | 74100 | 1.20075201994885 | 5.66585248409012 | 10.8875611357383 | 4.9431958341961 | 1.84126601050907e-06 |
| Rap2b | ENSMUSG00000036894 | 74012 | 1.26698616515446 | 2.40251562071566 | 10.8859158274627 | 4.9522660972433 | 1.84126601050907e-06 |
| Mmp2 | ENSMUSG00000031740 | 17390 | 1.66799917733294 | 6.47838457549236 | 10.8750702181646 | 5.0125018887033 | 1.85734434391242e-06 |
| Col5a3 | ENSMUSG00000004098 | 53867 | 1.23858114508784 | 5.46364080616565 | 10.8630285645923 | 5.0802973275420 | 1.87463788656748e-06 |
| Tnni2 | ENSMUSG00000031097 | 21953 | -1.10067939450458 | 10.8258829551942 | -10.8607083679206 | 5.0934722560657 | 1.87463788656748e-06 |
| Lrtm1 | ENSMUSG00000045776 | 319476 | -1.83314113970471 | 5.23898662028015 | -10.8383923735857 | 5.2220680659256 | 1.91551765196757e-06 |
| Gpnmb | ENSMUSG00000029816 | 93695 | 2.58680261082181 | 1.79377644432882 | 10.8167515561482 | 5.3500857322777 | 1.95087202730587e-06 |
| S100a6 | ENSMUSG00000001025 | 20200 | 0.91661751249695 | 4.393647791038 | 10.8160743920912 | 5.3541451668809 | 1.95087202730587e-06 |
| Gas1 | ENSMUSG00000052957 | 14451 | 1.2295289488099 | 3.93548710399598 | 10.8105379861571 | 5.3874580907059 | 1.95648851792382e-06 |
| Dach2 | ENSMUSG00000025592 | 93837 | -3.85334964391857 | 1.4286049513064 | -10.8001622752319 | 5.4504863752660 | 1.97282339629249e-06 |
| Adam12 | ENSMUSG00000054555 | 11489 | 2.62634793743137 | 1.83202942053677 | 10.7946153560056 | 5.4845039371211 | 1.97858457216736e-06 |
| Capn3 | ENSMUSG00000079110 | 12335 | -0.675336010745421 | 6.17916285222535 | -10.7913288225362 | 5.5047660319681 | 1.97936175971855e-06 |
| Pex11a | ENSMUSG00000030545 | 18631 | -1.38847265116631 | 2.27628987115417 | -10.7882990983849 | 5.5235155950109 | 1.97959176947754e-06 |
| Prxl2a | ENSMUSG00000021792 | 70564 | -1.1748457752857 | 4.76202478881903 | -10.7669695693564 | 5.6574576749727 | 2.02096960278193e-06 |
| Alad | ENSMUSG00000028393 | 17025 | -1.03530955783121 | 3.37999525893052 | -10.7502067143092 | 5.7651531712119 | 2.04560249280216e-06 |
| Mpc2 | ENSMUSG00000026568 | 70456 | -1.01536813187485 | 6.56849559088902 | -10.7478765824605 | 5.7802957250492 | 2.04560249280216e-06 |
| Prune2 | ENSMUSG00000039126 | 353211 | 1.46083232884588 | 6.37047327052513 | 10.7475293672117 | 5.7825557613746 | 2.04560249280216e-06 |
| Rmdn1 | ENSMUSG00000028229 | 66302 | -1.1230680371654 | 4.01758882360342 | -10.7325025447098 | 5.8812746637303 | 2.07381333384632e-06 |
| Sln | ENSMUSG00000042045 | 66402 | 2.68452216167858 | 5.61035824176785 | 10.7264058470657 | 5.9218382465147 | 2.08140237532646e-06 |
| Tmem176a | ENSMUSG00000023367 | 66058 | 1.42042808549976 | 1.93152994819078 | 10.6921718726203 | 6.1552215281401 | 2.14565155577823e-06 |
| Fibin | ENSMUSG00000074971 | 67606 | 1.70100858668614 | 2.54257235178138 | 10.6912996305687 | 6.1612945313523 | 2.14565155577823e-06 |
| Ncf1 | ENSMUSG00000015950 | 17969 | 1.83357762220735 | 1.43498084715801 | 10.6909799393688 | 6.1635219880556 | 2.14565155577823e-06 |
| Akr1b8 | ENSMUSG00000029762 | 14187 | 2.02835485858176 | 1.15718365034863 | 10.6684496576399 | 6.3226862364544 | 2.19407248414867e-06 |
| Ucp2 | ENSMUSG00000033685 | 22228 | 1.38742323004456 | 4.44866966830415 | 10.6384335665096 | 6.5415778683277 | 2.26284771135098e-06 |
| Fam20c | ENSMUSG00000025854 | 80752 | 0.884257467173798 | 4.07389335061396 | 10.6334451013631 | 6.5787327048533 | 2.26852136267356e-06 |
| Pdpn | ENSMUSG00000028583 | 14726 | 1.3889895753231 | 2.57802863613511 | 10.6227364327937 | 6.6592547806522 | 2.2890664782173e-06 |
| Bckdk | ENSMUSG00000030802 | 12041 | -0.755557205004812 | 5.53279059700797 | -10.6157338736164 | 6.7124773186623 | 2.30012819969586e-06 |
| Ccdc88a | ENSMUSG00000032740 | 108686 | 0.891491940122679 | 3.53761408070869 | 10.5985827885944 | 6.8447587555699 | 2.33186981360136e-06 |
| Cycs | ENSMUSG00000063694 | 13063 | -0.901229953198493 | 7.16748443896668 | -10.5981959356385 | 6.8477743131098 | 2.33186981360136e-06 |
| Gprc5b | ENSMUSG00000008734 | 64297 | 0.912347137671898 | 3.90343585987802 | 10.5947135871424 | 6.8749834354020 | 2.33386471839688e-06 |
| Capn6 | ENSMUSG00000067276 | 12338 | 1.78461903376902 | 1.89707551938713 | 10.5598142007281 | 7.1541240872607 | 2.42110620426771e-06 |
| Ablim1 | ENSMUSG00000025085 | 226251 | 0.860584668015358 | 8.66561482114606 | 10.5518915315675 | 7.2191612530588 | 2.43557566843166e-06 |
| Dusp26 | ENSMUSG00000039661 | 66959 | -1.49063970213294 | 3.13602322564147 | -10.5419900672655 | 7.3013299950199 | 2.44986995634133e-06 |
| Clpx | ENSMUSG00000015357 | 270166 | -0.775297597491357 | 5.4685298857129 | -10.5398171736049 | 7.3194951695351 | 2.44986995634133e-06 |
| Cpeb2 | ENSMUSG00000039782 | 231207 | 0.996331656376781 | 3.38947114917853 | 10.5387103544631 | 7.3287665879024 | 2.44986995634133e-06 |
| Acad11 | ENSMUSG00000090150 | 102632 | -1.05998734137323 | 4.85035551228163 | -10.5188408156414 | 7.4973542991211 | 2.49858475133212e-06 |
| Rnh1 | ENSMUSG00000038650 | 107702 | 0.829128429075457 | 3.98883205156206 | 10.4996071258427 | 7.6644908489109 | 2.53860834329863e-06 |
| Fcgr2b | ENSMUSG00000026656 | 14130 | 1.85386347927017 | 2.62778916033851 | 10.495459376672 | 7.7010520029103 | 2.53860834329863e-06 |
| Slc9a9 | ENSMUSG00000031129 | 331004 | 1.73288645505648 | 1.81110747718533 | 10.4947328894886 | 7.7074748838769 | 2.53860834329863e-06 |
| Ttc9 | ENSMUSG00000042734 | 69480 | 0.916647549545508 | 5.06365238959794 | 10.493367470253 | 7.7195620313528 | 2.53860834329863e-06 |
| Casp3 | ENSMUSG00000031628 | 12367 | 1.4883278939291 | 1.78994200233315 | 10.4917878723948 | 7.7335703807377 | 2.53860834329863e-06 |
| Cd68 | ENSMUSG00000018774 | 12514 | 1.79670917293441 | 2.13549340010919 | 10.488146008663 | 7.7659709162869 | 2.5416116193393e-06 |
| Stat3 | ENSMUSG00000004040 | 20848 | 0.757539348811641 | 5.94771454549359 | 10.4847447222043 | 7.7963618478879 | 2.54394123460489e-06 |
| Idh3a | ENSMUSG00000032279 | 67834 | -0.955813994705185 | 7.97927906063862 | -10.4805892768108 | 7.8336634924737 | 2.54805225161963e-06 |
| C1qc | ENSMUSG00000036896 | 12262 | 1.55279401219799 | 3.79009458604102 | 10.478157420075 | 7.8555814545404 | 2.54805225161963e-06 |
| Ndufa9 | ENSMUSG00000000399 | 66108 | -0.734225962482065 | 6.41592798152175 | -10.4719323201319 | 7.9119858202201 | 2.55875494085287e-06 |
| Fyb | ENSMUSG00000022148 | 23880 | 2.22761982422125 | 2.03228994747435 | 10.4680019831222 | 7.9478200771642 | 2.56276168918828e-06 |
| Qrsl1 | ENSMUSG00000019863 | 76563 | -0.770141048123081 | 3.35119054728884 | -10.4617623638285 | 8.0050647619254 | 2.57362832095904e-06 |
| H2-Eb1 | ENSMUSG00000060586 | 14969 | 2.10784861962735 | 3.89308640463348 | 10.4510428999839 | 8.1044381366820 | 2.59793587308128e-06 |
| Fam129b | ENSMUSG00000026796 | 227737 | 1.05465120270692 | 3.7637780480178 | 10.4477743560324 | 8.1349997790027 | 2.6001076779029e-06 |
| Man1a | ENSMUSG00000003746 | 17155 | 0.923625144783396 | 4.08466146617772 | 10.4324876162114 | 8.2795745376134 | 2.63860143646217e-06 |
| Dcun1d2 | ENSMUSG00000038506 | 102323 | -0.568110998929037 | 4.74370621648349 | -10.410736359047 | 8.4900286530538 | 2.69106613385091e-06 |
| Hmcn2 | ENSMUSG00000055632 | 665700 | 1.34131364732765 | 7.39182028828966 | 10.4103886374847 | 8.4934389916619 | 2.69106613385091e-06 |
| Pcdhgc3 | ENSMUSG00000102918 | 93706 | 0.973462491762196 | 4.85983732875197 | 10.3544157828474 | 9.0619125518585 | 2.85064168621879e-06 |
| Selenbp1 | ENSMUSG00000068874 | 20341 | -0.943345923960992 | 3.66146973613646 | -10.3535746254826 | 9.0707587825349 | 2.85064168621879e-06 |

| | | | | | | | |
|---|---|---|---|---|---|---|---|
| Lamc1 | ENSMUSG00000026478 | 226519 | 0.856251961015811 | 7.49126471618202 | 10.3525399442436 | 9.0816528816263 | 2.85064168621879e-06 |
| Ankrd23 | ENSMUSG00000067653 | 78321 | -0.966175540204176 | 9.8702256732822 | -10.3506678785964 | 9.1013992177326 | 2.85064168621879e-06 |
| Gusb | ENSMUSG00000025534 | 110006 | 0.919206948301102 | 3.41107381419957 | 10.3379461226625 | 9.2368051694893 | 2.88478620879108e-06 |
| C3ar1 | ENSMUSG00000040552 | 12267 | 2.21037352090099 | 2.17834989178417 | 10.3291783143224 | 9.3313757979118 | 2.90601905546936e-06 |
| Dock2 | ENSMUSG00000020143 | 94176 | 1.70446998015898 | 2.62780268367453 | 10.3241567262585 | 9.3860036650089 | 2.91472744494923e-06 |
| Kcnn3 | ENSMUSG00000000794 | 140493 | 1.42887203722582 | 4.21884252727033 | 10.3167637869352 | 9.4670498209661 | 2.93156718393713e-06 |
| Vdac3 | ENSMUSG00000008892 | 22335 | -0.908535717872684 | 7.05237306046127 | -10.3120841153785 | 9.5187367118802 | 2.93924607337749e-06 |
| Chchd10 | ENSMUSG00000049422 | 103172 | -0.897354512719411 | 8.07634515885684 | -10.2932728951866 | 9.7295613260153 | 2.99588267196263e-06 |
| AI506816 | ENSMUSG00000105987 | 433855 | 0.964671997642731 | 4.34152307691377 | 10.276559674804 | 9.9210507576037 | 3.04626420874624e-06 |
| Nes | ENSMUSG00000004891 | 18008 | 1.13770181826364 | 6.60277148941734 | 10.2711259932632 | 9.9841685691955 | 3.05705732856797e-06 |
| Khdrbs3 | ENSMUSG00000022332 | 13992 | -0.878692332849565 | 5.16829069950432 | -10.2604680915481 | 1.0109217626656 | 3.08095610406441e-06 |
| Vat1 | ENSMUSG00000034993 | 26949 | 0.926560156195264 | 4.19883374853789 | 10.2596748242429 | 1.0118591541113 | 3.08095610406441e-06 |
| Mb | ENSMUSG00000018893 | 17189 | -0.911010172337472 | 11.4399643437103 | -10.2568829134668 | 1.0151656865450 | 3.08243781100672e-06 |
| Igdcc4 | ENSMUSG00000032816 | 56741 | 0.785963836124696 | 5.8977706682626 | 10.2441898579217 | 1.0303442756088 | 3.11985963342935e-06 |
| Ptprc | ENSMUSG00000026395 | 19264 | 1.72816197380999 | 2.96400117707241 | 10.2362632475634 | 1.0399456229876 | 3.13428649722127e-06 |
| Fn1 | ENSMUSG00000026193 | 14268 | 2.03713673142502 | 8.93935624063367 | 10.2355260000872 | 1.0408434713121 | 3.13428649722127e-06 |
| Tgfb2 | ENSMUSG00000039239 | 21808 | 1.1550991318738 | 4.15998265128061 | 10.2282935771902 | 1.0496953315312 | 3.15225815081552e-06 |
| Bdh1 | ENSMUSG00000046598 | 71911 | -1.08392336350552 | 7.22776861787937 | -10.2243777048349 | 1.0545214730303 | 3.15352414530289e-06 |
| Paip2b | ENSMUSG00000045896 | 232164 | -0.822959091394029 | 5.25847361056222 | -10.2232733785609 | 1.0558867781363 | 3.15352414530289e-06 |
| St3gal6 | ENSMUSG00000022747 | 54613 | -0.849827544869224 | 4.35337061439012 | -10.2182104086216 | 1.0621704135894 | 3.16364708200168e-06 |
| Plin4 | ENSMUSG00000002831 | 57435 | -0.923187645006003 | 7.34104739592722 | -10.2152166165714 | 1.0659047408372 | 3.16614258752512e-06 |
| Coro6 | ENSMUSG00000020836 | 216961 | -0.632066481845531 | 7.82906115901703 | -10.2065485421294 | 1.0767960510408 | 3.18982591705362e-06 |
| Sfrp4 | ENSMUSG00000021319 | 20379 | 3.15992315768623 | 2.2933915189627 | 10.2035943159767 | 1.0805350441276 | 3.19225096415126e-06 |
| Mcu | ENSMUSG00000009647 | 215999 | 0.750155812898912 | 5.47725863298141 | 10.1995907876997 | 1.0856241761378 | 3.19864093513829e-06 |
| Ube2b | ENSMUSG00000020390 | 22210 | -0.698044829424445 | 6.8998086492533 | -10.1772294473132 | 1.1145228706616 | 3.27039183743291e-06 |
| Ckap4 | ENSMUSG00000046841 | 216197 | 1.3706165239771 | 4.40159556848291 | 10.176133406264 | 1.1159602555696 | 3.27039183743291e-06 |
| Jpt1 | ENSMUSG00000020737 | 15374 | 0.750201163519978 | 3.42328479100331 | 10.1536396458853 | 1.1458998685653 | 3.34915279767054e-06 |
| Fbn1 | ENSMUSG00000027204 | 14118 | 1.11543762105892 | 7.85003897813707 | 10.1450505032371 | 1.1575570181010 | 3.37420153729657e-06 |
| Lbp | ENSMUSG00000016024 | 16803 | 1.4130848338585 | 3.30410388040711 | 10.1404242860407 | 1.1638880104298 | 3.38363293670457e-06 |
| P3h2 | ENSMUSG00000038168 | 210530 | 1.30773778233985 | 5.17561713659808 | 10.1324011340605 | 1.1749553745291 | 3.40674726763334e-06 |
| Arhgap10 | ENSMUSG00000037148 | 78514 | -0.85891180910351 | 5.35648165212987 | -10.1197034709353 | 1.1927007163586 | 3.44905066944875e-06 |
| Chrng | ENSMUSG00000026253 | 11449 | 4.41083160006177 | -0.273183721747016 | 10.1045755817021 | 1.2142159757706 | 3.50200391323182e-06 |
| Igf1 | ENSMUSG00000020053 | 16000 | 1.37055987515974 | 5.0960707746583 | 10.1022188976229 | 1.2176047764906 | 3.5025362662683e-06 |
| Ndufb9 | ENSMUSG00000022354 | 66218 | -0.711574282027379 | 7.74308096322795 | -10.0947273762193 | 1.2284443395426 | 3.52444227704473e-06 |
| Alas1 | ENSMUSG00000032786 | 11655 | -0.878813345600846 | 6.07392578974539 | -10.0799280013645 | 1.2501611007660 | 3.5773588985533e-06 |
| Acsl1 | ENSMUSG00000018796 | 14081 | -0.778053449755939 | 7.88916404592122 | -10.0764985636289 | 1.2552516491639 | 3.582547200264e-06 |
| Nt5c1a | ENSMUSG00000054958 | 230718 | -1.99127680849517 | 1.13017699844594 | -10.0736102087848 | 1.2595561671688 | 3.58547095399013e-06 |
| Serping1 | ENSMUSG00000023224 | 12258 | 1.09215289231999 | 5.95850056566756 | 10.0566897854261 | 1.2850907228606 | 3.64865628352987e-06 |
| Pxmp2 | ENSMUSG00000029499 | 19301 | -1.14563214838309 | 3.43242507717255 | -10.0537507744524 | 1.2895819226002 | 3.65192227874188e-06 |
| Akap1 | ENSMUSG00000018428 | 11640 | -0.853804879652476 | 6.2161008517827 | -10.03775892677 | 1.3143144720529 | 3.7069247707109e-06 |
| Arfgap3 | ENSMUSG00000054277 | 66251 | 1.05684544033227 | 3.34005757259258 | 10.0368169150036 | 1.3157870378152 | 3.7069247707109e-06 |
| Oat | ENSMUSG00000030934 | 18242 | -0.708047239586266 | 5.64097283687145 | -10.0209639393826 | 1.3408341482209 | 3.76414432365921e-06 |
| Ppm1k | ENSMUSG00000037826 | 243382 | -1.31849178655866 | 4.02394711319566 | -10.0196177000825 | 1.3429844353006 | 3.76414432365921e-06 |
| Egln3 | ENSMUSG00000035105 | 112407 | -1.2043663856531 | 5.75857041296283 | -10.0092981710984 | 1.3595898873449 | 3.80094042418607e-06 |
| Cyth1 | ENSMUSG00000017132 | 19157 | -0.752218182974111 | 4.86166173668328 | -10.0029905782766 | 1.3698472318916 | 3.81984696219569e-06 |
| Slc44a1 | ENSMUSG00000028412 | 100434 | 0.708192973758938 | 4.90560334616724 | 9.99389517350323 | 1.3847835103187 | 3.84624265264393e-06 |
| Mcee | ENSMUSG00000033429 | 73724 | -1.02457676221077 | 4.32340468025672 | -9.99294709309591 | 1.3863503843579 | 3.84624265264393e-06 |
| Mvp | ENSMUSG00000030681 | 78388 | 0.713869920224748 | 6.27462140709455 | 9.9872136737874 | 1.3958662363093 | 3.86283894407527e-06 |
| Mtmr11 | ENSMUSG00000045934 | 194126 | 1.4851646544298 | 2.40786153251563 | 9.98418325863311 | 1.4009239572587 | 3.86704539818074e-06 |
| Gmpr | ENSMUSG00000000253 | 66355 | -0.897934305583486 | 5.0554959546256 | -9.97762071739589 | 1.4119437940016 | 3.88764675371087e-06 |
| Pabpc1 | ENSMUSG00000022283 | 18458 | 0.617416581448219 | 6.41891739129712 | 9.97177747077471 | 1.4218335290326 | 3.90504078036592e-06 |
| 1810014B01Rik | ENSMUSG00000097412 | 66263 | -1.13507203526337 | 2.63994510099333 | -9.96905576447009 | 1.4264652056679 | 3.90794264740759e-06 |
| Sdha | ENSMUSG00000021577 | 66945 | -0.791890895369519 | 8.51796118423453 | -9.95915760834683 | 1.4434454387574 | 3.93807328545755e-06 |
| Egln1 | ENSMUSG00000031987 | 112405 | -0.766003389189276 | 7.53138238245607 | -9.95799183208029 | 1.4454594517089 | 3.93807328545755e-06 |
| Klhl13 | ENSMUSG00000036782 | 67455 | 1.34446908482104 | 3.52488962727899 | 9.95636707986294 | 1.4482713939748 | 3.93807328545755e-06 |
| Grn | ENSMUSG00000034708 | 14824 | 0.96571032591146 | 4.82796721688111 | 9.95250346415404 | 1.4549815482212 | 3.94060131506666e-06 |
| Glis3 | ENSMUSG00000052942 | 226075 | 1.74651996417709 | 1.59544984106746 | 9.95168282279507 | 1.4564110614645 | 3.94060131506666e-06 |
| Sod3 | ENSMUSG00000072941 | 20657 | 0.901555690451879 | 4.41994316254347 | 9.93754858756029 | 1.4812683936998 | 3.99796168186e-06 |
| Fstl1 | ENSMUSG00000022816 | 14314 | 1.31365843103123 | 7.01704031188569 | 9.93288140597137 | 1.4895753337234 | 4.00168032723112e-06 |
| Itpripl2 | ENSMUSG00000095115 | 319622 | 0.84838112599719 | 4.91731550459009 | 9.93266154204151 | 1.4899678832522 | 4.00168032723112e-06 |
| Ltbp3 | ENSMUSG00000024940 | 16998 | 0.927088829471917 | 6.00255654791327 | 9.92385425088097 | 1.5057835829611 | 4.03424518268339e-06 |
| Soat1 | ENSMUSG00000026600 | 20652 | 1.39017383655493 | 2.42764707010476 | 9.91870575734994 | 1.5151117467063 | 4.04931210348345e-06 |
| Hk2 | ENSMUSG00000000628 | 15277 | -0.803464408157375 | 8.25526864461724 | -9.91295847875871 | 1.5255975697222 | 4.06427925617479e-06 |
| Man2a1 | ENSMUSG00000024085 | 17158 | 0.881543381434231 | 4.07421904154802 | 9.91156685271262 | 1.5281481788380 | 4.06427925617479e-06 |
| Arfgef1 | ENSMUSG00000067851 | 211673 | -0.623104997697129 | 6.91893650714204 | -9.90673811478823 | 1.5370337048301 | 4.07798918143168e-06 |
| St8sia5 | ENSMUSG00000025425 | 225742 | -0.938020993215658 | 3.70412627967003 | -9.90168984888228 | 1.5463820688994 | 4.09285772279402e-06 |
| Mmp19 | ENSMUSG00000025355 | 58223 | 2.27844544591077 | 1.50326551041331 | 9.87567782101727 | 1.5955207383619 | 4.21271429735126e-06 |
| Cspg4 | ENSMUSG00000032911 | 121021 | 1.08907348269194 | 5.18956842444913 | 9.87180230871788 | 1.6029831026062 | 4.2222188661661e-06 |
| Slc7a1 | ENSMUSG00000041313 | 11987 | 1.04223741721993 | 3.29805306333296 | 9.86624866993775 | 1.6137416085847 | 4.24033882774997e-06 |
| Idh2 | ENSMUSG00000030541 | 269951 | -0.871553832625976 | 9.30179787994177 | -9.86356127549033 | 1.6189752184469 | 4.24388923569389e-06 |
| Spon1 | ENSMUSG00000038156 | 233744 | 1.70638946162495 | 2.11893300254878 | 9.85825265980055 | 1.6293667757606 | 4.25215280794742e-06 |
| Anxa7 | ENSMUSG00000021814 | 11750 | 0.744979846160566 | 5.11647175138996 | 9.85797734970819 | 1.6299076265025 | 4.25215280794742e-06 |
| Mettl11b | ENSMUSG00000040113 | 240879 | -2.85652600024234 | 1.70065554943255 | -9.84410692828802 | 1.6574053322460 | 4.31359468732889e-06 |
| Slc2a4 | ENSMUSG00000018566 | 20528 | -0.830162580291303 | 7.08157204390169 | -9.82910659704981 | 1.6877001099011 | 4.37668585938629e-06 |
| Loxl1 | ENSMUSG00000032334 | 16949 | 1.51510871891781 | 5.346691508632 | 9.828148712773 | 1.6896545903037 | 4.37668585938629e-06 |
| Paqr8 | ENSMUSG00000025931 | 74229 | 1.78603252144075 | 1.40155242878304 | 9.81864249222431 | 1.7091824205724 | 4.40110771962583e-06 |
| Alox12 | ENSMUSG00000000320 | 11684 | 1.47609759936742 | 2.67983562750762 | 9.81809074160456 | 1.7103231920242 | 4.40110771962583e-06 |
| Lyz1 | ENSMUSG00000069515 | 17110 | 2.21911339761892 | 1.67132651122769 | 9.8176854674407 | 1.7111616328249 | 4.40110771962583e-06 |
| Piezo2 | ENSMUSG00000041482 | 667742 | 1.71478327095005 | 2.62466431730041 | 9.80978996583256 | 1.7275836799463 | 4.43291483697027e-06 |
| C1qtnf9 | ENSMUSG00000071347 | 239126 | -0.969604555322533 | 3.90748893525335 | -9.80180396757237 | 1.7443648280349 | 4.46171793607686e-06 |
| Mmp14 | ENSMUSG00000000957 | 17387 | 1.46902835867058 | 4.49552741057391 | 9.80057050604413 | 1.7469721678171 | 4.46171793607686e-06 |
| Tceal5 | ENSMUSG00000054034 | 331532 | 0.979071271608534 | 3.38309741192509 | 9.79449505808011 | 1.7598754050769 | 4.48419535032535e-06 |
| Fcgr3 | ENSMUSG00000059498 | 14131 | 1.85996933563565 | 2.23725755360264 | 9.79065758335588 | 1.7680778387706 | 4.4946183385121e-06 |
| Dpyd | ENSMUSG00000033308 | 99586 | -1.63878593713309 | 1.26371479674738 | -9.78287421967805 | 1.7848397454222 | 4.52670145178905e-06 |
| Arhgap30 | ENSMUSG00000048865 | 226652 | 1.50588300316128 | 1.90093646045307 | 9.77532300587923 | 1.8012635362751 | 4.5577804895889e-06 |
| Galnt5 | ENSMUSG00000026828 | 241391 | 2.83203943947025 | 2.37271486419035 | 9.76757075563692 | 1.8182921380635 | 4.57587367937984e-06 |
| Isca1 | ENSMUSG00000044792 | 69046 | -0.741152790476463 | 6.71073223023273 | -9.7673075137393 | 1.8188733768926 | 4.57587367937984e-06 |
| Ndufs8 | ENSMUSG00000059734 | 225887 | -0.741046863451259 | 5.98835371124226 | -9.76540875685444 | 1.8230717030497 | 4.57587367937984e-06 |
| Tmem94 | ENSMUSG00000020747 | 71947 | -0.710412799947393 | 5.28138657092444 | -9.76446663381706 | 1.8251586535629 | 4.57587367937984e-06 |
| Aldh1a1 | ENSMUSG00000053279 | 11668 | -1.16883662691997 | 5.43167469348254 | -9.7569514302811 | 1.8418973887595 | 4.60727239279865e-06 |
| Tgfbi | ENSMUSG00000035493 | 21810 | 0.876840753168965 | 4.52074158677498 | 9.75117778626588 | 1.8548681488562 | 4.6291241404446e-06 |
| Rab40b | ENSMUSG00000025170 | 217371 | 1.89861670355591 | 1.97283857913394 | 9.74068299006428 | 1.8786949684792 | 4.67790767663922e-06 |
| Mrps18a | ENSMUSG00000023967 | 68565 | -0.621820489929729 | 4.88555112772295 | -9.72352564637249 | 1.9183528554377 | 4.76579887790685e-06 |
| Cox6a2 | ENSMUSG00000030785 | 12862 | -0.935490997352867 | 8.79981646962106 | -9.7179449137441 | 1.9314436840509 | 4.78744011572805e-06 |
| Nectin3 | ENSMUSG00000022656 | 58998 | 0.999985447068646 | 2.35961166257738 | 9.70928187401782 | 1.9519535450318 | 4.82733126713645e-06 |
| Rgs10 | ENSMUSG00000030844 | 67865 | 1.66794537388601 | 1.44934845187385 | 9.7053689537429 | 1.9612933597383 | 4.83601219513027e-06 |
| C2cd2 | ENSMUSG00000045975 | 207781 | -0.616779206400693 | 5.26098828693663 | -9.70410854696426 | 1.9643119702111 | 4.83601219513027e-06 |
| Mymk | ENSMUSG00000009214 | 66139 | 5.39655482608958 | 0.393702988057036 | 9.69858461786346 | 1.9775999773731 | 4.85778547250927e-06 |
| Mrpl14 | ENSMUSG00000023939 | 68463 | -0.912826141934862 | 3.83155176264184 | -9.68818560075092 | 2.0028755993206 | 4.90884151932162e-06 |
| Mgat2 | ENSMUSG00000043998 | 217664 | 0.856515025653709 | 3.62292453068758 | 9.6862434211974 | 2.0076342251649 | 4.90949658059923e-06 |
| Pitpnc1 | ENSMUSG00000040430 | 71795 | -0.724771897750327 | 6.04947049359706 | -9.67538284486028 | 2.0344670816346 | 4.96400885476535e-06 |
| Macf1 | ENSMUSG00000028649 | 11426 | 0.988912446064318 | 8.31812865005744 | 9.66305891020195 | 2.0653786611885 | 5.02820805021211e-06 |
| L2hgdh | ENSMUSG00000020988 | 217666 | -0.967187339936709 | 3.61489668912468 | -9.65151653231782 | 2.0947837255953 | 5.08846242321845e-06 |
| Gpx1 | ENSMUSG00000063856 | 14775 | 0.914500050820792 | 4.64826002906949 | 9.64633487303909 | 2.1081290731385 | 5.10878500531966e-06 |
| Ugp2 | ENSMUSG00000001891 | 216558 | -0.89398519000768 | 8.36673347769599 | -9.64425207945155 | 2.1135187831052 | 5.10878500531966e-06 |
| Ndufs3 | ENSMUSG00000005510 | 68349 | -0.687624097852122 | 6.3245186431201 | -9.64149912795636 | 2.1206652054406 | 5.10878500531966e-06 |
| Acot13 | ENSMUSG00000006717 | 66834 | -1.0266859389431 | 5.20094211665147 | -9.64104572910019 | 2.1218446550316 | 5.10878500531966e-06 |
| Lrrc39 | ENSMUSG00000027961 | 109245 | -0.873317421616182 | 5.83592113882151 | -9.6345662164458 | 2.1387766731784 | 5.13823468450853e-06 |
| Lrp1 | ENSMUSG00000040249 | 16971 | 1.21272477095561 | 8.09217440694881 | 9.629686535871 | 2.1516230185882 | 5.14975465538492e-06 |
| Cxcl14 | ENSMUSG00000021508 | 57266 | 1.53584548173023 | 4.99941359785958 | 9.62916755473644 | 2.1529941245182 | 5.14975465538492e-06 |
| Mllt11 | ENSMUSG00000053192 | 56772 | 0.868345530076371 | 5.07170789536836 | 9.6065136330075 | 2.2137599227551 | 5.28353923922184e-06 |
| Mccc1 | ENSMUSG00000027709 | 72039 | -0.605278577960508 | 4.48279957514839 | -9.60295252704429 | 2.2234769794489 | 5.28413372008359e-06 |
| C1qtnf6 | ENSMUSG00000022440 | 72709 | 1.60769187803221 | 2.19096333327313 | 9.60210930327735 | 2.2257844956737 | 5.28413372008359e-06 |
| Mest | ENSMUSG00000051855 | 17294 | 1.88602165824103 | 2.59803138186137 | 9.60030792462225 | 2.2307225877078 | 5.28413372008359e-06 |
| Ecm1 | ENSMUSG00000028108 | 13601 | 1.04342827111812 | 4.13686959807162 | 9.59561966057097 | 2.2436292193477 | 5.28413372008359e-06 |
| Aqp4 | ENSMUSG00000024411 | 11829 | -2.6704501209533 | 2.6043581442297 | -9.59528461185787 | 2.2445546360968 | 5.28413372008359e-06 |

| | | | | | | | |
|---|---|---|---|---|---|---|---|
| Nup133 | ENSMUSG00000039509 | 234865 | -1.12474853577842 | 4.16216258087084 | -9.59523748275438 | 2.244684840996€ | 5.28413372008359e-06 |
| Galnt17 | ENSMUSG00000034040 | 212996 | 1.11000099742117 | 4.03737643355177 | 9.59409359081472 | 2.2478475709805 | 5.28413372008359e-06 |
| Spi1 | ENSMUSG00000002111 | 20375 | 1.82865647997096 | 1.34805076372415 | 9.59102153902645 | 2.2563649457379 | 5.29277365276003e-06 |
| Ubash3b | ENSMUSG00000032020 | 72828 | 1.60166578395653 | 1.35513656779811 | 9.58696952984056 | 2.2676518749026 | 5.30202932497751e-06 |
| Fndc3b | ENSMUSG00000039286 | 72007 | 0.859726585079157 | 5.27780685841943 | 9.5861250878152 | 2.2700116403705 | 5.30202932497751e-06 |
| F13a1 | ENSMUSG00000039109 | 74145 | 1.07026986432628 | 4.34379104926553 | 9.58341884414678 | 2.2775917808364 | 5.30839141925862e-06 |
| Itga7 | ENSMUSG00000025348 | 16404 | 1.00329983487686 | 6.08552141672602 | 9.56828374886594 | 2.3204847055398 | 5.39685496090546e-06 |
| Plxna1 | ENSMUSG00000030084 | 18844 | 0.545882698400737 | 4.99322502758001 | 9.56512546614555 | 2.3295432733611 | 5.40641985586216e-06 |
| Rtn4 | ENSMUSG00000020458 | 68585 | 0.767442485721656 | 8.26418117528719 | 9.54947115440478 | 2.3750020206622 | 5.50024302708886e-06 |
| Syk | ENSMUSG00000021457 | 20963 | 1.14508677539277 | 2.65531419179354 | 9.5396716856192 | 2.4039387353945 | 5.55548717052799e-06 |
| Zdhhc23 | ENSMUSG00000036304 | 332175 | -1.02053058068854 | 5.19991461457068 | -9.53508081518217 | 2.4176239458155 | 5.57532644550834e-06 |
| Ltbp2 | ENSMUSG00000002020 | 16997 | 2.93264315069205 | 3.39750614277125 | 9.52331549337362 | 2.4530759642836 | 5.63605032462732e-06 |
| Slc16a1 | ENSMUSG00000032902 | 20501 | -0.770254840641277 | 5.97913695997702 | -9.52237693778706 | 2.4559278172050 | 5.63605032462732e-06 |
| Dock7 | ENSMUSG00000028556 | 67299 | 0.832791483196533 | 4.03997267375227 | 9.52122802357227 | 2.4594236619222 | 5.63605032462732e-06 |
| Pvalb | ENSMUSG00000005716 | 19293 | -1.70707804245894 | 6.0989786623219 | -9.5118282378251 | 2.4882244884555 | 5.69012173290955e-06 |
| Nsmf | ENSMUSG00000006476 | 56876 | 0.860315400421587 | 3.28892311161737 | 9.50413275804736 | 2.5120706174610 | 5.73266052598464e-06 |
| Mfap5 | ENSMUSG00000030116 | 50530 | 1.80896107747642 | 3.9680194082403 | 9.50062297377741 | 2.5230271228149 | 5.7456686415604e-06 |
| Iscu | ENSMUSG00000025825 | 66383 | -0.605721466635884 | 5.52986040673173 | -9.49087736279092 | 2.5537176097845 | 5.80346927080148e-06 |
| Angptl1 | ENSMUSG00000033544 | 72713 | 1.69023823073419 | 3.72844251993167 | 9.48708321257398 | 2.5657732535418 | 5.81876917727511e-06 |
| Prg4 | ENSMUSG00000006014 | 96875 | 2.09151317172222 | 6.02847997836622 | 9.46737486356676 | 2.6293773691004 | 5.94570230855603e-06 |
| Nab2 | ENSMUSG00000025402 | 17937 | 1.36908186359986 | 2.84056879713451 | 9.46638290144927 | 2.6326227402260 | 5.94570230855603e-06 |
| Clic4 | ENSMUSG00000037242 | 29876 | 1.0102767668269 | 6.66335374265553 | 9.46364331077127 | 2.6416079409555 | 5.95369410362587e-06 |
| Dnm3os | ENSMUSG00000078190 | 474332 | 1.97811391132698 | 1.96721033480604 | 9.45548750970676 | 2.6685507825119 | 6.00204292255939e-06 |
| Rtn4ip1 | ENSMUSG00000019864 | 170728 | -0.76088945907453 | 3.98064813513416 | -9.45381214825027 | 2.6741214981602 | 6.0022221963839e-06 |
| Mpst | ENSMUSG00000071711 | 246221 | -1.03716285407756 | 2.81101216254635 | -9.44436169221157 | 2.7057777647927 | 6.05190231815945e-06 |
| Slc25a3 | ENSMUSG00000061904 | 18674 | -0.813731858286181 | 8.54788589517011 | -9.44390192296395 | 2.7073279970542 | 6.05190231815945e-06 |
| Axl | ENSMUSG00000002602 | 26362 | 0.884549206460541 | 5.98035564468903 | 9.43364622092383 | 2.7421546520191 | 6.10620718416606e-06 |
| Dym | ENSMUSG00000035765 | 69190 | -0.582538082607841 | 4.97907799071644 | -9.43345935033607 | 2.7427936395805 | 6.10620718416606e-06 |
| Nampt | ENSMUSG00000020572 | 59027 | -0.691406886996334 | 6.32495044203972 | -9.42800194357273 | 2.7615248342434 | 6.13541218762506e-06 |
| Dmac2 | ENSMUSG00000057229 | 66349 | -0.906322998035607 | 3.77581501009508 | -9.40105840248128 | 2.8560184318190 | 6.33248224710228e-06 |
| Inpp5d | ENSMUSG00000026288 | 16331 | 1.28135694948786 | 2.4003397165178 | 9.39846153354547 | 2.8653060360745 | 6.34021463164601e-06 |
| Lamc2 | ENSMUSG00000026479 | 16782 | 1.21805999014615 | 2.36357878134508 | 9.38289717926996 | 2.9216493059203 | 6.44724215674865e-06 |
| Iqgap2 | ENSMUSG00000021676 | 544963 | 1.15355187924661 | 3.00015133569796 | 9.38080197368394 | 2.9293235808881 | 6.44724215674865e-06 |
| Zswim4 | ENSMUSG00000035671 | 212168 | 0.712099130823152 | 3.68350156706862 | 9.38024455134207 | 2.9313689066911 | 6.44724215674865e-06 |
| Pdp2 | ENSMUSG00000048371 | 382051 | -1.27468437364789 | 2.41652861011546 | -9.37590009705727 | 2.9473620041327 | 6.46940041509548e-06 |
| Dnajc28 | ENSMUSG00000039763 | 246738 | -0.959516513551996 | 3.86494655281014 | -9.36604889040159 | 2.9839719580779 | 6.53663276027064e-06 |
| Acot1 | ENSMUSG00000072949 | 26897 | -2.20669572868619 | 0.831994941406673 | -9.35904635687831 | 3.0102895140901 | 6.58109493570396e-06 |
| Akr1b10 | ENSMUSG00000061758 | 67861 | -0.905033295695468 | 3.77967316452822 | -9.33842414581085 | 3.0892387367178 | 6.74021329961333e-06 |
| Plbd1 | ENSMUSG00000030214 | 66857 | -1.15143949199385 | 4.10883797979815 | -9.33472845304838 | 3.1036185166563 | 6.75809840748409e-06 |
| Hibadh | ENSMUSG00000029776 | 58875 | -0.750313304382814 | 5.15632529582024 | -9.33202550830683 | 3.1141806473519 | 6.76761603503056e-06 |
| Timp1 | ENSMUSG00000001131 | 21857 | 2.7849260432757 | 1.73458073545329 | 9.32606004342393 | 3.1376271322667 | 6.79770490878549e-06 |
| Ipo13 | ENSMUSG00000033365 | 230673 | -0.644123219572748 | 6.39763222838223 | -9.32453421403415 | 3.1436543258588 | 6.79770490878549e-06 |
| Lum | ENSMUSG00000036446 | 17022 | 1.35408510511965 | 4.78780136394189 | 9.32376877349621 | 3.1466825394249 | 6.79770490878549e-06 |
| S100a13 | ENSMUSG00000042312 | 20196 | 1.33396487502252 | 2.85599180320873 | 9.31564764348309 | 3.1790029684454 | 6.85398056175088e-06 |
| Fam57b | ENSMUSG00000058966 | 68952 | -2.40869523743758 | 1.26903943293519 | -9.30404754071774 | 3.2257830668148 | 6.93537739629332e-06 |
| Adi1 | ENSMUSG00000020629 | 104923 | -0.942591465980269 | 3.11328321601741 | -9.30314688178349 | 3.2294457000396 | 6.93537739629332e-06 |
| Pdgfra | ENSMUSG00000029231 | 18595 | 0.873263929481449 | 4.99938277339441 | 9.29847580297229 | 3.2485122989881 | 6.96264469416449e-06 |
| Cpd | ENSMUSG00000020841 | 12874 | 0.690629804215408 | 6.08323326465387 | 9.28887341399949 | 3.2880849536252 | 7.03367057300924e-06 |
| Dpp3 | ENSMUSG00000063904 | 75221 | 0.768783147124418 | 3.74653240725028 | 9.27598303039584 | 3.3420162042469 | 7.13507404855928e-06 |
| Itga5 | ENSMUSG00000000555 | 16402 | 1.2796789646282 | 4.82774190623845 | 9.26670658803351 | 3.3814093089985 | 7.2051043190377e-06 |
| Cdkn1a | ENSMUSG00000023067 | 12575 | 1.54458607636476 | 4.14879091128232 | 9.26121756139831 | 3.4049512856496 | 7.24115223802264e-06 |
| Dlg1 | ENSMUSG00000022770 | 13383 | -0.546224250106013 | 5.24798672661311 | -9.25280871658599 | 3.4413549490274 | 7.29220168104621e-06 |
| Thbs3 | ENSMUSG00000028047 | 21827 | 1.73962431640187 | 3.5994625871588 | 9.25259211570672 | 3.4422981130910 | 7.29220168104621e-06 |
| Idh3g | ENSMUSG00000002010 | 15929 | -0.797715528776489 | 6.78632596780949 | -9.24912413195568 | 3.4574365681431 | 7.30142652748472e-06 |
| Plaat3 | ENSMUSG00000060675 | 225845 | -0.747450725200935 | 6.73682625105624 | -9.24853578580342 | 3.4600118390239 | 7.30142652748472e-06 |
| Bicc1 | ENSMUSG00000014329 | 83675 | 1.01666102301607 | 4.4677776851771 | 9.24166361598915 | 3.4902438581183 | 7.351031910037e-06 |
| Sec24d | ENSMUSG00000039234 | 69608 | 1.06176415941252 | 3.89740530482275 | 9.23062004278503 | 3.5394169052785 | 7.44005636013771e-06 |
| Rhot2 | ENSMUSG00000025733 | 214952 | -0.64322542445988 | 5.32534154005687 | -9.22912613770097 | 3.5461251153890 | 7.44005636013771e-06 |
| Uqcrc1 | ENSMUSG00000025651 | 22273 | -0.814987658506448 | 8.58939428579953 | -9.22247928489082 | 3.5761364514940 | 7.48864895618415e-06 |
| Ankrd40 | ENSMUSG00000020864 | 71452 | -0.546442813875014 | 6.83684969389495 | -9.21594762353646 | 3.6058910466831 | 7.53651912644227e-06 |
| Celf2 | ENSMUSG00000002107 | 14007 | 0.568170506096178 | 5.07229335491908 | 9.19908436035565 | 3.6839351103456 | 7.66584177046536e-06 |
| Ppp1r9b | ENSMUSG00000038976 | 217124 | 0.728928100768554 | 5.50560751976505 | 9.19766309649112 | 3.6905944100259 | 7.66584177046536e-06 |
| Auh | ENSMUSG00000021460 | 11992 | -0.600093252386642 | 4.72618216685804 | -9.19674490791988 | 3.6949033700097 | 7.66584177046536e-06 |
| Coq9 | ENSMUSG00000031782 | 67914 | -0.796051128511469 | 7.01966407384589 | -9.19639075451573 | 3.6965668037702 | 7.66584177046536e-06 |
| Narf | ENSMUSG00000000056 | 67608 | -0.616755315636671 | 5.75164601077988 | -9.19505861702435 | 3.7028308981847 | 7.66584177046536e-06 |
| Ndufv2 | ENSMUSG00000024099 | 72900 | -0.773841861823479 | 7.57645600457253 | -9.18497441323328 | 3.7506176009081 | 7.75009470614882e-06 |
| Kcnn1 | ENSMUSG00000002908 | 84036 | -0.909727023127139 | 3.09419874784215 | -9.14734381042614 | 3.9348143461595 | 8.09008930502664e-06 |
| Atp9a | ENSMUSG00000027546 | 11981 | 0.801318742606734 | 5.03020827393887 | 9.14712626497094 | 3.9359067617899 | 8.09008930502664e-06 |
| Cstb | ENSMUSG00000005054 | 13014 | 1.24526041491304 | 2.13597801319387 | 9.14683709274199 | 3.9373593543812 | 8.09008930502664e-06 |
| Spcs3 | ENSMUSG00000054408 | 76687 | 0.544832251067663 | 4.8960438916935 | 9.14166082558407 | 3.9634578782840 | 8.12843490947894e-06 |
| Nt5c2 | ENSMUSG00000025041 | 76952 | 0.986310157571846 | 5.13372608634327 | 9.12265464431266 | 4.0608757261485 | 8.30746525034012e-06 |
| Fndc1 | ENSMUSG00000071984 | 68655 | 1.37420536359763 | 6.67269421471402 | 9.12167728564467 | 4.0659536263214 | 8.30746525034012e-06 |
| Sdhd | ENSMUSG00000000171 | 66925 | -0.697490165818364 | 6.20004258225869 | -9.11658758738666 | 4.0925067163125 | 8.3395799641183e-06 |
| Trps1 | ENSMUSG00000038679 | 83925 | 0.581421579235547 | 4.34635146545665 | 9.11574310095069 | 4.0969302358121 | 8.3395799641183e-06 |
| Mmp3 | ENSMUSG00000043613 | 17392 | 5.67592186210871 | -1.37137408943719 | 9.10884750768503 | 4.1332410366833 | 8.39785460445 83e-06 |
| Sh3rf3 | ENSMUSG00000037990 | 237353 | 1.60966394503288 | 0.822605097641845 | 9.09794787920275 | 4.1913361663487 | 8.50009195442628e-06 |
| Tnfrsf23 | ENSMUSG00000037613 | 79201 | 1.74130104409254 | 1.78960817280893 | 9.09453273565791 | 4.2097171251990 | 8.52155886954651e-06 |
| Mrc2 | ENSMUSG00000020695 | 17534 | 1.3630836170432 | 4.89026160308754 | 9.08878258957872 | 4.2408594923910 | 8.55896924858679e-06 |
| Utrn | ENSMUSG00000019820 | 22288 | 0.615416104700105 | 7.12945989700288 | 9.0882312893905 | 4.2438581399085 | 8.55896924858679e-06 |
| Pcp4l1 | ENSMUSG00000038370 | 66425 | -0.980885784343265 | 6.26226369611011 | -9.08273653513388 | 4.2738689100497 | 8.60362082058089e-06 |
| Ndufs1 | ENSMUSG00000025968 | 227197 | -0.723866016116492 | 7.82820068023967 | -9.0700566239153 | 4.3439877473699 | 8.72870037987158e-06 |
| Pfn1 | ENSMUSG00000018293 | 18643 | 0.595767830181303 | 5.65217707975847 | 9.06582790010928 | 4.3676434613116 | 8.76013039919093e-06 |
| Amot | ENSMUSG00000041688 | 27494 | 0.728489520768798 | 7.07082920129626 | 9.06223388376353 | 4.3878562929894 | 8.78455258949955e-06 |
| Ctsc | ENSMUSG00000030560 | 13032 | 1.1628783806031 | 4.0766070489615 | 9.0607332597945 | 4.3963252568287 | 8.78541707173579e-06 |
| Ghr | ENSMUSG00000055737 | 14600 | -0.713856714223223 | 5.70799409601551 | -9.0564151837077 | 4.4207919998740 | 8.81818929755902e-06 |
| Tep1 | ENSMUSG00000006281 | 21745 | 0.900075681030787 | 3.26019181471466 | 9.04377283745106 | 4.4932613607014 | 8.94641893148043e-06 |
| Timp2 | ENSMUSG00000017466 | 21858 | 1.00136637606037 | 6.21572674835005 | 9.03667192422227 | 4.5345189077291 | 9.01215021461593e-06 |
| Pde3b | ENSMUSG00000030671 | 18576 | 1.21461028949245 | 2.35048119768871 | 9.03301610146353 | 4.5559168057284 | 9.02808640599253e-06 |
| Hspa9 | ENSMUSG00000024359 | 15526 | -0.743981975893602 | 8.46983349021606 | -9.03248137267467 | 4.5590556180659 | 9.02808640599253e-06 |
| Mccc2 | ENSMUSG00000021646 | 78038 | -0.883986159869295 | 3.8054286485523 | -9.02831664604514 | 4.5835810557189 | 9.06023951538233e-06 |
| Ndufa8 | ENSMUSG00000026895 | 68375 | -0.709869973864709 | 6.17461527549351 | -9.02417342024222 | 4.6081191442927 | 9.09230151015588e-06 |
| Dap | ENSMUSG00000039168 | 223453 | 0.874204923430677 | 3.43368248413551 | 9.02171211932195 | 4.6227622120002 | 9.10475923231978e-06 |
| Ptgfrn | ENSMUSG00000027864 | 19221 | 1.25809308581672 | 4.72289467567683 | 9.01101942735882 | 4.6869532974137 | 9.2145839018038e-06 |
| Drp2 | ENSMUSG00000000223 | 13497 | 0.932699366475361 | 3.56040788204403 | 9.00675643444154 | 4.7128088686044 | 9.24878164142102e-06 |
| Pon2 | ENSMUSG00000032667 | 330260 | 0.697318520404107 | 4.40959405274314 | 9.00395330858162 | 4.7298927946278 | 9.26567350144745e-06 |
| Acss1 | ENSMUSG00000027452 | 68738 | -1.0645097797378 | 5.0490891780595 | -8.99518322664429 | 4.7837695825333 | 9.35445175432419e-06 |
| Rab3gap2 | ENSMUSG00000039318 | 98732 | 0.633148362048962 | 4.86500240072892 | 8.98814898047627 | 4.8274542596143 | 9.4230183056865e-06 |
| Ube2r2 | ENSMUSG00000036241 | 67615 | -0.544623308090275 | 5.74138675031053 | -8.98549266597533 | 4.8440608982872 | 9.43857926545064e-06 |
| Ano5 | ENSMUSG00000055489 | 233246 | -0.839255356977094 | 5.70346682434425 | -8.98108105206761 | 4.8717756097520 | 9.47569024736652e-06 |
| B3galt2 | ENSMUSG00000033849 | 26878 | 1.62697129350291 | 3.89069527860295 | 8.97391529041539 | 4.9171524400293 | 9.54696151366985e-06 |
| Zeb2 | ENSMUSG00000026872 | 24136 | 0.651614076550309 | 5.8142546093532 | 8.96978527802989 | 4.9435095092779 | 9.57324869452442e-06 |
| Serpinb6a | ENSMUSG00000060147 | 20719 | 1.11102473264035 | 7.03939239409811 | 8.96905160207156 | 4.9482074031710 | 9.57324869452442e-06 |
| Rorc | ENSMUSG00000028150 | 19885 | -0.862397214283809 | 5.23079666319204 | -8.9649743532486 | 4.9744015931126 | 9.59562869870423e-06 |
| Atl2 | ENSMUSG00000059811 | 56298 | -0.637349081851459 | 5.82696179958623 | -8.96451966149145 | 4.9773318746366 | 9.59562869870423e-06 |
| Me1 | ENSMUSG00000032418 | 17436 | -0.712213168262277 | 5.44505215486971 | -8.95973401235919 | 5.0082847625111 | 9.62712540891095e-06 |
| Chrna2 | ENSMUSG00000022041 | 110902 | -6.11982948832555 | -1.43580951227495 | -8.95927199594565 | 5.0112838328335 | 9.62712540891095e-06 |
| Mrpl15 | ENSMUSG00000033845 | 27395 | -0.616707057030816 | 5.01743313831737 | -8.95283363305982 | 5.0532762451515 | 9.69076537469322e-06 |
| Coq5 | ENSMUSG00000041733 | 52064 | -0.722733224056318 | 5.62377748461903 | -8.94579578364038 | 5.0996070055542 | 9.76248759679745e-06 |
| Myh13 | ENSMUSG00000060180 | 544791 | -1.15414778988767 | 3.84766172866679 | -8.93797022307628 | 5.1516541456159 | 9.84488312337906e-06 |
| Tnfrsf21 | ENSMUSG00000023915 | 94185 | 0.754302257954079 | 3.88738447027387 | 8.92008517421992 | 5.2727383346614 | 1.00586915769955e-05 |
| Tns3 | ENSMUSG00000020422 | 319939 | 0.914568017073643 | 4.62913040486656 | 8.91230564689383 | 5.3263478136196 | 1.01432592248565e-05 |
| Naaa | ENSMUSG00000029413 | 67111 | 1.14779685320513 | 2.83846660848021 | 8.89295135024287 | 5.4622502490587 | 1.0383975212602e-05 |
| C1qtnf3 | ENSMUSG00000058914 | 81799 | 4.58708769161992 | 2.49721435826985 | 8.89045429998593 | 5.4800507959776 | 1.03997283421583e-05 |
| Fbxo3 | ENSMUSG00000027180 | 57443 | -0.551871824024189 | 5.57879125895348 | -8.88048276914139 | 5.5517513708678 | 1.05065664427748e-05 |
| Vim | ENSMUSG00000026728 | 22352 | 1.01736179328013 | 7.66256211696221 | 8.8792018288256 | 5.5610340456826 | 1.05065664427748e-05 |
| Adgre1 | ENSMUSG00000004730 | 13733 | 1.92338802109487 | 2.34249492505352 | 8.87862998414352 | 5.5651833961820 | 1.05065664427748e-05 |

| | | | | | | | |
|---|---|---|---|---|---|---|---|
| Cers6 | ENSMUSG00000027035 | 241447 | 1.57822343579538 | 2.06603359397616 | 8.86681308601235 | 5.6516685879541 | 1.06514464370564e-05 |
| Uqcrb | ENSMUSG00000021520 | 67530 | -0.759883023404424 | 7.34732611943245 | -8.86309777974314 | 5.6791544057404 | 1.06848256125901e-05 |
| P2rx7 | ENSMUSG00000029468 | 18439 | 1.26194949507169 | 1.78416064747398 | 8.86047190952079 | 5.6986662673814 | 1.07031135685131e-05 |
| Tnip2 | ENSMUSG00000059866 | 231130 | 1.15221045366066 | 2.34492630256671 | 8.85270194136228 | 5.7568205528263 | 1.07937916746046e-05 |
| Dpt | ENSMUSG00000026574 | 56429 | 1.21471648940993 | 4.41342476571955 | 8.84424270852552 | 5.8208516149924 | 1.08951590759387e-05 |
| Aco2 | ENSMUSG00000022477 | 11429 | -0.767050372232365 | 9.54641114831146 | -8.83995869048889 | 5.8535673206186 | 1.09376657062704e-05 |
| Dglucy | ENSMUSG00000021185 | 217830 | -1.7553171142232 | 2.45010715258455 | -8.82957261930302 | 5.9336965568581 | 1.10684704885694e-05 |
| Gpihbp1 | ENSMUSG00000022579 | 68453 | -1.09660313450843 | 3.58594359711189 | -8.82284609304186 | 5.9862136545175 | 1.11474108104823e-05 |
| Cox5a | ENSMUSG00000000088 | 12858 | -0.732765235246789 | 6.86493430031028 | -8.80455040078845 | 6.1315717173699 | 1.13986752453352e-05 |
| Nmt2 | ENSMUSG00000026643 | 18108 | 1.04920336591077 | 2.74606215030795 | 8.79196231331474 | 6.2337589917854 | 1.15689676637024e-05 |
| Slc9a2 | ENSMUSG00000026062 | 226999 | -0.637733457735808 | 5.10995543856914 | -8.78662736770308 | 6.2776116185941 | 1.16306055259071e-05 |
| Asb11 | ENSMUSG00000031382 | 68854 | -0.965655699505191 | 5.30422044473635 | -8.78126492379411 | 6.3220208423169 | 1.16930642685899e-05 |
| Clpb | ENSMUSG00000001829 | 20480 | -0.66123322488008 | 5.23797082988318 | -8.77727312461129 | 6.3552957719596 | 1.173475305461e-05 |
| Uqcrfs1 | ENSMUSG00000038462 | 66694 | -0.716656203815018 | 7.49709049128006 | -8.77234841790006 | 6.3966039294978 | 1.17911091995516e-05 |
| Fh1 | ENSMUSG00000026526 | 14194 | -0.668522538537221 | 6.74396407323048 | -8.76907138881969 | 6.4242495157433 | 1.18221332418503e-05 |
| Sorl1 | ENSMUSG00000049313 | 20660 | -0.665133254119512 | 5.5142979356089 | -8.7634149629275 | 6.4722672976534 | 1.18904796354033e-05 |
| Adam8 | ENSMUSG00000025473 | 11501 | 2.41694679053813 | 0.816031659926118 | 8.7616327206906 | 6.4874757845768 | 1.18984224498673e-05 |
| Sox11 | ENSMUSG00000063632 | 20666 | 4.90601696236048 | 0.73243847995751 | 8.74866962277046 | 6.5992436289656 | 1.208313770657e-05 |
| Mipep | ENSMUSG00000021993 | 70478 | -0.679058299058266 | 4.21343982011703 | -8.74523885872198 | 6.6291648931295 | 1.21176256599998e-05 |
| Prkaa1 | ENSMUSG00000050697 | 105787 | 0.637187630250005 | 4.38514956577283 | 8.74360427103167 | 6.6434715602299 | 1.21235037771075e-05 |
| Btg1 | ENSMUSG00000036478 | 12226 | 0.774702326279391 | 3.89438267384268 | 8.73878653051471 | 6.6858298550689 | 1.21704444865557e-05 |
| Cav3 | ENSMUSG00000062694 | 12391 | 0.843360803722655 | 4.4051957919953 | 8.73814838430583 | 6.6914620221571 | 1.21704444865557e-05 |
| Slc25a4 | ENSMUSG00000031633 | 11739 | -0.842439709586024 | 10.9535517461635 | -8.73214049350859 | 6.7447344748394 | 1.22469588944302e-05 |
| Iffo1 | ENSMUSG00000038271 | 320678 | 0.938254830643323 | 3.2209944329782 | 8.72270877512055 | 6.8292774524026 | 1.23799057764865e-05 |
| Pcca | ENSMUSG00000041650 | 110821 | -0.881507458396233 | 4.14180756470862 | -8.71904749909387 | 6.8623987794390 | 1.2419351168551e-05 |
| Cox5b | ENSMUSG00000061518 | 12859 | -0.638257643828499 | 7.78190865214244 | -8.71583367298823 | 6.8916129947127 | 1.2434915802729e-05 |
| Ndufa5 | ENSMUSG00000023089 | 68202 | -0.759557166399028 | 6.63918308338173 | -8.71559906623317 | 6.8937507789349 | 1.2434915802729e-05 |
| Olfml2b | ENSMUSG00000038463 | 320078 | 1.18456062357719 | 3.70506556559113 | 8.71339251086084 | 6.9138918746843 | 1.24507005077717e-05 |
| Grsf1 | ENSMUSG00000044221 | 231413 | -0.6377378788965 | 6.44856880086737 | -8.71070315928802 | 6.9385244461493 | 1.24588785403949e-05 |
| Peg10 | ENSMUSG00000092035 | 170676 | 2.31937310476668 | 0.916211436903227 | 8.71040853671965 | 6.9412286443147 | 1.24588785403949e-05 |
| Itga9 | ENSMUSG00000039115 | 104099 | 0.989231056016409 | 3.93163511180566 | 8.68571843766447 | 7.1718752844526 | 1.28517653662871e-05 |
| Pcyt1a | ENSMUSG00000005615 | 13026 | -0.524448601121718 | 5.93840998278664 | -8.67241660764257 | 7.2994998582510 | 1.30590561293851e-05 |
| Gsta4 | ENSMUSG00000032348 | 14860 | -1.07423675801804 | 3.03032926754626 | -8.66627803357654 | 7.3592082467747 | 1.31443636185448e-05 |
| Scn4b | ENSMUSG00000046480 | 399548 | -0.877195886255177 | 7.24784063347229 | -8.65433810248101 | 7.4768363853037 | 1.33326751268769e-05 |
| Ostn | ENSMUSG00000052276 | 239790 | -1.0560870468122 | 3.77843102244588 | -8.65198816478484 | 7.5002215513463 | 1.33525931234148e-05 |
| Abca1 | ENSMUSG00000015243 | 11303 | 0.893822158606936 | 4.93433193116307 | 8.63077634971491 | 7.7148593626789 | 1.37123784867387e-05 |
| Gabbr1 | ENSMUSG00000024462 | 54393 | 1.23283406645179 | 3.23615055278181 | 8.62834791525255 | 7.7398453264101 | 1.37126733858956e-05 |
| Adamts14 | ENSMUSG00000059901 | 237360 | 1.544217487556 | 1.07521626767264 | 8.62832177078871 | 7.7401147919655 | 1.37126733858956e-05 |
| Cebpa | ENSMUSG00000034957 | 12606 | 1.41072366366871 | 2.51070762369847 | 8.62547393838262 | 7.7695265655797 | 1.37425072634874e-05 |
| Ptms | ENSMUSG00000030122 | 69202 | 0.700847033910076 | 5.55362266575941 | 8.62062088892513 | 7.8199216877575 | 1.38092995103196e-05 |
| Mrpl55 | ENSMUSG00000036860 | 67212 | -0.616951069042057 | 4.05433697741277 | -8.61647153592076 | 7.8632848762751 | 1.38634785455748e-05 |
| Dio2 | ENSMUSG00000007682 | 13371 | 2.28979805650259 | 3.97867453486311 | 8.60270618860758 | 8.0089798150803 | 1.4077549773942e-05 |
| Slc25a19 | ENSMUSG00000020744 | 67283 | -0.634033889439114 | 3.81755844028133 | -8.60256752606711 | 8.0104619516896 | 1.4077549773942e-05 |
| Sntb2 | ENSMUSG00000041308 | 20650 | 0.850754413000914 | 4.28863259973041 | 8.59569338688364 | 8.0843047884226 | 1.41765549467704e-05 |
| Ank | ENSMUSG00000022265 | 11732 | -0.502912379431795 | 6.73223726882518 | -8.59491271085118 | 8.0927365170475 | 1.41765549467704e-05 |
| Ehd4 | ENSMUSG00000027293 | 98878 | 0.918638236332003 | 5.65555874145337 | 8.58594224657237 | 8.1902956571633 | 1.43244994925525e-05 |
| Mrps9 | ENSMUSG00000060679 | 69527 | -0.608489564626444 | 4.8516301229973 | -8.58195988115031 | 8.2340059866328 | 1.43779424025373e-05 |
| Abhd5 | ENSMUSG00000032540 | 67469 | -0.561228712594421 | 4.38111622180886 | -8.57959504061274 | 8.2600796216008 | 1.44004673594448e-05 |
| Fmc1 | ENSMUSG00000019689 | 66117 | -0.885219761404574 | 3.74256614874453 | -8.57168450270576 | 8.3479368265456 | 1.45281420992e-05 |
| Net1 | ENSMUSG00000021215 | 56349 | 0.843986681245947 | 5.46303103765674 | 8.57061463532798 | 8.3598951426189 | 1.45281420992e-05 |
| Tnnc2 | ENSMUSG00000017300 | 21925 | -0.852098893137121 | 10.9259364047817 | -8.56362514157556 | 8.4384685131369 | 1.46414125900159e-05 |
| Slc25a11 | ENSMUSG00000014606 | 67863 | -0.691763673386572 | 6.34322460159355 | -8.55859442907528 | 8.4955072775962 | 1.47170190255792e-05 |
| Rmnd1 | ENSMUSG00000019763 | 66084 | -0.757251418330849 | 4.34836397345062 | -8.55601341220122 | 8.5249299541831 | 1.47446217292999e-05 |
| Pfkm | ENSMUSG00000033065 | 18642 | -0.620420110028229 | 9.29468647757686 | -8.54008005947525 | 8.7089777559801 | 1.50391525830362e-05 |
| 2310016D03Rik | ENSMUSG00000102098 | 69566 | -3.42891011343936 | 0.179655194311134 | -8.5386300744596 | 8.7259349531324 | 1.50446679767651e-05 |
| Fam89b | ENSMUSG00000024939 | 17826 | 0.69860396445862 | 2.94812168790933 | 8.52565271427685 | 8.8792725935678 | 1.52849336567386e-05 |
| Mdh2 | ENSMUSG00000019179 | 17448 | -0.666240697442779 | 8.85994898353278 | -8.52211990302844 | 8.9215092991931 | 1.53334934197295e-05 |
| Wdfy4 | ENSMUSG00000051506 | 545030 | 1.32424295092047 | 2.38369711402106 | 8.5193331382745 | 8.9549771496854 | 1.53668532532514e-05 |
| Lifr | ENSMUSG00000054263 | 16880 | -0.637586691793255 | 5.80986020811735 | -8.51683405111817 | 8.9851035356592 | 1.5394383502867e-05 |
| Smyd2 | ENSMUSG00000026603 | 226830 | -0.929298535851743 | 4.2193112924499 | -8.50743978114793 | 9.0993174757799 | 1.55656712562991e-05 |
| Klhl23 | ENSMUSG00000042155 | 277396 | -0.934501659183948 | 4.27417599922821 | -8.49961314984029 | 9.1956497839165 | 1.57058824668737e-05 |
| Pyurf | ENSMUSG00000043162 | 66459 | -0.661294797790883 | 4.61651913290889 | -8.49775459605408 | 9.2186841372147 | 1.57206608898432e-05 |
| Pygm | ENSMUSG00000032648 | 19309 | -0.792575247106202 | 9.55501381345189 | -8.49459358570466 | 9.2580015997795 | 1.57264301799072e-05 |
| Erbin | ENSMUSG00000021709 | 59079 | 0.686654460012138 | 7.01599686674622 | 8.4944705864193 | 9.2595350917861 | 1.57264301799072e-05 |
| Endog | ENSMUSG00000015337 | 13804 | -0.804781437735891 | 3.90015256793776 | -8.49326961019249 | 9.2745224316372 | 1.57264301799072e-05 |
| Cnst | ENSMUSG00000038949 | 226744 | -0.676514808008256 | 5.27022413688211 | -8.49286197531285 | 9.2796152831764 | 1.57264301799072e-05 |
| Mlec | ENSMUSG00000048578 | 109154 | 0.695505723165267 | 5.91938266521534 | 8.48694458192658 | 9.3538806196534 | 1.58277506274662e-05 |
| Kif21a | ENSMUSG00000022629 | 16564 | -0.480467545549168 | 5.65520410373908 | -8.48476020656358 | 9.3814545762748 | 1.58498732570727e-05 |
| Lipe | ENSMUSG00000003123 | 16890 | -0.668500361183333 | 3.8587725370132 | -8.48331264846385 | 9.3997750398495 | 1.58563180494752e-05 |
| Anxa5 | ENSMUSG00000027712 | 11747 | 0.838786286956157 | 5.06135940428988 | 8.48106481002388 | 9.4282992941233 | 1.58799290576367e-05 |
| Cacng1 | ENSMUSG00000020722 | 12299 | 0.750260196877489 | 5.16358545070106 | 8.47968315727617 | 9.4458775711215 | 1.58850596507583e-05 |
| Cib2 | ENSMUSG00000037493 | 56506 | -1.50153651899371 | 1.51500599342537 | -8.47567711480638 | 9.4970420676491 | 1.59283140032749e-05 |
| Ttll12 | ENSMUSG00000016757 | 223723 | 1.16919844415094 | 2.61152921677462 | 8.4752400759839 | 9.5026416330161 | 1.59283140032749e-05 |
| Ppara | ENSMUSG00000022383 | 19013 | -1.82131323995402 | 4.01885179493995 | -8.47425207229089 | 9.5153133694433 | 1.59283140032749e-05 |
| Uprt | ENSMUSG00000073016 | 331487 | 0.732934113831904 | 3.60468542849925 | 8.47108743590499 | 9.5560226545582 | 1.59720005561126e-05 |
| Nans | ENSMUSG00000028334 | 94181 | 0.792233843030021 | 2.79469614274239 | 8.46729250757672 | 9.6050839230626 | 1.60294919638165e-05 |
| Dgat2 | ENSMUSG00000030747 | 67800 | -1.31830388198995 | 5.47760131758328 | -8.45791168237954 | 9.7275117772947 | 1.61929543261732e-05 |
| Gcdh | ENSMUSG00000003809 | 270076 | -0.791390127257172 | 3.65031542392125 | -8.45746242943218 | 9.7334163412266 | 1.61929543261732e-05 |
| Sema3c | ENSMUSG00000028780 | 20348 | 0.658229213115844 | 5.13256617256798 | 8.45464449729632 | 9.7705395425698 | 1.61929543261732e-05 |
| Dnajc15 | ENSMUSG00000022013 | 66148 | -0.886434124787804 | 4.06253096581089 | -8.45459161914272 | 9.7712375904261 | 1.61929543261732e-05 |
| Fndc5 | ENSMUSG00000001334 | 384061 | -0.846014570490469 | 6.76364691543382 | -8.45414756343804 | 9.7771016881111 | 1.61929543261732e-05 |
| Coro1a | ENSMUSG00000030707 | 12721 | 1.62552895381123 | 2.15881598839806 | 8.44021449135091 | 9.9630099941426 | 1.64758944396329e-05 |
| Bgn | ENSMUSG00000031375 | 12111 | 1.62157257132591 | 7.02709140759737 | 8.43108440981982 | 1.0086865443241 | 1.66555175468382e-05 |
| Ppic | ENSMUSG00000024538 | 19038 | 1.04083678372519 | 3.91288920941139 | 8.42662829238591 | 1.0147908317922 | 1.67310385857028e-05 |
| Nfkb2 | ENSMUSG00000025225 | 18034 | 0.858310580330618 | 4.19231781588252 | 8.42381536308227 | 1.0186643571046 | 1.67461947351954e-05 |
| Trim35 | ENSMUSG00000022043 | 66854 | 0.639912426318389 | 6.56148625981332 | 8.42373584650446 | 1.0187740827833 | 1.67461947351954e-05 |
| Hivep3 | ENSMUSG00000028634 | 16656 | 1.18775321069787 | 2.52504537006572 | 8.42201219076503 | 1.0211556587572 | 1.67601388977113e-05 |
| Carhsp1 | ENSMUSG00000008393 | 52502 | 1.15611436477458 | 4.54743576799697 | 8.41852512058203 | 1.0259918413685 | 1.6814268092953e-05 |
| Hspb8 | ENSMUSG00000041548 | 80888 | 0.648662448113303 | 7.529343245488 | 8.41433727277296 | 1.0318320885203 | 1.68846655084071e-05 |
| Ndufaf4 | ENSMUSG00000028261 | 68493 | -0.563459659921438 | 4.76576322662608 | -8.40751862232391 | 1.0414168282450 | 1.70160349021625e-05 |
| Sh3d19 | ENSMUSG00000028082 | 27059 | 1.10717298833826 | 4.35980504556944 | 8.40057291454144 | 1.0512774227629 | 1.71515126988384e-05 |
| Evi2a | ENSMUSG00000078771 | 14017 | 1.4750751036296 | 1.91568405177867 | 8.38698503554225 | 1.0708554875571 | 1.74448902153315e-05 |
| Snrpd3 | ENSMUSG00000020180 | 67332 | -0.611542801710293 | 4.43054182680021 | -8.37592419032961 | 1.0870782546141 | 1.76752089470705e-05 |
| Pygo1 | ENSMUSG00000034910 | 72135 | -0.654747094728849 | 4.1694388468165 | -8.37514723065836 | 1.0882275749134 | 1.76752089470705e-05 |
| Tmem119 | ENSMUSG00000054675 | 231633 | 1.81778434102702 | 1.13912203203674 | 8.36901797163469 | 1.0973396603535 | 1.77967653224406e-05 |
| Mapkapk2 | ENSMUSG00000016528 | 17164 | -0.666212141254102 | 7.79504445385423 | -8.35981965859125 | 1.1111666820034 | 1.79943155570061e-05 |
| Rpl3l | ENSMUSG00000002500 | 66211 | -0.84636962569072 | 7.24972361858039 | -8.35711105127236 | 1.1152734586245 | 1.80341038109835e-05 |
| Chrna1 | ENSMUSG00000027107 | 11435 | 0.678332333379287 | 4.82631847114281 | 8.34545926081698 | 1.1331244796682 | 1.82956923002264e-05 |
| Gpc1 | ENSMUSG00000034220 | 14733 | 0.590947573413676 | 7.78330534900804 | 8.33960798435836 | 1.1422030563119 | 1.84150761187987e-05 |
| Epas1 | ENSMUSG00000024140 | 13819 | -0.798972050558764 | 7.68326833780682 | -8.33225884035202 | 1.1537151912150 | 1.85732853537142e-05 |
| Chchd3 | ENSMUSG00000053768 | 66075 | -0.754564435949316 | 7.24961717604653 | -8.32605905758992 | 1.1635226850356 | 1.87036271619476e-05 |
| Sucla2 | ENSMUSG00000022110 | 20916 | -0.708982661542537 | 7.5515373609552 | -8.32497524377366 | 1.1652462492287 | 1.87038278271946e-05 |
| Mt2 | ENSMUSG00000031762 | 17750 | 2.97911143021135 | 2.31991802410652 | 8.3238542380018 | 1.1670318106380 | 1.87050215866346e-05 |
| Chrnd | ENSMUSG00000026251 | 11447 | 1.54563390391211 | 2.91243271589328 | 8.32123964533625 | 1.1712076852722 | 1.87365906553123e-05 |
| Hook1 | ENSMUSG00000028572 | 77963 | -0.753154921015646 | 3.27666396326203 | -8.32047645010371 | 1.1724296046321 | 1.87365906553123e-05 |
| Slc7a5 | ENSMUSG00000040010 | 20539 | 1.17259809070735 | 1.94022644676896 | 8.31793911516882 | 1.1765017612890 | 1.87742200768627e-05 |
| Dhrs7c | ENSMUSG00000033044 | 68460 | -1.10388854717226 | 4.05672219701986 | -8.2974585574712 | 1.2099257870956 | 1.92540498568262e-05 |
| Ctsk | ENSMUSG00000028111 | 13038 | 1.75292782464616 | 2.6579732979967 | 8.29657205284211 | 1.2113951214399 | 1.92540498568262e-05 |
| Rrbp1 | ENSMUSG00000027422 | 81910 | 0.721930699512447 | 6.18086420002265 | 8.29629486411015 | 1.2118549356414 | 1.92540498568262e-05 |
| Cd63 | ENSMUSG00000025351 | 12512 | 0.602897454114379 | 5.66178292065127 | 8.29005957477357 | 1.2222474440461 | 1.93909823089526e-05 |
| Fam131a | ENSMUSG00000050821 | 78408 | -1.31907752678541 | 2.64034294579068 | -8.28559810204369 | 1.2297415258266 | 1.94816008968276e-05 |
| Rab11fip3 | ENSMUSG00000037098 | 215445 | -0.454203870731383 | 7.14536624926172 | -8.28427634716888 | 1.2319710740075 | 1.94886770043068e-05 |
| Cat | ENSMUSG00000027187 | 12359 | -0.635158920288686 | 6.90934029130128 | -8.28070870197807 | 1.2380104570147 | 1.95559137364569e-05 |
| Rai14 | ENSMUSG00000022246 | 75646 | 1.15219367726077 | 3.60939237373812 | 8.27499564808317 | 1.2477471752552 | 1.96756694024036e-05 |
| Pgm2 | ENSMUSG00000025791 | 72157 | -0.930516853589941 | 6.55488463710467 | -8.27415209563899 | 1.2491917084684 | 1.96756694024036e-05 |
| Rasgrp3 | ENSMUSG00000071042 | 240168 | -0.971423919546384 | 4.17345725115865 | -8.26616546563063 | 1.2629566236190 | 1.98638544644313e-05 |

| | | | | | | | |
|---|---|---|---|---|---|---|---|
| Tgfbr1 | ENSMUSG00000007613 | 21812 | 0.628083921411176 | 4.46067126342042 | 8.26187314464957 | 1.2704208433551 | 1.995254344643e-05 |
| Pgam2 | ENSMUSG00000020475 | 56012 | -0.628379926079656 | 8.78517205140849 | -8.25227865296222 | 1.2872752012809 | 2.018824279082e-05 |
| Thrb | ENSMUSG00000021779 | 21834 | -0.830562528704541 | 4.47540416576754 | -8.240115714551 | 1.3089833111927 | 2.04951908458179e-05 |
| Ndufv1 | ENSMUSG00000037916 | 17995 | -0.700063989950539 | 7.54561266481174 | -8.23922000482583 | 1.3105972373274 | 2.04951908458179e-05 |
| Txlnb | ENSMUSG00000039891 | 378431 | -0.639906489351244 | 9.16586389113831 | -8.23443388088563 | 1.3192569345792 | 2.05579074043136e-05 |
| Gstk1 | ENSMUSG00000029864 | 76263 | -0.747218133181029 | 4.13057330741283 | -8.23412101082702 | 1.3198251315354 | 2.05579074043136e-05 |
| Ddr1 | ENSMUSG00000003534 | 12305 | 0.787728074502743 | 3.62031113387541 | 8.23388724429068 | 1.3202498397061 | 2.05579074043136e-05 |
| Adck1 | ENSMUSG00000021044 | 72113 | -0.795346332903763 | 3.13631128972693 | -8.22999839172318 | 1.3273364420060 | 2.06388544062133e-05 |
| Ndufc1 | ENSMUSG00000037152 | 66377 | -0.800791471221961 | 5.86722829907935 | -8.22013079260426 | 1.3454998177480 | 2.08912501384407e-05 |
| 6530402F18Rik | ENSMUSG00000079499 | 76220 | 1.00075440666985 | 2.86955338149473 | 8.21911149079175 | 1.3473910298783 | 2.08912501384407e-05 |
| Snrpn | ENSMUSG00000102252 | 20646 | -0.81256306929586 | 4.26467069453204 | -8.21390228317012 | 1.3571003375193 | 2.09860239109941e-05 |
| Fam171b | ENSMUSG00000048388 | 241520 | 2.10902360175553 | 1.89288069283801 | 8.21377247199793 | 1.3573432353007 | 2.09860239109941e-05 |
| Phkb | ENSMUSG00000036879 | 102093 | -0.67901929733608 | 5.75749856481863 | -8.20410761237267 | 1.3755579837296 | 2.12376049719515e-05 |
| Golim4 | ENSMUSG00000034109 | 73124 | 0.629893157006213 | 4.86438499418312 | 8.20243173672011 | 1.3787427008542 | 2.12567510056949e-05 |
| Mrpl51 | ENSMUSG00000030335 | 66493 | -0.511886481638406 | 5.24473686660785 | -8.1932407319078 | 1.3963483276411 | 2.14735076057795e-05 |
| Tent5a | ENSMUSG00000032265 | 212943 | 0.861186359344023 | 2.92574722041543 | 8.19229618223798 | 1.3981711096745 | 2.14735076057795e-05 |
| Nras | ENSMUSG00000027852 | 18176 | 0.661572122413292 | 3.91505938776833 | 8.19186897815054 | 1.3989963527086 | 2.14735076057795e-05 |
| Camk1d | ENSMUSG00000039145 | 227541 | 1.3102705547737 | 1.85177123170867 | 8.19100873635817 | 1.4006596764176 | 2.14735076057795e-05 |
| Gm15543 | ENSMUSG00000086863 | 105242817 | -4.08626047486739 | -0.918822461376218 | -8.18837889900913 | 1.4057576345273 | 2.14921156411118e-05 |
| Gm30794 | ENSMUSG00000117545 | 102632818 | -1.9450485553978 | 0.911029181672125 | -8.18835412493127 | 1.4058057527577 | 2.14921156411118e-05 |
| Bckdha | ENSMUSG00000060376 | 12039 | -0.831861877397876 | 5.31836897121124 | -8.18582100871565 | 1.4107350145371 | 2.15295563094357e-05 |
| Rab31 | ENSMUSG00000056515 | 106572 | 1.36798358696191 | 3.65382549087274 | 8.18507308109554 | 1.4121939322903 | 2.15295563094357e-05 |
| Nmnat1 | ENSMUSG00000028992 | 66454 | -0.694134382573276 | 3.76496964710621 | -8.18107859775346 | 1.4200127799652 | 2.1589213261869e-05 |
| Ankrd1 | ENSMUSG00000024803 | 107765 | 1.64702430770807 | 3.68449460613925 | 8.18061385724068 | 1.4209254470209 | 2.1589213261869e-05 |
| Cd82 | ENSMUSG00000027215 | 12521 | 1.02469183528902 | 2.51622434998506 | 8.17907362621149 | 1.4239546369552 | 2.1589213261869e-05 |
| Far1 | ENSMUSG00000030759 | 67420 | 0.635246669600019 | 3.90886286248747 | 8.17893027053451 | 1.4242369245557 | 2.1589213261869e-05 |
| Glg1 | ENSMUSG00000003316 | 20340 | 0.423942350739257 | 6.25422174867042 | 8.17804460113599 | 1.4259822500292 | 2.1589213261869e-05 |
| Vgll3 | ENSMUSG00000091243 | 73569 | 1.20315552920346 | 2.51979771194483 | 8.17605658281411 | 1.4299081654528 | 2.16152195781429e-05 |
| Stard7 | ENSMUSG00000027367 | 99138 | -0.70730385939001 | 6.68843270205871 | -8.17440946473568 | 1.4331695623274 | 2.16152195781429e-05 |
| Atf5 | ENSMUSG00000038539 | 107503 | -0.650852351961864 | 4.85173955501672 | -8.1741761104602 | 1.4336322563492 | 2.16152195781429e-05 |
| Dab2 | ENSMUSG00000022150 | 13132 | 1.11339218155665 | 4.40507002696454 | 8.17211647732305 | 1.4377229694392 | 2.16470382630034e-05 |
| Rabgef1 | ENSMUSG00000025340 | 56715 | -0.525803309025358 | 5.04521967802138 | -8.16879951809916 | 1.4443369593566 | 2.17167088070527e-05 |
| Sema3f | ENSMUSG00000034684 | 20350 | 0.73070367670513 | 3.55536076323973 | 8.16325331622394 | 1.4554682655734 | 2.18429536215512e-05 |
| Smco1 | ENSMUSG00000046345 | 69576 | -2.13347479255525 | 2.36428395158799 | -8.16225988220844 | 1.4574716958916 | 2.18429536215512e-05 |
| Svip | ENSMUSG00000074093 | 75744 | -0.871382336224087 | 3.56817009722409 | -8.16068378605299 | 1.4606561702513 | 2.18429536215512e-05 |
| Kcna7 | ENSMUSG00000038201 | 16495 | -0.68317279775763 | 5.03020411543917 | -8.16064912166169 | 1.4607262919544 | 2.18429536215512e-05 |
| ATP8 | ENSMUSG00000064356 | 17706 | -0.788295334060463 | 10.3944457719988 | -8.14944899764525 | 1.4835708320395 | 2.2154252411236e-05 |
| Asap3 | ENSMUSG00000036995 | 230837 | 1.03725893705051 | 1.84435368378419 | 8.14338367903937 | 1.4961000204282 | 2.23027840040669e-05 |
| Tubb5 | ENSMUSG00000001525 | 22154 | 0.820357165821054 | 5.58905455742653 | 8.14187183013053 | 1.4992405083984 | 2.23027840040669e-05 |
| Pf4 | ENSMUSG00000029373 | 56744 | 1.24530640801375 | 2.16985080205098 | 8.14168057138763 | 1.4996382986908 | 2.23027840040669e-05 |
| Cs | ENSMUSG00000005683 | 12974 | -0.592512417143748 | 8.48165789833118 | -8.13366136327766 | 1.5164182708983 | 2.25216958141172e-05 |
| Oma1 | ENSMUSG00000035069 | 67013 | -0.651483695405986 | 3.37925646220105 | -8.12906829299408 | 1.5261187653839 | 2.26350125161632e-05 |
| Fastk | ENSMUSG00000028959 | 66587 | -0.553585836960521 | 5.90951982363257 | -8.12476527716617 | 1.5352664128335 | 2.27398335483518e-05 |
| Anks1 | ENSMUSG00000024219 | 224650 | -0.720318808927427 | 4.13077895365673 | -8.11975217180996 | 1.5459970337851 | 2.28630113655673e-05 |
| 5930412G12Rik | ENSMUSG00000072591 | 319616 | 1.04050631487604 | 3.26578706642615 | 8.11802322508808 | 1.5497162973805 | 2.28630113655673e-05 |
| Ndufs7 | ENSMUSG00000020153 | 75406 | -0.691938780332316 | 7.02875938126089 | -8.11795771069926 | 1.5498574166943 | 2.28630113655673e-05 |
| Bex1 | ENSMUSG00000050071 | 19716 | 1.29125089815933 | 3.63741737687704 | 8.10296863792896 | 1.5825053423059 | 2.33131615859121e-05 |
| Slc39a14 | ENSMUSG00000022094 | 213053 | 1.1397639457552 | 3.4074426365025 | 8.10152389165259 | 1.5856904931364 | 2.33286443882558e-05 |
| Atp5f1 | ENSMUSG00000000563 | 11950 | -0.650750738158719 | 7.92764408727377 | -8.09805470612059 | 1.5933666157501 | 2.34100678451139e-05 |
| Cd248 | ENSMUSG00000056481 | 70445 | 0.75430969320572 | 4.77835058068521 | 8.09490804617987 | 1.6003631960448 | 2.34813021422369e-05 |
| Nfatc2 | ENSMUSG00000027544 | 18019 | 0.748351610378664 | 5.16141727167908 | 8.09335347959869 | 1.6038317868902 | 2.35006504859208e-05 |
| Csf2ra | ENSMUSG00000059326 | 12982 | 1.60454607912955 | 1.07520147683779 | 8.09209381298283 | 1.6066482379670 | 2.35104041354986e-05 |
| Ms4a7 | ENSMUSG00000024672 | 109225 | 2.43216008755683 | 0.434781149203057 | 8.0904299387962 | 1.6103764893226 | 2.3533456423511e-05 |
| Chpf2 | ENSMUSG00000038181 | 100910 | 0.93875157568075 | 2.97195141769767 | 8.08683684889919 | 1.6184588878061 | 2.36007840483439e-05 |
| Igfbp5 | ENSMUSG00000026185 | 16011 | 0.825061526430239 | 7.94436188523598 | 8.08646322012551 | 1.6193018055308 | 2.36007840483439e-05 |
| Dipk2a | ENSMUSG00000045414 | 68861 | 0.712762037220999 | 3.62741113689753 | 8.08435390678745 | 1.6240692234542 | 2.3638749243114e-05 |
| Kif16b | ENSMUSG00000038844 | 16558 | -0.680207022778474 | 6.16409170941749 | -8.07953469480831 | 1.6350174446460 | 2.37664570311521e-05 |
| Klhl7 | ENSMUSG00000028986 | 52323 | -0.630304655130168 | 4.26492989812934 | -8.07857893073234 | 1.6371980246467 | 2.37665492794332e-05 |
| Sox6 | ENSMUSG00000051910 | 20679 | -0.808891461928256 | 5.5046811748387 | -8.07677056918679 | 1.6413322541182 | 2.37949640182574e-05 |
| Mrpl42 | ENSMUSG00000062981 | 67270 | -0.678567385715148 | 6.40856804185177 | -8.07100189955202 | 1.6545946092423 | 2.39554618193748e-05 |
| Sestd1 | ENSMUSG00000042272 | 228071 | 0.555122475169805 | 4.07790249742835 | 8.06866563557294 | 1.6599980524541 | 2.40019030573764e-05 |
| Vamp5 | ENSMUSG00000073002 | 53620 | -0.730532120826017 | 3.88252371639393 | -8.0634864702874 | 1.6720435712439 | 2.41441324666671e-05 |
| Pcolce | ENSMUSG00000029718 | 18542 | 1.08678880705029 | 4.83788177648869 | 8.06166451972579 | 1.6763030143915 | 2.41737048157178e-05 |
| Medag | ENSMUSG00000029659 | 70717 | 1.24604746375031 | 4.09223786482296 | 8.04867655001886 | 1.7070023707869 | 2.4583982760306e-05 |
| Slc30a4 | ENSMUSG00000005802 | 22785 | 0.666433816833855 | 4.59399324965537 | 8.04591953791315 | 1.7135954569545 | 2.46464630789081e-05 |
| Aldh2 | ENSMUSG00000029455 | 11669 | -0.489092652831107 | 6.3000143450189 | -8.03482690151441 | 1.7403964609431 | 2.49990456170423e-05 |
| Lrrc52 | ENSMUSG00000040485 | 240899 | -1.88131807728376 | 2.19245719382669 | -8.03183308355919 | 1.7477057830871 | 2.50710917518707e-05 |
| Ahnak | ENSMUSG00000069833 | 66395 | 0.672507041496003 | 9.98044383687548 | 8.02738868278149 | 1.7586167655975 | 2.51945476602192e-05 |
| Ro60 | ENSMUSG00000018199 | 20822 | 0.611722444557321 | 3.79871097075977 | 8.02308365846475 | 1.7692544821388 | 2.53137706076697e-05 |
| Fabp4 | ENSMUSG00000062515 | 11770 | -0.908058841985005 | 6.43801339841813 | -8.01324755809797 | 1.7938164152650 | 2.56316434447873e-05 |
| Tubb2b | ENSMUSG00000045136 | 73710 | 1.56684656642001 | 1.88474781424863 | 8.00870411306768 | 1.8052838108224 | 2.57618241985641e-05 |
| Prex1 | ENSMUSG00000039621 | 277360 | 0.834504624770778 | 4.36999345970967 | 8.00514111482435 | 1.8143309974250 | 2.58571735760801e-05 |
| Atp5c1 | ENSMUSG00000025781 | 11949 | -0.702981440647489 | 8.5541026395462 | -8.00319153755753 | 1.8193017177755 | 2.58942540065166e-05 |
| Snta1 | ENSMUSG00000027488 | 20648 | -0.580183308582882 | 6.01591504524843 | -7.99368499582784 | 1.8437477441319 | 2.62080709897352e-05 |
| Pdha1 | ENSMUSG00000031299 | 18597 | -0.69076385369581 | 8.76127945328017 | -7.99144908005849 | 1.8495478387015 | 2.62279690863983e-05 |
| Cfb | ENSMUSG00000090231 | 14962 | 1.83452725004672 | 3.01480625793361 | 7.99129570685927 | 1.8499464061488 | 2.62279690863983e-05 |
| Mgst1 | ENSMUSG00000008540 | 56615 | -1.15229600023163 | 3.63352721671276 | -7.98896916311137 | 1.8560035253596 | 2.62797597612782e-05 |
| Pamr1 | ENSMUSG00000027188 | 210622 | 2.29171351964 | 2.75445446315324 | 7.98715242271935 | 1.8607480021622 | 2.63128543488164e-05 |
| 2310065F04Rik | ENSMUSG00000087410 | 74184 | -2.73187268280903 | 1.73166923866653 | -7.9820045526873 | 1.8742618203486 | 2.64697105403503e-05 |
| Flrt2 | ENSMUSG00000047414 | 399558 | 0.936676919349725 | 4.36545133580852 | 7.96853777515514 | 1.9101084593434 | 2.69411555730112e-05 |
| Itgax | ENSMUSG00000030789 | 16411 | 2.68772672135347 | 0.296201307493433 | 7.96430994972172 | 1.9215117152124 | 2.70377911642468e-05 |
| Tbrg4 | ENSMUSG00000000384 | 21379 | -0.507916359154836 | 4.51235537213832 | -7.96416392986925 | 1.9219068460909 | 2.70377911642468e-05 |
| Ndufb8 | ENSMUSG00000025204 | 67264 | -0.738318206035614 | 6.70590566121855 | -7.96247253888903 | 1.9264900476872 | 2.70674327908348e-05 |
| B230118H07Rik | ENSMUSG00000027165 | 68170 | -0.790824652302494 | 3.55791057876347 | -7.96121217626286 | 1.9299128166647 | 2.70807150179232e-05 |
| Pcsk5 | ENSMUSG00000024713 | 18552 | 1.04989676155568 | 3.25602190554239 | 7.95725714488559 | 1.9406954316257 | 2.71971048244886e-05 |
| Cep85 | ENSMUSG00000037443 | 70012 | -0.535828891162996 | 5.67014586098909 | -7.94957446588174 | 1.9618237255196 | 2.74579963426994e-05 |
| E2f2 | ENSMUSG00000018983 | 242705 | 2.96538458161613 | 0.693182860874852 | 7.94800204546417 | 1.9661780621747 | 2.74837498690949e-05 |
| Cox4i1 | ENSMUSG00000031818 | 12857 | -0.673151317825154 | 7.76799092130298 | -7.94269123904845 | 1.9809606264923 | 2.76379761238598e-05 |
| Cxcl16 | ENSMUSG00000018920 | 66102 | 1.6052091542819 | 1.88865808304555 | 7.94222351600887 | 1.9822681622089 | 2.76379761238598e-05 |
| Fbp2 | ENSMUSG00000021456 | 14120 | -1.88434372561902 | 1.57475586436798 | -7.9322183748657 | 2.0104581362108 | 2.79953094100897e-05 |
| Srpx2 | ENSMUSG00000031253 | 68792 | 1.80932348480847 | 2.52407777843788 | 7.92965660108689 | 2.0177441981015 | 2.80610201392465e-05 |
| Fbln2 | ENSMUSG00000064080 | 14115 | 0.631930217497342 | 6.67243500461789 | 7.92800097373037 | 2.0224679356663 | 2.8090974593099e-05 |
| Gli3 | ENSMUSG00000021318 | 14634 | 0.970055284912088 | 1.85157758044587 | 7.92633088653763 | 2.0272448099257 | 2.81215901234753e-05 |
| Gys1 | ENSMUSG00000003865 | 14936 | -0.637282103287512 | 8.2409449201017 | -7.92364404722802 | 2.0349549737380 | 2.81927665626494e-05 |
| Ckm | ENSMUSG00000030399 | 12715 | -0.848697548638439 | 11.0927836213908 | -7.91889082488031 | 2.0486710884794 | 2.83200693071957e-05 |
| Hdhd2 | ENSMUSG00000025421 | 76987 | -0.514466617500598 | 5.14300012259779 | -7.91866495724864 | 2.0493252970443 | 2.83200693071957e-05 |
| Smim10l1 | ENSMUSG00000072704 | 381820 | -0.511473879064811 | 6.62105956911886 | -7.90925326234927 | 2.0767834235102 | 2.86632823262508e-05 |
| Myf6 | ENSMUSG00000035923 | 17878 | 0.70254345588594 | 5.58810554777557 | 7.90415251768672 | 2.0918273121145 | 2.88345073754399e-05 |
| Acot11 | ENSMUSG00000034853 | 329910 | -0.875281463958471 | 3.65366328140104 | -7.90280688448517 | 2.0958152870368 | 2.88530943357683e-05 |
| 1700021F05Rik | ENSMUSG00000019797 | 67851 | -0.558245927430164 | 4.36442216002326 | -7.89863394238933 | 2.1082337301797 | 2.898755082339e-05 |
| Pkd1 | ENSMUSG00000032855 | 18763 | 0.894901464485308 | 6.12519079091369 | 7.89172290492959 | 2.1289724829802 | 2.92222347527896e-05 |
| Bola3 | ENSMUSG00000045160 | 78653 | -0.663124007756174 | 6.70513149863873 | -7.89116743352514 | 2.1306487144793 | 2.92222347527896e-05 |
| Idh3b | ENSMUSG00000027406 | 170718 | -0.722454368825186 | 7.81354661474428 | -7.88890270592421 | 2.1374974181550 | 2.92794289196147e-05 |
| Gatm | ENSMUSG00000027199 | 67092 | 1.12907993509249 | 1.97273813952569 | 7.87788554634505 | 2.1711486388801 | 2.97031611659556e-05 |
| Fgf1 | ENSMUSG00000036585 | 14164 | -0.657386380234477 | 5.53835862218805 | -7.86657394498087 | 2.2062841735582 | 3.01461153764571e-05 |
| Gadl1 | ENSMUSG00000056880 | 73748 | 1.38038814034844 | 1.63940829037922 | 7.86099086161639 | 2.2238477701010 | 3.03170420478803e-05 |
| Aifm2 | ENSMUSG00000020085 | 71361 | -0.966076876164128 | 3.40086970217542 | -7.86083485860328 | 2.2243406570670 | 3.03170420478803e-05 |
| ND2 | ENSMUSG00000064345 | 17717 | -0.842071604751403 | 11.89051918289 | -7.85643613596524 | 2.2382860646336 | 3.04691220081074e-05 |
| Fgl2 | ENSMUSG00000039899 | 14190 | 1.25587763505924 | 4.20215434968696 | 7.85499128284528 | 2.2428869235056 | 3.04937773144777e-05 |
| Bmp1 | ENSMUSG00000022098 | 12153 | 1.01075971510679 | 4.27128587132115 | 7.85406008759215 | 2.2458574484565 | 3.04962332535138e-05 |
| Eepd1 | ENSMUSG00000036611 | 67484 | -1.31016873525382 | 4.25352224137777 | -7.85127722468023 | 2.2547596970460 | 3.05791293404598e-05 |
| Mpped2 | ENSMUSG00000016386 | 77015 | -1.37915654595837 | 1.35161904481747 | -7.82910696552606 | 2.3270314453726 | 3.15201743858346e-05 |
| Phb | ENSMUSG00000038845 | 18673 | -0.704311647606953 | 5.8810008217069 | -7.82571386191574 | 2.3383073501199 | 3.1633709955645e-05 |
| Gan | ENSMUSG00000052557 | 209239 | 0.738732195925268 | 3.74575390085844 | 7.82309835130088 | 2.3470386902260 | 3.17125833409898e-05 |
| Anxa2 | ENSMUSG00000032231 | 12306 | 1.04484424271271 | 5.42714534376741 | 7.81960540945812 | 2.3587530777480 | 3.18315183862514e-05 |
| Dld | ENSMUSG00000020664 | 13382 | -0.653107451172274 | 7.21666257516317 | -7.81621609434387 | 2.3701791808627 | 3.19214645941589e-05 |

| | | | | | | | |
|---|---|---|---|---|---|---|---|
| Stradb | ENSMUSG00000026027 | 227154 | -0.516878237869079 | 5.2015257319246 | -7.8158967623948 | 2.3712587366624 | 3.19214645941589e-05 |
| Mxra8 | ENSMUSG00000029070 | 74761 | 0.983999946005641 | 4.39208205224027 | 7.81322960226675 | 2.3802959034899 | 3.19646717274265e-05 |
| Galns | ENSMUSG00000015027 | 50917 | 1.0600492969571 | 2.3103029943089 | 7.81303758613318 | 2.3809479208043 | 3.19646717274265e-05 |
| Eif4a2 | ENSMUSG00000022884 | 13682 | -0.583670576032352 | 6.75203642366411 | -7.81236272477198 | 2.3832410079455 | 3.19646717274265e-05 |
| Ndufaf5 | ENSMUSG00000027384 | 69487 | -0.736555797867956 | 3.36319243839045 | -7.80945325206781 | 2.3931538312446 | 3.20582898643813e-05 |
| Ndufs4 | ENSMUSG00000021764 | 17993 | -0.641998802376368 | 6.89539013033106 | -7.79825530494932 | 2.4317158466407 | 3.25115906870301e-05 |
| Coq3 | ENSMUSG00000028247 | 230027 | -0.637380098706652 | 3.95973303470603 | -7.79790254437964 | 2.4329412846025 | 3.25115906870301e-05 |
| Taok3 | ENSMUSG00000061288 | 330177 | 0.646603593508637 | 3.94124804783327 | 7.77995837774804 | 2.4961481925508 | 3.3315501700578e-05 |
| Itm2a | ENSMUSG00000031239 | 16431 | 0.707571312268425 | 4.9608276520441 | 7.77742811055981 | 2.5051999100062 | 3.33799378374828e-05 |
| C1s1 | ENSMUSG00000038521 | 50908 | 0.917539461097546 | 4.66223522364072 | 7.77690281976627 | 2.5070834291989 | 3.33799378374828e-05 |
| Pakap | ENSMUSG00000089945 | 677884 | 1.00753812866764 | 4.36193378620701 | 7.77191519724471 | 2.5250423463973 | 3.35781482828099e-05 |
| Mrpl1 | ENSMUSG00000029486 | 94061 | -0.625939240214314 | 4.22868309209478 | -7.76969075319581 | 2.5330958065020 | 3.36443138042215e-05 |
| Cttn | ENSMUSG00000031078 | 13043 | 0.625894826051076 | 4.74563122225142 | 7.76237193685241 | 2.5597857839606 | 3.39250942423122e-05 |
| Ndufb10 | ENSMUSG00000040048 | 68342 | -0.646904093100678 | 7.43165463478774 | -7.76219273145507 | 2.5604430289916 | 3.39250942423122e-05 |
| Ccl9 | ENSMUSG00000019122 | 20308 | 1.84933120157291 | 2.3977642558824 | 7.75882742491853 | 2.5728187304931 | 3.4028581399003e-05 |
| Tlr2 | ENSMUSG00000027995 | 24088 | 1.89677127544402 | 0.852320097540544 | 7.75837708529653 | 2.5744796283025 | 3.4028581399003e-05 |
| Ndufb11 | ENSMUSG00000031059 | 104130 | -0.696191125019209 | 6.50873971126869 | -7.75651241449422 | 2.5813688226905 | 3.40784330927904e-05 |
| Pld3 | ENSMUSG00000003363 | 18807 | 0.83967873611992 | 3.62221549513316 | 7.75475582826332 | 2.5878765594154 | 3.40942400112255e-05 |
| Zhx3 | ENSMUSG00000035877 | 320799 | 0.467685233204119 | 5.234518553697 | 7.75450580586506 | 2.5888042456607 | 3.40942400112255e-05 |
| Ces5a | ENSMUSG00000058019 | 67935 | 1.90946914976384 | 0.840118995702858 | 7.75190091529306 | 2.5984904516495 | 3.41806247015412e-05 |
| Cd59a | ENSMUSG00000032679 | 12509 | -0.96589599872643 | 3.85947243777408 | -7.74584583987832 | 2.6211547529616 | 3.43750208209664e-05 |
| Pik3cg | ENSMUSG00000020573 | 30955 | 1.30478849504272 | 1.11967513766689 | 7.74542607510985 | 2.6227336874470 | 3.43750208209664e-05 |
| Ndufaf6 | ENSMUSG00000050323 | 76947 | -0.91863400440468 | 2.46033612013266 | -7.74503577164436 | 2.6242027081836 | 3.43750208209664e-05 |
| Pold4 | ENSMUSG00000024854 | 69745 | 0.883281401839255 | 2.27319475984085 | 7.74459896059812 | 2.6258478076577 | 3.43750208209664e-05 |
| Dusp4 | ENSMUSG00000031530 | 319520 | 1.05150530686619 | 1.72763785776012 | 7.73906834445086 | 2.6467717507589 | 3.46074904396478e-05 |
| Apbb1ip | ENSMUSG00000026786 | 54519 | 1.19479493500482 | 2.42784367688478 | 7.7348972522107 | 2.6626690457774 | 3.47686282774766e-05 |
| Myoc | ENSMUSG00000026697 | 17926 | 0.81399470940201 | 4.67969325438082 | 7.73416847515976 | 2.6654570027010 | 3.47686282774766e-05 |
| Dtnbp1 | ENSMUSG00000057531 | 94245 | -0.625748187735002 | 4.65970886137876 | -7.73306376118893 | 2.6696890160253 | 3.47823249513384e-05 |
| Arl6ip5 | ENSMUSG00000035199 | 65106 | 0.483983825032055 | 6.28714333763513 | 7.73082975556855 | 2.6782689673777 | 3.48525691457218e-05 |
| Hba-a2 | ENSMUSG00000069917 | 110257 | -1.24157395497548 | 6.24802736659211 | -7.72958457661519 | 2.6830638941129 | 3.4873449972115e-05 |
| B4galt5 | ENSMUSG00000017929 | 56336 | 1.09756562450575 | 3.69966521635879 | 7.72298912442173 | 2.7086137460955 | 3.51237866618992e-05 |
| Ddo | ENSMUSG00000063428 | 70503 | -0.627253609706415 | 4.15759644928821 | -7.72295398995389 | 2.7087505402965 | 3.51237866618992e-05 |
| Adsl | ENSMUSG00000022407 | 11564 | -0.539946280842214 | 5.9777874446078 | -7.71735874816303 | 2.7306289985516 | 3.53655279421428e-05 |
| Agl | ENSMUSG00000033400 | 77559 | -0.691169234845734 | 7.7921335433161 | -7.71355490285713 | 2.7456096094905 | 3.54911963560859e-05 |
| Cox7a2l | ENSMUSG00000024248 | 20463 | -0.498160259735564 | 5.56148333736635 | -7.71242827048222 | 2.7500632994934 | 3.54911963560859e-05 |
| Mtfr1 | ENSMUSG00000027601 | 67472 | -0.507791382297953 | 4.53635606059271 | -7.7124259261976 | 2.7500725746596 | 3.54911963560859e-05 |
| Sdhc | ENSMUSG00000058076 | 66052 | -0.642073788039709 | 6.82460775813781 | -7.70529290421393 | 2.7784484464727 | 3.58151178872572e-05 |
| Mrpl45 | ENSMUSG00000018882 | 67036 | -0.653625840145738 | 4.91586485220565 | -7.7004078559636 | 2.7980605073681 | 3.60254409965151e-05 |
| Cyba | ENSMUSG00000006519 | 13057 | 1.10957461317824 | 2.16545471673588 | 7.6948843092399 | 2.8204127556503 | 3.62705080376632e-05 |
| Ptprs | ENSMUSG00000013236 | 19280 | 0.84071932673949 | 5.40564681017901 | 7.69231762482035 | 2.8308637115026 | 3.63621283553886e-05 |
| Csf2rb2 | ENSMUSG00000071714 | 12984 | 1.26891295911467 | 1.60720534380984 | 7.68890706633723 | 2.8448142486043 | 3.64984325721768e-05 |
| Nkiras1 | ENSMUSG00000021772 | 69721 | -0.623726554991097 | 3.79228808699248 | -7.68393204881788 | 2.8652947957472 | 3.66807287776235e-05 |
| Loxl2 | ENSMUSG00000034205 | 94352 | 1.25644059942549 | 5.52878586412092 | 7.6838256838122 | 2.8657343679526 | 3.66807287776235e-05 |
| Trip10 | ENSMUSG00000019487 | 106628 | -0.626106252098984 | 5.96845535358147 | -7.67887831034701 | 2.8862594090473 | 3.69002357898208e-05 |
| Atp5j | ENSMUSG00000022890 | 11957 | -0.683907182723424 | 8.0768593738997 | -7.67727283139888 | 2.8929534395550 | 3.69426098688977e-05 |
| Msrb3 | ENSMUSG00000051236 | 320183 | -0.481434025857403 | 6.3531489305853 | -7.67489466824953 | 2.9028994107847 | 3.70016867195715e-05 |
| Gas7 | ENSMUSG00000033066 | 14457 | 0.731857807919163 | 4.19050018816677 | 7.67454859958076 | 2.9043497580635 | 3.70016867195715e-05 |
| Tgif1 | ENSMUSG00000047407 | 21815 | 1.23422867075067 | 1.54180607991925 | 7.66954469831581 | 2.9254067335200 | 3.72265669430827e-05 |
| Hmgn5 | ENSMUSG00000031245 | 50887 | -0.873811138592448 | 3.84230590021741 | -7.66301415301767 | 2.9531315552860 | 3.75356756172459e-05 |
| AW551984 | ENSMUSG00000038112 | 244810 | 2.70714384109802 | 0.998572801867656 | 7.65939774277 | 2.9686043081801 | 3.76885176454324e-05 |
| Tram2 | ENSMUSG00000041779 | 170829 | 1.24166425812517 | 3.25623111860404 | 7.65674257880549 | 2.9800190413041 | 3.7789545406608e-05 |
| Pik3ap1 | ENSMUSG00000025017 | 83490 | 1.88883270757775 | 1.4900708099062 | 7.65317130853666 | 2.9954455519349 | 3.79411533351111e-05 |
| Pdcd6ip | ENSMUSG00000032504 | 18571 | 0.447761530240203 | 6.14779005016194 | 7.64784898476896 | 3.0185931024967 | 3.81608666528097e-05 |
| Olfml3 | ENSMUSG00000027848 | 99543 | 1.12199300458219 | 3.90259032972385 | 7.64757861941382 | 3.0197740055512 | 3.81608666528097e-05 |
| Gng5 | ENSMUSG00000068523 | 14707 | -0.654459175224233 | 5.98361163163472 | -7.64531985569997 | 3.0296589863232 | 3.82141247494278e-05 |
| Bpgm | ENSMUSG00000038871 | 12183 | -0.637698341043384 | 3.84236307115456 | -7.64501851060779 | 3.0309803456000 | 3.82141247494278e-05 |
| Bmper | ENSMUSG00000031963 | 73230 | 1.25669127599256 | 1.8763418483597 | 7.63975460928018 | 3.0541605985522 | 3.84206920882128e-05 |
| Dpysl3 | ENSMUSG00000024501 | 22240 | 1.28358193529592 | 4.54881302152229 | 7.63949669609589 | 3.0553011663243 | 3.84206920882128e-05 |
| Ssc5d | ENSMUSG00000035279 | 269855 | 1.49818940983156 | 4.95210238230163 | 7.63890740141415 | 3.0579088936735 | 3.84206920882128e-05 |
| Atp5o | ENSMUSG00000022956 | 28080 | -0.63154665038862 | 7.54660476898893 | -7.63600424530326 | 3.0707903035028 | 3.85382420293793e-05 |
| Npr3 | ENSMUSG00000022206 | 18162 | 0.784754173170514 | 6.12740295800751 | 7.63179739915305 | 3.0895583122928 | 3.87293141188908e-05 |
| Arpc1b | ENSMUSG00000029622 | 11867 | 0.819007653710387 | 4.27057602644781 | 7.62723803504405 | 3.1100362734958 | 3.89413591129243e-05 |
| Ndufa10 | ENSMUSG00000026260 | 67273 | -0.664874456801201 | 7.5360834676737 | -7.62298623655867 | 3.1292623983649 | 3.91176080264908e-05 |
| Clcn5 | ENSMUSG00000004317 | 12728 | 1.16085658501116 | 2.30994310473915 | 7.62254396556055 | 3.1312695108571 | 3.91176080264908e-05 |
| Fat1 | ENSMUSG00000070047 | 14107 | 1.19043846813073 | 5.32302058125226 | 7.6196663746524 | 3.1443619230330 | 3.9236324407162e-05 |
| Ssr1 | ENSMUSG00000021427 | 107513 | 0.426383975006837 | 5.54813554424484 | 7.61301534484395 | 3.1748450323644 | 3.95715291320128e-05 |
| Sec31a | ENSMUSG00000035325 | 69162 | 0.440571511598834 | 6.12030735942058 | 7.60984463339652 | 3.1894871116271 | 3.97087512724327e-05 |
| Cox17 | ENSMUSG00000046516 | 12856 | -0.475509079802888 | 5.41176157378362 | -7.60650717965254 | 3.2049764772386 | 3.985619780739e-05 |
| Fermt3 | ENSMUSG00000024965 | 108101 | 1.74803970089136 | 0.965195607538751 | 7.59347943138073 | 3.2662058517749 | 4.05714729156271e-05 |
| Atp5j2 | ENSMUSG00000038690 | 57423 | -0.70046234705208 | 6.48155661147607 | -7.5921853985563 | 3.2723549828458 | 4.06017165919274e-05 |
| Dnajb5 | ENSMUSG00000036052 | 56323 | -0.551562201254862 | 6.61732239048944 | -7.58694492744424 | 3.2973828804388 | 4.08658642472534e-05 |
| Ciapin1 | ENSMUSG00000031781 | 109006 | -0.769578121663306 | 3.52685565093201 | -7.57757011510179 | 3.3426629629076 | 4.13440947247684e-05 |
| Notch2 | ENSMUSG00000027878 | 18129 | 0.874667359422205 | 4.81187821206287 | 7.57739091588853 | 3.3435348766531 | 4.13440947247684e-05 |
| Cox10 | ENSMUSG00000042148 | 70383 | -0.610733971065046 | 4.8762880127494 | -7.57188932390594 | 3.3704210231600 | 4.16294601176979e-05 |
| Paics | ENSMUSG00000029247 | 67054 | -0.508773291863108 | 4.95444165623332 | -7.56935842484272 | 3.3828662608438 | 4.17360170398237e-05 |
| Acox1 | ENSMUSG00000020777 | 11430 | -0.49298367700818 | 6.82203473219494 | -7.56422971493122 | 3.4082352207013 | 4.20015999971667e-05 |
| Pla2g12a | ENSMUSG00000027999 | 66350 | -0.84458889380455 | 7.05566327286607 | -7.56244078098419 | 3.4171314350916 | 4.20638104921021e-05 |
| Fap | ENSMUSG00000000392 | 14089 | 1.20412226978049 | 2.60925758182836 | 7.56071793081372 | 3.4257222657320 | 4.21207552192469e-05 |
| Adgra2 | ENSMUSG00000031486 | 78560 | 1.15742397567153 | 2.94031991721801 | 7.55996895227337 | 3.4294641062235 | 4.21207552192469e-05 |
| Ext1 | ENSMUSG00000061731 | 14042 | 0.728148380569502 | 4.10583153947889 | 7.55451391243022 | 3.4568479624309 | 4.24094333078932e-05 |
| Hspa1b | ENSMUSG00000090877 | 15511 | 0.816241205215485 | 5.24536445677682 | 7.55054244619439 | 3.4769299872143 | 4.26079839576681e-05 |
| Sdr39u1 | ENSMUSG00000022223 | 654795 | -0.537088916921867 | 4.4602020205363 | -7.5490775784273 | 3.4843683561041 | 4.26513219491315e-05 |
| Arhgap5 | ENSMUSG00000035133 | 11855 | -0.44091530065071 | 5.81693437233583 | -7.5474603635024 | 3.4925998929739 | 4.27042611074918e-05 |
| Mcur1 | ENSMUSG00000021371 | 76137 | -0.575209766775427 | 5.49660464535255 | -7.53994958873434 | 3.5311001031198 | 4.31216582617219e-05 |
| Plek | ENSMUSG00000020120 | 56193 | 1.84794681244515 | 1.68130635075988 | 7.53873498725212 | 3.5373682386183 | 4.31216582617219e-05 |
| Scn7a | ENSMUSG00000034810 | 20272 | 1.06156260603952 | 3.6551916536802 | 7.53850204485113 | 3.5385717190343 | 4.31216582617219e-05 |
| Rab8b | ENSMUSG00000036943 | 235442 | 0.75224613114683 | 3.4680764295694 | 7.53605006814148 | 3.5512660132733 | 4.32281612372954e-05 |
| Gcat | ENSMUSG00000006378 | 26912 | -1.62071890131093 | 1.55830522852532 | -7.534598420709 | 3.5588041501862 | 4.32717332210078e-05 |
| Sbno2 | ENSMUSG00000035673 | 216161 | 1.14103457102561 | 4.09586374350891 | 7.53375589273977 | 3.5631870103172 | 4.32768857886426e-05 |
| Atxn10 | ENSMUSG00000016541 | 54138 | 0.572915450302608 | 4.48088401581922 | 7.52589567179003 | 3.6043525885357 | 4.37282776307261e-05 |
| Sh2b2 | ENSMUSG00000005057 | 23921 | 1.29593457201709 | 1.37724429878456 | 7.51709119408421 | 3.6510620266868 | 4.42458525650926e-05 |
| Tmed9 | ENSMUSG00000058569 | 67511 | 0.557703289689801 | 4.78318361714331 | 7.51379071345499 | 3.6687364788014 | 4.44108066996437e-05 |
| Gm13889 | ENSMUSG00000087006 | 620695 | 1.58882214359694 | 3.11965476142342 | 7.50822443167009 | 3.6987500698231 | 4.47245984659697e-05 |
| Rassf2 | ENSMUSG00000027339 | 215653 | 1.21729055441608 | 3.11469315799913 | 7.50527047512371 | 3.7147833867412 | 4.48688366856001e-05 |
| Dlg5 | ENSMUSG00000021782 | 71228 | 0.688619578176718 | 3.90727818013691 | 7.49986780244611 | 3.7442983535622 | 4.51754142414887e-05 |
| Ddhd1 | ENSMUSG00000037697 | 114874 | 0.763461013702287 | 3.13081423250165 | 7.49475085987841 | 3.7724813433797 | 4.54652630258918e-05 |
| Tlr7 | ENSMUSG00000044583 | 170743 | 1.8651066500063 | 0.775557976046685 | 7.49070945656246 | 3.7948991979137 | 4.56850695290698e-05 |
| Epm2a | ENSMUSG00000055493 | 13853 | -0.639570732371759 | 5.24478918905716 | -7.48388627972341 | 3.8330683092373 | 4.60938060377046e-05 |
| Mrpl18 | ENSMUSG00000057388 | 67681 | -0.598475727863939 | 5.80646690462012 | -7.48174676848768 | 3.8451203444644 | 4.61085240141483e-05 |
| Grtp1 | ENSMUSG00000038515 | 66790 | -1.47180544240137 | 1.06842717934724 | -7.4814932407632 | 3.8465511421257 | 4.61085240141483e-05 |
| Sec61a1 | ENSMUSG00000030082 | 53421 | 0.726087646622976 | 4.23188684096595 | 7.48073973124836 | 3.8508069421716 | 4.61085240141483e-05 |
| Atp2a1 | ENSMUSG00000030730 | 11937 | -0.896434326867286 | 11.816551756446 | -7.48067641107369 | 3.8511647996448 | 4.61085240141483e-05 |
| Usp15 | ENSMUSG00000020124 | 14479 | -0.498890594438866 | 6.80392203928203 | -7.47784230360363 | 3.8672180184526 | 4.62097477869216e-05 |
| Higd2a | ENSMUSG00000025868 | 67044 | -0.529205065231278 | 4.46719075016702 | -7.47769149710643 | 3.8680742132497 | 4.62097477869216e-05 |
| Zfyve9 | ENSMUSG00000034557 | 230597 | -0.49166241617421 | 5.90917035101861 | -7.47630227461491 | 3.8759708846959 | 4.6253534651323e-05 |
| Cd300ld | ENSMUSG00000034641 | 217305 | 1.7414763572095 | 1.03471909322721 | 7.47515492450853 | 3.8825055623327 | 4.6280990514569e-05 |
| Pdzrn3 | ENSMUSG00000035357 | 55983 | -0.612918836600621 | 6.91882099734942 | -7.47279430892943 | 3.8959870429593 | 4.63911049744973e-05 |
| Myo9a | ENSMUSG00000039585 | 270163 | 0.587325716246832 | 5.2050729208563 | 7.47198994755976 | 3.9005920639031 | 4.6395399184467e-05 |
| Pccb | ENSMUSG00000032527 | 66904 | -0.621438410571565 | 4.82906771678306 | -7.46982747167297 | 3.9130009677096 | 4.64924060630806e-05 |
| Tcea3 | ENSMUSG00000001604 | 21401 | -0.829556606766875 | 7.35685470788786 | -7.46855308121065 | 3.9203333465732 | 4.6510674214888e-05 |
| Gpd1 | ENSMUSG00000023019 | 14555 | -1.06960722436564 | 4.19891001907822 | -7.46738573719508 | 3.9270625821057 | 4.6510674214888e-05 |
| Mrpl34 | ENSMUSG00000034880 | 94065 | -0.705281410518416 | 3.27769901775772 | -7.46734401939476 | 3.9273032934170 | 4.6510674214888e-05 |
| Cbl | ENSMUSG00000034342 | 12402 | 0.548364350423852 | 4.40949646229147 | 7.4653818354057 | 3.9386427371865 | 4.65944845889465e-05 |
| Esrra | ENSMUSG00000024955 | 26379 | -0.666043247534529 | 5.14003447672123 | -7.45936586195609 | 3.9736255178047 | 4.6957514092026e-05 |
| Actr3 | ENSMUSG00000026341 | 74117 | 0.484089793916041 | 5.65465559239116 | 7.45781985543904 | 3.9826685340934 | 4.70135526416578e-05 |
| Dmac2l | ENSMUSG00000054894 | 68055 | -1.05239559493691 | 1.94811733496739 | -7.45242132762324 | 4.0144172214937 | 4.73372110551761e-05 |

| | | | | | | | |
|---|---|---|---|---|---|---|---|
| Pvr | ENSMUSG00000040511 | 52118 | 1.1122051308392 | 3.02186863749291 | 7.44779593134947 | 4.0418322192591 | 4.76091249878462e-05 |
| Igsf10 | ENSMUSG00000036334 | 242050 | 1.0848540873965 | 4.33949853123499 | 7.44602983480257 | 4.0523352192239 | 4.7681659648341e-05 |
| Anp32a | ENSMUSG00000032249 | 11737 | -0.516960363688831 | 4.69777504436925 | -7.43655249923605 | 4.1093023104071 | 4.82997672635058e-05 |
| Ephb3 | ENSMUSG00000005958 | 13845 | 1.64253489275788 | 1.50283114524365 | 7.4348193058177 | 4.1198085228269 | 4.83712427100126e-05 |
| Shisal1 | ENSMUSG00000062760 | 72301 | 3.62546588778811 | -0.625753709155336 | 7.43377304307633 | 4.1261644796762 | 4.83938883340568e-05 |
| Smad1 | ENSMUSG00000031681 | 17125 | 0.689462335719317 | 3.86538901666136 | 7.43233351397844 | 4.1349264636036 | 4.84446743554678e-05 |
| Syngr1 | ENSMUSG00000022415 | 20972 | -1.07775354566111 | 1.90889545896368 | -7.43051105076839 | 4.1460475084114 | 4.85229607221042e-05 |
| Kbtbd13 | ENSMUSG00000054978 | 74492 | -0.882475078417781 | 3.18668639642758 | -7.42929393841539 | 4.1534922044369 | 4.85580997718717e-05 |
| Ccdc8 | ENSMUSG00000041117 | 434130 | 0.840557910187634 | 2.7470881117592 | 7.42782162723804 | 4.1625167846244 | 4.8611614287104e-05 |
| Ndufb2 | ENSMUSG00000002416 | 68198 | -0.673222839797621 | 5.00807969977428 | -7.42556504759848 | 4.1763888639438 | 4.87215652847068e-05 |
| Myc | ENSMUSG00000022346 | 17869 | 1.48519068230658 | 1.76845941867989 | 7.42369925715743 | 4.1878955585734 | 4.87912728334443e-05 |
| ND6 | ENSMUSG00000064368 | 17722 | -0.77369352522645 | 8.24042642659431 | -7.42314968041529 | 4.1912912991136 | 4.87912728334443e-05 |
| Mospd1 | ENSMUSG00000023074 | 70380 | -0.529384247133928 | 3.86567589452197 | -7.42212490656272 | 4.1976309861908 | 4.88130896915452e-05 |
| Patl1 | ENSMUSG00000046139 | 225929 | -0.624189083970138 | 4.51760040889802 | -7.42007165854845 | 4.2103638086488 | 4.89091251778331e-05 |
| Uqcrc2 | ENSMUSG00000030884 | 67003 | -0.583048631860085 | 7.36301175133131 | -7.41925157492777 | 4.2154608167099 | 4.891635051747e-05 |
| Tmod4 | ENSMUSG00000005628 | 50874 | -0.591809998396386 | 6.70459544630551 | -7.41806253031969 | 4.2228626029431 | 4.89502768958335e-05 |
| Sat1 | ENSMUSG00000025283 | 20229 | 0.917819804282433 | 3.41058348379299 | 7.41669905039771 | 4.2313671785296 | 4.89969010895212e-05 |
| Mrln | ENSMUSG00000019933 | 69563 | -1.12218849083461 | 1.78809803923319 | -7.40631149829993 | 4.2967566227411 | 4.9701425019242e-05 |
| Scn3a | ENSMUSG00000057182 | 20269 | 1.34096504769037 | 1.12986013587465 | 7.40400275808206 | 4.3114348327656 | 4.97675319652861e-05 |
| Hbb-b1 | ENSMUSG00000052305 | 15129 | -1.27061523758341 | 5.63284186821552 | -7.403980371404 | 4.3115774193693 | 4.97675319652861e-05 |
| Fgfr1 | ENSMUSG00000031565 | 14182 | 0.575570654924092 | 5.89803434235877 | 7.40075249551675 | 4.3321889286845 | 4.99526974466784e-05 |
| Slc41a2 | ENSMUSG00000034591 | 338365 | 1.88837270103215 | 0.993530392048209 | 7.39922538640783 | 4.3419765501347 | 5.00127983872742e-05 |
| Dse | ENSMUSG00000039497 | 212898 | 1.14681744961501 | 3.14640139314978 | 7.39796506422117 | 4.3500718927324 | 5.00335151331923e-05 |
| Mib1 | ENSMUSG00000024294 | 225164 | 0.488633138889124 | 5.47369251108358 | 7.3975207631557 | 4.3529295482266 | 5.00335151331923e-05 |
| Fbxo31 | ENSMUSG00000052934 | 76454 | -0.65517205982774 | 5.74179600602858 | -7.39578000278619 | 4.3641449231474 | 5.01097354568532e-05 |
| Pfkfb1 | ENSMUSG00000025271 | 18639 | -2.10430340033944 | 2.28130194024738 | -7.39227001623319 | 4.3868521647737 | 5.03176086181975e-05 |
| Fam220a | ENSMUSG00000118332 | 67238 | -0.6170417060326 | 5.63595716116509 | -7.3907505199622 | 4.3967210247459 | 5.03779428946521e-05 |
| Maged1 | ENSMUSG00000025151 | 94275 | 0.443900038552477 | 5.2766992227241 | 7.3878944701994 | 4.4153341960185 | 5.05382388446894e-05 |
| Pdpr | ENSMUSG00000033624 | 319518 | -0.643907408308898 | 6.1812747137499 | -7.38435453667279 | 4.4385200982792 | 5.07504845128565e-05 |
| Mrpl2 | ENSMUSG00000002767 | 27398 | -0.560812420149066 | 4.3694845840855 | -7.37839527130407 | 4.4778436429320 | 5.11287130872715e-05 |
| Pgm5 | ENSMUSG00000041731 | 226041 | 0.517203807751197 | 4.89805372293026 | 7.37792629354406 | 4.4809539051876 | 5.11287130872715e-05 |
| Hba-a1 | ENSMUSG00000069919 | 15122 | -1.49814922018228 | 3.74898523121236 | -7.37602346986599 | 4.4935969113232 | 5.12195076513808e-05 |
| Cox6b1 | ENSMUSG00000036751 | 110323 | -0.582443546390065 | 6.45953015063824 | -7.3743824047939 | 4.5045310185483 | 5.12788787604884e-05 |
| Ppia | ENSMUSG00000071866 | 268373 | 0.525100027288858 | 5.4132100047301 | 7.37383448798773 | 4.5081879506750 | 5.12788787604884e-05 |
| Lhfp | ENSMUSG00000048332 | 108927 | 0.991897064956622 | 3.58626650293863 | 7.37272481317245 | 4.5156038211991 | 5.13098392614637e-05 |
| Ubxn2a | ENSMUSG00000020634 | 217379 | -0.647497560496406 | 6.31295320610926 | -7.37198914291985 | 4.5205273626476 | 5.13124450686338e-05 |
| Abca9 | ENSMUSG00000041797 | 217262 | 0.774532821325006 | 4.39117400188811 | 7.36864627698926 | 4.5429714564847 | 5.15137147207832e-05 |
| Angptl2 | ENSMUSG00000004105 | 26360 | 0.968951804685586 | 5.05996493460524 | 7.3639427324817 | 4.5747510497500 | 5.18203147407436e-05 |
| Ndufb5 | ENSMUSG00000027673 | 66046 | -0.604100788228955 | 6.67896976693908 | -7.36114141819353 | 4.5937898743007 | 5.19821088157154e-05 |
| Eprs | ENSMUSG00000026615 | 107508 | -0.475782124037757 | 7.9250931108749 | -7.35840422724357 | 4.6124738844934 | 5.21395574264712e-05 |
| Fastkd1 | ENSMUSG00000027086 | 320720 | -0.581346031030224 | 3.18476962365176 | -7.35499797766993 | 4.6358372258860 | 5.23495214009922e-05 |
| Asb8 | ENSMUSG00000048175 | 78541 | -0.503255899505665 | 5.00065713222655 | -7.35415461450427 | 4.6416411391871 | 5.23609693420585e-05 |
| Slco3a1 | ENSMUSG00000025790 | 108116 | -0.588987628781044 | 4.93266995121341 | -7.34974105284525 | 4.6721403305578 | 5.26445954511078e-05 |
| Islr | ENSMUSG00000037206 | 26968 | 1.02814047799763 | 4.28906379111851 | 7.34847860494565 | 4.6809032132842 | 5.26445954511078e-05 |
| Scrn3 | ENSMUSG00000008226 | 74616 | -0.604568572723969 | 4.14416912377503 | -7.34814777559616 | 4.6832024347693 | 5.26445954511078e-05 |
| Sdc2 | ENSMUSG00000022261 | 15529 | 0.824891314716008 | 4.03248016730898 | 7.34773856301904 | 4.6860480627507 | 5.26445954511078e-05 |
| Arhgap45 | ENSMUSG00000035697 | 70719 | 1.55518981523218 | 1.87912679854753 | 7.33269353474238 | 4.7919502027080 | 5.37790633119113e-05 |
| Ezr | ENSMUSG00000052397 | 22350 | 0.970625609816187 | 2.68524891697771 | 7.33142773542725 | 4.8009749238678 | 5.3825083992615e-05 |
| S100a10 | ENSMUSG00000041959 | 20194 | 1.05154470688727 | 4.28915639406344 | 7.32642985614427 | 4.8367841489924 | 5.41709913244221e-05 |
| Esrrb | ENSMUSG00000021255 | 26380 | -0.851636867183588 | 4.46023512428551 | -7.32325783550629 | 4.8596577958078 | 5.43714630153278e-05 |
| Fam177a | ENSMUSG00000095595 | 73385 | 1.14263726213279 | 4.9497943008571 | 7.32090569999323 | 4.8766773060494 | 5.44698932238246e-05 |
| Gpsm2 | ENSMUSG00000027883 | 76123 | 1.35833535927203 | 1.69233325813872 | 7.32066753360044 | 4.8784215045397 | 5.44698932238246e-05 |
| Fzd1 | ENSMUSG00000044674 | 14362 | 1.05740746805542 | 2.77463994365068 | 7.31496645192917 | 4.9199896692295 | 5.48223654214654e-05 |
| Tram1 | ENSMUSG00000025935 | 72265 | 0.652367268157625 | 4.30369679093408 | 7.31496228747263 | 4.9200201700171 | 5.48223654214654e-05 |
| Slc39a13 | ENSMUSG00000002105 | 68427 | 0.57194162722715 | 3.49399084658338 | 7.31425727694287 | 4.9251866035976 | 5.48240476211056e-05 |
| Hjv | ENSMUSG00000038403 | 69585 | -0.584367075008191 | 7.74305912654548 | -7.30483451481329 | 4.9947917947376 | 5.55422880043515e-05 |
| Etv5 | ENSMUSG00000013089 | 104156 | 0.766851750103023 | 3.91665075952332 | 7.29120856191795 | 5.0972917200445 | 5.6597722400359e-05 |
| Slc25a12 | ENSMUSG00000027010 | 78830 | -0.679109860948599 | 7.63280974561252 | -7.2908445795091 | 5.1000600644363 | 5.6597722400359e-05 |
| Col6a2 | ENSMUSG00000020241 | 12834 | 1.09338876895327 | 6.86612033916297 | 7.28854813741902 | 5.1175628440511 | 5.67343604952567e-05 |
| Thap12 | ENSMUSG00000030753 | 72981 | -0.509282311214765 | 6.61270155048327 | -7.28615913544933 | 5.1358385307304 | 5.68433698669868e-05 |
| Mdh1 | ENSMUSG00000020321 | 17449 | -0.742035371389605 | 8.66455718872883 | -7.28586315226123 | 5.1381075815692 | 5.68433698669868e-05 |
| Fars2 | ENSMUSG00000021420 | 69955 | -0.713481215176357 | 3.35341591698284 | -7.28522591681246 | 5.1429963222440 | 5.68433698669868e-05 |
| Oxnad1 | ENSMUSG00000021906 | 218885 | -0.697676374532756 | 3.77031564841491 | -7.2821570818202 | 5.1666087952126 | 5.6991111122287e-05 |
| Arhgap22 | ENSMUSG00000063506 | 239027 | 1.51880242799694 | 1.45361303364333 | 7.28213347438763 | 5.1667908811807 | 5.6991111122287e-05 |
| Popdc3 | ENSMUSG00000019848 | 78977 | 0.560748738107814 | 4.56574060079575 | 7.27939038743704 | 5.1879948090038 | 5.71673097351018e-05 |
| Rap1gap2 | ENSMUSG00000038807 | 380711 | 0.800051901011265 | 3.34539572969006 | 7.27667753312078 | 5.2090555814517 | 5.73415776040774e-05 |
| Tcf15 | ENSMUSG00000068079 | 21407 | -1.25194840590146 | 3.01898216338048 | -7.27539389441226 | 5.2190523859773 | 5.73938245785895e-05 |
| Lgr4 | ENSMUSG00000050199 | 107515 | -0.846643718253881 | 3.90486977136987 | -7.27291657548398 | 5.2384028233611 | 5.75487248865934e-05 |
| Slc39a1 | ENSMUSG00000052310 | 30791 | 0.470858916050775 | 4.79788688844817 | 7.26478006959094 | 5.3024930752705 | 5.8141134583336e-05 |
| Eif1 | ENSMUSG00000035530 | 20918 | -0.576353558087958 | 7.86071002497201 | -7.26472053456881 | 5.3029650699465 | 5.8141134583336e-05 |
| Tec | ENSMUSG00000029217 | 21682 | 0.906474773137068 | 2.12589503081237 | 7.25967653323923 | 5.3431158117751 | 5.84855648487791e-05 |
| Tnip1 | ENSMUSG00000020400 | 57783 | 0.480171775601383 | 4.69164996635636 | 7.25943069531262 | 5.3450808968923 | 5.84855648487791e-05 |
| Mmab | ENSMUSG00000029575 | 77697 | -0.690628755747085 | 3.38076069398283 | -7.25695802301524 | 5.3648886264388 | 5.86435975756033e-05 |
| Gpt2 | ENSMUSG00000031700 | 108682 | -1.32042342966664 | 3.88439404108341 | -7.25596381804883 | 5.3728747296330 | 5.86722214481709e-05 |
| Tmem158 | ENSMUSG00000054871 | 72309 | 2.18118060069299 | 0.468425131773777 | 7.25320450485028 | 5.3951052727640 | 5.88561833698442e-05 |
| Sulf1 | ENSMUSG00000016918 | 240725 | 0.988183717811079 | 4.44038659623954 | 7.2521690403325 | 5.4034726265799 | 5.88886932015412e-05 |
| Lilr4b | ENSMUSG00000112023 | 14727 | 2.05028137188439 | 1.17799207052678 | 7.24712363546363 | 5.4444402082486 | 5.92760716298468e-05 |
| Ethe1 | ENSMUSG00000064254 | 66071 | -0.782582004182538 | 2.88635243609134 | -7.24606941165362 | 5.4530416879274 | 5.93106454634174e-05 |
| Sh3pxd2a | ENSMUSG00000053617 | 14218 | 0.680116836520706 | 6.41197307444286 | 7.24317304997127 | 5.4767473492470 | 5.9484685445612e-05 |
| Gpr34 | ENSMUSG00000040229 | 23890 | 2.01162226300078 | 0.388969271420889 | 7.2427856106632 | 5.4799266529806 | 5.9484685445612e-05 |
| Cd109 | ENSMUSG00000046186 | 235505 | 1.25844366012476 | 1.91317613790054 | 7.24193166538883 | 5.4869409704117 | 5.9501737844812e-05 |
| Fcrls | ENSMUSG00000015852 | 80891 | 2.60181601663375 | 1.19806688065787 | 7.23705218894746 | 5.5272036020163 | 5.98789520055901e-05 |
| Slc11a1 | ENSMUSG00000026177 | 18173 | 1.43386151344745 | 1.22125589421914 | 7.23480213174633 | 5.5458749208141 | 6.00217413459603e-05 |
| Arpc5 | ENSMUSG00000008475 | 67771 | 0.652827640893818 | 3.45774637665107 | 7.23276329888822 | 5.5628510723896 | 6.01459199528107e-05 |
| C1ra | ENSMUSG00000055172 | 50909 | 1.08289410464808 | 3.82154442172165 | 7.22821264503959 | 5.6009401809011 | 6.04979022899505e-05 |
| Tgfb3 | ENSMUSG00000021253 | 21809 | 0.401415385550358 | 5.71245585836478 | 7.2247126916877 | 5.6304224774102 | 6.07563159926662e-05 |
| Agap1 | ENSMUSG00000055013 | 347722 | 0.491088936384458 | 5.48869357103166 | 7.22344412305683 | 5.6411489052842 | 6.08120302600216e-05 |
| Tbc1d31 | ENSMUSG00000022364 | 210544 | 1.31630079575103 | 1.35297122629167 | 7.21988944490511 | 5.6713208806060 | 6.10770527545858e-05 |
| Lrrc17 | ENSMUSG00000039883 | 74511 | 1.57303864709796 | 1.47116664559628 | 7.21767669403173 | 5.6901887615030 | 6.11648820989346e-05 |
| Npepps | ENSMUSG00000001441 | 19155 | -0.378284068583765 | 6.16802952377893 | -7.21760569188554 | 5.6907952868882 | 6.11648820989346e-05 |
| Sept9 | ENSMUSG00000059248 | 53860 | 0.726082259961432 | 4.77583867991702 | 7.21696599114212 | 5.6962629198349 | 6.11648820989346e-05 |
| Flad1 | ENSMUSG00000042642 | 319945 | -0.554602307391522 | 4.15162892355508 | -7.21555277570146 | 5.7083616169056 | 6.12346426245295e-05 |
| Tuba1c | ENSMUSG00000043091 | 22146 | 1.31955785927899 | 2.43730979931408 | 7.21363227446974 | 5.7248468155832 | 6.13512750403334e-05 |
| Adcy7 | ENSMUSG00000031659 | 11513 | 1.16836679327241 | 4.25674081983232 | 7.21182751186633 | 5.7403844219005 | 6.14575339038153e-05 |
| Il10ra | ENSMUSG00000032089 | 16154 | 1.68491446126421 | 1.2887849034521 | 7.20932608437902 | 5.7619935424302 | 6.16285238867952e-05 |
| Nipal3 | ENSMUSG00000028803 | 74552 | -0.699175735716688 | 3.08101054923259 | -7.20414335149636 | 5.8070398855888 | 6.20496119153192e-05 |
| Hbb-bt | ENSMUSG00000073940 | 101488143 | -1.26975671657947 | 3.25934554005102 | -7.19747504818368 | 5.8655466933934 | 6.26135653373864e-05 |
| Fbln7 | ENSMUSG00000027386 | 70370 | 3.47734817993338 | 2.21196178710171 | 7.19617593908906 | 5.8770172658519 | 6.2674805593198e-05 |
| Ogdh | ENSMUSG00000020456 | 18293 | -0.601691752453839 | 9.2230942021583 | -7.19350829948537 | 5.9006457903001 | 6.28657245391e-05 |
| Mlycd | ENSMUSG00000074064 | 56690 | -0.721131875188394 | 5.48679211452743 | -7.18723919789612 | 5.9565704005203 | 6.33476805969198e-05 |
| Antxr1 | ENSMUSG00000033420 | 69538 | 1.20325157607908 | 5.23195360480075 | 7.18713568897599 | 5.9574984588448 | 6.33476805969198e-05 |
| Dlst | ENSMUSG00000004789 | 78920 | -0.566820483097882 | 7.41914178075372 | -7.18517222952629 | 5.9751317700385 | 6.34185338910819e-05 |
| Pcyt2 | ENSMUSG00000025137 | 68671 | -0.679194787398831 | 3.58721806912014 | -7.18510180632821 | 5.9757652463465 | 6.34185338910819e-05 |
| Jph1 | ENSMUSG00000042686 | 57339 | -0.616980470071504 | 6.89252717650256 | -7.18117572274692 | 6.0111940288537 | 6.37326497860331e-05 |
| Dock10 | ENSMUSG00000038608 | 210293 | 0.876603945278353 | 2.96395963413428 | 7.17850690363252 | 6.0354040581280 | 6.39273272862377e-05 |
| Lrrc2 | ENSMUSG00000032495 | 74249 | -0.476782267510147 | 6.68794377385702 | -7.17601081141739 | 6.0581405114866 | 6.4106034783214e-05 |
| Cdk18 | ENSMUSG00000026437 | 18557 | -1.05797822166414 | 1.57391931239525 | -7.17443697787734 | 6.0725228102306 | 6.41573253689383e-05 |
| Prkaa2 | ENSMUSG00000028518 | 108079 | -0.560841307207307 | 7.00120058640005 | -7.17419621054541 | 6.0747262166637 | 6.41573253689383e-05 |
| Zfp697 | ENSMUSG00000050064 | 242109 | 1.19959447571821 | 2.7876959758064 | 7.17299873018814 | 6.0856976415135 | 6.42111591885951e-05 |
| Gpt | ENSMUSG00000022546 | 76282 | -0.889928614530132 | 3.57081746404185 | -7.17117919044573 | 6.1024085478959 | 6.43253884638863e-05 |
| Kdelr2 | ENSMUSG00000079111 | 66913 | 0.598532119932299 | 4.2245318181927 | 7.16985256321559 | 6.1146230521016 | 6.43920468039724e-05 |
| Scn5a | ENSMUSG00000032511 | 20271 | 3.04738374216606 | 0.371841376278533 | 7.1684405887396 | 6.1276517628853 | 6.44671428489887e-05 |
| Mki67 | ENSMUSG00000031004 | 17345 | 1.8074263748833 | 2.79507889613062 | 7.16285050732427 | 6.1795219218213 | 6.49503405071432e-05 |
| Spg21 | ENSMUSG00000032388 | 27965 | 0.946824798048419 | 2.18116019174211 | 7.16161482160036 | 6.1910503241838 | 6.50090020111946e-05 |
| Tsc1 | ENSMUSG00000026812 | 64930 | -0.492823754845664 | 5.92891172242327 | -7.15399506729427 | 6.2626434865881 | 6.56976544643902e-05 |
| Hes6 | ENSMUSG00000067071 | 55927 | 0.972932998959424 | 2.13761066627497 | 7.14906702096374 | 6.3094117326414 | 6.61248127032628e-05 |

| | | | | | | | |
|---|---|---|---|---|---|---|---|
| Raph1 | ENSMUSG00000026014 | 77300 | 0.580406419237646 | 6.07367080075735 | 7.14807954744728 | 6.3188273958126 | 6.61600596394906e-05 |
| B4galnt1 | ENSMUSG00000006731 | 14421 | 1.45223636017812 | 1.11807107679573 | 7.14501900665872 | 6.3481044475669 | 6.64029949438801e-05 |
| Tlr13 | ENSMUSG00000033777 | 279572 | 2.31217891276235 | 0.29562157047278 | 7.14254282336426 | 6.3718964812083 | 6.65881457323981e-05 |
| Hcls1 | ENSMUSG00000022831 | 15163 | 1.14789586504196 | 1.49183594512724 | 7.13223983425646 | 6.4719078776808 | 6.75686962855106e-05 |
| Shc1 | ENSMUSG00000042626 | 20416 | 0.647901839997126 | 3.75991950692762 | 7.12706837103728 | 6.5227315972150 | 6.80339553685451e-05 |
| Ndufab1 | ENSMUSG00000030869 | 70316 | -0.659372301636813 | 7.21421047734812 | -7.12644164265676 | 6.5289195116278 | 6.80339553685451e-05 |
| Ndufs2 | ENSMUSG00000013593 | 226646 | -0.621859916302149 | 8.08590895046863 | -7.12138118302378 | 6.5791111372801 | 6.84328445653249e-05 |
| Slc22a4 | ENSMUSG00000020334 | 30805 | 1.29236086556969 | 1.54982532379931 | 7.12132004364914 | 6.5797200291058 | 6.84328445653249e-05 |
| Itga11 | ENSMUSG00000032243 | 319480 | 1.29383416748996 | 3.676797900206 | 7.11706883471242 | 6.6222045533412 | 6.88092376165149e-05 |
| Npr2 | ENSMUSG00000028469 | 230103 | -0.445614182025031 | 5.29105384973747 | -7.11331376804387 | 6.6599720465748 | 6.91359491368561e-05 |
| Stn1 | ENSMUSG00000042694 | 108689 | 0.795785783875776 | 2.08063945057344 | 7.11075153864086 | 6.6858729087600 | 6.93389722634311e-05 |
| 2210408F21Rik | ENSMUSG00000087380 | 73652 | -0.687914119245862 | 3.24110128815446 | -7.1056470987974 | 6.7377900968065 | 6.98111692399927e-05 |
| Armcx2 | ENSMUSG00000033436 | 67416 | 1.01742741009409 | 1.93947504537377 | 7.10389197767031 | 6.7557397065260 | 6.99308624356399e-05 |
| Bmp6 | ENSMUSG00000039004 | 12161 | -0.630092688842933 | 3.432448070557 | -7.09916355338363 | 6.8043491989815 | 7.03673993321355e-05 |
| Hmox1 | ENSMUSG00000005413 | 15368 | 1.26469627781865 | 1.86269592408712 | 7.09848211962644 | 6.8113849409851 | 7.03735810868707e-05 |
| Atp5a1 | ENSMUSG00000025428 | 11946 | -0.659460656580096 | 10.1557073549435 | -7.09418536940033 | 6.8559259523735 | 7.0766880628324e-05 |
| Cep170 | ENSMUSG00000057335 | 545389 | 0.595996210978741 | 4.39213995200322 | 7.08890617159887 | 6.9110730400994 | 7.1268810755969e-05 |
| Micall1 | ENSMUSG00000033039 | 27008 | 0.766838712485809 | 5.59581575410686 | 7.08704179228219 | 6.9306603990341 | 7.14034390403792e-05 |
| Eid1 | ENSMUSG00000091337 | 58521 | 0.572585328545527 | 5.41449629210618 | 7.08541806584014 | 6.9477672321752 | 7.15122821232654e-05 |
| Gstm7 | ENSMUSG00000004035 | 68312 | -1.02118381558866 | 1.8912660498513 | -7.08439434975291 | 6.9585755633223 | 7.15561519122074e-05 |
| Gnal | ENSMUSG00000024524 | 14680 | 1.60210598788267 | 1.66957312514486 | 7.07883987624083 | 7.0175299211326 | 7.20432996064999e-05 |
| Npdc1 | ENSMUSG00000015094 | 18146 | 0.925589426653748 | 3.06524787265775 | 7.07868977579886 | 7.0191303705902 | 7.20432996064999e-05 |
| Hectd2os | ENSMUSG00000087579 | 668215 | 0.719231736734683 | 3.56558362759401 | 7.07468669236434 | 7.0619559669921 | 7.24148599204423e-05 |
| Adgrf5 | ENSMUSG00000056492 | 224792 | -0.886363067147499 | 6.7524596519768 | -7.07376755679763 | 7.0718279419876 | 7.24481267421433e-05 |
| Ugcg | ENSMUSG00000028381 | 22234 | 0.781532690335007 | 3.30526996003934 | 7.07060244180291 | 7.1059345799325 | 7.27293734955451e-05 |
| Copg1 | ENSMUSG00000030058 | 54161 | 0.457807369970276 | 5.22637272989478 | 7.06894691208397 | 7.1238434290507 | 7.28444644742315e-05 |
| Ndufv3 | ENSMUSG00000024038 | 78330 | -0.580623212079337 | 5.9290181513372 | -7.06599551284388 | 7.1558888766940 | 7.30466992221043e-05 |
| Gm12295 | ENSMUSG00000085162 | 327946 | 1.39638515385393 | 1.2263944331943 | 7.06589470716349 | 7.1569860824145 | 7.30466992221043e-05 |
| Pld4 | ENSMUSG00000052160 | 104759 | 1.3670571968624 | 2.06203793771427 | 7.06330505178667 | 7.1852338635121 | 7.32665964198242e-05 |
| 1700025G04Rik | ENSMUSG00000032666 | 69399 | 0.429019475783897 | 5.67490833918118 | 7.06118346835453 | 7.2084637376715 | 7.34349646938374e-05 |
| Timm23 | ENSMUSG00000013701 | 53600 | -0.549470841381037 | 6.31997629273128 | -7.05301362763205 | 7.2986617095375 | 7.42846100064753e-05 |
| Ddost | ENSMUSG00000028757 | 13200 | 0.501053045000963 | 4.14946012383678 | 7.05094187960501 | 7.3217235796651 | 7.43979836631713e-05 |
| Ubr2 | ENSMUSG00000023977 | 224826 | -0.400014122458438 | 6.36165506711324 | -7.0504071058659 | 7.3276889606146 | 7.43979836631713e-05 |
| Rps6ka2 | ENSMUSG00000023809 | 20112 | -0.600352623810956 | 6.90327259117284 | -7.05018039934186 | 7.3302194131584 | 7.43979836631713e-05 |
| Prkcd | ENSMUSG00000021948 | 18753 | 0.939571070567659 | 2.58393967370149 | 7.04824285438281 | 7.3518836022188 | 7.45486453208294e-05 |
| Rab12 | ENSMUSG00000023460 | 19328 | -0.522578689365145 | 6.62203343274993 | -7.04510843643821 | 7.3870737010926 | 7.48360543342386e-05 |
| Cdr2 | ENSMUSG00000030878 | 12585 | 0.960909280563902 | 2.88487076530712 | 7.04153059471862 | 7.4274597465542 | 7.51755208236897e-05 |
| Plcd4 | ENSMUSG00000026173 | 18802 | -1.60037433531095 | 1.89691585662324 | -7.03116863017916 | 7.5457454115055 | 7.63020750168061e-05 |
| Rbms3 | ENSMUSG00000039607 | 207181 | 0.807071297264244 | 3.59947188694459 | 7.02998248170033 | 7.5594121423053 | 7.63696248868204e-05 |
| Chchd2 | ENSMUSG00000070493 | 14004 | -0.515826039016519 | 6.92949189108846 | -7.02316388679231 | 7.6384845124817 | 7.70972060996659e-05 |
| P2ry1 | ENSMUSG00000027765 | 18441 | -0.929832807171127 | 2.74466744379529 | -7.0197898439133 | 7.6779346008052 | 7.74238958684523e-05 |
| Bnip3 | ENSMUSG00000078566 | 12176 | -0.831895574140358 | 7.77000069690296 | -7.01888014061516 | 7.6886078623320 | 7.74600668600475e-05 |
| Tor3a | ENSMUSG00000060519 | 30935 | 0.758329816034877 | 3.25248103641413 | 7.01779514303211 | 7.7013582963509 | 7.75170787637309e-05 |
| Kdr | ENSMUSG00000062960 | 16542 | -1.04964722486355 | 5.94580930838207 | -7.0171654543609 | 7.7087683744576 | 7.75202825217999e-05 |
| Mgp | ENSMUSG00000030218 | 17313 | 1.36722121742003 | 5.84661173135921 | 7.01467163238407 | 7.7381893510040 | 7.77446211358691e-05 |
| Nipsnap2 | ENSMUSG00000029432 | 14467 | -0.574902042544523 | 8.15810645227627 | -7.01367846945951 | 7.7499392420457 | 7.77463228412277e-05 |
| Plod2 | ENSMUSG00000032374 | 26432 | 1.86336926471 | 2.86060209884915 | 7.0134551692606 | 7.7525836517188 | 7.77463228412277e-05 |
| Nbl1 | ENSMUSG00000041120 | 17965 | 1.14032554584534 | 4.16904139620309 | 7.01136229699115 | 7.7774146609203 | 7.79238493661968e-05 |
| Klc1 | ENSMUSG00000021288 | 16593 | 0.429917903851468 | 5.80467741918502 | 7.00704253462228 | 7.8289328044187 | 7.82972826979528e-05 |
| Tmbim1 | ENSMUSG00000006301 | 69660 | 0.460825704148013 | 5.1437154112091 | 7.00703591866535 | 7.8290119832460 | 7.82972826979528e-05 |
| Creb3l2 | ENSMUSG00000038648 | 208647 | 0.733100297717962 | 4.82170630122072 | 7.00514495289105 | 7.8516775029672 | 7.83868356927713e-05 |
| Cx3cr1 | ENSMUSG00000052336 | 13051 | 1.67454997414534 | 1.43474894496445 | 7.00509238522998 | 7.8523085795978 | 7.83868356927713e-05 |
| Serpine2 | ENSMUSG00000026249 | 20720 | -0.403282768410377 | 5.77404600407276 | -7.00351888286239 | 7.8712233589964 | 7.85039621689695e-05 |
| Selplg | ENSMUSG00000048163 | 20345 | 1.3896841693301 | 1.46714769179872 | 7.00266288464131 | 7.8815333628332 | 7.85351332626524e-05 |
| 2510002D24Rik | ENSMUSG00000071632 | 72307 | -0.646235443827412 | 3.53777202523766 | -6.99681400957733 | 7.9523625420684 | 7.91687385677138e-05 |
| Hspe1 | ENSMUSG00000073676 | 15528 | -0.59359333250639 | 4.87634524030784 | -6.99412793567962 | 7.9851155874176 | 7.94224735996924e-05 |
| Map2k6 | ENSMUSG00000020623 | 26399 | -0.898747323075269 | 3.97209614223587 | -6.99305174435624 | 7.9982782440879 | 7.94810722601138e-05 |
| Zc3h8 | ENSMUSG00000027387 | 57432 | -0.923639032297965 | 1.69797774556187 | -6.99096621207182 | 8.0238512120253 | 7.96627771104891e-05 |
| Mfap4 | ENSMUSG00000042436 | 76293 | 1.51542297704694 | 4.20888879335095 | 6.98606494175722 | 8.0842915974397 | 8.01900103916644e-05 |
| Cmss1 | ENSMUSG00000022748 | 66497 | -1.09011019677413 | 2.08028722499928 | -6.98136960344189 | 8.1426436709766 | 8.06955919922442e-05 |
| Fam214b | ENSMUSG00000036002 | 230088 | 0.568797296873071 | 4.38956990779083 | 6.98024516968867 | 8.1566836724619 | 8.07615119779728e-05 |
| Spred1 | ENSMUSG00000027351 | 114715 | 0.659099059153675 | 4.26280584548537 | 6.97628020247642 | 8.2063957914688 | 8.11646383212101e-05 |
| Alas2 | ENSMUSG00000025270 | 11656 | -1.46655935684398 | 1.88076003831251 | -6.97581505701574 | 8.2122486490950 | 8.11646383212101e-05 |
| Crybg3 | ENSMUSG00000022723 | 224273 | -0.501961698090825 | 6.58076944279354 | -6.97468135281372 | 8.2265323608529 | 8.12323624539144e-05 |
| Cbfb | ENSMUSG00000031885 | 12400 | 0.522427545482159 | 4.1280283478957 | 6.97207822906488 | 8.2594289052413 | 8.14835896779726e-05 |
| Vegfa | ENSMUSG00000023951 | 22339 | -1.22849543695051 | 7.16830586397519 | -6.97095057132304 | 8.2737225979520 | 8.15510024510492e-05 |
| Fmn1 | ENSMUSG00000044042 | 14260 | 1.44489156648269 | 0.959840309704387 | 6.96940774661829 | 8.2933211047234 | 8.16705342303894e-05 |
| Ccbe1 | ENSMUSG00000046318 | 320924 | 1.88851936130537 | 1.40187902738318 | 6.96667231603857 | 8.3281899226955 | 8.19400936498513e-05 |
| Pde5a | ENSMUSG00000053965 | 242202 | 0.886512513170488 | 3.48208926299978 | 6.95613681792647 | 8.4639404361246 | 8.32008389453945e-05 |
| A930003A15Rik | ENSMUSG00000075330 | 68162 | 3.3390347584633 | -0.300897062389519 | 6.95410046309234 | 8.4904474361644 | 8.33864159251691e-05 |
| Atp5b | ENSMUSG00000025393 | 11947 | -0.5577763133139 | 10.0328406727432 | -6.95090781607958 | 8.5321823883732 | 8.36317371029409e-05 |
| Asnsd1 | ENSMUSG00000026095 | 70396 | -0.520695975382032 | 5.84201620088105 | -6.95059086403296 | 8.5363374582300 | 8.36317371029409e-05 |
| ND1 | ENSMUSG00000064341 | 17716 | -0.756431231177404 | 12.2920419597814 | -6.95043520892435 | 8.5383787948844 | 8.36317371029409e-05 |
| Ginm1 | ENSMUSG00000040006 | 215751 | -0.685563091498707 | 5.45584352131406 | -6.94692313472797 | 8.5845751515801 | 8.40089444779974e-05 |
| Tgtp2 | ENSMUSG00000078921 | 100039796 | -1.48028595166201 | 3.14154229816336 | -6.94187798020234 | 8.6513994963570 | 8.45871626964923e-05 |
| Epb41 | ENSMUSG00000028906 | 269587 | -0.567521405794219 | 5.5946062789645 | -6.93839066610714 | 8.6979107978524 | 8.49357203111202e-05 |
| Mrps36 | ENSMUSG00000061474 | 66128 | -0.636018656384028 | 4.67135721301332 | -6.93748874367569 | 8.7099829233613 | 8.49357203111202e-05 |
| Csrp2 | ENSMUSG00000020186 | 13008 | 1.61804511204237 | 1.91898059745184 | 6.93746061148987 | 8.7103597537979 | 8.49357203111202e-05 |
| Fkbp4 | ENSMUSG00000030357 | 14228 | -0.429309419222503 | 5.79818435092414 | -6.93647259683748 | 8.7236051297139 | 8.49890620970611e-05 |
| Fahd2a | ENSMUSG00000027371 | 68126 | -0.679336542515616 | 2.35547726659053 | -6.93215695334654 | 8.7817107305786 | 8.54535675735077e-05 |
| Fam210a | ENSMUSG00000038121 | 108654 | -0.645287995072837 | 7.12366521045629 | -6.93177160718991 | 8.7869188502993 | 8.54535675735077e-05 |
| Tango2 | ENSMUSG00000013539 | 27883 | -0.738697039233288 | 3.60868465502383 | -6.92938690371206 | 8.8192217834350 | 8.56914785019809e-05 |
| Acot7 | ENSMUSG00000028937 | 70025 | -0.555245287207868 | 3.3241404623296 | -6.92773488367235 | 8.8416733798884 | 8.58333318965904e-05 |
| Gata2 | ENSMUSG00000015053 | 14461 | -1.12602157702851 | 2.43367866789115 | -6.92601890902978 | 8.8650580625264 | 8.59506300504666e-05 |
| Zfp641 | ENSMUSG00000022987 | 239652 | 0.754709363574212 | 3.58584177906296 | 6.92569479715306 | 8.8694822703253 | 8.59506300504666e-05 |
| ATP6 | ENSMUSG00000064357 | 17705 | -0.768324474639744 | 12.3575961450874 | -6.92394988585806 | 8.8933408423787 | 8.61054993339612e-05 |
| Dcaf11 | ENSMUSG00000022214 | 28199 | -0.563444335575137 | 6.27614943686564 | -6.92295098753681 | 8.9070295026697 | 8.61617163660913e-05 |
| Fitm2 | ENSMUSG00000048486 | 228859 | -0.426034397858885 | 6.06674384655775 | -6.92178660700702 | 8.9230139790179 | 8.61786064766171e-05 |
| Dbn1 | ENSMUSG00000034675 | 56320 | 1.09410535952735 | 3.54865338926521 | 6.9216753217827 | 8.9245432743143 | 8.61786064766171e-05 |
| Cyc1 | ENSMUSG00000022551 | 66445 | -0.555022610008083 | 7.20432167065999 | -6.91690355484202 | 8.9903785458853 | 8.67377121668778e-05 |
| Hs3st5 | ENSMUSG00000044499 | 319415 | -1.36803665853931 | 1.39636264352314 | -6.9153852635388 | 9.0114335474725 | 8.68641799889084e-05 |
| Clec7a | ENSMUSG00000079293 | 56644 | 1.99972496408788 | 0.721667449615165 | 6.91342579623721 | 9.0386835747440 | 8.70500882427555e-05 |
| Serpinb8 | ENSMUSG00000026315 | 20725 | 1.22568150207738 | 1.89276686107237 | 6.90903806105297 | 9.1000192255269 | 8.75636533047841e-05 |
| Cd14 | ENSMUSG00000051439 | 12475 | 1.77107700577547 | 0.59328327512872 | 6.90778130054252 | 9.1176681909930 | 8.76563157394415e-05 |
| Ppp5c | ENSMUSG00000003099 | 19060 | -0.41857742754139 | 5.0982838438231 | -6.90357433160037 | 9.1770110460263 | 8.81493038173233e-05 |
| Atp5d | ENSMUSG00000003072 | 66043 | -0.505729064896137 | 7.38062227042279 | -6.90248949135777 | 9.1923796814153 | 8.82194050022399e-05 |
| Rab43 | ENSMUSG00000030055 | 69834 | 0.837319834951576 | 3.04570687036828 | 6.90156605346006 | 9.2054831853852 | 8.82676637714441e-05 |
| Cox7a2 | ENSMUSG00000032330 | 12866 | -0.595235431422714 | 5.23683807192513 | -6.89932976676332 | 9.2372976796731 | 8.84950928453173e-05 |
| Myom3 | ENSMUSG00000037139 | 242702 | 0.694553845482746 | 9.19289971747011 | 6.89427403090644 | 9.3096517001873 | 8.91101600129138e-05 |
| P3h1 | ENSMUSG00000028641 | 56401 | 0.78613619695658 | 2.98915179155768 | 6.89366537529057 | 9.3184026075276 | 8.91158870541427e-05 |
| Col6a1 | ENSMUSG00000001119 | 12833 | 0.996506970880911 | 6.56739926550289 | 6.89280813516811 | 9.3307422553245 | 8.91558947490845e-05 |
| Pdk3 | ENSMUSG00000035232 | 236900 | 1.37034557789767 | 1.14357530425188 | 6.8916362666169 | 9.3476387801763 | 8.92393358131944e-05 |
| Cdc37l1 | ENSMUSG00000024780 | 67072 | -0.460524587825009 | 5.94209301246599 | -6.88993040741011 | 9.3722923833587 | 8.93966213285295e-05 |
| Apobec1 | ENSMUSG00000040613 | 11810 | 1.33948632265476 | 1.47955782925527 | 6.8888039997605 | 9.3886091986852 | 8.94741823459711e-05 |
| Mid2 | ENSMUSG00000000266 | 23947 | 0.604573884768527 | 3.38734133897396 | 6.88662379816162 | 9.4202762321201 | 8.96977695934712e-05 |
| Zbtb44 | ENSMUSG00000047412 | 235132 | -0.533002502048251 | 6.05858208366909 | -6.88512408146133 | 9.4421247977709 | 8.98275597601695e-05 |
| Rb1 | ENSMUSG00000022105 | 19645 | 0.525513594920925 | 4.28521549350631 | 6.88411203395159 | 9.4568990020486 | 8.98898808620815e-05 |
| Tceal8 | ENSMUSG00000051579 | 66684 | 0.558465406591927 | 3.38379055650473 | 6.88196162844373 | 9.4883724176727 | 9.00340408673704e-05 |
| Ptcd3 | ENSMUSG00000063884 | 69956 | -0.428384458476367 | 5.03348040733697 | -6.88195029634523 | 9.4885385673049 | 9.00340408673704e-05 |
| Dkk3 | ENSMUSG00000030772 | 50781 | 1.91195934801186 | 1.6871176607187 | 6.88040651622903 | 9.5112019913485 | 9.01708143689773e-05 |
| Enpp2 | ENSMUSG00000022425 | 18606 | -0.703053649525052 | 3.99233787924459 | -6.87973370465357 | 9.5210970586817 | 9.01864055012567e-05 |
| Hspa2 | ENSMUSG00000059970 | 15512 | 0.759846093306704 | 4.29479958549281 | 6.87660538330604 | 9.5672483646648 | 9.05451011897415e-05 |
| Olfm1 | ENSMUSG00000026833 | 56177 | 0.94615146348521 | 3.77396386122913 | 6.87530364002128 | 9.5865221737329 | 9.0570697571609e-05 |
| Mrps33 | ENSMUSG00000029918 | 14548 | -0.524570680127 | 3.5924697730507 | -6.87530349677012 | 9.5865242969857 | 9.0570697571609e-05 |
| Sos2 | ENSMUSG00000034801 | 20663 | -0.460319362080297 | 5.47128194083681 | -6.87266539830195 | 9.6257102182392 | 9.08623820341736e-05 |
| Capns1 | ENSMUSG00000001794 | 12336 | 0.362327866945553 | 6.16053312548906 | 6.87168347099831 | 9.6403387727717 | 9.09219526532943e-05 |

| | | | | | | | |
|---|---|---|---|---|---|---|---|
| Fsd2 | ENSMUSG00000038663 | 244091 | -0.575607890907344 | 7.5604183087641 | -6.87013512432286 | 9.6634534305408 | 9.10613874562427e-05 |
| Zfp36l2 | ENSMUSG00000045817 | 12193 | 0.912932145938936 | 4.59683992560179 | 6.85941887976923 | 9.825044480638 | 9.25043593607705e-05 |
| Prdx6 | ENSMUSG00000026701 | 11758 | -0.448760987152999 | 5.64447898683415 | -6.85400579588323 | 9.907750817990 | 9.31705891635564e-05 |
| Rasa1 | ENSMUSG00000021549 | 218397 | 0.449418184433536 | 4.74883316599107 | 6.853673430114 | 9.912852913476 | 9.31705891635564e-05 |
| St3gal3 | ENSMUSG00000028538 | 20441 | -0.487038085382618 | 4.93096237187292 | -6.85101452706514 | 9.953769302049 | 9.3458880451593e-05 |
| Adamts5 | ENSMUSG00000022894 | 23794 | 0.659066908031281 | 4.18767878800872 | 6.85057012355034 | 9.960625352310 | 9.3458880451593e-05 |
| Gm20300 | ENSMUSG00000111080 | 100504586 | -0.619117491682047 | 4.09652494010517 | -6.84977387777371 | 9.972921931102 | 9.34940048275105e-05 |
| Dapp1 | ENSMUSG00000028159 | 26377 | -1.25972692901638 | 1.76232910622168 | -6.84715509106205 | 1.001347750836 | 9.37937640479595e-05 |
| Myl3 | ENSMUSG00000059741 | 17897 | -0.81201026823188 | 10.4731111938688 | -6.84659851214539 | 1.002211932183 | 9.37943375916373e-05 |
| Gmnn | ENSMUSG00000006715 | 57441 | -1.9560843737209 | 1.03933164047554 | -6.84462122654894 | 1.005288368661 | 9.40017720944244e-05 |
| Polq | ENSMUSG00000034206 | 77782 | -1.45106115684985 | 1.18754755411184 | -6.84383877091338 | 1.006508534013 | 9.4035425515393e-05 |
| Mmachc | ENSMUSG00000028690 | 67096 | -0.582376163493016 | 4.21135585952074 | -6.84147162976495 | 1.010209393659 | 9.43005882330793e-05 |
| Rhobtb1 | ENSMUSG00000019944 | 69288 | -0.712094698871118 | 4.8631409187472 | -6.84007151479394 | 1.012405133559 | 9.43505411623221e-05 |
| Saraf | ENSMUSG00000031532 | 67887 | -0.419456751212809 | 5.63650003704579 | -6.84002968239385 | 1.012470814961 | 9.43505411623221e-05 |
| Tagap1 | ENSMUSG00000052031 | 380608 | -0.657558660494087 | 3.49584381990838 | -6.83381442134959 | 1.022279603806 | 9.51169108389146e-05 |
| Itm2c | ENSMUSG00000026223 | 64294 | 0.767885389048069 | 3.3426686466836 | 6.83371646079061 | 1.022435003528 | 9.51169108389146e-05 |
| E2f6 | ENSMUSG00000057469 | 50496 | -0.491054808176164 | 6.26430200203511 | -6.83224545879548 | 1.024771528129 | 9.52532106631418e-05 |
| Afg1l | ENSMUSG00000038302 | 215951 | -0.543889541902598 | 3.39897816848926 | -6.82965898441645 | 1.028893535028 | 9.55550996720026e-05 |
| Map1lc3b | ENSMUSG00000031812 | 67443 | -0.413990496074663 | 6.26970485629191 | -6.82906753018788 | 1.029838578697 | 9.55616766021824e-05 |
| Col1a1 | ENSMUSG00000001506 | 12842 | 1.39332333571393 | 9.40044118965437 | 6.82501713305239 | 1.036335097935 | 9.60829427950128e-05 |
| Ttc7b | ENSMUSG00000033530 | 104718 | -0.486676817647351 | 6.36990242571444 | -6.82021098181928 | 1.044099945084 | 9.67208177942632e-05 |
| Dele1 | ENSMUSG00000024442 | 66839 | -0.538170020394082 | 5.51124900681572 | -6.81865454859769 | 1.046627671318 | 9.68728795527524e-05 |
| Apobec2 | ENSMUSG00000040694 | 11811 | 0.629415948832843 | 8.60858549752897 | 6.81809898302103 | 1.047531501371 | 9.68745079652866e-05 |
| Eya4 | ENSMUSG00000010461 | 14051 | 0.725055039469784 | 3.0717454572418 | 6.81419726225995 | 1.053902326750 | 9.73812876898241e-05 |
| Oxsr1 | ENSMUSG00000036737 | 108737 | -0.409884313314996 | 5.54948093167893 | -6.81305396007201 | 1.055776875734 | 9.74721032825599e-05 |
| Cd163 | ENSMUSG00000008845 | 93671 | 1.00798233871343 | 3.33826872083436 | 6.80486901401004 | 1.069300044669 | 9.86372893526251e-05 |
| Atf6 | ENSMUSG00000026663 | 226641 | 0.468794233395349 | 5.7287204253218 | 6.79895753356201 | 1.079180615322 | 9.94647833566218e-05 |
| Entpd7 | ENSMUSG00000025192 | 93685 | 1.2298066105934 | 0.930151021405429 | 6.79742712192466 | 1.081754246000 | 9.95613053005391e-05 |
| Timm44 | ENSMUSG00000002949 | 21856 | -0.512182219981199 | 5.50855872433551 | -6.79725179542883 | 1.082049498646 | 9.95613053005391e-05 |
| Pttg1 | ENSMUSG00000020415 | 30939 | -0.726010268806691 | 3.7292046142621 | -6.79605319341092 | 1.084070243128 | 9.96633459011121e-05 |
| Xbp1 | ENSMUSG00000020484 | 22433 | 0.602687344273758 | 5.13105481846328 | 6.79544735465531 | 1.085093152315 | 9.96735567055599e-05 |
| Nsmaf | ENSMUSG00000028245 | 18201 | -0.464330678106865 | 4.617299991152 | -6.79362262907342 | 1.088180210758 | 9.98731980167795e-05 |
| Klhl33 | ENSMUSG00000090799 | 546611 | -1.4415237224469 | 4.50086287350039 | -6.79016450601051 | 1.094056093830 | 0.000100328248000484 |
| Dnaja3 | ENSMUSG00000004069 | 83945 | -0.507565722048154 | 6.18581546960777 | -6.78224349627575 | 1.107641649275 | 0.000101488942734501 |
| Flt1 | ENSMUSG00000029648 | 14254 | -0.67340754608253 | 6.08081891187601 | -6.77936714588111 | 1.112618922363 | 0.000101859610053231 |
| C530008M17Rik | ENSMUSG00000036377 | 320827 | -0.98297273788094 | 2.6203144728769 | -6.77728290391286 | 1.116240264464 | 0.000102105626199715 |
| Creb5 | ENSMUSG00000053007 | 231991 | 0.929016000774288 | 3.6198201038348 | 6.77417989776872 | 1.121654755091 | 0.000102515118126252 |
| Tcaf1 | ENSMUSG00000036667 | 77574 | 0.844083034084584 | 3.00917698516506 | 6.77185690800802 | 1.125726317026 | 0.000102801289652589 |
| Ptafr | ENSMUSG00000056529 | 19204 | 1.40755673203884 | 1.04734287326557 | 6.77103911534975 | 1.127163391506 | 0.000102846602942895 |
| Tbc1d10b | ENSMUSG00000042492 | 68449 | 0.434517483956519 | 4.82134346702297 | 6.76826304540026 | 1.132056124459 | 0.000103091843019041 |
| Slc25a42 | ENSMUSG00000002346 | 73095 | -0.748629311098717 | 3.88675480471203 | -6.76808960794417 | 1.132362545151 | 0.000103091843019041 |
| Cd276 | ENSMUSG00000035914 | 102657 | 0.880319223045326 | 2.66447673233531 | 6.7673266556505 | 1.133711531023 | 0.000103091843019041 |
| Krt18 | ENSMUSG00000023043 | 16668 | 4.65038341439221 | -0.458300873812413 | 6.76663382229675 | 1.134938007789 | 0.000103091843019041 |
| Tab2 | ENSMUSG00000015755 | 68652 | -0.484747603796259 | 6.09697475677165 | -6.76639981849326 | 1.135352565354 | 0.000103091843019041 |
| Gbp6 | ENSMUSG00000104713 | 100702 | -1.46019167039489 | 5.87955513003153 | -6.76631107417321 | 1.135509825221 | 0.000103091843019041 |
| Col1a2 | ENSMUSG00000029661 | 12843 | 1.2521733718831 | 8.99693574782272 | 6.76448253028855 | 1.138755227339 | 0.000103300692033614 |
| Timm21 | ENSMUSG00000024645 | 67105 | -0.700560087475268 | 3.50537021185187 | -6.76364868092676 | 1.140238437694 | 0.000103349472325352 |
| Apoo | ENSMUSG00000079508 | 68316 | -0.711184297927656 | 4.81747992328946 | -6.76219612024922 | 1.142827051595 | 0.000103498280869857 |
| 4930511M06Rik | ENSMUSG00000086607 | 75084 | -1.23448414695336 | 2.40132185079408 | -6.75987538897407 | 1.146975710885 | 0.000103788009070271 |
| As3mt | ENSMUSG00000003559 | 57344 | -0.701850827139092 | 3.94094003530018 | -6.75856853007141 | 1.149318910303 | 0.000103914019921617 |
| Got1 | ENSMUSG00000025190 | 14718 | -0.669670202128811 | 7.46345764959159 | -6.75667512457042 | 1.152722752218 | 0.000104135639706605 |
| Bak1 | ENSMUSG00000057789 | 12018 | 0.76923942103831 | 3.230317613695 | 6.75557016821584 | 1.154714084529 | 0.000104229394368263 |
| Mpc1 | ENSMUSG00000023861 | 55951 | -0.690281652346296 | 7.51649346851579 | -6.75301459033209 | 1.159333618273 | 0.000104560031199258 |
| Tnfrsf1a | ENSMUSG00000030341 | 21937 | 0.474568489924476 | 4.54875625538037 | 6.75223797773748 | 1.160741304723 | 0.000104600685918687 |
| Polr2m | ENSMUSG00000032199 | 28015 | -0.496970055786083 | 6.96653075443068 | -6.74963529889122 | 1.165472078815 | 0.000104940488414568 |
| Mmaa | ENSMUSG00000037022 | 109136 | -0.599747972178505 | 2.93212837712183 | -6.74469305347225 | 1.174511457362 | 0.000105600828574461 |
| Sgpl1 | ENSMUSG00000020097 | 20397 | 0.72674127912709 | 3.61304983845386 | 6.74456972428665 | 1.174737970419 | 0.000105600828574461 |
| Eci2 | ENSMUSG00000021417 | 23986 | -0.691624390899665 | 5.07280388660566 | -6.74247788653903 | 1.178586981268 | 0.000105859772327368 |
| Usp20 | ENSMUSG00000026854 | 74270 | 0.732660040279428 | 3.96827013241524 | 6.74100150487217 | 1.181311551829 | 0.000105953004364945 |
| Kiz | ENSMUSG00000074749 | 228730 | -0.5698193759807 | 4.88691815488986 | -6.74036917144172 | 1.182480515583 | 0.000105953004364945 |
| Serinc5 | ENSMUSG00000021703 | 218442 | 1.03763909302176 | 2.72581329822365 | 6.74034087986231 | 1.182532845350 | 0.000105953004364945 |
| Pbxip1 | ENSMUSG00000042613 | 229534 | 0.4577824212561 | 7.19108421765224 | 6.73413438001586 | 1.194072020992 | 0.000106899273230669 |
| Clybl | ENSMUSG00000025545 | 69634 | -0.645459315241725 | 3.7017446644836 | -6.73208693048307 | 1.197904664265 | 0.000107154630810875 |
| Shc4 | ENSMUSG00000035109 | 271849 | 1.18731998688454 | 1.22446726753992 | 6.72905099135201 | 1.203611578324 | 0.000107577090455119 |
| Ndrg2 | ENSMUSG00000004558 | 29811 | -0.571533804330635 | 9.09425839039088 | -6.72602283572106 | 1.209332439847 | 0.000107954041020985 |
| Mrps5 | ENSMUSG00000027374 | 77721 | -0.555448961238437 | 4.88106940027499 | -6.72577378410657 | 1.209804228805 | 0.000107954041020985 |
| Serp1 | ENSMUSG00000027808 | 28146 | 0.727635260098694 | 3.89688589527454 | 6.71850838543895 | 1.223653178968 | 0.000109038884357935 |
| Lonp2 | ENSMUSG00000047866 | 66887 | -0.452965840436792 | 5.01270833979622 | -6.71835010581903 | 1.223956738698 | 0.000109038884357935 |
| Rtkn | ENSMUSG00000034930 | 20166 | 0.587170368827322 | 2.72581542769294 | 6.71682906648039 | 1.226877947317 | 0.000109210120864202 |
| Emb | ENSMUSG00000021728 | 13723 | 1.83077654363372 | 1.80024222503013 | 6.71295489637597 | 1.234351696883 | 0.000109785991852147 |
| Plat | ENSMUSG00000031538 | 18791 | 1.16708515880286 | 2.88438079018253 | 6.70822528432527 | 1.243540854269 | 0.000110513374618067 |
| Scai | ENSMUSG00000035236 | 320271 | 0.952996781358099 | 2.25354277209968 | 6.70521715918477 | 1.249422856689 | 0.000110945907769922 |
| Ndufa7 | ENSMUSG00000041881 | 66416 | -0.571091977013233 | 4.61900550755716 | -6.70365276169624 | 1.252493427913 | 0.000111128292699042 |
| H2-DMb1 | ENSMUSG00000079547 | 14999 | 2.03441905189009 | 0.916429380058371 | 6.69994877799247 | 1.259795317406 | 0.000111685503767778 |
| Zbtb18 | ENSMUSG00000063659 | 30928 | -0.66793475768814 | 5.69402575319512 | -6.69849033770372 | 1.262682738225 | 0.000111850769947667 |
| Numbl | ENSMUSG00000063160 | 18223 | 1.0302302718792 | 2.57104764490121 | 6.69694679847344 | 1.265746242408 | 0.00011192658614072 |
| Esyt2 | ENSMUSG00000021171 | 52635 | 0.38888987662166 | 5.32350235890878 | 6.69646906884983 | 1.266695993250 | 0.00011192658614072 |
| Ubtd1 | ENSMUSG00000025171 | 226122 | 0.727747712346151 | 2.50817210276482 | 6.69626985567282 | 1.267092261246 | 0.00011192658614072 |
| Ppip5k1 | ENSMUSG00000033526 | 327655 | -0.48453826443935 | 7.39748542012039 | -6.69599742833254 | 1.267634376015 | 0.00011192658614072 |
| Grhpr | ENSMUSG00000035637 | 76238 | -0.706079212206573 | 2.74330014156626 | -6.69318031270207 | 1.273254640509 | 0.00011233209423252 |
| Timm17a | ENSMUSG00000062580 | 21854 | -0.565479969014824 | 5.83264944966892 | -6.69046722895921 | 1.278692205709 | 0.000112720842746851 |
| Atp5g1 | ENSMUSG00000006057 | 11951 | -0.618696065926444 | 7.3243718484515 | -6.68593719453168 | 1.287825963821 | 0.0001134193651856 |
| Cygb | ENSMUSG00000020810 | 114886 | 0.76681477020362 | 3.97124099955357 | 6.68527026381012 | 1.289176474405 | 0.0001134193651856 |
| Rpl3 | ENSMUSG00000060036 | 27367 | 0.966325331946839 | 5.82994444436217 | 6.68499764653636 | 1.289728944522 | 0.0001134193651856 |
| Man1c1 | ENSMUSG00000037306 | 230815 | 1.08373932263072 | 3.20707411229807 | 6.68446961591273 | 1.290799730633 | 0.000113422281797022 |
| Pecam1 | ENSMUSG00000020717 | 18613 | -0.606631736444201 | 6.10607141447263 | -6.68018149665379 | 1.299530351948 | 0.00011409772110162 |
| Ccng1 | ENSMUSG00000020326 | 12450 | -0.555657098018433 | 8.65539238182302 | -6.67860748404736 | 1.302750652418 | 0.000114288662773532 |
| 1810055G02Rik | ENSMUSG00000035372 | 72056 | 0.695505274168978 | 2.607386446343 | 6.67443043303305 | 1.311337425088 | 0.000114949714463866 |
| Eno1 | ENSMUSG00000063524 | 13806 | 0.547653906607784 | 5.89659043139758 | 6.67328021783827 | 1.313712391433 | 0.000115065626208043 |
| Rbl2 | ENSMUSG00000031666 | 19651 | -0.487403506344368 | 5.49494787122216 | -6.66783833601514 | 1.325010458606 | 0.000115962284411745 |
| Cox11 | ENSMUSG00000020544 | 69802 | -0.604845846204852 | 2.7451254852915 | -6.66477879224764 | 1.331407435039 | 0.000116428917379312 |
| Aff2 | ENSMUSG00000031189 | 14266 | 1.77462504385082 | 0.710087560422365 | 6.66113525787277 | 1.339067947547 | 0.000117005209709356 |
| Tspan9 | ENSMUSG00000030352 | 109246 | 0.839791134686605 | 4.41489798724043 | 6.65979898590174 | 1.341889089963 | 0.000117158064236367 |
| Uqcr11 | ENSMUSG00000020163 | 66594 | -0.489095001823808 | 6.70632460298867 | -6.65699149568324 | 1.347836711207 | 0.000117583424502862 |
| Crlf1 | ENSMUSG00000007888 | 12931 | 1.69843675090386 | 2.39247661525273 | 6.65634455253834 | 1.349211186444 | 0.000117609469529716 |
| Frem1 | ENSMUSG00000059049 | 329872 | 1.56853147360645 | 1.54834551661273 | 6.65236134587943 | 1.357706413530 | 0.000118255687699651 |
| Trf | ENSMUSG00000032554 | 22041 | 0.904721210188057 | 3.29971720171154 | 6.65067837293277 | 1.361312726083 | 0.000118433625452738 |
| Ampd1 | ENSMUSG00000070385 | 229665 | -0.528296882289759 | 6.89826088305232 | -6.65034583045808 | 1.362026499983 | 0.000118433625452738 |
| Bckdhb | ENSMUSG00000032263 | 12040 | -0.844546266187001 | 3.5896649754022 | -6.64989270368127 | 1.362999733048 | 0.000118433625452738 |
| Cnpy3 | ENSMUSG00000023973 | 72029 | 0.699491540038999 | 3.11463434011655 | 6.64039988352504 | 1.383558062913 | 0.000120081965740276 |
| B4galt6 | ENSMUSG00000056124 | 56386 | 0.969773045991939 | 3.22993003557 | 6.64012105428309 | 1.384166835904 | 0.000120081965740276 |
| Fsd1l | ENSMUSG00000054752 | 319636 | 0.744902473213235 | 5.08575261390536 | 6.63508256089225 | 1.395216271855 | 0.000120863779809207 |
| Igsf3 | ENSMUSG00000042035 | 78908 | 1.17836005302416 | 2.18987067649358 | 6.63500363968995 | 1.395390084340 | 0.000120863779809207 |
| Dennd4a | ENSMUSG00000053641 | 102442 | 0.871577413305367 | 3.28147679707062 | 6.63135723227789 | 1.403445715310 | 0.000121465281979901 |
| Rrad | ENSMUSG00000031880 | 56437 | 2.02455781522626 | 5.20797224536833 | 6.62592474666098 | 1.415538210495 | 0.000122395227397577 |
| Mlf1 | ENSMUSG00000048416 | 17349 | -0.880673299896284 | 5.67721195836476 | -6.62547946503205 | 1.416534248902 | 0.000122395227397577 |
| Impdh1 | ENSMUSG00000003500 | 23917 | 0.646274081065076 | 3.20574370598685 | 6.62457826534059 | 1.418552379922 | 0.000122395227397577 |
| Nek7 | ENSMUSG00000026393 | 59125 | -0.373143032937455 | 6.22816843231603 | -6.62452604722801 | 1.418669409136 | 0.000122395227397577 |
| Atg4d | ENSMUSG00000002820 | 235040 | -0.532155763746565 | 4.25816828566272 | -6.62287683760746 | 1.422370796532 | 0.000122617785306765 |
| Mreg | ENSMUSG00000039395 | 381269 | -0.718486713279621 | 5.87140509274777 | -6.62017382029782 | 1.428459322657 | 0.000123045617462364 |
| Fkbp10 | ENSMUSG00000001555 | 14230 | 1.12400600756415 | 2.91760417919118 | 6.61635393204215 | 1.437110487229 | 0.000123606779595836 |
| Lrrc30 | ENSMUSG00000073375 | 240131 | -1.10426714829122 | 4.16451758733548 | -6.6162988934317 | 1.437235539898 | 0.000123606779595836 |
| Cfl1 | ENSMUSG00000056201 | 12631 | 0.568632376874396 | 5.4206004783994 | 6.60903933618156 | 1.453830881875 | 0.000124935734039171 |
| Slc39a6 | ENSMUSG00000024270 | 106957 | 0.787512816514879 | 2.06333654399305 | 6.60368482997107 | 1.466200667865 | 0.000125899760411942 |
| Arhgef40 | ENSMUSG00000004562 | 268739 | 0.689389570696815 | 4.28468644519765 | 6.6002047493142 | 1.474299676402 | 0.000126495838012191 |
| Sugt1 | ENSMUSG00000022024 | 67955 | -0.468603941084436 | 5.29718841965213 | -6.59899587810559 | 1.477124044054 | 0.000126638768043631 |

| | | | | | | | |
|---|---|---|---|---|---|---|---|
| Col16a1 | ENSMUSG00000040690 | 107581 | 1.34627733863872 | 4.10283986435298 | 6.59176727004161 | 1.4941321904659 | 0.000127933642382719 |
| Hikeshi | ENSMUSG00000062797 | 67669 | -0.6499103042515 | 2.81513697525449 | -6.59158306043736 | 1.4945683041142 | 0.000127933642382719 |
| Ghitm | ENSMUSG00000041028 | 66092 | -0.524647762680746 | 7.28667239075282 | -6.59004497704977 | 1.4982149368419 | 0.000128073547607453 |
| Csdc2 | ENSMUSG00000042109 | 105859 | 1.2543317836222 | 1.02019158015842 | 6.58990552016137 | 1.4985460377818 | 0.000128073547607453 |
| Rab27b | ENSMUSG00000024511 | 80718 | 1.71023197362814 | 0.558529810222475 | 6.58762271630078 | 1.5039768890835 | 0.000128437276363842 |
| Cdc42se2 | ENSMUSG00000052298 | 72729 | 0.433919602858918 | 4.41076274840147 | 6.58472048633179 | 1.5109113434838 | 0.000128928742354579 |
| Serpina3n | ENSMUSG00000021091 | 20716 | 4.54598023795376 | 2.01397742554573 | 6.58407288761632 | 1.5124632802576 | 0.000128960500128672 |
| Rnf114 | ENSMUSG00000006418 | 81018 | -0.546828412277212 | 3.95925094585498 | -6.58259130960317 | 1.5160201282339 | 0.000129163024331448 |
| Slc30a9 | ENSMUSG00000029221 | 109108 | -0.457211059601993 | 5.96272233775943 | -6.57805684150977 | 1.5269609903856 | 0.000129993851915155 |
| Pcdh19 | ENSMUSG00000051323 | 279653 | 0.560131107386393 | 3.35966724834895 | 6.57675607454099 | 1.5301148475434 | 0.000130160975863797 |
| Rpn2 | ENSMUSG00000027642 | 20014 | 0.411496394924437 | 4.87354295089102 | 6.57403119152176 | 1.5367439140728 | 0.000130623232696194 |
| Manba | ENSMUSG00000028164 | 110173 | 0.671994904802294 | 2.81345086541305 | 6.57199007927072 | 1.5417293374209 | 0.000130945170064866 |
| Stard10 | ENSMUSG00000030688 | 56018 | -0.631157744340177 | 4.49184383300885 | -6.56732490687987 | 1.5531881545370 | 0.000131815991593516 |
| Hpn | ENSMUSG00000001249 | 15451 | -1.29666647532415 | 2.17446315743833 | -6.56469854376968 | 1.5596786134578 | 0.000132264134396487 |
| Prkdc | ENSMUSG00000022672 | 19090 | 0.678813784996982 | 3.67684130077901 | 6.56247482911526 | 1.5651963665612 | 0.000132629158782025 |
| Asrgl1 | ENSMUSG00000024654 | 66514 | -0.515759815152344 | 4.57906630399941 | -6.55834061264911 | 1.5755094434940 | 0.000133399641571132 |
| Cdc34 | ENSMUSG00000020307 | 216150 | -0.450043160192026 | 5.67619125571045 | -6.55774936203466 | 1.5769902008963 | 0.000133421670944256 |
| Lpar1 | ENSMUSG00000038668 | 14745 | 1.01018978586272 | 3.84083812256764 | 6.55665109471355 | 1.5797446502529 | 0.00013355134394366 |
| Dync1h1 | ENSMUSG00000018707 | 13424 | 0.492475892398472 | 8.39990553447928 | 6.55460706087196 | 1.5848845867055 | 0.000133882329345269 |
| Cpxm1 | ENSMUSG00000027408 | 56264 | 1.77996366713038 | 0.879431749684953 | 6.55264480110214 | 1.5898354775513 | 0.000134196846371539 |
| Atp5mpl | ENSMUSG00000021290 | 70257 | -0.619750663009427 | 6.91790226981596 | -6.55137835024912 | 1.5930394562116 | 0.000134363536233403 |
| Ndufa11 | ENSMUSG00000002379 | 69875 | -0.701356549174369 | 6.01218596955974 | -6.54875040082585 | 1.5997096041740 | 0.000134822094704908 |
| Fer1l6 | ENSMUSG00000037106 | 631797 | -1.2745121393312 | 1.32319286694056 | -6.54799613176487 | 1.6016294889353 | 0.000134879907115195 |
| Tmem126b | ENSMUSG00000030614 | 68472 | -0.500134044071674 | 3.6055845198084 | -6.54501526071684 | 1.6092406665286 | 0.000135416549082561 |
| Chchd7 | ENSMUSG00000042198 | 66433 | -0.647218054929701 | 3.47735404381876 | -6.54353783165485 | 1.6130271647216 | 0.000135630768750558 |
| Wdtc1 | ENSMUSG00000037622 | 230796 | -0.458212813881159 | 5.42641112415302 | -6.54268194250927 | 1.6152250144384 | 0.000135711180882603 |
| Pex19 | ENSMUSG00000003464 | 19298 | -0.442039453876911 | 5.24125018135662 | -6.53835179408062 | 1.6263929386779 | 0.000136544556165044 |
| Pgd | ENSMUSG00000028961 | 110208 | 0.624352137473866 | 2.98563054920865 | 6.53785594338553 | 1.6276769744345 | 0.000136547482790054 |
| Orai2 | ENSMUSG00000039747 | 269717 | 1.8098757872033 | 0.808537797346312 | 6.53644626233011 | 1.6313332690718 | 0.000136749263529328 |
| Gnpat | ENSMUSG00000031985 | 14712 | -0.494044070534002 | 6.7853657954967 | -6.53586644623766 | 1.6328396512719 | 0.000136770653088534 |
| Ecpas | ENSMUSG00000050812 | 230249 | -0.441398724783099 | 7.44483995757908 | -6.52934251645378 | 1.6498904394607 | 0.000137975546496991 |
| Ikzf1 | ENSMUSG00000018654 | 22778 | 1.55554723661552 | 0.919361011906897 | 6.52887119515642 | 1.6511295190242 | 0.000137975546496991 |
| Mfn2 | ENSMUSG00000029020 | 170731 | -0.552177332147124 | 8.6952239817453 | -6.52869984267 41 | 1.6515802389503 | 0.000137975546496991 |
| Dhrs9 | ENSMUSG00000027068 | 241452 | 3.22232418592733 | 1.29380757324383 | 6.52843646836161 | 1.6522732628722 | 0.000137975546496991 |
| Lipa | ENSMUSG00000024781 | 16889 | 0.69360806023981 | 2.93837837015027 | 6.52404903054743 | 1.6638632278887 | 0.000138837320183599 |
| Prdx3 | ENSMUSG00000024997 | 11757 | -0.581530172481245 | 5.64609338864309 | -6.52300924331672 | 1.6666224927597 | 0.000138961483358935 |
| Homer1 | ENSMUSG00000007617 | 26556 | -0.518591468785486 | 5.43065492470396 | -6.51920218697373 | 1.6767664269832 | 0.000139700715040804 |
| Tshz3 | ENSMUSG00000021217 | 243931 | 0.893059567528402 | 2.49836878370989 | 6.51822990821321 | 1.6793674871723 | 0.000139783387465116 |
| Igbp1 | ENSMUSG00000031221 | 18518 | -0.477141070438707 | 4.56743878496465 | -6.51756892078325 | 1.6811382017716 | 0.000139783387465116 |
| Trak2 | ENSMUSG00000026028 | 70827 | -0.615208139070256 | 7.6505709524651 | -6.51739850499329 | 1.6815950463516 | 0.000139783387465116 |
| Rnase4 | ENSMUSG00000021876 | 58809 | 0.751360095263825 | 4.00789248492577 | 6.51555948662021 | 1.6865333521671 | 0.000140087356174312 |
| Atl3 | ENSMUSG00000024759 | 109168 | 0.559682335770479 | 4.10708247283284 | 6.51216075242505 | 1.6957002139816 | 0.000140741830212804 |
| Cluh | ENSMUSG00000020741 | 74148 | -0.489188369358081 | 7.30808093391499 | -6.50528797968235 | 1.7143978358012 | 0.000142185756776504 |
| Pdgfrl | ENSMUSG00000031595 | 68797 | 1.8657074147791 | 2.50746019198448 | 6.50391877097579 | 1.7181486966915 | 0.00014238880518222 |
| Mrpl35 | ENSMUSG00000052962 | 66223 | -0.425517058166649 | 4.36020700745758 | -6.50301629218849 | 1.7206257073396 | 0.000142486057628255 |
| Foxn3 | ENSMUSG00000033713 | 71375 | -0.407385042356609 | 7.6679471645736 | -6.49945958520055 | 1.7304244039444 | 0.00014318901710459 |
| Ak1 | ENSMUSG00000026817 | 11636 | -0.570093707624718 | 8.49742295684595 | -6.49764962325863 | 1.7354333888408 | 0.000143494874231614 |
| Ttll1 | ENSMUSG00000022442 | 319953 | 1.12922786283632 | 2.12276144532229 | 6.49586036327462 | 1.7404000991776 | 0.000143796776145963 |
| Krt80 | ENSMUSG00000037185 | 74127 | 1.4098724598646 | 2.81162086306084 | 6.49493032659658 | 1.7429876509856 | 0.000143816276304411 |
| Fam114a1 | ENSMUSG00000029185 | 68303 | 0.815808974981748 | 3.49843444127389 | 6.49465439504688 | 1.7437561282613 | 0.000143816276304411 |
| Fcer1g | ENSMUSG00000058715 | 14127 | 1.75452274718265 | 1.7441592754763 | 6.49435759118838 | 1.7445831340192 | 0.000143816276304411 |
| Zbtb1 | ENSMUSG00000033454 | 268564 | 0.480948245982718 | 4.1819859615641 | 6.4928992741456 | 1.7486525541837 | 0.000144043112809213 |
| Nfam1 | ENSMUSG00000058099 | 74039 | 1.48386754051561 | 1.34383739396769 | 6.492200474286 | 1.7506060917691 | 0.00014409544570127 |
| Asb12 | ENSMUSG00000031204 | 70392 | -0.697431209961599 | 3.02670378036442 | -6.4895570379948 | 1.7580168039531 | 0.000144596551422215 |
| Dusp18 | ENSMUSG00000047205 | 75219 | -0.924565929625646 | 6.08867965944051 | -6.48756062777628 | 1.7636355206169 | 0.000144949623126794 |
| Mrs2 | ENSMUSG00000021339 | 380836 | -0.367200819028503 | 5.35936973904393 | -6.48660718592668 | 1.7663255689013 | 0.000144954734399216 |
| Tmem106a | ENSMUSG00000034947 | 217203 | 1.30039456750262 | 1.17894117469751 | 6.48642522459479 | 1.7668394470431 | 0.000144954734399216 |
| Acads | ENSMUSG00000029545 | 11409 | -0.414800217110162 | 5.79025024837239 | -6.48612913360566 | 1.7676759761609 | 0.000144954734399216 |
| Tmem201 | ENSMUSG00000044700 | 230917 | -0.431838965447147 | 5.08936163337638 | -6.48377890964713 | 1.7743307495245 | 0.000145391374985405 |
| Csf2rb | ENSMUSG00000071713 | 12983 | 1.43960837154699 | 2.05305002217079 | 6.48050798064273 | 1.7836365046774 | 0.000146044424214453 |
| Dapk1 | ENSMUSG00000021559 | 69635 | 0.93595053790089 | 2.77839252366224 | 6.479371099843 | 1.7868829464523 | 0.000146194270782618 |
| Gck | ENSMUSG00000041798 | 103988 | 1.57313837057495 | 0.897291752300462 | 6.47893097195783 | 1.7881414329554 | 0.000146194270782618 |
| Map1s | ENSMUSG00000019261 | 270058 | 0.53179369289776 | 3.24893591367246 | 6.47747800234514 | 1.7923026375690 | 0.000146351081548666 |
| Myom2 | ENSMUSG00000031461 | 17930 | 0.611204222870239 | 8.55996439159777 | 6.47732648736109 | 1.7927371529929 | 0.000146351081548666 |
| Uqcr10 | ENSMUSG00000059534 | 66152 | -0.67522587378665 | 6.31829080847927 | -6.47569940319367 | 1.7974103137617 | 0.000146424300077535 |
| Cit | ENSMUSG00000029516 | 12704 | 1.21273717361219 | 1.63740642316223 | 6.47548821033594 | 1.7980178226392 | 0.000146424300077535 |
| Abcd3 | ENSMUSG00000028127 | 19299 | -0.446997356495728 | 5.78167295807439 | -6.47522118920733 | 1.7987862345226 | 0.000146424300077535 |
| Tie1 | ENSMUSG00000033191 | 21846 | -0.553061311111357 | 4.35228076138134 | -6.47514964471745 | 1.7989921782465 | 0.000146424300077535 |
| Pnpla2 | ENSMUSG00000025509 | 66853 | -0.54521271948303 | 7.60754888836072 | -6.47384446627001 | 1.8027535504119 | 0.000146621272764528 |
| Calu | ENSMUSG00000029767 | 12321 | 0.454069407595834 | 7.06776055564023 | 6.47336730708252 | 1.8041307348033 | 0.000146624186335576 |
| Fam49a | ENSMUSG00000020589 | 76820 | 0.667965234569811 | 2.87651822684756 | 6.47210115531151 | 1.8077904970159 | 0.000146812465994664 |
| Pnmal2 | ENSMUSG00000070802 | 434128 | 1.24228423945521 | 1.50581299584451 | 6.46991659622407 | 1.8141232785296 | 0.000147217383501171 |
| Kat14 | ENSMUSG00000027425 | 228714 | -0.398106117762759 | 4.59072584386372 | -6.46893677325301 | 1.8169712581233 | 0.000147339115894258 |
| Fastkd2 | ENSMUSG00000025962 | 75619 | -0.628120019122758 | 3.66905965053991 | -6.45515625471402 | 1.8575288322432 | 0.000150474512113459 |
| Cdk5rap3 | ENSMUSG00000018669 | 80280 | 0.700060358363642 | 3.32395699665219 | 6.4548671570979 | 1.8583898211798 | 0.000150474512113459 |
| Apba3 | ENSMUSG00000004931 | 57267 | -0.481189939151186 | 5.10862825483136 | -6.45361864926748 | 1.8621129508231 | 0.000150664372061053 |
| Cul3 | ENSMUSG00000004364 | 26554 | -0.434134151760978 | 6.99855602524344 | -6.45120348189865 | 1.8693374499435 | 0.000151125889734406 |
| Slc38a3 | ENSMUSG00000010064 | 76257 | -0.754633894213861 | 5.38848605837439 | -6.45046551973347 | 1.8715508028338 | 0.000151125889734406 |
| Rab3ip | ENSMUSG00000064181 | 216363 | -0.496419820238835 | 4.16322116718371 | -6.45032764306125 | 1.8719646391033 | 0.000151125889734406 |
| Rasgrf2 | ENSMUSG00000021708 | 19418 | -0.685373033591106 | 2.80312961240944 | -6.44879867753560 | 1.8765602889776 | 0.000151290844012853 |
| Aqp7 | ENSMUSG00000028427 | 11832 | -1.09735871235213 | 2.82995160118762 | -6.44872700798808 | 1.8767759992812 | 0.000151290844012853 |
| Gfm1 | ENSMUSG00000027774 | 28030 | -0.410336296129324 | 6.15807313789288 | -6.44785425608469 | 1.8794048958762 | 0.000151391119504959 |
| Kitl | ENSMUSG00000019966 | 17311 | -0.803703554055565 | 4.65423465514071 | -6.44536855434825 | 1.8869136167112 | 0.000151847234282762 |
| Asb16 | ENSMUSG00000034768 | 217217 | -0.511062041611969 | 5.47325737875508 | -6.44506077724079 | 1.8878454980356 | 0.000151847234282762 |
| P2rx4 | ENSMUSG00000029470 | 18438 | 0.925323956593822 | 2.21482967053631 | 6.4444762508383 | 1.8896167125984 | 0.000151877943275096 |
| Trrap | ENSMUSG00000045482 | 100683 | 0.472800000484686 | 5.42823049898432 | 6.44338595538269 | 1.8929251722973 | 0.000152032072434841 |
| Ift122 | ENSMUSG00000030323 | 81896 | 0.827392823635209 | 2.38820378683806 | 6.44136984583382 | 1.8990590611455 | 0.000152258502353157 |
| Foxp4 | ENSMUSG00000023991 | 74123 | 0.701113374263996 | 3.45642068336064 | 6.44100248456779 | 1.9001789880943 | 0.000152258502353157 |
| Prxl2c | ENSMUSG00000021482 | 66129 | -0.38780245170792 | 4.94273913814212 | -6.4407127412122 | 1.9010627819881 | 0.000152258502353157 |
| Lilrb4a | ENSMUSG00000112148 | 14728 | 1.68554632229316 | 1.23476044853899 | 6.44062974101182 | 1.9013160343249 | 0.000152258502353157 |
| Hsd3b7 | ENSMUSG00000042289 | 101502 | 2.03549974033147 | 1.29612108200966 | 6.43788465332147 | 1.9097119690002 | 0.000152818898485665 |
| Kpna3 | ENSMUSG00000021929 | 16648 | -0.353931719470678 | 6.34022849072336 | -6.43343461169762 | 1.9234056180612 | 0.00015380209810554 |
| Dync1li1 | ENSMUSG00000032435 | 235661 | 0.46052220780207 | 5.16126254841795 | 6.42998520982471 | 1.9340912341338 | 0.000154543503511088 |
| Zfp568 | ENSMUSG00000074221 | 243905 | 0.887349593015442 | 4.5812531828717 | 6.42878676271846 | 1.9378184232374 | 0.000154728219024166 |
| Snx7 | ENSMUSG00000028007 | 76561 | 0.947916943236263 | 2.21748050921415 | 6.42610530061512 | 1.9461852355915 | 0.000155282852629574 |
| Rras2 | ENSMUSG00000055723 | 66922 | 0.565902267763519 | 3.84946930497062 | 6.41906971133549 | 1.9683193458135 | 0.000156921442608067 |
| Tmem177 | ENSMUSG00000036975 | 66343 | -0.750716814236287 | 2.21591029470335 | -6.41866738152898 | 1.9695930771042 | 0.000156921442608067 |
| Vps36 | ENSMUSG00000031479 | 70160 | -0.440956789417564 | 4.85729258381803 | -6.4176569205244 | 1.9727958969351 | 0.000157062140927884 |
| Ndufb4 | ENSMUSG00000022820 | 68194 | -0.76566342859322 | 6.10973655665286 | -6.41416931643714 | 1.9838926222707 | 0.000157830642314715 |
| Nnt | ENSMUSG00000025453 | 18115 | -0.878528554004156 | 7.83990748053357 | -6.41316221206148 | 1.9871092047970 | 0.00015786470924324 |
| Atp5md | ENSMUSG00000071528 | 66477 | -0.769881104115506 | 6.28330862436744 | -6.41297773775295 | 1.9876989906726 | 0.00015786470924324 |
| Dact3 | ENSMUSG00000078794 | 629378 | 1.01256418013797 | 1.76594072499012 | 6.41267933371086 | 1.9886534134840 | 0.00015786470924324 |
| Lpin2 | ENSMUSG00000024052 | 64898 | 0.875937279020326 | 2.21760450465579 | 6.41207302621384 | 1.9905941289861 | 0.000157904095964788 |
| Prr12 | ENSMUSG00000046574 | 233210 | 0.670365212826592 | 4.51326578632442 | 6.4099250922123 | 1.9974854640288 | 0.00015831005642105 |
| Gda | ENSMUSG00000058624 | 14544 | 0.649679633601826 | 3.8510984620907 | 6.40957584610492 | 1.9986083419728 | 0.00015831005642105 |
| Lgi1 | ENSMUSG00000067242 | 56839 | -0.854010451133044 | 4.34070104970575 | -6.40343042407735 | 2.0184759119661 | 0.000159767995609718 |
| Slc41a3 | ENSMUSG00000030089 | 71699 | -1.28476643168138 | 4.28957156411368 | -6.39763318179812 | 2.0374085323163 | 0.000161149874578506 |
| Dnajc19 | ENSMUSG00000027679 | 67713 | -0.491197949782823 | 4.6202255391056 | -6.39270518042682 | 2.0536494158639 | 0.000162317004807008 |
| Me3 | ENSMUSG00000030621 | 109264 | -0.768288243131235 | 4.64688702677886 | -6.39215832893747 | 2.0554600217875 | 0.000162332260617295 |
| Atp5h | ENSMUSG00000034566 | 71679 | -0.578366174680182 | 8.0371746153277 | -6.39175015695829 | 2.0568125601953 | 0.000162332260617295 |
| Tmed10 | ENSMUSG00000021248 | 68581 | 0.359137794032299 | 6.2346325234319 | 6.3887018524847 | 2.0669432483776 | 0.000163014117229502 |
| Mrps25 | ENSMUSG00000014551 | 64658 | -0.521927837375297 | 3.87628166332775 | -6.38755863586193 | 2.0707561313869 | 0.00016319708199128 |
| Capn2 | ENSMUSG00000026509 | 12334 | 0.524044591362649 | 6.25073448277192 | 6.38610818887026 | 2.0756043575922 | 0.000163461320121331 |
| Lama2 | ENSMUSG00000019899 | 16773 | 0.603667245329842 | 6.82041279138758 | 6.38240224847038 | 2.0880460942776 | 0.000164322763546072 |
| Qsox1 | ENSMUSG00000033684 | 104009 | 0.464237400987617 | 4.2923406202546 | 6.38055224129237 | 2.0942863744192 | 0.000164695283156664 |
| Vcl | ENSMUSG00000021823 | 22330 | 0.432593549254306 | 6.34341544229653 | 6.37660935787608 | 2.1076517688750 | 0.000165627185374361 |

| | | | | | | | |
|---|---|---|---|---|---|---|---|
| Mycbp2 | ENSMUSG00000033004 | 105689 | 0.496152134785415 | 7.14625354511068 | 6.37396471113345 | 2.11666675839800 | 0.0001662161231038 |
| Hdx | ENSMUSG00000034551 | 245596 | 1.66720084459334 | 0.684309295770345 | 6.36782304442343 | 2.1377592454512 | 0.000167751947681459 |
| Ly86 | ENSMUSG00000021423 | 17084 | 1.89277338149733 | 0.787023728064455 | 6.36503371428545 | 2.1474116758993 | 0.000168388500927228 |
| Smim3 | ENSMUSG00000038059 | 106878 | 1.04624177091285 | 1.9515142531457 | 6.36399299301963 | 2.1510248179258 | 0.000168550912435463 |
| Tnni1 | ENSMUSG00000026418 | 21952 | 0.766758221507599 | 9.81009111308441 | 6.36286807306939 | 2.1549374766606 | 0.000168736544107291 |
| Sdhaf2 | ENSMUSG00000024668 | 66072 | -0.55977784810632 | 4.47531763239771 | -6.36069344227411 | 2.1625224608099 | 0.000169137262926244 |
| Zfp91 | ENSMUSG00000024695 | 109910 | -0.445880537348493 | 7.36951794836029 | -6.36011991413343 | 2.1645275731805 | 0.000169137262926244 |
| Des | ENSMUSG00000026208 | 13346 | 0.622661246426711 | 11.6932968824711 | 6.36007147350147 | 2.1646970161359 | 0.000169137262926244 |
| Fth1 | ENSMUSG00000024661 | 14319 | -0.512433101496049 | 9.1470538349917 | -6.35949593746765 | 2.1667112818153 | 0.000169173721582307 |
| Alox5ap | ENSMUSG00000060063 | 11690 | 1.13853377064794 | 1.54394626973226 | 6.35530788928624 | 2.1814281943572 | 0.000170201224786005 |
| Mast4 | ENSMUSG00000034751 | 328329 | 0.862220602040526 | 5.60509353366674 | 6.35479225777039 | 2.1832473991986 | 0.00017022166419858 |
| Rras | ENSMUSG00000038387 | 20130 | 0.692918706159025 | 3.61298195269314 | 6.35422970950533 | 2.1852339532459 | 0.00017025511292182 |
| Dnm1 | ENSMUSG00000026825 | 13429 | 0.753611286119839 | 3.36154143387158 | 6.35166960866984 | 2.1942986250856 | 0.000170839588823438 |
| Adgre5 | ENSMUSG00000002885 | 26364 | 0.51691272588581 | 4.83147724788695 | 6.35057986947479 | 2.1981691190128 | 0.000171019121992381 |
| Uqcrq | ENSMUSG00000044894 | 22272 | -0.605976775719987 | 8.07948717996158 | -6.34537812355399 | 2.216743757308 | 0.000172341579026556 |
| Myh9 | ENSMUSG00000022443 | 17886 | 0.532754236707761 | 7.88783757345583 | 6.34463770916448 | 2.2194010659478 | 0.0001724255369714 |
| Cacna1g | ENSMUSG00000020866 | 12291 | 1.19479622463035 | 2.57941776247095 | 6.34199467993507 | 2.2289141399587 | 0.000173041622612849 |
| Wls | ENSMUSG00000028173 | 68151 | 0.572601660647912 | 4.0231158221179 | 6.33911492884702 | 2.2393281251771 | 0.000173726726304548 |
| Emcn | ENSMUSG00000054690 | 59308 | -1.08545245511533 | 3.09794749166872 | -6.33831329224274 | 2.2422361708029 | 0.000173742449252319 |
| Tek | ENSMUSG00000006386 | 21687 | -0.646996119850257 | 4.62285527772852 | -6.33784094563163 | 2.2439515331355 | 0.000173742449252319 |
| Zfp768 | ENSMUSG00000047371 | 233890 | -0.477478833284232 | 4.13431672869309 | -6.33774527565226 | 2.2442991340616 | 0.000173742449252319 |
| Aldh3b1 | ENSMUSG00000024885 | 67689 | 1.43688678886932 | 0.709323140499556 | 6.33428192266199 | 2.2569208895226 | 0.000174595911134978 |
| Igfn1 | ENSMUSG00000051985 | 226438 | 0.778401382757025 | 3.14965440017389 | 6.33355655823888 | 2.2595738390919 | 0.000174677522171952 |
| Ung | ENSMUSG00000029591 | 22256 | -1.44326562009464 | 1.80675282494676 | -6.33175123427938 | 2.2661908956644 | 0.0001750652486255 |
| Sirt3 | ENSMUSG00000025486 | 64384 | -0.585699149513593 | 2.88845113078174 | -6.32752718230108 | 2.2817531347556 | 0.000176142962683718 |
| Ctdsp1 | ENSMUSG00000026176 | 227292 | -0.342139780413561 | 5.54622163158739 | -6.32699270977623 | 2.2837302383487 | 0.00017617117315025 |
| Gpc3 | ENSMUSG00000055653 | 14734 | -0.938058449119205 | 2.38132866989299 | -6.32631516849684 | 2.2862391692236 | 0.000176240341035142 |
| Slc2a12 | ENSMUSG00000037490 | 353169 | -0.826766507091437 | 2.22654840845036 | -6.32545374342183 | 2.2894332100953 | 0.000176362187593741 |
| Tnfrsf1b | ENSMUSG00000028599 | 21938 | 0.97030500037807 | 2.99573577883873 | 6.32166236921334 | 2.3035470079720 | 0.000177324453127763 |
| Surf4 | ENSMUSG00000014867 | 20932 | 0.371374257393864 | 5.79764510573083 | 6.31888673198751 | 2.3139376582154 | 0.000177998962293832 |
| Lmod1 | ENSMUSG00000048096 | 93689 | -1.27868828787601 | 2.02011903693906 | -6.31638312122252 | 2.3233522875962 | 0.000178597495469161 |
| Nos3 | ENSMUSG00000028978 | 18127 | -0.728956081806505 | 3.36341308952656 | -6.31323390702564 | 2.3352519376841 | 0.000179386078220838 |
| Ecsit | ENSMUSG00000066839 | 26940 | -0.555768756452099 | 5.36276740227943 | -6.31265809876881 | 2.3374346192666 | 0.000179427653252832 |
| Mmrn2 | ENSMUSG00000041445 | 105450 | -0.621030759337283 | 4.41825246977708 | -6.31106792931828 | 2.3434735373925 | 0.000179764977103421 |
| Rab28 | ENSMUSG00000029128 | 100972 | -0.476412379771331 | 4.9995005974515 | -6.3103799659001 | 2.3460912727951 | 0.000179839577159351 |
| Themis2 | ENSMUSG00000037731 | 230787 | 1.01044603009713 | 1.5963050987135 | 6.30626828360049 | 2.3618007203833 | 0.000180916914327329 |
| Ndufb6 | ENSMUSG00000071014 | 230075 | -0.544783119530837 | 4.71531317013037 | -6.30554747248378 | 2.3645660984308 | 0.000180926706041528 |
| Mrps15 | ENSMUSG00000028861 | 66407 | -0.462427026702069 | 4.95834212035449 | -6.30537224011495 | 2.3652388887873 | 0.000180926706041528 |
| Msr1 | ENSMUSG00000025044 | 20288 | 2.07362684221002 | 0.240716494156544 | 6.29723612262516 | 2.3966997228710 | 0.000183164629811372 |
| 6230400D17Rik | ENSMUSG00000097577 | 76133 | -1.1953561046886 | 1.27143209267598 | -6.29694171691402 | 2.3978463567841 | 0.000183164629811372 |
| St6galnac4 | ENSMUSG00000079442 | 20448 | 0.603090956029664 | 4.23917829147572 | 6.29595099831851 | 2.4017091904783 | 0.00018333158632066 |
| Scarf2 | ENSMUSG00000012017 | 224024 | 1.0425398305832 | 3.27565263411627 | 6.29535033164261 | 2.4040543912396 | 0.000183382543968183 |
| Zfp770 | ENSMUSG00000040321 | 228491 | -0.755680166771242 | 5.17392193159311 | -6.29403139159075 | 2.4092124270508 | 0.000183647845467871 |
| Smc4 | ENSMUSG00000034349 | 70099 | 0.669441789378442 | 3.39212529531919 | 6.29160572580282 | 2.4187290087653 | 0.00018424478602658 |
| Trerf1 | ENSMUSG00000064043 | 224829 | 0.961512383614783 | 1.68374332040943 | 6.29112969210624 | 2.4206012639226 | 0.000184259000110994 |
| Trim72 | ENSMUSG00000042828 | 434246 | 0.576558850100735 | 7.37871599163525 | 6.28965509492041 | 2.4264105845101 | 0.000184572679883653 |
| Cd53 | ENSMUSG00000040747 | 12508 | 1.77755729826457 | 1.28184131878973 | 6.2880554879619 | 2.4327289852891 | 0.00018492462126701 |
| Zfp62 | ENSMUSG00000046311 | 22720 | -0.48201838442852 | 4.34515088162016 | -6.28558428873006 | 2.4425242046324 | 0.000185540181242792 |
| Blcap | ENSMUSG00000067787 | 53619 | -0.558427785361888 | 3.5145259372254 | -6.28502086400443 | 2.4447632841670 | 0.000185581301800212 |
| Fem1a | ENSMUSG00000043683 | 14154 | -0.405407323335647 | 7.62864590207701 | -6.28027856452965 | 2.4636952417404 | 0.000186888637664569 |
| Pde7a | ENSMUSG00000069094 | 18583 | -0.745330091087687 | 3.9419132613746 | -6.27971365346808 | 2.4659607103962 | 0.000186930766472547 |
| Ndufa13 | ENSMUSG00000036199 | 67184 | -0.531891441220401 | 7.1169559278663 | -6.27901153176591 | 2.4687794917872 | 0.000186950271977339 |
| Dnase2a | ENSMUSG00000003812 | 13423 | 0.71948623555115 | 2.09867315276981 | 6.27878251090531 | 2.4696996662511 | 0.000186950271977339 |
| Ciao3 | ENSMUSG00000002280 | 67563 | -0.45736300407695 | 4.18050566678955 | -6.27808030854658 | 2.4725232693233 | 0.000186950271977339 |
| Ctsh | ENSMUSG00000032359 | 13036 | 0.732496734954033 | 3.18704485881356 | 6.27794714101931 | 2.4730591279776 | 0.000186950271977339 |
| Cfp | ENSMUSG00000001128 | 18636 | 1.49904900042433 | 0.99280628281275 | 6.27615152558276 | 2.4802965275963 | 0.000187318252505982 |
| Slc25a39 | ENSMUSG00000018677 | 68066 | -0.498802273384121 | 4.29169974969033 | -6.27588957180655 | 2.4813542185404 | 0.000187318252505982 |
| Trpt1 | ENSMUSG00000047656 | 107328 | -0.681513541065406 | 2.24145443660557 | -6.27116537529898 | 2.5005107814189 | 0.000188634115608633 |
| Prickle3 | ENSMUSG00000031145 | 54630 | -0.738407976564655 | 3.50562594918143 | -6.26398012739018 | 2.5299457407249 | 0.000190723013047338 |
| Zfp639 | ENSMUSG00000027667 | 67778 | -0.598795255036519 | 3.50604040909198 | -6.2617335224518 | 2.5392237756580 | 0.000191164927144374 |
| Sept5 | ENSMUSG00000072214 | 18951 | 1.11822539636118 | 2.48104167667327 | 6.2617137940106 | 2.5393054085960 | 0.000191164927144374 |
| Ramp1 | ENSMUSG00000034353 | 51801 | -0.820899680582112 | 5.11513576640173 | -6.25842157306966 | 2.5529668095503 | 0.000191916648808964 |
| Kirrel | ENSMUSG00000041734 | 170643 | 0.820305149565248 | 3.58744342504486 | 6.25812285135053 | 2.5542102057068 | 0.000191916648808964 |
| Csnk1g3 | ENSMUSG00000073563 | 70425 | 0.485826063955392 | 4.32234087306418 | 6.25803934641372 | 2.5545578997076 | 0.000191916648808964 |
| Pcmtd1 | ENSMUSG00000051285 | 319263 | -0.434508137104262 | 5.70756440736348 | -6.25753076499403 | 2.5566765829352 | 0.000191943899231218 |
| Arhgap44 | ENSMUSG00000033389 | 216831 | 0.562936139999212 | 4.18047675624103 | 6.25332765848572 | 2.5742570985357 | 0.000193131121098794 |
| 2310002L09Rik | ENSMUSG00000028396 | 71886 | -0.577711013328949 | 5.07361959853058 | -6.24672072329209 | 2.6021497471783 | 0.000195089841470555 |
| Cercam | ENSMUSG00000039787 | 99151 | 1.32089082935075 | 1.95015989829149 | 6.24415642761225 | 2.6130609750293 | 0.000195744259461083 |
| Morc4 | ENSMUSG00000031434 | 75746 | 0.687562151616462 | 3.05733962651426 | 6.24382845964945 | 2.6144599653570 | 0.000195744259461083 |
| Calcoco1 | ENSMUSG00000023055 | 67488 | -0.369332214638027 | 5.74487099237543 | -6.23936771053911 | 2.6335663057653 | 0.000197039789790015 |
| Apip | ENSMUSG00000010911 | 56369 | -0.665524689352213 | 2.90646167330743 | -6.23817208924791 | 2.6387123267266 | 0.000197289770475026 |
| Thbs2 | ENSMUSG00000023885 | 21826 | 0.902272254744441 | 5.59080329720254 | 6.23143818900327 | 2.6678936309611 | 0.000199335237730937 |
| Prox1 | ENSMUSG00000010175 | 19130 | 0.472090135344052 | 4.97239013733476 | 6.23061590808192 | 2.6714801653223 | 0.000199466869447668 |
| Pcnx3 | ENSMUSG00000054874 | 104401 | 0.506860258957019 | 3.61345333654964 | 6.22584863050763 | 2.6923737273713 | 0.000200842007530636 |
| Il17ra | ENSMUSG00000002897 | 16172 | 1.00531366480573 | 1.83168151553749 | 6.22557634433157 | 2.6935722535899 | 0.000200842007530636 |
| Zfp521 | ENSMUSG00000024420 | 225207 | 0.61553778102499 | 3.72458116681807 | 6.22432772304054 | 2.6990755203022 | 0.000201115163683873 |
| Rrm1 | ENSMUSG00000030978 | 20133 | -0.523320007372635 | 3.53904243707854 | -6.22120718020226 | 2.7128810077684 | 0.00020200614642995 |
| Micu1 | ENSMUSG00000020111 | 216001 | 0.440484505087984 | 6.00659406568819 | 6.22000934848536 | 2.7182000025683 | 0.000202264426331345 |
| Mgst3 | ENSMUSG00000026688 | 66447 | -0.560743757314719 | 5.86029513178877 | -6.21852501326041 | 2.7248064432721 | 0.000202608896524634 |
| Fads1 | ENSMUSG00000010663 | 76267 | 0.661558984976534 | 2.93112169791366 | 6.21813696642823 | 2.7265363350813 | 0.000202608896524634 |
| Ybx2 | ENSMUSG00000018554 | 53422 | -1.84764205747466 | 0.289535465251673 | -6.21665285512987 | 2.7331630871286 | 0.000202963353976876 |
| Leng1 | ENSMUSG00000078813 | 69757 | -0.570086141563632 | 3.10667899949687 | -6.21593923769215 | 2.7363555115438 | 0.000203062471803708 |
| Prrx2 | ENSMUSG00000039476 | 20204 | 1.64293480040614 | 0.783527171637653 | 6.21520781340882 | 2.7396316663503 | 0.000203129077161803 |
| Cep68 | ENSMUSG00000044066 | 216543 | -0.395361244964247 | 5.08933638265944 | -6.21490937120339 | 2.7409696168114 | 0.000203129077161803 |
| Aldh4a1 | ENSMUSG00000028737 | 212647 | -0.524702877533037 | 4.97132622286856 | -6.20712865330079 | 2.7760950681496 | 0.000205592785839728 |
| Ddr2 | ENSMUSG00000026674 | 18214 | 0.696660556195336 | 4.98492392364935 | 6.20530308447112 | 2.7844048230765 | 0.00020606857901862 |
| Mfsd4b1 | ENSMUSG00000038522 | 215929 | -1.27971982369884 | 1.47215298191731 | -6.20416351052206 | 2.7896052750043 | 0.000206313770372615 |
| Barx2 | ENSMUSG00000032033 | 12023 | -0.934614712296865 | 4.3607586946767 | -6.20363328092549 | 2.7920284617657 | 0.000206353367921309 |
| Fahd1 | ENSMUSG00000045316 | 68636 | -0.692561129764569 | 3.24977703201714 | -6.20262122191509 | 2.7966597899617 | 0.000206556001108595 |
| Myo1f | ENSMUSG00000024300 | 17916 | 1.75392231801673 | 1.58885238753456 | 6.20152287851473 | 2.8016951107731 | 0.000206788178635122 |
| C5ar1 | ENSMUSG00000049130 | 12273 | 1.55086853647723 | 0.780017027539236 | 6.19528801002038 | 2.8304599175909 | 0.000208770292572109 |
| Islr2 | ENSMUSG00000051243 | 320563 | 1.13143023103214 | 1.7869862692171 | 6.19132092991013 | 2.8489237250363 | 0.000209990460137375 |
| Plxdc2 | ENSMUSG00000026748 | 67448 | 0.767754768201821 | 4.99410456897732 | 6.19076921380872 | 2.8515015678835 | 0.0002100388385346 |
| March1 | ENSMUSG00000036469 | 72925 | 1.85833064797007 | 0.383083549767486 | 6.18871553726465 | 2.8611187614662 | 0.000210605314354125 |
| Pnpla8 | ENSMUSG00000036257 | 67452 | -0.440036190435287 | 6.16953892190227 | -6.18665754061135 | 2.8707904068837 | 0.000211118602075921 |
| Syt17 | ENSMUSG00000058420 | 110058 | 1.70470594526554 | 0.416114787050817 | 6.18641029994671 | 2.8719546362354 | 0.000211118602075921 |
| Kbtbd12 | ENSMUSG00000033182 | 74589 | -0.469345657337487 | 4.09868988895269 | -6.18436132076012 | 2.8816221833327 | 0.000211686909180177 |
| Ccr5 | ENSMUSG00000079227 | 12774 | 1.64154606710679 | 1.62862113049995 | 6.18309157842583 | 2.8876302828141 | 0.000211868803218466 |
| Ltbp1 | ENSMUSG00000001870 | 268977 | 0.679992575999186 | 4.57206377815397 | 6.18301886952984 | 2.8879747213934 | 0.000211868803218466 |
| Bzw2 | ENSMUSG00000020547 | 66912 | -0.542826868075476 | 6.22613612766583 | -6.17891037486652 | 2.9075079552576 | 0.000213158748886122 |
| Vav1 | ENSMUSG00000034116 | 22324 | 1.75124724586436 | 1.03670515307206 | 6.17650447885237 | 2.919010882667 | 0.000213858632429175 |
| Adipor2 | ENSMUSG00000030168 | 68465 | -0.45138587200219 | 6.01019618590086 | -6.17420254960009 | 2.9300615300939 | 0.000214524464738489 |
| Nob1 | ENSMUSG00000003848 | 67619 | -0.741173002014842 | 3.4279447102836 | -6.16984523656356 | 2.9510998656834 | 0.000215920164871389 |
| Slit3 | ENSMUSG00000056427 | 20564 | 0.737497551813794 | 4.24202268151122 | 6.16905439626245 | 2.9549352643058 | 0.0002160561697266 |
| Cenpf | ENSMUSG00000026605 | 108000 | 1.25123953212612 | 1.43858434359768 | 6.16644154831942 | 2.9676442975431 | 0.000216820729261407 |
| Atp1b4 | ENSMUSG00000016327 | 67821 | 1.16253128803611 | 3.41913837286055 | 6.16608992500954 | 2.9693589946420 | 0.000216820729261407 |
| Rab15 | ENSMUSG00000021062 | 104886 | 0.714103869949364 | 4.02828717426633 | 6.16539011988184 | 2.9727747066012 | 0.000216925235766746 |
| Ipo5 | ENSMUSG00000030662 | 70572 | -0.435696392234148 | 7.5931891073012 | -6.16282377795684 | 2.9853362808995 | 0.000217607845233874 |
| Amz2 | ENSMUSG00000020610 | 13929 | -0.45205282661915 | 4.57924663273392 | -6.16266592160763 | 2.9861107661770 | 0.000217607845233874 |
| Acyp2 | ENSMUSG00000060923 | 75572 | -0.711561534437352 | 4.87532748000576 | -6.16085738440547 | 2.9949990203020 | 0.000218110155169363 |
| Capg | ENSMUSG00000056737 | 12332 | 1.14580656399596 | 2.82380492719494 | 6.15824960113276 | 3.0078642564688 | 0.000218901226281365 |
| Fdx1 | ENSMUSG00000032051 | 14148 | -0.626635755694976 | 3.07079880866412 | -6.15616074437804 | 3.0182113251419 | 0.00021950810375999 |
| Atp5k | ENSMUSG00000050856 | 11958 | -0.537015074076456 | 5.83512249939338 | -6.15511558401157 | 3.0234025247041 | 0.000219739448121951 |
| Crim1 | ENSMUSG00000024074 | 50766 | -0.769900084537011 | 5.93807874784962 | -6.14871904469705 | 3.0553786483465 | 0.000221915907010469 |
| Car14 | ENSMUSG00000038526 | 23831 | -1.55837821844637 | 2.88172517148981 | -6.14647597691722 | 3.0666757107222 | 0.000222542246656566 |
| Ptgr2 | ENSMUSG00000072946 | 77219 | -0.498527756776152 | 6.02551195659977 | -6.14619895225511 | 3.0680739704642 | 0.000222542246656566 |

| | | | | | | | |
|---|---|---|---|---|---|---|---|
| Slc39a7 | ENSMUSG00000024327 | 14977 | 0.370985741722 79 | 4.55439381642507 | 6.143437570502 | 3.0820485154159 | 0.000223407641392651 |
| Car2 | ENSMUSG00000027562 | 12349 | -1.64578729790066 | 2.33724706613483 | -6.14093640251165 | 3.0947639498581 | 0.00022418068082107 |
| Ndufa4 | ENSMUSG00000029632 | 17992 | -0.595139103121123 | 5.95210109742149 | -6.13878371156639 | 3.1057519529254 | 0.000224827646340582 |
| Syt12 | ENSMUSG00000049303 | 171180 | 1.5493695354198 | 1.71965860829432 | 6.13636974034435 | 3.1181224064446 | 0.00022549950810914 |
| 0610040J01Rik | ENSMUSG00000060512 | 76261 | -2.91370643457537 | -0.480583194684401 | -6.136167941696 | 3.1191588716587 | 0.00022549950810914 |
| Iqgap1 | ENSMUSG00000030536 | 29875 | 0.780348589724759 | 6.23421120473242 | 6.1343758458082 | 3.1283792231337 | 0.000226016611289322 |
| Slf1 | ENSMUSG00000021597 | 105377 | -0.582977628835278 | 2.50997135143836 | -6.13310316648209 | 3.1349445658605 | 0.000226263765752585 |
| Spry2 | ENSMUSG00000022114 | 24064 | 0.711820266686947 | 3.24061282772822 | 6.13291044148997 | 3.1359400339874 | 0.000226263765752585 |
| Cep170b | ENSMUSG00000072825 | 217882 | 0.589297908588786 | 4.09403138627222 | 6.13037418875534 | 3.1490713546316 | 0.000226990556687834 |
| Adgrl1 | ENSMUSG00000013033 | 330814 | 0.521861597934471 | 4.22905793594129 | 6.13016321417293 | 3.1501662656247 | 0.000226990556687834 |
| Cdc42ep3 | ENSMUSG00000036533 | 260409 | -0.565222773138752 | 3.66841515776169 | -6.12349934962974 | 3.1849569760036 | 0.000229346276052014 |
| Ccn4 | ENSMUSG00000005124 | 22402 | 1.94751490201693 | 1.66504050467849 | 6.12303289829406 | 3.1874073012306 | 0.000229371620867362 |
| Sv2b | ENSMUSG00000053025 | 64176 | 1.5365377304206 | 1.7845301488641 | 6.11541831766913 | 3.2276890225065 | 0.000232117557269861 |
| Krt222 | ENSMUSG00000035849 | 268481 | -0.906863175223289 | 2.37755991416688 | -6.11457393796397 | 3.2321886978203 | 0.000232288327783527 |
| Cplane1 | ENSMUSG00000039801 | 73692 | 0.673763680239177 | 3.20221182640416 | 6.11374382138565 | 3.2366187925324 | 0.000232453876617422 |
| Arl2bp | ENSMUSG00000031776 | 107566 | 0.38367746549406 | 4.86894620683572 | 6.11241658454169 | 3.2437151392980 | 0.000232810572473192 |
| Stard9 | ENSMUSG00000033705 | 668880 | 0.592644161429311 | 4.10357966852587 | 6.11145882931257 | 3.2488461324983 | 0.000233025833821123 |
| Carns1 | ENSMUSG00000075289 | 107239 | 1.02940986281857 | 1.62357169603185 | 6.10789872585123 | 3.2679936245539 | 0.000234245497114749 |
| Tpd52 | ENSMUSG00000027506 | 21985 | 0.437312832219612 | 4.34704278289164 | 6.1043615827028 | 3.2871351155453 | 0.00023546313203163 |
| Sned1 | ENSMUSG00000047793 | 208777 | 0.756954072883928 | 3.57468710346796 | 6.10159206601286 | 3.3022047757648 | 0.00023638769092263 |
| Skp1a | ENSMUSG00000036309 | 21402 | -0.424278178049583 | 6.65073912415464 | -6.10103165920889 | 3.3052629170535 | 0.000236451760119846 |
| Ndufa6 | ENSMUSG00000022450 | 67130 | -0.513414854403761 | 6.86134070683161 | -6.0968196281257 | 3.3283434473479 | 0.000237947169541929 |
| Nfs1 | ENSMUSG00000027618 | 18041 | -0.433279033281376 | 4.97462707743896 | -6.09517869785741 | 3.3373809895651 | 0.000238421798452938 |
| Abi2 | ENSMUSG00000026782 | 329165 | 0.577596219288791 | 3.65692761483302 | 6.09482276097021 | 3.3393447391039 | 0.000238421798452938 |
| App | ENSMUSG00000022892 | 11820 | 0.536592921984234 | 7.19082772435798 | 6.09408153625006 | 3.3434380693836 | 0.000238506075936337 |
| Cdnf | ENSMUSG00000039496 | 227526 | -0.52178194664836 | 4.42612094719855 | -6.09381903265203 | 3.3448889800604 | 0.000238506075936337 |
| Wdfy3 | ENSMUSG00000043940 | 72145 | 0.408332281770001 | 6.25422501869058 | 6.09227503832023 | 3.3534363518786 | 0.000238959665986864 |
| Gpbp1 | ENSMUSG00000032745 | 73274 | -0.347545529281613 | 5.1126859038582 | -6.09162911161055 | 3.3570189331412 | 0.000239059113733987 |
| Elmod3 | ENSMUSG00000056698 | 232089 | -0.70856621867723 | 2.6135727332409 | -6.08787414049765 | 3.3779253876780 | 0.000240391291749405 |
| Kcnn2 | ENSMUSG00000054477 | 140492 | -1.24218101848408 | 0.788652770082168 | -6.08515556933354 | 3.3931468617033 | 0.000241317425798824 |
| Basp1 | ENSMUSG00000045763 | 70350 | 1.24616642253125 | 1.6449342769965 | 6.0835474233024 | 3.4021848986411 | 0.00024180288119016 |
| Ndufs6 | ENSMUSG00000021606 | 407785 | -0.53811141734103 | 4.95170095683422 | -6.08270667282959 | 3.4069201165858 | 0.000241982090931774 |
| Tti1 | ENSMUSG00000027650 | 75425 | -0.448000832071918 | 3.75235832806796 | -6.08106771230053 | 3.4161708786556 | 0.000242481583601201 |
| Tgfbr3 | ENSMUSG00000029287 | 21814 | 0.458269425770475 | 5.33330023976126 | 6.06731703261178 | 3.4948311022079 | 0.000247903950540135 |
| Ammecr1 | ENSMUSG00000042225 | 56068 | 0.838497311452979 | 2.50915141001143 | 6.06444178740681 | 3.5115180766544 | 0.000248926096601233 |
| Sdk1 | ENSMUSG00000039683 | 330222 | 0.999136111513576 | 1.45905199805716 | 6.06322041964351 | 3.5186318051118 | 0.00024925205726146 |
| Tmem233 | ENSMUSG00000079278 | 545798 | -1.69994380057933 | 2.21523572743052 | -6.06286979836104 | 3.5206767579516 | 0.00024925205726146 |
| Sh3bp2 | ENSMUSG00000054520 | 24055 | 1.28407736310567 | 1.45189034507558 | 6.05912446577745 | 3.5425989915571 | 0.000250607265088883 |
| Pgghg | ENSMUSG00000062031 | 212974 | 0.938654057524008 | 1.56559763775412 | 6.05881712479471 | 3.5444042798226 | 0.000250607265088883 |
| Relb | ENSMUSG00000002983 | 19698 | 0.838854932414558 | 2.56960776267074 | 6.05829350658182 | 3.5474821839257 | 0.000250662752116953 |
| Slc28a2 | ENSMUSG00000027219 | 269346 | -0.879733852697447 | 2.80847854329318 | -6.04953071126379 | 3.5994099871987 | 0.000254167639341537 |
| Ctnnal1 | ENSMUSG00000038816 | 54366 | -0.727297410231874 | 3.19084921495064 | -6.04778659406969 | 3.6098404704981 | 0.000254739613834829 |
| Immt | ENSMUSG00000052337 | 76614 | -0.424097748023643 | 7.53076081635274 | -6.04450371076355 | 3.6295595842776 | 0.000255965908488593 |
| Sec31b | ENSMUSG00000051984 | 240667 | -0.637591030255467 | 3.31304240610782 | -6.03891303353908 | 3.6634016689008 | 0.000258077517540698 |
| Col4a5 | ENSMUSG00000031274 | 12830 | 0.946865367575276 | 2.21712751534281 | 6.03877788872835 | 3.6642238333470 | 0.000258077517540698 |
| Tpbg | ENSMUSG00000035274 | 21983 | 1.82600491180155 | 0.423312448943215 | 6.03827233396089 | 3.6673011359933 | 0.000258127937653208 |
| Me2 | ENSMUSG00000024556 | 107029 | 0.776892085087434 | 1.79073616167764 | 6.0360364305895 | 3.6809436281916 | 0.000258921459457937 |
| Cox6c | ENSMUSG00000014313 | 12864 | -0.61157819083587 | 8.65870895905935 | -6.03476722978393 | 3.6887114392076 | 0.000259300995125265 |
| Stk17b | ENSMUSG00000026094 | 98267 | 0.925692092198382 | 2.36318475322821 | 6.03402608319347 | 3.6932553941628 | 0.000259402280383972 |
| Lrrc59 | ENSMUSG00000020869 | 98238 | 0.416684231451607 | 4.97245931586754 | 6.03375832511907 | 3.6948984590416 | 0.000259402280383972 |
| Popdc2 | ENSMUSG00000022803 | 64082 | 0.840979549881282 | 3.98637286895241 | 6.03032651328732 | 3.7160254732405 | 0.00026071806449289 |
| Eln | ENSMUSG00000029675 | 13717 | 1.07052326211931 | 4.86443536452776 | 6.02843935697579 | 3.7276972743464 | 0.000261369204014629 |
| Sacs | ENSMUSG00000048279 | 50720 | 0.94965363076583 | 6.91435530731081 | 6.02761243557078 | 3.7328237964122 | 0.000261560877683219 |
| Scn3b | ENSMUSG00000049281 | 235281 | 1.34349025492888 | 2.0788178875463 | 6.02621493876996 | 3.7415044560343 | 0.000262001186476047 |
| Dupd1 | ENSMUSG00000063821 | 435391 | 0.832527590728183 | 3.08128278422369 | 6.02091572794761 | 3.7746140332908 | 0.00026415048654227 |
| Pik3r5 | ENSMUSG00000020901 | 320207 | 1.05354258071207 | 1.92582270400278 | 6.0174746430582 | 3.7962786850705 | 0.000265496623842013 |
| Rtf1 | ENSMUSG00000027304 | 76246 | -0.55193489098068 | 5.11133380716184 | -6.01642082159138 | 3.8029395082888 | 0.000265792402590191 |
| Unc45b | ENSMUSG00000018845 | 217012 | -0.386972348950647 | 6.60533793951555 | -6.01481424662057 | 3.8131177161613 | 0.000266333480864921 |
| Olfr1033 | ENSMUSG00000045392 | 258571 | 1.08354555524034 | 1.74440135684256 | 6.01429025003412 | 3.8164435999369 | 0.000266395561883212 |
| Coq2 | ENSMUSG00000029319 | 71883 | -0.505771878734943 | 4.74937526523619 | -6.01302680132111 | 3.8244754313354 | 0.000266785838799795 |
| Cd200 | ENSMUSG00000022661 | 17470 | -0.633998670096563 | 3.56761631078679 | -6.01183969519134 | 3.8320381245975 | 0.000267142912882501 |
| Aldh18a1 | ENSMUSG00000025007 | 56454 | 0.913642204294 | 2.22818766700559 | 6.00974131844407 | 3.8454446906003 | 0.000267906666111872 |
| Rfx7 | ENSMUSG00000037674 | 319758 | 0.441228886668919 | 5.31758007240854 | 6.00753507645045 | 3.8595935448713 | 0.000268721127636868 |
| Zfp27 | ENSMUSG00000062040 | 22689 | -0.57017831624704 | 3.16732833590871 | -6.00616315188117 | 3.8684193970411 | 0.000268954892027637 |
| Rnf128 | ENSMUSG00000031438 | 66889 | -0.650105294551847 | 3.16123713520805 | -6.00557436790515 | 3.8722136543149 | 0.000268954892027637 |
| Glo1 | ENSMUSG00000024026 | 109801 | -0.346523438806669 | 5.24188216797982 | -6.00555810146357 | 3.8723185344107 | 0.000268954892027637 |
| Sgms2 | ENSMUSG00000050931 | 74442 | 1.29237147454123 | 2.33252400847668 | 6.00548452365987 | 3.8727929745814 | 0.000268954892027637 |
| Cmtm4 | ENSMUSG00000096188 | 97487 | -0.55807191239774 | 2.82275413881818 | -6.00497290062043 | 3.8760936814352 | 0.000269013206550912 |
| Angpt2 | ENSMUSG00000031465 | 11601 | -1.69712862727584 | 0.739723750808071 | -6.00187382203431 | 3.8961504591069 | 0.000270233633683361 |
| Txndc5 | ENSMUSG00000038991 | 105245 | 0.599808380261949 | 4.90522783552965 | 5.99930616917073 | 3.9128504808783 | 0.000271219838975785 |
| Cdkn1c | ENSMUSG00000037664 | 12577 | 0.682299961526288 | 3.66692649294076 | 5.99598109456463 | 3.9345885539012 | 0.000272553786328863 |
| Actb | ENSMUSG00000029580 | 11461 | 0.446007277453662 | 8.57624970852755 | 5.99452550653273 | 3.9441445103925 | 0.000273042708316029 |
| Arsb | ENSMUSG00000042082 | 11881 | 0.66731887533822 | 2.97604429616603 | 5.9911832147656 | 3.9661790510029 | 0.000274394324091854 |
| Pla2g15 | ENSMUSG00000031903 | 192654 | 0.763822021055145 | 2.01431739688945 | 5.98567764594017 | 4.0027575972477 | 0.000276749799465626 |
| Atg9a | ENSMUSG00000033124 | 245860 | -0.370159794932587 | 5.42465649868055 | -5.98454868636795 | 4.0103019814900 | 0.000277096150187535 |
| Akap7 | ENSMUSG00000039166 | 432442 | -0.484846603607109 | 4.06893931170972 | -5.97987537538587 | 4.0416910317328 | 0.0002790885955014 |
| Ppm1a | ENSMUSG00000021096 | 19042 | -0.371332997330275 | 7.04530776231184 | -5.97829605004788 | 4.0523570595025 | 0.000279648453392819 |
| Hmga1 | ENSMUSG00000046711 | 15361 | 0.390808610921315 | 4.74755218843768 | 5.97750807782485 | 4.0576897043654 | 0.000279839786488448 |
| Gpd2 | ENSMUSG00000026827 | 14571 | -0.810280860764578 | 3.64661676207414 | -5.97558635011971 | 4.0707260216430 | 0.000280561829398361 |
| Rtn2 | ENSMUSG00000030401 | 20167 | -0.632025138571507 | 6.63164928826494 | -5.9751148000679 | 4.0739315637276 | 0.000280605834424117 |
| Ace | ENSMUSG00000020681 | 11421 | 0.531965889698999 | 5.78416886459846 | 5.97199405802122 | 4.0952129245591 | 0.000281894033239017 |
| Sgsh | ENSMUSG00000005043 | 27029 | 0.719206926419716 | 1.92629470731213 | 5.97047705865532 | 4.1055999684466 | 0.000282431172152864 |
| Xirp2 | ENSMUSG00000027022 | 241431 | 0.60433057314466 | 12.1284906317503 | 5.96739870544764 | 4.1267628635342 | 0.000283708458247124 |
| Mgll | ENSMUSG00000033174 | 23945 | -0.703544405155898 | 5.40593345612854 | -5.96638572770083 | 4.1337518475130 | 0.000284010317065775 |
| Plekhb2 | ENSMUSG00000026123 | 226971 | -0.419582114074132 | 5.1899580896839 | -5.96399691882332 | 4.1502825541579 | 0.000284966951001884 |
| Ccl8 | ENSMUSG00000009185 | 20307 | 5.33519653393384 | -0.670115320066419 | 5.96272347208201 | 4.1591232318059 | 0.000285315486807133 |
| Mast3 | ENSMUSG00000031833 | 546071 | 0.772156393434479 | 2.01409141105369 | 5.96251405585948 | 4.1605789586549 | 0.000285315486807133 |
| Ttyh3 | ENSMUSG00000036565 | 78339 | 0.627806405106085 | 3.03303723242259 | 5.96189405337465 | 4.1648919540997 | 0.000285432187775955 |
| Ppt1 | ENSMUSG00000028657 | 19063 | 0.537564090641179 | 3.87312196115431 | 5.96112048774349 | 4.1702797814463 | 0.000285499947031625 |
| Tbxas1 | ENSMUSG00000029925 | 21391 | 1.5353183755076 | 0.63297279453741 | 5.96100219305232 | 4.1711043400375 | 0.000285499947031625 |
| 4833439L19Rik | ENSMUSG00000025871 | 97820 | -0.380392348020293 | 6.50872171902538 | -5.9582540313782 | 4.1903083168155 | 0.000286541814784935 |
| Kcmf1 | ENSMUSG00000055239 | 74287 | -0.406697338046806 | 6.5755319367418 | -5.95807414505185 | 4.1915685833053 | 0.000286541814784935 |
| Gzf1 | ENSMUSG00000027439 | 74533 | -0.519101083308582 | 5.23049478304044 | -5.95678915404246 | 4.2005826897159 | 0.000286978558633029 |
| Musk | ENSMUSG00000057280 | 18198 | 0.498791039903079 | 5.10549483369515 | 5.95631424118769 | 4.2039193129533 | 0.000287016200585651 |
| Cth | ENSMUSG00000028179 | 107869 | -2.53878905027197 | 0.533023638423289 | -5.95577041644436 | 4.2077435120571 | 0.000287016200585651 |
| Mrpl30 | ENSMUSG00000026087 | 107734 | -0.497384793325751 | 5.48962463686006 | -5.95559031603674 | 4.2090107907675 | 0.000287016200585651 |
| Cyth4 | ENSMUSG00000018008 | 72318 | 1.34108471095623 | 2.50750978307945 | 5.95459509319402 | 4.2160209168550 | 0.000287314991534554 |
| Zfp90 | ENSMUSG00000031907 | 22751 | 0.953601985347547 | 1.37839805075116 | 5.95421249669383 | 4.2187191013548 | 0.000287319741413766 |
| Htt | ENSMUSG00000029104 | 15194 | 0.449156154944465 | 4.49558089314888 | 5.95371151202818 | 4.2222549357181 | 0.000287381498769212 |
| Baz1a | ENSMUSG00000035021 | 217578 | 0.645678147030259 | 3.21305464097064 | 5.95182413883095 | 4.2356035430818 | 0.000288110655441364 |
| Phldb1 | ENSMUSG00000048537 | 102693 | 0.439404524866873 | 5.77542779225041 | 5.94991812898314 | 4.2491289277311 | 0.000288850922319835 |
| Ankrd33b | ENSMUSG00000022237 | 67434 | -0.435234212569001 | 6.07779085838734 | -5.94834585339221 | 4.2603201994296 | 0.000289431697327317 |
| Zmynd11 | ENSMUSG00000021156 | 66505 | -0.345551445179482 | 6.28783686184458 | -5.94541793661325 | 4.2812433045584 | 0.000290672487963533 |
| Cox7c | ENSMUSG00000017778 | 12867 | -0.568015444457707 | 6.76723753880381 | -5.94112549787206 | 4.3121125776877 | 0.000292586608235286 |
| Jund | ENSMUSG00000071076 | 16478 | 0.511184463670231 | 6.1591642600219 | 5.94064493551058 | 4.3155830783976 | 0.000292640438151144 |
| Mia2 | ENSMUSG00000021000 | 338320 | -0.382062333032531 | 5.61802737553662 | -5.93968128987264 | 4.3225511224876 | 0.000292931223310056 |
| Azin1 | ENSMUSG00000037458 | 54375 | 0.711064093061979 | 5.26227654346259 | 5.93810179558037 | 4.3339978727879 | 0.000293524973652073 |
| Emp3 | ENSMUSG00000040212 | 13732 | 0.829017057615552 | 1.60054177761894 | 5.93736411800718 | 4.3393547906569 | 0.000293705803199201 |
| P4hb | ENSMUSG00000025130 | 18453 | 0.391353737406849 | 6.99117017583736 | 5.93623163475002 | 4.3475922647894 | 0.00029408125647533 |
| 2210416O15Rik | ENSMUSG00000018378 | 70393 | -0.48847569140523 | 6.10206321374741 | -5.93554944404199 | 4.3525623143366 | 0.000294235365850428 |
| Dlat | ENSMUSG00000000168 | 235339 | -0.511507361635507 | 6.92806811198437 | -5.93313221990371 | 4.3702208775776 | 0.000295075841119695 |
| Cdc42se1 | ENSMUSG00000046722 | 57912 | 0.479603711128314 | 4.12930162817689 | 5.93310854922541 | 4.3703941704582 | 0.000295075841119695 |
| Pla2g4a | ENSMUSG00000056220 | 18783 | 1.03772085314748 | 1.14878898213102 | 5.93176970667488 | 4.3802075786499 | 0.000295543123933612 |
| Dusp27 | ENSMUSG00000026564 | 240892 | 0.534158818804368 | 6.89598332807323 | 5.9313473664192 | 4.3833080322517 | 0.000295543123933612 |
| 0610012G03Rik | ENSMUSG00000107002 | 106264 | -0.548335885054724 | 4.5712365555084 | -5.93105900093695 | 4.3854262832340 | 0.000295543123933612 |
| Gfm2 | ENSMUSG00000021666 | 320806 | -0.488783039651266 | 5.30946048709817 | -5.93021446756253 | 4.3916361691402 | 0.000295779266573458 |

| | | | | | | | |
|---|---|---|---|---|---|---|---|
| Txnrd1 | ENSMUSG00000020250 | 50493 | -0.405882302524041 | 5.55553186216742 | -5.92729727121113 | 4.4131575216075 | 0.000297045719634801 |
| COX1 | ENSMUSG00000064351 | 17708 | -0.702692633857961 | 14.0151416235388 | -5.92662084625849 | 4.4181635772889 | 0.000297199668082125 |
| Nfxl1 | ENSMUSG00000072889 | 100978 | 0.898450674360716 | 2.04441790326021 | 5.91854350521926 | 4.4784046434116 | 0.00030106667378306 |
| Tspan13 | ENSMUSG00000020577 | 66109 | -0.481440612229153 | 3.96267017035705 | -5.91292075712758 | 4.5208477581028 | 0.000303733170521341 |
| Fam102b | ENSMUSG00000040339 | 329739 | 0.766514208416064 | 4.19981326510394 | 5.91159290490856 | 4.5309324878251 | 0.000304150379605565 |
| Trim54 | ENSMUSG00000062077 | 58522 | -0.470327930383191 | 6.52489487547497 | -5.91137068168027 | 4.5326225265526 | 0.000304150379605565 |
| Paqr9 | ENSMUSG00000064225 | 75552 | -0.712597811454643 | 2.87020896261804 | -5.9101251552758 | 4.5421071821248 | 0.000304560956072096 |
| Dpf3 | ENSMUSG00000021221 | 70127 | -0.519712381445529 | 3.07491689958692 | -5.90947517421626 | 4.5470650418438 | 0.000304560956072096 |
| Mrm3 | ENSMUSG00000038046 | 67390 | -0.65545215515402 | 3.02017661347545 | -5.90947061860375 | 4.5470998107186 | 0.000304560956072096 |
| Aifm1 | ENSMUSG00000036932 | 26926 | -0.498899177586348 | 4.93408693057778 | -5.90895430704567 | 4.5510421586734 | 0.000304638345599872 |
| Mrpl50 | ENSMUSG00000044018 | 28028 | -0.512930522581676 | 3.85800860446867 | -5.90559957161186 | 4.5767450989879 | 0.000306171362772567 |
| 1700017B05Rik | ENSMUSG00000032300 | 74211 | 0.991757470113011 | 2.03066586586273 | 5.90311252413298 | 4.5958983880908 | 0.000307264619450895 |
| Nt5c | ENSMUSG00000020736 | 50773 | 1.03803163951604 | 1.88660925454618 | 5.89837205206191 | 4.6326391152545 | 0.000309345474962123 |
| Rasa3 | ENSMUSG00000031453 | 19414 | 0.574800963417398 | 3.64144799490589 | 5.89836645602266 | 4.6326826686762 | 0.000309345474962123 |
| Ndufs5 | ENSMUSG00000028648 | 595136 | -0.468877194029083 | 6.1226937662845 | -5.89252877907912 | 4.6783515851000 | 0.000312204280688211 |
| Tgtp1 | ENSMUSG00000078922 | 21822 | -1.64561874577303 | 2.2815832068934 | -5.88864994706612 | 4.7089572346080 | 0.000314054981888352 |
| Btbd2 | ENSMUSG00000003344 | 208198 | -0.538904413657978 | 3.94762934358669 | -5.88671517224842 | 4.7243019012177 | 0.000314886244403729 |
| Hmgn3 | ENSMUSG00000066456 | 94353 | 0.962514690767144 | 1.36926207793816 | 5.88593394376241 | 4.7305126723460 | 0.000315108068381564 |
| Fcgr1 | ENSMUSG00000015947 | 14129 | 2.33655976936894 | 0.0907077505903283 | 5.88443836338391 | 4.7424264564595 | 0.00031558100023489 |
| Aqp1 | ENSMUSG00000004655 | 11826 | -0.676900056239019 | 6.31228239752282 | -5.88431801151258 | 4.7433865463903 | 0.00031558100023489 |
| Dynll1 | ENSMUSG00000009013 | 56455 | 0.652556476201752 | 4.79759861927454 | 5.88336929803617 | 4.7509619169061 | 0.000315892729402072 |
| Stk11 | ENSMUSG00000003068 | 20869 | -0.343411883959428 | 5.81304422054218 | -5.88091166025766 | 4.7706449710252 | 0.000317008633302595 |
| Pkn1 | ENSMUSG00000057672 | 320795 | 0.42686392391484 | 5.64804847662321 | 5.87968659388196 | 4.7804883808581 | 0.000317469735669264 |
| Dsel | ENSMUSG00000038702 | 319901 | 0.856325019517493 | 2.85362620508892 | 5.87861924542978 | 4.7890818979546 | 0.000317847323779858 |
| Rnf145 | ENSMUSG00000019189 | 74315 | 0.635956030475656 | 2.79220564482126 | 5.87511658769284 | 4.8173968095104 | 0.000319532551727907 |
| Sesn3 | ENSMUSG00000032009 | 75747 | 0.370460853552878 | 5.55967167637558 | 5.87250644903662 | 4.8386109220975 | 0.000320745033289558 |
| Tbc1d24 | ENSMUSG00000036473 | 224617 | -0.539271149497719 | 3.08565228655634 | -5.87178819240775 | 4.8444657905964 | 0.000320938518527329 |
| Tfb2m | ENSMUSG00000026492 | 15278 | -0.467379648497618 | 4.57282795082716 | -5.87122506280881 | 4.8490613391055 | 0.000321048391870156 |
| Tspan5 | ENSMUSG00000028152 | 56224 | 0.97251860691468 | 1.92173024826315 | 5.86947253265411 | 4.8633925521568 | 0.000321802324380302 |
| Ipmk | ENSMUSG00000060733 | 69718 | -0.413764039926822 | 4.74145574880764 | -5.86779294149429 | 4.8771690150952 | 0.000322373997026124 |
| Fbxo40 | ENSMUSG00000047746 | 207215 | -0.471628466691803 | 7.76536185463659 | -5.86770023936317 | 4.8779305743409 | 0.000322373997026124 |
| F2rl1 | ENSMUSG00000021678 | 14063 | 2.22498116477488 | 1.71160242437646 | 5.86304231887948 | 4.9163570247384 | 0.000324717212310672 |
| Pnkd | ENSMUSG00000026179 | 56695 | -0.411472939547067 | 4.09744486492535 | -5.86034625483695 | 4.9387437267147 | 0.000325998838627531 |
| March6 | ENSMUSG00000039100 | 223455 | -0.408121427040363 | 6.79571034520601 | -5.85867951297877 | 4.9526369452103 | 0.000326718614653554 |
| Tstd3 | ENSMUSG00000028251 | 77032 | -0.719642872734954 | 2.92428914135402 | -5.85559991052269 | 4.9784151103387 | 0.000328221083058583 |
| Rassf9 | ENSMUSG00000044921 | 237504 | -1.07708225520421 | 2.77932849387576 | -5.85340315832628 | 4.9968892596013 | 0.000329240485212187 |
| Wsb2 | ENSMUSG00000029364 | 59043 | 0.336446260671774 | 6.01290177851969 | 5.85081156637665 | 5.0187764167796 | 0.000330483403679068 |
| Scn2a | ENSMUSG00000075318 | 110876 | 0.843087591074047 | 3.29308920918785 | 5.84867152532585 | 5.0369258258927 | 0.000331478845290992 |
| Gpr157 | ENSMUSG00000047875 | 269604 | -1.42586176120914 | 4.00331789224225 | -5.84775236958006 | 5.0447421888326 | 0.000331793482949033 |
| Bcl7c | ENSMUSG00000030814 | 12055 | 0.579549227531684 | 3.27516961213244 | 5.84497010045708 | 5.0684798289255 | 0.000333154257426251 |
| Emilin1 | ENSMUSG00000029163 | 100952 | 1.09642308949679 | 3.55902166840275 | 5.84461208736404 | 5.0715428061631 | 0.000333155254892847 |
| Cav2 | ENSMUSG00000000058 | 12390 | -0.783093612940155 | 3.99332139258321 | -5.83931365883199 | 5.1171011459352 | 0.00033594614189921 |
| Kif5b | ENSMUSG00000006740 | 16573 | 0.473915503905455 | 8.1198672193822 | 5.83734247979313 | 5.1341596330637 | 0.000336863739189797 |
| Cdr2l | ENSMUSG00000050910 | 237988 | 0.851461382103259 | 1.84734061881619 | 5.83560559647679 | 5.1492399691767 | 0.000337278510343156 |
| Cd300lg | ENSMUSG00000017309 | 52685 | -1.00592862277782 | 3.84264009641206 | -5.83522807785758 | 5.1525238833327 | 0.000337278510343156 |
| Timp3 | ENSMUSG00000020044 | 21859 | -0.694072508213316 | 6.07035873186665 | -5.83501487288101 | 5.1543794559683 | 0.000337278510343156 |
| Pgpep1l | ENSMUSG00000030553 | 78444 | 2.5334573030949 | -0.0592367126216085 | 5.83474352363509 | 5.1567420854451 | 0.000337278510343156 |
| Zfp991 | ENSMUSG00000067916 | 666532 | 0.94049222319118 | 1.33169268452052 | 5.83461751516347 | 5.1578396234789 | 0.000337278510343156 |
| Efhd2 | ENSMUSG00000040659 | 27984 | 0.787853036896065 | 3.62968599318546 | 5.83448497448993 | 5.1589943215968 | 0.000337278510343156 |
| Ifi207 | ENSMUSG00000073490 | 226691 | 1.20765170401676 | 2.38534606644312 | 5.83395084338906 | 5.1636504355746 | 0.000337381129176728 |
| Tns1 | ENSMUSG00000055322 | 21961 | -0.441424334995332 | 8.5871744869173 | -5.83139346047462 | 5.1860047370405 | 0.000338639293790859 |
| Ppp1r14b | ENSMUSG00000056612 | 18938 | 0.445397869495977 | 4.81520602483139 | 5.82810684364997 | 5.2148824704549 | 0.000340321673340553 |
| Lrrc20 | ENSMUSG00000037151 | 216011 | -0.462021496447714 | 5.87049092560012 | -5.82643253564967 | 5.2296585073875 | 0.000341082321863084 |
| Enah | ENSMUSG00000022995 | 13800 | 0.641908005551077 | 7.03910912054441 | 5.82324574051394 | 5.2579039370741 | 0.000342650229503512 |
| Ndufaf1 | ENSMUSG00000027305 | 69702 | -0.583594374465524 | 3.88936144978767 | -5.8230135840946 | 5.2599678447250 | 0.000342650229503512 |
| Tgfbr2 | ENSMUSG00000032440 | 21813 | 0.630994684507806 | 5.99581128569622 | 5.82234265091212 | 5.2659373251247 | 0.000342834787974622 |
| Itgb1 | ENSMUSG00000025809 | 16412 | 0.421211034958366 | 7.58310444223667 | 5.82183939810627 | 5.2704195692651 | 0.000342922358997844 |
| Pdhx | ENSMUSG00000010914 | 27402 | -0.484015731860506 | 5.49576006754753 | -5.82045483292232 | 5.2827719210169 | 0.000343521593507654 |
| Cisd1 | ENSMUSG00000037710 | 52637 | -0.453021361471015 | 5.68561530099503 | -5.81752647091384 | 5.3089973223896 | 0.000345021698757679 |
| Atpif1 | ENSMUSG00000054428 | 11983 | 0.52622128241904 | 4.17903007837659 | 5.81690367286783 | 5.3145924725952 | 0.000345180096957447 |
| Fam171a2 | ENSMUSG00000034685 | 217219 | 1.40590744360563 | 1.01380965257152 | 5.81548068112389 | 5.3273996945980 | 0.000345806449297219 |
| Arl4c | ENSMUSG00000049866 | 320982 | 1.31835903176769 | 1.89896146391069 | 5.81484425396386 | 5.3331381423007 | 0.000345973489812992 |
| Coa7 | ENSMUSG00000048351 | 69893 | -0.634224298960592 | 3.02491641337941 | -5.81105192183139 | 5.3674669740909 | 0.000347993959037888 |
| Myh10 | ENSMUSG00000020900 | 77579 | 0.57346088847635 | 7.53622484405959 | 5.81063222027279 | 5.3712803996648 | 0.000348034772073127 |
| Eml1 | ENSMUSG00000058070 | 68519 | -0.379236669786913 | 5.11698991879606 | -5.80999691942817 | 5.3770581752082 | 0.000348202742376783 |
| Prdx5 | ENSMUSG00000024953 | 54683 | -0.413238072703737 | 5.23120030723268 | -5.80653945272298 | 5.4086167335405 | 0.000349939589177745 |
| Cdh5 | ENSMUSG00000031871 | 12562 | -0.690830700201375 | 6.63275873114772 | -5.80635761851495 | 5.4102818196906 | 0.000349939589177745 |
| Ddb1 | ENSMUSG00000024740 | 13194 | 0.412890039410977 | 7.07353373638557 | 5.80593533126814 | 5.4141508484285 | 0.000349982749403738 |
| Mcub | ENSMUSG00000027994 | 66815 | 1.45731368634418 | 0.686598425571906 | 5.8055294395275 | 5.4178723948246 | 0.000350016330660923 |
| Nid1 | ENSMUSG00000005397 | 18073 | 0.600420554142031 | 7.06992345495615 | 5.80480091125224 | 5.4245588656406 | 0.000350241305140685 |
| Plet1os | ENSMUSG00000101304 | 69631 | -1.38365969149252 | 0.823939652018656 | -5.80318209373215 | 5.4394474523480 | 0.000350995278049682 |
| Adpgk | ENSMUSG00000025236 | 72141 | 0.682621966042542 | 3.09307436056147 | 5.80101050670306 | 5.4594872959257 | 0.000352080564199203 |
| Scamp2 | ENSMUSG00000040188 | 24044 | 0.488818236442792 | 3.48960811434094 | 5.80051142196159 | 5.4641038817373 | 0.000352089956716541 |
| Msh2 | ENSMUSG00000024151 | 17685 | -0.729584811351441 | 2.90558869564295 | -5.80029846685098 | 5.4660749844293 | 0.000352089956716541 |
| Nol3 | ENSMUSG00000014776 | 78688 | 0.435536790642281 | 5.93965164592525 | 5.79251067835648 | 5.5386729874548 | 0.000356556150917958 |
| Kyat3 | ENSMUSG00000040213 | 229905 | -0.6757779462481 | 2.53649642601765 | -5.79147394303572 | 5.5484134457840 | 0.000356972968663126 |
| Hint3 | ENSMUSG00000019791 | 66847 | -0.521566344723594 | 3.90408646520797 | -5.78888283504594 | 5.5728363495126 | 0.000358333377273663 |
| 0610009L18Rik | ENSMUSG00000043644 | 66838 | -0.882875514513726 | 2.05860534407592 | -5.78831401506525 | 5.5782129254798 | 0.00035846822744515 |
| Zfp516 | ENSMUSG00000058881 | 329003 | 0.398287496261687 | 4.29543549332426 | 5.78653126222209 | 5.5950990468823 | 0.00035934211328714 |
| Hspa13 | ENSMUSG00000032932 | 110920 | 0.594505257608926 | 3.19259934579504 | 5.78501434104428 | 5.6095094280371 | 0.000359977190784619 |
| Spg7 | ENSMUSG00000000738 | 234847 | -0.520992697094947 | 5.81445426177308 | -5.78479736678446 | 5.6115738093220 | 0.000359977190784619 |
| Cpeb4 | ENSMUSG00000020300 | 67579 | -0.461924913432043 | 6.38972755726869 | -5.77864508351143 | 5.6704418423998 | 0.000363540174658492 |
| Trp53bp1 | ENSMUSG00000043909 | 27223 | 0.487517237763601 | 4.41880559328515 | 5.77704949581006 | 5.6858146824568 | 0.000364312076752261 |
| Stx11 | ENSMUSG00000039232 | 74732 | 1.59917231629829 | 0.633186866134746 | 5.77343125176338 | 5.7208370940575 | 0.000366341360721401 |
| Kcnj8 | ENSMUSG00000030247 | 16523 | -0.96956649243457 | 3.60479852574731 | -5.77173350983524 | 5.7373480893213 | 0.000367183559510371 |
| Sbk1 | ENSMUSG00000042978 | 104175 | 0.583376099353865 | 4.94259304788706 | 5.77013122470037 | 5.7529765654559 | 0.000367895547107159 |
| Tmem150c | ENSMUSG00000050640 | 231503 | -1.13427283117223 | 1.15132175220637 | -5.76990319802254 | 5.7552043322042 | 0.000367895547107159 |
| Tmcc1 | ENSMUSG00000030126 | 330401 | -0.403729938051132 | 4.94669692461887 | -5.76778275837683 | 5.7759638825014 | 0.000369006786672259 |
| Hmox2 | ENSMUSG00000004070 | 15369 | -0.484661611628415 | 4.51003154635083 | -5.76565802739402 | 5.7968441940063 | 0.00037012443857876 |
| Adam19 | ENSMUSG00000011256 | 11492 | 1.11038887193387 | 3.88242266941374 | 5.76450380575749 | 5.8082201975331 | 0.000370634296434527 |
| Lamp2 | ENSMUSG00000016534 | 16784 | 0.425196881089739 | 6.05598463941972 | 5.76137119067838 | 5.8392133398835 | 0.000372394638379621 |
| Atat1 | ENSMUSG00000024426 | 73242 | 1.02500562115603 | 1.14060174138406 | 5.7580638907313 | 5.8721229547950 | 0.000374117673034165 |
| Cdk14 | ENSMUSG00000028926 | 18647 | 0.73254699524839 | 2.77403090979305 | 5.75796841249718 | 5.8730759027227 | 0.000374117673034165 |
| Src | ENSMUSG00000027646 | 20779 | 0.966529975912707 | 1.64001183106737 | 5.75705302047075 | 5.8822204669226 | 0.0003744819564585 |
| Ak3 | ENSMUSG00000024782 | 56248 | -0.429700949790958 | 4.86352142220035 | -5.75555056764096 | 5.8972620033821 | 0.000375221018387487 |
| B3gnt9 | ENSMUSG00000069920 | 97440 | 1.36148645656956 | 0.949681868015536 | 5.75487279597273 | 5.9040605829379 | 0.000375435056614864 |
| Rnf38 | ENSMUSG00000035696 | 73469 | 0.446366646002539 | 3.69004144119901 | 5.75436138278203 | 5.9091958946389 | 0.000375543141420342 |
| Osgepl1 | ENSMUSG00000026096 | 72085 | -0.545406223922282 | 3.77098955876771 | -5.75188969627568 | 5.9340811979280 | 0.000376905529195538 |
| Fbxo32 | ENSMUSG00000022358 | 67731 | -0.471247227041227 | 7.97775986878128 | -5.75016994151532 | 5.9514607262471 | 0.000377603630027552 |
| Cmc2 | ENSMUSG00000014633 | 66531 | -0.642684348351119 | 2.57665344207719 | -5.75011852976823 | 5.9519811045418 | 0.000377603630027552 |
| Gnl1 | ENSMUSG00000024429 | 14670 | 0.506907077075362 | 3.94718366110075 | 5.7487588916072 | 5.9657603818442 | 0.000378258275718909 |
| Ptpn1 | ENSMUSG00000027540 | 19246 | 0.521032517461686 | 4.10219577242172 | 5.74013338619463 | 6.0539581791734 | 0.00038362792380606 |
| Mrps14 | ENSMUSG00000058267 | 64659 | -0.537710171379666 | 4.4095939269244 | -5.73959379379062 | 6.0595208593135 | 0.000383757951988162 |
| Sox4 | ENSMUSG00000076431 | 20677 | 1.07264384911535 | 3.44982480396779 | 5.73825775370599 | 6.0733172153109 | 0.000384408977883981 |
| Togaram2 | ENSMUSG00000045761 | 320159 | 0.914012936967461 | 3.62346353833356 | 5.73376926796628 | 6.1199082535840 | 0.000387133779629209 |
| Ikbkb | ENSMUSG00000031537 | 16150 | 0.52693034366345 | 4.11556912412996 | 5.733326670669 | 6.1245227134559 | 0.000387201606597957 |
| Tor1aip1 | ENSMUSG00000026466 | 208263 | -0.393981962831774 | 6.78034923510819 | -5.73198709889454 | 6.1385111400488 | 0.000387861648970369 |
| Nek9 | ENSMUSG00000034290 | 217718 | -0.37096675468245 | 6.34281265543652 | -5.72991295788153 | 6.1602363707963 | 0.000389009496066865 |
| Msn | ENSMUSG00000031207 | 17698 | 0.560091200082869 | 6.69267120556824 | 5.72873800959321 | 6.1725788991495 | 0.000389563856504642 |
| Fmnl1 | ENSMUSG00000055805 | 57778 | 0.659125328362801 | 3.62423702965217 | 5.72789768777743 | 6.1814221557865 | 0.000389896858539545 |
| Ncf2 | ENSMUSG00000026480 | 17970 | 0.976200746860633 | 1.73045751153875 | 5.7247675849603 | 6.2144792826049 | 0.000391755899874017 |
| Plekho2 | ENSMUSG00000050721 | 102595 | 0.655180135720397 | 3.26621493065977 | 5.72272805290926 | 6.2361184895083 | 0.000392893436362051 |
| Pmepa1 | ENSMUSG00000038400 | 65112 | 1.03852208510552 | 5.05371255176402 | 5.71994767161635 | 6.2657452897841 | 0.000394532613840038 |
| Naxe | ENSMUSG00000028070 | 246703 | -0.44906703593883 | 3.31094384674665 | -5.71767094054118 | 6.2901151869997 | 0.000395839085256728 |
| Cox7b | ENSMUSG00000031231 | 66142 | -0.583448866344962 | 7.447567050501 | -5.71554174143553 | 6.3129957978596 | 0.000396997382935675 |
| Col12a1 | ENSMUSG00000032332 | 12816 | 1.26885240943727 | 6.25098903956353 | 5.71528274797112 | 6.3157849137786 | 0.000396997382935675 |

| | | | | | | | |
|---|---|---|---|---|---|---|---|
| Fibp | ENSMUSG00000024911 | 58249 | 0.524400245581781 | 3.02086536276612 | 5.71472664808166 | 6.3217779497061 | 0.00039714571705884 |
| Ptprb | ENSMUSG00000020154 | 19263 | -0.646517917350821 | 6.83275053119086 | -5.71371126189983 | 6.3327360484480 | 0.000397404782625248 |
| Snd1 | ENSMUSG00000001424 | 56463 | 0.428790841793306 | 5.13553728672984 | 5.71346024058808 | 6.3354481501782 | 0.000397404782625248 |
| G3bp2 | ENSMUSG00000029405 | 23881 | -0.438205266768789 | 5.86523231410813 | -5.71333437529852 | 6.3368084906761 | 0.000397404782625248 |
| Ciita | ENSMUSG00000022504 | 12265 | 1.86272720519529 | 1.27033673258565 | 5.70929231337173 | 6.3806578550823 | 0.000399925292510925 |
| Cd33 | ENSMUSG00000004609 | 12489 | 1.17549993847061 | 1.44586987138813 | 5.70715466815436 | 6.4039760141056 | 0.000401156801204518 |
| Pid1 | ENSMUSG00000045658 | 98496 | 0.885860282048 | 2.95512119443979 | 5.70674975391486 | 6.4084030048233 | 0.000401204199574592 |
| Lcp1 | ENSMUSG00000021998 | 18826 | 0.9752869867749 | 4.09734740855221 | 5.70132687889878 | 6.4680021469990 | 0.000404703671830832 |
| Bmi1 | ENSMUSG00000026739 | 12151 | -0.39726255559067 | 5.43247690171151 | -5.7005253054978 | 6.4768608396408 | 0.000405026120355343 |
| Arhgap42 | ENSMUSG00000050730 | 71544 | 0.521217173734903 | 3.43723649556917 | 5.70013172544635 | 6.4812151929154 | 0.000405066685384557 |
| Ptpre | ENSMUSG00000041836 | 19267 | 0.969609383041496 | 2.12988263009431 | 5.69970596735453 | 6.4859289998553 | 0.00040512965655668 |
| Chodl | ENSMUSG00000022860 | 246048 | 0.884106889311706 | 3.22679931274738 | 5.69935715650477 | 6.4897935573722 | 0.000405139539552461 |
| Klhl40 | ENSMUSG00000074001 | 72330 | 0.718624631023037 | 6.99814900570256 | 5.69638707884717 | 6.5227976330337 | 0.000406967471042761 |
| 4930453N24Rik | ENSMUSG00000059920 | 67609 | -0.495850634146295 | 4.38196446507994 | -5.69602634343031 | 6.5268181534594 | 0.000406986019597631 |
| Usp28 | ENSMUSG00000032267 | 235323 | 0.464081208046005 | 6.19463744875853 | 5.69185938540952 | 6.5734489913512 | 0.000409660039478111 |
| Rassf4 | ENSMUSG00000042129 | 213391 | 1.24757563255817 | 3.02826009189888 | 5.68607272004482 | 6.6387850092376 | 0.000413496062313258 |
| Kdelr3 | ENSMUSG00000010830 | 105785 | 1.18654841689352 | 1.04897610540082 | 5.68127108906661 | 6.6935155730838 | 0.000416667532627446 |
| Tomm40l | ENSMUSG00000005674 | 641376 | -0.390496126689106 | 4.32470348892875 | -5.68015225754649 | 6.7063361480895 | 0.000417030670436537 |
| Psat1 | ENSMUSG00000024640 | 107272 | 0.88616475341453 | 1.3076147425391 | 5.68009618066947 | 6.7069794037821 | 0.000417030670436537 |
| Sept2 | ENSMUSG00000026276 | 18000 | 0.341866870784679 | 5.24566729451526 | 5.67825194641646 | 6.7281705985065 | 0.000417801062260131 |
| Edem1 | ENSMUSG00000030104 | 192193 | 0.795002620750256 | 3.22513062701446 | 5.67805780598687 | 6.7304054478251 | 0.000417801062260131 |
| Marcks | ENSMUSG00000069662 | 17118 | 0.700858966661684 | 5.54321265003673 | 5.67787718839933 | 6.7324853270523 | 0.000417801062260131 |
| Rnf139 | ENSMUSG00000037075 | 75841 | -0.382148852919283 | 4.18045444271525 | -5.67768857042886 | 6.7346580523497 | 0.000417801062260131 |
| Exoc6 | ENSMUSG00000053799 | 107371 | -0.517539543619387 | 5.67949834974627 | -5.67687969323794 | 6.7439839903638 | 0.000418037369634375 |
| Mcm6 | ENSMUSG00000026355 | 17219 | 0.741441087751995 | 2.72000532193485 | 5.67669495555828 | 6.7461158177205 | 0.000418037369634375 |
| Rfesd | ENSMUSG00000043190 | 218341 | -0.664695232071318 | 2.17291405556023 | -5.67214010718726 | 6.7989015129880 | 0.000420957862440659 |
| Poldip3 | ENSMUSG00000041815 | 73826 | 0.36597911660257 | 4.10785874269341 | 5.67196429566971 | 6.80094762666e- | 0.000420957862440659 |
| Ubl7 | ENSMUSG00000055720 | 69459 | -0.381765430477987 | 4.70436561419311 | -5.67074978476149 | 6.8150998662423 | 0.00042159511396658 |
| Dok2 | ENSMUSG00000022102 | 13449 | 1.55648617755995 | 0.602733903482539 | 5.66936680769114 | 6.8312527471851 | 0.000422355338119235 |
| Atp5e | ENSMUSG00000016252 | 67126 | -0.621664758679937 | 5.77715264919529 | -5.66865808409184 | 6.8395460250226 | 0.00042244397869632 |
| Pank4 | ENSMUSG00000029056 | 269614 | -0.624685004074703 | 3.84911480687377 | -5.66858381424589 | 6.8404157194445 | 0.00042244397869632 |
| Comp | ENSMUSG00000031849 | 12845 | 1.37160275653973 | 4.86067255534148 | 5.66428971691977 | 6.8908967499680 | 0.00042521178714753 |
| Atp8b2 | ENSMUSG00000060671 | 54667 | 0.701685863790326 | 3.64045492548438 | 5.66411039361952 | 6.8930133274670 | 0.00042521178714753 |
| Rpap3 | ENSMUSG00000022466 | 71919 | -0.450478245376771 | 3.43934818496815 | -5.66368842039152 | 6.8979966220862 | 0.000425279193886208 |
| Mpzl3 | ENSMUSG00000070305 | 319742 | 1.99123210144674 | 0.632523558556392 | 5.66253562933597 | 6.9116297568891 | 0.000425879508864463 |
| Cd81 | ENSMUSG00000037706 | 12520 | 0.478625287239798 | 5.86692253036927 | 5.65915883231183 | 6.9517270668758 | 0.000428108893340959 |
| Crtc1 | ENSMUSG00000003575 | 382056 | 0.487219854755389 | 3.0008137248694 | 5.65543348004979 | 6.9962460323584 | 0.000430607913174044 |
| Kif11 | ENSMUSG00000012443 | 16551 | 1.31719596622701 | 1.11277664076448 | 5.65407366476185 | 7.0125704889929 | 0.000431235599924446 |
| Prkcsh | ENSMUSG00000003402 | 19089 | 0.423251543147778 | 3.73343800544809 | 5.6539269281745 | 7.0143344311194 | 0.000431235599924446 |
| Tnfsf10 | ENSMUSG00000039304 | 22035 | -0.665332549827788 | 2.98935890469478 | -5.64867147375023 | 7.0778183545194 | 0.000434762471628707 |
| Tppp3 | ENSMUSG00000014846 | 67971 | 0.96085907308266 | 6.16982380756937 | 5.64849180422237 | 7.0799993028811 | 0.000434762471628707 |
| Palmd | ENSMUSG00000033377 | 114301 | -0.549996731963048 | 6.21719211720368 | -5.64819255476487 | 7.0836333544115 | 0.000434762471628707 |
| Eaf1 | ENSMUSG00000021890 | 74427 | 0.381488646132531 | 4.27379856281481 | 5.64632087506716 | 7.1064071814984 | 0.000435915470824685 |
| Abcb7 | ENSMUSG00000031333 | 11306 | -0.409886725738921 | 5.79638719859908 | -5.64473319751951 | 7.1257854904853 | 0.000436859008393132 |
| Clcn1 | ENSMUSG00000029862 | 12723 | -0.651260744490467 | 5.31543379161637 | -5.63893852240556 | 7.1969829046784 | 0.000440976570241253 |
| Apool | ENSMUSG00000025525 | 68117 | -0.570146910384948 | 3.73743353638243 | -5.63772639391531 | 7.2119699397713 | 0.000441647302025996 |
| Mt1 | ENSMUSG00000031765 | 17748 | 1.04311736791566 | 4.53785279378465 | 5.63707560355942 | 7.2200299407132 | 0.000441893321847348 |
| Ogdhl | ENSMUSG00000021913 | 239017 | -1.0919659158483 | 0.889940654098485 | -5.63517505712028 | 7.2436221369974 | 0.000443089163847335 |
| Taco1 | ENSMUSG00000001983 | 70207 | -0.504155194524254 | 4.09846434028576 | -5.63366941736216 | 7.26236950709e- | 0.000443969848716362 |
| Tmem38a | ENSMUSG00000031791 | 74166 | -0.486104015385962 | 8.6188105819112 | -5.6333668758989 | 7.2661427074702 | 0.000443969848716362 |
| Tmem242 | ENSMUSG00000004945 | 70544 | -0.483141347154965 | 4.34177941508101 | -5.63279652813736 | 7.2732614857917 | 0.000444156543582065 |
| Htra3 | ENSMUSG00000029096 | 78558 | 0.6405383944384 | 4.92384721864665 | 5.63235650176909 | 7.2787586517950 | 0.000444244058195264 |
| Sytl2 | ENSMUSG00000030616 | 83671 | 0.909701739491673 | 1.70510254755785 | 5.63180536458925 | 7.2856500415172 | 0.000444416521226702 |
| Man1a2 | ENSMUSG00000008763 | 17156 | 0.384252824936176 | 4.20460295740195 | 5.62925817913928 | 7.3175887756702 | 0.00044611579981512 |
| Dhrs11 | ENSMUSG00000034449 | 192970 | -0.875860934862094 | 3.84168447961965 | -5.62734386164394 | 7.3416885536605 | 0.000447335549498683 |
| Fbxo44 | ENSMUSG00000029001 | 230903 | -1.00903388826344 | 1.5634013602178 | -5.62329499840761 | 7.3929347779736 | 0.000450207075532197 |
| Canx | ENSMUSG00000020368 | 12330 | 0.382565401573269 | 7.32641116135188 | 5.62002444007568 | 7.4346034441380 | 0.000452492484676355 |
| Dnaja2 | ENSMUSG00000031701 | 56445 | -0.371024055688971 | 6.73032838780014 | -5.61398553911083 | 7.5121897259645 | 0.00045696018861724 |
| Pdlim3 | ENSMUSG00000031636 | 53318 | 0.586709489547985 | 7.00834743046166 | 5.61119091679323 | 7.5483807872390 | 0.00045890628690384 |
| Hist1h1b | ENSMUSG00000058773 | 56702 | 1.89248925255434 | 0.377999080860777 | 5.60987112620573 | 7.5655358178391 | 0.000459625552238823 |
| Top2a | ENSMUSG00000020914 | 21973 | 1.84776969611189 | 1.26251253059941 | 5.6096340829677 | 7.5686212975014 | 0.000459625552238823 |
| Pros1 | ENSMUSG00000022912 | 19128 | 0.5832275396644 | 3.41710688426735 | 5.60719466693165 | 7.6004506624556 | 0.000461302199840656 |
| Mamstr | ENSMUSG00000042918 | 74490 | 1.07076182732702 | 2.49924979239157 | 5.60182375898448 | 7.6710248664221 | 0.000465327263123533 |
| Psme1 | ENSMUSG00000022216 | 19186 | -0.599860922369093 | 4.86548331536038 | -5.60046293740223 | 7.6890148773226 | 0.000466159853710562 |
| D430041D05Rik | ENSMUSG00000068373 | 241589 | -0.98542982285375 | 1.41659362784531 | -5.5997753534682 | 7.6981215008623 | 0.000466453249035066 |
| Osbpl5 | ENSMUSG00000037606 | 79196 | 0.662378803398396 | 3.51914348330608 | 5.59728309409638 | 7.7312246713974 | 0.000468199539518258 |
| Cd47 | ENSMUSG00000055447 | 16423 | 0.495855516679545 | 4.8592625155249 | 5.59588588587931 | 7.7498480918312 | 0.000469067494417541 |
| Usp11 | ENSMUSG00000031066 | 236733 | 0.658377493296725 | 2.18166075031414 | 5.59397283659333 | 7.7754233652041 | 0.000470355023824273 |
| Bap1 | ENSMUSG00000021901 | 104416 | -0.396011786440268 | 5.13909139372139 | -5.59322306440133 | 7.7854710421020 | 0.000470702344918239 |
| Nt5c3 | ENSMUSG00000029780 | 107569 | -0.36482773904988 | 4.34810736960988 | -5.59284047954687 | 7.7906032871685 | 0.000470752263858702 |
| Cthrc1 | ENSMUSG00000054196 | 68588 | 2.79821474527775 | 0.879795147870974 | 5.59229600814602 | 7.7979132916045 | 0.000470933647461486 |
| Bcas2 | ENSMUSG00000005687 | 68183 | -0.409943785906012 | 4.23679884939454 | -5.59177334965849 | 7.8049371924816 | 0.000471097561849901 |
| Coro7 | ENSMUSG00000039637 | 78885 | 0.571312445596905 | 2.81796476751934 | 5.59083385229228 | 7.8175795613284 | 0.000471600232808393 |
| Plekha1 | ENSMUSG00000040268 | 101476 | 0.378704473586763 | 4.95443304596035 | 5.5887727842182 | 7.8453895206581 | 0.000472950096702937 |
| 1110008P14Rik | ENSMUSG00000039195 | 73737 | 0.609556464824029 | 2.35577382207166 | 5.5885346569473 | 7.8486092344627 | 0.000472950096702937 |
| Cd84 | ENSMUSG00000038147 | 12523 | 1.42809705254059 | 1.70834737603803 | 5.58785330832845 | 7.8578293769110 | 0.000473244809471153 |
| Gbp9 | ENSMUSG00000029298 | 236573 | -0.770063856319199 | 3.50744937335186 | -5.5853995936553 | 7.8911275581968 | 0.00047498852058728 |
| Ccdc134 | ENSMUSG00000068114 | 76457 | 0.78896357591887 | 1.88346901723902 | 5.58442042200323 | 7.9044566217695 | 0.000475528978164904 |
| Parm1 | ENSMUSG00000034981 | 231440 | -0.801384623109963 | 1.46948519473622 | -5.58361730724279 | 7.9154066839175 | 0.00047592580012048 |
| Eno3 | ENSMUSG00000060600 | 13808 | -0.646062400849038 | 8.95927276455303 | -5.58154840508238 | 7.9436883064248 | 0.00047736369916179 |
| Hexb | ENSMUSG00000021665 | 15212 | 0.741579958529568 | 2.8773476272886 | 5.58013700318471 | 7.9630426549441 | 0.000478064019493077 |
| C2cd5 | ENSMUSG00000030279 | 74741 | 0.614016882528538 | 3.50907054926071 | 5.5800607942867 | 7.9640890997794 | 0.000478064019493077 |
| Ccdc88b | ENSMUSG00000047810 | 78317 | 1.28960351668982 | 0.998253152676074 | 5.57818786550052 | 7.9898520711273 | 0.000479152722524959 |
| Epsti1 | ENSMUSG00000022014 | 108670 | 1.45555976710974 | 0.565931692506973 | 5.57810508894129 | 7.9909927103009 | 0.000479152722524959 |
| Junb | ENSMUSG00000052837 | 16477 | 1.19765807862267 | 3.26896470211942 | 5.57481299471588 | 8.0364953478857 | 0.000481490335363349 |
| Sertad4 | ENSMUSG00000016262 | 214791 | 1.05098239490325 | 2.15840249494754 | 5.57464767477378 | 8.0387875037792 | 0.000481490335363349 |
| Lhfpl2 | ENSMUSG00000045312 | 218454 | 1.0629491564947 | 2.43808419275229 | 5.57075564233906 | 8.0929485343811 | 0.000484468896107997 |
| Cad | ENSMUSG00000013629 | 69719 | 0.502768806596376 | 3.48683725933476 | 5.56966504445276 | 8.1081935529496 | 0.00048511583868246 |
| Gyg | ENSMUSG00000019528 | 27357 | -0.426055920657866 | 7.18752285527296 | -5.56888484515875 | 8.1191180752076 | 0.000485503718162406 |
| Nrros | ENSMUSG00000052384 | 224109 | 0.942479381307488 | 1.45670135792535 | 5.56853388201706 | 8.1240373602448 | 0.000485532271103533 |
| Lrp10 | ENSMUSG00000022175 | 65107 | 0.338132945934339 | 4.70915442146379 | 5.56795416366073 | 8.1321698315971 | 0.000485740073684635 |
| Pnpt1 | ENSMUSG00000020464 | 71701 | -0.380811940017155 | 3.76890443312345 | -5.56765273407664 | 8.1364017465608 | 0.000485740073684635 |
| Ldlrad3 | ENSMUSG00000048058 | 241576 | 0.598279106677938 | 5.22515993390812 | 5.56550688446992 | 8.1665950399651 | 0.000487276475883512 |
| Iqsec1 | ENSMUSG00000034312 | 232227 | -0.372335149831543 | 5.93275791561463 | -5.56419516256082 | 8.1851094156643 | 0.000488114735529882 |
| Cops3 | ENSMUSG00000019373 | 26572 | -0.390177189954241 | 5.34807243118006 | -5.56363595188355 | 8.1930157765147 | 0.000488319822535891 |
| Pdgfc | ENSMUSG00000028019 | 54635 | 1.03804229019261 | 1.24867464526019 | 5.56170278051092 | 8.2204094058594 | 0.000489663114540581 |
| Opa1 | ENSMUSG00000038084 | 74143 | -0.426588763332322 | 6.6625704883432 | -5.56141379439244 | 8.2245126548029 | 0.000489663114540581 |
| Asah1 | ENSMUSG00000031591 | 11886 | 0.464097465742781 | 4.07797967446946 | 5.56026187924897 | 8.2408897487188 | 0.000490371071547338 |
| Heatr1 | ENSMUSG00000050244 | 217995 | 0.394253359454049 | 3.80962189797679 | 5.55806849119597 | 8.2721681685508 | 0.000491964473614957 |
| Trmt2b | ENSMUSG00000067369 | 215201 | -0.563853417437662 | 3.09990658600416 | -5.55587551830708 | 8.3035648775274 | 0.000493563173878482 |
| Mavs | ENSMUSG00000037523 | 228607 | -0.526508804105338 | 6.26062345575625 | -5.55251142475937 | 8.3519710472885 | 0.000496042715443549 |
| Emilin2 | ENSMUSG00000024053 | 246707 | 0.808672802774752 | 3.44782420355295 | 5.5523462145763 | 8.3543558606858 | 0.000496042715443549 |
| Phb2 | ENSMUSG00000004264 | 12034 | -0.410427587359269 | 5.98934460409784 | -5.55120176815929 | 8.3708955503781 | 0.000496754936271353 |
| Syngr2 | ENSMUSG00000048277 | 20973 | -0.318733669990922 | 5.55891886354738 | -5.55042510281215 | 8.3821395320263 | 0.000496900297632089 |
| Myoz1 | ENSMUSG00000068697 | 59011 | -0.68874744586404 | 6.70137721752533 | -5.55029477506288 | 8.3840278677937 | 0.000496900297632089 |
| Evi2b | ENSMUSG00000093938 | 216984 | 1.9648413000647 | 0.446095828222163 | 5.55009092108544 | 8.3869824273278 | 0.000496900297632089 |
| Ms4a6c | ENSMUSG00000079419 | 73656 | 1.66377585971808 | 0.990263258615512 | 5.54692172537911 | 8.4330555750833 | 0.00049925818384885 |
| Thop1 | ENSMUSG00000004929 | 50492 | -0.385365600282283 | 4.05135883788197 | -5.54672563583239 | 8.4359149718125 | 0.00049925818384885 |
| Irf8 | ENSMUSG00000041515 | 15900 | 1.16003265856025 | 1.76058012441028 | 5.54601422433002 | 8.4462973619866 | 0.000499602145367257 |
| Tmem167 | ENSMUSG00000012422 | 66074 | 0.502062623150338 | 3.1169324708791 | 5.54533523724683 | 8.4562190172934 | 0.000499918496906623 |
| Jak2 | ENSMUSG00000024789 | 16452 | -0.380346085031441 | 4.83901449123005 | -5.54312955136191 | 8.4885336907066 | 0.000501557631206026 |
| Trmt10c | ENSMUSG00000044763 | 52575 | -0.439439796808432 | 3.46541842506114 | -5.54228326678168 | 8.5009665592109 | 0.000501850762922749 |
| Atp11a | ENSMUSG00000031441 | 50770 | 0.364315761198488 | 5.24529078523994 | 5.54207050998145 | 8.5040951908194 | 0.000501850762922749 |
| Calr | ENSMUSG00000003814 | 12317 | 0.408830218811721 | 7.06748279752048 | 5.54185483344445 | 8.5072679873374 | 0.000501850762922749 |
| Gss | ENSMUSG00000027610 | 14854 | 0.689537572633157 | 2.31188921542838 | 5.54139523324764 | 8.5140332532189 | 0.000501978950868048 |
| Klf15 | ENSMUSG00000030087 | 66277 | -0.799565590586049 | 4.65552503320805 | -5.54011978651453 | 8.5328371930574 | 0.000502816406238872 |

| | | | | | | | |
|---|---|---|---|---|---|---|---|
| Tst | ENSMUSG00000044986 | 22117 | -1.21717891892751 | 1.34530968595471 | -5.53951038756813 | 8.5418369096962 | 0.00050307553480544 |
| Ppib | ENSMUSG00000032383 | 19035 | 0.576523278108486 | 4.31688855568987 | 5.53796279788085 | 8.5647366617154 | 0.000504152592618262 |
| Slc29a3 | ENSMUSG00000020100 | 71279 | 0.833203305580264 | 2.04451913304333 | 5.53656843016592 | 8.5854241863999 | 0.000505098341127763 |
| Gps2 | ENSMUSG00000023170 | 56310 | -0.36448084657236 | 5.00086712322677 | -5.5362262149004 | 8.5905094452879 | 0.000505125652213245 |
| Phf20l1 | ENSMUSG00000072501 | 239510 | -0.367899570865406 | 5.92848274557542 | -5.53514863332037 | 8.6065427119775 | 0.00050579633540122 |
| Mecom | ENSMUSG00000027684 | 14013 | -0.71673806615543 | 3.00677350677732 | -5.53131123218056 | 8.6638938615691 | 0.000508893196135477 |
| Nfkbie | ENSMUSG00000023947 | 18037 | 1.35061383588867 | 1.01580295745789 | 5.53043721607143 | 8.6770121152521 | 0.000509390007689692 |
| Eif2ak3 | ENSMUSG00000031668 | 13666 | 0.702273296440479 | 2.8715139300544 | 5.52952936163758 | 8.6906602568380 | 0.000509917376637128 |
| Hsd17b10 | ENSMUSG00000025260 | 15108 | -0.52059359417536 | 4.29045998258416 | -5.52911312936734 | 8.6969251508009 | 0.000510011206134147 |
| Polr2a | ENSMUSG00000005198 | 20020 | 0.414804499024176 | 6.04225912285937 | 5.52619463853994 | 8.7409855433035 | 0.000512060113222508 |
| Adam22 | ENSMUSG00000040537 | 11496 | -0.786797462838057 | 1.82676409581168 | -5.52617829569922 | 8.7412329272088 | 0.000512060113222508 |
| Fxyd5 | ENSMUSG00000009687 | 18301 | 0.797715902015971 | 2.80977932393003 | 5.52414738016342 | 8.7720322629755 | 0.000513589098374857 |
| Rac2 | ENSMUSG00000033220 | 19354 | 1.24507673008983 | 0.928025487937648 | 5.52293815493219 | 8.7904243593920 | 0.000514243600721563 |
| Tmem63b | ENSMUSG00000036026 | 224807 | -0.461157888994046 | 5.68617297584223 | -5.52279397647148 | 8.7926199775738 | 0.000514243600721563 |
| Atg10 | ENSMUSG00000021619 | 66795 | -0.709467930870161 | 2.28338962456995 | -5.52193941756065 | 8.8056453889456 | 0.000514729998644735 |
| Eif4ebp2 | ENSMUSG00000020091 | 13688 | -0.37480419894699 | 6.39008254914103 | -5.51968203358979 | 8.8401502006491 | 0.000516470774149097 |
| Coq7 | ENSMUSG00000030652 | 12850 | -0.572644227183314 | 5.40300631406409 | -5.51789032132898 | 8.8676377390191 | 0.000517799936566337 |
| Pcnt | ENSMUSG00000001151 | 18541 | -0.4755580424307 | 7.42846504564203 | -5.51727758537407 | 8.8770585246844 | 0.000518073287417649 |
| Pm20d2 | ENSMUSG00000054659 | 242377 | -1.00897408503684 | 4.01488052849781 | -5.51623881140249 | 8.8930535493160 | 0.000518729820424621 |
| Thra | ENSMUSG00000058756 | 21833 | -0.42392963770151 | 5.7923919598459 | -5.51434017672608 | 8.9223666465737 | 0.000520162079006385 |
| Dop1b | ENSMUSG00000022946 | 70028 | 0.503812308792058 | 3.57254389001159 | 5.51227111160678 | 8.9544261320807 | 0.000521752836086221 |
| Iqsec3 | ENSMUSG00000040797 | 243621 | 1.55546289294099 | 1.07695448163655 | 5.50577764897058 | 9.0558249305628 | 0.000527379980372839 |
| Yap1 | ENSMUSG00000053110 | 22601 | 0.374872444876857 | 5.52450225194327 | 5.50494194281915 | 9.0689618750978 | 0.000527863803283782 |
| Mapk9 | ENSMUSG00000020366 | 26420 | -0.368571916592951 | 4.34337405012704 | -5.50459873985327 | 9.0743626514174 | 0.000527897063026311 |
| Bfar | ENSMUSG00000022684 | 67118 | -0.382929642794359 | 4.91470500157346 | -5.50377199526021 | 9.0873864440429 | 0.000528373517126772 |
| Psd3 | ENSMUSG00000030465 | 234353 | 0.434784122784426 | 4.51913554030105 | 5.50237705252953 | 9.1094054586351 | 0.000529372201320259 |
| Rrm2 | ENSMUSG00000020649 | 20135 | 0.926956533365067 | 1.09964134748302 | 5.49930320671928 | 9.1581229471882 | 0.000531920520381054 |
| Pde8a | ENSMUSG00000025584 | 18584 | -0.51175658454551 | 5.07273466010906 | -5.49816786927773 | 9.1761857888728 | 0.000532363996882207 |
| Eef1a2 | ENSMUSG00000016349 | 13628 | -0.399583512703856 | 9.77158197387972 | -5.49813588027301 | 9.1766952626567 | 0.000532363996882207 |
| Rab7b | ENSMUSG00000052688 | 226421 | 0.910047855430674 | 1.95788312146152 | 5.49790526816018 | 9.1803689884087 | 0.000532363996882207 |
| Zfp777 | ENSMUSG00000071477 | 72306 | -0.416993970725391 | 3.77868689827327 | -5.4971629144508 | 9.1922053718057 | 0.000532767746125176 |
| Spats2 | ENSMUSG00000051934 | 72572 | 0.637457212457575 | 2.32486130854516 | 5.49115836891094 | 9.2885328342937 | 0.000538065460581159 |
| Cnot7 | ENSMUSG00000031601 | 18983 | -0.387461033170469 | 4.44712952345272 | -5.49046068344917 | 9.2997936563952 | 0.000538295805416945 |
| Itgav | ENSMUSG00000027087 | 16410 | 0.398390412394402 | 6.04647861886491 | 5.4903019149217 | 9.3023582145513 | 0.000538295805416945 |
| Srpk3 | ENSMUSG00000002007 | 56504 | 0.398279591588422 | 4.79089884509971 | 5.48810319840551 | 9.3379498045523 | 0.000539551735133199 |
| Card14 | ENSMUSG00000013483 | 170720 | 2.5036949194429 | 0.77550654581403 | 5.4879151771349 | 9.3409998947030 | 0.000539551735133199 |
| Pofut1 | ENSMUSG00000046020 | 140484 | 0.543932390795142 | 3.39384812274937 | 5.48788900511619 | 9.3414246490132 | 0.000539551735133199 |
| Ndufb7 | ENSMUSG00000033938 | 66916 | -0.367830648780339 | 6.2591275581991 | -5.48774226367193 | 9.3438060068351 | 0.000539551735133199 |
| Megf10 | ENSMUSG00000024593 | 70417 | 1.37152911496361 | 1.62560082372457 | 5.48659482910669 | 9.3624487725242 | 0.000540342806401595 |
| Gid4 | ENSMUSG00000018415 | 66771 | -0.372615346751047 | 4.87285333082733 | -5.48486384833124 | 9.3906462451395 | 0.00054168419053098 |
| Sfrp1 | ENSMUSG00000031548 | 20377 | 1.66463447094754 | 1.22768358764562 | 5.48341706521026 | 9.4142822979915 | 0.000542761180376294 |
| Uckl1os | ENSMUSG00000010492 | 100271841 | -0.82748805569017 | 3.82570928776596 | -5.48254828555139 | 9.4285054116972 | 0.000543214916501493 |
| Wdr1 | ENSMUSG00000005103 | 22388 | 0.365596315987955 | 6.72800090127437 | 5.48232946128184 | 9.4320914053593 | 0.000543214916501493 |
| Scd2 | ENSMUSG00000025203 | 20250 | 0.616802137206222 | 4.51750518822621 | 5.47980439728595 | 9.4735743520646 | 0.000545316699538765 |
| Sla | ENSMUSG00000022372 | 20491 | 1.55250312795792 | 0.529108927819187 | 5.47767394533423 | 9.5087227955992 | 0.000547051836203659 |
| Mrpl46 | ENSMUSG00000030612 | 67308 | -0.466177892891135 | 3.93052980895525 | -5.4742573788495 | 9.5653746520647 | 0.000550021621892267 |
| Psmd8 | ENSMUSG00000030591 | 57296 | 0.418547055590537 | 6.33756916425024 | 5.47232043328897 | 9.5976487927662 | 0.000551587271050091 |
| Bloc1s6 | ENSMUSG00000005804 | 18457 | 0.627632072763555 | 3.71293598916693 | 5.47025643637345 | 9.6321652430065 | 0.000553280074993718 |
| Zdbf2 | ENSMUSG00000027520 | 73884 | 1.46899004990893 | 2.23687619156165 | 5.46960926460318 | 9.6430146759750 | 0.000553612360415354 |
| Oxa1l | ENSMUSG00000000959 | 69089 | -0.376963500016902 | 4.78471519328095 | -5.46875058307715 | 9.6574296650050 | 0.000554148890646565 |
| Naa38 | ENSMUSG00000059278 | 78304 | -0.553397744637649 | 3.16907644880572 | -5.46744072889822 | 9.6794620582350 | 0.000555121719614728 |
| Lrig2 | ENSMUSG00000032913 | 269473 | 0.481150856964658 | 3.34514910354133 | 5.46630520373091 | 9.6986046768097 | 0.000555927885276388 |
| Tmem126a | ENSMUSG00000030615 | 66271 | -0.519758382166356 | 3.73723085189271 | -5.46233473102969 | 9.7658504119065 | 0.000559489050589889 |
| Hccs | ENSMUSG00000031352 | 15159 | -0.476802958067263 | 4.4319042003113 | -5.46076700814167 | 9.7925361482013 | 0.000560724005426865 |
| Steap2 | ENSMUSG00000015653 | 74051 | 1.35918189480121 | 0.550756692522897 | 5.45425697747602 | 9.9041679037355 | 0.000566618228162191 |
| Msln | ENSMUSG00000063011 | 56047 | 1.55450846127587 | 0.729437430776724 | 5.45416001614534 | 9.9058405819956 | 0.000566618228162191 |
| B3gnt2 | ENSMUSG00000051650 | 53625 | 0.706665901721442 | 2.47051039689156 | 5.45316004416979 | 9.9231083185736 | 0.000567309084886657 |
| Serinc2 | ENSMUSG00000023232 | 230779 | 0.553678085982798 | 3.75519898087941 | 5.45089083623923 | 9.9624103251332 | 0.000569119475589637 |
| Tarsl2 | ENSMUSG00000030515 | 272396 | -0.501798665261006 | 5.58330056381739 | -5.45073082650914 | 9.9651877804278 | 0.000569119475589637 |
| Tyrobp | ENSMUSG00000030579 | 22177 | 1.82028216890166 | 0.897638956835467 | 5.44911045635907 | 9.9933598750584 | 0.000570430374904771 |
| Gpam | ENSMUSG00000024978 | 14732 | -0.649516177661837 | 7.08238912235866 | -5.44596905167804 | 0.0001004821440 | 0.000573262169559362 |
| Zim1 | ENSMUSG00000002266 | 22776 | 0.909524269901573 | 1.75281857219108 | 5.44551244886508 | 0.0001005621366 | 0.000573419257177081 |
| Baiap2 | ENSMUSG00000025372 | 108100 | 0.922441148502701 | 1.63302983167097 | 5.44455549745215 | 0.0001007300017 | 0.000574076980811485 |
| Slc38a1 | ENSMUSG00000023169 | 105727 | 0.805914766498055 | 3.04312582563382 | 5.43977428999904 | 0.0001015731028 | 0.000578580295790164 |
| Acsl3 | ENSMUSG00000032883 | 74205 | 0.556480902971705 | 4.05479579451049 | 5.4394539155102 | 0.0001016298596 | 0.000578602080947576 |
| Rtl8b | ENSMUSG00000067924 | 553127 | 0.420001354137613 | 4.10356422712122 | 5.43909593614675 | 0.0001016933175 | 0.000578661975023237 |
| Ss18l2 | ENSMUSG00000032526 | 26901 | -0.585222597883216 | 3.88030411077316 | -5.43760730282032 | 0.0001019576476 | 0.000579864227966595 |
| Tnks2 | ENSMUSG00000024811 | 74493 | -0.304087815943221 | 7.10858195441465 | -5.43692075960033 | 0.0001020797959 | 0.000580257020080979 |
| Clmp | ENSMUSG00000032024 | 71566 | 0.814430357025147 | 3.26915181908452 | 5.43644556550374 | 0.0001021644310 | 0.00058043627648238 |
| Mrpl3 | ENSMUSG00000032563 | 94062 | -0.45014102944282 | 4.8940865119829 | -5.43489184707419 | 0.0001024416705 | 0.000581709039356128 |
| Rhbdf2 | ENSMUSG00000020806 | 217344 | -0.504298322969814 | 3.30335496804004 | -5.43335168408094 | 0.0001027172673 | 0.000582971157788555 |
| Synj2bp | ENSMUSG00000090935 | 24071 | -0.423721091803769 | 5.27321921405136 | -5.43266932453778 | 0.0001028396164 | 0.000583138157531068 |
| Vars | ENSMUSG00000007029 | 22321 | 0.3484276874837 | 4.78115990459021 | 5.43259258006626 | 0.0001028533864 | 0.000583138157531068 |
| Gm5860 | ENSMUSG00000085022 | 545641 | -0.924723433840627 | 1.46688992835676 | -5.43176956441231 | 0.0001030011789 | 0.000583204182817612 |
| Ptges2 | ENSMUSG00000026820 | 96979 | -0.518289338601929 | 4.08899551209656 | -5.43176103502346 | 0.0001030027117 | 0.000583204182817612 |
| Sf3b3 | ENSMUSG00000033732 | 101943 | 0.344738162310581 | 4.4657240231455 | 5.43163651708565 | 0.0001030250916 | 0.000583204182817612 |
| Lnpep | ENSMUSG00000023845 | 240028 | 0.410197461424123 | 6.86995328842092 | 5.42746346888871 | 0.0001037780698 | 0.000587162568065106 |
| Gtf2e1 | ENSMUSG00000022828 | 74197 | -0.584584932278151 | 2.85345673927734 | -5.42644187061525 | 0.0001039632803 | 0.000587906165037976 |
| Mtus2 | ENSMUSG00000029651 | 77521 | 3.60635887012276 | -0.0270251354260735 | 5.42406689453976 | 0.0001043951873 | 0.000590043326441668 |
| Eif3h | ENSMUSG00000022312 | 68135 | -0.369174783185444 | 5.85996070720698 | -5.42152611710803 | 0.0001048593233 | 0.000592360343147213 |
| Lrpprc | ENSMUSG00000024120 | 72416 | -0.505579391238636 | 6.4335278318451 | -5.42084366780732 | 0.0001049843567 | 0.000592681098662996 |
| Cdk6 | ENSMUSG00000040274 | 12571 | 0.901850639196017 | 2.40100708081675 | 5.42052631966092 | 0.0001050425519 | 0.000592681098662996 |
| Srp54b | ENSMUSG00000112449 | 665155 | 1.71297816065919 | 3.94459808833977 | 5.42032894232118 | 0.0001050787639 | 0.000592681098662996 |
| Atad3a | ENSMUSG00000029036 | 108888 | -0.482551299335792 | 4.19431761595491 | -5.41958906867481 | 0.0001052146217 | 0.000593141325757828 |
| Rwdd4a | ENSMUSG00000031568 | 192174 | -0.452846295100291 | 4.27501343132888 | -5.41876604417554 | 0.0001053659636 | 0.000593688324061245 |
| Larp4 | ENSMUSG00000023025 | 207214 | -0.418402231977928 | 5.60977690155963 | -5.41487251630349 | 0.0001060850149 | 0.000597431890158082 |
| 2900026A02Rik | ENSMUSG00000051339 | 243219 | 0.439070571785912 | 5.66037895006291 | 5.41386393834618 | 0.0001062721126 | 0.000598177375690074 |
| Dpf2 | ENSMUSG00000024826 | 19708 | -0.384312837139396 | 4.51174778966546 | -5.41233363635551 | 0.0001065566522 | 0.000599470286147929 |
| Tsga10 | ENSMUSG00000060771 | 211484 | -0.717684734041891 | 3.67696328872275 | -5.41190218694447 | 0.0001066370184 | 0.000599613810855819 |
| Dusp23 | ENSMUSG00000026544 | 68440 | -0.734247641505344 | 2.31950581269563 | -5.41150519114429 | 0.0001067110228 | 0.000599721434750185 |
| Itpkc | ENSMUSG00000003752 | 233011 | 0.686223912772053 | 2.53005666169305 | 5.40811873196595 | 0.0001073444819 | 0.000602932562846611 |
| Lpp | ENSMUSG00000033306 | 210126 | 0.585529671051845 | 5.4272125769167 | 5.40786176795532 | 0.0001073927087 | 0.000602932562846611 |
| Zfp955b | ENSMUSG00000096910 | 100043468 | -0.504918314119632 | 3.52217848669486 | -5.40483978655572 | 0.0001079615746 | 0.000605815180745548 |
| Adamts4 | ENSMUSG00000006403 | 240913 | 1.87514856250534 | 0.668483390283088 | 5.402990232251 | 0.0001083112920 | 0.000607462523294191 |
| Ptprf | ENSMUSG00000033295 | 19268 | 0.892786855733721 | 2.3359200350076 | 5.4026583266349 | 0.0001083741745 | 0.000607462523294191 |
| Itih5 | ENSMUSG00000025780 | 209378 | 0.607640906219967 | 5.55317871221113 | 5.4024067531699 | 0.0001084218628 | 0.000607462523294191 |
| Mrps35 | ENSMUSG00000040112 | 232536 | -0.461113140445553 | 4.0002672911478 | -5.40136838107522 | 0.0001086189289 | 0.00060825487294325 |
| Map4k3 | ENSMUSG00000024242 | 225028 | -0.415552953952505 | 4.09730838749663 | -5.39958849433169 | 0.0001089575935 | 0.000609838942542738 |
| Gkap1 | ENSMUSG00000021552 | 56278 | -0.438034497083209 | 3.88895300960035 | -5.39743912949945 | 0.0001093680305 | 0.000611822897787294 |
| Mylk4 | ENSMUSG00000044951 | 238564 | -1.60408374395808 | 5.71027423759487 | -5.39431075545366 | 0.0001099683057 | 0.000614866266022038 |
| Nsd2 | ENSMUSG00000057406 | 107823 | 0.431410776582753 | 4.09379008402476 | 5.39147057796314 | 0.0001105162627 | 0.000617614145236301 |
| Emp1 | ENSMUSG00000030208 | 13730 | 1.17019920499739 | 5.86407430390485 | 5.39082905513392 | 0.0001106404265 | 0.000617992081092759 |
| Mrps21 | ENSMUSG00000054312 | 66292 | -0.374630657386997 | 4.70062482107326 | -5.38912394128155 | 0.0001109711525 | 0.000619522813109093 |
| Sys1 | ENSMUSG00000045503 | 66460 | -0.650471827922095 | 2.76725409708821 | -5.38676611931414 | 0.0001114301809 | 0.000621522680938533 |
| Atp1b1 | ENSMUSG00000026576 | 11931 | -0.379223059372784 | 8.27003101041152 | -5.38669994028898 | 0.0001114430934 | 0.000621522680938533 |
| Uggt1 | ENSMUSG00000037470 | 320011 | 0.345325121459775 | 6.26208255985865 | 5.38530505333864 | 0.0001117156210 | 0.000622724861535821 |
| Pcgf5 | ENSMUSG00000024805 | 76073 | -0.393123852023063 | 4.24379416317518 | -5.3843230061592 | 0.0001119079071 | 0.000623478762928802 |
| Morf4l2 | ENSMUSG00000031422 | 56397 | 0.321900190188919 | 5.51173996223899 | 5.3838948832905 | 0.0001119918423 | 0.000623628542286572 |
| Ifrd2 | ENSMUSG00000010048 | 15983 | -0.450885473625335 | 3.44143555307442 | -5.38321146325111 | 0.0001121259658 | 0.000624057501471447 |
| Litaf | ENSMUSG00000022500 | 56722 | 0.813266486621863 | 3.5084116415356 | 5.38141362867538 | 0.0001124795990 | 0.000625707123108656 |
| Cav1 | ENSMUSG00000007655 | 12389 | -0.717633072215301 | 5.94987573892479 | -5.38044566897488 | 0.0001126704744 | 0.000626450130214415 |
| Phospho1 | ENSMUSG00000050860 | 237928 | -0.327967919780453 | 5.9347445548746 | -5.37737693723497 | 0.0001132778656 | 0.000629057054447578 |
| Socs3 | ENSMUSG00000053113 | 12702 | 1.95337228541401 | 2.19041755113118 | 5.37717743380059 | 0.0001133174711 | 0.000629057054447578 |
| Mapkapk3 | ENSMUSG00000032577 | 102626 | -0.639944574107858 | 3.53783384442113 | -5.37702961999752 | 0.0001133468244 | 0.000629057054447578 |
| Apaf1 | ENSMUSG00000019979 | 11783 | 0.594297473822225 | 2.01266256857678 | 5.3769152801814 | 0.0001133695359 | 0.000629057054447578 |
| Tnxb | ENSMUSG00000033327 | 81877 | 0.598972830121357 | 7.37232018288655 | 5.37426250929837 | 0.0001138977952 | 0.000631667580076122 |

| | | | | | | | |
|---|---|---|---|---|---|---|---|
| Aebp1 | ENSMUSG00000020473 | 11568 | 0.842798014257861 | 5.81258958487627 | 5.37285403341971 | 0.0001141793164 | 0.000632907762517273 |
| Mad2l1bp | ENSMUSG00000034509 | 66591 | -1.01210586107341 | 2.61369080768501 | -5.37134638068838 | 0.0001144814661 | 0.000634260976562407 |
| Car3 | ENSMUSG00000027559 | 12350 | 0.579240753435009 | 10.3020241185831 | 5.37054027093267 | 0.0001146433617 | 0.000634836163761689 |
| Etfrf1 | ENSMUSG00000040370 | 67636 | -0.477931104162535 | 4.05826664571269 | -5.36719956899985 | 0.0001153168470 | 0.000638242255975453 |
| Akap13 | ENSMUSG00000066406 | 75547 | 0.375195541304797 | 7.54230065519804 | 5.36420343378977 | 0.0001159243830 | 0.000641280076246177 |
| Pick1 | ENSMUSG00000068206 | 18693 | -0.677549930550505 | 3.85474297991725 | -5.36070777438719 | 0.0001166374368 | 0.000644898240871703 |
| Cavin2 | ENSMUSG00000045954 | 20324 | -0.793830349093385 | 5.79759268202636 | -5.35849955005851 | 0.0001170902358 | 0.000646696697625923 |
| Midn | ENSMUSG00000035621 | 59090 | 0.440426758496295 | 5.60147631758998 | 5.35834236723013 | 0.0001171225362 | 0.000646696697625923 |
| Mrpl47 | ENSMUSG00000037531 | 74600 | -0.580957535199235 | 3.43791088098969 | -5.35825645860713 | 0.0001171401940 | 0.000646696697625923 |
| Timm8b | ENSMUSG00000039016 | 30057 | -0.582294149709029 | 4.65488207431191 | -5.35533524484461 | 0.0001177422829 | 0.00064969252652483 |
| Sh2d4a | ENSMUSG00000053886 | 72281 | 0.741558335933747 | 2.56016806340621 | 5.35447605301646 | 0.0001179199846 | 0.000650333611457845 |
| Anapc13 | ENSMUSG00000035048 | 69010 | -0.504210406233205 | 3.94622750048814 | -5.35419847606814 | 0.0001179774541 | 0.000650333611457845 |
| Krba1 | ENSMUSG00000042810 | 77827 | -0.484286994335707 | 4.48577349935765 | -5.35265998786558 | 0.0001182965133 | 0.000651763703151723 |
| Jmjd6 | ENSMUSG00000056962 | 107817 | -0.431014702796861 | 3.74515440717507 | -5.34905073008194 | 0.0001190485604 | 0.000655576732561706 |
| Pkd2 | ENSMUSG00000034462 | 18764 | 0.807073813062964 | 5.52979124450673 | 5.34779971066052 | 0.0001193103942 | 0.000656687774454052 |
| Afg3l2 | ENSMUSG00000024527 | 69597 | -0.303347628868424 | 5.90226271594044 | -5.346693034165 | 0.0001195425189 | 0.000657634260271094 |
| Lias | ENSMUSG00000029199 | 79464 | -0.478327585822858 | 4.05059139173077 | -5.34612440060724 | 0.0001196619728 | 0.000657960274499214 |
| Msrb2 | ENSMUSG00000023094 | 76467 | -0.907725018821475 | 1.94596012912948 | -5.34535141691402 | 0.0001198245549 | 0.000658522981324583 |
| Gga2 | ENSMUSG00000030872 | 74105 | 0.584251957963548 | 3.50104793909017 | 5.34451434933907 | 0.0001200008759 | 0.000659064872930256 |
| Cop1 | ENSMUSG00000040782 | 26374 | -0.322916614118672 | 5.33486469436148 | -5.34431103276398 | 0.0001200437436 | 0.000659064872930256 |
| Ppp1r16b | ENSMUSG00000037754 | 228852 | -0.750824920066017 | 3.21372445822532 | -5.34351122395832 | 0.0001202125324 | 0.000659660237264811 |
| Nbeal2 | ENSMUSG00000056724 | 235627 | 0.772329899917079 | 2.73641574792181 | 5.34223067993831 | 0.0001204832901 | 0.000660814272464521 |
| Miga2 | ENSMUSG00000026858 | 108958 | -0.658181140818123 | 3.60609518342572 | -5.34097753620909 | 0.0001207488709 | 0.000661668045689053 |
| Nsmce4a | ENSMUSG00000040331 | 67872 | -0.438764919337274 | 3.91687579602799 | -5.3409250040861 | 0.0001207600174 | 0.000661668045689053 |
| Cdon | ENSMUSG00000038119 | 57810 | 0.536299744927204 | 4.29752962096559 | 5.33325622750354 | 0.0001223988040 | 0.000670311285904716 |
| Mrgprf | ENSMUSG00000031070 | 211577 | 0.627557660976912 | 1.70851728685442 | 5.3274131300571 | 0.0001236630279 | 0.000676895622500155 |
| Dcaf6 | ENSMUSG00000026571 | 74106 | -0.367297473349614 | 6.947096818693 | -5.32623244744156 | 0.0001239201342 | 0.000677963457154688 |
| Tefm | ENSMUSG00000046909 | 68550 | -0.44682609022014 | 2.96577912597534 | -5.32244302641716 | 0.0001247490980 | 0.000682157273845552 |
| Spon2 | ENSMUSG00000037379 | 100689 | 2.39209312042572 | 1.28013601482244 | 5.31783886317459 | 0.0001257640892 | 0.000687363629748813 |
| Coa6 | ENSMUSG00000051671 | 67892 | -0.618470119718635 | 2.36648376420449 | -5.31442232794488 | 0.0001265228370 | 0.000691164983406592 |
| Tmem11 | ENSMUSG00000043284 | 216821 | -0.41393030003056 | 4.0859810450991 | -5.31292244954562 | 0.0001268574396 | 0.000692646689540819 |
| Sphk1 | ENSMUSG00000061878 | 20698 | 1.50453462887874 | 0.684622003611343 | 5.31218825297815 | 0.0001270215655 | 0.000693196571734207 |
| Hoxd8 | ENSMUSG00000027102 | 15437 | -0.638612742334924 | 2.57118077211607 | -5.31155919283353 | 0.0001271623651 | 0.00069331940753098 |
| Fam104a | ENSMUSG00000041629 | 28081 | 0.334157849592681 | 4.50289202011711 | 5.31152096003864 | 0.0001271709278 | 0.00069331940753098 |
| Tgm2 | ENSMUSG00000037820 | 21817 | 0.376933834671484 | 6.1623495211854 | 5.30982277697183 | 0.0001275518650 | 0.000694390808666709 |
| Cxcl12 | ENSMUSG00000061353 | 20315 | -0.823448053371781 | 6.51087580541164 | -5.30971913057655 | 0.0001275751535 | 0.000694390808666709 |
| H2-Q6 | ENSMUSG00000073409 | 110557 | -1.51719741332774 | 2.7247394934477 | -5.30954009252637 | 0.0001276153924 | 0.000694390808666709 |
| Ripor2 | ENSMUSG00000036006 | 193385 | 0.918853424421471 | 2.47526230229871 | 5.30951271216281 | 0.0001276215473 | 0.000694390808666709 |
| Fgd2 | ENSMUSG00000024013 | 26382 | 1.77720682767016 | 0.521839613786055 | 5.30908610759183 | 0.0001277174852 | 0.000694567080402603 |
| Tagln | ENSMUSG00000032085 | 21345 | -0.777970206657874 | 3.98139556305629 | -5.30840001921104 | 0.0001278719356 | 0.000695061227618619 |
| Dmxl2 | ENSMUSG00000041268 | 235380 | 0.647470963848249 | 3.02369371839086 | 5.30737026836804 | 0.0001281041164 | 0.000695977185282807 |
| Tmem64 | ENSMUSG00000043252 | 100201 | -0.453256568770661 | 6.61221802342171 | -5.30603096613035 | 0.0001284067511 | 0.000697274812261351 |
| Rhbdl1 | ENSMUSG00000025735 | 214951 | -1.33754368657393 | 0.92600437734341 | -5.30235238430108 | 0.0001292418259 | 0.000701460973085985 |
| Rpp21 | ENSMUSG00000024446 | 67676 | -0.652455755854417 | 1.99488217766729 | -5.30171004359302 | 0.0001293882241 | 0.000701907036104833 |
| Zfp148 | ENSMUSG00000022811 | 22661 | -0.308627062492786 | 4.96908538473064 | -5.30072260655624 | 0.0001296136120 | 0.000702630199072219 |
| Thoc7 | ENSMUSG00000053453 | 66231 | -0.359844269483014 | 4.18779639022414 | -5.30056297186443 | 0.0001296500879 | 0.000702630199072219 |
| Dis3l | ENSMUSG00000032396 | 213550 | -0.421700207384617 | 4.55111062478028 | -5.29925057920984 | 0.0001299503713 | 0.000703908577681661 |
| Fgfr3 | ENSMUSG00000054252 | 14184 | 0.949489968145993 | 2.58087249098986 | 5.29868778188821 | 0.0001300793651 | 0.000704258316315591 |
| Rdh14 | ENSMUSG00000020621 | 105014 | -0.480136470234509 | 4.07663146301516 | -5.29787217147492 | 0.0001302665412 | 0.000704922555400503 |
| Gsn | ENSMUSG00000026879 | 227753 | 0.44452237441841 | 9.07188457399502 | 5.2955786798173 | 0.0001307943872 | 0.000707428722201405 |
| Zbtb11 | ENSMUSG00000022601 | 271377 | -0.429432940374867 | 3.66224631623649 | -5.29442285021147 | 0.0001310612465 | 0.000708521506178401 |
| Btbd6 | ENSMUSG00000002803 | 399566 | -0.536352931007596 | 3.21805688349132 | -5.29377560557614 | 0.0001312109311 | 0.000708980073609514 |
| Pdia6 | ENSMUSG00000020571 | 71853 | 0.633567978272064 | 5.18333788637371 | 5.29237716432993 | 0.0001315349505 | 0.000710379715500128 |
| Mgl2 | ENSMUSG00000040950 | 216864 | 1.11636907611231 | 1.45363060678766 | 5.2888075575972 | 0.0001323658216 | 0.000714513973741527 |
| Mrpl28 | ENSMUSG00000024181 | 68611 | -0.388000854036742 | 4.49058576330235 | -5.28637155454545 | 0.0001329359752 | 0.000717237485386616 |
| Tmem186 | ENSMUSG00000043140 | 66690 | -0.559364997086108 | 3.07914572250798 | -5.28309348032268 | 0.0001337072659 | 0.000721042982005531 |
| Pcbp2 | ENSMUSG00000056851 | 18521 | -0.287353050371621 | 6.32031837578184 | -5.28206560656085 | 0.0001339500727 | 0.000721996175938888 |
| Lmbrd1 | ENSMUSG00000073725 | 68421 | -0.319038654325957 | 4.47919346804327 | -5.27999101891331 | 0.0001344415392 | 0.000724288055819482 |
| Slc1a5 | ENSMUSG00000001918 | 20514 | 0.663030591628478 | 2.97795512885491 | 5.2787207802977 | 0.0001347433851 | 0.000725556622460324 |
| Phpt1 | ENSMUSG00000036504 | 75454 | -0.629676635361448 | 3.87926968538284 | -5.27435928166524 | 0.0001357851993 | 0.000730567212500858 |
| Nfat5 | ENSMUSG00000003847 | 54446 | 0.424309901988201 | 5.68780680967079 | 5.27426600072644 | 0.0001358075725 | 0.000730567212500858 |
| Dnajc21 | ENSMUSG00000044224 | 78244 | -0.420032197098733 | 6.21333282128327 | -5.2730998598218 | 0.0001360875939 | 0.000731713472332578 |
| Tmod2 | ENSMUSG00000032186 | 50876 | 0.676188587203559 | 3.41762397159945 | 5.27197129915039 | 0.0001363591649 | 0.000732813191907396 |
| Zfp467 | ENSMUSG00000068551 | 68910 | -0.818547710870901 | 1.56013323308518 | -5.2713070649249 | 0.0001365192671 | 0.000733313076001786 |
| Wwtr1 | ENSMUSG00000027803 | 97064 | 0.504661446729657 | 5.87182589233328 | 5.26843846802807 | 0.0001372129490 | 0.000736677183522899 |
| Slc25a26 | ENSMUSG00000045100 | 67582 | -0.585746914469515 | 2.23997511402577 | -5.26587053337895 | 0.0001378370447 | 0.00073966457357279 |
| Farp1 | ENSMUSG00000025555 | 223254 | 0.69717256791745 | 3.01048817187675 | 5.26476644751898 | 0.0001381062851 | 0.000740745732327973 |
| Myo1g | ENSMUSG00000020437 | 246177 | 1.42489959315646 | 1.00896721215334 | 5.26366833381551 | 0.0001383746134 | 0.000741820941218199 |
| Palld | ENSMUSG00000058056 | 72333 | 0.38963959420071 | 6.25192325722568 | 5.26240401427353 | 0.0001386842285 | 0.000743116324585013 |
| Gaa | ENSMUSG00000025579 | 14387 | -0.296743411628632 | 6.29963240950612 | -5.26118114674112 | 0.0001389843803 | 0.000744359755593774 |
| Sntb1 | ENSMUSG00000060429 | 20649 | 0.850262647113665 | 1.91118459965575 | 5.25840699411537 | 0.0001396678091 | 0.000747653683643525 |
| Atp5g3 | ENSMUSG00000018770 | 228033 | -0.507548604370862 | 7.19962984872292 | -5.25768961452578 | 0.0001398451097 | 0.000748236365734006 |
| Rgs5 | ENSMUSG00000026678 | 19737 | -0.792141175331902 | 8.04871742570648 | -5.25373280793733 | 0.0001408272643 | 0.000752968772601502 |
| Ttc13 | ENSMUSG00000037300 | 234875 | 0.426471534472979 | 3.7070659124542 | 5.2535718767676 | 0.0001408673625 | 0.000752968772601502 |
| Repin1 | ENSMUSG00000052751 | 58887 | -0.491336608963776 | 4.23708128174464 | -5.24883559156576 | 0.0001420528204 | 0.000758934203561159 |
| Dand5 | ENSMUSG00000053226 | 23863 | -0.628187169604232 | 2.63987074360586 | -5.24824109067885 | 0.0001422023531 | 0.000759361955425417 |
| Rnf146 | ENSMUSG00000038876 | 68031 | -0.332734078207355 | 4.79998300875247 | -5.24745480685877 | 0.0001424003766 | 0.000760048104150858 |
| Pitpnm2 | ENSMUSG00000029406 | 19679 | -0.32255234821468 | 4.9961632719139 | -5.24589632550032 | 0.0001427937266 | 0.000761775610470784 |
| Mkrn2 | ENSMUSG00000000439 | 67027 | -0.389746570239469 | 4.59538124464827 | -5.24158687559927 | 0.0001438873135 | 0.000767043990378037 |
| Shkbp1 | ENSMUSG00000089832 | 192192 | 0.8872069460275 | 1.12582901250076 | 5.24145224021784 | 0.0001439216196 | 0.000767043990378037 |
| Tbc1d22b | ENSMUSG00000042203 | 381085 | -0.379135989474549 | 3.42956310753046 | -5.24110490674812 | 0.0001440101622 | 0.000767141853762872 |
| Ndst1 | ENSMUSG00000054008 | 15531 | -0.27789243478779 | 5.96496462031438 | -5.24052185333182 | 0.0001441589226 | 0.000767560245505467 |
| Gatb | ENSMUSG00000028085 | 229487 | -0.579718249984461 | 2.98854778277596 | -5.23881062649318 | 0.0001445964518 | 0.000769379534552674 |
| Lbh | ENSMUSG00000024063 | 77889 | 0.90983898413737 | 4.54942911464893 | 5.2386352067144 | 0.0001446413817 | 0.000769379534552674 |
| Ddx56 | ENSMUSG00000004393 | 52513 | -0.533067446117197 | 3.63007916225334 | -5.23335395179007 | 0.0001460008970 | 0.000776190360002165 |
| Mylk | ENSMUSG00000022836 | 107589 | -0.763273959860012 | 4.6238329614413 | -5.23311077729082 | 0.0001460638158 | 0.000776190360002165 |
| Atf6b | ENSMUSG00000015461 | 12915 | 0.660841872231456 | 2.79350761858722 | 5.23265880040126 | 0.0001461808349 | 0.000776434746054969 |
| Alkbh5 | ENSMUSG00000042650 | 268420 | -0.338787456019787 | 6.76620788062795 | -5.2295152787755 | 0.0001469974191 | 0.000780392806364484 |
| Aldh5a1 | ENSMUSG00000035936 | 214579 | -0.723234136946962 | 2.50548037794794 | -5.22516582557272 | 0.0001481351181 | 0.000785889406478074 |
| Fhod3 | ENSMUSG00000034295 | 225288 | -0.453228366223377 | 5.37812473423121 | -5.22500801937179 | 0.0001481765681 | 0.000785889406478074 |
| Cenpq | ENSMUSG00000023919 | 83815 | -0.830690797131745 | 1.46625320115457 | -5.22355415103482 | 0.0001485590170 | 0.000787535701004067 |
| Lrrn1 | ENSMUSG00000034648 | 16979 | 0.935285794397378 | 5.38193739404687 | 5.22232443058689 | 0.0001488833060 | 0.000788872234042519 |
| Tuba8 | ENSMUSG00000030137 | 53857 | -0.470315834862183 | 6.91933216763979 | -5.22113819404886 | 0.0001491968275 | 0.000789836315635327 |
| Tfam | ENSMUSG00000003923 | 21780 | -0.397542348108862 | 4.31367690704146 | -5.22108928262478 | 0.0001492097696 | 0.000789836315635327 |
| Abhd18 | ENSMUSG00000037818 | 269423 | -1.61540692877605 | 4.14521884713545 | -5.22026180965511 | 0.0001494288976 | 0.000790613397722803 |
| Megf8 | ENSMUSG00000045039 | 269878 | 0.462623615967828 | 4.7493574680859 | 5.21871561178226 | 0.0001498392552 | 0.000792401015632523 |
| Fez1 | ENSMUSG00000032118 | 235180 | 0.800552555766503 | 2.35125193194132 | 5.21791100424996 | 0.0001500532611 | 0.000793149031707773 |
| Mdfic | ENSMUSG00000041390 | 16543 | 0.442949827866549 | 3.44948449699047 | 5.21431751587483 | 0.0001510129388 | 0.000797705055670779 |
| Clip3 | ENSMUSG00000013921 | 76686 | 0.691969502507325 | 2.41491856819458 | 5.21403990332879 | 0.0001510873439 | 0.000797705055670779 |
| Fn3k | ENSMUSG00000025175 | 63828 | -0.825861103243993 | 1.24040174722177 | -5.21386541476883 | 0.0001511341295 | 0.000797705055670779 |
| Bsg | ENSMUSG00000023175 | 12215 | -0.373664686069045 | 7.35682795255116 | -5.21243527147794 | 0.0001515181635 | 0.000799346064806504 |
| Upf2 | ENSMUSG00000043241 | 326622 | -0.434377804572112 | 5.23314767077714 | -5.20948874809018 | 0.0001523126001 | 0.000803149557379474 |
| Tecrl | ENSMUSG00000049537 | 243078 | 0.772529320002577 | 3.02520710578697 | 5.20833818162156 | 0.0001526239919 | 0.000804403498800619 |
| Camkk2 | ENSMUSG00000029471 | 207565 | 0.651955232272581 | 2.34024861175 | 5.2073791216877 | 0.0001528840613 | 0.000805385867259268 |
| Prdx4 | ENSMUSG00000025289 | 53381 | 0.897418369687515 | 1.11746279729949 | 5.20633199510084 | 0.0001531685395 | 0.000806495812220844 |
| Dock5 | ENSMUSG00000044447 | 68813 | 0.809670674479604 | 1.68085679838436 | 5.20483596518917 | 0.0001535759316 | 0.000807590231487126 |
| Rab3il1 | ENSMUSG00000024663 | 74760 | 0.834134571349884 | 2.95673449706585 | 5.20457179191493 | 0.0001536479873 | 0.000807590231487126 |
| Sh3bgrl3 | ENSMUSG00000028843 | 73723 | 0.547345044676927 | 3.08484052777425 | 5.20447544844949 | 0.0001536742747 | 0.000807590231487126 |
| Gdf11 | ENSMUSG00000025352 | 14561 | 1.05709311954218 | 1.99448988009887 | 5.20400811845807 | 0.0001538018525 | 0.000807590231487126 |
| Atad1 | ENSMUSG00000013662 | 67979 | -0.315296710228022 | 5.15065410093077 | -5.20396815024846 | 0.0001538127686 | 0.000807590231487126 |
| Clpp | ENSMUSG00000002660 | 53895 | -0.533768029406239 | 3.54091435129284 | -5.2039428670181 | 0.0001538196744 | 0.000807590231487126 |
| Dnajc11 | ENSMUSG00000039768 | 230935 | -0.336057862058589 | 4.94651835663798 | -5.20331246126115 | 0.0001539919666 | 0.000808106666811314 |
| Micos10 | ENSMUSG00000050608 | 433771 | -0.477291149436813 | 5.92971668613342 | -5.19999295585509 | 0.0001549025243 | 0.000812494958723918 |
| Gas2l1 | ENSMUSG00000034201 | 78926 | 0.837269471750922 | 3.19886455815278 | 5.19593897569859 | 0.0001560221722 | 0.000817975235076993 |
| Zfp131 | ENSMUSG00000094870 | 72465 | -0.411341722936533 | 4.162587958164 | -5.194403795 | 0.0001564483656 | 0.000819267939790534 |
| Loxl3 | ENSMUSG00000000693 | 16950 | 1.07168757937532 | 2.77916123081488 | 5.19433414458907 | 0.0001564677306 | 0.000819267939790534 |

| | | | | | | | |
|---|---|---|---|---|---|---|---|
| Rbmx | ENSMUSG00000031134 | 19655 | -0.36733122282691 | 3.91401227602804 | -5.19422539435536 | 0.0001564979714 | 0.000819267939790534 |
| Entpd2 | ENSMUSG00000015085 | 12496 | 0.613838433489802 | 2.5621897505699 | 5.19397170083947 | 0.0001565685414 | 0.000819267939790534 |
| Mtg1 | ENSMUSG00000039018 | 212508 | -0.542501387090426 | 2.92970160064598 | -5.19146192395325 | 0.0001572684762 | 0.000822536705262203 |
| Hk1 | ENSMUSG00000037012 | 15275 | 0.502519044265154 | 3.80642018765689 | 5.1908180771493 | 0.0001574485596 | 0.000823084746655881 |
| Selenoo | ENSMUSG00000035757 | 223776 | -0.448669427779979 | 3.04752995043722 | -5.1905266839447 | 0.0001575301326 | 0.000823117533676617 |
| Lilrb4a | ENSMUSG00000062593 | 14728 | 4.72465074906315 | -1.12832035216184 | 5.18900879372602 | 0.0001579557663 | 0.000824947196349605 |
| Acad12 | ENSMUSG00000042647 | 338350 | -0.475366903411303 | 3.91038802227104 | -5.18760900639469 | 0.0001583493452 | 0.000826607780963515 |
| Ric8b | ENSMUSG00000035620 | 237422 | -0.452163204460258 | 5.04568233089125 | -5.18542033323983 | 0.0001589667859 | 0.00082943481473692 |
| Cstf2t | ENSMUSG00000053536 | 83410 | -0.591804561595657 | 4.60745638933791 | -5.18502108769908 | 0.0001590796863 | 0.000829627887410968 |
| Ppp1r3a | ENSMUSG00000042717 | 140491 | -0.610709523283638 | 7.33041840921205 | -5.18290955471121 | 0.0001596781870 | 0.000832352056423718 |
| Tusc2 | ENSMUSG00000010054 | 80385 | -0.433342872558353 | 3.403721098332 | -5.18251509798423 | 0.0001597902532 | 0.000832539207903832 |
| Sh2d3c | ENSMUSG00000059013 | 27387 | -0.735122940591637 | 2.31620583759967 | -5.18022683615982 | 0.0001604419739 | 0.000835536549613385 |
| Mrvi1 | ENSMUSG00000005611 | 17540 | -0.877020584040297 | 2.63766503242972 | -5.17918887249529 | 0.0001607385097 | 0.000836682214320996 |
| Actr1b | ENSMUSG00000037351 | 226977 | 0.3672729895418 | 7.18743447936696 | 5.17857840647769 | 0.0001609131805 | 0.000837192753979855 |
| Trappc8 | ENSMUSG00000033382 | 75964 | -0.292773670310189 | 5.04293756480709 | -5.17810001788104 | 0.0001610501986 | 0.000837507003440598 |
| Prpsap1 | ENSMUSG00000015869 | 67763 | 0.528637734225924 | 2.89628922160243 | 5.17605852310613 | 0.0001616362836 | 0.000840155119410732 |
| Kbtbd11 | ENSMUSG00000055675 | 74901 | 1.40652365813805 | 0.758722624530474 | 5.17565054503973 | 0.0001617536747 | 0.000840365693480496 |
| Spp1 | ENSMUSG00000029304 | 20750 | 1.64459125305452 | 2.06918695539849 | 5.17482888251423 | 0.0001619903688 | 0.000841195592596455 |
| 9530068E07Rik | ENSMUSG00000036275 | 213673 | 0.376076806918295 | 5.33285619863664 | 5.17431086230579 | 0.0001621397791 | 0.000841571664523553 |
| Otud5 | ENSMUSG00000031154 | 54644 | 0.364759615415044 | 3.67683126277311 | 5.17383600295702 | 0.0001622768668 | 0.00084188345102831 |
| Mul1 | ENSMUSG00000041241 | 68350 | -0.362650523717706 | 4.22536068908478 | -5.16745376076881 | 0.0001641311328 | 0.00085109934201664 |
| Usp24 | ENSMUSG00000028514 | 329908 | -0.387677076883866 | 7.21067941847036 | -5.16620432134359 | 0.0001644967165 | 0.000852590615575614 |
| Speg | ENSMUSG00000026207 | 11790 | -0.364292004820882 | 8.3117600607257 | -5.16526167742474 | 0.0001647730940 | 0.000853618336949776 |
| Nup210 | ENSMUSG00000030091 | 54563 | -0.502473324033107 | 3.49223488786923 | -5.16364272496107 | 0.0001652488911 | 0.000855677701890912 |
| Prkce | ENSMUSG00000045038 | 18754 | -0.441179520359297 | 3.25148242759268 | -5.16116264357349 | 0.0001659805458 | 0.000859059349752705 |
| Slc30a7 | ENSMUSG00000054414 | 66500 | 0.474769763276274 | 2.92158892017991 | 5.16058417145748 | 0.0001661516873 | 0.000859538142294059 |
| Cdc14a | ENSMUSG00000033502 | 229776 | -0.534779000416848 | 3.2399753641231 | -5.15951345049962 | 0.0001664689465 | 0.000860772022336539 |
| Uqcrh | ENSMUSG00000063882 | 66576 | -0.432947617644866 | 6.47785347790279 | -5.15871985952502 | 0.0001667044983 | 0.0008613031888108 |
| Hspg2 | ENSMUSG00000028763 | 15530 | 0.609095250641897 | 8.75138556293824 | 5.15863650163196 | 0.0001667292605 | 0.0008613031888108 |
| Ddx20 | ENSMUSG00000027905 | 53975 | -0.543981712031999 | 2.85921840508423 | -5.15659582773255 | 0.0001673366589 | 0.000864032602077629 |
| Nox4 | ENSMUSG00000030562 | 50490 | 1.07020643274941 | 3.06027186580558 | 5.15519029163396 | 0.0001677563518 | 0.000865790690229369 |
| Abcb8 | ENSMUSG00000028973 | 74610 | -0.455539341991817 | 3.98514057736094 | -5.15409812996026 | 0.0001680832284 | 0.000867068320225119 |
| Ninj1 | ENSMUSG00000037966 | 18081 | 0.56737054895752 | 3.59929130340257 | 5.15324540094711 | 0.0001683389059 | 0.000867977632501045 |
| Smim4 | ENSMUSG00000058351 | 66487 | -1.02990179344331 | 1.45002292613567 | -5.15288136367063 | 0.0001684481802 | 0.000868131569203297 |
| Myo5c | ENSMUSG00000033590 | 208943 | -1.70541266742891 | 0.823030678418928 | -5.15135629881402 | 0.0001689067682 | 0.000870000707144391 |
| Mfsd12 | ENSMUSG00000034854 | 73822 | 0.845865530789196 | 1.48414878355724 | 5.15114622498133 | 0.0001689700394 | 0.000870000707144391 |
| Cisd3 | ENSMUSG00000078695 | 217149 | -0.462993424056128 | 3.32999723074778 | -5.14807706269157 | 0.0001698972484 | 0.000874362911140354 |
| Ralgds | ENSMUSG00000026821 | 19730 | 0.549801981091712 | 3.25078539461484 | 5.14541528914325 | 0.0001707056762 | 0.000878109998626431 |
| Yars2 | ENSMUSG00000022792 | 70120 | -0.53817622871119 | 2.68310161242857 | -5.14485524389189 | 0.0001708762817 | 0.000878574146600751 |
| Sap30 | ENSMUSG00000031609 | 60406 | 0.51331943215103 | 2.49743666662595 | 5.14237710210003 | 0.0001716333283 | 0.000882051674812083 |
| Fblim1 | ENSMUSG00000006219 | 74202 | 1.65194633085224 | 1.47533096455674 | 5.13901822023606 | 0.0001726650147 | 0.000886936690048779 |
| Srd5a3 | ENSMUSG00000029233 | 57357 | 0.794920325764104 | 1.89329282705673 | 5.13642125346195 | 0.0001734671058 | 0.000890638296649609 |
| Mapk1ip1 | ENSMUSG00000041775 | 69546 | -0.533021307993201 | 3.16935444876281 | -5.13600659082183 | 0.0001735955362 | 0.000890879251759188 |
| Napb | ENSMUSG00000027438 | 17957 | 0.849000689008452 | 2.38439898871004 | 5.13473172897866 | 0.0001739910106 | 0.00089248978752079 |
| Ppargc1a | ENSMUSG00000029167 | 19017 | -1.36339866471371 | 5.65284558816436 | -5.13299527578315 | 0.0001745311857 | 0.00089484070896935 |
| Tmem132a | ENSMUSG00000024736 | 98170 | 0.85128357823271 | 2.85211692861714 | 5.13243398025231 | 0.0001747061667 | 0.000895203941253691 |
| Abra | ENSMUSG00000042895 | 223513 | -0.530340735505161 | 7.64514005460773 | -5.13224275199001 | 0.0001747658229 | 0.000895203941253691 |
| Ruvbl2 | ENSMUSG00000003868 | 20174 | -0.475953747712194 | 3.26301563995471 | -5.13110247280281 | 0.0001751219888 | 0.000896578371832793 |
| Cdh15 | ENSMUSG00000031962 | 12555 | 0.661084971483416 | 3.81935881829481 | 5.1308588294514 | 0.0001751981888 | 0.000896578371832793 |
| Pkp4 | ENSMUSG00000026991 | 227937 | -0.553499055870915 | 4.44400869593826 | -5.13002235914856 | 0.0001754600596 | 0.000897498320798258 |
| Chd4 | ENSMUSG00000063870 | 107932 | 0.464470897440643 | 6.80569094613985 | 5.12844254670838 | 0.0001759557588 | 0.000899612909222482 |
| Ptpn6 | ENSMUSG00000004266 | 15170 | 1.44817863912344 | 1.51534020618209 | 5.1274339045958 | 0.0001762730043 | 0.000900813562663828 |
| Mrpl39 | ENSMUSG00000022889 | 27393 | -0.353796973094528 | 4.17271620163226 | -5.12644055822149 | 0.0001765860213 | 0.000901991495186721 |
| Slc25a33 | ENSMUSG00000028982 | 70556 | -0.844784183458135 | 3.51340726816071 | -5.12488026279172 | 0.0001770788614 | 0.000904086424427195 |
| Mrps31 | ENSMUSG00000031533 | 57312 | -0.498272644853374 | 3.66305251289506 | -5.12345528411076 | 0.0001775302119 | 0.000905967668773933 |
| Cars | ENSMUSG00000010755 | 27267 | -0.305043695405719 | 4.67455948993612 | -5.12299062230913 | 0.0001776776487 | 0.000906296956927003 |
| Mtpn | ENSMUSG00000029840 | 14489 | 0.602712945684507 | 4.40471142112125 | 5.12146298097434 | 0.0001781632679 | 0.000908350131365769 |
| Cd300ld3 | ENSMUSG00000069607 | 382551 | 2.08120118161077 | 0.688964709428312 | 5.11990188942074 | 0.0001786609491 | 0.000910462859979178 |
| Tpi1 | ENSMUSG00000023456 | 21991 | -0.479372404332999 | 7.40103149157672 | -5.11728264408726 | 0.0001794992292 | 0.000914308515564568 |
| Acvr2b | ENSMUSG00000061393 | 11481 | -0.625115600489627 | 2.41948953221604 | -5.11660210604904 | 0.0001797177028 | 0.000914994974196916 |
| Zfp839 | ENSMUSG00000021271 | 72805 | -0.655310050696624 | 3.54113758936142 | -5.1161857335563 | 0.0001798515077 | 0.000915249921459358 |
| Neo1 | ENSMUSG00000032340 | 18007 | -0.302448606062973 | 5.31083614082919 | -5.1158488670574 | 0.0001799598386 | 0.000915375056608479 |
| Lair1 | ENSMUSG00000055541 | 52855 | 1.16068200830226 | 0.951516432561717 | 5.11555587194876 | 0.0001800541165 | 0.000915428626904609 |
| Ccn5 | ENSMUSG00000027656 | 22403 | 1.92732605025713 | 0.649572001498852 | 5.11345852487289 | 0.0001807304897 | 0.000918440252882815 |
| Metap1d | ENSMUSG00000041921 | 66559 | -0.429696456048432 | 3.23870559855907 | -5.11263058849445 | 0.0001809982190 | 0.000919373388561116 |
| Fzd9 | ENSMUSG00000049551 | 14371 | -0.763899402682495 | 5.45209338588413 | -5.11085345450448 | 0.0001815742856 | 0.000921871117660173 |
| Helb | ENSMUSG00000020228 | 117599 | 0.743053931254505 | 2.6397194495086 | 5.11005657030425 | 0.0001818332193 | 0.000922757159288799 |
| Ecm2 | ENSMUSG00000043631 | 407800 | 0.675512004301972 | 4.40745115255196 | 5.10916347518724 | 0.0001821238723 | 0.000923803270747499 |
| Slc3a2 | ENSMUSG00000010095 | 17254 | 0.440123253818794 | 3.62732554135979 | 5.10849396819892 | 0.0001823420770 | 0.000924084978205209 |
| Vcam1 | ENSMUSG00000027962 | 22329 | 0.565296273989343 | 2.18360013367216 | 5.1082202168745 | 0.0001824313761 | 0.000924084978205209 |
| Uaca | ENSMUSG00000034485 | 72565 | -0.484869772870024 | 8.19370799373703 | -5.10821516718848 | 0.0001824330237 | 0.000924084978205209 |
| Syne1 | ENSMUSG00000096054 | 64009 | 0.433996144059535 | 7.69048626268746 | 5.10768102482925 | 0.0001826073965 | 0.000924539810640361 |
| Csk | ENSMUSG00000032312 | 12988 | 0.590731096358887 | 2.89303753355049 | 5.10593697954889 | 0.0001831779562 | 0.000926999184852366 |
| Mlip | ENSMUSG00000032355 | 69642 | 0.397194058234376 | 6.16067256299922 | 5.10524137323473 | 0.0001834060396 | 0.000927723932974912 |
| BC048679 | ENSMUSG00000061877 | 210321 | -5.25214043496802 | -0.472772360295476 | -5.103673445365 | 0.0001839212353 | 0.000929549606180333 |
| Btnl9 | ENSMUSG00000040283 | 237754 | -1.01290360188358 | 2.14383030504994 | -5.10362541456863 | 0.0001839370412 | 0.000929549606180333 |
| Ppp3cb | ENSMUSG00000021816 | 19056 | -0.353918507743634 | 5.97979552383116 | -5.10255372747987 | 0.0001842900789 | 0.000930484480313661 |
| Tcp11l2 | ENSMUSG00000020034 | 216198 | 0.428393836785008 | 4.71873822085206 | 5.10254705619331 | 0.0001842922788 | 0.000930484480313661 |
| Fjx1 | ENSMUSG00000075012 | 14221 | 1.9581373067623 | 0.0800329852446794 | 5.10054986456427 | 0.0001849520909 | 0.000933384721165216 |
| Ndufb3 | ENSMUSG00000026032 | 66495 | -0.472035914110926 | 5.07652938004038 | -5.09928601245199 | 0.0001853709003 | 0.000935066595242543 |
| Tpm1 | ENSMUSG00000032366 | 22003 | -0.563641808446068 | 11.5812216072989 | -5.09634549963375 | 0.0001863491405 | 0.000939567553257854 |
| Shox2 | ENSMUSG00000027833 | 20429 | 1.25519428815613 | 0.658142733593269 | 5.09574181665108 | 0.0001865506361 | 0.000940149840405136 |
| Enpp1 | ENSMUSG00000037370 | 18605 | 1.1536286719048 | 3.35144603683886 | 5.09086545077805 | 0.0001881865979 | 0.000947957466259689 |
| Arrb2 | ENSMUSG00000060216 | 216869 | 1.10648316760702 | 1.85753538835137 | 5.0894371752295 | 0.0001886685906 | 0.000949947657448665 |
| Mrpl13 | ENSMUSG00000022370 | 68537 | -0.488907445079571 | 3.95419837438301 | -5.08799258735533 | 0.0001891573960 | 0.000951970301982738 |
| Prelid1 | ENSMUSG00000021486 | 66494 | 0.588962314033165 | 3.86999614647782 | 5.08693759617482 | 0.0001895152066 | 0.000953332132329752 |
| Man2b1 | ENSMUSG00000005142 | 17159 | 0.7297088001845 | 3.70570739183369 | 5.08619018285517 | 0.0001897691256 | 0.000954170336946488 |
| Actr2 | ENSMUSG00000020152 | 66713 | 0.265021842312134 | 5.66236850454124 | 5.08412438762775 | 0.0001904727835 | 0.000957268044469099 |
| Ulk1 | ENSMUSG00000029512 | 22241 | -0.309804883837629 | 5.40052417771868 | -5.08348761465299 | 0.0001906902305 | 0.000957920455210909 |
| Llgl1 | ENSMUSG00000020536 | 16897 | 0.46852150123484 | 3.62264148301237 | 5.08312702708005 | 0.0001908134796 | 0.000958099286354708 |
| Selenom | ENSMUSG00000075702 | 114679 | 0.62570633960031 | 2.50108280439802 | 5.08283767419661 | 0.0001909124406 | 0.000958156055652805 |
| Hectd2 | ENSMUSG00000041180 | 226098 | 0.464548457896994 | 3.21777954126905 | 5.08254884904251 | 0.0001910112745 | 0.000958212134790741 |
| P2ry6 | ENSMUSG00000048779 | 233571 | 1.64390691147631 | 0.72233437155656 | 5.08005531564147 | 0.0001918667617 | 0.000962062189461004 |
| Cgrrf1 | ENSMUSG00000055128 | 68755 | -0.464321986376693 | 3.69972221634394 | -5.0788049768368 | 0.0001922972317 | 0.000963778560647161 |
| Ybx1 | ENSMUSG00000028639 | 22608 | -0.456412935919429 | 9.07808428227286 | -5.07833237120376 | 0.0001924602035 | 0.000964153292700271 |
| Arhgap35 | ENSMUSG00000058230 | 232906 | 0.334202271953332 | 5.94603756576589 | 5.07514626799172 | 0.0001935626480 | 0.000969231930978684 |
| Stac3 | ENSMUSG00000040287 | 237611 | -0.413816752786189 | 6.40523140689419 | -5.07381169836839 | 0.0001940263831 | 0.000970889252553744 |
| C3 | ENSMUSG00000024164 | 12266 | 0.867227272066262 | 5.40184391462517 | 5.07368270561438 | 0.0001940712667 | 0.000970889252553744 |
| Piezo1 | ENSMUSG00000014444 | 234839 | 0.683514475018059 | 4.48988065221364 | 5.07153668481315 | 0.0001948195715 | 0.00097418697946484 |
| Tmem70 | ENSMUSG00000025940 | 70397 | -0.468015893175947 | 4.61314468961035 | -5.06942364815807 | 0.0001955593105 | 0.000977438876998645 |
| Adamts12 | ENSMUSG00000047497 | 239337 | 0.745650230503401 | 3.10264952334179 | 5.06915517148623 | 0.0001956535089 | 0.000977462754346793 |
| Ccpg1 | ENSMUSG00000034563 | 72278 | -0.279981330540521 | 5.92070792579547 | -5.06763021508638 | 0.0001961894558 | 0.000979692527076028 |
| Spr | ENSMUSG00000033735 | 20751 | -0.387836911222628 | 3.50169008468227 | -5.05983957178968 | 0.0001989514310 | 0.000992728216828876 |
| H3f3b | ENSMUSG00000016559 | 15081 | 0.340788663847063 | 5.62855033977333 | 5.05975518672619 | 0.0001989815682 | 0.000992728216828876 |
| Osbpl1a | ENSMUSG00000044252 | 64291 | -0.482997524304689 | 5.51269178396571 | -5.05942895107078 | 0.0001990981245 | 0.000992856568933539 |
| Gm33543 | ENSMUSG00000110353 | 102636491 | -1.4936943231302 | 0.783455912602117 | -5.05616398215818 | 0.0002002685425 | 0.000998237774006576 |
| Cdk2ap2 | ENSMUSG00000024856 | 52004 | 0.520493484345624 | 2.59480898056246 | 5.05437263115613 | 0.0002009137422 | 0.00100099731836438 |
| Tbcc | ENSMUSG00000036430 | 72726 | 0.44919464065931 | 2.68502586929208 | 5.05356548132611 | 0.0002012051641 | 0.00100194308229027 |
| Bin2 | ENSMUSG00000098112 | 668218 | 0.88271046846758 | 1.16296183440497 | 5.05333934043593 | 0.0002012868912 | 0.00100194308229027 |
| Coq8b | ENSMUSG00000003762 | 76889 | 0.894528923041341 | 1.75317690680932 | 5.04966268814224 | 0.0002026204897 | 0.00100812224546349 |
| Vdac2 | ENSMUSG00000021771 | 22334 | -0.35681009111756 | 6.70888105366497 | -5.04853144377395 | 0.0002030326632 | 0.00100971339497269 |
| Zfp593 | ENSMUSG00000028840 | 68040 | 0.780514304118309 | 1.87494015741185 | 5.04812158817309 | 0.0002031822111 | 0.00100999761245804 |
| Ap2b1 | ENSMUSG00000035152 | 71770 | 0.299666808240526 | 4.99388787414148 | 5.04704909930855 | 0.0002035740829 | 0.00101124430542156 |
| Pdss2 | ENSMUSG00000038240 | 71365 | -0.553431611311567 | 3.1905869809391 | -5.04692895000649 | 0.0002036180327 | 0.00101124430542156 |
| Aldoa | ENSMUSG00000030695 | 11674 | -0.515653151328008 | 10.7128188844751 | -5.04622547452977 | 0.0002038755575 | 0.00101206345100893 |
| Snrk | ENSMUSG00000038145 | 20623 | -0.463069258608546 | 5.17919937020425 | -5.0427174917526 | 0.0002051648115 | 0.0010180011600286 |

| | | | | | | | |
|---|---|---|---|---|---|---|---|
| Eif1a | ENSMUSG00000057561 | 13664 | 0.595906105942313 | 2.87923307637112 | 5.04131389634167 | 0.0002056830352 | 0.00101966223718256 |
| Stat5b | ENSMUSG00000020919 | 20851 | -0.666183636428431 | 5.83471610674918 | -5.04124243755393 | 0.0002057094550 | 0.00101966223718256 |
| Csrp3 | ENSMUSG00000030470 | 13009 | 0.603931537971325 | 8.71805106557677 | 5.04105323305499 | 0.0002057794250 | 0.00101966223718256 |
| Zfp691 | ENSMUSG00000045268 | 195522 | -0.616416753207283 | 3.18294165104944 | -5.03904214695096 | 0.0002065246801 | 0.00102289138157984 |
| Sdc1 | ENSMUSG00000020592 | 20969 | 0.799760349360932 | 1.77651329463647 | 5.03841034213925 | 0.0002067593897 | 0.00102359007650247 |
| Agpat2 | ENSMUSG00000026922 | 67512 | -0.802397102695886 | 2.16484882118923 | -5.03658838611584 | 0.0002074377849 | 0.00102648367019577 |
| Acss3 | ENSMUSG00000035948 | 380660 | -0.889736191672667 | 1.42954491429528 | -5.03438801157723 | 0.0002082601715 | 0.00102968122103885 |
| Abcc9 | ENSMUSG00000030249 | 20928 | -0.37008450756003 | 7.22241323117385 | -5.034355465226 | 0.0002082723611 | 0.00102968122103885 |
| Aars | ENSMUSG00000031960 | 234734 | -0.414908079211331 | 6.12504618644703 | -5.03401355944092 | 0.0002084004599 | 0.00102984874678894 |
| Nrbp2 | ENSMUSG00000075590 | 223649 | 0.495604721848047 | 2.85322104985777 | 5.03157770348368 | 0.0002093154567 | 0.00103390296319609 |
| Mgme1 | ENSMUSG00000027424 | 74528 | -0.631918084789896 | 2.22576403084501 | -5.03063234580669 | 0.0002096716929 | 0.00103514855846525 |
| Ube2z | ENSMUSG00000014349 | 268470 | 0.432937271551971 | 4.32256893014975 | 5.03040614093941 | 0.0002097570265 | 0.00103514855846525 |
| Agpat1 | ENSMUSG00000034254 | 55979 | -0.375479530500805 | 4.39745852270419 | -5.02996012093269 | 0.0002099253891 | 0.00103551192632098 |
| Kctd11 | ENSMUSG00000046731 | 216858 | 1.06187500856119 | 1.23232581561037 | 5.02969544321572 | 0.0002100253655 | 0.00103553778573809 |
| Sec63 | ENSMUSG00000019802 | 140740 | 0.285412289601661 | 5.34743122912251 | 5.02906608126629 | 0.0002102632927 | 0.00103624348668696 |
| Chd9 | ENSMUSG00000056608 | 109151 | 0.478246399243628 | 5.75493697574864 | 5.02610254045893 | 0.0002113874207 | 0.00104131405861644 |
| Cux1 | ENSMUSG00000029705 | 13047 | 0.398632625714802 | 6.42547730788963 | 5.02350564795011 | 0.0002123776128 | 0.00104572057950899 |
| Lrrk2 | ENSMUSG00000036273 | 66725 | 0.462658573025877 | 3.86811509305962 | 5.02285675045053 | 0.0002126257896 | 0.00104647118704599 |
| Gxylt2 | ENSMUSG00000030074 | 232313 | 0.993205741803776 | 2.56930274497089 | 5.02238783724224 | 0.0002128053175 | 0.00104685266315951 |
| Dusp10 | ENSMUSG00000039384 | 63953 | 0.723363489743303 | 4.80349404432414 | 5.02215417139575 | 0.0002128948376 | 0.00104685266315951 |
| Churc1 | ENSMUSG00000090258 | 211151 | -0.521199344635527 | 3.76273372379699 | -5.02167256032819 | 0.0002130794720 | 0.00104728943742177 |
| Kcnj2 | ENSMUSG00000041695 | 16518 | -0.792757369016607 | 6.01609728720422 | -5.02054295988458 | 0.0002135131789 | 0.00104855257612824 |
| Efs | ENSMUSG00000022203 | 13644 | 1.04256178345509 | 0.893888536109069 | 5.0205035753365 | 0.0002135283171 | 0.00104855257612824 |
| Zfand3 | ENSMUSG00000044477 | 21769 | -0.276192377601115 | 5.70397021287528 | -5.01956373713763 | 0.0002138898920 | 0.00104985649307766 |
| Pcnx | ENSMUSG00000021140 | 54604 | -0.323911109198321 | 6.69762912583099 | -5.01731476130296 | 0.0002147577074 | 0.00105364295361082 |
| Pdhb | ENSMUSG00000021748 | 68263 | -0.424566944608476 | 6.88519698911027 | -5.01660036455739 | 0.0002150341388 | 0.00105446391676609 |
| Klhdc3 | ENSMUSG00000063576 | 71765 | -0.377558984153601 | 4.44227481617478 | -5.01638390279303 | 0.0002151179703 | 0.00105446391676609 |
| Reep6 | ENSMUSG00000035504 | 70335 | -0.958642906121889 | 2.07895879607508 | -5.0159611456585 | 0.0002152817943 | 0.00105471471987959 |
| Pmvk | ENSMUSG00000027952 | 68603 | -0.733151225076633 | 1.46883232165704 | -5.01575400167267 | 0.0002153621127 | 0.00105471471987959 |
| Myo9b | ENSMUSG00000004677 | 17925 | 0.581301670213116 | 4.44579913738187 | 5.01494686718241 | 0.0002156753702 | 0.00105523669940931 |
| Akap11 | ENSMUSG00000022016 | 219181 | 0.330228690166969 | 5.01387075099551 | 5.01483864451053 | 0.0002157174085 | 0.00105523669940931 |
| Baiap2l1 | ENSMUSG00000038859 | 66898 | 1.02797598911756 | 0.892730539411086 | 5.0147333848701 | 0.0002157583041 | 0.00105523669940931 |
| Nfyb | ENSMUSG00000020248 | 18045 | -0.411661315800681 | 3.84197272577443 | -5.01431444909966 | 0.0002159211495 | 0.00105556086138214 |
| Ppfia1 | ENSMUSG00000037519 | 233977 | -0.334782342451748 | 5.19839284741354 | -5.01405300191644 | 0.0002160228420 | 0.00105558591258266 |
| Scara3 | ENSMUSG00000034463 | 219151 | 0.659147723061981 | 3.06240378926371 | 5.01365093423468 | 0.0002161793275 | 0.00105587856522442 |
| Polb | ENSMUSG00000031536 | 18970 | -0.289622921388136 | 4.95547379667209 | -5.0130888539384 | 0.0002163982878 | 0.00105647596456768 |
| 4632415L05Rik | ENSMUSG00000048106 | 70808 | -0.505702417325704 | 2.78258977487726 | -5.01275422959944 | 0.0002165287515 | 0.0010566409744821 |
| Ncapg2 | ENSMUSG00000042029 | 76044 | 0.740173484268023 | 1.46888393654785 | 5.01181639426672 | 0.0002168948318 | 0.00105795511260202 |
| Hip1 | ENSMUSG00000039959 | 215114 | 0.311896549492352 | 5.24310311554589 | 5.01155486013356 | 0.0002169970354 | 0.00105798153212414 |
| Lyrm9 | ENSMUSG00000072640 | 66274 | -0.587901455945772 | 2.75287859714716 | -5.01126462441641 | 0.0002171105139 | 0.00105806287454092 |
| Rab3d | ENSMUSG00000019066 | 19340 | 0.779221662172795 | 2.47894279693089 | 5.01010410374067 | 0.0002175648798 | 0.00105980467995592 |
| Gdi2 | ENSMUSG00000021218 | 14569 | 0.299935620513628 | 6.2882200682953 | 5.00788721831475 | 0.0002184355847 | 0.00106357210533653 |
| Creb3l1 | ENSMUSG00000027230 | 26427 | 0.756451420659115 | 2.8198622488388 | 5.00550911860752 | 0.0002193736394 | 0.00106766395931863 |
| Mfap2 | ENSMUSG00000060572 | 17150 | 1.30376571952129 | 1.03917241186839 | 5.0039304443714 | 0.0002199986699 | 0.00107022939964933 |
| Ndufa4l2 | ENSMUSG00000040280 | 407790 | -0.90811112562833 | 1.87641781234418 | -5.00268082624699 | 0.0002204947336 | 0.0010720413681349 |
| ND3 | ENSMUSG00000064360 | 17718 | -0.566952110050787 | 8.62418409244999 | -5.00249829932827 | 0.0002205672890 | 0.0010720413681349 |
| Fmr1 | ENSMUSG00000000838 | 14265 | 0.423790689161283 | 3.52670520498881 | 5.00075388120505 | 0.0002212619590 | 0.00107479819931131 |
| CYTB | ENSMUSG00000064370 | 17711 | -0.626154905280526 | 12.8029593877205 | -5.00058045716856 | 0.0002213311450 | 0.00107479819931131 |
| Ywhaz | ENSMUSG00000022285 | 22631 | 0.36234232690422 | 6.33767772636105 | 4.99992457148152 | 0.0002215930083 | 0.00107559199567437 |
| Dennd4c | ENSMUSG00000038024 | 329877 | -0.33462107063632 | 5.58821441450802 | -4.9988389939024 | 0.0002220271351 | 0.00107722086730195 |
| Prickle1 | ENSMUSG00000036158 | 106042 | -0.541128768034231 | 2.9915262041749 | -4.99658454072894 | 0.0002229315287 | 0.00108112889993928 |
| Lcp2 | ENSMUSG00000002699 | 16822 | 1.33923652901846 | 0.634461903342973 | 4.99565902900429 | 0.0002233039145 | 0.00108210533565916 |
| Pkia | ENSMUSG00000027499 | 18767 | -0.48300419497312 | 7.31669901526613 | -4.99541950124981 | 0.0002234003954 | 0.00108210533565916 |
| Arfgef2 | ENSMUSG00000074582 | 99371 | -0.343319428296963 | 5.92839172334016 | -4.99534638582078 | 0.0002234298547 | 0.00108210533565916 |
| Med28 | ENSMUSG00000015804 | 66999 | -0.36975698874718 | 4.16801300194042 | -4.99507385671253 | 0.0002235396963 | 0.00108215784805825 |
| Nap1l4 | ENSMUSG00000059119 | 17955 | -0.37023969354039 | 6.40571796269668 | -4.99461598410792 | 0.0002237243662 | 0.00108257239817078 |
| Gabarapl1 | ENSMUSG00000030161 | 57436 | -0.938833841188785 | 4.75312198146488 | -4.99118515675038 | 0.0002251131520 | 0.00108881055953969 |
| Arf6 | ENSMUSG00000044147 | 11845 | 0.506699396116252 | 3.9033781961598 | 4.98893409142121 | 0.0002260292440 | 0.00109275792415054 |
| Cx3cl1 | ENSMUSG00000031778 | 20312 | 1.16912133771026 | 2.53834139511474 | 4.98867570406366 | 0.0002261346449 | 0.00109278417519323 |
| Egf | ENSMUSG00000028017 | 13645 | 0.628111938829979 | 4.51674853476112 | 4.98767521407611 | 0.0002265432456 | 0.00109427495286881 |
| Phactr2 | ENSMUSG00000062866 | 215789 | -0.35155198440949 | 5.18636923636019 | -4.98680662915637 | 0.0002268985989 | 0.00109550732554136 |
| Apol10b | ENSMUSG00000050014 | 328561 | -1.20014465058439 | 1.4236734114326 | -4.98587645577328 | 0.0002272797920 | 0.00109686331437373 |
| Cox14 | ENSMUSG00000023020 | 66379 | -0.371855223254846 | 4.60784279446676 | -4.97723891725505 | 0.0002308515011 | 0.00111360889622948 |
| Serf1 | ENSMUSG00000021643 | 20365 | 1.16053740882414 | 0.910131778133659 | 4.97189073725538 | 0.0002330922442 | 0.00112392206501853 |
| Micu2 | ENSMUSG00000021973 | 68514 | 0.403954761665591 | 6.13292822996001 | 4.97131416819276 | 0.0002333351580 | 0.00112459727167634 |
| Rps29 | ENSMUSG00000034892 | 20090 | -0.325427945539679 | 6.00463318601995 | -4.97081676309585 | 0.0002335449308 | 0.00112511222509491 |
| Rmnd5a | ENSMUSG00000002222 | 68477 | -0.338247522314248 | 6.63991763525486 | -4.96872272953307 | 0.0002344302096 | 0.00112846835927149 |
| Rpia | ENSMUSG00000053604 | 19895 | -0.692674581347017 | 2.38932438013044 | -4.96868061258904 | 0.0002344480508 | 0.00112846835927149 |
| Rbfa | ENSMUSG00000024570 | 68731 | -0.42376830597196 | 4.25671368271304 | -4.9641972754601 | 0.0002363553416 | 0.00113714799264829 |
| Tbc1d2b | ENSMUSG00000037410 | 67016 | 0.574645381970864 | 3.23076077039242 | 4.96070114760117 | 0.0002378538485 | 0.00114384618408489 |
| Zc2hc1a | ENSMUSG00000043542 | 67306 | 0.667711097413828 | 2.78362548229301 | 4.96046170160952 | 0.0002379568404 | 0.00114384618408489 |
| Arf5 | ENSMUSG00000020440 | 11844 | 0.336489285671097 | 4.34908493130467 | 4.95946482115606 | 0.0002383861229 | 0.00114504693431764 |
| B3glct | ENSMUSG00000051950 | 381694 | 0.644938153432508 | 1.89773042135408 | 4.95939518619865 | 0.0002384161396 | 0.00114504693431764 |
| Agpat4 | ENSMUSG00000023827 | 68262 | 1.11785896093133 | 0.902648142976301 | 4.95882893346402 | 0.0002386603732 | 0.00114571653042457 |
| Runx2 | ENSMUSG00000039153 | 12393 | 1.29994486519455 | 0.717950906963365 | 4.95855288120255 | 0.0002387795331 | 0.00114578537177785 |
| Snx2 | ENSMUSG00000034484 | 67804 | 0.35440685967643 | 4.21130545474398 | 4.9579846875228 | 0.0002390249927 | 0.00114645993663006 |
| Hlf | ENSMUSG00000003949 | 217082 | -1.17421188438234 | 1.89927924146519 | -4.9563382208036 | 0.0002397377487 | 0.00114937426812073 |
| Boc | ENSMUSG00000022687 | 117606 | 0.917853550763753 | 3.34610954643162 | 4.95366336950202 | 0.0002409004075 | 0.00115444206690913 |
| Tmem184c | ENSMUSG00000031617 | 234463 | 0.750047069372759 | 2.03023570909931 | 4.95320537383505 | 0.0002411000683 | 0.00115489257101575 |
| Cyyr1 | ENSMUSG00000041134 | 224405 | -0.757868740546519 | 4.7729564524264 | -4.95266267358205 | 0.0002413368784 | 0.00115552055102336 |
| Alcam | ENSMUSG00000022636 | 11658 | 1.07210464009918 | 0.718944200342909 | 4.95124887779912 | 0.0002419549317 | 0.00115797257408834 |
| Itgb8 | ENSMUSG00000025321 | 320910 | 0.91583454729843 | 1.38170090391778 | 4.94944257800616 | 0.0002427469667 | 0.00116117548067915 |
| Anxa6 | ENSMUSG00000018340 | 11749 | 0.39950474224883 | 6.95524864868345 | 4.9492384911325 | 0.0002428366250 | 0.00116117548067915 |
| Hist1h1d | ENSMUSG00000052565 | 14957 | 0.904496472101914 | 2.50799240403726 | 4.9484088721385 | 0.0002432014432 | 0.00116206013982516 |
| Arsi | ENSMUSG00000036412 | 545260 | 1.30309754357346 | 0.773343701542384 | 4.94833432826113 | 0.0002432342512 | 0.00116206013982516 |
| Cd274 | ENSMUSG00000016496 | 60533 | -1.2196718873463 | 2.21592188397099 | -4.94803492575207 | 0.0002433660693 | 0.00116218195906321 |
| Clptm1l | ENSMUSG00000021610 | 218335 | 0.344714187250688 | 4.48394085587943 | 4.94732383150776 | 0.0002436794410 | 0.00116317029286539 |
| H6pd | ENSMUSG00000028980 | 100198 | 0.403778863483462 | 4.24780469396938 | 4.94686187255333 | 0.0002438832462 | 0.00116363499134001 |
| Ccdc136 | ENSMUSG00000029769 | 232664 | 0.731320736425509 | 4.37245715797477 | 4.94403267837683 | 0.0002451352905 | 0.0011690985433595 |
| Cul5 | ENSMUSG00000032030 | 75717 | -0.326298343677276 | 6.17518729972006 | -4.93865217333027 | 0.0002475348746 | 0.00118002778642786 |
| Nceh1 | ENSMUSG00000027698 | 320024 | -0.497939273406358 | 5.16073249445056 | -4.93815047735319 | 0.0002477598607 | 0.00118058545654936 |
| Sbk3 | ENSMUSG00000085272 | 381835 | 1.00524876561083 | 3.67000235177714 | 4.93773634185932 | 0.0002479457403 | 0.00118095637812727 |
| Tbc1d8 | ENSMUSG00000003134 | 54610 | 0.35758404198225 | 3.28947141565228 | 4.93604656313249 | 0.0002487056805 | 0.00118406001473807 |
| P4ha3 | ENSMUSG00000051048 | 320452 | 3.85856275032529 | -0.46869205132839 | 4.93439901903335 | 0.0002494489578 | 0.00118708165345511 |
| Ppard | ENSMUSG00000002250 | 19015 | -0.396091650221948 | 4.21750057247671 | -4.93386911608615 | 0.0002496885100 | 0.0011877045705897 |
| E2f8 | ENSMUSG00000046179 | 108961 | 1.78532462030757 | 0.674371997305544 | 4.93349114113329 | 0.0002498595265 | 0.00118800108077148 |
| Cytip | ENSMUSG00000026832 | 227929 | 1.57970183366447 | 0.264181270646827 | 4.92965082101758 | 0.0002516040152 | 0.00119577543067479 |
| Rgs4 | ENSMUSG00000038530 | 19736 | -0.735609091745754 | 3.4100927497872 | -4.92929282128133 | 0.0002517672825 | 0.00119603136271559 |
| Washc4 | ENSMUSG00000034560 | 319277 | 0.367752185277748 | 4.28800577576649 | 4.92252686521148 | 0.0002548737183 | 0.00121026264779064 |
| Mark1 | ENSMUSG00000026620 | 226778 | -0.594096373849096 | 3.24850863846985 | -4.92183112649654 | 0.0002551954018 | 0.0012107111563578 |
| Eef1d | ENSMUSG00000055762 | 66656 | -0.355053805507109 | 5.76153906053812 | -4.92163577900881 | 0.0002552857990 | 0.0012107111563578 |
| Sars | ENSMUSG00000068739 | 20226 | -0.360407239825008 | 5.59986879458526 | -4.92160412592602 | 0.0002553004496 | 0.0012107111563578 |
| Asap2 | ENSMUSG00000052632 | 211914 | 0.483455165712332 | 4.44794869868211 | 4.92049736836325 | 0.0002558132619 | 0.00121261698441244 |
| Casc4 | ENSMUSG00000060227 | 319996 | 0.357076800234578 | 4.34109492013983 | 4.9195470411485 | 0.0002562544468 | 0.00121370200613695 |
| Azi2 | ENSMUSG00000039285 | 27215 | -0.308569937549385 | 5.04338005255075 | -4.91952600067106 | 0.0002562642237 | 0.00121370200613695 |
| Rbbp8 | ENSMUSG00000041238 | 225182 | 0.521751742632156 | 1.99322076268537 | 4.91915618454435 | 0.0002564361296 | 0.00121376255555659 |
| Arhgdia | ENSMUSG00000025132 | 192662 | 0.315762158904384 | 5.77738721958442 | 4.91902081398972 | 0.0002564990854 | 0.00121376255555659 |
| Tmem170b | ENSMUSG00000087370 | 621976 | -0.395909521278444 | 4.21808188657252 | -4.91743543147408 | 0.0002572375845 | 0.00121673043581975 |
| Bicd2 | ENSMUSG00000037933 | 76895 | 0.355039385830179 | 5.596951250252 | 4.91650671334613 | 0.0002576712238 | 0.00121825438917375 |
| Copb2 | ENSMUSG00000032458 | 50797 | 0.326354285640085 | 6.08401535568392 | 4.91192257827596 | 0.0002598228147 | 0.0012278958875421 |
| Eif4b | ENSMUSG00000058655 | 75705 | -0.291449981581985 | 7.27589837164239 | -4.91138131126111 | 0.0002600780910 | 0.00122857113800275 |
| Apopt1 | ENSMUSG00000037787 | 68020 | -0.357040241095151 | 3.66443153809235 | -4.90770064345814 | 0.0002618209206 | 0.00123626975534279 |
| Cd9 | ENSMUSG00000030342 | 12527 | 0.629983262202869 | 4.10030672381049 | 4.90678799769592 | 0.0002622549413 | 0.00123772398811958 |
| Zfp24 | ENSMUSG00000051469 | 59057 | -0.358258289365422 | 4.49562040603663 | -4.90657705616044 | 0.0002623553636 | 0.00123772398811958 |
| Lrrn2 | ENSMUSG00000026443 | 16980 | 1.01831212049589 | 1.4566315549234 | 4.90326828603806 | 0.0002639357997 | 0.00124460200301238 |
| Siglec1 | ENSMUSG00000027322 | 20612 | 1.2694221817119 | 1.59498374203995 | 4.90304879609003 | 0.0002640409884 | 0.00124460200301238 |

| | | | | | | | |
|---|---|---|---|---|---|---|---|
| Uba5 | ENSMUSG00000032557 | 66663 | 0.404611028075774 | 3.78144596161454 | 4.90278707435324 | 0.0002641664733 | 0.00124465677610435 |
| Itgbl1 | ENSMUSG00000032925 | 223272 | 0.773310230199912 | 4.26283557178978 | 4.90188839823336 | 0.0002645978238 | 0.00124615200875201 |
| Abcb1a | ENSMUSG00000040584 | 18671 | -0.669298275754435 | 3.48344358770506 | -4.90084354976701 | 0.0002651002548 | 0.00124798057106075 |
| Dysf | ENSMUSG00000033788 | 26903 | 0.372332876167488 | 5.79314742224839 | 4.90043725023165 | 0.0002652958976 | 0.0012483639506366 |
| Mars | ENSMUSG00000040354 | 216443 | -0.370917077922331 | 4.65448282087269 | -4.89891107799389 | 0.0002660321265 | 0.00125128966226385 |
| Fam102a | ENSMUSG00000039157 | 98952 | -0.706864217585723 | 2.39473266189455 | -4.89836011062294 | 0.0002662984356 | 0.00125198091156695 |
| Capn7 | ENSMUSG00000021893 | 12339 | -0.309860116911582 | 5.03775368279905 | -4.89777290312167 | 0.0002665825662 | 0.00125198091156695 |
| Srpx | ENSMUSG00000090084 | 51795 | 0.865256127705755 | 0.987490231640212 | 4.89772128571198 | 0.0002666075572 | 0.00125198091156695 |
| Tulp3 | ENSMUSG00000001521 | 22158 | 0.509976154849441 | 3.33797946273027 | 4.89766000485975 | 0.0002666372300 | 0.00125198091156695 |
| Casq2 | ENSMUSG00000027861 | 12373 | 0.549288678080765 | 7.32277268990475 | 4.89452323357387 | 0.0002681606800 | 0.00125859355666274 |
| Hibch | ENSMUSG00000041426 | 227095 | -0.418362414649096 | 4.78273770428099 | -4.89272072530421 | 0.0002690402008 | 0.00126217958625801 |
| Pla2g7 | ENSMUSG00000023913 | 27226 | 0.95809788123694 | 2.27219261693931 | 4.8924747110622 | 0.0002691604739 | 0.00126220211969349 |
| Ubqln4 | ENSMUSG00000008604 | 94232 | -0.294674660618315 | 5.80904119249415 | -4.8907854706354 | 0.0002699878310 | 0.00126553901409931 |
| Tnfaip3 | ENSMUSG00000019850 | 21929 | 0.979396535895985 | 2.46179087968463 | 4.88991295931907 | 0.0002704162043 | 0.00126700365602683 |
| Samm50 | ENSMUSG00000022437 | 68653 | -0.415819168562021 | 6.38023401966004 | -4.8858650304703 | 0.0002724128553 | 0.00127581187749346 |
| R3hdm2 | ENSMUSG00000025404 | 71750 | -0.352730253312961 | 7.34432455734327 | -4.88544351058401 | 0.0002726216496 | 0.00127624293463056 |
| Timp4 | ENSMUSG00000030317 | 110595 | -1.52159262621523 | 3.63817661621983 | -4.88335339752561 | 0.0002736594178 | 0.0012805526955654 |
| Wbp2 | ENSMUSG00000034341 | 22378 | -0.416328505064543 | 4.60619117842328 | -4.88300585385786 | 0.0002738323753 | 0.00128081373318246 |
| Slc2a13 | ENSMUSG00000036298 | 239606 | 0.980958378634929 | 0.837397881541641 | 4.8813749892538 | 0.0002746455037 | 0.00128406757990382 |
| Pink1 | ENSMUSG00000028756 | 68943 | -0.600954505261798 | 8.75757602370595 | -4.88104098628581 | 0.0002748123426 | 0.00128429829723987 |
| Zyx | ENSMUSG00000029860 | 22793 | 0.570978809597416 | 4.71396470362763 | 4.87761393930405 | 0.0002765302911 | 0.0012917746208513 |
| Ndufa2 | ENSMUSG00000014294 | 17991 | -0.435796645528518 | 5.88122596891809 | -4.87601453693889 | 0.0002773358715 | 0.00129464497480526 |
| Lgalsl | ENSMUSG00000042363 | 216551 | -0.43484136134211 | 5.12801600964524 | -4.8758840789748 | 0.0002774016875 | 0.00129464497480526 |
| Bcs1l | ENSMUSG00000026172 | 66821 | -0.574401602279354 | 2.7780651993377 | -4.87560552686477 | 0.0002775422712 | 0.00129464497480526 |
| Ankrd13a | ENSMUSG00000041870 | 68420 | 0.328908566405655 | 4.54197046326312 | 4.87545454906923 | 0.0002776184997 | 0.00129464497480526 |
| Rxra | ENSMUSG00000015846 | 20181 | -0.341983707802765 | 5.6817646035664 | -4.87200304959115 | 0.0002793671023 | 0.00130224383600156 |
| Atxn2 | ENSMUSG00000042605 | 20239 | -0.285925537137325 | 5.95822524001751 | -4.87113568272548 | 0.0002798083245 | 0.0013037445846868 |
| Ppp1r3c | ENSMUSG00000067279 | 53412 | -0.670884557043903 | 7.12874246166391 | -4.87036561347311 | 0.0002802006588 | 0.00130446601859255 |
| Ssr3 | ENSMUSG00000027828 | 67437 | 0.30438431348372 | 6.41989592864068 | 4.87019364731956 | 0.0002802883499 | 0.00130446601859255 |
| Hspb1 | ENSMUSG00000004951 | 15507 | 0.517435361098952 | 7.18535078887615 | 4.87012930591343 | 0.0002803211671 | 0.00130446601859255 |
| Rdh13 | ENSMUSG00000008435 | 108841 | -0.437722821655681 | 3.70876217125954 | -4.86970660220789 | 0.0002805368649 | 0.00130491424302457 |
| Anln | ENSMUSG00000036777 | 68743 | 1.08644655441791 | 0.900645101271914 | 4.86920187512654 | 0.0002807946436 | 0.0013055577413446 |
| Rbp1 | ENSMUSG00000046402 | 19659 | 1.201728857258 | 1.62585530731571 | 4.86790406320883 | 0.0002814586037 | 0.00130808843436747 |
| Pnpla7 | ENSMUSG00000036833 | 241274 | -0.363342406247123 | 4.09386390534916 | -4.86284011068709 | 0.0002840649522 | 0.00131964045603912 |
| Zfp366 | ENSMUSG00000050919 | 238803 | -0.792585745612242 | 3.69440819535669 | -4.86250951643397 | 0.0002842359738 | 0.00131987401440346 |
| Cog1 | ENSMUSG00000018661 | 16834 | 0.365845335400073 | 4.14025095691056 | 4.861507332724 | 0.0002847550727 | 0.00132172301512006 |
| Cipc | ENSMUSG00000034157 | 217732 | -0.380341670949585 | 4.5581774739938 | -4.85883812858526 | 0.0002861424429 | 0.00132759891518536 |
| Jam2 | ENSMUSG00000053062 | 67374 | -0.638955304114119 | 4.92091640204013 | -4.85784358356057 | 0.0002866611701 | 0.00132944134518329 |
| Atp6v0e | ENSMUSG00000015575 | 11974 | 0.547125723642455 | 3.15744231375728 | 4.85340434930511 | 0.0002889884857 | 0.0013396663008037 |
| Ccdc28b | ENSMUSG00000028795 | 66264 | -0.488465233211822 | 3.04807783286407 | -4.8478407407129 | 0.0002919329969 | 0.00135274251346633 |
| Zdhhc13 | ENSMUSG00000030471 | 243983 | 0.706522042063083 | 1.29394687353081 | 4.84725654539236 | 0.0002922439812 | 0.00135360972824723 |
| Dtx4 | ENSMUSG00000039982 | 207521 | 0.538279453944287 | 3.23477250262012 | 4.84691738337467 | 0.0002924246848 | 0.00135387303283296 |
| Rbpj | ENSMUSG00000039191 | 19664 | 0.355597423042206 | 5.08883773529782 | 4.84628146888219 | 0.0002927638093 | 0.00135486926352561 |
| Opa3 | ENSMUSG00000052214 | 403187 | -0.27656231389453 | 5.50020673454525 | -4.84505835449457 | 0.0002934172279 | 0.00135731854387023 |
| Inpp5a | ENSMUSG00000025477 | 212111 | -0.385352540722123 | 6.37197178154885 | -4.84155951606923 | 0.0002952947693 | 0.00136542602531712 |
| Evc | ENSMUSG00000029122 | 59056 | -0.746623305746394 | 4.65744196038908 | -4.84098208011982 | 0.0002956058292 | 0.00136628639300383 |
| Hat1 | ENSMUSG00000027018 | 107435 | -0.499002420504367 | 2.96916017535571 | -4.84036346505933 | 0.0002959394494 | 0.00136725026268525 |
| Phospho2 | ENSMUSG00000027088 | 73373 | -0.506534725011337 | 3.46925522171296 | -4.83942932763808 | 0.0002964439722 | 0.00136900256067998 |
| Vtn | ENSMUSG00000017344 | 22370 | -0.961956283688712 | 3.62577675462608 | -4.83873354996128 | 0.0002968203384 | 0.00137016179037379 |
| Tnfaip1 | ENSMUSG00000017615 | 21927 | -0.353448052550307 | 5.09322280470054 | -4.83834232049804 | 0.0002970321835 | 0.00137056091117371 |
| Ostc | ENSMUSG00000041084 | 66357 | 0.518621785309555 | 3.75023425241301 | 4.83745053808173 | 0.0002975156577 | 0.00137221251261891 |
| Rabl3 | ENSMUSG00000022827 | 67657 | -0.653360420223279 | 2.26384647004443 | -4.83684492122894 | 0.0002978444548 | 0.00137314961426983 |
| Mrpl37 | ENSMUSG00000028622 | 56280 | -0.419768194401041 | 5.1571029650561 | -4.83521266030111 | 0.0002987325095 | 0.00137666317975281 |
| Arpc2 | ENSMUSG00000006304 | 76709 | 0.335095253466181 | 5.44430742693228 | 4.83385426173044 | 0.0002994736610 | 0.0013794970878148 |
| Prcc | ENSMUSG00000004895 | 94315 | -0.356441937442214 | 4.63673618898156 | -4.82929572254484 | 0.0003019748118 | 0.00139043246339863 |
| Mterf2 | ENSMUSG00000049038 | 74238 | -0.725488503588889 | 2.62416364302443 | -4.82723883704546 | 0.0003031104605 | 0.00139507387138176 |
| H2-DMa | ENSMUSG00000037649 | 14998 | 1.20124216292547 | 0.936106853158543 | 4.82546100734797 | 0.0003040956083 | 0.00139901897945182 |
| Brd7 | ENSMUSG00000031660 | 26992 | -0.339906638860411 | 5.80022771782201 | -4.825030924831 | 0.0003043344286 | 0.00139952866603788 |
| Zswim6 | ENSMUSG00000032846 | 67263 | 0.92880112551304 | 1.73033043813569 | 4.82420005021322 | 0.0003047963559 | 0.00140106348459627 |
| Tubb2a | ENSMUSG00000058672 | 22151 | 0.859225642383114 | 3.69048869352614 | 4.82087717471665 | 0.0003066510055 | 0.00140899627640659 |
| Arid1a | ENSMUSG00000007880 | 93760 | 0.32122297840827 | 5.94734948400889 | 4.82036297094647 | 0.0003069390511 | 0.00140972721338661 |
| Itga4 | ENSMUSG00000027009 | 16401 | 0.668492457166917 | 3.19259964635778 | 4.81985837559243 | 0.0003072219872 | 0.00141029550201337 |
| Zcrb1 | ENSMUSG00000022635 | 67197 | -0.332142049328528 | 5.61626402204994 | -4.81968222814935 | 0.0003073208202 | 0.00141029550201337 |
| Aktip | ENSMUSG00000031667 | 14339 | -0.314235375421113 | 5.04825877484817 | -4.81859871454333 | 0.0003079294849 | 0.00141208241616243 |
| Csgalnact1 | ENSMUSG00000036356 | 234356 | 1.0045514775966 | 1.70238393068547 | 4.81852920663247 | 0.0003079685737 | 0.00141208241616243 |
| Sh3glb1 | ENSMUSG00000037062 | 54673 | -0.332799887333806 | 7.92559409875843 | -4.8174331998937 | 0.0003085856107 | 0.00141431836963487 |
| Camk2b | ENSMUSG00000057897 | 12323 | 0.465867685988623 | 7.10928146292515 | 4.80984066588092 | 0.0003128954870 | 0.00143347048172171 |
| Dapk2 | ENSMUSG00000032380 | 13143 | 0.610943217927473 | 4.86873849366739 | 4.80773309965679 | 0.0003141028822 | 0.00143839908058269 |
| St13 | ENSMUSG00000022403 | 70356 | -0.263233288402954 | 6.35073138259459 | -4.80674193556017 | 0.0003146723767 | 0.00143981828364971 |
| Tmem88 | ENSMUSG00000045377 | 67020 | -1.17210157026551 | 1.34038143411846 | -4.80673523400641 | 0.0003146762308 | 0.00143981828364971 |
| Acsf3 | ENSMUSG00000015016 | 257633 | -0.437782259842192 | 2.78270582026148 | -4.80607826484875 | 0.0003150543016 | 0.00144094500913575 |
| Sgk3 | ENSMUSG00000025915 | 170755 | -0.567679349928123 | 2.89830942943852 | -4.80383884202718 | 0.0003163465841 | 0.00144519440473404 |
| Plpp7 | ENSMUSG00000051373 | 227721 | -0.413896401998749 | 3.89889059861616 | -4.80379745596059 | 0.0003163705181 | 0.00144519440473404 |
| S1pr3 | ENSMUSG00000067586 | 13610 | -1.03742621255083 | 2.23393259844298 | -4.80378099288015 | 0.0003163800393 | 0.00144519440473404 |
| Dram1 | ENSMUSG00000020057 | 71712 | 0.984589378099079 | 1.33677335065771 | 4.80308777551305 | 0.0003167812256 | 0.0014464225468526 |
| Smim19 | ENSMUSG00000031534 | 102032 | -0.426029539247309 | 3.5280181699369 | -4.80284246455948 | 0.0003169233210 | 0.00144646714912975 |
| Qk | ENSMUSG00000062078 | 19317 | -0.329398097027685 | 8.93375718950216 | -4.80233395926646 | 0.0003172180814 | 0.00144720820033239 |
| Tmem25 | ENSMUSG00000002032 | 71687 | -0.649202998300659 | 2.0307378425898 | -4.80195870953184 | 0.0003174357812 | 0.00144759721514564 |
| Nfyc | ENSMUSG00000032897 | 18046 | -0.322241974466137 | 4.99059094673258 | -4.80074970008777 | 0.0003181382387 | 0.00145000493855441 |
| Gimap6 | ENSMUSG00000047867 | 231931 | -0.805261446019397 | 3.27624651607679 | -4.80059358974225 | 0.0003182290593 | 0.00145000493855441 |
| Vwa7 | ENSMUSG00000007030 | 27762 | -0.436600052831882 | 3.59694039957167 | -4.80029875060702 | 0.0003184006618 | 0.00145018234757746 |
| Ptp4a1 | ENSMUSG00000117310 | 19243 | -0.360659938174556 | 6.31346978854997 | -4.79947542309784 | 0.0003188803640 | 0.00145176229031036 |
| Ms4a6b | ENSMUSG00000024677 | 69774 | 1.99845786280334 | 0.411891620520774 | 4.79572679758585 | 0.0003210739551 | 0.00146114046795086 |
| Dnmt1 | ENSMUSG00000004099 | 13433 | 0.51230424373084 | 3.80152798148492 | 4.79204695627702 | 0.0003232425265 | 0.00147039702767127 |
| Mob3a | ENSMUSG00000003348 | 208228 | 0.821860196624872 | 1.95556089694631 | 4.78880706571595 | 0.0003251644085 | 0.00147852418873211 |
| Ptk2b | ENSMUSG00000059456 | 19229 | 1.11938475493279 | 1.09017894199062 | 4.78838649965977 | 0.0003254147534 | 0.00147904726403001 |
| Msh3 | ENSMUSG00000014850 | 17686 | -0.458612437549269 | 3.94164251474004 | -4.78791763885071 | 0.0003256940818 | 0.00147970158316808 |
| Nup93 | ENSMUSG00000032939 | 71805 | 0.464197653084756 | 2.94779260726742 | 4.78657646693668 | 0.0003264944724 | 0.00148272167770915 |
| Vldlr | ENSMUSG00000024924 | 22359 | -0.37017078732657 | 7.32042625931442 | -4.78494648773647 | 0.0003274699663 | 0.0014865341370853 |
| Prcp | ENSMUSG00000061119 | 72461 | 0.698010978234813 | 2.53028877684201 | 4.78138617213611 | 0.0003296112378 | 0.00149512756202394 |
| Ppp1cb | ENSMUSG00000014956 | 19046 | -0.40174768375098 | 7.77954119320872 | -4.78134418406081 | 0.0003296365771 | 0.00149512756202394 |
| Lamtor2 | ENSMUSG00000028062 | 83409 | -0.422639775954288 | 3.6391981909087 | -4.78085346801233 | 0.0003299328676 | 0.00149558897687686 |
| Scube2 | ENSMUSG00000007279 | 56788 | 0.613758311104517 | 3.68245635403332 | 4.78072257129815 | 0.0003300119487 | 0.00149558897687686 |
| Kyat1 | ENSMUSG00000039648 | 70266 | -0.655595967456879 | 2.60735247794512 | -4.78021145231051 | 0.0003303209290 | 0.00149636886670952 |
| Atp5l | ENSMUSG00000038717 | 27425 | -0.477050851330374 | 6.71787011233152 | -4.77977593245001 | 0.0003305844450 | 0.00149694224082876 |
| Gucy1b1 | ENSMUSG00000028005 | 54195 | -0.680349165801444 | 2.49311680198138 | -4.77941385590249 | 0.0003308036894 | 0.00149731475346079 |
| Sfxn3 | ENSMUSG00000025212 | 94280 | 0.70014852555199 | 2.64493368973709 | 4.77840984139902 | 0.0003314124290 | 0.0014994491978056 |
| Irak3 | ENSMUSG00000020227 | 73914 | 0.722968883320308 | 2.49874087211193 | 4.77583961033157 | 0.0003329760747 | 0.00150590048531288 |
| Swi5 | ENSMUSG00000044627 | 72931 | -0.380137156412846 | 5.09803086730638 | -4.7748994213588 | 0.0003335499652 | 0.00150787207192828 |
| Mdn1 | ENSMUSG00000058006 | 100019 | 0.359415371042002 | 4.49866440763357 | 4.77212401928504 | 0.0003352500598 | 0.00151493113038343 |
| Hist1h3b | ENSMUSG00000069267 | 319150 | 1.71042697160839 | 0.665665538907474 | 4.77177267333075 | 0.0003354659196 | 0.00151528015182886 |
| Epb41l1 | ENSMUSG00000027624 | 13821 | 0.518869337219878 | 3.41334816163921 | 4.77139067936735 | 0.0003357007724 | 0.00151571464015147 |
| Adgrl4 | ENSMUSG00000039167 | 170757 | -0.53788698681045 | 4.74084016044637 | -4.77080815507989 | 0.0003360592412 | 0.00151663817421975 |
| Hmgcs1 | ENSMUSG00000093930 | 208715 | -0.3364325579276 | 4.43403655573807 | -4.77060750089214 | 0.0003361828100 | 0.00151663817421975 |
| Fnbp1 | ENSMUSG00000075415 | 14269 | 0.510124916569304 | 4.43047977711722 | 4.7683807531946 | 0.0003375572742 | 0.00152221062895577 |
| Gbe1 | ENSMUSG00000022707 | 74185 | -0.436829234558572 | 6.17644187185726 | -4.76712133996193 | 0.0003383372284 | 0.00152509865741074 |
| Asb15 | ENSMUSG00000029685 | 78910 | -0.684582450145521 | 4.50953542421215 | -4.76651449454795 | 0.0003387137146 | 0.00152616637056474 |
| Ushbp1 | ENSMUSG00000034911 | 234395 | -0.82438057242785 | 3.3184310277697 | -4.76430782873319 | 0.0003400863940 | 0.00153171997238047 |
| Eif4g3 | ENSMUSG00000028760 | 230861 | -0.265272869684427 | 6.51604450965754 | -4.76369461656799 | 0.0003404688724 | 0.00153281105626585 |
| Sympk | ENSMUSG00000023118 | 68188 | -0.317347590269037 | 6.09595762228108 | -4.76194885420424 | 0.0003415602003 | 0.00153709121029386 |
| Asb18 | ENSMUSG00000067081 | 208372 | -0.46890416502839 | 4.08193114390321 | -4.76027959063108 | 0.0003426070994 | 0.00154116798526938 |
| Ehhadh | ENSMUSG00000022853 | 74147 | -1.00176305576475 | 1.63540857379631 | -4.75928961909759 | 0.0003432295436 | 0.00154333284301505 |
| Abi3bp | ENSMUSG00000035258 | 320712 | 0.888632139522241 | 5.41839524887253 | 4.7544823666043 | 0.0003462688174 | 0.00155579946237369 |
| A430005L14Rik | ENSMUSG00000047613 | 97159 | -0.456300391346481 | 2.89730753707232 | -4.75445418613772 | 0.0003462867159 | 0.00155579946237369 |
| Mrps28 | ENSMUSG00000040269 | 66230 | -0.427742633943547 | 4.09702286414906 | -4.751070391959 | 0.0003484428815 | 0.00156484352415651 |
| Nlgn2 | ENSMUSG00000051790 | 216856 | 0.820580900049803 | 2.39842770288765 | 4.7507784838641 | 0.0003486295374 | 0.00156503879826655 |

| | | | | | | | |
|---|---|---|---|---|---|---|---|
| Slc12a2 | ENSMUSG00000024597 | 20496 | -0.342862585944385 | 5.59601770501226 | -4.75030607132064 | 0.0003489318337 | 0.00156575282202493 |
| Il1rl1 | ENSMUSG00000026069 | 17082 | 1.39390687505812 | 0.449160399547754 | 4.7491133515355 | 0.0003496962637 | 0.00156853912971333 |
| Rnf11 | ENSMUSG00000028557 | 29864 | -0.358893155685894 | 6.17884300313161 | -4.7474305528853 | 0.0003507777462 | 0.00157274468579924 |
| Zbtb16 | ENSMUSG00000066687 | 235320 | -0.524805261432492 | 6.31561554017605 | -4.74678378495691 | 0.0003511943259 | 0.00157396686203782 |
| Sec23b | ENSMUSG00000027429 | 27054 | 0.470599166637281 | 3.2946306210646 | 4.7460629058051 | 0.0003516592447 | 0.00157540459194403 |
| Mrps24 | ENSMUSG00000020477 | 64660 | -0.38651107137233 | 4.97517580871394 | -4.74394795280268 | 0.0003530269338 | 0.00158088382392921 |
| Ptcd2 | ENSMUSG00000021650 | 68927 | -0.443697165103675 | 3.77439392506854 | -4.7421307449216 | 0.0003542064831 | 0.00158551640737519 |
| Tnrc18 | ENSMUSG00000039477 | 231861 | 0.372980081737672 | 5.50438409541656 | 4.74085118306637 | 0.0003550394973 | 0.00158859465623046 |
| Gse1 | ENSMUSG00000031822 | 382034 | 0.386288015394412 | 5.02277474247333 | 4.73964309135461 | 0.0003558278474 | 0.0015914706221703 |
| Ulk2 | ENSMUSG00000004798 | 29869 | -0.301712399727213 | 6.21371339003644 | -4.7376881460929 | 0.0003571074095 | 0.00159654032470857 |
| Dchs1 | ENSMUSG00000036862 | 233651 | 0.730329158247028 | 4.29003742271819 | 4.73462604909893 | 0.0003591212264 | 0.00160488721434716 |
| Nf1 | ENSMUSG00000020716 | 18015 | 0.332699766550761 | 5.41456649863244 | 4.73422223051387 | 0.0003593876788 | 0.00160542162530423 |
| Shb | ENSMUSG00000044813 | 230126 | 0.880739641218374 | 1.42239873251213 | 4.73373621556579 | 0.0003597086387 | 0.00160619899102174 |
| Ubac2 | ENSMUSG00000041765 | 68889 | -0.387082023250372 | 3.56549436265719 | -4.73264126389743 | 0.0003604328242 | 0.00160877550109172 |
| Mob1a | ENSMUSG00000043131 | 232157 | 0.532857411592493 | 4.44988536771364 | 4.73163927316574 | 0.0003610968510 | 0.0016105391978627 |
| Prelp | ENSMUSG00000041577 | 116847 | 0.961607163591918 | 5.77961034890101 | 4.7315494146493 | 0.0003611564629 | 0.0016105391978627 |
| 2410004B18Rik | ENSMUSG00000036873 | 66421 | -0.448226758486 | 3.23388020016412 | -4.73131502129625 | 0.0003613120067 | 0.0016105391978627 |
| Xxylt1 | ENSMUSG00000047434 | 268880 | 0.561969569009386 | 2.44582260169199 | 4.73115639113806 | 0.0003614173133 | 0.0016105391978627 |
| Tmed3 | ENSMUSG00000032353 | 66111 | 0.634888464175797 | 3.3301108491819 | 4.73034124995851 | 0.0003619589467 | 0.0016122955368149 |
| Trim2 | ENSMUSG00000027993 | 80890 | 0.679698444613375 | 2.3326514954312 | 4.728918200081 | 0.0003629065322 | 0.0016158579650369 |
| Wdr92 | ENSMUSG00000078970 | 103784 | -0.616023148352442 | 2.0805731248406 | -4.72824023396584 | 0.0003633588827 | 0.00161721333348076 |
| Fbn2 | ENSMUSG00000024598 | 14119 | 1.16918710789235 | 2.36153997168591 | 4.72637760969349 | 0.0003646046681 | 0.00162209752840964 |
| Serpinf1 | ENSMUSG00000000753 | 20317 | 0.824358603220096 | 5.70141225579397 | 4.72594340015766 | 0.0003648957177 | 0.00162273193292722 |
| Fundc2 | ENSMUSG00000031198 | 67391 | -0.422522723081515 | 6.0054741202561 | -4.72393808527752 | 0.0003662430032 | 0.0016280611097591 |
| Dusp13 | ENSMUSG00000021768 | 27389 | -0.502712064472513 | 4.09586322740036 | -4.71959658546806 | 0.0003691775659 | 0.00164043901341073 |
| Hyou1 | ENSMUSG00000032115 | 12282 | 0.482087838498146 | 5.02386157979251 | 4.71918262074663 | 0.0003694586480 | 0.00164102091889425 |
| Hs2st1 | ENSMUSG00000040151 | 23908 | 0.368777412525011 | 3.6641764596831 | 4.71697679432367 | 0.0003709601486 | 0.00164702087116801 |
| Gm34455 | ENSMUSG00000118107 | 102637715 | -1.81006330347362 | 0.570305818638866 | -4.71623021414774 | 0.0003714697752 | 0.00164861393141748 |
| Zfp612 | ENSMUSG00000044676 | 234725 | -0.493314247748487 | 2.24316675974605 | -4.71279116921201 | 0.0003738267039 | 0.00165840085268281 |
| Itga10 | ENSMUSG00000090210 | 213119 | 1.28684633528038 | 0.645995016712033 | 4.71238810736009 | 0.0003741039527 | 0.00165895752859059 |
| Srebf2 | ENSMUSG00000022463 | 20788 | 0.412369292800167 | 4.27487405221575 | 4.71014397687192 | 0.0003756514939 | 0.00166514455809879 |
| Gtf2a2 | ENSMUSG00000033543 | 235459 | -0.355276840480944 | 3.92370513873098 | -4.70692786249537 | 0.0003778808853 | 0.00167397148698511 |
| Slc35e4 | ENSMUSG00000048807 | 103710 | 0.52342632171543 | 3.16052704474088 | 4.7068297641529 | 0.0003779491016 | 0.00167397148698511 |
| Sft2d2 | ENSMUSG00000040848 | 108735 | 0.396012042155828 | 3.7611851140047 | 4.70537566288253 | 0.0003789617618 | 0.00167777684043352 |
| Uba52 | ENSMUSG00000090137 | 22186 | -0.357612955819904 | 7.64666013143616 | -4.70451779095117 | 0.0003795605163 | 0.0016793979634221 |
| Slc36a4 | ENSMUSG00000043885 | 234967 | 0.555294401102783 | 2.34758902701967 | 4.70441088600429 | 0.0003796351996 | 0.0016793979634221 |
| Myct1 | ENSMUSG00000046916 | 68632 | -0.914006618736609 | 2.71009017280776 | -4.702631013159 | 0.0003808808503 | 0.00168412697261783 |
| Bag1 | ENSMUSG00000028416 | 12017 | 0.44308543658569 | 7.36275664582408 | 4.70244346864747 | 0.0003810123504 | 0.00168412697261783 |
| Syngap1 | ENSMUSG00000067629 | 240057 | 1.09494069824486 | 1.48050493731642 | 4.70191717758657 | 0.0003813816200 | 0.00168507780453705 |
| Lsm3 | ENSMUSG00000034192 | 67678 | -0.54830511194087 | 2.9922182844632 | -4.70111001575069 | 0.0003819486817 | 0.00168690143049271 |
| Mical3 | ENSMUSG00000051586 | 194401 | 0.320722970257943 | 6.62736652132665 | 4.70083375438723 | 0.0003821429664 | 0.00168707785392457 |
| Tmem164 | ENSMUSG00000047045 | 209497 | 0.659895506979637 | 1.86032495058372 | 4.70022247567415 | 0.0003825732208 | 0.00168829546905503 |
| Daglb | ENSMUSG00000039206 | 231871 | 0.396064133314012 | 3.9630394069493 | 4.69972872301111 | 0.0003829211197 | 0.00168862290099237 |
| Ache | ENSMUSG00000023328 | 11423 | 0.610121928148018 | 3.37633940518459 | 4.69967870857236 | 0.0003829563783 | 0.00168862290099237 |
| Scpep1 | ENSMUSG00000000278 | 74617 | 0.62123809956175 | 3.0406901744502 | 4.69826500533339 | 0.0003839543852 | 0.00169234088115753 |
| Malsu1 | ENSMUSG00000029815 | 75593 | -0.512542200981759 | 3.46238951421831 | -4.69780883952637 | 0.0003842769912 | 0.00169308012553973 |
| Ivns1abp | ENSMUSG00000023150 | 117198 | -0.369993620194929 | 6.69930534385107 | -4.69711283739465 | 0.0003847697536 | 0.00169456816169551 |
| Ctsb | ENSMUSG00000021939 | 13030 | 0.36641929825346 | 7.2251206815837 | 4.69502247684175 | 0.0003862536471 | 0.00170041829113678 |
| Tigd4 | ENSMUSG00000047819 | 403175 | -0.626666641357554 | 2.5386615986855 | -4.69479994744235 | 0.0003864119638 | 0.00170043042555798 |
| Ppp2r5e | ENSMUSG00000021051 | 26932 | -0.300284437332447 | 4.8913362311365 | -4.69444462740768 | 0.0003866648928 | 0.00170085872970558 |
| Grb10 | ENSMUSG00000020176 | 14783 | 0.40664790956788 | 6.20091818546013 | 4.69258940356416 | 0.0003879882930 | 0.00170599357676278 |
| Wfikkn2 | ENSMUSG00000044177 | 278507 | -0.731771733937638 | 2.24040231761503 | -4.69068256617222 | 0.0003893534022 | 0.00171130761545632 |
| Myot | ENSMUSG00000024471 | 58916 | -0.455521638196252 | 10.1843523060251 | -4.68461787655833 | 0.0003937283008 | 0.00172984085862898 |
| Zswim7 | ENSMUSG00000014243 | 69747 | -0.733368102344247 | 2.45637160555074 | -4.68398922587795 | 0.0003941846955 | 0.00173115022387372 |
| Setbp1 | ENSMUSG00000024548 | 240427 | 0.385459014031861 | 4.62274083616658 | 4.68325364536591 | 0.0003947194162 | 0.00173280238514399 |
| Cmklr1 | ENSMUSG00000042190 | 14747 | 0.825903243980344 | 3.68412360147004 | 4.68239789225597 | 0.0003953424404 | 0.00173458088445751 |
| Xdh | ENSMUSG00000024066 | 22436 | -0.59896539083165 | 5.36769505823367 | -4.68226138977995 | 0.0003954419141 | 0.00173458088445751 |
| Plpbp | ENSMUSG00000031485 | 114863 | -0.424826239244946 | 4.71086766159317 | -4.6792439362483 | 0.0003976474623 | 0.00174355572020058 |
| ND5 | ENSMUSG00000064367 | 17721 | -0.635785975902312 | 12.6380489156593 | -4.67716819150475 | 0.0003991720813 | 0.00174953890173841 |
| Tbx18 | ENSMUSG00000032419 | 76365 | 0.811645860490457 | 1.45118288859015 | 4.67684092608959 | 0.0003994130073 | 0.00174989321962271 |
| Myod1 | ENSMUSG00000009471 | 17927 | 1.10756344458305 | 3.25327618046527 | 4.67464608220889 | 0.0004010327028 | 0.0017562854467937 |
| 1600012H06Rik | ENSMUSG00000050088 | 67912 | 0.550986078124944 | 2.6700359117726 | 4.67091337429511 | 0.0004038029132 | 0.00176712487416821 |
| Shc2 | ENSMUSG00000020312 | 216148 | 1.27751721118299 | 1.3351043803546 | 4.6708755083417 | 0.0004038311166 | 0.00176712487416821 |
| Robo4 | ENSMUSG00000032125 | 74144 | -0.87343992508562 | 3.41556159637113 | -4.66866002298765 | 0.0004054848128 | 0.00177309082394616 |
| Plin3 | ENSMUSG00000024197 | 66905 | -0.617515862259287 | 4.64576414598014 | -4.6686144587262 | 0.0004055188967 | 0.00177309082394616 |
| Cdip1 | ENSMUSG00000004071 | 66626 | -0.413443131201087 | 5.00921830549832 | -4.66773647016243 | 0.0004061762465 | 0.00177525491838773 |
| Zfp654 | ENSMUSG00000047141 | 72020 | -0.400381341960208 | 3.58778319163859 | -4.66736992969093 | 0.0004064510017 | 0.00177574576334694 |
| Rapgef3 | ENSMUSG00000022469 | 223864 | -1.0987661669823 | 2.89064994822937 | -4.66591499101769 | 0.0004075435094 | 0.00177980747165013 |
| Klhl30 | ENSMUSG00000026308 | 70788 | 0.363214919363227 | 5.0747832539629 | 4.66329994919294 | 0.0004095147856 | 0.00178770212548167 |
| Tppp | ENSMUSG00000021573 | 72948 | -0.770212601562822 | 2.90284650322524 | -4.66159556993416 | 0.0004108048971 | 0.00179261809592107 |
| Gadd45gip1 | ENSMUSG00000033751 | 102060 | -0.386363362333908 | 5.06876953341386 | -4.66073193458213 | 0.0004114602224 | 0.00179476124970811 |
| Ebag9 | ENSMUSG00000022339 | 55960 | -0.555710972937054 | 3.28923917062166 | -4.65836981246655 | 0.0004132581327 | 0.00180188458289873 |
| Arhgef15 | ENSMUSG00000052921 | 442801 | -0.799707770028252 | 4.2126209293415 | -4.65792967258359 | 0.0004135940398 | 0.00180263016328634 |
| Acy1 | ENSMUSG00000023262 | 109652 | -0.875270699850696 | 1.28609401955984 | -4.65746061329969 | 0.0004139523289 | 0.00180347266133064 |
| Aldh1b1 | ENSMUSG00000035561 | 72535 | 1.21190541430772 | 0.781645209021962 | 4.65682412893588 | 0.0004144390189 | 0.00180487367181038 |
| Actn1 | ENSMUSG00000015143 | 109711 | 0.676766155117548 | 4.4394940019153 | 4.65457171519404 | 0.0004161660988 | 0.00181167328827529 |
| Skap2 | ENSMUSG00000059182 | 54353 | 0.600333966015647 | 2.27437160498617 | 4.654033609593 | 0.0004165798030 | 0.00181275231980549 |
| Ntn1 | ENSMUSG00000020902 | 18208 | 0.711170235428335 | 3.54050636959219 | 4.65327106469086 | 0.0004171667891 | 0.00181458423104778 |
| Snai3 | ENSMUSG00000006587 | 30927 | -0.77996995403866 | 3.00554049506038 | -4.65227048562594 | 0.0004179383068 | 0.00181721703740126 |
| Hrc | ENSMUSG00000038239 | 15464 | -0.474428136421585 | 9.07694239954456 | -4.65158846839657 | 0.0004184650372 | 0.00181878382580049 |
| Fuom | ENSMUSG00000025466 | 69064 | -0.533591444653744 | 2.90182851111978 | -4.64863918519919 | 0.0004207507300 | 0.00182799134766074 |
| Pex10 | ENSMUSG00000029047 | 668173 | -0.685222772198905 | 2.10033296068554 | -4.64819493913975 | 0.0004210961390 | 0.00182821236848301 |
| Arhgap28 | ENSMUSG00000024043 | 268970 | 1.08255772097813 | 0.90947010503973 | 4.64814356384943 | 0.0004211361031 | 0.00182821236848301 |
| Sec16b | ENSMUSG00000026589 | 89867 | 1.19020392127577 | 1.23181178549221 | 4.64414103095037 | 0.0004242617179 | 0.00184104995601418 |
| Rbfox2 | ENSMUSG00000033565 | 93686 | 0.424590831096053 | 6.30578839402603 | 4.64374309557076 | 0.0004245737791 | 0.00184167300796342 |
| Mrps18b | ENSMUSG00000024436 | 66973 | -0.45543895879569 | 3.9852367048528 | -4.64249718934334 | 0.0004255523572 | 0.00184518556804213 |
| Ppa2 | ENSMUSG00000028013 | 74776 | -0.353696553858922 | 4.3295932210386 | -4.64228088574767 | 0.0004257224873 | 0.00184519132005644 |
| Trim3 | ENSMUSG00000036989 | 55992 | 0.370014166706601 | 3.70783102975765 | 4.64131651715235 | 0.0004264818531 | 0.00184704082513521 |
| Ehmt2 | ENSMUSG00000013787 | 110147 | 0.382910628323788 | 4.8532252609423 | 4.64116908887328 | 0.0004265980649 | 0.00184704082513521 |
| Lsm14b | ENSMUSG00000039108 | 241846 | -0.378953751327352 | 4.8977039543557 | -4.64109545077582 | 0.0004266561232 | 0.00184704082513521 |
| Ptpn13 | ENSMUSG00000034573 | 19249 | 0.741486611484473 | 3.14216218620368 | 4.63758364968672 | 0.0004294344410 | 0.00185833249212358 |
| Smc2 | ENSMUSG00000028312 | 14211 | 0.906554778202859 | 1.34279152258213 | 4.63707917499445 | 0.0004298350856 | 0.00185933016275581 |
| Sigmar1 | ENSMUSG00000036078 | 18391 | 0.514363231995076 | 2.47879390123649 | 4.63615285391614 | 0.0004305717608 | 0.00186178003064146 |
| Asl | ENSMUSG00000025533 | 109900 | 0.613989224295114 | 1.62410620387483 | 4.63503978877003 | 0.0004314586768 | 0.00186487734170892 |
| Cabin1 | ENSMUSG00000020196 | 104248 | 0.32818022158114 | 4.22441073268096 | 4.6339969694346 | 0.0004322913353 | 0.00186711581611609 |
| Tnpo1 | ENSMUSG00000009470 | 238799 | 0.272304205806476 | 5.47550726161209 | 4.63396337188911 | 0.0004323181896 | 0.00186711581611609 |
| Cic | ENSMUSG00000005442 | 71722 | 0.335136708043231 | 5.57729203278493 | 4.6333141419949 | 0.0004328374526 | 0.00186862014012378 |
| Nedd1 | ENSMUSG00000019988 | 17997 | -0.352159584381875 | 4.14532664646458 | -4.63259132309517 | 0.0004334163327 | 0.00187038055000735 |
| St8sia6 | ENSMUSG00000003418 | 241230 | -0.736469054705303 | 1.46837272768884 | -4.62425042559181 | 0.0004401545333 | 0.00189870923599698 |
| Pex6 | ENSMUSG00000002763 | 224824 | -0.517851312671538 | 3.39641810930087 | -4.62304877731509 | 0.0004411341838 | 0.00190218452211362 |
| Mamdc2 | ENSMUSG00000033207 | 71738 | 1.07842218468171 | 2.84120150907811 | 4.62267761147892 | 0.0004414372348 | 0.00190274069942084 |
| Sh3bgrl | ENSMUSG00000031246 | 56726 | 0.498693452486794 | 3.84546963447291 | 4.6222760651191 | 0.0004417653336 | 0.00190340436030095 |
| Hdac4 | ENSMUSG00000026313 | 208727 | 0.445371887460855 | 5.43825788088463 | 4.62090850964195 | 0.0004428846418 | 0.00190689204436909 |
| Herc4 | ENSMUSG00000020064 | 67345 | 0.27976726485499 | 4.25358364010868 | 4.62067338567559 | 0.0004430773801 | 0.00190689204436909 |
| Cep164 | ENSMUSG00000043987 | 214552 | 0.596870050748026 | 2.2168192734072 | 4.62064806410531 | 0.0004430981422 | 0.00190689204436909 |
| Yars | ENSMUSG00000028811 | 107271 | -0.390686638429925 | 4.42461501343429 | -4.6183117449177 | 0.0004450181106 | 0.00191440101036437 |
| Sparc | ENSMUSG00000018593 | 20692 | 0.73061304618717 | 8.27870599459549 | 4.61512449091192 | 0.0004476512643 | 0.00192497087740808 |
| Dock11 | ENSMUSG00000031093 | 75974 | 0.560855265550166 | 3.30776869120271 | 4.61468509176221 | 0.0004480155370 | 0.00192559725415856 |
| Shroom3 | ENSMUSG00000029381 | 27428 | 0.880005487819415 | 0.957250809059761 | 4.61452389906405 | 0.0004481492465 | 0.00192559725415856 |
| Eif4e | ENSMUSG00000028156 | 13684 | -0.32085852507141 | 5.3128586227287 | -4.61332801718441 | 0.0004491425229 | 0.00192910684434486 |
| Cst6 | ENSMUSG00000024846 | 73720 | 0.690773045597612 | 1.77359883098197 | 4.61112873503252 | 0.0004509751557 | 0.00193587345319077 |
| Ranbp6 | ENSMUSG00000074909 | 240614 | -0.530524825416122 | 3.53559908388427 | -4.61101259173952 | 0.0004510721512 | 0.00193587345319077 |
| Fkbp7 | ENSMUSG00000002732 | 14231 | 0.515492392377292 | 2.13622336017877 | 4.61041420953343 | 0.0004515722243 | 0.00193725902035007 |
| Dbnl | ENSMUSG00000020476 | 13169 | 0.383871980895818 | 4.13947451807147 | 4.60421400862552 | 0.0004567876769 | 0.0019588646906104 |
| Wipf1 | ENSMUSG00000075284 | 215280 | 0.60132759911226 | 4.05939950153185 | 4.60326056925353 | 0.0004575951980 | 0.00196155808211516 |
| Cavin3 | ENSMUSG00000037060 | 109042 | 0.50582411413913 | 3.83507441298421 | 4.60235819994485 | 0.0004583608254 | 0.00196406984843397 |

| | | | | | | | |
|---|---|---|---|---|---|---|---|
| Dennd1b | ENSMUSG00000056268 | 329260 | -0.277999306159527 | 4.33879767104148 | -4.60174547211039 | 0.0004588814581 | 0.0019655302581916 |
| Rilp | ENSMUSG00000038195 | 280408 | -0.468834138729164 | 4.47703833144655 | -4.600495988252 | 0.0004599450369 | 0.00196887440744218 |
| Lims2 | ENSMUSG00000024395 | 225341 | -0.669967012241745 | 3.5995015413674 | -4.60040517836034 | 0.0004600224349 | 0.00196887440744218 |
| Cbx4 | ENSMUSG00000039989 | 12418 | 0.717258850342778 | 2.38097282463714 | 4.59977940966231 | 0.0004605561494 | 0.00197038718962778 |
| Nfu1 | ENSMUSG00000029993 | 56748 | -0.301056019767866 | 4.43064329688406 | -4.59944433114731 | 0.0004608421992 | 0.00197083962447767 |
| Hspa12b | ENSMUSG00000074793 | 72630 | -0.955763720366431 | 3.43550178745486 | -4.59904386341828 | 0.0004611843116 | 0.00197153136918261 |
| Ilrun | ENSMUSG00000056692 | 224647 | -0.337964876584675 | 7.66228606272774 | -4.59794514162836 | 0.0004621242800 | 0.00197477736716221 |
| Mrpl4 | ENSMUSG00000003299 | 66163 | -0.408345450676395 | 4.17596331866951 | -4.5963847249004 | 0.0004634626409 | 0.00197972259799602 |
| Tceal9 | ENSMUSG00000042712 | 22381 | 0.436610806735581 | 3.27870329923229 | 4.5927605658768 | 0.0004665865565 | 0.00199173139925348 |
| Myd88 | ENSMUSG00000032508 | 17874 | 0.673285502094367 | 2.52120885208298 | 4.59270065752445 | 0.0004666383783 | 0.00199173139925348 |
| Lrp12 | ENSMUSG00000022305 | 239393 | 0.732195267030624 | 2.20480078207432 | 4.59163916272126 | 0.0004675575786 | 0.00199487583604648 |
| Nexn | ENSMUSG00000039103 | 68810 | -0.43388960624773 | 8.66739407834772 | -4.5900290878423 | 0.0004689553979 | 0.0020000590928128 |
| Mcc | ENSMUSG00000071856 | 328949 | -0.832007493653129 | 4.12498703988074 | -4.58793822671233 | 0.0004707770761 | 0.00200704532717247 |
| Calcrl | ENSMUSG00000059588 | 54598 | -0.560402426627995 | 4.27408592941635 | -4.58680894951653 | 0.0004717640099 | 0.00201046876923325 |
| Podxl2 | ENSMUSG00000033152 | 319655 | -0.340932773071117 | 3.79912442464741 | -4.58658593379217 | 0.0004719591679 | 0.00201051662678602 |
| Crispld2 | ENSMUSG00000031825 | 78892 | 0.605581088856738 | 4.2773162887729 | 4.58558746674831 | 0.0004728339367 | 0.00201345841945093 |
| Spryd3 | ENSMUSG00000036966 | 223918 | 0.48444612051635 | 2.56176042266529 | 4.58436286348246 | 0.0004739091165 | 0.00201725099419184 |
| Sfxn1 | ENSMUSG00000021474 | 14057 | 0.759704813756183 | 1.62478944940699 | 4.58249442520921 | 0.0004755544469 | 0.00202346658620238 |
| Gpbp1l1 | ENSMUSG00000034042 | 77110 | -0.251286592301476 | 5.09703399344831 | -4.58100961185538 | 0.0004768661732 | 0.00202825842018475 |
| Timm50 | ENSMUSG00000003438 | 66525 | -0.341987139341786 | 3.89036856841307 | -4.57857612520964 | 0.0004790240826 | 0.00203664420351697 |
| Cacng6 | ENSMUSG00000078815 | 54378 | -0.400697311569477 | 4.2816967743509 | -4.57655739635106 | 0.0004808218660 | 0.00204349293085041 |
| Tcaim | ENSMUSG00000046603 | 382117 | -0.468907951633272 | 4.79988971260191 | -4.57617122059867 | 0.0004811665701 | 0.0020438874941087 |
| Nudt7 | ENSMUSG00000031767 | 67528 | -0.529626189110173 | 3.91620170997207 | -4.57603450363742 | 0.0004812886661 | 0.0020438874941087 |
| Oaf | ENSMUSG00000032014 | 102644 | 0.695990779398182 | 3.11024015014757 | 4.57470650815451 | 0.0004824763105 | 0.0020481353594597 |
| Bop1 | ENSMUSG00000022557 | 12181 | 0.449955346441397 | 3.72461751405219 | 4.57405379786676 | 0.0004830611487 | 0.00204982197855587 |
| Deptor | ENSMUSG00000022419 | 97998 | 0.411613172963227 | 4.25302680292844 | 4.57320494959641 | 0.0004838228263 | 0.00205167524814218 |
| Car4 | ENSMUSG00000000805 | 12351 | -1.45200727507478 | 1.41814572050762 | -4.57314877403178 | 0.0004838732768 | 0.00205167524814218 |
| Fhl2 | ENSMUSG00000008136 | 14200 | 0.475302590520836 | 3.81180857640963 | 4.57286774659035 | 0.0004841257456 | 0.00205194979659548 |
| Kpna2 | ENSMUSG00000018362 | 16647 | 0.498096552714237 | 2.7700578809938 | 4.57247568405086 | 0.0004844781934 | 0.00205264772563835 |
| Mgat4b | ENSMUSG00000036620 | 103534 | 0.379910837630557 | 4.33846069936267 | 4.57135274283572 | 0.0004854891396 | 0.0020561339732725 |
| Immp1l | ENSMUSG00000042670 | 66541 | -0.357199221553281 | 3.66017519351433 | -4.56915362335041 | 0.0004874752570 | 0.00206374594690863 |
| Nol7 | ENSMUSG00000063200 | 70078 | -0.356159204920152 | 5.59805714523416 | -4.56789867470602 | 0.0004886124176 | 0.00206758934846829 |
| Zfp358 | ENSMUSG00000047264 | 140482 | -0.35712053530319 | 3.90476973765956 | -4.56773448316834 | 0.0004887614011 | 0.00206758934846829 |
| Il17d | ENSMUSG00000050222 | 239114 | 0.905617061909098 | 1.19290555478416 | 4.56752010287926 | 0.0004889559954 | 0.00206761237397232 |
| Stc2 | ENSMUSG00000020303 | 20856 | 0.934606804822725 | 1.5811445409376 | 4.56570157971378 | 0.0004906099056 | 0.00207380389744558 |
| Ap1s2 | ENSMUSG00000031367 | 108012 | 0.288028830977648 | 5.17667547700789 | 4.56296486563879 | 0.0004931098031 | 0.00208332908142136 |
| Lrfn4 | ENSMUSG00000045045 | 225875 | 1.06080464059444 | 0.814442625066633 | 4.56281781313458 | 0.0004932445030 | 0.00208332908142136 |
| Glrx5 | ENSMUSG00000021102 | 73046 | -0.412312008657629 | 4.76115750374813 | -4.56190667794766 | 0.0004940799502 | 0.00208605173258725 |
| Fndc3a | ENSMUSG00000033487 | 319448 | 0.487244954438222 | 4.30811886121572 | 4.56101021019168 | 0.0004949033764 | 0.00208872154735001 |
| Ccl2 | ENSMUSG00000035385 | 20296 | 1.98287474698781 | 0.191095834350259 | 4.56003846873791 | 0.0004957975460 | 0.0020916877560067 |
| Pex2 | ENSMUSG00000040374 | 19302 | -0.41604172215486 | 3.69216266905281 | -4.55954193526912 | 0.0004962550870 | 0.00209281032285253 |
| Zmat3 | ENSMUSG00000027663 | 22401 | 0.49224787820912 | 4.0885282102668 | 4.55907862653242 | 0.0004966824062 | 0.00209380462104734 |
| Poldip2 | ENSMUSG00000001100 | 67811 | -0.361735649345605 | 5.49530171208316 | -4.55664734262484 | 0.0004989310739 | 0.00210247323402186 |
| Ldlrad4 | ENSMUSG00000024544 | 52662 | 0.581497742697603 | 4.03815920870412 | 4.55258287858791 | 0.0005027137865 | 0.00211759707166226 |
| 5730455P16Rik | ENSMUSG00000057181 | 70591 | -0.322240555482356 | 4.77582170085058 | -4.55061044997435 | 0.0005045601584 | 0.00212358870737183 |
| Ttpal | ENSMUSG00000017679 | 76080 | 0.568438215109417 | 2.14011282060186 | 4.55050197686369 | 0.0005046619022 | 0.00212358870737183 |
| Car8 | ENSMUSG00000041261 | 12319 | -1.19156683015548 | 2.21500953669379 | -4.55044110554812 | 0.0005047190066 | 0.00212358870737183 |
| P2ry2 | ENSMUSG00000032860 | 18442 | -0.343738460178705 | 5.00752170102655 | -4.54860697614818 | 0.0005064427694 | 0.00213002151321605 |
| Egfl7 | ENSMUSG00000026921 | 353156 | -0.669788738869648 | 4.56431428791011 | -4.54622551743102 | 0.0005086900119 | 0.00213843951425947 |
| Efnb2 | ENSMUSG00000001300 | 13642 | -0.378069910670222 | 3.6611635929696 | -4.54607168363838 | 0.0005088355298 | 0.00213843951425947 |
| Zzef1 | ENSMUSG00000055670 | 195018 | 0.301291627690561 | 4.83631230608408 | 4.54457924941478 | 0.0005102495220 | 0.00214355784990557 |
| Cmtm8 | ENSMUSG00000041012 | 70031 | -1.52398451135094 | 1.37375736987533 | -4.54403674728918 | 0.0005107645163 | 0.00214489701450643 |
| Cops8 | ENSMUSG00000034432 | 108679 | -0.341717255095621 | 4.92827211789903 | -4.54310677733495 | 0.0005116485821 | 0.00214778442854428 |
| Ric8a | ENSMUSG00000025485 | 101489 | 0.505446208955196 | 3.21594052037172 | 4.5413279559255 | 0.0005133440083 | 0.0021532795077042 |
| Adgrg1 | ENSMUSG00000031785 | 14766 | -0.604096978280467 | 4.06646448402325 | -4.54121741789833 | 0.0005134495555 | 0.0021532795077042 |
| Tspan17 | ENSMUSG00000025875 | 74257 | 0.721180003284439 | 1.97048841935641 | 4.54107912654425 | 0.0005135816345 | 0.0021532795077042 |
| Shisa4 | ENSMUSG00000041889 | 77552 | 0.350442458370507 | 6.16047448342032 | 4.54090751904551 | 0.0005137455819 | 0.0021532795077042 |
| Smarca2 | ENSMUSG00000024921 | 67155 | -0.302111937721396 | 6.19097095099033 | -4.53790258718872 | 0.0005166251648 | 0.00216451884906733 |
| Nr2f2 | ENSMUSG00000030551 | 11819 | -0.506604694847097 | 3.69698943025637 | -4.53672720795635 | 0.0005177560480 | 0.00216836261418105 |
| Adamts3 | ENSMUSG00000043635 | 330119 | 1.09078879776227 | 0.821343561570021 | 4.53653696737604 | 0.0005179393272 | 0.00216836261418105 |
| Slc4a2 | ENSMUSG00000028962 | 20535 | 0.393775641297363 | 3.7543736570507 | 4.53611630651351 | 0.0005183448329 | 0.00216922946751405 |
| Chrd | ENSMUSG00000006958 | 12667 | 0.952215562735621 | 1.60208218791484 | 4.53422436843197 | 0.0005201726710 | 0.00217604572024067 |
| Ndc1 | ENSMUSG00000028614 | 72787 | -0.462186976670445 | 2.39351471102373 | -4.53381013694023 | 0.0005205737562 | 0.00217689048575906 |
| Spg20 | ENSMUSG00000036580 | 229285 | -0.245831061704978 | 5.19113443315957 | -4.53302893105618 | 0.0005213310394 | 0.00217908236516075 |
| Kptn | ENSMUSG00000006021 | 70394 | -0.820096005841625 | 1.06380207252155 | -4.53285828415276 | 0.0005214966121 | 0.00217908236516075 |
| Bnip1 | ENSMUSG00000024191 | 224630 | -0.490600194240661 | 2.68987601408102 | -4.53159445880676 | 0.0005227245502 | 0.00218337870054013 |
| Def8 | ENSMUSG00000001482 | 23854 | -0.365646654084903 | 4.79357637698081 | -4.53109572311577 | 0.0005232099459 | 0.0021845713977652 |
| Ttc32 | ENSMUSG00000066637 | 75516 | -0.62803426260497 | 2.38378462280031 | -4.53019434486858 | 0.0005240883965 | 0.00218740368926107 |
| Adgrd1 | ENSMUSG00000044017 | 243277 | 0.765485921389743 | 3.64869761539699 | 4.52970669525372 | 0.0005245642773 | 0.00218855424265431 |
| Hdac9 | ENSMUSG00000004698 | 79221 | 0.56380364005091 | 3.76232508865042 | 4.52774802606392 | 0.0005264801801 | 0.00219570959508258 |
| Rin2 | ENSMUSG00000001768 | 74030 | 0.689823702152023 | 4.17330140646675 | 4.52576534345332 | 0.0005284269378 | 0.0022029881227955 |
| Zadh2 | ENSMUSG00000049090 | 225791 | -0.412788803121933 | 3.80165661991698 | -4.52376874084284 | 0.0005303948817 | 0.00220993623246818 |
| Dkk2 | ENSMUSG00000028031 | 56811 | -1.5962915014035 | 2.97518912727001 | -4.52366442133136 | 0.0005304979118 | 0.00220993623246818 |
| Mapk7 | ENSMUSG00000001034 | 23939 | 0.711335203901208 | 1.80244350678526 | 4.5232017503587 | 0.0005309551133 | 0.00221099822621134 |
| Rasgrp2 | ENSMUSG00000032946 | 19395 | -1.00805871824066 | 2.09430988607913 | -4.52296763679245 | 0.0005311866144 | 0.00221111990957534 |
| Tmem250-ps | ENSMUSG00000087679 | 109299 | -0.426979063351088 | 3.91495239432119 | -4.52228797066535 | 0.0005318592873 | 0.00221307722510411 |
| Aspa | ENSMUSG00000020774 | 11484 | -0.737601440426453 | 1.15701890304239 | -4.51786284911471 | 0.0005362604538 | 0.00223054148437112 |
| C1qbp | ENSMUSG00000018446 | 12261 | -0.306172630634279 | 5.06245812101774 | -4.51748121996888 | 0.0005366417754 | 0.00223127852708125 |
| Cntln | ENSMUSG00000038070 | 338349 | 0.479492107210854 | 2.38080391456788 | 4.51582282539545 | 0.0005383020905 | 0.00223733085618226 |
| Npepl1 | ENSMUSG00000039263 | 228961 | -0.326584255548663 | 5.23371543881436 | -4.51466720473478 | 0.0005394621865 | 0.00224130032726438 |
| Vezf1 | ENSMUSG00000018377 | 22344 | -0.355470711878675 | 6.0210081885221 | -4.5140005854127 | 0.0005401325619 | 0.00224323291597223 |
| Mtfr1l | ENSMUSG00000046671 | 76824 | -0.355482809191993 | 5.86256723941959 | -4.51213247759566 | 0.0005420157851 | 0.00225019922065669 |
| Rp9 | ENSMUSG00000032239 | 55934 | -0.401225507099005 | 4.01561591157647 | -4.51178167448129 | 0.0005423701826 | 0.00225060872621259 |
| Myl9 | ENSMUSG00000067818 | 98932 | -0.809484365128194 | 3.62089434725318 | -4.51162730536661 | 0.0005425262092 | 0.00225060872621259 |
| Nckap5l | ENSMUSG00000023009 | 380969 | 0.728286200625376 | 2.08711089398465 | 4.51041248675425 | 0.0005437556913 | 0.0022548533619608 |
| Pdgfd | ENSMUSG00000032006 | 71785 | -0.995107144988938 | 2.4313167195428 | -4.50848900584664 | 0.0005457082799 | 0.002262092291446 |
| Lmbr1 | ENSMUSG00000010721 | 56873 | -0.526055080306731 | 2.59302266712481 | -4.50782370420889 | 0.0005463853343 | 0.00226404021589843 |
| Ptpn23 | ENSMUSG00000036057 | 104831 | 0.420453057414214 | 4.04477343323181 | 4.50555417465489 | 0.0005487014922 | 0.00227277605606433 |
| Khk | ENSMUSG00000029162 | 16548 | -0.809573306609683 | 1.85222826449032 | -4.50532875450793 | 0.0005489320968 | 0.00227286998125256 |
| Extl3 | ENSMUSG00000021978 | 54616 | 0.273083447607142 | 4.78397425843957 | 4.50199405483049 | 0.0005523552107 | 0.00228617751173756 |
| Tmem159 | ENSMUSG00000030917 | 233806 | 0.417365782474004 | 3.45841931139883 | 4.50037360459919 | 0.0005540265782 | 0.00229222730004339 |
| Slc9a3r2 | ENSMUSG00000002504 | 65962 | -1.11173876641321 | 4.97227616656036 | -4.49796984823962 | 0.0005565154928 | 0.00230165374656309 |
| Snhg12 | ENSMUSG00000086290 | 100039864 | 0.643853812719801 | 1.84447350287693 | 4.49746108509213 | 0.0005570437592 | 0.00230296722118061 |
| Snap29 | ENSMUSG00000022765 | 67474 | -0.36430983781699 | 4.01125794884503 | -4.49662905852378 | 0.0005579087978 | 0.00230567148166313 |
| Prdm16 | ENSMUSG00000039410 | 70673 | -0.796316809869255 | 2.22194058591164 | -4.49410417587998 | 0.0005605423630 | 0.0023150633706077 |
| Nudt13 | ENSMUSG00000021809 | 67725 | -0.504871802608739 | 3.50624166699345 | -4.49404431802297 | 0.0005606049530 | 0.0023150633706077 |
| Vamp8 | ENSMUSG00000050732 | 22320 | -0.532952375968343 | 2.89075876020156 | -4.49254587914903 | 0.0005621741406 | 0.00232029002426247 |
| Tmem109 | ENSMUSG00000034659 | 68539 | -0.300048427715397 | 5.80519628498531 | -4.49243050602998 | 0.0005622951490 | 0.00232029002426247 |
| Akap2 | ENSMUSG00000038729 | 11641 | 0.413010037954935 | 5.91029826023427 | 4.49205837099115 | 0.0005626856439 | 0.00232102519785569 |
| Egflam | ENSMUSG00000042961 | 268780 | -0.785816339306116 | 2.73126849025939 | -4.48993805012911 | 0.0005649159248 | 0.00232934589758522 |
| Ddit4l | ENSMUSG00000046818 | 73284 | -0.978169931318935 | 4.64690086867513 | -4.48675660697914 | 0.0005682794987 | 0.00234230409434963 |
| Gpm6b | ENSMUSG00000031342 | 14758 | 0.790249885722935 | 2.27214145118554 | 4.48656086106326 | 0.0005684871249 | 0.00234230409434963 |
| Trp53 | ENSMUSG00000059552 | 22059 | 0.487339870531663 | 2.63701885253654 | 4.48487978992013 | 0.0005702734525 | 0.00234877886576149 |
| Tkt | ENSMUSG00000021957 | 21881 | 0.448128680856767 | 3.0331438238009 | 4.48443188936601 | 0.0005707503746 | 0.00234985775719372 |
| Scx | ENSMUSG00000034161 | 20289 | 0.807553424173359 | 4.13071290674417 | 4.48369081463136 | 0.0005715403717 | 0.00235135382298425 |
| Arap3 | ENSMUSG00000024451 | 106952 | -0.800111958616258 | 4.06074967797406 | -4.48359867595887 | 0.0005716386719 | 0.00235135382298425 |
| Atrx | ENSMUSG00000031229 | 22589 | 0.305117128460564 | 6.12666523472669 | 4.48348584099221 | 0.0005717590761 | 0.00235135382298425 |
| Rela | ENSMUSG00000024927 | 19697 | 0.36623654577842 | 4.1125332503929 | 4.47672237847624 | 0.0005790242921 | 0.0023803364188148 |
| Trdmt1 | ENSMUSG00000026723 | 13434 | -0.523113596029499 | 1.81032003048726 | -4.4740065387296 | 0.0005819683716 | 0.0023915399512934 |
| Ptpn18 | ENSMUSG00000026126 | 19253 | 0.925219848860179 | 0.978877152714874 | 4.47055409817799 | 0.0005857332830 | 0.00240610692106792 |
| Zscan18 | ENSMUSG00000070822 | 232875 | 0.591019209750989 | 1.55993188547887 | 4.46908062458827 | 0.0005873477647 | 0.00241075952330547 |
| 2610507B11Rik | ENSMUSG00000010277 | 72503 | -0.375529438952414 | 8.13351087507821 | -4.46876554853496 | 0.0005876935886 | 0.00241075952330547 |
| Prmt2 | ENSMUSG00000020230 | 15468 | 1.14562182579485 | 1.10077018333272 | 4.46875870495769 | 0.0005877011024 | 0.00241075952330547 |
| Trpc4ap | ENSMUSG00000038324 | 56407 | -0.237682962056453 | 5.96523302647489 | -4.46871593068193 | 0.0005877480678 | 0.00241075952330547 |
| Usp38 | ENSMUSG00000038250 | 74841 | -0.31503819227812 | 4.10382854130968 | -4.46836046032512 | 0.0005881385178 | 0.00241145616621434 |
| Zfp943 | ENSMUSG00000053347 | 74670 | -0.384406425090394 | 3.86131005189628 | -4.46745619294871 | 0.0005891329763 | 0.00241462788303222 |

| | | | | | | | |
|---|---|---|---|---|---|---|---|
| Urod | ENSMUSG00000028684 | 22275 | -0.310847974045931 | 4.46671323097134 | -4.46686128828339 | 0.0005897881629 | 0.00241640719983256 |
| Pfdn1 | ENSMUSG00000024346 | 67199 | -0.365646040554505 | 4.96175801355072 | -4.46654571675382 | 0.0005901360167 | 0.00241692648916083 |
| Mif4gd | ENSMUSG00000020743 | 69674 | -0.383230314140345 | 3.51611374128151 | -4.46614352409564 | 0.0005905796599 | 0.00241783755165447 |
| H2-Q7 | ENSMUSG00000060550 | 15018 | -1.18717934197132 | 2.60602107172747 | -4.46462809834375 | 0.0005922543625 | 0.0024237860117948 |
| Ifi30 | ENSMUSG00000031838 | 65972 | 0.844651779496834 | 2.0256231938234 | 4.46377904581941 | 0.0005931947958 | 0.00242672616508375 |
| Myh4 | ENSMUSG00000057003 | 17884 | -3.19291826788309 | 8.28898924676014 | -4.46328273773899 | 0.0005937452336 | 0.00242806926656932 |
| Abca8a | ENSMUSG00000041828 | 217258 | 0.455475189710453 | 5.43506386736915 | 4.46303668659319 | 0.0005940183158 | 0.00242827756569854 |
| Smpx | ENSMUSG00000041476 | 66106 | 0.43430968088528 | 7.25348706905239 | 4.46167884985708 | 0.0005955276574 | 0.00243353750423892 |
| Arhgef17 | ENSMUSG00000032875 | 207212 | 0.431474093338772 | 5.47223211042851 | 4.45916685785417 | 0.0005983303783 | 0.00244289974823767 |
| Fryl | ENSMUSG00000070733 | 72313 | 0.431247003692286 | 4.95340999562298 | 4.45907923094058 | 0.0005984283923 | 0.00244289974823767 |
| Tent4a | ENSMUSG00000034575 | 210106 | 0.442947501317311 | 3.47752639273254 | 4.45902486822808 | 0.0005984892073 | 0.00244289974823767 |
| Nav2 | ENSMUSG00000052512 | 78286 | 0.405596130260748 | 6.13554904744217 | 4.45789301522592 | 0.0005997568505 | 0.00244716018413494 |
| Rubcn | ENSMUSG00000035629 | 100502698 | 0.359573308985442 | 4.12538150974167 | 4.45400817767779 | 0.0006041288790 | 0.0024640793943172 |
| Slc16a2 | ENSMUSG00000033965 | 20502 | 0.624897071032018 | 3.30912597865478 | 4.4529199020823 | 0.0006053595223 | 0.00246694499271311 |
| Serac1 | ENSMUSG00000015659 | 321007 | -0.485420143541014 | 2.36992184480662 | -4.45284713433817 | 0.0006054419017 | 0.00246694499271311 |
| Dnajb12 | ENSMUSG00000020109 | 56709 | -0.263305719832151 | 5.43102069026517 | -4.45278831381328 | 0.0006055085001 | 0.00246694499271311 |
| Ibtk | ENSMUSG00000035941 | 108837 | -0.341920241251306 | 6.34690608102676 | -4.45124915250686 | 0.0006072538762 | 0.00247240397943926 |
| Tbkbp1 | ENSMUSG00000038517 | 73174 | -0.743600095574797 | 2.10131713470307 | -4.45110452358131 | 0.0006074181490 | 0.00247240397943926 |
| Nf2 | ENSMUSG00000009073 | 18016 | -0.311558525885617 | 6.01154695064696 | -4.45100875551773 | 0.0006075269498 | 0.00247240397943926 |
| Selenon | ENSMUSG00000050989 | 74777 | 0.620327856052601 | 3.03369635651665 | 4.44977927833286 | 0.0006089255297 | 0.00247717341495705 |
| Cemip2 | ENSMUSG00000024754 | 83921 | 0.494750817896831 | 4.05262401291241 | 4.44660646477262 | 0.0006125501066 | 0.0024909915236763 |
| Tacc1 | ENSMUSG00000065954 | 320165 | 0.287877666290974 | 5.66493159159531 | 4.44566747090128 | 0.0006136270643 | 0.00249382165025132 |
| Stk11ip | ENSMUSG00000026213 | 71728 | 0.493621682711745 | 2.75193498970733 | 4.44560190514843 | 0.0006137023363 | 0.00249382165025132 |
| Zfand5 | ENSMUSG00000024750 | 22682 | -0.35438576680486 | 7.20134151919238 | -4.44422667135787 | 0.0006152833575 | 0.00249931712442183 |
| Zfp787 | ENSMUSG00000046792 | 67109 | -0.343216312245561 | 3.42300585146422 | -4.44250113344108 | 0.0006172730493 | 0.00250646794310736 |
| 2310057M21Rik | ENSMUSG00000040177 | 68277 | -0.464713939574716 | 3.8740840084198 | -4.44184883969306 | 0.0006180269278 | 0.00250859723313209 |
| D17H6S53E | ENSMUSG00000043311 | 114585 | -0.456905550224712 | 2.5897306454667 | -4.43987537213907 | 0.0006203135237 | 0.00251694399709064 |
| Hagh | ENSMUSG00000024158 | 14651 | -0.31956122682197 | 4.06375095684243 | -4.43939373575126 | 0.0006208729051 | 0.00251827893351845 |
| 4632428C04Rik | ENSMUSG00000097184 | 100043102 | -1.13178474276442 | 1.32242783010877 | -4.43839683719346 | 0.0006220323757 | 0.00252204595681688 |
| Zfp438 | ENSMUSG00000050945 | 240186 | -0.474951925830772 | 2.84965169651239 | -4.43616132257185 | 0.0006246405836 | 0.0025316819502836 |
| Eif6 | ENSMUSG00000027613 | 16418 | 0.36339610985035 | 3.93778410082044 | 4.43450520490911 | 0.0006265800753 | 0.00253847025884392 |
| Fam149a | ENSMUSG00000070044 | 212326 | 0.645103931586773 | 1.4345156032286 | 4.4343348657241 | 0.0006267799129 | 0.00253847025884392 |
| Cd302 | ENSMUSG00000060703 | 66205 | 1.00894414384856 | 1.80937278442108 | 4.43306705354164 | 0.0006282693457 | 0.0025435600808905 |
| Hspa1l | ENSMUSG00000007033 | 15482 | -0.396958092508269 | 5.33015188829805 | -4.43202601920917 | 0.0006294950889 | 0.00254757897725222 |
| Ddt | ENSMUSG00000001666 | 13202 | -0.501057401346809 | 3.21405925621683 | -4.43164231559159 | 0.0006299474945 | 0.00254846634447726 |
| Adcyap1r1 | ENSMUSG00000029778 | 11517 | 1.10917702793636 | 0.880357167656568 | 4.42830842763227 | 0.0006338924615 | 0.00256347706133658 |
| Dpy19l3 | ENSMUSG00000043671 | 233115 | 0.751318945036535 | 2.5597337034688 | 4.42781566321187 | 0.0006344777047 | 0.00256489489304051 |
| Tpm2 | ENSMUSG00000028464 | 22004 | -0.42773213148087 | 12.4690021814673 | -4.42703329131621 | 0.0006354080525 | 0.00256770625612431 |
| Ints10 | ENSMUSG00000031864 | 70885 | -0.458448555230679 | 3.77958652851649 | -4.42563593464829 | 0.0006370732052 | 0.00257348381599149 |
| Slc31a1 | ENSMUSG00000066150 | 20529 | -0.428338748161729 | 3.23442513790292 | -4.42526636957066 | 0.0006375143477 | 0.00257431449411026 |
| Ndufc2 | ENSMUSG00000030647 | 68197 | -0.593988952946323 | 5.80115481807836 | -4.42404701278678 | 0.0006389721065 | 0.0025792481894937 |
| Ncor1 | ENSMUSG00000018501 | 20185 | -0.265006622813022 | 7.50457187785676 | -4.42265428708394 | 0.0006406413382 | 0.00258503154986472 |
| Atg13 | ENSMUSG00000027244 | 51897 | 0.353432492061388 | 4.86674056444932 | 4.41996538338914 | 0.0006438768244 | 0.00259712825392532 |
| Eng | ENSMUSG00000026814 | 13805 | -0.75791013304586 | 5.51869926373423 | -4.4180589721501 | 0.0006461809668 | 0.00260546077025627 |
| Alkbh8 | ENSMUSG00000025899 | 67667 | -0.391223633429458 | 3.85325388891321 | -4.4173480884449 | 0.0006470423365 | 0.00260797189570042 |
| Wdr13 | ENSMUSG00000031166 | 73447 | 0.355754165802205 | 4.00137286377431 | 4.41617552778551 | 0.0006484657040 | 0.00261274552550658 |
| Atp1a2 | ENSMUSG00000007097 | 98660 | -0.401694242083144 | 10.5506875634839 | -4.41583199500275 | 0.0006488833287 | 0.0026134648735218 |
| Lbr | ENSMUSG00000004880 | 98386 | -0.330691813057073 | 3.69421624072413 | -4.41445546409611 | 0.0006505595300 | 0.0026192509109368 |
| Crebl2 | ENSMUSG00000032652 | 232430 | -0.534763481025492 | 2.62034115378821 | -4.41348412638381 | 0.0006517450149 | 0.00262305771675619 |
| Lactb2 | ENSMUSG00000025937 | 212442 | -0.375792817952371 | 3.86064669396439 | -4.40928743373264 | 0.0006568926033 | 0.00264227315770773 |
| Efemp2 | ENSMUSG00000024909 | 58859 | 0.744328022457528 | 2.03848888453241 | 4.40919790247478 | 0.0006570028764 | 0.00264227315770773 |
| Prmt7 | ENSMUSG00000060098 | 214572 | -0.319496500389704 | 4.50824815540402 | -4.40870395024271 | 0.0006576116069 | 0.00264374861198023 |
| Sirt2 | ENSMUSG00000015149 | 64383 | -0.311295424111552 | 5.2870868114843 | -4.40779303772869 | 0.0006587357116 | 0.0026472941411361 |
| Dynll2 | ENSMUSG00000020483 | 68097 | -0.31364326441525 | 6.31433310961994 | -4.40712290483189 | 0.0006595639491 | 0.00264964848516864 |
| Tgfb1 | ENSMUSG00000002603 | 21803 | 0.729194552420947 | 3.51329477901542 | 4.40667425543942 | 0.0006601190486 | 0.00265090423246769 |
| Stx8 | ENSMUSG00000020903 | 55943 | -0.386306362276554 | 3.09401931829119 | -4.40622766396904 | 0.0006606720804 | 0.00265215075711777 |
| Pcdh18 | ENSMUSG00000037892 | 73173 | 0.728578198783127 | 2.82679013834684 | 4.40432767113007 | 0.0006630302626 | 0.00266064016190934 |
| Ampd2 | ENSMUSG00000027889 | 109674 | 0.933877912254762 | 1.51038322006296 | 4.40262864417709 | 0.0006651463671 | 0.00266815227149202 |
| Kalrn | ENSMUSG00000061751 | 545156 | -0.311113587977519 | 4.76438708665143 | -4.40189849986555 | 0.0006660578860 | 0.00267004650386598 |
| 9530020I12Rik | ENSMUSG00000112041 | 77442 | -1.03368205890824 | 2.33793070158994 | -4.40169688053725 | 0.0006663098163 | 0.00267004650386598 |
| Ppp2r2d | ENSMUSG00000041769 | 52432 | -0.356741477265595 | 4.41275283774018 | -4.40166363009621 | 0.0006663513733 | 0.00267004650386598 |
| Pxdn | ENSMUSG00000020674 | 69675 | 0.595142910918407 | 4.94658131231812 | 4.40128588148828 | 0.0006668236782 | 0.00267095992204879 |
| Zfp275 | ENSMUSG00000031365 | 27081 | 0.430185075906851 | 2.79962524806912 | 4.40076522076747 | 0.0006674752342 | 0.00267194829507109 |
| Ephx1 | ENSMUSG00000038776 | 13849 | -0.482958576854867 | 4.46558893596004 | -4.4006980754846 | 0.0006675593078 | 0.00267194829507109 |
| Tle3 | ENSMUSG00000032280 | 21887 | 0.663913616140505 | 3.70480543840221 | 4.40041135385927 | 0.0006679184388 | 0.00267240719436056 |
| Rnps1 | ENSMUSG00000034681 | 19826 | -0.269215768123959 | 5.13777712616884 | -4.39913795532001 | 0.0006695158335 | 0.00267781835966732 |
| Tmx3 | ENSMUSG00000024614 | 67988 | 0.294452698056326 | 4.30847240968571 | 4.39852948909202 | 0.0006702805049 | 0.00267916276877105 |
| Tmem173 | ENSMUSG00000024349 | 72512 | 0.942761097713919 | 1.64842734102703 | 4.39848045836624 | 0.0006703421619 | 0.00267916276877105 |
| Aff3 | ENSMUSG00000037138 | 16764 | 0.684996779996337 | 2.18558330354738 | 4.39824348312666 | 0.0006706402452 | 0.0026793744594189 |
| Ubxn6 | ENSMUSG00000019578 | 66530 | -0.283371347168214 | 4.56381651041858 | -4.39768124623801 | 0.0006713480113 | 0.00268122218194539 |
| Eps8 | ENSMUSG00000015766 | 13860 | 0.520183437129332 | 3.48485117491066 | 4.39592272994057 | 0.0006735666774 | 0.002689100566506 |
| Nek6 | ENSMUSG00000026749 | 59126 | 0.907195964580791 | 1.76048632724818 | 4.39484666031113 | 0.0006749280489 | 0.00269355184493392 |
| Tnmd | ENSMUSG00000031250 | 64103 | 0.902086226923317 | 3.85381868826199 | 4.39372164261632 | 0.0006763543803 | 0.0026982590259664 |
| Letm1 | ENSMUSG00000005299 | 56384 | -0.356296943106226 | 6.31150837517381 | -4.39352335477325 | 0.0006766060976 | 0.00269827845786863 |
| Zfp217 | ENSMUSG00000052056 | 228913 | 0.533226090349187 | 3.04503784461561 | 4.39274452870604 | 0.0006775957166 | 0.00270123952557483 |
| Kif3a | ENSMUSG00000018395 | 16568 | 0.440753605516588 | 2.69573500055265 | 4.39087281944927 | 0.0006799801209 | 0.0027097567268452 |
| Epb41l3 | ENSMUSG00000024044 | 13823 | 0.691683340949749 | 1.93546795098142 | 4.39035265220333 | 0.0006806443063 | 0.00271141505570964 |
| Pdrg1 | ENSMUSG00000027472 | 68559 | -0.360270405997515 | 4.54610475325114 | -4.38888160042575 | 0.0006825262699 | 0.00271792155057943 |
| Gbp4 | ENSMUSG00000079363 | 17472 | -0.911902457066632 | 3.28951305802457 | -4.38817099741259 | 0.0006834372880 | 0.002720502288 |
| Usp10 | ENSMUSG00000031826 | 22224 | -0.259602138484702 | 4.34136864427471 | -4.38798799577525 | 0.0006836721054 | 0.002720502288 |
| Cux2 | ENSMUSG00000042589 | 13048 | -0.905689807785154 | 2.00397143200917 | -4.38626780125995 | 0.0006858834330 | 0.00272830851783596 |
| Sorbs1 | ENSMUSG00000025006 | 20411 | -0.309900318749179 | 8.03725601092573 | -4.38541078532225 | 0.0006869878894 | 0.00273170775532208 |
| Arl3 | ENSMUSG00000025035 | 56350 | -0.369292766281752 | 3.93708292926318 | -4.38517779848888 | 0.0006872884582 | 0.00273190914072749 |
| Cluap1 | ENSMUSG00000014232 | 76779 | -0.477637648576139 | 2.5583238221421 | -4.38478896311045 | 0.0006877903889 | 0.00273291048402713 |
| 1110038B12Rik | ENSMUSG00000092203 | 68763 | 0.727457863933665 | 1.46085486407624 | 4.38371223724888 | 0.0006891822606 | 0.00273744596343097 |
| Nucb2 | ENSMUSG00000030659 | 53322 | 0.765372470261887 | 2.66646451505283 | 4.38211850271238 | 0.0006912477960 | 0.00274379346484156 |
| Mlxipl | ENSMUSG00000005373 | 58805 | -0.509729868283126 | 5.97903000035412 | -4.38209189488849 | 0.0006912823348 | 0.00274379346484156 |
| Cacnb3 | ENSMUSG00000003352 | 12297 | 1.13145364602958 | 1.40524259758328 | 4.38095447445103 | 0.0006927604492 | 0.00274866223970269 |
| Cdk5rap1 | ENSMUSG00000027487 | 66971 | -0.524824890447545 | 2.39450083268607 | -4.3799475543793 | 0.0006940716929 | 0.00275286562985402 |
| Mat2b | ENSMUSG00000042032 | 108645 | -0.385647832235924 | 4.61855464222095 | -4.3789121697428 | 0.0006954226724 | 0.00275722351546726 |
| Aatk | ENSMUSG00000025375 | 11302 | -0.949413289164498 | 1.67564896263489 | -4.37844073135465 | 0.0006960387074 | 0.00275866537741282 |
| Gucd1 | ENSMUSG00000033416 | 68778 | -0.575461075367556 | 4.86437842570944 | -4.37548328329244 | 0.0006999161002 | 0.00277302750690086 |
| Btbd1 | ENSMUSG00000025103 | 83962 | -0.400094324100092 | 7.56951748852354 | -4.37118794770139 | 0.0007055872367 | 0.00279442488076734 |
| Pgm3 | ENSMUSG00000056131 | 109785 | 0.541499092583297 | 1.91152308930395 | 4.3710062964924 | 0.0007058281123 | 0.00279442488076734 |
| Rab21 | ENSMUSG00000020132 | 216344 | -0.277794785743346 | 5.77450892163178 | -4.36903415271346 | 0.0007084487023 | 0.00280324497818957 |
| Pskh1 | ENSMUSG00000048310 | 244631 | -0.495485432905299 | 3.04986946486374 | -4.36894393048536 | 0.0007085688294 | 0.00280324497818957 |
| Igfbp3 | ENSMUSG00000020427 | 16009 | -0.778305164304304 | 2.96277648031008 | -4.36808132020638 | 0.0007097184196 | 0.00280677715088596 |
| Fermt2 | ENSMUSG00000037712 | 218952 | -0.256452400270598 | 7.1942839557427 | -4.3677415172694 | 0.0007101717984 | 0.00280755440470925 |
| Gmip | ENSMUSG00000036246 | 78816 | 0.880345834730558 | 1.61466058531456 | 4.36678340208591 | 0.0007114517626 | 0.00281159769233681 |
| Col8a2 | ENSMUSG00000056174 | 329941 | 1.05064691133343 | 2.80148088810551 | 4.36355621982662 | 0.0007157805344 | 0.00282768233552853 |
| Ddx24 | ENSMUSG00000041645 | 27225 | -0.269640462762399 | 6.04068358162262 | -4.36277286424977 | 0.0007168353744 | 0.00283082640092309 |
| Brms1 | ENSMUSG00000080268 | 107392 | 0.431057675621035 | 3.11963056875149 | 4.36158551124245 | 0.0007184372788 | 0.00283612780600091 |
| Jkamp | ENSMUSG00000005078 | 104771 | 0.59727194302149 | 1.90239455977654 | 4.36095096581756 | 0.0007192948809 | 0.00283848821088704 |
| Cyfip2 | ENSMUSG00000020340 | 76884 | 0.567681628112758 | 5.79102464800805 | 4.35936411231316 | 0.0007214441695 | 0.00284581971767367 |
| Nuak1 | ENSMUSG00000020032 | 77976 | 0.349592409906557 | 6.68964607187732 | 4.35919513376472 | 0.0007216734294 | 0.00284581971767367 |
| Gpx8 | ENSMUSG00000021760 | 69590 | 0.647488451880862 | 2.34751455699007 | 4.35850089147289 | 0.0007226161235 | 0.00284779743638715 |
| Tmbim6 | ENSMUSG00000023010 | 110213 | -0.244715584486497 | 5.40603918444443 | -4.35844210216438 | 0.0007226960102 | 0.00284779743638715 |
| Mtif2 | ENSMUSG00000020459 | 76784 | -0.396328399067773 | 4.69156681497299 | -4.35526322225151 | 0.0007270292595 | 0.0028637475513563 |
| Cdh23 | ENSMUSG00000012819 | 22295 | -0.811591685979634 | 1.64023111407889 | -4.35508887075045 | 0.0007272676976 | 0.0028637475513563 |
| Araf | ENSMUSG00000001127 | 11836 | -0.26661228932078 | 4.88888737653755 | -4.35323372203052 | 0.0007298097408 | 0.00287178618646774 |
| Ift88 | ENSMUSG00000040040 | 21821 | -0.526130635128377 | 2.30325899521506 | -4.35321561213341 | 0.0007298346012 | 0.00287178618646774 |
| Pnp | ENSMUSG00000115338 | 18950 | 0.615844330107065 | 2.65046945557904 | 4.35292951527362 | 0.0007302274572 | 0.00287229806954831 |
| Prkar2a | ENSMUSG00000032601 | 19087 | -0.319675110893372 | 7.21143025235571 | -4.34948060601259 | 0.0007349805343 | 0.00288995403637243 |
| D5Ertd579e | ENSMUSG00000029190 | 320661 | -0.301632435810418 | 7.07472052649968 | -4.34763324945822 | 0.0007375395470 | 0.00289897331510677 |
| Ssh3 | ENSMUSG00000034616 | 245857 | 0.891267222068728 | 1.17036934236236 | 4.34669375340173 | 0.0007388444842 | 0.0029030586115597 |
| Zfp598 | ENSMUSG00000041130 | 213753 | 0.314597976409334 | 3.93741015427209 | 4.34505384763073 | 0.0007411279761 | 0.00291067321449459 |

| | | | | | | | |
|---|---|---|---|---|---|---|---|
| Tob1 | ENSMUSG00000037573 | 22057 | -0.368033690203992 | 6.65213029577234 | -4.34491976820683 | 0.0007413149967 | 0.00291067321449459 |
| Snora73a | ENSMUSG00000064387 | 100306944 | 0.977179468251477 | 2.77589728324762 | 4.34319884981312 | 0.0007437197394 | 0.00291906659685957 |
| Tmsb4x | ENSMUSG00000049775 | 19241 | 0.427776625537311 | 6.19973673908671 | 4.33879154688099 | 0.0007499150421 | 0.0029423263910653 |
| Atf4 | ENSMUSG00000042406 | 11911 | -0.291444876592843 | 6.44062766103573 | -4.33732534627446 | 0.0007519878306 | 0.0029494004223438 |
| Ergic1 | ENSMUSG00000001576 | 67458 | 0.375796400474735 | 4.81122373602691 | 4.33711391810368 | 0.0007522872161 | 0.00294951634146918 |
| Sgca | ENSMUSG00000001508 | 20391 | -0.490566604551811 | 5.70405355545657 | -4.33679740598069 | 0.0007527356316 | 0.00295021627449607 |
| Slc38a9 | ENSMUSG00000047789 | 268706 | 0.544618651990021 | 2.95053859458258 | 4.33656392841 23 | 0.0007530665851 | 0.00295045549912371 |
| Hbegf | ENSMUSG00000024486 | 15200 | 0.961558067398028 | 3.00552054103583 | 4.3350863363 2944 | 0.0007551645406 | 0.0029576150460089 |
| B4galt1 | ENSMUSG00000028413 | 14595 | 0.405931493234982 | 4.68836635722821 | 4.33442962316529 | 0.0007560989108 | 0.00296021389475904 |
| Ifi205 | ENSMUSG00000054203 | 226695 | 1.26454383317074 | 2.23163451827468 | 4.33392910579489 | 0.0007568118423 | 0.00296194423510593 |
| Them4 | ENSMUSG00000028145 | 75778 | -0.603484781346239 | 1.73112536885818 | -4.33284201128633 | 0.0007583626740 | 0.00296695146399567 |
| Mtm1 | ENSMUSG00000031337 | 17772 | -0.308209661490584 | 3.46250114496355 | -4.33197102818645 | 0.0007596075663 | 0.00297075860760483 |
| Cdkn2aipnl | ENSMUSG00000020392 | 52626 | -0.533191896311592 | 1.93365421809359 | -4.32931969197855 | 0.0007634100811 | 0.00298456208769399 |
| Pim3 | ENSMUSG00000035828 | 223775 | -0.646582782887279 | 3.81811239086539 | -4.32894329946484 | 0.0007639514858 | 0.00298561090145156 |
| Mcoln1 | ENSMUSG00000004567 | 94178 | -0.356122134973538 | 3.81212643943991 | -4.32581289933794 | 0.0007684696164 | 0.00300132731317694 |
| Osbpl8 | ENSMUSG00000020189 | 237542 | 0.33398752312671 | 5.5628044032727 | 4.32577664271693 | 0.0007685221068 | 0.00300132731317694 |
| Gpx3 | ENSMUSG00000018339 | 14778 | 0.446105495825874 | 5.54951494994206 | 4.32536931620596 | 0.0007691120658 | 0.00300255856862962 |
| Faf1 | ENSMUSG00000010517 | 14084 | -0.328952296312538 | 5.42885049048776 | -4.32499542756279 | 0.0007696540051 | 0.00300360154590717 |
| Zc3h12c | ENSMUSG00000035164 | 244871 | 0.297747304348162 | 3.98453069433427 | 4.32328029144479 | 0.0007721450827 | 0.00301224764426891 |
| Slc6a17 | ENSMUSG00000027894 | 229706 | 1.37707091056259 | 0.533906329182542 | 4.32179378538076 | 0.0007743108124 | 0.00301961879815647 |
| Ap2a2 | ENSMUSG00000002957 | 11772 | 0.243312881879165 | 5.96992788276344 | 4.32045827554917 | 0.0007762618864 | 0.00302614788882301 |
| Echdc1 | ENSMUSG00000019883 | 52665 | -0.408723468718335 | 3.10014688126905 | -4.32018981523623 | 0.0007766546963 | 0.00302659981671075 |
| Fxyd6 | ENSMUSG00000066705 | 59095 | -0.384995454848334 | 5.66806242617645 | -4.31974928668496 | 0.0007772997193 | 0.00302803393892292 |
| Ccdc85a | ENSMUSG00000032878 | 216613 | -0.476486660758951 | 3.46136668141896 | -4.31761734092619 | 0.0007804291235 | 0.00303894700442554 |
| Lix1l | ENSMUSG00000049288 | 280411 | 0.752601347371602 | 2.94713934285805 | 4.31739134761062 | 0.0007807616103 | 0.00303894700442554 |
| Atp7a | ENSMUSG00000033792 | 11977 | 0.393812234827527 | 3.2155747096978 | 4.31727343037765 | 0.0007809351509 | 0.00303894700442554 |
| Ccnk | ENSMUSG00000021258 | 12454 | -0.356998744338593 | 5.09622134015777 | -4.31538677742934 | 0.0007837171750 | 0.00304868770111716 |
| Cbr4 | ENSMUSG00000031641 | 234309 | -0.545364158314983 | 3.05002863379276 | -4.31459051382284 | 0.0007848943913 | 0.00305138978838916 |
| Emc9 | ENSMUSG00000022217 | 85308 | -0.484304668301534 | 2.7076574369703 | -4.31453935173734 | 0.0007849700928 | 0.00305138978838916 |
| Lims1 | ENSMUSG00000019920 | 110829 | 0.288489489678587 | 6.3327247002113 | 4.3131565690629 | 0.0007870189654 | 0.00305826673006525 |
| Myo18a | ENSMUSG00000000631 | 360013 | 0.383523335796941 | 8.80378545511574 | 4.31263101051581 | 0.0007877991303 | 0.00305936193344263 |
| Tmem140 | ENSMUSG00000057137 | 68487 | -0.609382284635036 | 3.43760639653327 | -4.31258964736507 | 0.0007878605656 | 0.00305936193344263 |
| Eral1 | ENSMUSG00000020832 | 57837 | -0.461689602523714 | 3.0861695937153 | -4.31063485846318 | 0.0007907695792 | 0.0030695675676435 |
| Rcn3 | ENSMUSG00000019539 | 52377 | 0.860106960554441 | 3.48242906774692 | 4.31039295893931 | 0.0007911303294 | 0.00306959598847563 |
| Ociad2 | ENSMUSG00000029153 | 433904 | 0.548912734970459 | 2.18882782156008 | 4.31025340115438 | 0.0007913385322 | 0.00306959598847563 |
| Mtrf1 | ENSMUSG00000022022 | 211253 | -0.51186521080975 | 2.55695830417921 | -4.30799258354672 | 0.0007947192596 | 0.00308161625673822 |
| Spsb4 | ENSMUSG00000046997 | 211949 | 0.726644818438065 | 2.58483377414641 | 4.30702908334762 | 0.0007961645511 | 0.00308583192784802 |
| Dclre1c | ENSMUSG00000026648 | 227525 | 0.666407866325297 | 1.66203210587586 | 4.30689157325745 | 0.0007963710427 | 0.00308583192784802 |
| Arhgef2 | ENSMUSG00000028059 | 16800 | 0.330915901621494 | 5.60465928226717 | 4.30652850282085 | 0.0007969165112 | 0.00308635928181089 |
| Ccn3 | ENSMUSG00000037362 | 18133 | 0.719762334726053 | 2.56892050445187 | 4.30642516280843 | 0.0007970718372 | 0.00308635928181089 |
| Chd1 | ENSMUSG00000023852 | 12648 | 0.259340356090265 | 4.95306195179467 | 4.30360329859646 | 0.0008013253271 | 0.00310173057772215 |
| Mical2 | ENSMUSG00000038244 | 320878 | 0.593031453661014 | 4.42504170417845 | 4.30339273349088 | 0.0008016436543 | 0.00310186434905257 |
| Rasip1 | ENSMUSG00000044562 | 69903 | -0.556215005457521 | 3.74082588785615 | -4.30177844843051 | 0.0008040884198 | 0.00311022311320285 |
| Dip2b | ENSMUSG00000023026 | 239667 | 0.333476010971414 | 4.33839157845801 | 4.30122939360065 | 0.0008049216880 | 0.00311190822213171 |
| Dtx2 | ENSMUSG00000004947 | 74198 | 0.514331639890937 | 2.1708919958252 | 4.30111629237799 | 0.0008050934454 | 0.00311190822213171 |
| Abcb9 | ENSMUSG00000029408 | 56325 | -0.746130862781015 | 1.59121969652264 | -4.30088114722278 | 0.0008054506616 | 0.00311218847026997 |
| Ugdh | ENSMUSG00000029201 | 22235 | 0.628411921347854 | 3.16167214794681 | 4.29990248106703 | 0.0008069391368 | 0.00311629578951951 |
| Tlr4 | ENSMUSG00000039005 | 21898 | 0.719309267845231 | 3.91633199517788 | 4.29980744295377 | 0.0008070838331 | 0.00311629578951951 |
| Anxa11 | ENSMUSG00000021866 | 11744 | 0.392758524760422 | 5.55275915483556 | 4.29949972167482 | 0.0008075525246 | 0.00311700446591636 |
| Dap3 | ENSMUSG00000068921 | 65111 | -0.390853567409787 | 4.14320098485269 | -4.29928728676112 | 0.0008078762486 | 0.00311715329123858 |
| Vwa8 | ENSMUSG00000058997 | 219189 | -0.290031643470137 | 6.8760899297932 | -4.29851132263907 | 0.0008090598554 | 0.00312014020137546 |
| Map3k10 | ENSMUSG00000040390 | 269881 | -0.333905866185274 | 4.15584358895825 | -4.29840560485679 | 0.0008092212488 | 0.00312014020137546 |
| Stom | ENSMUSG00000026880 | 13830 | -0.295223723802713 | 4.73441250585771 | -4.29789759415245 | 0.0008099972623 | 0.00312203105612162 |
| Hemk1 | ENSMUSG00000032579 | 69536 | -0.515782401473699 | 2.30885026305673 | -4.29737046375033 | 0.0008108032921 | 0.00312403623066736 |
| Crtap | ENSMUSG00000032431 | 56693 | 0.651990548674683 | 2.84733293448206 | 4.29711221189542 | 0.0008111984837 | 0.00312445758473992 |
| Prps1l3 | ENSMUSG00000079104 | 328099 | -0.373881113120704 | 3.12216296683499 | -4.29560125656465 | 0.0008135146096 | 0.00313227481458974 |
| Ccdc124 | ENSMUSG00000007721 | 234388 | -0.381419276706318 | 5.00761678303987 | -4.29537814307655 | 0.0008138571941 | 0.00313236233386509 |
| Mprip | ENSMUSG00000005417 | 26936 | -0.265620010162717 | 7.11700753589701 | -4.29521326466835 | 0.0008141104556 | 0.00313236233386509 |
| Zdhhc20 | ENSMUSG00000021969 | 75965 | 0.564629847979981 | 2.92862325790317 | 4.29365125935697 | 0.0008165138025 | 0.00314050400279714 |
| Arfrp1 | ENSMUSG00000038671 | 76688 | -0.470296132302054 | 3.02194677462355 | -4.29270286357286 | 0.0008179765938 | 0.00314502361852508 |
| Osmr | ENSMUSG00000022146 | 18414 | 0.745578643118293 | 3.60049583315612 | 4.29037510543504 | 0.0008215783381 | 0.00315776118657588 |
| Clec3b | ENSMUSG00000025784 | 21922 | 0.491897891304677 | 3.62028375021778 | 4.28940563111058 | 0.0008230832185 | 0.0031624332729239 |
| Hist2h2be | ENSMUSG00000068854 | 319190 | -0.659741125291614 | 1.41068312732643 | -4.28826124743557 | 0.0008248632558 | 0.00316815890703693 |
| Gucy1a1 | ENSMUSG00000033910 | 60596 | -0.75766672130935 | 3.09524143528991 | -4.28678474814132 | 0.0008271657390 | 0.00317588643973798 |
| Fitm1 | ENSMUSG00000022215 | 68680 | -0.391514628237497 | 5.53805271933776 | -4.28635741323133 | 0.0008278333673 | 0.00317733375647805 |
| 1810058I24Rik | ENSMUSG00000073155 | 67705 | 0.345390664224201 | 4.56787158203436 | 4.28524326632172 | 0.0008295766156 | 0.00318290697992929 |
| Tmem165 | ENSMUSG00000029234 | 21982 | 0.383560401788194 | 3.28161227586868 | 4.28202823366107 | 0.0008346282218 | 0.00320116529563682 |
| Ccnd3 | ENSMUSG00000034165 | 12445 | 0.5183422565741 | 3.72238819255151 | 4.28033364652878 | 0.0008373035633 | 0.00321029998272546 |
| Gvin1 | ENSMUSG00000045868 | 74558 | -1.05968053472512 | 5.05251863309949 | -4.27885289411543 | 0.0008396485358 | 0.00321816204238445 |
| Appl2 | ENSMUSG00000020263 | 216190 | -0.400847141954465 | 3.94714509466945 | -4.27864375149663 | 0.0008399802857 | 0.00321830511870199 |
| Cbr2 | ENSMUSG00000025150 | 12409 | 0.802729621454745 | 1.79927243336075 | 4.27723563795739 | 0.0008422174027 | 0.00322574577053153 |
| Pgap1 | ENSMUSG00000073678 | 241062 | 0.605321376939429 | 2.0286579852558 | 4.27621065406053 | 0.0008438496836 | 0.00323086546110611 |
| Gprc5c | ENSMUSG00000051043 | 70355 | 0.58612602983617 | 3.58710888597681 | 4.27472135365496 | 0.0008462271886 | 0.00323883382341553 |
| Ulk3 | ENSMUSG00000032308 | 71742 | 0.711118856734779 | 1.67351413918482 | 4.2741122249697 | 0.0008472015809 | 0.00324142823972568 |
| Oser1 | ENSMUSG00000035399 | 66680 | -0.431830392434347 | 3.1645742081946 | -4.27080895348939 | 0.0008525057973 | 0.00326058113028074 |
| 2810013P06Rik | ENSMUSG00000099881 | 100503178 | -0.549722843453843 | 1.5999053319694 | -4.2697820347083 | 0.0008541617211 | 0.00326577186897647 |
| Sgsm3 | ENSMUSG00000042303 | 105835 | -0.382603101645465 | 2.85985935478464 | -4.26879073878793 | 0.0008557633415 | 0.00327075142871291 |
| Igf1r | ENSMUSG00000005533 | 16001 | 0.379394673624476 | 4.67606105093237 | 4.26812645230291 | 0.0008568383468 | 0.00327371547337006 |
| Prkx | ENSMUSG00000035725 | 19108 | 0.521681249233672 | 3.18488439198644 | 4.26767674731885 | 0.0008575668865 | 0.00327535417065516 |
| Ust | ENSMUSG00000047712 | 338362 | 0.702313023564722 | 1.27887924448567 | 4.26353009325129 | 0.0008643147322 | 0.00329893207564059 |
| Hhipl1 | ENSMUSG00000021260 | 214305 | 1.01992036932771 | 1.93048073598754 | 4.2635123371427 | 0.0008643437439 | 0.00329893207564059 |
| Gjc1 | ENSMUSG00000034520 | 14615 | -0.63569908108395 | 2.09939780583471 | -4.26325190539738 | 0.0008647693775 | 0.00329940456057398 |
| Maip1 | ENSMUSG00000025971 | 68115 | -0.546335104988363 | 3.32410198185312 | -4.26292357100383 | 0.0008653062944 | 0.00330030115285751 |
| Glb1 | ENSMUSG00000045594 | 12091 | 0.940356253678728 | 1.01893002343865 | 4.25859553321936 | 0.0008724159624 | 0.00332625702318198 |
| Enpp5 | ENSMUSG00000023960 | 83965 | -0.760405980101578 | 1.7936148412814 | -4.25502174679072 | 0.0008783319012 | 0.00334764505329546 |
| Cpm | ENSMUSG00000020183 | 70574 | -0.635189245117898 | 2.41759013051417 | -4.25454586576247 | 0.0008791227676 | 0.00334890100078722 |
| Ptpmt1 | ENSMUSG00000063235 | 66461 | -0.447259284166873 | 3.45793230477378 | -4.25441173179034 | 0.0008793458170 | 0.00334890100078722 |
| Kank2 | ENSMUSG00000032194 | 235041 | 0.260158983310709 | 4.88215611872815 | 4.25427062159508 | 0.0008795805299 | 0.00334890100078722 |
| Abcd1 | ENSMUSG00000031378 | 11666 | -0.355042529712392 | 4.10533712763488 | -4.25379713396773 | 0.0008803685674 | 0.00335073426565812 |
| Ankrd6 | ENSMUSG00000040183 | 140577 | -0.669709100358133 | 1.36050814532912 | -4.25191376795678 | 0.0008835102968 | 0.00336152142511021 |
| Col27a1 | ENSMUSG00000045672 | 373864 | 1.20070835467672 | 1.15654252045724 | 4.25043343790136 | 0.0008859877955 | 0.00336977473667057 |
| Myh1 | ENSMUSG00000056328 | 17879 | -0.804036892464524 | 12.0634396905189 | -4.24996116031169 | 0.0008867797076 | 0.00337058208787186 |
| Vasn | ENSMUSG00000039646 | 246154 | 0.718013217932181 | 2.57462524116989 | 4.2499390693658 | 0.0008868167674 | 0.00337058208787186 |
| Lrig3 | ENSMUSG00000020105 | 320398 | 0.607931771448659 | 2.11164469108736 | 4.2461378879616 | 0.0008932174064 | 0.00339372939506745 |
| Atp8b1 | ENSMUSG00000039529 | 54670 | 0.835485794483667 | 2.91936540445018 | 4.24540116529462 | 0.0008944634267 | 0.00339728273706721 |
| Rhobtb2 | ENSMUSG00000022075 | 246710 | -0.370446783936582 | 4.17513069167625 | -4.24486993867113 | 0.0008953630001 | 0.00339951821984437 |
| Oard1 | ENSMUSG00000040771 | 106821 | -0.575296484676894 | 2.74960942149304 | -4.24384447161142 | 0.0008971021473 | 0.00340477753712044 |
| Adamts7 | ENSMUSG00000032363 | 108153 | 1.0847689816366 | 0.769442990834784 | 4.24368603769946 | 0.0008973711540 | 0.00340477753712044 |
| Gfra1 | ENSMUSG00000025089 | 14585 | -0.978386729075948 | 0.841940070470589 | -4.24331613103205 | 0.0008979995454 | 0.00340597953884539 |
| Cep192 | ENSMUSG00000024542 | 70799 | 0.603913645250925 | 2.18471525528213 | 4.24248182370099 | 0.0008994185148 | 0.00341017821218343 |
| Cox16 | ENSMUSG00000091803 | 66272 | -0.41912435748158 | 3.18105509802162 | -4.23903226735192 | 0.0009053099526 | 0.0034313256214385 |
| Pde8b | ENSMUSG00000021684 | 218461 | -1.00070567105423 | 1.01596245165229 | -4.23828833167986 | 0.0009065857005 | 0.00343496994559675 |
| Gsap | ENSMUSG00000039934 | 212167 | 0.845149548297812 | 1.13177428029589 | 4.23796519114958 | 0.0009071404180 | 0.00343557617953406 |
| Podn | ENSMUSG00000028600 | 242608 | 0.634445230437495 | 3.82684142706735 | 4.23782901070424 | 0.0009073742960 | 0.00343557617953406 |
| Mettl22 | ENSMUSG00000039345 | 239706 | -0.37119056910514 | 4.62267060120615 | -4.23737531575867 | 0.0009081539247 | 0.00343733744869778 |
| Smndc1 | ENSMUSG00000025024 | 76479 | -0.253155362338138 | 4.09737392872048 | -4.2362164940676 | 0.0009101483700 | 0.00344293067762697 |
| Tbx4 | ENSMUSG00000000094 | 21387 | 0.760185656175286 | 1.77620149670935 | 4.23615077723662 | 0.0009102616099 | 0.00344293067762697 |
| Cyhr1 | ENSMUSG00000053929 | 54151 | -0.2769011493088 | 5.62865776604034 | -4.2351879452821 | 0.0009119223752 | 0.00344801919200391 |
| Naa10 | ENSMUSG00000031388 | 56292 | -0.471809866944078 | 3.48370540397748 | -4.23421958252703 | 0.0009135958229 | 0.00345315212319535 |
| Eif3j2 | ENSMUSG00000043424 | 100042807 | -0.360743542634974 | 5.1649446288214 | -4.2333700114438 | 0.0009150665841 | 0.0034575156695026 |
| Sufu | ENSMUSG00000025231 | 24069 | 0.400063314215222 | 2.77165601192481 | 4.23293572640157 | 0.0009158193501 | 0.00345916424222406 |
| Pitpnm1 | ENSMUSG00000024851 | 18739 | 0.532160156328415 | 2.08758869662763 | 4.2324057218848 | 0.0009167388937 | 0.00346144139823765 |
| Cct3 | ENSMUSG00000001416 | 12462 | 0.2748707907342 | 5.33104776035952 | 4.23074557046779 | 0.0009196253598 | 0.00347096158924249 |
| Gab2 | ENSMUSG00000004508 | 14389 | 0.447737067669752 | 3.98183528301844 | 4.23043890244994 | 0.0009201595778 | 0.00347096158924249 |
| Nav3 | ENSMUSG00000020181 | 260315 | 0.522344299504652 | 2.92236633799553 | 4.23040833039965 | 0.0009202128520 | 0.00347096158924249 |
| Robo1 | ENSMUSG00000022883 | 19876 | 0.663160030604232 | 2.00568930524339 | 4.22918419240052 | 0.0009223486171 | 0.00347781743173206 |

| | | | | | | | |
|---|---|---|---|---|---|---|---|
| Plcg2 | ENSMUSG00000034330 | 234779 | 0.801806850524437 | 1.70491080603161 | 4.22470822757302 | 0.0009302013524 | 0.00350621758072523 |
| Atp13a1 | ENSMUSG00000031862 | 170759 | 0.409206844630338 | 3.45646112283285 | 4.22416650785241 | 0.0009311564091 | 0.00350860762085126 |
| D3Ertd751e | ENSMUSG00000025766 | 73852 | -0.830683678533352 | 1.65761089665249 | -4.22336806566888 | 0.0009325659077 | 0.00351270776610447 |
| Vsir | ENSMUSG00000020101 | 74048 | 0.551049799153062 | 3.59000913965613 | 4.22184748900066 | 0.0009352562665 | 0.00352162805721503 |
| Txnrd2 | ENSMUSG00000075704 | 26462 | -0.584652401669398 | 2.16345654856028 | -4.22163496189364 | 0.0009356329261 | 0.00352183316639236 |
| Clta | ENSMUSG00000028478 | 12757 | 0.260925400681116 | 5.53510582149436 | 4.22077522358489 | 0.0009371582229 | 0.00352636025299546 |
| Scaf11 | ENSMUSG00000033228 | 72193 | -0.223970612262604 | 6.23031766729708 | -4.21972727427917 | 0.0009390208895 | 0.0035321532496829 |
| Klhl2 | ENSMUSG00000031605 | 77113 | 0.44719468798002 | 2.64800273850565 | 4.21887143105259 | 0.0009405449218 | 0.00353666891649388 |
| Bzw1 | ENSMUSG00000051223 | 66882 | 0.293664911296388 | 5.34495016868677 | 4.21840049122633 | 0.0009413846262 | 0.00353860912973368 |
| Fancm | ENSMUSG00000055884 | 104806 | 0.699694821197513 | 1.56165838176738 | 4.21618074177868 | 0.0009453529149 | 0.00355230412989415 |
| Kank3 | ENSMUSG00000042099 | 80880 | -0.680246026137986 | 3.24907159395089 | -4.21477200738577 | 0.0009478802504 | 0.0035605769819494 |
| Ilk | ENSMUSG00000030890 | 16202 | 0.314608592821074 | 4.66801041358524 | 4.21288988392764 | 0.0009512677085 | 0.0035720739682766 |
| Efemp1 | ENSMUSG00000020467 | 216616 | -0.571370969053036 | 2.12331911221859 | -4.2117523178147 | 0.0009533211371 | 0.00357855540884738 |
| H2-T24 | ENSMUSG00000053835 | 15042 | -1.00741410637357 | 1.79322456587102 | -4.20992080986887 | 0.0009566367811 | 0.0035897688483009 |
| Gstz1 | ENSMUSG00000021033 | 14874 | -0.376182091928653 | 4.03429072427344 | -4.20917149986847 | 0.0009579966975 | 0.00359363826383086 |
| Rab4a | ENSMUSG00000019478 | 19341 | -0.630063296682492 | 2.41305812354766 | -4.2078217238228 | 0.0009604514092 | 0.00360161041322327 |
| Hgs | ENSMUSG00000025793 | 15239 | 0.313273939922781 | 4.63725360468567 | 4.20726813298954 | 0.0009614600414 | 0.00360415628007368 |
| Ftl1 | ENSMUSG00000050708 | 14325 | 0.457967356955476 | 6.49192942584745 | 4.20595115948525 | 0.0009638639195 | 0.00361192886700101 |
| Mgat5 | ENSMUSG00000036155 | 107895 | 0.83562663462426 | 2.39908765797028 | 4.20547360995868 | 0.0009647371180 | 0.00361396211027328 |
| Tmem229b | ENSMUSG00000046157 | 268567 | 0.527419698917255 | 2.33334522748202 | 4.20410048656597 | 0.0009672524034 | 0.00362214320738196 |
| Art3 | ENSMUSG00000034842 | 109979 | -0.576455938611762 | 4.75683668580642 | -4.20348113698856 | 0.0009683891313 | 0.0036251580802206 |
| Txn2 | ENSMUSG00000005354 | 56551 | -0.308158243948347 | 6.36451528595406 | -4.20173456736378 | 0.0009716021098 | 0.00363594065842 |
| Naga | ENSMUSG00000022453 | 17939 | 0.457184151341609 | 2.28457673948112 | 4.20061734806796 | 0.0009736630816 | 0.00364169145294833 |
| Dgkd | ENSMUSG00000070738 | 227333 | 0.260927335923569 | 5.30015588340595 | 4.20054042357562 | 0.0009738051520 | 0.00364169145294833 |
| Shld2 | ENSMUSG00000041471 | 75698 | -0.637133262733671 | 2.18037048873424 | -4.20033463256925 | 0.0009741853282 | 0.00364186724438162 |
| Eif2ak1 | ENSMUSG00000029613 | 15467 | -0.304892673558787 | 4.6867546839624 | -4.20003127026134 | 0.0009747460346 | 0.00364271757437597 |
| Armcx3 | ENSMUSG00000049047 | 71703 | 0.333650921576821 | 4.86844991471537 | 4.19644459300148 | 0.0009814005029 | 0.00366633250092031 |
| Mto1 | ENSMUSG00000032342 | 68291 | -0.363944511432397 | 3.50039130612695 | -4.19615516899135 | 0.0009819395108 | 0.00366709285726409 |
| Rps6ka3 | ENSMUSG00000031309 | 110651 | 0.272818023194088 | 5.60484535525141 | 4.19355438112163 | 0.0009867967471 | 0.00368397378509461 |
| Txlng | ENSMUSG00000038344 | 353170 | -0.350864000240035 | 3.07986065100934 | -4.19243556239622 | 0.0009888938416 | 0.00369054236345009 |
| Sept6 | ENSMUSG00000050379 | 56526 | 0.704559716349539 | 1.53201916393982 | 4.19118150843972 | 0.0009912498612 | 0.0036978553012432 |
| Lyrm7 | ENSMUSG00000020268 | 75530 | -0.598011174971692 | 2.14328135213088 | -4.19103262723048 | 0.0009915299504 | 0.0036978553012432 |
| Exoc3 | ENSMUSG00000034152 | 211446 | -0.267313051629655 | 5.34946230778675 | -4.19028564226855 | 0.0009929364757 | 0.00370183786353655 |
| Cfap298 | ENSMUSG00000022972 | 68001 | -0.47212695620635 | 3.49095829322742 | -4.188561952911 | 0.0009961899001 | 0.00371270091995811 |
| Fbxl14 | ENSMUSG00000030019 | 101358 | 0.402673630534689 | 2.76241616880143 | 4.18828469752791 | 0.0009967142347 | 0.00371338899125337 |
| Fut11 | ENSMUSG00000039357 | 73068 | 0.703190042218454 | 2.14683365671604 | 4.18693326689938 | 0.0009992740695 | 0.00372165753142255 |
| Rpa1 | ENSMUSG00000000751 | 68275 | -0.28904448171874 | 4.89402626551589 | -4.18617753031029 | 0.0010007085037 | 0.00372468764815737 |
| Plagl1 | ENSMUSG00000019817 | 22634 | 0.635425326173503 | 1.23572077972482 | 4.18611122882976 | 0.0010008344488 | 0.00372468764815737 |
| Dcun1d3 | ENSMUSG00000048787 | 233805 | 0.384332835347086 | 3.9407096699959 | 4.18596625287398 | 0.0010011098993 | 0.00372468764815737 |
| Rnf2 | ENSMUSG00000026484 | 19821 | -0.558525710259856 | 3.77810936078234 | -4.1847569931823 | 0.0010034104967 | 0.00373197691060355 |
| Fyco1 | ENSMUSG00000025241 | 17281 | -0.325272901505784 | 9.2447696515203 | -4.18328394604798 | 0.0010062202836 | 0.00374115439488125 |
| Copb1 | ENSMUSG00000030754 | 70349 | 0.289851432001353 | 4.52821098924002 | 4.18305334845003 | 0.0010066608717 | 0.00374151988735363 |
| Rock2 | ENSMUSG00000020580 | 19878 | 0.283313924377049 | 7.53476672697623 | 4.1823954093077 | 0.0010079190433 | 0.00374398946201408 |
| Wbp1 | ENSMUSG00000030035 | 22377 | -0.431098492048958 | 3.27295895302392 | -4.18234770201694 | 0.0010080103363 | 0.00374398946201408 |
| Hhatl | ENSMUSG00000032523 | 74770 | -0.357768409343328 | 4.62120404340517 | -4.18165248918672 | 0.0010093416650 | 0.00374766091723451 |
| Tmem62 | ENSMUSG00000054484 | 96957 | 0.71267390362732 | 1.63729010326256 | 4.18121370935917 | 0.0010101828554 | 0.00374951062559563 |
| Map4k2 | ENSMUSG00000024948 | 26412 | -0.561752296644088 | 2.56943983071554 | -4.1807866003055 | 0.0010110023631 | 0.00375127862579703 |
| Gnai2 | ENSMUSG00000032562 | 14678 | 0.354072404420573 | 6.2537179635069 | 4.17971212671522 | 0.0010130670107 | 0.00375766389351799 |
| AI597479 | ENSMUSG00000010290 | 98404 | -0.330203591403325 | 4.04763262161107 | -4.17914206268926 | 0.0010141641711 | 0.0037604574474153 |
| Gm14326 | ENSMUSG00000078862 | 665211 | -0.389759853288777 | 3.42992048067552 | -4.17786844032492 | 0.0010166198298 | 0.00376828462527743 |
| F3 | ENSMUSG00000028128 | 14066 | -0.751389623613574 | 2.01503205084534 | -4.17743680944955 | 0.0010174534378 | 0.00377009611139729 |
| Alpk2 | ENSMUSG00000032845 | 225638 | -0.408195951918132 | 6.26864619974686 | -4.17682769695299 | 0.0010186310101 | 0.0037731804718202 |
| Rab7 | ENSMUSG00000079477 | 19349 | -0.299846011128804 | 6.69024583656344 | -4.1752748092848 | 0.0010216394815 | 0.00378304240259756 |
| Efnb1 | ENSMUSG00000031217 | 13641 | -0.6114506485266 | 2.94599893014711 | -4.17294943076394 | 0.0010261616027 | 0.00379850067024979 |
| Tecta | ENSMUSG00000037705 | 21683 | -1.64979883512576 | 0.995943506745638 | -4.17247834491127 | 0.0010270802158 | 0.00379989025233617 |
| Ntrk2 | ENSMUSG00000055254 | 18212 | 0.548842656050319 | 2.54267843518949 | 4.17240042139767 | 0.0010272322473 | 0.00379989025233617 |
| Cybc1 | ENSMUSG00000039294 | 217370 | 0.631443093895207 | 1.99141678881667 | 4.17137964892603 | 0.0010292259457 | 0.00380597727108454 |
| Dnajc30 | ENSMUSG00000061118 | 66114 | -0.509024289476643 | 3.21484108433324 | -4.16969857750116 | 0.0010325179562 | 0.00381640192959964 |
| Mxd4 | ENSMUSG00000037235 | 17122 | 0.428842521325598 | 4.91469342054617 | 4.16958371152209 | 0.0010327432904 | 0.00381640192959964 |
| Prkag1 | ENSMUSG00000067713 | 19082 | -0.372084986376882 | 4.3940092255717 | -4.16887321527429 | 0.0010341382030 | 0.00382026519018743 |
| Kars | ENSMUSG00000031948 | 85305 | 0.303673447195287 | 6.04857339012171 | 4.1680860731852 | 0.0010356858530 | 0.00382468988851344 |
| Sod1 | ENSMUSG00000022982 | 20655 | -0.275311842818349 | 5.82609027339802 | -4.16680716959112 | 0.0010382054593 | 0.00383269972193233 |
| Pdlim4 | ENSMUSG00000020388 | 30794 | 0.696007593069681 | 2.06550829555688 | 4.16533535675435 | 0.0010411129128 | 0.00384213546596948 |
| Mtx2 | ENSMUSG00000027099 | 53375 | -0.445662824435991 | 3.89897343800197 | -4.16273851381247 | 0.0010462631810 | 0.00385983895799279 |
| Zdhhc14 | ENSMUSG00000034265 | 224454 | 0.647471055996061 | 1.52087001960333 | 4.16210908800825 | 0.0010475154428 | 0.00386275380671569 |
| Ostf1 | ENSMUSG00000024725 | 20409 | 0.383756047598149 | 4.0227407449992 | 4.16198623424447 | 0.0010477600436 | 0.00386275380671569 |
| Htatip2 | ENSMUSG00000039745 | 53415 | 0.970984388375402 | 1.43710481667058 | 4.16067213866104 | 0.0010503800671 | 0.0038695958872591 |
| Ephb2 | ENSMUSG00000028664 | 13844 | 1.14080824218564 | 1.20525335037295 | 4.16064233495326 | 0.0010504395672 | 0.0038695958872591 |
| Rmdn2 | ENSMUSG00000036368 | 381110 | -0.545878369812563 | 2.04806133898288 | -4.16052294957853 | 0.0010506779428 | 0.0038695958872591 |
| Msl3 | ENSMUSG00000031358 | 17692 | -0.368593239484033 | 4.36891565013831 | -4.16031716636469 | 0.0010510889585 | 0.00386980579535006 |
| Gnl3l | ENSMUSG00000025266 | 237107 | 0.255683875089281 | 4.90975465517044 | 4.1582036210298 | 0.0010553199594 | 0.00388407490784718 |
| Gpx7 | ENSMUSG00000028597 | 67305 | 0.916651310816333 | 1.15620956675764 | 4.15552798993126 | 0.0010607012606 | 0.00390193036524013 |
| Parp6 | ENSMUSG00000025237 | 67287 | -0.313119401888335 | 3.67626680265952 | -4.15543673820456 | 0.0010608852845 | 0.00390193036524013 |
| Phc1 | ENSMUSG00000040669 | 13619 | -0.267800453104531 | 3.9167245050292 | -4.1538327335588 | 0.0010641253760 | 0.00391253094025944 |
| Ick | ENSMUSG00000009828 | 56542 | 0.37050278892992 | 3.55450771503295 | 4.15350971823771 | 0.0010647790952 | 0.00391361812037208 |
| Tjap1 | ENSMUSG00000012296 | 74094 | 0.280523847657288 | 4.47211446004697 | 4.1531397015985 | 0.0010655284424 | 0.00391436123728361 |
| Pon3 | ENSMUSG00000029759 | 269823 | -0.399971119880109 | 2.94321384552061 | -4.15305627528854 | 0.0010656974697 | 0.00391436123728361 |
| Kera | ENSMUSG00000019932 | 16545 | -1.37229680792114 | 1.51004395712827 | -4.15263986583718 | 0.0010665415552 | 0.0039161456970098 |
| Odc1 | ENSMUSG00000011179 | 18263 | 0.472424241984744 | 5.68602593643651 | 4.15237207448513 | 0.0010670847459 | 0.00391682449887612 |
| Mdm4 | ENSMUSG00000054387 | 17248 | 0.323559979663564 | 4.31228097834295 | 4.14890031899766 | 0.0010741526499 | 0.00394144431577565 |
| Zkscan5 | ENSMUSG00000055991 | 22757 | -0.38744664577676 | 3.52050029894698 | -4.14840915548032 | 0.0010751564482 | 0.00394269262090481 |
| Smim11 | ENSMUSG00000051989 | 68936 | -0.316132288836272 | 4.73831748423808 | -4.14838089892153 | 0.0010752142258 | 0.00394269262090481 |
| Irf5 | ENSMUSG00000029771 | 27056 | 0.69246270355211 | 2.58788853809831 | 4.14775658914738 | 0.0010764915974 | 0.00394605286770706 |
| Cd151 | ENSMUSG00000025510 | 12476 | 0.274510154132861 | 5.3510326418134 | 4.14697010018817 | 0.0010781030137 | 0.00395063494566467 |
| Mrpl48 | ENSMUSG00000030706 | 52443 | -0.678728121010495 | 4.84409043915528 | -4.14632800375897 | 0.0010794204235 | 0.00395413694700167 |
| Cmtm3 | ENSMUSG00000031875 | 68119 | 0.833019921669238 | 1.70634525749088 | 4.14526506734988 | 0.0010816049168 | 0.00396081184132328 |
| Ascc1 | ENSMUSG00000044475 | 69090 | -0.530048231169318 | 2.28646943671704 | -4.1441880096591 | 0.0010838230599 | 0.0039676054480262 |
| Bmpr1b | ENSMUSG00000052430 | 12167 | -0.614819763156415 | 3.14253520020193 | -4.14371557739934 | 0.0010847974815 | 0.00396984307679373 |
| Ywhah | ENSMUSG00000018965 | 22629 | 0.297301820050702 | 4.959250674989 | 4.14297047075191 | 0.0010863361390 | 0.00397316497094398 |
| Traf4 | ENSMUSG00000017386 | 22032 | 0.838627585946442 | 1.55663063166974 | 4.14292400080713 | 0.0010864321743 | 0.00397316497094398 |
| Klhl21 | ENSMUSG00000073700 | 242785 | -0.430807558748753 | 4.76653567784622 | -4.14211912319968 | 0.0010880969275 | 0.00397792224588127 |
| Fam13b | ENSMUSG00000036501 | 225358 | 0.475685072954987 | 3.52385025110025 | 4.14148799234271 | 0.0010894041460 | 0.00398136968255986 |
| Usp31 | ENSMUSG00000063317 | 76179 | 0.326762096602208 | 4.4663072333707 | 4.13827664710129 | 0.0010960805944 | 0.00400225139123467 |
| Taf7 | ENSMUSG00000051316 | 24074 | -0.411814070159316 | 3.59693660516287 | -4.13823838256931 | 0.0010961603997 | 0.00400225139123467 |
| Cep76 | ENSMUSG00000073542 | 225659 | 0.529725557654155 | 2.52105121720596 | 4.13797823532491 | 0.0010967031265 | 0.00400225139123467 |
| Pabpc4 | ENSMUSG00000011257 | 230721 | -0.231497553720811 | 7.35535747849347 | -4.13791999465089 | 0.0010968246677 | 0.00400225139123467 |
| Pum3 | ENSMUSG00000041360 | 52874 | 0.30174398665623 | 4.14378359112658 | 4.13786061409766 | 0.0010969486019 | 0.00400225139123467 |
| Nav1 | ENSMUSG00000009418 | 215690 | 0.589371859708961 | 5.08457690048858 | 4.13634780176504 | 0.0011001108704 | 0.00401140360712508 |
| Cebpg | ENSMUSG00000056216 | 12611 | -0.281775045319966 | 4.87730465593099 | -4.13630951776932 | 0.0011001910176 | 0.00401140360712508 |
| Stmn2 | ENSMUSG00000027500 | 20257 | -0.886761293890736 | 1.42591104866444 | -4.13527307585539 | 0.0011023630782 | 0.00401798293051351 |
| S1pr2 | ENSMUSG00000043895 | 14739 | 0.876350275934006 | 1.42688625856604 | 4.13271031146227 | 0.0011077527486 | 0.00403628176504939 |
| Rfc4 | ENSMUSG00000022881 | 106344 | -0.783004563609205 | 1.17974967670024 | -4.13029569489472 | 0.0011128555930 | 0.00405294017078861 |
| Snhg1 | ENSMUSG00000108414 | 83673 | 0.580736041265237 | 1.62449773903244 | 4.13019629232615 | 0.0011130661780 | 0.00405294017078861 |
| Pycr1 | ENSMUSG00000025140 | 209027 | 0.983598984688301 | 0.608357135170407 | 4.1291602686333 | 0.0011152634366 | 0.00405958862015145 |
| Mrpl16 | ENSMUSG00000024683 | 94063 | -0.395320978254788 | 4.08141567238561 | -4.12848124445315 | 0.0011167059639 | 0.00406348631550495 |
| Zfp319 | ENSMUSG00000046556 | 79233 | 0.393963298387841 | 3.27385989639522 | 4.12793915020629 | 0.0011178589696 | 0.00406632825199875 |
| Rps13 | ENSMUSG00000090862 | 68052 | -0.310621377051495 | 5.82957538628542 | -4.1270174332322 | 0.0011198222174 | 0.00407211465687077 |
| Snx27 | ENSMUSG00000028136 | 76742 | -0.245275388445987 | 4.37505355942583 | -4.1250118468512 | 0.0011241063280 | 0.00408633397788112 |
| Mrps7 | ENSMUSG00000046756 | 50529 | -0.355742882331676 | 4.84813606278634 | -4.12424234603237 | 0.0011257545101 | 0.00409096494346155 |
| Id2 | ENSMUSG00000020644 | 15902 | 0.878587434222437 | 1.67325348004842 | 4.12020650185893 | 0.0011344395111 | 0.00412115596416797 |
| Itsn1 | ENSMUSG00000022957 | 16443 | -0.249764990552268 | 6.54548659518486 | -4.12001880936123 | 0.0011348450877 | 0.00412125968564655 |
| Hnrnph3 | ENSMUSG00000020069 | 432467 | -0.335456680382861 | 3.61223438813617 | -4.11862107626872 | 0.0011378700704 | 0.00413087271338299 |
| 4931406P16Rik | ENSMUSG00000066571 | 233103 | -0.401543899561748 | 5.41597869456038 | -4.11839824782313 | 0.0011383530810 | 0.00413125415952258 |
| Sema4a | ENSMUSG00000028064 | 20351 | 0.919182740961282 | 0.97437494071745 | 4.11801095939139 | 0.0011391930811 | 0.00413293049123255 |
| Atad2 | ENSMUSG00000022360 | 70472 | 0.528361089853124 | 2.1638139621505 | 4.11739995425775 | 0.0011405195993 | 0.00413635004172495 |
| Hist1h2ap | ENSMUSG00000094777 | 319171 | 1.22781768876233 | 1.62667158639773 | 4.11722834724804 | 0.0011408924504 | 0.00413635004172495 |

| | | | | | | | |
|---|---|---|---|---|---|---|---|
| Dnajc18 | ENSMUSG00000024350 | 76594 | 0.313222277100754 | 4.47730548403486 | 4.11537900120577 | 0.0011449184726 | 0.00414957023355214 |
| Zfp869 | ENSMUSG00000054648 | 66869 | -0.340702432589761 | 3.58248511538981 | -4.11083096053256 | 0.0011548816481 | 0.00418429277300302 |
| Dusp3 | ENSMUSG00000003518 | 72349 | 0.345454537258827 | 5.85171814499113 | 4.10906208620023 | 0.0011587806149 | 0.00419702813201932 |
| R3hcc1l | ENSMUSG00000025184 | 52013 | -0.394380188872556 | 3.63375553667698 | -4.10829729492289 | 0.0011604705516 | 0.00420175674084473 |
| Arl2 | ENSMUSG00000024944 | 56327 | -0.479027000381279 | 3.5531985215432 | -4.10811249997613 | 0.0011608792666 | 0.0042018447893118 |
| Tcirg1 | ENSMUSG00000001750 | 27060 | 0.628674163798714 | 2.33092990206782 | 4.10474234567732 | 0.0011683590599 | 0.00422751833321555 |
| Nudt3 | ENSMUSG00000024213 | 56409 | -0.322358960385041 | 5.91359253946127 | -4.10323807052966 | 0.0011717136164 | 0.0042382533226787 |
| Capn1 | ENSMUSG00000024942 | 12333 | 0.355228055959712 | 5.2195397394342 | 4.09993354460056 | 0.0011791174820 | 0.00426362328679477 |
| Trim63 | ENSMUSG00000028834 | 433766 | -0.367913814789945 | 6.40215702187146 | -4.09954901859276 | 0.0011799821315 | 0.00426533884921227 |
| Map1lc3a | ENSMUSG00000027602 | 66734 | 0.359402149878548 | 6.1798840242304 | 4.09807432263673 | 0.0011833041816 | 0.00427593322648364 |
| Tmem86a | ENSMUSG00000010307 | 67893 | 0.687405038751534 | 2.23184731696117 | 4.09712361786653 | 0.0011854509184 | 0.00428227494688844 |
| Nudt8 | ENSMUSG00000110949 | 66387 | -0.493357059709819 | 2.64631225400571 | -4.09670862019818 | 0.0011863892545 | 0.0042842487417853 |
| Nfatc1 | ENSMUSG00000033016 | 18018 | -0.283704640829495 | 5.34065110366392 | -4.09605895225068 | 0.0011878597245 | 0.00428814222228408 |
| Hsbp1 | ENSMUSG00000031839 | 68196 | -0.222941515993775 | 5.36756583819604 | -4.0958029978096 | 0.0011884395694 | 0.00428881906013951 |
| Vps37c | ENSMUSG00000048832 | 107305 | 0.334635312944172 | 4.34609361277575 | 4.09299355405998 | 0.0011948232522 | 0.00431043332359957 |
| Il15ra | ENSMUSG00000023206 | 16169 | -0.697430985735489 | 2.65442915340035 | -4.09099073548589 | 0.0011993955577 | 0.00432550077258204 |
| Rpn1 | ENSMUSG00000030062 | 103963 | 0.282666879867699 | 5.38966467218461 | 4.09080293732025 | 0.0011998252076 | 0.0043256231348169 |
| Arf4 | ENSMUSG00000021877 | 11843 | 0.463669730815935 | 4.573400035323 | 4.0902260613365 | 0.0012011459876 | 0.00432789258988822 |
| Rnf122 | ENSMUSG00000039328 | 68867 | 0.936397431311458 | 2.14188202837901 | 4.09018216228198 | 0.0012012465572 | 0.00432789258988822 |
| Mmadhc | ENSMUSG00000026766 | 109129 | -0.331796110585736 | 4.76853378556911 | -4.08968037761786 | 0.0012023967236 | 0.00433060908925325 |
| Mettl14 | ENSMUSG00000028114 | 210529 | -0.355765775313834 | 3.58933743576641 | -4.08891938307253 | 0.0012041431929 | 0.00433547076495388 |
| Itga8 | ENSMUSG00000026768 | 241226 | 0.907590034367434 | 2.33915216670076 | 4.08680255769065 | 0.0012090149609 | 0.00435157805023528 |
| Pmm1 | ENSMUSG00000022474 | 29858 | 0.405373801078656 | 3.0491917482322 | 4.08620844453362 | 0.0012103859106 | 0.00435507846907342 |
| Tet3 | ENSMUSG00000034832 | 194388 | 0.416341039549545 | 4.84527992423644 | 4.0818783953573 | 0.0012204259944 | 0.00438975865256635 |
| Maf1 | ENSMUSG00000022553 | 68877 | -0.340283432323 | 4.54668753009113 | -4.08103805646965 | 0.0012223843578 | 0.00439535638681944 |
| Pcdh12 | ENSMUSG00000024440 | 53601 | -0.637834552438875 | 2.71692969099284 | -4.08008078053706 | 0.0012246191615 | 0.00440062380676367 |
| St3gal1 | ENSMUSG00000013846 | 20442 | 0.594092883149749 | 3.72612802296315 | 4.07994349419552 | 0.0012249400058 | 0.00440062380676367 |
| Srpr | ENSMUSG00000032042 | 67398 | -0.262616667717421 | 5.74333418325849 | -4.07989343570656 | 0.0012250570161 | 0.00440062380676367 |
| Pnpo | ENSMUSG00000018659 | 103711 | -0.617782488450417 | 1.52369695101506 | -4.07808328329118 | 0.0012292959041 | 0.00441439997644029 |
| Arhgap29 | ENSMUSG00000039831 | 214137 | -0.528971042677733 | 4.89849677461823 | -4.07720647361304 | 0.0012313545568 | 0.00441937018357153 |
| Slc41a1 | ENSMUSG00000013275 | 98396 | -0.472758740073597 | 6.13273524984311 | -4.07714944681075 | 0.0012314885718 | 0.00441937018357153 |
| Pkm | ENSMUSG00000032294 | 18746 | -0.32314244167307 | 9.19243403391451 | -4.07661989854445 | 0.0012327337429 | 0.00442238678822227 |
| Cops4 | ENSMUSG00000035297 | 26891 | -0.297396344747689 | 5.43842240636161 | -4.07545301054799 | 0.0012354821003 | 0.00442999972414173 |
| Trim8 | ENSMUSG00000025034 | 93679 | 0.320357217188385 | 5.21001912414224 | 4.07523872304193 | 0.0012359874912 | 0.00442999972414173 |
| Gng10 | ENSMUSG00000038607 | 14700 | 0.702285882800195 | 1.52388384796125 | 4.0751250308004 | 0.0012362557171 | 0.00442999972414173 |
| B3galnt2 | ENSMUSG00000039242 | 97884 | -0.307057129782383 | 4.27790543440011 | -4.07490706510419 | 0.0012367701146 | 0.00442999972414173 |
| 4632404H12Rik | ENSMUSG00000042579 | 74034 | -0.621291888958132 | 2.09142117894328 | -4.07485959010101 | 0.0012368821844 | 0.00442999972414173 |
| Pias2 | ENSMUSG00000025423 | 17344 | -0.340379587434544 | 4.29382222171833 | -4.07394287652366 | 0.0012390482256 | 0.00443630401411946 |
| Chd7 | ENSMUSG00000041235 | 320790 | -0.353535904217907 | 5.62000442912643 | -4.07368539697845 | 0.0012396573058 | 0.00443703143766071 |
| Cul2 | ENSMUSG00000024231 | 71745 | -0.287861165296286 | 4.99234398280459 | -4.073443457051 | 0.0012402299059 | 0.00443762785668958 |
| Rbms2 | ENSMUSG00000040043 | 56516 | -0.289939912227285 | 4.65526593945319 | -4.07288995518755 | 0.0012415409003 | 0.00443925396526429 |
| Zrsr1 | ENSMUSG00000044068 | 22183 | -0.321268007857459 | 3.70109375956742 | -4.07287682419751 | 0.0012415720189 | 0.00443925396526429 |
| Spire1 | ENSMUSG00000024533 | 68166 | 0.621188165810942 | 3.46575100877434 | 4.07273730064291 | 0.0012419027194 | 0.00443925396526429 |
| Sipa1l3 | ENSMUSG00000030583 | 74206 | 0.390281541924973 | 3.40225765319293 | 4.07205588208685 | 0.0012435191234 | 0.00444105282371334 |
| Smo | ENSMUSG00000001761 | 319757 | 0.427871423155901 | 3.06938361993883 | 4.07203645835808 | 0.0012435652304 | 0.00444105282371334 |
| Ndrg1 | ENSMUSG00000005125 | 17988 | -0.371166367819424 | 4.05511552113544 | -4.07192005096224 | 0.0012438415883 | 0.00444105282371334 |
| Ate1 | ENSMUSG00000030850 | 11907 | -0.268604510457124 | 4.56316643639391 | -4.07184024824804 | 0.0012440310809 | 0.00444105282371334 |
| Cbwd1 | ENSMUSG00000024878 | 226043 | -0.448082158567385 | 3.06649005624587 | -4.06908555121107 | 0.0012505903143 | 0.00446301101096892 |
| Ttyh2 | ENSMUSG00000034714 | 117160 | 0.682346912565028 | 2.67898833430831 | 4.06813400288567 | 0.0012528642806 | 0.00446966692286465 |
| Zfp961 | ENSMUSG00000052446 | 234413 | -0.933178914687194 | 1.05235732008656 | -4.06792579332661 | 0.0012533624149 | 0.00446998517377976 |
| Lrig1 | ENSMUSG00000030029 | 16206 | -0.305970518709058 | 4.70359147960845 | -4.06727467116506 | 0.0012549215126 | 0.00447408579740319 |
| Ankrd28 | ENSMUSG00000014496 | 105522 | -0.303524313240266 | 5.9516702407225 | -4.06667474639838 | 0.0012563597783 | 0.00447775309321334 |
| Fkbp3 | ENSMUSG00000020949 | 30795 | -0.33976730285567 | 6.76548750880564 | -4.06614253378057 | 0.0012576371240 | 0.00448084465533252 |
| Kif13b | ENSMUSG00000060012 | 16554 | 0.45982479974917 | 4.03349092219751 | 4.06571221039406 | 0.0012586709016 | 0.0044830666751286 |
| Tiam2 | ENSMUSG00000023800 | 24001 | 0.654368852337607 | 3.95014657958456 | 4.06529621293116 | 0.0012596710914 | 0.00448516765488941 |
| Myh2 | ENSMUSG00000033196 | 17882 | -0.535831268581294 | 13.7886696824854 | -4.06386408383806 | 0.0012631206156 | 0.00449598549316621 |
| Cacna1s | ENSMUSG00000026407 | 12292 | -0.371111523732775 | 9.02998089431234 | -4.06302812965561 | 0.0012651386242 | 0.00450170257212056 |
| Pmpcb | ENSMUSG00000029017 | 73078 | -0.339048687013841 | 5.4809663205696 | -4.06265287161235 | 0.0012660455779 | 0.00450346378531407 |
| Map3k14 | ENSMUSG00000020941 | 53859 | 0.696237360890375 | 3.13883852923648 | 4.0621258892414 | 0.0012673203558 | 0.00450653181836201 |
| Serhl | ENSMUSG00000058586 | 68607 | -0.542385017815085 | 2.71118525795046 | -4.06145484679635 | 0.0012689455208 | 0.0045108434107762 |
| Rp2 | ENSMUSG00000060090 | 19889 | 0.395364825081192 | 3.11672917872823 | 4.05967371189931 | 0.0012732695275 | 0.00452247642798623 |
| Atp8a1 | ENSMUSG00000037685 | 11980 | -0.394156023044787 | 3.71042233900264 | -4.05963690389211 | 0.0012733590442 | 0.00452247642798623 |
| Pak1 | ENSMUSG00000030774 | 18479 | 0.506528152817537 | 2.18630260710197 | 4.05959572703992 | 0.0012734591936 | 0.00452247642798623 |
| Smpdl3a | ENSMUSG00000019872 | 57319 | -0.366083514406641 | 4.14473422122588 | -4.05680106927483 | 0.0012802751853 | 0.00454520560276057 |
| Pop5 | ENSMUSG00000060152 | 117109 | -0.539858594356564 | 2.51827396331265 | -4.05579029737703 | 0.0012827495866 | 0.00455251160116395 |
| Tmem106b | ENSMUSG00000029571 | 71900 | -0.384703998136104 | 5.73656118630337 | -4.05451941899003 | 0.0012858676905 | 0.00456209663277594 |
| Jmy | ENSMUSG00000021690 | 57748 | -0.270747764098221 | 5.93246105131052 | -4.05386409082632 | 0.0012874785731 | 0.00456519563364368 |
| Lnpk | ENSMUSG00000009207 | 69605 | -0.301583957044651 | 4.74070739195988 | -4.05382430167574 | 0.0012875764466 | 0.00456519563364368 |
| Arhgef37 | ENSMUSG00000045094 | 328967 | 0.731498606030557 | 1.50967357470884 | 4.05220558706003 | 0.0012915646354 | 0.00457785117717652 |
| Lmod3 | ENSMUSG00000044086 | 320502 | -0.338567574762632 | 7.38612052389472 | -4.04816198606159 | 0.0013015826370 | 0.00461186379430355 |
| Pin1 | ENSMUSG00000032171 | 23988 | -0.315302261232487 | 4.51023235896281 | -4.04386118579057 | 0.0013123252220 | 0.00464842093395535 |
| Smim20 | ENSMUSG00000061461 | 66278 | -0.448284350832404 | 3.9691303731285 | -4.04239154969003 | 0.0013160168759 | 0.00465998719502137 |
| Trak1 | ENSMUSG00000032536 | 67095 | -0.28036358757995 | 7.85927745073861 | -4.04178166915506 | 0.0013175519869 | 0.00466391216619045 |
| Sfmbt1 | ENSMUSG00000006527 | 54650 | -0.359200344744635 | 3.28294991473387 | -4.04122070151299 | 0.0013189655999 | 0.004667404652919 |
| Asph | ENSMUSG00000028207 | 65973 | 0.261809941418044 | 6.92473960935719 | 4.04072424015562 | 0.0013202179555 | 0.00467032442468482 |
| Dcxr | ENSMUSG00000039450 | 67880 | -0.720635454247157 | 1.03299660778948 | -4.03855398151249 | 0.0013257068852 | 0.00468822451077033 |
| Sema6d | ENSMUSG00000027200 | 214968 | -0.373071191978291 | 3.65381478088564 | -4.03736563602451 | 0.0013287222960 | 0.0046973685050463 |
| Tm2d1 | ENSMUSG00000028563 | 94043 | -0.318832851849386 | 3.61440569218229 | -4.03697869521789 | 0.0013297056681 | 0.00469805545751643 |
| Ap5b1 | ENSMUSG00000049562 | 381201 | 0.648650072000726 | 1.31109388364602 | 4.03684394901696 | 0.0013300482873 | 0.00469805545751643 |
| Bcl3 | ENSMUSG00000053175 | 12051 | 1.23840892195637 | 1.03958523975459 | 4.0367819405205 | 0.0013302059867 | 0.00469805545751643 |
| Prmt9 | ENSMUSG00000037134 | 102182 | -0.377410911970331 | 3.29675820676446 | -4.03652835768067 | 0.0013308510953 | 0.00469881567288322 |
| Dnajc7 | ENSMUSG00000014195 | 56354 | -0.306539271675206 | 5.50981590911238 | -4.03630510480377 | 0.0013314193104 | 0.00469930399820072 |
| Tmem223 | ENSMUSG00000117924 | 66836 | -0.431296476651516 | 3.13848879638572 | -4.03465007203268 | 0.0013356393939 | 0.00471210416161289 |
| Il4ra | ENSMUSG00000030748 | 16190 | 0.933177889789854 | 3.36413421924605 | 4.03454489531684 | 0.0013359080410 | 0.00471210416161289 |
| Clasp1 | ENSMUSG00000064302 | 76707 | 0.253743614269832 | 6.09174712350207 | 4.03304538446614 | 0.0013397441828 | 0.00472411085907049 |
| Esam | ENSMUSG00000001946 | 69524 | -0.508933287476197 | 3.59703200086441 | -4.02952386818847 | 0.0013487975539 | 0.00475450050383386 |
| Cab39 | ENSMUSG00000036707 | 12283 | -0.372163032329889 | 7.29522232900564 | -4.02911393343916 | 0.0013498555064 | 0.0047557705051186 |
| Vhl | ENSMUSG00000033933 | 22346 | 0.591822587402915 | 1.50770987409713 | 4.02904713371072 | 0.0013500279825 | 0.0047557705051186 |
| Tk2 | ENSMUSG00000035824 | 57813 | 0.636571378044849 | 1.61087522754424 | 4.02749329429413 | 0.0013540463494 | 0.00476838938342269 |
| 2310040G24Rik | ENSMUSG00000101655 | 381792 | -0.562501554279878 | 3.86301067063767 | -4.02451154482489 | 0.0013617917480 | 0.0047941209656597 |
| Slc25a38 | ENSMUSG00000032519 | 208638 | -0.464674412697549 | 2.66292625558823 | -4.02401858455168 | 0.0013630766251 | 0.00479709935287444 |
| Cryz | ENSMUSG00000028199 | 12972 | -0.62913172258022 | 1.74775919773313 | -4.02281867298334 | 0.0013662093260 | 0.00480657680820801 |
| A730036I17Rik | ENSMUSG00000056738 | 329513 | -1.06473511624937 | 1.63146575736771 | -4.0220515567044 | 0.0013682159563 | 0.00481168344674451 |
| Usp22 | ENSMUSG00000042506 | 216825 | 0.2702294968433 | 4.43072347965743 | 4.02184086189894 | 0.0013687676218 | 0.00481168344674451 |
| Strn3 | ENSMUSG00000020954 | 94186 | -0.253795334350431 | 7.23892971030769 | -4.021704958082 | 0.0013691235817 | 0.00481168344674451 |
| Raf1 | ENSMUSG00000000441 | 110157 | -0.284193167313624 | 5.98419102912099 | -4.02159121461312 | 0.0013694215719 | 0.00481168344674451 |
| Bri3 | ENSMUSG00000047843 | 55950 | -0.403958699741664 | 2.74026208790071 | -4.02071220431811 | 0.0013717266826 | 0.00481823405139699 |
| Mpp5 | ENSMUSG00000021112 | 56217 | -0.257694424568458 | 4.8972054314608 | -4.02017324662546 | 0.0013731420055 | 0.00482165604322175 |
| Med24 | ENSMUSG00000017210 | 23989 | -0.275849592295275 | 4.73619412770529 | -4.01991737171038 | 0.0013738144657 | 0.00482246818409991 |
| Polr2c | ENSMUSG00000031783 | 20021 | -0.309319401795785 | 4.82886119751172 | -4.01947991966495 | 0.0013749649064 | 0.00482495710831673 |
| H2afy | ENSMUSG00000015937 | 26914 | -0.295321221218305 | 4.63915495528033 | -4.01740028239368 | 0.0013804475780 | 0.00484264200106239 |
| Eif2s2 | ENSMUSG00000074656 | 67204 | -0.370124767923754 | 6.02502196877251 | -4.01445633876149 | 0.0013882471569 | 0.00486844070330297 |
| Tmem136 | ENSMUSG00000048503 | 235300 | -0.875776715390682 | 1.05189033974495 | -4.01394990636993 | 0.0013895934208 | 0.00487159900026272 |
| Fv1 | ENSMUSG00000070583 | 14349 | -0.943291720127159 | 0.792660277627656 | -4.01295310460204 | 0.0013922471524 | 0.0048793374875439 |
| Mustn1 | ENSMUSG00000042485 | 66175 | 0.441708001406528 | 5.22038953081034 | 4.00884677376773 | 0.0014032340244 | 0.00491626638507711 |
| Sun2 | ENSMUSG00000042524 | 223697 | -0.26494146987425 | 6.19808046628219 | -4.00722449676608 | 0.0014075989934 | 0.0049299790444743 |
| Myh7 | ENSMUSG00000053093 | 140781 | 0.50167718703957 | 13.9361118337124 | 4.00688149248429 | 0.0014085236738 | 0.00493163750106416 |
| Galt | ENSMUSG00000036073 | 14430 | -0.408188824912989 | 3.99149394728739 | -4.00541784876176 | 0.0014124763952 | 0.00494389352427893 |
| Zcchc24 | ENSMUSG00000055538 | 71918 | 0.378073610701897 | 5.21601656223861 | 4.00496456777758 | 0.0014137028280 | 0.00494660230894571 |
| Trim65 | ENSMUSG00000054517 | 338364 | 0.521219980910266 | 2.3540295137431 | 4.00473293928049 | 0.0014143299609 | 0.00494721305715012 |
| Sptlc2 | ENSMUSG00000021036 | 20773 | 0.44642157652881 | 3.19912742251966 | 4.0044647624781 | 0.0014150564039 | 0.00494817068175961 |
| Cobll1 | ENSMUSG00000034903 | 319876 | -0.812017887225233 | 4.32314446406766 | -4.00403545102039 | 0.0014162201266 | 0.00495065628549098 |
| Best3 | ENSMUSG00000020169 | 382427 | -0.557296853612211 | 3.12055042195025 | -4.00336705312423 | 0.0014180338840 | 0.00495541188815207 |
| Slc20a2 | ENSMUSG00000037656 | 20516 | -0.284543540511644 | 5.80746459569236 | -4.00129448685287 | 0.0014236731037 | 0.0049735285064526 |
| Rab4b | ENSMUSG00000053291 | 19342 | 0.841675636587695 | 1.73153635398681 | 4.00069011446177 | 0.0014253218503 | 0.00497639611238337 |
| Glipr2 | ENSMUSG00000028480 | 384009 | 0.792228580893779 | 1.89814636568461 | 4.00063478725588 | 0.0014254728823 | 0.00497639611238337 |

| | | | | | | | |
|---|---|---|---|---|---|---|---|
| Vav3 | ENSMUSG00000033721 | 57257 | 0.703929240948816 | 1.92317868829953 | 4.00049310433496 | 0.0014258597222 | 0.00497639611238337 |
| Mrps17 | ENSMUSG00000034211 | 66258 | -0.296333240447479 | 3.8854911744448 | -3.99854092049275 | 0.0014312007612 | 0.00499264242851918 |
| Tsc22d4 | ENSMUSG00000029723 | 78829 | 0.436007441246174 | 5.5826774744512 | 3.99842006145998 | 0.0014315320951 | 0.00499264242851918 |
| Ccdc71l | ENSMUSG00000090946 | 72123 | 0.565038391455959 | 2.71183998298793 | 3.99829139635991 | 0.0014318849156 | 0.00499264242851918 |
| Atp5g2 | ENSMUSG00000062683 | 67942 | -0.316383177699184 | 4.96721573362389 | -3.99780595498872 | 0.0014332181109 | 0.00499569743960846 |
| Rbfox1 | ENSMUSG00000008658 | 268859 | -0.29674458982324 | 5.22060618201304 | -3.99743038937207 | 0.0014342482268 | 0.00499769441102579 |
| Ercc6l2 | ENSMUSG00000021470 | 76251 | -0.294956617144318 | 3.8570817527766 | -3.9952522401728 | 0.0014402446929 | 0.00501699003774287 |
| Trp53inp1 | ENSMUSG00000028211 | 60599 | 0.588590938762254 | 3.12132948789274 | 3.99466878796699 | 0.0014418552937 | 0.00501899775763301 |
| Usf2 | ENSMUSG00000058239 | 22282 | -0.270465741904717 | 5.52048375296401 | -3.99461150357091 | 0.0014420135247 | 0.00501899775763301 |
| Supt7l | ENSMUSG00000053134 | 72195 | -0.413909117937997 | 3.24439015639249 | -3.99454454004241 | 0.0014421985140 | 0.00501899775763301 |
| Gm21541 | ENSMUSG00000094065 | 100862177 | 1.27577003506933 | 0.472354870304666 | 3.99236186398353 | 0.0014482415491 | 0.00503755036603613 |
| Klf16 | ENSMUSG00000035397 | 118445 | 0.926426843996566 | 0.943063582966229 | 3.99228628362838 | 0.0014484512670 | 0.00503755036603613 |
| Prep | ENSMUSG00000019849 | 19072 | 0.262423755881524 | 4.75425511477599 | 3.99079105136299 | 0.0014526065812 | 0.00505039520990721 |
| Snord17 | ENSMUSG00000077714 | 100313519 | 1.16427630581148 | 4.41896288139339 | 3.98870931601068 | 0.0014584121289 | 0.00506896756164114 |
| Supv3l1 | ENSMUSG00000020079 | 338359 | -0.362950148418608 | 3.89577383576156 | -3.98716186728734 | 0.0014627430314 | 0.00508240434746501 |
| Nutf2 | ENSMUSG00000028450 | 68051 | -0.228453461189679 | 4.39652028558117 | -3.98603996736428 | 0.0014658911589 | 0.00509172426401001 |
| Slc16a10 | ENSMUSG00000019838 | 72472 | -0.559165675739492 | 3.33606374821263 | -3.98525702452853 | 0.0014680922548 | 0.00509774982150719 |
| Rangrf | ENSMUSG00000032892 | 57785 | -0.360331432339574 | 2.93349957232246 | -3.98271225394978 | 0.0014752697742 | 0.00511950638584403 |
| Hipk3 | ENSMUSG00000027177 | 15259 | -0.283350044957931 | 8.30352109927594 | -3.98270347913397 | 0.0014752945856 | 0.00511950638584403 |
| Npas2 | ENSMUSG00000026077 | 18143 | 1.2477806421923 | 2.05112564019517 | 3.98243629002288 | 0.0014760502839 | 0.00511994003919245 |
| Fxr1 | ENSMUSG00000027680 | 14359 | -0.295173865890122 | 8.25575025656962 | -3.98232812366451 | 0.0014763563263 | 0.00511994003919245 |
| Tcim | ENSMUSG00000056313 | 69068 | -1.15708156022746 | 3.05303119855092 | -3.98215301880558 | 0.0014768519000 | 0.00512003429099217 |
| Pigs | ENSMUSG00000041958 | 276846 | 0.399895561226402 | 2.98919101056334 | 3.98155665373825 | 0.0014785409817 | 0.00512426489260595 |
| Jak1 | ENSMUSG00000028530 | 16451 | 0.213899719339373 | 6.59542062536861 | 3.97935033553407 | 0.0014848071197 | 0.00514435075294483 |
| Ogn | ENSMUSG00000021390 | 18295 | 0.537979598942806 | 4.71748867837116 | 3.97865411927879 | 0.0014867900672 | 0.00514958879133872 |
| Sec23ip | ENSMUSG00000055319 | 207352 | 0.236603748843907 | 4.74415598178371 | 3.97797824036783 | 0.0014887176812 | 0.00515463192073821 |
| Zfp775 | ENSMUSG00000007216 | 243372 | -0.409470938354414 | 2.94475347550531 | -3.97747644578096 | 0.0014901504578 | 0.00515795904189938 |
| Zfhx4 | ENSMUSG00000025255 | 80892 | 0.382243549895203 | 5.37948745535244 | 3.97714682820399 | 0.0014910923838 | 0.00515958558007650 |
| Ier5l | ENSMUSG00000089762 | 72500 | 0.819421192933512 | 1.8447476448945 | 3.97503714076304 | 0.0014971355327 | 0.00517885712286981 |
| Ralgapb | ENSMUSG00000027652 | 228850 | -0.286646474255159 | 5.28866613789164 | -3.97346402004343 | 0.0015016579951 | 0.00519285781247512 |
| Sec11c | ENSMUSG00000024516 | 66286 | 0.442169835352758 | 3.17641047344072 | 3.97090666042407 | 0.0015090398167 | 0.00521536914640797 |
| Nol12 | ENSMUSG00000033099 | 97961 | 0.4614517552589 | 3.72657814338338 | 3.9707827997897 | 0.0015093982812 | 0.00521536914640797 |
| Fmo1 | ENSMUSG00000040181 | 14261 | -0.462432356067541 | 3.78338866926307 | -3.97071341693273 | 0.0015095991198 | 0.00521536914640797 |
| Ggnbp1 | ENSMUSG00000048731 | 70772 | -0.716222738100735 | 2.14680573270131 | -3.97032460408956 | 0.0015107251002 | 0.00521761013285005 |
| Chac2 | ENSMUSG00000020309 | 68044 | -0.561065246955179 | 2.63418642110549 | -3.96867309549995 | 0.0015155173554 | 0.00523250796364152 |
| Nrep | ENSMUSG00000042834 | 27528 | -0.47468815703324 | 4.76638277568525 | -3.96738979917327 | 0.0015192518822 | 0.00524374560311590 |
| Zfp317 | ENSMUSG00000057551 | 244713 | -0.260785595533685 | 3.59959162290741 | -3.96513334232457 | 0.0015258412405 | 0.00526482657849203 |
| Cnn3 | ENSMUSG00000053931 | 71994 | 0.484099533195163 | 4.93144718011254 | 3.96480613253962 | 0.0015267991891 | 0.00526646952859418 |
| Lamb3 | ENSMUSG00000026639 | 16780 | -0.65430187291403 | 1.44462005479155 | -3.96446068261731 | 0.0015278112052 | 0.00526829788146464 |
| Znrd1as | ENSMUSG00000036214 | 76416 | -0.543191615344344 | 2.68349728501969 | -3.96347290585631 | 0.0015307087416 | 0.00527650430739032 |
| Vcp | ENSMUSG00000028452 | 269523 | 0.252136548470987 | 7.94970916480848 | 3.96332043576722 | 0.0015311564964 | 0.00527650430739032 |
| Foxp1 | ENSMUSG00000030067 | 108655 | 0.402238469977998 | 4.44510350103565 | 3.96132984171104 | 0.0015370145206 | 0.00529502228979278 |
| Tent2 | ENSMUSG00000042167 | 100715 | 0.456101362908983 | 3.17128729037848 | 3.96072736688377 | 0.0015387920278 | 0.00529804913983610 |
| Ap1ar | ENSMUSG00000074238 | 211556 | -0.286143627361376 | 4.26677429579965 | -3.96070349447509 | 0.0015388625028 | 0.00529804913983610 |
| Setd1b | ENSMUSG00000038384 | 208043 | 0.380262592538728 | 4.68292452849298 | 3.95948791448242 | 0.0015424554484 | 0.00530874701086159 |
| Fbln1 | ENSMUSG00000006369 | 14114 | 0.701646100313709 | 3.34330483038654 | 3.95875932603951 | 0.0015446130797 | 0.00531449970887358 |
| Pdia4 | ENSMUSG00000025823 | 12304 | 0.568066120220875 | 4.77924572583707 | 3.95736338221264 | 0.0015487556300 | 0.00532707608321291 |
| Herc3 | ENSMUSG00000029804 | 73998 | -0.420356911617757 | 5.05678372920934 | -3.95679942204388 | 0.0015504324361 | 0.00533116607740980 |
| Colec12 | ENSMUSG00000036103 | 140792 | 0.504498712801632 | 4.19825608994008 | 3.95572225852023 | 0.0015536402935 | 0.00533850023395671 |
| Pdk1 | ENSMUSG00000006494 | 228026 | -0.264751646258283 | 6.80385234176719 | -3.95563121350785 | 0.0015539117417 | 0.00533850023395671 |
| Rpl10 | ENSMUSG00000008682 | 110954 | 0.289263919735181 | 6.81155512063804 | 3.95544907521248 | 0.0015544549276 | 0.00533850023395671 |
| Agt | ENSMUSG00000031980 | 11606 | -0.728548726886145 | 1.49487890047354 | -3.95542762406163 | 0.0015545189136 | 0.00533850023395671 |
| Tmem69 | ENSMUSG00000055900 | 230657 | -0.412469693663355 | 2.68886183025383 | -3.95524211787321 | 0.0015550723662 | 0.00533872362923845 |
| Polr3gl | ENSMUSG00000028104 | 69870 | -0.42229905239936 | 4.04727774655459 | -3.95398535904449 | 0.0015588271798 | 0.00534993403533074 |
| Ap3s1 | ENSMUSG00000024480 | 11777 | 0.631027537510594 | 2.05310185253863 | 3.95320768075892 | 0.0015611552861 | 0.00535624244613578 |
| Mylk2 | ENSMUSG00000027470 | 228785 | -0.586561513626283 | 6.3673936128179 | -3.95288955803905 | 0.0015621086622 | 0.00535783174977275 |
| Rgmb | ENSMUSG00000048027 | 68799 | 0.281250939772574 | 4.80158647393804 | 3.95097366480755 | 0.0015678629635 | 0.00537535472802906 |
| Vps8 | ENSMUSG00000033653 | 209018 | 0.301066222665737 | 3.22019206437758 | 3.95072796509127 | 0.0015686024775 | 0.00537535472802906 |
| Snupn | ENSMUSG00000055334 | 66069 | -0.527974350809693 | 2.18975842348343 | -3.9505359853947 | 0.0015691805515 | 0.00537535472802906 |
| Akna | ENSMUSG00000039158 | 100182 | 0.695590776664079 | 2.49822583092694 | 3.95043483018124 | 0.0015694852296 | 0.00537535472802906 |
| Mrpl36 | ENSMUSG00000021607 | 94066 | -0.372726317380395 | 3.1892629626278 | -3.95037138383919 | 0.0015696763600 | 0.00537535472802906 |
| Dcn | ENSMUSG00000019929 | 13179 | 0.356218081424302 | 7.6797430020362 | 3.94971195759978 | 0.0015716642726 | 0.00538047671907658 |
| Kbtbd4 | ENSMUSG00000005505 | 67136 | -0.420149232390495 | 3.14505011580198 | -3.94891083961577 | 0.0015740827924 | 0.00538706919370667 |
| Ppat | ENSMUSG00000029246 | 231327 | -0.398652992502642 | 3.41547716723319 | -3.94870737700625 | 0.0015746976374 | 0.00538748665897482 |
| Kcnc1 | ENSMUSG00000058975 | 16502 | -0.368126287562715 | 5.69144015655773 | -3.9485148322308 | 0.0015752797158 | 0.00538779179412292 |
| Vps13a | ENSMUSG00000046230 | 271564 | -0.392618025002141 | 4.93253511381379 | -3.94592411321936 | 0.0015831330858 | 0.00541295832371205 |
| Slc16a3 | ENSMUSG00000025161 | 80879 | -0.893202946884817 | 2.90532552687154 | -3.94542952299142 | 0.0015846369030 | 0.00541640587482817 |
| BC034090 | ENSMUSG00000033722 | 207792 | 1.28969698674779 | 1.03637058217157 | 3.9450790947838 | 0.0015857032746 | 0.00541835651614592 |
| Cbfa2t3 | ENSMUSG00000006362 | 12398 | -0.411223779985575 | 4.91388289467925 | -3.94477160439158 | 0.0015866395881 | 0.00541986166826995 |
| Marcksl1 | ENSMUSG00000047945 | 17357 | 1.08366128134189 | 0.992515545369523 | 3.94376526303956 | 0.0015897078669 | 0.00542864626472671 |
| Dcaf8 | ENSMUSG00000026554 | 98193 | -0.28983197206038 | 6.87619748262221 | -3.943601648386 | 0.0015902072916 | 0.00542865581033876 |
| Pip4k2a | ENSMUSG00000026737 | 18718 | 0.609650066560226 | 2.38642072140909 | 3.94298000635583 | 0.0015921062818 | 0.00543344170047980 |
| Mical1 | ENSMUSG00000019823 | 171580 | 0.575636316600087 | 2.46119369253201 | 3.94128594044119 | 0.0015972930797 | 0.00544944152768368 |
| Otulin | ENSMUSG00000046034 | 432940 | 0.347457191037796 | 5.22419016065577 | 3.93681965899842 | 0.0016110506180 | 0.00549466280994859 |
| Zfp180 | ENSMUSG00000057101 | 210135 | -0.343404064349899 | 3.5229236602367 | -3.93309361686883 | 0.0016226205895 | 0.00553239727517392 |
| Tm9sf3 | ENSMUSG00000025016 | 107358 | 0.259006358738915 | 6.01588256091499 | 3.93195070049928 | 0.0016261865179 | 0.00554282657558362 |
| Klf13 | ENSMUSG00000052040 | 50794 | 0.369299789501852 | 4.78773502718474 | 3.93177950188288 | 0.0016267213510 | 0.00554292116215504 |
| Myo1e | ENSMUSG00000032220 | 71602 | 0.45924124968216 | 4.19068217749563 | 3.93047210886079 | 0.0016308116428 | 0.00555512685207495 |
| Rad23a | ENSMUSG00000003813 | 19358 | -0.288190961251505 | 5.85422909812168 | -3.92936027941842 | 0.0016342983630 | 0.00556526959701259 |
| Rpl24 | ENSMUSG00000098274 | 68193 | -0.308235155806879 | 6.32098219292136 | -3.927749358901 | 0.0016393637579 | 0.00558078020510761 |
| Icam2 | ENSMUSG00000001029 | 15896 | -0.831217967178319 | 2.03704767006953 | -3.92702844002881 | 0.0016416358142 | 0.00558677493328582 |
| Pdcd5 | ENSMUSG00000030417 | 56330 | -0.366441026329081 | 5.38909039078997 | -3.92628958929169 | 0.0016439677224 | 0.00559127654695840 |
| Dhx35 | ENSMUSG00000027655 | 71715 | -0.427471715995996 | 2.92405735444535 | -3.92628519685557 | 0.0016439815956 | 0.00559127654695840 |
| Srp54a | ENSMUSG00000073079 | 24067 | 0.443849098444934 | 6.35354036559327 | 3.9257364432141 | 0.0016457157372 | 0.00559543350664833 |
| Rbx1 | ENSMUSG00000022400 | 56438 | -0.311816458272372 | 4.43827222418077 | -3.92532844238335 | 0.0016470062903 | 0.00559808014927857 |
| Trim23 | ENSMUSG00000021712 | 81003 | -0.298330191032543 | 3.93742694882141 | -3.92342245936555 | 0.0016530488394 | 0.00561687188822233 |
| Frmd6 | ENSMUSG00000048285 | 319710 | 0.441711461816873 | 4.68077024598159 | 3.9229143878091 | 0.0016546634008 | 0.00562060692891283 |
| Zfp979 | ENSMUSG00000066000 | 112422 | -0.651458454615816 | 1.82802803319292 | -3.92275304909991 | 0.0016551764435 | 0.00562060692891283 |
| Cdpf1 | ENSMUSG00000064284 | 72355 | -0.466042973861746 | 2.4628584609421 | -3.92116442467237 | 0.0016602368114 | 0.00563603993366058 |
| Ppp1r14c | ENSMUSG00000040653 | 76142 | -0.317628046139166 | 4.32900404370123 | -3.92008236330119 | 0.0016636926291 | 0.00564557334101091 |
| Bag4 | ENSMUSG00000037316 | 67384 | -0.364994371483867 | 2.85431627141384 | -3.91996182132099 | 0.0016640780628 | 0.00564557334101091 |
| Abcc4 | ENSMUSG00000032849 | 239273 | -0.501484574946123 | 3.00847877019617 | -3.91962286681737 | 0.0016651623606 | 0.00564720121254141 |
| Nsun4 | ENSMUSG00000028706 | 72181 | -0.465913550856313 | 2.86727405972305 | -3.91948889261494 | 0.0016655911361 | 0.00564720121254141 |
| Pxmp4 | ENSMUSG00000000876 | 59038 | -0.334147331366348 | 2.90804614257212 | -3.91861872150457 | 0.0016683788063 | 0.00565489883184647 |
| Cap2 | ENSMUSG00000021373 | 67252 | -0.258176491784893 | 7.40543348391853 | -3.91492510976539 | 0.0016802647573 | 0.00569342035409255 |
| Usp25 | ENSMUSG00000022867 | 30940 | -0.286635722741015 | 6.29404173743866 | -3.91431589336498 | 0.0016822334922 | 0.00569832485393160 |
| Prkar1a | ENSMUSG00000020612 | 19084 | 0.332125248587164 | 6.95081077070347 | 3.91335620082686 | 0.0016853395999 | 0.00570707780886290 |
| Minpp1 | ENSMUSG00000024896 | 17330 | 0.517417356481589 | 3.25096107867965 | 3.91076038229086 | 0.0016937705125 | 0.00573385118399827 |
| Tinagl1 | ENSMUSG00000028776 | 94242 | -1.06976151301513 | 3.45187051561587 | -3.90875370565197 | 0.0017003174658 | 0.00575334864937394 |
| Avpi1 | ENSMUSG00000018821 | 69534 | 0.642744820136287 | 2.34765084373832 | 3.90867257653815 | 0.0017005826993 | 0.00575334864937394 |
| Agbl1 | ENSMUSG00000025754 | 244071 | 0.445297807299482 | 4.77751641804742 | 3.9080701685465 | 0.0017025534587 | 0.00575823386683037 |
| Ino80c | ENSMUSG00000047989 | 225280 | 0.414565414659525 | 4.05545564923584 | 3.90781363095529 | 0.0017033934214 | 0.00575929275912614 |
| Scmh1 | ENSMUSG00000000085 | 29871 | -0.337699244598704 | 4.32055198472087 | -3.90719087438707 | 0.0017054342295 | 0.00576428381103708 |
| Traf3 | ENSMUSG00000021277 | 22031 | 0.401938822365059 | 5.44008000584518 | 3.90704145272324 | 0.0017059242639 | 0.00576428381103708 |
| Furin | ENSMUSG00000030530 | 18550 | 0.256029032422911 | 5.38159244588043 | 3.90560138569124 | 0.0017106543971 | 0.00577672715381268 |
| Fam20b | ENSMUSG00000033557 | 215015 | -0.265488227533653 | 5.34756107192381 | -3.90559853268651 | 0.0017106637816 | 0.00577672715381268 |
| Sclt1 | ENSMUSG00000059834 | 67161 | -0.640116689910995 | 2.1036248860132 | -3.90539423112293 | 0.0017113359307 | 0.00577721218606665 |
| Lmnb2 | ENSMUSG00000062075 | 16907 | 0.698462244784261 | 1.87125285336926 | 3.90360471702595 | 0.0017172349415 | 0.00579533656880621 |
| Tmem52 | ENSMUSG00000023153 | 69671 | -0.55162062349045 | 4.13097574318843 | -3.9032625442079 | 0.0017183652529 | 0.00579736128999060 |
| Apol6 | ENSMUSG00000033576 | 71939 | -0.419255060832761 | 5.55090691393606 | -3.90275233859014 | 0.0017200520435 | 0.00580126161318493 |
| Nmnat3 | ENSMUSG00000032456 | 74080 | -0.460021803335052 | 2.56512221810796 | -3.90228900518356 | 0.0017215853345 | 0.00580464197783406 |
| Bnip3l | ENSMUSG00000022051 | 12177 | -0.264592347579919 | 4.97411866606315 | -3.900572777778 | 0.0017272769448 | 0.00582203647357618 |
| Abi1 | ENSMUSG00000058835 | 11308 | 0.261926634128691 | 3.84176792822631 | 3.89757400659792 | 0.0017372680644 | 0.00585390789543973 |
| Asb4 | ENSMUSG00000042607 | 65255 | -1.12920390754548 | 1.30189896429712 | -3.89667982521859 | 0.0017402586454 | 0.00586217789018835 |
| Tmem150a | ENSMUSG00000055912 | 232086 | -0.534955876875558 | 2.11527850562534 | -3.89574711150867 | 0.0017433836970 | 0.00587089562319124 |
| Cyb561 | ENSMUSG00000019590 | 13056 | 0.675271117997263 | 1.92052977007065 | 3.89362947002584 | 0.0017505001248 | 0.00589304492288684 |

| | | | | | | | |
|---|---|---|---|---|---|---|---|
| Dixdc1 | ENSMUSG00000032064 | 330938 | -0.549623408221905 | 2.43664570873269 | -3.89221732687741 | 0.0017552621657 | 0.00590725699928135 |
| Ybx3 | ENSMUSG00000030189 | 56449 | -0.338565785909934 | 8.97079057056529 | -3.89048917560804 | 0.0017611078473 | 0.00592510614940124 |
| Dennd5a | ENSMUSG00000035901 | 19347 | 0.218320172990464 | 6.08280724153806 | 3.88919434237987 | 0.0017655007956 | 0.00593805821467202 |
| Slc35a5 | ENSMUSG00000022664 | 74102 | 0.546529143326336 | 1.86762736923775 | 3.88855928475498 | 0.0017676594256 | 0.00594348975128059 |
| Trim24 | ENSMUSG00000029833 | 21848 | -0.297909946594874 | 4.7789471183457 | -3.88784114880348 | 0.0017701036931 | 0.00594987806563327 |
| Golgb1 | ENSMUSG00000034243 | 224139 | 0.303470252084257 | 5.90943485201911 | 3.88755058101358 | 0.0017710936576 | 0.00595137558292266 |
| Cog6 | ENSMUSG00000027742 | 67542 | -0.277646752677131 | 5.01270100343376 | -3.88735613978891 | 0.0017717564344 | 0.00595177307451428 |
| Dennd6b | ENSMUSG00000015377 | 69440 | -0.48722182462498 | 2.39647325566066 | -3.88630340849369 | 0.0017753491956 | 0.00596200984886202 |
| Tapt1 | ENSMUSG00000046985 | 231225 | -0.258979200668989 | 4.88859725858635 | -3.88595779388145 | 0.0017765303295 | 0.0059641440517112 |
| Cmbl | ENSMUSG00000022235 | 69574 | -0.402226197270587 | 3.07866814152387 | -3.88439568872634 | 0.0017818788164 | 0.0059802632308347 |
| Fads6 | ENSMUSG00000044788 | 328035 | -0.43478504960305 | 3.76508029118917 | -3.88389570405486 | 0.0017835941814 | 0.00598418293337613 |
| Stat6 | ENSMUSG00000002147 | 20852 | 0.383711915926548 | 4.01678515798918 | 3.88348355609479 | 0.0017850094600 | 0.00598709371202873 |
| Pde4c | ENSMUSG00000031842 | 110385 | -0.876664585958972 | 1.20064028647107 | -3.88305050226581 | 0.0017864977621 | 0.00599024755758409 |
| Iffo2 | ENSMUSG00000041025 | 212632 | 0.450645321484586 | 2.44716819972564 | 3.88211281410789 | 0.0017897247120 | 0.00599922748448457 |
| Eid2 | ENSMUSG00000046058 | 386655 | 0.511411177112591 | 1.70807703780074 | 3.88138949974445 | 0.0017922179811 | 0.00600574333301376 |
| Adamts20 | ENSMUSG00000022449 | 223838 | 0.754375509917902 | 2.67671648602008 | 3.88089155751952 | 0.0017939364506 | 0.0060096596204848 |
| Sult1a1 | ENSMUSG00000030711 | 20887 | -0.745556750225347 | 2.49886888335577 | -3.88045935028155 | 0.0017954294222 | 0.00601281832541825 |
| Snx9 | ENSMUSG00000002365 | 66616 | 0.242438406939558 | 5.23432300400179 | 3.87950608769755 | 0.0017987267562 | 0.00602106497936251 |
| Os9 | ENSMUSG00000040462 | 216440 | 0.366880061865564 | 4.41051380529928 | 3.87942904252557 | 0.0017989935250 | 0.00602106497936251 |
| Edil3 | ENSMUSG00000034488 | 13612 | 0.941877723757421 | 1.01257319200804 | 3.87680442002185 | 0.0018081054302 | 0.00604970935369613 |
| Ptpn12 | ENSMUSG00000028771 | 19248 | 0.341170227877299 | 4.38355047154215 | 3.87648518424346 | 0.0018092169288 | 0.0060515759635505 |
| Hexa | ENSMUSG00000025232 | 15211 | 0.406359149454009 | 4.70317919192659 | 3.87534350045218 | 0.0018131976915 | 0.00606303578058841 |
| Fbxl3 | ENSMUSG00000022124 | 50789 | -0.246168399878571 | 5.13539070541427 | -3.87194290186642 | 0.0018251077466 | 0.00610099473339032 |
| Ppp1r14a | ENSMUSG00000037166 | 68458 | -0.845413013237789 | 1.07329320511598 | -3.8699967850245 | 0.0018319595854 | 0.00612202697283623 |
| D630024D03Rik | ENSMUSG00000085772 | 414116 | -0.799123845330254 | 1.48405387321476 | -3.86924355496815 | 0.0018346185768 | 0.00612903901713201 |
| Cpeb3 | ENSMUSG00000039652 | 208922 | -0.398983194512946 | 5.37587546942689 | -3.86694035779386 | 0.0018427735848 | 0.00615440209467663 |
| Phlda1 | ENSMUSG00000020205 | 21664 | 0.690564684729502 | 2.83321992090119 | 3.86631752852096 | 0.0018449852021 | 0.00615990630562002 |
| Ddah2 | ENSMUSG00000007039 | 51793 | 0.499915375145719 | 2.23841457249584 | 3.86461122173123 | 0.0018510580405 | 0.00617652217603132 |
| Nubp1 | ENSMUSG00000022503 | 26425 | 0.615090832569789 | 2.56895970470355 | 3.86460169616952 | 0.0018510919996 | 0.00617652217603132 |
| Kif1c | ENSMUSG00000020821 | 16562 | -0.367462860771125 | 9.36300307048287 | -3.86193348327245 | 0.0018606293876 | 0.00620645097249029 |
| Tmem147 | ENSMUSG00000006315 | 69804 | -0.351020291371738 | 3.68044223195919 | -3.86163461251317 | 0.0018617008022 | 0.00620813040553047 |
| Dusp5 | ENSMUSG00000034765 | 240672 | 1.52452133009887 | 0.584425043234275 | 3.86131168515432 | 0.0018628591639 | 0.0062100986645823 |
| Chd1l | ENSMUSG00000028089 | 68058 | -0.332606170947789 | 3.76445810697608 | -3.86083190041962 | 0.0018645815408 | 0.00621394537294903 |
| Mmp11 | ENSMUSG00000000901 | 17385 | 0.907322020056441 | 1.13138624691314 | 3.8590003633913 | 0.0018711715139 | 0.00623400664995008 |
| Ddx59 | ENSMUSG00000026404 | 67997 | -0.442052242574134 | 2.90978732247952 | -3.85763754786543 | 0.0018760904088 | 0.00624849002405179 |
| Mamld1 | ENSMUSG00000059401 | 333639 | 0.523288702396248 | 2.28908209118622 | 3.85743127392478 | 0.0018768360743 | 0.00624906948789422 |
| Etv6 | ENSMUSG00000030199 | 14011 | 0.285079014502495 | 4.02104489847127 | 3.85565873215066 | 0.0018832561558 | 0.00626853624828383 |
| Tle5 | ENSMUSG00000054452 | 14797 | -0.287474802732481 | 8.08874221320491 | -3.8544258701637 | 0.0018877347376 | 0.00628125146563363 |
| Ppp1r27 | ENSMUSG00000025129 | 68701 | -0.868693852331099 | 5.91669516304558 | -3.85429097036975 | 0.0018882254428 | 0.00628125146563363 |
| Sqor | ENSMUSG00000005803 | 59010 | -0.437286531322142 | 3.26599707233006 | -3.85407730702996 | 0.0018890029209 | 0.00628192605076581 |
| Atrnl1 | ENSMUSG00000054843 | 226255 | 0.421751673647895 | 4.29219992513367 | 3.85334894786517 | 0.0018916557278 | 0.00628883478127072 |
| Prkag3 | ENSMUSG00000006542 | 241113 | 0.795150320344955 | 3.18580889668885 | 3.85279676736073 | 0.0018936693936 | 0.00629361512361629 |
| Cyth3 | ENSMUSG00000018001 | 19159 | 0.401871532767795 | 4.55057343003942 | 3.85178821685913 | 0.0018973529714 | 0.00630394083018342 |
| Ntmt1 | ENSMUSG00000026857 | 66617 | 0.372124280570771 | 2.63049496160971 | 3.85095446361303 | 0.0019004036446 | 0.00631083553702812 |
| Vav2 | ENSMUSG00000009621 | 22325 | 0.473241645981975 | 2.45632908402936 | 3.8509055426179 | 0.0019005828001 | 0.00631083553702812 |
| Tuba1a | ENSMUSG00000072235 | 22142 | 0.615808327794859 | 5.00663938016336 | 3.85071927652833 | 0.0019012650897 | 0.00631118393454595 |
| Bet1l | ENSMUSG00000025484 | 54399 | 0.600638747503957 | 1.63799123762631 | 3.84992257020682 | 0.0019041862318 | 0.0063189616576591 |
| Stt3a | ENSMUSG00000032116 | 16430 | 0.417654487580013 | 4.61065444062328 | 3.84697624289345 | 0.0019150288179 | 0.00635301365979737 |
| Sema4b | ENSMUSG00000030539 | 20352 | 0.660730898960588 | 1.47809513282705 | 3.8459599696778 | 0.0019187833209 | 0.00636353776734078 |
| She | ENSMUSG00000046280 | 214547 | -0.655220548014606 | 3.15976037138949 | -3.8401559619344 | 0.0019403698189 | 0.00643317636983267 |
| Nectin1 | ENSMUSG00000032012 | 58235 | 0.661796037259516 | 2.11576904173806 | 3.83772010730552 | 0.0019495029441 | 0.00646149687158053 |
| Wdr60 | ENSMUSG00000042050 | 217935 | 0.3524637337318 | 3.39846330967127 | 3.83575242188891 | 0.0019569126768 | 0.00648408986679497 |
| Prrc2c | ENSMUSG00000040225 | 226562 | 0.247807932036879 | 6.85428995889516 | 3.83395067119859 | 0.0019637227217 | 0.00650033218546811 |
| Rhbdd3 | ENSMUSG00000034175 | 279766 | -0.464097616089207 | 2.37362617395436 | -3.83385254524394 | 0.0019640942994 | 0.00650033218546811 |
| Ctf1 | ENSMUSG00000042340 | 13019 | -0.680366266006997 | 1.06494722029398 | -3.83382741607811 | 0.0019641894686 | 0.00650033218546811 |
| Gatc | ENSMUSG00000029536 | 384281 | -0.308003570702045 | 3.54636028041306 | -3.8338263984891 | 0.0019641933225 | 0.00650033218546811 |
| Csf1 | ENSMUSG00000014599 | 12977 | 0.666983999548788 | 3.70186446870088 | 3.83192180283967 | 0.0019714201171 | 0.00652077250765106 |
| Fcgrt | ENSMUSG00000003420 | 14132 | 0.290020813287407 | 4.26450492325618 | 3.83188426683965 | 0.0019715628156 | 0.00652077250765106 |
| Ssx2ip | ENSMUSG00000036825 | 99167 | -0.326010975683808 | 4.8481169084504 | -3.83128172879702 | 0.0019738548918 | 0.00652637865190954 |
| Sp3 | ENSMUSG00000027109 | 20687 | 0.314723094603808 | 4.45313232653373 | 3.82896817074789 | 0.0019826810099 | 0.00655357911090577 |
| Gbp7 | ENSMUSG00000040253 | 229900 | -0.828506489205847 | 4.1233211835095 | -3.8287057569122 | 0.0019836846430 | 0.00655491439925673 |
| Fiz1 | ENSMUSG00000061374 | 23877 | -0.335751198946797 | 2.97564172973036 | -3.82816029114891 | 0.0019857724988 | 0.00655983051811874 |
| Med15 | ENSMUSG00000012114 | 94112 | 0.25937402638379 | 4.7721434121946 | 3.82793781936075 | 0.0019866246872 | 0.00655989586743908 |
| Ppan | ENSMUSG00000004100 | 235036 | 0.571012229537598 | 1.95429622632596 | 3.82784182367254 | 0.0019869925182 | 0.00655989586743908 |
| Scnm1 | ENSMUSG00000092607 | 69269 | -0.331881225599977 | 3.39656815363817 | -3.82699724563352 | 0.0019902317187 | 0.00656860595326632 |
| Ftsj1 | ENSMUSG00000031171 | 54632 | -0.351150832863407 | 2.63252080445224 | -3.82675685385582 | 0.0019911546738 | 0.00656966849995834 |
| Galk2 | ENSMUSG00000027207 | 69976 | 0.676335616209514 | 1.22899592214736 | 3.82316373153928 | 0.0020050021265 | 0.00661336096734492 |
| Eif2s3y | ENSMUSG00000069049 | 26908 | -0.351079029044388 | 4.21774987662039 | -3.82241280405027 | 0.0020079084833 | 0.00662094951171223 |
| Frem2 | ENSMUSG00000037016 | 242022 | 0.575850530427374 | 3.94229003450325 | 3.82204485279233 | 0.0020093341522 | 0.00662365247844057 |
| Zmpste24 | ENSMUSG00000043207 | 230709 | 0.297153222635651 | 3.85435341167689 | 3.82154590606374 | 0.0020112690256 | 0.00662803187207654 |
| Tspan2 | ENSMUSG00000027858 | 70747 | -0.731443142399083 | 2.44516866249334 | -3.82038898248501 | 0.0020157627850 | 0.00664083875935972 |
| Sik3 | ENSMUSG00000034135 | 70661 | -0.214220757855004 | 5.52239449036408 | -3.81806728847851 | 0.0020248116692 | 0.00666834775958493 |
| Mpdz | ENSMUSG00000028402 | 17475 | -0.237796775679207 | 6.55316071443919 | -3.81793386241126 | 0.0020253329578 | 0.00666834775958493 |
| Zfp110 | ENSMUSG00000058638 | 65020 | -0.298120949739968 | 3.6233552852066 | -3.81774023408686 | 0.0020260896965 | 0.0066688306152013 |
| Wasf2 | ENSMUSG00000028868 | 242687 | 0.359226192365445 | 5.36857175408431 | 3.81674982840413 | 0.0020299649099 | 0.00667957448230413 |
| Leprot | ENSMUSG00000035212 | 230514 | 0.51043296424113 | 2.66710806317122 | 3.81551042657466 | 0.0020348250227 | 0.00669355170731479 |
| Slf2 | ENSMUSG00000036097 | 226151 | 0.295171332093923 | 4.73318901069541 | 3.81478629011341 | 0.0020376700904 | 0.00670089403096317 |
| Ttc39c | ENSMUSG00000024424 | 72747 | 0.713550266065913 | 1.20180793815462 | 3.81354902991397 | 0.0020425405557 | 0.00671489047075903 |
| Ophn1 | ENSMUSG00000031214 | 94190 | 0.585327063188593 | 2.44672107800901 | 3.81250420875365 | 0.0020466627207 | 0.00672641918237345 |
| Samd4b | ENSMUSG00000109336 | 233033 | 0.325807616897781 | 3.78635561100658 | 3.81050809753637 | 0.0020545616146 | 0.00675034956725775 |
| Cdk16 | ENSMUSG00000031065 | 18555 | -0.254701035752125 | 6.13782758306845 | -3.80944541629532 | 0.0020587794531 | 0.00676070268546687 |
| COX3 | ENSMUSG00000064358 | 17710 | -0.516528536250755 | 11.6896819142967 | -3.8094025677041 | 0.0020589497063 | 0.00676070268546687 |
| Tsc22d2 | ENSMUSG00000027806 | 72033 | 0.265847035464792 | 4.57173694510644 | 3.80722755546065 | 0.0020676106941 | 0.00678699298559005 |
| Kdm5b | ENSMUSG00000042207 | 75605 | 0.323729109250968 | 4.04813520682578 | 3.80708037319577 | 0.0020681981186 | 0.00678699298559005 |
| F2r | ENSMUSG00000048376 | 14062 | 0.471531873130282 | 5.01232442227988 | 3.80683455810979 | 0.0020691795792 | 0.0067881758646876 |
| Elmod2 | ENSMUSG00000035151 | 244548 | 0.421582888162019 | 3.09535841471912 | 3.80560674067911 | 0.0020740889662 | 0.00680224017090405 |
| Itgb3 | ENSMUSG00000020689 | 16416 | 0.836752808799798 | 1.463224167889 | 3.80480368557138 | 0.0020773063704 | 0.00681074863099252 |
| Mrpl12 | ENSMUSG00000039640 | 56282 | -0.383688064473213 | 6.38287742253088 | -3.80458559033029 | 0.0020781810366 | 0.00681157328674981 |
| Prpf4 | ENSMUSG00000066148 | 70052 | -0.333389742634833 | 3.73332720561461 | -3.80419436568298 | 0.0020797509743 | 0.00681467562974999 |
| Noc3l | ENSMUSG00000024999 | 57753 | 0.340177942837748 | 3.20238704882422 | 3.80380813291232 | 0.0020813020651 | 0.00681771437632519 |
| Myo1c | ENSMUSG00000017774 | 17913 | 0.355669437877015 | 5.97515673677342 | 3.80322127254147 | 0.0020836611211 | 0.00682328366144583 |
| Col7a1 | ENSMUSG00000025650 | 12836 | 1.12998572681985 | 0.997106081166046 | 3.80307471502123 | 0.0020842506765 | 0.00682328366144583 |
| Cacna1a | ENSMUSG00000034656 | 12286 | 0.410262952083622 | 3.45257418652961 | 3.80030379132938 | 0.0020954292656 | 0.00685782553984391 |
| Tmed5 | ENSMUSG00000063406 | 73130 | -0.265284512097831 | 4.6808494790436 | -3.79903896311925 | 0.0021005521705 | 0.00687088110003838 |
| Nbn | ENSMUSG00000028224 | 27354 | -0.270727204516997 | 3.9398422150425 | -3.79900853539918 | 0.0021006755682 | 0.00687088110003838 |
| Rnaset2b | ENSMUSG00000094724 | 68195 | -0.285731013920441 | 4.14109606451876 | -3.7965860813014 | 0.0021105233997 | 0.0069010264083114 |
| Ppp6c | ENSMUSG00000026753 | 67857 | -0.236446135931507 | 4.74227641549024 | -3.79638860080964 | 0.0021113282725 | 0.00690159370446971 |
| Prpf38b | ENSMUSG00000027881 | 66921 | 0.287505376937455 | 4.42740622999916 | 3.794637159219 | 0.0021184803286 | 0.00692231806426122 |
| Flcn | ENSMUSG00000032633 | 216805 | -0.607133192099881 | 4.86020018284387 | -3.79442196088211 | 0.0021193607964 | 0.00692231806426122 |
| Birc3 | ENSMUSG00000032000 | 11796 | 0.522969822204139 | 2.84546310692453 | 3.79437131391851 | 0.0021195680688 | 0.00692231806426122 |
| Srp54c | ENSMUSG00000079108 | 100101806 | 0.264378385234832 | 5.66367563994587 | 3.79292497167873 | 0.0021254959345 | 0.00693960455816632 |
| Ptgfr | ENSMUSG00000028036 | 19220 | 1.16602588704295 | 1.119397068564 | 3.79263171181912 | 0.0021266999232 | 0.00694146218604652 |
| Tsr1 | ENSMUSG00000038335 | 104662 | 0.317407284688043 | 3.83799306535735 | 3.79162703681414 | 0.0021308299184 | 0.00695286622031888 |
| Gm4070 | ENSMUSG00000078606 | 100042856 | -0.690297128237122 | 5.12869808783914 | -3.79021700545756 | 0.0021366400188 | 0.00696974397091338 |
| Ctps | ENSMUSG00000028633 | 51797 | 0.381125730489798 | 3.52713477424015 | 3.78842678030605 | 0.0021440399699 | 0.00699179621479054 |
| Mogs | ENSMUSG00000030036 | 57377 | 0.554903713348727 | 2.54235783970529 | 3.78803272476801 | 0.0021456723111 | 0.00699503251807539 |
| Tbl2 | ENSMUSG00000005374 | 27368 | 0.452429000603694 | 2.87539328939609 | 3.78782577142496 | 0.0021465301041 | 0.00699574256672452 |
| Mxra7 | ENSMUSG00000020814 | 67622 | 0.654044675497152 | 3.52147555230538 | 3.78555645635498 | 0.0021559590341 | 0.0070243780035557 |
| Ski | ENSMUSG00000029050 | 20481 | 0.423574351535543 | 6.09613032954255 | 3.78505782542688 | 0.0021580364727 | 0.00702905145513943 |
| Fhl1 | ENSMUSG00000023092 | 14199 | 0.395828243828676 | 10.883997510281 | 3.78451828275984 | 0.0021602866566 | 0.00703428461241419 |
| AW209491 | ENSMUSG00000039182 | 105351 | -0.37632997361457 | 3.16306152645827 | -3.78262507465629 | 0.0021682012652 | 0.00705795355273749 |
| Nrap | ENSMUSG00000049134 | 18175 | -0.380106441072517 | 11.5390510234082 | -3.78242079457771 | 0.0021690570259 | 0.00705863719879048 |
| Cdc23 | ENSMUSG00000024370 | 52563 | -0.283239017108869 | 3.38950540437228 | -3.78187707634004 | 0.0021713364210 | 0.00706395191035 |
| Dhdds | ENSMUSG00000012117 | 67422 | 0.318598554786121 | 2.96317293720913 | 3.78087139873258 | 0.0021755588860 | 0.00707558291671132 |
| Kank4 | ENSMUSG00000035407 | 242553 | -1.20401583306506 | 1.0288985854414 | -3.77953640213478 | 0.0021811769520 | 0.00709174457535953 |
| Pnrc1 | ENSMUSG00000040128 | 108767 | -0.282126517956284 | 6.9011232585647 | -3.77893983669697 | 0.0021836922494 | 0.00709781147158463 |

| | | | | | | | |
|---|---|---|---|---|---|---|---|
| BC067074 | ENSMUSG00000021763 | 408066 | -0.47058512900189 | 2.95702475173852 | -3.77721569881083 | 0.0021909783353 | 0.00711791103760652 |
| Tmem120b | ENSMUSG00000054434 | 330189 | -0.498922811046087 | 2.68377684912719 | -3.77716844869676 | 0.0021911783589 | 0.00711791103760652 |
| Airn | ENSMUSG00000078247 | 104103 | 0.699574967811412 | 1.73213799827135 | 3.77586031915861 | 0.0021967234408 | 0.00713380390131802 |
| Bbip1 | ENSMUSG00000084957 | 100503572 | -0.358027781049772 | 3.3673787294164 | -3.77456776733981 | 0.0022022165164 | 0.00714893114117784 |
| Golph3 | ENSMUSG00000022200 | 66629 | -0.277179597551045 | 5.06782669201612 | -3.77445660018086 | 0.0022026896060 | 0.00714893114117784 |
| Mrpl38 | ENSMUSG00000020775 | 60441 | -0.303095991725955 | 4.74723936248502 | -3.77411941625306 | 0.0022041251786 | 0.00715146700127309 |
| Zfp397 | ENSMUSG00000024276 | 69256 | -0.401051188678896 | 4.12413759355061 | -3.77386560584447 | 0.0022052064134 | 0.00715285201954208 |
| Cask | ENSMUSG00000031012 | 12361 | 0.347795678017578 | 3.52251993252841 | 3.77359005546522 | 0.0022063808719 | 0.00715453850823551 |
| Scap | ENSMUSG00000032485 | 235623 | 0.311635107997985 | 4.11259227628692 | 3.77329775115174 | 0.0022076274352 | 0.007156457738592 |
| Rad17 | ENSMUSG00000021635 | 19356 | -0.336517193608396 | 3.51492446242473 | -3.77197350329787 | 0.0022132838172 | 0.00717266688596911 |
| Amy1 | ENSMUSG00000074264 | 11722 | -1.29745922599829 | 2.33923013610761 | -3.77153347806134 | 0.0022151666001 | 0.00717664081386988 |
| Mbp | ENSMUSG00000041607 | 17196 | -0.968679279867844 | 3.19519615463802 | -3.77050295422176 | 0.0022195823926 | 0.00718848911160043 |
| P3h3 | ENSMUSG00000023191 | 14789 | 0.912639896622597 | 2.7673813199356 | 3.77023943275506 | 0.0022207130189 | 0.00718848911160043 |
| Cdc42ep1 | ENSMUSG00000049521 | 104445 | -0.746835000957686 | 3.20496886641154 | -3.77021995542944 | 0.0022207966087 | 0.00718848911160043 |
| ND4 | ENSMUSG00000064363 | 17719 | -0.522535656324637 | 12.4230938841468 | -3.76989977936093 | 0.0022221711500 | 0.00718868286317701 |
| Klhl42 | ENSMUSG00000040102 | 232539 | 0.659605171866205 | 2.23212333522774 | 3.76982705406628 | 0.0022224834861 | 0.00718868286317701 |
| Trim28 | ENSMUSG00000005566 | 21849 | 0.238058948672235 | 4.73129798529911 | 3.76974652457349 | 0.0022228293914 | 0.00718868286317701 |
| Zfp992 | ENSMUSG00000070605 | 433791 | 1.1842704554004 | 0.707870584481901 | 3.76904266719175 | 0.0022258550629 | 0.00719633886204249 |
| Ybey | ENSMUSG00000033126 | 216119 | -0.56236035971858 | 2.40379393390311 | -3.76651284231204 | 0.0022367646605 | 0.00722947205922333 |
| Smtnl2 | ENSMUSG00000045667 | 276829 | -0.54436042710778 | 5.36657194963039 | -3.76619158207453 | 0.0022381539483 | 0.00723182406411183 |
| Fundc1 | ENSMUSG00000025040 | 72018 | -0.324987833757617 | 3.58099210074268 | -3.76581182268663 | 0.0022397973475 | 0.00723499551017647 |
| Kif1bp | ENSMUSG00000036955 | 72320 | -0.307576015341712 | 4.38645725253363 | -3.76546129288505 | 0.0022413153461 | 0.00723776013248816 |
| Nfkbib | ENSMUSG00000030595 | 18036 | -0.361757905555395 | 4.05609620584421 | -3.7626498776107 | 0.0022535283393 | 0.00727504969800292 |
| Golt1b | ENSMUSG00000030245 | 66964 | 0.306378151001915 | 3.13138274813057 | 3.7621722781581 | 0.0022556097838 | 0.00727961929357146 |
| Shtn1 | ENSMUSG00000041362 | 71653 | 0.572990406458058 | 1.90753173429144 | 3.76050686624731 | 0.0022628831919 | 0.00730093747673408 |
| Pola2 | ENSMUSG00000024833 | 18969 | -0.344851239096382 | 3.07898794267147 | -3.7600986148167 | 0.0022646698000 | 0.00730454576122966 |
| Lvrn | ENSMUSG00000024481 | 74574 | -0.976462238191185 | 0.708337557230219 | -3.7597228319303 | 0.0022663155861 | 0.00730769783851588 |
| Cttnbp2nl | ENSMUSG00000062127 | 80281 | 0.427258046101183 | 3.60685090589526 | 3.75773688690627 | 0.0022750334962 | 0.00733364528099543 |
| Qpctl | ENSMUSG00000030407 | 67369 | 0.653265575789307 | 1.13447303671941 | 3.75733501321358 | 0.0022768017888 | 0.0073371817082642 |
| Smc1a | ENSMUSG00000041133 | 24061 | 0.284936183084257 | 5.32421173155579 | 3.7565453667241 | 0.0022802804018 | 0.00734622607501381 |
| Sephs2 | ENSMUSG00000049091 | 20768 | -0.431827762413871 | 3.69320939253668 | -3.75632356481322 | 0.0022812584732 | 0.00734721165910653 |
| Rit1 | ENSMUSG00000028057 | 19769 | 0.364633240825665 | 3.64581617520458 | 3.75517653967574 | 0.0022863232794 | 0.00736135486528011 |
| A4galt | ENSMUSG00000047878 | 239559 | 0.95180901264944 | 1.11347429231337 | 3.75467438325427 | 0.0022885442003 | 0.00736633588172329 |
| Cfap74 | ENSMUSG00000078490 | 544678 | -0.467296011459718 | 1.61078231058299 | -3.75262814186972 | 0.0022976169614 | 0.00739336208581726 |
| Msi2 | ENSMUSG00000069769 | 76626 | -0.250410148691394 | 7.17173339727356 | -3.7513134617235 | 0.0023034653711 | 0.00740999999176303 |
| Kif26b | ENSMUSG00000026494 | 269152 | 0.771194218530593 | 1.01096534302695 | 3.74955281707699 | 0.0023113213949 | 0.0074319779805095 |
| Ggact | ENSMUSG00000041625 | 223267 | -0.536958887759434 | 1.57312038249336 | -3.74947768843115 | 0.0023116572256 | 0.0074319779805095 |
| Limk1 | ENSMUSG00000029674 | 16885 | 0.495894578214125 | 1.87165206116632 | 3.74715812246898 | 0.0023220503132 | 0.00746319669923987 |
| Ubxn1 | ENSMUSG00000071655 | 225896 | 0.31502681607551 | 6.65011306079307 | 3.74670668103198 | 0.0023240785669 | 0.0074675199340932 |
| Akt1s1 | ENSMUSG00000011096 | 67605 | -0.284342798084289 | 5.13569492401097 | -3.74610045061292 | 0.0023268050970 | 0.00747276790338695 |
| Tmem268 | ENSMUSG00000045917 | 230279 | -0.586970229863785 | 2.14851337227806 | -3.7460395612449 | 0.0023270791275 | 0.00747276790338695 |
| Kat8 | ENSMUSG00000030801 | 67773 | -0.413361748019839 | 2.8803560484842 | -3.74552622679036 | 0.0023293906765 | 0.00747799397499035 |
| Sh3glb2 | ENSMUSG00000026860 | 227700 | 0.358256638255841 | 3.38538992855008 | 3.74410465637342 | 0.0023358042120 | 0.0074963816328568 |
| Fam168a | ENSMUSG00000029461 | 319604 | 0.289620889660957 | 5.29978771263992 | 3.7430969247222 | 0.0023403615519 | 0.00750880308890268 |
| Dpm1 | ENSMUSG00000078919 | 13480 | -0.414190405499141 | 3.21604077316898 | -3.74266924878909 | 0.0023422983943 | 0.00751086068420318 |
| Larp4b | ENSMUSG00000033499 | 217980 | -0.241735387137365 | 6.67722228292547 | -3.74265187747474 | 0.0023423770993 | 0.00751086068420318 |
| Arhgef9 | ENSMUSG00000025656 | 236915 | -0.43461920201073 | 3.44313880805902 | -3.74233338705135 | 0.0023438205745 | 0.00751328524945435 |
| Nfkbiz | ENSMUSG00000035356 | 80859 | 0.728743810326531 | 1.80046400211594 | 3.73846649396144 | 0.0023614185873 | 0.00756687991446903 |
| Rnasel | ENSMUSG00000066800 | 24014 | 0.861603656420648 | 0.935821933848297 | 3.73835581001847 | 0.0023619242766 | 0.00756687991446903 |
| Zfp644 | ENSMUSG00000049606 | 52397 | -0.281209210523678 | 6.16379931558254 | -3.73652475124182 | 0.0023703059539 | 0.00759150729048848 |
| Mapk6 | ENSMUSG00000042688 | 50772 | 0.242697748613615 | 4.97486654929276 | 3.73617564683106 | 0.0023719074093 | 0.00759441121619118 |
| Tlk1 | ENSMUSG00000041997 | 228012 | -0.213610965482606 | 5.24884521349648 | -3.73466769029964 | 0.0023788375432 | 0.00761372239400052 |
| Fkbp15 | ENSMUSG00000066151 | 338355 | 0.263309550665039 | 4.33679069016018 | 3.7345603177192 | 0.0023793317809 | 0.00761372239400052 |
| Wdr47 | ENSMUSG00000040389 | 99512 | 0.510896202750206 | 2.44222767434745 | 3.73408797747637 | 0.0023815072107 | 0.00761845341560221 |
| Lrrk1 | ENSMUSG00000015133 | 233328 | 0.64725335155239 | 3.77257160945386 | 3.73094175366782 | 0.0023960492685 | 0.00766273099910031 |
| Arrdc3 | ENSMUSG00000074794 | 105171 | 0.635938812547125 | 4.58569017459984 | 3.7297181325807 | 0.0024017292765 | 0.00767864952394852 |
| Zfp112 | ENSMUSG00000052675 | 57745 | -0.478435508211328 | 1.82307077638371 | -3.72904524170649 | 0.0024048586439 | 0.0076864063852034 |
| Gstm2 | ENSMUSG00000040562 | 14863 | -0.364696157518467 | 3.46727778620572 | -3.72876144489821 | 0.0024061797205 | 0.00768838074409683 |
| C7 | ENSMUSG00000079105 | 109828 | 1.02657340867319 | 2.12665492646739 | 3.72729167880725 | 0.0024130333055 | 0.00770802661112528 |
| Xk | ENSMUSG00000015342 | 22439 | 0.489640225004714 | 2.72078154963512 | 3.72693959939894 | 0.0024146780111 | 0.00771102697644682 |
| Abraxas1 | ENSMUSG00000035234 | 70681 | -0.53436509272078 | 1.87114964930758 | -3.72636087644067 | 0.0024173839366 | 0.00771538777354349 |
| Alms1 | ENSMUSG00000063810 | 236266 | 0.512486294793527 | 1.81305408211401 | 3.72634563755149 | 0.0024174552304 | 0.00771538777354349 |
| Prag1 | ENSMUSG00000050271 | 244418 | -0.55479491828873 | 2.69608096530808 | -3.72593926274067 | 0.0024193572091 | 0.0077177010083307 |
| Eef2k | ENSMUSG00000035064 | 13631 | -0.400821550221895 | 4.53599394999659 | -3.72575866976738 | 0.0024202029365 | 0.0077177010083307 |
| Ifi209 | ENSMUSG00000043263 | 236312 | 1.51721628454093 | 0.258855177166885 | 3.7257383427199 | 0.0024202981480 | 0.0077177010083307 |
| Fgd5 | ENSMUSG00000034037 | 232237 | -0.54255013468704 | 4.31944135420897 | -3.72524345687895 | 0.0024226173600 | 0.00772284350046329 |
| Arhgap20 | ENSMUSG00000053199 | 244867 | -0.523852454833801 | 5.49173814908584 | -3.72412138451355 | 0.0024278841618 | 0.00773737661034186 |
| Mtrr | ENSMUSG00000034617 | 210009 | -0.325554941735197 | 3.55353070282405 | -3.72301203633459 | 0.0024331026781 | 0.00775174741312841 |
| Abtb2 | ENSMUSG00000032724 | 99382 | -0.403161889382992 | 3.82182689520152 | -3.72024452306883 | 0.0024461711611 | 0.00779111216847069 |
| Wbp4 | ENSMUSG00000022023 | 22380 | -0.272704412320825 | 4.75312772564991 | -3.71996926958165 | 0.0024474748302 | 0.00779299369914881 |
| Podxl | ENSMUSG00000025608 | 27205 | -0.654945815266649 | 5.98501789719242 | -3.71959732813841 | 0.0024492375592 | 0.00779633539889035 |
| Calhm5 | ENSMUSG00000049872 | 103511 | 1.28328087060171 | 0.817694521788891 | 3.71915025284456 | 0.0024513580747 | 0.00780081371631039 |
| Dyrk1a | ENSMUSG00000022897 | 13548 | -0.239578585181063 | 6.57533849821246 | -3.71816686646871 | 0.0024560289257 | 0.00781340284856659 |
| Lmtk2 | ENSMUSG00000038970 | 231876 | -0.247239321881961 | 5.1247171965944 | -3.71721125533772 | 0.0024605765147 | 0.00782559263390285 |
| Abcc1 | ENSMUSG00000023088 | 17250 | 0.385566896438793 | 5.53237333813659 | 3.71563309422206 | 0.0024681054520 | 0.00784725442021195 |
| Sh3bgr | ENSMUSG00000040666 | 50795 | -0.308384285954212 | 6.37645619271339 | -3.71439035632952 | 0.0024740506551 | 0.00786386964553951 |
| Igfbp6 | ENSMUSG00000023046 | 16012 | 0.443414189226716 | 4.83644382507253 | 3.71405740496841 | 0.0024756459495 | 0.00786665287061222 |
| Rcn1 | ENSMUSG00000005973 | 19672 | 0.462340784462582 | 3.35735596416699 | 3.71355977589167 | 0.0024780322251 | 0.00787194718183262 |
| Rrs1 | ENSMUSG00000061024 | 59014 | -0.298462183230698 | 3.70378157377321 | -3.71183206881837 | 0.0024863352367 | 0.00789457366187243 |
| A830010M20Rik | ENSMUSG00000111375 | 231570 | 0.433078131907104 | 3.58779272413526 | 3.71177721500478 | 0.0024865993155 | 0.00789457366187243 |
| 2310015D24Rik | ENSMUSG00000099411 | 70100 | 0.765164628596809 | 2.30535904220908 | 3.71108905237443 | 0.0024899147098 | 0.00790280420828136 |
| Ubap1 | ENSMUSG00000028437 | 67123 | -0.250569643328879 | 4.67437388643245 | -3.70771744844572 | 0.0025062232646 | 0.00795225733128127 |
| Olfml1 | ENSMUSG00000051041 | 244198 | 0.810413698093252 | 1.14033172448967 | 3.70750602019387 | 0.0025072495582 | 0.0079532051424334 |
| Sumo1 | ENSMUSG00000026021 | 22218 | -0.265027913752152 | 4.70830274813804 | -3.70649081310915 | 0.0025121834227 | 0.00796504371894788 |
| Wnk1 | ENSMUSG00000045962 | 232341 | 0.303817912157708 | 8.90815848615533 | 3.70643827930119 | 0.0025124390031 | 0.00796504371894788 |
| Sdhaf4 | ENSMUSG00000026154 | 68002 | -0.406559158848578 | 3.40941123322373 | -3.70466251363862 | 0.0025210937703 | 0.00799016410669865 |
| Nfia | ENSMUSG00000028565 | 18027 | 0.235147241893609 | 5.4656541949907 | 3.7039136120457 | 0.0025247528570 | 0.00799944158860431 |
| Homer2 | ENSMUSG00000025813 | 26557 | -0.304024874451982 | 7.68844361190025 | -3.70183721103757 | 0.0025349262599 | 0.00802934771018694 |
| Zfp9 | ENSMUSG00000072623 | 22750 | 0.441069088696694 | 2.82316519248964 | 3.70018580333151 | 0.0025430471105 | 0.00805273695415372 |
| Mthfd1 | ENSMUSG00000021048 | 108156 | -0.33019740706405 | 4.62018273525608 | -3.6979952451002 | 0.0025538600368 | 0.00808297209786489 |
| Auts2 | ENSMUSG00000029673 | 319974 | 0.459993131240855 | 4.74756466462705 | 3.6979519435341 | 0.0025540742499 | 0.00808297209786489 |
| Oxld1 | ENSMUSG00000039670 | 66431 | -0.676980636660548 | 2.25118232896792 | -3.69714322562405 | 0.0025580783354 | 0.00809330080591079 |
| Slc7a8 | ENSMUSG00000022180 | 50934 | 0.619347017629946 | 1.70747603722096 | 3.69631499851212 | 0.0025621856214 | 0.00810394995015465 |
| Mrpl58 | ENSMUSG00000018858 | 68572 | -0.395322349970625 | 4.06573036653419 | -3.69604725468357 | 0.0025635148298 | 0.00810580867938081 |
| Erg | ENSMUSG00000040732 | 13876 | -0.523446925427701 | 2.94902830549319 | -3.69492285363301 | 0.0025691045437 | 0.00812018831847113 |
| Tyk2 | ENSMUSG00000032175 | 54721 | 0.544245920949045 | 2.90290888408997 | 3.69483371588163 | 0.0025695482017 | 0.00812018831847113 |
| Eya1 | ENSMUSG00000025932 | 14048 | -0.679655612553423 | 3.84596742661875 | -3.69200285235722 | 0.0025836785675 | 0.00816248278077104 |
| Fgd1 | ENSMUSG00000025265 | 14163 | 0.670359755043755 | 1.75015078794436 | 3.69079573131551 | 0.0025897279309 | 0.00817923028401293 |
| Slc12a4 | ENSMUSG00000017765 | 20498 | 0.307232217991925 | 4.54780623548945 | 3.68990255417348 | 0.0025942132522 | 0.00819102976915146 |
| Coq8a | ENSMUSG00000026489 | 67426 | -0.438544748638256 | 10.1970696859193 | -3.68885361678412 | 0.0025994908361 | 0.00820532322541742 |
| Dnajc3 | ENSMUSG00000022136 | 100037258 | 0.315589406852557 | 5.09712971462225 | 3.68858832271778 | 0.0026008273529 | 0.0082071719961846 |
| Mrpl19 | ENSMUSG00000030045 | 56284 | -0.309252132715447 | 3.6190583235926 | -3.68562087475166 | 0.0026158245985 | 0.0082521150612344 |
| Stam | ENSMUSG00000026718 | 20844 | 0.34817863889034 | 3.49456032221156 | 3.68505074516506 | 0.0026187160273 | 0.00825675380067481 |
| Pcdhb16 | ENSMUSG00000047910 | 93887 | 0.895805296844308 | 0.856896300342668 | 3.68503306802176 | 0.0026188057292 | 0.00825675380067481 |
| Lman1 | ENSMUSG00000041891 | 70361 | 0.235095288795803 | 5.52995602569399 | 3.68454770080557 | 0.0026212699256 | 0.00826214001079996 |
| A330074K22Rik | ENSMUSG00000097960 | 434353 | 0.401016114508048 | 3.52292459061063 | 3.68436097490642 | 0.0026222185549 | 0.00826274748452039 |
| Snx18 | ENSMUSG00000042364 | 170625 | 0.394141055687602 | 3.9805679131633 | 3.68369693738822 | 0.0026255949092 | 0.00827100229181471 |
| Tut4 | ENSMUSG00000034610 | 230594 | -0.273783214599754 | 4.35304343250828 | -3.68318700952208 | 0.0026281906755 | 0.00827679408665496 |
| Get4 | ENSMUSG00000025858 | 67604 | 0.29280456188734 | 4.35041431623536 | 3.68298572380525 | 0.0026292160290 | 0.00827763836794067 |
| Fbxo4 | ENSMUSG00000022184 | 106052 | 0.605337144608062 | 1.26168552584525 | 3.68180571195186 | 0.0026352352065 | 0.00829419983943221 |
| Rgs19 | ENSMUSG00000002458 | 56470 | 0.706337872852474 | 1.20819089215158 | 3.68081824297518 | 0.0026402829825 | 0.00830769524531012 |
| Sar1a | ENSMUSG00000020088 | 20224 | -0.247392698723305 | 6.05046788833393 | -3.67874164436117 | 0.0026509302526 | 0.00833879671701138 |
| Mrpl21 | ENSMUSG00000024829 | 353242 | -0.271890323768929 | 3.5984945053932 | -3.67490795039142 | 0.0026707012489 | 0.0083985717352751 |
| Dpy19l1 | ENSMUSG00000043067 | 244745 | 0.485250908854862 | 3.01430985455251 | 3.67341872961394 | 0.0026784217075 | 0.00842042786445063 |
| Aida | ENSMUSG00000042901 | 108909 | 0.227197758394564 | 4.79934159924085 | 3.6718419548391 | 0.0026866207547 | 0.00844377558078758 |
| Myo18b | ENSMUSG00000072720 | 74376 | 0.375280031760388 | 9.67807798222363 | 3.67157301794485 | 0.0026880217361 | 0.00844575038737939 |

| | | | | | | | |
|---|---|---|---|---|---|---|---|
| Kifap3 | ENSMUSG00000026585 | 16579 | -0.252502108030304 | 4.24821569170585 | -3.66936884704121 | 0.0026995319666 | 0.00847947813992767 |
| Cyb5b | ENSMUSG00000031924 | 66427 | -0.285260012224561 | 5.41144228827316 | -3.66688736825259 | 0.0027125501677 | 0.00851792182811865 |
| Nck1 | ENSMUSG00000032475 | 17973 | -0.238528267120146 | 4.38474062118697 | -3.66616471671722 | 0.0027163532669 | 0.0085274145780904 |
| Plekha6 | ENSMUSG00000041757 | 240753 | 0.466812394827323 | 3.83031173174144 | 3.66575415643724 | 0.0027185163325 | 0.00852891529780576 |
| Wdr26 | ENSMUSG00000038733 | 226757 | 0.250059368147467 | 7.49166730843645 | 3.66569237817924 | 0.0027188419669 | 0.00852891529780576 |
| Psma5 | ENSMUSG00000068749 | 26442 | 0.341723243376889 | 4.49175234660286 | 3.66562976125269 | 0.0027191720622 | 0.00852891529780576 |
| Eif4ebp1 | ENSMUSG00000031490 | 13685 | 0.351277415706893 | 4.64428483884636 | 3.66537356315218 | 0.0027205230760 | 0.00853070503270498 |
| Slc9a1 | ENSMUSG00000028854 | 20544 | 0.358240879233473 | 3.37421190551687 | 3.66436106806187 | 0.0027258689510 | 0.00854501677765791 |
| Mapkap1 | ENSMUSG00000038696 | 227743 | -0.235000148398127 | 4.60882134058003 | -3.66309393779331 | 0.0027325742935 | 0.00856358073476179 |
| Pcgf3 | ENSMUSG00000033623 | 69587 | 0.2304843023112 | 5.20571048757522 | 3.66252015900347 | 0.0027356160923 | 0.00857065620680844 |
| Dhodh | ENSMUSG00000031730 | 56749 | -0.474009129815793 | 2.06973170594618 | -3.66218021587633 | 0.0027374198681 | 0.00857385002262827 |
| Rdm1 | ENSMUSG00000010362 | 66599 | -0.709401711810868 | 1.68895159372913 | -3.66194054525104 | 0.0027386923122 | 0.00857537830558571 |
| 2310061I04Rik | ENSMUSG00000050705 | 69662 | -0.3342531624115 | 4.55552744955713 | -3.66118851066626 | 0.0027426888632 | 0.00858543297932821 |
| Zfp759 | ENSMUSG00000057396 | 268670 | -0.602874144381344 | 1.60072526782446 | -3.65942109526118 | 0.0027521047796 | 0.00861244126719195 |
| Nudt19 | ENSMUSG00000034875 | 110959 | -0.443016473617739 | 3.57613106794271 | -3.65896118415407 | 0.0027545603314 | 0.00861765855268368 |
| Gphn | ENSMUSG00000047454 | 268566 | -0.279192762714269 | 4.51726384664791 | -3.65867006698331 | 0.0027561158093 | 0.00862005777153314 |
| Gtdc1 | ENSMUSG00000036890 | 227835 | -0.30602073740891 | 3.31388298803393 | -3.6584032268814 | 0.0027575423542 | 0.00862205248111516 |
| Mid1 | ENSMUSG00000035299 | 17318 | 0.617704612211869 | 3.55586115336411 | 3.65714657755283 | 0.0027642705613 | 0.00864061810286035 |
| Zfp451 | ENSMUSG00000042197 | 98403 | 0.309133424060764 | 4.03321680011994 | 3.65606652398848 | 0.0027700665653 | 0.00865626004178938 |
| Eml2 | ENSMUSG00000040811 | 72205 | 0.375848865705672 | 2.19482977271276 | 3.65474772381848 | 0.0027771604993 | 0.00867594781880335 |
| Sugp2 | ENSMUSG00000036054 | 234373 | -0.260127659297479 | 4.85888003497328 | -3.65404777175081 | 0.0027809330814 | 0.00868525128962439 |
| Fgf13 | ENSMUSG00000031137 | 14168 | -0.260967803327572 | 3.70422200955895 | -3.65350841123304 | 0.0027838436574 | 0.00869185804620496 |
| Prodh | ENSMUSG00000003526 | 19125 | -0.953042075315842 | 2.26496654235077 | -3.65265403595299 | 0.0027884604871 | 0.0087037868604364 |
| Cyp1b1 | ENSMUSG00000024087 | 13078 | 1.2471885177543 | 0.676265100574029 | 3.65239944971027 | 0.0027898377086 | 0.00870559976962348 |
| Tafa5 | ENSMUSG00000054863 | 106014 | -0.544196055961342 | 1.57915011540063 | -3.65224137928431 | 0.0027906931608 | 0.00870578394432166 |
| Itga3 | ENSMUSG00000001507 | 16400 | 0.440086673836952 | 2.12235436334141 | 3.65049674691974 | 0.0028001525268 | 0.00873280093325982 |
| Myh11 | ENSMUSG00000018830 | 17880 | -0.555681469560557 | 5.99260124938592 | -3.64912550482773 | 0.0028076101968 | 0.0087535616264144 |
| Bax | ENSMUSG00000003873 | 12028 | 0.475687393624657 | 1.84520173759395 | 3.64829292838509 | 0.0028121480921 | 0.00876520980749122 |
| Fam136a | ENSMUSG00000057497 | 66488 | -0.259108500896308 | 3.9353048203932 | -3.64783639185145 | 0.0028146395670 | 0.00876910449761135 |
| Gba2 | ENSMUSG00000028467 | 230101 | -0.491085000610876 | 2.82690631867132 | -3.64777000022993 | 0.0028150020750 | 0.00876910449761135 |
| H2-Q4 | ENSMUSG00000035929 | 15015 | -0.594008606591733 | 3.63830114638999 | -3.6475220597394 | 0.0028163562852 | 0.00877082351954726 |
| Lamp1 | ENSMUSG00000031447 | 16783 | -0.225788105446449 | 7.90528422900906 | -3.64728363633317 | 0.0028176591377 | 0.00877238166754861 |
| Mrpl54 | ENSMUSG00000034932 | 66047 | -0.343103113477481 | 4.66318749641721 | -3.64463855345559 | 0.0028321541265 | 0.00881499907676886 |
| Kidins220 | ENSMUSG00000036333 | 77480 | 0.406277656263707 | 5.72605065362069 | 3.64443497134575 | 0.0028332728805 | 0.00881522127590196 |
| Glud1 | ENSMUSG00000021794 | 14661 | -0.424917246294968 | 6.26989305861106 | -3.64433209332053 | 0.0028338384012 | 0.00881522127590196 |
| Mrps6 | ENSMUSG00000039680 | 121022 | -0.451187696806676 | 2.74861507946975 | -3.64407616793625 | 0.0028352457200 | 0.00881708989077371 |
| Zfpm1 | ENSMUSG00000049577 | 22761 | -0.588963738768404 | 3.07414302800913 | -3.64216737044565 | 0.0028457644655 | 0.00884526401698958 |
| Lrrc38 | ENSMUSG00000028584 | 242735 | -1.1425012645322 | 1.89865307993231 | -3.64213850353409 | 0.0028459238448 | 0.00884526401698958 |
| Calm1 | ENSMUSG00000001175 | 12313 | -0.266331158363604 | 7.48589366438609 | -3.64190467794665 | 0.0028472151705 | 0.0088467620889108 |
| Isyna1 | ENSMUSG00000019139 | 71780 | 0.54889918813742 | 1.55231281899771 | 3.64154119547787 | 0.0028492237192 | 0.0088504872052295 |
| Prkd2 | ENSMUSG00000041187 | 101540 | -0.557163627906041 | 2.72923775296708 | -3.64112737201684 | 0.0028515121936 | 0.00885507948544757 |
| Smad4 | ENSMUSG00000024515 | 17128 | -0.208931293538375 | 5.10125153263554 | -3.63875341674547 | 0.0028646764005 | 0.00889264173370035 |
| Ptpn9 | ENSMUSG00000032290 | 56294 | 0.48657869824466 | 2.91601540196844 | 3.63856745647212 | 0.0028657101979 | 0.00889264173370035 |
| Gja5 | ENSMUSG00000057123 | 14613 | -0.855575104190764 | 1.58445934180686 | -3.63850660612468 | 0.0028660485616 | 0.00889264173370035 |
| Nhsl1 | ENSMUSG00000039835 | 215819 | -0.378855250226378 | 3.14017126914911 | -3.63696734789619 | 0.0028746212267 | 0.00891550492827897 |
| Gm15787 | ENSMUSG00000086247 | 100504007 | -0.81471153900886 | 1.15507906777756 | -3.636890755596 | 0.0028750484742 | 0.00891550492827897 |
| H13 | ENSMUSG00000019188 | 14950 | 0.289895128089975 | 4.45697064607173 | 3.6363085782968 | 0.0028782980833 | 0.00892263592300032 |
| Rcan2 | ENSMUSG00000039601 | 53901 | -0.315184438554454 | 7.07229840531479 | -3.63616416172054 | 0.0028791047670 | 0.00892263592300032 |
| Dync2h1 | ENSMUSG00000047193 | 110350 | 0.602850593136846 | 2.69122867829856 | 3.63604029146009 | 0.0028797968654 | 0.00892263592300032 |
| Rspo3 | ENSMUSG00000019880 | 72780 | -0.485269170325946 | 3.46452286858313 | -3.63587744238669 | 0.0028807070062 | 0.00892292668894356 |
| Atp10a | ENSMUSG00000025324 | 11982 | 0.768800208818611 | 1.91755490566259 | 3.63526243932242 | 0.0028841468035 | 0.00893105062607052 |
| A730008H23Rik | ENSMUSG00000044783 | 212427 | 0.449380757914353 | 3.46957172346822 | 3.63486094358765 | 0.0028863946710 | 0.00893548007631561 |
| Scand1 | ENSMUSG00000046229 | 19018 | -0.259803999271385 | 4.42686378911983 | -3.63274194351689 | 0.0028982878429 | 0.00896975776075161 |
| Slco2a1 | ENSMUSG00000032548 | 24059 | 0.812590816754971 | 1.72755931358667 | 3.63209984055747 | 0.0029019015299 | 0.00897839955384448 |
| Tusc3 | ENSMUSG00000039530 | 80286 | 0.765117373163852 | 1.82636798758785 | 3.63191873237924 | 0.0029029216130 | 0.00897901419121548 |
| Pgk1 | ENSMUSG00000062070 | 18655 | -0.278259221502244 | 7.12032320441896 | -3.63167451606445 | 0.0029042977251 | 0.00898072940135166 |
| Selenof | ENSMUSG00000037072 | 93684 | 0.253069006197769 | 5.29088353570676 | 3.63074598306269 | 0.0029095358700 | 0.00899248634895248 |
| Ube2t | ENSMUSG00000026429 | 67196 | -0.733506921803708 | 1.2454038864278 | -3.63070892082956 | 0.0029097451483 | 0.00899248634895248 |
| Miga1 | ENSMUSG00000054942 | 215708 | -0.257962657631565 | 3.29628749572805 | -3.62998682681439 | 0.0029138256223 | 0.00900187367898707 |
| Slc22a5 | ENSMUSG00000018900 | 20520 | -0.376109470866561 | 2.53604099044783 | -3.62985096715559 | 0.0029145939984 | 0.00900187367898707 |
| Map11 | ENSMUSG00000036948 | 231807 | 0.80904808090955 | 0.859894591681181 | 3.62973443848736 | 0.0029152532086 | 0.00900187367898707 |
| Bet1 | ENSMUSG00000032757 | 12068 | 0.365622953647988 | 2.87399257947985 | 3.62937805251901 | 0.0029172702450 | 0.00900497151939024 |
| Rpp30 | ENSMUSG00000024800 | 54364 | -0.475548510304741 | 2.48378103088243 | -3.62913772560886 | 0.0029186312209 | 0.00900497151939024 |
| C030006K11Rik | ENSMUSG00000116138 | 223665 | -0.42709008685048 | 3.63784472416708 | -3.62912066732381 | 0.0029187278467 | 0.00900497151939024 |
| Tnnt1 | ENSMUSG00000064179 | 21955 | 0.453505246509676 | 11.2348224333651 | 3.62773823768492 | 0.0029265693428 | 0.0090266166724114 |
| Sgcd | ENSMUSG00000020354 | 24052 | 0.31655622405324 | 5.00553252909905 | 3.62621584693123 | 0.0029352294419 | 0.00905077377431628 |
| 5031439G07Rik | ENSMUSG00000036046 | 223739 | 0.220862909010613 | 6.32971495341978 | 3.62562209606353 | 0.0029386140217 | 0.00905865478606459 |
| Msantd4 | ENSMUSG00000041124 | 78100 | -0.28742965607727 | 5.32363271303389 | -3.62280579896773 | 0.0029547218166 | 0.00910574123995331 |
| Npc2 | ENSMUSG00000021242 | 67963 | 0.432520756753997 | 3.5044623838819 | 3.62213282529597 | 0.0029585841131 | 0.00911507410961856 |
| Crlf2 | ENSMUSG00000033467 | 57914 | 0.728406637327477 | 1.08382830657741 | 3.62114226727245 | 0.0029642783832 | 0.00912947761554818 |
| Mpp1 | ENSMUSG00000031402 | 17524 | 0.474621901786179 | 2.80152910586568 | 3.62102910522677 | 0.0029649296070 | 0.00912947761554818 |
| Ranbp9 | ENSMUSG00000038546 | 56705 | -0.258545961538966 | 5.13086175491918 | -3.62003505928851 | 0.0029706563670 | 0.00914453527135887 |
| Crocc | ENSMUSG00000040860 | 230872 | 0.700197734381537 | 1.30906740418882 | 3.61937440465886 | 0.0029744686380 | 0.00915277091999851 |
| Msantd3 | ENSMUSG00000039693 | 66665 | 0.287896980435502 | 3.62271708955854 | 3.61928127935166 | 0.0029750064110 | 0.00915277091999851 |
| Mff | ENSMUSG00000026150 | 75734 | -0.256965577984145 | 5.6146381447922 | -3.61755440020666 | 0.0029849965326 | 0.00918092208720976 |
| Zfp609 | ENSMUSG00000040524 | 214812 | -0.270275035162854 | 5.14030739078547 | -3.61686600547411 | 0.0029889884160 | 0.00919061388914082 |
| Acbd4 | ENSMUSG00000056938 | 67131 | 0.552458176876858 | 1.99210808565111 | 3.6132236770121 | 0.0030101998548 | 0.00925323245607503 |
| Szrd1 | ENSMUSG00000040842 | 213491 | -0.282845945876143 | 5.65469207374712 | -3.61259013572021 | 0.0030139048822 | 0.00926201694341834 |
| Ldlr | ENSMUSG00000032193 | 16835 | 0.560519061275681 | 3.32549798734071 | 3.61186531549301 | 0.0030181493868 | 0.0092701958685703 |
| Naalad2 | ENSMUSG00000043943 | 72560 | 0.673183857458337 | 2.82804350296561 | 3.61184601819637 | 0.0030182624733 | 0.0092701958685703 |
| Sept2 | ENSMUSG00000116048 | 18000 | 0.343617559910624 | 4.32427462818004 | 3.61112554630583 | 0.0030224876761 | 0.00928056538977992 |
| Kcnma1 | ENSMUSG00000063142 | 16531 | -0.248463590096897 | 7.25076474499014 | -3.60993777563349 | 0.0030294664357 | 0.00929938152460607 |
| Hacd4 | ENSMUSG00000028497 | 66775 | 0.603787218694707 | 1.47385149218803 | 3.6096407735323 | 0.0030312140245 | 0.00930213377389226 |
| Col18a1 | ENSMUSG00000001435 | 12822 | 0.463150807329405 | 4.40188893664868 | 3.60948132682712 | 0.0030321526455 | 0.00930240262931151 |
| Tomm20 | ENSMUSG00000093904 | 67952 | -0.31241206335536 | 6.27280210871085 | -3.60793581692489 | 0.0030412659290 | 0.009327743510369 |
| Cnpy4 | ENSMUSG00000036968 | 66455 | 0.642818528193256 | 1.20936907656369 | 3.60722790902998 | 0.0030454494525 | 0.00933795454856368 |
| Vgll2 | ENSMUSG00000049641 | 215031 | -0.608050116859546 | 7.32659352025379 | -3.60694217412559 | 0.0030471397110 | 0.00934051715679115 |
| Chpf | ENSMUSG00000032997 | 74241 | 0.760674386968815 | 2.78309233866306 | 3.60549842349758 | 0.0030556947276 | 0.00936411524180064 |
| Cpne8 | ENSMUSG00000052560 | 66871 | 0.455564306065434 | 2.45389016065835 | 3.60380078819881 | 0.0030657852783 | 0.00939240439396929 |
| Bicd1 | ENSMUSG00000003452 | 12121 | 0.69321236139477 | 1.98921016151211 | 3.60254932206336 | 0.0030732454547 | 0.00941262148098677 |
| Fam162a | ENSMUSG00000003955 | 70186 | -0.287410522464483 | 4.59359172380556 | -3.60200558668785 | 0.0030764924693 | 0.00941981501733571 |
| Dusp8 | ENSMUSG00000037887 | 18218 | -0.396540153587901 | 4.84484560679302 | -3.60186749404945 | 0.0030773176678 | 0.00941981501733571 |
| Rnf31 | ENSMUSG00000047098 | 268749 | -0.364656676275988 | 3.61021409943032 | -3.60166884971009 | 0.0030785050974 | 0.00942081165177485 |
| 1700120C14Rik | ENSMUSG00000100599 | 73600 | 0.438547622689789 | 2.89721473796547 | 3.60104278629917 | 0.0030822505325 | 0.00942963352121062 |
| Stub1 | ENSMUSG00000039615 | 56424 | -0.259790762786358 | 5.14429870484516 | -3.6008447274768 | 0.0030834363835 | 0.00943062202245647 |
| Mios | ENSMUSG00000042447 | 252875 | -0.264358246650934 | 3.64167277791872 | -3.59984116549148 | 0.0030894521838 | 0.00944637813205641 |
| Otud6b | ENSMUSG00000040550 | 72201 | -0.340040464744434 | 5.70080269277582 | -3.59796566764135 | 0.0031007266408 | 0.00947819991908532 |
| Anapc11 | ENSMUSG00000025135 | 66156 | -0.368187040495866 | 3.25077774388425 | -3.59721753339059 | 0.0031052356275 | 0.00948932922699984 |
| Snrnp27 | ENSMUSG00000001158 | 66618 | -0.300408711860231 | 4.58785094468087 | -3.59688507575704 | 0.0031072414719 | 0.0094928050669003 |
| Kpna6 | ENSMUSG00000003731 | 16650 | -0.216226571387636 | 5.62910374759524 | -3.59488254242081 | 0.0031193513068 | 0.00952638382263108 |
| Rnf149 | ENSMUSG00000048234 | 67702 | 0.433621328795909 | 3.0291651609771 | 3.59477950805164 | 0.0031199756733 | 0.00952638382263108 |
| Lysmd2 | ENSMUSG00000032184 | 70082 | 0.625004797204098 | 2.5205871602429 | 3.59452905881135 | 0.0031214938704 | 0.00952835786038262 |
| Senp2 | ENSMUSG00000022855 | 75826 | -0.220279264248933 | 4.50133805261946 | -3.59353568250106 | 0.0031275229852 | 0.00954409652469961 |
| Pcdhgb6 | ENSMUSG00000103088 | 93703 | 0.729827212127764 | 2.09110404309923 | 3.59336100401216 | 0.0031285843825 | 0.00954467091417154 |
| Tnfrsf12a | ENSMUSG00000023905 | 27279 | 0.803110008858865 | 4.40065222396124 | 3.59312775586774 | 0.0031300022347 | 0.00954511473841502 |
| Ptpro | ENSMUSG00000030223 | 19277 | 1.05849446942648 | 0.757950936883342 | 3.59283673458525 | 0.0031317721866 | 0.00954511473841502 |
| Adam10 | ENSMUSG00000054693 | 11487 | 0.258424090296876 | 4.39788514440592 | 3.59276323578003 | 0.0031322193566 | 0.00954511473841502 |
| Zfp870 | ENSMUSG00000095325 | 240066 | -0.430813111472028 | 2.75251617357264 | -3.59276268278388 | 0.0031322227213 | 0.00954511473841502 |
| Arid5b | ENSMUSG00000019947 | 71371 | 0.405359792812312 | 5.37161339527558 | 3.59190624111684 | 0.0031374381241 | 0.00955834340442018 |
| 2310022B05Rik | ENSMUSG00000031983 | 69551 | 0.339411542099568 | 3.84781826510538 | 3.59137806206556 | 0.0031406589153 | 0.00956548971962188 |
| Zbtb20 | ENSMUSG00000022708 | 56490 | 0.32827111428181 | 8.41782855794507 | 3.59046966954435 | 0.0031462060477 | 0.00957971540620687 |
| Usf3 | ENSMUSG00000068284 | 207806 | -0.255694425951322 | 6.71580655444479 | -3.5902121074878 | 0.0031477806642 | 0.00958184083558795 |
| 1700020I14Rik | ENSMUSG00000085438 | 66602 | -0.34244601308068 | 8.77312980197877 | -3.58937077888778 | 0.0031529297238 | 0.00959484265337936 |
| Hist2h3b | ENSMUSG00000074403 | 319154 | 0.700101109979222 | 1.21922005084384 | 3.58806210953364 | 0.0031609559334 | 0.00961659040050677 |
| Uchl3 | ENSMUSG00000022111 | 50933 | -0.295367045086588 | 3.60177757372562 | -3.5873655759312 | 0.0031652362801 | 0.00962432595543349 |
| Ago3 | ENSMUSG00000028842 | 214150 | 0.343184536067805 | 3.74350134121729 | 3.58736179680593 | 0.0031652595196 | 0.00962432595543349 |

| | | | | | | | |
|---|---|---|---|---|---|---|---|
| AI837181 | ENSMUSG00000047423 | 107242 | 0.392920419365817 | 2.93222983368372 | 3.58684988742819 | 0.0031684090752 | 0.00963122347097439 |
| Cdkl2 | ENSMUSG00000029403 | 53886 | 0.550720054892042 | 2.12550606751708 | 3.5860813523922 | 0.0031731434947 | 0.00964293342812063 |
| Fech | ENSMUSG00000024588 | 14151 | -0.320680086762835 | 6.3566791035701 | -3.58582842855508 | 0.0031747031514 | 0.00964499170344651 |
| 2300009A05Rik | ENSMUSG00000032403 | 69478 | -0.442954341490526 | 2.06944697375716 | -3.58363671452783 | 0.0031882509127 | 0.0096834594963925 |
| A930033H14Rik | ENSMUSG00000090622 | 320700 | -0.662194113770237 | 1.15164412157357 | -3.58229825961006 | 0.0031965531439 | 0.00970591396418913 |
| Lancl2 | ENSMUSG00000062190 | 71835 | -0.412272076207591 | 3.06013784221198 | -3.58215873824953 | 0.0031974198321 | 0.00970591396418913 |
| Mtx1 | ENSMUSG00000064068 | 17827 | -0.451944851157609 | 3.21474508970039 | -3.58155420074375 | 0.0032011778876 | 0.00971205783449377 |
| Kmt2b | ENSMUSG00000006307 | 75410 | 0.255037384938061 | 4.57764593253248 | 3.58154730457027 | 0.0032012207828 | 0.00971205783449377 |
| Tmem218 | ENSMUSG00000032121 | 66279 | -0.595163934971491 | 1.27658490181813 | -3.57998241732393 | 0.0032109696805 | 0.00973893162559322 |
| Aars2 | ENSMUSG00000023938 | 224805 | -0.385853330275406 | 3.45689110760868 | -3.57874997225755 | 0.0032186686856 | 0.00975957486906104 |
| Dag1 | ENSMUSG00000039952 | 13138 | 0.239938808926144 | 8.0558156287181 | 3.5785330655517 | 0.0032200256201 | 0.0097609817120112 |
| Cbr1 | ENSMUSG00000051483 | 12408 | -0.441990046722002 | 2.68676943430655 | -3.57767111881358 | 0.0032254235483 | 0.00977463399133763 |
| Acaca | ENSMUSG00000020532 | 107476 | -0.315666397042367 | 5.1037007091118 | -3.57489370332854 | 0.0032428794473 | 0.00982481021026184 |
| Jpt2 | ENSMUSG00000024165 | 52009 | 0.660587302816288 | 1.54521526900977 | 3.57469098433694 | 0.0032441572621 | 0.00982595816928454 |
| Slc30a2 | ENSMUSG00000028836 | 230810 | -0.791556622041018 | 1.04960091634375 | -3.57432888174632 | 0.0032464410005 | 0.00983015140631471 |
| Cdh22 | ENSMUSG00000053166 | 104010 | -0.966699017821142 | 1.61818847740039 | -3.57416154641366 | 0.0032474969142 | 0.00983062552482763 |
| Galnt16 | ENSMUSG00000021130 | 108760 | 0.499194586491548 | 2.56264202393995 | 3.57356458502811 | 0.0032512666721 | 0.00983709811140008 |
| Btbd9 | ENSMUSG00000062202 | 224671 | 0.300196001303078 | 3.35781364419926 | 3.57353795310646 | 0.0032514349534 | 0.00983709811140008 |
| Atp1b3 | ENSMUSG00000032412 | 11933 | 0.313368448361415 | 4.23596980233186 | 3.57295279041448 | 0.0032551346951 | 0.00984556650614184 |
| Spg11 | ENSMUSG00000033396 | 214585 | 0.415155443822521 | 3.87626916268678 | 3.57210766403491 | 0.0032604856003 | 0.00985902298694142 |
| Drosha | ENSMUSG00000022191 | 14000 | -0.251950200647417 | 5.78393435877567 | -3.57084520698377 | 0.0032684954118 | 0.00988050977516678 |
| Urb1 | ENSMUSG00000039929 | 207932 | 0.373823247870337 | 2.65442970272642 | 3.57009523901474 | 0.0032732630985 | 0.00989218659936198 |
| Homer3 | ENSMUSG00000003573 | 26558 | 0.676822550387267 | 1.46193213047296 | 3.56991225395741 | 0.0032744274362 | 0.00989297023382421 |
| Gabarapl2 | ENSMUSG00000031950 | 93739 | -0.236454972040869 | 4.7066164717671 | -3.56732505002705 | 0.0032909347894 | 0.00994009620984265 |
| Synpo2l | ENSMUSG00000039376 | 68760 | 0.335209726065663 | 7.98073295846405 | 3.56605272204865 | 0.0032990835888 | 0.00996195654971093 |
| Mtch1 | ENSMUSG00000024012 | 56462 | 0.232570934448773 | 5.06480529815082 | 3.56573277425402 | 0.0033011359486 | 0.00996540100943527 |
| Mcrip1 | ENSMUSG00000061111 | 192173 | 0.374464925577849 | 3.55018346090002 | 3.56498032265041 | 0.0033059677706 | 0.00997723183354953 |
| Scyl1 | ENSMUSG00000024941 | 78891 | -0.268311917172618 | 5.39543008852743 | -3.56431444811097 | 0.0033102496103 | 0.00998739676811753 |
| Frmd4b | ENSMUSG00000030064 | 232288 | 0.546199212758141 | 3.28407305364169 | 3.56387767883791 | 0.0033130612580 | 0.00999312158165332 |
| Ilkap | ENSMUSG00000026309 | 67444 | -0.236572391876201 | 4.39390715465004 | -3.56275286996285 | 0.0033203132022 | 0.0100103486819698 |
| Bmp2k | ENSMUSG00000034663 | 140780 | 0.424847492037215 | 4.29717636637798 | 3.56270778947105 | 0.0033206041826 | 0.0100103486819698 |
| Ago2 | ENSMUSG00000036698 | 239528 | 0.232150509730787 | 6.56785762626249 | 3.56193878214226 | 0.0033255718594 | 0.0100225602413827 |
| Wdr46 | ENSMUSG00000024312 | 57315 | 0.339314066146726 | 2.98785171807239 | 3.56114347267532 | 0.0033307173523 | 0.0100353008209083 |
| Clcn3 | ENSMUSG00000004319 | 12725 | -0.315491693143733 | 4.84027400779141 | -3.56087975058393 | 0.0033324253588 | 0.010037680241706 |
| Prdm5 | ENSMUSG00000029913 | 70779 | 0.487276418883635 | 1.42350272107453 | 3.55993759454544 | 0.0033385345091 | 0.010053311492981 |
| Ap1g2 | ENSMUSG00000040701 | 11766 | 0.609198223072861 | 1.7614819026755 | 3.55876445738248 | 0.0033461572303 | 0.0100734906872284 |
| Habp4 | ENSMUSG00000021476 | 56541 | 0.334144247595683 | 4.78175821673885 | 3.55659545031636 | 0.0033602972343 | 0.0101132734218727 |
| Pltp | ENSMUSG00000017754 | 18830 | 0.384923166869916 | 4.0863650695642 | 3.55549074378491 | 0.0033675221550 | 0.0101311553119965 |
| Ndufaf3 | ENSMUSG00000070283 | 66706 | -0.440910731946718 | 2.2897786984979 | -3.55540364885489 | 0.0033680924347 | 0.0101311553119965 |
| Esd | ENSMUSG00000021996 | 13885 | 0.293624040414167 | 5.25606560170971 | 3.55488913404958 | 0.0033714633647 | 0.0101385051004792 |
| C330018D20Rik | ENSMUSG00000024592 | 77422 | -0.515901431197433 | 2.22750832330856 | -3.55425302206715 | 0.0033756356765 | 0.0101482600605088 |
| Atg12 | ENSMUSG00000032905 | 67526 | 0.286878033695036 | 3.27342898536603 | 3.55374193452524 | 0.0033789917255 | 0.0101555563793449 |
| Trpc3 | ENSMUSG00000027716 | 22065 | -0.665775974695468 | 2.49800201519699 | -3.55322243146875 | 0.0033824064930 | 0.0101630251171008 |
| Mxi1 | ENSMUSG00000025025 | 17859 | -0.279792914872735 | 5.20688847669467 | -3.55221951874634 | 0.0033890086565 | 0.0101800642002332 |
| Mrps11 | ENSMUSG00000030611 | 67994 | -0.431467120432419 | 2.54760774254557 | -3.55121731431794 | 0.0033956191759 | 0.0101971190143136 |
| Rpl18a | ENSMUSG00000045128 | 76808 | -0.281591001120407 | 7.96327265802088 | -3.55092587874462 | 0.0033975439249 | 0.0101981651126319 |
| Hspb6 | ENSMUSG00000036854 | 243912 | -0.336637971655056 | 10.3640273392843 | -3.55088205621317 | 0.0033978334406 | 0.0101981651126319 |
| Tubg1 | ENSMUSG00000035198 | 103733 | -0.351277985328145 | 3.77247525516034 | -3.5503153787215 | 0.0034015794714 | 0.0102066058749637 |
| Gns | ENSMUSG00000034707 | 75612 | 0.283205484666173 | 5.31848230726062 | 3.55016133039257 | 0.0034025985322 | 0.0102068618429079 |
| Rab9 | ENSMUSG00000079316 | 56382 | -0.341208008238908 | 2.67751707025633 | -3.5496970179665 | 0.0034056719204 | 0.0102132783984195 |
| Yipf3 | ENSMUSG00000071074 | 28064 | 0.343154292849785 | 3.47066757544596 | 3.54921671225668 | 0.0034088541256 | 0.0102200176762809 |
| Emc4 | ENSMUSG00000027131 | 68032 | -0.297743022099889 | 3.65608955540617 | -3.54857933094334 | 0.0034130816584 | 0.0102298863748241 |
| Chchd1 | ENSMUSG00000063787 | 66121 | -0.371819662090022 | 4.78406392551314 | -3.54807758312224 | 0.0034164133095 | 0.010234586382683 |
| Nlrc3 | ENSMUSG00000049871 | 268857 | 0.711716594154507 | 4.92212624516457 | 3.54806117129776 | 0.0034165223410 | 0.010234586382683 |
| Galnt4 | ENSMUSG00000090035 | 14426 | 0.539774913026877 | 1.78567651379826 | 3.54635201033182 | 0.0034278963914 | 0.0102658453301475 |
| Ndufaf8 | ENSMUSG00000078572 | 208501 | -0.33897061617963 | 3.40688340886634 | -3.5449243295579 | 0.0034374266024 | 0.0102915667465163 |
| Lacc1 | ENSMUSG00000044350 | 210808 | 0.931066299881167 | 0.672927929779094 | 3.54329347995651 | 0.0034483458117 | 0.0103214315631784 |
| Snord15b | ENSMUSG00000064966 | 449631 | 0.540503953714286 | 5.82828551918913 | 3.54256062048327 | 0.0034532640210 | 0.0103333230261189 |
| Chst12 | ENSMUSG00000036599 | 59031 | 0.796782390350591 | 1.73615661073363 | 3.5405812591158 | 0.0034665829582 | 0.0103703388935804 |
| Nol9 | ENSMUSG00000028948 | 74035 | 0.291686846293886 | 3.06662697433402 | 3.54034912180429 | 0.0034681483871 | 0.0103721833158717 |
| Slk | ENSMUSG00000025060 | 20874 | 0.291543839883489 | 6.29421326696876 | 3.53913014961635 | 0.0034763803194 | 0.0103939587722561 |
| Rad52 | ENSMUSG00000030166 | 19365 | 0.34879701570378 | 2.49986051424758 | 3.53838311399494 | 0.0034814349401 | 0.010406225138345 |
| Nploc4 | ENSMUSG00000039703 | 217365 | 0.250375570349551 | 6.00444710728429 | 3.53561199181022 | 0.0035002499914 | 0.0104596043347317 |
| Rph3al | ENSMUSG00000020847 | 380714 | -0.675113612498586 | 1.5501712491282 | -3.53484902614661 | 0.0035054483110 | 0.0104709653075454 |
| Pprc1 | ENSMUSG00000055491 | 226169 | 0.320312632441047 | 3.88511280722271 | 3.53477285614395 | 0.0035059677088 | 0.0104709653075454 |
| Smc6 | ENSMUSG00000020608 | 67241 | -0.286202706938726 | 5.90821039806517 | -3.53424663030069 | 0.0035095581311 | 0.0104788254388903 |
| Hes1 | ENSMUSG00000022528 | 15205 | -0.403616497145822 | 4.07640946233221 | -3.5329919090001 | 0.0035181340569 | 0.010501562909983 |
| Vps13d | ENSMUSG00000020220 | 230895 | -0.24980638904937 | 6.56894677469767 | -3.53176315050695 | 0.0035265530560 | 0.0105238196712879 |
| Snora73b | ENSMUSG00000065353 | 100306945 | 1.01119489174192 | 2.80627431397855 | 3.53139671134334 | 0.0035290677011 | 0.01052844952007 |
| Pot1a | ENSMUSG00000029676 | 101185 | -0.375520688883328 | 2.86251397594757 | -3.53101411579237 | 0.0035316951513 | 0.0105311475569744 |
| Mief2 | ENSMUSG00000018599 | 237781 | -0.355462661072967 | 5.69528921088317 | -3.5308698167029 | 0.0035326866293 | 0.0105311475569744 |
| Akap5 | ENSMUSG00000021057 | 238276 | -1.00205552859001 | 0.998922426366924 | -3.53080502645929 | 0.0035331318941 | 0.0105311475569744 |
| 2210016L21Rik | ENSMUSG00000029559 | 72357 | -0.367888774018319 | 3.46706859106981 | -3.53070408076827 | 0.0035338257468 | 0.0105311475569744 |
| Irs1 | ENSMUSG00000055980 | 16367 | 0.534888537056549 | 6.48138605153043 | 3.52956500580897 | 0.0035416647573 | 0.0105516319058152 |
| Txlna | ENSMUSG00000053841 | 109658 | 0.234240847938309 | 4.4353493315743 | 3.52938174888955 | 0.0035429275548 | 0.0105525180114501 |
| Ablim3 | ENSMUSG00000032735 | 319713 | -0.400812553712813 | 3.97023004965205 | -3.5291709753355 | 0.0035443805282 | 0.0105539699139493 |
| Hsd17b12 | ENSMUSG00000027195 | 56348 | 0.467141664291416 | 2.64404528242569 | 3.52796592401892 | 0.0035526991536 | 0.0105758590546285 |
| Zfp672 | ENSMUSG00000049755 | 319475 | -0.237657179833614 | 4.40006237373573 | -3.5266197455792 | 0.0035620153309 | 0.0106007050318977 |
| Wars2 | ENSMUSG00000004233 | 70560 | -0.524341660865547 | 1.68427462613491 | -3.52607367498574 | 0.0035658014262 | 0.0106084162516954 |
| Naip2 | ENSMUSG00000078945 | 17948 | 0.939530711119294 | 0.864484418677539 | 3.52596615061411 | 0.0035665474087 | 0.0106084162516954 |
| Srf | ENSMUSG00000015605 | 20807 | -0.219791658770675 | 5.68139399625343 | -3.52545582195415 | 0.0035700901175 | 0.0106160650365607 |
| G0s2 | ENSMUSG00000009633 | 14373 | -1.0896108837539 | 4.17907674718246 | -3.52363637535663 | 0.0035827497038 | 0.0106508123503531 |
| Pdia3 | ENSMUSG00000027248 | 14827 | 0.530130276821451 | 6.0406145428634 | 3.5228903648004 | 0.0035879534974 | 0.0106633821861904 |
| Tspyl2 | ENSMUSG00000041096 | 52808 | 0.321833193324118 | 3.18671655262995 | 3.52268782340692 | 0.0035893676418 | 0.0106646854288728 |
| Camta2 | ENSMUSG00000040712 | 216874 | -0.243542826655736 | 5.13468675060482 | -3.52145975072582 | 0.0035979541143 | 0.0106872925065134 |
| Satb1 | ENSMUSG00000023927 | 20230 | -0.322182960201213 | 5.77155115742184 | -3.51981095558332 | 0.0036095148501 | 0.0107187195945054 |
| Nagk | ENSMUSG00000034744 | 56174 | 0.445348742574214 | 1.43257762698503 | 3.51940976510877 | 0.0036123335201 | 0.0107241764554385 |
| Ap1b1 | ENSMUSG00000009090 | 11764 | 0.276197039099734 | 5.86777417307563 | 3.51839156741208 | 0.0036194971252 | 0.0107425259505335 |
| Zbtb17 | ENSMUSG00000006215 | 22642 | -0.239097128740006 | 4.98409884104013 | -3.5179292316406 | 0.0036227546588 | 0.0107492755634031 |
| Cyb5d1 | ENSMUSG00000044795 | 327951 | -0.40328211821295 | 2.43072698517284 | -3.51763913219089 | 0.0036248001587 | 0.010752426196773 |
| Kcna2 | ENSMUSG00000040724 | 16490 | -0.728097886333336 | 1.26685789587055 | -3.51654570521114 | 0.0036325204369 | 0.0107724039327272 |
| Hivep1 | ENSMUSG00000021366 | 110521 | 0.211306639595294 | 5.47787799018959 | 3.51580009120582 | 0.0036377944611 | 0.0107851183225306 |
| Cped1 | ENSMUSG00000062980 | 214642 | 0.290733067041516 | 5.25742712722188 | 3.51555760149231 | 0.0036395113536 | 0.0107872827024628 |
| Jade1 | ENSMUSG00000025764 | 269424 | -0.303621824492365 | 4.49544141344709 | -3.51463783668435 | 0.0036460309797 | 0.0108036770504772 |
| Prkcq | ENSMUSG00000026778 | 18761 | -0.332269498964687 | 5.8859474612871 | -3.51419291775008 | 0.0036491889584 | 0.0108101042019904 |
| Gsk3a | ENSMUSG00000057177 | 606496 | -0.197857632795954 | 4.82960082331449 | -3.5130357818626 | 0.0036574151068 | 0.0108315373971316 |
| Bag5 | ENSMUSG00000049792 | 70369 | -0.399056848990091 | 3.7215105071973 | -3.51222705195437 | 0.0036631755337 | 0.0108456586565149 |
| Cstf1 | ENSMUSG00000027498 | 67337 | -0.313556508345193 | 3.22876265637155 | -3.51161723976252 | 0.0036675251675 | 0.0108555964273784 |
| Pstk | ENSMUSG00000063179 | 214580 | -0.474906498377529 | 2.90604564452397 | -3.511438996113 | 0.0036687975186 | 0.0108564227602038 |
| Dazap2 | ENSMUSG00000000346 | 23994 | -0.22874300289669 | 5.02514480581563 | -3.51116701684797 | 0.0036707398389 | 0.010859230630521 |
| Ppfibp2 | ENSMUSG00000036528 | 19024 | -0.484728818978541 | 4.13274223596803 | -3.50997439005521 | 0.0036792691667 | 0.0108815181984435 |
| Chchd4 | ENSMUSG00000034203 | 72170 | -0.317996408108576 | 3.58611976341964 | -3.50908942000494 | 0.0036856111554 | 0.0108973263567104 |
| Cdh11 | ENSMUSG00000031673 | 12552 | 0.645201936881114 | 2.8013321194132 | 3.50873374048797 | 0.0036881631821 | 0.0109019231323962 |
| Caskin2 | ENSMUSG00000034471 | 140721 | -0.400593125932127 | 4.5810089093231 | -3.50633185968273 | 0.0037054436326 | 0.0109500417269831 |
| Slc25a30 | ENSMUSG00000022003 | 67554 | 0.497983056448385 | 1.67302036772974 | 3.50497666543929 | 0.0037152297022 | 0.0109759934797457 |
| Pcm1 | ENSMUSG00000031592 | 18536 | -0.235398537944408 | 6.72333019315029 | -3.50323558167389 | 0.0037278406544 | 0.0110102745727547 |
| Dact1 | ENSMUSG00000044548 | 59036 | 0.712065382886654 | 2.75407255746065 | 3.5021857503955 | 0.0037354656330 | 0.0110298149201624 |
| Cyp4b1 | ENSMUSG00000028713 | 13120 | -1.03503765424078 | 1.0124080997806 | -3.50121101657133 | 0.0037425592727 | 0.0110477762382314 |
| Retreg1 | ENSMUSG00000022270 | 66270 | -0.56125107243386 | 7.49793817658347 | -3.50002688558072 | 0.0037511951040 | 0.0110688198487735 |
| Ap2a1 | ENSMUSG00000060279 | 11771 | 0.226378723624924 | 5.59043618817428 | 3.49990101904895 | 0.0037521142269 | 0.0110688198487735 |
| Tbc1d10a | ENSMUSG00000034412 | 103724 | 0.480621116956733 | 1.81407807827047 | 3.49981727824916 | 0.0037527258585 | 0.0110688198487735 |
| Dzip1 | ENSMUSG00000042156 | 66573 | 0.870517567851432 | 1.15769929973631 | 3.498971999973 | 0.0037589052965 | 0.0110837156175599 |
| Slc39a3 | ENSMUSG00000046822 | 106947 | -0.464609604724039 | 3.12394238347384 | -3.49884918440147 | 0.0037598039987 | 0.0110837156175599 |
| Psmd13 | ENSMUSG00000025487 | 23997 | 0.257623732554887 | 5.07252428214432 | 3.49864362988003 | 0.0037613086279 | 0.0110851616641723 |
| Dcaf4 | ENSMUSG00000021222 | 73828 | 0.272385762174539 | 3.85935825733016 | 3.49778012693946 | 0.0037676359878 | 0.0111008164376033 |
| Ctnnb1 | ENSMUSG00000006932 | 12387 | -0.179524043005941 | 7.00655153429144 | -3.49627091215374 | 0.0037787206477 | 0.0111304757209074 |

| | | | | | | | |
|---|---|---|---|---|---|---|---|
| Selenop | ENSMUSG00000064373 | 20363 | 0.273173222661512 | 7.0111221180105 | 3.49556562894653 | 0.0037839119904 | 0.0111427645387963 |
| Spop | ENSMUSG00000057522 | 20747 | -0.27311199200572 | 6.87021191944994 | -3.49264921967195 | 0.0038054552095 | 0.0112005200924171 |
| Gpr153 | ENSMUSG00000042804 | 100129 | 0.66015974673989 | 3.53270911700807 | 3.49253688401422 | 0.0038062874935 | 0.0112005200924171 |
| Gatad1 | ENSMUSG00000007415 | 67210 | -0.264789106174602 | 5.04402350955239 | -3.49249486434042 | 0.0038065988604 | 0.0112005200924171 |
| Dzip3 | ENSMUSG00000064061 | 224170 | 0.461017498169022 | 2.34137320967055 | 3.49151537599896 | 0.0038138641842 | 0.0112188776634886 |
| Ppp4r3b | ENSMUSG00000020463 | 104570 | -0.227647641224491 | 4.51993021613342 | -3.48895657246441 | 0.0038329101401 | 0.0112718699870309 |
| Aldh7a1 | ENSMUSG00000053644 | 110695 | -0.318015597568517 | 3.8260328073214 | -3.48820242170915 | 0.0038385418095 | 0.0112853955136429 |
| Cnot9 | ENSMUSG00000026174 | 58184 | -0.339445335969376 | 3.2357255393544 | -3.48790557701382 | 0.0038407608062 | 0.0112888831334477 |
| Ahsa2 | ENSMUSG00000020288 | 268390 | 0.31248770090217 | 3.61273757226177 | 3.48765332554965 | 0.0038426474736 | 0.0112913923480994 |
| Serpinh1 | ENSMUSG00000070436 | 12406 | 0.629500434819735 | 6.21822449580547 | 3.48747882671492 | 0.0038439531522 | 0.0112921934713793 |
| Napepld | ENSMUSG00000044968 | 242864 | -0.34413465426662 | 3.53130429380561 | -3.4861806402063 | 0.0038536808323 | 0.0113177284198022 |
| Tasor | ENSMUSG00000040651 | 218850 | 0.320534155970014 | 4.1867227934545 | 3.48556552037695 | 0.0038582987709 | 0.0113282470764306 |
| Armc10 | ENSMUSG00000038525 | 67211 | 0.415347085254606 | 2.11151646188782 | 3.48444783282827 | 0.0038667039740 | 0.0113498767830914 |
| Rnf220 | ENSMUSG00000028677 | 66743 | -0.268780735784969 | 4.32879281067693 | -3.48272377397357 | 0.0038797054430 | 0.0113849826035116 |
| Sox13 | ENSMUSG00000070643 | 20668 | -0.689863977284683 | 2.88978188368015 | -3.48113126262223 | 0.0038917540771 | 0.0114172742398549 |
| Clcc1 | ENSMUSG00000027884 | 229725 | -0.277807166462318 | 3.84146220677546 | -3.48090087364536 | 0.0038935002792 | 0.0114193323189885 |
| Tbcb | ENSMUSG00000006095 | 66411 | 0.329968670783609 | 3.55617680254329 | 3.47850391563123 | 0.0039117146644 | 0.0114696762332483 |
| Lrch3 | ENSMUSG00000022801 | 70144 | 0.261183830739794 | 4.20796014079411 | 3.47805296128489 | 0.0039151510449 | 0.0114766736584722 |
| Eif3j1 | ENSMUSG00000027236 | 78655 | -0.277737563707697 | 5.00266920896407 | -3.47784749619225 | 0.0039167177481 | 0.0114781881247557 |
| Rapgef1 | ENSMUSG00000039844 | 107746 | 0.226944827650264 | 8.3669327511065 | 3.47717823988493 | 0.0039218253184 | 0.0114900757319515 |
| Selenot | ENSMUSG00000075700 | 69227 | -0.271508910286361 | 6.46326081709442 | -3.47533740412753 | 0.0039359087319 | 0.0115282471458904 |
| Cnp | ENSMUSG00000006782 | 12799 | 0.410604087027902 | 2.18190180718097 | 3.4741768292313 | 0.0039448139855 | 0.0115512353808026 |
| Dnajb11 | ENSMUSG00000004460 | 67838 | 0.309408967909563 | 4.85046686061928 | 3.47245616555748 | 0.0039580543004 | 0.0115869018633421 |
| Poglut3 | ENSMUSG00000034487 | 68304 | 0.469701170072682 | 3.38220069466076 | 3.47161195389376 | 0.0039645667986 | 0.0116028593511192 |
| Gbp5 | ENSMUSG00000105504 | 229898 | -0.652224444516986 | 2.34465023324732 | -3.47037931052273 | 0.0039740951923 | 0.0116276323734295 |
| Uxs1 | ENSMUSG00000057363 | 67883 | 0.502440960908851 | 3.21911588790687 | 3.47018571769656 | 0.0039755937717 | 0.0116289043401956 |
| Dhrs3 | ENSMUSG00000066026 | 20148 | -0.442040293404243 | 4.27359534534537 | -3.4699340674961 | 0.0039775426186 | 0.0116314923394258 |
| Rev1 | ENSMUSG00000026082 | 56210 | -0.341534755121755 | 3.75868659458938 | -3.46969237272845 | 0.0039794152752 | 0.011633450057202 |
| Itpkb | ENSMUSG00000038855 | 320404 | -0.608071646035381 | 5.34827329185151 | -3.46957298649467 | 0.0039803406105 | 0.011633450057202 |
| Ttc39b | ENSMUSG00000038172 | 69863 | 0.371529206786282 | 3.63614040038905 | 3.4670987977458 | 0.0039995664112 | 0.0116865170921723 |
| Uxt | ENSMUSG00000001134 | 22294 | -0.443546619537406 | 2.32674236488119 | -3.46394407998494 | 0.0040242162064 | 0.0117554001477447 |
| Smarca5 | ENSMUSG00000031715 | 93762 | -0.202998020168305 | 6.09149983896818 | -3.46360242872721 | 0.0040268949209 | 0.0117600823885732 |
| Lrrn4cl | ENSMUSG00000071656 | 68852 | 0.475606878176212 | 3.17032347439407 | 3.46298542685203 | 0.0040317370768 | 0.0117710785222922 |
| Psmd5 | ENSMUSG00000026869 | 66998 | 0.368277196824407 | 4.09052779174184 | 3.46247073124099 | 0.0040357808374 | 0.0117797384068592 |
| Gfod1 | ENSMUSG00000051335 | 328232 | 0.308434038962329 | 4.1574870713187 | 3.46224489258876 | 0.0040375564534 | 0.0117817751182061 |
| Rcc2 | ENSMUSG00000040945 | 108911 | 0.347785213599331 | 3.19993988171928 | 3.4615094926222 | 0.0040433438654 | 0.0117955142229703 |
| Inca1 | ENSMUSG00000057054 | 103844 | -0.705413813366439 | 1.32076232392484 | -3.46110377034443 | 0.0040465403772 | 0.0118016896648821 |
| Wipi1 | ENSMUSG00000041895 | 52639 | -0.212260406817744 | 5.99099442854872 | -3.45990623835629 | 0.0040559900865 | 0.0118260943281113 |
| Arpc5l | ENSMUSG00000026755 | 74192 | -0.281673740740364 | 5.30598450762363 | -3.45976247033242 | 0.0040571260531 | 0.0118262519700033 |
| Chmp1b | ENSMUSG00000109901 | 67064 | 0.275120780844126 | 3.79244091053667 | 3.45857708306735 | 0.0040665044908 | 0.0118490364712348 |
| Fxr2 | ENSMUSG00000018765 | 23879 | -0.227047362716633 | 5.88306929619574 | -3.45850058181267 | 0.0040671104967 | 0.0118490364712348 |
| Commd1 | ENSMUSG00000051355 | 17846 | -0.324631416831755 | 4.55844182258739 | -3.45800450377246 | 0.0040710423972 | 0.0118573313202236 |
| Gfod2 | ENSMUSG00000013150 | 70575 | -0.639859086100074 | 1.80507066212351 | -3.45782805007629 | 0.0040724418882 | 0.011858247810 33 |
| Pcdhgb1 | ENSMUSG00000103037 | 93699 | 0.726774380713943 | 0.862329199049029 | 3.45688037202605 | 0.0040799664223 | 0.0118769943131 1264 |
| Ramp2 | ENSMUSG00000001240 | 54409 | -0.607634040049709 | 3.32220472256686 | -3.45554095267794 | 0.0040906252831 | 0.0119048522285244 |
| Tbc1d7 | ENSMUSG00000021368 | 67046 | -0.3424140645381 | 2.87174115409693 | -3.45442419261418 | 0.0040995337202 | 0.0119246418493608 |
| Rab3a | ENSMUSG00000031840 | 19339 | -0.243949796833872 | 3.86951891609372 | -3.45441501696753 | 0.0040996069956 | 0.0119246418493608 |
| Pitpnm3 | ENSMUSG00000040543 | 327958 | -1.04695908433153 | 0.898309104459239 | -3.45400552814042 | 0.0041028784610 | 0.011930982829794 |
| Snx13 | ENSMUSG00000020590 | 217463 | -0.204492896456617 | 5.81146972273861 | -3.45321709942852 | 0.0041091847456 | 0.011946143206 13 |
| Mxd1 | ENSMUSG00000001156 | 17119 | 0.468837717969454 | 2.22764294465235 | 3.45163091740675 | 0.0041219015187 | 0.0119799270144566 |
| Dpcd | ENSMUSG00000041035 | 226162 | -0.436236081935826 | 1.70772983837228 | -3.45146385948845 | 0.0041232431662 | 0.0119806408958695 |
| Gas2l3 | ENSMUSG00000074802 | 237436 | 0.71011385007515 | 1.0684907421131 | 3.45058070159237 | 0.0041303431541 | 0.011998081588701 |
| Camsap2 | ENSMUSG00000041570 | 67886 | 0.226678943176921 | 4.84239611824536 | 3.45018338019599 | 0.0041335413657 | 0.0120023778130952 |
| Rragd | ENSMUSG00000028278 | 52187 | -0.295329417217294 | 6.39349839164853 | -3.45012417392086 | 0.0041340181562 | 0.0120023778130952 |
| Ipo9 | ENSMUSG00000041879 | 226432 | 0.197352221067949 | 4.84558956174401 | 3.44994860080668 | 0.0041354323797 | 0.0120032956299823 |
| Tenm4 | ENSMUSG00000048078 | 23966 | 0.726681546215377 | 1.36956368325245 | 3.44951489520053 | 0.0041389279244 | 0.0120102524932663 |
| Rnf150 | ENSMUSG00000047747 | 330812 | -0.272643522000065 | 4.5547706910412 | -3.44937348471872 | 0.0041400682967 | 0.0120103732886657 |
| Rxrb | ENSMUSG00000039656 | 20182 | -0.310852336619392 | 3.70435696971695 | -3.44894818437292 | 0.0041434999425 | 0.0120171392602504 |
| Prepl | ENSMUSG00000024127 | 213760 | 0.354943465765863 | 5.06508509373674 | 3.44801970489159 | 0.0041510015846 | 0.0120357024725602 |
| Smoc2 | ENSMUSG00000023886 | 64074 | 0.462774533811572 | 5.71149109288505 | 3.44732489206207 | 0.0041566242729 | 0.0120472041509199 |
| Ephb4 | ENSMUSG00000029710 | 13846 | -0.512771214506212 | 4.5317451384342 | -3.44725718016567 | 0.0041571726335 | 0.0120472041509199 |
| Larp1b | ENSMUSG00000025762 | 214048 | -0.308636557706622 | 5.0184453899996 | -3.44651463468106 | 0.0041631908786 | 0.0120614469904608 |
| Adamtsl1 | ENSMUSG00000066113 | 77739 | 0.528337042247816 | 2.31089368644879 | 3.44488780641031 | 0.0041764068456 | 0.0120965297376839 |
| Ngrn | ENSMUSG00000047084 | 83485 | -0.294699748560557 | 3.11131713871452 | -3.4430181369351 | 0.0041916478280 | 0.0121374575916768 |
| Ptpra | ENSMUSG00000027303 | 19262 | 0.264122825512996 | 4.44524310849161 | 3.44124692710822 | 0.0042061378971 | 0.0121752792461483 |
| Sec61a2 | ENSMUSG00000025816 | 57743 | -0.393276623674792 | 3.06231745756285 | -3.44114941161421 | 0.0042069371249 | 0.0121752792461483 |
| Rassf5 | ENSMUSG00000026430 | 54354 | 0.901775827381118 | 0.663152553812535 | 3.43992414965006 | 0.0042169923080 | 0.012201149528695 |
| Pak4 | ENSMUSG00000030602 | 70584 | 0.374954153822121 | 2.08969522188957 | 3.43908511577598 | 0.0042238918549 | 0.0122178782392093 |
| Ehd2 | ENSMUSG00000074364 | 259300 | -0.446221591200429 | 5.96765793590021 | -3.43807081155025 | 0.0042322478749 | 0.0122388099261247 |
| Man1b1 | ENSMUSG00000036646 | 227619 | 0.284652676575013 | 3.52798574084039 | 3.43751624624925 | 0.0042368235242 | 0.0122488013603189 |
| Dusp28 | ENSMUSG00000047067 | 67446 | -0.431250204044165 | 2.42541908322935 | -3.43644855461087 | 0.0042456469561 | 0.012271064748313 |
| Ubac1 | ENSMUSG00000036352 | 98766 | -0.255657052339763 | 6.42285830697144 | -3.4361995613067 | 0.0042477070430 | 0.012273590423037 |
| Mapre1 | ENSMUSG00000027479 | 13589 | 0.21646457641005 | 5.12999213376793 | 3.43607160973215 | 0.0042487664587 | 0.012273590423037 |
| Rap2a | ENSMUSG00000051615 | 76108 | 0.563709311657287 | 2.82062041547066 | 3.43581752523053 | 0.0042508705021 | 0.0122764241634799 |
| Hp | ENSMUSG00000031722 | 15439 | 1.90671330881196 | 1.07430698176425 | 3.43472288912 | 0.0042599470631 | 0.012299387571789 |
| Atxn1l | ENSMUSG00000069895 | 52335 | 0.318772157698318 | 4.56358852047577 | 3.43400401259556 | 0.0042659185009 | 0.0123133760585687 |
| Gm15446 | ENSMUSG00000090015 | 100642166 | -0.812077427066167 | 1.79729636030221 | -3.43331000893251 | 0.0042716913297 | 0.0123267840352062 |
| Amacr | ENSMUSG00000022244 | 17117 | -0.30251564200573 | 3.30477905598714 | -3.43307321665436 | 0.0042736628044 | 0.0123292182938586 |
| Grcc10 | ENSMUSG00000072772 | 14790 | -0.219642299727509 | 4.98754698039957 | -3.43069358005316 | 0.0042935260422 | 0.0123801745563047 |
| Cgref1 | ENSMUSG00000029161 | 68567 | 0.717315922265974 | 2.0340925435406 | 3.43068583241173 | 0.0042935908647 | 0.0123801745563047 |
| Map4k4 | ENSMUSG00000026074 | 26921 | 0.302468848155579 | 6.27349596846482 | 3.43013613960122 | 0.0042981925232 | 0.012390174702476 |
| Sdf2l1 | ENSMUSG00000022769 | 64136 | 0.955285445655189 | 1.16130379588775 | 3.42897692172856 | 0.0043079130077 | 0.0124149214574 7 |
| Rapgef5 | ENSMUSG00000041992 | 217944 | -0.449442839702532 | 2.95265164219289 | -3.42879933901358 | 0.0043094040613 | 0.0124159451225398 |
| Tmem50a | ENSMUSG00000028822 | 71817 | 0.350093486876506 | 3.70427860093301 | 3.42745949026196 | 0.0043206707236 | 0.0124451256073988 |
| Frzb | ENSMUSG00000027004 | 20378 | -0.734306253733437 | 1.31182352503096 | -3.42583064532226 | 0.0043344074953 | 0.0124814036701702 |
| Clasp2 | ENSMUSG00000033392 | 76499 | -0.220795820407389 | 5.49557613021372 | -3.42558912004072 | 0.0043364481267 | 0.0124839911703727 |
| Smg9 | ENSMUSG00000002210 | 71997 | 0.318523165976222 | 2.9925365101187 | 3.42443072455403 | 0.0043462487929 | 0.0125089114154034 |
| Cmc1 | ENSMUSG00000039163 | 67899 | -0.457643660865457 | 2.41037831250284 | -3.42361659570494 | 0.0043531501192 | 0.0125254761656906 |
| Etl4 | ENSMUSG00000036617 | 208618 | -0.218160697761389 | 7.33351114689579 | -3.42259048850683 | 0.0043618640981 | 0.0125451817071748 |
| Stam2 | ENSMUSG00000055371 | 56324 | 0.415082442838615 | 3.02961956434458 | 3.42253991971628 | 0.0043622939958 | 0.0125451817071748 |
| Wnt4 | ENSMUSG00000036856 | 22417 | -0.888087322359273 | 1.0618381007746 | -3.42154591425486 | 0.0043707529489 | 0.0125662021265465 |
| A930016O22Rik | ENSMUSG00000040705 | 100503495 | -0.546027948083944 | 2.22080079201658 | -3.42096121928339 | 0.0043757363976 | 0.0125772218151096 |
| Tufm | ENSMUSG00000073838 | 233870 | -0.250784721844632 | 5.72171861548734 | -3.4199173517839 | 0.0043846476785 | 0.0125995225484185 |
| 1110032A03Rik | ENSMUSG00000037971 | 68721 | -0.383120028857969 | 2.02788118517697 | -3.41703214293697 | 0.0044093732868 | 0.0126672429431641 |
| Tmbim4 | ENSMUSG00000020225 | 68212 | 0.295731073063353 | 3.08704930264224 | 3.41641578782157 | 0.0044146734887 | 0.0126791371269875 |
| Tmem80 | ENSMUSG00000025505 | 71448 | -0.599368009228254 | 1.68891304088101 | -3.41437391850992 | 0.0044322779362 | 0.0127263541164623 |
| Slc35a1 | ENSMUSG00000028293 | 24060 | -0.37696486570823 | 2.4553660252437 | -3.41347404102012 | 0.0044400588606 | 0.0127453475329498 |
| Pafah1b1 | ENSMUSG00000020745 | 18472 | -0.191815354633975 | 6.17679564456509 | -3.41277049437414 | 0.0044461517590 | 0.0127564587435829 |
| Retnla | ENSMUSG00000061100 | 57262 | 0.663179964724969 | 1.48739961429787 | 3.41275758494115 | 0.0044462636367 | 0.0127564587435829 |
| Rnf26 | ENSMUSG00000053128 | 213211 | 0.475455651537482 | 1.82051062680255 | 3.41198352206326 | 0.0044529771202 | 0.0127723675940662 |
| Denr | ENSMUSG00000023106 | 68184 | -0.249518215482111 | 4.70145752244312 | -3.41159546297598 | 0.0044563466143 | 0.0127746827726584 |
| Rabgap1l | ENSMUSG00000026721 | 29809 | 0.341705547858817 | 3.69550413521302 | 3.4115641284685 | 0.0044566188019 | 0.0127746827726584 |
| Rpl37rt | ENSMUSG00000072692 | 100502825 | 0.390435433411121 | 3.07744489360104 | 3.41144590747894 | 0.0044576458810 | 0.0127746827726584 |
| Slc19a2 | ENSMUSG00000040918 | 116914 | -0.526073676068934 | 1.63570906989738 | -3.41135233941455 | 0.0044584589495 | 0.0127746827726584 |
| Zfp950 | ENSMUSG00000074733 | 414758 | 0.559295688027317 | 2.49290547716985 | 3.41054742799079 | 0.0044654594727 | 0.0127913882327339 |
| Ttc38 | ENSMUSG00000035944 | 239570 | -1.16708141666289 | 1.4367444458584 | -3.4099449190405 | 0.0044707068898 | 0.0128030644517911 |
| Zfp830 | ENSMUSG00000046010 | 66983 | -0.328638178554046 | 2.97692619226993 | -3.40956863314276 | 0.0044739872162 | 0.0128091027399529 |
| Kctd3 | ENSMUSG00000026608 | 226823 | 0.232925998762207 | 3.86177213170857 | 3.40863202728019 | 0.0044821627376 | 0.0128287928616592 |
| Uspl1 | ENSMUSG00000041264 | 231915 | -0.235419439017206 | 4.19102321302505 | -3.40851196316297 | 0.0044832118499 | 0.0128287928616592 |
| Ubtd2 | ENSMUSG00000044949 | 327900 | 0.655681144956215 | 1.62847744875063 | 3.40812320953719 | 0.0044866104499 | 0.0128318474015867 |
| Mrpl44 | ENSMUSG00000026248 | 69163 | -0.274932161917628 | 3.55710684901194 | -3.40812130632846 | 0.0044866270943 | 0.0128318474015867 |
| Vars2 | ENSMUSG00000038838 | 68915 | -0.321737948666633 | 3.10723657486605 | -3.40607025001101 | 0.0045046010879 | 0.0128798834665328 |
| Birc6 | ENSMUSG00000024073 | 12211 | 0.250196186493223 | 6.97140151249887 | 3.40536418407591 | 0.0045108053010 | 0.0128942501948855 |
| Adh5 | ENSMUSG00000028138 | 11532 | -0.281676731200499 | 4.02014624215482 | -3.4045589027537 | 0.0045178918205 | 0.0129111308472223 |
| Mkln1 | ENSMUSG00000025609 | 27418 | -0.215632120129345 | 5.62894611395536 | -3.40427327792978 | 0.0045204080257 | 0.012914945146197 |
| Ndfip1 | ENSMUSG00000024425 | 65113 | 0.221206134667172 | 5.3635784510374 | 3.40329033666443 | 0.0045290780076 | 0.0129363343875578 |

| | | | | | | | |
|---|---|---|---|---|---|---|---|
| Nudt16l1 | ENSMUSG00000022516 | 66911 | 0.440474403048931 | 2.20502782576548 | 3.40314401435347 | 0.0045303700662 | 0.0129366445124901 |
| Amigo3 | ENSMUSG00000032593 | 320844 | -1.14562850841429 | 0.510645936383213 | -3.39899159399065 | 0.0045671910898 | 0.0130383846322975 |
| Pcmtd2 | ENSMUSG00000027589 | 245867 | -0.266230272796466 | 4.33985488421572 | -3.39874928275498 | 0.0045693498955 | 0.0130411393494544 |
| Zfp629 | ENSMUSG00000045639 | 320683 | -0.515662546949496 | 3.89669742370011 | -3.39843700699239 | 0.0045721324887 | 0.0130449884223359 |
| Csad | ENSMUSG00000023044 | 246277 | 0.672411213433606 | 1.36241931920763 | 3.39833011970336 | 0.0045730853201 | 0.0130449884223359 |
| Mettl1 | ENSMUSG00000006732 | 17299 | 0.391891608846837 | 2.72573822961168 | 3.39752338847828 | 0.0045802832579 | 0.0130621122600467 |
| Fam118b | ENSMUSG00000050471 | 109229 | -0.366803552652714 | 2.29553299948971 | -3.39591926849994 | 0.0045946296657 | 0.0130993165997143 |
| Eid2b | ENSMUSG00000070705 | 434156 | -0.543502279619237 | 1.54332785557519 | -3.39579690888129 | 0.0045957258402 | 0.0130993165997143 |
| Zdhhc18 | ENSMUSG00000037553 | 503610 | 0.323598217604191 | 3.72046511366582 | 3.39540894921382 | 0.0045992031673 | 0.0131058106938894 |
| Rny3 | ENSMUSG00000064945 | 19874 | 1.40981322762382 | 3.58468543928909 | 3.3948333956548 | 0.0046043667975 | 0.0131146751161795 |
| Efna1 | ENSMUSG00000027954 | 13636 | -0.499008846244284 | 2.70384524573217 | -3.39479477619331 | 0.0046047134842 | 0.0131146751161795 |
| Ednra | ENSMUSG00000031616 | 13617 | 0.436548229249974 | 2.52566573201391 | 3.3941818821301 | 0.0046102189482 | 0.0131269349630847 |
| Crebzf | ENSMUSG00000051451 | 233490 | 0.363323363168955 | 3.01901474015217 | 3.39313145832088 | 0.0046196700319 | 0.0131504200830076 |
| Ptbp1 | ENSMUSG00000006498 | 19205 | 0.382418710677084 | 4.08924950958126 | 3.39199343378303 | 0.0046299312952 | 0.0131761986430788 |
| Fam50a | ENSMUSG00000001962 | 108160 | -0.303203989054012 | 3.28640989141908 | -3.39184641327841 | 0.0046312586113 | 0.0131765455180812 |
| Abhd16a | ENSMUSG00000007036 | 193742 | -0.266269776379089 | 4.79163745458495 | -3.39118123506888 | 0.0046372686938 | 0.0131902118377368 |
| Scrn2 | ENSMUSG00000020877 | 217140 | -0.465083891897036 | 1.54398535692646 | -3.38897453026488 | 0.0046572631752 | 0.0132436378171295 |
| Hsf2 | ENSMUSG00000019878 | 15500 | 0.361259862374443 | 4.81201398301912 | 3.38816714359994 | 0.0046646003843 | 0.0132610524839923 |
| Irak2 | ENSMUSG00000060477 | 108960 | 0.675622898484973 | 1.9456684205664 | 3.3876388642043 | 0.0046694074694 | 0.0132712670433205 |
| Bag2 | ENSMUSG00000042215 | 213539 | 0.328213914568125 | 4.73713750358112 | 3.38737677842227 | 0.0046717941714 | 0.0132745989311092 |
| Papss1 | ENSMUSG00000028032 | 23971 | 0.406610011007479 | 2.36601864615078 | 3.38556481845932 | 0.0046883285106 | 0.0133181182130127 |
| Enho | ENSMUSG00000028445 | 69638 | 0.474865584478089 | 1.51492582711966 | 3.38518133576958 | 0.0046918353733 | 0.0133246174241948 |
| 2410006H16Rik | ENSMUSG00000086841 | 69221 | 0.610254872245229 | 1.44568882900618 | 3.38455431766767 | 0.0046975749970 | 0.0133374525437037 |
| Derl1 | ENSMUSG00000022365 | 67819 | 0.264652176829501 | 4.86710386868789 | 3.38287317613184 | 0.0047129987737 | 0.0133777693055555 |
| Fry | ENSMUSG00000056602 | 320365 | -0.348633900270022 | 6.49470654783322 | -3.38264601003264 | 0.0047150868310 | 0.0133802217420056 |
| Tspan3 | ENSMUSG00000032324 | 56434 | 0.302874204517732 | 5.98282564313296 | 3.38206403104693 | 0.0047204404909 | 0.0133919374530982 |
| Alkbh7 | ENSMUSG00000002661 | 66400 | -0.568324903223349 | 2.02396024629815 | -3.38165974757152 | 0.0047241631172 | 0.0133990210260369 |
| Smtn | ENSMUSG00000020439 | 29856 | -0.227094914080779 | 5.96013657293363 | -3.38041335475737 | 0.0047356584454 | 0.0134281407176899 |
| Amn1 | ENSMUSG00000068250 | 232566 | 0.430281389860921 | 2.12539422911861 | 3.37745580544952 | 0.0047630482543 | 0.0135023030260055 |
| Heatr3 | ENSMUSG00000031657 | 234549 | -0.24370833967696 | 4.47188293998042 | -3.37686535571401 | 0.0047685354452 | 0.0135143533709133 |
| Klhl8 | ENSMUSG00000029312 | 246293 | -0.357281887463491 | 2.94960390889054 | -3.37623668739536 | 0.0047743847975 | 0.0135274235931366 |
| Stoml2 | ENSMUSG00000028455 | 66592 | -0.267687223164793 | 4.79032519963157 | -3.37552052939918 | 0.0047810569759 | 0.013541118626473 |
| Arhgap19 | ENSMUSG00000025154 | 71085 | -0.421470932085703 | 1.95253431416054 | -3.37545200309676 | 0.0047816959013 | 0.013541118626473 |
| Wnt9a | ENSMUSG00000000126 | 216795 | 0.410341984776777 | 3.14316420246492 | 3.37434162475041 | 0.0047920608191 | 0.0135669559218827 |
| Ldha | ENSMUSG00000063229 | 16828 | -0.262216030432047 | 8.09380318271339 | -3.3732269846936 | 0.0048024882382 | 0.0135929567407322 |
| Psmb2 | ENSMUSG00000028837 | 26445 | -0.236783583908069 | 4.25579794780091 | -3.37257356476285 | 0.0048086115623 | 0.0136067649463066 |
| Ankrd44 | ENSMUSG00000052331 | 329154 | 0.4277894827593 | 2.41773760696289 | 3.37230797341431 | 0.0048111027131 | 0.0136102908273792 |
| Zfp354c | ENSMUSG00000044807 | 30944 | 0.503581261648147 | 2.13881232696014 | 3.37179028750655 | 0.0048159621496 | 0.0136192241858164 |
| Dhx9 | ENSMUSG00000042699 | 13211 | 0.226652678184275 | 5.10459634832718 | 3.37170614801461 | 0.0048167524199 | 0.0136192241858164 |
| Sdhaf1 | ENSMUSG00000074211 | 68332 | -0.416765601560553 | 3.17716311082454 | -3.37088014637237 | 0.0048245174687 | 0.0136376520432877 |
| Elf2 | ENSMUSG00000037174 | 69257 | -0.219434559024618 | 4.95648367328659 | -3.37049219906447 | 0.0048281688130 | 0.0136444450092963 |
| Abca2 | ENSMUSG00000026944 | 11305 | 0.22102693337349 | 4.82664745653288 | 3.37001096096847 | 0.0048327020672 | 0.0136537261042981 |
| Arpc4 | ENSMUSG00000079426 | 68089 | 0.266454448421694 | 4.03715280882164 | 3.36944992253269 | 0.0048379924415 | 0.0136651409246987 |
| Sec61b | ENSMUSG00000053317 | 66212 | 0.434653394237204 | 2.07016671513198 | 3.36848002555699 | 0.0048471519285 | 0.0136841499728416 |
| Snrnp35 | ENSMUSG00000029402 | 76167 | -0.597675299695749 | 1.87104155562509 | -3.36847217717554 | 0.0048472261179 | 0.0136841499728416 |
| Hnrnph1 | ENSMUSG00000007850 | 59013 | -0.211726702482498 | 5.68545334769702 | -3.3673932077355 | 0.0048574363019 | 0.0137094335496652 |
| Coq4 | ENSMUSG00000026798 | 227683 | -0.514609632873721 | 1.80712853163049 | -3.36560536913577 | 0.0048744020576 | 0.013753765847099 |
| Sgsm2 | ENSMUSG00000038351 | 97761 | 0.545865095082956 | 2.00899438916981 | 3.36511205686133 | 0.0048790938374 | 0.0137634515449711 |
| Gbp3 | ENSMUSG00000028268 | 55932 | -0.915660707714943 | 3.6628696708984 | -3.36445825877158 | 0.0048853189560 | 0.0137774565296004 |
| S100a11 | ENSMUSG00000027907 | 20195 | 0.573991652412733 | 3.32685893329734 | 3.3633821011889 | 0.0048955829469 | 0.0138028416798914 |
| Plekhb1 | ENSMUSG00000030701 | 27276 | -0.498795224551323 | 2.78574020141095 | -3.36223446426221 | 0.0049065525633 | 0.0138274825445465 |
| Necap2 | ENSMUSG00000028923 | 66147 | 0.324651594872567 | 3.62413072899403 | 3.36218427365448 | 0.0049070328705 | 0.0138274825445465 |
| Ppip5k2 | ENSMUSG00000040648 | 227399 | 0.350983202540441 | 2.66107946607791 | 3.36207095256125 | 0.0049081174890 | 0.0138274825445465 |
| Dab2ip | ENSMUSG00000026883 | 69601 | -0.282593771792874 | 6.19584931197935 | -3.36125648368609 | 0.0049159200232 | 0.0138458958450435 |
| Hus1 | ENSMUSG00000020413 | 15574 | -0.36384294925341 | 2.26831273637097 | -3.3610247266448 | 0.0049181425110 | 0.0138485872715743 |
| Mrpl41 | ENSMUSG00000036850 | 107733 | -0.394476490185062 | 3.59559323614987 | -3.35896316696802 | 0.0049379568089 | 0.0139007998657184 |
| Cwc15 | ENSMUSG00000004096 | 66070 | -0.223175240726755 | 5.34682472883539 | -3.35865097133541 | 0.0049409644054 | 0.0139056853542606 |
| Fam91a1 | ENSMUSG00000037119 | 210998 | 0.319729247640138 | 3.52760713317298 | 3.35849287795532 | 0.0049424881303 | 0.0139063932440852 |
| Fam234a | ENSMUSG00000024187 | 106581 | 0.218565872155304 | 5.39222558923256 | 3.35806725009418 | 0.0049465927354 | 0.0139143605740616 |
| Adamts6 | ENSMUSG00000046169 | 108154 | 0.689275705020745 | 2.1231575697056 | 3.3568084041208 | 0.0049587526601 | 0.0139449769030433 |
| Ippk | ENSMUSG00000021385 | 75678 | 0.425146165507942 | 2.74126692015641 | 3.35608113156024 | 0.0049657914930 | 0.0139611797350891 |
| Akt2 | ENSMUSG00000004056 | 11652 | -0.22232212253873 | 6.4751114437105 | -3.35564358762161 | 0.0049700310530 | 0.0139695061561278 |
| Zfp51 | ENSMUSG00000023892 | 22709 | -0.36844682499944 | 2.74545831656327 | -3.35517754758491 | 0.0049745507246 | 0.0139786154166335 |
| Syvn1 | ENSMUSG00000024807 | 74126 | 0.353422628010089 | 3.71831730908604 | 3.35473481824444 | 0.0049788481532 | 0.0139870956471647 |
| Fam20a | ENSMUSG00000020614 | 208659 | 0.781445568116527 | 1.79355534711529 | 3.35434861075964 | 0.0049825999888 | 0.0139940391773052 |
| Fam135a | ENSMUSG00000026153 | 68187 | -0.281417728582975 | 4.41426028845761 | -3.35405367662557 | 0.0049854670568 | 0.0139984948365902 |
| Tubb4b | ENSMUSG00000036752 | 227613 | 0.443663103634441 | 6.89082986392697 | 3.35280618014401 | 0.0049976123708 | 0.0140289935351233 |
| Mob1b | ENSMUSG00000006262 | 68473 | 0.261751730670896 | 4.97337669775603 | 3.3516398330126 | 0.0050089945225 | 0.0140573348206428 |
| Cand1 | ENSMUSG00000020114 | 71902 | 0.294239201700627 | 4.47436305443622 | 3.35082772073756 | 0.0050169351427 | 0.0140760056586079 |
| Snx6 | ENSMUSG00000005656 | 72183 | 0.276748613937948 | 3.43915373972171 | 3.35047318423328 | 0.0050204056718 | 0.0140821284061401 |
| Hint2 | ENSMUSG00000028470 | 68917 | -0.326212279340333 | 2.55827823209914 | -3.35012032326748 | 0.0050238621939 | 0.0140865132445873 |
| Scaf1 | ENSMUSG00000038406 | 233208 | 0.204741998836151 | 5.58740896068551 | 3.3500505191688 | 0.0050245462574 | 0.0140865132445873 |
| Urm1 | ENSMUSG00000069020 | 68205 | -0.342848392790196 | 2.99735688531104 | -3.34837180369915 | 0.0050410254833 | 0.014129089630268 |
| Tom1l1 | ENSMUSG00000020541 | 71943 | -0.318061981974888 | 2.52749199161397 | -3.34789279899764 | 0.0050457376144 | 0.0141386715876239 |
| Zbtb10 | ENSMUSG00000069114 | 229055 | -0.449020748432627 | 3.38574024478826 | -3.34610657380854 | 0.0050633483612 | 0.014181155840987 |
| Tnfrsf11a | ENSMUSG00000026321 | 21934 | 0.846516674922378 | 1.2676775880519 | 3.34609184340725 | 0.0050634938475 | 0.014181155840987 |
| Sco2 | ENSMUSG00000091780 | 100126824 | -0.575171620405401 | 1.97314888829796 | -3.34574438693321 | 0.0050669267547 | 0.0141871353372138 |
| 0610009B22Rik | ENSMUSG00000007777 | 66050 | -0.468785785199741 | 1.67109866846615 | -3.34547836288315 | 0.0050695566809 | 0.0141908640408722 |
| Acp6 | ENSMUSG00000028093 | 66659 | -0.335543266201082 | 2.87379001460904 | -3.3452724080126 | 0.0050715937020 | 0.0141929315183921 |
| Chmp2b | ENSMUSG00000004843 | 68942 | -0.221640333948891 | 4.2625999090767 | -3.34477549917687 | 0.0050765123181 | 0.0142030601867832 |
| Foxred1 | ENSMUSG00000039048 | 235169 | -0.354801641064297 | 2.97742474679978 | -3.34339317495657 | 0.0050902184944 | 0.0142377631429484 |
| Usp40 | ENSMUSG00000005501 | 227334 | -0.3329937571612 | 3.71023578688419 | -3.34137333119015 | 0.0051103134632 | 0.014290313754496 |
| Fubp3 | ENSMUSG00000026843 | 320267 | 0.295400847041031 | 3.0045967246391 | 3.34085702040257 | 0.0051154628956 | 0.0143010549647731 |
| Slfn3 | ENSMUSG00000018986 | 20557 | -0.859449167000766 | 1.45492050146572 | -3.34010638522472 | 0.0051229586619 | 0.014318348538405 |
| Cnnm3 | ENSMUSG00000001138 | 94218 | 0.462865011029912 | 2.76208746496008 | 3.33992710424281 | 0.0051247505747 | 0.0143196954327003 |
| Ccdc25 | ENSMUSG00000022035 | 67179 | -0.230613886539351 | 3.9891878491541 | -3.33939736546497 | 0.0051300489905 | 0.0143301189837031 |
| Srr | ENSMUSG00000001323 | 27364 | -0.309054385013902 | 3.44635439828622 | -3.33929206137343 | 0.0051311028910 | 0.0143301189837031 |
| Ddx1 | ENSMUSG00000037149 | 104721 | -0.242363994731067 | 6.30391823903906 | -3.33878019277862 | 0.0051362288533 | 0.0143407707779575 |
| U2af1l4 | ENSMUSG00000078765 | 233073 | -0.387795164387874 | 3.60357781378247 | -3.33857641687515 | 0.0051382709398 | 0.0143428088977348 |
| Mrpl9 | ENSMUSG00000028140 | 78523 | -0.24755638343092 | 4.30839048717447 | -3.33841488487204 | 0.0051398902695 | 0.0143436662078914 |
| Tada2b | ENSMUSG00000029196 | 231151 | -0.321308450777716 | 3.85662353461792 | -3.338206236765 | 0.0051419826883 | 0.014343955072384 |
| Cited2 | ENSMUSG00000039910 | 17684 | 0.457058798560431 | 5.62221725069669 | 3.33808982472384 | 0.0051431504932 | 0.014343955072384 |
| Zfp617 | ENSMUSG00000066880 | 170938 | -0.30549500730125 | 4.1880539696161 | -3.33799455240307 | 0.0051441064301 | 0.014343955072384 |
| Prrg1 | ENSMUSG00000047996 | 546336 | 0.50427269013146 | 1.46424775726808 | 3.33788133157685 | 0.0051452426894 | 0.014343955072384 |
| Rbms1 | ENSMUSG00000026970 | 56878 | 0.328974237246276 | 4.84990089566305 | 3.33743101296069 | 0.0051497644833 | 0.0143529004506867 |
| D1Ertd622e | ENSMUSG00000044768 | 52392 | 0.673947476995386 | 1.69608488512368 | 3.3358534664437 | 0.0051656366321 | 0.0143934677608873 |
| Qser1 | ENSMUSG00000074994 | 99003 | -0.286150940765022 | 6.67312941371144 | -3.33536963875146 | 0.0051705143972 | 0.0144033875831802 |
| Bicral | ENSMUSG00000036568 | 210982 | 0.298890143952054 | 3.82151364172856 | 3.33459755086703 | 0.0051783078712 | 0.0144214225071101 |
| Mlh3 | ENSMUSG00000021245 | 217716 | -0.264144129528144 | 3.08252926205004 | -3.33285063213251 | 0.0051959848718 | 0.0144669665395833 |
| Hist2h2aa1 | ENSMUSG00000064220 | 15267 | -0.301165129421094 | 6.17224486367458 | -3.33263164362382 | 0.0051982050769 | 0.01446726901443 |
| Pi4kb | ENSMUSG00000038861 | 107650 | 0.2302403058939 | 3.76986746365603 | 3.33257884400935 | 0.0051987405259 | 0.01446726901443 |
| Chd3 | ENSMUSG00000018474 | 216848 | 0.456926430698265 | 3.99820686338873 | 3.33225064470682 | 0.0052020700874 | 0.0144728501210042 |
| Adcy4 | ENSMUSG00000022220 | 104110 | -0.626176660390959 | 2.85695392247961 | -3.33127508714752 | 0.0052119797029 | 0.0144967303138178 |
| Irf3 | ENSMUSG00000003184 | 54131 | 0.303105004298608 | 3.12960495919987 | 3.32967049982004 | 0.0052283201764 | 0.0145384807552028 |
| Rbm3 | ENSMUSG00000031167 | 19652 | 0.448848998770185 | 5.39351040583198 | 3.32953038874912 | 0.0052297494504 | 0.0145387566742276 |
| Zfp932 | ENSMUSG00000066613 | 69504 | 0.436630153737311 | 1.701598062485 | 3.32904361728955 | 0.0052347180586 | 0.0145488693362 |
| Actr1a | ENSMUSG00000025228 | 54130 | 0.201094843351572 | 5.48181063799243 | 3.32889187619266 | 0.0052362678897 | 0.0145494774537713 |
| Mtrex | ENSMUSG00000016018 | 72198 | -0.264652255060308 | 5.02319978789476 | -3.32861489987296 | 0.0052390980179 | 0.0145536417876601 |
| BC025920 | ENSMUSG00000074862 | 268319 | -0.383301954582191 | 2.36966146162433 | -3.32750096303221 | 0.0052504956749 | 0.0145815976176322 |
| Slc35b1 | ENSMUSG00000020873 | 110172 | 0.287050889545456 | 4.05781670207786 | 3.32666022908474 | 0.0052591144340 | 0.0146018236928207 |
| Dram2 | ENSMUSG00000027900 | 67171 | -0.231125433319288 | 3.86722967736916 | -3.32642657483791 | 0.0052615122544 | 0.0146047715725886 |
| Slc9a4 | ENSMUSG00000026065 | 110895 | -1.04997202502765 | 0.674039201272475 | -3.32559379102586 | 0.0052700674263 | 0.0146248050361141 |
| Zc3h14 | ENSMUSG00000021012 | 75553 | -0.195423904660374 | 5.28743755780775 | -3.32454368146574 | 0.0052808750874 | 0.0146510775585775 |
| Adap2 | ENSMUSG00000020709 | 216991 | 0.556871685249813 | 2.02053283903982 | 3.32426087307384 | 0.0052837895338 | 0.014655443642364 |
| Ptprg | ENSMUSG00000021745 | 19270 | -0.285891238552454 | 5.39384942269379 | -3.32297604125674 | 0.0052970505470 | 0.014684998867969 |
| Srgn | ENSMUSG00000020077 | 19073 | -0.387813796271555 | 3.1946371388576 | -3.32296815372582 | 0.0052971320589 | 0.014684998867969 |

| | | | | | | | |
|---|---|---|---|---|---|---|---|
| Rwdd1 | ENSMUSG00000019782 | 66521 | -0.310334320067257 | 5.63864224135548 | -3.32099079604523 | 0.0053176063796 | 0.0147319217163742 |
| Zc3h11a | ENSMUSG00000102976 | 70579 | 0.714691806180346 | 3.73409892432961 | 3.32086800154183 | 0.0053188804570 | 0.0147319217163742 |
| Actg1 | ENSMUSG00000062825 | 11465 | 0.286996286404698 | 7.53553570660044 | 3.32077764873318 | 0.0053198181251 | 0.0147319217163742 |
| Lsmem1 | ENSMUSG00000071342 | 380755 | -0.385399810989212 | 2.8390666589422 | -3.32071533500777 | 0.0053204649044 | 0.0147319217163742 |
| Hist1h2ac | ENSMUSG00000069270 | 319164 | 0.836056211179294 | 0.850934134272583 | 3.32068338702355 | 0.0053207965361 | 0.0147319217163742 |
| Ror1 | ENSMUSG00000035305 | 26563 | 0.572048629723254 | 1.884938425109 | 3.32045642810079 | 0.0053231530489 | 0.0147347140994014 |
| Ncbp1 | ENSMUSG00000028330 | 433702 | 0.290395652662016 | 4.14176883286506 | 3.3202776473653 | 0.0053250100655 | 0.0147361227914619 |
| Phf14 | ENSMUSG00000029629 | 75725 | -0.243442243899159 | 5.0431072113089 | -3.3201452127946 | 0.0053263860981 | 0.0147362000604339 |
| Incenp | ENSMUSG00000024660 | 16319 | 0.480789802928583 | 1.50609714665259 | 3.31981659100362 | 0.0053298021128 | 0.0147419197610818 |
| Nudt12 | ENSMUSG00000024228 | 67993 | -0.42002292765334 | 2.08519287788427 | -3.31945772425736 | 0.0053335350344 | 0.014748512891912 |
| Marc2 | ENSMUSG00000073481 | 67247 | -0.276996803710738 | 3.99187387255685 | -3.31915802519924 | 0.0053366545058 | 0.0147534067787029 |
| Mtln | ENSMUSG00000051319 | 67885 | -0.454618714855944 | 3.50717731083896 | -3.31883276372035 | 0.0053400421199 | 0.0147590392954489 |
| Lsm14a | ENSMUSG00000066568 | 67070 | -0.241899349211565 | 5.48123552068635 | -3.31861729292843 | 0.0053422874469 | 0.0147615126598442 |
| Frat2 | ENSMUSG00000047604 | 212398 | -0.921755141578654 | 0.740972743170619 | -3.31685292493621 | 0.0053607088291 | 0.0148086702580103 |
| Dtx3l | ENSMUSG00000049502 | 209200 | 0.292448458796276 | 4.54759486072047 | 3.31656601271889 | 0.0053637104364 | 0.0148132184892404 |
| Ppt2 | ENSMUSG00000015474 | 54397 | -0.328358956593539 | 3.03225502978261 | -3.31254836388858 | 0.0054059196907 | 0.0149247943868225 |
| 5430431A17Rik | ENSMUSG00000108322 | 71368 | 0.597262305839829 | 1.54443901679784 | 3.31246096233426 | 0.0054068416221 | 0.0149247943868225 |
| AW549877 | ENSMUSG00000041935 | 106064 | -0.348549088422621 | 3.7295811340953 | -3.31225329959975 | 0.0054090327284 | 0.0149270731519459 |
| Dph3 | ENSMUSG00000021905 | 105638 | -0.265163750304119 | 4.58741969303882 | -3.3117186055855 | 0.0054146785318 | 0.0149388821381158 |
| Ikzf5 | ENSMUSG00000040167 | 67143 | -0.233963726696373 | 4.07186598537245 | -3.3105241727606 | 0.0054273118419 | 0.01496995855265 |
| Mybpc1 | ENSMUSG00000020061 | 109272 | -0.347815007615914 | 11.5603181579269 | -3.30990178993381 | 0.0054339063914 | 0.014984366994097 |
| Ciao2a | ENSMUSG00000032381 | 68250 | -0.343206428539765 | 4.19710045489278 | -3.30923742823319 | 0.0054409546040 | 0.0150000188591704 |
| Lipo3 | ENSMUSG00000024766 | 381236 | -0.538387686265983 | 1.55480665076082 | -3.30830358345902 | 0.0054508772697 | 0.0150235853342737 |
| Ppp2r3d | ENSMUSG00000093803 | 19054 | 0.344901011527182 | 2.77140778499186 | 3.30802158406926 | 0.0054538772507 | 0.0150259032350801 |
| Vma21 | ENSMUSG00000073131 | 67048 | 0.37520648799577 | 4.3299247492491 | 3.30784860778112 | 0.0054557182357 | 0.0150259032350801 |
| Prkci | ENSMUSG00000037643 | 18759 | 0.357155215386105 | 2.71322511305173 | 3.30783697225311 | 0.0054558420949 | 0.0150259032350801 |
| Gem | ENSMUSG00000028214 | 14579 | 0.80708029386851 | 1.75506123113837 | 3.30759475554933 | 0.0054584211114 | 0.0150292194380367 |
| Phf7 | ENSMUSG00000021902 | 71838 | -0.465554295190465 | 1.64665029452536 | -3.30695903044384 | 0.0054651958495 | 0.0150440835637596 |
| Epha3 | ENSMUSG00000052504 | 13837 | 1.06727842147603 | 0.982786711076088 | 3.30541822709206 | 0.0054816507571 | 0.0150855801677564 |
| Atxn3 | ENSMUSG00000021189 | 110616 | 0.307535404354694 | 2.72122641368806 | 3.30443131803711 | 0.0054922164849 | 0.0151093098423951 |
| Tmem131l | ENSMUSG00000033767 | 229473 | 0.334832704050354 | 4.4369266035361 | 3.30435467197133 | 0.0054930379026 | 0.0151093098423951 |
| Creg1 | ENSMUSG00000040713 | 433375 | -0.310323177510663 | 4.6100599354301 | -3.30277633207951 | 0.0055099804269 | 0.0151520996343925 |
| Man2b2 | ENSMUSG00000029119 | 17160 | 0.591367015478874 | 2.74488982660899 | 3.30131335977021 | 0.0055257313481 | 0.0151915918928882 |
| Zfp821 | ENSMUSG00000031728 | 75871 | 0.513225431037544 | 2.19736273635833 | 3.30083368751396 | 0.0055309055067 | 0.0152008790141885 |
| Amfr | ENSMUSG00000031751 | 23802 | -0.205414095271265 | 6.14961656096649 | -3.30074241025581 | 0.0055318906521 | 0.0152008790141885 |
| Ppp4r1 | ENSMUSG00000061950 | 70351 | 0.277910550821293 | 3.88806702326195 | 3.29881762299792 | 0.0055527056940 | 0.01525424125201 |
| Map2k7 | ENSMUSG00000002948 | 26400 | -0.281094854435776 | 5.56564994321795 | -3.29852504830845 | 0.0055558765253 | 0.0152591171605653 |
| Mfsd8 | ENSMUSG00000025759 | 72175 | -0.464810414283778 | 2.02331278160602 | -3.29815926411883 | 0.0055598433320 | 0.0152661762025073 |
| Arl8b | ENSMUSG00000030105 | 67166 | -0.209198435095785 | 5.98008306822265 | -3.29768774661198 | 0.0055649609711 | 0.0152763908526199 |
| Creb3 | ENSMUSG00000028466 | 12913 | 0.317912195159238 | 3.48313698248895 | 3.29734600857471 | 0.0055686729940 | 0.0152827427813635 |
| Inpp5e | ENSMUSG00000026925 | 64436 | -0.320893120214004 | 2.90840523131923 | -3.29560525789619 | 0.0055876199227 | 0.0153297505350338 |
| Megf11 | ENSMUSG00000036466 | 214058 | -1.50222957297309 | 0.173703248901227 | -3.29551479304814 | 0.0055886063381 | 0.0153297505350338 |
| Eif4a1 | ENSMUSG00000059796 | 13681 | 0.305685950937901 | 5.92195304779009 | 3.29405909398985 | 0.0056045030958 | 0.0153694990818388 |
| Rbp7 | ENSMUSG00000028996 | 63954 | -1.71002313881436 | -0.0118769458207512 | -3.29261839923072 | 0.0056202806362 | 0.0154089008363906 |
| Ddrgk1 | ENSMUSG00000068290 | 77006 | -0.359402615819811 | 5.54391383732985 | -3.29155023985784 | 0.0056320071528 | 0.0154371790840999 |
| Gemin5 | ENSMUSG00000037275 | 216766 | 0.283890971208902 | 3.45163517899948 | 3.29104673542915 | 0.0056375432510 | 0.0154471356763373 |
| Bcam | ENSMUSG00000002980 | 57278 | 0.237350742982515 | 5.73812283826322 | 3.29096286435783 | 0.0056384659544 | 0.0154471356763373 |
| Rab5b | ENSMUSG00000000711 | 19344 | -0.23671260574041 | 5.42088981099371 | -3.29058837873648 | 0.0056425876870 | 0.0154533839871082 |
| Tatdn2 | ENSMUSG00000056952 | 381801 | -0.218807647127599 | 4.68283646890527 | -3.29049879427699 | 0.0056435741356 | 0.0154533839871082 |
| Lman2 | ENSMUSG00000021484 | 66890 | 0.257923101281258 | 5.65306505085571 | 3.28892293749637 | 0.0056609547875 | 0.0154970941104581 |
| Id1 | ENSMUSG00000042745 | 15901 | -1.223320950732 | 2.15457233523402 | -3.28860806857329 | 0.0056644340084 | 0.0155027361408579 |
| Gdpd1 | ENSMUSG00000061666 | 66569 | -0.50077048863848 | 1.42022048636733 | -3.28809114659973 | 0.0056701505121 | 0.0155144969332272 |
| Abr | ENSMUSG00000017631 | 109934 | 0.367900872138489 | 4.79144517545526 | 3.28695320473194 | 0.0056827550950 | 0.0155450940801195 |
| Tcof1 | ENSMUSG00000024613 | 21453 | -0.185530927148923 | 4.68546518833598 | -3.284042689498 | 0.0057151217721 | 0.0156297213138929 |
| Zfp661 | ENSMUSG00000034800 | 72180 | -0.563465314010255 | 1.15194799080533 | -3.28294276305005 | 0.0057274016622 | 0.0156593865857725 |
| Pmpca | ENSMUSG00000026926 | 66865 | -0.222570901032209 | 6.09099322312524 | -3.28228457680084 | 0.0057347624815 | 0.0156755910692267 |
| Gm20498 | ENSMUSG00000021139 | 105940408 | -0.422649857309889 | 2.62567482226055 | -3.28123190163158 | 0.0057465547745 | 0.0157019260130543 |
| Ecscr | ENSMUSG00000073599 | 68545 | -0.409449171703671 | 2.30239102795988 | -3.28116814399851 | 0.0057472697811 | 0.0157019260130543 |
| Cdkal1 | ENSMUSG00000006191 | 68916 | -0.343696614617895 | 3.59587025676653 | -3.28065821190761 | 0.0057529915964 | 0.015713630969621 |
| Fam107a | ENSMUSG00000021750 | 268709 | -1.15450733472801 | 0.71721814258045 | -3.28007086520891 | 0.0057595891349 | 0.0157277214175759 |
| Kdm1b | ENSMUSG00000038080 | 218214 | -0.316751605213245 | 3.74040886751109 | -3.27818740765753 | 0.0057807967854 | 0.01578169072464 |
| Csde1 | ENSMUSG00000068823 | 229663 | -0.263854700553651 | 9.65622869770288 | -3.27758366736192 | 0.0057876114216 | 0.0157963496754791 |
| Crebrf | ENSMUSG00000048249 | 77128 | -0.303488311873917 | 4.73491999907907 | -3.27675773051999 | 0.0057969471105 | 0.0158178803956326 |
| Kmt2d | ENSMUSG00000048154 | 381022 | 0.259513987179235 | 7.38563042340432 | 3.27602188448639 | 0.0058052771974 | 0.015836657111407 |
| Chm | ENSMUSG00000025531 | 12662 | -0.306523519672737 | 3.79381611761008 | -3.27535048233413 | 0.0058128882153 | 0.0158502982315094 |
| Bcl2l13 | ENSMUSG00000009112 | 94044 | -0.234119109914512 | 6.10329831008039 | -3.27532496159773 | 0.0058131777156 | 0.0158502982315094 |
| Ccdc50 | ENSMUSG00000038127 | 67501 | -0.37127450114638 | 5.93933086644337 | -3.27452344822874 | 0.0058222772206 | 0.0158711501993646 |
| Xylt1 | ENSMUSG00000030657 | 233781 | 0.633765148100666 | 2.40088784228484 | 3.27326525546483 | 0.0058365901535 | 0.0159061996928716 |
| Zbtb40 | ENSMUSG00000060862 | 230848 | -0.308841525456033 | 3.08323729611102 | -3.27262573582432 | 0.0058438786986 | 0.0159220932340459 |
| Crip1 | ENSMUSG00000006360 | 12925 | 0.485897425632047 | 4.18087968020184 | 3.27003972330412 | 0.0058734443335 | 0.0159971806477664 |
| Csnk2a1 | ENSMUSG00000074698 | 12995 | -0.221877658359679 | 6.2646927163573 | -3.26995941056571 | 0.0058743649364 | 0.0159971806477664 |
| Abhd14a | ENSMUSG00000042210 | 68644 | -0.630960243652431 | 1.65965565908196 | -3.26412042134228 | 0.0059416843376 | 0.0161764760883728 |
| Vmn2r90 | ENSMUSG00000092120 | 626942 | -1.52558401672552 | 3.03523476720024 | -3.26316007449707 | 0.0059528302080 | 0.0162027856086414 |
| Pi4k2a | ENSMUSG00000025178 | 84095 | 0.23393083328761 | 3.9869256569229 | 3.26235153823975 | 0.0059622303842 | 0.0162243316729398 |
| Arl8a | ENSMUSG00000026426 | 68724 | 0.390250560905961 | 3.61339006567499 | 3.26203567831278 | 0.0059659066603 | 0.016230295098084 |
| Pepd | ENSMUSG00000063931 | 18624 | 0.219665615513811 | 3.94652233899386 | 3.26176766251846 | 0.0059690278613 | 0.0162347458452662 |
| Nucb1 | ENSMUSG00000030824 | 18220 | 0.229561392799684 | 5.55179516126486 | 3.26020808638444 | 0.0059872225065 | 0.0162801813976931 |
| Pigb | ENSMUSG00000079469 | 55981 | 0.498615455483153 | 1.46044496014202 | 3.25990365666775 | 0.0059907805792 | 0.0162858051509341 |
| Rfk | ENSMUSG00000024712 | 54391 | -0.272303148600487 | 5.11578901699689 | -3.25916597010743 | 0.0059994111965 | 0.0162992051415476 |
| Tuba4a | ENSMUSG00000026202 | 22145 | -0.402605496730946 | 8.18806071109319 | -3.25910084943291 | 0.0060001736782 | 0.0162992051415476 |
| Scarb1 | ENSMUSG00000037936 | 20778 | 0.410815708914333 | 3.51979329396328 | 3.25910004462852 | 0.0060001831021 | 0.0162992051415476 |
| Tatdn1 | ENSMUSG00000050891 | 69694 | -0.308285412129226 | 2.7654005122473 | -3.25876050991641 | 0.0060041602079 | 0.0163059565796091 |
| Mef2d | ENSMUSG00000001419 | 17261 | -0.196045336642113 | 7.29891002668442 | -3.25815004661005 | 0.0060113174485 | 0.0163213390536604 |
| Dll4 | ENSMUSG00000027314 | 54485 | -0.493137427794159 | 2.5782186392634 | -3.25796534215144 | 0.0060134846574 | 0.0163231688082965 |
| Ptbp3 | ENSMUSG00000028382 | 230257 | 0.421547526320212 | 4.28308051263901 | 3.25779709886449 | 0.0060154594019 | 0.01632447535312 |
| Lactb | ENSMUSG00000032370 | 80907 | -0.311412468381576 | 3.06736872461639 | -3.25583667734165 | 0.0060385176335 | 0.0163829824405252 |
| Pofut2 | ENSMUSG00000020260 | 80294 | 0.291716576737415 | 4.11046405520359 | 3.25497970372264 | 0.0060486250243 | 0.0164063325413337 |
| Dmap1 | ENSMUSG00000009640 | 66233 | -0.374567545089898 | 2.89232679000387 | -3.25413244856294 | 0.0060586344421 | 0.0164294053800246 |
| Dipk2b | ENSMUSG00000037358 | 75905 | -0.55807168301041 | 1.65428186104635 | -3.25258222917143 | 0.0060769915779 | 0.0164750979509426 |
| Rapgef6 | ENSMUSG00000037533 | 192786 | 0.238980062678655 | 3.70639108224315 | 3.2523291413807 | 0.0060799938341 | 0.016479150161446 |
| Epha4 | ENSMUSG00000026235 | 13838 | -0.502253024206571 | 2.13755686609719 | -3.25115248448991 | 0.0060939714459 | 0.016512940474788 |
| Fam149b | ENSMUSG00000039599 | 105428 | -0.289708406849701 | 3.42613337125095 | -3.25091476743067 | 0.0060967992110 | 0.0165165085937959 |
| Art1 | ENSMUSG00000030996 | 11870 | -0.30869573638306 | 6.35342753471232 | -3.25000641095773 | 0.0061076166755 | 0.0165417140438423 |
| Arhgap1 | ENSMUSG00000027247 | 228359 | 0.36118585151992 | 3.02528461277365 | 3.24952584238054 | 0.0061133474540 | 0.0165531337676682 |
| Plce1 | ENSMUSG00000024998 | 74055 | 0.333039201665981 | 4.16590017396102 | 3.24895517314844 | 0.0061201596729 | 0.0165674753304965 |
| Irf2bpl | ENSMUSG00000034168 | 238330 | 0.451763650857544 | 5.06623235290709 | 3.2476973533508 | 0.0061352014163 | 0.0166040818721788 |
| Plpp5 | ENSMUSG00000031570 | 71910 | 0.616032200093921 | 1.40039947706291 | 3.246962556013 | 0.0061440056663 | 0.0166237935491443 |
| Macrod1 | ENSMUSG00000036278 | 107227 | -0.284098286941377 | 6.50885640543324 | -3.24648483851426 | 0.0061497363991 | 0.0166351815339826 |
| Ankfy1 | ENSMUSG00000020790 | 11736 | 0.324833660514178 | 4.62070830898162 | 3.24504928997086 | 0.0061669895503 | 0.0166777245854876 |
| Slc1a4 | ENSMUSG00000020142 | 55963 | 0.807465401135638 | 1.02668847374169 | 3.24474251039534 | 0.0061706828601 | 0.0166835850469184 |
| Smarcd3 | ENSMUSG00000028949 | 66993 | -0.337052699210245 | 6.05772644497043 | -3.24428988161938 | 0.0061761360864 | 0.0166941996935966 |
| Kxd1 | ENSMUSG00000055553 | 75620 | -0.283098642421983 | 4.90963076100368 | -3.24352005361072 | 0.0061854219755 | 0.0167151662829553 |
| Fbrsl1 | ENSMUSG00000043323 | 381668 | -0.242417727426935 | 4.88449281751281 | -3.24292707740179 | 0.0061925841507 | 0.016730384911478 |
| Rap1gds1 | ENSMUSG00000028149 | 229877 | -0.22391880844864 | 5.2992001012951 | -3.24083547667257 | 0.0062179135619 | 0.0167946659614969 |
| Rnu1a1 | ENSMUSG00000095969 | 19842 | 1.12332087245636 | 5.92004888930825 | 3.2401384582999 | 0.0062263775043 | 0.016810071717594 |
| Pibf1 | ENSMUSG00000022064 | 52023 | -0.289589953375158 | 3.96773728529216 | -3.24011250110082 | 0.0062266929269 | 0.016810071717594 |
| Usp46 | ENSMUSG00000054814 | 69727 | -0.402714256125315 | 2.98417997697732 | -3.23926744406773 | 0.0062369704949 | 0.0168336603655356 |
| Akap8 | ENSMUSG00000024045 | 56399 | -0.286160255346569 | 4.57061662476046 | -3.23828401507267 | 0.0062489523101 | 0.0168618360164594 |
| Ube2e3 | ENSMUSG00000027011 | 22193 | -0.404734856020216 | 3.09151220097961 | -3.23762106228393 | 0.0062570425276 | 0.0168794994743218 |
| Spag9 | ENSMUSG00000020859 | 70834 | -0.210559070113594 | 7.88888161627317 | -3.23744813745094 | 0.0062591545052 | 0.0168810308160301 |
| Commd7 | ENSMUSG00000056941 | 99311 | 0.433026071524283 | 2.60056870426841 | 3.23697428505995 | 0.0062649454476 | 0.0168924811762284 |
| Hoxc10 | ENSMUSG00000022484 | 209448 | -0.223561220698233 | 5.23786288177637 | -3.23659594022285 | 0.0062695730418 | 0.0169007898693035 |
| Armt1 | ENSMUSG00000061759 | 73419 | -0.453394256885895 | 3.13778317579489 | -3.23512218472039 | 0.0062876313976 | 0.016945290633131 |
| Zfp414 | ENSMUSG00000073423 | 328801 | 0.446972962545608 | 1.40568544629085 | 3.23408154969518 | 0.0063004139341 | 0.0169755545264472 |
| Gtpbp10 | ENSMUSG00000040464 | 207704 | -0.393513097955234 | 2.658797174742 | -3.23375389410327 | 0.0063044440371 | 0.0169787745877203 |
| Cyp2j6 | ENSMUSG00000052914 | 13110 | 0.566441906672942 | 1.94997245428178 | 3.23373182430132 | 0.0063047155842 | 0.0169787745877203 |

| | | | | | | | |
|---|---|---|---|---|---|---|---|
| Washc2 | ENSMUSG00000024104 | 28006 | 0.198529273845247 | 4.9629748925826 | 3.23337342799153 | 0.0063091269353 | 0.0169864695886378 |
| Nap1l1 | ENSMUSG00000058799 | 53605 | 0.210266103642938 | 5.93955637217132 | 3.23295406349036 | 0.0063142926384 | 0.0169961912905591 |
| Prmt1 | ENSMUSG00000109324 | 15469 | 0.284484710740223 | 4.90057206299196 | 3.23255605395312 | 0.0063191992054 | 0.0170052108602977 |
| Mrpl27 | ENSMUSG00000024414 | 94064 | -0.376889550741634 | 3.46457799564771 | -3.23220909181263 | 0.0063234795833 | 0.0170125413058527 |
| Ubiad1 | ENSMUSG00000047719 | 71707 | -0.556962612520534 | 3.26018793847398 | -3.23134988684236 | 0.0063340918489 | 0.0170368991143439 |
| Spns2 | ENSMUSG00000040447 | 216892 | -0.425119377101132 | 2.79467111964298 | -3.23119332222736 | 0.0063360275373 | 0.0170379131636704 |
| Tpcn1 | ENSMUSG00000032741 | 252972 | -0.253879639536416 | 5.6205135013232 | -3.23020678079632 | 0.0063482382701 | 0.0170665500568468 |
| Tnfrsf11b | ENSMUSG00000063727 | 18383 | 0.968571080548771 | 1.29374579679858 | 3.229474659312 | 0.0063573151788 | 0.017081482410779 |
| Usp53 | ENSMUSG00000039701 | 99526 | -0.31965958821325 | 4.05841597902559 | -3.22941349096594 | 0.0063580741367 | 0.017081482410779 |
| Gcnt2 | ENSMUSG00000021360 | 14538 | -0.445535737931508 | 3.07939713571407 | -3.22938073137956 | 0.0063584806449 | 0.017081482410779 |
| Slc25a17 | ENSMUSG00000022404 | 20524 | -0.37806248973513 | 2.3029540061618 | -3.22781617604373 | 0.0063779252387 | 0.0171295087923561 |
| Ctsa | ENSMUSG00000017760 | 19025 | 0.228352472546242 | 5.01490352761369 | 3.22746804566219 | 0.0063822599515 | 0.0171357089221664 |
| Synm | ENSMUSG00000030554 | 233335 | -0.321301716373059 | 8.14400648740218 | -3.22737901210135 | 0.0063833690175 | 0.0171357089221664 |
| Zfp385a | ENSMUSG00000000552 | 29813 | -0.490249241587487 | 4.45589253409417 | -3.22681419075389 | 0.0063904093292 | 0.0171478589606645 |
| Med9 | ENSMUSG00000061650 | 192191 | 0.328559476834018 | 2.96570252460446 | 3.22676421734329 | 0.0063910326050 | 0.0171478589606645 |
| Ube2h | ENSMUSG00000039159 | 22214 | -0.197455192499597 | 6.21533394240484 | -3.22550076973972 | 0.0064068107247 | 0.0171812721262707 |
| Bccip | ENSMUSG00000030983 | 66165 | -0.265372386381354 | 4.79874815308462 | -3.22542406002204 | 0.0064077699402 | 0.0171812721262707 |
| Baz1b | ENSMUSG00000002748 | 22385 | -0.196762484402184 | 6.35509413703642 | -3.2253895816385 | 0.0064082011214 | 0.0171812721262707 |
| Cacng4 | ENSMUSG00000020723 | 54377 | -1.21136608361302 | 0.366681488341026 | -3.22466568365078 | 0.0064172607774 | 0.0171984972094607 |
| Mis12 | ENSMUSG00000040599 | 67139 | 0.346198403232214 | 2.57090197368754 | 3.22462483363051 | 0.0064177724011 | 0.0171984972094607 |
| Mob2 | ENSMUSG00000025147 | 101513 | -0.240964501630121 | 4.01463355434958 | -3.22433007448371 | 0.0064214653040 | 0.0172041757936341 |
| Eri3 | ENSMUSG00000033423 | 140546 | -0.205632047153637 | 4.99167945746557 | -3.22321343898133 | 0.0064354744016 | 0.0172374836276639 |
| Cep128 | ENSMUSG00000061533 | 75216 | 0.588791775111725 | 1.67834814843853 | 3.22206711419086 | 0.0064498877636 | 0.0172718577031452 |
| Rsu1 | ENSMUSG00000026727 | 20163 | 0.366091710135033 | 3.5932934616755 | 3.22136150859397 | 0.0064587757677 | 0.0172907253945805 |
| AU040320 | ENSMUSG00000028830 | 100317 | -0.224309115728102 | 4.42764077392953 | -3.22125668480222 | 0.0064600972016 | 0.0172907253945805 |
| Add3 | ENSMUSG00000025026 | 27360 | 0.243582605291786 | 5.28141486261256 | 3.22030976007017 | 0.0064720466184 | 0.0173184679526086 |
| Coa5 | ENSMUSG00000026112 | 76178 | -0.231462058823722 | 5.82529506383768 | -3.2201553337693 | 0.0064739974469 | 0.0173194483828093 |
| Gpat3 | ENSMUSG00000029314 | 231510 | -0.599878249178597 | 1.71658305654506 | -3.2192616840266 | 0.0064852982481 | 0.0173409284565313 |
| Trap1 | ENSMUSG00000005981 | 68015 | -0.24660185183243 | 5.97949702052497 | -3.21917354104482 | 0.0064864139433 | 0.0173409284565313 |
| Hnrnpd | ENSMUSG00000000568 | 11991 | -0.243446514759022 | 4.92313202907551 | -3.21914416036025 | 0.0064867858804 | 0.0173409284565313 |
| Aen | ENSMUSG00000030609 | 68048 | -0.217900272323795 | 4.42188954988698 | -3.21800543280799 | 0.0065012177427 | 0.0173748643160854 |
| Emc1 | ENSMUSG00000078517 | 230866 | 0.230324985168813 | 5.02911939438077 | 3.21789181943314 | 0.0065026594014 | 0.0173748643160854 |
| Ptprm | ENSMUSG00000033278 | 19274 | -0.208101377761442 | 5.04804113567206 | -3.21760457590273 | 0.0065063057071 | 0.0173803586718839 |
| Ap3m1 | ENSMUSG00000021824 | 55946 | -0.231521266981673 | 4.18330105758077 | -3.21583604660737 | 0.0065288007160 | 0.0174361887678389 |
| Sergef | ENSMUSG00000030839 | 27414 | 0.726299796753282 | 1.20856252967788 | 3.21557440855224 | 0.0065321352458 | 0.0174400526995766 |
| Slc26a10 | ENSMUSG00000040441 | 216441 | -0.61465164251901 | 2.05420822662017 | -3.21547219042664 | 0.0065334384598 | 0.0174400526995766 |
| Eml3 | ENSMUSG00000071647 | 225898 | 0.608516542889328 | 1.57941137367171 | 3.21529973747698 | 0.0065356377108 | 0.0174416640179549 |
| Pcbp3 | ENSMUSG00000001120 | 59093 | 0.655997876778694 | 1.28405059624505 | 3.214726847124 | 0.0065429489573 | 0.0174569136080767 |
| Hdac5 | ENSMUSG00000008855 | 15184 | -0.304970845936164 | 5.64455154016238 | -3.21211332012806 | 0.0065764067298 | 0.0175380792822806 |
| Polr1d | ENSMUSG00000029642 | 20018 | 0.174216646036814 | 4.79831412787944 | 3.21209988405265 | 0.0065765791764 | 0.0175380792822806 |
| Gpc4 | ENSMUSG00000031119 | 14735 | 0.316213263275751 | 5.01299171207544 | 3.21179271581746 | 0.0065805227897 | 0.0175443157596393 |
| Ggct | ENSMUSG00000002797 | 110175 | -0.527495513894989 | 1.75057338574961 | -3.21133483226776 | 0.0065864057658 | 0.0175557184654282 |
| Smarcb1 | ENSMUSG00000000902 | 20587 | 0.243311942583134 | 3.77908760817011 | 3.20971358141842 | 0.0066072781942 | 0.0176070594688538 |
| Fbxo6 | ENSMUSG00000055401 | 50762 | -0.330608540775222 | 3.41483521532083 | -3.20860506918136 | 0.0066215875213 | 0.0176386884145098 |
| Mhrt | ENSMUSG00000097652 | 791403 | 0.566399097539484 | 0.967400779810571 | 3.20854416185332 | 0.0066223746457 | 0.0176386884145098 |
| Acap2 | ENSMUSG00000049076 | 78618 | 0.348014722288062 | 4.27077643304278 | 3.20811535788876 | 0.0066279188606 | 0.0176491549489124 |
| Timm8a1 | ENSMUSG00000048007 | 30058 | -0.446062214862031 | 3.0868332977526 | -3.20797422204402 | 0.0066297446888 | 0.0176497172903115 |
| Chst3 | ENSMUSG00000057337 | 53374 | 0.633465390696759 | 1.06706408505283 | 3.2078285357233 | 0.0066316299123 | 0.0176504374413331 |
| Ppp2r5a | ENSMUSG00000026626 | 226849 | -0.198439384204071 | 6.34542633005497 | -3.20767496026233 | 0.0066336178035 | 0.0176514304309419 |
| Adcy3 | ENSMUSG00000020654 | 104111 | 0.666044317469104 | 1.57489723527888 | 3.20697138514403 | 0.0066427325429 | 0.0176713821919626 |
| Kif26a | ENSMUSG00000021294 | 668303 | -0.475681859695616 | 3.32024155924504 | -3.20654918182526 | 0.0066482081479 | 0.0176787423728602 |
| Arhgap39 | ENSMUSG00000033697 | 223666 | 0.388849681316516 | 2.35231654862476 | 3.20650866393039 | 0.0066487338665 | 0.0176787423728602 |
| Pycr2 | ENSMUSG00000026520 | 69051 | 0.45654563068052 | 2.56967948339644 | 3.20603938313848 | 0.0066548257986 | 0.0176906373554266 |
| Dleu2 | ENSMUSG00000097589 | 668253 | 0.455689250233986 | 3.44328581351836 | 3.20576766010353 | 0.0066583556994 | 0.0176957175177677 |
| Farsb | ENSMUSG00000026245 | 23874 | -0.204447426161292 | 4.32167656387344 | -3.20337894039962 | 0.0066894677108 | 0.0177740815623708 |
| Rpl36 | ENSMUSG00000057863 | 54217 | -0.306341914705031 | 6.44253751589032 | -3.202546110742 | 0.0067003490594 | 0.0177963337239776 |
| Ltbr | ENSMUSG00000030339 | 17000 | 0.263883806271747 | 4.06843636390796 | 3.20248543264789 | 0.0067011425408 | 0.0177963337239776 |
| Mrpl57 | ENSMUSG00000021967 | 67840 | -0.23652788523467 | 3.90374730595183 | -3.20236430962655 | 0.0067027267351 | 0.0177963337239776 |
| Rfc1 | ENSMUSG00000029191 | 19687 | -0.256410546800367 | 4.75348371071449 | -3.20125581967497 | 0.0067172423076 | 0.0178305429006337 |
| Mpp6 | ENSMUSG00000038388 | 56524 | 0.253506810222435 | 6.13759778873283 | 3.2011042003423 | 0.0067192301886 | 0.0178314894857667 |
| Cep63 | ENSMUSG00000032534 | 28135 | -0.267897085746486 | 3.83162543630676 | -3.20082703958154 | 0.0067228655623 | 0.0178368066656713 |
| Caprin1 | ENSMUSG00000027184 | 53872 | 0.175738345384266 | 6.26342719159956 | 3.20006896407827 | 0.0067328188849 | 0.0178588796969417 |
| Slc25a15 | ENSMUSG00000031482 | 18408 | -0.32696492946487 | 3.17291560217916 | -3.19869762110134 | 0.0067508616554 | 0.0179003138039799 |
| Gga1 | ENSMUSG00000033128 | 106039 | -0.195123243210669 | 4.78830625767974 | -3.19863287304377 | 0.0067517147391 | 0.0179003138039799 |
| Exoc1 | ENSMUSG00000036435 | 69940 | 0.283651285141878 | 3.63150980563458 | 3.19849825001811 | 0.0067534888006 | 0.0179006755770573 |
| Tbrg1 | ENSMUSG00000011114 | 21376 | 0.233186719627159 | 3.7985549279894 | 3.19678871113061 | 0.0067760576344 | 0.0179561420610563 |
| Ckb | ENSMUSG00000001270 | 12709 | 0.491375518424746 | 3.20997344429101 | 3.19628339765563 | 0.0067827430487 | 0.0179695017608848 |
| Cldn1 | ENSMUSG00000022512 | 12737 | -0.612065567171223 | 1.19505953892806 | -3.19499585725675 | 0.0067997578032 | 0.01801021384709 |
| Atxn1 | ENSMUSG00000046876 | 20238 | -0.213843878224012 | 6.17982743708186 | -3.19450975711314 | 0.0068062609100 | 0.0180230712227981 |
| Nfatc4 | ENSMUSG00000023411 | 73181 | 0.586527867422268 | 2.08778570608381 | 3.19394502944863 | 0.0068137660840 | 0.018036157692721 |
| Ssr2 | ENSMUSG00000041355 | 66256 | 0.299372485724205 | 3.30553082252144 | 3.19388962065621 | 0.0068145029064 | 0.018036157692721 |
| Tpp2 | ENSMUSG00000041763 | 22019 | -0.175278625991809 | 5.66857690603947 | -3.19343234331134 | 0.0068205867889 | 0.0180478901452506 |
| 9930111J21Rik2 | ENSMUSG00000069892 | 245240 | -0.681923526834643 | 4.84153037865519 | -3.19218793196877 | 0.0068371706383 | 0.0180873940579572 |
| Gna12 | ENSMUSG00000000149 | 14673 | 0.181975878101803 | 5.04710092210365 | 3.19203628398165 | 0.0068391943504 | 0.018088370056767 |
| Dok4 | ENSMUSG00000040631 | 114255 | -0.748497773046752 | 2.125532090213 | -3.19153204908154 | 0.0068459275665 | 0.0181017983140906 |
| Pcdhga11 | ENSMUSG00000102742 | 93723 | 0.794520613067548 | 1.35076058619019 | 3.19131810824278 | 0.0068487863894 | 0.0181049779984131 |
| Rhog | ENSMUSG00000073982 | 56212 | 0.34527413625559 | 3.06828000005002 | 3.19000542497683 | 0.0068663534716 | 0.0181470284813838 |
| Rnft1 | ENSMUSG00000020521 | 76892 | 0.268637054808406 | 4.12713461984565 | 3.18883428852767 | 0.0068820642674 | 0.0181841538571599 |
| Glrx3 | ENSMUSG00000031068 | 30926 | -0.324701463425065 | 5.83153311436206 | -3.18815201024409 | 0.0068912335720 | 0.0182039811929611 |
| Ccl11 | ENSMUSG00000020676 | 20292 | -0.491518422023892 | 2.79027755479713 | -3.18720571021146 | 0.0069039713247 | 0.0182332231337502 |
| Lyar | ENSMUSG00000067367 | 17089 | -0.422886545664412 | 3.71913182687038 | -3.18590352063308 | 0.0069215379990 | 0.0182752009341007 |
| Pias1 | ENSMUSG00000032405 | 56469 | -0.219526830237912 | 3.76541813766886 | -3.18371210870202 | 0.0069512011714 | 0.0183490895930479 |
| Crnkl1 | ENSMUSG00000001767 | 66877 | 0.19006489857421 | 4.40264314489115 | 3.18310600732167 | 0.0069594278022 | 0.0183663701849868 |
| Nup50 | ENSMUSG00000016619 | 18141 | -0.275928676642756 | 3.84436366412462 | -3.18293527566195 | 0.0069617469017 | 0.0183680558492501 |
| Lysmd4 | ENSMUSG00000043831 | 75099 | -0.333453105587708 | 2.3263157405041 | -3.18198635844415 | 0.0069746503963 | 0.0183976601067598 |
| Coro1c | ENSMUSG00000004530 | 23790 | 0.397103997844988 | 4.07396320019687 | 3.18172655217181 | 0.0069781874356 | 0.0184025493025482 |
| Zranb2 | ENSMUSG00000028180 | 53861 | -0.231199339449458 | 4.85172711376944 | -3.18118464553561 | 0.0069855707924 | 0.0184175770218757 |
| Map4k5 | ENSMUSG00000034761 | 399510 | -0.228013994382517 | 4.84409841466352 | -3.18063742891207 | 0.0069930344096 | 0.0184312270758752 |
| Foxo4 | ENSMUSG00000042903 | 54601 | -0.40287159257382 | 5.230064241916 | -3.18055785620226 | 0.0069941203833 | 0.0184312270758752 |
| Pcdhga6 | ENSMUSG00000103793 | 93714 | 0.598016098662909 | 1.14814644744636 | 3.18027284951198 | 0.0069980114121 | 0.0184347246737673 |
| Pbdc1 | ENSMUSG00000031226 | 67683 | 0.306838250591386 | 2.66093765084249 | 3.18021360259067 | 0.0069988205467 | 0.0184347246737673 |
| Zfp433 | ENSMUSG00000096795 | 73610 | -0.515617616612233 | 1.59347276625659 | -3.17991347546506 | 0.0070029208147 | 0.0184410810468988 |
| Wdr73 | ENSMUSG00000025722 | 71968 | 0.535538033120377 | 1.25904233431618 | 3.17907340934341 | 0.0070144103579 | 0.0184668881556895 |
| Ccndbp1 | ENSMUSG00000023572 | 17151 | 0.286190308328542 | 3.23734749238307 | 3.17812080081434 | 0.0070274619044 | 0.0184959167495724 |
| Polr3c | ENSMUSG00000028099 | 74414 | -0.263817894315243 | 3.45532750104334 | -3.17802173227648 | 0.0070288206182 | 0.0184959167495724 |
| Prrc2a | ENSMUSG00000024393 | 53761 | 0.16515308290676 | 7.47767635322138 | 3.17747211320668 | 0.0070363633453 | 0.0185113087190457 |
| Laptm4b | ENSMUSG00000022257 | 114128 | -0.434351170317016 | 2.17006304578281 | -3.17631373360041 | 0.0070522868958 | 0.0185487362989309 |
| H2afv | ENSMUSG00000041126 | 77605 | -0.319148872171642 | 3.92692432780841 | -3.17555208718337 | 0.0070627763848 | 0.0185718567867226 |
| Cd83 | ENSMUSG00000015396 | 12522 | 1.02114000924253 | 0.686202583986673 | 3.1752346087496 | 0.0070671533336 | 0.0185788968470003 |
| Yipf7 | ENSMUSG00000029158 | 75581 | -0.30376757272226 | 5.14663966103035 | -3.1740898485685 | 0.0070829581760 | 0.0186159691807253 |
| Gpank1 | ENSMUSG00000092417 | 81845 | -0.417348164222868 | 2.16310283298928 | -3.1732245238438 | 0.0070949284669 | 0.0186429478538635 |
| Slc4a4 | ENSMUSG00000060961 | 54403 | -0.531733588734217 | 3.28909937809083 | -3.17250829248952 | 0.0071048515732 | 0.0186618117687121 |
| Gm5532 | ENSMUSG00000073535 | 433367 | -0.681843991103386 | 3.49658557061029 | -3.17245994098038 | 0.0071055219633 | 0.0186618117687121 |
| Adamts8 | ENSMUSG00000031994 | 30806 | -0.933896863417245 | 3.1147893030226 | -3.17081318165856 | 0.0071283918806 | 0.0187173796895426 |
| Shisa5 | ENSMUSG00000025647 | 66940 | 0.431681681989939 | 3.46735128686979 | 3.17054633221869 | 0.0071321047498 | 0.0187201893505124 |
| Rap1gap | ENSMUSG00000041351 | 110351 | -0.711006120321115 | 2.38043670460131 | -3.1704901228418 | 0.0071328870775 | 0.0187201893505124 |
| Ppil4 | ENSMUSG00000015757 | 67418 | -0.260900802357196 | 3.88522726163023 | -3.16950695836342 | 0.0071465847130 | 0.0187516364614451 |
| Cbx5 | ENSMUSG00000009575 | 12419 | 0.182577730043488 | 5.36366873750758 | 3.1688504263477 | 0.0071557462536 | 0.0187711692580273 |
| Ifi211 | ENSMUSG00000026536 | 381308 | 0.801067083325348 | 2.29203546967782 | 3.16868105525273 | 0.0071581116341 | 0.0187728690674289 |
| Tsku | ENSMUSG00000049580 | 244152 | 1.02804279530122 | 0.811217190114614 | 3.16837563484306 | 0.0071623790061 | 0.0187795550291394 |
| Tomm5 | ENSMUSG00000078713 | 68512 | -0.336872726612276 | 3.92337387588496 | -3.16785883650689 | 0.0071696055585 | 0.0187939948106579 |
| Zfp65 | ENSMUSG00000071281 | 235907 | -0.448577414234639 | 2.27524123182636 | -3.16614670232355 | 0.0071935988735 | 0.0188523685656132 |
| Fpgs | ENSMUSG00000009566 | 14287 | 0.501758730148734 | 1.5920588930573 | 3.16546070030694 | 0.0072032347537 | 0.0188730966188099 |
| Det1 | ENSMUSG00000030610 | 76375 | -0.36385892451509 | 2.68248110432161 | -3.16464025045942 | 0.0072147760559 | 0.0188988059111088 |
| Tmem39a | ENSMUSG00000002845 | 67846 | 0.60142433763771 | 1.72980486250201 | 3.1645074374108 | 0.0072166460765 | 0.0188991754342009 |
| Ndufa1 | ENSMUSG00000016427 | 54405 | -0.253708262086301 | 4.41278031297408 | -3.16418517114862 | 0.0072211856269 | 0.0189065341527789 |

| | | | | | | | |
|---|---|---|---|---|---|---|---|
| Mrpl11 | ENSMUSG00000024902 | 66419 | -0.355289764361259 | 3.71968822947648 | -3.16258902764366 | 0.0072437114634 | 0.0189609698293116 |
| Stk39 | ENSMUSG00000027030 | 53416 | 0.399507840676112 | 2.76864139396763 | 3.16240716324726 | 0.0072462824983 | 0.0189631587238023 |
| Osgep | ENSMUSG00000006289 | 66246 | -0.268068428025973 | 3.97871420634101 | -3.16049766714582 | 0.0072733322417 | 0.0190293907932715 |
| Flnc | ENSMUSG00000068699 | 68794 | 0.327528798874088 | 10.8270427932639 | 3.15810680952707 | 0.0073073429367 | 0.0191137989093137 |
| Parl | ENSMUSG00000033918 | 381038 | -0.27979053277376 | 3.66733630712393 | -3.15781457983418 | 0.0073115108580 | 0.0191201256434935 |
| Tmed4 | ENSMUSG00000004394 | 103694 | -0.251310601809945 | 4.81518840155465 | -3.15758160682928 | 0.0073148353301 | 0.0191242441984756 |
| Zfp652 | ENSMUSG00000075595 | 268469 | -0.241779219274992 | 6.82721976074245 | -3.15668118832769 | 0.0073276982759 | 0.0191532926480535 |
| Cfd | ENSMUSG00000061780 | 11537 | 1.26283790352739 | 3.1818984117897 | 3.15633721892409 | 0.0073326180093 | 0.0191615700359883 |
| Gmppb | ENSMUSG00000070284 | 331026 | 0.532761768292034 | 1.7922082745644 | 3.1560417516557 | 0.0073368466551 | 0.0191680379510492 |
| Tmem189 | ENSMUSG00000090213 | 407243 | 0.346163953337124 | 3.1599399100174 | 3.15584482673384 | 0.0073396663404 | 0.0191708226445412 |
| B230354K17Rik | ENSMUSG00000097119 | 105246278 | 0.217132185298286 | 3.58774370418776 | 3.15544469312399 | 0.0073453990146 | 0.0191812127636708 |
| Map3k5 | ENSMUSG00000071369 | 26408 | -0.329843756190917 | 3.77079686247986 | -3.15510200129238 | 0.0073503122753 | 0.0191894586773566 |
| Pum2 | ENSMUSG00000020594 | 80913 | -0.185028630561572 | 6.47415619015827 | -3.15470355885383 | 0.0073560289670 | 0.019196438845166 |
| Marveld1 | ENSMUSG00000044345 | 277010 | 0.436878894832922 | 3.52913576634411 | 3.15467086475709 | 0.0073564982455 | 0.019196438845166 |
| Oxr1 | ENSMUSG00000022307 | 170719 | -0.188561139323633 | 5.030391834806 | -3.15335267509051 | 0.0073754439189 | 0.0192412834075783 |
| Tra2a | ENSMUSG00000029817 | 101214 | -0.292224257531752 | 3.9495383995778 | -3.15179056415962 | 0.0073979583429 | 0.0192927044442264 |
| Klhdc10 | ENSMUSG00000029775 | 76788 | -0.246983715011648 | 4.93090637352828 | -3.15174028090988 | 0.0073986842036 | 0.0192927044442264 |
| Adcy9 | ENSMUSG00000005580 | 11515 | -0.332395932267817 | 5.15760409584584 | -3.15118983101015 | 0.0074066348329 | 0.019308830278727 |
| Rffl | ENSMUSG00000020696 | 67338 | -0.365094456798759 | 2.02373858435843 | -3.14868925109319 | 0.0074428602663 | 0.0193986422441148 |
| Resf1 | ENSMUSG00000032712 | 67246 | 0.572078838238911 | 4.90030157979029 | 3.1485180925776 | 0.0074453462565 | 0.0194004958117739 |
| Helz | ENSMUSG00000020721 | 78455 | -0.196635822640459 | 5.26584310033349 | -3.14801608505386 | 0.0074526424350 | 0.0194148795180124 |
| Csnk2b | ENSMUSG00000024387 | 13001 | -0.24757576550816 | 5.48430174652032 | -3.14602142077239 | 0.0074817033791 | 0.0194859422534866 |
| Serbp1 | ENSMUSG00000036371 | 66870 | -0.20273850004395 | 7.89756451924755 | -3.14557697838862 | 0.0074881939842 | 0.0194982011534974 |
| Polr3d | ENSMUSG00000000776 | 67065 | -0.254362016298082 | 3.98566657309123 | -3.14543914408839 | 0.0074902080466 | 0.0194988007043814 |
| Zfp871 | ENSMUSG00000024298 | 208292 | 0.234087795732631 | 4.44781381421894 | 3.14459084897648 | 0.0075026153969 | 0.0195264497389373 |
| Slc15a4 | ENSMUSG00000029416 | 100561 | 0.461606979746936 | 2.59591055201736 | 3.14314811240963 | 0.0075237642415 | 0.019576830974596 |
| Cox19 | ENSMUSG00000045438 | 68033 | -0.378399824078532 | 4.08912669989575 | -3.14299643943481 | 0.0075259910400 | 0.0195779647926105 |
| Cuta | ENSMUSG00000024194 | 67675 | -0.235651690547877 | 3.87961530553477 | -3.1412859467116 | 0.0075511492888 | 0.0196387373010845 |
| Tlk2 | ENSMUSG00000020694 | 24086 | -0.191244682013686 | 5.14644864573498 | -3.13862899763343 | 0.0075903944963 | 0.0197361090008766 |
| Mrps18c | ENSMUSG00000016833 | 68735 | -0.378899032585019 | 2.83869093299556 | -3.13797412296089 | 0.0076000986861 | 0.0197531736524269 |
| Ablim2 | ENSMUSG00000029095 | 231148 | 0.237509995978927 | 5.23216026523946 | 3.13794222890458 | 0.0076005716203 | 0.0197531736524269 |
| Spryd4 | ENSMUSG00000051346 | 66701 | -0.3897277771132946 | 3.16536678892015 | -3.13762209919286 | 0.0076053202201 | 0.0197608165738199 |
| Frmd8 | ENSMUSG00000024816 | 67457 | 0.454565615386308 | 1.66383375640896 | 3.137466955547 | 0.0076076225852 | 0.019762101349642 |
| Skiv2l | ENSMUSG00000040356 | 108077 | 0.274705060958485 | 3.96891395568855 | 3.13705678185454 | 0.0076137129969 | 0.0197732232763255 |
| Mrps30 | ENSMUSG00000021731 | 59054 | -0.247372664578377 | 4.4843211475532 | -3.13635453223782 | 0.0076241515417 | 0.0197956295732305 |
| Hdac3 | ENSMUSG00000024454 | 15183 | -0.290641542756775 | 4.06111367981744 | -3.13591778899658 | 0.0076306506744 | 0.019807799221549 |
| Rnf130 | ENSMUSG00000020376 | 59044 | -0.210857300019318 | 4.32839729301205 | -3.13437164683902 | 0.0076537030524 | 0.0198629221430959 |
| Ints2 | ENSMUSG00000018068 | 70422 | 0.421951818340619 | 2.32163944849335 | 3.13387992729652 | 0.0076610489312 | 0.0198772669992843 |
| Lgals4 | ENSMUSG00000053964 | 16855 | -0.978275141847803 | 1.07572243950671 | -3.13288915081314 | 0.0076758716449 | 0.019910999975118 |
| Nde1 | ENSMUSG00000022678 | 67203 | -0.221022535835302 | 3.99252446459217 | -3.13235857881981 | 0.0076838211185 | 0.0199268917312539 |
| Lpin3 | ENSMUSG00000027412 | 64899 | 0.488867415030345 | 2.20721722285738 | 3.13157571752656 | 0.0076955655822 | 0.0199503613357034 |
| Cep104 | ENSMUSG00000039523 | 230967 | -0.205810774146221 | 5.56368694013638 | -3.13146951665188 | 0.0076971601817 | 0.0199503613357034 |
| Rnf19b | ENSMUSG00000028793 | 75234 | 0.375661847484906 | 3.96068226356687 | 3.13138835292674 | 0.0076983790718 | 0.0199503613357034 |
| Huwe1 | ENSMUSG00000025261 | 59026 | 0.249978268634502 | 7.98687338501667 | 3.13126902274282 | 0.0077001714825 | 0.0199503613357034 |
| Snx10 | ENSMUSG00000038301 | 71982 | 0.26157973799693 | 3.1683380361081 | 3.13097379511367 | 0.0077046077645 | 0.019957124993868 |
| Irs2 | ENSMUSG00000038894 | 384783 | -0.924556196372947 | 5.20278180466073 | -3.13026577219032 | 0.0077152573538 | 0.019979975866985 |
| Cnot8 | ENSMUSG00000020515 | 69125 | -0.21366320197783 | 4.40600366539672 | -3.12987471145357 | 0.0077211457025 | 0.0199904887907728 |
| Tssc4 | ENSMUSG00000045752 | 56844 | 0.496625199207499 | 1.51729273879378 | 3.12956717638105 | 0.0077257795181 | 0.0199977494465258 |
| Akr1a1 | ENSMUSG00000028692 | 58810 | 0.204330103710288 | 5.80393241937362 | 3.12789555905928 | 0.0077510152703 | 0.0200583210038174 |
| Trio | ENSMUSG00000022263 | 223435 | 0.210025468292936 | 6.15694075446429 | 3.12649289432343 | 0.0077722540887 | 0.020108522945236 |
| Gtf2i | ENSMUSG00000060261 | 14886 | -0.178270220656971 | 6.43160148918612 | -3.12635712826079 | 0.0077743129004 | 0.0201090899940755 |
| Fam189b | ENSMUSG00000032657 | 68521 | 0.74517813129446 | 1.22200849649078 | 3.12532629821687 | 0.0077899625541 | 0.0201414009987133 |
| Myo7a | ENSMUSG00000030761 | 17921 | 0.524836956623139 | 3.04417627136171 | 3.1252916083828 | 0.0077904897468 | 0.0201414009987133 |
| Hspa4l | ENSMUSG00000025757 | 18415 | 0.380236493039998 | 3.91743807421977 | 3.1238557171461 | 0.0078123426523 | 0.0201931230865693 |
| Elf4 | ENSMUSG00000031103 | 56501 | 0.500157927522688 | 2.56450464958156 | 3.1236832172237 | 0.0078149720418 | 0.0201951440636025 |
| Stat1 | ENSMUSG00000026104 | 20846 | -0.641454771739144 | 4.70797319829822 | -3.12242722953139 | 0.0078341434530 | 0.0202399012254395 |
| Birc2 | ENSMUSG00000057367 | 11797 | -0.272224951886815 | 4.1033477748209 | -3.12150077029398 | 0.0078483149443 | 0.0202717227450882 |
| Rheb | ENSMUSG00000028945 | 19744 | -0.313382471243054 | 4.49339425484387 | -3.12136841000279 | 0.0078503416632 | 0.0202721674275924 |
| Lypla1 | ENSMUSG00000025903 | 18777 | -0.270437145936267 | 5.52697890864157 | -3.11994665733896 | 0.0078721446242 | 0.0203236686083394 |
| Clic1 | ENSMUSG00000007041 | 114584 | 0.509075749087704 | 3.07945349326792 | 3.11897465655112 | 0.0078870852081 | 0.0203547388589783 |
| Zfp954 | ENSMUSG00000062116 | 232853 | 0.542516137979288 | 1.3383439203026 | 3.11892146943444 | 0.0078879035592 | 0.0203547388589783 |
| Rnf40 | ENSMUSG00000030816 | 233900 | -0.165305804160477 | 5.33026128527342 | -3.11860046037029 | 0.0078928444855 | 0.0203626818672729 |
| Ggh | ENSMUSG00000073987 | 14590 | 0.373996370624823 | 2.32197351570602 | 3.11836444523184 | 0.0078964791627 | 0.0203672519415713 |
| Fgd4 | ENSMUSG00000022788 | 224014 | -0.947866529418631 | 5.10198485032529 | -3.11781659951718 | 0.0079049225138 | 0.0203842198629674 |
| Ttf2 | ENSMUSG00000033222 | 74044 | 0.394054006845778 | 2.19653115569315 | 3.11728583017298 | 0.0079131112575 | 0.0204005233859395 |
| Emg1 | ENSMUSG00000004268 | 14791 | -0.317137433377704 | 3.5130967529821 | -3.11659148591532 | 0.0079238363968 | 0.0204233566739227 |
| Rtraf | ENSMUSG00000021807 | 68045 | -0.233520577154766 | 5.72699859569548 | -3.11582454556929 | 0.0079356997072 | 0.0204491120932629 |
| Cd164 | ENSMUSG00000019818 | 53599 | 0.199744722002005 | 5.91364389983121 | 3.1152034460882 | 0.0079453200546 | 0.0204629699029365 |
| Abcb4 | ENSMUSG00000042476 | 18670 | -0.29861913534 | 6.80991419212916 | -3.11516132747364 | 0.0079459728594 | 0.0204629699029365 |
| Txnl4a | ENSMUSG00000024571 | 27366 | -0.202129216702732 | 4.91510354830874 | -3.11511483264882 | 0.0079466935539 | 0.0204629699029365 |
| Fxn | ENSMUSG00000059363 | 14297 | -0.2995230076545 | 3.39630064098972 | -3.11445283180133 | 0.0079569619804 | 0.0204845858238518 |
| Ppm1j | ENSMUSG00000002228 | 71887 | -0.296101649865944 | 3.29606796709071 | -3.11325606116465 | 0.0079755588598 | 0.0205276274775958 |
| Eif4e3 | ENSMUSG00000093661 | 66892 | -0.335404527782118 | 3.5067099100081 | -3.11251525653183 | 0.0079870920566 | 0.0205524725213586 |
| Pcgf2 | ENSMUSG00000018537 | 22658 | 0.883546429267216 | 0.832394970322883 | 3.11184287435649 | 0.0079975743906 | 0.0205730430182871 |
| Adcy5 | ENSMUSG00000022840 | 224129 | -0.470861707207 | 1.70597133991761 | -3.11176109074738 | 0.0079988503181 | 0.0205730430182871 |
| Ifitm2 | ENSMUSG00000060591 | 80876 | 0.414626462324139 | 4.0283277031472 | 3.11147146050715 | 0.0080033705448 | 0.0205798267291148 |
| Cyp2d22 | ENSMUSG00000061740 | 56448 | -0.645749644648591 | 1.76292970004048 | -3.11123373461504 | 0.0080070826059 | 0.0205845296249339 |
| 3830406C13Rik | ENSMUSG00000033111 | 218734 | -0.290904500442855 | 2.68865643714826 | -3.11105824570124 | 0.0080098239434 | 0.0205867353693484 |
| Ube2d-ps | ENSMUSG00000083844 | 76508 | -0.281372087682016 | 4.3372214999989 | -3.10854070151528 | 0.0080492538375 | 0.0206832143154338 |
| Tom1 | ENSMUSG00000042870 | 21968 | -0.24060014586137 | 5.50792283331309 | -3.10806836960491 | 0.0080566730138 | 0.020697413093955 |
| Pigyl | ENSMUSG00000010607 | 66268 | -0.377940871687719 | 2.59478204721368 | -3.10770249072069 | 0.0080624247513 | 0.0207073225933603 |
| Slc35b4 | ENSMUSG00000018999 | 58246 | 0.336717443043302 | 2.71721224729614 | 3.10750367915948 | 0.0080655518483 | 0.0207104879621351 |
| Ctif | ENSMUSG00000052928 | 269037 | 0.316738931024933 | 3.54550921799788 | 3.10501682228338 | 0.0081047696158 | 0.0208063026470877 |
| Tcf12 | ENSMUSG00000032228 | 21406 | 0.17415321033986 | 5.36085905804802 | 3.10392444428121 | 0.0081220563572 | 0.0208457849357093 |
| Arl6ip6 | ENSMUSG00000026960 | 65103 | -0.508410771436406 | 1.71805773723238 | -3.10371533405253 | 0.0081253696808 | 0.0208493934229412 |
| Pgs1 | ENSMUSG00000017715 | 74451 | 0.272578902342723 | 3.39433403159792 | 3.10336814077607 | 0.0081308738823 | 0.0208586206073371 |
| Gltp | ENSMUSG00000011884 | 56356 | 0.63924341963396 | 2.47025034512004 | 3.10322256528217 | 0.0081331828557 | 0.0208596484740239 |
| Ctns | ENSMUSG00000005949 | 83429 | 0.373943220480099 | 2.19780510538208 | 3.10239822604444 | 0.0081462700151 | 0.0208883128161489 |
| Stau2 | ENSMUSG00000025920 | 29819 | -0.222452449336061 | 5.72733540162799 | -3.10225540016519 | 0.0081485396386 | 0.0208892323616523 |
| Bub1b | ENSMUSG00000040084 | 12236 | 1.04213127534157 | 0.290843420279434 | 3.10046755934209 | 0.0081770032168 | 0.0209550898956711 |
| Msmo1 | ENSMUSG00000031604 | 66234 | 0.460230670972263 | 1.91273897341473 | 3.10040107311274 | 0.0081780636259 | 0.0209550898956711 |
| Vmn1r6 | ENSMUSG00000115701 | 171193 | -1.51641087086937 | 2.85112348844135 | -3.09899607166923 | 0.0082005044405 | 0.0210076667539894 |
| Fasn | ENSMUSG00000025153 | 14104 | 0.924155035348704 | 4.80275139904357 | 3.09864009715198 | 0.0082061998184 | 0.021017177758083 |
| Ccdc66 | ENSMUSG00000046753 | 320234 | -0.272596394033539 | 4.03138345860775 | -3.09852372472012 | 0.0082080625605 | 0.021017177758083 |
| Ap1s1 | ENSMUSG00000004849 | 11769 | 0.45384069874078 | 1.82891165013077 | 3.09805270091068 | 0.0082156064091 | 0.0210315676014461 |
| Cebpzos | ENSMUSG00000062691 | 68554 | -0.430432372652472 | 2.98656641035137 | -3.09626914777543 | 0.0082442340883 | 0.0210999116880031 |
| Gtf2f1 | ENSMUSG00000002658 | 98053 | 0.258946441679737 | 4.18939225887 | 3.09614583781185 | 0.0082462169906 | 0.0211000463306053 |
| Cnrip1 | ENSMUSG00000044629 | 380686 | 0.830580928296217 | 0.662414826948523 | 3.09555699694145 | 0.0082556924678 | 0.0211193480848023 |
| Apod | ENSMUSG00000022548 | 11815 | 0.677652446939072 | 4.28693754424258 | 3.09539086554384 | 0.0082583677701 | 0.0211212489694817 |
| Smap2 | ENSMUSG00000032870 | 69780 | 0.279634172468333 | 4.24122291625394 | 3.0940956529523 | 0.0082792548556 | 0.0211697157491412 |
| Atxn7l1 | ENSMUSG00000020564 | 380753 | -0.338521736178179 | 3.48022854127619 | -3.09369122673516 | 0.0082857875429 | 0.0211814648344595 |
| Cdh2 | ENSMUSG00000024304 | 12558 | 0.443439476263291 | 2.13501922251263 | 3.09306291432747 | 0.0082959468289 | 0.0212024771539907 |
| Gpr161 | ENSMUSG00000040836 | 240888 | 0.701940098364677 | 1.13163597415508 | 3.09020687521085 | 0.0083422829490 | 0.0213159174651185 |
| Cmip | ENSMUSG00000034390 | 74440 | 0.495507230729724 | 3.12247931859538 | 3.08991622203832 | 0.0083470128840 | 0.0213195855140714 |
| Gpd1l | ENSMUSG00000050627 | 333433 | -0.227554706594606 | 3.87121308440716 | -3.08978936901575 | 0.0083490780583 | 0.0213195855140714 |
| Tmem263 | ENSMUSG00000060935 | 103266 | 0.268682128615415 | 3.75667285952608 | 3.08975917912512 | 0.0083495696263 | 0.0213195855140714 |
| Pkn3 | ENSMUSG00000026785 | 263803 | -0.509822768953214 | 2.13659766759241 | -3.0895249443028 | 0.0083533845411 | 0.0213243452638588 |
| Erlec1 | ENSMUSG00000020311 | 66753 | -0.298444692968991 | 4.28853803999052 | -3.08872459343793 | 0.0083664327077 | 0.0213526677395324 |
| Rint1 | ENSMUSG00000028999 | 72772 | 0.364188065356173 | 3.0615482377856 | 3.0877407356832 | 0.0083825003694 | 0.0213886814981661 |
| Ebf1 | ENSMUSG00000057098 | 13591 | 0.332647435876753 | 4.31273539352061 | 3.08748911731923 | 0.0083866145451 | 0.0213941852025357 |
| Snapc3 | ENSMUSG00000028483 | 77634 | 0.377332001151792 | 2.81579191071624 | 3.08658208945418 | 0.0084014618974 | 0.0214270601961015 |
| Sox7 | ENSMUSG00000063060 | 20680 | -0.946304935468843 | 2.94370803663406 | -3.08634790239982 | 0.0084052996033 | 0.0214318474373779 |
| Dapk3 | ENSMUSG00000034974 | 13144 | 0.247969055981787 | 3.25525076147631 | 3.08542861687394 | 0.0084203811161 | 0.0214652952379146 |
| Mtch2 | ENSMUSG00000027282 | 56428 | -0.214550698301004 | 5.62493945385664 | -3.08471664949776 | 0.0084320799157 | 0.0214888891747752 |
| Celsr2 | ENSMUSG00000068740 | 53883 | -0.445347306337864 | 2.61247927992252 | -3.0846261425337 | 0.0084335682517 | 0.0214888891747752 |
| Gnai3 | ENSMUSG00000000001 | 14679 | 0.346667258001519 | 3.72359831968509 | 3.08317912270019 | 0.0084573991749 | 0.0215445887627588 |

| | | | | | | | |
|---|---|---|---|---|---|---|---|
| Slc9a7 | ENSMUSG00000037341 | 236727 | 0.285330780695096 | 2.83228135968735 | 3.08262869194765 | 0.0084664817661 | 0.0215627008821524 |
| Tcap | ENSMUSG00000007877 | 21393 | -0.488603794020912 | 9.63905667802442 | -3.08169446555062 | 0.0084819195225 | 0.0215969863267169 |
| 1600014C10Rik | ENSMUSG00000054676 | 72244 | -0.309915541640785 | 3.0187056901359 | -3.08149806460916 | 0.0084851683535 | 0.0216002275877275 |
| Mrm1 | ENSMUSG00000018405 | 217038 | -0.515997148228198 | 1.89463578007149 | -3.08001659195303 | 0.0085097160695 | 0.0216576731912991 |
| Pet100 | ENSMUSG00000087687 | 100503890 | -0.350311488555649 | 3.59700460015091 | -3.07963393936935 | 0.0085160679645 | 0.0216687939757934 |
| Nop10 | ENSMUSG00000027133 | 66181 | -0.367731667309436 | 3.09963977287496 | -3.07841769825547 | 0.0085362884379 | 0.0217151894148626 |
| Chkb | ENSMUSG00000022617 | 12651 | -0.326680528138658 | 3.4582787590138 | -3.07779988162653 | 0.0085465781445 | 0.0217363065841223 |
| Tshz2 | ENSMUSG00000047907 | 228911 | 0.265378726303111 | 4.68482497616139 | 3.07701841634663 | 0.0085596110708 | 0.0217643890707243 |
| Gria3 | ENSMUSG00000001986 | 53623 | 0.830634884873526 | 1.01403127747967 | 3.07665443919285 | 0.0085656880583 | 0.0217747758523921 |
| Lrrc61 | ENSMUSG00000073096 | 243371 | -0.419140299291502 | 2.37950314228466 | -3.07649472342719 | 0.0085683560370 | 0.0217763997808655 |
| Hars | ENSMUSG00000001380 | 15115 | 0.28066314212786 | 4.22431881960854 | 3.07637771029869 | 0.0085703112119 | 0.0217763997808655 |
| Aldh1l1 | ENSMUSG00000030088 | 107747 | -0.633209145981847 | 1.49893787707626 | -3.0727685681838 | 0.0086308346617 | 0.0219250880054218 |
| Ube2d1 | ENSMUSG00000019927 | 216080 | -0.206290314244015 | 5.41005199986266 | -3.07258147011805 | 0.0086339837531 | 0.0219279917299386 |
| Spats2l | ENSMUSG00000038305 | 67198 | 0.431962547615325 | 2.83785252271809 | 3.07138542319437 | 0.0086541416931 | 0.0219740819624536 |
| Got2 | ENSMUSG00000031672 | 14719 | -0.262179413379926 | 8.57406563809072 | -3.07076439733686 | 0.0086646267774 | 0.0219914058192455 |
| Apbb1 | ENSMUSG00000037032 | 11785 | 0.353452613306959 | 2.80484589465755 | 3.07074301002207 | 0.0086649880947 | 0.0219914058192455 |
| Tcf25 | ENSMUSG00000001472 | 66855 | 0.17495556443706 | 6.21895658146713 | 3.07052715982724 | 0.0086686355056 | 0.0219955558755415 |
| Impad1 | ENSMUSG00000066324 | 242291 | -0.202575330834777 | 6.13437042117585 | -3.06866691168134 | 0.0087001330446 | 0.022070353750545 |
| Dgkz | ENSMUSG00000040479 | 104418 | 0.337056737064071 | 5.40695091351601 | 3.06824322095599 | 0.0087073228167 | 0.0220834676820035 |
| Ikbip | ENSMUSG00000019975 | 67454 | 0.493570394591355 | 2.91348997115493 | 3.0667432125098 | 0.0087328245133 | 0.0221430073662362 |
| Sh3bp1 | ENSMUSG00000022436 | 20401 | 0.75530789500838 | 0.836216413049733 | 3.06636630475078 | 0.0087392439906 | 0.0221541456543072 |
| Csrnp2 | ENSMUSG00000044636 | 207785 | 0.590936119315173 | 1.91365554642323 | 3.06623897356019 | 0.0087414137492 | 0.0221545081598155 |
| Epha2 | ENSMUSG00000006445 | 13836 | -0.538840718343334 | 1.72205981912591 | -3.06470847047602 | 0.0087675358537 | 0.0222155619883378 |
| Ice1 | ENSMUSG00000034525 | 218333 | 0.222243970522916 | 5.63948460832921 | 3.06380937098189 | 0.0087829175159 | 0.0222493792274874 |
| Hcfc1r1 | ENSMUSG00000023904 | 353502 | 0.213298348690166 | 5.85348813572429 | 3.06330689838511 | 0.0087915254156 | 0.0222660250968372 |
| Spin1 | ENSMUSG00000021395 | 20729 | 0.22889327893593 | 4.04588366060812 | 3.06169616254443 | 0.0088191755861 | 0.022330879854518 |
| Xylt2 | ENSMUSG00000020868 | 217119 | 0.396676554533656 | 2.44243898724404 | 3.0614458855946 | 0.0088234796292 | 0.0223366039434349 |
| Mylk3 | ENSMUSG00000031698 | 213435 | 0.662398334148413 | 1.50032010127466 | 3.06114556664605 | 0.0088286470043 | 0.0223445103969485 |
| n-R5s100 | ENSMUSG00000094720 | 100861471 | 1.12466911973163 | 9.05336764947015 | 3.06038566154472 | 0.0088417355795 | 0.0223708360908324 |
| Rufy3 | ENSMUSG00000029291 | 52822 | 0.223420872577015 | 3.48885804928268 | 3.06029095947263 | 0.0088433680740 | 0.0223708360908324 |
| Ddx3x | ENSMUSG00000000787 | 13205 | -0.171635340545586 | 7.03195036195983 | -3.06018538597816 | 0.0088451883253 | 0.0223708360908324 |
| Usp1 | ENSMUSG00000028560 | 230484 | -0.225002469815155 | 4.3403706364538 | -3.05953173888298 | 0.0088564665042 | 0.022394178893861 |
| Spry4 | ENSMUSG00000024427 | 24066 | -0.757657960454908 | 2.80847807110628 | -3.05711971329039 | 0.0088982078660 | 0.0224945213190744 |
| Dgcr6 | ENSMUSG00000003531 | 13353 | -0.349753186053352 | 3.57887921221608 | -3.05625750217153 | 0.0089131762266 | 0.0225271512909307 |
| Polrmt | ENSMUSG00000020329 | 216151 | -0.197782443743904 | 4.17831965681072 | -3.05439252982984 | 0.0089456385351 | 0.0226039701404626 |
| Lsm12 | ENSMUSG00000020922 | 268490 | -0.231329566672053 | 5.43977959430722 | -3.05178034715971 | 0.0089913044617 | 0.0227141088678069 |
| Rasa4 | ENSMUSG00000004952 | 54153 | 0.412717128085473 | 4.05273816219365 | 3.05138101306606 | 0.0089983059460 | 0.0227265439685749 |
| Slc25a51 | ENSMUSG00000045973 | 230125 | -0.220862380338137 | 4.86165439781171 | -3.05049033149726 | 0.0090139416797 | 0.022760775352611 |
| Rabep2 | ENSMUSG00000030727 | 70314 | -0.40167076352705 | 3.86863528151036 | -3.05007908835506 | 0.0090211700675 | 0.022773766745468 |
| Fnip1 | ENSMUSG00000035992 | 216742 | -0.403523609873399 | 6.65029897728738 | -3.04939160241565 | 0.0090332668043 | 0.0227990393531928 |
| Mtrf1l | ENSMUSG00000019774 | 108853 | -0.37307724245897 | 2.24232015043391 | -3.04873840650913 | 0.0090447751029 | 0.0228228154779574 |
| Klf11 | ENSMUSG00000020653 | 194655 | 0.503705033521346 | 2.4139932439341 | 3.04753727017284 | 0.0090659752589 | 0.0228710305920551 |
| Ceacam1 | ENSMUSG00000074272 | 26365 | -0.645127854240449 | 2.64483905276073 | -3.04736447013525 | 0.0090690292492 | 0.022873456096921 |
| Itgb5 | ENSMUSG00000022817 | 16419 | -0.212411201261657 | 5.73969096397877 | -3.04692874630252 | 0.0090767345710 | 0.0228876091341614 |
| Olfr552 | ENSMUSG00000073973 | 259106 | -0.906653932323046 | 2.89082861826752 | -3.04494098090348 | 0.0091119686431 | 0.0229711552670584 |
| Zfp267 | ENSMUSG00000033883 | 241944 | 0.390947304330401 | 2.56109029854959 | 3.04362410294612 | 0.0091353856351 | 0.0230248790356049 |
| 2310069B03Rik | ENSMUSG00000100291 | 69652 | -1.13876378415641 | 0.442507522533836 | -3.04192795829955 | 0.0091656348388 | 0.0230957940119164 |
| Anapc7 | ENSMUSG00000029466 | 56317 | 0.206807519602192 | 3.63135445182376 | 3.0414041353015 | 0.0091749968191 | 0.0231140562870753 |
| Gm15417 | ENSMUSG00000074466 | 545539 | -0.497858786205229 | 1.32662165852199 | -3.04071894525789 | 0.0091872571420 | 0.0231396100966292 |
| Coa4 | ENSMUSG00000044881 | 68185 | -0.625650023603961 | 1.0605194221789 | -3.0404567649907 | 0.0091919527156 | 0.0231458744071334 |
| Rsl24d1 | ENSMUSG00000032215 | 225215 | -0.260238604728007 | 3.69580013232834 | -3.04034365581778 | 0.0091939792037 | 0.0231458744071334 |
| Sike1 | ENSMUSG00000027854 | 66641 | 0.268484709843873 | 4.2542509843312 | 3.0396633324097 | 0.0092061773779 | 0.0231712468152601 |
| Sap18 | ENSMUSG00000021963 | 20220 | -0.353232496704338 | 3.16514781540735 | -3.03781290633358 | 0.0092394367426 | 0.0232496047500588 |
| Gm7967 | ENSMUSG00000100009 | 666182 | -1.07436414620576 | 0.3840493424767 | -3.03686738338453 | 0.0092564774638 | 0.0232871243169308 |
| Epn1 | ENSMUSG00000035203 | 13854 | -0.208615256661584 | 6.52857103905948 | -3.03669994612238 | 0.0092594983557 | 0.0232893641341132 |
| 6430571L13Rik | ENSMUSG00000037977 | 235599 | -0.689980561683839 | 1.9540676421384 | -3.03620935449325 | 0.0092683552093 | 0.0233062780754053 |
| Sema3b | ENSMUSG00000057969 | 20347 | -0.299250393089757 | 3.22093323640138 | -3.03426705009612 | 0.0093035030096 | 0.0233892804503294 |
| Dcaf10 | ENSMUSG00000035572 | 242418 | 0.238469277208461 | 3.57337659295425 | 3.03331815185061 | 0.0093207222462 | 0.0234271820817351 |
| Mettl23 | ENSMUSG00000090266 | 74319 | -0.360760643857331 | 3.30356581102025 | -3.03318962944454 | 0.0093230569149 | 0.0234276632498735 |
| Gng2 | ENSMUSG00000043004 | 14702 | 0.738936783406262 | 0.927829555448116 | 3.03132313744916 | 0.0093570280523 | 0.0235076243713964 |
| Klf10 | ENSMUSG00000037465 | 21847 | 0.718306034639484 | 3.96370938767462 | 3.0307128313702 | 0.0093681625569 | 0.0235301895473937 |
| Usp16 | ENSMUSG00000025616 | 74112 | -0.182454726866245 | 5.42704423658467 | -3.03018904813264 | 0.0093777290050 | 0.0235464296063147 |
| Ttc3 | ENSMUSG00000040785 | 22129 | 0.185677497724178 | 6.56255229673139 | 3.03012297478106 | 0.0093789364656 | 0.0235464296063147 |
| Rpp14 | ENSMUSG00000023156 | 67053 | -0.280932470843304 | 3.54573164618074 | -3.02940738088186 | 0.0093920235019 | 0.0235738711596249 |
| Bahd1 | ENSMUSG00000040007 | 228536 | 0.419197552076952 | 2.72724335037197 | 3.02848557046243 | 0.0094089085926 | 0.0236108309977154 |
| Yipf5 | ENSMUSG00000024487 | 67180 | 0.370665203716658 | 3.067692286461 | 3.02364917671473 | 0.0094979928310 | 0.0238289097167689 |
| Eapp | ENSMUSG00000054302 | 66266 | -0.262804011909621 | 4.07754582174312 | -3.02282871608829 | 0.0095131881037 | 0.0238607505471707 |
| Xpnpep1 | ENSMUSG00000025027 | 170750 | 0.26867650813006 | 4.98667372141929 | 3.02272827294011 | 0.0095150500077 | 0.0238607505471707 |
| Cbx7 | ENSMUSG00000053411 | 52609 | -0.461594772778764 | 3.04918858498288 | -3.02094548025246 | 0.0095481576251 | 0.0239382823395001 |
| Zc3h3 | ENSMUSG00000075600 | 223642 | 0.415511008834192 | 2.40111978610941 | 3.01961388573743 | 0.0095729606793 | 0.0239949628951554 |
| Mapk4 | ENSMUSG00000024558 | 225724 | 0.589112202728338 | 1.61702522271383 | 3.01806332848577 | 0.0096019227542 | 0.0240620398044035 |
| Sorcs2 | ENSMUSG00000029093 | 81840 | 0.559039807946192 | 2.27057279205022 | 3.01768474422009 | 0.0096090073198 | 0.0240742743554601 |
| Dtd1 | ENSMUSG00000027430 | 66044 | 0.302862062286569 | 2.53146170869409 | 3.0162790102197 | 0.0096353586233 | 0.0241347628578519 |
| Pcdhga10 | ENSMUSG00000102222 | 93722 | 0.524924548950412 | 1.33888394849709 | 3.01375153691924 | 0.0096829178603 | 0.0242475927060251 |
| Gapdh | ENSMUSG00000057666 | 14433 | -0.301486605171105 | 10.6700823447392 | -3.01364962815404 | 0.0096848403398 | 0.0242475927060251 |
| Zbtb5 | ENSMUSG00000049657 | 230119 | 0.385420756151624 | 1.9885309375292 | 3.0135290307859 | 0.0096871158631 | 0.0242477360887778 |
| Gbp2 | ENSMUSG00000028270 | 14469 | -0.651860867797155 | 3.4191969552436 | -3.01335109508613 | 0.0096904742574 | 0.024250589310568 |
| Arid1b | ENSMUSG00000069729 | 239985 | -0.177916063779707 | 6.17894106600457 | -3.01283562099748 | 0.0097002099338 | 0.0242693968384638 |
| Rgcc | ENSMUSG00000022018 | 66214 | 0.538716761724844 | 5.10626979445943 | 3.01210362476973 | 0.0097140516859 | 0.0242984665855919 |
| Nme2 | ENSMUSG00000020857 | 18103 | -0.281239640008941 | 6.14572572102354 | -3.01170640163139 | 0.0097215712050 | 0.0243117123866199 |
| Jmjd8 | ENSMUSG00000025736 | 72106 | 0.356142203113807 | 2.26114132854412 | 3.01156869857035 | 0.0097241792995 | 0.0243126724435451 |
| Ralgps1 | ENSMUSG00000038831 | 241308 | 0.417242728682408 | 2.12535896971672 | 3.01122137334025 | 0.0097307607159 | 0.024323564003189 |
| Rab34 | ENSMUSG00000002059 | 19376 | 0.460364693976771 | 1.73455144418094 | 3.01025830506162 | 0.0097490328553 | 0.0243636666989869 |
| Dars2 | ENSMUSG00000026709 | 226539 | -0.358124978795193 | 2.4891813083439 | -3.0094529013299 | 0.0097643397512 | 0.0243963423591043 |
| Fopnl | ENSMUSG00000022677 | 66086 | -0.258287248749112 | 4.14539488590688 | -3.00796616544548 | 0.0097926581062 | 0.0244615049725144 |
| Crispld1 | ENSMUSG00000025776 | 83691 | 0.505876290956492 | 3.55040998627092 | 3.00674638257809 | 0.0098159524700 | 0.0245140910326594 |
| Tspyl1 | ENSMUSG00000047514 | 22110 | -0.197547096840665 | 5.31805446633818 | -3.00618218998101 | 0.0098267454943 | 0.0245346222157814 |
| Cby1 | ENSMUSG00000022428 | 73739 | -0.391253981829081 | 1.99065141863058 | -3.00598060814043 | 0.0098306046120 | 0.0245346222157814 |
| Mlxip | ENSMUSG00000038342 | 208104 | -0.194077757300634 | 5.85475851381299 | -3.00596481191018 | 0.0098309070812 | 0.0245346222157814 |
| Mrpl22 | ENSMUSG00000020514 | 216767 | -0.289497577925077 | 3.40672439864569 | -3.00527308528375 | 0.0098441614383 | 0.0245620928286367 |
| 4930521O11Rik | ENSMUSG00000071591 | 74715 | -2.41539739261321 | -1.56769608877693 | -3.00491144039615 | 0.0098510980570 | 0.0245737911595544 |
| Stk38l | ENSMUSG00000001630 | 232533 | -0.228137271388233 | 3.54007610373538 | -3.00356524960937 | 0.0098769615829 | 0.0246326869868486 |
| Card19 | ENSMUSG00000037960 | 68480 | 0.238447541835417 | 4.73908432981556 | 3.00319061101039 | 0.0098841712374 | 0.0246450446615147 |
| Paip1 | ENSMUSG00000025451 | 218693 | -0.2085398317422 | 4.82093991122619 | -3.0019376490238 | 0.0099083214936 | 0.0246996265101637 |
| Nkap | ENSMUSG00000016409 | 67050 | -0.246425517724766 | 3.27088942408103 | -3.00056381301185 | 0.0099348687163 | 0.0247601573046805 |
| Ccpg1os | ENSMUSG00000086158 | 546143 | -0.486707790525235 | 1.33254875454255 | -3.00038703365833 | 0.0099382898111 | 0.0247630375940601 |
| Igfbp7 | ENSMUSG00000036256 | 29817 | 0.40661704067727 | 6.06453471580132 | 2.99926865203552 | 0.0099599601710 | 0.0248113775365062 |
| Rfx5 | ENSMUSG00000005774 | 53970 | 0.377065141218482 | 2.19739535967976 | 2.99851866400016 | 0.0099745185459 | 0.024841982735367 |
| Ky | ENSMUSG00000035606 | 16716 | -0.348243210937919 | 4.87118830793222 | -2.99734850328428 | 0.0099972751926 | 0.0248929875013003 |
| Pcyox1 | ENSMUSG00000029998 | 66881 | 0.217098486714236 | 5.64788843841593 | 2.99666620013448 | 0.0100105679725 | 0.0249202346544182 |
| Manf | ENSMUSG00000032575 | 74840 | 0.701908657261077 | 4.13531241186671 | 2.9965528757634 | 0.0100127774771 | 0.0249202346544182 |
| Phip | ENSMUSG00000032253 | 83946 | 0.268615031690181 | 5.67252317014492 | 2.99629428783504 | 0.0100178210181 | 0.0249271116661428 |
| Vps37a | ENSMUSG00000031600 | 52348 | -0.19198724918248 | 4.95614190219622 | -2.99519384687375 | 0.0100393123432 | 0.0249749028731294 |
| Uba3 | ENSMUSG00000030061 | 22200 | 0.249189390542978 | 4.93356200763889 | 2.99314307543068 | 0.0100794852538 | 0.0250691361341892 |
| Dnajc9 | ENSMUSG00000021811 | 108671 | -0.333406001280852 | 2.90254351800338 | -2.99289973192428 | 0.0100842627019 | 0.0250753129196142 |
| Taf6 | ENSMUSG00000036980 | 21343 | -0.204925872316533 | 4.00355860973666 | -2.99235542898965 | 0.0100949568610 | 0.0250961959171768 |
| Nrip1 | ENSMUSG00000048490 | 268903 | 0.440492326143206 | 4.85800980446795 | 2.99000858925432 | 0.0101411949627 | 0.0252054120368864 |
| Adcy6 | ENSMUSG00000022994 | 11512 | -0.244312369520336 | 4.41897185801792 | -2.98956775815491 | 0.0101499037018 | 0.0252213224289945 |
| Cish | ENSMUSG00000032578 | 12700 | -1.6992050944766 | 2.87802031876467 | -2.98882306236751 | 0.0101646321899 | 0.0252521805609742 |
| Cyp4v3 | ENSMUSG00000079057 | 102294 | 0.753030848410513 | 0.713228519052719 | 2.9885869516846 | 0.0101693063692 | 0.0252580522432149 |
| Nxpe4 | ENSMUSG00000044229 | 244853 | -0.527783768702555 | 2.00740715586019 | -2.98845380530168 | 0.0101719431477 | 0.0252588620054003 |
| Luc7l3 | ENSMUSG00000020863 | 67684 | -0.196974469584783 | 5.2995062882397 | -2.98829491969136 | 0.0101750905413 | 0.0252598305745551 |
| Cst3 | ENSMUSG00000027447 | 13010 | 0.224235926517891 | 6.52906198240467 | 2.98820082748226 | 0.0101769548852 | 0.0252598305745551 |
| Shroom4 | ENSMUSG00000068270 | 208431 | -0.649373600232455 | 3.83826822767401 | -2.98766972452361 | 0.0101874845063 | 0.0252775635152269 |
| Nsun3 | ENSMUSG00000050312 | 106338 | -0.301827853176895 | 2.73186368540387 | -2.98760722859301 | 0.0101887242565 | 0.0252775635152269 |
| Stt3b | ENSMUSG00000032437 | 68292 | 0.200049723149632 | 6.0430427169896 | 2.98728894024526 | 0.0101950405532 | 0.0252842118311668 |

| | | | | | | | |
|---|---|---|---|---|---|---|---|
| Atxn7 | ENSMUSG00000021738 | 246103 | -0.230901835469398 | 4.68743482571546 | -2.98723908972144 | 0.0101960301666 | 0.0252842118311668 |
| Eif3c | ENSMUSG00000030738 | 56347 | -0.196759132761516 | 7.28289543525688 | -2.98644559215812 | 0.0102117951819 | 0.0253125495147857 |
| Nme6 | ENSMUSG00000032478 | 54369 | -0.330756304342531 | 2.35789814621659 | -2.98637043148643 | 0.0102132897059 | 0.0253125495147857 |
| Usp13 | ENSMUSG00000056900 | 72607 | -0.211896006205309 | 8.38557658485894 | -2.98631437332203 | 0.0102144045292 | 0.0253125495147857 |
| Zfp597 | ENSMUSG00000039789 | 71063 | -0.337217220066893 | 2.47985077376296 | -2.98588566658193 | 0.0102229341621 | 0.025327944997034 |
| Rnf13 | ENSMUSG00000036503 | 24017 | 0.269244262148198 | 4.93190681973834 | 2.98526540729865 | 0.0102352874410 | 0.0253528046720142 |
| Plpp1 | ENSMUSG00000021759 | 19012 | 0.284463385644954 | 4.91702369564222 | 2.98476727730069 | 0.0102452190501 | 0.0253716559666956 |
| Hlx | ENSMUSG00000039377 | 15284 | 0.385953279865817 | 2.97286382469385 | 2.98390170809925 | 0.0102624992726 | 0.0254086929897175 |
| Aldh9a1 | ENSMUSG00000026687 | 56752 | -0.44989472340835 | 2.53856792772942 | -2.98318507328237 | 0.0102768279926 | 0.0254384073342741 |
| Arhgap31 | ENSMUSG00000022799 | 12549 | -0.354102479551234 | 5.32174862309073 | -2.98171734787697 | 0.0103062361801 | 0.0255054262361734 |
| Dstyk | ENSMUSG00000042046 | 213452 | 0.206887588305466 | 4.19808205865915 | 2.98068499894052 | 0.0103269707837 | 0.025550954648456 |
| Nenf | ENSMUSG00000037499 | 66208 | -0.249883479460512 | 4.86155460977372 | -2.98009247312904 | 0.0103388902313 | 0.0255746569628138 |
| Scrn1 | ENSMUSG00000019124 | 69938 | 0.550094010214671 | 1.45069660734296 | 2.97926741368555 | 0.0103555100796 | 0.0256078860543007 |
| Psmd14 | ENSMUSG00000026914 | 59029 | 0.219200508269712 | 4.71722094120785 | 2.9791930723154 | 0.0103570088963 | 0.0256078860543007 |
| Ppp2ca | ENSMUSG00000020349 | 19052 | -0.230920125855379 | 7.20933593622633 | -2.9790562244569 | 0.0103597684846 | 0.0256089166226959 |
| Tlcd2 | ENSMUSG00000038217 | 380712 | -0.424203944136612 | 1.74999894773921 | -2.97840189420135 | 0.0103729733573 | 0.0256314318342628 |
| Snrpa1 | ENSMUSG00000030512 | 68981 | -0.3011194665471 | 2.86923158371531 | -2.97837252645824 | 0.0103735664106 | 0.0256314318342628 |
| Adgrl2 | ENSMUSG00000028184 | 99633 | -0.391911884642235 | 3.76150442191605 | -2.97817289613334 | 0.0103775986418 | 0.0256356001703706 |
| Chrac1 | ENSMUSG00000068391 | 93696 | -0.33066474242176 | 4.20440014881463 | -2.97781688390383 | 0.0103847934002 | 0.0256475771933787 |
| Plcl2 | ENSMUSG00000038910 | 224860 | -0.300813591707843 | 4.10415846658401 | -2.97764626684274 | 0.0103882432008 | 0.0256503018812034 |
| Pim1 | ENSMUSG00000024014 | 18712 | 0.59353483186214 | 2.52735949186559 | 2.97665973275909 | 0.0104082127201 | 0.0256938060623347 |
| Ncoa7 | ENSMUSG00000039697 | 211329 | 0.385562307791893 | 2.25813050445336 | 2.97648080239016 | 0.0104118387125 | 0.0256969539325488 |
| Cilp2 | ENSMUSG00000044006 | 68709 | 0.795636766582495 | 4.16831717040817 | 2.97615867128951 | 0.0104183697929 | 0.0257072686696532 |
| Foxj2 | ENSMUSG00000003154 | 60611 | -0.196983444378976 | 5.33618129167505 | -2.97559874343425 | 0.0104297317541 | 0.0257294962320535 |
| Atpaf2 | ENSMUSG00000042709 | 246782 | -0.30391496124341 | 3.50633076697265 | -2.9750316107942 | 0.0104412524085 | 0.0257521051619285 |
| Ppp1ca | ENSMUSG00000040385 | 19045 | 0.219955864890544 | 6.08569046074297 | 2.97476071812926 | 0.0104467597263 | 0.0257598760589215 |
| C1qtnf1 | ENSMUSG00000017446 | 56745 | 0.459242985148471 | 2.21654308084078 | 2.97441461317392 | 0.0104538003106 | 0.0257714233638857 |
| Nova2 | ENSMUSG00000030411 | 384569 | -0.615575767241685 | 1.97050105563505 | -2.97304788663707 | 0.0104816486156 | 0.0258342505114251 |
| Hacl1 | ENSMUSG00000021884 | 56794 | -0.369469505388798 | 1.91450165205417 | -2.97202477886632 | 0.0105025444083 | 0.0258799172515805 |
| Arap1 | ENSMUSG00000032812 | 69710 | 0.282469770643524 | 4.45334559643964 | 2.97141696225267 | 0.0105149762321 | 0.0259047115603733 |
| Vwf | ENSMUSG00000001930 | 22371 | -0.265356396659663 | 4.97060581014709 | -2.97056263696345 | 0.0105324760078 | 0.0259419772964524 |
| Dnajb2 | ENSMUSG00000026203 | 56812 | -0.247275246395346 | 4.48834762459778 | -2.9701942588238 | 0.0105400306540 | 0.0259547364449637 |
| Eif2b5 | ENSMUSG00000003235 | 224045 | -0.207593408097627 | 5.25150068255827 | -2.96869210926344 | 0.0105708920411 | 0.0260248695724167 |
| Cbx6 | ENSMUSG00000089715 | 494448 | 0.339585640774085 | 3.83550366716388 | 2.9683751780329 | 0.0105774147419 | 0.0260283689431517 |
| Nfatc3 | ENSMUSG00000031902 | 18021 | -0.179682792612399 | 5.03668799107314 | -2.96831789114524 | 0.0105785941770 | 0.0260283689431517 |
| Yipf1 | ENSMUSG00000057375 | 230584 | 0.497193489167861 | 1.49447265804097 | 2.96827599213998 | 0.0105794568854 | 0.0260283689431517 |
| Xpc | ENSMUSG00000030094 | 22591 | -0.201084402429414 | 3.80012359055397 | -2.96774927781884 | 0.0105903079677 | 0.0260492026093252 |
| Atp6v1b2 | ENSMUSG00000006273 | 11966 | -0.173788279376479 | 4.70596870448506 | -2.9671724972505 | 0.0106022031125 | 0.026072594425876 |
| Fam122a | ENSMUSG00000074922 | 68034 | -0.280041498103236 | 2.73176791925015 | -2.96555684366914 | 0.0106355936735 | 0.0261488246615704 |
| Galc | ENSMUSG00000021003 | 14420 | 0.362630842350766 | 2.28079741098597 | 2.96471092930802 | 0.0106531175563 | 0.0261848862448896 |
| Dennd2c | ENSMUSG00000007379 | 329727 | 0.613368255877711 | 1.09636558822684 | 2.96461763134236 | 0.0106550520553 | 0.0261848862448896 |
| Slc30a5 | ENSMUSG00000021629 | 69048 | 0.277042590479257 | 3.32524556975523 | 2.96364731404768 | 0.0106751918283 | 0.026228483226749 |
| Gnb1 | ENSMUSG00000029064 | 14688 | 0.211040564773986 | 5.78447890448506 | 2.96305546002446 | 0.0106874947439 | 0.0262528101228256 |
| Fzd8 | ENSMUSG00000036904 | 14370 | 0.439681237800753 | 2.21789818665361 | 2.96280332610191 | 0.0106927401282 | 0.0262597938310673 |
| Cdk15 | ENSMUSG00000026023 | 271697 | -1.19091416076556 | 2.43654680472146 | -2.96152979792948 | 0.0107192734767 | 0.0263190427614779 |
| Smad7 | ENSMUSG00000025880 | 17131 | -0.516216506078759 | 3.28600886227866 | -2.95946576733466 | 0.0107624148546 | 0.0264164664149508 |
| Dgke | ENSMUSG00000000276 | 56077 | -0.429118621007664 | 2.70101719067391 | -2.95940031589265 | 0.0107637856931 | 0.0264164664149508 |
| Uhrf1bp1l | ENSMUSG00000019951 | 75089 | 0.208544893149534 | 5.10378198547651 | 2.95893657325349 | 0.0107735034303 | 0.0264343806952326 |
| Rpl12 | ENSMUSG00000038900 | 269261 | 0.568865950087598 | 2.26331441798534 | 2.95842795955845 | 0.0107841714127 | 0.0264546179785325 |
| Isca2 | ENSMUSG00000021241 | 74316 | -0.362399596942474 | 2.93487312392891 | -2.95608594391504 | 0.0108334291984 | 0.0265694894700403 |
| Eif3b | ENSMUSG00000056076 | 27979 | 0.225700374882833 | 6.08744732888136 | 2.9558504385238 | 0.0108383946865 | 0.0265757048719648 |
| Prkar2b | ENSMUSG00000002997 | 19088 | 0.95667289932715 | 0.574038363221814 | 2.95529885541019 | 0.0108500332865 | 0.0265982762626115 |
| Vps4b | ENSMUSG00000009907 | 20479 | 0.24195227919168 | 4.05582247262727 | 2.95504028999292 | 0.0108554933599 | 0.0266056946003571 |
| Cd34 | ENSMUSG00000016494 | 12490 | 0.293300169977627 | 5.487220864369 | 2.95386402016572 | 0.0108803667045 | 0.0266606788718881 |
| Scd1 | ENSMUSG00000037071 | 20249 | 0.879795948169007 | 5.89377800205111 | 2.95223133817051 | 0.0109149846807 | 0.0267395108797214 |
| Kmt5b | ENSMUSG00000045098 | 225888 | 0.207312054776989 | 4.72725108743123 | 2.95169549488656 | 0.0109263699421 | 0.0267614048481861 |
| Zrsr2 | ENSMUSG00000031370 | 22184 | 0.276388689523178 | 3.44505944677 | 2.95149097603935 | 0.0109307185234 | 0.02676605828405 |
| Jtb | ENSMUSG00000027937 | 23922 | -0.234901962652561 | 4.14890556471452 | -2.94962079211253 | 0.0109705626987 | 0.0268576082552741 |
| Alg10b | ENSMUSG00000075470 | 380959 | 0.205536426669629 | 4.30474856050631 | 2.9489206407864 | 0.0109855162754 | 0.0268881948962762 |
| Guf1 | ENSMUSG00000029208 | 231279 | -0.230594251274636 | 3.9020696174603 | -2.9485003148264 | 0.0109945031154 | 0.0269041669027316 |
| March2 | ENSMUSG00000079557 | 224703 | -0.187648834282854 | 4.76102724014061 | -2.94795942659061 | 0.0110060783419 | 0.0269206498659187 |
| Pkib | ENSMUSG00000019876 | 18768 | -1.19598211472135 | 2.92151208542078 | -2.9479554022645 | 0.0110061645092 | 0.0269206498659187 |
| Cacul1 | ENSMUSG00000033417 | 78832 | -0.223995088740366 | 4.82861946073858 | -2.94658581936981 | 0.0110355282269 | 0.0269826131764246 |
| Tmco1 | ENSMUSG00000052428 | 68944 | -0.215790894555283 | 4.25308519038291 | -2.94654361481785 | 0.0110364343163 | 0.0269826131764246 |
| Kctd1 | ENSMUSG00000036225 | 106931 | 0.429585738744391 | 2.02298219982217 | 2.94586120060371 | 0.0110510952694 | 0.0270124155614114 |
| Mak16 | ENSMUSG00000031578 | 67920 | 0.205009751875052 | 3.7552272243501 | 2.94502727773523 | 0.0110690373483 | 0.0270502229497722 |
| Rere | ENSMUSG00000039852 | 68703 | -0.214755519590621 | 6.49061014636205 | -2.94471611728107 | 0.0110757394192 | 0.027056531100007 |
| Hdlbp | ENSMUSG00000034088 | 110611 | -0.217547367752998 | 8.42742192721358 | -2.94451768506332 | 0.0110800155347 | 0.027056531100007 |
| Ifngr2 | ENSMUSG00000022965 | 15980 | -0.238797713121634 | 5.15909876179047 | -2.94448699691486 | 0.0110806769946 | 0.027056531100007 |
| 2610044O15Rik8 | ENSMUSG00000071302 | 72139 | -0.270081676416308 | 3.73335917401433 | -2.94444791151879 | 0.0110815195073 | 0.027056531100007 |
| Rufy1 | ENSMUSG00000020375 | 216724 | -0.198176242576629 | 4.56592593298315 | -2.94307222266908 | 0.0111112137704 | 0.0271202294279034 |
| Tcf4 | ENSMUSG00000053477 | 21413 | 0.200381550795002 | 6.30320805849033 | 2.94300697672241 | 0.0111126240554 | 0.0271202294279034 |
| Atp6v0a1 | ENSMUSG00000019302 | 11975 | 0.225130918551899 | 4.99688313209279 | 2.94289469214933 | 0.0111150514899 | 0.0271202294279034 |
| Ece1 | ENSMUSG00000057530 | 230857 | -0.234388603440805 | 6.13678650555618 | -2.94235661305477 | 0.0111266912729 | 0.0271425713689813 |
| Lnx1 | ENSMUSG00000029228 | 16924 | -0.605604238931584 | 1.2356659420245 | -2.94210503390368 | 0.0111321375886 | 0.0271497983002178 |
| Col4a4 | ENSMUSG00000067158 | 12829 | -0.704741467429958 | 0.776905540931709 | -2.94195538389746 | 0.0111353785382 | 0.0271516446133743 |
| Inpp5f | ENSMUSG00000042105 | 101490 | 0.382438932952186 | 2.42249579656602 | 2.94140996311073 | 0.0111471985304 | 0.0271744039109219 |
| Eftud2 | ENSMUSG00000020929 | 20624 | 0.188183527962269 | 4.42290781712649 | 2.94095935266697 | 0.0111569732033 | 0.0271921681350541 |
| Rusc2 | ENSMUSG00000035969 | 100213 | 0.217286382373435 | 4.86530052813433 | 2.94048952358175 | 0.0111671737820 | 0.0272109622407538 |
| Sap30l | ENSMUSG00000020519 | 50724 | -0.226592312407095 | 4.02057367214126 | -2.93980428662429 | 0.0111820676498 | 0.0272411815199424 |
| Ube4b | ENSMUSG00000028960 | 63958 | 0.170761654052386 | 5.42087782201972 | 2.93755932093313 | 0.0112310003099 | 0.0273542924215307 |
| Atp1b2 | ENSMUSG00000041329 | 11932 | 0.287358299059116 | 4.14758992558367 | 2.93662708664136 | 0.0112513819939 | 0.027397829256059 |
| Gnl2 | ENSMUSG00000028869 | 230737 | -0.187909538840167 | 4.50027498294909 | -2.93593544202013 | 0.0112665271022 | 0.0274285986089854 |
| Zfp518b | ENSMUSG00000046572 | 100515 | 0.443968680426962 | 2.15958876134891 | 2.93435783547555 | 0.0113011478112 | 0.0275067572310864 |
| Lin52 | ENSMUSG00000085793 | 217708 | -0.473815038360071 | 1.96148434510954 | -2.93408170998904 | 0.0113072181995 | 0.0275154056409427 |
| Zfp970 | ENSMUSG00000078866 | 628308 | -0.323056537019129 | 3.60235411137834 | -2.93392800290635 | 0.0113105987159 | 0.0275175060237119 |
| Tmem222 | ENSMUSG00000028857 | 52174 | -0.322297274235074 | 2.67672097034131 | -2.93222667993992 | 0.0113480830217 | 0.0276025579686359 |
| Ifi27 | ENSMUSG00000064215 | 52668 | 0.439746597397702 | 3.32649975188465 | 2.93064604459041 | 0.0113830181599 | 0.0276813729714656 |
| Calhm2 | ENSMUSG00000033033 | 72691 | 0.59937228278461 | 1.27573932936345 | 2.93040183317632 | 0.0113884251702 | 0.027688361996553 |
| Cnpy2 | ENSMUSG00000025381 | 56530 | 0.259753171622932 | 3.67323296608134 | 2.92979800145261 | 0.0114018053098 | 0.0277147284504348 |
| Suds3 | ENSMUSG00000066900 | 71954 | -0.224419166039565 | 6.15119873584454 | -2.92925482097707 | 0.0114138547511 | 0.0277378493296189 |
| Pisd | ENSMUSG00000023452 | 320951 | 0.244715644466993 | 3.3631446773903 | 2.92906721681953 | 0.0114180193172 | 0.0277418024354866 |
| Zfp53 | ENSMUSG00000057409 | 24132 | 0.554401577689286 | 1.14111860551405 | 2.92827136720757 | 0.0114357028164 | 0.0277785927748627 |
| Gtpbp8 | ENSMUSG00000022668 | 66067 | -0.356575468464915 | 2.44088104465396 | -2.92770315641626 | 0.0114483447965 | 0.0278031230772925 |
| Ufm1 | ENSMUSG00000027746 | 67890 | 0.197847298643596 | 5.43083607386916 | 2.92698011456101 | 0.0114644515181 | 0.02783605498548 |
| Dnm2 | ENSMUSG00000033335 | 13430 | 0.164652782920133 | 5.41437394144244 | 2.92622116721011 | 0.0114813821393 | 0.0278709722773849 |
| Inpp5b | ENSMUSG00000028894 | 16330 | -0.343837514496027 | 2.97105669596992 | -2.9249857969345 | 0.0115089936015 | 0.0279317959720927 |
| Akt1 | ENSMUSG00000001729 | 11651 | -0.183025157078835 | 5.6971239920594 | -2.9240802077432 | 0.0115292758186 | 0.027974808873149 |
| Lmcd1 | ENSMUSG00000057604 | 30937 | 0.300794907789824 | 8.49610839278064 | 2.92365355161838 | 0.0115388437302 | 0.0279918110997474 |
| Taldo1 | ENSMUSG00000025503 | 21351 | 0.250534591784053 | 5.20177593291414 | 2.92247673052591 | 0.0115652749779 | 0.0280497050773561 |
| Fkrp | ENSMUSG00000048920 | 243853 | -0.230725418952533 | 4.47937511736644 | -2.92099913550674 | 0.0115985461707 | 0.0281241588713842 |
| Sbno1 | ENSMUSG00000038095 | 243272 | -0.18439759597526 | 6.11423963890497 | -2.92053755736684 | 0.0116089589278 | 0.0281431647905619 |
| Pygo2 | ENSMUSG00000047824 | 68911 | 0.325622007163278 | 3.0267248529113 | 2.91932043162059 | 0.0116364603519 | 0.0281987825424801 |
| Vps4a | ENSMUSG00000031913 | 116733 | -0.21402473665717 | 5.1575457757493 | -2.91929390374015 | 0.0116370604747 | 0.0281987825424801 |
| Ripor1 | ENSMUSG00000038604 | 75687 | 0.256091959713083 | 5.0465887172924 | 2.91907746176525 | 0.0116419580400 | 0.0282043979024803 |
| Endod1 | ENSMUSG00000037419 | 71946 | 0.528166999717833 | 2.97084504543733 | 2.91836486569978 | 0.0116580967600 | 0.0282372381306571 |
| Pck1 | ENSMUSG00000027513 | 18534 | 1.36454115803909 | 0.217933333558868 | 2.9180961237573 | 0.0116641888980 | 0.028243562246344 |
| Tex261 | ENSMUSG00000014748 | 21766 | -0.194569954057268 | 4.42174053729788 | -2.91802175355545 | 0.0116658753583 | 0.028243562246344 |
| Sec22c | ENSMUSG00000061536 | 215474 | 0.413330810787224 | 1.89900277388435 | 2.91785563377951 | 0.0116696432520 | 0.0282464283409333 |
| Ctdnep1 | ENSMUSG00000018559 | 67181 | -0.201953239489829 | 5.29177472405996 | -2.9176875289502 | 0.0116734573922 | 0.0282494050818858 |
| Ccdc43 | ENSMUSG00000020925 | 52715 | -0.230388595097116 | 5.06446709376619 | -2.91669192132833 | 0.0116960720125 | 0.0282978670139915 |
| Vmn1r81 | ENSMUSG00000115027 | 171244 | -1.17752924882853 | 4.0065499332575 | -2.91608987472225 | 0.0117097680952 | 0.0283247344653173 |
| Olfr46 | ENSMUSG00000093942 | 18345 | -1.4349009139904 | 4.36118974901032 | -2.9157650741104 | 0.0117171636194 | 0.0283311994009025 |
| Ube2l3 | ENSMUSG00000038965 | 22195 | -0.178238378097681 | 5.89772550574552 | -2.91574484329322 | 0.0117176244160 | 0.0283311994009025 |
| Bri3bp | ENSMUSG00000037905 | 76809 | 0.353464398635347 | 2.63529766731298 | 2.91491503656892 | 0.0117365403006 | 0.0283706594486169 |
| Gmppa | ENSMUSG00000033021 | 69080 | 0.303421644122795 | 3.31293823227157 | 2.91449509086326 | 0.0117461246603 | 0.0283875500026799 |

| | | | | | | | |
|---|---|---|---|---|---|---|---|
| Card6 | ENSMUSG00000041849 | 239319 | 0.365545666677861 | 2.17093915218475 | 2.91401871055307 | 0.0117570063640 | 0.0284074726699845 |
| Acly | ENSMUSG00000020917 | 104112 | 0.245655576036139 | 3.97301256450156 | 2.91390672619305 | 0.0117595658065 | 0.0284074726699845 |
| Slc25a44 | ENSMUSG00000050144 | 229517 | -0.238386593045381 | 3.80109646952297 | -2.91369071827096 | 0.0117645042970 | 0.0284131233917742 |
| Vps72 | ENSMUSG00000008958 | 21427 | -0.215037228304491 | 4.07930410221366 | -2.91342895475492 | 0.0117704916180 | 0.0284213041476694 |
| Aarsd1 | ENSMUSG00000075528 | 69684 | -0.226880575073232 | 3.91319878292265 | -2.91281963668807 | 0.0117844402010 | 0.0284487004941103 |
| Tiam1 | ENSMUSG00000002489 | 21844 | 0.487458049466892 | 2.6435396058729 | 2.91164777128161 | 0.0118113125247 | 0.0285072769283105 |
| Spag5 | ENSMUSG00000002055 | 54141 | 0.631161550654324 | 1.20490869121967 | 2.91079553671466 | 0.0118308932616 | 0.0285439240552608 |
| Zfp512 | ENSMUSG00000062761 | 269639 | -0.242347168248927 | 3.52156158237768 | -2.91075962970853 | 0.0118317189547 | 0.0285439240552608 |
| Fbf1 | ENSMUSG00000020776 | 217335 | 0.33385110992548 | 3.30795122944874 | 2.91059446708941 | 0.0118355176539 | 0.0285467880572996 |
| Zbtb25 | ENSMUSG00000056459 | 109929 | 0.429569194814942 | 1.83186327673222 | 2.91011450479664 | 0.0118465634942 | 0.0285671267494985 |
| Haghl | ENSMUSG00000061046 | 68977 | 0.483841284501229 | 1.62500402364004 | 2.90550505584509 | 0.0119531641276 | 0.0288178290868324 |
| Ankmy2 | ENSMUSG00000036188 | 217473 | -0.436875046885393 | 2.48539196048835 | -2.90515360004341 | 0.0119613307799 | 0.0288311591523823 |
| Kin | ENSMUSG00000037262 | 16588 | -0.365060014223519 | 1.98044720341026 | -2.90415972232247 | 0.0119844549417 | 0.0288805284322044 |
| Fam76a | ENSMUSG00000028878 | 230789 | -0.223264253314168 | 4.30869034812649 | -2.90380218723953 | 0.0119927843431 | 0.0288942309136046 |
| Arl5a | ENSMUSG00000036093 | 75423 | 0.211927385121149 | 5.4721537921551 | 2.90330403638036 | 0.0120043991485 | 0.0289095924816944 |
| Kcna5 | ENSMUSG00000045534 | 16493 | -0.58415886304779 | 2.30330864335735 | -2.90330186693542 | 0.0120044497552 | 0.0289095924816944 |
| Nr1h3 | ENSMUSG00000002108 | 22259 | -0.428787344461764 | 2.14441779286562 | -2.90256228994429 | 0.0120217141387 | 0.0289447923460356 |
| Vmn1r55 | ENSMUSG00000118215 | 384522 | -1.24021083011384 | 3.67656370365388 | -2.90229836723496 | 0.0120278809660 | 0.0289476216067321 |
| Zfp282 | ENSMUSG00000025821 | 101095 | 0.279093675615638 | 3.18926322880934 | 2.90228533109138 | 0.0120281856497 | 0.0289476216067321 |
| Stk4 | ENSMUSG00000018209 | 58231 | 0.252517889689979 | 3.94863732799949 | 2.90073408415559 | 0.0120644961184 | 0.0290286170086489 |
| Gabpa | ENSMUSG00000008976 | 14390 | -0.18996554001333 | 5.32161943353611 | -2.90022625845097 | 0.0120764063919 | 0.0290508799010699 |
| Ankrd13b | ENSMUSG00000037907 | 268445 | 0.253634179831959 | 3.61296991551809 | 2.89946355564654 | 0.0120943161921 | 0.0290875622214939 |
| Prkab1 | ENSMUSG00000029513 | 19079 | 0.322953074215486 | 2.1972017337988 | 2.89931571588859 | 0.0120977907953 | 0.0290895185183697 |
| Copa | ENSMUSG00000026553 | 12847 | 0.1788447242546 | 6.75507725006192 | 2.89681260962941 | 0.0121567695870 | 0.0292249061702124 |
| Babam1 | ENSMUSG00000031820 | 68251 | -0.191738116618467 | 4.69942903612543 | -2.89613325922621 | 0.0121728254297 | 0.0292570700906104 |
| Spag7 | ENSMUSG00000018287 | 216873 | -0.232760829494768 | 5.53621734449988 | -2.89576724113969 | 0.0121814846013 | 0.0292714460711034 |
| Pias4 | ENSMUSG00000004934 | 59004 | -0.202877190455558 | 3.48500677761884 | -2.89540549114769 | 0.0121900487629 | 0.0292855874786565 |
| Prss23 | ENSMUSG00000039405 | 76453 | 0.251205594296775 | 5.24301509406528 | 2.89511419326123 | 0.0121969493352 | 0.0292947093634689 |
| Pag1 | ENSMUSG00000027508 | 94212 | 0.553299404609533 | 1.37786491644151 | 2.89501898111356 | 0.0121992056557 | 0.0292947093634689 |
| Cap1 | ENSMUSG00000028656 | 12331 | 0.333382500836393 | 4.77081256318422 | 2.89460747325324 | 0.0122089622263 | 0.0293116991206249 |
| Vmn1r218 | ENSMUSG00000115020 | 171256 | -1.21712606933624 | 2.79392447162295 | -2.89328695654333 | 0.0122403227245 | 0.0293805374839289 |
| Mrps16 | ENSMUSG00000049960 | 66242 | -0.27507167522453 | 2.95595878566597 | -2.89272021319571 | 0.0122538064453 | 0.0294022152914928 |
| Zdhhc8 | ENSMUSG00000060166 | 27801 | 0.268417927844713 | 3.96682524337372 | 2.89268126634847 | 0.0122547335896 | 0.0294022152914928 |
| Pik3r3 | ENSMUSG00000028698 | 18710 | 0.360977750078376 | 3.2392650151117 | 2.89252703659591 | 0.0122584057653 | 0.0294045717404704 |
| Snx5 | ENSMUSG00000027423 | 69178 | 0.187001936991126 | 5.38301811900881 | 2.8923281127648 | 0.0122631436979 | 0.0294094830544726 |
| Cnep1r1 | ENSMUSG00000036810 | 382030 | -0.268107180333817 | 3.45723405812304 | -2.89197440649638 | 0.0122715726673 | 0.0294232421204249 |
| Acp1 | ENSMUSG00000044573 | 11431 | -0.292035811781074 | 4.50072787330951 | -2.89179249795619 | 0.0122759098467 | 0.029425748407813 |
| Cry2 | ENSMUSG00000068742 | 12953 | 0.340166663920074 | 4.61252459214828 | 2.89170477746334 | 0.0122780018743 | 0.029425748407813 |
| Usp9x | ENSMUSG00000031010 | 22284 | 0.208004732938791 | 7.47902385596448 | 2.89096800290421 | 0.0122955869242 | 0.0294614337284642 |
| U2surp | ENSMUSG00000032407 | 67958 | -0.179961318518179 | 5.0538419166773 | -2.89074103270473 | 0.0123010091582 | 0.0294679664933584 |
| Cspp1 | ENSMUSG00000056763 | 211660 | 0.363093979077831 | 2.41278752114915 | 2.89029917604864 | 0.0123115717072 | 0.0294868076976066 |
| Tor2a | ENSMUSG00000009563 | 30933 | 0.472989956671337 | 1.22507808133682 | 2.89001939043745 | 0.0123182645796 | 0.0294963746156071 |
| 4930523C07Rik | ENSMUSG00000090394 | 67647 | -0.427879142635515 | 2.7533976407202 | -2.88972033819262 | 0.0123254223002 | 0.0295070501891934 |
| Hnrnpa0 | ENSMUSG00000007836 | 77134 | 0.252144484772267 | 4.52391743941078 | 2.88725483461446 | 0.0123845896700 | 0.0296422048792274 |
| Mindy3 | ENSMUSG00000026767 | 66960 | 0.323859704505296 | 1.81661637117672 | 2.88665777476866 | 0.0123989599875 | 0.0296701032451875 |
| Mlst8 | ENSMUSG00000024142 | 56716 | -0.276226587502608 | 2.81992945654138 | -2.88527807453425 | 0.0124322301122 | 0.0297432058123214 |
| Timmdc1 | ENSMUSG00000002846 | 76916 | -0.247044753562131 | 4.49693351298205 | -2.88516311274262 | 0.0124350062683 | 0.0297433377504376 |
| Vmn1r91 | ENSMUSG00000095201 | 667067 | -2.30850099500632 | -1.51277903080305 | -2.88372149308297 | 0.0124698711080 | 0.0298202058809123 |
| Sp4 | ENSMUSG00000025323 | 20688 | -0.280847899887559 | 3.29578949542205 | -2.8828718444984 | 0.0124904644939 | 0.0298629193752212 |
| Dpp10 | ENSMUSG00000036815 | 269109 | -1.36191812115145 | 1.39665381534338 | -2.88209824933113 | 0.0125092436563 | 0.0299012775874547 |
| Fkbp14 | ENSMUSG00000038074 | 231997 | 0.417678417993827 | 1.83660136178991 | 2.88165671046143 | 0.0125199745512 | 0.0299203851814989 |
| Ubl3 | ENSMUSG00000001687 | 24109 | -0.177985549130528 | 6.41753265523999 | -2.88138883748596 | 0.0125264891917 | 0.0299294105694197 |
| Polg | ENSMUSG00000039176 | 18975 | -0.18999525518209 | 4.79921521217259 | -2.8804722271597 | 0.0125488063112 | 0.0299761804606382 |
| Myorg | ENSMUSG00000046312 | 329828 | -0.356860993138303 | 4.76667328576517 | -2.87883412078714 | 0.0125887875491 | 0.03006511616774 |
| Psg16 | ENSMUSG00000066760 | 26436 | -0.272530336800466 | 4.85728994967975 | -2.87860007102006 | 0.0125945102278 | 0.0300694463788943 |
| Hacd3 | ENSMUSG00000033629 | 57874 | 0.289751757359139 | 2.98246931616719 | 2.87853497333236 | 0.0125961023665 | 0.0300694463788943 |
| Cdc42bpa | ENSMUSG00000026490 | 226751 | 0.211556759039391 | 5.67032822301549 | 2.87789564638807 | 0.0126117493552 | 0.0301002253717372 |
| Zfp862-ps | ENSMUSG00000107476 | 58894 | 0.394522065473886 | 2.12441611688931 | 2.87657997861973 | 0.0126440094437 | 0.0301706324448339 |
| Ehbp1l1 | ENSMUSG00000024937 | 114601 | 0.239762875478114 | 8.39187300980583 | 2.8757859828788 | 0.0126635174561 | 0.030210586929978 |
| Ndufaf7 | ENSMUSG00000024082 | 73694 | -0.310407904993679 | 3.02608744211116 | -2.87430205760329 | 0.0127000560921 | 0.0302911440416533 |
| Akip1 | ENSMUSG00000031023 | 57373 | 0.467162961073954 | 1.57389625091511 | 2.87407247746227 | 0.0127057182964 | 0.0302980381104291 |
| Fxyd1 | ENSMUSG00000036570 | 56188 | -0.217381355836968 | 6.67466657667817 | -2.87339488009715 | 0.0127224445772 | 0.0303313067991015 |
| Dlc1 | ENSMUSG00000031523 | 50768 | -0.454791521379499 | 5.31186666385226 | -2.87308953492433 | 0.0127299890095 | 0.0303426755044381 |
| Dlg4 | ENSMUSG00000020886 | 13385 | 0.579249784178921 | 2.26416705369395 | 2.87264932589565 | 0.0127408733922 | 0.0303616756446534 |
| Eps8l1 | ENSMUSG00000006154 | 67425 | -0.803041150707758 | 0.844007199828745 | -2.8725425243275 | 0.0127435154933 | 0.0303616756446534 |
| Snora74a | ENSMUSG00000065649 | 436583 | 0.857616104975663 | 2.47542704578878 | 2.87081091398046 | 0.0127864281000 | 0.0304572773068871 |
| Amer1 | ENSMUSG00000050332 | 72345 | 0.357401522693319 | 3.14414371693777 | 2.86986759961728 | 0.0128098650300 | 0.0305064563492465 |
| Rev3l | ENSMUSG00000019841 | 19714 | 0.358908570155538 | 5.03324900319965 | 2.86974151742979 | 0.0128130007809 | 0.0305072776162592 |
| Trappc9 | ENSMUSG00000047921 | 76510 | -0.164653712734538 | 4.55099102948748 | -2.86769564217604 | 0.0128639887993 | 0.0306220081806613 |
| Vmn2r59 | ENSMUSG00000092032 | 628444 | -1.53683133840377 | 2.69919707407331 | -2.86724481147584 | 0.0128752513748 | 0.0306421451727767 |
| Smim26 | ENSMUSG00000074754 | 228715 | -0.565143918467583 | 1.65195104851076 | -2.86533202861883 | 0.0129231441853 | 0.0307494316695383 |
| Pacsin2 | ENSMUSG00000016664 | 23970 | -0.18974899973156 | 6.29110933558271 | -2.86418212837355 | 0.0129520200259 | 0.0308114321878825 |
| Tax1bp3 | ENSMUSG00000040158 | 76281 | 0.354948683677506 | 3.0970695883537 | 2.86381282111458 | 0.0129613073826 | 0.0308268170147852 |
| Bcl6b | ENSMUSG00000000317 | 12029 | -0.95337065060788 | 2.31150470333906 | -2.86317030930206 | 0.0129774809162 | 0.0308585694790353 |
| Gas8 | ENSMUSG00000040220 | 104346 | 0.47471852016666 | 1.75482771002057 | 2.86292016866451 | 0.0129837829119 | 0.0308667331493968 |
| Armc5 | ENSMUSG00000042178 | 233912 | 0.309390367831756 | 2.75337869694626 | 2.86280991082559 | 0.0129865616827 | 0.0308667331493968 |
| Stard3nl | ENSMUSG00000003062 | 76205 | 0.416812647730723 | 1.71112585309827 | 2.86011893233795 | 0.0130545626613 | 0.0310216140112086 |
| Mettl5 | ENSMUSG00000051730 | 75422 | -0.34205827182971 | 2.48448837098103 | -2.85938228187542 | 0.0130732387825 | 0.0310592421500858 |
| 1190007I07Rik | ENSMUSG00000063320 | 544717 | -0.438471220382471 | 1.94602942250205 | -2.85892373377353 | 0.0130848775085 | 0.0310801382107118 |
| Vkorc1 | ENSMUSG00000096145 | 27973 | 0.55853909113639 | 1.49195542698505 | 2.85839332267081 | 0.0130983529500 | 0.0311053869425593 |
| Ahi1 | ENSMUSG00000019986 | 52906 | 0.583365875862123 | 1.02006836906801 | 2.85807901844911 | 0.0131063444933 | 0.0311176046169793 |
| Tpd52l1 | ENSMUSG00000000296 | 21987 | -0.288607262807767 | 4.1883480631349 | -2.85693946543466 | 0.0131353591553 | 0.0311770580822024 |
| Cenpv | ENSMUSG00000018509 | 73139 | -0.425165061340311 | 1.86778993359836 | -2.85687156917437 | 0.0131370898844 | 0.0311770580822024 |
| Olfml2a | ENSMUSG00000046618 | 241327 | 0.245045128672517 | 3.97651885261712 | 2.85654919412653 | 0.0131453105382 | 0.0311897958528459 |
| Mrps27 | ENSMUSG00000041632 | 218506 | -0.279087853830116 | 3.712518654725 | -2.85615000325681 | 0.0131554970273 | 0.0312071914074491 |
| Slc6a8 | ENSMUSG00000019558 | 102857 | -0.339883448804407 | 6.64725106470802 | -2.85573963445305 | 0.0131659768439 | 0.0312252750881632 |
| Tubgcp2 | ENSMUSG00000025474 | 74237 | -0.248212778551256 | 3.75997191255215 | -2.85558973658088 | 0.0131698069159 | 0.0312275833834872 |
| Asxl1 | ENSMUSG00000042548 | 228790 | 0.263028533363857 | 4.02725413269836 | 2.85515905924069 | 0.0131808173396 | 0.0312469126740156 |
| Bmp5 | ENSMUSG00000032179 | 12160 | -0.493236521402173 | 1.57566563325867 | -2.8540104540658 | 0.0132102261277 | 0.0313098399399129 |
| Dcbld2 | ENSMUSG00000035107 | 73379 | 0.41610536956732 | 3.36421647196405 | 2.85380708540474 | 0.0132154398799 | 0.0313154071813012 |
| Vipr1 | ENSMUSG00000032528 | 22354 | -0.788447009726788 | 0.936429775482296 | -2.85237276714699 | 0.0132522689193 | 0.0313958715988268 |
| Jakmip2 | ENSMUSG00000024502 | 76217 | -0.95558418397445 | 1.40484817097112 | -2.8518396137182 | 0.0132659844271 | 0.0314174756141257 |
| Peg13 | ENSMUSG00000106847 | 353342 | -0.330014812663907 | 3.88262203298776 | -2.85179486051531 | 0.0132671363493 | 0.0314174756141257 |
| Slc38a4 | ENSMUSG00000022464 | 69354 | -0.328287900754642 | 3.7376019636008 | -2.84957257818665 | 0.0133244604796 | 0.031546388889438 |
| Rb1cc1 | ENSMUSG00000025907 | 12421 | -0.210115126877083 | 6.60328239847032 | -2.84817777000998 | 0.0133605640693 | 0.0316250164231048 |
| Cdc123 | ENSMUSG00000039128 | 98828 | -0.238319218110345 | 3.83819074913308 | -2.84680175298867 | 0.0133962754755 | 0.0316959520770074 |
| Nkain1 | ENSMUSG00000078532 | 67149 | -0.337455618529746 | 2.13418956657567 | -2.84679960074151 | 0.0133963314057 | 0.0316959520770074 |
| Zfp322a | ENSMUSG00000046351 | 218100 | 0.326486674855608 | 3.47019542304591 | 2.84646103306793 | 0.0134051325705 | 0.0317099121679988 |
| Nrbp1 | ENSMUSG00000029148 | 192292 | -0.161123668950218 | 5.6468964693225 | -2.84633812443277 | 0.0134083290218 | 0.031710611107321 |
| Cops5 | ENSMUSG00000025917 | 26754 | -0.238224541927293 | 5.18115570004929 | -2.84543227909029 | 0.0134319102101 | 0.0317595090864803 |
| Ifnar1 | ENSMUSG00000022967 | 15975 | 0.307301326382274 | 3.99260764383499 | 2.84471864171728 | 0.0134505164912 | 0.0317902596289263 |
| Col4a2 | ENSMUSG00000031503 | 12827 | 0.409919763569197 | 7.70551477213855 | 2.84471038412376 | 0.0134507319352 | 0.0317902596289263 |
| Tinf2 | ENSMUSG00000007589 | 28113 | -0.329180894138637 | 2.79598672458355 | -2.84417378881659 | 0.0134647391950 | 0.0318164859795476 |
| Taf12 | ENSMUSG00000028899 | 66464 | -0.205837267472262 | 3.84972439590191 | -2.84351530833005 | 0.0134819477292 | 0.0318502638054724 |
| Haus1 | ENSMUSG00000041840 | 225745 | -0.363840443436493 | 2.45327851768935 | -2.84190078210242 | 0.0135242327878 | 0.0319432559645579 |
| Abhd11 | ENSMUSG00000040532 | 68758 | -0.250565037545988 | 3.5863779205057 | -2.84034901290629 | 0.0135649970161 | 0.0320187325506702 |
| Pdcd4 | ENSMUSG00000024975 | 18569 | 0.325955608430913 | 3.80616368467481 | 2.84034625557296 | 0.0135650695573 | 0.0320187325506702 |
| Gimap1 | ENSMUSG00000090019 | 16205 | -0.524056544697499 | 1.69775671807497 | -2.84032065577841 | 0.0135657430662 | 0.0320187325506702 |
| Josd1 | ENSMUSG00000022426 | 74158 | -0.327534283121044 | 3.90340298882066 | -2.84023849134117 | 0.0135679049652 | 0.0320187325506702 |
| Plod3 | ENSMUSG00000004846 | 26433 | 0.4070495348838 | 3.65178495670936 | 2.83844576780641 | 0.0136151592010 | 0.0321201338756987 |
| Hnrnpab | ENSMUSG00000020358 | 15384 | 0.159475458814557 | 6.29979132689608 | 2.83838550150506 | 0.0136167505607 | 0.0321201338756987 |
| Pcdhga9 | ENSMUSG00000102440 | 93717 | 1.042283152397 | 0.30999828129964 | 2.83790046275755 | 0.0136295648894 | 0.0321434247695995 |
| Nudt9 | ENSMUSG00000029310 | 74167 | -0.221994258619411 | 4.41509811946183 | -2.83745827006798 | 0.0136412575827 | 0.0321640609656346 |
| Ppp1r37 | ENSMUSG00000051403 | 232947 | -0.210303133034816 | 5.41591558167323 | -2.83557549283758 | 0.0136911533675 | 0.0322747460558633 |
| Sdhaf3 | ENSMUSG00000042505 | 71238 | -0.372550845909746 | 2.04492110011226 | -2.834876477911 | 0.0137097236715 | 0.0323115544315178 |
| Slc33a1 | ENSMUSG00000027822 | 11416 | 0.427848852034419 | 2.35406537391616 | 2.83372602237918 | 0.0137403410018 | 0.0323767336691357 |

| | | | | | | | |
|---|---|---|---|---|---|---|---|
| Atg16l2 | ENSMUSG00000047767 | 73683 | 0.39649928104236 | 1.74695553874705 | 2.83295309690544 | 0.0137609487447 | 0.0324183040363295 |
| Immp2l | ENSMUSG00000056899 | 93757 | -0.440912406315344 | 1.66681915512275 | -2.83270386514223 | 0.0137676002329 | 0.0324269851639629 |
| Pigh | ENSMUSG00000021120 | 110417 | 0.477815148156925 | 1.33630282124466 | 2.83254408662632 | 0.0137718660569 | 0.0324300447798972 |
| Kansl2 | ENSMUSG00000022992 | 69612 | -0.283966635611578 | 4.49241184736067 | -2.83219860806095 | 0.0137810942055 | 0.0324447858626636 |
| Apbb2 | ENSMUSG00000029207 | 11787 | -0.191926106434834 | 4.98557889509989 | -2.83161991766809 | 0.0137965653295 | 0.0324644531177907 |
| Ptpn21 | ENSMUSG00000021009 | 24000 | 0.230294195573034 | 5.84150116874725 | 2.83156496830661 | 0.0137980352715 | 0.0324644531177907 |
| Casp12 | ENSMUSG00000025887 | 12364 | 0.313598376800206 | 3.13827417231583 | 2.83155291206841 | 0.0137983578067 | 0.0324644531177907 |
| Ascc3 | ENSMUSG00000038774 | 77987 | 0.208867647945568 | 4.50204268051075 | 2.831043912906 | 0.0138119815911 | 0.0324895138311519 |
| Arfip1 | ENSMUSG00000102805 | 99889 | 0.270278986406046 | 2.76405119470797 | 2.83086274847605 | 0.0138168337940 | 0.0324939350693962 |
| Ago4 | ENSMUSG00000042500 | 76850 | -0.31758369895911 | 2.89835032899286 | -2.83049985307232 | 0.0138265584089 | 0.0325098107051241 |
| Rrp9 | ENSMUSG00000041506 | 27966 | -0.228666026770925 | 3.24426643728818 | -2.82985750078846 | 0.0138437881918 | 0.0325410597315565 |
| Tmem134 | ENSMUSG00000024845 | 66990 | -0.242011447571683 | 4.33447313451742 | -2.82978244986444 | 0.0138458026540 | 0.0325410597315565 |
| Dr1 | ENSMUSG00000029265 | 13486 | 0.242351914411036 | 3.72465342764849 | 2.82962729852773 | 0.0138499680256 | 0.032543852211713 |
| Ep400 | ENSMUSG00000029505 | 75560 | 0.172243456214634 | 5.53952500564626 | 2.82876635026988 | 0.0138731044026 | 0.0325912108801931 |
| Sgk1 | ENSMUSG00000019970 | 20393 | -0.646991015793503 | 4.37931079165095 | -2.82797026044378 | 0.0138945315960 | 0.0326345347822509 |
| Tle4 | ENSMUSG00000024642 | 21888 | -0.299778371122204 | 3.87438583043182 | -2.8275984933134 | 0.0139045490387 | 0.0326510473776146 |
| Ppp1r18 | ENSMUSG00000034595 | 76448 | 0.42670387129744 | 3.37783102276992 | 2.82608175615873 | 0.0139454917721 | 0.0327401569074297 |
| Mcam | ENSMUSG00000032135 | 84004 | -0.345725028446747 | 3.84180509425967 | -2.82582315560003 | 0.0139524842240 | 0.0327495394144782 |
| Naa15 | ENSMUSG00000063273 | 74838 | -0.204626819369217 | 5.52233935391227 | -2.82533721994887 | 0.0139656330346 | 0.032773365114177 |
| 4930470H14Rik | ENSMUSG00000116975 | 75815 | -0.893882711911054 | 1.28738578735798 | -2.82488983815587 | 0.0139777493807 | 0.0327947582058541 |
| Sumo2 | ENSMUSG00000020738 | 170930 | -0.200691668367875 | 5.1106562740558 | -2.82411301125971 | 0.0139988125384 | 0.0328371287249042 |
| Uso1 | ENSMUSG00000029407 | 56041 | 0.175163881267791 | 6.56473916896647 | 2.82351566571293 | 0.0140150303634 | 0.0328681177649273 |
| Ryr3 | ENSMUSG00000057378 | 20192 | 0.641860124402593 | 1.33464958998164 | 2.82314178144175 | 0.0140251906374 | 0.0328848903598731 |
| Stag1 | ENSMUSG00000037286 | 20842 | -0.167361602420887 | 4.98364833554442 | -2.8230071069942 | 0.0140288521752 | 0.0328864214299016 |
| Papolg | ENSMUSG00000020273 | 216578 | -0.322617442570738 | 2.5949178702029 | -2.82256288161618 | 0.0140409364573 | 0.0329076921988758 |
| Ly6c1 | ENSMUSG00000079018 | 17067 | -0.370480486572995 | 4.77618841380069 | -2.82145336640313 | 0.0140711632922 | 0.0329714653692857 |
| Cdk2ap1 | ENSMUSG00000029394 | 13445 | 0.224209203253208 | 5.66696001241159 | 2.82031628570009 | 0.0141022073659 | 0.0330371257429864 |
| Cfap36 | ENSMUSG00000020462 | 216618 | 0.338726740737382 | 3.02803334067892 | 2.81939926466026 | 0.0141272924228 | 0.0330888008301474 |
| Arsk | ENSMUSG00000021592 | 77041 | 0.341709740688645 | 1.96028147036278 | 2.81789897034294 | 0.0141684273435 | 0.0331736429874897 |
| Plekhf1 | ENSMUSG00000074170 | 72287 | -0.314527810857392 | 3.12810996870277 | -2.81785679132145 | 0.0141695855007 | 0.0331736429874897 |
| Lamb2 | ENSMUSG00000052911 | 16779 | 0.19893452960548 | 6.94558988772675 | 2.81771125377673 | 0.0141735824035 | 0.0331758949149409 |
| Nkd1 | ENSMUSG00000031661 | 93960 | -0.797177586980397 | 0.451962207401629 | -2.81720356857338 | 0.0141875336429 | 0.0332014408587737 |
| Tasor2 | ENSMUSG00000033799 | 105203 | 0.201461467866666 | 4.22331430118412 | 2.81668899986284 | 0.0142016877919 | 0.0332274506107452 |
| Bclaf1 | ENSMUSG00000037608 | 72567 | -0.195316447351534 | 5.75933358628905 | -2.81570840603832 | 0.0142286991808 | 0.0332835246621323 |
| Fbxl19 | ENSMUSG00000030811 | 233902 | 0.547686510046307 | 1.00967983335782 | 2.81548440113525 | 0.0142348766781 | 0.0332840216710695 |
| Gpr4 | ENSMUSG00000044317 | 319197 | -0.933103359297077 | 0.801039488704243 | -2.81541690503443 | 0.0142367385683 | 0.0332840216710695 |
| Tmed1 | ENSMUSG00000032180 | 17083 | -0.263168475098987 | 3.41952212437966 | -2.81536949905295 | 0.0142380464123 | 0.0332840216710695 |
| Man2a2 | ENSMUSG00000038886 | 140481 | 0.212558770560833 | 6.43419474829001 | 2.81498928256251 | 0.0142485401527 | 0.0333008500480095 |
| Mrrf | ENSMUSG00000026887 | 67871 | -0.275787500804597 | 3.51620071737928 | -2.81488795260286 | 0.0142513380774 | 0.0333008500480095 |
| Stx5a | ENSMUSG00000010110 | 56389 | 0.311261505869171 | 3.41338216239604 | 2.81465396845226 | 0.0142578009117 | 0.0333088313243031 |
| Swt1 | ENSMUSG00000052748 | 66875 | -0.368145206272651 | 2.82413898867045 | -2.8144386026523 | 0.0142637520332 | 0.0333122730914932 |
| Fbxo7 | ENSMUSG00000001786 | 69754 | -0.352299845846099 | 2.05331609049486 | -2.81438009532838 | 0.0142653691648 | 0.0333122730914932 |
| Fam219a | ENSMUSG00000028439 | 71901 | -0.183989215827147 | 3.85780884321475 | -2.81364150169973 | 0.0142857992306 | 0.0333512040318423 |
| Lrp5 | ENSMUSG00000024913 | 16973 | -0.225701783643305 | 5.56720195481161 | -2.8135568441078 | 0.0142881427573 | 0.0333512040318423 |
| Ripk1 | ENSMUSG00000021408 | 19766 | 0.370284889612027 | 3.05930700637527 | 2.81307227456615 | 0.0143015640728 | 0.033375404970222 |
| Senp6 | ENSMUSG00000034252 | 215351 | -0.184189413434039 | 5.49189411364559 | -2.81185398251376 | 0.0143353622527 | 0.0334471386947977 |
| Hdhd5 | ENSMUSG00000058979 | 214932 | -0.281577965500242 | 3.11713423752053 | -2.81159058036688 | 0.0143426799198 | 0.0334570708928252 |
| Olfr152 | ENSMUSG00000068816 | 258640 | -1.38325507682775 | 0.42909728772456 | -2.81119863122537 | 0.0143535755792 | 0.0334753434301555 |
| Ttc27 | ENSMUSG00000024078 | 74196 | -0.28243675417869 | 2.27468061403451 | -2.80835584271482 | 0.0144328446370 | 0.0336530342848948 |
| Vash1 | ENSMUSG00000021256 | 238328 | 0.38474914935865 | 3.75638298217291 | 2.80754818150031 | 0.0144554438892 | 0.0336985406596466 |
| Fntb | ENSMUSG00000033373 | 110606 | 0.297536924811428 | 2.61965762446657 | 2.80738428039674 | 0.0144600342571 | 0.0337020542569591 |
| Arvcf | ENSMUSG00000000325 | 11877 | 0.323936797833461 | 3.98954630091475 | 2.80655216217142 | 0.0144833614020 | 0.0337492269208731 |
| Herpud2 | ENSMUSG00000008429 | 80517 | 0.162143769455556 | 4.98363939953446 | 2.806379758838 | 0.0144881990754 | 0.0337512965393781 |
| Igf2bp2 | ENSMUSG00000033581 | 319765 | 0.784256088927099 | 1.43754744832288 | 2.80630045438321 | 0.0144904249070 | 0.0337512965393781 |
| Ctnna3 | ENSMUSG00000060843 | 216033 | -0.296922465502277 | 3.25387200299645 | -2.80518032609757 | 0.0145218992962 | 0.033817401194679 |
| 8030462N17Rik | ENSMUSG00000047466 | 212163 | -0.198058388138314 | 5.19756235867166 | -2.80319632233454 | 0.0145778122574 | 0.0339403760991933 |
| Sgf29 | ENSMUSG00000030714 | 75565 | -0.402698911680234 | 2.47038479044694 | -2.80206856901077 | 0.0146096885055 | 0.0340073477543284 |
| Wars | ENSMUSG00000021266 | 22375 | -0.233787446048582 | 4.52843925742016 | -2.80122018533613 | 0.0146337133280 | 0.0340560188182785 |
| Smim13 | ENSMUSG00000091264 | 108934 | -0.214656324038554 | 5.1344162958195 | -2.80012362326227 | 0.0146648235449 | 0.0341211549956891 |
| Fstl3 | ENSMUSG00000020325 | 83554 | 0.551564150035491 | 1.37587373425764 | 2.79971843400567 | 0.0146763354393 | 0.0341342004418636 |
| Olfr1220 | ENSMUSG00000101480 | 258902 | -1.42538553637161 | 0.0123372529163708 | -2.79970646244118 | 0.0146766757000 | 0.0341342004418636 |
| Arhgdib | ENSMUSG00000030220 | 11857 | 0.323936789049081 | 2.4994160337073 | 2.79956195627904 | 0.0146807835229 | 0.0341364911059655 |
| Apbb3 | ENSMUSG00000117679 | 225372 | -0.401354939630768 | 1.50029439447301 | -2.79921507541609 | 0.0146906487767 | 0.0341521654143558 |
| Trit1 | ENSMUSG00000028653 | 66966 | 0.422784086372686 | 1.86751773260101 | 2.79891710465504 | 0.0146991282210 | 0.0341646120740841 |
| Gm33180 | ENSMUSG00000111852 | 102635981 | -1.18839586527103 | 0.242698201211276 | -2.79809146173154 | 0.0147226488649 | 0.0342065168642287 |
| Ppp2r2a | ENSMUSG00000022052 | 71978 | -0.206818665536662 | 5.25416898356022 | -2.79806454986748 | 0.0147234161418 | 0.0342065168642287 |
| Tspan7 | ENSMUSG00000058254 | 21912 | -0.18054270580935 | 4.94611281374954 | -2.7971768255107 | 0.0147487477923 | 0.0342580879979942 |
| Arpc3 | ENSMUSG00000029465 | 56378 | 0.247681144080333 | 4.20901780216533 | 2.79592904377527 | 0.0147844260928 | 0.0343336650989909 |
| Ltv1 | ENSMUSG00000019814 | 353258 | -0.21152347639639 | 4.02686249531211 | -2.79490332111311 | 0.0148138183112 | 0.0343946151147125 |
| Nedd4 | ENSMUSG00000032216 | 17999 | 0.169705526665206 | 8.60701486978002 | 2.79468349134364 | 0.0148201250102 | 0.0344019508361336 |
| Map3k8 | ENSMUSG00000024235 | 26410 | 0.542207715333262 | 1.46396982116009 | 2.79384584938448 | 0.0148441802556 | 0.0344504743895516 |
| Snx19 | ENSMUSG00000031993 | 102607 | -0.201559002262742 | 4.82341347785 | -2.79339141102463 | 0.0148572467379 | 0.0344734799602977 |
| Ddx27 | ENSMUSG00000017999 | 228889 | 0.207665829382877 | 3.7484171046958 | 2.79291051592559 | 0.0148710862064 | 0.0344982689564475 |
| Ttc4 | ENSMUSG00000025413 | 72354 | -0.213250173069361 | 3.82422829477228 | -2.79234090964327 | 0.0148874949988 | 0.0345290065420787 |
| A630089N07Rik | ENSMUSG00000116673 | 320586 | -1.03406516099005 | 2.94298718629066 | -2.79104407586901 | 0.0149249194233 | 0.0346084629224186 |
| Cep290 | ENSMUSG00000019971 | 216274 | -0.316489225811383 | 3.17373973566155 | -2.79090771092431 | 0.0149288600366 | 0.0346102585493321 |
| Ifngr1 | ENSMUSG00000020009 | 15979 | 0.419770912292259 | 4.55635584246785 | 2.79057484909713 | 0.0149384832108 | 0.0346252247620502 |
| Vmn1r87 | ENSMUSG00000070815 | 171261 | -1.28604349117453 | 1.47072487609917 | -2.7902753075236 | 0.0149471482782 | 0.0346379643479156 |
| Myl1 | ENSMUSG00000061816 | 17901 | 0.285721526554637 | 11.2310499119928 | 2.78996533963967 | 0.0149561201400 | 0.0346514093368337 |
| Rnf6 | ENSMUSG00000029634 | 74132 | -0.167993174702104 | 5.26381210617885 | -2.78975090109988 | 0.0149623300414 | 0.0346584508758706 |
| Nr4a1 | ENSMUSG00000023034 | 15370 | -1.01276936819255 | 4.40503166334838 | -2.78943833737687 | 0.0149713860638 | 0.03467208073381 5 |
| Syne3 | ENSMUSG00000054150 | 212073 | 0.344705748019073 | 3.01998608880772 | 2.78855709339842 | 0.0149969475897 | 0.0347239216487378 |
| Samd5 | ENSMUSG00000060487 | 320825 | -0.499263162310187 | 2.00557246593319 | -2.78703786115403 | 0.0150411151493 | 0.0348188118801189 |
| Gm572 | ENSMUSG00000070577 | 230909 | -0.504216295477356 | 1.11123423710181 | -2.78519499762573 | 0.0150948624691 | 0.0349358335061985 |
| Plekhm1 | ENSMUSG00000034247 | 353047 | 0.212714274520069 | 4.25111736629593 | 2.78493321590258 | 0.0151025126062 | 0.0349443530952003 |
| Taz | ENSMUSG00000009995 | 66826 | -0.199295118007147 | 3.9488610160201 | -2.78485027579901 | 0.0151049371855 | 0.0349443530952003 |
| Ccdc47 | ENSMUSG00000078622 | 67163 | -0.17020936571539 | 5.99519001994362 | -2.7842319105846 | 0.0151230258115 | 0.0349787971109331 |
| Rab13 | ENSMUSG00000027935 | 68328 | 0.628133720144565 | 1.10986513129946 | 2.78261395894355 | 0.0151704551143 | 0.0350810757044882 |
| Rpl23a | ENSMUSG00000058546 | 268449 | -0.241623703874769 | 7.533710492455 | -2.78242958934982 | 0.0151758690384 | 0.0350861726859546 |
| Hpgd | ENSMUSG00000031613 | 15446 | 0.678381231856106 | 1.01943644677547 | 2.78192812777685 | 0.0151906037893 | 0.0351128124384764 |
| Pigg | ENSMUSG00000029263 | 433931 | -0.356535234934095 | 2.51814703467176 | -2.78179937457088 | 0.0151943892832 | 0.0351141372631644 |
| Trim13 | ENSMUSG00000035235 | 66597 | 0.570154834277547 | 1.82967822956491 | 2.78018959906846 | 0.0152417966049 | 0.0352162499870214 |
| Zbtb7a | ENSMUSG00000035011 | 16969 | 0.193594318712178 | 5.32544781581916 | 2.7799895782047 | 0.0152476972564 | 0.0352224384426289 |
| Smarcc1 | ENSMUSG00000032481 | 20588 | 0.17124654210591 | 5.01467220692231 | 2.77965957172196 | 0.0152574373972 | 0.0352374916944536 |
| Dtna | ENSMUSG00000024302 | 13527 | 0.250874648260014 | 5.72854907010133 | 2.77942962714351 | 0.0152642278125 | 0.035245727549407 |
| Fam124a | ENSMUSG00000035184 | 629059 | 0.43372912581775 | 2.06208347390582 | 2.77793857057974 | 0.0153083315602 | 0.0353400997434117 |
| Kctd5 | ENSMUSG00000016946 | 69259 | 0.26026646934511 | 2.72304339562768 | 2.77758273296828 | 0.0153188752571 | 0.0353569732761547 |
| Ttll4 | ENSMUSG00000033257 | 67534 | 0.263428429467507 | 3.18553836773103 | 2.77725241124752 | 0.0153286692653 | 0.0353721097191283 |
| Uckl1 | ENSMUSG00000089917 | 68556 | 0.330117504288614 | 2.84663325501922 | 2.77673633012806 | 0.0153439832951 | 0.0353999749682692 |
| Hif1a | ENSMUSG00000021109 | 15251 | 0.217748615520115 | 5.03460101271929 | 2.77607196712534 | 0.0153637194495 | 0.0354380285509353 |
| LOC102634835 | ENSMUSG00000114627 | 102634835 | -0.551822968233515 | 1.30541641771044 | -2.77493140490056 | 0.0153976599714 | 0.0355088230269454 |
| Smarca4 | ENSMUSG00000032187 | 20586 | 0.184429211809617 | 6.53415023045191 | 2.77480635870887 | 0.0154013855218 | 0.0355099230413677 |
| 2610001J05Rik | ENSMUSG00000052419 | 66520 | 0.281382210988606 | 2.8889070577089 | 2.77357381456659 | 0.0154381544408 | 0.0355871923646177 |
| Naa40 | ENSMUSG00000024764 | 70999 | -0.271494502329007 | 3.49043610386009 | -2.77318860864085 | 0.0154496634123 | 0.0356062135273197 |
| Pten | ENSMUSG00000013663 | 19211 | 0.182657058611027 | 6.4428982809502 | 2.77037245644138 | 0.0155340583654 | 0.0357931686325299 |
| Mir99ahg | ENSMUSG00000090386 | 77994 | 0.604342796766764 | 1.8899913738321 | 2.77019733666754 | 0.0155393212616 | 0.0357976957831454 |
| Txn1 | ENSMUSG00000028367 | 22166 | 0.32763002687273 | 3.46102720371453 | 2.77008916988237 | 0.0155425728832 | 0.0357976957831454 |
| Zfp106 | ENSMUSG00000027288 | 20402 | -0.261015525313892 | 10.5866711225297 | -2.76955596132908 | 0.0155586114977 | 0.0358270870615906 |
| Zfyve1 | ENSMUSG00000042628 | 217695 | 0.234812082370021 | 4.30622804399378 | 2.76907970318817 | 0.0155729507694 | 0.0358525536774875 |
| Zfp280b | ENSMUSG00000049764 | 64453 | -0.309532820769781 | 2.66834074590879 | -2.7678167834278 | 0.0156110376473 | 0.0359326705671114 |
| Myo1b | ENSMUSG00000018417 | 17912 | -0.368731744357909 | 3.95021923416754 | -2.76718305551845 | 0.0156301838159 | 0.0359691661667844 |
| Zfp1 | ENSMUSG00000055835 | 22640 | -0.268259077589965 | 2.47470037590743 | -2.76692539807833 | 0.0156379747232 | 0.035979520458862 |
| Snord22 | ENSMUSG00000065087 | 100127111 | 1.21667457658773 | 2.09388033065868 | 2.76538460749683 | 0.0156846436862 | 0.0360793013751888 |
| Pmf1 | ENSMUSG00000028066 | 67037 | -0.365318398890859 | 2.42947197968076 | -2.76516883668876 | 0.0156911900306 | 0.0360867658795517 |
| Phtf2 | ENSMUSG00000039987 | 68770 | -0.176599590404702 | 6.89416619548926 | -2.76486703102959 | 0.0157003511007 | 0.0361002393525243 |
| Cdc26 | ENSMUSG00000066149 | 66440 | -0.212493761374484 | 3.43924297644476 | -2.76472317825496 | 0.0157047194782 | 0.0361026895093699 |

| | | | | | | | |
|---|---|---|---|---|---|---|---|
| Vps26a | ENSMUSG00000020078 | 30930 | -0.217750149895675 | 4.59026489507431 | -2.76400361259483 | 0.0157265883875 | 0.0361387844969077 |
| Als2cl | ENSMUSG00000044037 | 235633 | -0.674998309267081 | 2.51402872089479 | -2.76398899301574 | 0.0157270330117 | 0.0361387844969077 |
| Kpna4 | ENSMUSG00000027782 | 16649 | -0.169700496639728 | 6.53260206174948 | -2.76367456226597 | 0.0157365987499 | 0.0361531653921739 |
| Hectd4 | ENSMUSG00000042744 | 269700 | 0.189814975173356 | 5.21914746628292 | 2.76343907173119 | 0.0157437666650 | 0.0361620326571196 |
| Zfc3h1 | ENSMUSG00000034163 | 216345 | 0.287401485772169 | 4.54136973803397 | 2.76276854576431 | 0.0157641937883 | 0.0362013450210878 |
| Efl1 | ENSMUSG00000038563 | 101592 | 0.29200941989665 | 3.33941444125633 | 2.76245542817376 | 0.0157737415840 | 0.0362156625194879 |
| Nck2 | ENSMUSG00000066877 | 17974 | 0.209393460718383 | 3.35936613030942 | 2.76141579680556 | 0.0158054833247 | 0.0362809194084029 |
| Olfr720 | ENSMUSG00000052417 | 258387 | -1.47914535588423 | 0.922915815187611 | -2.75993825295922 | 0.0158507028148 | 0.0363770800900932 |
| Mterf4 | ENSMUSG00000026273 | 69821 | -0.307856035442402 | 1.98019522869776 | -2.75945719520633 | 0.0158654526350 | 0.036396563957463 |
| Ifnar2 | ENSMUSG00000022971 | 15976 | 0.368031180428969 | 2.97140856087699 | 2.75944417905589 | 0.0158658519126 | 0.036396563957463 |
| Nae1 | ENSMUSG00000031878 | 234664 | -0.199089021176568 | 4.02365217568624 | -2.75900571800039 | 0.0158793076907 | 0.0364197885789522 |
| Zfp704 | ENSMUSG00000040209 | 170753 | 0.301399696777379 | 4.07134070920943 | 2.75840505459836 | 0.0158977593599 | 0.036454459316863 |
| H2-Ke6 | ENSMUSG00000073422 | 14979 | -0.273629480506889 | 3.60523848259787 | -2.75772294059062 | 0.0159187385235 | 0.0364949099834982 |
| Ppp1cc | ENSMUSG00000004455 | 19047 | -0.229231824954775 | 4.56434729001295 | -2.75661405435527 | 0.0159529013322 | 0.0365602884103971 |
| 1700094D03Rik | ENSMUSG00000027942 | 73545 | -0.35658317688261 | 3.99042103343973 | -2.75658020597598 | 0.0159539452673 | 0.0365602884103971 |
| Prdx1 | ENSMUSG00000028691 | 18477 | -0.219258845049122 | 5.70221309185059 | -2.75564093482251 | 0.0159829404704 | 0.0366172587947514 |
| N4bp1 | ENSMUSG00000031652 | 80750 | -0.209260398034217 | 4.35317509843138 | -2.75555792833714 | 0.0159855053488 | 0.0366172587947514 |
| Ccl21a | ENSMUSG00000094686 | 18829 | 0.727499540199067 | 1.94781146880724 | 2.75519572120571 | 0.0159967021641 | 0.0366352296995735 |
| Ubn2 | ENSMUSG00000038538 | 320538 | 0.183352497123018 | 5.66140343848662 | 2.75498857099991 | 0.0160031091915 | 0.0366422259264461 |
| Hmgb1 | ENSMUSG00000066551 | 15289 | -0.186126733516869 | 6.25008534200303 | -2.75221461730168 | 0.0160891480090 | 0.0368315134840667 |
| Cc2d1b | ENSMUSG00000028582 | 319965 | 0.215618766938451 | 3.8288465081191 | 2.75207800388186 | 0.0160933969679 | 0.0368335264356108 |
| Dpep1 | ENSMUSG00000019278 | 13479 | 0.333752908899292 | 2.51363290892353 | 2.7516437506625 | 0.0161069104252 | 0.036856738090528 |
| Trex1 | ENSMUSG00000049734 | 22040 | 0.381221614898012 | 1.89444813508707 | 2.75092226966909 | 0.0161293866083 | 0.0368937650765752 |
| Erdr1 | ENSMUSG00000096768 | 170942 | 0.735657169907819 | 4.59581859802115 | 2.75090766067763 | 0.0161298420364 | 0.0368937650765752 |
| Arhgef6 | ENSMUSG00000031133 | 73341 | -0.173960252650983 | 5.54841669567865 | -2.75059292137638 | 0.0161396569338 | 0.0369084916200042 |
| Slc8a1 | ENSMUSG00000054640 | 20541 | 0.373363083043457 | 3.18710722677127 | 2.74930116923195 | 0.0161800003956 | 0.0369930107352747 |
| Nudt14 | ENSMUSG00000002804 | 66174 | -0.442029933558534 | 1.89948812959506 | -2.74889315961369 | 0.0161927636545 | 0.0370144499180726 |
| Nol10 | ENSMUSG00000061458 | 217431 | 0.292268389040785 | 2.94789462532433 | 2.74876475224655 | 0.0161967824986 | 0.0370158957751264 |
| St3gal4 | ENSMUSG00000032038 | 20443 | 0.566601699761265 | 1.50316639652465 | 2.74845408436767 | 0.0162065096948 | 0.0370303840874569 |
| Neurl4 | ENSMUSG00000047284 | 216860 | -0.242976530043985 | 4.18692256710492 | -2.74786248869561 | 0.0162250486999 | 0.0370649963091076 |
| Col15a1 | ENSMUSG00000028339 | 12819 | 0.434421748806382 | 6.33893046975519 | 2.74694746098334 | 0.0162537640205 | 0.0371228362951806 |
| Tnfsf12 | ENSMUSG00000097328 | 21944 | 0.433751908258996 | 1.87787404649683 | 2.74655343529095 | 0.0162661445952 | 0.0371433521142302 |
| Serpine1 | ENSMUSG00000037411 | 18787 | 0.657492810970043 | 2.79619072188211 | 2.74577666696175 | 0.0162905782228 | 0.0371913764732449 |
| Fnbp1l | ENSMUSG00000039735 | 214459 | -0.389890525389458 | 2.98937338009228 | -2.74463961051964 | 0.0163264095845 | 0.0372653963600713 |
| Triobp | ENSMUSG00000033088 | 110253 | -0.229201148628462 | 4.36580100064674 | -2.74351515800594 | 0.0163619195043 | 0.0373386517958695 |
| Abhd17c | ENSMUSG00000038459 | 70178 | 0.334126961033846 | 2.24901851318392 | 2.74312676898733 | 0.0163742022567 | 0.037358882251766 |
| Hist1h2ab | ENSMUSG00000061615 | 319172 | 1.41336200422061 | 0.270700090408304 | 2.74244804599176 | 0.0163956884170 | 0.0374000980982326 |
| Mcfd2 | ENSMUSG00000024150 | 193813 | -0.215810704467712 | 4.36802103121287 | -2.74226234923053 | 0.0164015717701 | 0.0374057127100789 |
| Slc25a1 | ENSMUSG00000003528 | 13358 | 0.621670183545334 | 1.14561585502152 | 2.74213085540448 | 0.0164057390817 | 0.0374074121614831 |
| Braf | ENSMUSG00000002413 | 109880 | 0.227065323396116 | 3.17636587341995 | 2.74133612576857 | 0.0164309477399 | 0.037457078153261 |
| Fbxo10 | ENSMUSG00000048232 | 269529 | 0.424966862645108 | 1.73023976386727 | 2.74117625197496 | 0.0164360234738 | 0.0374608366541216 |
| Ypel3 | ENSMUSG00000042675 | 66090 | -0.20348785843515 | 4.49652067627809 | -2.74043209261879 | 0.0164596694961 | 0.0375069099985113 |
| Ces2b | ENSMUSG00000050097 | 234669 | -1.26238386860474 | 2.28728789794678 | -2.73998523660926 | 0.0164738845292 | 0.0375314780719155 |
| Cobl | ENSMUSG00000020173 | 12808 | 0.216036572590188 | 5.13871736361188 | 2.73941822089337 | 0.0164919392548 | 0.037564781828347 |
| Kcnb1 | ENSMUSG00000050556 | 16500 | -0.351964301634481 | 4.53846629895291 | -2.73855661414826 | 0.0165194112606 | 0.0376130614845198 |
| Rab11a | ENSMUSG00000004771 | 53869 | -0.188462186154521 | 4.63711772062159 | -2.73853762530484 | 0.0165200172159 | 0.0376130614845198 |
| Trappc2 | ENSMUSG00000079317 | 66226 | -0.328240554063536 | 2.26443535303945 | -2.73582649065234 | 0.0166067557888 | 0.0378026754535931 |
| Zfp942 | ENSMUSG00000071267 | 73233 | -0.285766688907519 | 2.65665019994847 | -2.73542598853451 | 0.0166196068901 | 0.0378240522414847 |
| Slc25a22 | ENSMUSG00000019082 | 68267 | -0.609885120600172 | 3.54365655040626 | -2.7349966099676 | 0.0166333953526 | 0.0378475529975162 |
| Vmn1r26 | ENSMUSG00000048697 | 171190 | -1.4408184633232 | 2.57576951640993 | -2.73247073239339 | 0.0167147343151 | 0.0380247160871613 |
| Brcc3 | ENSMUSG00000031201 | 210766 | -0.212614996494124 | 3.59379623842034 | -2.73015992601107 | 0.0167894876427 | 0.0381868267630507 |
| Txndc11 | ENSMUSG00000022498 | 106200 | 0.35691188913694 | 3.18855544783573 | 2.7292435318101 | 0.0168192228073 | 0.0382464997933482 |
| Tbx1 | ENSMUSG00000009097 | 21380 | -0.259089925142463 | 3.99875040203812 | -2.72854472021635 | 0.0168419323989 | 0.0382901753437967 |
| Nudc | ENSMUSG00000028851 | 18221 | -0.161071463754924 | 5.12312363253169 | -2.72762944750708 | 0.0168717217110 | 0.0383499251452189 |
| Klhl15 | ENSMUSG00000043929 | 236904 | 0.400623533428736 | 1.95641879772322 | 2.72724873423935 | 0.0168841279001 | 0.0383682834177342 |
| Ap4s1 | ENSMUSG00000020955 | 11782 | -0.257539649998122 | 3.82315057134261 | -2.72716620706356 | 0.0168868183627 | 0.0383682834177342 |
| Dusp7 | ENSMUSG00000053716 | 235584 | -0.324127257770438 | 3.67500077064182 | -2.72684647977039 | 0.0168972457217 | 0.0383839968796737 |
| Iars | ENSMUSG00000037851 | 105148 | -0.173649711690847 | 5.55512556079316 | -2.72673162179956 | 0.0169009931525 | 0.0383845327551902 |
| Med11 | ENSMUSG00000018923 | 66172 | 0.451147963468706 | 1.24072256602919 | 2.72657879503566 | 0.0169059806358 | 0.0383878841566908 |
| Crot | ENSMUSG00000003623 | 74114 | -0.358901915289964 | 4.23860549359958 | -2.72630087839547 | 0.0169150540947 | 0.0384004602735201 |
| 9130401M01Rik | ENSMUSG00000101892 | 75758 | 0.348076324720052 | 2.42023308201844 | 2.72619398844964 | 0.0169185451173 | 0.0384004602735201 |
| Kctd12b | ENSMUSG00000041633 | 207474 | -0.558408048578745 | 2.58687685374875 | -2.72555951564377 | 0.0169392814634 | 0.0384395444628426 |
| Slc35a4 | ENSMUSG00000033272 | 67843 | -0.186498442017471 | 4.82179546570233 | -2.72525405568732 | 0.0169492735947 | 0.0384542361279041 |
| Sh3gl1 | ENSMUSG00000003200 | 20405 | 0.434087581243103 | 2.88172960314891 | 2.72504519098004 | 0.0169561092376 | 0.0384617617922888 |
| Akap6 | ENSMUSG00000061603 | 238161 | 0.210981197171939 | 7.34128156068938 | 2.72447239693373 | 0.0169748692304 | 0.0384963268793662 |
| Twf2 | ENSMUSG00000023277 | 23999 | -0.205858817944849 | 7.09821172730822 | -2.72418572752369 | 0.0169842657534 | 0.0385096471585629 |
| Rab5a | ENSMUSG00000017831 | 271457 | 0.175454832896359 | 4.95702151824372 | 2.71912238152445 | 0.0171510725395 | 0.0388797955059967 |
| Rae1 | ENSMUSG00000027509 | 66679 | 0.241484030352569 | 2.93138233989119 | 2.71893566756491 | 0.0171572541065 | 0.0388818239771965 |
| Zbed4 | ENSMUSG00000034333 | 223773 | -0.280955781120409 | 3.90987378605684 | -2.71888048658983 | 0.0171590814075 | 0.0388818239771965 |
| Fbxl18 | ENSMUSG00000066640 | 231863 | 0.321583661406123 | 2.75320993391797 | 2.71819821342282 | 0.0171816903679 | 0.0389188609993502 |
| Fuca1 | ENSMUSG00000028673 | 71665 | 0.227035544929207 | 3.38518474710845 | 2.71817237465382 | 0.0171825471761 | 0.0389188609993502 |
| Zhx1 | ENSMUSG00000022361 | 22770 | -0.201193329354398 | 5.21691139270539 | -2.71760433643928 | 0.0172013937421 | 0.0389458688401515 |
| Tma16 | ENSMUSG00000025591 | 66282 | -0.286737187206342 | 3.41344605934338 | -2.71759821917076 | 0.0172015968127 | 0.0389458688401515 |
| Gpn3 | ENSMUSG00000029464 | 68080 | -0.415487183132646 | 2.03942008069868 | -2.71658453386316 | 0.0172352797216 | 0.0390140490034161 |
| Ncald | ENSMUSG00000051359 | 52589 | 0.315941212229633 | 2.34642719976124 | 2.71642248870404 | 0.0172406701481 | 0.0390181708879784 |
| Atp6ap1 | ENSMUSG00000019087 | 54411 | 0.187486500115019 | 4.89447931985821 | 2.71459149026488 | 0.0173016926489 | 0.0391481685667182 |
| Rbm20 | ENSMUSG00000043639 | 73713 | -0.217470239636648 | 5.69692791921609 | -2.71410083952475 | 0.0173180805483 | 0.0391771395848166 |
| Idnk | ENSMUSG00000050002 | 75731 | 0.513170761595635 | 1.34008173952049 | 2.71367136594327 | 0.0173324375369 | 0.0392015052174951 |
| Mvb12b | ENSMUSG00000038740 | 72543 | -0.190148047787442 | 4.35894236167485 | -2.71341245315409 | 0.0173410984113 | 0.0392129802927061 |
| Spsb1 | ENSMUSG00000039911 | 74646 | 0.604142804481447 | 2.87077686913326 | 2.71279079556995 | 0.0173619106776 | 0.0392519225682691 |
| Ercc5 | ENSMUSG00000026048 | 22592 | -0.194809310244859 | 3.99341871471197 | -2.71064303657463 | 0.0174340024416 | 0.0394067577935938 |
| Apobec3 | ENSMUSG00000009585 | 80287 | 0.596950787451588 | 1.95778986427308 | 2.71033654634372 | 0.0174443138800 | 0.0394219133806721 |
| Fes | ENSMUSG00000053158 | 14159 | 0.506500039826042 | 2.49831615446155 | 2.70934628002976 | 0.0174776706860 | 0.0394891315148981 |
| Mrpl32 | ENSMUSG00000015672 | 75398 | -0.268305689502803 | 3.38413916081628 | -2.70882744641329 | 0.0174951722561 | 0.0395205058755207 |
| Ublcp1 | ENSMUSG00000041231 | 79560 | -0.344081463816838 | 3.53316311921894 | -2.70764143558839 | 0.0175352436106 | 0.0396028404769545 |
| Phlpp1 | ENSMUSG00000044340 | 98432 | -0.240096021408787 | 3.66484368865655 | -2.70737324804762 | 0.0175443171735 | 0.0396151479082318 |
| Sirt7 | ENSMUSG00000025138 | 209011 | 0.377595485718069 | 1.87554039210569 | 2.70720622030523 | 0.0175499705201 | 0.039619728987145 |
| Angpt1 | ENSMUSG00000022309 | 11600 | -0.227003957064028 | 4.33802922384665 | -2.70558296477235 | 0.0176050050431 | 0.0397357650479089 |
| Mocs2 | ENSMUSG00000015536 | 17434 | -0.321417756473675 | 3.44508927614201 | -2.70458452116869 | 0.0176389396118 | 0.0398041389134784 |
| Tmem168 | ENSMUSG00000029569 | 101118 | 0.25753181966981 | 3.37603913056641 | 2.70397931794826 | 0.0176595399696 | 0.0398424007035214 |
| Wdr3 | ENSMUSG00000033285 | 269470 | 0.237799762317056 | 3.08900182207895 | 2.70284127701466 | 0.0176983409935 | 0.0399203212245085 |
| Tdrd7 | ENSMUSG00000035517 | 100121 | 0.338313796860874 | 3.1711972620445 | 2.70275221246863 | 0.0177013811156 | 0.0399203212245085 |
| COX2 | ENSMUSG00000064354 | 17709 | -0.398626849148393 | 10.8245092507125 | -2.70258913896125 | 0.0177069487731 | 0.0399246404784741 |
| Sbk2 | ENSMUSG00000030433 | 381836 | 0.322266580289442 | 5.46035844346306 | 2.70142311885608 | 0.0177468088158 | 0.0400062625625107 |
| Stx16 | ENSMUSG00000027522 | 228960 | 0.24356574674639 | 3.83418837413106 | 2.70012872408794 | 0.0177911597456 | 0.0400936724731484 |
| Slc1a1 | ENSMUSG00000024935 | 20510 | -0.476857090549036 | 1.98509074990595 | -2.70007742440654 | 0.0177929196932 | 0.0400936724731484 |
| Rpl39 | ENSMUSG00000079641 | 67248 | 0.295965764257731 | 4.81255627243137 | 2.69970121279971 | 0.0178058316405 | 0.040113378117273 |
| Tapbp | ENSMUSG00000024308 | 21356 | 0.22276734309603 | 4.76889475535361 | 2.69960881797957 | 0.0178090041170 | 0.040113378117273 |
| Supt16 | ENSMUSG00000035726 | 114741 | 0.171661558038505 | 4.87914552668456 | 2.6988640788449 | 0.0178345956675 | 0.0401627452084598 |
| Msl2 | ENSMUSG00000066415 | 77853 | -0.157906409917867 | 4.55600558053827 | -2.69865114393927 | 0.0178419193589 | 0.0401709620004401 |
| Acsf2 | ENSMUSG00000076435 | 264895 | -0.523114846047469 | 2.70296876028285 | -2.69766050058436 | 0.0178760301392 | 0.0402394739399744 |
| Purb | ENSMUSG00000094483 | 19291 | 0.172599762106779 | 5.4637856837763 | 2.69733649092057 | 0.0178872005364 | 0.0402563288169045 |
| Eif3a | ENSMUSG00000024991 | 13669 | 0.182261671941091 | 7.87551781718883 | 2.69712221424909 | 0.0178945915695 | 0.0402646727967694 |
| Abhd6 | ENSMUSG00000025277 | 66082 | -0.568082929765716 | 1.25541076718179 | -2.69699357638542 | 0.0178990300976 | 0.0402663712692412 |
| Tprn | ENSMUSG00000048707 | 97031 | 0.451083859358957 | 1.13698489179294 | 2.69514906443583 | 0.0179627913390 | 0.0404014963224424 |
| Rccd1 | ENSMUSG00000038930 | 269955 | 0.534809445923573 | 1.3781781518509 | 2.69452764873566 | 0.0179843222878 | 0.040441601918954 |
| Snrpb2 | ENSMUSG00000008333 | 20639 | -0.285169309672445 | 2.89298348119515 | -2.69421705910743 | 0.0179950930887 | 0.0404574994968079 |
| Vps13b | ENSMUSG00000037646 | 666173 | 0.217960856435079 | 4.60947487439245 | 2.6922891275853 | 0.0180620916979 | 0.0405997788093645 |
| Dusp14 | ENSMUSG00000018648 | 56405 | 0.641878339853585 | 1.79191598926046 | 2.69207625002462 | 0.0180695044044 | 0.0406080905931137 |
| C130074G19Rik | ENSMUSG00000039349 | 226777 | -0.385555610066122 | 3.36088462973653 | -2.69147935801161 | 0.0180903048616 | 0.0406464794331521 |
| Coq6 | ENSMUSG00000021235 | 217707 | -0.321238686177501 | 2.96955560947583 | -2.69096946655529 | 0.0181080920004 | 0.0406780833655731 |
| F11r | ENSMUSG00000038235 | 16456 | -0.448007979576285 | 2.4605009951826 | -2.68963846147904 | 0.0181546033202 | 0.0407741871570777 |
| A130077B15Rik | ENSMUSG00000112095 | 319272 | -1.10357307166266 | 3.4491835027394 | -2.68751483425557 | 0.0182290534714 | 0.0409329875712719 |
| Mien1 | ENSMUSG00000002580 | 103742 | 0.346743592386397 | 2.0968921744533 | 2.68662956714828 | 0.0182601769175 | 0.0409944534576783 |
| Umad1 | ENSMUSG00000089862 | 100036521 | -0.407377284204063 | 1.42226992795858 | -2.6856091379602 | 0.0182961164807 | 0.04106670415828 |
| Tfrc | ENSMUSG00000022797 | 22042 | -0.438051281770858 | 7.0900710492083 | -2.68533804106034 | 0.0183056761013 | 0.0410797260241287 |

| | | | | | | | |
|---|---|---|---|---|---|---|---|
| Srek1ip1 | ENSMUSG00000021716 | 67288 | -0.254648464885892 | 3.20519466744336 | -2.68411679681977 | 0.0183488008657 | 0.0411680505466363 |
| Ino80d | ENSMUSG00000040865 | 227195 | 0.167144491649911 | 4.67148474821645 | 2.68378536601329 | 0.0183605214575 | 0.0411858937107839 |
| Thpo | ENSMUSG00000022847 | 21832 | -0.417273420098761 | 1.9459852501874 | -2.68355136614018 | 0.0183688009292 | 0.0411960120963301 |
| Chordc1 | ENSMUSG00000001774 | 66917 | -0.229356518856585 | 4.47490512185896 | -2.68343299078041 | 0.0183729907153 | 0.0411969562071072 |
| 4933427D14Rik | ENSMUSG00000020807 | 74477 | 0.708645941150004 | 0.823183362610733 | 2.68271625585205 | 0.0183983794726 | 0.0412454237111247 |
| Ctnnbip1 | ENSMUSG00000028988 | 67087 | -0.295811242658215 | 3.30634264312326 | -2.68220327440519 | 0.0184165704633 | 0.041277387195226 |
| Enox2 | ENSMUSG00000031109 | 209224 | 0.424222718903266 | 1.74808821366759 | 2.6816373115855 | 0.0184366613438 | 0.0413142978985655 |
| Wdfy2 | ENSMUSG00000014547 | 268752 | 0.231711072132529 | 3.55327564428781 | 2.68144140769815 | 0.0184436206250 | 0.0413214218186021 |
| Amigo1 | ENSMUSG00000050947 | 229715 | 0.256601036050269 | 3.70208286901812 | 2.68116684352341 | 0.0184533785351 | 0.0413348116327078 |
| Mydgf | ENSMUSG00000019579 | 28106 | 0.357674450503694 | 2.73858697105407 | 2.68051380926767 | 0.0184766073301 | 0.0413783640084731 |
| Abcb6 | ENSMUSG00000026198 | 74104 | -0.281209695417639 | 3.11803747883767 | -2.68033074283804 | 0.0184831242124 | 0.0413844798784584 |
| Wdr91 | ENSMUSG00000058486 | 101240 | 0.353418804433757 | 2.13991189318126 | 2.68013445583571 | 0.0184901142126 | 0.0413916523569514 |
| Rgs7bp | ENSMUSG00000021719 | 52882 | -1.00856728635457 | 2.10068677531163 | -2.6797810248932 | 0.0185027067737 | 0.041411361126812 |
| Pld1 | ENSMUSG00000027695 | 18805 | 0.352959972306298 | 2.77418454794515 | 2.67872057705902 | 0.0185405401027 | 0.0414875422442632 |
| Trappc12 | ENSMUSG00000020628 | 217449 | -0.286142812061509 | 3.83240711122104 | -2.67751918601015 | 0.0185834928082 | 0.0415751452899738 |
| Dnajb9 | ENSMUSG00000014905 | 27362 | 0.302265359701072 | 4.64465615618933 | 2.67635295576326 | 0.0186252810630 | 0.0416573720155156 |
| Gpc6 | ENSMUSG00000058571 | 23888 | 0.526363943946267 | 1.48791048743737 | 2.67628081930812 | 0.0186278688511 | 0.0416573720155156 |
| Opcml | ENSMUSG00000062257 | 330908 | -0.579521788730849 | 1.32956962480234 | -2.6756384631875 | 0.0186509278587 | 0.0417004075319222 |
| Arf3 | ENSMUSG00000051853 | 11842 | 0.287195872388621 | 3.43077239632512 | 2.67536297597674 | 0.0186608256826 | 0.0417140052223117 |
| Gm14325 | ENSMUSG00000095362 | 329575 | -0.288449059946758 | 3.25397587411303 | -2.67445909828774 | 0.0186933365054 | 0.041777479774048 |
| Abcf3 | ENSMUSG00000003234 | 27406 | -0.224629525809616 | 4.66263581008416 | -2.67431506321406 | 0.0186985222717 | 0.041777479774048 |
| Psenen | ENSMUSG00000036835 | 66340 | -0.241141467456788 | 3.78436094591819 | -2.67425495191532 | 0.0187006869027 | 0.041777479774048 |
| Bsdc1 | ENSMUSG00000040859 | 100383 | -0.183952223083294 | 4.61471041257569 | -2.67393664520109 | 0.0187121533151 | 0.0417945541249214 |
| Cnot1 | ENSMUSG00000036550 | 234594 | -0.134414915703404 | 6.37243614360594 | -2.67379620700024 | 0.0187172145171 | 0.0417973180566137 |
| Mllt3 | ENSMUSG00000028496 | 70122 | -0.211635811634181 | 5.09451834195611 | -2.6729313305068 | 0.0187484128664 | 0.0418584356705782 |
| Sephs1 | ENSMUSG00000026662 | 109079 | -0.208830347166862 | 3.14046322022433 | -2.66977403640511 | 0.0188627347800 | 0.0421050753279146 |
| Nphp1 | ENSMUSG00000027378 | 53885 | -0.248528754851246 | 3.55380887711952 | -2.6693731930354 | 0.0188772972768 | 0.0421272493540657 |
| Mrpl10 | ENSMUSG00000001445 | 107732 | -0.18851961094892 | 3.914557944112 | -2.66927978549234 | 0.0188806923105 | 0.0421272493540657 |
| Fkbp5 | ENSMUSG00000024222 | 14229 | 0.712513330452508 | 4.20236465408207 | 2.66918246112135 | 0.0188842303389 | 0.0421272493540657 |
| Dusp11 | ENSMUSG00000030002 | 72102 | 0.278771165669072 | 4.35950870435427 | 2.6686604999293 | 0.0189032161780 | 0.0421609989884291 |
| Dctpp1 | ENSMUSG00000042462 | 66422 | -0.401263734147683 | 2.09697959801153 | -2.66755725305004 | 0.0189434068486 | 0.0422420196374914 |
| Kif3b | ENSMUSG00000027475 | 16569 | 0.298373958686854 | 3.17521460309391 | 2.66712767937282 | 0.0189590784518 | 0.0422683431688808 |
| Olfr633 | ENSMUSG00000073937 | 258351 | -1.52465996948493 | -0.469348696420239 | -2.66681291256259 | 0.0189705697035 | 0.0422823600208722 |
| Clip1 | ENSMUSG00000049550 | 56430 | -0.170370733883454 | 8.24484586779041 | -2.66674357833848 | 0.0189731018115 | 0.0422823600208722 |
| Pacs1 | ENSMUSG00000024855 | 107975 | -0.198952439333711 | 3.96555586631151 | -2.66587083522901 | 0.0190050027942 | 0.0423448197194417 |
| D230025D16Rik | ENSMUSG00000031889 | 234678 | -0.35183517779165 | 2.97155502941771 | -2.66548353194977 | 0.0190191764222 | 0.0423677639029509 |
| Tmed2 | ENSMUSG00000029390 | 56334 | 0.183533614500386 | 6.95142727139444 | 2.6652746776612 | 0.0190268238538 | 0.0423761637217006 |
| Zscan26 | ENSMUSG00000022228 | 432731 | -0.307810925416301 | 3.99068301552388 | -2.66455100490511 | 0.0190533450756 | 0.0424261170828963 |
| Pcdh1 | ENSMUSG00000051375 | 75599 | -0.39744214936802 | 3.95645057050371 | -2.66445093272657 | 0.0190570153578 | 0.0424261170828963 |
| Hsph1 | ENSMUSG00000029657 | 15505 | 0.25533238708083 | 5.69900202558115 | 2.6634267533034 | 0.0190946180709 | 0.0425011749407472 |
| Abcb10 | ENSMUSG00000031974 | 56199 | -0.268186381602327 | 3.85598111761933 | -2.6631695859158 | 0.0191040713641 | 0.0425135594628862 |
| Ube2g1 | ENSMUSG00000020794 | 67128 | -0.263469515650526 | 5.36957189476221 | -2.66222651173283 | 0.0191387767294 | 0.0425821226195666 |
| Dcaf15 | ENSMUSG00000037103 | 212123 | 0.433629569371533 | 1.63126693176436 | 2.66154832344744 | 0.0191637721686 | 0.0426290585217678 |
| Ap5s1 | ENSMUSG00000068264 | 69596 | -0.424014658345727 | 3.01333108353774 | -2.66105331751997 | 0.0191820362396 | 0.0426610047071496 |
| Ap3s2 | ENSMUSG00000063801 | 11778 | 0.201286797134133 | 3.74089274638043 | 2.65963645447886 | 0.0192344073707 | 0.0427687768449254 |
| Llgl2 | ENSMUSG00000020782 | 217325 | -0.433928742175393 | 1.76421566708032 | -2.65809998297605 | 0.0192913566694 | 0.0428866828866977 |
| Ddx52 | ENSMUSG00000020677 | 78394 | -0.227814107281028 | 3.38294078625081 | -2.65687464693349 | 0.0193368911936 | 0.042979169914194 |
| Acta2 | ENSMUSG00000035783 | 11475 | -0.47286034741169 | 5.41241013540175 | -2.65650936962442 | 0.0193504854104 | 0.0430006415983188 |
| Pdss1 | ENSMUSG00000026784 | 56075 | -0.461282645134249 | 2.54546488034443 | -2.65557914368145 | 0.0193851468086 | 0.0430689105214929 |
| Insig2 | ENSMUSG00000003721 | 72999 | -0.197004416435165 | 4.23143836521157 | -2.65543585225431 | 0.0193904913901 | 0.0430720303567828 |
| Abhd10 | ENSMUSG00000033157 | 213012 | -0.37116711051607 | 1.94751100435807 | -2.65379113483419 | 0.0194519398388 | 0.0431965037773229 |
| Med30 | ENSMUSG00000038622 | 69790 | 0.309462446368635 | 2.40700690124785 | 2.65372455651921 | 0.0194544312593 | 0.0431965037773229 |
| Col4a3bp | ENSMUSG00000021669 | 68018 | -0.157649784358467 | 5.4142305315002 | -2.6534209779997 | 0.0194657953723 | 0.0432129588170288 |
| Myl10 | ENSMUSG00000005474 | 59310 | -0.570292102160866 | 1.13358685154725 | -2.65327754208581 | 0.0194711669748 | 0.0432156888777913 |
| Rprd1b | ENSMUSG00000027651 | 70470 | -0.152800454633496 | 4.58795025206208 | -2.65308627498831 | 0.0194783320723 | 0.0432156888777913 |
| Sorbs2 | ENSMUSG00000031626 | 234214 | 0.286878145503943 | 3.70417397472479 | 2.65307150033166 | 0.0194788856555 | 0.0432156888777913 |
| Rnf7 | ENSMUSG00000051234 | 19823 | -0.196442582456114 | 4.80916814293108 | -2.652239832057 | 0.0195100716243 | 0.0432760943437563 |
| Brat1 | ENSMUSG00000000148 | 231841 | 0.384581600735065 | 2.39065935708994 | 2.65152111828106 | 0.0195370610942 | 0.0433128044188692 |
| Hist1h4n | ENSMUSG00000069305 | 319161 | 0.538912671220985 | 1.4467535567499 | 2.65150186717414 | 0.0195377845193 | 0.0433128044188692 |
| Sema3g | ENSMUSG00000021904 | 218877 | -0.338567429383491 | 2.99276270641923 | -2.65148259668017 | 0.0195385086990 | 0.0433128044188692 |
| Dock9 | ENSMUSG00000025558 | 105445 | -0.267017723253929 | 4.98338313097375 | -2.65087427989369 | 0.0195613824839 | 0.0433494881532732 |
| Trim55 | ENSMUSG00000060913 | 381485 | -0.179667734805842 | 5.16651942015031 | -2.65083160010845 | 0.0195629882956 | 0.0433494881532732 |
| Slc47a1 | ENSMUSG00000010122 | 67473 | -0.313166738996856 | 3.73964946075659 | -2.64836937537598 | 0.0196558457843 | 0.0435464228352729 |
| Hmbs | ENSMUSG00000032126 | 15288 | -0.278902464141603 | 3.00926206559205 | -2.6476380997992 | 0.0196835066443 | 0.0435984526730234 |
| Peak1 | ENSMUSG00000074305 | 244895 | 0.268231186868064 | 5.17902312011619 | 2.64753768392606 | 0.0196873078761 | 0.0435984526730234 |
| Cmtm7 | ENSMUSG00000032436 | 102545 | 0.521009924983156 | 0.972679329200601 | 2.64685597291199 | 0.0197131328323 | 0.0436468006867535 |
| Abca3 | ENSMUSG00000024130 | 27410 | 0.193304459281452 | 3.75366541143369 | 2.64668598463099 | 0.0197195775474 | 0.0436522280622244 |
| Bmt2 | ENSMUSG00000042742 | 101148 | 0.266019910289712 | 3.40912906864902 | 2.64650271745381 | 0.0197265279942 | 0.0436587725257985 |
| Myadml2 | ENSMUSG00000025141 | 68515 | -0.184138297553064 | 5.22713336481389 | -2.64515460528325 | 0.0197777286235 | 0.0437632290655338 |
| Rps6kc1 | ENSMUSG00000089872 | 320119 | 0.200488307818514 | 3.71582947217948 | 2.64480621051043 | 0.0197909814601 | 0.0437761221357213 |
| Sel1l | ENSMUSG00000020964 | 20338 | 0.149496763711438 | 6.10620667331492 | 2.64479087885202 | 0.0197915648700 | 0.0437761221357213 |
| Med16 | ENSMUSG00000013833 | 216154 | -0.17031680324465 | 4.50385550730455 | -2.64444258415175 | 0.0198048229015 | 0.0437965848951677 |
| Ccdc107 | ENSMUSG00000028461 | 622404 | 0.45419930449146 | 3.03234515250241 | 2.64378948251255 | 0.0198297068362 | 0.0438427438160512 |
| Vmn1r60 | ENSMUSG00000118298 | 636697 | -1.00095619836287 | 1.2329076313278 | -2.64135687519311 | 0.0199226594828 | 0.0440393510225906 |
| 6330415B21Rik | ENSMUSG00000108077 | 70753 | -0.918265724389186 | 0.217318439939131 | -2.64049051745466 | 0.0199558660564 | 0.0441030431095681 |
| Prom1 | ENSMUSG00000029086 | 19126 | -0.673677631735874 | 0.927123500057777 | -2.64039469678348 | 0.0199595420604 | 0.0441030431095681 |
| Aplp2 | ENSMUSG00000031996 | 11804 | -0.143650930142566 | 6.60028957994484 | -2.63975942294939 | 0.0199839299424 | 0.044148006912091 |
| Socs6 | ENSMUSG00000056153 | 54607 | 0.247903800883337 | 5.37758049454904 | 2.63820405696918 | 0.0200437619610 | 0.0442712390375143 |
| Slc43a2 | ENSMUSG00000038178 | 215113 | -0.22665550600658 | 6.04551166491686 | -2.63759483672452 | 0.0200672448982 | 0.0443121263827706 |
| Usp54 | ENSMUSG00000034235 | 78787 | 0.341107089779516 | 2.30615659457747 | 2.63751352177308 | 0.0200703812753 | 0.0443121263827706 |
| Spata13 | ENSMUSG00000021990 | 219140 | -0.501180820293334 | 2.71152221862593 | -2.63709784223957 | 0.0200864217750 | 0.0443385857072973 |
| Nhsl2 | ENSMUSG00000079481 | 100042480 | 0.290703907219728 | 3.05890788912148 | 2.63649867133599 | 0.0201095648548 | 0.0443783376978059 |
| Ddx54 | ENSMUSG00000029599 | 71990 | 0.178535245526856 | 4.60906714365271 | 2.63642143297552 | 0.0201125500827 | 0.0443783376978059 |
| Atg5 | ENSMUSG00000038160 | 11793 | 0.246795420297871 | 3.47582076030435 | 2.63505847121225 | 0.0201652987733 | 0.0444847769444916 |
| Aip | ENSMUSG00000024847 | 11632 | -0.215013388287793 | 3.82129222926998 | -2.63496481765809 | 0.0201689282350 | 0.0444847769444916 |
| Ccnl1 | ENSMUSG00000027829 | 56706 | 0.27605761885138 | 3.54221626866764 | 2.6333923833785 | 0.0202299613346 | 0.0446103908310541 |
| Copg2 | ENSMUSG00000025607 | 54160 | 0.299799193919576 | 3.27066594633882 | 2.63248733385648 | 0.0202651715374 | 0.0446790217982373 |
| Klhl28 | ENSMUSG00000020948 | 66689 | 0.206886263830569 | 3.09259732946689 | 2.63134067794942 | 0.0203098666597 | 0.0447685324577366 |
| Mrps26 | ENSMUSG00000037740 | 99045 | -0.240824533674796 | 4.34925177809423 | -2.6308721362528 | 0.0203281573112 | 0.0447998160422812 |
| Agmo | ENSMUSG00000050103 | 319660 | 0.480619781974413 | 1.27200864055971 | 2.63009011688163 | 0.0203587209289 | 0.0448581291019676 |
| Rbbp4 | ENSMUSG00000057236 | 19646 | 0.169871198956453 | 4.91022894879275 | 2.62878819904062 | 0.0204097026504 | 0.0449572205904056 |
| Ppp1r7 | ENSMUSG00000026275 | 66385 | -0.190077239874304 | 4.04583608726793 | -2.62873167369027 | 0.0204119189269 | 0.0449572205904056 |
| Dctn6 | ENSMUSG00000031516 | 22428 | -0.256491593854634 | 3.4074705991731 | -2.62749510719847 | 0.0204604613670 | 0.0450550570514619 |
| Erc1 | ENSMUSG00000030172 | 111173 | -0.165507400210321 | 6.3637680090425 | -2.62730654545079 | 0.0204678733497 | 0.0450623008228978 |
| Tcf7l2 | ENSMUSG00000024985 | 21416 | 0.316555890004954 | 2.78547963332717 | 2.62672424093639 | 0.0204907790158 | 0.0451036458763718 |
| Dda1 | ENSMUSG00000074247 | 66498 | 0.208354899009783 | 3.80369814690858 | 2.62596311809257 | 0.0205207562223 | 0.0451605367962062 |
| Kdm3a | ENSMUSG00000053470 | 104263 | 0.278290398854099 | 3.69498616001531 | 2.62582097978122 | 0.0205263591240 | 0.045163774473689 |
| Slc7a6 | ENSMUSG00000031904 | 330836 | 0.374847727669464 | 2.11648060122958 | 2.62542265865066 | 0.0205420683172 | 0.0451892430620759 |
| Hipk2 | ENSMUSG00000061436 | 15258 | 0.257203795562773 | 6.91339087704205 | 2.6250773978718 | 0.0205556943246 | 0.0452101196505398 |
| Rabl6 | ENSMUSG00000015087 | 227624 | -0.200511511859202 | 6.52942224602376 | -2.62474833309664 | 0.0205686893038 | 0.0452296002373481 |
| Kpnb1 | ENSMUSG00000001440 | 16211 | 0.177282999906486 | 5.92194053404677 | 2.62457590073114 | 0.0205755019503 | 0.0452354810577398 |
| Lonp1 | ENSMUSG00000041168 | 74142 | -0.187486575053515 | 6.00127288857673 | -2.62293411472335 | 0.0206404771588 | 0.0453692048709978 |
| Adnp | ENSMUSG00000051149 | 11538 | 0.365477673162421 | 3.34394519218072 | 2.62281334797939 | 0.0206452644656 | 0.0453706043172226 |
| Abcg1 | ENSMUSG00000024030 | 11307 | 0.579797835565073 | 2.14268633448432 | 2.62182426652412 | 0.0206845131311 | 0.0454477212133669 |
| Gch1 | ENSMUSG00000037580 | 14528 | 0.399729130150309 | 1.69247770044441 | 2.62084851298732 | 0.0207233038269 | 0.0455238010717105 |
| Ppa1 | ENSMUSG00000020089 | 67895 | -0.303393788408153 | 4.59518674517918 | -2.62038109630283 | 0.0207419107764 | 0.0455551275514179 |
| Smim12 | ENSMUSG00000042380 | 80284 | -0.35190143672552 | 2.46729910146308 | -2.62020945089913 | 0.0207487477127 | 0.0455551275514179 |
| Rabep1 | ENSMUSG00000020817 | 54189 | -0.16779246220453 | 5.83175166154947 | -2.62017634098904 | 0.0207500667897 | 0.0455551275514179 |
| Acot6 | ENSMUSG00000043487 | 217700 | 0.399877407519272 | 1.74041638760152 | 2.61982060956668 | 0.0207642440266 | 0.0455770986859001 |
| Lamtor1 | ENSMUSG00000030842 | 66508 | -0.184618179306744 | 5.37731380314287 | -2.61941502008928 | 0.0207804195556 | 0.0456034467675412 |
| 4930435F18Rik | ENSMUSG00000114290 | 74680 | -1.12635697379959 | 0.430769177395319 | -2.61783919472046 | 0.0208433829055 | 0.0457324404937866 |
| Nnt | ENSMUSG00000116207 | 18115 | -1.1597212239549 | 5.76902328618806 | -2.61764068972383 | 0.0208513274114 | 0.0457379570351775 |
| Ndst2 | ENSMUSG00000039308 | 17423 | 0.363082674672481 | 2.10214236409569 | 2.61747560978249 | 0.0208579364262 | 0.0457379570351775 |
| Lsm6 | ENSMUSG00000031683 | 78651 | -0.21795119030412 | 3.89043999975021 | -2.61746278644348 | 0.0208584498966 | 0.0457379570351775 |
| Klhl29 | ENSMUSG00000020627 | 208439 | 0.545425663822595 | 1.95757409079208 | 2.61666136051999 | 0.0208905648171 | 0.0457991905368303 |
| Mbtps2 | ENSMUSG00000046873 | 270669 | 0.275287346436617 | 3.07738098289156 | 2.61628131389325 | 0.0209058108840 | 0.0458234246589203 |

| | | | | | | | |
|---|---|---|---|---|---|---|---|
| Mpz | ENSMUSG00000056569 | 17528 | -0.927644480036544 | 2.67653432036354 | -2.61597197149826 | 0.0209182285218 | 0.0458414506761139 |
| Mdm2 | ENSMUSG00000020184 | 17246 | 0.178078484064984 | 5.24206238383193 | 2.6156785068856 | 0.0209300154025 | 0.0458580874655364 |
| Gpsm1 | ENSMUSG00000026930 | 67839 | 0.445815987106893 | 1.53844458516627 | 2.61480298670295 | 0.0209652185694 | 0.0459175404409872 |
| Tbc1d14 | ENSMUSG00000029192 | 100855 | 0.206476677502759 | 3.46596428129758 | 2.61479471132771 | 0.0209655515818 | 0.0459175404409872 |
| Ddah1 | ENSMUSG00000028194 | 69219 | 0.339576356752721 | 3.18232315098926 | 2.61363054199334 | 0.0210124504758 | 0.0460110368892693 |
| Naglu | ENSMUSG00000001751 | 27419 | 0.361911904022699 | 1.71195780564327 | 2.61313451282344 | 0.0210324640352 | 0.0460352683325009 |
| Tbc1d12 | ENSMUSG00000048720 | 209478 | 0.309419499632233 | 2.51745224025147 | 2.61305541194404 | 0.0210356572693 | 0.0460352683325009 |
| Lrba | ENSMUSG00000028080 | 80877 | -0.216640543392492 | 5.70636137307055 | -2.61303118645042 | 0.0210366353255 | 0.0460352683325009 |
| Exoc8 | ENSMUSG00000074030 | 102058 | 0.267690522230385 | 3.27464543342738 | 2.61293888230271 | 0.0210403623263 | 0.0460352683325009 |
| Sacm1l | ENSMUSG00000025240 | 83493 | -0.266377051441396 | 4.60893731724862 | -2.61231110329809 | 0.0210657273900 | 0.0460815421454548 |
| Zfp971 | ENSMUSG00000074519 | 626848 | -0.331495866992974 | 3.36807569822062 | -2.61217092969486 | 0.0210713950736 | 0.0460847177970812 |
| Senp8 | ENSMUSG00000051705 | 71599 | -0.299610156889396 | 2.13639765876094 | -2.61187933108879 | 0.0210831901079 | 0.0460968423720559 |
| Eya2 | ENSMUSG00000017897 | 14049 | 0.560933502091768 | 1.82209580923408 | 2.61182538355488 | 0.0210853729631 | 0.0460968423720559 |
| Znrf1 | ENSMUSG00000033545 | 170737 | -0.152977892833735 | 5.44115378954151 | -2.61170515397844 | 0.0210902385474 | 0.0460969294729625 |
| Ide | ENSMUSG00000056999 | 15925 | -0.175931925467887 | 6.08777924025174 | -2.61161600649786 | 0.0210938469695 | 0.0460969294729625 |
| Rbpms | ENSMUSG00000031586 | 19663 | -0.385119130199193 | 2.46432496393792 | -2.61113874539306 | 0.0211131752437 | 0.0461299457504202 |
| Tmem183a | ENSMUSG00000042305 | 57439 | 0.312737648859012 | 2.81713663322277 | 2.61037733982403 | 0.0211440464337 | 0.0461881637823744 |
| Vps28 | ENSMUSG00000115987 | 66914 | -0.215610236115978 | 3.90238283860762 | -2.60852778915759 | 0.0212192187770 | 0.0463361350162593 |
| Zfp182 | ENSMUSG00000054737 | 319535 | 0.398052113621524 | 1.46114535608067 | 2.60850214834829 | 0.0212202627290 | 0.0463361350162593 |
| Reps2 | ENSMUSG00000040855 | 194590 | 0.528668268203005 | 1.48478717334382 | 2.60800375062459 | 0.0212405646175 | 0.0463712026830897 |
| Cisd2 | ENSMUSG00000028165 | 67006 | -0.216077745787891 | 4.95349030086383 | -2.60769780727031 | 0.0212530363312 | 0.0463891653626936 |
| Ufsp2 | ENSMUSG00000031634 | 192169 | -0.237717399758433 | 3.68693632447133 | -2.60668055026448 | 0.0212945556309 | 0.0464705106013297 |
| Capza2 | ENSMUSG00000015733 | 12343 | -0.187724345068768 | 6.41260477707469 | -2.60624201600097 | 0.0213124786207 | 0.0465003400804151 |
| Hdac1 | ENSMUSG00000028800 | 433759 | 0.315421129916847 | 3.23272370447196 | 2.60584610790434 | 0.0213286720232 | 0.0465163606488477 |
| Slc36a1 | ENSMUSG00000020261 | 215335 | 0.344096096967315 | 2.50008211090424 | 2.6058422465397 | 0.0213288300191 | 0.0465163606488477 |
| Cbfa2t2 | ENSMUSG00000038533 | 12396 | 0.223319656852311 | 4.04541376820591 | 2.60565475107586 | 0.0213365031582 | 0.0465163606488477 |
| Fbxl17 | ENSMUSG00000023965 | 50758 | -0.240487048781937 | 4.6833442575704 | -2.60564644518857 | 0.0213368431335 | 0.0465163606488477 |
| Wdr41 | ENSMUSG00000042015 | 218460 | -0.205991329323663 | 4.01868168625038 | -2.60509240012621 | 0.0213595330762 | 0.0465500203004099 |
| Hdgf | ENSMUSG00000004897 | 15191 | -0.173062399294655 | 6.61916090405849 | -2.605061487716 | 0.0213607997280 | 0.0465500203004099 |
| Rasa2 | ENSMUSG00000032413 | 114713 | 0.388340836848263 | 2.19594010440548 | 2.60415451559769 | 0.0213979957796 | 0.0466183628837028 |
| Arf2 | ENSMUSG00000062421 | 11841 | -0.312941160477544 | 3.31567009431155 | -2.60408887454766 | 0.0214006902342 | 0.0466183628837028 |
| Mettl4 | ENSMUSG00000055660 | 76781 | -0.268292617215862 | 2.95520817739205 | -2.60375500802927 | 0.0214143999807 | 0.0466389332913005 |
| Ergic2 | ENSMUSG00000030304 | 67456 | -0.211052755977389 | 5.12585621008792 | -2.60351069960565 | 0.0214244375438 | 0.0466437006295349 |
| Ppwd1 | ENSMUSG00000021713 | 238831 | -0.301006907239675 | 2.37292087536104 | -2.60349401704709 | 0.0214251231233 | 0.0466437006295349 |
| Grk4 | ENSMUSG00000052783 | 14772 | -0.923231219408215 | 2.40930886255085 | -2.60115619856106 | 0.0215214075850 | 0.0468439877165391 |
| Paox | ENSMUSG00000025464 | 212503 | -0.62090507741803 | 1.74112225155681 | -2.59973254115125 | 0.0215802469564 | 0.0469627074418893 |
| Zfand6 | ENSMUSG00000030629 | 65098 | -0.221425756187839 | 4.57909407230063 | -2.59908350898414 | 0.0216071229538 | 0.0470102433177579 |
| Slc26a2 | ENSMUSG00000034320 | 13521 | 0.32597585283383 | 3.32154201237388 | 2.59899738189794 | 0.0216106918554 | 0.0470102433177579 |
| Dnajc25 | ENSMUSG00000070972 | 72429 | -0.267991599784949 | 3.57616168977902 | -2.59876607203079 | 0.0216202796117 | 0.0470217422275323 |
| Vmn2r117 | ENSMUSG00000091407 | 619788 | -1.02218133997573 | -0.0591172005757814 | -2.5984322608836 | 0.0216341232915 | 0.0470424908892905 |
| Tnks | ENSMUSG00000031529 | 21951 | 0.168908229530791 | 4.65012034113799 | 2.59790829930087 | 0.0216558701259 | 0.0470804129568361 |
| Smpd1 | ENSMUSG00000037049 | 20597 | -0.228519968180194 | 4.76671436263713 | -2.59774516758447 | 0.0216626451686 | 0.0470857773590039 |
| Snx4 | ENSMUSG00000022808 | 69150 | -0.252188054605831 | 4.52375614688269 | -2.59761915088245 | 0.0216678801883 | 0.0470877929103348 |
| Paip2 | ENSMUSG00000037058 | 67869 | -0.189077424066507 | 5.70789427559881 | -2.59737564033794 | 0.0216779996399 | 0.0470884614761164 |
| Hist2h3c2 | ENSMUSG00000081058 | 97114 | 1.28771349953545 | 0.965568601968552 | 2.59734160027323 | 0.0216794145911 | 0.0470884614761164 |
| Copz2 | ENSMUSG00000018672 | 56358 | 0.283593489847075 | 3.76579209385468 | 2.59730078691224 | 0.0216811112075 | 0.0470884614761164 |
| Stxbp2 | ENSMUSG00000004626 | 20911 | 0.478816063223423 | 1.34612636439947 | 2.59555517235915 | 0.0217537970810 | 0.0472369399867704 |
| Tomm7 | ENSMUSG00000028998 | 66169 | -0.204475417359235 | 4.07602853335846 | -2.59350713709782 | 0.0218393762964 | 0.0474133510022963 |
| Pfkfb2 | ENSMUSG00000026409 | 18640 | -0.556369773273469 | 3.01945585724509 | -2.59324227567325 | 0.0218504675706 | 0.0474272001285464 |
| Eif3f | ENSMUSG00000031029 | 66085 | -0.172136110466675 | 5.92638099443002 | -2.59314761232644 | 0.0218544329930 | 0.0474272001285464 |
| Retreg2 | ENSMUSG00000049339 | 227298 | 0.318668717190915 | 2.65450029752403 | 2.59286453525301 | 0.0218662951733 | 0.0474435237276286 |
| Slc26a6 | ENSMUSG00000023259 | 171429 | -0.421789245715172 | 1.49623793947291 | -2.59247448841785 | 0.0218826500704 | 0.0474695868068696 |
| Tmem214 | ENSMUSG00000038828 | 68796 | 0.20177264692671 | 4.35377587992136 | 2.5921833098118 | 0.0218948670824 | 0.047486665094785 |
| Fam114a2 | ENSMUSG00000020523 | 67726 | -0.196225437327574 | 4.88082547592787 | -2.59175257848835 | 0.0219129514306 | 0.0475164594502002 |
| Mrpl40 | ENSMUSG00000022706 | 18100 | -0.276958185026354 | 3.15473103554948 | -2.59088726245847 | 0.0219493256215 | 0.0475858941629593 |
| Pgam5 | ENSMUSG00000029500 | 72542 | -0.249196533491744 | 3.9058544626104 | -2.5893552776647 | 0.0220138670920 | 0.047711636376347 |
| Yif1b | ENSMUSG00000030588 | 77254 | -0.201561587696113 | 3.85892311159844 | -2.58930342793426 | 0.0220160546960 | 0.047711636376347 |
| Haus8 | ENSMUSG00000035439 | 76478 | -0.353543235950255 | 2.26798405785087 | -2.58913923722592 | 0.0220229834923 | 0.0477171917849741 |
| Ets1 | ENSMUSG00000032035 | 23871 | -0.464428068658414 | 4.80612406708672 | -2.58859286114 | 0.0220460555909 | 0.0477577157479708 |
| Vmn2r56 | ENSMUSG00000090762 | 629079 | -0.864638484959136 | 0.707072952791489 | -2.58693779439041 | 0.0221160877656 | 0.0478999317150976 |
| Hmgxb3 | ENSMUSG00000024622 | 106894 | -0.229306533136577 | 4.00970515656454 | -2.58658675855277 | 0.0221309690758 | 0.0479226669906051 |
| Timeless | ENSMUSG00000039994 | 21853 | -0.346221070592886 | 2.63915590646463 | -2.58638652189575 | 0.0221394619610 | 0.0479315624274244 |
| Utp14b | ENSMUSG00000079470 | 195434 | 0.359016792229512 | 2.56457395412549 | 2.58556463703111 | 0.0221743546301 | 0.0479975981112841 |
| Hcfc1 | ENSMUSG00000031386 | 15161 | 0.167301011874302 | 5.92282618998312 | 2.58461186179787 | 0.0222148707733 | 0.0480757775537461 |
| Cry1 | ENSMUSG00000020038 | 12952 | -0.256248196758239 | 3.41302529764729 | -2.58430470756041 | 0.0222279475666 | 0.0480945555919869 |
| Dpysl2 | ENSMUSG00000022048 | 12934 | 0.318180201729611 | 5.39619872532149 | 2.58408754249256 | 0.0222371976521 | 0.0481050479984362 |
| Mettl21c | ENSMUSG00000047343 | 433294 | 0.447153458232104 | 5.70555488841544 | 2.58215848215218 | 0.0223195290697 | 0.0482735995768635 |
| Metrn | ENSMUSG00000002274 | 70083 | -0.366928322899826 | 2.18048577691556 | -2.58184209733779 | 0.0223330603469 | 0.0482933101192245 |
| Nab1 | ENSMUSG00000002881 | 17936 | 0.230856343605955 | 4.28121600136903 | 2.58008869807975 | 0.0224081944206 | 0.0484461972334199 |
| Polr2e | ENSMUSG00000004667 | 66420 | 0.181721312874816 | 4.4366164888909 | 2.57994691865882 | 0.0224142804154 | 0.0484497724383631 |
| Dpp9 | ENSMUSG00000001229 | 224897 | -0.190468477880541 | 4.77936762994107 | -2.57963600941203 | 0.0224276320402 | 0.0484690482072475 |
| Ctdspl | ENSMUSG00000047409 | 69274 | -0.45780746162497 | 2.25777268766538 | -2.57714293424216 | 0.0225349726483 | 0.0486913986991562 |
| Fbxo45 | ENSMUSG00000035764 | 268882 | -0.215382328976928 | 3.80666687710166 | -2.57635026442388 | 0.0225692054177 | 0.0487557281464352 |
| Vmp1 | ENSMUSG00000018171 | 75909 | -0.23879034252448 | 4.33764507829929 | -2.57591480479342 | 0.0225880328633 | 0.0487831788959827 |
| Cmpk2 | ENSMUSG00000020638 | 22169 | -0.653918338882704 | 1.90985552662396 | -2.57584995193827 | 0.0225908381274 | 0.0487831788959827 |
| Aff4 | ENSMUSG00000049470 | 93736 | 0.152610651245076 | 6.73746606231093 | 2.57427159050897 | 0.0226592154181 | 0.0489211700050372 |
| Nop53 | ENSMUSG00000041560 | 68077 | -0.185456138667792 | 5.59975651628016 | -2.57359088062038 | 0.0226887666430 | 0.0489752978228794 |
| Twnk | ENSMUSG00000025209 | 226153 | -0.276268409448548 | 3.92314415819892 | -2.57318349808246 | 0.0227064698998 | 0.0490038346447299 |
| Lyn | ENSMUSG00000042228 | 17096 | 0.307115013501181 | 3.18747029828558 | 2.57189279731848 | 0.0227626469438 | 0.04911537578815 |
| Tmem234 | ENSMUSG00000028797 | 76799 | 0.213574018358059 | 4.56462555780124 | 2.5712117852636 | 0.0227923417429 | 0.0491697429626903 |
| Nod1 | ENSMUSG00000038058 | 107607 | -0.321447717973433 | 3.25278042826101 | -2.57042521641272 | 0.0228266858346 | 0.0492341165860784 |
| Akap12 | ENSMUSG00000038587 | 83397 | -0.337146869873275 | 4.98503970242866 | -2.56807269216518 | 0.0229297032886 | 0.0494465548723559 |
| Utp18 | ENSMUSG00000054079 | 217109 | 0.329941319002907 | 2.94218375995831 | 2.56796837221536 | 0.0229342818665 | 0.0494466735864432 |
| Sox17 | ENSMUSG00000025902 | 20671 | -0.604067335642022 | 2.42784561850531 | -2.5676216544793 | 0.0229495055750 | 0.0494697387971624 |
| Fmo5 | ENSMUSG00000028088 | 14263 | -0.342531123445142 | 2.13125701504861 | -2.56713926456891 | 0.0229707026339 | 0.0495056684724173 |
| Vmn1r74 | ENSMUSG00000047655 | 171240 | -1.36421638284343 | 0.832930914472193 | -2.56614416833603 | 0.0230144886932 | 0.0495902574227915 |
| Btd | ENSMUSG00000021900 | 26363 | 0.421687900862738 | 1.53482063137845 | 2.5642027177787 | 0.0231001483608 | 0.0497650220205202 |
| Tm4sf1 | ENSMUSG00000027800 | 17112 | -0.396655693597484 | 3.98370561219435 | -2.56307096688098 | 0.0231502249211 | 0.0498630755888781 |
| Tmem198b | ENSMUSG00000047090 | 73827 | 0.463198362191579 | 1.34282910093318 | 2.56265888447083 | 0.0231684843560 | 0.0498925733838286 |
| Lamb1 | ENSMUSG00000002900 | 16777 | 0.380812217609672 | 5.55058113397043 | 2.56213844539051 | 0.0231915649741 | 0.0499324397739118 |
| Cutc | ENSMUSG00000025193 | 66388 | -0.360431538851227 | 2.56532087842258 | -2.56192053118202 | 0.0232012357071 | 0.0499434240871897 |
| Pik3r2 | ENSMUSG00000031834 | 18709 | 0.231283531028383 | 3.80838203294331 | 2.56052493649971 | 0.0232632627051 | 0.0500670849833701 |
| Cyb5d2 | ENSMUSG00000057778 | 192986 | -0.294347656914163 | 2.81343411563105 | -2.56015954898686 | 0.0232795287385 | 0.0500882589292917 |
| Prnp | ENSMUSG00000079037 | 19122 | 0.226783423761464 | 4.99173341073753 | 2.56009809789609 | 0.0232822654486 | 0.0500882589292917 |
| Capn5 | ENSMUSG00000035547 | 12337 | 0.471556997668503 | 1.48423296112387 | 2.55943717506259 | 0.0233117191456 | 0.0501417556042047 |
| Fgfrl1 | ENSMUSG00000008090 | 116701 | 0.198711449720204 | 5.28609903852051 | 2.55884308155733 | 0.0233382253382 | 0.0501888926169741 |
| Acap3 | ENSMUSG00000029033 | 140500 | 0.325149413102904 | 3.29316858368456 | 2.55822985510322 | 0.0233656156985 | 0.0502379121166075 |
| Lrrcc1 | ENSMUSG00000027550 | 71710 | 0.297531343527574 | 4.21038016149345 | 2.55791966182675 | 0.0233794826064 | 0.0502578415676394 |
| Mrps10 | ENSMUSG00000034729 | 64657 | -0.196942886041822 | 3.76301446378074 | -2.55773200316442 | 0.0233878755742 | 0.0502659984077867 |
| Smyd4 | ENSMUSG00000018809 | 319822 | -0.380246916619872 | 1.67389780081652 | -2.5572964384148 | 0.0234073672707 | 0.0502980011079274 |
| Sf3a2 | ENSMUSG00000020211 | 20222 | -0.194130320924966 | 4.07869883652262 | -2.55671494586632 | 0.0234334137473 | 0.0503440734418888 |
| Vps53 | ENSMUSG00000017288 | 68299 | 0.182128232260756 | 4.04625050533708 | 2.55651341945273 | 0.0234424471384 | 0.0503535841364136 |
| Sh3kbp1 | ENSMUSG00000040990 | 58194 | 0.354519451781078 | 3.58753754083303 | 2.55561547213892 | 0.0234827383792 | 0.0504302187079775 |
| Rnaseh2b | ENSMUSG00000021932 | 67153 | 0.452339522188733 | 1.40890061626415 | 2.55468891100351 | 0.0235243836274 | 0.0505048448943047 |
| Lsm4 | ENSMUSG00000031848 | 50783 | -0.240916491113461 | 3.80735835338615 | -2.55463678529942 | 0.0235267285885 | 0.0505048448943047 |
| Steap4 | ENSMUSG00000012428 | 117167 | -0.343036704640434 | 4.30074346892139 | -2.55440889079655 | 0.0235369834509 | 0.0505127405230898 |
| Vmn2r79 | ENSMUSG00000090362 | 621430 | -0.970624030006831 | 3.38113649091101 | -2.55430948370333 | 0.0235414579516 | 0.0505127405230898 |
| Hspa1a | ENSMUSG00000091971 | 193740 | 0.323335943016723 | 5.80180712424653 | 2.55417580380287 | 0.0235474764309 | 0.0505127405230898 |
| Cebpz | ENSMUSG00000024081 | 12607 | -0.164993073661658 | 4.55201488204232 | -2.55414439247174 | 0.0235488908339 | 0.0505127405230898 |
| Scp2 | ENSMUSG00000028603 | 20280 | -0.2047042928867 | 5.86508862564735 | -2.55365913749318 | 0.0235707515294 | 0.0505497125698917 |
| Klf9 | ENSMUSG00000033863 | 16601 | -0.283229191077136 | 6.56431026282924 | -2.55343180397352 | 0.0235809996234 | 0.0505617706716751 |
| Wbp11 | ENSMUSG00000030216 | 60321 | -0.176417827426741 | 5.03123106913671 | -2.55249233864328 | 0.0236233959413 | 0.0506427419169737 |
| Ralgps2 | ENSMUSG00000026594 | 78255 | 0.689356678433539 | 0.724940126485555 | 2.55131268315208 | 0.0236767358118 | 0.0507471370901468 |
| Card10 | ENSMUSG00000033170 | 105844 | -0.449298370683665 | 3.16807262082943 | -2.54975315999764 | 0.0237474304286 | 0.0508886810459286 |
| Ccdc186 | ENSMUSG00000035173 | 213993 | 0.225503421981742 | 4.48007944582854 | 2.54868166424676 | 0.0237961203479 | 0.0509830246027804 |
| Hspa5 | ENSMUSG00000026864 | 14828 | 0.387215512029855 | 7.57971086694786 | 2.54752108151651 | 0.0238489671711 | 0.0510862355766309 |

| | | | | | | | |
|---|---|---|---|---|---|---|---|
| Vmn2r17 | ENSMUSG00000091879 | 384221 | -1.52187464845313 | -0.921109647366357 | -2.54723456700674 | 0.0238620309542 | 0.0511042046162239 |
| Metrnl | ENSMUSG00000039208 | 210029 | 0.450247997951662 | 2.9873592226753 | 2.54651429124726 | 0.0238949028235 | 0.051164580365185 |
| Dusp22 | ENSMUSG00000069255 | 105352 | -0.17086140440215 | 4.71211064422183 | -2.54594334797372 | 0.0239209905159 | 0.0512104087993213 |
| Castor2 | ENSMUSG00000015944 | 80909 | -0.279292883942121 | 5.69957068146051 | -2.54583938324762 | 0.0239257438568 | 0.051210555335765 |
| Dipk1a | ENSMUSG00000029270 | 67266 | -0.289444600090633 | 2.57461686217599 | -2.54566490633755 | 0.0239337231132 | 0.0512176051979843 |
| Mcmdc2 | ENSMUSG00000046101 | 240697 | -1.52951354443146 | 1.33171631760045 | -2.54519280843837 | 0.0239553261819 | 0.0512538012321082 |
| Apba1 | ENSMUSG00000024897 | 319924 | 0.530282931860848 | 2.25777352801247 | 2.54362643488196 | 0.0240271378542 | 0.0513898101492844 |
| Sik1 | ENSMUSG00000024042 | 17691 | 0.708884743158047 | 5.42602426452733 | 2.5436011780824 | 0.0240282974726 | 0.0513898101492844 |
| Txndc9 | ENSMUSG00000058407 | 98258 | -0.253217383892629 | 4.0137472316282 | -2.54298262092219 | 0.0240567142186 | 0.0514405209554835 |
| Nos2 | ENSMUSG00000020826 | 18126 | -0.694979259132864 | 1.25395516881022 | -2.54280120598481 | 0.0240650546355 | 0.0514482910660518 |
| Glcci1 | ENSMUSG00000029638 | 170772 | 0.60307174350664 | 1.98688134428893 | 2.54106894185089 | 0.0241448348055 | 0.051608758165798 |
| Uba52 | ENSMUSG00000068240 | 22186 | -0.686232473401045 | 3.55211704845471 | -2.54013935606053 | 0.0241877524083 | 0.0516875680879877 |
| Clec11a | ENSMUSG00000004473 | 20256 | 0.744555008169397 | 1.83918144948529 | 2.54006556039199 | 0.0241911625961 | 0.0516875680879877 |
| Dhx34 | ENSMUSG00000006019 | 71723 | 0.353488980084012 | 1.81550888094476 | 2.53981466550242 | 0.0242027602237 | 0.0517022419398344 |
| Lgals8 | ENSMUSG00000057554 | 56048 | 0.178036205592127 | 3.80065333813643 | 2.5393915234814 | 0.0242223321332 | 0.0517339414905903 |
| Zbtb41 | ENSMUSG00000033964 | 226470 | -0.191657070386938 | 4.58520297466716 | -2.53909533258924 | 0.0242360411572 | 0.0517531091795753 |
| Ccdc102a | ENSMUSG00000063605 | 234582 | 0.436014967843596 | 1.87588396247496 | 2.53889969057865 | 0.0242451004380 | 0.0517623423610955 |
| Rgma | ENSMUSG00000070509 | 244058 | 0.357257848148618 | 4.55056975862902 | 2.53867156447388 | 0.0242556680322 | 0.0517647843384845 |
| Srrm4 | ENSMUSG00000063919 | 68955 | 1.11415050084533 | 0.364469504042916 | 2.53867054146573 | 0.0242557154314 | 0.0517647843384845 |
| Chmp7 | ENSMUSG00000034190 | 105513 | -0.190644832771985 | 4.65443679176497 | -2.5364878978832 | 0.0243570480843 | 0.0519708945168922 |
| Nudt15 | ENSMUSG00000033405 | 214254 | -0.371688914066715 | 1.76274368524795 | -2.53633840180107 | 0.0243640035929 | 0.0519755900222601 |
| Mrpl53 | ENSMUSG00000030037 | 68499 | -0.254929198514534 | 3.90344846121468 | -2.53603582695716 | 0.0243780871607 | 0.0519954869765263 |
| Eef1akmt2 | ENSMUSG00000030960 | 72096 | 0.375023334082069 | 1.50217129987729 | 2.53589269459201 | 0.0243847520964 | 0.0519995562165408 |
| Ubr4 | ENSMUSG00000066036 | 69116 | 0.206109920070405 | 7.76576939947026 | 2.53529791339558 | 0.0244124668132 | 0.0520485029715448 |
| Cdkn2d | ENSMUSG00000096472 | 12581 | 0.41815079483308 | 1.64549644245785 | 2.53494398333933 | 0.0244289731235 | 0.0520735384580849 |
| Gm14295 | ENSMUSG00000078877 | 100039123 | -0.312688843201306 | 2.57915014024476 | -2.53395054073802 | 0.0244753620279 | 0.0521605707850141 |
| Olfr775 | ENSMUSG00000096220 | 258538 | -1.2624118577768 | 0.0437104284765288 | -2.53386531243457 | 0.0244793457256 | 0.0521605707850141 |
| Ankrd27 | ENSMUSG00000034867 | 245886 | -0.240531153821824 | 3.8540770966108 | -2.53331146847263 | 0.0245052484511 | 0.0521956455210052 |
| Pole | ENSMUSG00000007080 | 18973 | -0.468088546685978 | 1.86401535949792 | -2.5333091576057 | 0.0245053565834 | 0.0521956455210052 |
| Tfeb | ENSMUSG00000023990 | 21425 | -0.225279874323466 | 4.46873935939624 | -2.53298870559978 | 0.0245203559312 | 0.052217418796855 |
| Zyg11b | ENSMUSG00000034636 | 414872 | -0.205887153643523 | 6.90093327438616 | -2.53255678394705 | 0.0245405868268 | 0.052250322283758 |
| Ppm1g | ENSMUSG00000029147 | 14208 | -0.168995243901473 | 4.9445506994052 | -2.53242212802465 | 0.0245468973019 | 0.0522535802156829 |
| Rpl10a | ENSMUSG00000037805 | 19896 | 0.218479315834052 | 5.4953499641009 | 2.53202941848124 | 0.0245653100599 | 0.0522825942884423 |
| Pik3cb | ENSMUSG00000032462 | 74769 | -0.244715287839697 | 2.51326476537079 | -2.53132313259373 | 0.0245984588119 | 0.0523429537226802 |
| Plekhg5 | ENSMUSG00000039713 | 269608 | 0.255154606936663 | 3.66884977361422 | 2.53049189039971 | 0.0246375274888 | 0.0524158841924296 |
| LOC102633794 | ENSMUSG00000086735 | 102633794 | -1.0416377000759 | 0.612149456831123 | -2.52965191877731 | 0.0246770672051 | 0.052489788211718 |
| Ttc28 | ENSMUSG00000033209 | 209683 | 0.246203365119467 | 4.82673868935616 | 2.52926671382406 | 0.0246952202840 | 0.0525129161371172 |
| Tsen2 | ENSMUSG00000042389 | 381802 | 0.402792379704103 | 1.86687725817231 | 2.52921733079903 | 0.0246975484274 | 0.0525129161371172 |
| Rdx | ENSMUSG00000032050 | 19684 | -0.162486707235195 | 7.2019105466181 | -2.52855901440742 | 0.0247286047111 | 0.0525687238618293 |
| Gm10125 | ENSMUSG00000063087 | 791318 | -1.0617320902072 | 2.80972988757666 | -2.52840107568581 | 0.0247360611163 | 0.0525743503913803 |
| Myef2 | ENSMUSG00000027201 | 17876 | 0.303260245606078 | 2.29669694483452 | 2.52767691448174 | 0.0247702770613 | 0.0526368387553907 |
| Olfr1116 | ENSMUSG00000070855 | 257875 | -1.16630923172061 | -0.122038242481307 | -2.52711592218878 | 0.0247968147754 | 0.0526829897591103 |
| Tsen15 | ENSMUSG00000014980 | 66637 | -0.323435047298373 | 2.50250353167511 | -2.52693065025815 | 0.0248055850723 | 0.0526913817384088 |
| Mtmr12 | ENSMUSG00000039458 | 268783 | 0.354433200715395 | 2.767723642968 | 2.52650815999166 | 0.0248255958570 | 0.0527236425711874 |
| Rab22a | ENSMUSG00000027519 | 19334 | -0.159490779471736 | 5.41459442290096 | -2.52590567820819 | 0.0248541586510 | 0.0527740497697368 |
| Zfp952 | ENSMUSG00000053390 | 240067 | -0.369167234372178 | 1.80150699131334 | -2.52523824015842 | 0.0248858378988 | 0.0528310534224834 |
| Carf | ENSMUSG00000026017 | 241066 | -0.333881378768865 | 2.2113422369529 | -2.52410708169977 | 0.0249396160126 | 0.0529349403173237 |
| Mmp15 | ENSMUSG00000031790 | 17388 | -0.271558027227539 | 4.33363182951534 | -2.52113619561974 | 0.0250813931264 | 0.0532255306280963 |
| St6galnac2 | ENSMUSG00000057286 | 20446 | -0.574855287290324 | 1.18622640656823 | -2.51939142353872 | 0.0251650189626 | 0.0533926285482116 |
| Ccdc14 | ENSMUSG00000022833 | 239839 | -0.637362803899587 | 2.00037607114361 | -2.51871872478805 | 0.0251973326698 | 0.0534508137811238 |
| Txnrd3 | ENSMUSG00000000811 | 232223 | 0.258990226174327 | 2.26148075447596 | 2.51696653419508 | 0.0252816884781 | 0.0536155825241327 |
| Ift172 | ENSMUSG00000038564 | 67661 | -0.180986864702552 | 4.46234768294237 | -2.51690167139659 | 0.0252848163856 | 0.0536155825241327 |
| C1galt1 | ENSMUSG00000042460 | 94192 | 0.154353117190534 | 4.45050326323407 | 2.51640194037895 | 0.0253089276351 | 0.0536563010046378 |
| Nacc2 | ENSMUSG00000026932 | 67991 | -0.183433381281844 | 4.34691189092709 | -2.51612732771328 | 0.0253221866964 | 0.05366469902896 |
| Sae1 | ENSMUSG00000052833 | 56459 | 0.212119472941918 | 3.25015046787796 | 2.51611654039406 | 0.0253227076746 | 0.05366469902896 |
| Nacc1 | ENSMUSG00000001910 | 66830 | 0.180332503885633 | 5.51064322530917 | 2.51483393537013 | 0.0253847252549 | 0.0537857010587596 |
| Lig1 | ENSMUSG00000056394 | 16881 | 0.623255930383178 | 0.924721539349175 | 2.51375279524506 | 0.0254371150024 | 0.0538843167584548 |
| Commd3 | ENSMUSG00000051154 | 12238 | -0.220141329258885 | 5.35860535221504 | -2.51367009096282 | 0.0254411269591 | 0.0538843167584548 |
| Arfgap2 | ENSMUSG00000027255 | 77038 | -0.207312457093385 | 5.35533946785382 | -2.51159760988598 | 0.0255418614400 | 0.0540871924450495 |
| Vasp | ENSMUSG00000030403 | 22323 | 0.262819649146532 | 3.62754010072325 | 2.51131405422407 | 0.0255556736978 | 0.0541059595567174 |
| Zfp185 | ENSMUSG00000031351 | 22673 | 0.579393078684042 | 2.10035061201485 | 2.51053855473458 | 0.0255934857229 | 0.0541755213860928 |
| Fam118a | ENSMUSG00000022434 | 73225 | 0.339432672339067 | 2.25449806630946 | 2.51039224184542 | 0.0256006257243 | 0.0541801432319553 |
| Gon7 | ENSMUSG00000091931 | 100233175 | -0.450790770496223 | 1.28927544837432 | -2.50981635724985 | 0.0256287472548 | 0.0542291591642098 |
| Cyp2e1 | ENSMUSG00000025479 | 13106 | 0.824917185924825 | 1.27636730073637 | 2.50963567359414 | 0.0256375765042 | 0.0542319803305763 |
| Brwd3 | ENSMUSG00000063663 | 382236 | 0.326904289350755 | 2.79056961296019 | 2.5095860248433 | 0.0256400031422 | 0.0542319803305763 |
| Gm14430 | ENSMUSG00000078899 | 627914 | -0.156261488729232 | 5.48493383588408 | -2.50943079695995 | 0.0256475915043 | 0.0542375358356206 |
| Papss2 | ENSMUSG00000024899 | 23972 | -0.36622907578557 | 2.49312095035784 | -2.50776706287106 | 0.0257290594985 | 0.0543992938837021 |
| Hprt | ENSMUSG00000025630 | 15452 | 0.217550959231991 | 3.81654249408541 | 2.5058785758317 | 0.0258218347277 | 0.0545848917827509 |
| Mtg2 | ENSMUSG00000039069 | 52856 | -0.313072754626576 | 2.70420163718375 | -2.5050326151989 | 0.0258634983401 | 0.054662393727054 |
| Klhl11 | ENSMUSG00000048732 | 217194 | 0.315356247683728 | 1.80780525154015 | 2.5045773686011 | 0.0258859460239 | 0.0546992607748112 |
| Tomm70a | ENSMUSG00000022752 | 28185 | -0.15856222583964 | 5.72194483258718 | -2.50443485897416 | 0.0258929768605 | 0.0547035427257686 |
| Rgs12 | ENSMUSG00000029101 | 71729 | -0.338708810215969 | 2.43502162573285 | -2.50432469659956 | 0.0258984130763 | 0.0547044547512011 |
| Gtf2a1 | ENSMUSG00000020962 | 83602 | 0.181572834954157 | 4.74103640080877 | 2.50262072159936 | 0.0259826396171 | 0.0548717607525162 |
| Ppp2r5d | ENSMUSG00000059409 | 21770 | -0.188296106235145 | 4.58668220152116 | -2.50251460604617 | 0.0259878935571 | 0.0548722550653242 |
| Brd4 | ENSMUSG00000024002 | 57261 | 0.150374533004341 | 5.77329579432593 | 2.50191625040738 | 0.0260175381748 | 0.054924239047741 |
| Prx | ENSMUSG00000053198 | 19153 | -0.736717722798525 | 2.1263104655726 | -2.50159979191138 | 0.0260332297769 | 0.0549406792502351 |
| Cactin | ENSMUSG00000034889 | 70312 | -0.237931623264148 | 3.27191209046755 | -2.50155647921299 | 0.0260353781462 | 0.0549406792502351 |
| Fastkd3 | ENSMUSG00000021532 | 69577 | -0.453955406882236 | 1.33608023722348 | -2.49995646762141 | 0.0261148605320 | 0.0550821248007538 |
| Med12 | ENSMUSG00000079487 | 59024 | 0.18188729650666 | 4.6949075459501 | 2.49994200917513 | 0.0261155798346 | 0.0550821248007538 |
| Psip1 | ENSMUSG00000028484 | 101739 | 0.173920845238216 | 4.09636808091 | 2.49990293579262 | 0.0261175238168 | 0.0550821248007538 |
| Ube2c | ENSMUSG00000001403 | 68612 | -0.38755032240869 | 2.22644196855732 | -2.49922415500261 | 0.0261513167940 | 0.0551427553773418 |
| Kcnc4 | ENSMUSG00000027895 | 99738 | -0.497669253852757 | 1.69725610479168 | -2.49862965616048 | 0.0261809483498 | 0.0551945894719057 |
| Pigt | ENSMUSG00000017721 | 78928 | 0.198569195676526 | 4.33004841201851 | 2.49843403773714 | 0.0261907056008 | 0.0552045127116101 |
| Tsc2 | ENSMUSG00000002496 | 22084 | -0.142054537014635 | 6.24973725168094 | -2.49654021941327 | 0.0262853483400 | 0.0553933184316896 |
| Macrod2 | ENSMUSG00000068205 | 72899 | -0.481200659827022 | 1.80774066142421 | -2.49441625236268 | 0.0263918838817 | 0.0556071092349911 |
| Mfsd11 | ENSMUSG00000020818 | 69900 | 0.25197042882805 | 3.31134944177554 | 2.49255746555977 | 0.0264854588253 | 0.0557935152091263 |
| Rpgrip1l | ENSMUSG00000033282 | 244585 | 0.43382470001434 | 1.46600306436911 | 2.49233205529453 | 0.0264968280766 | 0.0558067105405833 |
| Arhgef1 | ENSMUSG00000040940 | 16801 | 0.182432283610864 | 4.610319101047 | 2.49202607547191 | 0.0265122686046 | 0.0558284739197188 |
| Sgta | ENSMUSG00000004937 | 52551 | -0.188061800719926 | 5.77124536464763 | -2.49130477524869 | 0.0265487014996 | 0.0558934823258469 |
| Pip4p1 | ENSMUSG00000035953 | 219024 | 0.439694729318223 | 2.33284949326384 | 2.4912124779096 | 0.0265533669122 | 0.0558934823258469 |
| Agrn | ENSMUSG00000041936 | 11603 | -0.397193997193554 | 3.92122621619714 | -2.49078395329795 | 0.0265750381446 | 0.0559283292180929 |
| Gcsh | ENSMUSG00000034424 | 68133 | -0.194275460887732 | 4.54458013971106 | -2.49015117003747 | 0.0266070701783 | 0.0559849632569266 |
| Ticam1 | ENSMUSG00000047123 | 106759 | 0.308749459633262 | 2.12510744854869 | 2.49000507594746 | 0.0266144708579 | 0.055989757688301 |
| Wfdc1 | ENSMUSG00000023336 | 67866 | -0.228767071483992 | 4.26719819544094 | -2.48869451616308 | 0.0266809483398 | 0.0561188082168564 |
| Zfp638 | ENSMUSG00000030016 | 18139 | -0.14637040749499 | 5.65387518958317 | -2.48662238499969 | 0.0267863818547 | 0.0563284320411729 |
| Rab5if | ENSMUSG00000027637 | 67388 | -0.209299800179626 | 3.9523837597834 | -2.48653342171063 | 0.0267909174075 | 0.0563284320411729 |
| Ids | ENSMUSG00000035847 | 15931 | 0.20466687562382 | 3.81869791051888 | 2.48574792130338 | 0.0268309960557 | 0.0564018495932935 |
| Slc35e1 | ENSMUSG00000019731 | 270066 | -0.203361348478134 | 5.69477929214348 | -2.48521730018318 | 0.0268581025721 | 0.0564479752385396 |
| Stxbp4 | ENSMUSG00000020546 | 20913 | -0.379464838225752 | 1.91515701819684 | -2.48449487066276 | 0.0268950498228 | 0.0565147615558395 |
| Zscan22 | ENSMUSG00000054715 | 232878 | -0.485132547016912 | 1.32396081766132 | -2.48123018468777 | 0.0270626255751 | 0.056855960054181 |
| Fam126b | ENSMUSG00000038174 | 213056 | -0.178843290503836 | 3.78278586973883 | -2.48112217871131 | 0.0270681866056 | 0.0568567155622889 |
| Cab39l | ENSMUSG00000021981 | 69008 | -0.192440750489081 | 3.81885889216897 | -2.48099190189671 | 0.0270748957812 | 0.0568583856999524 |
| Ahdc1 | ENSMUSG00000037692 | 230793 | 0.277897733986257 | 5.49572216168328 | 2.48082131459137 | 0.0270836833372 | 0.0568583856999524 |
| Wdyhv1 | ENSMUSG00000022359 | 76773 | 0.409369331406875 | 1.77163798064886 | 2.48080378630106 | 0.0270845864367 | 0.0568583856999524 |
| Agfg2 | ENSMUSG00000029722 | 231801 | 0.458541653096521 | 1.63743116550206 | 2.48049093339284 | 0.0271007102330 | 0.0568813102068022 |
| Olfr1120 | ENSMUSG00000081234 | 259031 | -1.23073163371482 | 0.969095842497218 | -2.47992715316633 | 0.0271297895854 | 0.0569314129312891 |
| Endov | ENSMUSG00000039850 | 338371 | 0.289995926270644 | 2.13906611249887 | 2.47945161819509 | 0.0271543406105 | 0.0569719956265508 |
| Pcmt1 | ENSMUSG00000019795 | 18537 | -0.184061209952312 | 5.43193179402228 | -2.47827906723165 | 0.0272149685305 | 0.0570882404542774 |
| Zbtb39 | ENSMUSG00000044617 | 320080 | -0.250791908980614 | 2.97123856205575 | -2.47734054548302 | 0.0272635893737 | 0.0571792585272887 |
| Fubp1 | ENSMUSG00000028034 | 51886 | 0.213306858433355 | 5.11137567037958 | 2.47706518610991 | 0.0272778703967 | 0.0571982354319429 |
| Vmn1r180 | ENSMUSG00000092473 | 232962 | -1.85803276587879 | -1.05732287529236 | -2.47678203539804 | 0.0272925629938 | 0.0572180679105994 |
| Sms | ENSMUSG00000071708 | 20603 | 0.177719454811455 | 4.59487596414863 | 2.47614139792761 | 0.0273258335103 | 0.0572768333847524 |
| Ass1 | ENSMUSG00000076441 | 11898 | 0.452682300357926 | 1.66266062415801 | 2.47541980850019 | 0.0273633547726 | 0.0573444844745994 |
| Gsr | ENSMUSG00000031584 | 14782 | 0.163281265278728 | 4.31179997475797 | 2.47470593556425 | 0.0274005234611 | 0.057408751373907 |
| Chtop | ENSMUSG00000001017 | 66511 | -0.144988310603409 | 5.01473427588858 | -2.47462913292601 | 0.0274045251732 | 0.057408751373907 |
| Fosl2 | ENSMUSG00000029135 | 14284 | 0.3935418609071 | 4.3965573415248 | 2.4742870632955 | 0.0274223551285 | 0.0574350955950533 |

| | | | | | | | |
|---|---|---|---|---|---|---|---|
| Etv1 | ENSMUSG00000004151 | 14009 | 0.37504441741895 | 2.16581979314689 | 2.47364039651328 | 0.0274560922625 | 0.0574947403299629 |
| Olfr338 | ENSMUSG00000068950 | 258949 | -0.916883018854453 | 2.1686623578541 | -2.47325444828553 | 0.0274762464786 | 0.0575203740666222 |
| Mrpl23 | ENSMUSG00000037772 | 19935 | -0.392733452856367 | 2.62568583521805 | -2.47320446418192 | 0.0274788576869 | 0.0575203740666222 |
| Npc1 | ENSMUSG00000024413 | 18145 | -0.165813369586787 | 4.32409780676776 | -2.47273708317408 | 0.0275032855626 | 0.0575604852546073 |
| Rlf | ENSMUSG00000049878 | 109263 | 0.217717934704241 | 3.89142893535197 | 2.47242491971256 | 0.0275196125329 | 0.057583630282771 |
| Cul4b | ENSMUSG00000031095 | 72584 | 0.192481619404453 | 3.60109738331206 | 2.4721279427778 | 0.0275351538427 | 0.0575950860643889 |
| Zscan21 | ENSMUSG00000037017 | 22697 | -0.223302026970847 | 3.33422567192717 | -2.47211893694265 | 0.0275356252650 | 0.0575950860643889 |
| Rnf34 | ENSMUSG00000029474 | 80751 | -0.290284928200972 | 3.26598600063474 | -2.47116032398094 | 0.0275858494451 | 0.057689098964048 |
| Asxl2 | ENSMUSG00000037486 | 75302 | 0.190577040219326 | 4.57457987255327 | 2.47038281963363 | 0.0276266494445 | 0.0577633712849823 |
| 2310039H08Rik | ENSMUSG00000062619 | 67101 | -0.327748867507651 | 2.23145469513732 | -2.46904349175248 | 0.0276970673004 | 0.0578995300557213 |
| Slc35f6 | ENSMUSG00000029175 | 74919 | 0.259968142313049 | 3.17919780081534 | 2.46763621406047 | 0.0277712433074 | 0.0580434915093813 |
| Gtpbp2 | ENSMUSG00000023952 | 56055 | -0.197896307595779 | 4.07982326812887 | -2.46669242032364 | 0.0278210963424 | 0.0581365712329268 |
| Olfr716 | ENSMUSG00000073896 | 258597 | -1.39173362168723 | -0.374728381278125 | -2.46552475263231 | 0.0278828936720 | 0.0582545700934189 |
| Arhgef25 | ENSMUSG00000019467 | 52666 | 0.30602234037043 | 2.882426333179 | 2.46478404647151 | 0.0279221628443 | 0.0583254657082207 |
| Pfdn6 | ENSMUSG00000024309 | 14976 | -0.221684603737276 | 2.99164682869599 | -2.46383479896765 | 0.0279725655807 | 0.0584195862367412 |
| Rnf141 | ENSMUSG00000030788 | 67150 | -0.176639331320353 | 3.62280706195684 | -2.4632694264667 | 0.0280026269443 | 0.0584711967772039 |
| Psmd11 | ENSMUSG00000017428 | 69077 | 0.18472777962698 | 6.12451359874426 | 2.46281800311673 | 0.0280266517754 | 0.0585101853623458 |
| Tbc1d23 | ENSMUSG00000022749 | 67581 | 0.259066720549904 | 3.40402298132103 | 2.46222722019377 | 0.0280581232159 | 0.058564702095293 |
| Rest | ENSMUSG00000029249 | 19712 | 0.201548981908867 | 4.38590671580387 | 2.46084334953657 | 0.0281319757617 | 0.0587076416670912 |
| Sgpp1 | ENSMUSG00000021054 | 81535 | 0.210060993376412 | 3.85957112146147 | 2.46073262590311 | 0.0281378927594 | 0.0587087813997446 |
| Ranbp3 | ENSMUSG00000002372 | 71810 | -0.165552397655869 | 4.18906862828256 | -2.46058712452651 | 0.0281456700712 | 0.0587138014406907 |
| Rab10 | ENSMUSG00000020671 | 19325 | -0.151769826587405 | 7.25100597920443 | -2.45761370814031 | 0.0283050567791 | 0.0590350268370159 |
| Slc25a32 | ENSMUSG00000022299 | 69906 | 0.215747663699139 | 3.30409426219122 | 2.45722569761368 | 0.0283259193800 | 0.0590672691231355 |
| Amotl2 | ENSMUSG00000032531 | 56332 | -0.428142671021985 | 4.53564447151263 | -2.45702373499443 | 0.0283367843658 | 0.0590701535195904 |
| Inpp4b | ENSMUSG00000037940 | 234515 | -0.248972489199153 | 6.70045498549988 | -2.45699909076746 | 0.0283381104250 | 0.0590701535195904 |
| Ppp4r1l-ps | ENSMUSG00000055897 | 100043911 | 0.47495899279771 | 2.2041637020198 | 2.45645916744365 | 0.0283671776131 | 0.0591194696834794 |
| Ubn1 | ENSMUSG00000039473 | 170644 | 0.162274180869332 | 5.41184203304631 | 2.45633316318222 | 0.0283739652622 | 0.0591223435534368 |
| Nprl2 | ENSMUSG00000010057 | 56032 | -0.386369589868355 | 1.96863419692725 | -2.45534734449871 | 0.0284271235142 | 0.059221819541594 |
| Zfp280c | ENSMUSG00000036916 | 208968 | 0.311753590280444 | 2.12797360761756 | 2.45483587759969 | 0.0284547409204 | 0.0592680588798017 |
| Vmn1r197 | ENSMUSG00000069294 | 171278 | -1.07254607968615 | 1.7808324998351 | -2.4546729538771 | 0.0284635436258 | 0.0592750991378705 |
| Pusl1 | ENSMUSG00000051557 | 433813 | 0.382877475218175 | 1.5133861938416 | 2.45399787680314 | 0.0285000456014 | 0.0593398092323587 |
| Xpo4 | ENSMUSG00000021952 | 57258 | 0.289498511890921 | 6.56150036025102 | 2.4535870130007 | 0.0285222833062 | 0.0593641119612461 |
| Oga | ENSMUSG00000025220 | 76055 | 0.160528555450101 | 5.65177695200666 | 2.4535815434329 | 0.0285225794548 | 0.0593641119612461 |
| Kbtbd3 | ENSMUSG00000025893 | 69149 | -0.313326677077649 | 1.96144370750055 | -2.45318447113778 | 0.0285440867205 | 0.059397565570511 |
| Dpp4 | ENSMUSG00000035000 | 13482 | 0.42801977551792 | 1.91173772811083 | 2.45110737610877 | 0.0286568453064 | 0.0596208557374744 |
| Vmn1r183 | ENSMUSG00000118318 | 209824 | -0.946794066375418 | 0.892416166042919 | -2.45074582945556 | 0.0286765160404 | 0.059650427561946 |
| Rbbp6 | ENSMUSG00000030779 | 19647 | -0.153534832715588 | 5.28896587911658 | -2.45051655905625 | 0.0286889966998 | 0.0596650348032722 |
| Nsd1 | ENSMUSG00000021488 | 18193 | 0.167412464158637 | 6.9195444205528 | 2.44889317818756 | 0.0287775167595 | 0.0598377469467112 |
| Lhcgr | ENSMUSG00000024107 | 16867 | -1.23411949508144 | -0.180325503031218 | -2.44829148221721 | 0.0288103925928 | 0.0598947130909786 |
| Mknk1 | ENSMUSG00000028708 | 17346 | 0.300248317458891 | 3.03602947703262 | 2.44808942010902 | 0.0288214410550 | 0.0598953801088583 |
| Cited4 | ENSMUSG00000070803 | 56222 | 0.435912180091699 | 1.37365170011127 | 2.44808519263175 | 0.0288216722507 | 0.0598953801088583 |
| Col22a1 | ENSMUSG00000079022 | 69700 | 0.449349050404918 | 5.29500272865773 | 2.44759915342023 | 0.0288482650002 | 0.0599392481880735 |
| Nbeal1 | ENSMUSG00000073664 | 269198 | 0.189618464328985 | 5.96206768725356 | 2.44735998466603 | 0.0288613593105 | 0.0599550586514687 |
| Rtl8a | ENSMUSG00000067925 | 66158 | 0.226806899339587 | 3.64258360022481 | 2.44703899510533 | 0.0288789421982 | 0.0599801856677485 |
| Rhbdf1 | ENSMUSG00000020282 | 13650 | 0.267944815471473 | 3.6834015871856 | 2.44611280186426 | 0.0289297339495 | 0.0600742632603981 |
| Zfp385b | ENSMUSG00000027016 | 241494 | 0.308137137125799 | 2.62969367058785 | 2.4457335416247 | 0.0289505569855 | 0.0601060851678086 |
| Uap1l1 | ENSMUSG00000026956 | 227620 | 0.382704931671016 | 2.12179283831207 | 2.44458363420225 | 0.0290137796927 | 0.0602259069162854 |
| Dach1 | ENSMUSG00000055639 | 13134 | -0.451348610222063 | 2.32577742870223 | -2.44375701526858 | 0.0290593094526 | 0.0603089636656032 |
| Rab11b | ENSMUSG00000077450 | 19326 | -0.139631670680414 | 5.93117381223436 | -2.44305628067939 | 0.0290979591491 | 0.0603777128814976 |
| Ptbp2 | ENSMUSG00000028134 | 56195 | 0.207077664441889 | 3.39398247349835 | 2.44290940061768 | 0.0291060667004 | 0.0603830736576667 |
| Fam185a | ENSMUSG00000047221 | 330050 | -0.319707120302751 | 2.48935816414534 | -2.44184038051366 | 0.0291651401552 | 0.0604941455479058 |
| Rpusd3 | ENSMUSG00000051169 | 101122 | -0.500045964000446 | 1.19865673784609 | -2.44133460672084 | 0.0291931289271 | 0.0605236618597527 |
| Sec14l1 | ENSMUSG00000020823 | 74136 | 0.22762760488482 | 5.37180883878232 | 2.44129894206257 | 0.0291951035269 | 0.0605236618597527 |
| Ntn4 | ENSMUSG00000020019 | 57764 | 0.230510955203258 | 4.4049813788455 | 2.441244727887 | 0.0291981053793 | 0.0605236618597527 |
| Agtrap | ENSMUSG00000029007 | 11610 | 0.303621582714455 | 3.53414411491848 | 2.44118310262544 | 0.0292015179442 | 0.0605236618597527 |
| Acbd3 | ENSMUSG00000026499 | 170760 | 0.223712925227116 | 4.41538570339349 | 2.44085096808053 | 0.0292199168277 | 0.060544717486456 |
| Anapc1 | ENSMUSG00000014355 | 17222 | -0.155349401404517 | 5.62418763507702 | -2.44074037705975 | 0.0292260455770 | 0.060544717486456 |
| Tm7sf3 | ENSMUSG00000040234 | 67623 | 0.305288479680462 | 2.97577718771356 | 2.44069982310574 | 0.0292282933104 | 0.060544717486456 |
| Itch | ENSMUSG00000027598 | 16396 | 0.165421157294549 | 4.85622720709404 | 2.44023102167116 | 0.0292542890014 | 0.0605870847054045 |
| Camk2g | ENSMUSG00000021820 | 12325 | -0.250177605174684 | 5.96154306950373 | -2.43951955743683 | 0.0292937829782 | 0.060657386197256 |
| Nup85 | ENSMUSG00000020739 | 445007 | -0.223766819485248 | 3.14046236250022 | -2.43935917056817 | 0.0293026932365 | 0.0606643446531216 |
| Acad9 | ENSMUSG00000027710 | 229211 | -0.746292196248599 | 5.11233316165931 | -2.43915213560294 | 0.0293141988565 | 0.0606766725432649 |
| Olfr1221 | ENSMUSG00000075102 | 258904 | -1.46593503456841 | -0.0478203840399204 | -2.43845629022495 | 0.0293529009996 | 0.0607452784603381 |
| Inmt | ENSMUSG00000003477 | 21743 | -1.03485608312246 | 0.610091632295596 | -2.43808169939709 | 0.0293737555735 | 0.0607769301863212 |
| Atxn7l3 | ENSMUSG00000059995 | 217218 | 0.217876213309289 | 3.18700006715753 | 2.43783689021693 | 0.0293873924845 | 0.0607936387677601 |
| Golga3 | ENSMUSG00000029502 | 269682 | 0.158509442791001 | 6.84054904480721 | 2.43755349362041 | 0.0294031864476 | 0.0608148024709326 |
| Cbx1 | ENSMUSG00000018666 | 12412 | -0.214768859680255 | 4.36136349755542 | -2.43690607989268 | 0.0294392979238 | 0.060877973062 |
| Mafk | ENSMUSG00000018143 | 17135 | -0.237547595020853 | 4.25736482452354 | -2.4355514470972 | 0.0295149940960 | 0.061022962069908 |
| Arpp19 | ENSMUSG00000007656 | 59046 | -0.236374287181627 | 5.85462813756564 | -2.43485998203447 | 0.0295537045840 | 0.0610914419078239 |
| Dffa | ENSMUSG00000028974 | 13347 | -0.243308235165962 | 6.00805699490459 | -2.43297769763383 | 0.0296593273814 | 0.061298186350378 |
| Ctu2 | ENSMUSG00000049482 | 66965 | 0.348069285080082 | 2.10246125182205 | 2.43264720861496 | 0.0296779097367 | 0.0613249964712134 |
| Zfp799 | ENSMUSG00000095253 | 240064 | -0.418010225239902 | 2.66593743060343 | -2.43174941585731 | 0.0297284460490 | 0.0614178120887691 |
| Traf5 | ENSMUSG00000026637 | 22033 | -0.639186269108843 | 2.74575841967945 | -2.43138803579181 | 0.0297488111893 | 0.0614482719407474 |
| Srgap1 | ENSMUSG00000020121 | 117600 | 0.548133444635924 | 1.28826037550658 | 2.43096448577424 | 0.0297726968337 | 0.0614859907595112 |
| Cpa6 | ENSMUSG00000042501 | 329093 | -1.1769234288893 | 1.41642202701355 | -2.43026533821485 | 0.0298121646517 | 0.0615558692498372 |
| Il17rd | ENSMUSG00000040717 | 171463 | 0.343382044654359 | 3.22041807622978 | 2.42920436335754 | 0.0298721537398 | 0.0616680854637428 |
| Dhrs7 | ENSMUSG00000021094 | 66375 | 0.256549233695073 | 4.22601874289653 | 2.42842567879756 | 0.0299162551799 | 0.0617474670263808 |
| Aasdhppt | ENSMUSG00000025894 | 67618 | -0.22006926138647 | 3.38919359732561 | -2.42815905919725 | 0.0299313697090 | 0.061767000621106[illegible] |
| Epb41l4a | ENSMUSG00000024376 | 13824 | -0.324787468367695 | 1.73475552979601 | -2.4277585404591 | 0.0299540886416 | 0.0618022164858939 |
| Cct6a | ENSMUSG00000029447 | 12466 | 0.135766722973689 | 5.84384841522018 | 2.42738112493286 | 0.0299755121679 | 0.0618347468405549 |
| Itga6 | ENSMUSG00000027111 | 16403 | -0.320563550053735 | 5.03232127025895 | -2.4270175929067 | 0.0299961614716 | 0.061865668121926 |
| Pla2g6 | ENSMUSG00000042632 | 53357 | -0.245417243434949 | 3.16750028224733 | -2.42675268455887 | 0.0300112173338 | 0.0618850437042645 |
| Rfc2 | ENSMUSG00000023104 | 19718 | -0.289063567621464 | 2.43046817520004 | -2.42635427913154 | 0.0300338740103 | 0.0619200823854173 |
| Supt3 | ENSMUSG00000038954 | 109115 | -0.267009219150259 | 2.41793920910726 | -2.42550162221709 | 0.0300824182240 | 0.0620084694714724 |
| Cox20 | ENSMUSG00000026500 | 66359 | -0.232676942723725 | 2.90773429655945 | -2.42512450636522 | 0.0301039124420 | 0.062041075962303 |
| Lratd2 | ENSMUSG00000072568 | 399603 | -0.440891570026319 | 2.5002398785394 | -2.42376530607983 | 0.0301815037943 | 0.0621892588080204 |
| Pfkl | ENSMUSG00000020277 | 18641 | -0.394736050148964 | 2.59273665837488 | -2.4229472161035 | 0.0302282974622 | 0.062273938854057 |
| Phtf1 | ENSMUSG00000058388 | 18685 | 0.393953116814199 | 2.42562496007658 | 2.42276063822548 | 0.0302389791816 | 0.0622842060362363 |
| Bud23 | ENSMUSG00000005378 | 66138 | -0.259988279959273 | 2.70729739618546 | -2.42227165017596 | 0.0302669912311 | 0.0623301584678526 |
| Mpi | ENSMUSG00000032306 | 110119 | -0.211585835218659 | 4.41385166683447 | -2.42133787004489 | 0.0303205524740 | 0.0624286982660028 |
| Vmn1r224 | ENSMUSG00000091151 | 665525 | -1.01632930421066 | 0.723429637339594 | -2.4204989224195 | 0.0303687514371 | 0.0625161623275331 |
| Uhmk1 | ENSMUSG00000026667 | 16589 | -0.299101668392219 | 4.78031030740525 | -2.42018220713538 | 0.0303869663063 | 0.0625365682556284 |
| Casp2 | ENSMUSG00000029863 | 12366 | 0.426319114101387 | 1.56577990157628 | 2.42012763031786 | 0.0303901061727 | 0.0625365682556284 |
| Trim68 | ENSMUSG00000073968 | 101700 | -0.356407739229051 | 1.44202467400638 | -2.41996511538667 | 0.0303994576784 | 0.0625440376214237 |
| Notch4 | ENSMUSG00000015468 | 18132 | -0.487825360474643 | 3.47242602218583 | -2.41972270072694 | 0.0304134119217 | 0.0625609720956926 |
| Ctxn3 | ENSMUSG00000069372 | 629147 | -0.325937235804706 | 3.46614878842622 | -2.41932372993134 | 0.0304363914218 | 0.0625964618310291 |
| Col6a6 | ENSMUSG00000043719 | 245026 | -0.45485383606912 | 1.65984634034278 | -2.41855917008339 | 0.0304804741222 | 0.0626753315707739 |
| Pcdhgb7 | ENSMUSG00000104063 | 93704 | 0.482682718372371 | 1.47497517277348 | 2.41837920084833 | 0.0304908595901 | 0.0626784022550357 |
| Mtmr14 | ENSMUSG00000030269 | 97287 | -0.272461489074989 | 2.96191891168983 | -2.41828608507192 | 0.0304962343412 | 0.0626784022550357 |
| Vmn2r1 | ENSMUSG00000027824 | 56544 | -1.00079789311476 | 2.30676571129984 | -2.41823524137751 | 0.0304991694807 | 0.0626784022550357 |
| Rnf115 | ENSMUSG00000028098 | 67845 | 0.264583140109667 | 6.36688988953786 | 2.41801153939238 | 0.0305120867097 | 0.0626931616210321 |
| Ttc30b | ENSMUSG00000075273 | 72421 | -0.386814122221685 | 1.82959413245012 | -2.41724924325613 | 0.0305561432744 | 0.0627718853847976 |
| Nub1 | ENSMUSG00000028954 | 53312 | -0.168533248371904 | 4.7357618682948 | -2.41642876056875 | 0.0306036306416 | 0.0628576261825701 |
| Depdc5 | ENSMUSG00000037426 | 277854 | 0.258855562478635 | 3.69347398074632 | 2.41632806579944 | 0.0306094634454 | 0.0628577954015218 |
| Med4 | ENSMUSG00000022109 | 67381 | -0.24709258823765 | 3.21877981903372 | -2.41512634480341 | 0.0306791558915 | 0.0629890783340443 |
| Ubxn4 | ENSMUSG00000026353 | 67812 | 0.184836717253861 | 6.26340050090905 | 2.41425657400123 | 0.0307296918271 | 0.063080988049321 |
| Gm30948 | ENSMUSG00000103168 | 102633020 | -2.19564741445943 | 1.40462495316457 | -2.41396317070085 | 0.0307467572540 | 0.0631041688967692 |
| Smim8 | ENSMUSG00000028295 | 66291 | -0.334463067224064 | 2.65926700506779 | -2.41349426087101 | 0.0307740495957 | 0.0631472758842642 |
| Ankrd11 | ENSMUSG00000035569 | 77087 | -0.15366947192894 | 7.08422199739595 | -2.41340385164785 | 0.0307793144187 | 0.0631472758842642 |
| Chst11 | ENSMUSG00000034612 | 58250 | 0.377716723451866 | 1.64262467626446 | 2.41222191292145 | 0.0308482217686 | 0.0632767709051442 |
| Evi5 | ENSMUSG00000011831 | 14020 | 0.153501021434369 | 4.23084017533573 | 2.4114373405193 | 0.0308940439169 | 0.0633553748353125 |
| Utp4 | ENSMUSG00000041438 | 21771 | -0.215329399886262 | 3.07260881975436 | -2.41136736290143 | 0.0308981340451 | 0.0633553748353125 |
| Rnaset2a | ENSMUSG00000095687 | 100037283 | -0.238878737756101 | 3.96162356462209 | -2.41126164995845 | 0.0309043138516 | 0.0633561617990035 |
| Lpcat3 | ENSMUSG00000004270 | 14792 | -0.226841825279997 | 2.87519059273912 | -2.41069458616244 | 0.0309374836379 | 0.0634122695756113 |
| Fcho2 | ENSMUSG00000041685 | 218503 | -0.139694787528886 | 5.28197865939597 | -2.41015081504302 | 0.0309693228849 | 0.0634656296316573 |
| Ctdp1 | ENSMUSG00000033323 | 67655 | 0.225493636575492 | 3.8403729334341 | 2.40975564658538 | 0.0309924806865 | 0.0635011820776585 |

| | | | | | | | |
|---|---|---|---|---|---|---|---|
| Hspa12a | ENSMUSG00000025092 | 73442 | -0.357405771233571 | 2.83297010495894 | -2.40920335880231 | 0.0310248737653 | 0.063555640016736 |
| Ksr1 | ENSMUSG00000018334 | 16706 | -0.199386051020785 | 5.54141330958049 | -2.40728704995657 | 0.0311375210885 | 0.0637744506312039 |
| Lrrc8c | ENSMUSG00000054720 | 100604 | -0.403838753334232 | 4.13888265927872 | -2.40665023689546 | 0.0311750415792 | 0.0638393367370447 |
| 1700123O20Rik | ENSMUSG00000040822 | 58248 | 0.345333755840522 | 1.89390128015821 | 2.40642488382476 | 0.0311883295329 | 0.0638545851516672 |
| Hars2 | ENSMUSG00000019143 | 70791 | -0.206520927788481 | 3.05438252150726 | -2.40561238278768 | 0.0312362836112 | 0.0639407895420706 |
| Erf | ENSMUSG00000040857 | 13875 | 0.380136132006966 | 2.39935923125262 | 2.40426752195341 | 0.0313158124912 | 0.0640915832880255 |
| Spry1 | ENSMUSG00000037211 | 24063 | 0.232376213269596 | 4.39546687627693 | 2.40220652542197 | 0.0314380656066 | 0.0643297445051144 |
| Gramd4 | ENSMUSG00000035900 | 223752 | -0.294651509296347 | 3.17365243865654 | -2.40157391040427 | 0.0314756820333 | 0.0643946622321695 |
| Cyb5r1 | ENSMUSG00000026456 | 72017 | 0.170965434142399 | 5.04709244730621 | 2.40101402165669 | 0.0315090098618 | 0.0644507834581311 |
| Lurap1 | ENSMUSG00000028701 | 68075 | 0.324778481231674 | 2.57798884644708 | 2.39933341998414 | 0.0316092514506 | 0.0646365826770105 |
| Lrch4 | ENSMUSG00000093445 | 231798 | 0.304460892373516 | 3.5480188213117 | 2.39929292511583 | 0.0316116705691 | 0.0646365826770105 |
| Rarg | ENSMUSG00000001288 | 19411 | -0.408791328016222 | 3.33250892595492 | -2.3985512443929 | 0.0316560090195 | 0.0647151364490008 |
| Ube2j2 | ENSMUSG00000023286 | 140499 | -0.142752526519562 | 4.39938659568466 | -2.39780674873384 | 0.0317005754349 | 0.0647854707037603 |
| Nln | ENSMUSG00000021710 | 75805 | 0.209356188594045 | 3.71703334711755 | 2.39767990312787 | 0.0317081745345 | 0.0647854707037603 |
| Vps29 | ENSMUSG00000029462 | 56433 | 0.204049677872468 | 3.4664206160565 | 2.39767957877633 | 0.0317081939680 | 0.0647854707037603 |
| Klhl24 | ENSMUSG00000062901 | 75785 | -0.154272822766599 | 7.45093542802922 | -2.39643716822766 | 0.0317827166189 | 0.0649255980867214 |
| Exosc10 | ENSMUSG00000017264 | 50912 | -0.148831186566278 | 4.21407026860832 | -2.3963201985246 | 0.0317897413332 | 0.0649278143711233 |
| Aldh16a1 | ENSMUSG00000007833 | 69748 | 0.441788758963769 | 1.68453473728072 | 2.39601711230246 | 0.0318079503258 | 0.0649528684871083 |
| Rnf187 | ENSMUSG00000020496 | 108660 | 0.167747715837717 | 5.2875965021279 | 2.39586140474474 | 0.0318173088854 | 0.0649598437479105 |
| Slc30a1 | ENSMUSG00000037434 | 22782 | 0.314000233188508 | 3.67276409566706 | 2.39502705431882 | 0.0318675009875 | 0.065050168682529 |
| Gcc1 | ENSMUSG00000029708 | 74375 | 0.24760221820275 | 3.28992611843989 | 2.39486725835103 | 0.0318771224653 | 0.065057659758922 |
| Tvp23b | ENSMUSG00000014177 | 67510 | 0.495632338346943 | 1.35229478146942 | 2.39458590661329 | 0.0318940696759 | 0.0650800962530191 |
| Cntfr | ENSMUSG00000028444 | 12804 | -0.44777776081458 | 1.20858670054282 | -2.39428998690475 | 0.0319119036624 | 0.0651043335077587 |
| Gart | ENSMUSG00000022962 | 14450 | -0.192409008438941 | 4.61883680398165 | -2.39314283664303 | 0.0319811281309 | 0.0652333852583733 |
| Xpo5 | ENSMUSG00000067150 | 72322 | 0.191367909658134 | 3.88472348771364 | 2.3930163584648 | 0.0319887691837 | 0.0652367977514678 |
| Tab3 | ENSMUSG00000035476 | 66724 | -0.235800476877421 | 2.81052539398012 | -2.39281574704902 | 0.0320008924940 | 0.0652493482283591 |
| Il33 | ENSMUSG00000024810 | 77125 | 0.623703393915932 | 1.93700079037691 | 2.3924984879594 | 0.0320200739807 | 0.0652762828578609 |
| Zc3h7a | ENSMUSG00000037965 | 106205 | 0.214902095356128 | 3.90068044045724 | 2.39117134453278 | 0.0321004320404 | 0.0654270512977198 |
| Ucp3 | ENSMUSG00000032942 | 22229 | -0.814235199876309 | 4.58961973083935 | -2.39107947913327 | 0.0321060015699 | 0.0654270512977198 |
| Ubap2 | ENSMUSG00000028433 | 68926 | -0.178739220298898 | 6.31708728286051 | -2.39060835595248 | 0.0321345788719 | 0.0654730813884607 |
| Camsap1 | ENSMUSG00000026933 | 227634 | 0.153964917325766 | 5.57028218706204 | 2.39004612746134 | 0.0321687141737 | 0.0655304164429248 |
| Tigd2 | ENSMUSG00000049232 | 68140 | -0.197809173202704 | 3.70588542690128 | -2.38985285603179 | 0.0321804564958 | 0.0655421222203349 |
| Nsmce3 | ENSMUSG00000070520 | 66647 | 0.25210174229233 | 2.55510637707195 | 2.38973133916053 | 0.0321878414195 | 0.0655449505508331 |
| Simc1 | ENSMUSG00000043183 | 319719 | 0.380713275439069 | 1.68246478689228 | 2.38903599812989 | 0.0322301303526 | 0.0656188405448286 |
| Gtf3c2 | ENSMUSG00000106864 | 71752 | 0.192109544146179 | 4.15361110169228 | 2.38853126156957 | 0.0322608603733 | 0.0656582132859488 |
| Tmem161b | ENSMUSG00000035762 | 72745 | -0.300293531367985 | 2.07931081129774 | -2.3884753229944 | 0.0322642678169 | 0.0656582132859488 |
| Fam117b | ENSMUSG00000041040 | 72750 | 0.258862296463394 | 3.33758333861942 | 2.38842244809234 | 0.0322674889554 | 0.0656582132859488 |
| Pgp | ENSMUSG00000043445 | 67078 | -0.24106825977142 | 4.86615773244941 | -2.38771618634025 | 0.0323105438288 | 0.0657335854444698 |
| Sept4 | ENSMUSG00000020486 | 18952 | -0.378189469600625 | 2.78747562020136 | -2.38712394099149 | 0.0323466902978 | 0.0657948774926415 |
| Arfgap1 | ENSMUSG00000027575 | 228998 | 0.200980238189833 | 3.87767760907578 | 2.38686378407434 | 0.0323625806222 | 0.0658149523315527 |
| Foxk1 | ENSMUSG00000056493 | 17425 | 0.209992880391602 | 4.83049643431139 | 2.38647532672476 | 0.0323863213619 | 0.0658509819209459 |
| Numa1 | ENSMUSG00000066306 | 101706 | 0.148820879434528 | 6.6651430461165 | 2.38570060830693 | 0.0324337182031 | 0.0659350890233199 |
| Gm7072 | ENSMUSG00000117284 | 631624 | -0.256352072186806 | 3.36883665224453 | -2.38411383653314 | 0.0325310025270 | 0.0661205631505442 |
| Zc3h18 | ENSMUSG00000017478 | 76014 | -0.144193569706749 | 4.96285286763988 | -2.38385217097368 | 0.0325470718286 | 0.0661409262241136 |
| Ddx21 | ENSMUSG00000020075 | 56200 | 0.165536591893388 | 5.73696386564824 | 2.3830818596101 | 0.0325944218202 | 0.0662248373451725 |
| Slc8a3 | ENSMUSG00000079055 | 110893 | 0.275268831247216 | 4.24526228116045 | 2.38229215693147 | 0.0326430319249 | 0.0663112770808086 |
| Spata5 | ENSMUSG00000027722 | 57815 | -0.232773320265685 | 2.78728868690687 | -2.38218904661127 | 0.0326493839725 | 0.0663118573401093 |
| Ganab | ENSMUSG00000071650 | 14376 | 0.233579739113851 | 3.88948484972162 | 2.3817967810921 | 0.0326735600153 | 0.0663486317161282 |
| Vrk2 | ENSMUSG00000064090 | 69922 | -0.417763929949371 | 1.34331296352106 | -2.38164810287467 | 0.0326827277845 | 0.0663549215104736 |
| Tfcp2 | ENSMUSG00000009733 | 21422 | -0.25416698488688 | 3.13627303543793 | -2.38154501308457 | 0.0326890859275 | 0.0663555057147781 |
| Afdn | ENSMUSG00000068036 | 17356 | -0.24428923452647 | 6.03890194011623 | -2.38119618941978 | 0.0327106086426 | 0.0663868665191784 |
| Msra | ENSMUSG00000054733 | 110265 | -0.30166850104599 | 2.71467832591666 | -2.37845381706159 | 0.0328802863722 | 0.0667188435743643 |
| Echdc2 | ENSMUSG00000028601 | 52430 | -0.476594791636984 | 1.34558625963741 | -2.37787564282191 | 0.0329161664095 | 0.0667792529738695 |
| Isoc1 | ENSMUSG00000024601 | 66307 | -0.219450660423036 | 3.48407376298568 | -2.37704157198594 | 0.0329679924742 | 0.0668719847348826 |
| Smcr8 | ENSMUSG00000049323 | 237782 | -0.207265038303885 | 4.99165983835876 | -2.37660630462679 | 0.0329950692329 | 0.0669052946587063 |
| Suv39h1 | ENSMUSG00000039231 | 20937 | -0.410013612467776 | 1.45905428106223 | -2.37656684723388 | 0.0329975248134 | 0.0669052946587063 |
| Arhgap27 | ENSMUSG00000034255 | 544817 | -0.383730264828621 | 2.42668072577185 | -2.37648247176311 | 0.0330027763973 | 0.0669052946587063 |
| Pfdn2 | ENSMUSG00000006412 | 18637 | -0.21680956390617 | 3.70692150165549 | -2.37442817972953 | 0.0331308828087 | 0.0671525458896518 |
| Epb41l2 | ENSMUSG00000019978 | 13822 | 0.280225619401693 | 5.17206133873656 | 2.37346106878316 | 0.0331913560553 | 0.0672626460957896 |
| Rcsd1 | ENSMUSG00000040723 | 226594 | -0.339846250867287 | 5.43067819651839 | -2.3727238321335 | 0.0332375259107 | 0.0673437248805138 |
| Dennd3 | ENSMUSG00000036661 | 105841 | -0.56205323812179 | 2.17954542864544 | -2.37239997223438 | 0.0332578271472 | 0.0673675860236086 |
| Slc35c1 | ENSMUSG00000049922 | 228368 | 0.320708199652927 | 2.23001705513383 | 2.37233935032594 | 0.0332616285581 | 0.0673675860236086 |
| Txndc17 | ENSMUSG00000020803 | 52700 | 0.220844345666519 | 3.41182987056432 | 2.37177872167519 | 0.0332968034467 | 0.0674263353977071 |
| Ccm2l | ENSMUSG00000027474 | 228788 | -0.491794342217732 | 1.34936401191006 | -2.3702485823977 | 0.0333929876068 | 0.0676085844657775 |
| Ssrp1 | ENSMUSG00000027067 | 20833 | 0.193526871906235 | 4.58708442277252 | 2.36927899883974 | 0.0334540722493 | 0.0677197155107061 |
| Polr2k | ENSMUSG00000045996 | 17749 | -0.300129490590575 | 2.44151986410146 | -2.36834431790841 | 0.0335130586946 | 0.0678265588948854 |
| Plagl2 | ENSMUSG00000051413 | 54711 | 0.293435747040241 | 2.32735938578438 | 2.36799591533308 | 0.0335350712307 | 0.0678554494041277 |
| Maml1 | ENSMUSG00000050567 | 103806 | 0.242800344702884 | 3.45808872174561 | 2.36792191078233 | 0.0335397487083 | 0.0678554494041277 |
| Tecpr1 | ENSMUSG00000066621 | 70381 | -0.219449355024037 | 4.14953746319724 | -2.36729904865721 | 0.0335791414958 | 0.0679225750723228 |
| Tmub1 | ENSMUSG00000028958 | 64295 | -0.330573215820514 | 2.00952138162902 | -2.36719670953012 | 0.0335856181242 | 0.0679231067005448 |
| Pafah1b2 | ENSMUSG00000003131 | 18475 | 0.209356493373334 | 4.30381676817032 | 2.36642531573445 | 0.0336344747860 | 0.0680093310925196 |
| Slc10a3 | ENSMUSG00000032806 | 214601 | 0.546288346506502 | 1.4579760766698 | 2.3653793031056 | 0.0337008326576 | 0.068130904712462 |
| Enoph1 | ENSMUSG00000029326 | 67870 | -0.266765628563947 | 1.94326147974802 | -2.36490278561517 | 0.0337311036898 | 0.0681794923139244 |
| Higd1a | ENSMUSG00000038412 | 56295 | -0.237105462657406 | 4.78671255046381 | -2.36476317439567 | 0.0337399774688 | 0.0681848204310977 |
| Zbtb26 | ENSMUSG00000050714 | 320633 | -0.414468111172275 | 2.27191902958377 | -2.36382036255122 | 0.0337999613847 | 0.0682934155077896 |
| Dus1l | ENSMUSG00000025155 | 68730 | -0.260532469487299 | 4.00775782103209 | -2.36340243625673 | 0.0338265832943 | 0.068331102698072 |
| 1810037I17Rik | ENSMUSG00000054091 | 67704 | -0.332633462219548 | 1.97133071195744 | -2.36333131209325 | 0.0338311158907 | 0.068331102698072 |
| Olfr66 | ENSMUSG00000058200 | 18367 | -2.28013846869079 | -1.41972909566935 | -2.36320286091366 | 0.0338393032843 | 0.0683350127842033 |
| Tmem204 | ENSMUSG00000024168 | 407831 | -0.622479584383858 | 2.99047017711362 | -2.36309681420979 | 0.0338460640522 | 0.0683360410333863 |
| Slc25a28 | ENSMUSG00000040414 | 246696 | -0.27169837234739 | 3.38618887969994 | -2.36282691797855 | 0.0338632764753 | 0.0683581671564444 |
| Gosr1 | ENSMUSG00000010392 | 53334 | -0.189390906612668 | 3.79660979817622 | -2.36255146581375 | 0.0338808518121 | 0.0683810175699305 |
| Tmem205 | ENSMUSG00000040883 | 235043 | 0.432150340792759 | 1.74916194641246 | 2.36034178897366 | 0.0340221555567 | 0.0686535319163283 |
| Zfp324 | ENSMUSG00000004500 | 243834 | 0.4169339062826 | 1.64019923294945 | 2.35999271085109 | 0.0340445295195 | 0.0686860007712584 |
| Elmsan1 | ENSMUSG00000042507 | 238317 | 0.359883645116915 | 4.41120662507564 | 2.3593856717927 | 0.0340834706811 | 0.0687394250417234 |
| Ubox5 | ENSMUSG00000027300 | 140629 | -0.329556378590682 | 2.37286524446009 | -2.35938386775653 | 0.0340835864720 | 0.0687394250417234 |
| Trmt1l | ENSMUSG00000053286 | 98685 | -0.320362725237603 | 3.49458220628064 | -2.35856822680503 | 0.0341359761992 | 0.0688323844003444 |
| Specc1 | ENSMUSG00000042331 | 432572 | 0.292604373915223 | 2.40876303778117 | 2.3583397669207 | 0.0341506642157 | 0.0688493010959622 |
| Vmn1r201 | ENSMUSG00000094898 | 171255 | -1.20351132254041 | 0.545164175274886 | -2.35737062243599 | 0.0342130387837 | 0.0689623320938062 |
| Ppp6r3 | ENSMUSG00000024908 | 52036 | -0.158969107826463 | 5.88315385768728 | -2.3571281117285 | 0.0342286638218 | 0.0689811069758108 |
| Bcl9 | ENSMUSG00000038256 | 77578 | 0.199384131441716 | 2.92529783682165 | 2.35680214258608 | 0.0342496768051 | 0.069010731273211 |
| Fam111a | ENSMUSG00000024691 | 107373 | 0.553684078679431 | 2.02933366234034 | 2.35640125961483 | 0.0342755357892 | 0.069050107208318 |
| Enc1 | ENSMUSG00000041773 | 13803 | 0.633734327414077 | 2.20271026705471 | 2.35582096623652 | 0.0343130005494 | 0.0691128448509508 |
| Vmn1r207-ps | ENSMUSG00000084136 | 432735 | -1.01526886183494 | 0.688265505550527 | -2.35536910837167 | 0.0343422002533 | 0.0691589150643722 |
| Smg6 | ENSMUSG00000038290 | 103677 | 0.15569637570651 | 4.81507544042549 | 2.35379815642277 | 0.0344439014158 | 0.0693509460999587 |
| Ptpa | ENSMUSG00000039515 | 110854 | 0.160507995715399 | 4.75987443275415 | 2.35351336505034 | 0.0344623690367 | 0.0693753510019851 |
| Cyb5r4 | ENSMUSG00000032872 | 266690 | -0.272574415691631 | 5.08685359424418 | -2.35267044270641 | 0.0345170844832 | 0.069472068250724 |
| Phkg2 | ENSMUSG00000030815 | 68961 | 0.296131917881783 | 2.32018195976109 | 2.35257746954652 | 0.0345231245757 | 0.069472068250724 |
| Tma7 | ENSMUSG00000091537 | 66167 | -0.278158124766001 | 4.33960079018437 | -2.34921461046576 | 0.0347422727215 | 0.0699001993592367 |
| Olfr639 | ENSMUSG00000066263 | 259088 | -0.9799772267265 | 3.31569030679752 | -2.34724564059365 | 0.0348711980811 | 0.0701466813075923 |
| Psph | ENSMUSG00000029446 | 100678 | -0.36610981097779 | 1.96485946739074 | -2.34711977489945 | 0.0348794550317 | 0.0701503814079715 |
| Zfp143 | ENSMUSG00000061079 | 20841 | -0.331779722433217 | 2.91849170552599 | -2.34666987067528 | 0.0349089845174 | 0.070196856100045 |
| Ahctf1 | ENSMUSG00000026491 | 226747 | -0.161760968670894 | 4.83985285971228 | -2.34583783588098 | 0.0349636577664 | 0.070293864823386 |
| Rprd1a | ENSMUSG00000040446 | 225283 | -0.148021784642085 | 4.22055746338983 | -2.34393536641135 | 0.0350889754602 | 0.0705328412571191 |
| Edem3 | ENSMUSG00000043019 | 66967 | 0.134477093743995 | 5.1073898027053 | 2.34362434201587 | 0.0351095035126 | 0.0705611294166574 |
| A330094K24Rik | ENSMUSG00000097666 | 105246501 | -0.722049748975977 | 0.823788050627407 | -2.34226481570453 | 0.0351993681900 | 0.0707266038762619 |
| Ckap5 | ENSMUSG00000040549 | 75786 | -0.164498196276998 | 5.77445344723791 | -2.34218304583442 | 0.0352047801382 | 0.0707266038762619 |
| Slc37a4 | ENSMUSG00000032114 | 14385 | -0.418345002670357 | 2.52541016513162 | -2.33969939151197 | 0.0353695381781 | 0.0710445464581178 |
| Lyst | ENSMUSG00000019726 | 17101 | 0.219987905663533 | 3.80545440527953 | 2.33954994600532 | 0.0353794752594 | 0.0710493677350768 |
| 2310020H05Rik | ENSMUSG00000100410 | 69603 | -0.387809781318205 | 1.94963101517033 | -2.33946780694928 | 0.0353849380614 | 0.0710493677350768 |
| Ccdc6 | ENSMUSG00000048701 | 76551 | -0.134474861841243 | 5.51064261308293 | -2.3382266601574 | 0.0354675802688 | 0.0712022258804519 |
| Snx16 | ENSMUSG00000027534 | 74718 | -0.380223542142442 | 2.06441567308948 | -2.33752361036644 | 0.0355144743769 | 0.0712767521928308 |
| Gcm1 | ENSMUSG00000023333 | 14531 | -1.91809865225551 | 0.0899080483322893 | -2.33744363124955 | 0.0355198127904 | 0.0712767521928308 |
| Gmcl1 | ENSMUSG00000001157 | 23885 | 0.35927205893996 | 1.44462472664716 | 2.33737693063517 | 0.0355242654785 | 0.0712767521928308 |
| Zfp97 | ENSMUSG00000095990 | 22759 | -0.344308171334631 | 2.8351428970566 | -2.33652235683036 | 0.0355813606036 | 0.0713756162275813 |
| Nop58 | ENSMUSG00000026020 | 55989 | 0.189946143258531 | 3.58003839173018 | 2.3364440232764 | 0.0355865985216 | 0.0713756162275813 |
| Mboat7 | ENSMUSG00000035596 | 77582 | 0.287374545988486 | 2.33738537134614 | 2.33584519811533 | 0.0356266642705 | 0.0714428668394647 |

| | | | | | | | |
|---|---|---|---|---|---|---|---|
| Ddx60 | ENSMUSG00000037921 | 234311 | 0.470324163508756 | 1.49486813151144 | 2.33545940718671 | 0.0356524991628 | 0.0714815605921894 |
| Pitx2 | ENSMUSG00000028023 | 18741 | 0.2894426345155 | 3.73774351163067 | 2.33457105274262 | 0.0357120563930 | 0.0715878394337442 |
| Fam43a | ENSMUSG00000046546 | 224093 | -0.424000384915058 | 3.18845011893512 | -2.33323763062555 | 0.0358016292410 | 0.0717542371165615 |
| Scamp3 | ENSMUSG00000028049 | 24045 | -0.161194289637258 | 4.07269957610323 | -2.33312771971466 | 0.0358090220304 | 0.0717558973079468 |
| Stxbp3 | ENSMUSG00000027882 | 20912 | 0.265103514463545 | 3.59852847616776 | 2.33270272238716 | 0.0358376216888 | 0.0717941214848822 |
| Slc25a29 | ENSMUSG00000021265 | 214663 | -0.501774131626514 | 1.52460838526324 | -2.33260495651866 | 0.0358442037861 | 0.0717941214848822 |
| Rars | ENSMUSG00000018848 | 104458 | 0.165407589772244 | 4.55870644627998 | 2.33255153005202 | 0.0358478012134 | 0.0717941214848822 |
| Maml3 | ENSMUSG00000061143 | 433586 | 0.329344505766543 | 2.99552891555898 | 2.33226995064096 | 0.0358667667952 | 0.0718113716629003 |
| Snord15a | ENSMUSG00000065822 | 449630 | 0.443344588273217 | 4.39326477760678 | 2.3322285887621 | 0.0358695534973 | 0.0718113716629003 |
| Ifi35 | ENSMUSG00000010358 | 70110 | 0.254908434625708 | 2.86475289523926 | 2.33184801199648 | 0.0358952040018 | 0.0718495650878143 |
| Kctd2 | ENSMUSG00000016940 | 70382 | -0.253021872640666 | 3.18018241332147 | -2.3310821850002 | 0.0359468727697 | 0.0719398143987223 |
| Acsl5 | ENSMUSG00000024981 | 433256 | 0.293726312494465 | 2.78923099583192 | 2.33087261729927 | 0.0359610241659 | 0.0719549615884456 |
| Arrb1 | ENSMUSG00000018909 | 109689 | 0.217453082302352 | 3.62197047910521 | 2.33063263079169 | 0.0359772361347 | 0.0719645841586895 |
| Enox1 | ENSMUSG00000022012 | 239188 | 0.559918128011028 | 1.13122527242033 | 2.33060652176556 | 0.0359790003135 | 0.0719645841586895 |
| Plp1 | ENSMUSG00000031425 | 18823 | -0.597602022507138 | 1.90648959692883 | -2.32939351647458 | 0.0360610532673 | 0.0721155091959973 |
| Nol6 | ENSMUSG00000028430 | 230082 | 0.181942557112455 | 3.59998464106765 | 2.32907154070469 | 0.0360828629125 | 0.0721459254612959 |
| Srm | ENSMUSG00000006442 | 20810 | 0.316477386526112 | 2.03291090748466 | 2.32820612986196 | 0.0361415452188 | 0.0722270321945901 |
| Snw1 | ENSMUSG00000021039 | 66354 | -0.188187733768982 | 5.18743595659397 | -2.32809899318839 | 0.0361488163035 | 0.0722270321945901 |
| Tmem192 | ENSMUSG00000025521 | 73067 | -0.312425662836631 | 2.78469070767397 | -2.32805440645689 | 0.0361518426968 | 0.0722270321945901 |
| Tnfaip6 | ENSMUSG00000053475 | 21930 | 1.0818961162655 | 0.204325778688801 | 2.32801615422176 | 0.0361544391380 | 0.0722270321945901 |
| Lcor | ENSMUSG00000025019 | 212391 | 0.234907938576124 | 4.94046314351262 | 2.32798631292308 | 0.0361564651147 | 0.0722270321945901 |
| Dmd | ENSMUSG00000045103 | 13405 | 0.227717540647474 | 8.16878785932867 | 2.32784097721075 | 0.0361663328668 | 0.0722335436811917 |
| Olfr107 | ENSMUSG00000063188 | 258504 | -0.992552811750103 | 1.4656368011857 | -2.32762890967976 | 0.0361807360445 | 0.072244247177828 |
| Cplx2 | ENSMUSG00000025867 | 12890 | -0.386275344222579 | 1.73475854146539 | -2.32756746592384 | 0.0361849101910 | 0.072244247177828 |
| Etnk1 | ENSMUSG00000030275 | 75320 | -0.204797130578122 | 4.62149118281776 | -2.32672189359303 | 0.0362424000917 | 0.0723458136235181 |
| Cxxc5 | ENSMUSG00000046668 | 67393 | 0.463977098817209 | 3.18766213580337 | 2.32647281083537 | 0.0362593515781 | 0.0723535954582886 |
| Gm2260 | ENSMUSG00000098428 | 100039484 | 0.367684678501915 | 2.70173805090963 | 2.32647009095159 | 0.0362595367231 | 0.0723535954582886 |
| Neu3 | ENSMUSG00000035239 | 50877 | -0.37531904848485 | 1.48308629767338 | -2.32608773862014 | 0.0362855727171 | 0.0723923335228052 |
| Tmem14c | ENSMUSG00000021361 | 66154 | -0.236252336222315 | 3.98353206076748 | -2.32503244852349 | 0.0363575239516 | 0.072522644948126 |
| Kcng4 | ENSMUSG00000045246 | 66733 | -1.00138355395783 | 1.4730215739897 | -2.32356302505823 | 0.0364579368509 | 0.0727051718454738 |
| Tnpo3 | ENSMUSG00000012535 | 320938 | 0.122343432198599 | 5.10827094730692 | 2.32349879370377 | 0.0364623320882 | 0.0727051718454738 |
| Lmo4 | ENSMUSG00000028266 | 16911 | -0.204171788357452 | 3.28240965441529 | -2.32302942036631 | 0.0364944657280 | 0.0727559738961983 |
| Taf3 | ENSMUSG00000025782 | 209361 | -0.154892399039491 | 4.03096172987302 | -2.32157138093134 | 0.0365944555392 | 0.0729420119437493 |
| Znhit2 | ENSMUSG00000075227 | 29805 | 0.332751480803844 | 1.33174937367533 | 2.32083199675844 | 0.0366452604405 | 0.0730190981118038 |
| Rfxap | ENSMUSG00000036615 | 170767 | -0.261018013922835 | 2.99451304865801 | -2.32081412777014 | 0.0366464890898 | 0.0730190981118038 |
| Rpa2 | ENSMUSG00000028884 | 19891 | -0.362691501717295 | 2.60366693357781 | -2.32053715934796 | 0.0366655380885 | 0.073043739173726 |
| Fam13a | ENSMUSG00000037709 | 58909 | 0.206005552427467 | 5.00809640861185 | 2.31994969556767 | 0.0367059730408 | 0.0731109678041309 |
| Map3k2 | ENSMUSG00000024383 | 26405 | 0.223439357870332 | 4.16349644615677 | 2.31877574456427 | 0.0367869024209 | 0.0732588140579203 |
| Gm19410 | ENSMUSG00000109372 | 100502846 | -0.517658133499785 | 1.3010443923231 | -2.31864203516703 | 0.0367961307510 | 0.0732638443060195 |
| Olfr802 | ENSMUSG00000093866 | 258934 | -1.05119980948414 | 0.359546174821878 | -2.31803773239504 | 0.0368378657863 | 0.0733335842125403 |
| Olfr1307 | ENSMUSG00000094747 | 257956 | -0.888008032955732 | 3.59799701706847 | -2.31744611732778 | 0.0368787680365 | 0.0734016411885806 |
| Tbc1d5 | ENSMUSG00000023923 | 72238 | 0.207920051700861 | 3.8400161604389 | 2.31653001166208 | 0.0369421894137 | 0.0735144861607948 |
| Plxdc1 | ENSMUSG00000017417 | 72324 | -0.45108309289878 | 2.22089658677484 | -2.31612160042835 | 0.0369704967526 | 0.0735574262838324 |
| Tnfaip8 | ENSMUSG00000062210 | 106869 | 0.398211696612498 | 1.59536200636515 | 2.31595128125677 | 0.0369823077965 | 0.0735675353091317 |
| Tmem246 | ENSMUSG00000039611 | 67063 | -0.305417381458119 | 2.96446725390172 | -2.31534600175449 | 0.0370243107750 | 0.0736376894253363 |
| Atp6v1e1 | ENSMUSG00000019210 | 11973 | -0.185689490302638 | 5.34404194238651 | -2.31494952760664 | 0.0370518483441 | 0.0736790529834006 |
| Rngtt | ENSMUSG00000028274 | 24018 | -0.282670401713876 | 3.25562696808817 | -2.31454606023248 | 0.0370798915623 | 0.0737214068147682 |
| Wac | ENSMUSG00000024283 | 225131 | -0.134254113839228 | 4.95649335766513 | -2.31400985551153 | 0.0371171918915 | 0.0737821466750885 |
| Mrtfb | ENSMUSG00000009569 | 239719 | 0.196933320404311 | 5.09826667177401 | 2.3138727445427 | 0.0371267355307 | 0.0737876992885124 |
| Nedd4l | ENSMUSG00000024589 | 83814 | 0.243275655470728 | 3.83209800521179 | 2.31304446350047 | 0.0371844376859 | 0.0738820290464973 |
| Ascc2 | ENSMUSG00000020412 | 75452 | 0.156627944485552 | 4.31794977878095 | 2.31299744148591 | 0.0371877160199 | 0.0738820290464973 |
| Il1r1 | ENSMUSG00000026072 | 16177 | 0.275716414842679 | 2.71337422822996 | 2.31285323589984 | 0.0371977716157 | 0.0738885774181675 |
| Zfp931 | ENSMUSG00000078861 | 353208 | -0.350048442558041 | 1.65829576590936 | -2.31236487931208 | 0.0372318443354 | 0.0739428216624794 |
| Vmn2r7 | ENSMUSG00000062200 | 319217 | -1.62034980889032 | -0.77032861483699 | -2.31169139954757 | 0.0372788816427 | 0.0740227893255527 |
| Zfp874b | ENSMUSG00000059839 | 408067 | -0.297137314515243 | 2.30116982477782 | -2.31127627312713 | 0.0373079030263 | 0.074056826765602 |
| Xpr1 | ENSMUSG00000026469 | 19775 | 0.179675551190744 | 5.45484125947998 | 2.31125239155578 | 0.0373095732319 | 0.074056826765602 |
| Arhgap12 | ENSMUSG00000041225 | 75415 | -0.151453318480249 | 4.46902827466503 | -2.31106908599406 | 0.0373223954328 | 0.0740688279731391 |
| Manbal | ENSMUSG00000063019 | 69161 | 0.369403979320225 | 2.00946794313618 | 2.31079306268712 | 0.0373417111047 | 0.0740937092187678 |
| Kbtbd2 | ENSMUSG00000059486 | 210973 | -0.179987776724932 | 4.2583458917217 | -2.31035123671762 | 0.0373726490964 | 0.0741416383436112 |
| Dtymk | ENSMUSG00000026281 | 21915 | 0.309675673583764 | 1.99539620231163 | 2.31002931482205 | 0.0373952063320 | 0.0741729269490188 |
| Parp4 | ENSMUSG00000054509 | 328417 | 0.239401926746996 | 5.10830456988423 | 2.30958760880549 | 0.0374261778932 | 0.0742208908838167 |
| Myoz2 | ENSMUSG00000028116 | 59006 | -0.20840831898923 | 8.98563590149359 | -2.30850383905947 | 0.0375022726868 | 0.0743583063196739 |
| Fat4 | ENSMUSG00000046743 | 329628 | 0.281518219227743 | 4.53845038069552 | 2.30776758497888 | 0.0375540509087 | 0.0744474665366654 |
| Metap2 | ENSMUSG00000036112 | 56307 | -0.199950167870546 | 6.36285667032583 | -2.30641095492693 | 0.0376496354086 | 0.0746234206077694 |
| Rpl27a | ENSMUSG00000046364 | 26451 | 0.133696335382743 | 5.78739863684291 | 2.30617818463937 | 0.0376660589159 | 0.074642438363053 |
| Tlnrd1 | ENSMUSG00000070462 | 80889 | 0.308767476740047 | 2.71880165751438 | 2.30562825990659 | 0.0377048867514 | 0.0747058396012491 |
| Smu1 | ENSMUSG00000028409 | 74255 | -0.151444110879474 | 4.61658831555619 | -2.3052842806076 | 0.0377291929249 | 0.0747342221786213 |
| D830005E20Rik | ENSMUSG00000112707 | 442820 | -0.947334458080073 | 0.169532197984378 | -2.30523204086131 | 0.0377328855734 | 0.0747342221786213 |
| Nrarp | ENSMUSG00000078202 | 67122 | -0.440713105969393 | 1.78361375624034 | -2.3043541865293 | 0.0377949893018 | 0.0748436645033567 |
| Erap1 | ENSMUSG00000021583 | 80898 | -0.252636069558422 | 3.75007938952381 | -2.30249064976088 | 0.0379271459771 | 0.0750917646579633 |
| Diablo | ENSMUSG00000029433 | 66593 | -0.167064622458309 | 4.27372945084043 | -2.3022138512066 | 0.0379468130124 | 0.0751170976165512 |
| Slc25a5 | ENSMUSG00000016319 | 11740 | 0.318912480418525 | 5.05759271962191 | 2.30209332767941 | 0.0379553794424 | 0.0751204513282135 |
| Aurkaip1 | ENSMUSG00000065990 | 66077 | -0.220250523348947 | 5.11712224365121 | -2.30163155568762 | 0.0379882176776 | 0.075171833351554 |
| Atn1 | ENSMUSG00000004263 | 13498 | 0.145147012002908 | 5.31236438555321 | 2.30133259950677 | 0.0380094918460 | 0.0752003177139244 |
| Aacs | ENSMUSG00000029482 | 78894 | 0.431344662438303 | 1.62100492029995 | 2.30115507823065 | 0.0380221298613 | 0.0752003598861663 |
| Racgap1 | ENSMUSG00000023015 | 26934 | 0.555972443349337 | 1.1145312571488 | 2.30113903413264 | 0.0380232722615 | 0.0752003598861663 |
| Evi5l | ENSMUSG00000011832 | 213027 | -0.204640832821382 | 3.92263107738883 | -2.30077756697946 | 0.0380490186975 | 0.075237667037509 |
| Acp2 | ENSMUSG00000002103 | 11432 | 0.222688496549527 | 3.51912043985675 | 2.30026526540981 | 0.0380855369741 | 0.0752962569478894 |
| Psmb10 | ENSMUSG00000031897 | 19171 | 0.493583415209979 | 2.52951496913481 | 2.29983008990241 | 0.0381165835469 | 0.0753376075655213 |
| Camk2a | ENSMUSG00000024617 | 12322 | -0.159967315829178 | 5.50093574435886 | -2.29971925964656 | 0.0381244942969 | 0.0753376075655213 |
| Fbh1 | ENSMUSG00000058594 | 50755 | 0.204962110008529 | 4.11131691608337 | 2.29968235469538 | 0.0381271288127 | 0.0753376075655213 |
| Dhx40 | ENSMUSG00000018425 | 67487 | 0.178807342758142 | 3.53631741146791 | 2.29948420718132 | 0.0381412768194 | 0.0753466860591716 |
| Fam8a1 | ENSMUSG00000069237 | 97863 | 0.198639243125388 | 3.72237825555291 | 2.29942494533841 | 0.0381455091621 | 0.0753466860591716 |
| Vmn2r31 | ENSMUSG00000096373 | 100042591 | -1.84602285071762 | -1.3893917698718 | -2.29865316662423 | 0.0382006684010 | 0.0754420065568115 |
| Mterf3 | ENSMUSG00000021519 | 66410 | -0.252987052253677 | 3.43726716583395 | -2.29696285221043 | 0.0383217394435 | 0.0756674374742904 |
| Ddx28 | ENSMUSG00000045538 | 71986 | -0.242056662585016 | 2.47479136845962 | -2.29652150658747 | 0.0383534111438 | 0.0757162971308717 |
| Wdr81 | ENSMUSG00000045374 | 192652 | 0.293451241771357 | 3.51606436462444 | 2.29523747034856 | 0.0384456966342 | 0.075884779687488 |
| Osgin1 | ENSMUSG00000074063 | 71839 | -0.403854196852873 | 2.4710384605844 | -2.29501881554188 | 0.0384614325811 | 0.0759021338768782 |
| Olfr914 | ENSMUSG00000047050 | 258782 | -0.983232256785063 | 0.91477328609193 | -2.29489404155389 | 0.0384704149304 | 0.0759061562463649 |
| Ogfrl1 | ENSMUSG00000026158 | 70155 | 0.22416157370792 | 2.96582295828664 | 2.29476159329795 | 0.0384799519137 | 0.075911271317295 |
| Ebf2 | ENSMUSG00000022053 | 13592 | 0.361036298306182 | 2.76116622348811 | 2.29406556838208 | 0.0385301062034 | 0.0759964978183809 |
| B4galt4 | ENSMUSG00000022793 | 56375 | -0.37740197380743 | 2.23630042577467 | -2.29376114940327 | 0.0385520615135 | 0.0760260841430832 |
| Entpd6 | ENSMUSG00000033068 | 12497 | -0.445483146688506 | 1.36953745312779 | -2.29352906835723 | 0.0385688076159 | 0.0760357146702983 |
| Hspbp1 | ENSMUSG00000063802 | 66245 | -0.264324969152981 | 3.08937730223892 | -2.29350067290593 | 0.0385708569981 | 0.0760357146702983 |
| Ahsa1 | ENSMUSG00000021037 | 217737 | -0.164897794406043 | 5.04135585745876 | -2.29310721435457 | 0.0385992646488 | 0.0760779952895883 |
| Olfr432 | ENSMUSG00000047048 | 258711 | -1.84840502893699 | 1.51988784819168 | -2.29170185021618 | 0.0387008934006 | 0.0762645512462379 |
| Mfap3 | ENSMUSG00000020522 | 216760 | 0.209109592011893 | 3.07638536512063 | 2.29093251679309 | 0.0387566347980 | 0.0763606299527415 |
| Sfrp5 | ENSMUSG00000018822 | 54612 | -0.559883768750109 | 1.91277300708486 | -2.29073128257787 | 0.0387712275741 | 0.0763756151763058 |
| Nhp2 | ENSMUSG00000001056 | 52530 | 0.282133879685081 | 2.98098427825294 | 2.29031166261701 | 0.0388016736090 | 0.076421818778524 |
| Lmo2 | ENSMUSG00000032698 | 16909 | 0.2451414592372 | 3.13007637815601 | 2.28993973896456 | 0.0388286778794 | 0.0764532467688561 |
| Tial1 | ENSMUSG00000030846 | 21843 | -0.177317099896329 | 4.34399001953052 | -2.28989924790202 | 0.0388316188876 | 0.0764532467688561 |
| Sppl3 | ENSMUSG00000029550 | 74585 | 0.241363507861955 | 3.32366700884454 | 2.28974580931683 | 0.0388427655828 | 0.0764614209590737 |
| Gm4737 | ENSMUSG00000048087 | 11615 | 0.649608424653084 | 1.25585297827595 | 2.28926053234003 | 0.0388780389201 | 0.0765170766000376 |
| Actr8 | ENSMUSG00000015971 | 56249 | -0.211198789036413 | 3.5870841927436 | -2.28766333308742 | 0.0389943485844 | 0.0767321736052109 |
| Hmg20a | ENSMUSG00000032329 | 66867 | -0.226068021317146 | 3.32479489145112 | -2.28693949986737 | 0.0390471671081 | 0.0768222792763887 |
| Tmed7 | ENSMUSG00000033184 | 66676 | 0.184831244261189 | 4.8010745157139 | 2.28653492383013 | 0.0390767187122 | 0.0768665848917946 |
| Ctla2a | ENSMUSG00000044258 | 13024 | -0.469004276496982 | 2.80218132460739 | -2.28573986503833 | 0.0391348541083 | 0.0769670907266294 |
| Thbs1 | ENSMUSG00000040152 | 21825 | 0.668045063874691 | 6.1960877850059 | 2.28514240577676 | 0.0391785946452 | 0.0770392549140689 |
| Dnajc17 | ENSMUSG00000034278 | 69408 | -0.300093205328396 | 1.9133599918644 | -2.28421010988263 | 0.0392469412005 | 0.0771485240116269 |
| Mtmr4 | ENSMUSG00000018401 | 170749 | -0.191439569621911 | 3.66213459095275 | -2.28419187132165 | 0.0392482793915 | 0.0771485240116269 |
| Agpat3 | ENSMUSG00000001211 | 28169 | -0.161784293163088 | 6.65440670699575 | -2.28401663331279 | 0.0392611390687 | 0.0771599264941535 |
| Flii | ENSMUSG00000002812 | 14248 | 0.134472007406517 | 6.69286055336417 | 2.28279370170445 | 0.0393509936853 | 0.0773226158500876 |
| Otud3 | ENSMUSG00000041161 | 73162 | 0.231092445787956 | 2.95121901246338 | 2.28254862861357 | 0.0393690237335 | 0.0773441406238277 |
| Tril | ENSMUSG00000043496 | 66873 | 0.529250112754324 | 2.66796590893463 | 2.28223397880195 | 0.0393921839928 | 0.0773757346318526 |
| 2700097O09Rik | ENSMUSG00000062198 | 72658 | -0.415930068499357 | 1.60926781587649 | -2.28149373470354 | 0.0394467215261 | 0.0774689387356527 |
| Gm10778 | ENSMUSG00000096856 | 100233208 | -0.317124345758818 | 1.54818121062688 | -2.28058710507193 | 0.0395136147282 | 0.0775863701445399 |

| | | | | | | | |
|---|---|---|---|---|---|---|---|
| Fam98b | ENSMUSG00000027349 | 68215 | -0.223253550609599 | 3.18883241534013 | -2.28025942655456 | 0.0395378179546 | 0.0776122290428895 |
| Ube2v1 | ENSMUSG00000078923 | 66589 | -0.175117763761478 | 6.07653087152348 | -2.2802165674512 | 0.0395409846802 | 0.0776122290428895 |
| Gab1 | ENSMUSG00000031714 | 14388 | -0.162957853290835 | 4.08192257822039 | -2.27792761870506 | 0.0397104565200 | 0.0779308797522804 |
| Atp6ap2 | ENSMUSG00000031007 | 70495 | 0.349897813275429 | 3.36153443948933 | 2.27775448482655 | 0.0397233030802 | 0.0779420976432162 |
| Gbp10 | ENSMUSG00000105096 | 626578 | -0.769914516065247 | 4.93710022972033 | -2.27756274692074 | 0.0397375346431 | 0.0779490820222375 |
| Meaf6 | ENSMUSG00000028863 | 70088 | 0.274316464059984 | 2.20286306876201 | 2.277502809866 | 0.0397419844017 | 0.0779490820222375 |
| Olfr90 | ENSMUSG00000056600 | 258469 | -0.857756355831213 | 2.52382611781115 | -2.27741834775599 | 0.0397482557123 | 0.0779490820222375 |
| Nup35 | ENSMUSG00000026999 | 69482 | -0.323054774349441 | 2.09777060961517 | -2.2768451001798 | 0.0397908440448 | 0.07801860381233 |
| Ptma | ENSMUSG00000026238 | 19231 | 0.209773228640388 | 6.02065799936434 | 2.27504846176654 | 0.0399246012383 | 0.0782668249886759 |
| Tfcp2l1 | ENSMUSG00000026380 | 81879 | -0.73643395768115 | 1.06804830286063 | -2.27475610089793 | 0.0399464072221 | 0.0782955311718563 |
| Sos1 | ENSMUSG00000024241 | 20662 | -0.1840346430436 | 5.1088011419532 | -2.27465415151198 | 0.0399540138481 | 0.0782964011068584 |
| Slc35b2 | ENSMUSG00000037089 | 73836 | 0.313036607481965 | 2.95221711526336 | 2.27398929367798 | 0.0400036536354 | 0.0783684104809219 |
| Stau1 | ENSMUSG00000039536 | 20853 | 0.139497459094602 | 5.13705035982677 | 2.27396995483427 | 0.0400050983883 | 0.0783684104809219 |
| Dnajc1 | ENSMUSG00000026740 | 13418 | 0.148335195310501 | 4.13880474927246 | 2.27314863541528 | 0.0400665024512 | 0.0784746350643689 |
| Dcaf7 | ENSMUSG00000049354 | 71833 | 0.149322912384113 | 5.19586664058419 | 2.27265929807189 | 0.0401031289283 | 0.0785323006657662 |
| Cpne5 | ENSMUSG00000024008 | 240058 | -0.637401547359295 | 1.11375800082336 | -2.272414977767 | 0.0401214279217 | 0.0785540620836777 |
| Calm3 | ENSMUSG00000019370 | 12315 | -0.121045116432735 | 6.24929444866608 | -2.27221651286651 | 0.0401362982599 | 0.0785691039183276 |
| Osgin2 | ENSMUSG00000041153 | 209212 | 0.251894746007121 | 2.76363024708546 | 2.27194713330128 | 0.0401564903310 | 0.0785945560982486 |
| Snhg4 | ENSMUSG00000117869 | 100503380 | 0.606762252366832 | 4.29886026858447 | 2.27158866984343 | 0.0401833747976 | 0.0786123824666644 |
| Depp1 | ENSMUSG00000048489 | 213393 | -1.01760504541597 | 2.17579751079862 | -2.27155264232817 | 0.0401860777713 | 0.0786123824666644 |
| Tspan12 | ENSMUSG00000029669 | 269831 | -0.245226274830343 | 2.78564825951918 | -2.27153803855568 | 0.0401871734721 | 0.0786123824666644 |
| Evl | ENSMUSG00000021262 | 14026 | 0.249642671604167 | 2.79634643260951 | 2.27133877472108 | 0.0402021267785 | 0.0786275626795654 |
| 2510009E07Rik | ENSMUSG00000043391 | 72190 | 0.14129976930815 | 4.76290204668007 | 2.26901585578406 | 0.0403768329517 | 0.0789551271905616 |
| Sgcb | ENSMUSG00000029156 | 24051 | 0.14964306878819 | 5.74393660772335 | 2.26835669227386 | 0.0404265388551 | 0.0790381856958453 |
| Ttc19 | ENSMUSG00000042298 | 72795 | -0.218568314354305 | 4.31235837299696 | -2.2666161977983 | 0.0405580629722 | 0.0792811492041347 |
| Nkd2 | ENSMUSG00000021567 | 72293 | 0.702947103216525 | 1.75970030711704 | 2.26619398064537 | 0.0405900295002 | 0.0793294497527749 |
| Pdik1l | ENSMUSG00000050890 | 230809 | -0.359756504393221 | 1.76906901764956 | -2.26508385372441 | 0.0406741919027 | 0.0794797267946001 |
| BC004004 | ENSMUSG00000052712 | 80748 | -0.169698819481188 | 4.8108219665539 | -2.26481327978112 | 0.0406947299520 | 0.0795056466678088 |
| Rhox8 | ENSMUSG00000064137 | 434768 | -1.00248995063092 | 0.960052141970978 | -2.26438513983515 | 0.0407272481179 | 0.0795549587522362 |
| Hist4h4 | ENSMUSG00000096010 | 320332 | 0.41927465920466 | 1.50349450380068 | 2.26422467688906 | 0.0407394419444 | 0.0795645595666094 |
| Gnpnat1 | ENSMUSG00000037722 | 54342 | 0.325380294270883 | 2.44483496893677 | 2.26278133246956 | 0.0408492788361 | 0.0797648208214277 |
| Rtf2 | ENSMUSG00000027502 | 66404 | 0.168490058534246 | 4.26948102858524 | 2.26267642885221 | 0.0408572727636 | 0.0797661812071728 |
| Sipa1l2 | ENSMUSG00000001995 | 244668 | -0.212263345827886 | 4.15924029066465 | -2.26199236839397 | 0.0409094361374 | 0.0798537582890405 |
| Olfr1459 | ENSMUSG00000057503 | 258684 | -1.42261867002972 | 0.395206959868455 | -2.26168109977026 | 0.0409331928722 | 0.0798858652537855 |
| Cryzl1 | ENSMUSG00000058240 | 66609 | -0.312179369161178 | 2.7373238783702 | -2.26115222851887 | 0.0409735873775 | 0.0799504254952373 |
| Dlg3 | ENSMUSG00000000881 | 53310 | 0.343257442691185 | 1.31336141757807 | 2.26086082483249 | 0.0409958604779 | 0.0799796092957437 |
| Kmt2a | ENSMUSG00000002028 | 214162 | 0.179811615463329 | 5.95711611162719 | 2.26000975242502 | 0.0410609765964 | 0.0800923510307403 |
| Erp29 | ENSMUSG00000029616 | 67397 | 0.204031696920729 | 3.8501250851097 | 2.25970424702571 | 0.0410843748003 | 0.0801185103724023 |
| Cd55 | ENSMUSG00000026399 | 13136 | 0.297825297986104 | 4.24535066077737 | 2.25964326706577 | 0.0410890466698 | 0.0801185103724023 |
| Cgnl1 | ENSMUSG00000032232 | 68178 | 0.270421841993826 | 3.66738944732128 | 2.25934372828209 | 0.0411120025679 | 0.0801489745087485 |
| Uba7 | ENSMUSG00000032596 | 74153 | -0.397660393572687 | 2.2163527518756 | -2.25921253505291 | 0.0411220606947 | 0.0801542877058814 |
| Emc6 | ENSMUSG00000047260 | 66048 | -0.211319806771847 | 4.95668731703895 | -2.25890394959388 | 0.0411457280131 | 0.0801861210397329 |
| Rny1 | ENSMUSG00000065701 | 19872 | 0.815875132360971 | 2.82000493723154 | 2.25869568358392 | 0.0411617084745 | 0.0802029653003968 |
| Wdr33 | ENSMUSG00000024400 | 74320 | -0.1415353017032 | 4.91177007037471 | -2.25854766705418 | 0.0411730694899 | 0.0802108042407856 |
| Lrrc8b | ENSMUSG00000070639 | 433926 | 0.344757634245254 | 1.76836519358994 | 2.25804980262321 | 0.0412113047991 | 0.0802709858801192 |
| Snx30 | ENSMUSG00000028385 | 209131 | 0.257491077174588 | 2.21772223143975 | 2.25682240117367 | 0.0413057106118 | 0.0804405349543145 |
| 9130023H24Rik | ENSMUSG00000062944 | 100043133 | 0.373594433869199 | 1.25273710557169 | 2.25659812043982 | 0.0413229832240 | 0.0804501842227457 |
| Zbed6 | ENSMUSG00000102049 | 667118 | 0.200052669774739 | 4.79134477555508 | 2.25656693979436 | 0.0413253850892 | 0.0804501842227457 |
| Grpel2 | ENSMUSG00000024580 | 17714 | -0.185871751118561 | 3.46648524099619 | -2.25512531584843 | 0.0414365781313 | 0.080652285533127 |
| Rftn2 | ENSMUSG00000025978 | 74013 | 0.418676621840099 | 2.25476856786657 | 2.25438612292365 | 0.0414937017017 | 0.0807490926299094 |
| Sfi1 | ENSMUSG00000023764 | 78887 | -0.390147143845704 | 2.54598046972191 | -2.25407851116399 | 0.0415174952796 | 0.08076899095775 |
| Polr2j | ENSMUSG00000039771 | 20022 | -0.194882235104151 | 3.48954447990365 | -2.25406288083503 | 0.0415187046191 | 0.08076899095775 |
| Slc10a2 | ENSMUSG00000023073 | 20494 | -1.30022849331273 | 1.72156359475224 | -2.25396605822186 | 0.0415261966540 | 0.0807691913924767 |
| H2afj | ENSMUSG00000060032 | 232440 | 0.188827571519611 | 3.35941061275794 | 2.25356507272522 | 0.0415572380705 | 0.0808151875732697 |
| Cast | ENSMUSG00000021585 | 12380 | 0.130311752443395 | 6.67775559533771 | 2.2527470782517 | 0.0416206291486 | 0.0809240656748213 |
| Gpx4 | ENSMUSG00000075706 | 625249 | -0.191174045247364 | 7.19212949567545 | -2.25115237513856 | 0.0417444739143 | 0.0811504258149477 |
| Orc3 | ENSMUSG00000040044 | 50793 | -0.170985170157461 | 3.78445558954219 | -2.25070367225914 | 0.0417793827628 | 0.081203846547098 |
| Zfp2 | ENSMUSG00000049321 | 22678 | -0.52134038727683 | 1.0525546773867 | -2.25046277554717 | 0.0417981357467 | 0.0812258527728826 |
| Tjp1 | ENSMUSG00000030516 | 21872 | -0.225297972600107 | 5.57654429873741 | -2.25027436956967 | 0.0418128080335 | 0.0812269898737226 |
| Olfr539 | ENSMUSG00000059136 | 258963 | -1.01619716793629 | 3.22370410906477 | -2.25026442441099 | 0.0418135826566 | 0.0812269898737226 |
| Stambp | ENSMUSG00000006906 | 70527 | 0.313182376393787 | 2.26290412178732 | 2.24998018142664 | 0.0418357279074 | 0.0812555688976347 |
| Psmb5 | ENSMUSG00000022193 | 19173 | 0.194066672976753 | 4.71961781748384 | 2.24954154303078 | 0.0418699237246 | 0.0813075388583414 |
| Ppp6r2 | ENSMUSG00000036561 | 71474 | -0.211669145613916 | 5.68547159776407 | -2.24942081583864 | 0.0418793401179 | 0.0813113795433965 |
| Vrk3 | ENSMUSG00000002205 | 101568 | -0.274832191170755 | 2.53078016852817 | -2.24874462422789 | 0.0419321181211 | 0.0813993932130043 |
| Bcl2l11 | ENSMUSG00000027381 | 12125 | 0.327613082790787 | 2.70563908367831 | 2.24825735986324 | 0.0419701888423 | 0.0814588306526821 |
| Klhl9 | ENSMUSG00000070923 | 242521 | -0.170441213302641 | 5.59595021665449 | -2.24729587533369 | 0.0420454066552 | 0.0815879496824472 |
| Tab1 | ENSMUSG00000022414 | 66513 | -0.209268807978412 | 3.48898989384059 | -2.24721623225594 | 0.0420516428973 | 0.0815879496824472 |
| Adamtsl5 | ENSMUSG00000043822 | 66548 | 0.15426369662529 | 5.34555023721711 | 2.2469148657446 | 0.0420752484975 | 0.0816192619923209 |
| Mtif3 | ENSMUSG00000016510 | 76366 | -0.211780482424219 | 3.72719368507462 | -2.24463616227773 | 0.0422541403993 | 0.0819517403664812 |
| Foxo3 | ENSMUSG00000048756 | 56484 | 0.17561517715765 | 5.25761493452057 | 2.24431002074993 | 0.0422798029875 | 0.0819862790239367 |
| Sema6b | ENSMUSG00000001227 | 20359 | 0.33781567425092 | 1.84289669307107 | 2.24421923457789 | 0.0422869491480 | 0.0819862790239367 |
| Tmem44 | ENSMUSG00000022537 | 224090 | -0.448282091741574 | 1.8412628588852 | -2.24356241054176 | 0.0423386844148 | 0.0820720268378627 |
| Baz2b | ENSMUSG00000026987 | 407823 | -0.171089632027947 | 5.33485298707006 | -2.24318778182651 | 0.0423682190281 | 0.0821147167015034 |
| Cers5 | ENSMUSG00000023021 | 71949 | 0.356549760173212 | 1.58853694811427 | 2.24296262258507 | 0.0423859792450 | 0.0821345752752642 |
| Msl1 | ENSMUSG00000052915 | 74026 | -0.148230654324757 | 5.9000987999795 | -2.24233238483176 | 0.0424357287135 | 0.0822164038582609 |
| Actr10 | ENSMUSG00000021076 | 56444 | -0.182875255744127 | 4.15868143126395 | -2.24170925664977 | 0.0424849710152 | 0.0822972210115394 |
| Qars | ENSMUSG00000032604 | 97541 | -0.180869251681453 | 4.94202185988769 | -2.24141313372248 | 0.0425083907902 | 0.0823279978257659 |
| Smarce1 | ENSMUSG00000037935 | 57376 | 0.152742924588448 | 5.24552958639198 | 2.24075902235219 | 0.0425601662425 | 0.0824136717799344 |
| Snrpb | ENSMUSG00000027404 | 20638 | -0.163529045861143 | 5.24772351313996 | -2.2402607831605 | 0.0425996436321 | 0.082475505586863 |
| Nr2c1 | ENSMUSG00000005897 | 22025 | -0.231382649610576 | 2.4387138158972 | -2.23869305102772 | 0.0427240859994 | 0.0827017857375426 |
| Sept10 | ENSMUSG00000019917 | 103080 | 0.324700615608077 | 2.76884209330543 | 2.23807714373454 | 0.0427730686155 | 0.0827724958211557 |
| Ifi44 | ENSMUSG00000028037 | 99899 | -1.13286109055951 | 1.03295902177913 | -2.2380433257303 | 0.0427757596645 | 0.0827724958211557 |
| Eda | ENSMUSG00000059327 | 13607 | 0.375103213253699 | 1.2269762463423 | 2.23671865002796 | 0.0428812954458 | 0.0829620248705007 |
| Trub2 | ENSMUSG00000039826 | 227682 | -0.228468627334876 | 4.2196841990085 | -2.23620724564298 | 0.0429221041135 | 0.0830262820854303 |
| Meg3 | ENSMUSG00000021268 | 17263 | 0.423649598529823 | 2.29563913312646 | 2.23602061147689 | 0.0429370061078 | 0.08304041290957 |
| Ret | ENSMUSG00000030110 | 19713 | -0.321147886350697 | 2.50441272980205 | -2.23513067074811 | 0.0430081313033 | 0.0831632554885867 |
| Smarcd2 | ENSMUSG00000078619 | 83796 | 0.257919159699088 | 2.84964480994301 | 2.23495263435855 | 0.0430223734984 | 0.0831760814841065 |
| Slc46a3 | ENSMUSG00000029650 | 71706 | -0.512914912295382 | 0.991245897474279 | -2.2346708498857 | 0.0430449241926 | 0.0831917577546988 |
| Itga1 | ENSMUSG00000042284 | 109700 | -0.244999016349992 | 5.40092325199648 | -2.234661117995 | 0.0430457032165 | 0.0831917577546988 |
| Cntn1 | ENSMUSG00000055022 | 12805 | -0.743047615129665 | 0.850612370700701 | -2.23404460599944 | 0.0430950811683 | 0.083272464601662 |
| Myl6 | ENSMUSG00000090841 | 17904 | 0.197822444913591 | 5.5698822058414 | 2.23372263941924 | 0.0431208894344 | 0.0833076073537974 |
| Ncbp3 | ENSMUSG00000020783 | 66874 | -0.158204812218066 | 3.76665273939654 | -2.23342099560855 | 0.0431450818551 | 0.0833396164973448 |
| Stk24 | ENSMUSG00000063410 | 223255 | -0.165898095803495 | 4.53041274062247 | -2.23177335204363 | 0.0432774515699 | 0.0835519412178796 |
| Nhlrc2 | ENSMUSG00000025078 | 66866 | 0.14702353781087 | 4.45063333912868 | 2.23174421151786 | 0.0432797961191 | 0.0835519412178796 |
| Tgs1 | ENSMUSG00000028233 | 116940 | -0.191469355397069 | 3.48016490565047 | -2.23173553705411 | 0.0432804940604 | 0.0835519412178796 |
| Pcnx4 | ENSMUSG00000034501 | 67708 | 0.231881446760105 | 2.14140384612312 | 2.23167236592764 | 0.0432855770850 | 0.0835519412178796 |
| Gimap4 | ENSMUSG00000054435 | 107526 | -0.402933574319986 | 2.7144260966389 | -2.23108612406579 | 0.0433327754548 | 0.083628278336297 |
| Ap4e1 | ENSMUSG00000001998 | 108011 | 0.22052455750302 | 3.38467681634396 | 2.23067544364881 | 0.0433658681875 | 0.0836773707252642 |
| Eif5b | ENSMUSG00000026083 | 226982 | -0.155688183390796 | 6.78776179113679 | -2.23010028997274 | 0.0434122541715 | 0.0837487857755187 |
| Sash1 | ENSMUSG00000015305 | 70097 | 0.194032440244733 | 4.99708315613902 | 2.22997385133342 | 0.0434224576669 | 0.0837487857755187 |
| Grpel1 | ENSMUSG00000029198 | 17713 | -0.18679914625497 | 5.30180522141858 | -2.22993164912544 | 0.0434258638528 | 0.0837487857755187 |
| Arih2 | ENSMUSG00000064145 | 23807 | -0.183542429348742 | 5.37682741632296 | -2.22853577993091 | 0.0435386676632 | 0.0839458615399789 |
| Cerk | ENSMUSG00000035891 | 223753 | 0.234776287521241 | 3.03981423313249 | 2.22840424719966 | 0.0435493113574 | 0.0839458615399789 |
| Appl1 | ENSMUSG00000040760 | 72993 | 0.169807941139573 | 4.77091802877614 | 2.22838225193469 | 0.0435510916400 | 0.0839458615399789 |
| Zkscan14 | ENSMUSG00000029627 | 67235 | -0.296636788572407 | 2.34579551855927 | -2.22814169287617 | 0.0435705647005 | 0.0839685900462199 |
| Rcor1 | ENSMUSG00000037896 | 217864 | 0.204684334748488 | 3.52983885193662 | 2.22682655430729 | 0.0436771698627 | 0.0841592003823706 |
| Clic5 | ENSMUSG00000023959 | 224796 | 0.169693184348679 | 7.64340012719816 | 2.22644393310055 | 0.0437082311666 | 0.0842023519768109 |
| Akirin2 | ENSMUSG00000028291 | 433693 | -0.154884975239324 | 4.53043856298828 | -2.22636094443535 | 0.0437149709512 | 0.0842023519768109 |
| Rnf214 | ENSMUSG00000042790 | 235315 | -0.122195426750243 | 4.83505255579979 | -2.2254413342508 | 0.0437897209674 | 0.0843314728496728 |
| Strn | ENSMUSG00000024077 | 268980 | 0.235793507702189 | 3.97254685893191 | 2.22478243357887 | 0.0438433532184 | 0.0844019618601075 |
| Atp13a2 | ENSMUSG00000036622 | 74772 | 0.20558243252335 | 3.69590753461632 | 2.22469831353554 | 0.0438502047437 | 0.0844019618601075 |
| Golga4 | ENSMUSG00000038708 | 54214 | -0.187537323147525 | 9.60603500770243 | -2.22466211557755 | 0.0438531533546 | 0.0844019618601075 |
| Lztr1 | ENSMUSG00000022761 | 66863 | 0.179912973142994 | 4.00999779205686 | 2.22461234032069 | 0.0438572082485 | 0.0844019618601075 |
| Rgs2 | ENSMUSG00000026360 | 19735 | 0.45651809054664 | 2.70688409094265 | 2.22447882589103 | 0.0438680866150 | 0.0844080364002072 |
| Gstcd | ENSMUSG00000028018 | 67553 | 0.532651968364274 | 0.859766068796276 | 2.22431650614579 | 0.0438813153706 | 0.0844186304675408 |
| Prdx2 | ENSMUSG00000005161 | 21672 | -0.146785219574976 | 5.932213661137 | -2.22398967230067 | 0.0439079630938 | 0.0844444867835111 |

| | | | | | | | |
|---|---|---|---|---|---|---|---|
| 6430548M08Rik | ENSMUSG00000031824 | 234797 | 0.277135788276242 | 6.27533720487187 | 2.22388171062379 | 0.043916768867 | 0.0844444867835111 |
| Rock1 | ENSMUSG00000024290 | 19877 | 0.134570783373985 | 5.57579385119851 | 2.22381254429284 | 0.043922411214 | 0.0844444867835111 |
| Nup188 | ENSMUSG00000052533 | 227699 | 0.207942356836785 | 3.74429137578607 | 2.22377276523476 | 0.043925656559 | 0.0844444867835111 |
| Anapc5 | ENSMUSG00000029472 | 59008 | -0.156013876702175 | 6.68996601502019 | -2.22313951622933 | 0.043977350066 | 0.0845289983435929 |
| Fam110b | ENSMUSG00000049119 | 242297 | 0.556191665429636 | 2.02499141463683 | 2.22240750089805 | 0.044037177328 | 0.0846291113527619 |
| Ttl | ENSMUSG00000027394 | 69737 | 0.185802618710652 | 3.94766279699603 | 2.22193296490772 | 0.044076001779 | 0.0846888337939231 |
| Olfr804 | ENSMUSG00000095401 | 258068 | -0.929490312490801 | 0.911601583883219 | -2.22177367429898 | 0.044089041446 | 0.0846990003614386 |
| Itgb4 | ENSMUSG00000020758 | 192897 | -0.372115517816548 | 2.84974791740251 | -2.22142836972138 | 0.044117320826 | 0.0847384350643833 |
| Gask1b | ENSMUSG00000027955 | 68659 | -0.486093741917794 | 3.92847121752343 | -2.22124672224963 | 0.044132204034 | 0.0847521297076125 |
| Elmo1 | ENSMUSG00000041112 | 140580 | 0.267609507595353 | 2.98003962319545 | 2.22086789029302 | 0.044163258648 | 0.084796869891008 |
| Dynlt3 | ENSMUSG00000031176 | 67117 | 0.134845675079506 | 4.85220886406431 | 2.21988474330525 | 0.044243947584 | 0.0849368793542908 |
| Fkbp9 | ENSMUSG00000029781 | 27055 | 0.155806804227375 | 5.30112543867802 | 2.21899270491495 | 0.044317278873 | 0.0850627173603598 |
| Aatf | ENSMUSG00000018697 | 56321 | 0.171237622642904 | 3.31466381997229 | 2.21754002474953 | 0.044436942936 | 0.0852774268324714 |
| Pa2g4 | ENSMUSG00000025364 | 18813 | 0.152989196976423 | 5.34269658542973 | 2.21700227865398 | 0.044481316517 | 0.0853402283850788 |
| Xrcc5 | ENSMUSG00000026187 | 22596 | 0.198126854945621 | 3.30518471976923 | 2.21689150380718 | 0.044490462573 | 0.0853402283850788 |
| Gid8 | ENSMUSG00000027573 | 76425 | -0.217861803314842 | 4.27223728424877 | -2.21685969078303 | 0.044493089522 | 0.0853402283850788 |
| Tdp2 | ENSMUSG00000035958 | 56196 | -0.237741673661497 | 2.17041079452789 | -2.21614508659214 | 0.044552136105 | 0.0854384911877036 |
| Xpa | ENSMUSG00000028329 | 22590 | 0.233908310015069 | 2.45126931064203 | 2.2147713256788 | 0.044665854516 | 0.0856415463469865 |
| Ppp1r21 | ENSMUSG00000034709 | 73825 | 0.239370389099086 | 2.53968291966548 | 2.213786094252 | 0.044747578664 | 0.0857831957877325 |
| Mnt | ENSMUSG00000000282 | 17428 | 0.361660783075203 | 2.96328855425477 | 2.2133192379776 | 0.044786353022 | 0.0858424732401511 |
| Sap18b | ENSMUSG00000061104 | 100041953 | -0.175580411125901 | 4.16730681983001 | -2.21200378192206 | 0.044895776861 | 0.0860371207702074 |
| Rab23 | ENSMUSG00000004768 | 19335 | 0.353990899563384 | 2.66878293976719 | 2.21109414121211 | 0.044971590415 | 0.0861673014609578 |
| Slc11a2 | ENSMUSG00000023030 | 18174 | 0.169161068048394 | 3.17180711886384 | 2.21047361427252 | 0.045023376860 | 0.0862514077227358 |
| Twf1 | ENSMUSG00000022451 | 19230 | 0.230740469236877 | 3.45963571575042 | 2.2100805133531 | 0.045056212288 | 0.0862991863548381 |
| Slc12a6 | ENSMUSG00000096764 | 107723 | -0.719593735496931 | 1.15947859144728 | -2.20986536730915 | 0.045074192793 | 0.0863185006006869 |
| Rab8a | ENSMUSG00000003037 | 17274 | 0.237023835658838 | 3.21368771711342 | 2.20861725505258 | 0.045178634817 | 0.0865033556118091 |
| Lbx1 | ENSMUSG00000025216 | 16814 | -0.318557279773145 | 4.88761485306487 | -2.2083220598593 | 0.045203369938 | 0.0865355581073216 |
| Clu | ENSMUSG00000022037 | 12759 | 0.338832630577638 | 5.04487899833627 | 2.20810925218727 | 0.045221209491 | 0.0865545510333408 |
| Rsbn1l | ENSMUSG00000039968 | 242860 | -0.180537836598887 | 3.76961838844718 | -2.20752397529285 | 0.045270307019 | 0.086633355396735 |
| Ddit3 | ENSMUSG00000025408 | 13198 | -0.406319124023964 | 4.11778172891474 | -2.20735835538658 | 0.045284209568 | 0.0866378463845425 |
| Esf1 | ENSMUSG00000045624 | 66580 | -0.165569207488645 | 4.45435160447137 | -2.20730718728632 | 0.045288505556 | 0.0866378463845425 |
| Exosc1 | ENSMUSG00000034321 | 66583 | 0.277135632610896 | 2.31236136033264 | 2.20700196352634 | 0.045314139580 | 0.0866682700233269 |
| Cadm3 | ENSMUSG00000005338 | 94332 | 0.344112382337861 | 2.76637364821076 | 2.20692903622057 | 0.045320266348 | 0.0866682700233269 |
| 1810013L24Rik | ENSMUSG00000022507 | 69053 | -0.192513404612926 | 5.5858513596882 | -2.20679089211402 | 0.045331874240 | 0.08667530476186 |
| Vmn2r76 | ENSMUSG00000091239 | 675969 | -0.899772495394337 | 2.23731983306912 | -2.206551003329 | 0.045352038103 | 0.0866962226505492 |
| Olfr1385 | ENSMUSG00000059729 | 258027 | -0.989421032456619 | 3.66246477970081 | -2.20647204354078 | 0.045358676913 | 0.0866962226505492 |
| Ncoa4 | ENSMUSG00000056234 | 27057 | -0.145882580704782 | 4.95950922382062 | -2.2062897266813 | 0.045374009299 | 0.0867103663721991 |
| Zfp715 | ENSMUSG00000012640 | 69930 | -0.16057001041486 | 3.27654880458454 | -2.20602727079184 | 0.045396089713 | 0.0867373984717905 |
| Inpp1 | ENSMUSG00000026102 | 16329 | -0.234232091624218 | 3.19076168285544 | -2.20587183663094 | 0.045409171142 | 0.0867472299470231 |
| Anapc2 | ENSMUSG00000026965 | 99152 | 0.156868852314382 | 4.71666815531779 | 2.20495879131162 | 0.045486084997 | 0.0868789787011599 |
| Snap23 | ENSMUSG00000027287 | 20619 | -0.147720603331233 | 4.56256714194947 | -2.20297679963686 | 0.045653466418 | 0.0871834453915194 |
| Nrd1 | ENSMUSG00000053510 | 230598 | -0.173305837782274 | 7.62369257387886 | -2.20253060441684 | 0.045691227695 | 0.087240316146376 |
| Eif3e | ENSMUSG00000022336 | 16341 | -0.186793659672978 | 6.03287199844159 | -2.20229345862337 | 0.045711309144 | 0.0872634160431 |
| Poll | ENSMUSG00000025218 | 56626 | -0.233196927593387 | 2.70226229289742 | -2.20139398600547 | 0.045787551537 | 0.0873937010398339 |
| Fam173a | ENSMUSG00000057411 | 214917 | 0.2549120752189 | 2.73716315894383 | 2.20091829504566 | 0.045827920936 | 0.0874332070796584 |
| Flot1 | ENSMUSG00000059714 | 14251 | 0.180299361822225 | 5.76975352507642 | 2.20090731893312 | 0.045828852815 | 0.0874332070796584 |
| Cdc42ep2 | ENSMUSG00000045664 | 104252 | -0.232854455113645 | 3.27380923696592 | -2.20086735897112 | 0.045832245592 | 0.0874332070796584 |
| Zfp574 | ENSMUSG00000045252 | 232976 | -0.222831015779431 | 2.88210996732522 | -2.2005837619404 | 0.045856330995 | 0.0874638900919051 |
| Cdk4 | ENSMUSG00000006728 | 12567 | 0.21387264852712 | 3.41759012012734 | 2.20023615472136 | 0.045885868864 | 0.0875049603199327 |
| Rrm2b | ENSMUSG00000022292 | 382985 | -0.197638493075866 | 4.06570154213132 | -2.2000887224731 | 0.045898402283 | 0.0875135941673739 |
| Rnf5 | ENSMUSG00000015478 | 54197 | -0.36014961628037 | 1.72241832346797 | -2.19944593280563 | 0.045953084239 | 0.0876025747861212 |
| Ccdc9b | ENSMUSG00000045838 | 214239 | -0.35979507053798 | 4.83067841743355 | -2.19927402321995 | 0.045967718894 | 0.0876151935896745 |
| Ppdpf | ENSMUSG00000016344 | 66496 | -0.185123241878132 | 3.34762810628085 | -2.19838744457033 | 0.046043262700 | 0.0877438815505624 |
| Kank1 | ENSMUSG00000032702 | 107351 | 0.160749837307005 | 4.48672567319434 | 2.19796301393523 | 0.046079468871 | 0.0877975726404612 |
| Chd6 | ENSMUSG00000057133 | 71389 | -0.180763623583416 | 4.80986941748577 | -2.19665743772056 | 0.046191008807 | 0.0879947572809705 |
| Snrpc | ENSMUSG00000024217 | 20630 | -0.165181532608299 | 3.67895241694561 | -2.19653895020045 | 0.046201144127 | 0.0879987291272012 |
| Zfp3 | ENSMUSG00000043602 | 193043 | -0.246803119998492 | 3.27137074142889 | -2.19546804189343 | 0.046292843308 | 0.0881580261675433 |
| Farp2 | ENSMUSG00000034066 | 227377 | 0.247459806370956 | 3.44673378424305 | 2.19495169603338 | 0.046337117631 | 0.0882269696629155 |
| Lmf2 | ENSMUSG00000022614 | 105847 | 0.192380108418021 | 3.44242211451725 | 2.19427905919147 | 0.046394852724 | 0.0883215143034862 |
| Tmf1 | ENSMUSG00000030059 | 232286 | 0.128415925739435 | 6.19106128763348 | 2.19395135175958 | 0.046423005572 | 0.088359720340535 |
| Strada | ENSMUSG00000069631 | 72149 | 0.252573420870593 | 2.3280905669224 | 2.19180933655676 | 0.046607417513 | 0.088695278697317 |
| Cyp27a1 | ENSMUSG00000026170 | 104086 | -0.352197067973742 | 2.71056105989279 | -2.19067209306978 | 0.046705604638 | 0.088856695893431 |
| Vmn1r58 | ENSMUSG00000078808 | 81014 | -0.853009769522452 | 2.70957865837506 | -2.19063863299272 | 0.046708496441 | 0.088856695893431 |
| Edem2 | ENSMUSG00000038312 | 108687 | 0.389595910517847 | 1.4708495732119 | 2.19045167122172 | 0.046724657787 | 0.0888719739468222 |
| Ahcyl2 | ENSMUSG00000029772 | 74340 | -0.146866863006665 | 4.39792773149126 | -2.19032105995648 | 0.046735951191 | 0.0888779892962451 |
| Ppp3cc | ENSMUSG00000022092 | 19057 | -0.262860070379889 | 3.64597769851078 | -2.1896241463055 | 0.046796253605 | 0.0889771870176096 |
| Olfr986 | ENSMUSG00000051095 | 258608 | -0.747207421783487 | 2.72780261266849 | -2.18920651965007 | 0.046832424843 | 0.0890220272046438 |
| Telo2 | ENSMUSG00000024170 | 71718 | 0.295997289540189 | 2.91044986890033 | 2.18916381948179 | 0.046836124641 | 0.0890220272046438 |
| Tada3 | ENSMUSG00000048930 | 101206 | 0.195812209193176 | 2.81297343179408 | 2.18838809315387 | 0.046903385886 | 0.0891343725875038 |
| Cp | ENSMUSG00000003617 | 12870 | 0.235726812958189 | 4.57183010808504 | 2.18770269817477 | 0.046962890003 | 0.0892319399658447 |
| Fut8 | ENSMUSG00000021065 | 53618 | 0.152834537114479 | 4.28102895639882 | 2.1867837383627 | 0.047042782528 | 0.0893682057391451 |
| Mllt6 | ENSMUSG00000038437 | 246198 | -0.164618054643201 | 6.13121399919022 | -2.18607903435991 | 0.047104134289 | 0.0894642981246307 |
| Thrap3 | ENSMUSG00000043962 | 230753 | -0.157428791632559 | 6.38218201558633 | -2.18601475974985 | 0.047109733785 | 0.0894642981246307 |
| Cul1 | ENSMUSG00000029686 | 26965 | 0.133729197432461 | 5.57445271480557 | 2.18585628687663 | 0.047123542337 | 0.0894749767743893 |
| Ocrl | ENSMUSG00000001173 | 320634 | 0.246225860070437 | 3.24802286983431 | 2.18487701973566 | 0.047208954997 | 0.0896215851126279 |
| Rbm18 | ENSMUSG00000026889 | 67889 | -0.170467472298672 | 3.86964652023043 | -2.18471093959657 | 0.047223455042 | 0.0896335452447917 |
| Eps15l1 | ENSMUSG00000006276 | 13859 | -0.184708238091305 | 4.53774037160932 | -2.18440108934961 | 0.047250518444 | 0.0896693432503092 |
| Emc8 | ENSMUSG00000031819 | 18117 | -0.209720102845258 | 3.36331694841201 | -2.18415359728762 | 0.047272145696 | 0.0896948142012596 |
| Tmem87a | ENSMUSG00000033808 | 211499 | 0.192396338196474 | 3.78893404538816 | 2.18385749206537 | 0.047298033218 | 0.0897065947701168 |
| Tmx1 | ENSMUSG00000021072 | 72736 | 0.218139956104078 | 3.2181302426106 | 2.18377775508634 | 0.047305006632 | 0.0897065947701168 |
| Phldb2 | ENSMUSG00000033149 | 208177 | -0.231836610726858 | 3.11843813383065 | -2.18372081651844 | 0.047309986795 | 0.0897065947701168 |
| Ap1g1 | ENSMUSG00000031731 | 11765 | -0.147719161159982 | 5.3203550493106 | -2.18370716461494 | 0.047311180939 | 0.0897065947701168 |
| Foxk2 | ENSMUSG00000039275 | 68837 | -0.138412129973396 | 5.20524987700792 | -2.18307449209182 | 0.047366552335 | 0.0897960082526041 |
| Brk1 | ENSMUSG00000033940 | 101314 | 0.178583314377721 | 3.36071290811771 | 2.18228578350195 | 0.047435664937 | 0.0899114363496898 |
| Taf5 | ENSMUSG00000025049 | 226182 | -0.392732978588724 | 1.32210881752918 | -2.181614771396 | 0.047494538388 | 0.0900074200970239 |
| Atrn | ENSMUSG00000027312 | 11990 | 0.148060575274053 | 4.68605859235651 | 2.18016918255621 | 0.047621604323 | 0.0902325804931691 |
| Armc6 | ENSMUSG00000002343 | 76813 | 0.313966362819213 | 1.49356775898135 | 2.17997711690795 | 0.047638510642 | 0.0902489705077707 |
| Prkaca | ENSMUSG00000005469 | 18747 | 0.170208540223835 | 6.73662678459655 | 2.17916271644454 | 0.047710259559 | 0.0903597888129109 |
| Zfp703 | ENSMUSG00000085795 | 353310 | 0.215705375099668 | 5.53464100550294 | 2.17912551343633 | 0.047713539568 | 0.0903597888129109 |
| Synj1 | ENSMUSG00000022973 | 104015 | 0.139187746009677 | 4.25880827496576 | 2.17831014856818 | 0.047785479367 | 0.0904679597464922 |
| Macc1 | ENSMUSG00000041886 | 238455 | -1.23596969672868 | 0.052104037050001 | -2.17829054042666 | 0.047787210646 | 0.0904679597464922 |
| Hax1 | ENSMUSG00000027944 | 23897 | -0.210090409044191 | 3.34826223993326 | -2.17763046235825 | 0.047845525761 | 0.0905618452329136 |
| Ptpn4 | ENSMUSG00000026384 | 19258 | -0.203993173267148 | 5.71873885564002 | -2.17754169796192 | 0.047853372798 | 0.0905618452329136 |
| Psma6 | ENSMUSG00000021024 | 26443 | -0.155449318381454 | 5.22208068196324 | -2.17651048621817 | 0.047944623280 | 0.0907188293377246 |
| S100a16 | ENSMUSG00000074457 | 67860 | 0.20298411872736 | 3.68131937916773 | 2.17611133682804 | 0.047979987129 | 0.0907700310342924 |
| Zswim8 | ENSMUSG00000021819 | 268721 | 0.221999741033553 | 4.73843665604129 | 2.17573727846733 | 0.048013150110 | 0.090817052059905 |
| Ube4a | ENSMUSG00000059890 | 140630 | -0.150294178970946 | 4.58048978848326 | -2.17532650202498 | 0.048049593115 | 0.0908551203924096 |
| Eloa | ENSMUSG00000028668 | 27224 | -0.162539396020701 | 6.27933880412724 | -2.17532305023343 | 0.048049899459 | 0.0908551203924096 |
| Dusp6 | ENSMUSG00000019960 | 67603 | 0.241396996335382 | 3.5711677192794 | 2.17442958201317 | 0.048129255443 | 0.0909894312095667 |
| Spata1 | ENSMUSG00000028188 | 70951 | -0.36938147777691 | 1.72405403862105 | -2.17413567445738 | 0.048155386482 | 0.0910230900299563 |
| Tmem267 | ENSMUSG00000074634 | 633640 | -0.506300252008325 | 1.19160363404441 | -2.17391488486057 | 0.048175025401 | 0.0910444679579054 |
| Rab14 | ENSMUSG00000026878 | 68365 | -0.13580186805186 | 6.03115776033585 | -2.17332336078141 | 0.048227677547 | 0.0911282183695292 |
| Rapgef2 | ENSMUSG00000062232 | 76089 | -0.183014675534921 | 4.92040371795459 | -2.17272097037593 | 0.048281352220 | 0.091213871607162 |
| Dync1li2 | ENSMUSG00000035770 | 234663 | 0.176272944859869 | 6.23517134596892 | 2.17241496941202 | 0.048308639154 | 0.0912496517370576 |
| Fscn1 | ENSMUSG00000029581 | 14086 | 0.37287912449405 | 3.5733441666901 | 2.17206482123997 | 0.048339880507 | 0.0912845831691451 |
| Cavin4 | ENSMUSG00000028348 | 68016 | 0.203048332509627 | 6.63274830814325 | 2.17196346277786 | 0.048348927562 | 0.0912845831691451 |
| Fgfr1op2 | ENSMUSG00000040242 | 67529 | -0.133840039750719 | 5.75933562261602 | -2.1719269701307 | 0.048352185211 | 0.0912845831691451 |
| Alkbh6 | ENSMUSG00000042831 | 233065 | -0.254703098484827 | 1.91720191321575 | -2.17159488732165 | 0.048381839225 | 0.0913247944345162 |
| Grik5 | ENSMUSG00000003378 | 14809 | 0.352685018539939 | 1.37301678671638 | 2.17035826401519 | 0.048492415662 | 0.0915177133297844 |
| B2m | ENSMUSG00000060802 | 12010 | -0.333759078686732 | 6.08111898292524 | -2.16949002351843 | 0.048570193164 | 0.0916486762448167 |
| Ptch1 | ENSMUSG00000021466 | 19206 | -0.268507866204255 | 3.10136641731637 | -2.16937706902017 | 0.048580320265 | 0.0916519642419447 |
| Gon4l | ENSMUSG00000054199 | 76022 | -0.143612912758061 | 4.4693868926081 | -2.16879553039716 | 0.048632490207 | 0.0917345557299097 |
| Glis2 | ENSMUSG00000014303 | 83396 | 0.398663757067216 | 2.31724107230333 | 2.16852805274016 | 0.048656503259 | 0.0917640160669696 |
| Tprgl | ENSMUSG00000029030 | 67808 | -0.173206221271138 | 5.09849997806383 | -2.1681868952569 | 0.048687147068 | 0.0917924966282479 |
| Smim7 | ENSMUSG00000044600 | 66818 | 0.188115732255354 | 4.01443708286185 | 2.16817295567827 | 0.048688399547 | 0.0917924966282479 |
| Ube2g2 | ENSMUSG00000009293 | 22213 | 0.196839973585456 | 5.74875452441136 | 2.16788158802079 | 0.048714585993 | 0.0918126131287701 |

| | | | | | | | |
|---|---|---|---|---|---|---|---|
| Rps12 | ENSMUSG00000061983 | 20042 | 0.199559672265128 | 5.27967512361851 | 2.16786732423386 | 0.048715868278 | 0.0918126131287701 |
| Elof1 | ENSMUSG00000013822 | 66126 | -0.218948519599107 | 2.66377065915692 | -2.1675622528373 | 0.048743301119 | 0.0918458275699316 |
| Fkbp2 | ENSMUSG00000056629 | 14227 | 0.243421193394807 | 2.27628726951368 | 2.16748448471023 | 0.048750296547 | 0.0918458275699316 |
| Ifi27l2a | ENSMUSG00000079017 | 76933 | 0.774450345563287 | 1.87067599705949 | 2.16695536273423 | 0.048797917247 | 0.0918954104978114 |
| Stk3 | ENSMUSG00000022329 | 56274 | 0.222868035973967 | 2.60240382228417 | 2.16683100886127 | 0.048809115338 | 0.0918954104978114 |
| Cacna1h | ENSMUSG00000024112 | 58226 | -0.460053488870313 | 1.81869320836011 | -2.16682902026172 | 0.048809294431 | 0.0918954104978114 |
| Pqlc1 | ENSMUSG00000034006 | 66943 | 0.262403797893961 | 4.74363859862553 | 2.16678423743206 | 0.048813327736 | 0.0918954104978114 |
| Mfsd14b | ENSMUSG00000038212 | 66631 | 0.262828686815047 | 3.1765548414419 | 2.16663047505142 | 0.048827178507 | 0.0918954104978114 |
| Rab11fip5 | ENSMUSG00000051343 | 52055 | 0.249399914839328 | 6.65492780134863 | 2.16648634743983 | 0.048840164722 | 0.0918954104978114 |
| Cpt1a | ENSMUSG00000024900 | 12894 | 0.458558866099958 | 3.697086429647 | 2.16636905569873 | 0.048850735348 | 0.0918954104978114 |
| Lamtor3 | ENSMUSG00000091512 | 56692 | -0.267202622570093 | 4.22697494183485 | -2.16635947656773 | 0.048851598738 | 0.0918954104978114 |
| Gm29683 | ENSMUSG00000109097 | 624549 | -1.20633369306248 | 0.706240696371324 | -2.16635196112883 | 0.048852276132 | 0.0918954104978114 |
| Ftsj3 | ENSMUSG00000020706 | 56095 | 0.150387991514852 | 4.59676503666969 | 2.16545565557292 | 0.048933126444 | 0.0920277824496485 |
| Vmn2r108 | ENSMUSG00000091805 | 627805 | -1.06155874078172 | 2.16085866509726 | -2.16538523472434 | 0.048939483979 | 0.0920277824496485 |
| Ppp2r3a | ENSMUSG00000043154 | 235542 | -0.167826386218398 | 8.05851772368558 | -2.16513268826118 | 0.048962290008 | 0.0920548318002036 |
| Gm13387 | ENSMUSG00000087028 | 329358 | -0.637263723732915 | 0.596354844154735 | -2.16470948925074 | 0.049000528960 | 0.0921103721358044 |
| Gfer | ENSMUSG00000040888 | 11692 | -0.171940066030325 | 3.8833997312443 | -2.16461927600494 | 0.049008683957 | 0.0921103721358044 |
| Myadm | ENSMUSG00000068566 | 50918 | 0.364810632233206 | 5.20762313501047 | 2.16413851207273 | 0.049052164913 | 0.0921640318996542 |
| Ptk2 | ENSMUSG00000022607 | 14083 | -0.201218327462565 | 4.82085552700065 | -2.16411715513698 | 0.049054097300 | 0.0921640318996542 |
| Gin1 | ENSMUSG00000026333 | 252876 | -0.686444493360593 | 1.96201587378038 | -2.16283831560064 | 0.049169937098 | 0.0923657986637082 |
| Fat3 | ENSMUSG00000074505 | 270120 | -0.415976704617064 | 1.43117373506512 | -2.16215430355617 | 0.049232001106 | 0.0924578842064116 |
| Angptl4 | ENSMUSG00000002289 | 57875 | 1.55703522194457 | 1.82778290503306 | 2.16211164150424 | 0.049235874482 | 0.0924578842064116 |
| Irak4 | ENSMUSG00000059883 | 266632 | 0.521585056918358 | 1.266601978064 | 2.1619325865366 | 0.0492521343571 | 0.0924725318889968 |
| Stmn1 | ENSMUSG00000028832 | 16765 | 0.487331534643457 | 1.255623732337 | 2.16156916172498 | 0.049285152171 | 0.0925186327295128 |
| Zranb1 | ENSMUSG00000030967 | 360216 | -0.205651466038624 | 5.42654524633341 | -2.1613260604201 | 0.049307249912 | 0.0925364310855203 |
| Vps41 | ENSMUSG00000041236 | 218035 | -0.137322857166279 | 4.31290131268741 | -2.16119332237609 | 0.049319319615 | 0.0925364310855203 |
| Ube2f | ENSMUSG00000034343 | 67921 | -0.147749213765684 | 4.31557531081765 | -2.16115617959172 | 0.049322697456 | 0.0925364310855203 |
| Slfn8 | ENSMUSG00000035208 | 276950 | 0.755467017643224 | 0.975322467865077 | 2.16109243013337 | 0.049328495465 | 0.0925364310855203 |
| Tnfrsf19 | ENSMUSG00000060548 | 29820 | 0.342644984130758 | 3.58316003610724 | 2.16085642842044 | 0.0493499653491 | 0.0925608221056434 |
| Trib1 | ENSMUSG00000032501 | 211770 | 0.358128202563459 | 4.69437146404599 | 2.16076231323231 | 0.049358529764 | 0.0925610034055343 |
| Ergic3 | ENSMUSG00000005881 | 66366 | 0.185157920934896 | 3.96921485217515 | 2.16058856662347 | 0.049374344228 | 0.0925747781752652 |
| Gtpbp3 | ENSMUSG00000007610 | 70359 | -0.23780107021557 | 2.71654962078343 | -2.16033304531122 | 0.049397610447 | 0.0926025175436752 |
| Rpl17-ps8 | ENSMUSG00000082035 | 102632403 | 0.237279087513637 | 3.4575612234205 | 2.15787656622744 | 0.049621805472 | 0.0930068511013283 |
| Klhdc1 | ENSMUSG00000051890 | 271005 | -0.232000071121239 | 3.81507733097729 | -2.15582612426966 | 0.049809669937 | 0.0933429628123626 |
| Ddx41 | ENSMUSG00000021494 | 72935 | 0.194981480282104 | 2.80869998696939 | 2.15524365769677 | 0.049863157347 | 0.0934271808306729 |
| Zmat5 | ENSMUSG00000009076 | 67178 | -0.305518219848237 | 1.98540673929766 | -2.15506959629607 | 0.049879151660 | 0.0934411322709236 |
| Dis3 | ENSMUSG00000033166 | 72662 | 0.166069170658855 | 3.81791981878142 | 2.15433642259648 | 0.049946574779 | 0.0935514065994021 |
| H2-T22 | ENSMUSG00000056116 | 15039 | -0.247825288522936 | 2.88750970772516 | -2.15412147682105 | 0.049966357449 | 0.0935724264656838 |
| Tfdp1 | ENSMUSG00000038482 | 21781 | 0.156609986820779 | 4.70103222389148 | 2.15262926682286 | 0.050103895683 | 0.0938139232137123 |
| Igsf11 | ENSMUSG00000022790 | 207683 | -0.694284120949391 | 1.19648230438214 | -2.15228272180884 | 0.050135887541 | 0.0938577473400654 |
| Psmb8 | ENSMUSG00000024338 | 16913 | -0.385289189591541 | 2.72424160331775 | -2.15217909749437 | 0.050145457487 | 0.0938595883220078 |
| Capza1 | ENSMUSG00000070372 | 12340 | 0.189277642157493 | 4.44392837773976 | 2.15181591723151 | 0.050179011487 | 0.0939063130578004 |
| Tlr3 | ENSMUSG00000031639 | 142980 | 0.323379076092438 | 1.28470685972316 | 2.15171915797313 | 0.050187954549 | 0.093906972129445 |
| Rpl26 | ENSMUSG00000060938 | 19941 | -0.184350444945821 | 7.43036406962751 | -2.1505535081265 | 0.050295807735 | 0.094092670606554 |
| Arhgap17 | ENSMUSG00000030766 | 70497 | 0.164137922738036 | 3.90987294461343 | 2.14970905310028 | 0.050374077016 | 0.094222969860015 |
| Lynx1 | ENSMUSG00000022594 | 23936 | -0.196043999782091 | 7.31691205181114 | -2.14927517194079 | 0.050414335966 | 0.0942821396837609 |
| Rtl6 | ENSMUSG00000055745 | 223732 | 0.231528195262024 | 2.40647899199075 | 2.14905823616404 | 0.050434476250 | 0.0942980938040287 |
| Zbtb6 | ENSMUSG00000066798 | 241322 | 0.25333231906778 | 2.41482659546781 | 2.14899745778303 | 0.050440120251 | 0.0942980938040287 |
| Leprotl1 | ENSMUSG00000031513 | 68192 | 0.266226120714253 | 2.81674758331685 | 2.14835091687429 | 0.0505001958091 | 0.0943942613525703 |
| Bcl10 | ENSMUSG00000028191 | 12042 | 0.18003564517842 | 2.95426774040207 | 2.14783347953117 | 0.050548323344 | 0.0944680667607675 |
| 4930402H24Rik | ENSMUSG00000027309 | 228602 | 0.289941719274423 | 3.30506322076465 | 2.1476502391293 | 0.050565377034 | 0.0944837839935239 |
| Zfhx3 | ENSMUSG00000038872 | 11906 | 0.16309891560791 | 5.57174902158775 | 2.14720770019509 | 0.050606585071 | 0.0945424127849978 |
| Srsf6 | ENSMUSG00000016921 | 67996 | 0.190719291728561 | 4.87065508073321 | 2.14712755050624 | 0.050614051744 | 0.0945424127849978 |
| Tbl1x | ENSMUSG00000025246 | 21372 | 0.17640115795505 | 4.12193542000881 | 2.14648829720378 | 0.050673640798 | 0.0946341236070078 |
| Vmn1r88 | ENSMUSG00000117853 | 100312474 | -0.656193362990188 | 1.49935163024335 | -2.14641513724386 | 0.050680464696 | 0.0946341236070078 |
| Rlim | ENSMUSG00000056537 | 19820 | -0.164621141810645 | 5.72897017332724 | -2.14602826408842 | 0.050716564025 | 0.0946853563387391 |
| 0610010F05Rik | ENSMUSG00000042208 | 71675 | -0.224528057669628 | 3.52949329271397 | -2.14572213207659 | 0.050745146351 | 0.0947225400910644 |
| Kif3c | ENSMUSG00000020668 | 16570 | 0.213621666512311 | 4.44566694699134 | 2.14517134225207 | 0.050796609234 | 0.0948024134441316 |
| BC017158 | ENSMUSG00000030780 | 233913 | 0.33871192658518 | 1.60240268675193 | 2.14474132820415 | 0.050836821295 | 0.0948612655476619 |
| Wdr37 | ENSMUSG00000021147 | 207615 | -0.217009433200905 | 4.28749003182841 | -2.14377777642547 | 0.050927034064 | 0.0950105045552519 |
| Fam98a | ENSMUSG00000002017 | 72722 | -0.157251468493943 | 4.17540961072818 | -2.14370148530543 | 0.050934183212 | 0.0950105045552519 |
| Tent5c | ENSMUSG00000044468 | 74645 | -0.385421807404423 | 1.48560962069014 | -2.14231304328035 | 0.051064456009 | 0.0952372579142453 |
| Zfp28 | ENSMUSG00000062861 | 22690 | 0.327537795953503 | 1.45733722323359 | 2.14219688583625 | 0.051075368736 | 0.0952413605691562 |
| Hoxc11 | ENSMUSG00000001656 | 109663 | 0.329927091557669 | 1.97994058491182 | 2.14176412944442 | 0.051116044353 | 0.0953009518733079 |
| Mras | ENSMUSG00000032470 | 17532 | 0.309139750169372 | 2.28320385201198 | 2.1410396396556 | 0.051184208162 | 0.095411763202913 |
| Msc | ENSMUSG00000025930 | 17681 | 0.66152380730102 | 0.522252005699239 | 2.14021286111598 | 0.051262099345 | 0.0955406663161159 |
| Dnm3 | ENSMUSG00000040265 | 103967 | 0.371885602463034 | 2.00463826329959 | 2.14004194143991 | 0.051278215540 | 0.0955544108543495 |
| Slx1b | ENSMUSG00000059772 | 75764 | 0.327144538434953 | 1.85746063505051 | 2.13954150393526 | 0.051325429474 | 0.0956260899245197 |
| Twist1 | ENSMUSG00000035799 | 22160 | 0.619693332538795 | 0.550971672059294 | 2.13938750826739 | 0.051339966394 | 0.0956368733227321 |
| H2afx | ENSMUSG00000049932 | 15270 | 0.412940494898684 | 1.54466752545642 | 2.13761447668044 | 0.051507614088 | 0.0959328215374575 |
| Gna13 | ENSMUSG00000020611 | 14674 | 0.16014811442909 | 5.40381468645333 | 2.13732540341727 | 0.051534995514 | 0.0959674677973792 |
| Ric1 | ENSMUSG00000038658 | 226089 | -0.181673033672815 | 4.7560465571081 | -2.13718430210743 | 0.051548365754 | 0.0959760153399806 |
| Fzd2 | ENSMUSG00000050288 | 57265 | 0.374913951722795 | 1.21596623820091 | 2.13692209679397 | 0.051573219946 | 0.0960059378808196 |
| Sybu | ENSMUSG00000022340 | 319613 | -0.920845408712371 | 0.412423129983547 | -2.13653141823177 | 0.051610272708 | 0.0960585545669539 |
| Zfand2a | ENSMUSG00000053581 | 100494 | 0.251347813903821 | 2.57664493914176 | 2.13640644920067 | 0.051622130263 | 0.0960642672646431 |
| Arhgap32 | ENSMUSG00000041444 | 330914 | 0.32999072933857 | 2.92416703929273 | 2.13520460002067 | 0.051736296113 | 0.0962603323937976 |
| Galnt18 | ENSMUSG00000038296 | 233733 | -0.320274160052125 | 2.20037456827829 | -2.13437912266855 | 0.051814845921 | 0.0963900750114159 |
| Edn1 | ENSMUSG00000021367 | 13614 | -0.676954484849589 | 0.804628577642655 | -2.13273059934619 | 0.051972046721 | 0.0966660613761724 |
| Otud4 | ENSMUSG00000036990 | 73945 | -0.126655082791265 | 5.75885141272411 | -2.13257116975318 | 0.051987273229 | 0.0966779318936409 |
| Gpr107 | ENSMUSG00000000194 | 277463 | 0.156847023806368 | 4.36413974875623 | 2.13246812374152 | 0.051997116968 | 0.0966797900297404 |
| Srxn1 | ENSMUSG00000032802 | 76650 | -0.261730886828516 | 4.64345538047356 | -2.1318578059475 | 0.052055454763 | 0.0967717986435441 |
| LTO1 | ENSMUSG00000031072 | 72284 | 0.265874411280676 | 2.35755803083683 | 2.13156992977952 | 0.052082992825 | 0.0968065285795675 |
| Ppm1b | ENSMUSG00000061130 | 19043 | -0.14635931014839 | 7.52861126622348 | -2.13126026269072 | 0.0521126305381 | 0.0968451486591363 |
| Ten1 | ENSMUSG00000020778 | 69535 | 0.431258539044946 | 1.29463953769735 | 2.13084380930817 | 0.052152513347 | 0.0969027916720192 |
| Zfp367 | ENSMUSG00000044934 | 238673 | 0.291964812860309 | 2.91784413659709 | 2.12745362167897 | 0.052478242939 | 0.097491446901847 |
| Pex16 | ENSMUSG00000027222 | 18633 | -0.244603891007724 | 2.18592419923463 | -2.12714739639866 | 0.052507758162 | 0.0975297033949564 |
| Dcun1d5 | ENSMUSG00000032002 | 76863 | -0.181098403883977 | 5.12021456490819 | -2.12588428004111 | 0.052629665771 | 0.0977395305046168 |
| Itm2b | ENSMUSG00000022108 | 16432 | 0.152019160133013 | 7.40493015955215 | 2.1257544949602 | 0.052642206678 | 0.0977462138953667 |
| Cdc25b | ENSMUSG00000027330 | 12531 | 0.632389343101042 | 0.944769088824668 | 2.12547212365291 | 0.052669501334 | 0.0977779087903425 |
| Vmn1r76 | ENSMUSG00000115267 | 171239 | -1.02749369596364 | 2.51340972016608 | -2.12539285177098 | 0.052677166303 | 0.0977779087903425 |
| Zfp93 | ENSMUSG00000055305 | 22755 | -0.378790037772788 | 1.46791157797424 | -2.1251091197746 | 0.052704609468 | 0.0978122387265274 |
| Vmn1r236 | ENSMUSG00000054142 | 171235 | -1.2064206694251 | -0.386917245862325 | -2.12478096228814 | 0.052736366153 | 0.0978545609269488 |
| Sem1 | ENSMUSG00000042541 | 20422 | -0.157197323659606 | 5.50065752293201 | -2.1238800905113 | 0.052823637500 | 0.0979998610853908 |
| Cct8 | ENSMUSG00000025613 | 12469 | 0.124531864348685 | 5.85988002141655 | 2.12337937486915 | 0.052872202073 | 0.0980657227695311 |
| Zfp748 | ENSMUSG00000095432 | 212276 | -0.392589737137343 | 1.97160709997815 | -2.12332909756384 | 0.052877080779 | 0.0980657227695311 |
| Arhgef7 | ENSMUSG00000031511 | 54126 | -0.145927228984567 | 5.82458999412702 | -2.12292016804309 | 0.052916777207 | 0.098122695785947 |
| Rnf217 | ENSMUSG00000063760 | 268291 | 0.193349137474887 | 4.14659841831033 | 2.12268647511751 | 0.052939475164 | 0.0981438986084454 |
| Adamtsl4 | ENSMUSG00000015850 | 229595 | 0.250145236840336 | 5.00075304974723 | 2.12261757778997 | 0.0529461687031 | 0.0981438986084454 |
| Smad3 | ENSMUSG00000032402 | 17127 | -0.292823298240251 | 5.35590832763574 | -2.12250547248795 | 0.052957061681 | 0.0981474468018427 |
| Top2b | ENSMUSG00000017485 | 21974 | 0.131686106285671 | 5.95354410598122 | 2.12218331291163 | 0.052988376687 | 0.0981888363403842 |
| Mafb | ENSMUSG00000074622 | 16658 | 0.269623347452403 | 4.61012174520089 | 2.12164755734952 | 0.053040492097 | 0.098268749002127 |
| Grb2 | ENSMUSG00000059923 | 14784 | 0.133426603660467 | 5.04080114834113 | 2.12106353793675 | 0.053097356667 | 0.0983574319154844 |
| Mcf2l | ENSMUSG00000031442 | 17207 | -0.536453320349046 | 4.33716236674206 | -2.12081283445463 | 0.053121784456 | 0.098386009132075 |
| Gorasp2 | ENSMUSG00000014959 | 70231 | 0.159286848901785 | 5.33353866710448 | 2.12059885120039 | 0.053142642606 | 0.0983989415979651 |
| Tanc1 | ENSMUSG00000035168 | 66860 | -0.156808420052347 | 4.32748181014567 | -2.1205565098155 | 0.053146770761 | 0.0983989415979651 |
| Iqce | ENSMUSG00000036555 | 74239 | 0.442804477630502 | 1.64810510863495 | 2.11983381246487 | 0.053217277545 | 0.0985127960795091 |
| Hdac2 | ENSMUSG00000019777 | 15182 | -0.147305392858284 | 4.91392944806957 | -2.11956829051146 | 0.053243203870 | 0.0985324552547343 |
| Tpd52l2 | ENSMUSG00000000827 | 66314 | 0.18491261697036 | 4.00864064453991 | 2.11954042273273 | 0.053245925641 | 0.0985324552547343 |
| Pold2 | ENSMUSG00000020471 | 18972 | -0.263880950651625 | 3.26729302709176 | -2.11940376030537 | 0.053259274966 | 0.0985404764145096 |
| Ccnd1 | ENSMUSG00000070348 | 12443 | 0.548784243607642 | 3.95202927632836 | 2.11914638476913 | 0.053284424120 | 0.0985703232469375 |
| Bub3 | ENSMUSG00000066979 | 12237 | -0.174890932422925 | 3.77450769511212 | -2.11846728698848 | 0.053350834402 | 0.098676475609713 |
| Snrnp70 | ENSMUSG00000063511 | 20637 | 0.189722006421464 | 4.94385697744871 | 2.11798203616543 | 0.053398335155 | 0.0987382508541438 |
| Psmd12 | ENSMUSG00000020720 | 66997 | -0.155106961990011 | 5.78997343697002 | -2.11794155297878 | 0.053402299794 | 0.0987382508541438 |
| Pomgnt2 | ENSMUSG00000066235 | 215494 | -0.274959610326069 | 1.91024570149698 | -2.1171417506625 | 0.053480682945 | 0.0988664544698664 |
| Surf6 | ENSMUSG00000036160 | 20935 | 0.248200470434562 | 2.48004229228261 | 2.1167409559939 | 0.053520002309 | 0.0989224121152457 |
| Hsd3b2 | ENSMUSG00000063730 | 15493 | -0.651497322990217 | 4.1563297485632 | -2.11587072504242 | 0.053605467569 | 0.0990636290793319 |

| | | | | | | | |
|---|---|---|---|---|---|---|---|
| Tomm40 | ENSMUSG00000002984 | 53333 | -0.189029125992421 | 4.43067015100234 | -2.11485662901941 | 0.0537052218644 | 0.0992219781125256 |
| Lnx2 | ENSMUSG00000016520 | 140887 | 0.267440650941589 | 2.80471737621042 | 2.11481512812506 | 0.0537093078850 | 0.0992219781125256 |
| Hnrnpf | ENSMUSG00000042079 | 98758 | 0.173346216200857 | 6.16030318861897 | 2.11409412720508 | 0.0537803410358 | 0.0993364156578761 |
| Glb1l2 | ENSMUSG00000036395 | 244757 | -0.659185418141301 | 2.257875076214 | -2.11379001211559 | 0.0538103286887 | 0.099375013160528 |
| 4833420G17Rik | ENSMUSG00000062822 | 67392 | 0.311411523022383 | 2.77999416590506 | 2.11298317454807 | 0.0538899632539 | 0.0995052682988473 |
| Astn2 | ENSMUSG00000028373 | 56079 | 0.314600855408889 | 2.37007403874911 | 2.11233164910873 | 0.0539543484872 | 0.0996073270249299 |
| Zdhhc3 | ENSMUSG00000025786 | 69035 | 0.142476213408377 | 5.38416887656887 | 2.11193785276795 | 0.0539932989961 | 0.0996624031285287 |
| Stk38 | ENSMUSG00000024006 | 106504 | 0.18535250144749 | 4.16236954424899 | 2.11148439026944 | 0.0540381834693 | 0.0997284118696545 |
| Fem1b | ENSMUSG00000032244 | 14155 | 0.190942890200763 | 4.09619585065395 | 2.1102227256641 | 0.0541632476236 | 0.0999423463494419 |
| Slc12a7 | ENSMUSG00000017756 | 20499 | 0.210668248141253 | 3.82804214902434 | 2.11008356049612 | 0.0541770590083 | 0.0999459255103627 |
| Atp6v0a4 | ENSMUSG00000038600 | 140494 | 0.599157897148402 | 0.747784288446138 | 2.10993686723257 | 0.0541916210578 | 0.0999459255103627 |
| Ggta1 | ENSMUSG00000035778 | 14594 | -0.280208183064266 | 3.02930973010659 | -2.10989692189341 | 0.0541955870096 | 0.0999459255103627 |
| BC005624 | ENSMUSG00000026851 | 227707 | -0.158199697268535 | 3.72887760650732 | -2.10983474526594 | 0.0542017607195 | 0.0999459255103627 |
| Epg5 | ENSMUSG00000039840 | 100502841 | -0.177979414838083 | 4.4019939027489 | -2.10839632809668 | 0.0543447681384 | 0.10019272398747 |
| Rnf44 | ENSMUSG00000034928 | 105239 | 0.146544148115991 | 4.98526883307525 | 2.10825720469932 | 0.0543586183587 | 0.100201358731861 |
| Nip7 | ENSMUSG00000031917 | 66164 | 0.268055751572829 | 2.56274125192904 | 2.10803962584578 | 0.0543802856677 | 0.100224397679007 |
| Wdr77 | ENSMUSG00000000561 | 70465 | 0.206442500439667 | 3.60962382474223 | 2.10789046297619 | 0.0543951444941 | 0.100234882748673 |
| Prkra | ENSMUSG00000002731 | 23992 | 0.45062065219604 | 1.20761365113969 | 2.10770129217521 | 0.0544139941257 | 0.100238553757171 |
| Pln | ENSMUSG00000038583 | 18821 | -0.637748164301804 | 1.1181640039989 | -2.10768641395708 | 0.0544154769001 | 0.100238553757171 |
| Ankib1 | ENSMUSG00000040351 | 70797 | 0.141595795897274 | 4.16228505806086 | 2.10732717301475 | 0.0544512905159 | 0.100287625379911 |
| Olfr45 | ENSMUSG00000066122 | 18344 | -0.704535453915732 | 2.21954738822513 | -2.1066342367378 | 0.0545204328908 | 0.100390382022451 |
| Snip1 | ENSMUSG00000050213 | 76793 | 0.255828644126811 | 2.25999980481475 | 2.10658398388734 | 0.0545254503766 | 0.100390382022451 |
| Egr1 | ENSMUSG00000038418 | 13653 | 0.874905946515285 | 4.14730085785593 | 2.10530317575665 | 0.0546534772472 | 0.100609154562099 |
| Acaa1a | ENSMUSG00000036138 | 113868 | -0.171151225748898 | 4.03916839218327 | -2.10513425581488 | 0.0546703829476 | 0.100623329853599 |
| Tns2 | ENSMUSG00000037003 | 209039 | -0.302338720831286 | 5.55296335644784 | -2.10470825637025 | 0.0547130390245 | 0.100684887134298 |
| Dusp1 | ENSMUSG00000024190 | 19252 | -0.678135424668472 | 5.47976320265383 | -2.10456230969446 | 0.0547276602290 | 0.100694841235075 |
| Tbcd | ENSMUSG00000039230 | 108903 | 0.209509577115507 | 3.17651968111042 | 2.10394556600871 | 0.0547894857588 | 0.100791630567093 |
| Tubd1 | ENSMUSG00000020513 | 56427 | -0.333033205959776 | 1.34811208616506 | -2.1033069217278 | 0.0548535752606 | 0.100892551097742 |
| 2310011J03Rik | ENSMUSG00000020133 | 66374 | 0.326543741758046 | 1.98287514764076 | 2.10304300175219 | 0.0548800805886 | 0.100924320476822 |
| Mrpl49 | ENSMUSG00000007338 | 18120 | -0.183065891047282 | 3.19400078024022 | -2.10235652393704 | 0.0549490788465 | 0.10103421040726 |
| Phf2 | ENSMUSG00000038025 | 18676 | -0.150870167435049 | 5.58062229470246 | -2.10212684636858 | 0.0549721818364 | 0.101059690490141 |
| Mllt1 | ENSMUSG00000024212 | 64144 | -0.181801270082214 | 4.31190831966495 | -2.10115078230728 | 0.0550704635594 | 0.101221148948589 |
| Ccdc174 | ENSMUSG00000034083 | 232236 | -0.19833527351881 | 2.97786367244029 | -2.10107075984973 | 0.0550785284005 | 0.101221148948589 |
| Dgcr8 | ENSMUSG00000022718 | 94223 | 0.228624602385648 | 3.4554940881545 | 2.10052539978082 | 0.0551335201989 | 0.101289044483743 |
| Poglut2 | ENSMUSG00000026047 | 72050 | 0.400063486604177 | 1.76873333430098 | 2.10052058873575 | 0.0551340055510 | 0.101289044483743 |
| Htra1 | ENSMUSG00000006205 | 56213 | 0.275207804161031 | 3.4734638498286 | 2.09864894858048 | 0.0553231231198 | 0.101619401583464 |
| Pcsk6 | ENSMUSG00000030513 | 18553 | 0.208584904942356 | 4.50300214724058 | 2.09798185716745 | 0.0553906739338 | 0.101726387226273 |
| Txnl4a | ENSMUSG00000057130 | 27366 | 0.177805219580419 | 3.27840430333692 | 2.09696358547126 | 0.0554939334418 | 0.101890947106012 |
| Atm | ENSMUSG00000034218 | 11920 | 0.179424005542467 | 3.93505618915886 | 2.09691445199534 | 0.0554989204161 | 0.101890947106012 |
| Ift52 | ENSMUSG00000017858 | 245866 | -0.242270946081879 | 2.61338417035499 | -2.09669331779867 | 0.0555213703535 | 0.101915046067893 |
| P3h4 | ENSMUSG00000006931 | 66180 | 0.273693400552077 | 1.99667838717123 | 2.09618540852163 | 0.0555729661090 | 0.10199262802848 |
| Rbmxl1 | ENSMUSG00000037070 | 19656 | 0.164743177528215 | 3.5218435014444 | 2.09571914162461 | 0.0556203707960 | 0.102062493397996 |
| Nr3c1 | ENSMUSG00000024431 | 14815 | -0.153976619800696 | 6.37482414963195 | -2.09555703108804 | 0.0556368611279 | 0.102075617487434 |
| Nxf1 | ENSMUSG00000010097 | 53319 | 0.165137590586552 | 4.14477145218841 | 2.0951562011198 | 0.0556776541320 | 0.102133317220509 |
| Gm14432 | ENSMUSG00000078878 | 668030 | -0.367700822213396 | 1.47706107213098 | -2.09368741467869 | 0.0558273717015 | 0.102383938488178 |
| Ube2e2 | ENSMUSG00000058317 | 218793 | 0.388129368983174 | 1.33309418618718 | 2.09362961372205 | 0.0558332711460 | 0.102383938488178 |
| Med29 | ENSMUSG00000003444 | 67224 | -0.21243194585401 | 3.13084720670931 | -2.09354039208109 | 0.0558423786722 | 0.102383938488178 |
| Rapsn | ENSMUSG00000002104 | 19400 | -0.190701880260007 | 3.35541111820146 | -2.09282494599942 | 0.0559154594870 | 0.102500735813033 |
| Armcx1 | ENSMUSG00000033460 | 78248 | 0.410968426777107 | 1.29646068527458 | 2.09217920409257 | 0.0559814963559 | 0.102604583612773 |
| Rbm34 | ENSMUSG00000033931 | 52202 | 0.226555820352228 | 2.59145808539081 | 2.09175289696577 | 0.0560251323256 | 0.102667346429414 |
| Hic1 | ENSMUSG00000043099 | 15248 | -0.401997220330872 | 3.23580922016201 | -2.09085200402098 | 0.0561174497401 | 0.102808736897972 |
| Golga1 | ENSMUSG00000026754 | 76899 | -0.192433000954065 | 4.13003057254056 | -2.09081642051384 | 0.0561210989909 | 0.102808736897972 |
| Ube2k | ENSMUSG00000029203 | 53323 | -0.142410854064501 | 4.57025729488033 | -2.09046348192432 | 0.0561573063671 | 0.102857827731069 |
| Pdlim1 | ENSMUSG00000055044 | 54132 | -0.20174329555346 | 7.14390937041806 | -2.09026501613773 | 0.0561776761501 | 0.102877898810067 |
| Ptprk | ENSMUSG00000019889 | 19272 | -0.215114992745782 | 3.18230735731112 | -2.08926675584665 | 0.0562802377514 | 0.103048455420594 |
| Mark4 | ENSMUSG00000030397 | 232944 | -0.221294040516026 | 4.00394646595374 | -2.08905407190559 | 0.0563021113893 | 0.103071240930728 |
| Rcc1l | ENSMUSG00000061979 | 94254 | -0.249611430709633 | 2.95430925342643 | -2.08846428306745 | 0.0563628098927 | 0.103165082876437 |
| Mrpl43 | ENSMUSG00000025208 | 94067 | -0.174989956588483 | 4.48423137325124 | -2.08834586755469 | 0.0563750040117 | 0.103170127047208 |
| Vamp4 | ENSMUSG00000026696 | 53330 | -0.215429578530206 | 3.13682293684991 | -2.08748424378073 | 0.0564638054221 | 0.103315342663143 |
| Acbd6 | ENSMUSG00000033701 | 72482 | -0.14495713651362 | 3.74725259792503 | -2.08738531134212 | 0.0564740099753 | 0.103316720174189 |
| Abcd4 | ENSMUSG00000021240 | 19300 | 0.255401687888553 | 2.26415633681714 | 2.08727569287561 | 0.0564853187530 | 0.103320117016297 |
| Drg1 | ENSMUSG00000020457 | 13494 | -0.164392836678136 | 4.52223996379804 | -2.08416568056269 | 0.0568070385308 | 0.10389120598639 |
| Crcp | ENSMUSG00000025532 | 12909 | 0.237849602324001 | 2.69629867218037 | 2.08383201208462 | 0.0568416560161 | 0.103937126448981 |
| Stox2 | ENSMUSG00000038143 | 71069 | 0.182911166925533 | 4.78042509411655 | 2.08373652430321 | 0.0568515662922 | 0.103937861037054 |
| Kcnj11 | ENSMUSG00000096146 | 16514 | -0.142446969137795 | 5.53365408367127 | -2.08351696536345 | 0.0568743594652 | 0.10396214436691 |
| Nr2f6 | ENSMUSG00000002393 | 13864 | 0.189094689668168 | 3.21207285344931 | 2.08327615556772 | 0.0568993684971 | 0.103990469326551 |
| Jmjd1c | ENSMUSG00000037876 | 108829 | 0.152968666938707 | 5.85240715536271 | 2.08230658189468 | 0.0570001656006 | 0.104147582392848 |
| Atox1 | ENSMUSG00000018585 | 11927 | 0.249514103942014 | 2.7542067684265 | 2.08226598325211 | 0.0570043898505 | 0.104147582392848 |
| Nudt6 | ENSMUSG00000050174 | 229228 | -0.243209599692284 | 2.52314419089162 | -2.08183352966557 | 0.0570494042308 | 0.104210505507698 |
| Usp47 | ENSMUSG00000059263 | 74996 | -0.165499893060092 | 8.55337026692696 | -2.08175197204224 | 0.0570578973070 | 0.104210505507698 |
| Zfp966 | ENSMUSG00000089756 | 667962 | -0.224354107579547 | 3.18984527706645 | -2.0811953565295 | 0.0571158922387 | 0.104299000678497 |
| Nfe2l2 | ENSMUSG00000015839 | 18024 | -0.248356018983158 | 5.08554093318508 | -2.08085008543249 | 0.0571518941854 | 0.104347311732251 |
| Rap1b | ENSMUSG00000052681 | 215449 | 0.240189548226075 | 4.81481505384318 | 2.08035793875136 | 0.0572032474034 | 0.104423630154885 |
| Tsnax | ENSMUSG00000056820 | 53424 | -0.164357975773782 | 6.02906096812543 | -2.08002571141949 | 0.0572379379366 | 0.104469510700607 |
| Bbs2 | ENSMUSG00000031755 | 67378 | -0.432132330638726 | 1.23186054232004 | -2.0799070538416 | 0.0572503326515 | 0.104471839934017 |
| Sssca1 | ENSMUSG00000079478 | 56390 | -0.244725258070614 | 2.41966278721764 | -2.07983051660136 | 0.0572583288852 | 0.104471839934017 |
| Ccdc88c | ENSMUSG00000021182 | 68339 | 0.14857583355674 | 4.10267384077284 | 2.07938023080853 | 0.0573053934699 | 0.104538860616117 |
| Fli1 | ENSMUSG00000016087 | 14247 | -0.23665359848518 | 3.41342231789816 | -2.07929612566757 | 0.0573141882419 | 0.104538860616117 |
| Scn1b | ENSMUSG00000019194 | 20266 | -0.152967867958512 | 6.86333270977059 | -2.07837256775894 | 0.0574108458109 | 0.104681629468811 |
| Dnajc5 | ENSMUSG00000000826 | 13002 | 0.15595726723872 | 5.25919386250765 | 2.07836521951974 | 0.0574116154666 | 0.104681629468811 |
| Ggcx | ENSMUSG00000053460 | 56316 | 0.306815007317937 | 2.89772905007076 | 2.07507555047115 | 0.0577571368252 | 0.105294073154835 |
| Trim44 | ENSMUSG00000027189 | 80985 | 0.163350689770872 | 4.9700632172131 | 2.07368905177288 | 0.0579033396355 | 0.105543005762168 |
| Lcorl | ENSMUSG00000015882 | 209707 | 0.214401847601037 | 2.60967085600783 | 2.07346273023737 | 0.0579272371630 | 0.105568961225246 |
| Olfr161 | ENSMUSG00000051003 | 258859 | -0.648222354102592 | 0.678656846906528 | -2.07301197311917 | 0.0579748602702 | 0.105631531269727 |
| Vmn2r20 | ENSMUSG00000094145 | 667180 | -0.986774205905675 | 0.128129393525389 | -2.07295485720904 | 0.0579808972297 | 0.105631531269727 |
| Mtus1 | ENSMUSG00000045636 | 102103 | -0.191805171201613 | 6.91403037473361 | -2.07242680429149 | 0.0580367382005 | 0.105715644937521 |
| Gba | ENSMUSG00000028048 | 14466 | 0.269627427909377 | 3.05808619810746 | 2.0721983272293 | 0.0580609147986 | 0.105742062589762 |
| Ube3b | ENSMUSG00000029577 | 117146 | -0.163552879843122 | 7.13422051445775 | -2.0720398464987 | 0.0580776901166 | 0.105754994280418 |
| Cwc27 | ENSMUSG00000021715 | 67285 | -0.212082028880853 | 3.72319644322491 | -2.07158608681011 | 0.0581257457629 | 0.10582487124168 |
| Galnt1 | ENSMUSG00000000420 | 14423 | -0.138856908157147 | 5.47671758812248 | -2.07077169636936 | 0.0582120864592 | 0.105964415834509 |
| Dus2 | ENSMUSG00000031901 | 66369 | -0.241467702516997 | 1.75696756175699 | -2.07048263074044 | 0.0582427613689 | 0.106002601485894 |
| Akap8l | ENSMUSG00000002625 | 54194 | -0.196497712960609 | 4.21300732104139 | -2.07008199128103 | 0.0582853009335 | 0.10606236465871 |
| Sertad1 | ENSMUSG00000008384 | 55942 | 0.328095646979811 | 1.9901183178737 | 2.06977554095547 | 0.0583178589713 | 0.106103947472713 |
| Lrrfip2 | ENSMUSG00000032497 | 71268 | 0.118125945951246 | 6.26213060650437 | 2.06930831885814 | 0.0583675301746 | 0.106143973503111 |
| Wdr35 | ENSMUSG00000066643 | 74682 | 0.199891165213376 | 3.25597217589513 | 2.06930247635015 | 0.0583681515496 | 0.106143973503111 |
| Proser3 | ENSMUSG00000036864 | 333193 | -0.249087779898451 | 1.78471999915964 | -2.06929459714002 | 0.0583689895459 | 0.106143973503111 |
| Snhg18 | ENSMUSG00000096956 | 100616095 | 0.474095937420468 | 1.66113794158262 | 2.06801134377071 | 0.0585056190209 | 0.10635760508178 |
| Olfr1152 | ENSMUSG00000045225 | 258103 | -0.811128835803329 | 1.16267275787294 | -2.06800846174756 | 0.0585059262059 | 0.10635760508178 |
| Atp2a3 | ENSMUSG00000020788 | 53313 | -0.236449674877032 | 3.19371547613571 | -2.06733564760536 | 0.0585776799878 | 0.106470339199818 |
| Rbm39 | ENSMUSG00000027620 | 170791 | -0.119090956394904 | 6.14811422673522 | -2.06713725500452 | 0.0585988535447 | 0.10649111689072 |
| Dnaja4 | ENSMUSG00000032285 | 58233 | -0.164425258871703 | 7.50966837677004 | -2.06703328997758 | 0.0586099520939 | 0.106493581505739 |
| Prex2 | ENSMUSG00000048960 | 109294 | -0.248772176538003 | 4.60028741702858 | -2.06679351015099 | 0.0586355566534 | 0.106522398168247 |
| Ptdss1 | ENSMUSG00000021518 | 19210 | 0.132183103780771 | 4.71316985606585 | 2.06633499460686 | 0.0586845474072 | 0.106593683567487 |
| Bcorl1 | ENSMUSG00000036959 | 320376 | 0.236071620036893 | 3.10791759634048 | 2.06592488014543 | 0.0587283987395 | 0.106655611666686 |
| Hdgfl2 | ENSMUSG00000002833 | 15193 | 0.141108607169887 | 4.66624571890919 | 2.06577853673597 | 0.0587440537854 | 0.106666320918329 |
| Ccdc3 | ENSMUSG00000026676 | 74186 | 0.35302583147591 | 2.60372408435419 | 2.06556994542249 | 0.0587663744526 | 0.106689127909361 |
| Mtpap | ENSMUSG00000024234 | 67440 | -0.194564330567359 | 3.62856799417723 | -2.06542193337906 | 0.0587822174879 | 0.106700169272886 |
| Hnrnpc | ENSMUSG00000060373 | 15381 | -0.129749024270907 | 5.49944549226813 | -2.06524121429038 | 0.0588015668010 | 0.106717570430344 |
| Tmem131 | ENSMUSG00000026116 | 56030 | 0.148219677830983 | 6.15043038013825 | 2.06500619350047 | 0.0588267389211 | 0.106745531730879 |
| Strip1 | ENSMUSG00000014601 | 229707 | 0.138968836879852 | 3.99680168704525 | 2.06480522815439 | 0.0588482714020 | 0.106766880447488 |
| Ube2q2 | ENSMUSG00000032307 | 109161 | -0.142523518598578 | 4.6516894622233 | -2.06458562462286 | 0.0588718092068 | 0.106791859681325 |
| Cyb5rl | ENSMUSG00000028621 | 230582 | -0.366384240844663 | 1.53948281668888 | -2.06440208775754 | 0.0588914879476 | 0.106792531136702 |
| Abcf2 | ENSMUSG00000028953 | 27407 | -0.160400312862011 | 5.95474195968233 | -2.06439993544879 | 0.0588917187532 | 0.106792531136702 |
| Sim1 | ENSMUSG00000019913 | 20464 | -0.35112579906084 | 3.0696830597241 | -2.06424844330239 | 0.0589079663131 | 0.106804275960982 |
| Mypop | ENSMUSG00000048481 | 232934 | 0.323335717401708 | 1.39348175800264 | 2.06385866424775 | 0.0589497892341 | 0.106862379124089 |
| St6galnac6 | ENSMUSG00000026811 | 50935 | -0.197712467890959 | 3.49461277568612 | -2.06228809708148 | 0.0591185878423 | 0.107150602504493 |

| | | | | | | | |
|---|---|---|---|---|---|---|---|
| Sncaip | ENSMUSG00000024534 | 67847 | -0.369184207477422 | 2.04355247834589 | -2.0621372168859 | 0.0591348273707 | 0.107162267571292 |
| Pear1 | ENSMUSG00000028073 | 73182 | -0.415052163529999 | 2.87145892004413 | -2.0617524720787 | 0.0591762568632 | 0.107219569662143 |
| Trim7 | ENSMUSG00000040350 | 94089 | 0.29170430351264 | 5.02236282221164 | 2.06038426541485 | 0.0593238033537 | 0.107452788256927 |
| Asb6 | ENSMUSG00000039483 | 72323 | 0.275819254301189 | 1.95983350525733 | 2.06037656925431 | 0.0593246342631 | 0.107452788256927 |
| Cltc | ENSMUSG00000047126 | 67300 | 0.148520424013335 | 7.33993050782273 | 2.0594627187077 | 0.0594233738348 | 0.10760171653575 |
| Mindy2 | ENSMUSG00000042444 | 235461 | -0.151694961518244 | 4.95139441319183 | -2.05943339289026 | 0.0594265449388 | 0.10760171653575 |
| Tmem123 | ENSMUSG00000050912 | 71929 | 0.14812873338673 | 4.45827255844728 | 2.05898656743035 | 0.0594748810937 | 0.107671401993282 |
| Taf1 | ENSMUSG00000031314 | 270627 | 0.165668004350406 | 4.58082666405349 | 2.05859238057695 | 0.0595175531240 | 0.107725834272549 |
| Prr14 | ENSMUSG00000030822 | 233895 | -0.166911817123633 | 3.82521791510039 | -2.0585216858886 | 0.0595252090482 | 0.107725834272549 |
| Ptp4a2 | ENSMUSG00000028788 | 19244 | -0.182824494790835 | 7.97002961983075 | -2.05843578190952 | 0.0595345132961 | 0.107725834272549 |
| Mob4 | ENSMUSG00000025979 | 19070 | -0.175269220883083 | 4.17020294388497 | -2.05829512796979 | 0.0595497504054 | 0.10773557128135 |
| Nr4a2 | ENSMUSG00000026826 | 18227 | 0.815352788079135 | 0.555030214099506 | 2.05796404701066 | 0.0595856307866 | 0.107782646058011 |
| Ubr1 | ENSMUSG00000027272 | 22222 | -0.183401050424473 | 5.21773878227901 | -2.05760154975895 | 0.0596249387609 | 0.107835904301198 |
| Rpusd1 | ENSMUSG00000041199 | 106707 | 0.484749780603548 | 0.957846594720294 | 2.05712769740922 | 0.0596763578243 | 0.107911045058301 |
| Ndor1 | ENSMUSG00000006471 | 78797 | 0.363558722377872 | 2.06380064880223 | 2.05691642250746 | 0.0596992970625 | 0.107934670226749 |
| ND4L | ENSMUSG00000065947 | 17720 | -0.32354917400589 | 9.87599422462148 | -2.05570866355508 | 0.0598305862854 | 0.108144193814859 |
| Vmn2r116 | ENSMUSG00000090966 | 619697 | -1.52662124474834 | -0.880274681651376 | -2.05566835622408 | 0.0598349724812 | 0.108144193814859 |
| Cdipt | ENSMUSG00000030682 | 52858 | 0.195428389309327 | 3.45564925149412 | 2.05544495268573 | 0.0598592883727 | 0.108161201991923 |
| Mettl16 | ENSMUSG00000010554 | 67493 | -0.190173529814567 | 3.90358368721755 | -2.05540008755366 | 0.0598641727244 | 0.108161201991923 |
| Suox | ENSMUSG00000049858 | 211389 | 0.183464563436853 | 3.04681609802605 | 2.05472526304507 | 0.0599376835499 | 0.108276122770519 |
| Snx33 | ENSMUSG00000032733 | 235406 | 0.305663060797853 | 2.46526171012883 | 2.05459562851853 | 0.0599518146006 | 0.108283755023027 |
| Arcn1 | ENSMUSG00000032096 | 213827 | 0.120036307750217 | 6.28982287637079 | 2.05417200608166 | 0.0599980137999 | 0.108349296026335 |
| Gm14496 | ENSMUSG00000098505 | 672125 | -0.735471460115007 | 1.31014752305564 | -2.0535410047075 | 0.0600668901763 | 0.108455760904808 |
| Gpi1 | ENSMUSG00000036427 | 14751 | -0.164294866555515 | 7.92139090122905 | -2.0528396951131 | 0.0601435266324 | 0.108576199771944 |
| Ncdn | ENSMUSG00000028833 | 26562 | 0.227875709087103 | 3.12498182907379 | 2.05199452793822 | 0.0602360032358 | 0.108725190120714 |
| Csgalnact2 | ENSMUSG00000042042 | 78752 | 0.128045674616802 | 4.78516371063218 | 2.05111599854838 | 0.0603322693870 | 0.108880970227757 |
| Mettl17 | ENSMUSG00000004561 | 52535 | -0.347572054006236 | 1.6704947631869 | -2.05042002996702 | 0.0604086319532 | 0.108996479787993 |
| Mitf | ENSMUSG00000035158 | 17342 | -0.148349511206818 | 4.53057015026135 | -2.05035094166824 | 0.0604162172752 | 0.108996479787993 |
| Ltbp4 | ENSMUSG00000040488 | 108075 | 0.23019086303388 | 7.1368845037362 | 2.04984273709759 | 0.0604720409414 | 0.109079188041343 |
| Praf2 | ENSMUSG00000031149 | 54637 | 0.456661241496988 | 1.57893109158903 | 2.0480756135755 | 0.0606665211274 | 0.10941193572738 |
| Vti1b | ENSMUSG00000021124 | 53612 | -0.149535911058747 | 5.67645783905418 | -2.04782723803939 | 0.0606939022257 | 0.109423890411954 |
| Rab30 | ENSMUSG00000030643 | 75985 | 0.375145302950895 | 2.57382162008772 | 2.0478268416484 | 0.0606939459332 | 0.109423890411954 |
| Bloc1s1 | ENSMUSG00000090247 | 14533 | -0.19140867477055 | 4.40332228169663 | -2.04772189063135 | 0.0607055192434 | 0.109423890411954 |
| Mppe1 | ENSMUSG00000062526 | 225651 | 0.244444433649726 | 1.93015674184564 | 2.04765232675047 | 0.0607131914141 | 0.109423890411954 |
| Tulp4 | ENSMUSG00000034377 | 68842 | 0.154388360950509 | 7.83861605524198 | 2.04755696161835 | 0.0607237106478 | 0.109424807301627 |
| Ddx23 | ENSMUSG00000003360 | 74351 | -0.152152161066778 | 5.81142016110922 | -2.04676826231787 | 0.0608107724630 | 0.109563631744441 |
| 3425401B19Rik | ENSMUSG00000071540 | 100504518 | -0.164709876567843 | 9.65437378742746 | -2.04664520937282 | 0.0608243662367 | 0.109570063832171 |
| Il16 | ENSMUSG00000001741 | 16170 | 0.284843738242186 | 2.29451444387423 | 2.04610562973449 | 0.0608840072063 | 0.109659430346431 |
| Slc35f5 | ENSMUSG00000026342 | 74150 | 0.187296221288775 | 3.98870646934299 | 2.04555996686295 | 0.0609443754281 | 0.109750077068313 |
| Mcat | ENSMUSG00000048755 | 223722 | -0.224492250826344 | 2.63335172382311 | -2.04529493585452 | 0.0609737164727 | 0.109784828654795 |
| P2rx6 | ENSMUSG00000022758 | 18440 | 0.331589348568059 | 1.40529598153801 | 2.04483065878663 | 0.0610251470504 | 0.109859335047426 |
| Ier2 | ENSMUSG00000053560 | 15936 | 0.384265410031432 | 2.68363409337874 | 2.0442889231203 | 0.0610852087325 | 0.109949352322673 |
| Trmt1 | ENSMUSG00000001909 | 212528 | -0.19110428127417 | 3.72402754512588 | -2.04378516518389 | 0.0611411087779 | 0.110031850518823 |
| Tagln2 | ENSMUSG00000026547 | 21346 | 0.34247064979211 | 4.5024292783784 | 2.04355290917803 | 0.0611668972034 | 0.110046262012221 |
| Saal1 | ENSMUSG00000006763 | 78935 | 0.346712099880324 | 1.36539263145666 | 2.04353171051661 | 0.0611692514853 | 0.110046262012221 |
| Vmn2r60 | ENSMUSG00000090619 | 637898 | -0.264299774689591 | 2.57770215615759 | -2.04335767405579 | 0.0611885827939 | 0.110062925542281 |
| P4ha2 | ENSMUSG00000018906 | 18452 | -0.197525467058546 | 3.63390500598855 | -2.04323003533604 | 0.0612027639987 | 0.110070321367238 |
| Cyp51 | ENSMUSG00000001467 | 13121 | 0.383575872001231 | 1.76666552665231 | 2.04276741471165 | 0.0612541885221 | 0.110144684115046 |
| Kdm1a | ENSMUSG00000036940 | 99982 | 0.136264729187398 | 4.45421617519906 | 2.04185981353996 | 0.0613551924902 | 0.11030815939328 |
| 2410131K14Rik | ENSMUSG00000032840 | 76792 | -0.214464636591679 | 3.22870440423617 | -2.04108742224495 | 0.0614412703200 | 0.110444750183841 |
| Mbtps1 | ENSMUSG00000031835 | 56453 | 0.162247801076403 | 5.16885654001616 | 2.04057499225926 | 0.0614984386245 | 0.110529337817316 |
| Rnf14 | ENSMUSG00000060450 | 56736 | -0.16917359378521 | 5.59879922507736 | -2.04036204975376 | 0.0615222095814 | 0.110553883435001 |
| Zfp68 | ENSMUSG00000058291 | 24135 | -0.206965698671207 | 4.25157994585056 | -2.0389684109762 | 0.0616779919180 | 0.110815603165077 |
| Mrto4 | ENSMUSG00000028741 | 69902 | -0.214311130332848 | 3.01442131022083 | -2.03850236359858 | 0.0617301682305 | 0.110891120612751 |
| Ano1 | ENSMUSG00000031075 | 101772 | -0.353323492745253 | 2.58139719852294 | -2.03837849693169 | 0.0617440425588 | 0.11089781945633 |
| Rgp1 | ENSMUSG00000028468 | 242406 | -0.156673150698913 | 4.69216403980642 | -2.03812207647075 | 0.0617727733956 | 0.110931195332406 |
| Hpcal1 | ENSMUSG00000071379 | 53602 | 0.345365493583613 | 1.64372622204128 | 2.03788039429722 | 0.0617998641480 | 0.110961615473423 |
| Mrtfa | ENSMUSG00000042292 | 223701 | 0.375071007486742 | 3.61542954283591 | 2.0376778244329 | 0.0618225791392 | 0.110984170236601 |
| Pla2r1 | ENSMUSG00000054580 | 18779 | 0.280740269273966 | 1.57563796542068 | 2.03748105085374 | 0.0618446515360 | 0.111001601998918 |
| Tmem47 | ENSMUSG00000025666 | 192216 | -0.207521224324682 | 2.87115764990412 | -2.03738432333976 | 0.0618555042731 | 0.111001601998918 |
| Trim30d | ENSMUSG00000057596 | 209387 | 0.513937119407736 | 0.845898719562374 | 2.03731972094247 | 0.0618627535794 | 0.111001601998918 |
| Pld2 | ENSMUSG00000020828 | 18806 | 0.28636182723166 | 2.06459488186051 | 2.03718838890056 | 0.0618774933133 | 0.111009827573907 |
| Recql | ENSMUSG00000030243 | 19691 | -0.2325575267511 | 2.34999417205473 | -2.03708206679717 | 0.0618894284877 | 0.111013019822656 |
| Eif1b | ENSMUSG00000006941 | 68969 | -0.194437586552599 | 3.73317066798566 | -2.03672098534097 | 0.0619299774790 | 0.111067528108803 |
| Pikfyve | ENSMUSG00000025949 | 18711 | 0.195434186180501 | 4.00124909303015 | 2.03598904064976 | 0.0620122490615 | 0.111196833085903 |
| Ccng2 | ENSMUSG00000029385 | 12452 | 0.225169906286135 | 2.84728239422785 | 2.03542340329168 | 0.0620758964411 | 0.111292705264658 |
| Olfr1284 | ENSMUSG00000108931 | 258379 | -0.698606914878737 | 0.347226297983465 | -2.03461167944042 | 0.0621673394474 | 0.111438371187263 |
| Tmod1 | ENSMUSG00000028328 | 21916 | 0.14637905384872 | 6.39942467798148 | 2.03447014479559 | 0.0621832964283 | 0.111448698681534 |
| Pex7 | ENSMUSG00000020003 | 18634 | -0.205853761204701 | 3.26725804325824 | -2.03279380121154 | 0.0623725789276 | 0.111754273012612 |
| Coro1b | ENSMUSG00000024835 | 23789 | 0.243060186830844 | 3.84333425448639 | 2.03277910921209 | 0.0623742402021 | 0.111754273012612 |
| Irgq | ENSMUSG00000041037 | 210146 | -0.254627828290849 | 4.21069812249422 | -2.03206318286001 | 0.0624552418325 | 0.111881063335284 |
| Apln | ENSMUSG00000037010 | 30878 | -1.09486355966768 | 0.405102470666537 | -2.03167014929293 | 0.0624997517771 | 0.111942452347322 |
| Armcx4 | ENSMUSG00000049804 | 100503043 | 0.232581144089769 | 5.46608949681192 | 2.03140351055635 | 0.0625299644927 | 0.111978217868578 |
| Rrp12 | ENSMUSG00000035049 | 107094 | 0.182342031515108 | 5.05888332720584 | 2.03072666707678 | 0.0626067177359 | 0.112097302468751 |
| Ncapd3 | ENSMUSG00000035024 | 78658 | -0.215609444842391 | 2.99045526423083 | -2.02978833440801 | 0.0627132670453 | 0.112269689170029 |
| Eif1ad | ENSMUSG00000024841 | 69860 | -0.171099567248369 | 3.69761046776783 | -2.02935128489344 | 0.0627629517508 | 0.11234023671011 |
| Prkab2 | ENSMUSG00000038205 | 108097 | 0.237471412359382 | 5.13560400893836 | 2.02906645205925 | 0.0627953516478 | 0.112379827908085 |
| Zfp956 | ENSMUSG00000045466 | 101197 | -0.508259703489492 | 1.23602448123301 | -2.02844031438984 | 0.0628666293008 | 0.112488971171614 |
| Cpeb1 | ENSMUSG00000025586 | 12877 | 0.485594641499447 | 2.08153215039188 | 2.02804339370444 | 0.0629118521375 | 0.112551465747302 |
| Plin2 | ENSMUSG00000028494 | 11520 | -0.188041036410208 | 4.10545538594789 | -2.02667561465587 | 0.0630679184224 | 0.112812210158007 |
| Arhgef12 | ENSMUSG00000059495 | 69632 | 0.12949103307616 | 7.6301804807603 | 2.02656513110456 | 0.0630805403551 | 0.112816326345301 |
| Plxnd1 | ENSMUSG00000030123 | 67784 | 0.20720473256623 | 5.99071950977024 | 2.02577781912116 | 0.0631705522904 | 0.112958826613236 |
| Ifi208 | ENSMUSG00000066677 | 100033459 | -0.917138687786197 | 0.696222941332565 | -2.02557774904393 | 0.0631934447634 | 0.112981279801999 |
| Mcrs1 | ENSMUSG00000037570 | 51812 | -0.218587800099352 | 3.24557924169388 | -2.02356925217644 | 0.0634236849180 | 0.113374374462682 |
| Zfp113 | ENSMUSG00000037007 | 56314 | -0.280936463185163 | 2.58891972059095 | -2.02332907691145 | 0.0634512685508 | 0.11340513677721 |
| Nup153 | ENSMUSG00000021374 | 218210 | -0.155589472662576 | 5.20341567012929 | -2.02252697108716 | 0.0635434687450 | 0.113539057068939 |
| Mast2 | ENSMUSG00000003810 | 17776 | 0.136040323293019 | 7.14876823644161 | 2.02249651507914 | 0.0635469720197 | 0.113539057068939 |
| Zbtb38 | ENSMUSG00000040433 | 245007 | -0.153975081808123 | 5.80832090669268 | -2.02163740530885 | 0.0636458663853 | 0.11369716709566 |
| Spns1 | ENSMUSG00000030741 | 73658 | -0.244156271151475 | 2.83712387421344 | -2.02139559120794 | 0.0636737277576 | 0.113719397087641 |
| Vapb | ENSMUSG00000054455 | 56491 | -0.151839837405127 | 6.28315069294817 | -2.02134882688535 | 0.0636791171506 | 0.113719397087641 |
| Trir | ENSMUSG00000041203 | 68544 | 0.152958207236343 | 4.63702586978725 | 2.02004244973381 | 0.0638298413394 | 0.113969943757209 |
| Zbtb49 | ENSMUSG00000029127 | 75079 | 0.294600150744043 | 1.36525195080174 | 2.01957274268682 | 0.0638841142664 | 0.114048220324405 |
| Mif | ENSMUSG00000033307 | 17319 | -0.196974335906592 | 4.29433130184258 | -2.01909372503032 | 0.0639395067049 | 0.114128469593619 |
| Fyn | ENSMUSG00000019843 | 14360 | 0.254593989944909 | 3.56556506439162 | 2.0187268766366 | 0.0639819580223 | 0.114185597247728 |
| Wdr5 | ENSMUSG00000026917 | 140858 | -0.190598214304054 | 3.66087549929713 | -2.01758960674401 | 0.0641137264798 | 0.114402080338153 |
| A930005H10Rik | ENSMUSG00000054426 | 68161 | -0.464449377170451 | 1.36953010137153 | -2.01741131161556 | 0.0641344070480 | 0.11442030413615 |
| Osbpl9 | ENSMUSG00000028559 | 100273 | -0.156515004415139 | 5.37183722298152 | -2.0168342292183 | 0.0642013852677 | 0.114521106782237 |
| Scyl2 | ENSMUSG00000069539 | 213326 | 0.151311096731668 | 3.83846370663882 | 2.01662269592753 | 0.0642259526695 | 0.114546237335819 |
| Gpr137b | ENSMUSG00000021306 | 83924 | 0.354407944955484 | 1.79985943599332 | 2.01634958023739 | 0.0642576850120 | 0.114584136193553 |
| Herpud1 | ENSMUSG00000031770 | 64209 | -0.212724827812334 | 5.34538393746197 | -2.0159534001551 | 0.0643037413862 | 0.114647561098279 |
| Dcaf1 | ENSMUSG00000040325 | 321006 | -0.152172915862882 | 4.4960757461226 | -2.01555040381373 | 0.0643506212881 | 0.114694221669141 |
| Mta1 | ENSMUSG00000021144 | 116870 | -0.122665649062242 | 4.99649695105047 | -2.01554802947145 | 0.0643508975845 | 0.114694221669141 |
| Lrrc42 | ENSMUSG00000028617 | 77809 | 0.310924510425888 | 1.93868654285079 | 2.01470147844516 | 0.0644494781856 | 0.114837379661716 |
| Zc3hav1 | ENSMUSG00000029826 | 78781 | 0.246224674254574 | 3.98727408389658 | 2.01467786609005 | 0.0644522298256 | 0.114837379661716 |
| Map3k1 | ENSMUSG00000021754 | 26401 | -0.183109773065993 | 4.57922880221636 | -2.01429200772889 | 0.0644972107098 | 0.114898795676286 |
| Hacd1 | ENSMUSG00000063275 | 30963 | 0.19763123512299 | 4.38904684316176 | 2.01326861116672 | 0.0646166514604 | 0.115092816867305 |
| Hspbap1 | ENSMUSG00000022849 | 66667 | 0.321993347918647 | 1.54766597283741 | 2.01215778051365 | 0.0647465266763 | 0.115299564530054 |
| Pomk | ENSMUSG00000037251 | 74653 | 0.218250507377318 | 2.09810925605108 | 2.01209544519069 | 0.0647538218506 | 0.115299564530054 |
| Ccdc144b | ENSMUSG00000047696 | 241943 | -0.696238477014803 | 0.582470283563634 | -2.01176482968588 | 0.0647925267890 | 0.115349692236336 |
| Rhot1 | ENSMUSG00000017686 | 59040 | -0.130155167334971 | 4.82927266071326 | -2.01144338603142 | 0.0648301783629 | 0.115397928624876 |
| Tmem42 | ENSMUSG00000066233 | 66079 | -0.38195026173685 | 2.1857520461957 | -2.01107777760579 | 0.0648730275104 | 0.115455399497986 |
| Ptx4 | ENSMUSG00000044172 | 68509 | 0.530187225389466 | 2.41295839133936 | 2.00958689227587 | 0.0650480281145 | 0.115748005098538 |
| Qrich1 | ENSMUSG00000006673 | 69232 | -0.12115518803548 | 5.49911508957969 | -2.00934877382304 | 0.0650760186618 | 0.115778381344258 |
| Maf | ENSMUSG00000055435 | 17132 | 0.269366607916106 | 4.34847165431183 | 2.00926148237578 | 0.0650862824408 | 0.115778381344258 |
| Kdm4a | ENSMUSG00000033326 | 230674 | -0.132341961688165 | 4.93658867036307 | -2.00909334437335 | 0.0651060563947 | 0.115781014056835 |
| Ehbp1 | ENSMUSG00000042302 | 216565 | -0.170872803334916 | 3.88002810253169 | -2.00906877433682 | 0.0651089464282 | 0.115781014056835 |

| | | | | | | | |
|---|---|---|---|---|---|---|---|
| Mocos | ENSMUSG00000039616 | 68591 | 0.318674548474636 | 1.68105011039408 | 2.00824380697066 | 0.0652060510614 | 0.115934831514843 |
| Chchd5 | ENSMUSG00000037938 | 66170 | -0.377486153768846 | 1.318294960642 | -2.00767241697352 | 0.0652733857454 | 0.116035677278142 |
| Ehd3 | ENSMUSG00000024065 | 57440 | 0.280278760461279 | 1.81028407421945 | 2.00621017978835 | 0.0654459919959 | 0.116298541731196 |
| Peli1 | ENSMUSG00000020134 | 67245 | -0.15285636999394 | 3.74855993463264 | -2.00620149008879 | 0.0654470190020 | 0.116298541731196 |
| Flna | ENSMUSG00000031328 | 192176 | 0.257921857862777 | 6.95799761873896 | 2.00600154021778 | 0.0654706544856 | 0.116298541731196 |
| Adcy2 | ENSMUSG00000021536 | 210044 | -0.142379542737864 | 5.19479079838632 | -2.00594116839593 | 0.0654777924006 | 0.116298541731196 |
| Vmn2r53 | ENSMUSG00000096002 | 637908 | -0.798078123603053 | 4.61993239970305 | -2.00590716630648 | 0.0654818128691 | 0.116298541731196 |
| Slc8b1 | ENSMUSG00000032754 | 170756 | 0.328624318081026 | 2.04257247101368 | 2.00587944791827 | 0.0654850905091 | 0.116298541731196 |
| Upf1 | ENSMUSG00000058301 | 19704 | 0.110723938302923 | 5.34277346764795 | 2.00536708070841 | 0.0655457039652 | 0.116387279084553 |
| Rnf157 | ENSMUSG00000052949 | 217340 | -0.251477602793589 | 2.85648215256103 | -2.00522071271896 | 0.0655630288733 | 0.116399134093647 |
| BC003965 | ENSMUSG00000067722 | 214489 | -0.207565937065567 | 3.32531531109425 | -2.00462142397611 | 0.0656340077874 | 0.116506225905247 |
| C2cd3 | ENSMUSG00000047248 | 277939 | 0.17255642228439 | 3.32968101079336 | 2.00447570646307 | 0.0656512770221 | 0.116517958942839 |
| Med8 | ENSMUSG00000006392 | 80509 | -0.18928793109669 | 3.56378671543138 | -2.00394681365851 | 0.0657139920726 | 0.116610332361427 |
| Gnb4 | ENSMUSG00000027669 | 14696 | -0.271802639273343 | 2.76436598824766 | -2.0035471672766 | 0.0657614178117 | 0.116675549115461 |
| Fem1c | ENSMUSG00000033319 | 240263 | -0.158253511935568 | 4.03533388550401 | -2.00285524762698 | 0.0658436016870 | 0.116792050456594 |
| Ints8 | ENSMUSG00000040738 | 72656 | 0.193932469234545 | 3.39443875920212 | 2.00281445363104 | 0.0658484500012 | 0.116792050456594 |
| Lingo3 | ENSMUSG00000051067 | 237403 | 0.373835863895601 | 2.04661696955594 | 2.00248841949834 | 0.0658872105157 | 0.116840689481004 |
| Inppl1 | ENSMUSG00000032737 | 16332 | 0.131966633128187 | 7.03217456087829 | 2.00240399267652 | 0.0658972509971 | 0.116840689481004 |
| Phf13 | ENSMUSG00000047777 | 230936 | 0.347847577679013 | 1.71197657581425 | 2.00109076071936 | 0.0660536083338 | 0.117098928429713 |
| Sypl2 | ENSMUSG00000027887 | 17306 | 0.154114501889763 | 6.98443739783307 | 2.00028827387014 | 0.0661493223232 | 0.117249593370349 |
| Ccdc77 | ENSMUSG00000030177 | 67200 | -0.215787394808435 | 2.47528644192595 | -2.00009696835269 | 0.0661721584377 | 0.117271054455691 |
| Mvk | ENSMUSG00000041939 | 17855 | 0.253166200808112 | 1.96306895713954 | 1.9995294096016 | 0.0662399504174 | 0.117372166965899 |
| Scamp4 | ENSMUSG00000113949 | 56214 | 0.233143960912576 | 2.32068129120357 | 1.99922564927813 | 0.0662762591992 | 0.117417469903934 |
| Thap4 | ENSMUSG00000026279 | 67026 | -0.208039002704457 | 3.48357796187456 | -1.99912755895028 | 0.0662879879356 | 0.117419218299322 |
| Smad6 | ENSMUSG00000036867 | 17130 | -0.393013489967142 | 1.57671927871946 | -1.99861752204697 | 0.0663490041384 | 0.11750825732952 |
| Med14 | ENSMUSG00000064127 | 26896 | -0.14462703702782 | 4.69554658276262 | -1.99795099989808 | 0.0664288184746 | 0.117630554794486 |
| Alg5 | ENSMUSG00000036632 | 66248 | 0.204589982859405 | 2.51766870648158 | 1.99633621524384 | 0.0666225500907 | 0.11795450195041 |
| Gm10845 | ENSMUSG00000095959 | 100038734 | -0.597408679756272 | 3.19286697568139 | -1.99559307855598 | 0.0667118808938 | 0.118093533611359 |
| F8a | ENSMUSG00000078317 | 14070 | 0.380777957818262 | 1.34843334923792 | 1.99544913526017 | 0.0667291966888 | 0.118105059748415 |
| Zfp146 | ENSMUSG00000037029 | 26465 | -0.159446387943973 | 4.11908399638993 | -1.99387015052981 | 0.0669194128829 | 0.118420043220024 |
| Fbxo22 | ENSMUSG00000032309 | 71999 | -0.163553256035165 | 3.97631245979488 | -1.99379209638549 | 0.0669288287451 | 0.118420043220024 |
| Pola1 | ENSMUSG00000006678 | 18968 | 0.252844177651823 | 1.51521713923988 | 1.99321010097574 | 0.0669990745736 | 0.118525147137804 |
| Gm14288 | ENSMUSG00000078889 | 13999 | -0.188297011820449 | 3.82078378633999 | -1.99204490240844 | 0.0671399151483 | 0.118755082926717 |
| Usp2 | ENSMUSG00000032010 | 53376 | -0.318274434094362 | 6.43294431337721 | -1.99142230752867 | 0.0672152809959 | 0.118869153303169 |
| Brix1 | ENSMUSG00000022247 | 67832 | -0.209112298785031 | 3.41824145560909 | -1.99110993127785 | 0.0672531237144 | 0.118916838454044 |
| Itpr3 | ENSMUSG00000042644 | 16440 | -0.270168530279568 | 3.00457468018828 | -1.99087892176789 | 0.0672811218440 | 0.118942534974653 |
| Cdk12 | ENSMUSG00000003119 | 69131 | 0.170443059555219 | 4.36175690524725 | 1.99081048187961 | 0.0672894187433 | 0.118942534974653 |
| Trim16 | ENSMUSG00000047821 | 94092 | -0.181542428731527 | 5.31497423851172 | -1.99049322077811 | 0.0673278922387 | 0.118991299929132 |
| Adam9 | ENSMUSG00000031555 | 11502 | 0.1369094885895 | 5.70180591963023 | 1.99011769319683 | 0.0673734576221 | 0.119052580871001 |
| Unk | ENSMUSG00000020770 | 217331 | -0.207859686582546 | 2.83540172286002 | -1.98925312279972 | 0.0674784694219 | 0.119218291490408 |
| Bod1 | ENSMUSG00000044502 | 69556 | -0.152212149618091 | 5.67748695452447 | -1.98916609384797 | 0.0674890483709 | 0.119218291490408 |
| Bin1 | ENSMUSG00000024381 | 30948 | 0.143331960921649 | 7.9418936428642 | 1.98788935580949 | 0.0676444191490 | 0.119473444129684 |
| Pdk4 | ENSMUSG00000019577 | 27273 | -0.550214834771682 | 9.35254109850468 | -1.98742809370145 | 0.0677006323020 | 0.119535772678324 |
| Tars2 | ENSMUSG00000028107 | 71807 | -0.138256765191292 | 4.12056092354137 | -1.98738957940287 | 0.0677053279029 | 0.119535772678324 |
| Prpf6 | ENSMUSG00000002455 | 68879 | -0.118591564836798 | 5.01100403414872 | -1.98733063185133 | 0.0677125152707 | 0.119535772678324 |
| Banf1 | ENSMUSG00000024844 | 23825 | -0.140950847098503 | 3.39570686715023 | -1.98651988487807 | 0.0678114391112 | 0.119672998989252 |
| Snhg5 | ENSMUSG00000097195 | 72655 | 0.350258421824213 | 1.24639955075198 | 1.98651410448254 | 0.0678121448851 | 0.119672998989252 |
| Atxn2l | ENSMUSG00000032637 | 233871 | 0.182016592605208 | 5.14076529126791 | 1.9863821745002 | 0.0678282550823 | 0.119682107555364 |
| Ppid | ENSMUSG00000027804 | 67738 | -0.175192043884224 | 4.08857264585673 | -1.98393894088521 | 0.0681272367968 | 0.120190255879062 |
| Elovl1 | ENSMUSG00000006390 | 54325 | 0.367716780839205 | 1.91473157900009 | 1.98319462030488 | 0.0682185598218 | 0.120331338647127 |
| Dhx29 | ENSMUSG00000042426 | 218629 | 0.133123266950269 | 4.88823346657874 | 1.98308734209104 | 0.0682317313595 | 0.120331338647127 |
| Kcnk6 | ENSMUSG00000046410 | 52150 | 0.433658814611147 | 1.04791952438796 | 1.98301812274789 | 0.0682402312938 | 0.120331338647127 |
| Ddx10 | ENSMUSG00000053289 | 77591 | 0.153073930655359 | 3.68589641008135 | 1.98267883282867 | 0.0682819091343 | 0.120385411088334 |
| Ino80 | ENSMUSG00000034154 | 68142 | 0.161852994582015 | 4.35723837908597 | 1.98097377761251 | 0.0684917084619 | 0.120735827317726 |
| Ncln | ENSMUSG00000020238 | 103425 | -0.133337160892104 | 4.31406229671166 | -1.9801265168607 | 0.0685961793623 | 0.120900489617779 |
| Wipf3 | ENSMUSG00000086040 | 330319 | 0.130841605341284 | 4.37279452784934 | 1.97995301936718 | 0.0686175903328 | 0.120918729635368 |
| Rab2b | ENSMUSG00000022159 | 76338 | 0.38623523085098 | 1.10534079466569 | 1.97921393345263 | 0.0687088679524 | 0.12105477475286 |
| Eif4g1 | ENSMUSG00000045983 | 208643 | -0.135791392498925 | 8.58130727476551 | -1.97914861176933 | 0.0687169405673 | 0.12105477475286 |
| Trappc5 | ENSMUSG00000040236 | 66682 | -0.168169215677668 | 4.11548138113671 | -1.97825139965223 | 0.0688279081210 | 0.121230722473664 |
| Rnf144a | ENSMUSG00000020642 | 108089 | -0.370057473526961 | 3.08565996743272 | -1.97783898846104 | 0.0688789702887 | 0.121301115551255 |
| Timm9 | ENSMUSG00000021079 | 30056 | -0.204668612871962 | 2.88342449154864 | -1.97706031826111 | 0.0689754748625 | 0.12145150059961 |
| Katnbl1 | ENSMUSG00000027132 | 72425 | 0.343055301759377 | 1.72950742386437 | 1.97674378585458 | 0.0690147396910 | 0.121501066123883 |
| Atp2b3 | ENSMUSG00000031376 | 320707 | -0.403668702892351 | 1.83108088649221 | -1.97638481169948 | 0.0690592940263 | 0.121559926409362 |
| Lmo7 | ENSMUSG00000033060 | 380928 | 0.174663524039779 | 7.19373728763181 | 1.97615878950086 | 0.0690873604317 | 0.121589749940369 |
| Hsp90b1 | ENSMUSG00000020048 | 22027 | 0.223089637741757 | 7.42940555178864 | 1.97595923620765 | 0.0691121487218 | 0.121613795505228 |
| Stim2 | ENSMUSG00000039156 | 116873 | 0.169507018584165 | 3.68447436364391 | 1.97476124466763 | 0.0692611330116 | 0.121839168651339 |
| Gpkow | ENSMUSG00000031148 | 209416 | -0.175731418164816 | 4.01429115571909 | -1.97475011179651 | 0.0692625188911 | 0.121839168651339 |
| Alg14 | ENSMUSG00000039887 | 66789 | 0.209100138678949 | 2.40357752725632 | 1.97463686498012 | 0.0692766179040 | 0.121844362077162 |
| Tor1b | ENSMUSG00000026848 | 30934 | 0.134681217431639 | 3.94685993054572 | 1.97449454327922 | 0.0692943404069 | 0.121855925834737 |
| Gm20826 | ENSMUSG00000094081 | 100040160 | -1.30632444109324 | -0.554155577551778 | -1.97367866971913 | 0.0693960164907 | 0.121988770134442 |
| Eri1 | ENSMUSG00000031527 | 67276 | 0.207109443261829 | 2.78303679966308 | 1.9736308826745 | 0.0694019760523 | 0.121988770134442 |
| Vmn2r114 | ENSMUSG00000091945 | 666002 | -0.860990260944134 | 2.53828731110338 | -1.9734657386992 | 0.0694225748967 | 0.121988770134442 |
| Loxl4 | ENSMUSG00000025185 | 67573 | 0.664090090777736 | 0.865369519247533 | 1.97345551082934 | 0.0694238508298 | 0.121988770134442 |
| Ppp1r8 | ENSMUSG00000028882 | 100336 | -0.181553190601457 | 3.38099688927133 | -1.97331064677376 | 0.0694419250130 | 0.121988770134442 |
| Timm10 | ENSMUSG00000027076 | 30059 | -0.265432222690886 | 3.00300068525264 | -1.97327171422954 | 0.0694467832244 | 0.121988770134442 |
| Casp9 | ENSMUSG00000028914 | 12371 | 0.371357834142661 | 1.71788499511142 | 1.97326194274396 | 0.0694480026115 | 0.121988770134442 |
| Zfp446 | ENSMUSG00000033961 | 269870 | -0.319626357325055 | 1.47464820883039 | -1.97251150085501 | 0.0695417090276 | 0.12213374379523 |
| Zdhhc6 | ENSMUSG00000024982 | 66980 | -0.151631433290027 | 3.63010807272752 | -1.97211574837838 | 0.0695911725207 | 0.122200981015942 |
| Dctn3 | ENSMUSG00000028447 | 53598 | 0.241646077458383 | 2.82930386939685 | 1.97177440296354 | 0.0696338617322 | 0.12224873691901 |
| Fnbp4 | ENSMUSG00000008200 | 55935 | 0.17475144419246 | 3.84151482542594 | 1.97165472181297 | 0.0696488349191 | 0.12224873691901 |
| Irf7 | ENSMUSG00000025498 | 54123 | -0.823997789934722 | 3.37784124222922 | -1.97163006850384 | 0.0696519196351 | 0.12224873691901 |
| Vmn1r72 | ENSMUSG00000095430 | 252905 | -1.00342971089002 | 0.707630256883505 | -1.9710687877525 | 0.0697221830236 | 0.122352413329869 |
| Rpl37 | ENSMUSG00000041841 | 67281 | -0.155022682884852 | 6.13233388458131 | -1.96983004368832 | 0.0698774833683 | 0.122605260144452 |
| Msto1 | ENSMUSG00000068922 | 229524 | 0.296643659894921 | 2.21978825976357 | 1.96949569333109 | 0.0699194547591 | 0.122615614048549 |
| Bace2 | ENSMUSG00000040605 | 56175 | 0.390054432523962 | 1.82120745369434 | 1.96944726086562 | 0.0699255364533 | 0.122615614048549 |
| Emx2 | ENSMUSG00000043969 | 13797 | 0.341173273295615 | 1.74861983420173 | 1.96942768456245 | 0.0699279947988 | 0.122615614048549 |
| Ndrg3 | ENSMUSG00000027634 | 29812 | -0.16787654537428 | 3.89674137460487 | -1.96942562525989 | 0.0699282534057 | 0.122615614048549 |
| Efcab2 | ENSMUSG00000026495 | 68226 | -0.227028232338405 | 4.22112386590724 | -1.96906766867532 | 0.0699732188235 | 0.1226747802662 1 |
| Olfr31 | ENSMUSG00000072707 | 18330 | -0.965149719968315 | -0.432284209461415 | -1.96890787736475 | 0.0699932998521 | 0.122684368691855 |
| Brpf3 | ENSMUSG00000063952 | 268936 | 0.211484361451765 | 3.27889259410022 | 1.96884554127851 | 0.0700011350774 | 0.122684368691855 |
| Ifi204 | ENSMUSG00000073489 | 15951 | 0.425303823985558 | 3.04795422299663 | 1.9680788836983 | 0.0700975643359 | 0.122833676780445 |
| Csnk1e | ENSMUSG00000022433 | 27373 | -0.139467868915636 | 4.59850945034271 | -1.96779631286509 | 0.0701331363578 | 0.122873595878192 |
| Tm2d3 | ENSMUSG00000078681 | 68634 | -0.339647777968718 | 1.51589529368656 | -1.96771935938555 | 0.0701428266654 | 0.122873595878192 |
| Sema5a | ENSMUSG00000022231 | 20356 | 0.312627677212064 | 2.99743437736894 | 1.96742763405452 | 0.0701795730647 | 0.122918268413784 |
| Mrps23 | ENSMUSG00000023723 | 64656 | -0.155678773698306 | 4.83621910991663 | -1.96704089579275 | 0.0702283146306 | 0.122983932590143 |
| Rhoa | ENSMUSG00000007815 | 11848 | 0.129319044891116 | 6.53652612069536 | 1.96687463111878 | 0.0702492788788 | 0.123000939840659 |
| Chic2 | ENSMUSG00000029229 | 74277 | 0.184238344300354 | 3.39846750422225 | 1.96432129783119 | 0.0705719466291 | 0.123546116047884 |
| Cdkl5 | ENSMUSG00000031292 | 382253 | -0.378100454148756 | 1.44109119862762 | -1.9642125966344 | 0.0705857133257 | 0.123550429521894 |
| Vbp1 | ENSMUSG00000031197 | 22327 | -0.168129706217488 | 4.0241987479812 | -1.96374438063421 | 0.0706450395897 | 0.123634474504563 |
| Zbtb46 | ENSMUSG00000027583 | 72147 | -0.358960943983274 | 2.69185049517165 | -1.96346892737563 | 0.0706799627311 | 0.123675791998477 |
| Ipo4 | ENSMUSG00000002319 | 75751 | 0.154493502421 | 4.98769854670733 | 1.96143827814625 | 0.0709379040666 | 0.124107271023196 |
| Ccnyl1 | ENSMUSG00000070871 | 227210 | 0.139733871987922 | 3.57934062265405 | 1.96101861501701 | 0.0709913184461 | 0.12417896548882 |
| Gm16249 | ENSMUSG00000084949 | 102633259 | -1.10640587092527 | 0.355948423692403 | -1.96093783644865 | 0.0710016040897 | 0.12417896548882 |
| Specc1l | ENSMUSG00000033444 | 74392 | -0.125300061441325 | 6.16776052230848 | -1.96080849582725 | 0.0710180760383 | 0.124187904203275 |
| Dnm1l | ENSMUSG00000022789 | 74006 | -0.147347491278702 | 5.69021340271481 | -1.9606063851437 | 0.0710438224765 | 0.124213055580797 |
| Pdlim5 | ENSMUSG00000028273 | 56376 | -0.161503739927063 | 8.11280240632969 | -1.96034085857563 | 0.0710776602777 | 0.124252343594482 |
| Pus10 | ENSMUSG00000020280 | 74467 | -0.192014368504265 | 3.22318022129535 | -1.9600875170992 | 0.0711099589683 | 0.124288928922822 |
| Rpl41 | ENSMUSG00000093674 | 67945 | 0.128696065998019 | 6.80391703567975 | 1.95962273388539 | 0.0711692493633 | 0.124372672228759 |
| Ppp1r13l | ENSMUSG00000040734 | 333654 | 0.297000659140273 | 1.92455290639129 | 1.9593069247878 | 0.0712095615171 | 0.124423228411678 |
| Parp8 | ENSMUSG00000021725 | 52552 | 0.183073798935968 | 3.37897800362848 | 1.95889855821485 | 0.0712617192772 | 0.124494462748933 |
| D8Ertd738e | ENSMUSG00000019362 | 101966 | -0.182595732357591 | 5.1252776673171 | -1.95878814375241 | 0.0712758277196 | 0.124499212656363 |
| Mettl3 | ENSMUSG00000022160 | 56335 | -0.226979772123634 | 2.83054288053595 | -1.95861221467877 | 0.0712983126981 | 0.124518590206733 |
| Nbea | ENSMUSG00000027799 | 26422 | 0.226199441771048 | 2.95139517115597 | 1.9581108545256 | 0.0713624255875 | 0.124610650814276 |
| Pcdhga5 | ENSMUSG00000103567 | 93713 | 0.42953831176404 | 0.968608074806914 | 1.95783812114479 | 0.0713973242561 | 0.124648273252678 |
| Cant1 | ENSMUSG00000025575 | 76025 | 0.213058639809262 | 3.03305894658471 | 1.95776426329678 | 0.0714067777063 | 0.124648273252678 |
| Xpo6 | ENSMUSG00000000131 | 74204 | 0.230911994730208 | 4.07587382620478 | 1.95691701661763 | 0.0715153029598 | 0.124817783275484 |

| | | | | | | | |
|---|---|---|---|---|---|---|---|
| Rbsn | ENSMUSG00000014550 | 78287 | 0.140960467557461 | 4.42568822076294 | 1.95666782108603 | 0.0715472514693 | 0.124853608845949 |
| Smco4 | ENSMUSG00000058173 | 170748 | -0.391487844063919 | 1.45311893443944 | -1.95640371265859 | 0.0715811261264 | 0.124892783669334 |
| Cmtm6 | ENSMUSG00000032434 | 67213 | 0.261096496346227 | 3.4263101432834 | 1.95609004642258 | 0.0716213761002 | 0.12494306769099 |
| Lrp6 | ENSMUSG00000030201 | 16974 | 0.122488179834091 | 6.51092716131849 | 1.95563008742297 | 0.0716804358575 | 0.125006504623313 |
| Lima1 | ENSMUSG00000023022 | 65970 | 0.305993593999456 | 4.59793467558041 | 1.95558816515191 | 0.0716858209792 | 0.125006504623313 |
| Pla1a | ENSMUSG00000002847 | 85031 | 0.362907732023598 | 1.47735035183033 | 1.95553969185359 | 0.0716920480727 | 0.125006504623313 |
| Cdk9 | ENSMUSG00000009555 | 107951 | 0.192304100657302 | 3.26257948318273 | 1.95484274415597 | 0.0717816356231 | 0.125142752631023 |
| Plxna4 | ENSMUSG00000029765 | 243743 | 0.177199552557014 | 3.89452713918226 | 1.95466203046751 | 0.0718048817206 | 0.12516331718837 |
| Usp19 | ENSMUSG00000006676 | 71472 | -0.114840339338511 | 6.38200254152859 | -1.95450650166929 | 0.0718248936595 | 0.125178238614878 |
| Sec24c | ENSMUSG00000039367 | 218811 | 0.117433200947706 | 5.35717268947275 | 1.95426018472007 | 0.0718565977484 | 0.125213529409831 |
| Tet1 | ENSMUSG00000047146 | 52463 | 0.529383609955395 | 2.21697406261226 | 1.953813144345 | 0.0719141700784 | 0.125282090860819 |
| Hnrnpa2b1 | ENSMUSG00000004980 | 53379 | -0.128122245102959 | 7.51065140621601 | -1.95377669853811 | 0.0719188656254 | 0.125282090860819 |
| Lemd3 | ENSMUSG00000048661 | 380664 | -0.200312544741676 | 3.30722939284326 | -1.95368109550876 | 0.0719311841058 | 0.125283584044088 |
| Serinc1 | ENSMUSG00000019877 | 56442 | 0.124603606255932 | 6.41436819715614 | 1.95345004105541 | 0.0719609634918 | 0.125315483818612 |
| 2310009A05Rik | ENSMUSG00000098332 | 66364 | -0.328854154278398 | 2.04322348278076 | -1.9529855487876 | 0.0720208634726 | 0.125383667912117 |
| Dalrd3 | ENSMUSG00000019039 | 67789 | -0.274827337880874 | 2.04190719120894 | -1.95296853538502 | 0.0720230583496 | 0.125383667912117 |
| Tmed8 | ENSMUSG00000034111 | 382620 | 0.492068415851119 | 1.71058854130224 | 1.95274108474051 | 0.0720524073233 | 0.125414787333109 |
| Arap2 | ENSMUSG00000037999 | 212285 | -0.395462523894496 | 2.21525835742798 | -1.95239815317823 | 0.0720966779378 | 0.125471865393853 |
| Fam53b | ENSMUSG00000030956 | 77938 | 0.181455412648151 | 4.21667414169359 | 1.95228411576796 | 0.0721114050478 | 0.125477518079961 |
| Ewsr1 | ENSMUSG00000009079 | 14030 | 0.172133393674199 | 5.51283366543046 | 1.95182719410981 | 0.0721704407120 | 0.12556025583693 |
| Nr6a1 | ENSMUSG00000063972 | 14536 | -0.387981252564043 | 1.76379006122246 | -1.95003499246646 | 0.0724024243177 | 0.125943809709946 |
| Rps6ka1 | ENSMUSG00000003644 | 20111 | 0.316902471681019 | 1.82819177067492 | 1.94955614532751 | 0.0724645215595 | 0.126031771705316 |
| Smarcal1 | ENSMUSG00000039354 | 54380 | 0.172257582990643 | 4.21965778710527 | 1.94942969189213 | 0.0724809282347 | 0.126040252391619 |
| Mea1 | ENSMUSG00000002768 | 17256 | -0.189126314221297 | 3.88585046968278 | -1.94919429429135 | 0.0725114788685 | 0.126073322015541 |
| Tspan4 | ENSMUSG00000025511 | 64540 | 0.292008498107564 | 2.35378756488321 | 1.94844399759427 | 0.0726089330673 | 0.126222685648789 |
| Prps1 | ENSMUSG00000031432 | 19139 | -0.216597839491282 | 3.63029717934995 | -1.94803660059251 | 0.0726618988668 | 0.126294675864847 |
| Hras | ENSMUSG00000025499 | 15461 | 0.155179755997993 | 4.64096673723764 | 1.94792856133325 | 0.0726759509886 | 0.126299017528975 |
| Pdzd8 | ENSMUSG00000074746 | 107368 | -0.125546152274089 | 5.11261492762857 | -1.94759275716202 | 0.0727196431604 | 0.126340780608563 |
| Cmya5 | ENSMUSG00000047419 | 76469 | -0.238447719948409 | 11.5467224760928 | -1.94756620704053 | 0.0727230986724 | 0.126340780608563 |
| Maml2 | ENSMUSG00000031925 | 270118 | 0.219106012437811 | 3.20690770255835 | 1.94651857127206 | 0.0728595685300 | 0.126557753631368 |
| Rps27a | ENSMUSG00000020460 | 78294 | 0.211063337362503 | 6.5447220625875 | 1.94531818080458 | 0.0730162238085 | 0.12679613989642 |
| 1110008L16Rik | ENSMUSG00000021023 | 66132 | -0.266603596951083 | 1.82434688673251 | -1.94528921777191 | 0.0730200073779 | 0.12679613989642 |
| Dmac1 | ENSMUSG00000028398 | 66928 | -0.161971782388515 | 3.87683626078826 | -1.94518281237837 | 0.0730339091199 | 0.126800136688348 |
| Lsm7 | ENSMUSG00000035215 | 66094 | 0.186586162514236 | 3.04285439949852 | 1.94456905929588 | 0.0731141423354 | 0.126919277412841 |
| Eml5 | ENSMUSG00000051166 | 319670 | 0.35339331785997 | 1.18202043523316 | 1.94382501289727 | 0.0732115159040 | 0.12706812962002 |
| Gm18852 | ENSMUSG00000099241 | 100417829 | -2.57477133958235 | -0.984179566286857 | -1.94299537597662 | 0.0733202300304 | 0.127236614456678 |
| Chad | ENSMUSG00000039084 | 12643 | -0.599659334779298 | 3.99407259538778 | -1.94140347213481 | 0.0735292421220 | 0.127575043026543 |
| Snu13 | ENSMUSG00000063480 | 20826 | -0.177897708457018 | 4.59905691015685 | -1.94133235890029 | 0.0735385917217 | 0.127575043026543 |
| Usp4 | ENSMUSG00000032612 | 22258 | 0.118903444834836 | 5.78670006943286 | 1.94085317866264 | 0.0736016200901 | 0.127664123961446 |
| Adprhl2 | ENSMUSG00000042558 | 100206 | 0.305855737617898 | 1.70496821523642 | 1.939263782554 | 0.0738110316592 | 0.128007042212831 |
| Snora21 | ENSMUSG00000064901 | 100302498 | 0.60548279420267 | 3.08274027515829 | 1.93842707169467 | 0.0739214906933 | 0.128171938619516 |
| Garem1 | ENSMUSG00000042680 | 381126 | 0.20717316740022 | 3.5487650244318 | 1.93836595606601 | 0.0739295648153 | 0.128171938619516 |
| Cenpb | ENSMUSG00000068267 | 12616 | 0.145753093950115 | 6.80194069359567 | 1.93791816764414 | 0.0739887476379 | 0.128238833100399 |
| Rpusd2 | ENSMUSG00000027324 | 271842 | -0.417620069869831 | 1.35146993589387 | -1.93789649697584 | 0.0739916128775 | 0.128238833100399 |
| Ccdc91 | ENSMUSG00000030301 | 67015 | -0.189939083316039 | 5.39972498906048 | -1.93772897138137 | 0.0740137660864 | 0.128256892373306 |
| Tbc1d8b | ENSMUSG00000042473 | 245638 | 0.282244746337277 | 2.35697787653597 | 1.93755276866934 | 0.0740370732528 | 0.12827694527285 |
| Pam16 | ENSMUSG00000014301 | 66449 | -0.245605163972252 | 3.39237098909326 | -1.93666874051539 | 0.0741541087142 | 0.128459360119686 |
| Plcl1 | ENSMUSG00000038349 | 227120 | -0.227200408283094 | 2.95733311542509 | -1.93631516000817 | 0.0742009659534 | 0.128520164607328 |
| Rbm14 | ENSMUSG00000006456 | 56275 | 0.238162345232226 | 2.69997829567014 | 1.93562984816673 | 0.0742918616728 | 0.128657214820375 |
| Pcbd2 | ENSMUSG00000021496 | 72562 | -0.313272943757843 | 1.33715528661492 | -1.93473768000082 | 0.0744103454207 | 0.128841990463144 |
| Olfr635 | ENSMUSG00000094520 | 259122 | -0.789547558276629 | 2.52159357314222 | -1.93420466721253 | 0.0744812138242 | 0.128944274360679 |
| Sharpin | ENSMUSG00000022552 | 106025 | -0.197979988361585 | 3.15735495372298 | -1.93309198507128 | 0.0746293519518 | 0.129180276513979 |
| Rcbtb1 | ENSMUSG00000035469 | 71330 | -0.161853977238334 | 3.59197520647805 | -1.93272702699578 | 0.0746779993846 | 0.129244016984505 |
| Pgm2l1 | ENSMUSG00000030729 | 70974 | -0.188401800699605 | 3.01201171499079 | -1.93213469498032 | 0.0747570162511 | 0.129353179446679 |
| Zfp960 | ENSMUSG00000096696 | 449000 | -0.3034760824668 | 1.98917525372428 | -1.93207681606234 | 0.0747647413543 | 0.129353179446679 |
| Arhgap24 | ENSMUSG00000057315 | 231532 | 0.173184408489425 | 5.68107816265475 | 1.93178206683262 | 0.0748040928204 | 0.129400781550883 |
| Shank3 | ENSMUSG00000022623 | 58234 | -0.319989042508761 | 4.39689599021884 | -1.93151313679359 | 0.0748400136462 | 0.129442434994773 |
| Bag6 | ENSMUSG00000024392 | 224727 | -0.128208423702231 | 6.93848781241861 | -1.93126704673867 | 0.0748728974954 | 0.129463148660496 |
| 2210418O10Rik | ENSMUSG00000078894 | 100504263 | -0.151867768838969 | 3.80362774019261 | -1.93124625013085 | 0.0748756770498 | 0.129463148660496 |
| Dpagt1 | ENSMUSG00000032123 | 13478 | 0.259236901457832 | 2.28272612042301 | 1.9296469810676 | 0.0750897074235 | 0.129812682563108 |
| Rpl38 | ENSMUSG00000057322 | 67671 | -0.162653286403561 | 5.71175907966993 | -1.92946647723424 | 0.0751138991823 | 0.12983397089853 |
| Zfp664 | ENSMUSG00000079215 | 269704 | -0.12659916124498 | 4.57652612564504 | -1.92902821052934 | 0.0751726667252 | 0.129915007110424 |
| Rsph3a | ENSMUSG00000073471 | 66832 | -0.18322670377195 | 3.75217188088211 | -1.92882361291716 | 0.0752001156989 | 0.129941900838607 |
| Edf1 | ENSMUSG00000015092 | 59022 | -0.147179990477039 | 5.66972822791179 | -1.92828965037774 | 0.0752717954588 | 0.130045202491038 |
| Palm | ENSMUSG00000035863 | 18483 | -0.277722251497756 | 4.3240928430911 | -1.92780620564751 | 0.0753367472249 | 0.130128096778467 |
| Supt4a | ENSMUSG00000020485 | 20922 | -0.218703207412241 | 2.29888164786475 | -1.92775533534235 | 0.0753435847141 | 0.130128096778467 |
| Brpf1 | ENSMUSG00000001632 | 78783 | 0.161701776049209 | 3.9469129712179 | 1.92701037575446 | 0.0754437795470 | 0.130280561489477 |
| Zxdc | ENSMUSG00000034430 | 80292 | 0.154995647166533 | 4.15223777868518 | 1.92681480338553 | 0.0754701035104 | 0.130296540814883 |
| Zfp532 | ENSMUSG00000042439 | 328977 | -0.13418046492932 | 4.16906684401766 | -1.9267645214552 | 0.0754768727874 | 0.130296540814883 |
| Zfp36l1 | ENSMUSG00000021127 | 12192 | 0.201639916300223 | 5.54512673731588 | 1.92650603957278 | 0.0755116799938 | 0.130336045162335 |
| Fzd6 | ENSMUSG00000022297 | 14368 | -0.278383376430842 | 2.51035007837993 | -1.92626826272825 | 0.0755437119407 | 0.130364845790173 |
| Ccnj | ENSMUSG00000025010 | 240665 | -0.282906974630509 | 1.99478733291282 | -1.92620513454635 | 0.0755522182856 | 0.130364845790173 |
| Fam189a2 | ENSMUSG00000071604 | 381217 | -0.196153174258095 | 3.46995835656184 | -1.92588043576318 | 0.0755959842898 | 0.130404099262287 |
| Hr | ENSMUSG00000022096 | 15460 | -0.239031764758125 | 4.24609058626868 | -1.92575671076178 | 0.0756126671899 | 0.130404099262287 |
| Stx2 | ENSMUSG00000029428 | 13852 | 0.246644324484605 | 3.1109670226004 | 1.92574797845111 | 0.0756138447684 | 0.130404099262287 |
| Cldnd1 | ENSMUSG00000022744 | 224250 | -0.201950486651765 | 4.01108982439866 | -1.9256824173272 | 0.0756226864169 | 0.130404099262287 |
| Ciart | ENSMUSG00000038550 | 229599 | -0.838103469545691 | 0.114179648080978 | -1.92554881970979 | 0.0756407064534 | 0.130414599722903 |
| Stag2 | ENSMUSG00000025862 | 20843 | 0.108363520938394 | 5.8378884756736 | 1.9252283933201 | 0.0756839424062 | 0.13045704182934 |
| Cep250 | ENSMUSG00000038241 | 16328 | 0.251280660237551 | 3.52560644964489 | 1.9251406460432 | 0.0756957862877 | 0.13045704182934 |
| Cept1 | ENSMUSG00000040774 | 99712 | -0.225694769243997 | 2.19858042565732 | -1.92510108458453 | 0.0757011267334 | 0.13045704182934 |
| Olfr607 | ENSMUSG00000081945 | 546989 | -0.744699701180826 | 1.51157204539126 | -1.9238293217831 | 0.0758729859506 | 0.130732599215957 |
| Flt4 | ENSMUSG00000020357 | 14257 | -0.330259291340211 | 2.86125557536047 | -1.92370747857674 | 0.0758894697858 | 0.130740393101442 |
| Fggy | ENSMUSG00000028573 | 75578 | 0.310124322453038 | 1.80779361401516 | 1.92308266948432 | 0.0759740496424 | 0.130865480088551 |
| Bptf | ENSMUSG00000040481 | 207165 | 0.144712878711529 | 5.93619910617324 | 1.92279275184688 | 0.0760133246210 | 0.130912502195092 |
| Nono | ENSMUSG00000031311 | 53610 | 0.123091799593814 | 5.30756391946483 | 1.92212976375578 | 0.0761032085752 | 0.131046656102137 |
| Akirin1 | ENSMUSG00000023075 | 68050 | -0.186503345775487 | 4.0955187232453 | -1.92123189995462 | 0.0762250894953 | 0.131235856556002 |
| Pigx | ENSMUSG00000023791 | 72084 | -0.387016247622242 | 1.63821646183521 | -1.92095279625473 | 0.0762630126600 | 0.131262182132191 |
| Glyr1 | ENSMUSG00000022536 | 74022 | -0.128266103033337 | 4.88517879284864 | -1.9209426135804 | 0.0762643965530 | 0.131262182132191 |
| Fam129a | ENSMUSG00000026483 | 63913 | 0.200120359286617 | 6.00536391142422 | 1.92033944810135 | 0.0763464114466 | 0.131375553678568 |
| Gm8995 | ENSMUSG00000063286 | 668139 | 0.274022399885691 | 3.60258241856988 | 1.92028143764839 | 0.0763543035879 | 0.131375553678568 |
| Dnajc8 | ENSMUSG00000054405 | 68598 | -0.118688108173425 | 5.41113798390866 | -1.92004833473407 | 0.0763860239842 | 0.131409447304361 |
| Sh3tc1 | ENSMUSG00000036553 | 231147 | -0.260736519580802 | 2.17289110897242 | -1.91994625206093 | 0.0763999190535 | 0.131412669579004 |
| Vps39 | ENSMUSG00000027291 | 269338 | 0.116323375616056 | 4.36061696252205 | 1.91983644203511 | 0.0764148685007 | 0.131417704150677 |
| Ciao1 | ENSMUSG00000003662 | 26371 | -0.142890524752384 | 4.0759904040265 | -1.91959368630532 | 0.0764479264959 | 0.131453875181181 |
| Ptger4 | ENSMUSG00000039942 | 19219 | 0.50467023560411 | 1.75412926466999 | 1.91899825249594 | 0.0765290664977 | 0.131572699887834 |
| Pcbp1 | ENSMUSG00000051695 | 23983 | 0.110415849495169 | 5.68238723598358 | 1.91790591261369 | 0.0766781234424 | 0.131808235154865 |
| S1pr1 | ENSMUSG00000045092 | 13609 | -0.322411419150185 | 4.47812787675275 | -1.91771109450881 | 0.0767047353336 | 0.131833248731495 |
| Cep95 | ENSMUSG00000018372 | 320162 | -0.286738707217639 | 1.5083016273545 | -1.91718738142357 | 0.0767763154115 | 0.131929355908856 |
| Kansl1l | ENSMUSG00000026004 | 68691 | -0.187541105969791 | 4.34077375104204 | -1.91712539149589 | 0.0767847920860 | 0.131929355908856 |
| Psmd10 | ENSMUSG00000031429 | 53380 | 0.295583254203909 | 1.8372775922592 | 1.91627175937739 | 0.0769016065444 | 0.132109297679897 |
| Tsc22d1 | ENSMUSG00000022010 | 21807 | -0.12769195435934 | 6.66389424858308 | -1.9161029914636 | 0.0769247205168 | 0.132128240095747 |
| Lrrc45 | ENSMUSG00000025145 | 217366 | 0.345587360881053 | 1.99553320603131 | 1.91424858143535 | 0.0771791108601 | 0.132544361478783 |
| Rnf121 | ENSMUSG00000070426 | 75212 | 0.161216491874722 | 2.73829571794458 | 1.9134082484316 | 0.0772946399977 | 0.132721914831192 |
| Fads2 | ENSMUSG00000024665 | 56473 | 0.394305639959575 | 1.38770406860399 | 1.91252958281662 | 0.0774156069114 | 0.13290874809938 |
| Cep19 | ENSMUSG00000035790 | 66994 | -0.216939194177992 | 2.81364472029323 | -1.91212930282624 | 0.0774707708484 | 0.132982568490012 |
| Prdm11 | ENSMUSG00000075028 | 100042784 | -0.586156462402366 | 2.15675484116295 | -1.91202811767611 | 0.0774847211613 | 0.132985631498681 |
| Smarcd1 | ENSMUSG00000023018 | 83797 | 0.217386710874993 | 2.82562526176991 | 1.91180247771092 | 0.0775158381646 | 0.133018151801832 |
| Zfp963 | ENSMUSG00000092260 | 620419 | -0.36391605909304 | 1.07121868970259 | -1.91127547418096 | 0.0775885590396 | 0.133112772567453 |
| Pias3 | ENSMUSG00000028101 | 229615 | 0.264586428684794 | 2.16933191706989 | 1.91122640536214 | 0.0775953331625 | 0.133112772567453 |
| Vmn2r97 | ENSMUSG00000091491 | 627367 | -0.734371055587977 | 0.698812621337264 | -1.91096460254108 | 0.0776314850307 | 0.133153443812757 |
| Zfp189 | ENSMUSG00000039634 | 230162 | -0.254520664621296 | 1.39210109379403 | -1.91087831142529 | 0.0776434041590 | 0.133153443812757 |
| Yaf2 | ENSMUSG00000022634 | 67057 | -0.268891681097395 | 3.16867948469993 | -1.90954962799243 | 0.0778271408167 | 0.133447604120444 |
| Cdc5l | ENSMUSG00000023932 | 71702 | -0.139691353690286 | 5.67026869139566 | -1.90899292939651 | 0.0779042408998 | 0.133558854654416 |
| Hcfc2 | ENSMUSG00000020246 | 67933 | -0.153727650880253 | 3.53705645655996 | -1.90819406614086 | 0.0780150006435 | 0.133727767294064 |
| Rnf167 | ENSMUSG00000040746 | 70510 | -0.30558603081212 | 3.78550501062112 | -1.90781043413232 | 0.0780682406847 | 0.133798046240995 |
| Mynn | ENSMUSG00000037730 | 80732 | -0.144530340423635 | 3.50520790165819 | -1.90741115000065 | 0.0781236878962 | 0.133872085341533 |

| | | | | | | | |
|---|---|---|---|---|---|---|---|
| Rreb1 | ENSMUSG00000039087 | 68750 | -0.155057912244857 | 5.69077097445294 | -1.90687074318227 | 0.0781987891939 | 0.133979775027983 |
| 1110004F10Rik | ENSMUSG00000030663 | 56372 | -0.130637947579511 | 4.68541895762879 | -1.90620279986924 | 0.0782917049086 | 0.134117948026447 |
| Spty2d1 | ENSMUSG00000049516 | 101685 | 0.248323030185393 | 3.05281305609303 | 1.90517910402593 | 0.0784343026004 | 0.134341172316789 |
| Chrdl1 | ENSMUSG00000031283 | 83453 | 0.496068520264765 | 1.37510757883005 | 1.90424654624822 | 0.0785644099036 | 0.134542936653144 |
| Vps50 | ENSMUSG00000001376 | 73288 | -0.143995939676236 | 4.19687734636799 | -1.90373123903154 | 0.0786363876957 | 0.134645105561172 |
| Vps35 | ENSMUSG00000031696 | 65114 | -0.142142224200154 | 6.00415027951672 | -1.90264185499994 | 0.0787887489186 | 0.134884857389154 |
| Mapk1ip1l | ENSMUSG00000021840 | 218975 | 0.150697583101664 | 4.4482351416129 | 1.90166746479603 | 0.0789252536756 | 0.135097392409813 |
| Recql5 | ENSMUSG00000020752 | 170472 | -0.249719091334658 | 2.20700992972238 | -1.90043804475591 | 0.0790977919034 | 0.135371530185794 |
| R3hdm4 | ENSMUSG00000035781 | 109284 | -0.112573616338269 | 5.35620284056432 | -1.89950034744411 | 0.0792296190236 | 0.135575918213367 |
| Rhbdl3 | ENSMUSG00000017692 | 246104 | 0.341740740097747 | 1.07491385092923 | 1.89857194510508 | 0.0793603354100 | 0.135778341895064 |
| Cpq | ENSMUSG00000039007 | 54381 | 0.240490363983116 | 3.21159879207562 | 1.89774322270794 | 0.0794771820868 | 0.135956976117613 |
| Mlx | ENSMUSG00000017801 | 21428 | -0.252432577021913 | 2.4333211242687 | -1.89752399069045 | 0.0795081190053 | 0.135970657909274 |
| Grwd1 | ENSMUSG00000053801 | 101612 | 0.204937787234051 | 2.41153632099584 | 1.89751025185106 | 0.0795100581242 | 0.135970657909274 |
| Vmn2r120 | ENSMUSG00000090655 | 224916 | -0.563735653770008 | 2.29549134778195 | -1.897261121253 | 0.0795452281920 | 0.136009524380974 |
| Fbxl20 | ENSMUSG00000020883 | 72194 | 0.183208970360928 | 4.09659709254256 | 1.89707713659903 | 0.0795712105698 | 0.1360326716827 |
| Rcn2 | ENSMUSG00000032320 | 26611 | 0.163888754140863 | 4.1151728296915 | 1.89580702292587 | 0.0797507861822 | 0.13631834929763 |
| Mpg | ENSMUSG00000020287 | 268395 | 0.382452255693766 | 1.24253451264758 | 1.89519036983455 | 0.0798381041977 | 0.136442723986213 |
| Nova1 | ENSMUSG00000021047 | 664883 | -0.226737856403143 | 2.19216934429537 | -1.89511689573423 | 0.0798485138907 | 0.136442723986213 |
| Eif4enif1 | ENSMUSG00000020454 | 74203 | -0.115645919615559 | 4.83283946569014 | -1.89421899530536 | 0.0799758262661 | 0.136609555510825 |
| Rmi1 | ENSMUSG00000035367 | 74386 | -0.218591840670908 | 2.20423790267179 | -1.8941948462429 | 0.0799792528724 | 0.136609555510825 |
| Ppil1 | ENSMUSG00000024007 | 68816 | 0.368670831069922 | 1.42290251594812 | 1.89416393800708 | 0.0799836387585 | 0.136609555510825 |
| Gipc1 | ENSMUSG00000019433 | 67903 | 0.159526776325623 | 2.99524366768139 | 1.89373008781213 | 0.0800452251809 | 0.136693384854392 |
| Brip1os | ENSMUSG00000085208 | 74038 | 0.252405124145813 | 2.44547386509242 | 1.89358647823485 | 0.0800656204716 | 0.136706856822142 |
| Psmc6 | ENSMUSG00000021832 | 67089 | 0.109883207054124 | 5.8277650933977 | 1.89234582265289 | 0.0802420130532 | 0.136986638245331 |
| Traf2 | ENSMUSG00000026942 | 22030 | 0.272653141189466 | 3.12069714020087 | 1.89198021299431 | 0.0802940613445 | 0.137054088781612 |
| Eif1ax | ENSMUSG00000067194 | 66235 | -0.142952382186745 | 5.25149999704557 | -1.89177863330665 | 0.0803227713460 | 0.137081688303467 |
| Fbxw5 | ENSMUSG00000015095 | 30839 | -0.129346013926767 | 5.00457035764981 | -1.89085812571388 | 0.0804539927823 | 0.137267615693104 |
| Ggnbp2 | ENSMUSG00000020530 | 217039 | -0.164979420385063 | 4.73949126065169 | -1.89083823465424 | 0.0804568304588 | 0.137267615693104 |
| 3110039I08Rik | ENSMUSG00000074415 | 73144 | 0.357520730264559 | 2.85923506456581 | 1.89052409302083 | 0.0805016581954 | 0.137305193621354 |
| Snhg6 | ENSMUSG00000098234 | 73824 | -0.338915436257043 | 1.68288252963628 | -1.89050783917478 | 0.0805039782196 | 0.137305193621354 |
| Gdf1 | ENSMUSG00000087408 | 14559 | 0.459289422187463 | 1.12590960055474 | 1.89031635781546 | 0.0805313142367 | 0.137330389379394 |
| Emc7 | ENSMUSG00000055943 | 73024 | -0.134143435700279 | 4.17628234907161 | -1.88994369601099 | 0.0805845397782 | 0.137399719905869 |
| Lsm11 | ENSMUSG00000044847 | 72290 | -0.241364119357083 | 1.47041008109779 | -1.88954825260115 | 0.0806410539049 | 0.137474635095925 |
| Snora44 | ENSMUSG00000064604 | 100217418 | 1.18773798365778 | -0.752105877155373 | 1.88938418720236 | 0.0806645115540 | 0.13749318194253 |
| Pax7 | ENSMUSG00000028736 | 18509 | 0.516380129581477 | 1.84975805056832 | 1.88650334472139 | 0.0810774137706 | 0.138175430297259 |
| Pfkfb3 | ENSMUSG00000026773 | 170768 | 0.342150880542905 | 3.23577438805595 | 1.88606916099016 | 0.0811398094200 | 0.138260211499764 |
| Dlx6 | | 13396 | -1.01940311986167 | -0.623544801244044 | -1.88594236267274 | 0.0811580395333 | 0.138265988310728 |
| Mospd3 | ENSMUSG00000037221 | 68929 | 0.29404908928079 | 1.59538122218723 | 1.88586963409597 | 0.0811684975747 | 0.138265988310728 |
| Mbd5 | | 109241 | -0.137701699767184 | 4.44382088218999 | -1.8857184874518 | 0.0811902356776 | 0.138281468711887 |
| Appbp2 | ENSMUSG00000018481 | 66884 | -0.156649242965639 | 4.41205367601224 | -1.88561294427349 | 0.0812054181524 | 0.138285780623907 |
| Scarf1 | ENSMUSG00000038188 | 380713 | -0.333886616993422 | 1.90132863589053 | -1.88542928581712 | 0.0812318436944 | 0.138301788603257 |
| Ell | ENSMUSG00000070002 | 13716 | -0.468130015083461 | 3.82549193851587 | -1.88537175539157 | 0.0812401230099 | 0.138301788603257 |
| Creld1 | ENSMUSG00000030284 | 171508 | 0.129910224573146 | 4.03904701956128 | 1.88499103793028 | 0.0812949320297 | 0.138373544381335 |
| Ube2d3 | ENSMUSG00000078578 | 66105 | -0.125277444716906 | 7.34620912709809 | -1.88486607571296 | 0.0813129291816 | 0.138382629438629 |
| Disp1 | ENSMUSG00000030768 | 68897 | 0.343999621235246 | 2.44899803163546 | 1.88394604492001 | 0.0814455436029 | 0.138569862945896 |
| Mettl21a | ENSMUSG00000025956 | 67099 | 0.240516445215404 | 2.31239767316252 | 1.88392693505453 | 0.0814483001946 | 0.138569862945896 |
| Inpp5k | ENSMUSG00000006127 | 19062 | -0.169464084525691 | 3.72757203674355 | -1.88336416560738 | 0.0815295173125 | 0.138686454052807 |
| Eea1 | ENSMUSG00000036499 | 216238 | -0.119008362764022 | 6.11445387725035 | -1.88297443992074 | 0.0815858041828 | 0.138760607674331 |
| Pparg | ENSMUSG00000000440 | 19016 | -0.410049893711584 | 1.18064152311631 | -1.88236601992214 | 0.0816737465976 | 0.138888569393066 |
| Bysl | ENSMUSG00000023988 | 53414 | 0.149372390548874 | 3.98921003809409 | 1.88102291239805 | 0.0818681857352 | 0.139197563893601 |
| Hectd1 | ENSMUSG00000035247 | 207304 | -0.132228411909352 | 7.962659034076 | -1.8803564229264 | 0.0819648273217 | 0.139335420839525 |
| Inpp4a | ENSMUSG00000026113 | 269180 | -0.133160903521219 | 4.89407867351005 | -1.88028796876082 | 0.0819747590722 | 0.139335420839525 |
| Eif4g2 | ENSMUSG00000005610 | 13690 | -0.120827955532485 | 9.01434720841412 | -1.87993674861612 | 0.0820257334600 | 0.139400387314236 |
| Tfpt | ENSMUSG00000006335 | 69714 | -0.22448064782953 | 2.58406558159902 | -1.8797691309045 | 0.0820500706042 | 0.139420071782211 |
| Slc43a3 | ENSMUSG00000027074 | 58207 | -0.320738591021665 | 3.82979611358999 | -1.87925259486575 | 0.0821251100927 | 0.139525890336374 |
| Lin37 | ENSMUSG00000036845 | 75660 | 0.302698567318637 | 1.46204688428965 | 1.87833267656809 | 0.0822589042231 | 0.139731481284075 |
| D130040H23Rik | ENSMUSG00000079038 | 211135 | -0.803676558911521 | 1.22649116830032 | -1.87748800473842 | 0.0823819277253 | 0.139918715345865 |
| Ficd | ENSMUSG00000053334 | 231630 | 0.246037628508433 | 2.63756063203532 | 1.87727967791708 | 0.0824122953293 | 0.139948547498142 |
| Zfp141 | ENSMUSG00000092416 | 434178 | 0.259265762675826 | 1.65118374150526 | 1.87627924979905 | 0.0825582676504 | 0.140174654191804 |
| Ilf3 | ENSMUSG00000032178 | 16201 | 0.117844062442344 | 4.23667137510536 | 1.87616289655097 | 0.0825752598803 | 0.140176804820968 |
| Elk4 | ENSMUSG00000026436 | 13714 | 0.121243169544556 | 5.80073960889793 | 1.87609496631725 | 0.0825851818723 | 0.140176804820968 |
| Vps33a | ENSMUSG00000029434 | 77573 | -0.133042487182708 | 4.72347567941612 | -1.87570478055803 | 0.0826421938394 | 0.140251796438358 |
| Mtmr2 | ENSMUSG00000031918 | 77116 | -0.150431668026835 | 4.00889785362276 | -1.87512380235879 | 0.0827271492702 | 0.140374180172778 |
| Git1 | ENSMUSG00000011877 | 216963 | -0.125485260122146 | 5.65956940005033 | -1.87473706021836 | 0.0827837455972 | 0.140448412715116 |
| Srpk1 | ENSMUSG00000004865 | 20815 | 0.122685263936076 | 4.26939382464277 | 1.87452306539734 | 0.0828150768833 | 0.140476834233107 |
| Amotl1 | ENSMUSG00000013076 | 75723 | 0.143607647301979 | 9.66966219817489 | 1.87444710855906 | 0.0828262004054 | 0.140476834233107 |
| Rrp1b | ENSMUSG00000058392 | 72462 | 0.196736312950453 | 2.74122445528928 | 1.87433274327506 | 0.0828429512079 | 0.140483447045288 |
| Ttc33 | ENSMUSG00000022151 | 67515 | -0.179420392392087 | 4.32989532124181 | -1.87271114841987 | 0.0830807914719 | 0.140864918811772 |
| Trim32 | ENSMUSG00000051675 | 69807 | 0.246880829103728 | 2.4765202900583 | 1.8721566816606 | 0.0831622566921 | 0.140959984661864 |
| Usp37 | ENSMUSG00000033364 | 319651 | 0.17462387787832 | 3.23209350231999 | 1.87215399725445 | 0.0831626512747 | 0.140959984661864 |
| Sdcbp | ENSMUSG00000028249 | 53378 | 0.172499322844022 | 4.44028225431386 | 1.87199677815245 | 0.0831857639572 | 0.140977300126626 |
| 2310009B15Rik | ENSMUSG00000079283 | 69549 | -0.235710924120714 | 2.18097226058 | -1.87184703995869 | 0.0832077822645 | 0.140992755841606 |
| Plaa | ENSMUSG00000028577 | 18786 | 0.144312171574323 | 5.24415973739733 | 1.87151922280135 | 0.0832560046163 | 0.141052601745376 |
| Invs | ENSMUSG00000028344 | 16348 | 0.347302664614207 | 1.92344089046348 | 1.8711235959314 | 0.0833142354454 | 0.141129382869041 |
| Thyn1 | ENSMUSG00000035443 | 77862 | 0.304661812225918 | 2.56043067748651 | 1.87100989647508 | 0.0833309772361 | 0.14113587111374 |
| Pqlc3 | ENSMUSG00000045679 | 217430 | 0.431373361035378 | 1.47442524416209 | 1.86984249432807 | 0.0835030481721 | 0.141405394201379 |
| Pcf11 | ENSMUSG00000041328 | 74737 | -0.186128968194844 | 4.69238009763886 | -1.86975080374738 | 0.0835165766042 | 0.141406396973468 |
| Gm10033 | ENSMUSG00000110444 | 378466 | -0.270101198853155 | 3.41835530425158 | -1.86858040110418 | 0.0836894367185 | 0.141677130675368 |
| Prune1 | ENSMUSG00000015711 | 229589 | -0.13948909120152 | 5.08486534949917 | -1.86820236618081 | 0.0837453386808 | 0.141749813737988 |
| Gpatch2l | ENSMUSG00000021254 | 70373 | -0.151306532678482 | 3.96544824257596 | -1.86780380921412 | 0.0838043118061 | 0.141827671830415 |
| Alpk3 | ENSMUSG00000038763 | 116904 | -0.159105704315184 | 9.8978238798409 | -1.86717345074885 | 0.0838976602686 | 0.141963672507195 |
| Naa30 | ENSMUSG00000036282 | 70646 | 0.146780208619751 | 3.6309166743621 | 1.86655518781156 | 0.0839893085928 | 0.142078156927463 |
| Galnt10 | ENSMUSG00000020520 | 171212 | 0.251897264535987 | 3.27551416231062 | 1.8665416680798 | 0.0839913137009 | 0.142078156927463 |
| Ddx6 | ENSMUSG00000032097 | 13209 | 0.103971740750051 | 6.33125440047837 | 1.86599848616043 | 0.0840719085862 | 0.142192485340562 |
| Sppl2a | ENSMUSG00000027366 | 66552 | 0.103382481625341 | 5.19147339855292 | 1.86383892885346 | 0.0843930239841 | 0.142713512557393 |
| Clip4 | ENSMUSG00000024059 | 78785 | -0.119814798764176 | 7.01693152206082 | -1.86350889800284 | 0.0844421951849 | 0.142774576267112 |
| Pwwp2a | ENSMUSG00000044950 | 70802 | -0.205781017562654 | 2.82385152943579 | -1.86295373279926 | 0.0845249673349 | 0.142892424673404 |
| Fbxl6 | ENSMUSG00000022559 | 30840 | 0.361118917083035 | 1.33144873505118 | 1.86279436476736 | 0.0845487417554 | 0.14291051432336 |
| Plcb4 | ENSMUSG00000039943 | 18798 | 0.163734178230257 | 5.10996696064156 | 1.8623651908533 | 0.0846127956099 | 0.142996671121326 |
| Rps6ka4 | ENSMUSG00000024952 | 56613 | 0.163689383767073 | 3.55305823851427 | 1.86218102443486 | 0.0846402957019 | 0.143021034521258 |
| Fam76b | ENSMUSG00000037808 | 72826 | 0.175364687449597 | 3.09349916869368 | 1.86195483732519 | 0.0846740814251 | 0.143056009900838 |
| Rchy1 | ENSMUSG00000029397 | 68098 | 0.166501383731634 | 4.16676353888123 | 1.86178356025732 | 0.0846996732817 | 0.14307713315449 |
| Ofd1 | ENSMUSG00000040586 | 237222 | -0.233950265516099 | 3.12232591318773 | -1.86124276600365 | 0.0847805233589 | 0.143191579239284 |
| Atg4b | ENSMUSG00000026280 | 66615 | -0.167269762494846 | 3.12179667637748 | -1.86056427961075 | 0.0848820570833 | 0.143340918581443 |
| Etaa1 | ENSMUSG00000016984 | 68145 | -0.277370299155114 | 2.28187817306609 | -1.85953843855245 | 0.0850357795463 | 0.143578329660296 |
| Rangap1 | ENSMUSG00000022391 | 19387 | 0.174823268281079 | 4.18600358567221 | 1.8594057100407 | 0.0850556872471 | 0.143589763289366 |
| Klhl38 | ENSMUSG00000022357 | 268807 | -0.431737398387819 | 3.98473597970589 | -1.85882452811258 | 0.0851429070758 | 0.14371481118682 |
| Oxsm | ENSMUSG00000021786 | 71147 | -0.189709548145196 | 2.94804949176713 | -1.85811621773881 | 0.0852493143761 | 0.143872202477294 |
| Ssbp1 | ENSMUSG00000029911 | 381760 | -0.170348531259596 | 3.66925345437902 | -1.85744331192006 | 0.0853505138659 | 0.144014599879683 |
| Tmem60 | ENSMUSG00000045435 | 212090 | -0.266085404480627 | 1.55074854543811 | -1.85738000960267 | 0.0853600395773 | 0.144014599879683 |
| Aebp2 | ENSMUSG00000030232 | 11569 | 0.142157367228323 | 4.9665604449583 | 1.85663700741657 | 0.0854719178731 | 0.144181100967807 |
| Osbpl11 | ENSMUSG00000022807 | 106326 | -0.138003421125796 | 4.06718168442308 | -1.85649598895447 | 0.0854931667691 | 0.144194693095708 |
| Gas5 | ENSMUSG00000053332 | 14455 | 0.186209779106405 | 3.92023498989344 | 1.85633109258982 | 0.0855180196590 | 0.144205952036369 |
| Kdm6b | ENSMUSG00000018476 | 216850 | -0.131474744478276 | 5.34436591637125 | -1.85627664737755 | 0.0855262269739 | 0.144205952036369 |
| Zmym5 | ENSMUSG00000040123 | 219105 | -0.199982246263678 | 3.88403077612591 | -1.8561545824371 | 0.0855446301661 | 0.144214736635758 |
| Sp1 | ENSMUSG00000001280 | 20683 | 0.129676543748679 | 4.86386737345406 | 1.85538735947169 | 0.0856603826282 | 0.144329119039636 |
| Ccny | ENSMUSG00000024286 | 67974 | -0.115002590328839 | 5.64257756540797 | -1.85538134332843 | 0.0856612908526 | 0.144329119039636 |
| C1d | ENSMUSG00000000581 | 57316 | -0.197403836949511 | 3.64328527019062 | -1.85536623867193 | 0.0856635711586 | 0.144329119039636 |
| Gm31517 | ENSMUSG00000114538 | 102633770 | -1.26303053879987 | -1.23691999613666 | -1.85535482953661 | 0.0856652935988 | 0.144329119039636 |
| Nsmce2 | ENSMUSG00000059586 | 68501 | -0.164591839121716 | 2.96343783068484 | -1.85512430328626 | 0.0857001028870 | 0.144365514664551 |
| Hist1h4h | ENSMUSG00000060981 | 69386 | -0.164393881712726 | 3.78766110283709 | -1.85401625612467 | 0.0858675948946 | 0.144625374390331 |
| Phka2 | ENSMUSG00000031295 | 110094 | 0.188796126757625 | 2.80195884686967 | 1.85391207906547 | 0.0858833574030 | 0.144629637925294 |
| Lrrc8d | ENSMUSG00000046079 | 231549 | 0.195071169980463 | 2.98649902722868 | 1.85346489838463 | 0.0859510476528 | 0.144721334241151 |
| Psap | ENSMUSG00000004207 | 19156 | -0.122283367133699 | 7.92346162459044 | -1.85263569956977 | 0.0860766914871 | 0.144910567479686 |
| Il10rb | ENSMUSG00000022969 | 16155 | 0.248773169975598 | 2.84596881255275 | 1.85233766575738 | 0.0861218912349 | 0.144964335246265 |
| Fis1 | ENSMUSG00000019054 | 66437 | -0.207066479578355 | 4.45792884689227 | -1.8520031561057 | 0.0861726483365 | 0.145027439409907 |

| | | | | | | | |
|---|---|---|---|---|---|---|---|
| Vamp1 | ENSMUSG00000030337 | 22317 | 0.260541002616486 | 3.25867764174028 | 1.85184927378026 | 0.0861960068372 | 0.145044419756429 |
| Eml4 | ENSMUSG00000032624 | 78798 | 0.242735524832362 | 2.26872008442758 | 1.85164414646922 | 0.0862271529062 | 0.145074497216885 |
| Stk19 | ENSMUSG00000061207 | 54402 | 0.290960993500581 | 1.48065241414097 | 1.85154995949563 | 0.0862414574358 | 0.145076234415442 |
| Mthfd2 | ENSMUSG00000005667 | 17768 | 0.279680002253615 | 1.91055655336545 | 1.85125090595224 | 0.0862868899730 | 0.145121610342429 |
| Olfr102 | ENSMUSG00000091531 | 258218 | -1.04630643022525 | 1.40130664658769 | -1.85119764471245 | 0.0862949837366 | 0.145121610342429 |
| Adipor1 | ENSMUSG00000026457 | 72674 | -0.133334898734455 | 7.04182682437425 | -1.85109295014991 | 0.0863108954780 | 0.145126041912078 |
| Anp32e | ENSMUSG00000015749 | 66471 | -0.14538617531907 | 5.29678711351489 | -1.85074639034375 | 0.0863635853341 | 0.145192302566568 |
| Ccnh | ENSMUSG00000021548 | 66671 | -0.157215531075763 | 3.18957478659431 | -1.85013993242191 | 0.0864558589007 | 0.145325079754557 |
| Zfp706 | ENSMUSG00000062397 | 68036 | -0.156354181127544 | 5.11337281974208 | -1.84903090400546 | 0.0866248287743 | 0.145586716379607 |
| Nipal2 | ENSMUSG00000038879 | 223473 | -0.767605563527036 | 1.20869238305815 | -1.84879505145004 | 0.0866608011838 | 0.145624783664922 |
| Ppl | ENSMUSG00000039457 | 19041 | -0.375453310971455 | 1.34784581292182 | -1.8484471804485 | 0.0867138832511 | 0.145676800664421 |
| Elob | ENSMUSG00000055839 | 67673 | -0.134811945273955 | 4.97189038213867 | -1.84835979024504 | 0.0867272228250 | 0.145676800664421 |
| Phykpl | ENSMUSG00000020359 | 72947 | -0.21669925449388 | 3.46701921244077 | -1.8483302187787 | 0.0867317371442 | 0.145676800664421 |
| Olfr365 | ENSMUSG00000059429 | 258656 | -0.865453273729224 | 2.82270281326695 | -1.84778103771111 | 0.0868156124058 | 0.145795277186643 |
| Rpap2 | ENSMUSG00000033773 | 231571 | 0.222808463362246 | 2.7895889551917 | 1.84757186080223 | 0.0868475787281 | 0.145826556540314 |
| Rogdi | ENSMUSG00000022540 | 66049 | 0.273929438533093 | 1.81229495556383 | 1.84661144401147 | 0.0869944850829 | 0.146050793494269 |
| Adamts1 | ENSMUSG00000022893 | 11504 | -0.389698199198701 | 5.43282433697801 | -1.84629867577274 | 0.0870423746415 | 0.146108752642261 |
| Tmem256 | ENSMUSG00000070394 | 69186 | -0.249716217431105 | 3.83207225898269 | -1.84613258104176 | 0.0870678158961 | 0.14612901820373 |
| Sec16a | ENSMUSG00000026924 | 227648 | 0.093776181102446 | 5.70021004230187 | 1.84441212794801 | 0.0873317366167 | 0.14654946468487 |
| Srp19 | ENSMUSG00000014504 | 66384 | -0.134308603833922 | 4.31292505161136 | -1.84412784062451 | 0.0873754159794 | 0.146600256649403 |
| Uchl5 | ENSMUSG00000018189 | 56207 | -0.186046251724445 | 3.58065751278175 | -1.84381499083177 | 0.0874235065169 | 0.146658433047408 |
| Terf2ip | ENSMUSG00000033430 | 57321 | -0.1468861757514 | 3.9051215971932 | -1.84295725614313 | 0.0875554775118 | 0.146857284744811 |
| Reck | ENSMUSG00000028476 | 53614 | 0.297621872283393 | 1.80676094800032 | 1.84241661421277 | 0.0876387525874 | 0.146974410023504 |
| Cnot2 | ENSMUSG00000020166 | 72068 | -0.116458356433505 | 4.18105111289577 | -1.84216356022878 | 0.0876777549365 | 0.147017263262994 |
| Zdhhc9 | ENSMUSG00000036985 | 208884 | 0.278988740331634 | 2.51374481840161 | 1.84167379407836 | 0.0877532852275 | 0.147121343684415 |
| Selenoh | ENSMUSG00000076437 | 72657 | -0.29873558046773 | 1.38129261555218 | -1.8411143845978 | 0.0878396272164 | 0.147243515580896 |
| Ncapd2 | ENSMUSG00000038252 | 68298 | 0.186948065620661 | 3.03696298671836 | 1.84003869349806 | 0.0880058691165 | 0.147499563832054 |
| A230050P20Rik | ENSMUSG00000038884 | 319278 | 0.363271923715953 | 1.36956417317691 | 1.83940358249019 | 0.0881041545126 | 0.147641654603407 |
| Nisch | ENSMUSG00000021910 | 64652 | 0.140083114318757 | 6.82383218248584 | 1.83874344257095 | 0.0882064176827 | 0.147790366598725 |
| Nr1h2 | ENSMUSG00000060601 | 22260 | -0.113254537322534 | 4.74943141589602 | -1.83764496206045 | 0.0883768210591 | 0.148053184827152 |
| Gm16576 | ENSMUSG00000087543 | 100504191 | -0.223252268816687 | 2.18797811706789 | -1.83667860595363 | 0.0885269727685 | 0.148259930058936 |
| Tbck | ENSMUSG00000028030 | 271981 | 0.121036355792783 | 3.76122401442002 | 1.83667612078546 | 0.0885273592072 | 0.148259930058936 |
| Ndfip2 | ENSMUSG00000053253 | 76273 | 0.185658760558964 | 2.79423583122266 | 1.83598100926163 | 0.0886355071293 | 0.148418310114948 |
| Ormdl3 | ENSMUSG00000038150 | 66612 | 0.153257642932667 | 3.63335100530332 | 1.83570735211829 | 0.0886781162056 | 0.148466915031974 |
| Rsf1 | ENSMUSG00000035623 | 233532 | 0.159641329514639 | 5.13577781754362 | 1.83412651314587 | 0.0889246169946 | 0.148856812417911 |
| Sec24b | ENSMUSG00000001052 | 99683 | 0.101686365116492 | 4.63568922493876 | 1.83402240880343 | 0.0889408712630 | 0.148861225505492 |
| Bcl2l2 | ENSMUSG00000089682 | 12050 | -0.126038482628827 | 4.19989867694082 | -1.83319819561918 | 0.0890696558093 | 0.14905395095714 |
| Tmem243 | ENSMUSG00000079659 | 652925 | -0.250600796701949 | 2.26093408332286 | -1.83294136365981 | 0.0891098203679 | 0.149098338656399 |
| Hdac11 | ENSMUSG00000034245 | 232232 | 0.327431577906014 | 2.00858752638641 | 1.83234149889325 | 0.0892036932769 | 0.149232563699166 |
| Eef2 | ENSMUSG00000034994 | 13629 | -0.152221536105504 | 9.54285029646748 | -1.83198511466387 | 0.0892595058913 | 0.149303084758709 |
| Sc5d | ENSMUSG00000032018 | 235293 | -0.200431790393229 | 3.2954101841239 | -1.83156353425351 | 0.0893255691950 | 0.149383868940784 |
| Engase | ENSMUSG00000033857 | 217364 | 0.379310699003244 | 1.38974402676388 | 1.83150251912063 | 0.0893351341383 | 0.149383868940784 |
| Osbpl7 | ENSMUSG00000038534 | 71240 | 0.403145653018663 | 1.24391931971092 | 1.83074395872676 | 0.0894541254255 | 0.149559964060323 |
| Ppp1r10 | ENSMUSG00000039220 | 52040 | -0.251217615961447 | 4.39181727458415 | -1.83046765236873 | 0.0894975034269 | 0.149604741486052 |
| Pygb | ENSMUSG00000033059 | 110078 | 0.177590218341974 | 3.07128939929898 | 1.83039902671385 | 0.0895082800584 | 0.149604741486052 |
| U2af1 | ENSMUSG00000061613 | 108121 | -0.17699952641329 | 3.28984159018348 | -1.83027112034672 | 0.0895283689442 | 0.149615441206216 |
| Snora81 | ENSMUSG00000087935 | 100217420 | 1.9168688586872 | -0.397675382669002 | 1.83014805493993 | 0.0895477013292 | 0.149624873621187 |
| Krr1 | ENSMUSG00000063334 | 52705 | -0.118737372771752 | 4.26647515649175 | -1.82964554550508 | 0.0896266795622 | 0.149733949915116 |
| Sin3b | ENSMUSG00000031622 | 20467 | 0.121680571797732 | 4.34930636055353 | 1.82941237190006 | 0.0896633481265 | 0.149772319433339 |
| Park7 | ENSMUSG00000028964 | 57320 | -0.144944497174748 | 6.50454480058657 | -1.82874174540651 | 0.0897688849377 | 0.149925696142726 |
| Nfic | ENSMUSG00000055053 | 18029 | 0.221275427667535 | 8.11546831458519 | 1.82799651046786 | 0.0898862935679 | 0.150098850441675 |
| Meox1 | ENSMUSG00000001493 | 17285 | 0.484148369190389 | 3.00154888771619 | 1.82763496710972 | 0.0899433027878 | 0.150171107801057 |
| Plekha7 | ENSMUSG00000045659 | 233765 | 0.414711504227399 | 1.08047964699349 | 1.82736767104536 | 0.0899854716575 | 0.15020323800657 |
| Pank2 | ENSMUSG00000037514 | 74450 | 0.158737032814847 | 3.56004465474624 | 1.82733879086618 | 0.0899900288816 | 0.15020323800657 |
| Rbm45 | ENSMUSG00000042369 | 241490 | 0.270040332583683 | 2.61068098358785 | 1.82678061288225 | 0.0900781487031 | 0.150327365415976 |
| Ly6e | ENSMUSG00000022587 | 17069 | -0.258382996129983 | 5.16167111399406 | -1.82616536576773 | 0.0901753676670 | 0.150466637760381 |
| Clip2 | ENSMUSG00000063146 | 269713 | 0.274001352153485 | 2.86026124995218 | 1.82595386423947 | 0.0902088100353 | 0.150499466193004 |
| Cacna1c | ENSMUSG00000051331 | 12288 | 0.302921070923199 | 2.79679634999871 | 1.82507069788759 | 0.0903485753959 | 0.15070964102753 |
| Fam172a | ENSMUSG00000064138 | 68675 | 0.187603331666752 | 3.24489326216379 | 1.82478556754903 | 0.0903937400804 | 0.150758622255206 |
| Ppargc1b | ENSMUSG00000033871 | 170826 | 0.268268072312368 | 4.80932736726479 | 1.82471120324308 | 0.0904055227227 | 0.150758622255206 |
| Anapc16 | ENSMUSG00000020107 | 52717 | -0.165824757227957 | 3.81711475169814 | -1.82398520928707 | 0.0905206250634 | 0.150927540050183 |
| 4930533N22Rik | ENSMUSG00000108803 | 75205 | -0.821517252609682 | 2.81871920352406 | -1.82314101058811 | 0.0906546332254 | 0.151127923713242 |
| Sptbn1 | ENSMUSG00000020315 | 20742 | -0.169805117367332 | 8.85310384319252 | -1.82259773644914 | 0.0907409665736 | 0.15122788396763 |
| Ctps2 | ENSMUSG00000031360 | 55936 | 0.227558442595305 | 1.85087224426497 | 1.82258957378461 | 0.0907422642890 | 0.15122788396763 |
| Zfand1 | ENSMUSG00000039795 | 66361 | -0.274477361047011 | 2.27629957573076 | -1.82187749193038 | 0.0908555363765 | 0.151393577459083 |
| Cln5 | ENSMUSG00000022125 | 211286 | -0.209872353484127 | 2.91287288559805 | -1.82122205335431 | 0.0909599100500 | 0.151544395177183 |
| Pgm1 | ENSMUSG00000029171 | 66681 | 0.316393861915886 | 1.92256746259096 | 1.82018756126563 | 0.0911248636784 | 0.151786918041184 |
| Fbxw4 | ENSMUSG00000040913 | 30838 | -0.160057995518801 | 3.00401737361618 | -1.82013501925315 | 0.0911332488431 | 0.151786918041184 |
| Kctd15 | ENSMUSG00000030499 | 233107 | -0.274473093271385 | 3.30326853573982 | -1.81964881738144 | 0.0912108744475 | 0.151893063465312 |
| Mgrn1 | ENSMUSG00000022517 | 17237 | -0.15065482817675 | 5.84011438446665 | -1.81819864335342 | 0.0914427568772 | 0.152256020628345 |
| Myom1 | ENSMUSG00000024049 | 17929 | -0.203831477522584 | 10.3240593476998 | -1.8176516843707 | 0.0915303524517 | 0.152378660165939 |
| Gtf2h2 | ENSMUSG00000021639 | 23894 | -0.252171635124209 | 2.61212280091562 | -1.81717677541521 | 0.0916064701208 | 0.152464313457699 |
| Zbtb21 | ENSMUSG00000046962 | 114565 | 0.19675750573293 | 3.52316313407107 | 1.81715664291353 | 0.0916096981785 | 0.152464313457699 |
| Cdc42 | ENSMUSG00000006699 | 12540 | 0.120969014103212 | 5.86757920996305 | 1.81669873272452 | 0.0916831472808 | 0.152563325152601 |
| Pdzd2 | ENSMUSG00000022197 | 68070 | -0.183252330834145 | 5.00606329507411 | -1.81652593009783 | 0.0917108786346 | 0.152586242672104 |
| Zfp30 | ENSMUSG00000047473 | 22693 | 0.241789386007842 | 1.88168757013485 | 1.81634213373174 | 0.0917403825021 | 0.152612101830955 |
| Wdr12 | ENSMUSG00000026019 | 57750 | -0.137116621785923 | 3.55799984797349 | -1.81621936806708 | 0.0917600941629 | 0.152621666052289 |
| Ighmbp2 | ENSMUSG00000024831 | 20589 | 0.298185033227806 | 1.59066609185829 | 1.8158006860863 | 0.0918273476369 | 0.152693867550809 |
| Top1mt | ENSMUSG00000000934 | 72960 | -0.235858134868197 | 2.867852803034 | -1.81577520988009 | 0.0918314413392 | 0.152693867550809 |
| Dstn | ENSMUSG00000015932 | 56431 | 0.206241047499166 | 4.65982333162837 | 1.81443663273265 | 0.0920467635233 | 0.153028619326851 |
| Matn2 | ENSMUSG00000022324 | 17181 | 0.297129977152729 | 2.7150729429894 | 1.81424850209652 | 0.0920770622116 | 0.153055712748782 |
| Polr2i | ENSMUSG00000019738 | 69920 | -0.280689940406172 | 2.19337743164623 | -1.81394210846587 | 0.0921264264136 | 0.153103260298527 |
| Itprid2 | ENSMUSG00000027007 | 70599 | -0.149932835956333 | 4.48331233689311 | -1.81389710488268 | 0.0921336791001 | 0.153103260298527 |
| Fgd6 | ENSMUSG00000020021 | 13998 | 0.226935082375406 | 2.23624545250762 | 1.81317755176716 | 0.0922497102261 | 0.153272774355082 |
| Nmt1 | ENSMUSG00000020936 | 18107 | -0.10175281030007 | 5.34713755020866 | -1.81216179335235 | 0.0924137282913 | 0.153521955008006 |
| 2310030G06Rik | ENSMUSG00000032062 | 66952 | -0.401811720177399 | 1.79298936122947 | -1.81198923275158 | 0.0924416181625 | 0.15354495185156 |
| Mfsd6 | ENSMUSG00000041439 | 98682 | 0.189475100486816 | 3.56319430859839 | 1.81178464256974 | 0.0924746945253 | 0.153576555128601 |
| Cep120 | ENSMUSG00000048799 | 225523 | -0.169593605497652 | 4.90681134640207 | -1.81127841654126 | 0.0925565822550 | 0.153689199548896 |
| Pitpna | ENSMUSG00000017781 | 18738 | 0.132021511495872 | 4.94708241714808 | 1.81115274078101 | 0.0925769217630 | 0.153699625120292 |
| Dok7 | ENSMUSG00000044716 | 231134 | 0.244768448461243 | 3.40684874690575 | 1.81078801161344 | 0.0926359725825 | 0.153774307714475 |
| Cdk2 | ENSMUSG00000025358 | 12566 | 0.263609522183519 | 1.32892527018072 | 1.81041094932878 | 0.0926970555832 | 0.153852340507256 |
| Gpatch11 | ENSMUSG00000050668 | 53951 | -0.225717539790609 | 3.34575067137663 | -1.80921492110956 | 0.0928910470805 | 0.154150908704581 |
| Hook2 | ENSMUSG00000052566 | 170833 | 0.36557261861384 | 1.71889607232296 | 1.80899809745985 | 0.0929262539423 | 0.154181170810191 |
| Zfp46 | ENSMUSG00000051351 | 22704 | -0.218823108827991 | 5.42067189172637 | -1.80890422689168 | 0.0929414999286 | 0.154181170810191 |
| Zfp958 | ENSMUSG00000058748 | 233987 | -0.239005493162734 | 1.63234182877741 | -1.80881813847062 | 0.0929554839436 | 0.154181170810191 |
| Schip1 | ENSMUSG00000027777 | 30953 | -0.126839326813404 | 5.51475351183417 | -1.80875523672815 | 0.0929657027545 | 0.154181170810191 |
| Lin7c | ENSMUSG00000027162 | 22343 | -0.120003720743291 | 4.45145419672622 | -1.80842301881093 | 0.0930196904498 | 0.154232449976391 |
| Wdr59 | ENSMUSG00000031959 | 319481 | 0.224901419484551 | 3.53769168143442 | 1.80833428304348 | 0.0930341153642 | 0.154232449976391 |
| Dclre1b | ENSMUSG00000027845 | 140917 | 0.405768220667208 | 1.074054091544 | 1.80830453808677 | 0.0930389511613 | 0.154232449976391 |
| Uros | ENSMUSG00000030979 | 22276 | -0.180348514948662 | 2.72153835496952 | -1.80715473292098 | 0.0932260534538 | 0.154519179727648 |
| Morc3 | ENSMUSG00000039456 | 338467 | 0.187783326582261 | 3.82272754541532 | 1.80568921173933 | 0.0934650181636 | 0.154886900192005 |
| Nr1d2 | ENSMUSG00000021775 | 353187 | -0.268194187317066 | 5.93253711903206 | -1.80553883975696 | 0.0934895684313 | 0.154886900192005 |
| Ddx31 | ENSMUSG00000026806 | 227674 | 0.254111876580884 | 2.10272300636316 | 1.80553363207059 | 0.0934904187601 | 0.154886900192005 |
| Rrp36 | ENSMUSG00000023971 | 224823 | 0.193169930978183 | 2.41572878563353 | 1.80540408938888 | 0.0935115731538 | 0.154898470396185 |
| Gli2 | ENSMUSG00000048402 | 14633 | 0.540321519594007 | 1.24194020190735 | 1.8049643756787 | 0.0935834105559 | 0.154993978907168 |
| Galnt2 | ENSMUSG00000089704 | 108148 | 0.11436437200306 | 5.97418939742449 | 1.80427488837877 | 0.0936961535318 | 0.15515719652429 |
| Acvrl1 | ENSMUSG00000000530 | 11482 | -0.27645911054666 | 3.60951127458149 | -1.803501140886 | 0.0938228189771 | 0.155343416273755 |
| Parp11 | ENSMUSG00000037997 | 101187 | 0.245329355185142 | 1.78400177906855 | 1.80318424927407 | 0.0938747394933 | 0.155405842405207 |
| Pdpk1 | ENSMUSG00000024122 | 18607 | 0.126636391647345 | 5.70360573602789 | 1.80273002014213 | 0.0939492065804 | 0.155505568917445 |
| Yae1d1 | ENSMUSG00000075054 | 67008 | -0.130782377529997 | 3.7418324681156 | -1.80227704584591 | 0.0940235204881 | 0.155605011726843 |
| Snx1 | ENSMUSG00000032382 | 56440 | 0.197982393880343 | 3.33503184871216 | 1.80178271638681 | 0.0941046789333 | 0.155693364542545 |
| Cnot10 | ENSMUSG00000056167 | 78893 | -0.176349814889196 | 3.69405911885871 | -1.80177836393724 | 0.0941053937912 | 0.155693364542545 |
| F630040K05Rik | ENSMUSG00000097727 | 433111 | -0.681030209151584 | 1.44246862535636 | -1.801175630803 | 0.0942044351432 | 0.155833638097835 |
| Cdyl | ENSMUSG00000059288 | 12593 | 0.171463478377682 | 2.83743972578863 | 1.80096350906645 | 0.0942393132104 | 0.155867745907659 |
| Bcas3 | ENSMUSG00000059439 | 192197 | -0.135751100047813 | 4.25765364256299 | -1.80031398174185 | 0.0943461833127 | 0.15602089709398 |
| Tm9sf1 | ENSMUSG00000002320 | 74140 | 0.168875081189957 | 3.72194194945448 | 1.79999391478683 | 0.0943988853797 | 0.156071581900686 |

| | | | | | | | |
|---|---|---|---|---|---|---|---|
| Gm20735 | ENSMUSG00000110656 | 319574 | -0.782463407784514 | 0.532206763863069 | -1.799954432698 | 0.0944053883018 | 0.156071581900686 |
| Svbp | ENSMUSG00000028643 | 69216 | 0.270148029491906 | 1.90357672925078 | 1.79944474958904 | 0.0944893718916 | 0.156186802381386 |
| Aar2 | ENSMUSG00000027628 | 68295 | -0.135530530953319 | 3.66485394545413 | -1.7991373390772 | 0.0945400580350 | 0.156246957118378 |
| Tpst1 | ENSMUSG00000034118 | 22021 | 0.187023342418009 | 2.66240089881599 | 1.7987257302073 | 0.0946079625115 | 0.156335546215271 |
| Pan3 | ENSMUSG00000029647 | 72587 | 0.192196950222097 | 3.74997559954304 | 1.7975723942509 | 0.0947984641231 | 0.156626664348547 |
| Pgrmc2 | ENSMUSG00000049940 | 70804 | 0.150089400169053 | 3.68012578805373 | 1.7971489779064 | 0.0948684875486 | 0.156718669698413 |
| Rnf41 | ENSMUSG00000025373 | 67588 | 0.184411041639786 | 3.30220795326263 | 1.79671179181498 | 0.0949408366097 | 0.156814488513392 |
| Ptprt | ENSMUSG00000053141 | 19281 | -0.611586720575136 | 1.69282390731977 | -1.79561146395236 | 0.0951231456408 | 0.157091872639337 |
| Tspan31 | ENSMUSG00000006736 | 67125 | 0.178714915344738 | 3.69431655397204 | 1.79515666077837 | 0.0951985913966 | 0.157192719419414 |
| Dbhos | ENSMUSG00000085008 | 329371 | -0.838607133437324 | 0.263870373019163 | -1.79501760506454 | 0.0952216695319 | 0.157207078938714 |
| Xpo7 | ENSMUSG00000022100 | 65246 | -0.128980011972492 | 4.88840001628712 | -1.7941275174448 | 0.0953695096063 | 0.157427379871227 |
| Snx24 | ENSMUSG00000024535 | 69226 | 0.197081672847701 | 2.70351537391609 | 1.79361431954128 | 0.0954548428031 | 0.157544449144038 |
| Relch | ENSMUSG00000026319 | 227446 | 0.161177336253045 | 3.4028077590527 | 1.79301211686242 | 0.0955550623201 | 0.157686048644596 |
| Slu7 | ENSMUSG00000020409 | 193116 | -0.113808793407533 | 4.94296621345322 | -1.79273184334135 | 0.0956017378751 | 0.15773925988126 |
| Setd1a | ENSMUSG00000042308 | 233904 | 0.168078573615077 | 4.34880612226118 | 1.7916231730329 | 0.0957865704489 | 0.158020374521142 |
| Ubl4a | ENSMUSG00000015290 | 27643 | -0.12384716078645 | 4.59621933938103 | -1.7913526124474 | 0.0958317254473 | 0.1580710111461 |
| Adprh | ENSMUSG00000002844 | 11544 | 0.131951026395497 | 4.0080379691761 | 1.79007932647408 | 0.0960444844541 | 0.158398047611436 |
| Heca | ENSMUSG00000039879 | 380629 | -0.148019032527168 | 4.45667755443421 | -1.78978697913697 | 0.0960933934719 | 0.158454802238976 |
| Cd93 | ENSMUSG00000027435 | 17064 | -0.202393161978784 | 5.87700313101715 | -1.78964670055581 | 0.0961168696284 | 0.158469608131031 |
| 0610030E20Rik | ENSMUSG00000058706 | 68364 | -0.188566019784709 | 3.1648446440727 | -1.78908372645652 | 0.0962111369338 | 0.158601106593941 |
| Cc2d1a | ENSMUSG00000036686 | 212139 | 0.146056713745379 | 3.50860937887248 | 1.78890954258514 | 0.0962403198738 | 0.158625291999628 |
| Zbtb22 | ENSMUSG00000051390 | 81630 | 0.208576288489235 | 2.86588903933995 | 1.78783600195021 | 0.0964203561666 | 0.158898072253484 |
| Borcs5 | ENSMUSG00000042992 | 67774 | -0.212880869406366 | 2.88333134522925 | -1.78751814627546 | 0.0964737191676 | 0.158962047666733 |
| Vmn2r112 | ENSMUSG00000094921 | 628185 | -0.801486647235551 | 1.42580593858061 | -1.78722808398322 | 0.0965224390590 | 0.159018354386472 |
| Hspb11 | ENSMUSG00000063172 | 72938 | -0.508411780378229 | 1.95588046850875 | -1.7868766414357 | 0.0965814979691 | 0.159091674849464 |
| Hexim1 | ENSMUSG00000048878 | 192231 | 0.213798389878544 | 4.48274376385201 | 1.78626012116584 | 0.0966851802189 | 0.15923846692385 |
| Usp21 | ENSMUSG00000053483 | 30941 | -0.195779740143506 | 2.69315156543169 | -1.78615595205332 | 0.0967027084728 | 0.159243342319515 |
| Map3k11 | ENSMUSG00000004054 | 26403 | -0.146499433754971 | 3.54380458017272 | -1.78424281745519 | 0.0970251302686 | 0.159750218250651 |
| Synpo | ENSMUSG00000043079 | 104027 | -0.121616712580539 | 7.40328669665324 | -1.78376404494556 | 0.0971059677547 | 0.159859236976908 |
| Tchp | ENSMUSG00000002486 | 77832 | 0.318923177569398 | 2.0620993613618 | 1.78356609308258 | 0.0971394081057 | 0.159890207800657 |
| Wdr36 | ENSMUSG00000038299 | 225348 | 0.135914063938515 | 3.69377512362975 | 1.78267273362537 | 0.0972904524809 | 0.160114714855322 |
| Madd | ENSMUSG00000040687 | 228355 | 0.271019036861804 | 2.7039729549224 | 1.78222331522889 | 0.0973665167480 | 0.160215775187894 |
| Prkd3 | ENSMUSG00000024070 | 75292 | 0.133402430574989 | 3.9628095937087 | 1.78084741194328 | 0.0975997183046 | 0.160575334254727 |
| Tor1a | ENSMUSG00000026849 | 30931 | 0.249518656188164 | 1.9124450741183 | 1.77817141364809 | 0.0980546969681 | 0.161289585504672 |
| Gm3055 | ENSMUSG00000094622 | 100040944 | -0.445889244610524 | 0.960644874035665 | -1.77812057008721 | 0.0980633597350 | 0.161289585504672 |
| 2310022A10Rik | ENSMUSG00000049643 | 66367 | -0.132083632639898 | 5.39606434716149 | -1.77767956492672 | 0.0981385271248 | 0.161388933353693 |
| Sac3d1 | ENSMUSG00000024790 | 66406 | 0.301766662278342 | 1.5411258725008 | 1.77752997803147 | 0.0981640351945 | 0.161406598783406 |
| Zfp655 | ENSMUSG00000007812 | 72611 | -0.230381742816759 | 3.85765687967285 | -1.77742458889707 | 0.0981820100526 | 0.161411874249639 |
| Ubr7 | ENSMUSG00000041712 | 66622 | -0.170058947456225 | 3.27910239849324 | -1.77718561875739 | 0.0982227789396 | 0.161454616028405 |
| Mgat4a | ENSMUSG00000026110 | 269181 | 0.253508368325139 | 2.80486453093596 | 1.77649366191361 | 0.0983409134178 | 0.161624496251743 |
| Cpped1 | ENSMUSG00000065979 | 223978 | -0.104104159997441 | 4.44081833901665 | -1.77505954361865 | 0.0985861560826 | 0.162003197856174 |
| Sirt4 | ENSMUSG00000029524 | 75387 | -0.277097093691084 | 1.63542949169864 | -1.77470706850105 | 0.0986465146165 | 0.162078017627097 |
| Zfp955a | ENSMUSG00000094441 | 77652 | -0.173666250435673 | 3.03886939145364 | -1.77424127234305 | 0.0987263288804 | 0.162184776223697 |
| Use1 | ENSMUSG00000002395 | 67023 | -0.153165499778548 | 5.4500776824308 | -1.77383169840542 | 0.0987965568991 | 0.162275757094644 |
| Riok3 | ENSMUSG00000024404 | 66878 | 0.133574807601871 | 6.24718119687095 | 1.77368299909815 | 0.0988220647692 | 0.162293267727342 |
| Fbxo8 | ENSMUSG00000038206 | 50753 | -0.207284365386099 | 3.61520739158806 | -1.7730141796058 | 0.0989368664424 | 0.162457396286934 |
| Urgcp | ENSMUSG00000049680 | 72046 | -0.153694230851402 | 5.13862534624112 | -1.77245269656666 | 0.0990333355482 | 0.162591377422304 |
| Mme | ENSMUSG00000027820 | 17380 | 0.309474674833082 | 2.19628536192466 | 1.77231621087366 | 0.0990567979574 | 0.162597516755601 |
| Mapkapk5 | ENSMUSG00000029454 | 17165 | -0.146710018653534 | 5.39869432370029 | -1.77223938554116 | 0.0990700066976 | 0.162597516755601 |
| Plin1 | ENSMUSG00000030546 | 103968 | 0.757175440496084 | 1.41153941633905 | 1.77217138388194 | 0.0990816996714 | 0.162597516755601 |
| 9930021J03Rik | ENSMUSG00000046138 | 240613 | -0.145380563131195 | 4.66461254321404 | -1.7717966937428 | 0.0991461501359 | 0.1626788602726 8 |
| Rnf185 | ENSMUSG00000020448 | 193670 | -0.170431869444862 | 3.53580952049929 | -1.7716834853797 | 0.0991656304404 | 0.162686403473588 |
| Trnau1ap | ENSMUSG00000028898 | 71787 | 0.200904389461111 | 2.76858522997884 | 1.7710019695719 | 0.0992829740327 | 0.162854470160923 |
| Pik3ip1 | ENSMUSG00000034614 | 216505 | 0.324094887251496 | 2.84233216873368 | 1.77086203124278 | 0.0993070839229 | 0.162869577548589 |
| Sarnp | ENSMUSG00000078427 | 66118 | -0.147810153944671 | 4.66377176025455 | -1.77005044834923 | 0.0994470136801 | 0.163074603441011 |
| Fchsd2 | ENSMUSG00000030691 | 207278 | 0.15990622975731 | 3.26186872478549 | 1.76960818873658 | 0.0995233399862 | 0.163175285644138 |
| Slc7a14 | ENSMUSG00000069072 | 241919 | -0.434476290614894 | 1.58684875279412 | -1.76945646520606 | 0.0995495368154 | 0.163193759287602 |
| B4galt3 | ENSMUSG00000052423 | 57370 | -0.29346653560176 | 1.63241916938481 | -1.76906483018245 | 0.0996171854731 | 0.163280170101592 |
| Hist1h2bc | ENSMUSG00000018102 | 68024 | -0.120242969124653 | 5.29644441480553 | -1.76842581866443 | 0.0997276521462 | 0.163436726478296 |
| Mrpl20 | ENSMUSG00000029066 | 66448 | -0.147273203507191 | 3.70875084649633 | -1.76722712347748 | 0.0999351655914 | 0.163752255296131 |
| Atp6v1a | ENSMUSG00000052459 | 11964 | 0.220790675349734 | 4.30041095145058 | 1.76710540388437 | 0.0999562586512 | 0.163762269681833 |
| Psmd4 | ENSMUSG00000005625 | 19185 | 0.134783505181294 | 6.79252344843534 | 1.7669413961987 | 0.0999846861632 | 0.163784295586748 |
| Zfp526 | ENSMUSG00000046541 | 210172 | -0.153150765617861 | 2.85780892078822 | -1.76683922565512 | 0.1000023990509 | 0.163788765961363 |
| Olfr731 | ENSMUSG00000048080 | 258360 | -0.642087522907469 | 1.41522831272847 | -1.76673049845062 | 0.1000212517013 | 0.163795101475285 |
| Rida | ENSMUSG00000022323 | 15473 | -0.336335004949567 | 1.22892255928992 | -1.76620283314518 | 0.1001127905697 | 0.163920448429909 |
| Rbm19 | ENSMUSG00000029594 | 74111 | -0.142141588991542 | 3.6656282907868 | -1.76483824236476 | 0.1003498635625 | 0.164284013569352 |
| Brwd1 | ENSMUSG00000022914 | 93871 | 0.218848599667237 | 5.99481543969978 | 1.76472183418132 | 0.1003701104404 | 0.164292554241542 |
| Ipo11 | ENSMUSG00000042590 | 76582 | -0.127006489323572 | 3.80793846973304 | -1.76451294441689 | 0.1004064517366 | 0.164327432839308 |
| Riox2 | ENSMUSG00000022724 | 67014 | -0.175100124140702 | 3.39991425774788 | -1.76415335291351 | 0.1004690385189 | 0.164405248510567 |
| Cops7a | ENSMUSG00000030127 | 26894 | -0.149553981454495 | 5.02804056473102 | -1.76395815364622 | 0.1005030273897 | 0.1644362509201 |
| Zfp318 | ENSMUSG00000015597 | 57908 | 0.149544718066301 | 4.35424148775857 | 1.76253695788286 | 0.1007507995178 | 0.164816969399821 |
| Tmem185a | ENSMUSG00000073139 | 236848 | -0.192874531848535 | 2.58517840244167 | -1.76190760276567 | 0.1008606947850 | 0.164972056665555 |
| Glrx | ENSMUSG00000021591 | 93692 | -0.224729687059286 | 2.7878207678738 | -1.76174414550149 | 0.1008892543842 | 0.164994081339691 |
| Susd2 | ENSMUSG00000006342 | 71733 | -0.425191615145623 | 1.17090802686536 | -1.76147830690451 | 0.1009357175953 | 0.16504537457512 |
| Thoc6 | ENSMUSG00000041319 | 386612 | 0.246416666291927 | 2.26708688841264 | 1.76103995128067 | 0.1010123747703 | 0.165146016843307 |
| Fam171a1 | ENSMUSG00000050530 | 269233 | 0.262679572603838 | 3.35417234335713 | 1.7605548100584 | 0.1010972737749 | 0.16526010163515 |
| Ttc17 | ENSMUSG00000027194 | 74569 | 0.13695029231235 | 4.21095790828523 | 1.76040354395515 | 0.1011237580681 | 0.165278678146387 |
| Slfn9 | ENSMUSG00000069793 | 237886 | -0.547424465644785 | 3.02146454970843 | -1.75991866469142 | 0.1012086942294 | 0.165392769714701 |
| Zfp651 | ENSMUSG00000013419 | 270210 | -0.153706071162165 | 6.30117597544521 | -1.75959776597617 | 0.1012649407672 | 0.165459950303007 |
| Armcx5 | ENSMUSG00000072969 | 494468 | 0.273518745279481 | 1.78785533345373 | 1.75872818645602 | 0.1014174984944 | 0.165684453152445 |
| Ndel1 | ENSMUSG00000018736 | 83431 | -0.122328833175835 | 5.0427062469745 | -1.75856341018084 | 0.1014464295641 | 0.16570695181802 |
| Tmem127 | ENSMUSG00000034850 | 69470 | 0.122897245681673 | 4.91126770144459 | 1.75814706101891 | 0.1015195638914 | 0.1658016364706 |
| Cmah | ENSMUSG00000016756 | 12763 | 0.344595819040313 | 2.97116449921393 | 1.75731335097767 | 0.1016661508575 | 0.166016237679043 |
| Il2rg | ENSMUSG00000031304 | 16186 | 0.401836322295849 | 1.2429406550319 | 1.75696894414122 | 0.1017267609199 | 0.166090399345168 |
| Arih1 | ENSMUSG00000025234 | 23806 | 0.119405640011518 | 6.31700580052305 | 1.7568730103831 | 0.1017436494233 | 0.166093164851602 |
| Topbp1 | ENSMUSG00000032555 | 235559 | 0.148983659718169 | 3.53931242422527 | 1.75662944853528 | 0.1017865380458 | 0.166138367534538 |
| Wdr45 | ENSMUSG00000039382 | 54636 | -0.165975390743658 | 3.18237098880264 | -1.7564758074596 | 0.1018136008331 | 0.166146134173916 |
| Rnf4 | ENSMUSG00000029110 | 19822 | 0.146554428364482 | 4.17469502457722 | 1.75642985974036 | 0.1018216954378 | 0.166146134173916 |
| Grk6 | ENSMUSG00000074886 | 26385 | 0.144350992369185 | 3.18870038937943 | 1.75569785146685 | 0.1019507302440 | 0.166310000133101 |
| Ccdc130 | ENSMUSG00000004994 | 67736 | -0.271165898341652 | 2.25522685208909 | -1.75568754077944 | 0.1019525487962 | 0.166310000133101 |
| Tctn1 | ENSMUSG00000038593 | 654470 | 0.367964293518624 | 1.46398318610994 | 1.75552252859159 | 0.1019816568094 | 0.166332660487024 |
| Reln | ENSMUSG00000042453 | 19699 | -0.332730179847455 | 4.13059532067188 | -1.75475473325556 | 0.1021171921969 | 0.166528871834201 |
| Nit1 | ENSMUSG00000013997 | 27045 | -0.187158871643477 | 2.95323113330379 | -1.75383398754584 | 0.1022799377702 | 0.166769391373278 |
| Eme2 | ENSMUSG00000073436 | 193838 | 0.21578943280545 | 2.6119133623806 | 1.75374447215672 | 0.1022957722378 | 0.166770333532017 |
| Arglu1 | ENSMUSG00000040459 | 234023 | -0.116025995831365 | 4.97641767330083 | -1.75189371851405 | 0.1026236411352 | 0.167279901766909 |
| Sfswap | ENSMUSG00000029439 | 231769 | -0.137567817118282 | 4.13224818507958 | -1.75174967024693 | 0.1026491989184 | 0.167296614489003 |
| Dnal1 | ENSMUSG00000042523 | 105000 | 0.238832616775586 | 2.70037507335126 | 1.75151071975014 | 0.1026916071806 | 0.167340780872314 |
| Adal | ENSMUSG00000027259 | 75894 | -0.199786798340065 | 3.16480106088204 | -1.7510491174262 | 0.1027735750268 | 0.16744938867471 |
| Zfp945 | ENSMUSG00000059142 | 240041 | 0.249082558310744 | 2.54588387286775 | 1.75092686471357 | 0.1027952934452 | 0.16745981410577 |
| Lsm1 | ENSMUSG00000037296 | 67207 | -0.233562126158692 | 1.68190703359865 | -1.7505934838161 | 0.1028545398376 | 0.167531362682887 |
| Naxd | ENSMUSG00000031505 | 69225 | -0.159477987212449 | 3.17075501693828 | -1.7489783344166 | 0.1031420027090 | 0.167974557749143 |
| Rsrc2 | ENSMUSG00000029422 | 208606 | -0.124754893298073 | 5.03039267029396 | -1.74867056341631 | 0.1031968601717 | 0.16803886169179 |
| Ube2s | ENSMUSG00000060860 | 77891 | -0.133677983061681 | 4.31409529741528 | -1.74838804307888 | 0.1032472396728 | 0.168095855952387 |
| Ccdc117 | ENSMUSG00000020482 | 104479 | -0.139414637265409 | 4.14189821317707 | -1.74783670651172 | 0.1033456176683 | 0.16823096749551 |
| Mdga2 | ENSMUSG00000034912 | 320772 | -0.583594617953673 | 2.01035338256334 | -1.74742013241227 | 0.1034200043062 | 0.168308942629985 |
| Zeb1 | ENSMUSG00000024238 | 21417 | 0.0879397632744849 | 5.8932048201519 | 1.74739601019193 | 0.1034243132051 | 0.168308942629985 |
| Tbp | ENSMUSG00000014767 | 21374 | -0.249451596618932 | 2.72134278697346 | -1.74681994400964 | 0.1035272618722 | 0.168451399155341 |
| Pak1ip1 | ENSMUSG00000038683 | 68083 | -0.146373473862816 | 4.33491350554994 | -1.74652965209887 | 0.1035791742521 | 0.168510783412784 |
| Zc3h13 | ENSMUSG00000022000 | 67302 | 0.126071190884855 | 5.40022693493351 | 1.74632593582833 | 0.1036156182264 | 0.168544988516928 |
| Mcph1 | ENSMUSG00000039842 | 244329 | 0.266274394899407 | 2.06914891597605 | 1.74536012342747 | 0.1037885524892 | 0.168801170549088 |
| Csnk2a2 | ENSMUSG00000046707 | 13000 | 0.175419233409135 | 3.3933358057959 | 1.74507331483571 | 0.1038399563948 | 0.168859649412614 |
| Khsrp | ENSMUSG00000007670 | 16549 | 0.103130074125261 | 4.87790838233061 | 1.74490232276119 | 0.1038706135498 | 0.168884378508545 |
| Emc2 | ENSMUSG00000022337 | 66736 | -0.139202877558358 | 5.2383700214765 | -1.7446895662149 | 0.1039087698387 | 0.168921291360441 |
| Anp32b | ENSMUSG00000028333 | 67628 | 0.11580089072604 | 5.61405844858922 | 1.74454977693511 | 0.1039338467493 | 0.168929966148768 |
| Nrp1 | ENSMUSG00000025810 | 18186 | -0.192625460566733 | 5.58929499504229 | -1.74448753339153 | 0.1039450143917 | 0.168929966148768 |
| Adamts15 | ENSMUSG00000033453 | 235130 | 0.402461815064171 | 1.99016561002141 | 1.74437289488001 | 0.1039655854394 | 0.16893828072522 |

| | | | | | | | |
|---|---|---|---|---|---|---|---|
| Lrrc58 | ENSMUSG00000034158 | 320184 | 0.110379012648239 | 6.27692896856055 | 1.74297164167374 | 0.1042173214682 | 0.16929805595533 |
| Zfp11 | ENSMUSG00000051034 | 22648 | -0.289018000277347 | 2.15950023940418 | -1.74296804145622 | 0.1042179689436 | 0.16929805595533 |
| Paxbp1 | ENSMUSG00000022974 | 67367 | 0.166440715536494 | 3.34681848770948 | 1.74266702558166 | 0.1042721172608 | 0.169360849001237 |
| Neurl1b | ENSMUSG00000034413 | 240055 | -0.281993828850383 | 2.64621030675114 | -1.7423666963693 | 0.1043261668638 | 0.16942346307966 |
| Nhlrc3 | ENSMUSG00000042997 | 212114 | 0.206651082966784 | 2.52548402272315 | 1.74197189834496 | 0.1043972555252 | 0.169513725511989 |
| Stard4 | ENSMUSG00000024378 | 170459 | -0.266238018009633 | 1.79353436726952 | -1.74042853303539 | 0.1046755704602 | 0.169940392202708 |
| Nr1d1 | ENSMUSG00000020889 | 217166 | -0.288913919304696 | 6.60183206941803 | -1.74028950567313 | 0.1047006734457 | 0.16994311840192 |
| Adipoq | ENSMUSG00000022878 | 11450 | 0.82120469626278 | 0.684818923885813 | 1.74023355040433 | 0.1047107783142 | 0.16994311840192 |
| Mdc1 | ENSMUSG00000061607 | 240087 | 0.176825636179143 | 3.08608647572318 | 1.74007740999854 | 0.1047389800111 | 0.16994311840192 |
| Ehd1 | ENSMUSG00000024772 | 13660 | -0.205506988172384 | 4.68805473357235 | -1.74007487872647 | 0.1047394372585 | 0.16994311840192 |
| Bicra | ENSMUSG00000070808 | 243842 | 0.200393408820331 | 4.10561391743581 | 1.73978249384252 | 0.1047922653707 | 0.169987720174496 |
| Cradd | ENSMUSG00000045867 | 12905 | -0.326763344865288 | 1.7336534210997 | -1.73968515368048 | 0.1048098580227 | 0.169987720174496 |
| Pomp | ENSMUSG00000029649 | 66537 | 0.130050037277696 | 5.22999983611877 | 1.73966456638561 | 0.1048135791762 | 0.169987720174496 |
| Wdr6 | ENSMUSG00000066357 | 83669 | 0.246985268811573 | 2.2607560786755 | 1.73907742853976 | 0.1049197535336 | 0.170134672285513 |
| Soga1 | ENSMUSG00000055485 | 320706 | 0.245200735750428 | 4.18546499623314 | 1.73722203853857 | 0.1052558961020 | 0.17065324107266 |
| Rdh10 | ENSMUSG00000025921 | 98711 | 0.391806265424416 | 1.00557016430108 | 1.73714005197321 | 0.1052707716176 | 0.17065324107266 |
| Irgm1 | ENSMUSG00000046879 | 15944 | -0.3991901990829 | 3.6675637698887 | -1.7367883815592 | 0.1053345992380 | 0.170731391499224 |
| Ppie | ENSMUSG00000028651 | 56031 | -0.265620333827986 | 1.66729177499845 | -1.73620118615873 | 0.1054412505172 | 0.170878919111103 |
| Lins1 | ENSMUSG00000053091 | 72635 | -0.281576419741934 | 1.73445831875104 | -1.73573597007711 | 0.1055258146571 | 0.17097196455496 |
| Pcolce2 | ENSMUSG00000015354 | 76477 | 0.233406060993843 | 2.3565069898976 | 1.73571324801943 | 0.1055299464689 | 0.17097196455496 |
| Nup107 | ENSMUSG00000052798 | 103468 | -0.209363127623614 | 2.43655640061738 | -1.73520833077433 | 0.1056217982479 | 0.171095417404923 |
| Prr13 | ENSMUSG00000023048 | 66151 | 0.155584317035979 | 3.03837082357128 | 1.73491692360955 | 0.1056748415910 | 0.171155977690341 |
| Rpl11 | ENSMUSG00000059291 | 67025 | 0.123418981572939 | 5.90313902364936 | 1.73461257673481 | 0.1057302654125 | 0.171220374996227 |
| Alg3 | ENSMUSG00000033809 | 208624 | -0.237854687659601 | 1.52567089902904 | -1.73393193609884 | 0.1058543080804 | 0.171395858632377 |
| Rnf169 | ENSMUSG00000058761 | 108937 | 0.150980687254101 | 3.73130265988474 | 1.7335791781893 | 0.1059186466654 | 0.171474633693736 |
| Ei24 | ENSMUSG00000062762 | 13663 | -0.114127170268603 | 4.90520232171186 | -1.73336020507981 | 0.1059586019707 | 0.171513916502707 |
| Hfe | ENSMUSG00000006611 | 15216 | -0.157678130078803 | 3.21279813923443 | -1.73301646034486 | 0.1060213508038 | 0.171590077826002 |
| Cr1l | ENSMUSG00000016481 | 12946 | -0.116636886833891 | 4.16122817870453 | -1.73276687984938 | 0.1060669310073 | 0.171618337302485 |
| Depdc7 | ENSMUSG00000027173 | 211896 | -0.369592395709349 | 1.17301273568267 | -1.73274891826995 | 0.1060702119490 | 0.171618337302485 |
| Atg4a | ENSMUSG00000079418 | 666468 | 0.173060444473042 | 3.56922302861763 | 1.73239564806827 | 0.1061347600749 | 0.171697360127192 |
| Cldn12 | ENSMUSG00000046798 | 64945 | -0.259125313836177 | 2.43201111473075 | -1.73178546222495 | 0.1062463325292 | 0.171852420964362 |
| Mmgt1 | ENSMUSG00000061273 | 236792 | -0.132191473559047 | 4.42720816143998 | -1.72964991430012 | 0.1066376334726 | 0.172459827117787 |
| Pmm2 | ENSMUSG00000022711 | 54128 | -0.157624998186424 | 3.64804579258265 | -1.7280050139612 | 0.1069398989710 | 0.172923082197112 |
| Bnip2 | ENSMUSG00000011958 | 12175 | -0.13360328887371 | 5.81214195544924 | -1.72719896117966 | 0.1070882944630 | 0.173137427418361 |
| Mcm2 | ENSMUSG00000002870 | 17216 | -0.272244543545857 | 1.7802590868222 | -1.72698313357411 | 0.1071280594734 | 0.173155848383687 |
| Sorbs3 | ENSMUSG00000022091 | 20410 | 0.119967229880714 | 4.65814877303397 | 1.72696517004428 | 0.1071313697391 | 0.173155848383687 |
| Itfg2 | ENSMUSG00000001518 | 101142 | -0.222070055228603 | 2.93473212752261 | -1.72641644537655 | 0.1072325305876 | 0.173293730315361 |
| Ppcs | ENSMUSG00000028636 | 106564 | -0.294126477973634 | 1.68279766341949 | -1.72620901556742 | 0.1072707935291 | 0.173329939995208 |
| Trmt10b | ENSMUSG00000035601 | 69934 | -0.324498799190188 | 1.32879083573802 | -1.726094510149 | 0.1072919206014 | 0.173338454640077 |
| Kat6a | ENSMUSG00000031540 | 244349 | -0.0974569700665153 | 5.84436395191662 | -1.72539169391204 | 0.1074216756912 | 0.173522437857484 |
| Snapc5 | ENSMUSG00000032398 | 330959 | -0.197053632678482 | 2.91073398840039 | -1.72513372052675 | 0.1074693378874 | 0.173573778434968 |
| Rusc1 | ENSMUSG00000041263 | 72296 | -0.230800113871975 | 2.82355463541324 | -1.72471221266849 | 0.1075472542475 | 0.173673960138823 |
| Klhdc8b | ENSMUSG00000032609 | 78267 | 0.318565630328833 | 1.23048132747375 | 1.72455303827707 | 0.1075766908332 | 0.173681962832955 |
| Iigp1 | ENSMUSG00000054072 | 60440 | -0.599911802625947 | 4.07999141482662 | -1.72450643341236 | 0.1075853109522 | 0.173681962832955 |
| Zfp316 | ENSMUSG00000046658 | 54201 | 0.180575860111778 | 2.67311961356905 | 1.72442769081097 | 0.1075998767088 | 0.173681962832955 |
| Dgka | ENSMUSG00000025357 | 13139 | -0.265404919889169 | 3.30454450659324 | -1.72371275527557 | 0.1077322046513 | 0.173869884695738 |
| Dhdh | ENSMUSG00000011382 | 71755 | -0.240678156633902 | 4.18469255961103 | -1.72352087685167 | 0.1077677439933 | 0.173879006094778 |
| Pde3a | ENSMUSG00000041741 | 54611 | -0.341303617851097 | 2.95806078006917 | -1.72351047811429 | 0.1077696703222 | 0.173879006094778 |
| Bcor | ENSMUSG00000040363 | 71458 | -0.158658514264629 | 4.85279233838523 | -1.72289780764557 | 0.1078832189546 | 0.174036521014326 |
| Nt5dc2 | ENSMUSG00000071547 | 70021 | 0.392985164709813 | 1.19037214159141 | 1.72244238425928 | 0.1079676927187 | 0.174147092977585 |
| Rita1 | ENSMUSG00000029600 | 100764 | -0.381388501743452 | 1.17176615333456 | -1.72030835937051 | 0.1083642977000 | 0.174761011826456 |
| Zfp606 | ENSMUSG00000030386 | 67370 | -0.226812312923715 | 2.94571963849686 | -1.72007660371975 | 0.1084074463242 | 0.174789920597261 |
| Rps3a1 | ENSMUSG00000028081 | 20091 | -0.132031366084091 | 7.07272796690179 | -1.72004031603786 | 0.1084142037919 | 0.174789920597261 |
| Stim1 | ENSMUSG00000030987 | 20866 | 0.11718991147093 | 7.47200316888907 | 1.71946946166824 | 0.1085205567832 | 0.174935585635969 |
| Alg12 | ENSMUSG00000035845 | 223774 | 0.265168866625288 | 1.54745772213621 | 1.71901658647762 | 0.1086049950747 | 0.175045886340682 |
| Pitrm1 | ENSMUSG00000021193 | 69617 | -0.150848662904218 | 4.71066266433699 | -1.71871962025837 | 0.1086603956673 | 0.175109359433788 |
| Polk | ENSMUSG00000021668 | 27015 | -0.234263718288875 | 1.9939033657639 | -1.71819618726265 | 0.1087581054547 | 0.175240986250836 |
| Strap | ENSMUSG00000030224 | 20901 | -0.119546056190114 | 6.00295570878052 | -1.71804055188419 | 0.1087871730042 | 0.1752530075172 |
| Ifrd1 | ENSMUSG00000001627 | 15982 | -0.159720900132307 | 5.22220966096391 | -1.71798456373345 | 0.1087976314163 | 0.1752530075172 |
| Nifk | ENSMUSG00000026377 | 67949 | 0.179041854099696 | 3.01585616894162 | 1.71787398576375 | 0.1088182896393 | 0.175260457351891 |
| Akr7a5 | ENSMUSG00000028743 | 110198 | -0.157721308699474 | 3.35040285803983 | -1.71679205778524 | 0.1090205983018 | 0.17556042428374 |
| Adgra3 | ENSMUSG00000029090 | 70693 | -0.111022505034781 | 4.15990238703273 | -1.71561859085451 | 0.1092403982852 | 0.17588846570276 |
| Sparcl1 | ENSMUSG00000029309 | 13602 | 0.186950132663706 | 7.50657302248538 | 1.71479862126262 | 0.1093942169033 | 0.176110189244633 |
| Ank1 | ENSMUSG00000031543 | 11733 | -0.136260833393614 | 6.95935574778992 | -1.71420655012575 | 0.1095054024712 | 0.176263224033677 |
| Rhoj | ENSMUSG00000046768 | 80837 | 0.222970998166745 | 4.2012135940872 | 1.71258678890883 | 0.1098100875537 | 0.176712370680908 |
| Prr14l | ENSMUSG00000054280 | 215476 | 0.177356240705239 | 3.69816049576537 | 1.71255129789805 | 0.1098167719362 | 0.176712370680908 |
| Sema7a | ENSMUSG00000038264 | 20361 | 0.215758598938927 | 3.42407760641587 | 1.71232085547961 | 0.1098601822121 | 0.176756204262735 |
| Olfr466 | ENSMUSG00000049806 | 258816 | -0.74949000603618 | 0.454777836897604 | -1.71209600054594 | 0.1099025544901 | 0.176796733842937 |
| Nars | ENSMUSG00000024587 | 70223 | -0.148841023050025 | 5.68102652213017 | -1.71201553815593 | 0.1099177205376 | 0.176796733842937 |
| Arpin | ENSMUSG00000039043 | 70420 | 0.276375105142382 | 2.28569029175353 | 1.71186862021661 | 0.1099454172900 | 0.176815265028339 |
| Thap2 | ENSMUSG00000020137 | 66816 | -0.315332289876078 | 2.91025717843413 | -1.71144579886787 | 0.1100251612296 | 0.176857285859275 |
| Lrrc8a | ENSMUSG00000007476 | 241296 | -0.224225306486455 | 4.94688814758854 | -1.7113996455845 | 0.1100338688018 | 0.176857285859275 |
| Enpep | ENSMUSG00000028024 | 13809 | 0.214919696331789 | 2.78434839182003 | 1.71138480157021 | 0.1100366694970 | 0.176857285859275 |
| Ipo8 | ENSMUSG00000040029 | 320727 | -0.151433509303076 | 4.23137683637668 | -1.71131876070787 | 0.1100491305206 | 0.176857285859275 |
| Ndufaf2 | ENSMUSG00000068184 | 75597 | -0.170924381110413 | 2.65966022546671 | -1.71130120442559 | 0.1100524433642 | 0.176857285859275 |
| Pip5k1a | ENSMUSG00000028126 | 18720 | -0.144603987724713 | 4.00180159218855 | -1.71065558601879 | 0.1101743315536 | 0.177027137764644 |
| Gopc | ENSMUSG00000019861 | 94221 | 0.115539528397712 | 3.97809865343937 | 1.71052655794837 | 0.1101987053812 | 0.177034861668119 |
| Tmem230 | ENSMUSG00000027341 | 70612 | -0.162327633437859 | 2.73623409561038 | -1.71045867854694 | 0.1102115299287 | 0.177034861668119 |
| Aak1 | ENSMUSG00000057230 | 269774 | 0.11301790346466 | 5.19598941540381 | 1.70926925650329 | 0.1104364622947 | 0.177358269339545 |
| Mef2c | ENSMUSG00000005583 | 17260 | 0.146156424011586 | 9.42897071223705 | 1.70922248726709 | 0.1104453151033 | 0.177358269339545 |
| Ttbk2 | ENSMUSG00000090100 | 140810 | 0.140639626417816 | 3.97095241261031 | 1.70905679768726 | 0.1104766830119 | 0.177382582550416 |
| Prkch | ENSMUSG00000021108 | 18755 | -0.206048263900688 | 3.09941607747254 | -1.70880059748267 | 0.1105252015814 | 0.177434421866132 |
| Rnf215 | ENSMUSG00000003581 | 71673 | 0.239996666731036 | 2.01606357561795 | 1.70763157205236 | 0.1107468267717 | 0.177764106232296 |
| Noct | ENSMUSG00000023087 | 12457 | -0.217992060476529 | 6.84488678804186 | -1.70721076797985 | 0.1108266989281 | 0.177866193801776 |
| Tmem100 | ENSMUSG00000069763 | 67888 | 0.432379117356324 | 2.07988644034163 | 1.70696392528989 | 0.1108735753243 | 0.177915304150135 |
| Thsd1 | ENSMUSG00000031480 | 56229 | -0.407820179947118 | 1.66376152694463 | -1.70650202628501 | 0.1109613386088 | 0.178030000342516 |
| Rpl9 | ENSMUSG00000047215 | 20005 | -0.127629110796049 | 5.53653754611744 | -1.70565358530403 | 0.1111227061343 | 0.178262738590317 |
| Tmem104 | ENSMUSG00000045980 | 320534 | 0.19709191501315 | 2.61121679025907 | 1.70481773989883 | 0.1112818798216 | 0.178484398205555 |
| Agap2 | ENSMUSG00000025422 | 216439 | -0.448878627011883 | 1.87958730339554 | -1.70473187384341 | 0.1112982430188 | 0.178484398205555 |
| Mid1ip1 | ENSMUSG00000008035 | 68041 | 0.417269998999771 | 4.70180295337849 | 1.70467089015066 | 0.1113098657549 | 0.178484398205555 |
| Pdgfrb | ENSMUSG00000024620 | 18596 | 0.204538489679971 | 5.5236729434236 | 1.70441372690485 | 0.1113588896217 | 0.178536817608619 |
| Senp3 | ENSMUSG00000005204 | 80886 | -0.117602704248119 | 3.99129035058834 | -1.70383406543467 | 0.1114694619972 | 0.178687885187193 |
| Ap4m1 | ENSMUSG00000019518 | 11781 | 0.290028256231857 | 2.21128652783827 | 1.70316689301625 | 0.1115968468425 | 0.178865855254544 |
| Parva | ENSMUSG00000030770 | 57342 | 0.233462696555318 | 4.32369434570827 | 1.70286670446945 | 0.1116542042958 | 0.178931550675504 |
| Sik2 | ENSMUSG00000037112 | 235344 | -0.267927928451721 | 3.71763084899809 | -1.70253389590094 | 0.1117178247892 | 0.179007262206299 |
| Hnrnpdl | ENSMUSG00000029328 | 50926 | -0.128134358890705 | 5.38176008648217 | -1.70230452902734 | 0.1117616896607 | 0.179027826886456 |
| Ubqln2 | ENSMUSG00000050148 | 54609 | 0.122167417927451 | 4.40569567142298 | 1.70229550832014 | 0.1117634151196 | 0.179027826886456 |
| Cndp2 | ENSMUSG00000024644 | 66054 | 0.150391134999483 | 3.58441703308443 | 1.70137115755561 | 0.1119403468352 | 0.17928497161256 |
| Uba6 | ENSMUSG00000035898 | 231380 | 0.182379009639366 | 2.80461410185904 | 1.70076003801581 | 0.1120574574846 | 0.179446244911382 |
| Ubr3 | ENSMUSG00000044308 | 68795 | -0.121299485575037 | 7.92880702946108 | -1.70023557648211 | 0.1121580474237 | 0.17958101895249 |
| Wdsub1 | ENSMUSG00000026988 | 72137 | 0.240671288396975 | 1.94979650980208 | 1.69991770546024 | 0.1122190526086 | 0.179652381966215 |
| Hspa8 | ENSMUSG00000015656 | 15481 | -0.136837153158371 | 10.108064748765 | -1.69925556325116 | 0.1123462232566 | 0.179829633389728 |
| Zdhhc5 | ENSMUSG00000034075 | 228136 | -0.114766991619343 | 6.59929659652999 | -1.69888216891387 | 0.1124179929635 | 0.179898108465134 |
| Taf13 | ENSMUSG00000048100 | 99730 | 0.236143925811839 | 2.24757012250799 | 1.69886175747444 | 0.1124219173840 | 0.179898108465134 |
| Emp2 | ENSMUSG00000022505 | 13731 | 0.201467353224927 | 3.34486456315983 | 1.69840858586209 | 0.1125090777567 | 0.179997871766857 |
| Trip11 | ENSMUSG00000021188 | 109181 | -0.101982451997476 | 6.56671915335971 | -1.69836639806081 | 0.1125171949302 | 0.179997871766857 |
| Atp6v0b | ENSMUSG00000033379 | 114143 | -0.146911575596887 | 3.57121656881816 | -1.69793849363418 | 0.1125995552506 | 0.180103268721777 |
| Aptx | ENSMUSG00000028411 | 66408 | -0.156599183168706 | 3.25970254113908 | -1.69782188725777 | 0.1126220080812 | 0.180112826676862 |
| Nosip | ENSMUSG00000003421 | 66394 | -0.140117202083406 | 3.84343231153876 | -1.69758970414326 | 0.1126667271922 | 0.180157986386492 |
| Blmh | ENSMUSG00000020840 | 104184 | 0.116826304790284 | 3.99121023594689 | 1.69557954035054 | 0.1130545425953 | 0.180751675458433 |
| Vmn2r115 | ENSMUSG00000091076 | 638102 | -0.524895780189672 | 1.84753411993742 | -1.69483200619874 | 0.1131990609687 | 0.180934998867986 |
| Klf6 | ENSMUSG00000000078 | 23849 | 0.317090186200493 | 4.75718854953222 | 1.6948152067816 | 0.1132023106081 | 0.180934998867986 |
| Supt6 | ENSMUSG00000002052 | 20926 | 0.114043157371492 | 6.0261390546344 | 1.69420472964099 | 0.1133204554234 | 0.181097353542975 |
| Wbp1l | ENSMUSG00000047731 | 226178 | -0.150849655927889 | 5.11889384983451 | -1.69401659972695 | 0.1133568857577 | 0.18112909197745 |
| Ramac | ENSMUSG00000038646 | 67148 | -0.206385196377991 | 3.0239111606252 | -1.69354302197617 | 0.1134486370173 | 0.181249203629976 |
| Ttll7 | ENSMUSG00000036745 | 70892 | 0.139063048567036 | 5.80422412085426 | 1.69338599714656 | 0.1134790734925 | 0.181271336014479 |

| | | | | | | | |
|---|---|---|---|---|---|---|---|
| Rsrp1 | ENSMUSG00000037266 | 27981 | -0.285473438670665 | 5.96753096653212 | -1.69319487570897 | 0.1135161286578 | 0.18130403307407 |
| Kif13a | ENSMUSG00000021375 | 16553 | 0.13333845371447 | 7.33311509601678 | 1.69215143427524 | 0.1137186213079 | 0.181600913004694 |
| Prpf18 | ENSMUSG00000039449 | 67229 | -0.0966446818332924 | 4.95433804882185 | -1.69096176428238 | 0.1139498779708 | 0.181943633669282 |
| Hsp90aa1 | ENSMUSG00000021270 | 15519 | 0.168258387332551 | 6.42036714166806 | 1.69069473550988 | 0.1140018416079 | 0.182000019076424 |
| Ctdsp2 | ENSMUSG00000078429 | 52468 | 0.123328002049059 | 6.49062730513549 | 1.68996514301004 | 0.1141439258514 | 0.18220024145482 |
| Vmn1r233 | ENSMUSG00000045575 | 171236 | -0.671703997040334 | 0.647085503797009 | -1.68883415556412 | 0.1143644865811 | 0.182525653791586 |
| Zfp933 | ENSMUSG00000059423 | 242747 | -0.272062631976211 | 2.79770408489336 | -1.68862922567922 | 0.1144044911691 | 0.182562845689038 |
| Lrp4 | ENSMUSG00000027253 | 228357 | 0.177697099919589 | 4.57790018480187 | 1.68716944250083 | 0.1146898121805 | 0.182991437300464 |
| Wdr83os | ENSMUSG00000059355 | 414077 | 0.170937578325006 | 2.87792451995854 | 1.68687115175586 | 0.1147481911052 | 0.183057862955633 |
| B230219D22Rik | ENSMUSG00000045767 | 78521 | -0.102904876252083 | 4.93694826180466 | -1.68610903696258 | 0.1148974640213 | 0.183269251308487 |
| Icmt | ENSMUSG00000039662 | 57295 | -0.119355466077567 | 4.0520417336012 | -1.68539763033751 | 0.1150369584430 | 0.183464982891933 |
| Stoml1 | ENSMUSG00000032333 | 69106 | 0.231361914336135 | 1.94897947055082 | 1.68507575849683 | 0.1151001206409 | 0.183538937815637 |
| Sec14l5 | ENSMUSG00000091712 | 665119 | 0.19243492795052 | 3.97574481682503 | 1.68484222880902 | 0.1151459661250 | 0.183585261918385 |
| Ccm2 | ENSMUSG00000000378 | 216527 | -0.177108591450197 | 3.20369316648301 | -1.68395631402433 | 0.1153200303860 | 0.183835970854536 |
| Tead3 | ENSMUSG00000002249 | 21678 | 0.264354863338033 | 2.18679695859646 | 1.68378003368881 | 0.1153546933772 | 0.183864414305366 |
| Trmt11 | ENSMUSG00000019792 | 73681 | -0.296363110766417 | 1.54838464581715 | -1.68352259550721 | 0.1154053312889 | 0.18387976560913 |
| Pigp | ENSMUSG00000022940 | 56176 | -0.157903012301745 | 4.04682826866891 | -1.68348440305382 | 0.1154128453765 | 0.18387976560913 |
| Rps19bp1 | ENSMUSG00000051518 | 66538 | 0.28393388641635 | 2.52742874927265 | 1.68347451822086 | 0.1154147902153 | 0.18387976560913 |
| Zranb3 | ENSMUSG00000036086 | 226409 | -0.31023076178229 | 1.45746495261528 | -1.68250868662662 | 0.1156049558738 | 0.184114091192971 |
| Nxn | ENSMUSG00000020844 | 18230 | 0.100290127749751 | 4.92778051826257 | 1.68244718559356 | 0.1156170742968 | 0.184114091192971 |
| Foxo1 | ENSMUSG00000044167 | 56458 | -0.18964792614283 | 4.39553222450468 | -1.68239205472182 | 0.1156279384620 | 0.184114091192971 |
| Srl | ENSMUSG00000022519 | 106393 | -0.160741520852687 | 9.60774455910527 | -1.6823012755597 | 0.1156458294765 | 0.184114091192971 |
| Tap1 | ENSMUSG00000037321 | 21354 | -0.281901396246982 | 2.41966649479499 | -1.68222588691814 | 0.1156606891264 | 0.184114091192971 |
| Rfxank | ENSMUSG00000036120 | 19727 | -0.231280157248035 | 2.28128181600293 | -1.68221452980852 | 0.1156629278402 | 0.184114091192971 |
| Isg20l2 | ENSMUSG00000048039 | 229504 | 0.220348795209255 | 2.65807999650563 | 1.68157579507193 | 0.1157888963561 | 0.184265160283557 |
| Mtmr9 | ENSMUSG00000035078 | 210376 | 0.139095429123051 | 3.29207418799053 | 1.68156236789734 | 0.1157915456946 | 0.184265160283557 |
| Mzt1 | ENSMUSG00000033186 | 76789 | -0.15306802368259 | 3.37592317327404 | -1.68029361042379 | 0.1160421259709 | 0.184637042065218 |
| Entr1 | ENSMUSG00000026927 | 68112 | -0.143532513074722 | 3.97453148331212 | -1.6800784035221 | 0.1160846765118 | 0.184668476528112 |
| Eif4e2 | ENSMUSG00000026254 | 26987 | 0.0983861431683675 | 4.72332888699175 | 1.68002281234089 | 0.1160956701766 | 0.184668476528112 |
| Lrch1 | ENSMUSG00000068015 | 380916 | 0.28225858740891 | 3.56772012002812 | 1.67888160533051 | 0.1163215557987 | 0.185000862277974 |
| Fam53a | ENSMUSG00000037339 | 74504 | 0.119536832932083 | 5.10638571871019 | 1.6786199517942 | 0.1163734005589 | 0.185056392422188 |
| Zfp128 | ENSMUSG00000060397 | 243833 | -0.278401970574958 | 1.26707118681092 | -1.67777589134302 | 0.1165407829207 | 0.185295606997352 |
| Pdzd11 | ENSMUSG00000015668 | 72621 | 0.245490993617316 | 1.68513394619939 | 1.67768131867663 | 0.1165595503797 | 0.185298494066548 |
| Gnao1 | ENSMUSG00000031748 | 14681 | 0.160164047460763 | 3.4958944462341 | 1.67753447911589 | 0.1165886951763 | 0.185317875086866 |
| Zfp52 | ENSMUSG00000051341 | 22710 | -0.285045890116487 | 3.16984753422474 | -1.67741965359232 | 0.1166114902521 | 0.185327159050038 |
| Brd2 | ENSMUSG00000024335 | 14312 | 0.127982802020166 | 6.11177073171546 | 1.67732163897742 | 0.1166309511243 | 0.185331142134092 |
| Fmod | ENSMUSG00000041559 | 14264 | 0.398831409679936 | 7.11371761137213 | 1.67610417130647 | 0.1168729171868 | 0.185688642117679 |
| Tipin | ENSMUSG00000032397 | 66131 | -0.190357248465611 | 1.98872691711328 | -1.67589878801107 | 0.1169137794310 | 0.18572656924299 |
| Wdr18 | ENSMUSG00000035754 | 216156 | -0.140724587069605 | 3.9264681344548 | -1.67581167594021 | 0.1169311146769 | 0.185727116322774 |
| Gosr2 | ENSMUSG00000020946 | 56494 | 0.113899986353021 | 4.85185213929533 | 1.67468835985598 | 0.1171548553734 | 0.186055458969498 |
| Wdr11 | ENSMUSG00000042055 | 207425 | -0.116867321243316 | 3.92863365583983 | -1.67416701775722 | 0.1172588228032 | 0.186193520055563 |
| Rab6a | ENSMUSG00000030704 | 19346 | 0.118443616234358 | 5.64872999899845 | 1.67383892661076 | 0.1173242929746 | 0.186270420697974 |
| Mtmr1 | ENSMUSG00000015214 | 53332 | 0.136997658798631 | 4.62027455718664 | 1.67355561786071 | 0.1173808525934 | 0.186333154182289 |
| BC005537 | ENSMUSG00000019132 | 79555 | 0.101908881746564 | 5.77217388186793 | 1.67325947562308 | 0.1174399997699 | 0.186379096955982 |
| Dnajc4 | ENSMUSG00000024963 | 57431 | -0.165281910343716 | 2.94059892899822 | -1.67323997564615 | 0.1174438953282 | 0.186379096955982 |
| Nfix | ENSMUSG00000001911 | 18032 | 0.119088320172057 | 8.81301122712195 | 1.67280509541477 | 0.1175308017835 | 0.186489939656779 |
| Nepro | ENSMUSG00000036208 | 212547 | 0.254687730770406 | 1.50088042258453 | 1.67143397459853 | 0.1178051747407 | 0.186898166196026 |
| Olfr1156 | ENSMUSG00000075144 | 258814 | -0.646540738385459 | 1.26998354807015 | -1.67041085069565 | 0.1180102751715 | 0.187196389188759 |
| Setd5 | ENSMUSG00000034269 | 72895 | 0.140926471098783 | 5.23933820268346 | 1.67014956491825 | 0.1180627038004 | 0.187233768557141 |
| Cables1 | ENSMUSG00000040957 | 63955 | -0.360406436001535 | 1.72869668998876 | -1.67012269361668 | 0.1180680968497 | 0.187233768557141 |
| Elmo2 | ENSMUSG00000017670 | 140579 | 0.153371362308722 | 3.46379640225984 | 1.67001390667889 | 0.1180899325036 | 0.187241231824392 |
| Kat2a | ENSMUSG00000020918 | 14534 | 0.165188305764028 | 3.26294041391541 | 1.66982173749491 | 0.1181285132191 | 0.18727523973877 |
| Pdf | ENSMUSG00000078931 | 68023 | -0.175126017291521 | 3.38413905694868 | -1.66964106812569 | 0.1181647952276 | 0.187305594059389 |
| Bphl | ENSMUSG00000038286 | 68021 | -0.20358463106276 | 1.99460710094595 | -1.66896206975692 | 0.1183012387087 | 0.187475142038855 |
| Micos13 | ENSMUSG00000049760 | 224904 | -0.199587582533638 | 3.73752565013752 | -1.66893809506091 | 0.1183060588952 | 0.187475142038855 |
| Extl2 | ENSMUSG00000027963 | 58193 | 0.201479892681578 | 2.57140861213031 | 1.66871033565959 | 0.1183518591836 | 0.18752053525681 |
| Cct7 | ENSMUSG00000030007 | 12468 | -0.106197918510887 | 5.98431229728829 | -1.6685220999683 | 0.1183897233148 | 0.187553342833951 |
| Bag3 | ENSMUSG00000030847 | 29810 | -0.139562511659819 | 7.94567792086342 | -1.66829474169606 | 0.1184354711729 | 0.187598628516323 |
| Bola2 | ENSMUSG00000047721 | 66162 | 0.195752446251727 | 2.55506666443705 | 1.66743281444444 | 0.1186090440080 | 0.187846343096495 |
| Abat | ENSMUSG00000057880 | 268860 | 0.189697128164481 | 3.34887110582994 | 1.66703975869077 | 0.1186882703717 | 0.18792155611488 |
| Lig3 | ENSMUSG00000020697 | 16882 | 0.120851188147914 | 4.48731903359168 | 1.66702662786797 | 0.1186909178864 | 0.18792155611488 |
| Sh3bp4 | ENSMUSG00000036206 | 98402 | 0.254940884444398 | 2.75715255781879 | 1.66611604318819 | 0.1188746412538 | 0.188185185162265 |
| Hdac6 | ENSMUSG00000031161 | 15185 | 0.194173131405171 | 2.21849787493671 | 1.66582055852844 | 0.1189343128222 | 0.188252385383753 |
| Trafd1 | ENSMUSG00000042726 | 231712 | -0.14249019936101 | 5.0516650627795 | -1.66509219150664 | 0.1190815144167 | 0.188446873718543 |
| Grina | ENSMUSG00000022564 | 66168 | -0.128875875650957 | 5.10160401991148 | -1.66504198834608 | 0.1190916662380 | 0.188446873718543 |
| Zfp169 | ENSMUSG00000050954 | 67911 | 0.272179305029324 | 1.70130134671583 | 1.66407234469084 | 0.1192878908137 | 0.188730052755154 |
| Fnta | ENSMUSG00000015994 | 14272 | -0.102002611236748 | 4.76907520198678 | -1.66393505193808 | 0.1193156972346 | 0.18874672742 |
| Usp30 | ENSMUSG00000029592 | 100756 | 0.196551880248379 | 2.35505528429769 | 1.66372143164018 | 0.1193589738135 | 0.18878786612014 |
| Ube2a | ENSMUSG00000016308 | 22209 | -0.137509847364714 | 3.85698973276562 | -1.66360052785543 | 0.1193834733543 | 0.188799297921989 |
| Dok5 | ENSMUSG00000027560 | 76829 | 0.236980259512284 | 2.68126343410353 | 1.6630161479454 | 0.1195019520486 | 0.188959328488894 |
| Pef1 | ENSMUSG00000028779 | 67898 | -0.207451703302431 | 2.87893154493306 | -1.66211285864816 | 0.1196852891718 | 0.189221853621202 |
| Heatr5a | ENSMUSG00000035181 | 320487 | -0.128912237657263 | 5.21927290433688 | -1.66169286155653 | 0.1197706179186 | 0.189329374471233 |
| Poglut1 | ENSMUSG00000034064 | 224143 | 0.216437866937575 | 2.24566305920657 | 1.6608906165932 | 0.1199337536365 | 0.189559841093258 |
| Gsdmd | ENSMUSG00000022575 | 69146 | 0.342760570931134 | 1.31927612535769 | 1.66033099098525 | 0.1200476677067 | 0.189712455645906 |
| Olfr1336 | ENSMUSG00000048620 | 258917 | -0.634276478096286 | 2.18442725422153 | -1.66003541541002 | 0.1201078714023 | 0.18978015933786 |
| Psmb4 | ENSMUSG00000005779 | 19172 | 0.127820869629931 | 5.06380549332189 | 1.65955356138154 | 0.1202060733038 | 0.18990787502301 |
| Ncaph2 | ENSMUSG00000008690 | 52683 | -0.115584277702395 | 5.18361521018972 | -1.65940812569414 | 0.1202357268684 | 0.189927273179065 |
| Olfr1487 | ENSMUSG00000094846 | 258629 | -0.984341764532186 | -0.650693012476632 | -1.65871197342642 | 0.1203777570043 | 0.190124152841336 |
| Mkks | ENSMUSG00000027274 | 59030 | -0.245988574241594 | 2.81465420448444 | -1.65819197802351 | 0.1204839427741 | 0.190264371347034 |
| Anks3 | ENSMUSG00000022515 | 72615 | 0.237163494518892 | 2.33571773609556 | 1.65750551256112 | 0.1206242476843 | 0.190458421412314 |
| Dcakd | ENSMUSG00000020935 | 68087 | 0.143276113925274 | 3.31694537697342 | 1.65728007947056 | 0.1206703544387 | 0.190503703692887 |
| Myo10 | ENSMUSG00000022272 | 17909 | -0.110738487687491 | 6.2523754069579 | -1.6570670553817 | 0.1207139373592 | 0.190539695992026 |
| Strn4 | ENSMUSG00000030374 | 97387 | 0.130531468637291 | 4.4456126780433 | 1.65697611929757 | 0.1207325462896 | 0.190539695992026 |
| Rpe | ENSMUSG00000026005 | 66646 | 0.310345717425725 | 2.02179941187716 | 1.65691308782854 | 0.1207454463577 | 0.190539695992026 |
| Mapk12 | ENSMUSG00000022610 | 29857 | -0.127480927480426 | 5.62183952613941 | -1.65652512838131 | 0.1208248728958 | 0.190630819093774 |
| St8sia4 | ENSMUSG00000040710 | 20452 | 0.279886055230611 | 3.05363441800131 | 1.65642133207728 | 0.1208461307281 | 0.190630819093774 |
| Zfp667 | ENSMUSG00000054893 | 384763 | -0.319177479365629 | 1.48530896695385 | -1.65637554222337 | 0.1208555096807 | 0.190630819093774 |
| Sp110 | ENSMUSG00000070034 | 109032 | -0.348031668512446 | 1.83881537079377 | -1.65551506182516 | 0.1210318764788 | 0.1908814661363 |
| Phf20 | ENSMUSG00000038116 | 228829 | -0.135933763685778 | 4.24270230482572 | -1.65485115295938 | 0.1211681066300 | 0.191054097470031 |
| Zw10 | ENSMUSG00000032264 | 26951 | 0.20002583702076 | 2.98125913395459 | 1.65481127839416 | 0.1211762929063 | 0.191054097470031 |
| Prpf38a | ENSMUSG00000063800 | 230596 | 0.166119654094322 | 2.76230674418636 | 1.6546462824068 | 0.1212101718227 | 0.191079952147993 |
| Tmtc2 | ENSMUSG00000036019 | 278279 | -0.377234665646295 | 1.3079675946817 | -1.65439996918625 | 0.1212607631093 | 0.191132141535502 |
| Surf2 | ENSMUSG00000014873 | 20931 | -0.190656598096948 | 2.40259156383221 | -1.65299514413987 | 0.1215496578094 | 0.191559877381071 |
| Borcs6 | ENSMUSG00000045176 | 71923 | -0.206507632956681 | 2.14632599469543 | -1.65245092652106 | 0.1216617342752 | 0.191708868006734 |
| Psma2 | ENSMUSG00000015671 | 19166 | -0.115170644310417 | 4.98134044266096 | -1.65187206074478 | 0.1217810449966 | 0.191869213441614 |
| Prrx1 | ENSMUSG00000026586 | 18933 | 0.132006795349373 | 6.47993618279377 | 1.65151033469303 | 0.1218556525202 | 0.191959091756546 |
| Slc25a10 | ENSMUSG00000025792 | 27376 | 0.250880316741626 | 1.50379620200047 | 1.65125974638778 | 0.1219073607548 | 0.19201287613998 |
| Itpripl1 | ENSMUSG00000074825 | 73338 | -0.227730021656087 | 2.07206378929006 | -1.6510677502748 | 0.1219469915748 | 0.192047624968219 |
| Tnnc1 | ENSMUSG00000091898 | 21924 | 0.223878960875769 | 9.75712307630857 | 1.65065390334672 | 0.1220324538252 | 0.192154530792889 |
| Tfdp2 | ENSMUSG00000032411 | 211586 | -0.170736430000528 | 5.13162518573827 | -1.65025481769763 | 0.1221149171664 | 0.192256684365017 |
| Zmynd19 | ENSMUSG00000026974 | 67187 | -0.157018556950836 | 3.08824455646831 | -1.65008440571075 | 0.1221501442937 | 0.192284450932395 |
| Rnf8 | ENSMUSG00000090083 | 58230 | -0.191035998184489 | 3.82678644192121 | -1.64989002823527 | 0.1221903363079 | 0.192320023928297 |
| Snora64 | ENSMUSG00000077709 | 104366 | 1.23215014660313 | -0.359265702291978 | 1.64972001842328 | 0.1222254991871 | 0.192347672273951 |
| Zfp967 | ENSMUSG00000095199 | 100303732 | -0.188033203066324 | 2.95572373095243 | -1.64945261196715 | 0.1222808242987 | 0.192403848769399 |
| Mdp1 | ENSMUSG00000002329 | 67881 | 0.194164658777219 | 3.49027580620353 | 1.64937735108215 | 0.1222963993458 | 0.192403848769399 |
| Qdpr | ENSMUSG00000015806 | 110391 | 0.182364343834197 | 3.78011065886245 | 1.64806443897518 | 0.1225683811399 | 0.192803996868702 |
| Ctsd | ENSMUSG00000007891 | 13033 | -0.131459084018538 | 7.59307963364046 | -1.64696501171572 | 0.1227965424972 | 0.193135108782428 |
| Socs2 | ENSMUSG00000020027 | 216233 | 0.261638371670609 | 2.67203971895451 | 1.64652277755327 | 0.1228884224264 | 0.193251812047695 |
| Cdk5 | ENSMUSG00000028969 | 12568 | 0.191837523870797 | 1.96235910715422 | 1.64605622217177 | 0.1229854202511 | 0.193376528878831 |
| Prkca | ENSMUSG00000050965 | 18750 | -0.10615215870466 | 5.10169132966815 | -1.6455539594195 | 0.1230899161673 | 0.193500709743119 |
| Fam3c | ENSMUSG00000029672 | 27999 | -0.167534192233752 | 3.44954045034346 | -1.64550646212754 | 0.1230998019900 | 0.193500709743119 |
| Fra10ac1 | ENSMUSG00000054237 | 70567 | -0.145826777701838 | 3.31344209093379 | -1.64527621971631 | 0.1231477331586 | 0.193548220151943 |
| Nt5dc1 | ENSMUSG00000039480 | 319638 | 0.337066626991319 | 1.72098100299967 | 1.64496061354512 | 0.1232134614785 | 0.193616596301421 |
| Mef2a | ENSMUSG00000030557 | 17258 | -0.105192645996098 | 6.60212882179392 | -1.64489723859684 | 0.1232266636601 | 0.193616596301421 |
| Zbed3 | ENSMUSG00000041995 | 72114 | -0.207520665635465 | 2.80142801728441 | -1.64420127361225 | 0.1233717271028 | 0.193816664122075 |
| Tspan15 | ENSMUSG00000037031 | 70423 | 0.221985862510689 | 2.63096784810348 | 1.644039462531 | 0.1234054754666 | 0.193841823872023 |

| | | | | | | | |
|---|---|---|---|---|---|---|---|
| Gga3 | ENSMUSG00000020740 | 260302 | -0.162920139852821 | 3.27971878737211 | -1.64379882782035 | 0.1234556786455 | 0.193892819435924 |
| Ube2l6 | ENSMUSG00000027078 | 56791 | -0.271952446721146 | 1.99387372736255 | -1.64329375204962 | 0.1235611094523 | 0.194030525416361 |
| Dock6 | ENSMUSG00000032198 | 319899 | -0.282973683466605 | 4.25122222945821 | -1.64300715635522 | 0.1236209689858 | 0.194096640618191 |
| Arl5b | ENSMUSG00000017418 | 75869 | -0.141444516815971 | 3.74332222139606 | -1.64210050041871 | 0.1238105029160 | 0.194366308685228 |
| Aph1a | ENSMUSG00000015750 | 226548 | -0.160390592879471 | 3.96860709967065 | -1.6419103104921 | 0.1238502936430 | 0.194380769697771 |
| Dnajc2 | ENSMUSG00000029014 | 22791 | 0.147369123693021 | 5.47147178608365 | 1.64188648274399 | 0.1238552795668 | 0.194380769697771 |
| Slc25a46 | ENSMUSG00000024259 | 67453 | -0.0957524971868411 | 5.0852639693461 | -1.64156018313395 | 0.1239235748750 | 0.194460034016552 |
| Cblb | ENSMUSG00000022637 | 208650 | 0.236315818704001 | 3.88306696809932 | 1.64146263380718 | 0.1239439985445 | 0.194463180569159 |
| Phf11d | ENSMUSG00000068245 | 219132 | -0.403533396894391 | 1.9696550804742 | -1.6413806755769 | 0.1239611602054 | 0.194463180569159 |
| Vmn1r18 | ENSMUSG00000091382 | 171199 | -0.713015708139555 | 1.62030895234635 | -1.63917499653072 | 0.1244237953305 | 0.195160927931967 |
| Olfr374 | ENSMUSG00000046881 | 258335 | -0.562150219299982 | 1.5240282182665 | -1.63903454639959 | 0.1244533051123 | 0.195179207773776 |
| Brf2 | ENSMUSG00000031487 | 66653 | -0.215785874635942 | 2.10116721749778 | -1.63877136786955 | 0.1245086175638 | 0.195237942704078 |
| Maea | ENSMUSG00000079562 | 59003 | 0.111841371289787 | 5.47872823499403 | 1.63821934113157 | 0.1246247068840 | 0.195370896767086 |
| Ralgapa1 | ENSMUSG00000021027 | 56784 | -0.109010374547918 | 5.95051545138038 | -1.63819821149497 | 0.1246291522419 | 0.195370896767086 |
| Aopep | ENSMUSG00000021458 | 72061 | 0.224788295437425 | 5.78811166689315 | 1.63795531280057 | 0.1246802643722 | 0.19542299539053 |
| Efr3b | ENSMUSG00000020658 | 668212 | -0.183706257253496 | 4.47698837040282 | -1.63761116571384 | 0.1247527129487 | 0.195508516880662 |
| Ercc1 | ENSMUSG00000003549 | 13870 | -0.172062572179131 | 2.45770284815636 | -1.63677990689015 | 0.1249278573683 | 0.195754932467584 |
| Dhps | ENSMUSG00000060038 | 330817 | -0.18363173021365 | 2.82042215297593 | -1.63663020290585 | 0.1249594223584 | 0.195776328765964 |
| Tnk2 | ENSMUSG00000022791 | 51789 | -0.350675085707017 | 4.43104692707147 | -1.63649327704055 | 0.1249882991558 | 0.195793507892309 |
| Akr1e1 | ENSMUSG00000045410 | 56043 | -0.180081179143344 | 2.74526664888089 | -1.63631542423729 | 0.1250258158389 | 0.195824214491373 |
| Nsfl1c | ENSMUSG00000027455 | 386649 | 0.122166005216361 | 4.93682751294299 | 1.63521557884604 | 0.1252580371067 | 0.19615982859803 |
| Ykt6 | ENSMUSG00000002741 | 56418 | 0.103038455002824 | 4.39518703528764 | 1.63493469324805 | 0.1253174032808 | 0.196224686329066 |
| Thy1 | ENSMUSG00000032011 | 21838 | 0.284789559589456 | 2.21107229077977 | 1.63459285226706 | 0.1253896855522 | 0.196309746888016 |
| Cdc42bpg | ENSMUSG00000024769 | 240505 | 0.242155929471935 | 2.16627136774591 | 1.63393417294933 | 0.1255290653696 | 0.196499815774773 |
| Atp6v0e2 | ENSMUSG00000039347 | 76252 | -0.212229305757125 | 3.98357865390377 | -1.63341112084865 | 0.1256398415906 | 0.196631574568726 |
| Snora16a | ENSMUSG00000065097 | 100310813 | 0.497427925353776 | 1.95287227412605 | 1.63336688977542 | 0.1256492130969 | 0.196631574568726 |
| Rpl36a | ENSMUSG00000079435 | 19982 | 0.143843284610722 | 5.10125585330378 | 1.6324891156063 | 0.1258353179936 | 0.196894626537171 |
| Crbn | ENSMUSG00000005362 | 58799 | -0.132799562040251 | 3.9416462524233 | -1.63196635139943 | 0.1259462676309 | 0.197040024541917 |
| Fbln5 | ENSMUSG00000021186 | 23876 | 0.23366989407325 | 3.4352030205363 | 1.63159016701426 | 0.1260261602020 | 0.197136799823836 |
| Olfr667 | ENSMUSG00000056782 | 259062 | -0.783805388628869 | -0.453177620188954 | -1.63109989437655 | 0.1261303483968 | 0.197271546476801 |
| Rora | ENSMUSG00000032238 | 19883 | 0.111268355405512 | 5.92062481556412 | 1.630792914885 | 0.1261956228756 | 0.197345401094888 |
| Ift57 | ENSMUSG00000032965 | 73916 | 0.267482814608202 | 1.5914708970219 | 1.62954710561736 | 0.1264608260225 | 0.19770382368582 |
| Slc44a2 | ENSMUSG00000057193 | 68682 | 0.139003065324496 | 6.29667605227202 | 1.62954631304431 | 0.1264609948962 | 0.19770382368582 |
| Chd2 | ENSMUSG00000078671 | 244059 | -0.163531087968686 | 5.55737364576365 | -1.62864011748904 | 0.1266542062080 | 0.1979775672901 |
| Gnaq | ENSMUSG00000024639 | 14682 | 0.13923863791705 | 3.92535917757191 | 1.62847008430672 | 0.1266904877512 | 0.197994501605187 |
| Vps25 | ENSMUSG00000078656 | 28084 | -0.127023811489468 | 3.73492006838521 | -1.62841958022582 | 0.1267012660075 | 0.197994501605187 |
| Fam174a | ENSMUSG00000051185 | 67698 | -0.123197881246582 | 4.82261797393248 | -1.62700607571395 | 0.1270032498104 | 0.19843803940516 |
| Ercc2 | ENSMUSG00000030400 | 13871 | 0.181360092867783 | 2.97412696144309 | 1.62682951408402 | 0.1270410145866 | 0.198468676639501 |
| Hsd17b7 | ENSMUSG00000026675 | 15490 | 0.458428340141242 | 0.953562832985694 | 1.62669349704836 | 0.1270701139163 | 0.198485769537015 |
| Ccser2 | ENSMUSG00000058690 | 72972 | 0.108964042769173 | 6.92270576909167 | 1.62614808693952 | 0.1271868563864 | 0.198639738128387 |
| Wipf2 | ENSMUSG00000038013 | 68524 | -0.183232314549396 | 2.90906746722726 | -1.62507579544014 | 0.1274166464573 | 0.198970194632135 |
| Dido1 | ENSMUSG00000038914 | 23856 | -0.123336353827579 | 5.41033043772353 | -1.62481843855168 | 0.1274718510771 | 0.199027968022314 |
| Oaz2 | ENSMUSG00000040652 | 18247 | -0.0934359786459185 | 5.89096744293284 | -1.62442846456958 | 0.1275555423759 | 0.199130196188465 |
| Acin1 | ENSMUSG00000022185 | 56215 | -0.0897068775785588 | 6.02093804634345 | -1.62322346716051 | 0.1278144446849 | 0.199505882457678 |
| Syt7 | ENSMUSG00000024743 | 54525 | -0.439100111279181 | 1.7533451455186 | -1.62290353476663 | 0.1278832608952 | 0.199584797950574 |
| Rai1 | ENSMUSG00000062115 | 19377 | 0.121866243956152 | 4.76231713411809 | 1.62252214931316 | 0.1279653373586 | 0.199684382964691 |
| Myzap | ENSMUSG00000041361 | 102371 | 0.174584237189891 | 3.14054473451189 | 1.62190076320491 | 0.1280991611674 | 0.199864677522299 |
| Pus7 | ENSMUSG00000057541 | 78697 | 0.158321289858676 | 3.3245426342366 | 1.62067061091551 | 0.1283644484959 | 0.200244068437908 |
| Ppp3r1 | ENSMUSG00000033953 | 19058 | 0.103680312135839 | 6.09436690654626 | 1.62060337537732 | 0.1283789618161 | 0.200244068437908 |
| Zfhx2 | ENSMUSG00000040721 | 239102 | 0.139829418337408 | 3.3859327375874 | 1.62038008573122 | 0.1284271708406 | 0.200290684043242 |
| Zfp579 | ENSMUSG00000051550 | 68490 | 0.21837416143472 | 2.74626086328767 | 1.61969128833438 | 0.1285759834169 | 0.20049416187317 |
| Phf23 | ENSMUSG00000018572 | 78246 | 0.162321326561346 | 3.04059578336041 | 1.61930479990374 | 0.1286595484374 | 0.2005958527984 |
| Fhod1 | ENSMUSG00000014778 | 234686 | 0.188610001467354 | 3.86010608617227 | 1.61809168982744 | 0.1289221478573 | 0.200976611270556 |
| Fmnl2 | ENSMUSG00000036053 | 71409 | 0.246629414935511 | 2.82369665740964 | 1.61731032173024 | 0.1290915345332 | 0.201211972619864 |
| Olfr648 | ENSMUSG00000042909 | 258746 | -1.33723511185049 | -1.57458449904601 | -1.61672624843683 | 0.1292182769819 | 0.201380807768767 |
| Adamts10 | ENSMUSG00000024299 | 224697 | 0.353736701205203 | 2.8832462049872 | 1.61650156121916 | 0.1292670622210 | 0.201428119335531 |
| Angel2 | ENSMUSG00000026634 | 52477 | -0.153246943395078 | 3.96135273584495 | -1.61574135695789 | 0.1294322398655 | 0.201656757977475 |
| Ube2d2a | ENSMUSG00000091896 | 56550 | 0.111131703759984 | 4.85034363149377 | 1.61551431695913 | 0.1294816066558 | 0.201704922667077 |
| Siae | ENSMUSG00000001942 | 22619 | 0.253591658440808 | 1.91629990280437 | 1.61532710718974 | 0.1295223251525 | 0.201739603340406 |
| Mylip | ENSMUSG00000038175 | 218203 | 0.272566461826125 | 3.81945522670386 | 1.61510210246862 | 0.1295712787845 | 0.201787099073097 |
| Sp100 | ENSMUSG00000026222 | 20684 | -0.306639984926087 | 3.59619797505041 | -1.6149225288005 | 0.1296103595967 | 0.201819208084302 |
| Gnai1 | ENSMUSG00000057614 | 14677 | -0.285318806184135 | 1.38886158389559 | -1.61338057334368 | 0.1299463569200 | 0.202298018475939 |
| Wnt5a | ENSMUSG00000021994 | 22418 | -0.212895875208973 | 2.04885356450365 | -1.61330129719971 | 0.1299636518029 | 0.202298018475939 |
| Olfr1381 | ENSMUSG00000095187 | 258461 | -1.1237070179154 | 0.576797046908848 | -1.61325672302479 | 0.1299733769789 | 0.202298018475939 |
| Slc9a3r1 | ENSMUSG00000020733 | 26941 | 0.261943821091768 | 1.59744158111875 | 1.6124324028063 | 0.1301533403060 | 0.202549282870998 |
| Slbp | ENSMUSG00000004642 | 20492 | -0.249915204569475 | 2.51177079861224 | -1.61115603763461 | 0.1304324179174 | 0.202954698968791 |
| Clec14a | ENSMUSG00000045930 | 66864 | -0.294990992466486 | 2.37264215083415 | -1.61071202013896 | 0.1305296236782 | 0.203077044751939 |
| Bhlhe41 | ENSMUSG00000030256 | 79362 | 0.283936223330559 | 4.41933495825957 | 1.61058231522761 | 0.1305580309163 | 0.203092334701448 |
| Rnf123 | ENSMUSG00000041528 | 84585 | 0.109396307847696 | 5.47130028043927 | 1.61046218038502 | 0.1305843469387 | 0.203104367727242 |
| Ogt | ENSMUSG00000034160 | 108155 | 0.158333196390461 | 5.59493864498632 | 1.61015180463105 | 0.1306523572254 | 0.203181237278584 |
| Polr3e | ENSMUSG00000030880 | 26939 | 0.166312285684739 | 2.82512289084583 | 1.6099436381059 | 0.1306979883282 | 0.20322328739917 |
| Atf2 | ENSMUSG00000027104 | 11909 | -0.146873777381575 | 4.03392486325636 | -1.60969991850224 | 0.1307514303391 | 0.203277469070879 |
| Znhit1 | ENSMUSG00000059518 | 70103 | -0.166998121664839 | 3.30559581878048 | -1.60845010935749 | 0.1310257810809 | 0.20365158113058 |
| Tmub2 | ENSMUSG00000034757 | 72053 | 0.187131056454594 | 2.67295544878946 | 1.60843397206754 | 0.1310293266939 | 0.20365158113058 |
| Ranbp3l | ENSMUSG00000048424 | 223332 | -0.737725135493412 | 0.669370923260899 | -1.60781546537724 | 0.1311652847003 | 0.203817002018611 |
| Plekha5 | ENSMUSG00000030231 | 109135 | 0.135901570764483 | 3.40473199306039 | 1.60778015597166 | 0.1311730499680 | 0.203817002018611 |
| Ube2m | ENSMUSG00000005575 | 22192 | -0.103138589521887 | 5.45152714789759 | -1.60664198909528 | 0.1314235691835 | 0.204177236319742 |
| Scaf4 | ENSMUSG00000022983 | 224432 | 0.108737719008222 | 4.20946670423679 | 1.60639110175663 | 0.1314788469551 | 0.204234087831015 |
| Zfp764 | ENSMUSG00000045757 | 233893 | 0.312609404294086 | 1.15818959719199 | 1.60563889123882 | 0.1316447011610 | 0.204462663880508 |
| Ier3ip1 | ENSMUSG00000090000 | 66191 | -0.116282435223595 | 4.50182140105814 | -1.6052279098698 | 0.1317353943639 | 0.204574456001228 |
| Zfp865 | ENSMUSG00000074405 | 319748 | 0.153378495701628 | 3.7663924170338 | 1.60407977192143 | 0.1319890453263 | 0.204939240690645 |
| Mtmr3 | ENSMUSG00000034354 | 74302 | -0.0911107404753597 | 6.07878033294421 | -1.60373404106856 | 0.1320655080395 | 0.205028840843728 |
| Inka2 | ENSMUSG00000048458 | 109050 | -0.296260922501061 | 2.6050690753688 | -1.60274873110162 | 0.1322836314908 | 0.205338309546558 |
| Fbxo11 | ENSMUSG00000005371 | 225055 | -0.10034399656573 | 5.22540653021298 | -1.60243455084628 | 0.1323532486385 | 0.20541720301974 |
| Srfbp1 | ENSMUSG00000024528 | 67222 | -0.185113018654538 | 3.03835436561293 | -1.60148250646061 | 0.1325643990255 | 0.205675623902166 |
| Tm6sf1 | ENSMUSG00000038623 | 107769 | 0.216951456960248 | 3.10234979324891 | 1.6014824577563 | 0.1325644098349 | 0.205675623902166 |
| Kdm8 | ENSMUSG00000030752 | 77035 | -0.389772951885551 | 1.3150589563887 | -1.60142933474136 | 0.1325762003489 | 0.205675623902166 |
| Idua | ENSMUSG00000033540 | 15932 | -0.261814860425073 | 2.67976079635704 | -1.60127191989364 | 0.1326111434754 | 0.205682910473457 |
| Pabpn1 | ENSMUSG00000022194 | 54196 | -0.125714654396251 | 4.1218392299736 | -1.60118919846287 | 0.1326295092573 | 0.205682910473457 |
| Ccz1 | ENSMUSG00000029617 | 231874 | -0.137174199892226 | 4.50771376311974 | -1.60115390114869 | 0.1326373466222 | 0.205682910473457 |
| N4bp2 | ENSMUSG00000037795 | 333789 | 0.139188777721863 | 3.45210690362877 | 1.6004875849099 | 0.1327853695025 | 0.205883244543548 |
| Lama5 | ENSMUSG00000015647 | 16776 | 0.180218142233866 | 4.51674069575826 | 1.59975565578505 | 0.1329481325835 | 0.206106373176959 |
| Phlpp2 | ENSMUSG00000031732 | 244650 | 0.215909807281206 | 1.94907628501689 | 1.59960870970573 | 0.1329808305215 | 0.206127830180261 |
| Edc4 | ENSMUSG00000036270 | 234699 | 0.0988147706373255 | 4.48054847530642 | 1.59917220203193 | 0.1330780016215 | 0.206249203987717 |
| Gnb2 | ENSMUSG00000029713 | 14693 | 0.101171511706085 | 4.91911636547581 | 1.59762969340424 | 0.1334218706002 | 0.206752830667844 |
| Slc52a2 | ENSMUSG00000022560 | 52710 | 0.275661003507333 | 1.0522700207623 | 1.59721512846658 | 0.1335144194183 | 0.206866919725291 |
| Cnot3 | ENSMUSG00000035632 | 232791 | 0.136991446339551 | 4.08015414245278 | 1.59706817923003 | 0.1335472381250 | 0.206888443869703 |
| Ezh1 | ENSMUSG00000006920 | 14055 | -0.0937682662835337 | 4.70918876400772 | -1.59650807364729 | 0.1336723923177 | 0.207035338393541 |
| Jph2 | ENSMUSG00000017817 | 59091 | -0.145062630647263 | 7.30045319046147 | -1.59647431213428 | 0.1336799394732 | 0.207035338393541 |
| Fkbp8 | ENSMUSG00000019428 | 14232 | 0.0970368476617338 | 5.295268098223 | 1.59529457876385 | 0.1339438915567 | 0.207414744100628 |
| Blzf1 | ENSMUSG00000026577 | 66352 | -0.14169579214155 | 3.95425668375529 | -1.59475009490154 | 0.1340658651129 | 0.207574216933437 |
| Igsf8 | ENSMUSG00000038034 | 140559 | -0.282279891817269 | 3.49311418461853 | -1.59360599293442 | 0.1343224751060 | 0.20794207270698 |
| Cdk11b | ENSMUSG00000029062 | 12537 | 0.126243785202677 | 5.39701218081072 | 1.59286434820727 | 0.1344890441518 | 0.208163737082544 |
| Mesd | ENSMUSG00000038503 | 67943 | 0.12158513580116 | 3.92057634771531 | 1.59279891539537 | 0.1345037485146 | 0.208163737082544 |
| Npr1 | ENSMUSG00000027931 | 18160 | -0.198963973654108 | 3.23856668025104 | -1.58982286597055 | 0.1351740053361 | 0.209151899387363 |
| Smg7 | ENSMUSG00000042772 | 226517 | 0.129356206309535 | 5.245599362802 | 1.58972659587549 | 0.1351957349255 | 0.209151899387363 |
| Dnajc13 | ENSMUSG00000032560 | 235567 | -0.100156114452859 | 5.42400112108203 | -1.58969881289196 | 0.1352020065164 | 0.209151899387363 |
| Serpinb9 | ENSMUSG00000045827 | 20723 | 0.273947950040774 | 1.69228527836611 | 1.58957337808386 | 0.1352303246567 | 0.209151899387363 |
| Wdcp | ENSMUSG00000051721 | 238037 | -0.226509792985288 | 2.04134333284395 | -1.58953977032866 | 0.1352379128048 | 0.209151899387363 |
| Znrf3 | ENSMUSG00000041961 | 407821 | 0.231204387307314 | 2.67820051126247 | 1.58924189409708 | 0.1353051849997 | 0.209226337138576 |
| Gucy1a2 | ENSMUSG00000041624 | 234889 | -0.298181650554111 | 1.73713341327166 | -1.58847501693148 | 0.1354785086411 | 0.209464721068801 |
| Olfr1250 | ENSMUSG00000075078 | 258967 | -0.889374565088756 | 1.39193229843823 | -1.58802311202554 | 0.1355807341429 | 0.209593127551463 |
| Mphosph10 | ENSMUSG00000030521 | 67973 | -0.131356128198396 | 3.90904124953683 | -1.58782820275322 | 0.1356248450946 | 0.209631671624652 |
| Csnk1d | ENSMUSG00000025162 | 104318 | 0.113786789400172 | 4.6613156723415 | 1.58735329112718 | 0.1357323765732 | 0.209768218340467 |
| Fam204a | ENSMUSG00000057858 | 76539 | -0.16717262668576 | 3.61025586008627 | -1.58696835793538 | 0.1358195885786 | 0.209873328068036 |
| Mfhas1 | ENSMUSG00000070056 | 52065 | 0.247403065355861 | 2.69863701664413 | 1.58681861521761 | 0.1358535279118 | 0.20989610086289 |

| | | | | | | | |
|---|---|---|---|---|---|---|---|
| Hmgcl | ENSMUSG00000028672 | 15356 | -0.142182411135903 | 3.70609039411401 | -1.58664249149945 | 0.1358934558625 | 0.209917229180776 |
| Lgals3bp | ENSMUSG00000033880 | 19039 | 0.308500066711046 | 3.71427626064669 | 1.58655612255627 | 0.1359130397362 | 0.209917229180776 |
| Tiparp | ENSMUSG00000034640 | 99929 | 0.292472656212455 | 3.40081433029623 | 1.58650419822124 | 0.1359248145770 | 0.209917229180776 |
| Sgo2a | ENSMUSG00000026039 | 68549 | -0.206785884922451 | 2.47315635585563 | -1.58607507384751 | 0.1360221603759 | 0.210037891661179 |
| Gja1 | ENSMUSG00000050953 | 14609 | 0.218684185595643 | 3.71288340996922 | 1.58598728844036 | 0.1360420816652 | 0.210038982299765 |
| Tsfm | ENSMUSG00000040521 | 66399 | -0.167639590126758 | 2.74397136925309 | -1.58583608100249 | 0.1360764013065 | 0.2100622994891 |
| Ctsl | ENSMUSG00000021477 | 13039 | 0.139402716312155 | 5.85753462327916 | 1.58573190318994 | 0.1361000509391 | 0.21006914103577 |
| Maff | ENSMUSG00000042622 | 17133 | -0.289740602299013 | 1.52885737293401 | -1.58538249372261 | 0.1361793969609 | 0.210161935363519 |
| Wasl | ENSMUSG00000029684 | 73178 | 0.100979026860591 | 4.34903714609119 | 1.58507103468892 | 0.1362501585051 | 0.210241457173886 |
| Mtf1 | ENSMUSG00000028890 | 17764 | -0.176852763918146 | 4.2104016861956 | -1.5849647932759 | 0.1362743031219 | 0.210249034216869 |
| Exoc5 | ENSMUSG00000061244 | 105504 | 0.107322728380463 | 4.8286942662267 | 1.58402355678788 | 0.1364883709931 | 0.210549588389251 |
| Nynrin | ENSMUSG00000075592 | 277154 | 0.258590280930811 | 2.40455639276706 | 1.58372109256215 | 0.1365572225841 | 0.210626075923163 |
| Slc2a1 | ENSMUSG00000028645 | 20525 | 0.205666166201218 | 3.40542291793796 | 1.58353561391415 | 0.1365994588764 | 0.210661496187733 |
| Wsb1 | ENSMUSG00000017677 | 78889 | 0.311913629366945 | 2.98635746163567 | 1.58253943547105 | 0.1368264959739 | 0.210981863096568 |
| Art5 | ENSMUSG00000070424 | 11875 | -0.264838812348089 | 3.44628062464774 | -1.58075619910353 | 0.1372337199978 | 0.211579942637006 |
| Srebf1 | ENSMUSG00000020538 | 20787 | 0.159781720013381 | 4.10832231392662 | 1.58059142950631 | 0.1372713996801 | 0.211608189240434 |
| Zfp408 | ENSMUSG00000075040 | 381410 | 0.253331680555269 | 1.80844072906636 | 1.58047031401826 | 0.1372991021642 | 0.211621049881168 |
| Rnf219 | ENSMUSG00000022120 | 72486 | -0.258518301008462 | 1.63129216593605 | -1.57973837324943 | 0.1374666196524 | 0.211849375358997 |
| Btbd7 | ENSMUSG00000041702 | 238386 | -0.100639221581793 | 4.90964588203984 | -1.57950545203401 | 0.1375199645784 | 0.211901710291344 |
| Mfsd5 | ENSMUSG00000045665 | 106073 | 0.230605175164735 | 2.46145864964213 | 1.57861625793688 | 0.1377237763756 | 0.212185849127877 |
| Dyrk2 | ENSMUSG00000028630 | 69181 | 0.192664741500777 | 5.08583433430544 | 1.57727677863978 | 0.1380312876284 | 0.212629651221367 |
| Mapk8ip3 | ENSMUSG00000024163 | 30957 | 0.198775233885994 | 4.05645694490006 | 1.57491636177286 | 0.1385746158609 | 0.213436540224885 |
| Gtf2h5 | ENSMUSG00000034345 | 66467 | -0.144734186976382 | 3.4406572792214 | -1.5746450450386 | 0.1386371858872 | 0.213502828815681 |
| Ncoa3 | ENSMUSG00000027678 | 17979 | -0.128953813457129 | 5.16085092547252 | -1.57446262996909 | 0.1386792673826 | 0.213537550607037 |
| Gatad2a | ENSMUSG00000036180 | 234366 | 0.0948439955830116 | 4.86803638712859 | 1.57367754095886 | 0.1388605054094 | 0.213786504877576 |
| Degs1 | ENSMUSG00000038633 | 13244 | 0.154368070896817 | 3.97490839159943 | 1.57321609504057 | 0.1389671251455 | 0.213920524569284 |
| Ankrd12 | ENSMUSG00000034647 | 106585 | 0.149182553951875 | 4.67867643150786 | 1.57260975547603 | 0.1391073301755 | 0.21408449389935 |
| Bcr | ENSMUSG00000009681 | 110279 | 0.167231179283283 | 3.09417528396092 | 1.57258605426049 | 0.1391128131156 | 0.21408449389935 |
| Tfe3 | ENSMUSG00000000134 | 209446 | 0.211801467740472 | 3.16756765115998 | 1.57249042593508 | 0.1391349372557 | 0.214088400780164 |
| Afap1l1 | ENSMUSG00000033032 | 106877 | 0.139559943729217 | 5.62203004728866 | 1.57214174948551 | 0.1392156310438 | 0.214182415614465 |
| Pfas | ENSMUSG00000020899 | 237823 | -0.171959087635128 | 3.21124043857375 | -1.57124664601993 | 0.1394229676440 | 0.214421295760329 |
| 1110002L01Rik | ENSMUSG00000071456 | 100043040 | -0.162731162785047 | 2.33476123163593 | -1.57122806792 | 0.1394272737719 | 0.214421295760329 |
| Uvrag | ENSMUSG00000035354 | 78610 | 0.127413377550369 | 3.73438382839994 | 1.57118836848129 | 0.1394364758952 | 0.214421295760329 |
| Trabd2b | ENSMUSG00000070867 | 666048 | -0.217964132936591 | 2.04902017444841 | -1.5711327733987 | 0.1394493634214 | 0.214421295760329 |
| St3gal2 | ENSMUSG00000031749 | 20444 | -0.119058385645076 | 5.45735483652545 | -1.57045575924026 | 0.1396063844257 | 0.214632544044746 |
| Erh | ENSMUSG00000021131 | 13877 | 0.161981570484166 | 3.29948414371265 | 1.56805047963059 | 0.1401654716063 | 0.215450602244606 |
| Slc38a6 | ENSMUSG00000044712 | 625098 | 0.204036995348354 | 5.37673438800052 | 1.56799708138212 | 0.1401779053301 | 0.215450602244606 |
| Atp6v0c | ENSMUSG00000024121 | 11984 | 0.107660178812066 | 5.5901971092565 | 1.56698573994095 | 0.1404135736435 | 0.215782479052072 |
| Nudt22 | ENSMUSG00000037349 | 68323 | 0.200429573254514 | 1.71349354071255 | 1.56605806087141 | 0.1406300449744 | 0.216084765478714 |
| Ankzf1 | ENSMUSG00000026199 | 52231 | -0.17658633195505 | 2.05143855000286 | -1.56592013232717 | 0.1406622546218 | 0.216095395124976 |
| Dctn5 | ENSMUSG00000030868 | 59288 | 0.176304034592586 | 2.81439269897325 | 1.56585913522716 | 0.1406765009339 | 0.216095395124976 |
| Snapin | ENSMUSG00000001018 | 20615 | -0.104909068506879 | 5.03325990435484 | -1.56534037630475 | 0.1407977107865 | 0.2162511980621 |
| Pole4 | ENSMUSG00000030042 | 66979 | 0.196380507375775 | 2.69135339677572 | 1.56416978417678 | 0.1410715525190 | 0.216641351585556 |
| Eef1g | ENSMUSG00000071644 | 67160 | -0.117631392621881 | 7.73006218911502 | -1.5632539368784 | 0.1412861186170 | 0.216940379632324 |
| Nqo2 | ENSMUSG00000046949 | 18105 | -0.15230008186276 | 3.57174836385969 | -1.56312238925478 | 0.1413169607357 | 0.21695726092728 |
| Nup98 | ENSMUSG00000063550 | 269966 | 0.103661973803579 | 5.05995896392039 | 1.56181254884047 | 0.1416243751716 | 0.217398686280178 |
| Kctd10 | ENSMUSG00000001098 | 330171 | -0.196753699257131 | 4.04226496411269 | -1.56149598114801 | 0.1416987582172 | 0.217482326042208 |
| Cetn2 | ENSMUSG00000031347 | 26370 | -0.209481789811411 | 2.85484064961341 | -1.56095959790426 | 0.1418248670555 | 0.21764532104223 |
| Myh6 | ENSMUSG00000040752 | 17888 | -0.183398944316563 | 3.99483112403229 | -1.559678833126 | 0.1421263756984 | 0.218077402127911 |
| Bdp1 | ENSMUSG00000049658 | 544971 | -0.129815431698299 | 5.25273016939615 | -1.559477221417 | 0.1421738876416 | 0.218096812797883 |
| Sdccag8 | ENSMUSG00000026504 | 76816 | 0.161442198622577 | 2.38300828744251 | 1.55939649670025 | 0.1421929150892 | 0.218096812797883 |
| Gmeb2 | ENSMUSG00000038705 | 229004 | -0.211090576415463 | 2.61271449202065 | -1.55937118154391 | 0.1421988825185 | 0.218096812797883 |
| Sbf1 | ENSMUSG00000036529 | 77980 | 0.104789478422194 | 5.73941553807621 | 1.55862167620607 | 0.1423756571900 | 0.218337304818202 |
| Dut | ENSMUSG00000027203 | 110074 | -0.320827633843041 | 1.18458357757383 | -1.55792381560422 | 0.1425404201679 | 0.218559311664522 |
| Med6 | ENSMUSG00000002679 | 69792 | -0.191083268873594 | 1.68213264061539 | -1.55567793672574 | 0.1430717734096 | 0.219343275615825 |
| Ankle2 | ENSMUSG00000029501 | 71782 | -0.0945950259887134 | 4.9095467134787 | -1.55532298771341 | 0.1431559057103 | 0.21944148160431 |
| Ppp1r11 | ENSMUSG00000036398 | 76497 | -0.157246127366798 | 3.34333642876415 | -1.55426714468613 | 0.1434064184892 | 0.219794666363635 |
| Tnfrsf10b | ENSMUSG00000022074 | 21933 | 0.410271428037748 | 1.35675353325815 | 1.5532696640801 | 0.1436434281128 | 0.22010983409482 |
| Abcc5 | ENSMUSG00000022822 | 27416 | 0.170636359696168 | 4.72985230252653 | 1.55323225109956 | 0.1436523242550 | 0.22010983409482 |
| Olfr393 | ENSMUSG00000094488 | 259010 | -0.755177910661131 | 1.70296319491073 | -1.55211451209742 | 0.1439183199591 | 0.22048649691295 |
| 9530082P21Rik | ENSMUSG00000097006 | 638247 | -0.361677229819001 | 1.25759829236554 | -1.55181102014284 | 0.1439906165366 | 0.220549513482167 |
| Eps8l2 | ENSMUSG00000025504 | 98845 | -0.286607150508168 | 1.17342675292344 | -1.55177245389379 | 0.1439998058477 | 0.220549513482167 |
| Atp2b1 | ENSMUSG00000019943 | 67972 | 0.0889056181512657 | 6.39294918027789 | 1.54996257293703 | 0.1444316160834 | 0.221167541862516 |
| Supt20 | ENSMUSG00000027751 | 56790 | 0.184559637314605 | 3.28909567119169 | 1.54991160951121 | 0.1444437911770 | 0.221167541862516 |
| Rnf113a2 | ENSMUSG00000098134 | 66381 | -0.233284725373199 | 1.91489363520312 | -1.54942002307643 | 0.1445612755176 | 0.221295449469867 |
| Usp5 | ENSMUSG00000038429 | 22225 | 0.132050615489211 | 5.24726197724006 | 1.549392662896 | 0.1445678167289 | 0.221295449469867 |
| Gorasp1 | ENSMUSG00000032513 | 74498 | 0.16542341811224 | 3.07825190679581 | 1.54901394125864 | 0.1446583866225 | 0.221403083754038 |
| Rad1 | ENSMUSG00000022248 | 19355 | -0.325334998990103 | 1.48542377640221 | -1.5481388808182 | 0.1448678394213 | 0.221692615527685 |
| Paxip1 | ENSMUSG00000002221 | 55982 | -0.188270848095992 | 3.54387658008114 | -1.54744967900275 | 0.1450329876479 | 0.221914276033006 |
| Cog8 | ENSMUSG00000031916 | 97484 | -0.163814862786497 | 3.77661422534545 | -1.54702743645253 | 0.1451342458022 | 0.22203813028195 |
| Mcm7 | ENSMUSG00000029730 | 17220 | 0.227909062474395 | 2.46275050672582 | 1.54640049741554 | 0.1452847035967 | 0.222237208930395 |
| Kif2a | ENSMUSG00000021693 | 16563 | -0.113246134314682 | 3.90199864316866 | -1.54607901260678 | 0.1453619076787 | 0.222324193764739 |
| Aga | ENSMUSG00000031521 | 11593 | -0.208213208007738 | 2.0613430189794 | -1.54599339724356 | 0.1453824739782 | 0.222324541557935 |
| Rab11fip2 | ENSMUSG00000040022 | 74998 | -0.13353653229131 | 3.10370873204924 | -1.54501620769584 | 0.1456173878412 | 0.222652632045451 |
| Thumpd1 | ENSMUSG00000030942 | 233802 | -0.123991652703762 | 4.18673464903088 | -1.54438404226793 | 0.1457695312309 | 0.222854090333608 |
| Zfp687 | ENSMUSG00000019338 | 78266 | 0.177323449144076 | 2.60484144169487 | 1.54297528374048 | 0.1461090652461 | 0.223341937100467 |
| Uqcc3 | ENSMUSG00000071654 | 107197 | 0.179751972477316 | 3.62736684121642 | 1.54230584875207 | 0.1462706463432 | 0.223557667111045 |
| Caml | ENSMUSG00000021501 | 12328 | 0.236115595268142 | 2.31075258162899 | 1.54160072231289 | 0.1464410069909 | 0.223786753448685 |
| Hist1h2be | ENSMUSG00000047246 | 319179 | 0.160200417098216 | 3.10073748173046 | 1.5412312800807 | 0.1465303328429 | 0.223866376857232 |
| Atf7ip | ENSMUSG00000030213 | 54343 | 0.120631302714167 | 5.89083943034696 | 1.54120785046498 | 0.1465359993549 | 0.223866376857232 |
| 2810004N23Rik | ENSMUSG00000031984 | 66523 | -0.127041816122625 | 5.13381037598389 | -1.54113115102221 | 0.1465545506166 | 0.223866376857232 |
| Dpy19l4 | ENSMUSG00000045205 | 381510 | -0.141709609319032 | 4.07602151467621 | -1.54097025454464 | 0.1465934730838 | 0.223894544401208 |
| Nup62 | ENSMUSG00000109511 | 18226 | -0.142823885180586 | 3.99561433206284 | -1.54015053850403 | 0.1467919074085 | 0.224166295038141 |
| Csnk1g1 | ENSMUSG00000032384 | 214897 | 0.135297907476112 | 3.1352250971233 | 1.5399987607323 | 0.1468286744083 | 0.224191121659189 |
| Ssna1 | ENSMUSG00000026966 | 68475 | 0.122595794467871 | 3.07997807137298 | 1.53980447116966 | 0.1468757510157 | 0.224231680775507 |
| Parp9 | ENSMUSG00000022906 | 80285 | 0.219433394726888 | 3.32669975088379 | 1.53922863071684 | 0.1470153534461 | 0.224407307953937 |
| Ip6k2 | ENSMUSG00000032599 | 76500 | 0.205948943490248 | 2.65076456836996 | 1.53916062035178 | 0.1470318488304 | 0.224407307953937 |
| Parg | ENSMUSG00000021911 | 26430 | -0.101547571634403 | 4.48640924612275 | -1.53891584496423 | 0.1470912302460 | 0.224466597489772 |
| Zfp825 | ENSMUSG00000069208 | 235956 | -0.213064672633631 | 1.60353243069596 | -1.53702660085752 | 0.1475502402011 | 0.225106187664403 |
| Timm13 | ENSMUSG00000020219 | 30055 | -0.126687047245698 | 3.62214129424233 | -1.53702127658594 | 0.1475515355059 | 0.225106187664403 |
| Zfp72 | ENSMUSG00000069184 | 238722 | -0.395668406848739 | 0.974925087713425 | -1.53662938092498 | 0.1476469036504 | 0.225220248925942 |
| Mir22hg | ENSMUSG00000085148 | 100042498 | 0.276983018006987 | 3.37260147027317 | 1.5362127844529 | 0.1477483403119 | 0.225341480535276 |
| Las1l | ENSMUSG00000057421 | 76130 | -0.151772845497634 | 4.65856141171567 | -1.53613367678287 | 0.1477676088625 | 0.225341480535276 |
| Prpsap2 | ENSMUSG00000020528 | 212627 | 0.291151885384327 | 1.39571381040525 | 1.53600923102802 | 0.1477979249104 | 0.225356272450275 |
| Hist1h3e | ENSMUSG00000069273 | 319151 | 0.347408312167968 | 1.37402886334222 | 1.53575493090944 | 0.1478598910575 | 0.225419312294299 |
| Pbx2 | ENSMUSG00000034673 | 18515 | 0.164797147429215 | 2.75541569232895 | 1.53528964122235 | 0.1479733269903 | 0.225556719228987 |
| Epc1 | ENSMUSG00000024240 | 13831 | -0.129587510696697 | 4.32289181002279 | -1.53521598894627 | 0.1479912899378 | 0.225556719228987 |
| E4f1 | ENSMUSG00000024137 | 13560 | -0.140276656260125 | 3.38616881298558 | -1.53503133507809 | 0.1480363330649 | 0.22556417432552 |
| Cacna2d1 | ENSMUSG00000040118 | 12293 | -0.123533121887402 | 7.67269274058721 | -1.53502674907377 | 0.1480374518901 | 0.22556417432552 |
| Notch3 | ENSMUSG00000038146 | 18131 | -0.203373905496217 | 5.14204775191367 | -1.5342614432873 | 0.1482242606803 | 0.22580321371136 |
| Mrps2 | ENSMUSG00000035772 | 118451 | -0.171750301145261 | 3.54739197617438 | -1.53421482153713 | 0.1482356473874 | 0.22580321371136 |
| Ppp1ccb | ENSMUSG00000100153 | 434233 | -0.182947578889454 | 3.28115940918966 | -1.53412452735063 | 0.1482577025906 | 0.225805343042867 |
| Igip | ENSMUSG00000110185 | 109169 | 0.173489962545476 | 2.81367444385003 | 1.53377729541396 | 0.1483425432761 | 0.225900133971894 |
| Clcn7 | ENSMUSG00000036636 | 26373 | 0.108742236194492 | 4.09089468306711 | 1.53370066707118 | 0.1483612717760 | 0.225900133971894 |
| Myo6 | ENSMUSG00000033577 | 17920 | -0.130107948591462 | 3.73301676658826 | -1.53328509135089 | 0.1484628764250 | 0.226023356852648 |
| Terf2 | ENSMUSG00000031921 | 21750 | -0.135987628272343 | 3.02202845954615 | -1.53296502367757 | 0.1485411705468 | 0.226111061864248 |
| Gpn1 | ENSMUSG00000064037 | 74254 | -0.14419764194105 | 3.23496523247386 | -1.53278001999192 | 0.1485864416884 | 0.226148481494932 |
| Olfr982 | ENSMUSG00000049010 | 258853 | -1.15523759473806 | -1.29421075823064 | -1.53258215720633 | 0.1486348724951 | 0.226190699045573 |
| Irak1 | ENSMUSG00000031392 | 16179 | 0.0933185400610439 | 5.47759842406855 | 1.53181491795142 | 0.1488227964205 | 0.226434226681036 |
| 4931406C07Rik | ENSMUSG00000031938 | 70984 | 0.189306151128422 | 2.17970186318965 | 1.53175969775685 | 0.1488363295858 | 0.226434226681036 |
| Ube3a | ENSMUSG00000025326 | 22215 | -0.13055432091699 | 6.66034181079745 | -1.53152541332865 | 0.1488937588091 | 0.226490074804179 |
| Apmap | ENSMUSG00000033096 | 71881 | -0.17477056503026 | 2.93482517312132 | -1.53012250239364 | 0.1492380433789 | 0.226982197325069 |
| Suz12 | ENSMUSG00000017548 | 52615 | -0.115190732250118 | 4.05635443439852 | -1.52975167336828 | 0.1493291605742 | 0.227089183950597 |
| Dhx32 | ENSMUSG00000030986 | 101437 | 0.117220544750596 | 5.53802467009622 | 1.52936692134907 | 0.1494237488170 | 0.227201418600555 |
| 6030458C11Rik | ENSMUSG00000022195 | 77877 | -0.0954622850336388 | 4.2360276391881 | -1.52848845759353 | 0.1496399031390 | 0.227498439667958 |
| Mbnl1 | ENSMUSG00000027763 | 56758 | -0.100102635535398 | 7.69956951774219 | -1.52776332196356 | 0.1498185298437 | 0.227738332599338 |

| | | | | | | | |
|---|---|---|---|---|---|---|---|
| Pes1 | ENSMUSG00000020430 | 64934 | 0.108864921450699 | 4.12645828172424 | 1.5272766809241 | 0.1499385084485 | 0.227889020557652 |
| Hps6 | ENSMUSG00000074811 | 20170 | 0.294097311268478 | 1.4346384246337 | 1.52703322111472 | 0.1499985627315 | 0.227921075425024 |
| Iba57 | ENSMUSG00000049287 | 216792 | -0.187689265589493 | 2.68949684340545 | -1.52702212387511 | 0.1500013005770 | 0.227921075425024 |
| Gmeb1 | ENSMUSG00000028901 | 56809 | -0.14386207156499 | 3.20400686443517 | -1.52685145971625 | 0.1500434111807 | 0.227953374234509 |
| Gm20604 | ENSMUSG00000041716 | 100859931 | -0.224364389339605 | 1.71423808113639 | -1.52657725236757 | 0.1501110916330 | 0.228024505647694 |
| Chp1 | ENSMUSG00000014077 | 56398 | -0.0943509154907023 | 5.1005770995295 | -1.52644632568811 | 0.1501434163894 | 0.228041918098175 |
| Cxcr2 | ENSMUSG00000026180 | 12765 | -0.542644411301535 | 1.11104569488288 | -1.52553628583008 | 0.1503682614689 | 0.228351690208047 |
| Gnptab | ENSMUSG00000035311 | 432486 | 0.114342685239241 | 3.61126209635734 | 1.5253256768075 | 0.1504203377388 | 0.228399043175882 |
| Abca6 | ENSMUSG00000044749 | 76184 | 0.219758496139413 | 2.31817279344194 | 1.52493527059882 | 0.1505169121105 | 0.228513939761243 |
| Prxl2b | ENSMUSG00000029059 | 66469 | 0.198187880245736 | 3.07922767114685 | 1.52378827286362 | 0.1508009484560 | 0.228913368639492 |
| Ap3b1 | ENSMUSG00000021686 | 11774 | 0.0984017652240015 | 4.74792906172814 | 1.52266593850108 | 0.1510793177899 | 0.229304085359844 |
| Siva1 | ENSMUSG00000064326 | 30954 | -0.17890899067348 | 2.9371418992257 | -1.5216863703459 | 0.1513226336909 | 0.229641497830844 |
| Derl2 | ENSMUSG00000018442 | 116891 | 0.162907679074884 | 2.69485015521902 | 1.52092667873907 | 0.1515115631914 | 0.229896293343426 |
| Malt1 | ENSMUSG00000032688 | 240354 | 0.208193215506509 | 2.41963157813404 | 1.52051691431187 | 0.1516135515654 | 0.230019116191772 |
| Wwc2 | ENSMUSG00000031563 | 52357 | -0.112182427371003 | 4.28565351411376 | -1.51998785230526 | 0.1517453186964 | 0.230177485978646 |
| Zcchc4 | ENSMUSG00000029179 | 78796 | -0.167588830413923 | 2.48308400680467 | -1.51988741269306 | 0.1517703449588 | 0.230177485978646 |
| Slc25a37 | ENSMUSG00000034248 | 67712 | 0.196307989040487 | 2.77384555889015 | 1.5198442109115 | 0.1517811105053 | 0.230177485978646 |
| Dynlrb1 | ENSMUSG00000047459 | 67068 | -0.145402908386077 | 4.67850247004611 | -1.51934502939436 | 0.1519055496446 | 0.230334243746077 |
| Zfp710 | ENSMUSG00000048897 | 209225 | -0.179536049959999 | 3.97218087541272 | -1.51923065083656 | 0.1519340748442 | 0.23034554398366 |
| Rps6ka5 | ENSMUSG00000021180 | 73086 | -0.307169334766247 | 3.42835351569454 | -1.51885488499487 | 0.1520278201670 | 0.230455706870899 |
| Lgi4 | ENSMUSG00000036560 | 243914 | 0.282514620123219 | 1.56165721271397 | 1.51869275887419 | 0.1520682822234 | 0.230485079448728 |
| Zfp101 | ENSMUSG00000055240 | 22643 | -0.222718838928892 | 2.17905389324755 | -1.51828437914656 | 0.1521702426467 | 0.230607642086782 |
| Srrm1 | ENSMUSG00000028809 | 51796 | -0.101308794418077 | 6.01999722809839 | -1.51766312979893 | 0.1523254615660 | 0.230810870582019 |
| Mpnd | ENSMUSG00000003199 | 68047 | -0.189141767466166 | 3.69637163947596 | -1.51687574374882 | 0.1525223826107 | 0.231077223051689 |
| Srrt | ENSMUSG00000037364 | 83701 | 0.135985659812261 | 4.60788362310274 | 1.51653716009669 | 0.1526071269238 | 0.231173573226789 |
| Acer2 | ENSMUSG00000038007 | 230379 | -0.204305857016169 | 1.65970691055724 | -1.51635165778477 | 0.1526535733399 | 0.231211890006805 |
| Srsf2 | ENSMUSG00000034120 | 20382 | -0.0916537349359396 | 5.11140349328809 | -1.5161054699785 | 0.1527152328121 | 0.231273234950103 |
| Adss | ENSMUSG00000015961 | 11566 | 0.25852332148737 | 2.70967501298324 | 1.51600148597467 | 0.1527412826815 | 0.231280642885757 |
| Zbtb7c | ENSMUSG00000044646 | 207259 | 0.367283143945116 | 1.0173860226542 | 1.51550940508549 | 0.1528646089269 | 0.231435324124773 |
| Rmdn3 | ENSMUSG00000070730 | 67809 | -0.188153156758681 | 2.21618491761834 | -1.51523205756048 | 0.1529341554793 | 0.231508551938086 |
| Tcp11l1 | ENSMUSG00000027175 | 320554 | -0.196313403934019 | 2.75562760871292 | -1.51512633470717 | 0.1529606731776 | 0.231516632304756 |
| Ifit2 | ENSMUSG00000045932 | 15958 | -0.390371951660726 | 3.40830262124465 | -1.5149343922914 | 0.1530088266664 | 0.231557453176047 |
| Romo1 | ENSMUSG00000067847 | 67067 | -0.150426128267774 | 3.45171187594746 | -1.51432283427886 | 0.1531623367743 | 0.231742956377618 |
| Olfr825 | ENSMUSG00000058084 | 258672 | -0.728156104316687 | -0.344798186553774 | -1.51427716864441 | 0.1531738047597 | 0.231742956377618 |
| Katnal1 | ENSMUSG00000041298 | 231912 | -0.134638255436739 | 3.40645940477177 | -1.5140272307385 | 0.1532365844188 | 0.231805854453711 |
| Txndc12 | ENSMUSG00000028567 | 66073 | 0.182744342909832 | 2.36731323532593 | 1.51393942536283 | 0.1532586446383 | 0.231807146055383 |
| Pxdc1 | ENSMUSG00000021411 | 66895 | -0.418602001609163 | 1.73060511565167 | -1.51364895965656 | 0.1533316404108 | 0.231885467810038 |
| Ryk | ENSMUSG00000032547 | 20187 | -0.0946625926054594 | 4.4768443402621 | -1.51250797794709 | 0.1536186611667 | 0.232287395934873 |
| Phc3 | ENSMUSG00000037652 | 241915 | 0.113655656151881 | 4.82278071185418 | 1.51191295422306 | 0.1537685234331 | 0.232475490658554 |
| Fbxw7 | ENSMUSG00000028086 | 50754 | -0.160171239804313 | 5.12644098639241 | -1.51184522719823 | 0.1537855889627 | 0.232475490658554 |
| S100a1 | ENSMUSG00000044080 | 20193 | -0.162768350900556 | 5.94308166597349 | -1.51145147444709 | 0.1538848366646 | 0.232593355860288 |
| Med20 | ENSMUSG00000092558 | 56771 | -0.203826549997803 | 2.4772746411704 | -1.51132374064672 | 0.1539170443739 | 0.232609873088784 |
| Olfr351 | ENSMUSG00000075383 | 258944 | -0.526156621594094 | 0.668219521391016 | -1.51076513871005 | 0.1540579613264 | 0.232790651819082 |
| Lrrc14 | ENSMUSG00000033728 | 223664 | 0.209655213952376 | 1.95305721944803 | 1.5106524516629 | 0.1540864018133 | 0.232801445503997 |
| Ankrd26 | ENSMUSG00000007827 | 232339 | -0.165770853131219 | 2.51885241667096 | -1.51004379323457 | 0.1542400947647 | 0.233001447743778 |
| Cog5 | ENSMUSG00000035933 | 238123 | -0.118388297161777 | 3.84679577141633 | -1.50937102367086 | 0.1544101274963 | 0.233226074846306 |
| Cmpk1 | ENSMUSG00000028719 | 66588 | 0.159306299997577 | 3.82316630400082 | 1.50827326214209 | 0.1546879114784 | 0.233584909309476 |
| Olfr1167 | ENSMUSG00000100899 | 258291 | -0.731193644459699 | 0.210548345112021 | -1.50826329232706 | 0.1546904362355 | 0.233584909309476 |
| Fip1l1 | ENSMUSG00000029227 | 66899 | 0.0932204739767331 | 4.78546093785884 | 1.50813177929539 | 0.1547237438761 | 0.233602934296995 |
| Acvr2a | ENSMUSG00000052155 | 11480 | 0.104543299931606 | 4.11810027159784 | 1.50788078103271 | 0.1547873297845 | 0.233661530230779 |
| Cmas | ENSMUSG00000030282 | 12764 | 0.106161150020322 | 3.49602909605969 | 1.50780983701672 | 0.1548053061871 | 0.233661530230779 |
| Pigl | ENSMUSG00000014245 | 327942 | -0.37962096108767 | 1.57478989785379 | -1.50746960257332 | 0.1548915422623 | 0.233759415776608 |
| Coq10b | ENSMUSG00000025981 | 67876 | -0.196524415634841 | 2.60544541247373 | -1.50716798948585 | 0.1549680233295 | 0.233842554253927 |
| Atg16l1 | ENSMUSG00000026289 | 77040 | -0.159780431016724 | 4.04508773654883 | -1.50651598146699 | 0.1551334644539 | 0.234041965366947 |
| Pttg1ip | ENSMUSG00000009291 | 108705 | 0.170216502385069 | 4.64409269314543 | 1.50647843827391 | 0.1551429952473 | 0.234041965366947 |
| Paqr4 | ENSMUSG00000023909 | 76498 | 0.245213696383211 | 1.79465228531603 | 1.50577010930351 | 0.1553229059829 | 0.234281038402081 |
| Mat2a | ENSMUSG00000053907 | 232087 | -0.1299346101837 | 5.45603888321729 | -1.50451574328793 | 0.1556419394138 | 0.234729861994033 |
| Tnfaip2 | ENSMUSG00000021281 | 21928 | -0.173917100863732 | 4.56387708083419 | -1.5041980088581 | 0.1557228393910 | 0.234819472669818 |
| Orc5 | ENSMUSG00000029012 | 26429 | -0.261934122655794 | 1.55765657511551 | -1.50389689129425 | 0.1557995412795 | 0.234902729065762 |
| Psmc4 | ENSMUSG00000030603 | 23996 | 0.128855344582128 | 5.4697837797257 | 1.50319241071951 | 0.1559791142364 | 0.235141042300156 |
| Tbx15 | ENSMUSG00000027868 | 21384 | -0.149122793571132 | 6.629439749475 | -1.50292484340031 | 0.1560473633100 | 0.235211490394651 |
| Sart1 | ENSMUSG00000039148 | 20227 | 0.135621881449498 | 4.4588351519129 | 1.50268408965576 | 0.1561087945534 | 0.235271643907884 |
| Il11ra1 | ENSMUSG00000073889 | 16157 | -0.117847334956242 | 5.28785496237037 | -1.50254247863598 | 0.1561449378741 | 0.235293674648859 |
| Utp6 | ENSMUSG00000035575 | 216987 | -0.107539363438642 | 4.79148603489626 | -1.50236725754331 | 0.1561896692708 | 0.235328638842171 |
| Aasdh | ENSMUSG00000055923 | 231326 | -0.22283759541365 | 1.82097747778818 | -1.50205781237066 | 0.1562686925795 | 0.235415253388542 |
| Alg2 | ENSMUSG00000039740 | 56737 | 0.228480349197175 | 2.05872399593515 | 1.5016006326958 | 0.1563855048330 | 0.235558764410907 |
| Rps24 | ENSMUSG00000025290 | 20088 | -0.115223294651064 | 7.41405612575443 | -1.50087042192038 | 0.1565722312945 | 0.235807531038974 |
| Parvb | ENSMUSG00000022438 | 170736 | 0.152234878075097 | 5.70480493793047 | 1.50036529742378 | 0.1567015098417 | 0.235969720909278 |
| Siah1a | ENSMUSG00000036840 | 20437 | -0.172715303783499 | 2.37363140206618 | -1.49990085109891 | 0.1568204570372 | 0.23611631072646 |
| Top1 | ENSMUSG00000070544 | 21969 | -0.0929204873535454 | 5.15298794587258 | -1.4992151696324 | 0.1569962031692 | 0.236348367558581 |
| Sub1 | ENSMUSG00000022205 | 20024 | -0.13685237419966 | 4.47273319667766 | -1.49816272388192 | 0.1572662778177 | 0.236689200719759 |
| Ifih1 | ENSMUSG00000026896 | 71586 | 0.199480054762076 | 3.28013745643294 | 1.49810189839617 | 0.1572818986105 | 0.236689200719759 |
| Taok1 | ENSMUSG00000017291 | 216965 | -0.106406534009737 | 7.30736727889803 | -1.49807984218198 | 0.1572875632630 | 0.236689200719759 |
| Olfr853 | ENSMUSG00000051160 | 258908 | -0.805052613880541 | -0.120750971706477 | -1.49679640047445 | 0.1576174837978 | 0.237139909911665 |
| Olfr715b | ENSMUSG00000062987 | 384732 | -0.730583527023212 | 0.818783820821971 | -1.49674595777091 | 0.1576304624845 | 0.237139909911665 |
| Tango6 | ENSMUSG00000041949 | 272538 | -0.145449746496937 | 2.8292670232393 | -1.49649249447261 | 0.1576956911387 | 0.23720539422569 |
| Igtp | ENSMUSG00000078853 | 16145 | -0.754633977765477 | 2.5979102316044 | -1.49628348406824 | 0.1577494970041 | 0.237253680758441 |
| Aspscr1 | ENSMUSG00000025142 | 68938 | 0.10701046716514 | 3.84933546880188 | 1.49566407417949 | 0.1579090436558 | 0.237460965222426 |
| Sf3b6 | ENSMUSG00000037361 | 66055 | 0.197190644138587 | 3.28131261557128 | 1.49486363490709 | 0.1581154212826 | 0.237738606608025 |
| Ppm1f | ENSMUSG00000026181 | 68606 | -0.156403934697249 | 3.89197440858436 | -1.49473905246458 | 0.1581475628934 | 0.237754230503127 |
| Creld2 | ENSMUSG00000023272 | 76737 | 0.285018923839499 | 2.48125803191584 | 1.49392124920065 | 0.1583586887214 | 0.238015994466264 |
| Olfr600 | ENSMUSG00000045540 | 259048 | -0.677557270431527 | 0.26647747777263 | -1.49389592651396 | 0.1583652298740 | 0.238015994466264 |
| G6pc3 | ENSMUSG00000034793 | 68401 | -0.145510146072683 | 3.27702654031479 | -1.49306500438059 | 0.1585799934861 | 0.238291106290765 |
| Eif3m | ENSMUSG00000027170 | 98221 | -0.111631209378256 | 5.0187287641408 | -1.49301905795597 | 0.1585918761563 | 0.238291106290765 |
| Trp53i13 | ENSMUSG00000044328 | 216964 | 0.324968357841731 | 1.08857040081961 | 1.49264585415698 | 0.1586884220004 | 0.238403400341746 |
| Mb21d2 | ENSMUSG00000051065 | 239796 | 0.165489158123781 | 2.27392892286981 | 1.49211236914258 | 0.1588265178241 | 0.23857807700094 |
| Nup88 | ENSMUSG00000040667 | 19069 | -0.102846064639622 | 4.82137542883361 | -1.49201484906519 | 0.1588517724402 | 0.238583226785442 |
| Dhx57 | ENSMUSG00000035051 | 106794 | -0.194874076284136 | 3.0946223724614 | -1.49158800781671 | 0.1589623506802 | 0.238714207951355 |
| Eloc | ENSMUSG00000079658 | 67923 | -0.122576281643813 | 3.88968429907974 | -1.49150964733237 | 0.1589826579348 | 0.238714207951355 |
| Zfp120 | ENSMUSG00000068134 | 104348 | -0.147569344876415 | 2.64079170265652 | -1.49142296604892 | 0.1590051240872 | 0.23871515058342 |
| Fam81a | ENSMUSG00000032224 | 76886 | -0.407182948190473 | 0.990160249534331 | -1.49110483049752 | 0.1590876017610 | 0.238806176167232 |
| Ralbp1 | ENSMUSG00000024096 | 19765 | 0.0818011909088519 | 5.30734610255279 | 1.49057780824045 | 0.1592243133669 | 0.238946301236998 |
| Sidt2 | ENSMUSG00000034908 | 214597 | -0.101089383781264 | 4.83414376395511 | -1.49057643729173 | 0.1592246691254 | 0.238946301236998 |
| Ppp1r12a | ENSMUSG00000019907 | 17931 | 0.120151351142353 | 5.57728277356883 | 1.4895602732339 | 0.1594885461033 | 0.239309443713919 |
| Zfp277 | ENSMUSG00000055917 | 246196 | -0.186980466335266 | 3.24102906572595 | -1.48763190003921 | 0.1599903185907 | 0.240029395074908 |
| Prpf4b | ENSMUSG00000021413 | 19134 | -0.140553376911344 | 5.2404925263649 | -1.48749042406786 | 0.1600271836719 | 0.240051755827966 |
| Ubb | ENSMUSG00000019505 | 22187 | -0.117420379645754 | 8.73728777727563 | -1.48696577237152 | 0.1601639571982 | 0.24022395940371 |
| Olfr63 | ENSMUSG00000054666 | 258939 | -0.410664673042924 | 1.58684623467956 | -1.48682711317221 | 0.1602001212615 | 0.240245236042008 |
| Fabp5 | ENSMUSG00000027533 | 16592 | -0.288637529283415 | 3.35510920651496 | -1.48671808081075 | 0.1602285631036 | 0.240254927748438 |
| Frmd4a | ENSMUSG00000026657 | 209630 | 0.217214135279626 | 3.22039425647097 | 1.48652083981657 | 0.1602800255820 | 0.240299130385024 |
| Bend3 | ENSMUSG00000038214 | 331623 | 0.240079083845771 | 2.65851465688306 | 1.48581140223789 | 0.1604652412374 | 0.240543822266436 |
| Tubgcp6 | ENSMUSG00000051786 | 328580 | 0.150847883793914 | 3.39167316274271 | 1.48532786738033 | 0.1605915828521 | 0.240700204601169 |
| Pnpla6 | ENSMUSG00000004565 | 50767 | 0.178549883776241 | 2.39089197556677 | 1.48518975343614 | 0.1606276856763 | 0.240721309587119 |
| Lzts3 | ENSMUSG00000037703 | 241638 | -0.256754005827751 | 1.87770102383764 | -1.48484345421258 | 0.1607182379427 | 0.240823997114776 |
| A130010J15Rik | ENSMUSG00000079144 | 319266 | 0.195691155073752 | 2.03697859147227 | 1.48474626829248 | 0.1607436583857 | 0.240829074810185 |
| Pak2 | ENSMUSG00000022781 | 224105 | 0.0897033795965241 | 4.72229011833851 | 1.48334193636657 | 0.1611113606069 | 0.24134689362677 |
| Csnk1g2 | ENSMUSG00000003345 | 103236 | -0.0839648337458296 | 6.08334198613852 | -1.48321603534003 | 0.1611443603656 | 0.24136325063809 |
| Kremen1 | ENSMUSG00000020393 | 84035 | 0.114489355728812 | 7.0808171659665 | 1.4829342017188 | 0.1612182520415 | 0.241440843001199 |
| Col2a1 | ENSMUSG00000022483 | 12824 | -0.513108486932467 | 0.948529015859274 | -1.48278730350972 | 0.1612567773980 | 0.241465456676427 |
| Tmtc3 | ENSMUSG00000036676 | 237500 | 0.144257474384117 | 3.32648483088674 | 1.48147697469877 | 0.1616007656336 | 0.241947400238574 |
| Fam107b | ENSMUSG00000026655 | 66540 | 0.244023258191609 | 2.06604368742454 | 1.48116002593935 | 0.1616840638644 | 0.242038962216196 |
| Unc5b | ENSMUSG00000020099 | 107449 | -0.362240895813808 | 2.8652360043829 | -1.48082363658789 | 0.1617725108449 | 0.242138205675265 |
| Glt8d1 | ENSMUSG00000021916 | 76485 | 0.201508409743114 | 1.94069552390457 | 1.48008439193417 | 0.1619670237337 | 0.242396157781221 |
| Gja4 | ENSMUSG00000050234 | 14612 | -0.372346803557165 | 1.42073191738098 | -1.4791870369597 | 0.1622034035015 | 0.242716687703631 |
| Unkl | ENSMUSG00000015127 | 74154 | 0.210840311084345 | 2.2358151044705 | 1.47907644799694 | 0.1622325547332 | 0.242727081274146 |
| Anxa3 | ENSMUSG00000029484 | 11745 | -0.239140403664585 | 3.40540130916791 | -1.47880190924671 | 0.1623049421450 | 0.242802151715812 |

| | | | | | | | |
|---|---|---|---|---|---|---|---|
| Scly | ENSMUSG00000026307 | 50880 | -0.167521761982047 | 2.32316377708632 | -1.47843271381078 | 0.1624023304256 | 0.242914596864049 |
| Fads3 | ENSMUSG00000024664 | 60527 | 0.228273259437187 | 2.25775783393485 | 1.47809274499908 | 0.1624920526030 | 0.243015546176558 |
| Pdxk | ENSMUSG00000032788 | 216134 | -0.278868091918482 | 3.10042565269031 | -1.47798427226053 | 0.1625206887333 | 0.243025122919844 |
| Grk2 | ENSMUSG00000024858 | 110355 | -0.0846618310667466 | 5.0441154004908 | -1.4778190489268 | 0.1625643148213 | 0.243043967384485 |
| Yif1a | ENSMUSG00000024875 | 68090 | -0.124731174779913 | 3.56705072981769 | -1.47776813780225 | 0.1625777595385 | 0.243043967384485 |
| Hmgn2 | ENSMUSG00000003038 | 15331 | -0.103173408149397 | 4.8491404089941 | -1.47698760695355 | 0.1627840008148 | 0.243319009012328 |
| Cand2 | ENSMUSG00000030319 | 67088 | 0.121922041198036 | 6.82107051021525 | 1.47593255299805 | 0.1630631293697 | 0.243702907730523 |
| Zfp507 | ENSMUSG00000044452 | 668501 | -0.110098760814052 | 4.06219329876155 | -1.47548219452522 | 0.1631824001171 | 0.243847821692456 |
| Rnf138 | ENSMUSG00000024317 | 56515 | 0.189504046901743 | 2.13967102616716 | 1.4751147486741 | 0.1632797669807 | 0.243959968953858 |
| Srcap | ENSMUSG00000053877 | 100043597 | 0.123139131575136 | 6.57047635283173 | 1.47427902098902 | 0.1635014022760 | 0.244257732442119 |
| Ccn2 | ENSMUSG00000019997 | 14219 | 0.617167480169762 | 5.377634930587 | 1.47384500369366 | 0.1636166034877 | 0.244396432457525 |
| Slc25a36 | ENSMUSG00000032449 | 192287 | -0.143260449744068 | 4.99869330555078 | -1.47327858012992 | 0.1637670518308 | 0.244572007228566 |
| Mfsd1 | ENSMUSG00000027775 | 66868 | 0.179964662694573 | 3.2775721729854 | 1.47323401315981 | 0.1637788942377 | 0.244572007228566 |
| Ola1 | ENSMUSG00000027108 | 67059 | 0.0955112807045957 | 4.52619974767838 | 1.47301451461009 | 0.1638372302397 | 0.244625701919166 |
| Rpl22l1 | ENSMUSG00000039221 | 68028 | -0.19940410744888 | 2.68662132320707 | -1.47244754569907 | 0.1639879939838 | 0.244816193928923 |
| Rxylt1 | ENSMUSG00000034620 | 216395 | -0.117694172523326 | 3.58465666117658 | -1.47236631162276 | 0.1640096046236 | 0.244816193928923 |
| Gtf2h3 | ENSMUSG00000029387 | 209357 | 0.189329271814912 | 2.52034258514301 | 1.47188872011502 | 0.1641367062897 | 0.244972465381397 |
| Golga2 | ENSMUSG00000002546 | 99412 | -0.0841150977519209 | 6.33392636825182 | -1.4707814509298 | 0.1644317025079 | 0.245379240971242 |
| Map2k3os | ENSMUSG00000042549 | 24082 | -0.266720816414274 | 1.72772349232796 | -1.47030461113463 | 0.1645588781290 | 0.245535503252542 |
| Trappc10 | ENSMUSG00000000374 | 216131 | 0.115782952257889 | 5.51812169804531 | 1.46870600876156 | 0.1649858359466 | 0.246138961748644 |
| Clmn | ENSMUSG00000021097 | 94040 | -0.267638244285886 | 1.76480591005422 | -1.46857052425677 | 0.1650220640923 | 0.246154001444158 |
| Abi3 | ENSMUSG00000018381 | 66610 | -0.370502329844872 | 1.88515585538992 | -1.46846947586866 | 0.1650490884692 | 0.246154001444158 |
| Prkn | ENSMUSG00000023826 | 50873 | -0.184934832580494 | 2.98908959815402 | -1.46841569303953 | 0.1650634736607 | 0.246154001444158 |
| Ip6k3 | ENSMUSG00000024210 | 271424 | 0.125913450909811 | 4.97664619865809 | 1.46759205526597 | 0.1652839019026 | 0.24644332527394 |
| Arfip2 | ENSMUSG00000030881 | 76932 | 0.183066751424414 | 2.13105619533573 | 1.4674854870038 | 0.1653124405465 | 0.24644332527394 |
| Clint1 | ENSMUSG00000006169 | 216705 | 0.096345563694018 | 5.58081886218673 | 1.46740188597315 | 0.1653348315308 | 0.24644332527394 |
| Nadk | ENSMUSG00000029063 | 192185 | 0.121551146150677 | 4.07631290743509 | 1.46730855182456 | 0.1653598323549 | 0.24644332527394 |
| Fbxo9 | ENSMUSG00000001366 | 71538 | -0.186584950659418 | 4.80169817482145 | -1.46726980436577 | 0.1653702123213 | 0.24644332527394 |
| Cog7 | ENSMUSG00000034951 | 233824 | -0.111836381670102 | 3.8586084884125 | -1.46615118190396 | 0.1656701131490 | 0.246856598532189 |
| Mapk14 | ENSMUSG00000053436 | 26416 | -0.0989224519827916 | 5.21619535154878 | -1.46603047231248 | 0.1657025024319 | 0.246871208134584 |
| Trub1 | ENSMUSG00000025086 | 72133 | -0.167251886200943 | 2.13367874073995 | -1.46570479287858 | 0.1657899165321 | 0.246967781086453 |
| Sirt5 | ENSMUSG00000054021 | 68346 | -0.191756790412638 | 4.16052428124871 | -1.46547323507115 | 0.1658520913964 | 0.247026735393685 |
| Rapgef4 | ENSMUSG00000049044 | 56508 | 0.165948639700776 | 2.54933686639025 | 1.4651399681031 | 0.1659416101784 | 0.247126088196959 |
| Letm2 | ENSMUSG00000037363 | 270035 | -0.205828267563148 | 1.44513543366311 | -1.46505659314866 | 0.1659640118428 | 0.247126088196959 |
| Afg3l1 | ENSMUSG00000031967 | 114896 | -0.120850785684249 | 4.25127938772858 | -1.46375734064566 | 0.1663134301435 | 0.24761265389589 |
| Gm32687 | ENSMUSG00000112640 | 102635315 | 0.364932335542971 | 1.49696478913773 | 1.46292851379774 | 0.1665366550318 | 0.247911231942347 |
| Ogfod1 | ENSMUSG00000033009 | 270086 | -0.113041146031374 | 3.9529589223792 | -1.46249156261132 | 0.1666544385009 | 0.248052786935415 |
| Flnb | ENSMUSG00000025278 | 286940 | -0.207168612744471 | 5.13786148106461 | -1.46184347293863 | 0.1668292644327 | 0.248279195304843 |
| Vps26b | ENSMUSG00000031988 | 69091 | 0.130113112859088 | 4.34765399224215 | 1.46055621515513 | 0.1671769652101 | 0.248762783380415 |
| Aplf | ENSMUSG00000030051 | 72103 | -0.233989120809835 | 1.80396188928424 | -1.46009726563659 | 0.1673010785620 | 0.248913582382135 |
| Matr3 | ENSMUSG00000037236 | 17184 | -0.111488524490887 | 6.01369340327938 | -1.45942731366236 | 0.1674823916284 | 0.24914943153104 |
| Rpl13a | ENSMUSG00000074129 | 22121 | 0.150097897219666 | 7.31212093637984 | 1.45908378369691 | 0.1675754269544 | 0.249253911013647 |
| Vti1a | ENSMUSG00000024983 | 53611 | -0.134684986995525 | 3.71467145650018 | -1.45882233941705 | 0.1676462606908 | 0.249325343620611 |
| Gsk3b | ENSMUSG00000022812 | 56637 | 0.104920390862025 | 6.83315158045463 | 1.45845964151823 | 0.1677445688129 | 0.249437611439822 |
| Fbxo25 | ENSMUSG00000038365 | 66822 | 0.185176255803872 | 2.06212996042489 | 1.45729680289156 | 0.1680600778979 | 0.249872784480722 |
| U2af2 | ENSMUSG00000030435 | 22185 | 0.0952487511256619 | 4.55588762660035 | 1.45699334362716 | 0.1681424961252 | 0.249961325329146 |
| Mtor | ENSMUSG00000028991 | 56717 | -0.0916962294036585 | 5.85907918675029 | -1.45631937221325 | 0.1683256648067 | 0.250199597770286 |
| Ip6k1 | ENSMUSG00000032594 | 27399 | 0.105726733949298 | 4.30262489231944 | 1.45520723284743 | 0.1686282815813 | 0.250615329159144 |
| Thtpa | ENSMUSG00000045691 | 105663 | -0.124341355829368 | 2.93450627102911 | -1.45506927684402 | 0.1686658515635 | 0.250637088559092 |
| Pde4dip | ENSMUSG00000038170 | 83679 | -0.173071582269513 | 11.7191795073247 | -1.45418355179032 | 0.1689072305546 | 0.250942159409682 |
| B4gat1 | ENSMUSG00000047379 | 108902 | 0.14662141365441 | 3.07327587329249 | 1.45414749694115 | 0.1689170623855 | 0.250942159409682 |
| Lrfn3 | ENSMUSG00000036957 | 233067 | -0.294040638078587 | 1.06500793580019 | -1.45323982521925 | 0.1691647339250 | 0.251253296436032 |
| Tob2 | ENSMUSG00000048546 | 57259 | -0.228769712928253 | 5.31030143286501 | -1.45321149440317 | 0.1691724692863 | 0.251253296436032 |
| Ambra1 | ENSMUSG00000040506 | 228361 | 0.111246063586823 | 4.23222087810794 | 1.45305698770196 | 0.1692146605370 | 0.251281816917547 |
| Rap2c | ENSMUSG00000050029 | 72065 | 0.108436933753207 | 4.26066670828637 | 1.45136706266859 | 0.1696767093918 | 0.251933720533814 |
| Khdrbs1 | ENSMUSG00000028790 | 20218 | 0.0829768213838911 | 4.76980892039841 | 1.45063989384566 | 0.1698758437085 | 0.25219514431325 |
| Rps17 | ENSMUSG00000061787 | 20068 | -0.125269644264125 | 7.07275067064172 | -1.44961289582837 | 0.1701574261720 | 0.252549487687545 |
| Ccdc90b | ENSMUSG00000030613 | 66365 | -0.130881996893844 | 3.73115400892214 | -1.44960084201259 | 0.1701607334021 | 0.252549487687545 |
| Cops2 | ENSMUSG00000027206 | 12848 | -0.0947932915617846 | 5.57769448784218 | -1.44938879613635 | 0.1702189216416 | 0.25260155205883 |
| Nvl | ENSMUSG00000026516 | 67459 | -0.0931558586771795 | 4.34636272333646 | -1.44852570306092 | 0.1704559374787 | 0.252918942932097 |
| Jam3 | ENSMUSG00000031990 | 83964 | 0.120529759465268 | 3.64705158927554 | 1.44826407501207 | 0.1705278381193 | 0.252991286439005 |
| Tceanc2 | ENSMUSG00000028619 | 66526 | 0.156722309901131 | 2.98648819719257 | 1.44815247421455 | 0.1705585159624 | 0.253002461390368 |
| Man2c1 | ENSMUSG00000032295 | 73744 | 0.135428087703925 | 3.39327136896256 | 1.44798583506589 | 0.1706043318248 | 0.253036085641383 |
| Ift140 | ENSMUSG00000024169 | 106633 | 0.19164545839767 | 1.88691968435534 | 1.44729979456412 | 0.1707930604226 | 0.253281636613714 |
| Abl1 | ENSMUSG00000026842 | 11350 | -0.127468332088452 | 4.83615222081713 | -1.44714804156256 | 0.1708348309499 | 0.253309215560739 |
| Hey1 | ENSMUSG00000040289 | 15213 | -0.300953301352666 | 1.76885510489425 | -1.44702520572373 | 0.1708686481743 | 0.253324995686192 |
| Gfpt2 | ENSMUSG00000020363 | 14584 | 0.30846241165178 | 3.27279804263001 | 1.44684123105188 | 0.1709193076280 | 0.253365737955283 |
| Ppp1r15a | ENSMUSG00000040435 | 17872 | -0.229222059415275 | 4.49671388498291 | -1.44669838439778 | 0.1709586506734 | 0.253389696340487 |
| Pecr | ENSMUSG00000026189 | 111175 | -0.207926085156408 | 2.05361516070289 | -1.44639188431999 | 0.1710430929035 | 0.253480483802771 |
| Impa1 | ENSMUSG00000027531 | 55980 | 0.134538915367188 | 2.64336650209205 | 1.44614533637703 | 0.1711110432930 | 0.253515687667288 |
| Trmt112 | ENSMUSG00000038812 | 67674 | 0.170011996256307 | 3.20146124778023 | 1.44613739480352 | 0.1711132324224 | 0.253515687667288 |
| Fam217b | ENSMUSG00000070476 | 71532 | 0.275361882670723 | 1.88692887251291 | 1.44580863164812 | 0.1712038779292 | 0.253615610468209 |
| Cacfd1 | ENSMUSG00000015488 | 381356 | 0.128225514725595 | 3.19172689708617 | 1.44552036362099 | 0.1712833912159 | 0.25369901753139 |
| Atpsckmt | ENSMUSG00000039065 | 68073 | -0.164233534259884 | 2.19636478610064 | -1.44507341525605 | 0.1714067343928 | 0.253847312511662 |
| Arfip1 | ENSMUSG00000074513 | 99889 | 0.179660025085314 | 2.70923179490181 | 1.44489240527544 | 0.1714567083236 | 0.253886924774516 |
| Gtf3c6 | ENSMUSG00000019837 | 67371 | -0.122476700854913 | 4.06248240526177 | -1.44451994318893 | 0.1715595773243 | 0.253993912682042 |
| Dek | ENSMUSG00000021377 | 110052 | -0.0946040471928011 | 5.29511409541019 | -1.44446255273094 | 0.1715754323707 | 0.253993912682042 |
| Pym1 | ENSMUSG00000064030 | 78428 | 0.214727482008192 | 2.05940633484065 | 1.44407939788585 | 0.1716813164432 | 0.254116245096925 |
| Usp14 | ENSMUSG00000047879 | 59025 | 0.111341245083549 | 4.39311132514221 | 1.44373924798241 | 0.1717753618496 | 0.254221023609318 |
| 1110002E22Rik | ENSMUSG00000090066 | 102634333 | -0.127692066651337 | 8.06612926612401 | -1.44359908504501 | 0.1718141269101 | 0.254243972012228 |
| Nfatc2ip | ENSMUSG00000030722 | 18020 | -0.182363117326918 | 1.48932801928153 | -1.44319606882777 | 0.1719256303205 | 0.254374535061472 |
| Gm14296 | ENSMUSG00000074527 | 102639598 | -0.354054321249582 | 0.917541052151726 | -1.44179429337512 | 0.1723139327448 | 0.254914548495602 |
| Epha5 | ENSMUSG00000029245 | 13839 | -0.490465696564394 | 1.10665063480109 | -1.44119101150285 | 0.1724812712804 | 0.255127574609751 |
| Trim12c | ENSMUSG00000057143 | 319236 | -0.206834896468847 | 2.51464026170134 | -1.44071534723931 | 0.1726133066793 | 0.255288331120475 |
| Gpr137 | ENSMUSG00000024958 | 107173 | -0.138841393291475 | 2.40565711431259 | -1.44061365579877 | 0.1726415452254 | 0.255295553417195 |
| Ncoa2 | ENSMUSG00000005886 | 17978 | -0.10415869798639 | 5.23457493655692 | -1.44010350026449 | 0.1727832676710 | 0.255470566605183 |
| Brox | ENSMUSG00000046836 | 71678 | 0.0973218579339479 | 4.51555959080619 | 1.44001629554479 | 0.1728075030532 | 0.255471844181097 |
| Mphosph9 | ENSMUSG00000038126 | 269702 | 0.23266714289901 | 1.85265876738401 | 1.43964083444805 | 0.1729118811715 | 0.255567802564279 |
| Atp9b | ENSMUSG00000024566 | 50771 | 0.108123552771774 | 4.21654399724414 | 1.43961029045166 | 0.1729203747070 | 0.255567802564279 |
| Stimate | ENSMUSG00000006526 | 69179 | -0.201892418196817 | 2.20973574784259 | -1.43953054366742 | 0.1729425519685 | 0.255567802564279 |
| Wdr20 | ENSMUSG00000037957 | 69641 | -0.133821255244065 | 3.24118666594396 | -1.43893614412674 | 0.1731079267671 | 0.255777608474094 |
| Wrnip1 | ENSMUSG00000021400 | 78903 | 0.139396033276742 | 2.54212999478264 | 1.43769772545926 | 0.1734529047674 | 0.256252696582406 |
| BC002059 | ENSMUSG00000060149 | 213811 | -0.200728209809041 | 1.53879863479096 | -1.43747647990781 | 0.1735145959013 | 0.256309195648327 |
| Erbb3 | ENSMUSG00000018166 | 13867 | 0.170795857620602 | 3.18951927160859 | 1.43725594562646 | 0.1735761068901 | 0.256357033134449 |
| Mcts1 | ENSMUSG00000000355 | 68995 | -0.191233018709418 | 2.57464777908106 | -1.43719221743741 | 0.1735938852128 | 0.256357033134449 |
| Add1 | ENSMUSG00000029106 | 11518 | 0.0819205243299068 | 6.14645939123027 | 1.43682973317454 | 0.1736950366861 | 0.256471760909879 |
| Ubfd1 | ENSMUSG00000030870 | 28018 | 0.100866805095216 | 3.87648467704594 | 1.43578085567383 | 0.1739880031151 | 0.256869646414378 |
| Zcchc9 | ENSMUSG00000021621 | 69085 | 0.145058401725396 | 2.492355778195 | 1.43556478520676 | 0.1740484057425 | 0.256924121967904 |
| Srprb | ENSMUSG00000032553 | 20818 | 0.151840019365695 | 3.39681502479157 | 1.43487820868445 | 0.1742404544731 | 0.257161317771828 |
| Inafm1 | ENSMUSG00000091811 | 66300 | -0.211226939815417 | 1.65689737414746 | -1.43482215585645 | 0.1742561413169 | 0.257161317771828 |
| Zfp628 | ENSMUSG00000074406 | 232816 | -0.140927192782975 | 3.2890756918615 | -1.43457708533511 | 0.1743247401163 | 0.257227825892569 |
| Uhrf2 | ENSMUSG00000024817 | 109113 | 0.17653002316337 | 3.56455181559241 | 1.43440913480614 | 0.1743717648858 | 0.257262486431012 |
| Ptov1 | ENSMUSG00000038502 | 84113 | -0.111943592130393 | 5.09779827300218 | -1.43399501879839 | 0.1744877590704 | 0.257398879136241 |
| Ston2 | ENSMUSG00000020961 | 108800 | -0.127120724231851 | 5.62559075630995 | -1.43384453588308 | 0.1745299253413 | 0.25742634110193 |
| Chfr | ENSMUSG00000014668 | 231600 | 0.139438521964484 | 2.95830601723507 | 1.43374957168892 | 0.1745565392760 | 0.257430859528736 |
| Elp4 | ENSMUSG00000027167 | 77766 | -0.167179879030151 | 2.44820429677257 | -1.43343232297445 | 0.1746454734433 | 0.257527272387643 |
| Gle1 | ENSMUSG00000019715 | 74412 | -0.121249856994893 | 4.20573445557264 | -1.43332094196503 | 0.1746767057628 | 0.257538585202827 |
| Ext2 | ENSMUSG00000027198 | 14043 | 0.114877055369022 | 4.41205276478232 | 1.43160379134281 | 0.1751587997645 | 0.258214543523386 |
| Cd38 | ENSMUSG00000029084 | 12494 | -0.191836816945764 | 2.28876037496025 | -1.43062589729575 | 0.1754338399065 | 0.258585127294821 |
| Mnat1 | ENSMUSG00000021103 | 17420 | 0.127639021217133 | 3.32179233062745 | 1.43049587115083 | 0.1754704377800 | 0.258604200535735 |
| Wrap53 | ENSMUSG00000041346 | 216853 | -0.230596610649957 | 1.25663508251273 | -1.43008724364415 | 0.1755854936420 | 0.258738882583047 |
| Mon2 | ENSMUSG00000034602 | 67074 | 0.0938691958564641 | 5.10129728182847 | 1.42974544802207 | 0.1756817800357 | 0.258845873779589 |
| Ccdc138 | ENSMUSG00000038010 | 76138 | -0.270652074285192 | 1.87975642332914 | -1.42961194411463 | 0.1757194010062 | 0.258866411374525 |
| Cd320 | ENSMUSG00000002308 | 54219 | -0.220656570708828 | 1.70366896739037 | -1.42951406239569 | 0.1757469880130 | 0.258872163585852 |
| Tmem72 | ENSMUSG00000048108 | 319776 | -0.431058769339856 | 5.05267491204441 | -1.42900245086724 | 0.1758912391060 | 0.259049735503173 |
| Wnt5b | ENSMUSG00000030170 | 22419 | -0.27264870909502 | 1.78412410339262 | -1.42798616876707 | 0.1761780767235 | 0.259437229781175 |

| | | | | | | | |
|---|---|---|---|---|---|---|---|
| Vwa1 | ENSMUSG00000042116 | 246228 | -0.209332012286868 | 3.07610183086094 | -1.42629697982431 | 0.176655696380 | 0.260105524937959 |
| Esco1 | ENSMUSG00000024293 | 77805 | -0.104687593303316 | 3.62002831141142 | -1.42620789810537 | 0.17668091413 | 0.260107619184709 |
| Atp2c1 | ENSMUSG00000032570 | 235574 | 0.151090419905901 | 4.49941890259236 | 1.42498642133175 | 0.17702699832 | 0.260582025137195 |
| Uevld | ENSMUSG00000043262 | 54122 | -0.16357078452874 | 2.72195421312759 | -1.42433834211897 | 0.17721084700 | 0.260817528900492 |
| Tpbgl | ENSMUSG00000096606 | 100503386 | -0.297161207267323 | 2.77872389101029 | -1.42392449828182 | 0.17732833546 | 0.260955308955486 |
| Vmn2r89 | ENSMUSG00000070448 | 22301 | -1.07794092377196 | -0.238108708090702 | -1.42367357015875 | 0.17739599193 | 0.261025039545596 |
| Sft2d1 | ENSMUSG00000073468 | 106489 | 0.351427025266535 | 0.961039106350708 | 1.42313368015657 | 0.17755301438 | 0.261198045587646 |
| Slc39a10 | ENSMUSG00000025986 | 227059 | 0.177484090195929 | 3.34659267161379 | 1.42309162621636 | 0.17756496905 | 0.261198045587646 |
| Wdr61 | ENSMUSG00000061559 | 66317 | -0.18464159057952 | 3.05068846787038 | -1.42201502804045 | 0.17787124105 | 0.261613365979394 |
| Rabgap1 | ENSMUSG00000035437 | 227800 | -0.0821665182028841 | 5.29748266371091 | -1.42163731502078 | 0.17797879712 | 0.261736342177473 |
| Nup214 | ENSMUSG00000001855 | 227720 | 0.128143002954663 | 4.04638111048189 | 1.42141898812472 | 0.17804099163 | 0.261792585363602 |
| Plekhj1 | ENSMUSG00000035278 | 78670 | 0.173412875143414 | 2.36652157767032 | 1.4210455915173 | 0.17814740244 | 0.261913820593702 |
| Kti12 | ENSMUSG00000073775 | 100087 | -0.145406148000356 | 3.11323156998122 | -1.42043729359982 | 0.17832086853 | 0.262133595209986 |
| Rfx3 | ENSMUSG00000040929 | 19726 | 0.229800038888638 | 1.55240970027707 | 1.42010020244286 | 0.17841705589 | 0.262239725424224 |
| Ints1 | ENSMUSG00000029547 | 68510 | 0.103553050136643 | 5.26860614409046 | 1.4198647678288 | 0.17848426155 | 0.262303235152089 |
| Pcnp | ENSMUSG00000071533 | 76302 | -0.0946278499229845 | 4.55987536023392 | -1.41939622650713 | 0.17861807082 | 0.262446841154221 |
| Zfp949 | ENSMUSG00000032425 | 71640 | -0.283845288786013 | 2.29380522552866 | -1.41935448007535 | 0.17862997089 | 0.262446841154221 |
| 0610010K14Rik | ENSMUSG00000020831 | 104457 | -0.175098575496148 | 2.16093329864716 | -1.41906908849688 | 0.17871154645 | 0.262531368668776 |
| H2-K1 | ENSMUSG00000061232 | 14972 | -0.213929719426618 | 5.49319075518883 | -1.41843763419218 | 0.17889209137 | 0.262761280682217 |
| Lincmd1 | ENSMUSG00000099906 | 105661730 | 0.43954135535572 | 3.85464162151016 | 1.41822270193392 | 0.17895357911 | 0.262816280175916 |
| Magt1 | ENSMUSG00000031232 | 67075 | 0.109682133630852 | 4.60668683314415 | 1.4176498933768 | 0.17911753353 | 0.263021730127402 |
| Hip1r | ENSMUSG00000000915 | 29816 | -0.208583038390395 | 3.02755705795664 | -1.41716687569163 | 0.17925588392 | 0.263189532193857 |
| Nat10 | ENSMUSG00000027185 | 98956 | 0.103865719207497 | 3.68278453833891 | 1.41679251280968 | 0.17936327650 | 0.263311841986002 |
| Cln8 | ENSMUSG00000026317 | 26889 | 0.146490635574848 | 3.31836561594999 | 1.41585932939521 | 0.17963084775 | 0.263669235499999 |
| Slc9a8 | ENSMUSG00000039463 | 77031 | -0.113935165732701 | 4.07551374472747 | -1.41564195696513 | 0.17969324638 | 0.263725413023299 |
| Tti2 | ENSMUSG00000031577 | 234138 | -0.175864701236394 | 2.10336737564525 | -1.4154381576018 | 0.17975176505 | 0.263775881834921 |
| Map2k5 | ENSMUSG00000058444 | 23938 | -0.134842879780886 | 3.12045375619943 | -1.41520417978563 | 0.17981896858 | 0.263792185503929 |
| Toe1 | ENSMUSG00000028688 | 68276 | 0.18943736216362 | 1.78500495956574 | 1.41516346191036 | 0.17983066577 | 0.263792185503929 |
| Gtf3a | ENSMUSG00000016503 | 66596 | -0.135619793135235 | 3.39634741970852 | -1.41514742565623 | 0.17983527274 | 0.263792185503929 |
| Srsf7 | ENSMUSG00000024097 | 225027 | -0.133954441991735 | 3.83057433782803 | -1.41493121888556 | 0.17989739520 | 0.263847903798757 |
| Mrpl33 | ENSMUSG00000106918 | 66845 | -0.110644540027762 | 4.26800250477887 | -1.41441896036048 | 0.18004465284 | 0.264028454552837 |
| Serf2 | ENSMUSG00000074884 | 378702 | -0.108067515028784 | 7.09072826815967 | -1.41423453766523 | 0.18009769279 | 0.264070808838907 |
| Slc25a23 | ENSMUSG00000046329 | 66972 | -0.109595444938528 | 4.13228059267996 | -1.41409601156169 | 0.18013754139 | 0.264093812358359 |
| Tet2 | ENSMUSG00000040943 | 214133 | 0.116421173489957 | 5.41428079385234 | 1.4137242213691 | 0.18024452713 | 0.264202374185367 |
| Lage3 | ENSMUSG00000015289 | 66192 | -0.229270470977886 | 1.23143356983864 | -1.41367070557513 | 0.18025993108 | 0.264202374185367 |
| Pigu | ENSMUSG00000038383 | 228812 | 0.132770816004458 | 3.17388568807959 | 1.41313322259787 | 0.18041470025 | 0.264393764375154 |
| Hspb7 | ENSMUSG00000006221 | 29818 | 0.144648045628044 | 10.7378029822212 | 1.41290395523932 | 0.18048075165 | 0.264455106745951 |
| Styx | ENSMUSG00000053205 | 56291 | 0.155487887231133 | 3.82563109708738 | 1.41264811373937 | 0.18055448263 | 0.264527683909783 |
| Sh2b1 | ENSMUSG00000030733 | 20399 | 0.0793639946209836 | 4.85971588851971 | 1.41187730877196 | 0.18077671680 | 0.264817862670135 |
| Prkag2 | ENSMUSG00000028944 | 108099 | -0.18316852387865 | 2.53469491343735 | -1.4111660939239 | 0.18098207663 | 0.265083087191671 |
| Pou2f1 | ENSMUSG00000026565 | 18986 | -0.127185258529349 | 3.98621762634743 | -1.41093427743692 | 0.18104903630 | 0.26514563449575 |
| Pwwp2b | ENSMUSG00000060260 | 101631 | -0.207203394073254 | 2.32387264143487 | -1.40916678094696 | 0.18156024782 | 0.265858683043656 |
| Kazn | ENSMUSG00000040606 | 71529 | -0.360980081290443 | 1.42196517447256 | -1.40905217711112 | 0.18159343576 | 0.265871664385518 |
| Pold1 | ENSMUSG00000038644 | 18971 | -0.167471392300766 | 2.00555521477211 | -1.40861871235152 | 0.18171900749 | 0.266019883615809 |
| Pds5b | ENSMUSG00000034021 | 100710 | -0.120702714786145 | 4.47496127496364 | -1.40800816657269 | 0.18189600015 | 0.266210666041539 |
| Slfn5 | ENSMUSG00000054404 | 327978 | -0.178136272520451 | 5.39404875623358 | -1.40800113622001 | 0.18189803903 | 0.266210666041539 |
| Ppp4c | ENSMUSG00000030697 | 56420 | 0.139230119333916 | 3.46426115742605 | 1.40766901608589 | 0.18199437890 | 0.266316004793301 |
| Haus3 | ENSMUSG00000079555 | 231123 | 0.208879072546586 | 1.61986214113005 | 1.40675081386446 | 0.18226094627 | 0.266670379302209 |
| Olfr344 | ENSMUSG00000096822 | 258621 | -0.646988609066798 | 1.96392422934496 | -1.40646413452427 | 0.18234423950 | 0.266756542031257 |
| Rpl36al | ENSMUSG00000049751 | 66483 | 0.144744637912919 | 5.23815011087071 | 1.40635446390376 | 0.18237611206 | 0.266767467017253 |
| Wtap | ENSMUSG00000060475 | 60532 | -0.100079233591835 | 4.41142275449852 | -1.40565140683334 | 0.18258054443 | 0.267030764138842 |
| Cracr2b | ENSMUSG00000048200 | 213573 | -0.283593337619889 | 1.55473535119371 | -1.40554065644052 | 0.18261276520 | 0.267042158834054 |
| Mex3d | ENSMUSG00000048696 | 237400 | 0.167305092725467 | 2.43299037964723 | 1.40511716947779 | 0.18273601458 | 0.267186647326443 |
| Slain2 | ENSMUSG00000036087 | 75991 | 0.100610887060284 | 7.23970997410988 | 1.40499361777453 | 0.18277198533 | 0.267203500292768 |
| Sirt1 | ENSMUSG00000020063 | 93759 | -0.125560481208475 | 3.3272626556586 | -1.40468513460362 | 0.18286182242 | 0.26729908812231 |
| N4bp3 | ENSMUSG00000001053 | 212706 | -0.272442311414529 | 2.13024186635121 | -1.40395493751914 | 0.18307461726 | 0.267542532804689 |
| Psmd9 | ENSMUSG00000029440 | 67151 | -0.149282027026152 | 2.67073606509529 | -1.40394568010877 | 0.18307731638 | 0.267542532804689 |
| Luc7l | ENSMUSG00000024188 | 66978 | 0.168586176127079 | 3.35170372067862 | 1.40289530484275 | 0.18338378026 | 0.267954565179013 |
| Dzip1l | ENSMUSG00000037784 | 72507 | 0.303700501967944 | 1.23340296915588 | 1.40252055146925 | 0.18349322306 | 0.268078644925493 |
| Nop56 | ENSMUSG00000027405 | 67134 | 0.129443842899447 | 4.35052471384591 | 1.40213768327089 | 0.18360509139 | 0.268206234670817 |
| Ddx18 | ENSMUSG00000001674 | 66942 | -0.117525984976042 | 3.42514065522873 | -1.40175176474016 | 0.18371790795 | 0.26831170651151 |
| Tnik | ENSMUSG00000027692 | 665113 | -0.123110093387586 | 6.24383767189944 | -1.4017227719409 | 0.18372638580 | 0.26831170651151 |
| Tpgs2 | ENSMUSG00000024269 | 66648 | -0.232807812459503 | 2.0731058782636 | -1.40150394606356 | 0.18379038359 | 0.268369313793169 |
| Ctcf | ENSMUSG00000005698 | 13018 | -0.113182278987247 | 4.73170226048229 | -1.40055006132398 | 0.18406957154 | 0.268741082749117 |
| Dnttip2 | ENSMUSG00000039756 | 99480 | -0.0974716629151002 | 4.88218183252361 | -1.39923362930723 | 0.18445546368 | 0.269268494158445 |
| St14 | ENSMUSG00000031995 | 19143 | -0.694405600746355 | -0.220782495079137 | -1.39860446325668 | 0.18464010414 | 0.269502066823557 |
| G530011O06Rik | ENSMUSG00000072844 | 654820 | 0.57731730641799 | 0.330618355602872 | 1.3982469214484 | 0.18474510919 | 0.269601819736175 |
| Nr4a3 | ENSMUSG00000028341 | 18124 | -0.521824268769537 | 2.72411269457905 | -1.39820380790386 | 0.18475777437 | 0.269601819736175 |
| Kcnq1ot1 | ENSMUSG00000101609 | 63830 | -0.353462894595153 | 5.63507431811589 | -1.397425381597 | 0.18498657059 | 0.269899653385292 |
| Carm1 | ENSMUSG00000032185 | 59035 | -0.11141536247873 | 5.3363693198814 | -1.39712054008737 | 0.18507623370 | 0.2699944362178 |
| Nudcd2 | ENSMUSG00000020328 | 52653 | 0.113884943028147 | 2.96862901811327 | 1.39696569702698 | 0.18512179144 | 0.269998013402675 |
| Asf1a | ENSMUSG00000019857 | 66403 | -0.171875585138656 | 2.83300784476567 | -1.39694430463868 | 0.18512808621 | 0.269998013402675 |
| Ywhaq | ENSMUSG00000076432 | 22630 | 0.168929188056937 | 4.57001928560296 | 1.39673280066297 | 0.18519033135 | 0.270052763083861 |
| Tln2 | ENSMUSG00000052698 | 70549 | 0.130171501627601 | 7.53165781071659 | 1.39571773683741 | 0.18548930246 | 0.270452656430302 |
| Axin1 | ENSMUSG00000024182 | 12005 | 0.106866726486327 | 4.13062744063032 | 1.3949895627908 | 0.18570401964 | 0.270729613057104 |
| A430033K04Rik | ENSMUSG00000056014 | 243308 | -0.193135767451647 | 1.81344560152221 | -1.39417096482675 | 0.18594564466 | 0.271016031044253 |
| Tbc1d15 | ENSMUSG00000020130 | 66687 | -0.108353357491112 | 4.27872951175645 | -1.39415597501788 | 0.18595007161 | 0.271016031044253 |
| Tsen54 | ENSMUSG00000020781 | 76265 | 0.289435987999376 | 1.49135331381357 | 1.39398078104804 | 0.18600181815 | 0.271055309191383 |
| Pomgnt1 | ENSMUSG00000028700 | 68273 | 0.13341519217203 | 5.4803057917566 | 1.39348984743324 | 0.18614688700 | 0.271230554760951 |
| Lrpap1 | ENSMUSG00000029103 | 16976 | 0.113931002148298 | 4.65128776369311 | 1.39336552205316 | 0.18618363942 | 0.271247949156 |
| Stat5a | ENSMUSG00000004043 | 20850 | 0.165190410445915 | 3.40325905529659 | 1.39302081144625 | 0.18628557223 | 0.271360286522047 |
| Pcif1 | ENSMUSG00000039849 | 228866 | -0.109297262956191 | 4.08259557728926 | -1.39243112584043 | 0.18646005240 | 0.271578258351344 |
| Nedd8 | ENSMUSG00000010376 | 18002 | -0.109263873637294 | 4.72893434444798 | -1.39226125226577 | 0.18651034024 | 0.271615311652461 |
| Pop4 | ENSMUSG00000030423 | 66161 | -0.183830649604653 | 2.82448574200358 | -1.39151046434066 | 0.18673273209 | 0.271889722728063 |
| Amdhd2 | ENSMUSG00000036820 | 245847 | 0.222855735283156 | 1.72894742158262 | 1.39145720333017 | 0.18674851690 | 0.271889722728063 |
| Tank | ENSMUSG00000064289 | 21353 | 0.0961339127822938 | 3.92317061968517 | 1.39084594345725 | 0.18692975276 | 0.272117342848496 |
| Ube2o | ENSMUSG00000020802 | 217342 | 0.090988201510017 | 5.05243397736503 | 1.39063136190711 | 0.18699340957 | 0.272173762985663 |
| Selenok | ENSMUSG00000042682 | 80795 | -0.105933543493475 | 5.44575647628863 | -1.39005696776567 | 0.18716389458 | 0.272385638631484 |
| B3gnt3 | ENSMUSG00000031803 | 72297 | 0.247421255102395 | 1.75305937055625 | 1.38970585020037 | 0.18726817222 | 0.272501116954375 |
| March5 | ENSMUSG00000023307 | 69104 | -0.089908703440309 | 4.93391000484348 | -1.38867307342097 | 0.18757517146 | 0.272911513288129 |
| Dctn2 | ENSMUSG00000025410 | 69654 | 0.0918091151898617 | 5.34304584081688 | 1.38804589639105 | 0.18776180567 | 0.273131395344267 |
| Sh3bp5 | ENSMUSG00000021892 | 24056 | -0.130116442286968 | 4.66619782545206 | -1.38795640939562 | 0.18778844750 | 0.273131395344267 |
| Vipas39 | ENSMUSG00000021038 | 104799 | -0.222830690818548 | 2.8183134470555 | -1.38791337907136 | 0.18780125948 | 0.273131395344267 |
| Slc24a3 | ENSMUSG00000063873 | 94249 | -0.340394568401266 | 1.02552519081026 | -1.38607696547998 | 0.18834870919 | 0.273890749923393 |
| Rhno1 | ENSMUSG00000048668 | 72440 | 0.210824946389222 | 1.43292405235049 | 1.38599393055776 | 0.18837349354 | 0.273890749923393 |
| Cdc14b | ENSMUSG00000033102 | 218294 | 0.240955904863298 | 2.05847486971474 | 1.38537840172939 | 0.18855730085 | 0.274121539525643 |
| Hmgxb4 | ENSMUSG00000034518 | 70823 | -0.209107785613476 | 3.00378476180538 | -1.38518926846983 | 0.18861380884 | 0.274164404371605 |
| Atg2a | ENSMUSG00000024773 | 329015 | -0.0999557171036392 | 4.45785079309399 | -1.38501022119217 | 0.18866731623 | 0.274164404371605 |
| Zfp423 | ENSMUSG00000045333 | 94187 | 0.0869541292634901 | 4.56248680858585 | 1.38495176810921 | 0.18868478735 | 0.274164404371605 |
| Pkn2 | ENSMUSG00000004591 | 109333 | -0.0751653849340954 | 5.544331384865 | -1.38494399308150 | 0.18868711134 | 0.274164404371605 |
| Zfp560 | ENSMUSG00000045519 | 434377 | -0.19507791712411 | 2.13524807480225 | -1.38426223700561 | 0.18889098254 | 0.274424153397184 |
| Olfr243 | ENSMUSG00000094822 | 436002 | -0.546181705662722 | 0.25625156294666 | -1.3841582393353 | 0.18892209773 | 0.274432883770126 |
| Asb5 | ENSMUSG00000031519 | 76294 | 0.151344280132129 | 6.34050579805381 | 1.38367846457683 | 0.18906569669 | 0.274584120991799 |
| Rnf10 | ENSMUSG00000041740 | 50849 | -0.0965147390066408 | 8.00147501082058 | -1.38364254790424 | 0.18907645034 | 0.274584120991799 |
| Hnrnpul2 | ENSMUSG00000071659 | 68693 | 0.0837139488991783 | 5.94225915568067 | 1.38292390846932 | 0.18929172049 | 0.274860228043429 |
| Exd2 | ENSMUSG00000032705 | 97827 | -0.167241024385217 | 3.71014898638572 | -1.38267740964969 | 0.18936560615 | 0.274930992284633 |
| Zkscan17 | ENSMUSG00000020472 | 268417 | -0.116021689781514 | 4.12174849509273 | -1.38239443210817 | 0.18945045517 | 0.275017652791685 |
| Myo1d | ENSMUSG00000035441 | 338367 | 0.209446446130772 | 3.71086142195045 | 1.38219455153179 | 0.18951040691 | 0.275041570883321 |
| Vps54 | ENSMUSG00000020128 | 245944 | -0.123517159765554 | 4.39172943634893 | -1.38217172390689 | 0.18951724577 | 0.275041570883321 |
| Olfr1260 | ENSMUSG00000042894 | 258983 | -0.83317434814667 | -0.33128536383258 | -1.3818959505302 | 0.18959999765 | 0.275125126031726 |
| Olfr77 | ENSMUSG00000051118 | 258336 | -0.361765312547798 | 1.42002003547601 | -1.38108641033089 | 0.18984306301 | 0.27541907700392 |
| Zfp36 | ENSMUSG00000044786 | 22695 | -0.367220853929288 | 3.41327556838426 | -1.38105345400916 | 0.18985296361 | 0.27541907700392 |
| Rbm12 | ENSMUSG00000089824 | 75710 | 0.274601736875546 | 2.27911804551052 | 1.38068523069138 | 0.18996361244 | 0.27554302648395 |
| Ctdspl2 | ENSMUSG00000033411 | 329506 | 0.114596564179977 | 3.27995731614477 | 1.38027233627693 | 0.19008774761 | 0.275686502483202 |
| Zfp946 | ENSMUSG00000071266 | 74149 | -0.209816056634638 | 1.75027855470191 | -1.3801799642688 | 0.19011552803 | 0.275690214505041 |
| Pyroxd1 | ENSMUSG00000041671 | 232491 | -0.153458125576329 | 2.67879028709547 | -1.37998665498988 | 0.19017367560 | 0.275737955708821 |

| | | | | | | | |
|---|---|---|---|---|---|---|---|
| Surf1 | ENSMUSG00000015790 | 20930 | -0.128580496531824 | 3.23437208379534 | -1.37938898806397 | 0.1903535466102 | 0.275962150927988 |
| Fbxo42 | ENSMUSG00000028920 | 213499 | 0.118829970914393 | 3.66344099111118 | 1.37853216450624 | 0.1906116556948 | 0.276299696114657 |
| Gstm5 | ENSMUSG00000004032 | 14866 | -0.180697892389798 | 4.10567888339349 | -1.37732142271886 | 0.1909768684307 | 0.276792382500193 |
| Ano6 | ENSMUSG00000064210 | 105722 | 0.0860147484713956 | 5.56132988790751 | 1.3771081895613 | 0.1910412482967 | 0.27684898384351 |
| Wdr74 | ENSMUSG00000042729 | 107071 | 0.176478990103769 | 2.41917503959269 | 1.37668151511453 | 0.1911701243138 | 0.276999022915527 |
| Rab2a | ENSMUSG00000047187 | 59021 | -0.085102002680169 | 6.16626142791672 | -1.37642056556334 | 0.1912489786464 | 0.277076552098713 |
| Rabac1 | ENSMUSG00000003380 | 14470 | 0.12367090498703 | 4.29503668588557 | 1.37603954103859 | 0.1913641654145 | 0.277206691246567 |
| Paqr3 | ENSMUSG00000055725 | 231474 | -0.319635814646322 | 1.09087340353532 | -1.37573010655497 | 0.1914577517918 | 0.277305510114788 |
| Rmc1 | ENSMUSG00000024410 | 76482 | -0.135208144789059 | 3.00981193696556 | -1.37516756392598 | 0.1916279851436 | 0.277515302809441 |
| Slc31a2 | ENSMUSG00000066152 | 20530 | 0.157885242760267 | 1.9713292375613 | 1.37468110955369 | 0.1917752931396 | 0.277686728791833 |
| Gprasp1 | ENSMUSG00000043384 | 67298 | 0.157142217525827 | 4.31617893880029 | 1.37460891083573 | 0.1917971642464 | 0.277686728791833 |
| Dpy30 | ENSMUSG00000024067 | 66310 | -0.138904089085214 | 3.3278381011984 | -1.37396471148147 | 0.1919924016192 | 0.277932584041785 |
| Tle1 | ENSMUSG00000008305 | 21885 | -0.175433737261192 | 3.8007209016461 | -1.37307188897701 | 0.1922632584190 | 0.278287828095683 |
| Cep78 | ENSMUSG00000041491 | 208518 | -0.216702354062271 | 1.36307991837907 | -1.37259463704708 | 0.1924081713587 | 0.278460707152393 |
| Polr2b | ENSMUSG00000029250 | 231329 | -0.102146350364478 | 4.9571042264451 | -1.37192781588215 | 0.1926107950041 | 0.278717050594412 |
| Zfp771 | ENSMUSG00000054716 | 244216 | 0.200667483683411 | 2.02780504461783 | 1.37111536549681 | 0.1928579064443 | 0.279037693625782 |
| Coa3 | ENSMUSG00000017188 | 52469 | -0.104436685215101 | 4.20878608871638 | -1.37036597510156 | 0.1930860679473 | 0.279324700836521 |
| Srsf4 | ENSMUSG00000028911 | 57317 | -0.102638884135172 | 4.48659479030673 | -1.37029601523276 | 0.1931073794000 | 0.279324700836521 |
| Pde7b | ENSMUSG00000019990 | 29863 | -0.127830229095019 | 5.25633777288994 | -1.37021132638868 | 0.1931331802281 | 0.279325058623238 |
| Tamm41 | ENSMUSG00000030316 | 68971 | -0.187371707373734 | 1.64465683775164 | -1.36996168820496 | 0.1932092500182 | 0.279398109796183 |
| Rnf126 | ENSMUSG00000035890 | 70294 | -0.118003228679084 | 4.16463487398074 | -1.36962384471979 | 0.1933122368005 | 0.27951006090837 |
| Ppil2 | ENSMUSG00000022771 | 66053 | -0.103254500497735 | 3.47916758697984 | -1.36726079417289 | 0.1940338360384 | 0.280516315531872 |
| Psmb7 | ENSMUSG00000026750 | 19177 | -0.105904837424506 | 5.59473293925092 | -1.36705495256056 | 0.1940967976204 | 0.280570232053571 |
| Med27 | ENSMUSG00000026799 | 68975 | -0.133093745452325 | 2.68691926881095 | -1.36572030834801 | 0.1945054360533 | 0.281123750032913 |
| Tap2 | ENSMUSG00000024339 | 21355 | -0.15498301726385 | 2.89239483851017 | -1.36533845158471 | 0.1946224814753 | 0.281255730434536 |
| Zfp513 | ENSMUSG00000043059 | 101023 | 0.248136703485328 | 1.51442190653319 | 1.36475059836283 | 0.1948027807880 | 0.281479074262284 |
| Aprt | ENSMUSG00000006589 | 11821 | 0.12811367400027 | 2.89035064369463 | 1.36447839534947 | 0.1948863138849 | 0.281541220020072 |
| Senp5 | ENSMUSG00000022772 | 320213 | 0.122241802597435 | 4.09706632868331 | 1.36444259519315 | 0.1948973023412 | 0.281541220020072 |
| Tbc1d20 | ENSMUSG00000027465 | 67231 | 0.0956419566116802 | 4.47113902348877 | 1.36293192173756 | 0.1953614483774 | 0.282174417575812 |
| Mrgbp | ENSMUSG00000027569 | 73247 | -0.242925350565594 | 1.63625326012691 | -1.36277556208892 | 0.1954095405537 | 0.282206591067868 |
| Gnas | ENSMUSG00000027523 | 14683 | -0.0926333674838169 | 9.08280632966453 | -1.36229721411954 | 0.1955567280610 | 0.282381848670491 |
| Psmc2 | ENSMUSG00000028932 | 19181 | -0.0838608655856579 | 5.53188108081683 | -1.36221296594679 | 0.1955826605740 | 0.282381992172107 |
| Mylpf | ENSMUSG00000030672 | 17907 | 0.163351887834009 | 9.09870460975779 | 1.36199162106424 | 0.1956508063474 | 0.282443075037517 |
| Shprh | ENSMUSG00000090112 | 268281 | 0.128336493838455 | 4.14308108697722 | 1.36130786284272 | 0.1958614385064 | 0.282709809099976 |
| Ap5m1 | ENSMUSG00000036291 | 74385 | -0.137249927122802 | 2.93130706962046 | -1.36109255407124 | 0.1959278028430 | 0.282768261536533 |
| Noa1 | ENSMUSG00000036285 | 56412 | -0.12474210353566 | 3.35560272296635 | -1.36096191571812 | 0.1959680782804 | 0.282789051311437 |
| Myg1 | ENSMUSG00000001285 | 60315 | -0.196089127782334 | 1.89642744841936 | -1.36085941611001 | 0.1959996833554 | 0.282797325601687 |
| Cd3eap | ENSMUSG00000047649 | 70333 | -0.12961019150461 | 3.68105523564233 | -1.35960483703327 | 0.1963868618780 | 0.283318567663915 |
| Hyal1 | ENSMUSG00000010051 | 15586 | 0.39758425398028 | 1.03751609830344 | 1.35944359958751 | 0.1964366669884 | 0.283353022809541 |
| Cpe | ENSMUSG00000037852 | 12876 | 0.103467317797485 | 5.8459022401301 | 1.35918746592914 | 0.1965158060700 | 0.28342977650763 |
| Dip2c | ENSMUSG00000048264 | 208440 | -0.106226143702836 | 4.94710742812622 | -1.35831496868063 | 0.1967855818801 | 0.283781424212602 |
| Pdcl | ENSMUSG00000009030 | 67466 | 0.163172251137515 | 3.39872958000648 | 1.35821185286679 | 0.1968174852122 | 0.283789992198307 |
| Mtcl1 | ENSMUSG00000052105 | 68617 | 0.121281148087632 | 4.03358212221499 | 1.35806503338067 | 0.1968629174472 | 0.283818062597697 |
| Vmn2r124 | ENSMUSG00000094396 | 637021 | -0.658097817585661 | 0.480758705327995 | -1.35743436781849 | 0.1970581697190 | 0.284046150220539 |
| Grip2 | ENSMUSG00000030098 | 243547 | -0.227607364552048 | 2.7706945035404 | -1.35731428590506 | 0.1970953646408 | 0.284046150220539 |
| Gtpbp1 | ENSMUSG00000042535 | 14904 | -0.123540523558401 | 4.24183740710124 | -1.35730229031769 | 0.1970990805436 | 0.284046150220539 |
| Alkbh1 | ENSMUSG00000079036 | 211064 | 0.214076710688474 | 2.04166721111479 | 1.35699102922365 | 0.1971955206493 | 0.284147671528907 |
| Matn4 | ENSMUSG00000016995 | 17183 | -0.399279447546987 | 0.974787737900262 | -1.35673845815989 | 0.1972738047209 | 0.284223007698066 |
| Harbi1 | ENSMUSG00000027243 | 241547 | 0.170497472788807 | 1.96090559340999 | 1.35664205537741 | 0.1973036913219 | 0.284228604354263 |
| Entpd1 | ENSMUSG00000048120 | 12495 | 0.183867911031353 | 3.78760328970689 | 1.35612705552659 | 0.1974634131371 | 0.284421210831724 |
| Mllt10 | ENSMUSG00000026743 | 17354 | -0.154328864595408 | 3.70222748260474 | -1.35512489760029 | 0.1977745242058 | 0.284831795005802 |
| Zmym4 | ENSMUSG00000042446 | 67785 | -0.08174191713357 | 4.55484018586039 | -1.35478696152644 | 0.1978795235459 | 0.284945471200194 |
| Gpr155 | ENSMUSG00000041762 | 68526 | 0.12979007867226 | 2.95000098745547 | 1.35414607766837 | 0.1980787759242 | 0.285194823449439 |
| Wdr55 | ENSMUSG00000042660 | 67936 | 0.164697593457878 | 2.24490859327653 | 1.35399990755928 | 0.1981242434719 | 0.285205937596298 |
| Kmt5a | ENSMUSG00000049327 | 67956 | -0.106675349346339 | 5.23538846625412 | -1.35395350856681 | 0.1981386780812 | 0.285205937596298 |
| Syncrip | ENSMUSG00000032423 | 56403 | 0.0934061650840139 | 6.20011595675669 | 1.35363309886164 | 0.1982383801318 | 0.285301622013801 |
| Sf3b4 | ENSMUSG00000068856 | 107701 | -0.169327078904467 | 2.72661966494077 | -1.35351921234182 | 0.1982738281072 | 0.285301622013801 |
| Mvb12a | ENSMUSG00000031813 | 73711 | -0.144872457838068 | 2.94727411970505 | -1.35348829322786 | 0.1982834527892 | 0.285301622013801 |
| Mrps12 | ENSMUSG00000045948 | 24030 | 0.13066431925562 | 3.24552045077864 | 1.35298350836679 | 0.1984406390256 | 0.285490211264711 |
| Snx32 | ENSMUSG00000056185 | 225861 | 0.275549323688975 | 1.26655113590859 | 1.3528593438642 | 0.1984793184820 | 0.285508281395891 |
| H3f3c | ENSMUSG00000082029 | 625328 | 2.804568553035 | -1.98227173339121 | 1.35267005142086 | 0.1985382982760 | 0.285555544533686 |
| Tbk1 | ENSMUSG00000020115 | 56480 | -0.149029808791063 | 3.58650471785621 | -1.35217722934733 | 0.1986919189146 | 0.285738898257656 |
| Tmem135 | ENSMUSG00000039428 | 72759 | -0.107392586839693 | 3.85061168984468 | -1.35197041774522 | 0.1987564142899 | 0.285794049540095 |
| Hs1bp3 | ENSMUSG00000020605 | 58240 | -0.203328073956334 | 2.04037154330957 | -1.35062348132764 | 0.1991768814537 | 0.286326245031397 |
| Sema6c | ENSMUSG00000038777 | 20360 | -0.279903871179874 | 5.01691406003216 | -1.35061695877266 | 0.1991789193320 | 0.286326245031397 |
| Ywhag | ENSMUSG00000051391 | 22628 | -0.0702275122654017 | 7.32095807570023 | -1.35044328680985 | 0.1992331868797 | 0.28636659642105 |
| Pan2 | ENSMUSG00000005682 | 103135 | -0.179879707056435 | 2.42126740991927 | -1.34976145738141 | 0.1994463556195 | 0.286613176587733 |
| Gadd45g | ENSMUSG00000021453 | 23882 | -0.329457202391371 | 2.78047851357214 | -1.34972685104737 | 0.1994571799746 | 0.286613176587733 |
| Tubgcp3 | ENSMUSG00000000759 | 259279 | 0.102896786659198 | 3.29758436607836 | 1.34798439487404 | 0.2000028133927 | 0.287359457570403 |
| Rab10os | ENSMUSG00000079179 | 74173 | -0.28073247393899 | 1.38273738515301 | -1.34783660872854 | 0.2000491470395 | 0.287388254210712 |
| Gas2 | ENSMUSG00000030498 | 14453 | 0.315422130691381 | 1.38117723993201 | 1.34731557147167 | 0.2002125713958 | 0.287585232316477 |
| 4921524J17Rik | ENSMUSG00000036934 | 66714 | -0.176305803017517 | 2.37975945124344 | -1.34713351943831 | 0.2002696979759 | 0.287629492652098 |
| Ccdc9 | ENSMUSG00000041375 | 243846 | 0.122198445106869 | 3.5321251607575 | 1.34691993144175 | 0.2003367372144 | 0.287687976154933 |
| Btrc | ENSMUSG00000025217 | 12234 | -0.114108988193038 | 3.66990334102135 | -1.34622018794647 | 0.2005564948280 | 0.287965722443957 |
| Ccdc58 | ENSMUSG00000075229 | 381045 | -0.222300541675793 | 1.90063239027033 | -1.34531353604372 | 0.2008415245550 | 0.288337103350607 |
| Med26 | ENSMUSG00000045248 | 70625 | -0.189201063220181 | 2.74031207719276 | -1.34518200133688 | 0.2008829032798 | 0.288358636342956 |
| Gskip | ENSMUSG00000044715 | 66787 | 0.206331926157253 | 2.03189002913508 | 1.34509274016324 | 0.2009109873724 | 0.288361082322532 |
| Pex13 | ENSMUSG00000020283 | 72129 | -0.118511262886343 | 3.67187594625023 | -1.34438914844414 | 0.2011324690341 | 0.288641068532596 |
| Tox4 | ENSMUSG00000016831 | 268741 | -0.0884355657375764 | 4.99174668525025 | -1.34418217595929 | 0.2011976590957 | 0.288696719818226 |
| Ampd3 | ENSMUSG00000005686 | 11717 | 0.111395972156508 | 5.41906263756881 | 1.34317000900144 | 0.2015167084610 | 0.289101905019837 |
| Keap1 | ENSMUSG00000003308 | 50868 | -0.125795076065564 | 4.82851933200459 | -1.34307084576317 | 0.2015479882239 | 0.289101905019837 |
| Kcnq5 | ENSMUSG00000028033 | 226922 | 0.139280257611303 | 3.76808459429401 | 1.34303472435463 | 0.2015593832363 | 0.289101905019837 |
| Eef1aknmt | ENSMUSG00000026694 | 71449 | -0.206890127139799 | 1.82821685114668 | -1.34287339024623 | 0.2016102847686 | 0.289118934431991 |
| Agfg1 | ENSMUSG00000026159 | 15463 | 0.0919939965261625 | 4.58482217535633 | 1.34282943415127 | 0.2016241548966 | 0.289118934431991 |
| Mafa | ENSMUSG00000047591 | 378435 | 0.37272219013819 | 2.6826428637305 | 1.34262241265732 | 0.2016894899304 | 0.289174687097245 |
| 4632427E13Rik | ENSMUSG00000074024 | 666737 | 0.293163508413467 | 2.27927849283448 | 1.34210291254825 | 0.2018535175397 | 0.289371908226461 |
| Srsf9 | ENSMUSG00000029538 | 108014 | 0.163094140310128 | 2.41833193534137 | 1.34189587005508 | 0.2019189195937 | 0.289427709163321 |
| Tubgcp4 | ENSMUSG00000027263 | 51885 | 0.11993222919063 | 3.41653540663517 | 1.34051397819934 | 0.2023558824595 | 0.290016015624196 |
| Galnt15 | ENSMUSG00000021903 | 78754 | -0.204475074731584 | 3.85337459959134 | -1.33959304104861 | 0.2026475147952 | 0.290395907738134 |
| A230077H06Rik | ENSMUSG00000100600 | 320210 | -0.547550254785853 | 0.75858843028743 | -1.33844698353082 | 0.2030109122847 | 0.290878526960835 |
| Zxdb | ENSMUSG00000073062 | 668166 | 0.354886459776797 | 1.27884357083476 | 1.33809776318091 | 0.2031217498642 | 0.290999193678331 |
| Tarbp1 | ENSMUSG00000090290 | 212728 | 0.130648506139935 | 2.97907485798528 | 1.3377168992575 | 0.2032426866512 | 0.291105655855872 |
| Slc2a8 | ENSMUSG00000026791 | 56017 | -0.179796732351871 | 1.72854750740022 | -1.33767373840111 | 0.2032563953243 | 0.291105655855872 |
| Unc119b | ENSMUSG00000046562 | 106840 | 0.192187701808962 | 3.05626870902179 | 1.33761215810865 | 0.2032759556443 | 0.291105655855872 |
| G6pdx | ENSMUSG00000031400 | 14381 | 0.255523342940547 | 1.86884684817839 | 1.33682886136412 | 0.2035248947762 | 0.291423974953975 |
| Malat1 | ENSMUSG00000092341 | 72289 | 0.254546133598172 | 11.1096355798203 | 1.33638121891472 | 0.2036672708250 | 0.291589644713605 |
| Cdc42ep4 | ENSMUSG0000041598 | 56699 | 0.157281727640618 | 2.74841834665517 | 1.33623007398141 | 0.2037153618511 | 0.2916203012565 |
| Ttn | ENSMUSG00000051747 | 22138 | 0.268079203960562 | 15.5748852695347 | 1.33607612992787 | 0.2037643529650 | 0.291652238085812 |
| Dmtf1 | ENSMUSG00000042508 | 23857 | 0.12471385707104 | 3.30998142872786 | 1.33563908172479 | 0.2039034911492 | 0.291813179072119 |
| Laptm4a | ENSMUSG00000020585 | 17775 | -0.0943043986377834 | 6.19422432629829 | -1.33554572020212 | 0.2039332236075 | 0.291817524185641 |
| Manea | ENSMUSG00000040520 | 242362 | 0.165640532527255 | 2.63459486054759 | 1.3353368296536 | 0.2039997608525 | 0.291874526947488 |
| Pot1b | ENSMUSG00000024174 | 72836 | -0.444277595394903 | 2.38636589925921 | -1.33482613536219 | 0.2041625048269 | 0.292069145434307 |
| Thrsp | ENSMUSG00000035686 | 21835 | -0.188619659261704 | 2.33347548424776 | -1.33462491515762 | 0.2042266569797 | 0.292122688752365 |
| Klhl26 | ENSMUSG00000055707 | 234378 | -0.114432694950036 | 2.94133161454149 | -1.33407374036841 | 0.2044024638413 | 0.292335906352227 |
| Ifit3b | ENSMUSG00000062488 | 667370 | -0.485206554077868 | 1.75007961350223 | -1.33363309842248 | 0.2045431025536 | 0.292498777343476 |
| Clasrp | ENSMUSG00000061028 | 53609 | -0.174236641400984 | 2.91852386163838 | -1.3326733153995 | 0.2048497058717 | 0.292879477995798 |
| Apex1 | ENSMUSG00000035960 | 11792 | 0.178896658387951 | 2.83116984386956 | 1.33260520299853 | 0.2048714785699 | 0.292879477995798 |
| Golm1 | ENSMUSG00000021556 | 105348 | 0.190024762641598 | 2.36413775775152 | 1.33254818821627 | 0.2048897052633 | 0.292879477995798 |
| Trappc13 | ENSMUSG00000021711 | 66975 | -0.112008087165294 | 4.01543506297158 | -1.33226907835886 | 0.2049789510832 | 0.292968738793284 |
| Arid5a | ENSMUSG00000037447 | 214855 | 0.437710699172909 | 3.48616793381956 | 1.33194852460655 | 0.2050814875146 | 0.293076969541487 |
| Bcl7a | ENSMUSG00000029438 | 77045 | -0.129407899047052 | 3.94772724241522 | -1.331758710609 | 0.2051422234404 | 0.29310883178832 |
| Dynlt1f | ENSMUSG00000095677 | 100040531 | -0.266016333337993 | 2.368489135805 | -1.33171124950746 | 0.2051574121317 | 0.29310883178832 |
| Herc1 | ENSMUSG00000038664 | 235439 | -0.114323220139156 | 6.71555374780962 | -1.3315730146945 | 0.2052016557801 | 0.293133729656734 |
| Zdhhc17 | ENSMUSG00000035798 | 320150 | -0.131971139407061 | 3.44155402753255 | -1.33099222818444 | 0.2053876276809 | 0.293361055557311 |
| Extl1 | ENSMUSG00000028838 | 56219 | 0.107997384607544 | 5.00195755183893 | 1.3305847909119 | 0.2055181734095 | 0.293488258583429 |
| Elovl5 | ENSMUSG00000032349 | 68801 | -0.20254274879882 | 3.29769464430167 | -1.33045159143572 | 0.2055608660197 | 0.293488258583429 |

| | | | | | | | |
|---|---|---|---|---|---|---|---|
| Slc45a4 | ENSMUSG00000079020 | 106068 | 0.290099428229222 | 3.61423311034622 | 1.33040391714904 | 0.2055761481536 | 0.293488258583429 |
| Asna1 | ENSMUSG00000052456 | 56495 | -0.122287582114228 | 4.16132353924643 | -1.33034578603559 | 0.2055947834992 | 0.293488258583429 |
| Plcb3 | ENSMUSG00000024960 | 18797 | -0.114688071378748 | 3.95379426801383 | -1.33029541967997 | 0.2056109307686 | 0.293488258583429 |
| Gtf3c4 | ENSMUSG00000035666 | 269252 | 0.0987175879847237 | 4.13100855305532 | 1.32966981579462 | 0.2058115827045 | 0.293736311599884 |
| Mpv17 | ENSMUSG00000107283 | 17527 | -0.107025118725709 | 3.43058082584218 | -1.32934895534913 | 0.2059145547998 | 0.293844908422546 |
| Fam92a | ENSMUSG00000028218 | 68099 | 0.18706411305222 | 2.69004553173256 | 1.32824590811337 | 0.2062688682991 | 0.294302906379445 |
| Ddx51 | ENSMUSG00000029504 | 69663 | 0.176997537912935 | 2.19781536826681 | 1.32818219608039 | 0.2062893485206 | 0.294302906379445 |
| Diaph2 | ENSMUSG00000034480 | 54004 | 0.167159786016474 | 3.42273860608501 | 1.32788038312158 | 0.2063863886077 | 0.294402924947381 |
| Cript | ENSMUSG00000024146 | 56724 | -0.140045782138139 | 3.25271524289149 | -1.32776009676644 | 0.2064250738435 | 0.294419687132553 |
| Elp3 | ENSMUSG00000022031 | 74195 | -0.109685484564874 | 4.54284231772685 | -1.32708789382131 | 0.2066413687438 | 0.29468973277738 |
| Nbas | ENSMUSG00000020576 | 71169 | -0.0883013042149967 | 4.57246823611517 | -1.32686388620935 | 0.2067134884513 | 0.294754127610477 |
| Mlf2 | ENSMUSG00000030120 | 30853 | -0.0935774621711003 | 5.85933954083252 | -1.32672382697263 | 0.2067585911546 | 0.294761892331193 |
| Olfr509 | ENSMUSG00000049280 | 258369 | -0.515847806179466 | 0.914328984250266 | -1.32667950610153 | 0.2067728652818 | 0.294761892331193 |
| Msh6 | ENSMUSG00000005370 | 17688 | -0.15750855614928 | 2.68420050583259 | -1.32477618293998 | 0.2073866084680 | 0.295598254943775 |
| Lars | ENSMUSG00000024493 | 107045 | -0.0921771582890623 | 5.2047919265006 | -1.32419671163211 | 0.2075737563103 | 0.295826431581308 |
| Hoxc4 | ENSMUSG00000075394 | 15423 | 0.187452357698258 | 1.5098645374348 | 1.32408303000833 | 0.2076104872948 | 0.29584020813718 |
| Btg3 | ENSMUSG00000022863 | 12228 | 0.175212013835617 | 2.01099259301135 | 1.32394199748843 | 0.2076560627656 | 0.295866582649998 |
| Smox | ENSMUSG00000027333 | 228608 | -0.181531686384263 | 2.70519597360333 | -1.32384629422703 | 0.2076869944317 | 0.295872088639872 |
| Yeats2 | ENSMUSG00000041215 | 208146 | 0.206547733974725 | 2.90461214770679 | 1.32354160643222 | 0.2077854955174 | 0.295973840435455 |
| Nelfb | ENSMUSG00000013465 | 58202 | 0.106517482119836 | 3.52613454982692 | 1.32318722910773 | 0.2079001079965 | 0.296098512118649 |
| Mfap3l | ENSMUSG00000031647 | 71306 | 0.197425991869292 | 1.8418469947668 | 1.32306569638721 | 0.2079394257931 | 0.29611592800219 |
| 5730409E04Rik | ENSMUSG00000073755 | 230757 | 0.112578187353861 | 3.94456262877535 | 1.32262434503733 | 0.2080822606206 | 0.296280733469337 |
| Pdp1 | ENSMUSG00000049225 | 381511 | -0.297873930453752 | 4.83055983312022 | -1.32184457186332 | 0.2083348129077 | 0.296601698345279 |
| Dcp2 | ENSMUSG00000024472 | 70640 | -0.0989653557097472 | 3.91155373219022 | -1.32138092177828 | 0.2084850968824 | 0.296777001435413 |
| Pkig | ENSMUSG00000035268 | 18769 | 0.0975679340754949 | 5.59922939042424 | 1.32013394217491 | 0.2088897177852 | 0.297314258477959 |
| Bcap29 | ENSMUSG00000020650 | 12033 | -0.172454373630541 | 3.46999464574011 | -1.3195410940261 | 0.2090823079090 | 0.297549629964038 |
| Ifi47 | ENSMUSG00000078920 | 15953 | -0.49780568967323 | 0.612270702576281 | -1.31907777087464 | 0.2092329208779 | 0.297725209335639 |
| Sars2 | ENSMUSG00000070699 | 71984 | -0.166025314111494 | 2.77963750913256 | -1.31839521661717 | 0.2094549590907 | 0.298002363376452 |
| Cxxc1 | ENSMUSG00000024560 | 74322 | -0.110754388926054 | 3.4130887867209 | -1.31800413840172 | 0.2095822644124 | 0.298144681453936 |
| Lonrf3 | ENSMUSG00000016239 | 74365 | 0.325630282061308 | 0.941228627415758 | 1.31744353829145 | 0.2097648620776 | 0.298365609287093 |
| Nop14 | ENSMUSG00000036693 | 75416 | 0.0954851630183953 | 4.5034988867065 | 1.31725122140087 | 0.2098275327574 | 0.298415919928614 |
| Eif2a | ENSMUSG00000027810 | 229317 | -0.0962479083818506 | 4.12763292610273 | -1.31687683417349 | 0.2099495783851 | 0.298550649320625 |
| Bmf | ENSMUSG00000040093 | 171543 | 0.349555994198929 | 1.53526737315998 | 1.31641443998089 | 0.2101003922429 | 0.298726247087343 |
| Iars2 | ENSMUSG00000026618 | 381314 | -0.0842954905891453 | 5.55849996185338 | -1.3159684123494 | 0.2102459508018 | 0.298894328029026 |
| Socs7 | ENSMUSG00000038485 | 192157 | 0.111654487976742 | 5.64249116631018 | 1.31582266391012 | 0.2102935325978 | 0.298923095556157 |
| Cox15 | ENSMUSG00000040018 | 226139 | 0.103061860536291 | 4.34080044233597 | 1.31564746154931 | 0.2103507415698 | 0.298965538434575 |
| Plscr3 | ENSMUSG00000019461 | 70310 | 0.174990175982026 | 2.76412838749081 | 1.31551203511602 | 0.2103949710759 | 0.298989525329014 |
| Ing5 | ENSMUSG00000026283 | 66262 | -0.16682819310304 | 2.3389463747583 | -1.31483781605822 | 0.2106152788290 | 0.299263695941822 |
| Parp14 | ENSMUSG00000034422 | 547253 | -0.225216397219886 | 4.29624605648065 | -1.31412968729774 | 0.2108468671538 | 0.299553821790842 |
| Nudt2 | ENSMUSG00000028443 | 66401 | -0.157865966889037 | 2.24706707562608 | -1.31397740382884 | 0.2108966971590 | 0.299585678576412 |
| Kctd17 | ENSMUSG00000033287 | 72844 | 0.207632415177056 | 2.99480470524892 | 1.31351100282512 | 0.2110493713734 | 0.299763601673963 |
| Casp7 | ENSMUSG00000025076 | 12369 | -0.187521430019785 | 1.65414391417395 | -1.31279947137462 | 0.2112824595161 | 0.300055679481797 |
| Aamdc | ENSMUSG00000035642 | 66273 | -0.101906705539178 | 4.52361891147252 | -1.31191911284868 | 0.2115711402152 | 0.300419100912293 |
| Pde12 | ENSMUSG00000043702 | 211948 | 0.142471233864458 | 3.1912466311594 | 1.31185149211733 | 0.2115933270445 | 0.300419100912293 |
| Cep112 | ENSMUSG00000020728 | 76380 | 0.222398392630708 | 1.49435797866083 | 1.3109851389429 | 0.2118777493845 | 0.300783854353638 |
| Vcpip1 | ENSMUSG00000045210 | 70675 | 0.092111832087177 | 4.51250066381007 | 1.31075186114011 | 0.2119543867084 | 0.300853577601681 |
| Lama4 | ENSMUSG00000019846 | 16775 | 0.185788503520155 | 5.47626800447546 | 1.31061519368554 | 0.2119992955982 | 0.300878252425867 |
| Bach2 | ENSMUSG00000040270 | 12014 | 0.241728518989233 | 1.93021952198474 | 1.30900209730321 | 0.2125299378256 | 0.301592204384349 |
| Bud13 | ENSMUSG00000032077 | 215051 | -0.169282265017414 | 2.45223416291471 | -1.3085606065652 | 0.2126753563036 | 0.301759387299456 |
| Ddx5 | ENSMUSG00000020719 | 13207 | -0.0953398757215194 | 7.61296337619924 | -1.30720090652233 | 0.2131237182638 | 0.302356309453897 |
| Ddx46 | ENSMUSG00000021500 | 212880 | -0.117956739843009 | 5.38512221399821 | -1.30702029483224 | 0.2131833322649 | 0.302401635736886 |
| Ccdc71 | ENSMUSG00000049305 | 72454 | -0.119718550703978 | 3.40941456238452 | -1.30648914203818 | 0.2133587261392 | 0.302611163283765 |
| Rps6kb2 | ENSMUSG00000024830 | 58988 | 0.102680024352942 | 3.6334205531044 | 1.30571647848715 | 0.2136140773718 | 0.302934026952761 |
| Tpp1 | ENSMUSG00000030894 | 12751 | -0.102902160845808 | 4.44914791321331 | -1.3055623694871 | 0.2136650369778 | 0.302966989130251 |
| Icam1 | ENSMUSG00000037405 | 15894 | 0.265014281369199 | 2.42770143879615 | 1.30533847738858 | 0.2137390893528 | 0.303032682972283 |
| Nsun2 | ENSMUSG00000021595 | 28114 | 0.103673839499926 | 5.17879975548355 | 1.30491840399807 | 0.213878084431 | 0.30319042159451 |
| Cpne3 | ENSMUSG00000028228 | 70568 | 0.0927253250093982 | 4.03027026688528 | 1.30412504435529 | 0.2141407917577 | 0.303523469230334 |
| Crebbp | ENSMUSG00000022521 | 12914 | 0.0773991191700887 | 6.49389792759002 | 1.30336340794924 | 0.2143932384611 | 0.303841888969134 |
| 1700024F13Rik | ENSMUSG00000097709 | 73270 | -0.590098141505942 | 0.24015156196513 | -1.30251285849301 | 0.2146754382927 | 0.304202387344838 |
| Bmp2 | ENSMUSG00000027358 | 12156 | -0.214692720645512 | 1.52013422358042 | -1.30230463542115 | 0.2147445691103 | 0.304257399445794 |
| Siah2 | ENSMUSG00000036432 | 20439 | 0.161457808139844 | 3.14952679929958 | 1.30222826448839 | 0.2147699290205 | 0.304257399445794 |
| Tmpo | ENSMUSG00000019961 | 21917 | -0.0747696034272849 | 4.32166635383307 | -1.30152639515218 | 0.2150031060686 | 0.304521040854277 |
| Tmem182 | ENSMUSG00000079588 | 381339 | -0.105134923255372 | 7.09029947399955 | -1.30142880581401 | 0.2150355435761 | 0.304521040854277 |
| Prdm15 | ENSMUSG00000014039 | 114604 | -0.149396986145226 | 2.53133938706079 | -1.30141658989052 | 0.2150396042772 | 0.304521040854277 |
| Fam13c | ENSMUSG00000043259 | 71721 | -0.191332578809196 | 2.58539830547286 | -1.30096143844566 | 0.2151909452910 | 0.30466752757082 |
| Aimp2 | ENSMUSG00000029610 | 231872 | -0.125541388351449 | 3.59164128052788 | -1.30093785094426 | 0.2151987906297 | 0.30466752757082 |
| Plcd3 | ENSMUSG00000020937 | 72469 | 0.299375055393433 | 1.29888733086077 | 1.30040441848909 | 0.2153762746778 | 0.304879313455504 |
| C2cd2l | ENSMUSG00000032120 | 71764 | 0.145414954172587 | 3.1629210391254 | 1.30017232563369 | 0.2154535334746 | 0.304949187415647 |
| Klf12 | ENSMUSG00000072294 | 16597 | -0.159533498564396 | 3.91801840467413 | -1.29951681844305 | 0.2156718581142 | 0.305218679576258 |
| Pde4b | ENSMUSG00000028525 | 18578 | 0.26924761801475 | 4.89616313673687 | 1.29903697954796 | 0.2158317868504 | 0.30540547081711 |
| Helz2 | ENSMUSG00000027580 | 229003 | -0.262361487752929 | 3.70439627801148 | -1.29846683861861 | 0.2160219366896 | 0.305597106640996 |
| Naa25 | ENSMUSG00000042719 | 231713 | 0.0900095857766257 | 4.70886640170766 | 1.29846325943252 | 0.2160231308219 | 0.305597106640996 |
| Smg5 | ENSMUSG00000001415 | 229512 | -0.0874354910367074 | 5.76150625654969 | -1.29831168508491 | 0.2160737057913 | 0.305629099120738 |
| Sf1 | ENSMUSG00000024949 | 22668 | -0.0940767850228115 | 5.73156080441573 | -1.29743537627783 | 0.2163662851593 | 0.306003346238481 |
| Asah2 | ENSMUSG00000024887 | 54447 | -0.159694127687978 | 3.11166044930252 | -1.29696659839793 | 0.2165229297637 | 0.306185271054041 |
| Tnrc6c | ENSMUSG00000025571 | 217351 | 0.0923976152926125 | 5.39350188154867 | 1.29662558921478 | 0.2166369369318 | 0.306306862967502 |
| Papln | ENSMUSG00000021223 | 170721 | -0.316770024894905 | 2.19695292807826 | -1.29598802391247 | 0.2168502187498 | 0.306568771489156 |
| Itpr2 | ENSMUSG00000030287 | 16439 | 0.14088348147387 | 4.30009827502873 | 1.29576234029878 | 0.2169257560028 | 0.306635903125228 |
| Lzts1 | ENSMUSG00000036306 | 211134 | -0.291267267534103 | 1.92140687457556 | -1.29559376178442 | 0.2169821937050 | 0.306676022677758 |
| Hmgn1 | ENSMUSG00000040681 | 15312 | 0.0928581496086895 | 4.90856527648063 | 1.29533326975701 | 0.2170694259034 | 0.306759650232851 |
| Elac1 | ENSMUSG00000036941 | 114615 | -0.102035416881602 | 2.87285175518306 | -1.29482073730194 | 0.2172411421593 | 0.306962632490212 |
| Taf9 | ENSMUSG00000078941 | 108143 | -0.137770808065668 | 4.24587123454797 | -1.29430245474465 | 0.2174148956190 | 0.307151770025362 |
| Lncpint | ENSMUSG00000044471 | 232685 | 0.205482035395018 | 3.08591786901052 | 1.29425385191252 | 0.2174311953578 | 0.307151770025362 |
| Dnajb6 | ENSMUSG00000029131 | 23950 | -0.0800810091053288 | 5.89026530327356 | -1.29416668024109 | 0.2174604322286 | 0.307153377011477 |
| Trpm7 | ENSMUSG00000027365 | 58800 | -0.0913045339641698 | 5.22880771834298 | -1.29404727267181 | 0.2175004859461 | 0.307170259932461 |
| Rp1 | ENSMUSG00000025900 | 19888 | 0.159104337377619 | 4.62019631834626 | 1.29327457426584 | 0.2177598205320 | 0.307493680389176 |
| Ranbp10 | ENSMUSG00000037415 | 74334 | 0.0953265925274147 | 5.06462257735038 | 1.29300869785517 | 0.2178491117811 | 0.307493680389176 |
| 2610005L07Rik | ENSMUSG00000071793 | 381598 | 0.15340664753117 | 3.76109998964742 | 1.29295758385718 | 0.2178662811361 | 0.307493680389176 |
| Carmil1 | ENSMUSG00000021338 | 68732 | 0.317431806294459 | 1.27154846514846 | 1.29294769153177 | 0.2178696041250 | 0.307493680389176 |
| Grrp1 | ENSMUSG00000050105 | 72690 | 0.29055877516375 | 1.45448296016663 | 1.29294608149671 | 0.2178701449651 | 0.307493680389176 |
| Rcor3 | ENSMUSG00000037395 | 214742 | -0.0979573875105446 | 4.50709827933494 | -1.2925572905403 | 0.2180007784328 | 0.307638330628653 |
| Preb | ENSMUSG00000045302 | 50907 | 0.097428303057157 | 4.37120800675558 | 1.29208147394488 | 0.2181607378311 | 0.307824322348273 |
| Ank2 | ENSMUSG00000032826 | 109676 | -0.119139952662714 | 6.19871517389768 | -1.29168725220862 | 0.2182933379800 | 0.307971667199272 |
| Zfp410 | ENSMUSG00000042472 | 52708 | -0.150034248391156 | 3.11305948192189 | -1.29113181593567 | 0.2184802736733 | 0.308195621567085 |
| Smyd3 | ENSMUSG00000055067 | 69726 | -0.209504363239934 | 1.29531414558985 | -1.28967178400319 | 0.2189722683022 | 0.308849789007953 |
| Rbm48 | ENSMUSG00000040302 | 269623 | -0.20494299619854 | 1.5633710346701 | -1.28912013821831 | 0.2191583899673 | 0.309072424297896 |
| Wapl | ENSMUSG00000041408 | 218914 | -0.0773304943591899 | 5.94438590090208 | -1.28894682633528 | 0.2192168903888 | 0.309115045000104 |
| Slc29a1 | ENSMUSG00000023942 | 63959 | 0.108075339410679 | 4.16955370619099 | 1.28811228266956 | 0.2194987607479 | 0.309472585287674 |
| Gclm | ENSMUSG00000028124 | 14630 | 0.148273926910375 | 2.91673743590739 | 1.28787156608614 | 0.2195801175688 | 0.309547364604798 |
| Gak | ENSMUSG00000062234 | 231580 | 0.137628713556244 | 3.87502051878888 | 1.28710010005438 | 0.2198410183877 | 0.309875199483718 |
| Lmln | ENSMUSG00000022802 | 239833 | 0.166390442463853 | 1.79010246474495 | 1.28582161770892 | 0.2202739320615 | 0.310445377947967 |
| Riox1 | ENSMUSG00000046791 | 71952 | -0.229597332451891 | 1.2909230101483 | -1.28472039525601 | 0.2206473689810 | 0.310931596020638 |
| Cyb5a | ENSMUSG00000024646 | 109672 | -0.130988940169011 | 3.0526862798889 | -1.28434108077725 | 0.2207761158949 | 0.311072921222945 |
| Lmod2 | ENSMUSG00000029683 | 93677 | -0.138050720783243 | 9.34125248568 | -1.28419452157619 | 0.2208258770935 | 0.311102933691176 |
| Mtdh | ENSMUSG00000022255 | 67154 | -0.0739013739075824 | 6.57490566128705 | -1.28340443413941 | 0.2210942896407 | 0.311440938152521 |
| Snora23 | ENSMUSG00000064451 | 100379145 | 1.02361506559446 | -0.997598092972392 | 1.28300484058283 | 0.2212301408291 | 0.311562198418652 |
| Zfp457 | ENSMUSG00000055341 | 431706 | -0.412327287537977 | 0.592138322250673 | -1.28298355722132 | 0.2212373784764 | 0.311562198418652 |
| Oas2 | ENSMUSG00000032690 | 246728 | -0.408724061228437 | 1.94238325344418 | -1.282745657435 | 0.2213182918578 | 0.311635997462025 |
| Tra2b | ENSMUSG00000022858 | 20462 | 0.0847422583734305 | 4.98049584813614 | 1.28247615999677 | 0.2214099806414 | 0.311724948273039 |
| Nek4 | ENSMUSG00000021918 | 23955 | -0.167554288845859 | 2.39181219179328 | -1.28221664629415 | 0.2214983014172 | 0.311809134937791 |
| Rpl17 | ENSMUSG00000062328 | 319195 | -0.113017258769241 | 6.50980017823159 | -1.28158272165049 | 0.2217141644728 | 0.312036084962714 |
| Klhl12 | ENSMUSG00000026455 | 240756 | -0.0889391505109121 | 4.16944757595215 | -1.28157554055229 | 0.2217166107314 | 0.312036084962714 |
| Kansl3 | ENSMUSG00000010453 | 226976 | 0.0977579610377868 | 5.68141392413221 | 1.27979953741716 | 0.2223222725449 | 0.312848192737975 |
| Ank3 | ENSMUSG00000069601 | 11735 | 0.0840783668685603 | 7.3566310957172 | 1.27958804465014 | 0.2223944848121 | 0.312909526770738 |
| Slc7a2 | ENSMUSG00000031596 | 11988 | 0.16448708845189 | 4.43825608052626 | 1.27891018384359 | 0.2226260603166 | 0.313195040581849 |
| Pik3ca | ENSMUSG00000027665 | 18706 | -0.110990256978075 | 6.27224751835616 | -1.27842947392497 | 0.2227904003366 | 0.31338590478447 |

| | | | | | | | |
|---|---|---|---|---|---|---|---|
| Elk3 | ENSMUSG00000008398 | 13713 | -0.171947907701701 | 3.88664505400982 | -1.27824243473523 | 0.2228543694635 | 0.313401233257133 |
| Klhl31 | ENSMUSG00000044938 | 244923 | -0.107509122061176 | 8.62710890554321 | -1.27819670049534 | 0.2228700132242 | 0.313401233257133 |
| Entpd4 | ENSMUSG00000095463 | 67464 | -0.0694605432213518 | 5.34422868035998 | -1.2781461362149 | 0.2228873101583 | 0.313401233257133 |
| Wdr31 | ENSMUSG00000028391 | 71354 | -0.227516808433382 | 3.38420191108106 | -1.27762885210931 | 0.2230643232511 | 0.313609790026833 |
| Cep131 | ENSMUSG00000039781 | 12009 | 0.139526986483473 | 2.97248483989445 | 1.27561247888667 | 0.2237553906994 | 0.314540917661422 |
| Chac1 | ENSMUSG00000027313 | 69065 | 0.616047142197376 | 0.802413515512579 | 1.27524142122513 | 0.2238827482628 | 0.31467948068159 |
| Col4a1 | ENSMUSG00000031502 | 12826 | 0.194902488169244 | 8.33613243053634 | 1.27498194497368 | 0.2239718422274 | 0.314764233400379 |
| Socs5 | ENSMUSG00000037104 | 56468 | -0.107784340686605 | 4.42782368363688 | -1.27457208398173 | 0.2241126299725 | 0.314921604091729 |
| Pus3 | ENSMUSG00000032103 | 67049 | -0.176322623542609 | 2.03853368823 | -1.27446342772851 | 0.2241499653496 | 0.314933582421166 |
| Eif2b1 | ENSMUSG00000029388 | 209354 | -0.101186263669862 | 3.62957041626587 | -1.27393209698948 | 0.2243326072884 | 0.315122045148009 |
| Rprd2 | ENSMUSG00000028106 | 75137 | -0.0719418269924468 | 5.16378690467946 | -1.27390548572829 | 0.2243417578759 | 0.315122045148009 |
| Itgb1bp2 | ENSMUSG00000031312 | 26549 | -0.101700051727939 | 6.35596433448711 | -1.27325038095488 | 0.2245671169434 | 0.315398066980422 |
| Fbxl4 | ENSMUSG00000040410 | 269514 | 0.161457922369093 | 2.25339027987894 | 1.27306262288169 | 0.2246317398231 | 0.315448297534266 |
| Crybg1 | ENSMUSG00000019866 | 11630 | 0.16499327890806 | 2.84913737418347 | 1.27296678944808 | 0.2246647296392 | 0.315454098867909 |
| Xkrx | ENSMUSG00000031258 | 331524 | -0.82791645685014 | -0.0739582189215572 | -1.27278516098443 | 0.2247272642259 | 0.315501377505035 |
| Ssbp2 | ENSMUSG00000003992 | 66970 | -0.21078225978106 | 3.73138712633355 | -1.27258623790081 | 0.2247957692455 | 0.315536646255475 |
| Dvl2 | ENSMUSG00000020888 | 13543 | 0.204919653214259 | 1.83900375148505 | 1.27254457816466 | 0.2248101181079 | 0.315536646255475 |
| Htra2 | ENSMUSG00000068329 | 64704 | 0.179958568326918 | 2.0109140801529 | 1.27049811993842 | 0.2255158779516 | 0.316486591589403 |
| Mplkip | ENSMUSG00000012429 | 66308 | 0.163189868353081 | 2.31407225242794 | 1.27038781408699 | 0.2255539690428 | 0.316499414070298 |
| Rap1a | ENSMUSG00000068798 | 109905 | -0.0850586726117895 | 5.04105103311007 | -1.26994055488626 | 0.2257084701744 | 0.316675559937959 |
| Zcchc8 | ENSMUSG00000029427 | 70650 | -0.130863414706444 | 3.38213834889648 | -1.26934713526952 | 0.2259135909018 | 0.316888348319601 |
| Cdan1 | ENSMUSG00000027284 | 68968 | 0.144815693198258 | 2.59919800212509 | 1.26926165741256 | 0.2259431492932 | 0.316888348319601 |
| Lama3 | ENSMUSG00000024421 | 16774 | 0.3157757713486 | 1.5545799166197 | 1.269250223603 | 0.2259471033576 | 0.316888348319601 |
| Tesk1 | ENSMUSG00000028458 | 21754 | 0.086336815157229 | 5.2164708729616 | 1.26869191521444 | 0.2261402457371 | 0.317118540879069 |
| Kdm5d | ENSMUSG00000056673 | 20592 | -0.130435930673574 | 3.67562211854877 | -1.26812221376397 | 0.2263374648000 | 0.317316524582828 |
| Ercc6 | ENSMUSG00000054051 | 319955 | -0.18101626685388 | 2.5889211499752 | -1.26811637175176 | 0.2263394878942 | 0.317316524582828 |
| Zbtb12 | ENSMUSG00000049823 | 193736 | 0.259963823533573 | 1.07341360787866 | 1.26787655913903 | 0.2264225476340 | 0.317392263168545 |
| Dennd5b | ENSMUSG00000030313 | 320560 | 0.126425795075907 | 3.86857034739345 | 1.26748661295537 | 0.2265576582841 | 0.317540936364215 |
| Clec16a | ENSMUSG00000068663 | 74374 | -0.0852889487772179 | 5.87785505034377 | -1.26690162063147 | 0.2267604697776 | 0.317784448094796 |
| Rpph1 | ENSMUSG00000092837 | 85029 | 0.295651884340945 | 11.2413474289899 | 1.26652063879151 | 0.2268926302881 | 0.317906136670699 |
| Ppp1r12c | ENSMUSG00000019254 | 232807 | -0.147821810682697 | 4.62837763481935 | -1.26648363865132 | 0.2269054686949 | 0.317906136670699 |
| 2410002F23Rik | ENSMUSG00000045411 | 668661 | 0.176656994517314 | 2.04524007703496 | 1.26630874232531 | 0.2269661625019 | 0.317950419365451 |
| 9330159F19Rik | ENSMUSG00000004360 | 212448 | -0.31067153735183 | 3.93625315466688 | -1.26536221515431 | 0.2272948572138 | 0.318370077422464 |
| Fpgt | ENSMUSG00000053870 | 75540 | -0.211296068244088 | 1.95317178237382 | -1.2651076768631 | 0.2273833136780 | 0.318453171276678 |
| Gabpb2 | ENSMUSG00000038766 | 213054 | 0.100543885909897 | 4.03337206078642 | 1.26491336941009 | 0.2274508572899 | 0.318493820515866 |
| Zer1 | ENSMUSG00000039686 | 227693 | -0.0938926073700221 | 4.20774222248244 | -1.26485654910325 | 0.2274706117251 | 0.318493820515866 |
| Clstn1 | ENSMUSG00000039953 | 65945 | 0.0800901354742427 | 5.35573285741896 | 1.26425812568793 | 0.2276787454158 | 0.318713475601162 |
| Ranbp2 | ENSMUSG00000003226 | 19386 | 0.0848757661246129 | 6.94120360000397 | 1.26423783659616 | 0.2276858046811 | 0.318713475601162 |
| Psen2 | ENSMUSG00000010609 | 19165 | 0.202739866541005 | 2.08668311208008 | 1.2629742767229 | 0.2281257827948 | 0.319288467571057 |
| Rbm15b | ENSMUSG00000074102 | 109095 | 0.165814136292967 | 3.25001837763741 | 1.26259623622884 | 0.2282575495830 | 0.31943199007705 |
| Csnk1a1 | ENSMUSG00000024576 | 93687 | 0.080538422237006 | 7.20335403081883 | 1.26236178427795 | 0.2283392986427 | 0.319505488154578 |
| B3gat3 | ENSMUSG00000071649 | 72727 | -0.103642019851621 | 3.52386929650375 | -1.26205085361797 | 0.2284477502632 | 0.31960925934014 |
| Prob1 | ENSMUSG00000073600 | 381148 | 0.0966014313773469 | 6.98894896211088 | 1.26198152385818 | 0.2284719378358 | 0.31960925934014 |
| Enpp4 | ENSMUSG00000023961 | 224794 | 0.0868229214913763 | 4.76421653005506 | 1.2618479810263 | 0.2285185336141 | 0.319633536908043 |
| Uqcc2 | ENSMUSG00000024208 | 67267 | -0.112662832630274 | 5.39901084720353 | -1.26156262755205 | 0.2286181244549 | 0.319731924055422 |
| Atrip | ENSMUSG00000025646 | 235610 | -0.126484700262965 | 3.00029862621798 | -1.26138194798804 | 0.2286812010005 | 0.319779225807486 |
| Dcp1a | ENSMUSG00000021962 | 75901 | -0.134725199828732 | 3.15602959197882 | -1.26084015697431 | 0.2288704269730 | 0.320002895528595 |
| Mrpl24 | ENSMUSG00000019710 | 67707 | -0.145035595456253 | 3.83840326058064 | -1.26052079981671 | 0.2289820238670 | 0.320117982208868 |
| C1qtnf4 | ENSMUSG00000040794 | 67445 | -0.350768546295068 | 1.02000081145917 | -1.26008946232245 | 0.2291328199534 | 0.320285800981335 |
| Usb1 | ENSMUSG00000031792 | 101985 | 0.235923502763536 | 1.22064001994581 | 1.26000983359004 | 0.2291606668625 | 0.320285800981335 |
| Msx1 | ENSMUSG00000048450 | 17701 | 0.231221114289376 | 1.74205219249712 | 1.25988302623417 | 0.2292050181173 | 0.320306833679449 |
| Ypel2 | ENSMUSG00000018427 | 77864 | 0.277247305985372 | 3.63982842449912 | 1.25975685894209 | 0.2292491522649 | 0.32032755763884 |
| Rpl35 | ENSMUSG00000062997 | 66489 | -0.111341652035678 | 6.93396284680077 | -1.25960949008852 | 0.2293007113929 | 0.320358649825712 |
| Sfr1 | ENSMUSG00000025066 | 67788 | 0.0903700779714782 | 5.67376297745056 | 1.25952426491029 | 0.2293305328532 | 0.320359368002466 |
| Mocs1 | ENSMUSG00000064120 | 56738 | -0.130534720716423 | 2.73731765814069 | -1.25931274203098 | 0.2294045609056 | 0.320421831748015 |
| Cep97 | ENSMUSG00000022604 | 74201 | 0.159311796743679 | 2.48117576571363 | 1.25907843201324 | 0.2294865860459 | 0.32049544807311 |
| Itfg1 | ENSMUSG00000031703 | 71927 | -0.0865014582891215 | 4.41764531180411 | -1.2586699472465 | 0.2296296403302 | 0.320643958032075 |
| Gm18853 | ENSMUSG00000098934 | 100417831 | -1.46371644752527 | 0.119368378505885 | -1.25860728437978 | 0.2296515915683 | 0.320643958032075 |
| Bhlhe40 | ENSMUSG00000030103 | 20893 | -0.173658387822952 | 6.26779957817339 | -1.25818397336881 | 0.2297999239479 | 0.320810085399066 |
| Spata2 | ENSMUSG00000047030 | 263876 | -0.122627496493603 | 2.93284896365318 | -1.2576964628628 | 0.2299708465684 | 0.321007703210333 |
| Zfat | ENSMUSG00000022335 | 380993 | -0.160636072166732 | 1.49939488804586 | -1.25680082095734 | 0.2302851238835 | 0.321405348464124 |
| Fgf9 | ENSMUSG00000021974 | 14180 | -0.217788397354217 | 1.40856452069368 | -1.2565805501703 | 0.2303624681847 | 0.321472250699222 |
| Ccdc93 | ENSMUSG00000026339 | 70829 | 0.116339225513649 | 3.75806231700343 | 1.25584861413773 | 0.2306196227933 | 0.321790030221381 |
| Zfp503 | ENSMUSG00000039081 | 218820 | 0.19894867419549 | 1.70244520877319 | 1.2555485098815 | 0.2307251255850 | 0.321896151597955 |
| Arsa | ENSMUSG00000022620 | 11883 | 0.231209759183503 | 1.87839770354708 | 1.25516977981059 | 0.2308583241482 | 0.322027481108867 |
| Etf1 | ENSMUSG00000024360 | 225363 | -0.0792159598897317 | 6.501375253813 | -1.25507265907252 | 0.2308924911163 | 0.322027481108867 |
| D11Wsu47e | ENSMUSG00000041623 | 276852 | 0.177646278871329 | 1.72379548573351 | 1.25502960564817 | 0.2309076385446 | 0.322027481108867 |
| Ints4 | ENSMUSG00000025133 | 101861 | -0.0879599028045934 | 4.34123664810276 | -1.25471185099306 | 0.2310194580786 | 0.322142326349998 |
| Creb1 | ENSMUSG00000025958 | 12912 | -0.110123408091926 | 4.55113613970391 | -1.25396475247304 | 0.2312825349223 | 0.322468034341395 |
| Utp23 | ENSMUSG00000022313 | 78581 | 0.153125943249588 | 1.92651547049336 | 1.25354114364994 | 0.2314318062339 | 0.322635004966647 |
| Mrm2 | ENSMUSG00000029557 | 68017 | -0.283902464078528 | 1.43729504847407 | -1.25336880913004 | 0.2314925553080 | 0.322678541452776 |
| D10Wsu102e | ENSMUSG00000020255 | 28109 | -0.370688243306156 | 6.67455721349197 | -1.25309747786317 | 0.2315882269820 | 0.322770739403422 |
| Myo19 | ENSMUSG00000020527 | 66196 | -0.368880487808118 | 1.22396626754 | -1.25198022316104 | 0.2319825018934 | 0.32327903215167 |
| Zc3h4 | ENSMUSG00000059273 | 330474 | 0.0900685836070263 | 4.79555278327495 | 1.25144398300906 | 0.2321719275483 | 0.323501764184991 |
| Naca | ENSMUSG00000061315 | 17938 | 0.109668322960433 | 8.71859826813407 | 1.2510603754166 | 0.2323075111543 | 0.323649427023787 |
| Hsd17b11 | ENSMUSG00000029311 | 114664 | 0.176417306784951 | 2.41033278713123 | 1.25079875103574 | 0.2324000165087 | 0.32373704351429 |
| Cerkl | ENSMUSG00000075256 | 228094 | -0.838612584536398 | -0.509106575531615 | -1.25062578084703 | 0.2324611914421 | 0.323780999446244 |
| Luc7l2 | ENSMUSG00000029823 | 192196 | -0.101655356022124 | 6.80504926999773 | -1.25032443819678 | 0.2325677987075 | 0.323888216036697 |
| Rpl4 | ENSMUSG00000032399 | 67891 | -0.0986984777268729 | 8.02692596993883 | -1.25015672259216 | 0.2326271489137 | 0.323929600608456 |
| Kif1a | ENSMUSG00000014602 | 16560 | -0.271980547223431 | 1.34065948395249 | -1.24873989157053 | 0.2331290067815 | 0.324587081025214 |
| Mcmbp | ENSMUSG00000048170 | 210711 | 0.0935999133322203 | 3.89604406733667 | 1.24781512970946 | 0.2334570294345 | 0.324978039175072 |
| Commd6 | ENSMUSG00000075486 | 66200 | 0.194323747930425 | 1.77303300080018 | 1.24778065353265 | 0.2334692655422 | 0.324978039175072 |
| Lhpp | ENSMUSG00000030946 | 76429 | -0.182574603360234 | 1.60303304866498 | -1.24727189131624 | 0.2336498917419 | 0.32518805279234 |
| Tmem175 | ENSMUSG00000013495 | 72392 | 0.154391452529593 | 1.56073369277779 | 1.24630856766636 | 0.2339922036363 | 0.325623014379183 |
| Shmt2 | ENSMUSG00000025403 | 108037 | -0.104173504389128 | 4.3545333782968 | -1.24335509066456 | 0.2350441750474 | 0.327045300081905 |
| Pfkp | ENSMUSG00000021196 | 56421 | 0.213639951487501 | 2.72209530027821 | 1.24221209413679 | 0.2354522886751 | 0.327571460800374 |
| Esrrg | ENSMUSG00000026610 | 26381 | -0.236564941790593 | 6.67288402127194 | -1.24161858851092 | 0.2356644237320 | 0.327824868390804 |
| Serinc3 | ENSMUSG00000017707 | 26943 | 0.095874008411819 | 6.22385631055215 | 1.23987984076968 | 0.2362867671872 | 0.328648766016488 |
| Cacybp | ENSMUSG00000014226 | 12301 | -0.088881704680742 | 4.93213561176017 | -1.23978099988339 | 0.2363221838163 | 0.328656207544036 |
| Ist1 | ENSMUSG00000031729 | 71955 | -0.0889364956745053 | 4.24519877242433 | -1.23941367811125 | 0.2364538390858 | 0.328797470429653 |
| Zfp142 | ENSMUSG00000026135 | 77264 | -0.139834646041573 | 4.47701659958172 | -1.23903258018231 | 0.2365904930901 | 0.328945647413929 |
| Lss | ENSMUSG00000033105 | 16987 | 0.231418343536434 | 1.20782246347122 | 1.23806903376009 | 0.2369362789366 | 0.329365400174065 |
| Mycbp | ENSMUSG00000028647 | 56309 | -0.178329763439296 | 2.36150794910812 | -1.2380234197151 | 0.2369526582168 | 0.329365400174065 |
| Snrpe | ENSMUSG00000090553 | 20643 | -0.15099733806072 | 3.1214232632421 | -1.23762622744301 | 0.2370953213663 | 0.329505286221951 |
| Gde1 | ENSMUSG00000033917 | 56209 | -0.107478758919025 | 3.89402474551848 | -1.23757539689702 | 0.2371135835167 | 0.329505286221951 |
| Vmn1r227 | ENSMUSG00000067951 | 171229 | -0.436916074783632 | 0.781361549676403 | -1.2373574247771 | 0.2371919080387 | 0.329572231698481 |
| Nid2 | ENSMUSG00000021806 | 18074 | 0.178206028237578 | 3.62720662401763 | 1.2365110785313 | 0.2374962211470 | 0.329953125744535 |
| Brd9 | ENSMUSG00000057649 | 105246 | -0.101980934081036 | 3.51108255627356 | -1.23625824730563 | 0.2375871890111 | 0.330037560437317 |
| Bcl7b | ENSMUSG00000029681 | 12054 | -0.102707275082648 | 3.81727050834543 | -1.23606813886686 | 0.2376556074853 | 0.330090653802044 |
| M6pr | ENSMUSG00000007458 | 17113 | 0.0797808416853019 | 3.98707878235765 | 1.23572938784786 | 0.2377775596545 | 0.330218079606601 |
| Gmps | ENSMUSG00000027823 | 229363 | -0.110065303461439 | 5.08012994410989 | -1.23559168823593 | 0.2378271463279 | 0.330244986853559 |
| Cul9 | ENSMUSG00000040327 | 78309 | 0.116852198860664 | 4.19665473480328 | 1.23422068085552 | 0.2383213005506 | 0.330854147852193 |
| Afap1l2 | ENSMUSG00000025083 | 226250 | -0.33993238584135 | 1.94284270091423 | -1.23420662504235 | 0.2383263708890 | 0.330854147852193 |
| Sec22b | ENSMUSG00000027879 | 20333 | 0.0979235074941602 | 3.74147889055236 | 1.23375096538249 | 0.2384907864876 | 0.331040353916979 |
| Fbxo33 | ENSMUSG00000035329 | 70611 | 0.185793542288109 | 1.80490098473484 | 1.23360616630217 | 0.2385430529955 | 0.331070862404045 |
| Nicn1 | ENSMUSG00000032606 | 66257 | -0.147692300955291 | 2.39282342307702 | -1.23340894332741 | 0.2386142568660 | 0.331127642732377 |
| Zfp260 | ENSMUSG00000049421 | 26466 | 0.0957357469793578 | 4.57217774998576 | 1.23329873856316 | 0.2386540516297 | 0.331140827413702 |
| Cnot4 | ENSMUSG00000038784 | 53621 | -0.0958850191982146 | 4.35860172648252 | -1.2322342754016 | 0.2390386963419 | 0.331632439359576 |
| Ppil3 | ENSMUSG00000026035 | 70225 | -0.170678862639294 | 1.93828016614906 | -1.23197687735107 | 0.2391317805113 | 0.331719478777803 |
| Fam199x | ENSMUSG00000042595 | 245622 | 0.159523870895607 | 3.48556664337874 | 1.23178927505305 | 0.2391996420410 | 0.331771512136843 |
| Ly6a | ENSMUSG00000075602 | 110454 | -0.179390819883168 | 4.02142211377746 | -1.2316413238407 | 0.2392531712298 | 0.331803655761657 |
| Nupl2 | ENSMUSG00000048439 | 231042 | -0.147742196974232 | 2.14503074525576 | -1.23144680859802 | 0.2393235617527 | 0.331859172081615 |
| Tmem160 | ENSMUSG00000019158 | 69094 | -0.190706754341156 | 2.37832342077006 | -1.23034228861688 | 0.2397235704543 | 0.332332348066726 |
| Cacna2d2 | ENSMUSG00000010066 | 56808 | -0.269513620874741 | 0.952754333182603 | -1.23033668044724 | 0.2397256028273 | 0.332332348066726 |
| Trip6 | ENSMUSG00000023348 | 22051 | 0.272855761685805 | 1.43107578255174 | 1.22989718669702 | 0.2398849153048 | 0.332511033375279 |
| Sgip1 | ENSMUSG00000028524 | 73094 | -0.196521389505878 | 1.27034546252701 | -1.22940175298624 | 0.2400646052389 | 0.332717915540357 |

| | | | | | | | |
|---|---|---|---|---|---|---|---|
| Zfp523 | ENSMUSG00000024220 | 224656 | -0.150753941963967 | 3.2077561517362 | -1.22889777447084 | 0.2402475027832 | 0.332929190279368 |
| Wdr4 | ENSMUSG00000024037 | 57773 | 0.197206411997247 | 1.72594212342332 | 1.22870591066244 | 0.2403171603426 | 0.332983506110455 |
| Elf1 | ENSMUSG00000036461 | 13709 | 0.124417277715335 | 3.64689712043393 | 1.22856163891642 | 0.2403695497035 | 0.333013884386523 |
| Dcp1b | ENSMUSG00000041477 | 319618 | 0.187029776682338 | 1.68894645458526 | 1.22782572904572 | 0.2406369200381 | 0.333342057145672 |
| Smyd5 | ENSMUSG00000033706 | 232187 | 0.187218654293775 | 1.93857799577267 | 1.22756250827053 | 0.2407326099357 | 0.33343235671658 |
| Slc35f1 | ENSMUSG00000038602 | 215085 | -0.570088144087301 | 0.0436392426267437 | -1.22747599506117 | 0.2407640670157 | 0.333433677505319 |
| Phactr1 | ENSMUSG00000054728 | 218194 | 0.225636753371198 | 1.38393839634212 | 1.22681124131304 | 0.2410058858964 | 0.333726290693493 |
| Eva1b | ENSMUSG00000050212 | 230752 | 0.211344143652854 | 1.97700452395348 | 1.22646252975811 | 0.2411328136519 | 0.33385975757179 |
| Zscan29 | ENSMUSG00000050619 | 99334 | 0.123991602217635 | 2.90482058440385 | 1.22594616966818 | 0.2413208602677 | 0.334077801872664 |
| Cnnm4 | ENSMUSG00000037408 | 94220 | 0.14142217160219 | 3.91727152126983 | 1.22499954757668 | 0.2416658973660 | 0.334498191376327 |
| Stx7 | ENSMUSG00000019998 | 53331 | 0.101597117451647 | 4.29572685683456 | 1.22494516802711 | 0.2416857300786 | 0.334498191376327 |
| Ildr2 | ENSMUSG00000040612 | 100039795 | 0.208950863105148 | 1.38029667954031 | 1.22423894223274 | 0.2419434129769 | 0.334812437935269 |
| Nelfa | ENSMUSG00000029111 | 24116 | 0.142311341359797 | 2.66374302347337 | 1.22381936791031 | 0.2420966064278 | 0.334982025931995 |
| Apold1 | ENSMUSG00000090698 | 381823 | -0.792777182693939 | 1.55754971031654 | -1.22297304242437 | 0.2424058451542 | 0.33536745897746 |
| Map3k7cl | ENSMUSG00000025610 | 224419 | -0.184395071850667 | 2.04109273412124 | -1.22281592984245 | 0.2424632865596 | 0.335404478028822 |
| Wdr82 | ENSMUSG00000020257 | 77305 | 0.0967873715981344 | 4.53157997153079 | 1.22232982364404 | 0.2426410778007 | 0.335607949062315 |
| Snai2 | ENSMUSG00000022676 | 20583 | 0.20915028503714 | 1.63257137523329 | 1.2215974002889 | 0.2429091514218 | 0.335936226491882 |
| Dbndd2 | ENSMUSG00000017734 | 52840 | -0.10268132291741 | 3.44282249499904 | -1.22136531778298 | 0.2429941441387 | 0.336011257378948 |
| Traf3ip1 | ENSMUSG00000034292 | 74019 | -0.182747992648301 | 2.15606029298592 | -1.22126911648775 | 0.2430293815844 | 0.336017476612681 |
| Zfp563 | ENSMUSG00000067424 | 240068 | -0.16861305067862 | 1.94854900847104 | -1.22060102327744 | 0.2432742070252 | 0.336313438344858 |
| Traf7 | ENSMUSG00000052752 | 224619 | 0.111892308247757 | 4.23254341520531 | 1.22036189722285 | 0.2433618826657 | 0.336392101595792 |
| Sepsecs | ENSMUSG00000029173 | 211006 | -0.169268079019115 | 2.54101413329499 | -1.2198087305705 | 0.2435647962354 | 0.336630014875419 |
| Ankrd52 | ENSMUSG00000014498 | 237615 | -0.129350036457335 | 6.04892018657786 | -1.21966091941026 | 0.2436190390136 | 0.33666241661931 |
| Ppme1 | ENSMUSG00000030718 | 72590 | 0.0782802631233608 | 4.55605499829558 | 1.21925311622883 | 0.2437687410263 | 0.336826710676081 |
| Ccnt2 | ENSMUSG00000026349 | 72949 | -0.122199615191606 | 4.40174672857198 | -1.21906586161555 | 0.2438375051570 | 0.336842808319025 |
| Atp6v0d1 | ENSMUSG00000013160 | 11972 | 0.121916293337076 | 4.25319681910397 | 1.21905356311334 | 0.2438420219768 | 0.336842808319025 |
| Cstf3 | ENSMUSG00000027176 | 228410 | 0.134789790211233 | 2.6114128367759 | 1.21884430396617 | 0.2439188857531 | 0.336906411444035 |
| Eogt | ENSMUSG00000035245 | 101351 | -0.295916129808129 | 2.36012714123652 | -1.21817878890249 | 0.2441634647277 | 0.3372016213441 |
| Mon1a | ENSMUSG00000032583 | 72825 | -0.159587269967405 | 2.60850604756121 | -1.21768231606009 | 0.2443460451114 | 0.337411144405431 |
| Pcdhgb4 | ENSMUSG00000103585 | 93701 | 0.272610497983282 | 1.07995928461301 | 1.217022840964 | 0.2445887355982 | 0.337703608541668 |
| Aplnr | ENSMUSG00000044338 | 23796 | 0.281529104111466 | 3.51517109230786 | 1.21637387453288 | 0.2448277429146 | 0.337990914094411 |
| Mapk1 | ENSMUSG00000063358 | 26413 | -0.069965645794948 | 5.76203279890854 | -1.21618701220058 | 0.2448965961569 | 0.338043274730567 |
| Ms4a4d | ENSMUSG00000024678 | 66607 | 0.271721640900637 | 1.13582286815442 | 1.21567657161294 | 0.2450847556125 | 0.3382555243442 |
| Ajuba | ENSMUSG00000022178 | 16475 | -0.389399095416697 | 1.07737925957987 | -1.21560200938185 | 0.2451122503275 | 0.3382555243442 |
| Slco2b1 | ENSMUSG00000030737 | 101488 | 0.177041132894074 | 3.39614132394386 | 1.21527679530587 | 0.2452322007544 | 0.338378337092519 |
| Rps5 | ENSMUSG00000012848 | 20103 | -0.0899078811666127 | 6.35951053172453 | -1.21504121349023 | 0.2453191203061 | 0.338455547654498 |
| Ermp1 | ENSMUSG00000046324 | 226090 | 0.135688792787127 | 3.12288444403178 | 1.21413201995645 | 0.2456547994820 | 0.338875661727835 |
| H19 | ENSMUSG00000000031 | 14955 | 0.205897019988781 | 9.53336840741985 | 1.21404856418929 | 0.2456856297862 | 0.338875661727835 |
| Klhl18 | ENSMUSG00000054792 | 270201 | 0.136679168284655 | 3.18540820455521 | 1.21364269133838 | 0.2458356108205 | 0.339039750426314 |
| Zzz3 | ENSMUSG00000039068 | 108946 | 0.0957049822700764 | 4.80528491175572 | 1.21311778156526 | 0.2460296853302 | 0.339264600775193 |
| 4732491K20Rik | ENSMUSG00000097350 | 224523 | -0.213925833207594 | 1.62963549112484 | -1.21302534648273 | 0.2460638736808 | 0.339268945913875 |
| Vrk1 | ENSMUSG00000021115 | 22367 | -0.175104965713804 | 1.59042267463051 | -1.21276686590684 | 0.2461594958805 | 0.33935798328542 |
| Nectin2 | ENSMUSG00000062300 | 19294 | 0.174875835974085 | 1.60388215882695 | 1.21260598760524 | 0.2462190257916 | 0.339397247279718 |
| Ap4b1 | ENSMUSG00000032952 | 67489 | -0.103275211311215 | 3.13019026616905 | -1.21127617860906 | 0.2467115267489 | 0.340033249135434 |
| Zfp719 | ENSMUSG00000030469 | 210105 | 0.191929588639529 | 1.91321495917952 | 1.21119044147251 | 0.2467433062790 | 0.340034175609686 |
| Rps4x | ENSMUSG00000031320 | 20102 | -0.0752403356678855 | 6.53157503320145 | -1.2107082257708 | 0.2469221050466 | 0.340237681868827 |
| Arel1 | ENSMUSG00000042350 | 68497 | -0.0778762395312356 | 4.24802703527999 | -1.20997791254101 | 0.2471930874704 | 0.340568142064436 |
| Mbip | ENSMUSG00000021028 | 217588 | -0.243593496747391 | 1.09503049106101 | -1.20978144187479 | 0.2472660274428 | 0.340625702076607 |
| Ak2 | ENSMUSG00000028792 | 11637 | 0.147726022712893 | 2.64882118754739 | 1.20948914067701 | 0.2473745757032 | 0.340732294230418 |
| Gtpbp6 | ENSMUSG00000033434 | 107999 | 0.153592779296824 | 2.67179610932333 | 1.20917204555177 | 0.2474923734289 | 0.340831273320602 |
| Ddx3y | ENSMUSG00000069045 | 26900 | -0.0929796310407562 | 5.8884282093181 | -1.20912784798438 | 0.2475087958667 | 0.340831273320602 |
| Cc2d2a | ENSMUSG00000039765 | 231214 | 0.147559344630454 | 3.72385601771049 | 1.20816222009328 | 0.2478678053574 | 0.341282652772709 |
| Ywhae | ENSMUSG00000020849 | 22627 | -0.0774732730190832 | 7.56371855244466 | -1.20806844635073 | 0.2479026910015 | 0.341287697145796 |
| Hsf4 | ENSMUSG00000033249 | 26386 | 0.179692332302612 | 1.66809136429375 | 1.20749560259685 | 0.2481158831153 | 0.341538183898024 |
| Zfp84 | ENSMUSG00000046185 | 74352 | -0.130689443565453 | 3.69613122726996 | -1.20734050030482 | 0.2481736312775 | 0.341568523867961 |
| Sumo3 | ENSMUSG00000020265 | 20610 | 0.094396523104054 | 3.58495444811283 | 1.20726856111292 | 0.2482004194568 | 0.341568523867961 |
| Ap2m1 | ENSMUSG00000022841 | 11773 | 0.0744849117294493 | 6.16589101573275 | 1.20662114170691 | 0.2484416021393 | 0.341857395894473 |
| Nxt2 | ENSMUSG00000042271 | 237082 | -0.295191503290489 | 1.27241069377833 | -1.20624091728095 | 0.2485833320216 | 0.34200936467322 |
| Ccdc22 | ENSMUSG00000031143 | 54638 | 0.123463718547784 | 2.76863259857511 | 1.2049784747363 | 0.2490543636625 | 0.342614302692599 |
| Cpxm2 | ENSMUSG00000030862 | 55987 | -0.258027897513191 | 4.37995421194343 | -1.2043271602274 | 0.2492976483442 | 0.342905825349337 |
| Ep300 | ENSMUSG00000055024 | 328572 | -0.0717195585461381 | 6.01279492752089 | -1.20406451386543 | 0.2493958066789 | 0.342997680272683 |
| Slc6a6 | ENSMUSG00000030096 | 21366 | -0.0873822584071213 | 6.79678633203429 | -1.20358986197736 | 0.2495732738131 | 0.343198572908781 |
| Ctbp1 | ENSMUSG00000037373 | 13016 | -0.0798563174046203 | 6.2836891115202 | -1.20344581647492 | 0.2496271502976 | 0.343229481748855 |
| Rab1b | ENSMUSG00000024870 | 76308 | -0.075662042781687 | 5.59551225510858 | -1.20327998080263 | 0.2496891880475 | 0.343271602886139 |
| Tcp1 | ENSMUSG00000068039 | 21454 | -0.100272766274156 | 7.18801040168809 | -1.20314392508184 | 0.2497400943310 | 0.343298411862782 |
| Dnajc10 | ENSMUSG00000027006 | 66861 | 0.0788739293787808 | 4.42145697244929 | 1.20294169069526 | 0.2498157768022 | 0.343359267726027 |
| Lpgat1 | ENSMUSG00000026623 | 226856 | -0.0782655585650023 | 6.11633354489886 | -1.20279138644239 | 0.2498720369525 | 0.343389252917235 |
| Tada2a | ENSMUSG00000018651 | 217031 | -0.178018879471406 | 2.06942313940517 | -1.20271556458355 | 0.2499004214579 | 0.343389252917235 |
| Tmem209 | ENSMUSG00000029782 | 72649 | 0.145976336856915 | 2.15835158317922 | 1.20225449279965 | 0.2500730813420 | 0.343583314749892 |
| A630019I02Rik | ENSMUSG00000114968 | 408254 | -0.13439129526617 | 4.45242046233571 | -1.20158349732505 | 0.2503245183286 | 0.343885548554777 |
| Daam2 | ENSMUSG00000040260 | 76441 | 0.178507628096407 | 3.29318546344693 | 1.20136255777773 | 0.2504073523628 | 0.343956115692169 |
| Zfp64 | ENSMUSG00000027551 | 22722 | 0.150912592281812 | 2.5314522366908 | 1.19951461862924 | 0.2511010121407 | 0.344858592636119 |
| Tcf20 | ENSMUSG00000041852 | 21411 | -0.0770240896570158 | 6.04529359424113 | -1.19944420688543 | 0.2511274720870 | 0.344858592636119 |
| Rab3gap1 | ENSMUSG00000036104 | 226407 | 0.0803906793005007 | 5.76042800811312 | 1.19892065175106 | 0.2513242862753 | 0.345085513537937 |
| Pbx1 | ENSMUSG00000052534 | 18514 | -0.0799892683877559 | 5.03159072945951 | -1.19857095684838 | 0.2514558099490 | 0.345222740335719 |
| Pigm | ENSMUSG00000050229 | 67556 | 0.172676344453381 | 2.63035484665328 | 1.19781862585165 | 0.2517389504223 | 0.345568060663854 |
| Slirp | ENSMUSG00000021040 | 380773 | -0.133707891258626 | 2.86858569936484 | -1.19732560409632 | 0.2519246340113 | 0.345779529680804 |
| Clk4 | ENSMUSG00000020385 | 12750 | -0.113831552227083 | 3.87782051849452 | -1.19650097930149 | 0.2522354450147 | 0.346162667853929 |
| Map2k4 | ENSMUSG00000033352 | 26398 | -0.0992738167820315 | 5.3736948555019 | -1.19531211928815 | 0.2526840649942 | 0.346734812258604 |
| Smn1 | ENSMUSG00000021645 | 20595 | -0.137856496336981 | 1.93090057903295 | -1.19488025196166 | 0.2528471848365 | 0.346915096956051 |
| Usp32 | ENSMUSG00000000804 | 237898 | -0.0822416284799088 | 4.33758152466408 | -1.19458132761582 | 0.2529601389061 | 0.347026515861279 |
| Dcun1d1 | ENSMUSG00000027708 | 114893 | 0.100128789677194 | 4.76950119299136 | 1.19367644073652 | 0.2533023059204 | 0.347452315976473 |
| Nmd3 | ENSMUSG00000027787 | 97112 | -0.0948084346586122 | 3.59189033129006 | -1.1932242795059 | 0.2534734172892 | 0.347643403311022 |
| Klc4 | ENSMUSG00000003546 | 74764 | 0.0875991877394151 | 3.59330274616346 | 1.19285173299279 | 0.2536144674636 | 0.347793218397402 |
| Vapa | ENSMUSG00000024091 | 30960 | -0.0730582668410467 | 5.50635486656 | -1.19266946453956 | 0.2536834984769 | 0.347818023619553 |
| Yes1 | ENSMUSG00000014932 | 22612 | -0.108583229138894 | 3.5083917342878 | -1.19263594628334 | 0.2536961945218 | 0.347818023619553 |
| Trmt6 | ENSMUSG00000037376 | 66926 | 0.143399338291699 | 2.49645824666361 | 1.19208112170344 | 0.2539064226117 | 0.348062591618941 |
| Hnrnpl | ENSMUSG00000015165 | 15388 | 0.0863799779670158 | 5.49538716300064 | 1.19179678975638 | 0.2540142109484 | 0.348166688385952 |
| Slc22a23 | ENSMUSG00000038267 | 73102 | 0.088447164566535 | 4.2328545151059 | 1.19148011721165 | 0.2541343011709 | 0.348287618618363 |
| Sec61g | ENSMUSG00000078974 | 20335 | -0.145373390550147 | 2.97054279770423 | -1.1910020574497 | 0.2543156769213 | 0.348492498988055 |
| Herc6 | ENSMUSG00000029798 | 67138 | -0.331304035162822 | 1.80219401660254 | -1.19006899078794 | 0.2546699713550 | 0.348934251301366 |
| Als2 | ENSMUSG00000026024 | 74018 | -0.0758854879565174 | 4.523161976321 | -1.18992323443942 | 0.2547253509945 | 0.348966388234247 |
| Lamtor4 | ENSMUSG00000050552 | 66096 | -0.120777471711043 | 2.77854825036615 | -1.18806417277456 | 0.2554325144538 | 0.349891330262536 |
| Mbd4 | ENSMUSG00000030322 | 17193 | 0.232535927819338 | 1.08071970523858 | 1.18736260931365 | 0.2556997752947 | 0.350203953663023 |
| Gcc2 | ENSMUSG00000038039 | 70297 | -0.0929864803751794 | 4.36306154354375 | -1.18729690755416 | 0.2557248154915 | 0.350203953663023 |
| Chsy1 | ENSMUSG00000032640 | 269941 | 0.154406567349159 | 2.65674535408346 | 1.18711146656633 | 0.2557955008439 | 0.350256873321512 |
| Actn2 | ENSMUSG00000052374 | 11472 | -0.105637869480715 | 11.5751331535063 | -1.18667992450444 | 0.2559600522032 | 0.350438292914977 |
| Zfp329 | ENSMUSG00000057894 | 67230 | 0.166905567525174 | 2.8698159632436 | 1.18600073737604 | 0.2562191991577 | 0.350749162929619 |
| Gramd1b | ENSMUSG00000040111 | 235283 | -0.162812978126723 | 3.90737452471001 | -1.1857609279923 | 0.2563107480359 | 0.350830551813305 |
| Eif4a3 | ENSMUSG00000025580 | 192170 | 0.0920497300689043 | 4.13508166899272 | 1.18542468931809 | 0.2564391521013 | 0.350962360288011 |
| Pnisr | ENSMUSG00000028248 | 66625 | -0.109747562164003 | 5.24038950491953 | -1.1848918960627 | 0.2566427191127 | 0.351196990813922 |
| Etv3 | ENSMUSG00000003382 | 27049 | 0.0948378845331495 | 3.94957478737377 | 1.18382946731138 | 0.2570490199497 | 0.351696116986116 |
| Zfp488 | ENSMUSG00000044519 | 382867 | -0.358807998852503 | 2.31965287388278 | -1.1837699078278 | 0.2570718117938 | 0.351696116986116 |
| Brf1 | ENSMUSG00000011158 | 72308 | 0.0856453428800941 | 4.01055458880637 | 1.18337490902084 | 0.2572230069926 | 0.351858927472996 |
| Ints7 | ENSMUSG00000037461 | 77065 | 0.164807707875286 | 2.99828355498466 | 1.18324170029518 | 0.2572740113119 | 0.351884661868207 |
| Ubr5 | ENSMUSG00000037487 | 70790 | -0.070356543265869 | 7.36485422039838 | -1.18262407595653 | 0.2575105958598 | 0.352164184079053 |
| Suco | ENSMUSG00000040297 | 226551 | 0.0993250673815331 | 3.30850511736978 | 1.18179725607743 | 0.2578275772927 | 0.352553571101727 |
| Rexo1 | ENSMUSG00000047417 | 66932 | -0.120608675060303 | 3.88709737147434 | -1.18169755808369 | 0.2578658193063 | 0.352561759954633 |
| Scamp1 | ENSMUSG00000021687 | 107767 | -0.079931035974 | 4.05706595141189 | -1.18112381573115 | 0.2580859798096 | 0.352777416796984 |
| Sdad1 | ENSMUSG00000029415 | 231452 | 0.127717811220875 | 3.56167691169777 | 1.18111829683753 | 0.2580880982641 | 0.352777416796984 |
| Dohh | ENSMUSG00000078440 | 102115 | -0.0949399950556216 | 3.59307487619006 | -1.18101135500893 | 0.2581291510649 | 0.352789416140375 |
| Rhbdd2 | ENSMUSG00000039917 | 215160 | -0.166298698746939 | 1.91846078439778 | -1.18038357686507 | 0.2583702441530 | 0.353074776701724 |
| Cdo1 | ENSMUSG00000033022 | 12583 | 0.652036503508739 | 0.112650187392668 | 1.17991129253253 | 0.2585517357755 | 0.353278627970325 |
| Rflnb | ENSMUSG00000020846 | 76566 | -0.235258455167728 | 5.07280165005104 | -1.1797665395764 | 0.2586073818325 | 0.353310497539309 |
| Zbtb45 | ENSMUSG00000049600 | 232879 | 0.152147363691822 | 1.91199468250983 | 1.17945972211959 | 0.2587253594549 | 0.353427506148671 |
| Larp7 | ENSMUSG00000027968 | 28036 | -0.121371137172578 | 3.84983433305236 | -1.17906207419674 | 0.2588783251150 | 0.353550930990132 |

| | | | | | | | |
|---|---|---|---|---|---|---|---|
| Cachd1 | ENSMUSG00000028532 | 320508 | 0.196215301306884 | 1.75296447930277 | 1.17902079324339 | 0.2588942089134 | 0.353550930990132 |
| Pih1d1 | ENSMUSG00000003423 | 68845 | -0.10540921424604 | 2.96853030801595 | -1.17897262473497 | 0.2589127438089 | 0.353550930990132 |
| Ptcd1 | ENSMUSG00000029624 | 71799 | -0.16561748230451 | 2.98682353853863 | -1.17840032925916 | 0.2591330374214 | 0.353807548345533 |
| Megf9 | ENSMUSG00000039270 | 230316 | -0.101974397663565 | 3.25613254698669 | -1.17826737672717 | 0.2591842355324 | 0.353820042897224 |
| Adam23 | ENSMUSG00000025964 | 23792 | 0.179895923255643 | 1.71575457738154 | 1.17817838382574 | 0.2592185097846 | 0.353820042897224 |
| Rab24 | ENSMUSG00000034789 | 19336 | -0.0982588121351466 | 3.76234825459049 | -1.17812442195177 | 0.2592392940777 | 0.353820042897224 |
| Naa20 | ENSMUSG00000002728 | 67877 | -0.117387623825941 | 3.87226483476686 | -1.17769551986555 | 0.2594045384498 | 0.353984207147199 |
| Gemin6 | ENSMUSG00000055760 | 67242 | -0.250164141715728 | 1.17091570438469 | -1.17764413241993 | 0.2594243420964 | 0.353984207147199 |
| Nsd3 | ENSMUSG00000054823 | 234135 | 0.110418797121099 | 4.6210149762154 | 1.17724049213927 | 0.2595799372082 | 0.354152308240524 |
| Psme3 | ENSMUSG00000078652 | 19192 | -0.0859964151562185 | 5.13999289018569 | -1.17694479956942 | 0.2596939663808 | 0.354196495233026 |
| Rbbp5 | ENSMUSG00000026439 | 213464 | -0.139147079452781 | 3.28247201673059 | -1.17692463260298 | 0.2597017448612 | 0.354196495233026 |
| Trim25 | ENSMUSG00000000275 | 217069 | 0.135323686485151 | 3.76997193262743 | 1.17690444009746 | 0.2597095333723 | 0.354196495233026 |
| Rps16 | ENSMUSG00000037563 | 20055 | 0.105897558801916 | 5.32747831951331 | 1.17666403384044 | 0.2598022750315 | 0.35427877599431 |
| Rn7s1 | ENSMUSG00000099021 | 103948 | 0.252381988450162 | 13.4982580252974 | 1.17629643726468 | 0.2599441323875 | 0.354428004381665 |
| Pter | ENSMUSG00000026730 | 19212 | -0.129206537545337 | 3.18810375472887 | -1.17572290026301 | 0.2601655825458 | 0.354657293220627 |
| Patz1 | ENSMUSG00000020453 | 56218 | -0.113728654300463 | 2.99638972714675 | -1.17569285533764 | 0.2601771872963 | 0.354657293220627 |
| Nudt5 | ENSMUSG00000025817 | 53893 | 0.22544979902486 | 1.69298976304669 | 1.17433903664689 | 0.2607005096513 | 0.355317445594223 |
| Slc4a1ap | ENSMUSG00000029141 | 20534 | -0.0828923297949808 | 3.81798968365664 | -1.17427188826684 | 0.2607264871568 | 0.355317445594223 |
| Apex2 | ENSMUSG00000025269 | 77622 | -0.23619652331012 | 1.69047735453315 | -1.17219249749113 | 0.2615319234391 | 0.356370662567073 |
| Olfr472 | ENSMUSG00000073893 | 258770 | -0.204124469475454 | 1.6646299162595 | -1.17167648180516 | 0.2617320948395 | 0.356579327094298 |
| Cdkn1b | ENSMUSG00000003031 | 12576 | -0.0845759676626381 | 4.3169843537115 | -1.17162956841784 | 0.2617502992045 | 0.356579327094298 |
| Ilvbl | ENSMUSG00000032763 | 216136 | -0.191896847776728 | 1.91840659851934 | -1.17055925532387 | 0.2621658906643 | 0.357100978299341 |
| Kdm3b | ENSMUSG00000038773 | 277250 | -0.0818984204426876 | 5.3660560145333 | -1.16858798973145 | 0.2629326420552 | 0.358056656549751 |
| Slc25a25 | ENSMUSG00000026819 | 227731 | -0.410532495446559 | 2.69040014095271 | -1.16855494938515 | 0.2629455082514 | 0.358056656549751 |
| Katna1 | ENSMUSG00000019794 | 23924 | -0.129898035201298 | 2.97300040953433 | -1.16850291818503 | 0.2629657706322 | 0.358056656549751 |
| Magi3 | ENSMUSG00000052539 | 99470 | 0.167609626242331 | 2.596844114721 | 1.16779400510788 | 0.2632419606032 | 0.358388077139632 |
| Dock1 | ENSMUSG00000058325 | 330662 | 0.0890973651013844 | 4.78800886216069 | 1.16765027690911 | 0.2632979837896 | 0.358396106069317 |
| Gm14391 | ENSMUSG00000078903 | 665001 | 0.1371412070863 | 3.91359267969245 | 1.16761064784453 | 0.2633134322424 | 0.358396106069317 |
| Arhgef28 | ENSMUSG00000021662 | 110596 | -0.143963250617474 | 2.37264745992634 | -1.16745995269312 | 0.2633721835474 | 0.358431441528517 |
| Rbm42 | ENSMUSG00000036733 | 68035 | 0.0895989816078604 | 4.37139262076943 | 1.16715141658544 | 0.2634925035384 | 0.358550548509713 |
| Nctc1 | ENSMUSG00000087090 | 330677 | -0.121716534845359 | 4.95620564371972 | -1.16683395487685 | 0.2636163483756 | 0.358674421719499 |
| Gtf2e2 | ENSMUSG00000031585 | 68153 | 0.125776347493518 | 2.19372633126435 | 1.16514590709015 | 0.2642756238283 | 0.359526676299673 |
| Eif3d | ENSMUSG00000016554 | 55944 | 0.0774519678346 | 5.48721793810158 | 1.16471563911527 | 0.2644438696363 | 0.35971079380228 |
| Lemd2 | ENSMUSG00000044857 | 224640 | -0.098590428051899 | 3.81113069782548 | -1.16441201886543 | 0.2645626424634 | 0.359827578047541 |
| Ess2 | ENSMUSG00000003527 | 27886 | -0.130018095902753 | 2.59660536211615 | -1.16385390271063 | 0.2647810781453 | 0.360062835568374 |
| Galnt7 | ENSMUSG00000031608 | 108150 | 0.147921227966952 | 2.14658832278743 | 1.16380175607992 | 0.2648014943860 | 0.360062835568374 |
| Nckap5 | ENSMUSG00000049690 | 210356 | -0.217781613792169 | 2.51045974298987 | -1.16369835241228 | 0.2648419821539 | 0.360073097876165 |
| Clec2d | ENSMUSG00000030157 | 93694 | 0.165759889266123 | 4.58850812864051 | 1.16257746389992 | 0.2652811719212 | 0.360625356332747 |
| Sept8 | ENSMUSG00000018398 | 20362 | 0.180105844106164 | 4.52292145714031 | 1.16208697717674 | 0.2654735316633 | 0.360841976450094 |
| Atg7 | ENSMUSG00000030314 | 74244 | 0.154067424467737 | 2.46414156119771 | 1.16166452775515 | 0.2656392942649 | 0.3609590920359 |
| Zfand2b | ENSMUSG00000026197 | 68818 | 0.151832590319849 | 2.42030552735896 | 1.16163550286482 | 0.2656506860980 | 0.3609590920359 |
| Plod1 | ENSMUSG00000019055 | 18822 | 0.130678744474394 | 4.70583513228315 | 1.16161493456608 | 0.2656587590731 | 0.3609590920359 |
| Faap100 | ENSMUSG00000025384 | 71885 | -0.148381864943667 | 2.45267515449081 | -1.16076634869155 | 0.2659919898158 | 0.361366945149981 |
| Etfbkmt | ENSMUSG00000039958 | 320204 | -0.165781951967027 | 1.43035251707282 | -1.16047464835483 | 0.2661066115208 | 0.361477739596679 |
| Safb2 | ENSMUSG00000042625 | 224902 | 0.140665293791037 | 4.19409012189744 | 1.16021420643159 | 0.2662089824616 | 0.361571867207852 |
| Ptpn14 | ENSMUSG00000026604 | 19250 | -0.102230111308343 | 5.12914377434008 | -1.15960694817025 | 0.2664477925665 | 0.36185126357869 |
| Rps15a | ENSMUSG00000008683 | 267019 | 0.0911070057308123 | 5.46509808110221 | 1.15951355094799 | 0.2664845364915 | 0.361856207253274 |
| Arl1 | ENSMUSG00000060904 | 104303 | -0.0869008673016083 | 5.00690099377649 | -1.15722843148538 | 0.2673847493306 | 0.363033498314975 |
| Ntng2 | ENSMUSG00000035513 | 171171 | 0.127742699175601 | 3.36634379383084 | 1.15710061577161 | 0.2674351705239 | 0.363056861524692 |
| Pmp22 | ENSMUSG00000018217 | 18858 | -0.169790263113856 | 4.60612242559092 | -1.15638118509042 | 0.2677191100113 | 0.363397192541163 |
| Sdc4 | ENSMUSG00000017009 | 20971 | -0.130748991655151 | 5.20484809215756 | -1.15613849346226 | 0.2678149457992 | 0.363482142107172 |
| Ppp2r3c | ENSMUSG00000021022 | 59032 | -0.130291611069919 | 3.61106052553897 | -1.15565355689657 | 0.2680065197031 | 0.36369699154259 |
| Tmod3 | ENSMUSG00000058587 | 50875 | -0.139471746970272 | 4.74916078839157 | -1.15531261609892 | 0.2681412709920 | 0.363790969988018 |
| Cd2ap | ENSMUSG00000061665 | 12488 | 0.083143106454135 | 4.38260086524812 | 1.15530992884847 | 0.2681423332900 | 0.363790969988018 |
| Lfng | ENSMUSG00000029570 | 16848 | 0.39316158258132 | 1.61566650423127 | 1.15452632961968 | 0.2684522357077 | 0.364166218481187 |
| Insyn1 | ENSMUSG00000066607 | 319477 | 0.149048362455785 | 2.93394523722908 | 1.15423470012821 | 0.2685676410454 | 0.364237135875352 |
| Exoc7 | ENSMUSG00000020792 | 53413 | -0.0718459295962104 | 4.55794012077943 | -1.15422581727595 | 0.2685711568159 | 0.364237135875352 |
| Ptdss2 | ENSMUSG00000025495 | 27388 | -0.10752459606362 | 2.77799501507083 | -1.15341538038911 | 0.2688920703103 | 0.364627120774339 |
| Fam193a | ENSMUSG00000037210 | 231128 | -0.0688835816146005 | 4.70377383827458 | -1.1530186808197 | 0.2690492607353 | 0.364795022215133 |
| Washc5 | ENSMUSG00000022350 | 223593 | -0.0789003172002118 | 4.79797973353735 | -1.15272681259494 | 0.2691649571028 | 0.364906628561503 |
| Smyd1 | ENSMUSG00000055027 | 12180 | 0.0945024588818972 | 7.55490425127975 | 1.15229742919032 | 0.2693352335843 | 0.365092192250868 |
| Slc4a8 | ENSMUSG00000023032 | 59033 | -0.25617821496533 | 1.30773264982254 | -1.15211881487085 | 0.2694060892265 | 0.365142958628053 |
| Zfp729a | ENSMUSG00000021510 | 212281 | -0.0888325197408948 | 3.00526205685259 | -1.15196221520753 | 0.2694682234663 | 0.365181893219775 |
| Timm22 | ENSMUSG00000020843 | 56322 | -0.118912701036156 | 3.6784515238639 | -1.15135507279367 | 0.2697092237378 | 0.365440410701638 |
| Adat1 | ENSMUSG00000031949 | 30947 | -0.203605717418312 | 2.1597305127315 | -1.15131320977235 | 0.2697258469985 | 0.365440410701638 |
| Pbrm1 | ENSMUSG00000042323 | 66923 | 0.0713486682087172 | 6.0810684792124 | 1.15042641266747 | 0.2700781666946 | 0.365839919835941 |
| Gemin7 | ENSMUSG00000044709 | 69731 | 0.111075311314198 | 3.43140433636334 | 1.15040254346801 | 0.2700876546588 | 0.365839919835941 |
| Atp6v1c1 | ENSMUSG00000022295 | 66335 | -0.0945452106138633 | 4.23584791573244 | -1.14881409302024 | 0.2707196337191 | 0.366650516191814 |
| Rbl1 | ENSMUSG00000027641 | 19650 | 0.185331996348112 | 1.51970106833481 | 1.14796489964791 | 0.2710579570526 | 0.367063246846158 |
| Ttc12 | ENSMUSG00000040219 | 235330 | 0.22467576078652 | 1.50067765038802 | 1.14742277315126 | 0.2712741103167 | 0.367277222019403 |
| Aig1 | ENSMUSG00000019806 | 66253 | 0.124952078951315 | 3.04577418132557 | 1.1474000675236 | 0.2712831662779 | 0.367277222019403 |
| Ptpn2 | ENSMUSG00000024539 | 19255 | 0.134042829642263 | 2.98878392852546 | 1.14678821850208 | 0.2715272843903 | 0.367480993326826 |
| Pip5k1c | ENSMUSG00000034902 | 18717 | 0.0854066038279511 | 4.80473842865375 | 1.14677848031567 | 0.2715311711287 | 0.367480993326826 |
| Zfp12 | ENSMUSG00000029587 | 231866 | 0.13935999500074 | 3.16675633405098 | 1.14677005660506 | 0.2715345332632 | 0.367480993326826 |
| Dctn1 | ENSMUSG00000031865 | 13191 | -0.0749759326736861 | 7.31631200558338 | -1.14662390256528 | 0.2715928724189 | 0.367514445210649 |
| Max | ENSMUSG00000059436 | 17187 | -0.106486332907575 | 3.48709443620774 | -1.14620173375924 | 0.2717614399511 | 0.367697029348497 |
| Tmem8b | ENSMUSG00000078716 | 242409 | -0.0857100320136288 | 5.10306453761119 | -1.14599365579497 | 0.2718445526537 | 0.367763961022112 |
| Cfap410 | ENSMUSG00000020284 | 67884 | 0.193435319892791 | 1.53349677502498 | 1.14550847173681 | 0.2720384253614 | 0.367980698877072 |
| AW554918 | ENSMUSG00000033632 | 225289 | -0.180692633687446 | 2.73082471662847 | -1.14526749700628 | 0.2721347546734 | 0.368065454508201 |
| Lancl1 | ENSMUSG00000026000 | 14768 | 0.0964916124908877 | 3.74778812453697 | 1.14479757284351 | 0.2723226810657 | 0.368274059969091 |
| G2e3 | ENSMUSG00000035293 | 217558 | 0.212891595525064 | 1.37491060572986 | 1.14469896560416 | 0.2723621274459 | 0.368281842542327 |
| Zbtb2 | ENSMUSG00000075327 | 381990 | -0.125855379095817 | 2.33258172886425 | -1.14456057131694 | 0.2724174974031 | 0.368311152024001 |
| Spast | ENSMUSG00000024068 | 50850 | 0.0787176641418323 | 5.02115144056869 | 1.14440889938106 | 0.2724781894560 | 0.368346322148874 |
| Dolk | ENSMUSG00000075419 | 227697 | -0.122229189360213 | 2.74205904688835 | -1.14427720445757 | 0.2725308960035 | 0.368346322148874 |
| Zfp462 | ENSMUSG00000060206 | 242466 | -0.142007267935223 | 5.53463329367436 | -1.14424295915326 | 0.2725446028304 | 0.368346322148874 |
| Tfg | ENSMUSG00000022757 | 21787 | -0.0772864708602343 | 5.09710438361198 | -1.14395666582919 | 0.2726592135102 | 0.36845566360243 |
| Atp2b4 | ENSMUSG00000026463 | 381290 | 0.148221504835604 | 4.42975859949991 | 1.14362670775943 | 0.2727913499163 | 0.368588658335646 |
| Txndc15 | ENSMUSG00000021497 | 69672 | -0.0880403407580608 | 4.01695818925567 | -1.14352319160368 | 0.2728328144962 | 0.368599121895689 |
| Rpl8 | ENSMUSG00000003970 | 26961 | -0.0927907582122334 | 7.45233493781447 | -1.14343580963646 | 0.2728678200830 | 0.368600857801309 |
| Wdfy1 | ENSMUSG00000073643 | 69368 | -0.0998389694658623 | 4.14600705756848 | -1.14324319717364 | 0.2729449935475 | 0.3686595483094 |
| Lyrm4 | ENSMUSG00000046573 | 380840 | -0.172825230668441 | 1.60016725119094 | -1.14142290985725 | 0.2736751446031 | 0.369600074827862 |
| Cir1 | ENSMUSG00000041777 | 66935 | 0.142824746934515 | 3.9997252325256 | 1.14070947235498 | 0.2739617232836 | 0.369941395579222 |
| Stx17 | ENSMUSG00000061455 | 67727 | -0.0836856551217497 | 4.14472592550035 | -1.14004939779528 | 0.2742270702913 | 0.370253965582349 |
| Neurl1a | ENSMUSG00000006435 | 18011 | -0.0838358737854135 | 5.8273694845942 | -1.13919551654131 | 0.2745706170236 | 0.370672028490215 |
| Spcs2 | ENSMUSG00000035227 | 66624 | 0.101555277784692 | 4.616639392086 | 1.13880465843068 | 0.2747279823949 | 0.370834364456394 |
| Herc2 | ENSMUSG00000030451 | 15204 | 0.0952876582346605 | 6.27766832242917 | 1.13872834492716 | 0.2747587153720 | 0.370834364456394 |
| Elp2 | ENSMUSG00000024271 | 58523 | 0.106403775339269 | 3.39066070002663 | 1.13847097136587 | 0.2748623841590 | 0.370928484104022 |
| Arhgap26 | ENSMUSG00000036452 | 71302 | 0.141417189267602 | 2.77611101722139 | 1.13832005036805 | 0.2749231882301 | 0.37096474145706 |
| Slco5a1 | ENSMUSG00000025938 | 240726 | -0.0915049611138423 | 6.30863755581606 | -1.13682223644651 | 0.2755271926448 | 0.371733861120845 |
| Set | ENSMUSG00000054766 | 56086 | -0.0755757787709852 | 6.03275575516 | -1.13667081002527 | 0.2755883125986 | 0.371770436260192 |
| Riok1 | ENSMUSG00000021428 | 71340 | 0.117803525464635 | 2.87983945265617 | 1.13597816955508 | 0.2758680132018 | 0.372101832713454 |
| Cpsf3 | ENSMUSG00000054309 | 54451 | -0.104276950605393 | 4.30568281386172 | -1.13552939381331 | 0.2760493520766 | 0.372300489518784 |
| Bpnt1 | ENSMUSG00000026617 | 23827 | -0.0976920681438354 | 3.17046358191428 | -1.13530875761682 | 0.2761385387820 | 0.372374829438263 |
| Gpatch2 | ENSMUSG00000039210 | 67769 | -0.118868448850884 | 3.28761181003883 | -1.13467172232494 | 0.2763961674111 | 0.372676268159797 |
| Zfp982 | ENSMUSG00000078496 | 195531 | -0.181147278738335 | 2.00688819695128 | -1.13434334625046 | 0.2765290399891 | 0.372809439580898 |
| Gripap1 | ENSMUSG00000031153 | 54645 | -0.0743711681921013 | 4.27108413388714 | -1.13373489103031 | 0.2767753707921 | 0.373095520795325 |
| Tbccd1 | ENSMUSG00000004462 | 70573 | -0.130453074242997 | 1.91960273727524 | -1.13362795777004 | 0.2768186795393 | 0.373107889771235 |
| Cflar | ENSMUSG00000026031 | 12633 | 0.103659149174218 | 5.18429546639037 | 1.13274701587554 | 0.2771756633396 | 0.373542988036766 |
| Nop9 | ENSMUSG00000019297 | 67842 | 0.123326070290163 | 3.02526377945543 | 1.13232551366387 | 0.2773465921676 | 0.373686889666079 |
| Ctbp2 | ENSMUSG00000030970 | 13017 | -0.13185684641373 | 3.4383847154463 | -1.13231510802691 | 0.2773508129046 | 0.373686889666079 |
| Ankrd9 | ENSMUSG00000037904 | 74251 | -0.217781378276717 | 3.80495685987337 | -1.13207795653906 | 0.2774470195724 | 0.373761091911417 |
| Rpl15 | ENSMUSG00000012405 | 66480 | -0.0830519675819623 | 6.5126702248962 | -1.13201079150961 | 0.2774742714171 | 0.373761091911417 |
| Abhd12 | ENSMUSG00000032046 | 76192 | 0.0885642489720029 | 4.08775044532221 | 1.13080854548331 | 0.2779624200212 | 0.374372500400741 |
| Asb7 | ENSMUSG00000030509 | 117589 | 0.102749893045572 | 3.02660563303968 | 1.13039915033428 | 0.2781287956346 | 0.374550433051965 |
| L3mbtl3 | ENSMUSG00000039089 | 237339 | 0.138828932651103 | 2.76373593278435 | 1.12991716744633 | 0.2783247672873 | 0.374768173345423 |
| Tacc2 | ENSMUSG00000030852 | 57752 | -0.0865101500008105 | 9.51066666974548 | -1.12966311540604 | 0.2784281056360 | 0.374861143331395 |

| | | | | | | | |
|---|---|---|---|---|---|---|---|
| 9130221H12Rik | ENSMUSG00000090330 | 77124 | 0.167514717467633 | 1.51610181822079 | 1.12933188263156 | 0.2785628817725 | 0.374979799895654 |
| Ccdc157 | ENSMUSG00000051427 | 216516 | 0.164947438543672 | 2.06222946514799 | 1.12919457784762 | 0.2786187645672 | 0.374979799895654 |
| Fam45a | ENSMUSG00000024993 | 67894 | 0.142059511516155 | 2.15295668184271 | 1.12919362953519 | 0.2786191505582 | 0.374979799895654 |
| Rps23 | ENSMUSG00000049517 | 66475 | -0.0911490563113606 | 6.73168181408008 | -1.12880793983444 | 0.2787761712107 | 0.375144937523678 |
| Sf3b5 | ENSMUSG00000078348 | 66125 | -0.13004580473148 | 2.53795968721184 | -1.12840964290569 | 0.2789383948994 | 0.375317035283784 |
| Trim27 | ENSMUSG00000021326 | 19720 | 0.0901949362392087 | 3.4577692761503 | 1.12769079060289 | 0.2792313598116 | 0.375664983889421 |
| Diaph1 | ENSMUSG00000024456 | 13367 | -0.0692820753418175 | 7.34426337765722 | -1.12699530255798 | 0.2795150246696 | 0.376000336532601 |
| Dennd6a | ENSMUSG00000040818 | 211922 | 0.110419178322475 | 3.31492369285255 | 1.1265370041943 | 0.2797020676205 | 0.376205647983294 |
| 2610008E11Rik | ENSMUSG00000060301 | 72128 | -0.117804772137374 | 3.70296678813291 | -1.12642072407466 | 0.2797495395074 | 0.376223205752489 |
| Tepsin | ENSMUSG00000025377 | 78777 | 0.129849541300852 | 2.35340091166572 | 1.12522375314177 | 0.2802385632127 | 0.376834510330801 |
| Spint2 | ENSMUSG00000074227 | 20733 | 0.31056481212735 | 0.950410835218717 | 1.12477578834199 | 0.2804217460890 | 0.37703445344394 |
| Srpk2 | ENSMUSG00000062604 | 20817 | 0.104222532116663 | 4.74407041661617 | 1.12433659945224 | 0.2806014283046 | 0.377229641224722 |
| Mchr1 | ENSMUSG00000050164 | 207911 | -0.454689096836934 | 0.459901929564171 | -1.12383942290205 | 0.2808049397682 | 0.377448227908859 |
| Lsm5 | ENSMUSG00000091625 | 66373 | -0.175560032268597 | 1.72208218051242 | -1.12369732040771 | 0.2808631277322 | 0.377448227908859 |
| Jmjd4 | ENSMUSG00000036819 | 194952 | -0.124830103228731 | 3.14711586677715 | -1.1236863659207 | 0.2808676137417 | 0.377448227908859 |
| Rps7 | ENSMUSG00000061477 | 20115 | 0.103151257055656 | 6.78254386422758 | 1.12327931984064 | 0.281034343023 | 0.377625863993167 |
| Iqcc | ENSMUSG00000040795 | 230767 | 0.143236420784365 | 2.080848433271 | 1.123126233492 | 0.2810970677758 | 0.377663722696344 |
| Ddx58 | ENSMUSG00000040296 | 230073 | 0.175508837799967 | 2.91261107755497 | 1.12240585126362 | 0.2813923754233 | 0.378014016929118 |
| Cers2 | ENSMUSG00000015714 | 76893 | 0.132203264298818 | 3.39845320797587 | 1.12212199205828 | 0.2815088028040 | 0.378106037257646 |
| Them6 | ENSMUSG00000056665 | 223626 | -0.19334822270508 | 1.6442638922372 | -1.12207018575528 | 0.2815300555520 | 0.378106037257646 |
| Gins4 | ENSMUSG00000031546 | 109145 | -0.148856398405962 | 2.60535098243047 | -1.12141617304022 | 0.2817984587007 | 0.378420018680303 |
| Zc3h7b | ENSMUSG00000022390 | 20286 | -0.0661540409713961 | 5.48646293736764 | -1.12090414536452 | 0.2820087270055 | 0.378655864746109 |
| Snrpd2 | ENSMUSG00000040824 | 107686 | -0.143544275461823 | 3.26775967200003 | -1.12011472177126 | 0.2823331425839 | 0.379044900710494 |
| Ptpdc1 | ENSMUSG00000038042 | 218232 | -0.147434607236674 | 2.01258296564841 | -1.11986511020196 | 0.2824357797528 | 0.379136130231956 |
| Ccdc59 | ENSMUSG00000019897 | 52713 | -0.102533986419755 | 3.56377631663384 | -1.119026530674 | 0.2827807997128 | 0.379552667198051 |
| Nt5c3b | ENSMUSG00000017176 | 68106 | 0.110663158800772 | 2.81122427879239 | 1.1185681162924 | 0.2829695414529 | 0.379759368646005 |
| Cdk19 | ENSMUSG00000038481 | 78334 | -0.13235532713767 | 3.2733208730737 | -1.11777047114633 | 0.2832981805976 | 0.380153745655904 |
| Mn1 | ENSMUSG00000070576 | 433938 | -0.175841172429812 | 6.34725754085915 | -1.11744156712413 | 0.2834337768080 | 0.380289016115007 |
| Spsb3 | ENSMUSG00000024160 | 79043 | -0.0974344137551628 | 3.36879111805465 | -1.11725362924902 | 0.2835112793533 | 0.380346317453582 |
| Fez2 | ENSMUSG00000056121 | 225020 | -0.0788645956562615 | 5.35640501243364 | -1.11711259335026 | 0.2835694507892 | 0.380377674141333 |
| Scarb2 | ENSMUSG00000029426 | 12492 | -0.0703522349600298 | 5.9627009287164 | -1.1164219336371 | 0.2838544492784 | 0.380713249700934 |
| Abcd2 | ENSMUSG00000055782 | 26874 | -0.150278138333229 | 4.24901883541759 | -1.1163134761189 | 0.2838992235556 | 0.380726587251482 |
| Emc3 | ENSMUSG00000030286 | 66087 | 0.0794060876305602 | 4.48284506387291 | 1.11594462438652 | 0.2840515356893 | 0.380884118820018 |
| A530064D06Rik | ENSMUSG00000043939 | 328830 | -0.419006836335175 | 0.347421634181784 | -1.1151709975893 | 0.2843711942769 | 0.38122164015028 |
| Fas | ENSMUSG00000024778 | 14102 | -0.275167898588827 | 1.0341761743557 | -1.11516663134342 | 0.2843729991570 | 0.38122164015028 |
| Ppp2r1a | ENSMUSG00000007564 | 51792 | -0.0752819918203071 | 5.70541561861655 | -1.11497641706383 | 0.2844516366405 | 0.381246906147112 |
| Pdcd2l | ENSMUSG00000002635 | 68079 | -0.12914962233401 | 2.32055175191976 | -1.11495231602246 | 0.2844616015493 | 0.381246906147112 |
| Sspn | ENSMUSG00000030255 | 16651 | 0.0829276044303233 | 6.17863940055617 | 1.11325990906326 | 0.2851620092281 | 0.38211804495728 |
| Znhit3 | ENSMUSG00000020526 | 448850 | -0.187910032434627 | 1.98499857361972 | -1.11321285045903 | 0.2851815031343 | 0.38211804495728 |
| Cog3 | ENSMUSG00000034893 | 338337 | 0.096174314514986 | 3.60299971940523 | 1.11238919340114 | 0.2855228636379 | 0.382528547913474 |
| Fam210b | ENSMUSG00000027495 | 67017 | -0.177420176510008 | 1.96449783014152 | -1.11223398816619 | 0.2855872221283 | 0.382539313778982 |
| 4933415F23Rik | ENSMUSG00000073730 | 66755 | -0.3979277037589 | 1.59257408324161 | -1.11220102801936 | 0.2856008910209 | 0.382539313778982 |
| Prpf40b | ENSMUSG00000023007 | 54614 | -0.0930219512435029 | 4.06532747105022 | -1.11191888158098 | 0.2857179200585 | 0.382649177182043 |
| Dera | ENSMUSG00000030225 | 232449 | -0.174950629201851 | 1.64196697387437 | -1.11181149851165 | 0.2857624700279 | 0.382661957598397 |
| Pacrgl | ENSMUSG00000029089 | 66768 | 0.24745748283213 | 0.94675376222603 | 1.11066583045005 | 0.2862380986656 | 0.383210737272713 |
| Naa60 | ENSMUSG00000005982 | 74763 | -0.0833647988052628 | 3.95522531061739 | -1.11065547718311 | 0.2862423995820 | 0.383210737272713 |
| Rnf181 | ENSMUSG00000055850 | 66510 | 0.0817447712153183 | 3.55301161940377 | 1.1102659643641 | 0.2864042449393 | 0.383373198530539 |
| Slc17a5 | ENSMUSG00000049624 | 235504 | 0.167871662706186 | 1.55452841979958 | 1.11019462224669 | 0.2864338955629 | 0.383373198530539 |
| Pdcl3 | ENSMUSG00000026078 | 68833 | 0.0723104609451344 | 4.21100229321213 | 1.1098448282551 | 0.2865793075106 | 0.383520863173253 |
| Mndal | ENSMUSG00000090272 | 100040462 | -0.239794229055667 | 3.04832316129709 | -1.1088114319191 | 0.2870092226096 | 0.38404918745827 |
| Orc6 | ENSMUSG00000031697 | 56452 | 0.195761198239309 | 1.42576315487058 | 1.10860044310586 | 0.2870970581279 | 0.384097004742641 |
| Dyrk1b | ENSMUSG00000002409 | 13549 | -0.09739893146293 | 5.52846684369857 | -1.10848899032416 | 0.2871434645471 | 0.384097004742641 |
| Tmem59 | ENSMUSG00000028618 | 56374 | 0.0738316989800856 | 5.03524091192811 | 1.10847240066754 | 0.2871503725877 | 0.384097004742641 |
| Slc49a4 | ENSMUSG00000022848 | 224132 | -0.077091158102486 | 4.89587687658216 | -1.10797138433384 | 0.2873590579570 | 0.384329115689297 |
| Pspc1 | ENSMUSG00000021938 | 66645 | -0.115813526517662 | 3.02809730299844 | -1.10750978472196 | 0.2875514262761 | 0.384539349232307 |
| Ttf1 | ENSMUSG00000026803 | 22130 | -0.129486349132167 | 2.88869057966328 | -1.10682415596604 | 0.2878373358887 | 0.384847680573698 |
| Sh3rf2 | ENSMUSG00000057719 | 269016 | -0.329759202845674 | 4.62435517662644 | -1.10678803411386 | 0.2878524047544 | 0.384847680573698 |
| Cep57 | ENSMUSG00000031922 | 74360 | -0.135361683328442 | 2.5042654120341 | -1.10635180963988 | 0.2880344303023 | 0.385043947368804 |
| Hectd3 | ENSMUSG00000046861 | 76608 | 0.0940926326225817 | 3.91661199743983 | 1.10549837211383 | 0.2883907986468 | 0.385441610952488 |
| Mlh1 | ENSMUSG00000032498 | 17350 | 0.14361870468352 | 2.44781558789145 | 1.10547053626465 | 0.2884024275894 | 0.385441610952488 |
| 2510039O18Rik | ENSMUSG00000044496 | 77034 | 0.0847795645482678 | 3.72824359930917 | 1.10505911584573 | 0.2885743472054 | 0.385624228521142 |
| Klf2 | ENSMUSG00000055148 | 16598 | -0.471200507705705 | 4.08350319083804 | -1.10413844761656 | 0.2889593443749 | 0.386091503894702 |
| Zfp948 | ENSMUSG00000067931 | 381066 | 0.214022870008945 | 1.41764860575793 | 1.10373518028277 | 0.2891281007240 | 0.386269771329096 |
| Snrnp200 | ENSMUSG00000003660 | 320632 | -0.0739832487097516 | 6.18673397220103 | -1.10298935461835 | 0.2894404033835 | 0.386639746961394 |
| Dnajc14 | ENSMUSG00000025354 | 74330 | -0.0720643396530516 | 4.66985899887201 | -1.10201453426392 | 0.2898489757045 | 0.387138215228035 |
| Kctd20 | ENSMUSG00000005936 | 66989 | -0.0901443701879776 | 4.48021914218563 | -1.10175547103911 | 0.2899576285465 | 0.387236021703395 |
| Tysnd1 | ENSMUSG00000020087 | 71767 | 0.176765651484427 | 2.06089706423514 | 1.1009981457561 | 0.2902754310883 | 0.387582575258424 |
| Tsg101 | ENSMUSG00000014402 | 22088 | -0.0827585871225072 | 4.77320920797253 | -1.1009681161871 | 0.2902880380240 | 0.387582575258424 |
| C9orf72 | ENSMUSG00000028300 | 73205 | -0.116395322056624 | 2.51197668638742 | -1.10057735340756 | 0.2904521244908 | 0.387754295653269 |
| Xiap | ENSMUSG00000025860 | 11798 | -0.081602894551847 | 4.75904019337572 | -1.10037154160193 | 0.2905385755609 | 0.387822343321075 |
| Actr5 | ENSMUSG00000037761 | 109275 | 0.174263832092592 | 1.25038040162671 | 1.09953919293574 | 0.2908883997502 | 0.388210573374216 |
| Iqsec2 | ENSMUSG00000041115 | 245666 | 0.085055541806458 | 4.04591378535944 | 1.09951053744726 | 0.2909004488617 | 0.388210573374216 |
| Selenow | ENSMUSG00000041571 | 20364 | -0.0947896582280832 | 7.75745060727992 | -1.0993622632821 | 0.2909628014285 | 0.388246384572141 |
| Ptpn11 | ENSMUSG00000043733 | 19247 | -0.0876513367422831 | 6.87453808653221 | -1.09642728785927 | 0.2921990862150 | 0.389848432981449 |
| Slc35e3 | ENSMUSG00000060181 | 215436 | 0.127343428974273 | 2.6938041759756 | 1.09553627280881 | 0.2925751813409 | 0.390302575938199 |
| Rpl5 | ENSMUSG00000058558 | 100503670 | 0.0971087022938972 | 4.87848996056808 | 1.09517461116944 | 0.2927279411317 | 0.390458709519307 |
| Fam207a | ENSMUSG00000032977 | 108707 | 0.121247087799439 | 2.76400270899652 | 1.0946498939975 | 0.2929496789627 | 0.390696740171195 |
| Snf8 | ENSMUSG00000006058 | 27681 | -0.0956256726541218 | 3.02453410179692 | -1.09458318664515 | 0.2929778775211 | 0.390696740171195 |
| Dll1 | ENSMUSG00000014773 | 13388 | 0.241675464802151 | 2.36847432980776 | 1.09435946047801 | 0.2930724659643 | 0.390756497025869 |
| Mtss2 | ENSMUSG00000033763 | 244654 | -0.0684913699358649 | 5.94024340193758 | -1.09430810066455 | 0.2930941834338 | 0.390756497025869 |
| Id3 | ENSMUSG00000007872 | 15903 | -0.208440736280402 | 2.75290188458709 | -1.09280507693623 | 0.2937302695239 | 0.391556777581235 |
| Ldb1 | ENSMUSG00000025223 | 16825 | -0.108868121473432 | 3.73442076495477 | -1.0920910861238 | 0.2940327949283 | 0.391912264524086 |
| Chd8 | ENSMUSG00000053754 | 67772 | 0.071951986733958 | 5.42380148391704 | 1.09183942030334 | 0.2941394838826 | 0.392006668900478 |
| Phc2 | ENSMUSG00000028796 | 54383 | 0.117382921743351 | 4.21039309607886 | 1.09159667769908 | 0.2942424174218 | 0.392096045939135 |
| Tax1bp1 | ENSMUSG00000004535 | 52440 | -0.0666623000919682 | 6.37303675540004 | -1.09044951026505 | 0.2947292311262 | 0.392696882671961 |
| Krcc1 | ENSMUSG00000053012 | 57896 | 0.0983781972037132 | 4.49930692490324 | 1.09018963344687 | 0.2948395963961 | 0.392796054626026 |
| Brca2 | ENSMUSG00000041147 | 12190 | -0.191851317891457 | 1.67905332665678 | -1.08988520018139 | 0.2949689232935 | 0.392846865850115 |
| Impa2 | ENSMUSG00000024525 | 114663 | -0.107001241023672 | 2.93916725761213 | -1.08984438845293 | 0.2949862638248 | 0.392846865850115 |
| Atraid | ENSMUSG00000013622 | 381629 | -0.108106551251654 | 3.311088975822 | -1.08980234268214 | 0.2950041294857 | 0.392846865850115 |
| Nrde2 | ENSMUSG00000021179 | 217827 | -0.128876280415001 | 3.20448354679259 | -1.0897614844283 | 0.2950214913332 | 0.392846865850115 |
| Ccdc85b | ENSMUSG00000095098 | 240514 | -0.112351278144346 | 3.34147287843467 | -1.08943648550483 | 0.2951596199034 | 0.392971875874928 |
| Exoc6b | ENSMUSG00000033769 | 75914 | 0.0885879104683251 | 4.20228384693126 | 1.08937143673647 | 0.2951872722357 | 0.392971875874928 |
| Mtmr10 | ENSMUSG00000030522 | 233315 | -0.100099229229869 | 3.84110632924382 | -1.08909565873657 | 0.2953045273774 | 0.393044713622933 |
| Wrn | ENSMUSG00000031583 | 22427 | 0.113183499271007 | 2.57318639920195 | 1.08907362004747 | 0.2953138992759 | 0.393044713622933 |
| Pja2 | ENSMUSG00000024083 | 224938 | 0.0772098228650062 | 7.0986988405932 | 1.08884870687394 | 0.2954095557487 | 0.393124160444357 |
| Acad8 | ENSMUSG00000031969 | 66948 | -0.102929313843047 | 3.43471496639727 | -1.08867633361866 | 0.2954828824408 | 0.393173875588683 |
| Ankrd16 | ENSMUSG00000047909 | 320816 | 0.21175159575602 | 1.98826223731184 | 1.08844109919492 | 0.2955829718721 | 0.393259185191578 |
| Timm17b | ENSMUSG00000031158 | 21855 | -0.135407893992983 | 3.37990958901072 | -1.08827265641661 | 0.2956546578055 | 0.393306689116773 |
| Gdf10 | ENSMUSG00000021943 | 14560 | 0.217306633629005 | 1.46901277118204 | 1.08809423180668 | 0.2957306059624 | 0.393359850787961 |
| Ezh2 | ENSMUSG00000029687 | 14056 | -0.12142038822809 | 2.28984319610927 | -1.08792561691508 | 0.2958023918995 | 0.393407463907281 |
| Zkscan3 | ENSMUSG00000021327 | 72739 | -0.183838197159824 | 3.33128645879261 | -1.08745949786446 | 0.2960009046263 | 0.393623587405148 |
| Uap1 | ENSMUSG00000026670 | 107652 | 0.182643159863924 | 4.79723226849893 | 1.08720849171161 | 0.2961078453445 | 0.393717900189972 |
| Cers4 | ENSMUSG00000008206 | 67260 | -0.23641925907546 | 1.77172505247775 | -1.08699994222777 | 0.2961967193982 | 0.393763350825983 |
| Usp3 | ENSMUSG00000032376 | 235441 | 0.116277929945871 | 3.10904644290046 | 1.0869212526133 | 0.2962302584087 | 0.393763350825983 |
| Tars | ENSMUSG00000022241 | 110960 | 0.0770358920023426 | 4.95121720575943 | 1.08687471302375 | 0.2962500958002 | 0.393763350825983 |
| Hbp1 | ENSMUSG00000002996 | 73389 | -0.0891673505741259 | 4.97103427858816 | -1.0865918439262 | 0.2963706894104 | 0.393875745403715 |
| Angptl7 | ENSMUSG00000028989 | 654812 | -0.308823714796292 | 3.67949304045476 | -1.08604102902711 | 0.2966056195482 | 0.394091087584429 |
| Grk5 | ENSMUSG00000003228 | 14773 | 0.166976399032176 | 3.23576902476467 | 1.08598571359671 | 0.2966292200124 | 0.394091087584429 |
| Topors | ENSMUSG00000036822 | 106021 | 0.094798213375392 | 4.0654929712719 | 1.08594879474629 | 0.2966449723099 | 0.394091087584429 |
| Lcmt2 | ENSMUSG00000074890 | 329504 | -0.1562575762899 | 1.63283715682575 | -1.08587389148623 | 0.2966769334674 | 0.394091087584429 |
| Kpna1 | ENSMUSG00000022905 | 16646 | -0.0712675339622786 | 6.68968639243446 | -1.08553750685011 | 0.2968205001663 | 0.394233886672948 |
| Ing3 | ENSMUSG00000029670 | 71777 | -0.125338107338985 | 2.1914016485943 | -1.08536112173618 | 0.2968958008446 | 0.394285991864025 |
| Mapk8ip1 | ENSMUSG00000027223 | 19099 | 0.213460319130237 | 1.83335088241686 | 1.08474039016202 | 0.2971609109943 | 0.394590126102935 |
| Hmgb2 | ENSMUSG00000054717 | 97165 | 0.167970137182637 | 1.85115272255342 | 1.08371247390119 | 0.2976003148461 | 0.395125597180036 |
| Smg1 | ENSMUSG00000030655 | 233789 | 0.0820988947865811 | 6.66557280324975 | 1.08347840439627 | 0.2977004402827 | 0.395210531057935 |
| Ebna1bp2 | ENSMUSG00000028729 | 69072 | 0.116447033322896 | 3.74597216892013 | 1.08337738721051 | 0.2977436590873 | 0.395219907405455 |

| | | | | | | | |
|---|---|---|---|---|---|---|---|
| Mms19 | ENSMUSG00000025159 | 72199 | 0.140644348427646 | 2.91816455737655 | 1.08216351392364 | 0.2982633635272 | 0.395861683671198 |
| Setd7 | ENSMUSG00000037111 | 73251 | 0.0859798920544179 | 6.74995930840564 | 1.08106206020985 | 0.2987355211707 | 0.396440206618616 |
| Armh3 | ENSMUSG00000039901 | 71617 | -0.118469386694374 | 3.14446066435021 | -1.08036247945688 | 0.2990356975759 | 0.396790387254474 |
| Ttc5 | ENSMUSG00000006288 | 219022 | 0.134913110772185 | 1.95385057271319 | 1.08007352745406 | 0.2991597466913 | 0.396906807996462 |
| 4932438A13Rik | ENSMUSG00000037270 | 229227 | 0.0807920962988078 | 6.01407214959727 | 1.07950970761064 | 0.2994019087102 | 0.397179886421286 |
| Zfp236 | ENSMUSG00000041258 | 329002 | -0.0685015899786823 | 4.45698832845743 | -1.07876495971666 | 0.2997220032089 | 0.397556269515437 |
| Tead1 | ENSMUSG00000055320 | 21676 | -0.0974755190061671 | 8.37224855731655 | -1.07833996128575 | 0.2999047825899 | 0.397750446310461 |
| Srp72 | ENSMUSG00000036323 | 66661 | 0.0677489628593353 | 5.2122539986989 | 1.07751472664561 | 0.3002599284743 | 0.398173150327989 |
| Zfp334 | ENSMUSG00000017667 | 228876 | 0.171197540872574 | 2.01051205060473 | 1.07732228952254 | 0.3003427901477 | 0.398234720900629 |
| Arrdc2 | ENSMUSG00000002910 | 70807 | -0.542588716497202 | 1.3241170982468 | -1.07709054800528 | 0.3004425984862 | 0.398318743972504 |
| Mcm3 | ENSMUSG00000041859 | 17215 | 0.132683649240908 | 1.91595029927628 | 1.07659359087655 | 0.3006567151445 | 0.398519629471451 |
| Asb13 | ENSMUSG00000033781 | 142688 | 0.238321917664123 | 2.11726439337755 | 1.07656964026678 | 0.3006670372565 | 0.398519629471451 |
| Washc1 | ENSMUSG00000024101 | 68767 | -0.0899829364625497 | 3.76478697417741 | -1.07528079655107 | 0.3012228848482 | 0.399207972147989 |
| Polr1b | ENSMUSG00000027395 | 20017 | 0.111574129338368 | 2.6267933668586 | 1.07476413375332 | 0.3014459233140 | 0.39945513247012 |
| Gpr108 | ENSMUSG00000005823 | 78308 | 0.112941785365719 | 3.20413952384765 | 1.07452418493065 | 0.3015495486395 | 0.399544014082307 |
| Sox18 | ENSMUSG00000046470 | 20672 | -0.318616587351786 | 3.01467330650856 | -1.07380420792178 | 0.3018606396742 | 0.399907726612398 |
| Naa50 | ENSMUSG00000022698 | 72117 | -0.0909617232145408 | 7.53269044256988 | -1.0736285127806 | 0.3019365909877 | 0.399959873495789 |
| C330007P06Rik | ENSMUSG00000006423 | 77644 | -0.117179764574716 | 3.31793509699326 | -1.07319613729942 | 0.3021235631136 | 0.400136439046366 |
| C230072F16Rik | ENSMUSG00000112121 | 320784 | -0.544560894291021 | 0.24339675292168 | -1.07315098176203 | 0.3021430946746 | 0.400136439046366 |
| Elp1 | ENSMUSG00000028431 | 230233 | -0.105314927478499 | 4.61201199435001 | -1.07300163368907 | 0.3022077003120 | 0.40017351570086 |
| Bloc1s2 | ENSMUSG00000057506 | 73689 | -0.157166413795094 | 2.0029013493148 | -1.07274143933259 | 0.3023202807914 | 0.40023736080304 |
| Vps37b | ENSMUSG00000066278 | 330192 | -0.162868497276879 | 1.65398260971102 | -1.07272095467723 | 0.3023291453801 | 0.40023736080304 |
| Cebpd | ENSMUSG00000071637 | 12609 | 0.417211218843232 | 3.195714634252 | 1.07222592564056 | 0.3025434242920 | 0.400472532203504 |
| Bst2 | ENSMUSG00000046718 | 69550 | -0.31214224501557 | 1.45338555816943 | -1.07151780748171 | 0.3028501369707 | 0.400829985134628 |
| Rbm22 | ENSMUSG00000024604 | 66810 | -0.0975191903670054 | 3.62675027575311 | -1.07138783883614 | 0.3029064563200 | 0.40085598959249 |
| Slc38a7 | ENSMUSG00000036534 | 234595 | 0.191232058403813 | 2.16052089650061 | 1.07024856275165 | 0.3034004714530 | 0.4014611491893 |
| Usp48 | ENSMUSG00000043411 | 170707 | 0.101399668494149 | 4.29944258325704 | 1.07008885155068 | 0.3034697734209 | 0.401504247550803 |
| Ttc1 | ENSMUSG00000041278 | 66827 | -0.0833126720833217 | 4.32094017982795 | -1.06976328926824 | 0.3036110778942 | 0.401642586525674 |
| Slfn2 | ENSMUSG00000072620 | 20556 | 0.243944153549804 | 2.28814576785273 | 1.06962108967364 | 0.3036728123705 | 0.401675642790683 |
| Crk | ENSMUSG00000017776 | 12928 | -0.0816050394692054 | 5.37365917508694 | -1.06942388125877 | 0.3037584437427 | 0.401740296255572 |
| Nedd9 | ENSMUSG00000021365 | 18003 | -0.173952508159818 | 4.22076933021186 | -1.06923957218539 | 0.3038384901601 | 0.401761221997775 |
| Slc38a2 | ENSMUSG00000022462 | 67760 | 0.122868705798034 | 6.68068999012197 | 1.06913955557286 | 0.3038819344489 | 0.401761221997775 |
| Tyr | ENSMUSG00000004651 | 22173 | -0.473677159117152 | 1.43988963108232 | -1.06913358341823 | 0.3038845287235 | 0.401761221997775 |
| Rnf111 | ENSMUSG00000032217 | 93836 | -0.0739524197102179 | 4.20715535901286 | -1.06837605256569 | 0.3042137294073 | 0.402147814264269 |
| Zgrf1 | ENSMUSG00000051278 | 71643 | -0.210021795846721 | 1.10179043151523 | -1.06713039616248 | 0.3047556290185 | 0.402815451124798 |
| Rbm4 | ENSMUSG00000094936 | 19653 | -0.125899824906471 | 3.50548834638361 | -1.06683387169171 | 0.3048847316588 | 0.402937371752293 |
| Sccpdh | ENSMUSG00000038936 | 109232 | 0.0862014568158473 | 4.40127601942411 | 1.06651297809062 | 0.3050244898637 | 0.403073343653328 |
| Zmiz1 | ENSMUSG00000007817 | 328365 | 0.10516101749006 | 5.87484713578453 | 1.06605775048365 | 0.3052228356083 | 0.403286693585746 |
| Rnasek | ENSMUSG00000093989 | 52898 | 0.0875832766634153 | 4.23586647849889 | 1.06592353254319 | 0.3052813334772 | 0.403315235223542 |
| Dhcr24 | ENSMUSG00000034926 | 74754 | 0.205971707758934 | 2.13692099596894 | 1.0657156742944 | 0.3053719432775 | 0.403386188757371 |
| Lzic | ENSMUSG00000028990 | 69151 | 0.128764191886826 | 2.72745386113157 | 1.06559626344437 | 0.3054240059826 | 0.40340621186516 |
| Med7 | ENSMUSG00000020397 | 66213 | 0.0957535384195958 | 2.93006148996466 | 1.06532963854675 | 0.3055402771764 | 0.403511026919853 |
| Vps26c | ENSMUSG00000022898 | 13185 | 0.13878885922882 | 3.03279729039064 | 1.06480107668751 | 0.3057708719655 | 0.403766779591172 |
| Kmt2e | ENSMUSG00000029004 | 69188 | 0.0955354639786812 | 5.59410532774629 | 1.06424190086129 | 0.3060149626723 | 0.404040289524332 |
| Rad51d | ENSMUSG00000018841 | 19364 | 0.199706912817008 | 1.30407390847729 | 1.06384703347942 | 0.3061874164011 | 0.404219160468654 |
| Map3k4 | ENSMUSG00000014426 | 26407 | -0.0621558007016079 | 5.53928603693692 | -1.06313797423253 | 0.3064972700020 | 0.404579357371442 |
| Pex14 | ENSMUSG00000028975 | 56273 | -0.0905088454912955 | 3.83227376904973 | -1.06298618650781 | 0.3065636301957 | 0.404608632862582 |
| LOC108168669 | ENSMUSG00000094484 | 108168669 | -0.27400141708796 | 2.26224886459159 | -1.06291792303664 | 0.3065934778154 | 0.404608632862582 |
| 2610021A01Rik | ENSMUSG00000091474 | 668572 | -0.125520500245415 | 2.40440570017444 | -1.06257704480828 | 0.3067425560304 | 0.404756504100504 |
| Gclc | ENSMUSG00000032350 | 14629 | 0.11143960258893 | 3.09914170041475 | 1.06177810935897 | 0.3070921687706 | 0.405168919352089 |
| Pfdn5 | ENSMUSG00000001289 | 56612 | -0.0845024488387169 | 5.43597181412223 | -1.06142471602502 | 0.3072469068805 | 0.405324153887379 |
| Gm34589 | ENSMUSG00000115816 | 102637893 | -0.537239445494711 | 0.510412546345017 | -1.06118781023932 | 0.3073506715421 | 0.405401117584986 |
| Kif24 | ENSMUSG00000028438 | 109242 | 0.191726801721153 | 1.60869585672547 | 1.06112218018523 | 0.3073794220605 | 0.405401117584986 |
| Atic | ENSMUSG00000026192 | 108147 | 0.0790849715524926 | 3.62944894401157 | 1.06007641531863 | 0.3078378074734 | 0.405956698454798 |
| Sqstm1 | ENSMUSG00000015837 | 18412 | -0.0752656254502924 | 7.67220523698118 | -1.05992262647051 | 0.3079052595625 | 0.405996669756108 |
| Chtf8 | ENSMUSG00000046691 | 214987 | 0.0942188785725886 | 3.97326169201474 | 1.059479515846 | 0.3080996697110 | 0.406204015150402 |
| Xpo1 | ENSMUSG00000020290 | 103573 | 0.0915169168524299 | 5.00101055764404 | 1.05918266032691 | 0.3082299625972 | 0.406326787403558 |
| Kdelr1 | ENSMUSG00000002778 | 68137 | 0.0854692987903025 | 3.73561516103436 | 1.0590347701724 | 0.3082948882425 | 0.406349689271359 |
| Nptn | ENSMUSG00000032336 | 20320 | 0.0858769261515908 | 4.75465826570485 | 1.05892597154361 | 0.3083426586526 | 0.406349689271359 |
| Ntpcr | ENSMUSG00000031851 | 66566 | -0.149791373878051 | 2.20666472795409 | -1.05888908073108 | 0.3083588576073 | 0.406349689271359 |
| Zfp874a | ENSMUSG00000069206 | 238692 | -0.177315085267174 | 1.63315498007573 | -1.05872037678039 | 0.3084329444180 | 0.406398326354167 |
| Pdcd10 | ENSMUSG00000027835 | 56426 | -0.14665703055735 | 2.68422430666004 | -1.05810496510728 | 0.3087033154314 | 0.406674996149872 |
| Stxbp5 | ENSMUSG00000019790 | 78808 | 0.101903128425781 | 3.56123919988448 | 1.0580730797015 | 0.3087173285199 | 0.406674996149872 |
| Ehmt1 | ENSMUSG00000036893 | 77683 | 0.0833611941663781 | 4.36453019971117 | 1.05793550510996 | 0.3087777955672 | 0.406705637226812 |
| Ubxn8 | ENSMUSG00000052906 | 108159 | 0.160871447933749 | 1.27250454579153 | 1.05742681491294 | 0.3090014518664 | 0.406951189199009 |
| Relt | ENSMUSG00000008318 | 320100 | 0.26857308626349 | 3.15345990573669 | 1.05623932263956 | 0.3095240222777 | 0.407590300869565 |
| Polr3h | ENSMUSG00000022476 | 78929 | 0.132504407050605 | 1.90689255031368 | 1.05552604758513 | 0.3098382203981 | 0.407954900887959 |
| Pycrl | ENSMUSG00000022571 | 66194 | -0.165879511523332 | 1.72616857012158 | -1.05543571613359 | 0.3098780282093 | 0.407958174919424 |
| Ednrb | ENSMUSG00000022122 | 13618 | 0.19406607441752 | 3.71889836050948 | 1.05522914055893 | 0.3099690773372 | 0.408028899852318 |
| Meox2 | ENSMUSG00000036144 | 17286 | 0.124245813777121 | 3.9383087223069 | 1.05476802203622 | 0.3101723884799 | 0.40822563908347 |
| Blvrb | ENSMUSG00000040466 | 233016 | 0.145205201893293 | 1.56129453692581 | 1.05472077320516 | 0.3101932264410 | 0.40822563908347 |
| Thap7 | ENSMUSG00000022760 | 69009 | 0.129104730227817 | 1.79297554013364 | 1.054485321809 | 0.3102970819812 | 0.408305663605735 |
| Cdc73 | ENSMUSG00000026361 | 214498 | 0.0816764138963242 | 5.43013077625081 | 1.05435836623726 | 0.3103530915862 | 0.408305663605735 |
| Jcad | ENSMUSG00000033960 | 240185 | -0.134510343712069 | 5.73419834250691 | -1.0543288994327 | 0.3103660926630 | 0.408305663605735 |
| Cavin1 | ENSMUSG00000004044 | 19285 | -0.0693282912932016 | 7.21940558359369 | -1.05418956764217 | 0.3104275728031 | 0.408337400518761 |
| St6gal1 | ENSMUSG00000022885 | 20440 | 0.157577244652369 | 2.609846346575 | 1.05371445301066 | 0.3106372844754 | 0.408564090554826 |
| Col20a1 | ENSMUSG00000016356 | 73368 | -0.25568013406138 | 1.55398327427984 | -1.05351184103367 | 0.3107267474251 | 0.408584555982928 |
| Rplp0 | ENSMUSG00000067274 | 11837 | 0.0959809544309564 | 8.7072561052929 | 1.05350990657569 | 0.3107276016728 | 0.408584555982928 |
| Mttp | ENSMUSG00000028158 | 17777 | -0.180995566670397 | 1.45125858904821 | -1.05212400395555 | 0.3113400539647 | 0.409340645885133 |
| Stx6 | ENSMUSG00000026470 | 58244 | 0.105578974730335 | 3.11568471851913 | 1.05176076523582 | 0.3115007216111 | 0.4095026323429 |
| Dedd2 | ENSMUSG00000054499 | 67379 | -0.179752275616341 | 2.63710471933636 | -1.05091882400921 | 0.3118733631634 | 0.4099432098051 |
| Pml | ENSMUSG00000036986 | 18854 | 0.165456990233907 | 2.73779046467668 | 1.05069542870785 | 0.3119722924621 | 0.410023942395509 |
| Ncor2 | ENSMUSG00000029478 | 20602 | -0.069294256099151 | 7.21903551998396 | -1.05031522150799 | 0.3121407180055 | 0.410195983231419 |
| Ruvbl1 | ENSMUSG00000030079 | 56505 | 0.0832625980826616 | 3.60281239560278 | 1.04993715790775 | 0.3123082601993 | 0.410366822002598 |
| Rps20 | ENSMUSG00000028234 | 67427 | 0.0823301199478914 | 6.32145267752089 | 1.04894392380873 | 0.3127487352921 | 0.41089620498534 |
| Pnn | ENSMUSG00000020994 | 18949 | -0.0672075847207729 | 5.03077882143709 | -1.04879064087716 | 0.3128167531444 | 0.410904327005081 |
| Sap30bp | ENSMUSG00000020755 | 57230 | 0.114195850745493 | 2.93545155245021 | 1.0487109306884 | 0.3128521280795 | 0.410904327005081 |
| Aldh1l2 | ENSMUSG00000020256 | 216188 | 0.150808988607067 | 2.10447550442817 | 1.04867586835635 | 0.3128676894761 | 0.410904327005081 |
| Tmcc3 | ENSMUSG00000020023 | 319880 | -0.0672578435844686 | 5.03227799714399 | -1.04827007259613 | 0.3130478314721 | 0.411091524005502 |
| Vac14 | ENSMUSG00000010936 | 234729 | 0.0957694403557836 | 4.24388202353575 | 1.0468877839283 | 0.3136620322835 | 0.411848609596534 |
| Pigk | ENSMUSG00000039047 | 329777 | 0.0659202138819888 | 5.33052491453911 | 1.04654237142797 | 0.3138156495099 | 0.412000824500742 |
| Golga5 | ENSMUSG00000021192 | 27277 | -0.0632560643546443 | 4.92046392558879 | -1.04614011917258 | 0.3139946149401 | 0.412186277880475 |
| Ldlrap1 | ENSMUSG00000037295 | 100017 | 0.126354667306706 | 2.93947964073106 | 1.04598917506375 | 0.3140617905601 | 0.412224955885371 |
| Pds5a | ENSMUSG00000029202 | 71521 | -0.0626312253956414 | 5.44391238784776 | -1.0456031271286 | 0.3142336438784 | 0.412401003870208 |
| Pdlim7 | ENSMUSG00000021493 | 67399 | -0.0755058040732628 | 5.96366027541126 | -1.04526237746939 | 0.3143853894679 | 0.412550623322236 |
| Camk1 | ENSMUSG00000030272 | 52163 | -0.0778878661285836 | 3.92653311429039 | -1.04417870687524 | 0.3148683360939 | 0.413134771557211 |
| Frrs1 | ENSMUSG00000033386 | 20321 | 0.202611688587332 | 1.85986531017236 | 1.04393392754758 | 0.3149774992211 | 0.413228401822619 |
| Tmem87b | ENSMUSG00000014353 | 72477 | 0.105451377706427 | 3.61790641914916 | 1.04326076571358 | 0.3152778491239 | 0.413572803165006 |
| Homez | ENSMUSG00000057156 | 239099 | 0.180244284097905 | 1.3101588330249 | 1.0428905097315 | 0.3154431385392 | 0.413739974486781 |
| Nkiras2 | ENSMUSG00000017837 | 71966 | 0.0818940060780156 | 3.74001690535763 | 1.04229747098505 | 0.3157080146855 | 0.414037709481446 |
| Aven | ENSMUSG00000003604 | 74268 | 0.0984019781784267 | 2.65581013647001 | 1.04204242397177 | 0.3158219795134 | 0.414137482972797 |
| Zcchc17 | ENSMUSG00000028772 | 619605 | -0.0873402171688342 | 4.50591510229189 | -1.04186702654469 | 0.3159003713138 | 0.414190591199674 |
| Tsn | ENSMUSG00000026374 | 22099 | -0.0791076848908379 | 4.2871723737831 | -1.0414355192753 | 0.3160932889917 | 0.414360431653654 |
| Ikzf2 | ENSMUSG00000025997 | 22779 | -0.111185974395969 | 2.63463278213799 | -1.04140771800227 | 0.3161057213027 | 0.414360431653654 |
| Evc2 | ENSMUSG00000050248 | 68525 | 0.0796386514883629 | 4.52898637632674 | 1.04003961686405 | 0.3167179576835 | 0.415113188901536 |
| Elp5 | ENSMUSG00000018565 | 54351 | -0.112329570391901 | 3.41951964746549 | -1.03983813336125 | 0.3168081964140 | 0.415181680254388 |
| Rims4 | ENSMUSG00000035226 | 241770 | -0.208314208045385 | 1.32863867188787 | -1.0394869192851 | 0.3169655401625 | 0.415338086731768 |
| Gnl3 | ENSMUSG00000042354 | 30877 | 0.110164615812686 | 3.84880506303085 | 1.03875960079176 | 0.3172915601954 | 0.415715455411338 |
| Dnajc16 | ENSMUSG00000040697 | 214063 | -0.0804979008364768 | 3.87462864397434 | -1.03799533683648 | 0.3176344050558 | 0.416114774887941 |
| 2700049A03Rik | ENSMUSG00000034601 | 76967 | -0.146798054275506 | 2.26499627316327 | -1.0378732034375 | 0.3176892185460 | 0.416136710356789 |
| Ier5 | ENSMUSG00000056708 | 15939 | 0.148156439805391 | 3.75737337021095 | 1.03766510712974 | 0.3177826281250 | 0.416209190993853 |
| Top3a | ENSMUSG00000002814 | 21975 | 0.118942252288726 | 2.37381337962804 | 1.03733409116128 | 0.3179312548178 | 0.416353965066997 |
| Syf2 | ENSMUSG00000028821 | 68592 | -0.0848621374642411 | 4.71707979123032 | -1.03676956517391 | 0.3181848450168 | 0.41661119310614 |
| Atp1a1 | ENSMUSG00000033161 | 11928 | -0.137255847667547 | 7.50667792440721 | -1.03672715534604 | 0.3182039018610 | 0.41661119310614 |
| Fgfr1op | ENSMUSG00000069135 | 75296 | -0.0883617405517736 | 3.42567045966948 | -1.03629641309743 | 0.3183975029976 | 0.416814743145749 |
| Ago1 | ENSMUSG00000041530 | 236511 | 0.105069323736005 | 3.65902033254004 | 1.03581893289796 | 0.3186122114440 | 0.4170458727452 |

| | | | | | | | |
|---|---|---|---|---|---|---|---|
| Zfp788 | ENSMUSG00000074165 | 67607 | -0.213379531200546 | 1.57106625861386 | -1.03536296304042 | 0.3188173459863 | 0.417264416783674 |
| Tsc22d3 | ENSMUSG00000031431 | 14605 | 0.177470925269394 | 5.1963750127802 | 1.0352123530402 | 0.3188851245212 | 0.41730316007924 |
| Dnajb14 | ENSMUSG00000074212 | 70604 | 0.0988827375708219 | 4.44110699679585 | 1.03469048287853 | 0.3191200614268 | 0.41753286057156 |
| Dgkq | ENSMUSG00000004815 | 110524 | -0.176144807162439 | 1.8823729503994 | -1.03458785243538 | 0.3191662787323 | 0.41753286057156 |
| Dop1a | ENSMUSG00000034973 | 320615 | -0.0794725306368949 | 3.86005596006159 | -1.03456794723511 | 0.3191752431557 | 0.41753286057156 |
| Gcn1 | ENSMUSG00000041638 | 231659 | -0.062370528694589 | 5.74048250120868 | -1.0342822920722 | 0.3193039098666 | 0.417651195255693 |
| Abtb1 | ENSMUSG00000030083 | 80283 | 0.0957101193399428 | 3.23284498212016 | 1.03393756842815 | 0.3194592329799 | 0.417804364166527 |
| Lpar6 | ENSMUSG00000033446 | 67168 | -0.263089813044961 | 1.37665408737861 | -1.0329243826977 | 0.3199160659720 | 0.418351778578784 |
| Haus7 | ENSMUSG00000031371 | 73738 | 0.156248524343187 | 1.93168675128335 | 1.03194278521637 | 0.3203591103071 | 0.418833033582568 |
| Kbtbd7 | ENSMUSG00000043881 | 211255 | -0.0969870495219678 | 2.71630775543475 | -1.03193922802915 | 0.3203607166575 | 0.418833033582568 |
| Zdhhc7 | ENSMUSG00000031823 | 102193 | 0.0855401737963493 | 3.91696987029655 | 1.03070776321874 | 0.3209171728860 | 0.419510358385281 |
| Parp12 | ENSMUSG00000038507 | 243771 | -0.137268952527652 | 3.34218954218293 | -1.03046388552457 | 0.3210274562296 | 0.419604343422988 |
| Rpp25l | ENSMUSG00000036114 | 69961 | -0.17199571541527 | 1.78691039673159 | -1.0301506182393 | 0.3211691585694 | 0.419693790589885 |
| Neat1 | ENSMUSG00000092274 | 66961 | 0.242097311062595 | 7.92655652213845 | 1.03014283592167 | 0.3211726793783 | 0.419693790589885 |
| Dnaja1 | ENSMUSG00000028410 | 15502 | 0.0967398989103563 | 4.94067655742127 | 1.02951032504906 | 0.3214589284562 | 0.420017636499566 |
| St3gal5 | ENSMUSG00000056091 | 20454 | -0.110022265056763 | 7.55588666022284 | -1.02879271804728 | 0.3217839134741 | 0.420392011256858 |
| Tmem219 | ENSMUSG00000060538 | 68742 | 0.164495260534899 | 1.75916383510469 | 1.02860974453226 | 0.3218668155300 | 0.420450066988451 |
| Vps13c | ENSMUSG00000035284 | 320528 | 0.0835220405123787 | 5.9185359449742 | 1.02727514742555 | 0.3224719679555 | 0.421190235598273 |
| Sdc3 | ENSMUSG00000025743 | 20970 | 0.155665970353475 | 4.86702718466429 | 1.02698735650128 | 0.3226025706953 | 0.421310477929598 |
| Mark3 | ENSMUSG00000007411 | 17169 | -0.0673781283306392 | 5.22183812290324 | -1.02658420061129 | 0.3227855920235 | 0.421499140653407 |
| Commd9 | ENSMUSG00000027163 | 76501 | -0.0991552751705886 | 2.76326463873184 | -1.02645230195561 | 0.3228454866553 | 0.421526996491784 |
| Eif2ak4 | ENSMUSG00000005102 | 27103 | 0.148789319066473 | 1.98939032989571 | 1.02613094463731 | 0.3229914475016 | 0.421667205618152 |
| Slc23a2 | ENSMUSG00000027340 | 54338 | 0.0839041534559404 | 3.60947285733727 | 1.02599032737907 | 0.3230553310997 | 0.421700241730515 |
| Rer1 | ENSMUSG00000029048 | 67830 | -0.072372774887957 | 4.56282416410465 | -1.02551008893473 | 0.3232735766886 | 0.421934742302492 |
| Egr3 | ENSMUSG00000033730 | 13655 | 0.444127599566357 | 1.23502521254157 | 1.02481504649626 | 0.3235896302199 | 0.422296829982625 |
| Cwf19l1 | ENSMUSG00000025200 | 72502 | -0.240398673172717 | 1.2876368044946 | -1.02471260624684 | 0.3236362313902 | 0.422307227566782 |
| Armc1 | ENSMUSG00000027599 | 74252 | -0.0872235446454592 | 4.43040586061754 | -1.02454036617874 | 0.3237145962376 | 0.422359065585322 |
| Gimap8 | ENSMUSG00000064262 | 243374 | -0.19258537872939 | 1.8547240035179 | -1.02383016610146 | 0.3240378647805 | 0.422664733802537 |
| Cog4 | ENSMUSG00000031753 | 102339 | -0.0833484999320335 | 5.10153216250446 | -1.02374741002304 | 0.3240755488987 | 0.422664733802537 |
| Cbll1 | ENSMUSG00000020659 | 104836 | -0.112883570652169 | 2.70244920795556 | -1.02367974236855 | 0.3241063646604 | 0.422664733802537 |
| Setd2 | ENSMUSG00000044791 | 235626 | 0.0602292429043398 | 6.08672141818479 | 1.02361766301206 | 0.3241346373897 | 0.422664733802537 |
| Ccdc97 | ENSMUSG00000002608 | 52132 | -0.0732919316408589 | 4.6855485036279 | -1.02351157619579 | 0.3241829565303 | 0.422664733802537 |
| Selenos | ENSMUSG00000075701 | 109815 | -0.189212971089384 | 4.71493462927644 | -1.02350575957829 | 0.3241856059641 | 0.422664733802537 |
| Xirp1 | ENSMUSG00000079243 | 22437 | 0.189256563932402 | 9.39457763159996 | 1.02335140966859 | 0.3242559171516 | 0.422664733802537 |
| Ttc37 | ENSMUSG00000033991 | 218343 | 0.0829296252753675 | 3.61025219144801 | 1.02331112351077 | 0.3242742705721 | 0.422664733802537 |
| Cystm1 | ENSMUSG00000046727 | 66060 | -0.0973229903204276 | 5.36321840385959 | -1.02326151223303 | 0.3242968733328 | 0.422664733802537 |
| Tprkb | ENSMUSG00000054226 | 69786 | 0.12934156341068 | 2.55132984615033 | 1.02302661953669 | 0.3244039053220 | 0.422731748584875 |
| Smarcad1 | ENSMUSG00000029920 | 13990 | 0.10524575257858 | 3.28410577702461 | 1.02297893384095 | 0.3244256370760 | 0.422731748584875 |
| Dph5 | ENSMUSG00000033554 | 69740 | -0.159578632497624 | 3.00296625673318 | -1.02251010366744 | 0.3246393528467 | 0.422959805240462 |
| Polr2f | ENSMUSG00000033020 | 69833 | 0.100731606374767 | 2.97343903573793 | 1.0213892750837 | 0.3251506954969 | 0.423575527645962 |
| Samd8 | ENSMUSG00000021770 | 67630 | 0.0820703595332004 | 4.94532091450143 | 1.02070750555276 | 0.3254620169490 | 0.423930565689938 |
| Rars2 | ENSMUSG00000028292 | 109093 | -0.139828298495356 | 2.46132281771795 | -1.02042215976094 | 0.3255923806502 | 0.424049840687183 |
| Ssh2 | ENSMUSG00000037926 | 237860 | 0.0726925303454067 | 7.5759028867598 | 1.01997793756931 | 0.3257954043102 | 0.424263707947925 |
| Exosc3 | ENSMUSG00000028322 | 66362 | -0.128294698646561 | 2.2573903376657 | -1.01885766536515 | 0.3263078117913 | 0.424880368158629 |
| Ipw | ENSMUSG00000100826 | 16353 | -0.224224871497603 | 3.95152220002301 | -1.01864034949649 | 0.3264072786825 | 0.424959261943673 |
| Rc3h2 | ENSMUSG00000075376 | 319817 | 0.0681978177268121 | 6.05768563380753 | 1.01836810507912 | 0.3265319177216 | 0.425070905396567 |
| Ift74 | ENSMUSG00000028576 | 67694 | -0.158619288054601 | 2.20004648092736 | -1.01672098332955 | 0.3272867388499 | 0.426002779515196 |
| Klf3 | ENSMUSG00000029178 | 16599 | 0.162516670541587 | 4.44365718533192 | 1.01629455276447 | 0.3274823634681 | 0.426206657348509 |
| Sphk2 | ENSMUSG00000057342 | 56632 | -0.116947916216428 | 3.19026524851782 | -1.01577785676958 | 0.3277195106311 | 0.426411725770362 |
| Hbs1l | ENSMUSG00000019977 | 56422 | -0.0620738700154631 | 5.27127533565241 | -1.01574787027579 | 0.3277332772997 | 0.426411725770362 |
| Hist1h4c | ENSMUSG00000060678 | 319155 | 0.149961747643517 | 3.22092070575079 | 1.01569628882001 | 0.3277569591000 | 0.426411725770362 |
| Ube2j1 | ENSMUSG00000028277 | 56228 | 0.0737501667443965 | 4.22126432355551 | 1.01553903629175 | 0.3278291636746 | 0.426454907548224 |
| Psma3 | ENSMUSG00000060073 | 19167 | -0.0782259359459694 | 5.4462884243136 | -1.01517404785374 | 0.3279967972836 | 0.426622202654449 |
| Tmem41b | ENSMUSG00000047554 | 233724 | 0.0994169206627507 | 3.64671922111148 | 1.01498039237416 | 0.3280857654562 | 0.426687150767643 |
| Chmp4b | ENSMUSG00000038467 | 75608 | -0.0764561221607822 | 6.02978226354341 | -1.01260343573071 | 0.3291791961857 | 0.428058267131401 |
| Irf2 | ENSMUSG00000031627 | 16363 | -0.0953333275839192 | 3.79646542622852 | -1.01245936059353 | 0.3292455571459 | 0.428093634490627 |
| Ifit3 | ENSMUSG00000074896 | 15959 | -0.429422525765221 | 3.40305739221432 | -1.01230039504743 | 0.3293187878373 | 0.428121842149991 |
| Rabepk | ENSMUSG00000070953 | 227746 | -0.179080806894836 | 1.34107987983293 | -1.01224223525176 | 0.3293455832622 | 0.428121842149991 |
| 2900097C17Rik | ENSMUSG00000102869 | 347740 | -0.0619251637740215 | 7.6687552440165 | -1.01188899252747 | 0.3295083633467 | 0.428282511265549 |
| Tgds | ENSMUSG00000022130 | 76355 | 0.159520809626502 | 2.09475702136127 | 1.01135195604802 | 0.3297559499376 | 0.428553357361644 |
| Mir6236 | ENSMUSG00000098973 | 102465959 | 0.104870239166322 | 5.80775304366606 | 1.01056998988222 | 0.3301166950801 | 0.428922591695656 |
| Dnajc12 | ENSMUSG00000036764 | 30045 | -0.198552241896275 | 1.48854923906778 | -1.0105245354097 | 0.3301376733893 | 0.428922591695656 |
| F830016B08Rik | ENSMUSG00000090942 | 240328 | 0.45871121095969 | 5.47795813219943 | 1.01048097295352 | 0.3301577793913 | 0.428922591695656 |
| Bad | ENSMUSG00000024959 | 12015 | 0.106149613263029 | 2.44469632909289 | 1.00928242757725 | 0.3307113078213 | 0.429590648341664 |
| Mfap1a | ENSMUSG00000068479 | 67532 | -0.106340934760105 | 4.62635310238657 | -1.0091891819098 | 0.3307543998481 | 0.429595573281922 |
| Ppm1d | ENSMUSG00000020525 | 53892 | -0.140810354396729 | 2.33969087071199 | -1.00885852039176 | 0.3309072425435 | 0.429709239929323 |
| Zfp935 | ENSMUSG00000113450 | 71508 | 0.191622132726958 | 1.52025319140627 | 1.00882976585768 | 0.3309205362478 | 0.429709239929323 |
| Drc3 | ENSMUSG00000056598 | 74665 | -0.281355516965322 | 1.18914645924278 | -1.00870846500228 | 0.3309766199109 | 0.429731017014703 |
| Fos | ENSMUSG00000021250 | 14281 | 0.437346685914309 | 3.09258444029291 | 1.00692036813761 | 0.3318041449337 | 0.430754288393163 |
| Txndc16 | ENSMUSG00000021830 | 70561 | -0.110627392245204 | 4.19005405047151 | -1.00682981884018 | 0.3318460904638 | 0.430757583999616 |
| Mex3c | ENSMUSG00000037253 | 240396 | 0.0756079728862509 | 3.82203344052036 | 1.00639020505435 | 0.3320497889013 | 0.430970819577339 |
| Alg6 | ENSMUSG00000073792 | 320438 | -0.181728375829064 | 1.88772770130141 | -1.00573944687509 | 0.3323514880552 | 0.431311185555208 |
| Ifitm3 | ENSMUSG00000025492 | 66141 | 0.143487212365363 | 4.20059620615901 | 1.00491151365356 | 0.3327356129315 | 0.431758426514122 |
| Poc5 | ENSMUSG00000021671 | 67463 | -0.134829467270279 | 2.67118343607501 | -1.00480457750846 | 0.3327852499332 | 0.431771580655223 |
| Lmnb1 | ENSMUSG00000024590 | 16906 | 0.104425080799495 | 2.56999620415579 | 1.00376983110767 | 0.3332658278731 | 0.432343788806299 |
| Wnt11 | ENSMUSG00000015957 | 22411 | -0.185366706242579 | 1.71383584881594 | -1.00333886041371 | 0.3334661352732 | 0.432552310985092 |
| Ssh1 | ENSMUSG00000042121 | 231637 | 0.0981656170498852 | 3.56611236103731 | 1.00310555558122 | 0.3335746072750 | 0.432641674433248 |
| Ppp1r15b | ENSMUSG00000046062 | 108954 | 0.0878171196994533 | 4.59713188204536 | 1.00193805493716 | 0.3341178025526 | 0.433294779891242 |
| Mapk8 | ENSMUSG00000021936 | 26419 | -0.0672227225648028 | 4.67991253755684 | -1.00184165528686 | 0.3341626821844 | 0.433301575202679 |
| Cyp20a1 | ENSMUSG00000049439 | 77951 | 0.151150296382764 | 2.63273119765532 | 1.00175336823783 | 0.3342037887313 | 0.433303477004197 |
| Tmem161a | ENSMUSG00000002342 | 234371 | -0.0916464697439823 | 4.04682858858095 | -1.00141049854424 | 0.3343634637056 | 0.433459087021677 |
| Cnot6 | ENSMUSG00000020362 | 104625 | 0.0749646942434365 | 3.82491963528493 | 1.00096797952469 | 0.3345696265730 | 0.433674918542585 |
| Togaram1 | ENSMUSG00000035614 | 328108 | 0.0826582255930339 | 3.71978856554126 | 1.00048382576224 | 0.3347952910531 | 0.43391597421173 |
| Ywhab | ENSMUSG00000018326 | 54401 | 0.0639481120809897 | 5.16700722613261 | 0.999787737311396 | 0.3351199300629 | 0.434285234797591 |
| Mrpl52 | ENSMUSG00000010406 | 68836 | 0.0908587363659112 | 4.37911741680274 | 0.998885209908528 | 0.3355411837651 | 0.434779596934146 |
| Olfr613 | ENSMUSG00000078624 | 259104 | -0.475020220248428 | 0.311344903268047 | -0.9978934614187 | 0.3360045195205 | 0.435328363503508 |
| Rnf24 | ENSMUSG00000048911 | 51902 | 0.237186049270932 | 1.09464137915168 | 0.997314704462351 | 0.3362751214773 | 0.435627323165304 |
| Vps35l | ENSMUSG00000030982 | 71517 | 0.0759746477796805 | 3.82958428163174 | 0.996434087865899 | 0.3366871599047 | 0.436063408300217 |
| Frs2 | ENSMUSG00000020170 | 327826 | 0.106814247832808 | 3.72839099328549 | 0.996424742066817 | 0.3366915347227 | 0.436063408300217 |
| Lcmt1 | ENSMUSG00000030763 | 30949 | -0.141785142081596 | 2.63703656311942 | -0.995853882860074 | 0.3369588342219 | 0.436357897983679 |
| Yeats4 | ENSMUSG00000020171 | 64050 | 0.0980219469703671 | 3.16384335644533 | 0.995617762476299 | 0.3370694398522 | 0.436449425139224 |
| Exosc8 | ENSMUSG00000027752 | 69639 | 0.129395510355468 | 1.9190724066372 | 0.994964848118893 | 0.3373754193159 | 0.436793877596003 |
| Gpat4 | ENSMUSG00000031545 | 102247 | -0.0785544985910622 | 5.13826427211005 | -0.994857737409845 | 0.3374256342959 | 0.436807154013429 |
| Tmem199 | ENSMUSG00000051232 | 195040 | -0.14455774467218 | 1.62114662303167 | -0.994071114085456 | 0.3377945782551 | 0.437210582271592 |
| Mapre3 | ENSMUSG00000029166 | 100732 | 0.0683597397782902 | 4.86883558129514 | 0.994022753337389 | 0.3378172699538 | 0.437210582271592 |
| Thbd | ENSMUSG00000074743 | 21824 | -0.221665358091997 | 4.08596409827878 | -0.993905348285592 | 0.3378723629756 | 0.437218393767738 |
| Dph1 | ENSMUSG00000078789 | 116905 | 0.155803030921545 | 1.26095372991581 | 0.993839422917056 | 0.3379033016695 | 0.437218393767738 |
| Map3k7 | ENSMUSG00000028284 | 26409 | 0.0654953036971841 | 5.92990524558294 | 0.993709560984874 | 0.3379642516482 | 0.437245500623409 |
| Mboat1 | ENSMUSG00000038732 | 218121 | 0.161658225018831 | 1.43435963071253 | 0.993537327399414 | 0.3380451006725 | 0.437298342657063 |
| Reps1 | ENSMUSG00000019854 | 19707 | 0.0734937920307512 | 4.09849033033059 | 0.993368769551119 | 0.3381242376591 | 0.437348957739032 |
| Dhx15 | ENSMUSG00000029169 | 13204 | -0.0509844159426427 | 5.06648322970596 | -0.992336961043597 | 0.3386089552199 | 0.437924099563379 |
| Upf3a | ENSMUSG00000038398 | 67031 | -0.0807914026654819 | 3.30960348703585 | -0.991469738080308 | 0.3390167391210 | 0.438361772105948 |
| Hif1an | ENSMUSG00000036450 | 319594 | -0.0723755433210833 | 6.45305086285359 | -0.991446704075222 | 0.3390275749138 | 0.438361772105948 |
| Abcg2 | ENSMUSG00000029802 | 26357 | 0.0911849905832902 | 3.16119530533125 | 0.99120819558166 | 0.3391397900539 | 0.438455002374885 |
| Mecr | ENSMUSG00000028910 | 26922 | -0.107554581843043 | 2.54514115136414 | -0.991017735927182 | 0.3392294179170 | 0.438519012210413 |
| Pcsk7 | ENSMUSG00000035382 | 18554 | -0.0878562984143159 | 3.51449662940401 | -0.990319936670423 | 0.3395579379974 | 0.438891784350267 |
| Tmem259 | ENSMUSG00000013858 | 216157 | 0.0737196562217965 | 5.42251168708461 | 0.988649808343049 | 0.3403451488096 | 0.439857273780888 |
| Stard3 | ENSMUSG00000018167 | 59045 | -0.186678794272922 | 2.23168336246844 | -0.988163162328438 | 0.3405747733195 | 0.440102003446756 |
| Etv4 | ENSMUSG00000017724 | 18612 | 0.157576564873474 | 1.41373056719634 | 0.987873887714867 | 0.3407113203209 | 0.44022641163454 |
| Zfp174 | ENSMUSG00000054939 | 385674 | -0.193364636145401 | 0.992066188735998 | -0.987363623077182 | 0.3409522770777 | 0.440451127652857 |
| Sema4d | ENSMUSG00000021451 | 20354 | -0.100347062744098 | 4.71650717411274 | -0.987334939055198 | 0.3409658258346 | 0.440451127652857 |
| Xaf1 | ENSMUSG00000040483 | 327959 | -0.218811796158122 | 1.60187453763875 | -0.98695808250572 | 0.3411438678751 | 0.440629046407073 |
| Cpsf7 | ENSMUSG00000034820 | 269061 | 0.102015627533683 | 3.91636039097478 | 0.986576497347118 | 0.3413242115058 | 0.440809895554223 |
| Eef1b2 | ENSMUSG00000025967 | 55949 | -0.0720064172034807 | 6.28107864757474 | -0.985653071480073 | 0.3417609197917 | 0.441321750058368 |
| Gemin4 | ENSMUSG00000049396 | 276919 | 0.137501967954816 | 1.70266043369781 | 0.985190821950958 | 0.3419796773508 | 0.441552073366677 |
| Phf6 | ENSMUSG00000025626 | 70998 | -0.126561985921912 | 2.08760162578728 | -0.984660813356637 | 0.3422306244672 | 0.441823899380151 |

| | | | | | | | |
|---|---|---|---|---|---|---|---|
| Cdk8 | ENSMUSG00000029635 | 264064 | 0.116875908145635 | 3.68079684130009 | 0.984291511049887 | 0.3424055583836 | 0.441997538815791 |
| Oaz1 | ENSMUSG00000035242 | 18245 | -0.0873341935145007 | 6.14439275326654 | -0.983901264015971 | 0.3425904828203 | 0.442182508437006 |
| Cpne1 | ENSMUSG00000074643 | 266692 | 0.200663829349376 | 2.14270136941817 | 0.983818408105207 | 0.3426297545020 | 0.442182508437006 |
| Fzd4 | ENSMUSG00000049791 | 14366 | -0.133639764745785 | 5.67324420371935 | -0.983539733621527 | 0.3427618629568 | 0.442300781959824 |
| Ints3 | ENSMUSG00000027933 | 229543 | 0.0810744027836257 | 3.96416994502782 | 0.983423965151283 | 0.3428167548411 | 0.442319398863177 |
| Adarb1 | ENSMUSG00000020262 | 110532 | 0.113866604362827 | 3.64251328292206 | 0.98294303411898 | 0.3430448564514 | 0.442548942360595 |
| Rundc1 | ENSMUSG00000035007 | 217201 | 0.0943609423721199 | 2.73939387719405 | 0.982734398960446 | 0.3431438440035 | 0.442548942360595 |
| Fam53c | ENSMUSG00000034300 | 66306 | 0.081759172835449 | 4.06795043882802 | 0.982726085730905 | 0.3431477886607 | 0.442548942360595 |
| Rpl21 | ENSMUSG00000041453 | 19933 | -0.095568445976066 | 7.33247429866553 | -0.982707508506732 | 0.3431566037369 | 0.442548942360595 |
| Sqle | ENSMUSG00000022351 | 20775 | 0.303818201766868 | 0.68812505763475 | 0.982342998187397 | 0.3433296001162 | 0.442719813480026 |
| Klc2 | ENSMUSG00000024862 | 16594 | -0.0651373728853164 | 5.29394452452314 | -0.981590775314476 | 0.3436868009757 | 0.44312814596196 |
| Zfp60 | ENSMUSG00000037640 | 22718 | -0.139685584466675 | 2.84183742888361 | -0.981161681350744 | 0.3438906791892 | 0.443338720865346 |
| Ctc1 | ENSMUSG00000020898 | 68964 | -0.124428100272286 | 2.54521657888749 | -0.980955458826236 | 0.3439886936668 | 0.443385778117336 |
| Arhgef5 | ENSMUSG00000033542 | 54324 | 0.10599155198115 | 3.24349566396711 | 0.980914201155529 | 0.3440083052065 | 0.443385778117336 |
| Ocel1 | ENSMUSG00000002396 | 77090 | -0.119687586126052 | 2.6704144319098 | -0.980516479424439 | 0.3441974002089 | 0.443577196614469 |
| Gtf3c5 | ENSMUSG00000026816 | 70239 | 0.129783296227655 | 2.4023766564585 | 0.980402110097233 | 0.3442517902802 | 0.443594992284981 |
| Gtf2h1 | ENSMUSG00000006599 | 14884 | -0.0897145162235495 | 3.72835018376069 | -0.979815086446703 | 0.3445310545474 | 0.443902517357189 |
| Tm9sf4 | ENSMUSG00000068040 | 99237 | 0.0885920751070106 | 4.39909797856607 | 0.979566735384822 | 0.3446492508898 | 0.444002470415653 |
| Pex26 | ENSMUSG00000067825 | 74043 | 0.142355774599963 | 1.68929797157519 | 0.979257269227994 | 0.3447965737574 | 0.444130251086958 |
| Zhx2 | ENSMUSG00000071757 | 387609 | -0.0709456619361664 | 5.4555467876795 | -0.979187699119956 | 0.3448296991103 | 0.444130251086958 |
| Mau2 | ENSMUSG00000031858 | 74549 | 0.0609044726180104 | 4.68334247647683 | 0.978806544014452 | 0.3450112237915 | 0.444311697368631 |
| Usp18 | ENSMUSG00000030107 | 24110 | -0.371395009097584 | 0.922509523713418 | -0.978308947981267 | 0.3452483055641 | 0.444564639901211 |
| Nckipsd | ENSMUSG00000032598 | 80987 | 0.121544877758478 | 2.3338270399585 | 0.977843674382038 | 0.3454700919215 | 0.444797829775599 |
| Men1 | ENSMUSG00000024947 | 17283 | -0.140820718311443 | 2.32332489725846 | -0.977369597597425 | 0.3456961787037 | 0.445036500931718 |
| Clptm1 | ENSMUSG00000002981 | 56457 | -0.0719031493784517 | 5.47798766112802 | -0.97623818792194 | 0.3462361716958 | 0.445679179558125 |
| Slc25a40 | ENSMUSG00000054099 | 319653 | -0.194750546532024 | 1.54738793940014 | -0.976134600735025 | 0.3462856411356 | 0.445690373631717 |
| Ifi213 | ENSMUSG00000073491 | 623121 | -0.460485531513629 | 0.690802618730038 | -0.976036925785485 | 0.3463322917051 | 0.445697937441593 |
| Cltb | ENSMUSG00000047547 | 74325 | 0.0777218329481943 | 5.82960612257671 | 0.975645219218319 | 0.3465194196567 | 0.445886259713678 |
| Armc8 | ENSMUSG00000032468 | 74125 | -0.0720817794985917 | 4.51561684990354 | -0.975351834587762 | 0.3466596237850 | 0.446014165206506 |
| Baz2a | ENSMUSG00000040054 | 116848 | 0.0590359603579333 | 5.04566335197167 | 0.974823077599172 | 0.3469124105141 | 0.446286872935212 |
| Pank3 | ENSMUSG00000018846 | 211347 | 0.0588920951877103 | 4.84567709867325 | 0.974652661979458 | 0.3469939102260 | 0.446339189536435 |
| Prmt3 | ENSMUSG00000030505 | 71974 | 0.0827677704771581 | 3.08358127632071 | 0.974058621697434 | 0.3472781107745 | 0.446652197773418 |
| Zfp263 | ENSMUSG00000022529 | 74120 | 0.101714920379571 | 2.89009906686536 | 0.973818079934309 | 0.3473932376386 | 0.446747703603292 |
| Hdgfl3 | ENSMUSG00000025104 | 29877 | 0.101816625188365 | 2.54961921184137 | 0.97318811903178 | 0.3476948745653 | 0.447083010689843 |
| Ttc9c | ENSMUSG00000071660 | 70387 | 0.0854309090711816 | 3.13294104454756 | 0.972890121406087 | 0.3478376260233 | 0.44717607897764 |
| Kctd9 | ENSMUSG00000034327 | 105440 | -0.0761890656424375 | 5.05261250661815 | -0.972866232606024 | 0.3478490714067 | 0.44717607897764 |
| Dusp16 | ENSMUSG00000030203 | 70686 | -0.0740340411156302 | 4.56589834988211 | -0.972276641591732 | 0.3481316354734 | 0.447486701241764 |
| Ipo7 | ENSMUSG00000066232 | 233726 | 0.0602227736807031 | 6.97636706167503 | 0.972088625065357 | 0.3482217774164 | 0.447549941086316 |
| Dmxl1 | ENSMUSG00000037416 | 240283 | -0.0704243820041235 | 6.10178111887737 | -0.971737634709201 | 0.3483900991842 | 0.447713634397226 |
| Trabd | ENSMUSG00000015363 | 67976 | -0.088773124873148 | 4.28304902163996 | -0.971337922349108 | 0.3485818563919 | 0.447907402400407 |
| Kdm4b | ENSMUSG00000024201 | 193796 | 0.0719225296547217 | 4.13204736380243 | 0.970994363288072 | 0.3487467344709 | 0.44806659079711 |
| Cldn34c1 | ENSMUSG00000079450 | 73061 | -0.305152102731955 | 0.672771111763084 | -0.97010750276535 | 0.3491726046894 | 0.448561022665431 |
| Polh | ENSMUSG00000023953 | 80905 | 0.0833492264488234 | 2.99409145340632 | 0.96995098284723 | 0.3492478037285 | 0.448604905118244 |
| Zfp111 | ENSMUSG00000087598 | 56707 | 0.12146310122989 | 2.0513483207924 | 0.969289318108305 | 0.3495658231842 | 0.448960640726916 |
| Agps | ENSMUSG00000042410 | 228061 | 0.08532882244885 | 4.76783042441528 | 0.96706882551839 | 0.3506345679725 | 0.450232075057606 |
| N6amt1 | ENSMUSG00000044442 | 67768 | -0.11105205622967 | 2.88845533820392 | -0.967061384237098 | 0.3506381534137 | 0.450232075057606 |
| Mrfap1 | ENSMUSG00000055302 | 67568 | -0.0639932770962339 | 6.99080676195713 | -0.966968354504331 | 0.3506829802250 | 0.450236746163954 |
| Rc3h1 | ENSMUSG00000040423 | 381305 | -0.0900329735838188 | 5.39282671783881 | -0.966806112141114 | 0.3507611671584 | 0.450284241715705 |
| Ar | ENSMUSG00000046532 | 11835 | 0.0952654185097055 | 4.66283555899669 | 0.966358761699338 | 0.3509768156075 | 0.450471191498165 |
| Ddx39 | ENSMUSG00000005481 | 68278 | 0.0995435572700001 | 2.88121264877818 | 0.966333040302356 | 0.3509892176369 | 0.450471191498165 |
| Pfn2 | ENSMUSG00000027805 | 18645 | -0.067115311984451 | 5.69827418002839 | -0.966179405650704 | 0.3510633017685 | 0.450513377745004 |
| Tsta3 | ENSMUSG00000022570 | 22122 | -0.101487250498124 | 2.28320260097259 | -0.962825804060761 | 0.3526831943023 | 0.452539030041002 |
| Hsf1 | ENSMUSG00000022556 | 15499 | -0.0778765141773015 | 4.35448826135611 | -0.962467882933974 | 0.3528563921629 | 0.452708124733921 |
| Fbxo28 | ENSMUSG00000047539 | 67948 | -0.103390911266465 | 3.69973450427244 | -0.961309335694208 | 0.3534174239213 | 0.453374704950669 |
| Tbc1d32 | ENSMUSG00000038122 | 544696 | -0.127820566767498 | 2.01457485981115 | -0.960877997343974 | 0.3536264620290 | 0.453589633470985 |
| Myh14 | ENSMUSG00000030739 | 71960 | 0.0984739653539997 | 6.54580407500701 | 0.960452219340709 | 0.3538328908871 | 0.453801165116414 |
| Ash1l | ENSMUSG00000028053 | 192195 | -0.063245875948029 | 6.51864109428487 | -0.960293837874696 | 0.3539097002294 | 0.453846425763464 |
| Ganc | ENSMUSG00000062646 | 76051 | -0.0979097789802283 | 3.41241557882775 | -0.960112284942142 | 0.3539977613536 | 0.453906103150369 |
| Jag2 | ENSMUSG00000002799 | 16450 | -0.0880899314722434 | 4.42106274653513 | -0.960021234185046 | 0.3540419307703 | 0.453909493930426 |
| Tpt1 | ENSMUSG00000060126 | 22070 | -0.0802649445246306 | 8.91681383135853 | -0.95984637604269 | 0.3541267666755 | 0.453927904152173 |
| Eif2ak2 | ENSMUSG00000024079 | 19106 | -0.164243128478862 | 3.19062488161782 | -0.959776836856729 | 0.3541605089738 | 0.453927904152173 |
| Ythdf1 | ENSMUSG00000038848 | 228994 | -0.0673925318994728 | 4.55915655928778 | -0.959734876409879 | 0.3541808704156 | 0.453927904152173 |
| Map4 | ENSMUSG00000032479 | 17758 | 0.0758702207410131 | 9.52424889300162 | 0.959512380782338 | 0.3542888509120 | 0.454013063226272 |
| Spcs1 | ENSMUSG00000021917 | 69019 | 0.116296394929626 | 3.83203957162383 | 0.958622212754467 | 0.3547210948483 | 0.454513689812158 |
| Adh1 | ENSMUSG00000074207 | 11522 | -0.268742028071616 | 2.62618761973222 | -0.957962407399577 | 0.3550417197444 | 0.454871195326591 |
| Mbd6 | ENSMUSG00000025409 | 110962 | 0.110597362832718 | 3.7687269357746 | 0.956862538499351 | 0.3555766417212 | 0.455503137308622 |
| Nfkbil1 | ENSMUSG00000042419 | 18038 | 0.15551416481717 | 1.67020357717461 | 0.956268828766837 | 0.3558656283130 | 0.455819918337171 |
| Nat9 | ENSMUSG00000015542 | 66176 | -0.130083721178935 | 2.0352900668511 | -0.955869891276785 | 0.3560599027865 | 0.456015324822411 |
| P2rx5 | ENSMUSG00000005950 | 94045 | 0.11799591807672 | 3.60198610485119 | 0.953992636009845 | 0.3569750887876 | 0.457133867799645 |
| Cep89 | ENSMUSG00000023072 | 72140 | -0.130950784195629 | 2.6407364012668 | -0.952456832894438 | 0.3577250401775 | 0.458040577975958 |
| Smad2 | ENSMUSG00000024563 | 17126 | 0.0755965424632561 | 3.76907145207821 | 0.952180015421109 | 0.3578603310512 | 0.458160140398329 |
| Ccdc167 | ENSMUSG00000024018 | 68597 | -0.169062050762927 | 2.26844384005639 | -0.950774977355825 | 0.3585475783102 | 0.458986248800764 |
| Arid2 | ENSMUSG00000033237 | 77044 | 0.0675400809741156 | 4.71052839407195 | 0.950457038401607 | 0.3587032203327 | 0.459131721482144 |
| Cdc16 | ENSMUSG00000038416 | 69957 | -0.0655864782777069 | 4.11534961707845 | -0.950218398781273 | 0.3588200737246 | 0.459178216536454 |
| Sec24a | ENSMUSG00000036391 | 77371 | -0.0773570034276796 | 4.29256480977661 | -0.950211281524612 | 0.3588235592035 | 0.459178216536454 |
| Sh3rf1 | ENSMUSG00000031642 | 59009 | 0.172536178632127 | 2.8692033659818 | 0.950006555309177 | 0.3589238283442 | 0.459241675023811 |
| Klhdc2 | ENSMUSG00000020978 | 69554 | -0.0758646526570015 | 6.2153233065448 | -0.949938479684421 | 0.3589571742204 | 0.459241675023811 |
| Dpp8 | ENSMUSG00000032393 | 74388 | 0.0659610554540508 | 6.66390312906943 | 0.949561191704121 | 0.3591420227306 | 0.459375081869517 |
| Pde2a | ENSMUSG00000030653 | 207728 | -2.79120780775469 | 0.133831185603708 | -0.949554097696885 | 0.3591454990080 | 0.459375081869517 |
| Zswim9 | ENSMUSG00000070814 | 321008 | -0.141453607257263 | 2.78004241285729 | -0.949403357753502 | 0.3592193716972 | 0.459415812802428 |
| Eny2 | ENSMUSG00000022338 | 223527 | -0.0828994236804763 | 3.27487577134088 | -0.949114704442637 | 0.3593608609038 | 0.459518449133682 |
| Anapc10 | ENSMUSG00000036977 | 68999 | -0.107338410866992 | 1.83601301597521 | -0.949068118246299 | 0.3593836997204 | 0.459518449133682 |
| Sptb | ENSMUSG00000021061 | 20741 | -0.100580741086889 | 8.32277049206192 | -0.948860760896585 | 0.3594853687017 | 0.459594685997575 |
| Khdc4 | ENSMUSG00000028060 | 74200 | 0.0813448444156086 | 3.92263962944519 | 0.948761160870265 | 0.3595342105602 | 0.459603374533265 |
| Vta1 | ENSMUSG00000019868 | 66201 | 0.0836890434409527 | 3.36063626453672 | 0.94866460354388 | 0.3595815647765 | 0.459610159561835 |
| Snapc2 | ENSMUSG00000011837 | 102209 | 0.134442022676899 | 1.96350119134168 | 0.947904834540329 | 0.3599543276094 | 0.460014473843128 |
| Ccdc191 | ENSMUSG00000022701 | 212153 | -0.175204304471356 | 1.44026321522746 | -0.947848348636319 | 0.3599820518940 | 0.460014473843128 |
| Sap130 | ENSMUSG00000024260 | 269003 | 0.0751165147336699 | 3.60646536884298 | 0.947583289287853 | 0.3601121676949 | 0.460126955590099 |
| Parn | ENSMUSG00000022685 | 74108 | 0.108247314610884 | 2.9126726798 | 0.947490349347451 | 0.3601577990635 | 0.460131475171121 |
| Atg2b | ENSMUSG00000041341 | 76559 | -0.07965843571517 | 4.96979455279039 | -0.946946842012496 | 0.3604247296763 | 0.460418688803627 |
| Csrp1 | ENSMUSG00000026421 | 13007 | -0.0937692177063294 | 4.53811025963593 | -0.946359902281436 | 0.3607131465132 | 0.46070804564251 |
| Fnip2 | ENSMUSG00000061175 | 329679 | 0.170031024218364 | 3.52023074616418 | 0.946314349052369 | 0.3607355377050 | 0.46070804564251 |
| Rad21 | ENSMUSG00000022314 | 19357 | -0.0571224359089282 | 6.01626249639133 | -0.944916878886995 | 0.3614229213398 | 0.461532003874557 |
| Rpl13 | ENSMUSG00000000740 | 270106 | -0.0699507542263236 | 7.23101818832261 | -0.943859000201817 | 0.3619438762407 | 0.462135818959544 |
| Insig1 | ENSMUSG00000045294 | 231070 | 0.161231466418017 | 2.91576153949394 | 0.943785034653959 | 0.3619803203669 | 0.462135818959544 |
| Tor4a | ENSMUSG00000059555 | 227612 | 0.199826517142127 | 0.936369573888424 | 0.943442809886232 | 0.3621489738994 | 0.46229714278813 |
| Larp1 | ENSMUSG00000037331 | 73158 | -0.0548560168393335 | 6.74114422007261 | -0.943241602352346 | 0.3622481576286 | 0.462369758411758 |
| Ssbp4 | ENSMUSG00000070003 | 76900 | 0.0975073253794094 | 2.91754721011262 | 0.943012715333515 | 0.3623610088151 | 0.462459800041863 |
| Ccdc12 | ENSMUSG00000019659 | 72654 | -0.089687614878748 | 3.61989829778928 | -0.942698786132193 | 0.3625158293969 | 0.462603377438421 |
| Bmpr2 | ENSMUSG00000067336 | 12168 | 0.0598497783040477 | 6.92024255183115 | 0.940993681951138 | 0.3633575428905 | 0.463623357223851 |
| Prrc1 | ENSMUSG00000024594 | 73137 | -0.0752760867481292 | 4.92750259636994 | -0.940779653813311 | 0.3634632927177 | 0.463704161145902 |
| Rps26 | ENSMUSG00000025362 | 27370 | -0.0714685219448956 | 6.42790057448804 | -0.940046208450693 | 0.3638258458413 | 0.46411253598921 |
| Gabpb1 | ENSMUSG00000027361 | 14391 | -0.118105035032749 | 2.30708607552777 | -0.93992555354404 | 0.3638855115306 | 0.464134483843823 |
| Pfkfb4 | ENSMUSG00000025648 | 270198 | 0.117973283248143 | 2.65434775883769 | 0.939723485651513 | 0.3639854524616 | 0.464149879190578 |
| Ano8 | ENSMUSG00000034863 | 382014 | -0.0904261989328874 | 3.45573101025005 | -0.939715598570237 | 0.3639893537279 | 0.464149879190578 |
| Pelp1 | ENSMUSG00000018921 | 75273 | -0.0728639359181864 | 4.16827221631282 | -0.939643602930371 | 0.3640249670021 | 0.464149879190578 |
| Zbtb14 | ENSMUSG00000049672 | 22666 | -0.0869488855184568 | 3.31966558644432 | -0.939299577495938 | 0.3641951757339 | 0.464304673255116 |
| Zfp511 | ENSMUSG00000025470 | 69752 | 0.15981835865773 | 1.54525470870667 | 0.939226534595819 | 0.3642313213029 | 0.464304673255116 |
| Fadd | ENSMUSG00000031077 | 14082 | 0.152216662680937 | 1.85798457929819 | 0.938672115727486 | 0.3645057591934 | 0.464600332759218 |
| Actn4 | ENSMUSG00000054808 | 60595 | -0.115847147298584 | 5.95408635986108 | -0.938564043260282 | 0.3645592719483 | 0.464614364191156 |
| Slc38a10 | ENSMUSG00000061306 | 72055 | 0.0827390116473632 | 5.67210642866537 | 0.937276734869223 | 0.3651971116149 | 0.46537300385439 |
| Psmc5 | ENSMUSG00000020708 | 19184 | 0.0758502380222419 | 5.38800733091941 | 0.937171714308097 | 0.3652491818003 | 0.465385103888581 |
| Irgm2 | ENSMUSG00000069874 | 54396 | -0.279494261003274 | 3.63764358933478 | -0.936605522566395 | 0.3655299940601 | 0.465688620637671 |
| Wrb | ENSMUSG00000023147 | 71446 | -0.206358146886852 | 2.10680164171064 | -0.936197808846248 | 0.3657322995388 | 0.465892059930009 |
| Cep295 | ENSMUSG00000046111 | 319675 | -0.100868546397078 | 4.11915232770496 | -0.936033470045219 | 0.3658138656466 | 0.465915090854878 |
| Tmem260 | ENSMUSG00000036339 | 218989 | 0.130066886344603 | 2.4625425463117 | 0.935961391243863 | 0.3658496444368 | 0.465915090854878 |

| | | | | | | | |
|---|---|---|---|---|---|---|---|
| Zfp369 | ENSMUSG00000021514 | 170936 | 0.0709142146530759 | 3.46413557101926 | 0.935903768923386 | 0.3658782490072 | 0.465915090854878 |
| Isy1 | ENSMUSG00000030056 | 57905 | -0.0794121139303723 | 3.68973994121168 | -0.934846601775639 | 0.3664033185373 | 0.466529373900084 |
| Cenpx | ENSMUSG00000025144 | 20892 | 0.122640420477462 | 2.64590743627939 | 0.934525204878593 | 0.3665630524775 | 0.466678398163591 |
| Fhl3 | ENSMUSG00000032643 | 14201 | -0.11814123788205 | 6.87641804995694 | -0.934242303504584 | 0.3667036942402 | 0.466803083933927 |
| Nufip1 | ENSMUSG00000022009 | 27275 | 0.0871398698494952 | 2.58782045638218 | 0.933745150415858 | 0.3669509401046 | 0.467062271708631 |
| Cep83 | ENSMUSG00000020024 | 77048 | -0.0838177628549142 | 3.87679182751872 | -0.933661084941449 | 0.3669927592814 | 0.467062271708631 |
| Plekhm2 | ENSMUSG00000028917 | 69582 | 0.0579215003114166 | 4.82333885495748 | 0.933315019738518 | 0.3671649476168 | 0.467227013085005 |
| Map3k3 | ENSMUSG00000020700 | 26406 | -0.0847918225432744 | 4.45659899499004 | -0.932991135737469 | 0.3673261503102 | 0.467377738219254 |
| Rab1a | ENSMUSG00000020149 | 19324 | -0.0608236300108187 | 5.71198687305144 | -0.932727172277985 | 0.3674575658695 | 0.467490532183494 |
| Pomt2 | ENSMUSG00000034126 | 217734 | 0.11456746198335 | 2.64590332544902 | 0.932145994755428 | 0.3677470231690 | 0.467804341936524 |
| Usp7 | ENSMUSG00000022710 | 252870 | -0.0579517852009088 | 5.97810738444202 | -0.931537559787353 | 0.3680502257302 | 0.468135561724184 |
| Gm4841 | ENSMUSG00000068606 | 225594 | -0.798571445467594 | 2.2794284896661 | -0.931297672425263 | 0.3681698169437 | 0.468233190111974 |
| Cldn5 | ENSMUSG00000041378 | 12741 | -0.304087143740795 | 2.25266902764112 | -0.93096099794174 | 0.3683377049938 | 0.468392211876255 |
| Cdk10 | ENSMUSG00000033862 | 234854 | 0.127424653192042 | 2.0747019238576 | 0.930791878375996 | 0.3684220591812 | 0.468444984170047 |
| Has2 | ENSMUSG00000022367 | 15117 | 0.366705533007386 | 0.306055467674291 | 0.930630193374531 | 0.3685027176575 | 0.468493045675142 |
| Vkorc1l1 | ENSMUSG00000066735 | 69568 | 0.0754112676026284 | 3.87954906747839 | 0.930475868731141 | 0.3685797157597 | 0.468536442954948 |
| Eya3 | ENSMUSG00000028886 | 14050 | 0.0884760077013329 | 3.76904684520162 | 0.930052823051041 | 0.3687908456640 | 0.468750317901631 |
| Ercc8 | ENSMUSG00000021694 | 71991 | -0.159742477011173 | 1.32250079107233 | -0.929663190776656 | 0.3689853740982 | 0.468943044328373 |
| Aco1 | ENSMUSG00000028405 | 11428 | 0.076093054246301 | 3.68445738965232 | 0.929346995873505 | 0.3691432903686 | 0.469089201002109 |
| Rcc1 | ENSMUSG00000028896 | 100088 | -0.129897877592768 | 1.68999876126348 | -0.928834983765107 | 0.3693991024023 | 0.469359710375455 |
| Cwc22 | ENSMUSG00000027014 | 80744 | 0.0836733232627687 | 3.64354502048911 | 0.928268157874341 | 0.3696824438467 | 0.469580127849451 |
| Vps52 | ENSMUSG00000024319 | 224705 | -0.0577063431980687 | 4.31796489083461 | -0.928251204218882 | 0.3696909208550 | 0.469580127849451 |
| Adamtsl3 | ENSMUSG00000070469 | 269959 | 0.135161540675333 | 3.22878513453005 | 0.92823014120491 | 0.3697014527739 | 0.469580127849451 |
| Rfng | ENSMUSG00000025158 | 19719 | -0.0880159956539339 | 3.3131835332273 | -0.927712962451451 | 0.3699601174920 | 0.46985407741439 |
| Slc35d1 | ENSMUSG00000028521 | 242585 | 0.137957325484165 | 2.01348753637527 | 0.926692332394683 | 0.3704709489738 | 0.470416435661742 |
| Aldh3a2 | ENSMUSG00000010025 | 11671 | 0.0919239676780719 | 3.38365311470942 | 0.926656314706036 | 0.3704889849612 | 0.470416435661742 |
| Fdxacb1 | ENSMUSG00000037845 | 382137 | 0.134845218165051 | 1.33621004333058 | 0.926268475952079 | 0.3706832351340 | 0.470558558161766 |
| Zpr1 | ENSMUSG00000032078 | 22687 | -0.0657213704292703 | 3.96671632175022 | -0.926260933527249 | 0.3706870134782 | 0.470558558161766 |
| Pex12 | ENSMUSG00000018733 | 103737 | -0.172650794461233 | 2.07043542549226 | -0.925723622015421 | 0.3709562458937 | 0.470845648381827 |
| Zfp580 | ENSMUSG00000055633 | 68992 | -0.169935855249315 | 1.27492175236961 | -0.925470452030349 | 0.3710831494883 | 0.470952038436941 |
| Nsa2 | ENSMUSG00000060739 | 59050 | 0.0688452646508351 | 5.43282162513715 | 0.924782473669509 | 0.3714281561701 | 0.471335172404878 |
| Ankrd24 | ENSMUSG00000054708 | 70615 | -0.154710196595748 | 1.57225992948164 | -0.92432353794715 | 0.3716584261157 | 0.471572635620615 |
| Tbpl1 | ENSMUSG00000071359 | 237336 | 0.103458291509768 | 3.05615123058769 | 0.92302776173344 | 0.3723091116021 | 0.472342264287983 |
| Fbl | ENSMUSG00000046865 | 14113 | 0.102693521135487 | 3.05323436191098 | 0.922943577613234 | 0.3723514126218 | 0.472342264287983 |
| Emd | ENSMUSG00000001964 | 13726 | -0.0882028897720715 | 3.17271831113156 | -0.922622458424516 | 0.3725127997840 | 0.472492157628174 |
| Brd1 | ENSMUSG00000022387 | 223770 | -0.0696108812751639 | 5.05391085324543 | -0.922471830619515 | 0.3725885185238 | 0.472533367674169 |
| Chmp6 | ENSMUSG00000025371 | 208092 | -0.0817166250885308 | 3.60093125673508 | -0.922067756556741 | 0.3727916940869 | 0.472730912503819 |
| Zmym3 | ENSMUSG00000031310 | 56364 | -0.0836862781150405 | 2.72803256454488 | -0.921958006665254 | 0.3728468914799 | 0.472730912503819 |
| Ctso | ENSMUSG00000028015 | 229445 | 0.0953510245284082 | 2.28987040449967 | 0.921904067965885 | 0.3728740213711 | 0.472730912503819 |
| Rspry1 | ENSMUSG00000050079 | 67610 | -0.157947637532664 | 3.55019262611604 | -0.921364234956624 | 0.3731456197186 | 0.473020383757946 |
| Akr1b3 | ENSMUSG00000001642 | 11677 | -0.0832541939313023 | 5.71510028624073 | -0.920445571827862 | 0.3736081272853 | 0.473524508713404 |
| Daam1 | ENSMUSG00000034574 | 208846 | -0.101110562652102 | 4.76874116997516 | -0.920402262903281 | 0.3736299412362 | 0.473524508713404 |
| BC024978 | ENSMUSG00000078786 | 414069 | -0.129530921951897 | 3.28637995231749 | -0.919380323187017 | 0.3741449295142 | 0.474122214760086 |
| Hace1 | ENSMUSG00000038822 | 209462 | 0.119113310746101 | 1.87391265836856 | 0.919237538913759 | 0.3742169220620 | 0.474158476302307 |
| Tpm4 | ENSMUSG00000031799 | 326618 | 0.138274919791148 | 5.8183968239119 | 0.918741037172003 | 0.3744673350953 | 0.47442077421498 |
| Ccnq | ENSMUSG00000049489 | 69109 | -0.139869568903709 | 2.23606068456611 | -0.918569218351299 | 0.3745540196429 | 0.474475604208767 |
| E130307A14Rik | ENSMUSG00000087177 | 327744 | 0.157260231108895 | 1.42952637156324 | 0.917233441662972 | 0.3752284059396 | 0.475274821011176 |
| Mib2 | ENSMUSG00000029060 | 76580 | -0.0931952447749937 | 3.4369041302451 | -0.91711713834949 | 0.3752871629230 | 0.475294169610913 |
| Nmrk1 | ENSMUSG00000037847 | 225994 | -0.19462805383445 | 2.18401566142036 | -0.916904359525545 | 0.3753946761935 | 0.475375255499439 |
| Med25 | ENSMUSG00000002968 | 75613 | 0.0640772033970256 | 5.3681415129123 | 0.916557484731013 | 0.3755699911963 | 0.475542172334909 |
| Sppl2b | ENSMUSG00000035206 | 73218 | -0.0860781327335056 | 3.28463584444974 | -0.916010679335813 | 0.3758464681822 | 0.47583712574711 |
| Retreg3 | ENSMUSG00000017802 | 67998 | -0.0650839686538931 | 3.91919695475349 | -0.914663579694477 | 0.3765281894023 | 0.476645007337249 |
| Pcx | ENSMUSG00000024892 | 18563 | -0.0897272066829817 | 4.24231851888273 | -0.914504132996593 | 0.3766089362111 | 0.476692019653113 |
| Zfp219 | ENSMUSG00000049295 | 69890 | -0.108423614770614 | 3.09521442049491 | -0.914190641231765 | 0.3767677290700 | 0.476832436954683 |
| Ahcyl1 | ENSMUSG00000027893 | 229709 | -0.050964216425978 | 4.96659019184483 | -0.914112895921947 | 0.3768071164957 | 0.476832436954683 |
| E130309D02Rik | ENSMUSG00000039244 | 231868 | -0.0965050105187801 | 2.74100898305141 | -0.913611997256195 | 0.3770609502922 | 0.477023242433908 |
| Llph | ENSMUSG00000020224 | 66225 | 0.0919365195446851 | 3.39595301522842 | 0.913510483387118 | 0.3771124074580 | 0.477023242433908 |
| Prkacb | ENSMUSG00000005034 | 18749 | -0.0507190317283737 | 4.97902071289653 | -0.913502642408577 | 0.3771163822343 | 0.477023242433908 |
| Taf1b | ENSMUSG00000059669 | 21340 | 0.126795390243798 | 2.25326048264293 | 0.913470937307002 | 0.3771324545891 | 0.477023242433908 |
| Pald1 | ENSMUSG00000020092 | 27355 | 0.0976001657470991 | 3.05188232798792 | 0.9133469037616 | 0.3771953357818 | 0.477045779036284 |
| Lman2l | ENSMUSG00000001143 | 214895 | 0.0821955002246377 | 3.3989458916695 | 0.913263634013979 | 0.3772375550260 | 0.477045779036284 |
| Ube2e1 | ENSMUSG00000021774 | 22194 | -0.085165089677127 | 3.73550518686655 | -0.91264668581204 | 0.3775504599311 | 0.47738624378345 |
| Tardbp | ENSMUSG00000041459 | 230908 | -0.0682414719982489 | 5.17059316703784 | -0.912043618377321 | 0.3778564970926 | 0.47769034914223 |
| Cyp39a1 | ENSMUSG00000023963 | 56050 | -0.173024742738621 | 1.11390075522466 | -0.912000531569536 | 0.3778783687707 | 0.47769034914223 |
| Eif3l | ENSMUSG00000033047 | 223691 | 0.0631273746002625 | 5.58083691859568 | 0.911734148708393 | 0.3780136090545 | 0.477769102856988 |
| Zfp386 | ENSMUSG00000042063 | 56220 | -0.113672014463823 | 3.59697303268266 | -0.911638056861062 | 0.3780624022155 | 0.477769102856988 |
| Foxj3 | ENSMUSG00000032998 | 230700 | -0.0582642619204101 | 5.0869980829119 | -0.91161956922477 | 0.3780717902948 | 0.477769102856988 |
| Ovca2 | ENSMUSG00000038268 | 246257 | 0.16604334927076 | 1.48341004468528 | 0.910419972988546 | 0.3786812912274 | 0.478484012762334 |
| Dhcr7 | ENSMUSG00000058454 | 13360 | -0.159690056916369 | 1.39124387149544 | -0.91008634352526 | 0.3788509242880 | 0.478643025126322 |
| Fbxl16 | ENSMUSG00000025738 | 214931 | 0.191428024961047 | 1.34019693437219 | 0.909669791053163 | 0.3790627925267 | 0.47885535480299 |
| Asb1 | ENSMUSG00000026311 | 65247 | 0.0884426172489432 | 3.41888613415441 | 0.909189913030561 | 0.3793069703501 | 0.479108446140208 |
| Dync1i2 | ENSMUSG00000027012 | 13427 | 0.0535301993604769 | 5.27307221514703 | 0.90805298793893 | 0.3798859059383 | 0.479784267800338 |
| Zkscan7 | ENSMUSG00000063488 | 382118 | -0.149583976301626 | 1.99708794775544 | -0.907860989580321 | 0.3799837335032 | 0.47985237880363 |
| LOC105244208 | ENSMUSG00000081406 | 105244208 | -4.17215914898332 | -0.576082411482134 | -0.907306141499358 | 0.3802665383539 | 0.480154040747076 |
| Trp53inp2 | ENSMUSG00000038375 | 68728 | -0.0745034134676299 | 8.5203125423741 | -0.907203292117988 | 0.3803189762977 | 0.480164787469489 |
| Rpl34 | ENSMUSG00000062006 | 68436 | 0.0782723271551511 | 6.40513868129417 | 0.906875237460391 | 0.3804862686607 | 0.480320522315617 |
| Mta3 | ENSMUSG00000055817 | 116871 | 0.102988612528204 | 2.54886461880195 | 0.906196832049537 | 0.3808323832171 | 0.480701937753675 |
| Slc25a35 | ENSMUSG00000018740 | 71998 | -0.150860712284812 | 1.45438364172165 | -0.905846440796489 | 0.3810112331673 | 0.480872161384686 |
| Zfp637 | ENSMUSG00000059689 | 232337 | -0.0828913547396102 | 2.99755773095304 | -0.90467117025822 | 0.3816115454658 | 0.481574209592127 |
| Nrbf2 | ENSMUSG00000075000 | 641340 | 0.0872281662590195 | 3.47470349906666 | 0.903988363558314 | 0.3819606109308 | 0.481959071693972 |
| Hnrnpa1 | ENSMUSG00000046434 | 15382 | -0.0553626087554502 | 6.03664116394453 | -0.903576156802533 | 0.3821714455590 | 0.482169444991394 |
| Kcnq4 | ENSMUSG00000028631 | 60613 | 0.0790454444431736 | 4.24153758756323 | 0.901874394918985 | 0.3830427012475 | 0.48321289871165 |
| Itgb6 | ENSMUSG00000026971 | 16420 | 0.0905722602818306 | 5.61724881471156 | 0.901484273547834 | 0.3832426237889 | 0.483409314636135 |
| Ankrd17 | ENSMUSG00000055204 | 81702 | -0.0578980216918694 | 6.78152779306073 | -0.900946745764586 | 0.3835182032942 | 0.483701105366205 |
| Uck1 | ENSMUSG00000002550 | 22245 | 0.0928220582903405 | 2.69861132870877 | 0.900280128509063 | 0.3838601522169 | 0.484076525597991 |
| Api5 | ENSMUSG00000027193 | 11800 | 0.0710704968384448 | 4.44190751734214 | 0.899228199266663 | 0.3844001744329 | 0.48470161572573 |
| Prpf39 | ENSMUSG00000035597 | 328110 | 0.125585365207416 | 3.65715141812555 | 0.899121205571871 | 0.3844551301251 | 0.484714997393074 |
| Egln2 | ENSMUSG00000058709 | 112406 | -0.0678928460362682 | 4.9295151302662 | -0.898152207064447 | 0.3849530855514 | 0.485286838676345 |
| Zmym1 | ENSMUSG00000043872 | 68310 | -0.120134528437114 | 1.69061315368105 | -0.897057569610909 | 0.3855161337850 | 0.485846236910202 |
| Erlin1 | ENSMUSG00000025198 | 226144 | 0.124183185195584 | 2.12487368619099 | 0.897039624179326 | 0.3855253690416 | 0.485846236910202 |
| D930016D06Rik | ENSMUSG00000097392 | 100662 | 0.134523554069016 | 1.67132409031474 | 0.897030300552886 | 0.3855301673185 | 0.485846236910202 |
| Trib2 | ENSMUSG00000020601 | 217410 | 0.126676216361748 | 4.26979685377296 | 0.895901065961236 | 0.3861116133079 | 0.486522887039677 |
| Rpl14 | ENSMUSG00000025794 | 67115 | -0.0834972785767052 | 6.56620353362628 | -0.895657342395032 | 0.3862371855359 | 0.486596943567084 |
| Xab2 | ENSMUSG00000019470 | 67439 | 0.0744124593302944 | 3.3453253935515 | 0.895585777799737 | 0.3862740626114 | 0.486596943567084 |
| Terf1 | ENSMUSG00000025925 | 21749 | -0.213596633007214 | 2.06336525624785 | -0.895460052515841 | 0.3863388543717 | 0.486596943567084 |
| Lats1 | ENSMUSG00000040021 | 16798 | 0.0784434906436786 | 4.11860373981613 | 0.89544143986998 | 0.3863484469141 | 0.486596943567084 |
| Lrrc47 | ENSMUSG00000029028 | 72946 | 0.0657102479809941 | 4.39500076320837 | 0.894634189309923 | 0.3867646418179 | 0.487065011487578 |
| Zwint | ENSMUSG00000019923 | 52696 | -0.0818604986415563 | 3.8397029185422 | -0.894496792432759 | 0.3868355100159 | 0.48709814076542 |
| Cfap97 | ENSMUSG00000031631 | 66756 | -0.0928885466991893 | 3.70070320983324 | -0.894244201390295 | 0.3869658174782 | 0.487206098923664 |
| Eif2s3x | ENSMUSG00000035150 | 26905 | -0.0500799449092071 | 5.39507897959973 | -0.893580090870483 | 0.3873085633248 | 0.487581470195106 |
| Hoxa9 | ENSMUSG00000038227 | 15405 | 0.165422228970706 | 1.96139946954532 | 0.892786249521337 | 0.3877185336858 | 0.488041373988966 |
| Cnih1 | ENSMUSG00000015759 | 12793 | -0.0752768449111541 | 4.06479643468959 | -0.892131047042207 | 0.3880571276566 | 0.488410405666997 |
| Ralgapa2 | ENSMUSG00000037110 | 241694 | -0.0712621898525683 | 3.74822061895336 | -0.892046046302904 | 0.3881010688522 | 0.488410405666997 |
| Casc3 | ENSMUSG00000078676 | 192160 | -0.0614818666231054 | 4.03873057026676 | -0.891907443329168 | 0.3881727270117 | 0.488444351210476 |
| Plau | ENSMUSG00000021822 | 18792 | 0.109237441609534 | 3.87002932406553 | 0.891428219730686 | 0.3884205563216 | 0.488699942582003 |
| Rpl29 | ENSMUSG00000048758 | 19944 | -0.0755731955783014 | 7.50897425673368 | -0.890526004117288 | 0.3888874264150 | 0.489219620984924 |
| B3galt1 | ENSMUSG00000034780 | 26877 | -0.149690088803276 | 2.5403573921218 | -0.890457078270228 | 0.3889231091719 | 0.489219620984924 |
| Ndn | ENSMUSG00000033585 | 17984 | 0.144870150873541 | 2.2167074023453 | 0.890296088602242 | 0.3890064618193 | 0.489268166395973 |
| Pwwp3a | ENSMUSG00000020156 | 68114 | -0.0945979188268606 | 3.71613469832766 | -0.890137688744291 | 0.3890884854198 | 0.489315029236596 |
| Nmi | ENSMUSG00000026946 | 64685 | -0.10914775526758 | 2.33314964035711 | -0.889427911258105 | 0.3894561709187 | 0.489721086427316 |
| Notch1 | ENSMUSG00000026923 | 18128 | -0.112768114584246 | 4.83186754690229 | -0.889215213041445 | 0.3895664006986 | 0.48980335013077 |
| Nfkb1 | ENSMUSG00000028163 | 18033 | 0.0634233929988568 | 4.72822238070413 | 0.888934205924598 | 0.3897120636631 | 0.489930140069253 |
| Cdc27 | ENSMUSG00000020687 | 217232 | 0.0748269149347811 | 4.52296338882862 | 0.888360286480596 | 0.3900096755110 | 0.49024790283024 |
| Bbs4 | ENSMUSG00000025235 | 102774 | -0.177323013909609 | 2.10016798199956 | -0.88725642957977 | 0.3905825249187 | 0.490911530399742 |
| Snx21 | ENSMUSG00000050373 | 101113 | 0.102616199185799 | 3.27025495546648 | 0.887059157619093 | 0.3906849597255 | 0.490932173421821 |
| Zfp512b | ENSMUSG00000000823 | 269401 | -0.0854047991995746 | 3.79546856554341 | -0.887051814767509 | 0.3906887729024 | 0.490932173421821 |

| | | | | | | | |
|---|---|---|---|---|---|---|---|
| Morf4l1 | ENSMUSG00000062270 | 21761 | 0.0683475962350633 | 5.84901981222604 | 0.886213044163033 | 0.3911245162766 | 0.491423228439029 |
| Osbp | ENSMUSG00000024687 | 76303 | -0.0715922402170133 | 5.83266343206654 | -0.885885654825953 | 0.3912946850857 | 0.491580531280562 |
| Hist1h1e | ENSMUSG00000051627 | 50709 | 0.0697473673892102 | 4.93844741244953 | 0.885639050302186 | 0.3914228970527 | 0.491685093964983 |
| Phf3 | ENSMUSG00000048874 | 213109 | 0.0518884001509692 | 5.63429835148102 | 0.8854856776302 | 0.3915026512594 | 0.491728769495258 |
| Napg | ENSMUSG00000024581 | 108123 | 0.0679784032194406 | 3.75535872489457 | 0.885355569313293 | 0.3915702521611 | 0.491757172147643 |
| Tbc1d9b | ENSMUSG00000036644 | 76795 | -0.0593529214486006 | 4.93578755978295 | -0.885151967070873 | 0.3916762196524 | 0.491833745780679 |
| 2010315B03Rik | ENSMUSG00000074829 | 630836 | -0.148118277969359 | 2.23423074845176 | -0.884754643552513 | 0.3918829420906 | 0.492036806799109 |
| Klhl41 | ENSMUSG00000075307 | 228003 | 0.0788584838593209 | 9.34191899050403 | 0.884642537991205 | 0.3919412825550 | 0.49205353848738 |
| Med31 | ENSMUSG00000020801 | 67279 | -0.10903237577575 | 1.80886229942838 | -0.884113440564749 | 0.3922167075950 | 0.492342768801295 |
| Ccdc181 | ENSMUSG00000026578 | 74895 | -0.093082069608696 | 3.55032499221558 | -0.883696436302257 | 0.3924338740302 | 0.49255881008438 |
| Emc10 | ENSMUSG00000008140 | 69683 | 0.0769024028785985 | 4.08993455319204 | 0.883164054013391 | 0.3927112448660 | 0.49285035793696 |
| Cox8a | ENSMUSG00000035885 | 12868 | -0.0653139554507516 | 3.93188412948673 | -0.882638321965585 | 0.3929852809276 | 0.493137654209698 |
| Xrcc4 | ENSMUSG00000021615 | 108138 | -0.111701326241378 | 2.08534259824274 | -0.882483031644516 | 0.3930662501981 | 0.493179445866605 |
| Mtap | ENSMUSG00000062937 | 66902 | 0.166925509500806 | 1.49469893124302 | 0.882401395954491 | 0.3931088200380 | 0.493179445866605 |
| Usp45 | ENSMUSG00000040455 | 77593 | -0.0867166261005341 | 3.41542531425625 | -0.881869762749116 | 0.3933861222774 | 0.493470702618214 |
| Plcg1 | ENSMUSG00000016933 | 18803 | 0.0742046433030028 | 4.78077638053755 | 0.880945267431177 | 0.3938686576754 | 0.493999339032852 |
| Comt | ENSMUSG00000000326 | 12846 | -0.0594726049458973 | 4.31463575071228 | -0.880889214766127 | 0.3938979269060 | 0.493999339032852 |
| Pign | ENSMUSG00000056536 | 27392 | 0.13809724701534 | 2.83814059089832 | 0.880222856229099 | 0.3942459943087 | 0.49437914004692 |
| Gm31166 | ENSMUSG00000109695 | 102633307 | -0.106932903254344 | 2.33119210734418 | -0.879868821958936 | 0.3944310058578 | 0.494554407551305 |
| Top3b | ENSMUSG00000022779 | 21976 | -0.0715174788066015 | 3.33582764543959 | -0.879482314009169 | 0.3946330543939 | 0.494750994102623 |
| Scfd2 | ENSMUSG00000062110 | 212986 | 0.0990130347318616 | 2.32881218694119 | 0.878689708367509 | 0.3950476103826 | 0.495213925354715 |
| Irf2bp1 | ENSMUSG00000044030 | 272359 | 0.0887786986306547 | 3.44442925883771 | 0.878563434634847 | 0.3951136823486 | 0.49523995662799 |
| Smdt1 | ENSMUSG00000022452 | 69029 | -0.0709716149063073 | 4.29188327369458 | -0.877497328337212 | 0.3956718129080 | 0.495882663024275 |
| Sp2 | ENSMUSG00000018678 | 78912 | 0.0701192538264164 | 3.28454604462977 | 0.875437029865329 | 0.3967519295685 | 0.497173175147552 |
| Abca5 | ENSMUSG00000018800 | 217265 | 0.0691063152155715 | 4.64681373502138 | 0.875359722816903 | 0.3967924965682 | 0.497173175147552 |
| Cntrob | ENSMUSG00000032782 | 216846 | 0.207558144885043 | 0.946115325556962 | 0.875184593739341 | 0.3968844061586 | 0.497231340254406 |
| Zfp940 | ENSMUSG00000050855 | 233057 | -0.217410389195173 | 1.2331744224279 | -0.874952722063146 | 0.3970061168967 | 0.497326823722009 |
| Napa | ENSMUSG00000006024 | 108124 | -0.0607007362140717 | 5.43381238641313 | -0.874544140239885 | 0.3972206448683 | 0.497538543492101 |
| Prorsd1 | ENSMUSG00000032673 | 67939 | -0.112390331004503 | 2.18205171017903 | -0.874414736818639 | 0.3972886050470 | 0.497566652356685 |
| Xpnpep3 | ENSMUSG00000022401 | 321003 | 0.113737277212236 | 2.32144702314306 | 0.874104870827345 | 0.3974513724582 | 0.497713478329785 |
| Fmnl3 | ENSMUSG00000023008 | 22379 | -0.155483154667687 | 4.01037640843037 | -0.873366536943995 | 0.3978393875251 | 0.498142307564409 |
| Gpatch4 | ENSMUSG00000028069 | 66614 | -0.0903121571715775 | 3.91686245247233 | -0.872903969978987 | 0.3980826090375 | 0.498389760553102 |
| Map6 | ENSMUSG00000055407 | 17760 | 0.15389028090636 | 2.92420750737466 | 0.871835172645918 | 0.3986449734501 | 0.498965552047982 |
| Mier1 | ENSMUSG00000028522 | 71148 | 0.0604309655759439 | 4.70549808610725 | 0.871783217780651 | 0.3986723239029 | 0.498965552047982 |
| Fgf2 | ENSMUSG00000037225 | 14173 | -0.0997388098103928 | 3.4960310354681 | -0.871769670395525 | 0.3986794558217 | 0.498965552047982 |
| Polr2d | ENSMUSG00000024258 | 69241 | -0.116326555121009 | 1.83751262171842 | -0.870826200438902 | 0.3991763491582 | 0.499530243005009 |
| Lsm8 | ENSMUSG00000044155 | 76522 | -0.16567600330233 | 1.42036741231992 | -0.870286009501434 | 0.3994610362434 | 0.499829279667762 |
| Rhbdd1 | ENSMUSG00000026142 | 76867 | 0.133210679320073 | 2.26788612759236 | 0.869934058446918 | 0.3996465919165 | 0.500004222987262 |
| Zdhhc16 | ENSMUSG00000025157 | 74168 | 0.0991324844897088 | 2.21078833174998 | 0.869769159182159 | 0.3997335500085 | 0.500055783376418 |
| Arl6ip1 | ENSMUSG00000030654 | 54208 | 0.136016832604617 | 2.06899983572216 | 0.869210401084427 | 0.4000283002190 | 0.500367244501028 |
| Rbm12b1 | ENSMUSG00000046667 | 72397 | 0.143522537190346 | 1.58988151745569 | 0.86906672181618 | 0.4001041159433 | 0.500404815946943 |
| Ldah | ENSMUSG00000037669 | 68832 | -0.0874048342120939 | 2.99927526624906 | -0.868209971906261 | 0.4005563999913 | 0.500913168779877 |
| Plekhh3 | ENSMUSG00000035172 | 217198 | -0.178379482742986 | 3.20489011644614 | -0.867716941197809 | 0.4008168294433 | 0.501181510254503 |
| Timm10b | ENSMUSG00000089847 | 14356 | 0.0759460740045186 | 3.67739084767498 | 0.867606517774189 | 0.4008751730181 | 0.50119713099176 |
| Tmem251 | ENSMUSG00000046675 | 320351 | -0.151504582559257 | 1.66815569792958 | -0.866525481688028 | 0.4014466521131 | 0.501854226240658 |
| Sipa1l1 | ENSMUSG00000042700 | 217692 | -0.0740301363827656 | 3.82436581077529 | -0.866359640056237 | 0.4015343708813 | 0.501906484631626 |
| Brd3 | ENSMUSG00000026918 | 67382 | -0.0541076259710849 | 5.06508353189538 | -0.866094425583655 | 0.4016746776755 | 0.501990534906857 |
| Xrcc6 | ENSMUSG00000022471 | 14375 | -0.0853910404433158 | 3.94484494513624 | -0.866058927479496 | 0.4016934597777 | 0.501990534906857 |
| Mapt | ENSMUSG00000018411 | 17762 | 0.0790686374769969 | 6.18190219753416 | 0.865942942184346 | 0.4017548318746 | 0.502009838502679 |
| Tmem200a | ENSMUSG00000049420 | 77220 | -0.502465263780046 | 0.077851385365595 | -0.865749270729641 | 0.4018573245664 | 0.502080513754231 |
| Acvr1b | ENSMUSG00000000532 | 11479 | -0.149477820885226 | 2.81565426699483 | -0.865425939193419 | 0.4020284735250 | 0.502236942183144 |
| Zscan2 | ENSMUSG00000038797 | 22691 | -0.154322308796224 | 1.47515225513126 | -0.865184654928548 | 0.4021562241948 | 0.502316372988097 |
| Ythdf3 | ENSMUSG00000047213 | 229096 | -0.0639521249384867 | 5.32759351782725 | -0.865066465937046 | 0.4022188105916 | 0.502316372988097 |
| Zfp934 | ENSMUSG00000074865 | 77117 | -0.163114367940264 | 1.20939997366045 | -0.865045495452196 | 0.4022299160886 | 0.502316372988097 |
| Acer3 | ENSMUSG00000030760 | 66190 | 0.138804991136939 | 3.53342367899222 | 0.864367780530475 | 0.4025889290508 | 0.502707286206891 |
| Chaf1a | ENSMUSG00000002835 | 27221 | -0.146884922542101 | 1.39069869288039 | -0.863890189950172 | 0.4028420566882 | 0.5029659076709 |
| Cd40 | ENSMUSG00000017652 | 21939 | -0.298418443048946 | 1.53892849171594 | -0.863767357742468 | 0.4029071761398 | 0.502989760436859 |
| Tspan6 | ENSMUSG00000067377 | 56496 | 0.226078635096986 | 0.97276894383062 | 0.863528469465671 | 0.4030338428166 | 0.503090434604118 |
| Traf6 | ENSMUSG00000027164 | 22034 | -0.0648125933164243 | 3.72629045812408 | -0.863090628966046 | 0.4032660697772 | 0.503322837261412 |
| Fastkd5 | ENSMUSG00000079043 | 380601 | -0.11620562586087 | 1.88157498817537 | -0.861908471188906 | 0.4038935227457 | 0.504048418442008 |
| Mink1 | ENSMUSG00000020827 | 50932 | 0.0533003895035407 | 5.06984489504659 | 0.861681556633215 | 0.4040140365166 | 0.504069874177026 |
| Trmu | ENSMUSG00000022386 | 72026 | -0.11317917700314 | 2.47587714521048 | -0.861659637426441 | 0.4040256790251 | 0.504069874177026 |
| Plgrkt | ENSMUSG00000016495 | 67759 | -0.096195473460214 | 2.21247018661358 | -0.861615626055148 | 0.4040490565857 | 0.504069874177026 |
| Gorab | ENSMUSG00000040124 | 98376 | 0.145021045301395 | 1.36271406792602 | 0.861331398935102 | 0.4042000515044 | 0.504200703297431 |
| Taok2 | ENSMUSG00000059981 | 381921 | 0.0707609533023703 | 5.03926364403612 | 0.860290258740574 | 0.4047534760192 | 0.504833437513259 |
| Gigyf1 | ENSMUSG00000029714 | 57330 | -0.0663144189699461 | 4.38838274289151 | -0.860008410942578 | 0.4049033808285 | 0.504955050330639 |
| Mindy1 | ENSMUSG00000038712 | 75007 | -0.0589529060446674 | 4.64749042618348 | -0.859933236996032 | 0.4049433694262 | 0.504955050330639 |
| Ing1 | ENSMUSG00000045969 | 26356 | 0.090271198292966 | 3.46361453448057 | 0.8597497607344 | 0.4050409802310 | 0.505019157625867 |
| Coro2b | ENSMUSG00000041729 | 235431 | 0.185933424269544 | 2.81938594310139 | 0.855052165988168 | 0.4075454769736 | 0.508083896988881 |
| Olfr1247 | ENSMUSG00000075081 | 258968 | -0.327657934942708 | 2.34753919413992 | -0.854799294062076 | 0.4076805856807 | 0.508194375878219 |
| Phf21a | ENSMUSG00000058318 | 192285 | 0.071392818424818 | 3.7460815459223 | 0.854635208542416 | 0.4077682720018 | 0.508245721921542 |
| Jade2 | ENSMUSG00000020387 | 76901 | 0.132221499727113 | 2.82987172367661 | 0.854318495660196 | 0.4079375569116 | 0.508353167832156 |
| Zfp87 | ENSMUSG00000097333 | 170763 | 0.145565582019521 | 1.4279080851068 | 0.854299918962 55 | 0.4079474877160 | 0.508353167832156 |
| Ints13 | ENSMUSG00000040250 | 71177 | 0.093139834655815 | 3.6098655577226 | 0.854091637657936 | 0.4080588425534 | 0.508433968762294 |
| Ldb3 | ENSMUSG00000021798 | 24131 | -0.0715102498455877 | 10.1531329562317 | -0.853311492754592 | 0.4084761161168 | 0.508895877054199 |
| Rps27 | ENSMUSG00000090733 | 57294 | -0.0852458170586594 | 5.89372869698914 | -0.853064722559812 | 0.4086081642511 | 0.509002375319567 |
| Mroh1 | ENSMUSG00000022558 | 223658 | 0.0655692881066408 | 3.95324930020523 | 0.852864186770527 | 0.4087154929622 | 0.509078059887183 |
| Ldb2 | ENSMUSG00000039706 | 16826 | 0.166235353860491 | 1.32054162659369 | 0.852402882510739 | 0.4089624585333 | 0.509327632930123 |
| Ints11 | ENSMUSG00000029034 | 71957 | -0.100349376444883 | 3.28998624559536 | -0.852177098878123 | 0.4090833709917 | 0.509420178663824 |
| Mettl26 | ENSMUSG00000025731 | 68347 | -0.132854529144959 | 1.79715639817442 | -0.851594358625998 | 0.4093955517365 | 0.509750857276668 |
| Mri1 | ENSMUSG00000004996 | 67873 | 0.0886155609789472 | 2.67375804251691 | 0.849310474046229 | 0.4106205779995 | 0.511217942837484 |
| Jag1 | ENSMUSG00000027276 | 16449 | -0.121270459988309 | 3.78649370007665 | -0.847597090122373 | 0.4115411924675 | 0.512305748190718 |
| Limk2 | ENSMUSG00000020451 | 16886 | 0.088432559012154 | 3.29339312623379 | 0.847438254651704 | 0.4116266051210 | 0.512353725868641 |
| Atad2b | ENSMUSG00000052812 | 320817 | 0.0526631367678485 | 4.50884697526587 | 0.846435591183915 | 0.4121660509795 | 0.512966765713007 |
| Zfp202 | ENSMUSG00000025602 | 80902 | -0.150442626710059 | 1.33486214511374 | -0.846070747397169 | 0.4123624575754 | 0.513152780482437 |
| Eif2d | ENSMUSG00000026427 | 16865 | -0.0780133379809258 | 3.59024612484772 | -0.844790165626143 | 0.4130523235638 | 0.513952754567631 |
| Ccnd2 | ENSMUSG00000000184 | 12444 | 0.104766062006194 | 4.71002994004268 | 0.844411659908427 | 0.4132563753797 | 0.514092199470478 |
| Ddx39b | ENSMUSG00000019432 | 53817 | 0.0513811409818237 | 5.22233478563373 | 0.844278713553675 | 0.4133280623369 | 0.514092199470478 |
| Nmrk2 | ENSMUSG00000004939 | 69564 | -0.190396609576904 | 4.31582654638813 | -0.844243734455553 | 0.4133469250316 | 0.514092199470478 |
| Cirbp | ENSMUSG00000045193 | 12696 | 0.121892278657333 | 3.05312483162605 | 0.844233368639997 | 0.4133525149708 | 0.514092199470478 |
| Pik3c2b | ENSMUSG00000026447 | 240752 | 0.0853134445991772 | 4.11891232996401 | 0.843645056152727 | 0.4136698541117 | 0.514428347587623 |
| Vps33b | ENSMUSG00000030534 | 233405 | 0.105662820458835 | 1.92457377463648 | 0.843388573705108 | 0.4138082525412 | 0.51449221411732 |
| Pdcd7 | ENSMUSG00000041837 | 50996 | 0.0678190998567825 | 4.19125785556872 | 0.843375429608457 | 0.4138153459445 | 0.51449221411732 |
| Wdr24 | ENSMUSG00000025737 | 268933 | -0.0895844326508537 | 2.95930978599659 | -0.842678109638676 | 0.4141917798688 | 0.514901665614317 |
| Mysm1 | ENSMUSG00000062627 | 320713 | 0.0844951104677927 | 3.37175081506469 | 0.842220732052634 | 0.4144388084349 | 0.515092053289138 |
| Ap3d1 | ENSMUSG00000020198 | 11776 | -0.0692811402030999 | 5.53962380632296 | -0.842150066350979 | 0.4144769834789 | 0.515092053289138 |
| 9930104L06Rik | ENSMUSG00000044730 | 194268 | -0.129581777715816 | 1.99878345178851 | -0.842132828767424 | 0.4144862959227 | 0.515092053289138 |
| Map2k2 | ENSMUSG00000035027 | 26396 | -0.0583732954356657 | 6.19787610632543 | -0.841348383537738 | 0.4149102311368 | 0.515560274702363 |
| Xrcc1 | ENSMUSG00000051768 | 22594 | -0.0922230900240328 | 2.85057803606938 | -0.840807852393532 | 0.4152025148964 | 0.515864820451532 |
| Hlcs | ENSMUSG00000040820 | 110948 | -0.0642385572440434 | 3.79560307219855 | -0.838706159016165 | 0.4163402614584 | 0.517219615638437 |
| Fam222b | ENSMUSG00000037750 | 216971 | -0.0803775685953023 | 2.62147381098783 | -0.838051912346503 | 0.4166948546392 | 0.517589497459587 |
| Fau | ENSMUSG00000038274 | 14109 | 0.0568535735065074 | 6.13510837014439 | 0.837853109204785 | 0.4168026426946 | 0.517589497459587 |
| Nus1 | ENSMUSG00000023068 | 52014 | 0.0841897906063878 | 4.38849951250943 | 0.837848827396424 | 0.4168049644280 | 0.517589497459587 |
| Plekha4 | ENSMUSG00000040428 | 69217 | 0.155378602626967 | 1.53208351424891 | 0.837807445671588 | 0.4168274033607 | 0.517589497459587 |
| Upf3b | ENSMUSG00000036572 | 68134 | -0.0944047041917706 | 3.04814375768341 | -0.837124603109268 | 0.4171977843670 | 0.517990570299473 |
| Zfp608 | ENSMUSG00000052713 | 269023 | 0.122322615742142 | 3.39027904835144 | 0.836806642578241 | 0.4173703233134 | 0.518145940276995 |
| Crlf3 | ENSMUSG00000017561 | 54394 | 0.117701629367596 | 2.10516324298165 | 0.83598831394369 | 0.4178145989613 | 0.518638585197239 |
| Ino80e | ENSMUSG00000030689 | 233875 | 0.109059772709216 | 2.51755536139741 | 0.835839447210466 | 0.4178954530115 | 0.518680049604824 |
| Zfp800 | ENSMUSG00000039841 | 627049 | 0.0747956349739112 | 2.81545785005185 | 0.835593361111106 | 0.4180291323901 | 0.518787062460984 |
| Arid4b | ENSMUSG00000039219 | 94246 | 0.0558863112368453 | 4.70798376877365 | 0.834582081458235 | 0.4185787765023 | 0.51941021749883 |
| Eef1akmt1 | ENSMUSG00000021951 | 68043 | 0.101905360540492 | 2.87735284731085 | 0.833787524230136 | 0.4190109616365 | 0.519887493944309 |
| Crtc2 | ENSMUSG00000027936 | 74343 | -0.0891472658289518 | 3.30325922171669 | -0.833490118136928 | 0.4191728056022 | 0.52002927454758 |
| Ncbp2 | ENSMUSG00000022774 | 68092 | -0.114332941731409 | 2.92338927178035 | -0.832899664693772 | 0.4194942430480 | 0.520368993504093 |
| Spaca6 | ENSMUSG00000080316 | 75202 | 0.196096722296487 | 2.02614091401984 | 0.832683229599099 | 0.4196121087950 | 0.52039776914102 |
| Rabif | ENSMUSG00000042229 | 98710 | -0.107985348293376 | 2.00698273512821 | -0.832542470330422 | 0.4196887748162 | 0.52039776914102 |
| Ssr4 | ENSMUSG00000002014 | 20832 | 0.0612945322523615 | 3.99640083522804 | 0.832533748214667 | 0.4196935257107 | 0.52039776914102 |

| | | | | | | | |
|---|---|---|---|---|---|---|---|
| Elac2 | ENSMUSG00000020549 | 68626 | 0.0766275275510413 | 4.31432345599996 | 0.832483356451771 | 0.4197209745514 | 0.52039776914102 |
| Wipi2 | ENSMUSG00000029578 | 74781 | -0.0549859244971769 | 4.39309482578869 | -0.83242001651645 | 0.4197554780455 | 0.52039776914102 |
| Mgat1 | ENSMUSG00000020346 | 17308 | -0.057732736477818 | 4.47597912885995 | -0.832285376421544 | 0.4198288274238 | 0.520429679356967 |
| Vamp3 | ENSMUSG00000028955 | 22319 | -0.0728972739680396 | 4.52357977307086 | -0.831071454738624 | 0.4204905282160 | 0.521190833873473 |
| Rnmt | ENSMUSG00000009535 | 67897 | 0.0780890571742616 | 3.58709878984116 | 0.830504857904349 | 0.4207996101092 | 0.52151480023857 |
| Tpra1 | ENSMUSG00000002871 | 24100 | -0.089767993133048 | 3.2236556055818 | -0.829949199532762 | 0.4211028694086 | 0.52183147778091 |
| Limch1 | ENSMUSG00000037736 | 77569 | 0.0951201083926234 | 7.83079483170341 | 0.82967181416359 | 0.4212543103411 | 0.521959971246722 |
| Szt2 | ENSMUSG00000033253 | 230676 | 0.123590515935948 | 3.31427168542507 | 0.828805368076519 | 0.4217275834884 | 0.522487160275717 |
| Ube2i | ENSMUSG00000015120 | 22196 | 0.060456867830875 | 4.39135309204386 | 0.828586670150528 | 0.4218470963533 | 0.522575998440424 |
| Ppp2cb | ENSMUSG00000009630 | 19053 | -0.0659934090176177 | 4.12550344106947 | -0.82805253359407 | 0.4221390814930 | 0.522878447569426 |
| Alyref | ENSMUSG00000025134 | 21681 | -0.0541920581175268 | 3.96962972533154 | -0.827747924484143 | 0.4223056548443 | 0.523025505676762 |
| Secisbp2l | ENSMUSG00000035093 | 70354 | -0.0573300846380134 | 5.41694946670525 | -0.827620780590151 | 0.4223751952927 | 0.523052368839342 |
| Ebp | ENSMUSG00000031168 | 13595 | 0.159926915223504 | 1.43141293160373 | 0.827331652812065 | 0.4225333595247 | 0.523142592021546 |
| Fgf6 | ENSMUSG00000000183 | 14177 | -0.172907722084477 | 1.87998238563174 | -0.827312625530423 | 0.4225437695506 | 0.523142592021546 |
| Tex264 | ENSMUSG00000040813 | 21767 | -0.0759925380424402 | 3.38834883147933 | -0.827167495390988 | 0.4226231772888 | 0.523181647898567 |
| Rpl28 | ENSMUSG00000030432 | 19943 | -0.0732033420073313 | 5.60175161283808 | -0.82650307557581 | 0.4229868380430 | 0.523572542933862 |
| Slc9a6 | ENSMUSG00000060681 | 236794 | -0.0632487932033872 | 4.19990403222138 | -0.825817595324136 | 0.4233622399938 | 0.523937364102268 |
| Nars2 | ENSMUSG00000018995 | 244141 | -0.131569059093155 | 1.92087522280701 | -0.825789858562143 | 0.4233774345545 | 0.523937364102268 |
| Otud7b | ENSMUSG00000038495 | 229603 | -0.0569969894106857 | 5.59693888795099 | -0.825546156089801 | 0.4235109532535 | 0.524043268056916 |
| Focad | ENSMUSG00000038368 | 230393 | -0.0848720824880007 | 3.41118959701397 | -0.825267743448268 | 0.4236635224363 | 0.524172718025127 |
| Mafg | ENSMUSG00000051510 | 17134 | 0.072862903206941 | 5.12732136050438 | 0.824421079297247 | 0.4241277118057 | 0.524687643475362 |
| Chmp1a | ENSMUSG00000000743 | 234852 | 0.0718230817558574 | 4.15094422061479 | 0.823512882720178 | 0.4246260052518 | 0.52524463770605 |
| Dcaf17 | ENSMUSG00000041966 | 75763 | 0.114342841525011 | 2.05327445175987 | 0.822833092433997 | 0.4249992302695 | 0.525646818972175 |
| Wdr70 | ENSMUSG00000039828 | 545085 | 0.0609445026077955 | 3.38220904539619 | 0.822738258284425 | 0.4250513140083 | 0.525651760767598 |
| Deaf1 | ENSMUSG00000058886 | 54006 | -0.121306467800527 | 2.51646938218111 | -0.822483914811663 | 0.4251910221788 | 0.525760040304282 |
| Dars | ENSMUSG00000026356 | 226414 | -0.056724930527257 | 5.21955845835076 | -0.822403741176254 | 0.4252350669042 | 0.525760040304282 |
| Srek1 | ENSMUSG00000032621 | 218543 | 0.0688040564679575 | 4.40292089018428 | 0.822134841581716 | 0.4253828130786 | 0.52586325796697 |
| Ube2w | ENSMUSG00000025939 | 66799 | 0.102446623265872 | 2.59410640438591 | 0.822014745620517 | 0.4254488102841 | 0.52586325796697 |
| Pno1 | ENSMUSG00000020116 | 66249 | -0.0797930560796871 | 4.46143621450593 | -0.821989158965173 | 0.4254628719659 | 0.52586325796697 |
| Zbtb4 | ENSMUSG00000018750 | 75580 | 0.0521280663797441 | 5.13933586818487 | 0.820875638161164 | 0.4260751236970 | 0.526560449647558 |
| Trip4 | ENSMUSG00000032386 | 56404 | 0.0731714446535654 | 3.81397222014253 | 0.820419548744419 | 0.4263260626048 | 0.526811009533494 |
| Efhd1 | ENSMUSG00000026255 | 98363 | -0.14595650961166 | 1.11140960700382 | -0.820259494059308 | 0.4264141469491 | 0.526860296179609 |
| Thumpd3 | ENSMUSG00000030264 | 14911 | 0.066011353560818 | 3.28303801533707 | 0.819966187900276 | 0.4265755954965 | 0.527000207320644 |
| Tmem38b | ENSMUSG00000028420 | 52076 | 0.0980003159969272 | 5.72226073707075 | 0.819525339537618 | 0.4268183324779 | 0.527183446111387 |
| Fktn | ENSMUSG00000028414 | 246179 | -0.0857179752404971 | 3.84341457237529 | -0.819521634184286 | 0.4268203730752 | 0.527183446111387 |
| Hp1bp3 | ENSMUSG00000028759 | 15441 | -0.0461131818740288 | 6.19829207739714 | -0.818886328955737 | 0.4271703395633 | 0.527543619617749 |
| Rrp8 | ENSMUSG00000030888 | 101867 | 0.0886255179048745 | 2.81791528050006 | 0.818817074339606 | 0.4272085006729 | 0.527543619617749 |
| Scoc | ENSMUSG00000063253 | 56367 | -0.0849830801731279 | 3.59063648285912 | -0.818194471153766 | 0.4275516708041 | 0.52790775031738 |
| Aaas | ENSMUSG00000036678 | 223921 | -0.162792546904034 | 1.42965273231139 | -0.817200213925403 | 0.4281000625113 | 0.528525161882957 |
| Nckap1 | ENSMUSG00000027002 | 50884 | -0.0508986061468617 | 6.1123607734217 | -0.816928615105844 | 0.4282499446846 | 0.528650496368991 |
| Scara5 | ENSMUSG00000022032 | 71145 | 0.112217529736122 | 3.64174733771255 | 0.816606262970256 | 0.4284278793259 | 0.528796369518266 |
| Tbc1d1 | ENSMUSG00000029174 | 57915 | -0.120959888482603 | 4.93684647305022 | -0.816466008549932 | 0.4285053131114 | 0.528796369518266 |
| B4galt7 | ENSMUSG00000021504 | 218271 | -0.109560131025066 | 2.12940435566085 | -0.816451649202929 | 0.4285132413496 | 0.528796369518266 |
| Fam98c | ENSMUSG00000030590 | 73833 | -0.0910081147163502 | 2.36840478712461 | -0.815949004201532 | 0.4287908271087 | 0.52907918852304 |
| Lrrc49 | ENSMUSG00000047766 | 102747 | 0.129704889880986 | 2.11021972852608 | 0.81537400034011 | 0.4291085160119 | 0.529411420826916 |
| Ranbp1 | ENSMUSG00000005732 | 19385 | 0.0731978162399778 | 3.51805672311162 | 0.815194500418735 | 0.4292077207317 | 0.52947405431882 |
| Pum1 | ENSMUSG00000028580 | 80912 | -0.0555113316230083 | 5.53879318555491 | -0.815084721634594 | 0.4292683998020 | 0.529489153490919 |
| Tfip11 | ENSMUSG00000029345 | 54723 | 0.0659842087445992 | 3.68372659250791 | 0.814949011292002 | 0.4293434199532 | 0.529521936534857 |
| Tsen34 | ENSMUSG00000035585 | 66078 | -0.0601527984501611 | 3.70557061343205 | -0.814821426673766 | 0.4294139559831 | 0.529549182406611 |
| Eed | ENSMUSG00000030619 | 13626 | -0.0808302767434568 | 3.01572174715167 | -0.81417490260854 | 0.4297715066618 | 0.529908030674928 |
| Atg14 | ENSMUSG00000037526 | 100504663 | -0.100755069731715 | 2.65811903173641 | -0.814119956228817 | 0.4298019028418 | 0.529908030674928 |
| Rbm7 | ENSMUSG00000042396 | 67010 | -0.0908068846795946 | 3.15647502565401 | -0.813987135857643 | 0.4298753844414 | 0.52993885500502 |
| Cog2 | ENSMUSG00000031979 | 76332 | 0.0688677771669144 | 3.97782799371446 | 0.813879052477674 | 0.4299351865411 | 0.529952810563941 |
| Galnt11 | ENSMUSG00000038072 | 231050 | 0.105679355973717 | 3.19333753755567 | 0.813785078997751 | 0.4299871860679 | 0.529957146342141 |
| Dagla | ENSMUSG00000035735 | 269060 | -0.164611920873868 | 2.11601449577695 | -0.812901385333964 | 0.4304763705746 | 0.530500248788274 |
| Glt28d2 | ENSMUSG00000031286 | 320302 | -0.173807341905196 | 1.39992307838052 | -0.812644843427127 | 0.4306184513890 | 0.530551525431484 |
| Dpm2 | ENSMUSG00000026810 | 13481 | 0.143151597176325 | 1.35714254910045 | 0.812600736015947 | 0.4306428824892 | 0.530551525431484 |
| Mpzl1 | ENSMUSG00000026566 | 68481 | -0.132217021545733 | 1.99312594063831 | -0.812368060579276 | 0.4307717762843 | 0.530551525431484 |
| Taf5l | ENSMUSG00000038697 | 102162 | -0.0715759344979694 | 3.01633383204132 | -0.812258533513348 | 0.4308324589645 | 0.530551525431484 |
| Abhd4 | ENSMUSG00000040997 | 105501 | -0.0590462573411057 | 3.54628653613096 | -0.812257268443292 | 0.4308331598998 | 0.530551525431484 |
| Pigq | ENSMUSG00000025728 | 14755 | 0.0620042871637621 | 3.70642075779725 | 0.812222839150337 | 0.4308522363633 | 0.530551525431484 |
| Atf1 | ENSMUSG00000023027 | 11908 | -0.0519335734106621 | 4.84621764968108 | -0.812212917023636 | 0.4308577340824 | 0.530551525431484 |
| Pgpep1 | ENSMUSG00000056204 | 66522 | -0.0877010264455915 | 3.24045913305673 | -0.811596283706016 | 0.4311994914773 | 0.530912552527451 |
| Cherp | ENSMUSG00000052488 | 27967 | 0.0539414506255556 | 4.41795924612253 | 0.811277349243367 | 0.4313763236492 | 0.531070457687753 |
| Sh2b3 | ENSMUSG00000042594 | 16923 | -0.0855364992593382 | 4.03422121257028 | -0.811054778950208 | 0.4314997547908 | 0.531162592299404 |
| Rassf3 | ENSMUSG00000025795 | 192678 | 0.0608626533190977 | 6.46368119873557 | 0.810605026665021 | 0.4317492442536 | 0.531409862508357 |
| Pex1 | ENSMUSG00000005907 | 71382 | -0.0829071239865673 | 2.56492321292784 | -0.810491899624157 | 0.4318120134779 | 0.531427282067888 |
| Ppcdc | ENSMUSG00000063849 | 66812 | -0.148072969192666 | 3.10299577530183 | -0.809905314634717 | 0.4321375782552 | 0.531768081493631 |
| Irx3 | ENSMUSG00000031734 | 16373 | 0.184209400048019 | 1.63481619063523 | 0.809751016675326 | 0.4322232426111 | 0.531793946876669 |
| Pip4p2 | ENSMUSG00000028221 | 72519 | 0.0628075022784951 | 3.84462296032303 | 0.809692203747472 | 0.4322558977220 | 0.531793946876669 |
| Unc13b | ENSMUSG00000028456 | 22249 | 0.0927647262942881 | 2.98059257774381 | 0.809268986626339 | 0.4324909304866 | 0.532023223176817 |
| Tmem167b | ENSMUSG00000068732 | 67495 | 0.063501074322927 | 3.81684382177459 | 0.808995134716497 | 0.4326430575434 | 0.532150473951511 |
| Rpl27 | ENSMUSG00000063316 | 19942 | 0.0820298361583669 | 5.68126257077748 | 0.808486091942321 | 0.4329259268774 | 0.532437923312295 |
| Ssb | ENSMUSG00000068882 | 20823 | 0.058529195969049 | 5.1854961986564 | 0.80839928865087 | 0.4329741743960 | 0.532437923312295 |
| Mmd | ENSMUSG00000003948 | 67468 | 0.181710931175736 | 1.44425256632699 | 0.808238448217841 | 0.4330635828457 | 0.532487966713993 |
| Zfp445 | ENSMUSG00000047036 | 235682 | -0.070864878715355 | 4.72157307045928 | -0.808127621374372 | 0.4331251965261 | 0.532503826704267 |
| Smc5 | ENSMUSG00000024943 | 226026 | 0.048636458832064 | 4.46839459441523 | 0.807946914896255 | 0.4332256715989 | 0.532567455718472 |
| Adamts9 | ENSMUSG00000030022 | 101401 | -0.153799210977601 | 3.52208912671531 | -0.807190459279474 | 0.4336464335984 | 0.533024757187118 |
| Becn1 | ENSMUSG00000035086 | 56208 | -0.0509096625852807 | 4.70205030542428 | -0.80709366790742 | 0.4337002907027 | 0.533031018402525 |
| Ssbp3 | ENSMUSG00000061887 | 72475 | -0.0576010078741471 | 3.54205201715065 | -0.807000968935476 | 0.4337518747370 | 0.53303448485109 |
| Ppp1r2 | ENSMUSG00000047714 | 66849 | -0.056101691288557 | 6.38271370752626 | -0.806694146373015 | 0.4339226399573 | 0.533184394938618 |
| Rgl1 | ENSMUSG00000026482 | 19731 | 0.125431870165559 | 4.02739693762526 | 0.806168863191685 | 0.4342150922287 | 0.533483778032214 |
| Pitpnb | ENSMUSG00000050017 | 56305 | -0.0584134723192635 | 4.35849408507874 | -0.805598872072022 | 0.4345325793417 | 0.533813848593506 |
| Mapre2 | ENSMUSG00000024277 | 212307 | 0.0698392010859958 | 5.10830824585459 | 0.805500049198627 | 0.4345876392429 | 0.533821495062832 |
| Ctr9 | ENSMUSG00000005609 | 22083 | -0.0653947203291896 | 3.89811080793893 | -0.804769864467488 | 0.4349946063368 | 0.53426135301883 |
| Nr3c2 | ENSMUSG00000031618 | 110784 | 0.0886208124449288 | 3.8796493432853 | 0.804672032539497 | 0.4350491514083 | 0.534268315250513 |
| Atp6v1d | ENSMUSG00000021114 | 73834 | -0.0629598044519559 | 5.37388723159502 | -0.804463159725437 | 0.4351656207860 | 0.534351314402678 |
| Med10 | ENSMUSG00000021598 | 28077 | -0.110283875191579 | 2.06357964422886 | -0.804336751481718 | 0.4352361169160 | 0.534377849489963 |
| Ireb2 | ENSMUSG00000032293 | 64602 | -0.0639158624523111 | 4.61577654501322 | -0.804067548886203 | 0.4353862719690 | 0.534455002092609 |
| 2810006K23Rik | ENSMUSG00000047635 | 72650 | -0.105301042490594 | 2.58728201631758 | -0.804048779000155 | 0.4353967426250 | 0.534455002092609 |
| Fdxr | ENSMUSG00000018861 | 14149 | -0.10664270508741 | 1.51064444466728 | -0.80349505082051 | 0.4357057090856 | 0.534774209074303 |
| Secisbp2 | ENSMUSG00000035139 | 75420 | -0.061904576490615 | 3.65664334257928 | -0.803224717030417 | 0.4358565997826 | 0.534864489852367 |
| Pwwp3b | ENSMUSG00000042515 | 245631 | 0.177158895245735 | 1.43697469095877 | 0.803187945327733 | 0.4358771270336 | 0.534864489852367 |
| Zfp746 | ENSMUSG00000057691 | 69228 | 0.0724088601424215 | 2.98282223273821 | 0.801698465064029 | 0.4367091294755 | 0.535825288393485 |
| Zfp626 | ENSMUSG00000030604 | 71163 | -0.0852087788112112 | 2.26882850811272 | -0.801473682566702 | 0.4368347783145 | 0.53590542205143 |
| Osbpl2 | ENSMUSG00000039050 | 228983 | -0.063997582025892 | 3.94159299774501 | -0.801406220912073 | 0.4368724925350 | 0.53590542205143 |
| Pde2a | ENSMUSG00000110195 | 207728 | 0.312842834553809 | 3.80255268732804 | 0.801165236289415 | 0.4370072312864 | 0.53601055264729 |
| Pip4k2b | ENSMUSG00000018547 | 108083 | -0.0624173676375408 | 5.42293612309375 | -0.801008875810576 | 0.4370946694586 | 0.536057649708563 |
| Akt3 | ENSMUSG00000019699 | 23797 | -0.103756585645104 | 3.7472432773276 | -0.800830095049384 | 0.4371946590148 | 0.536110879402007 |
| D430042O09Rik | ENSMUSG00000032743 | 233865 | -0.0877471469711333 | 2.79140739475994 | -0.800755895157096 | 0.4372361622787 | 0.536110879402007 |
| Dazap1 | ENSMUSG00000069565 | 70248 | -0.0693218377587224 | 3.7373372502055 | -0.798948189738122 | 0.4382480726432 | 0.537291350612763 |
| Taf6l | ENSMUSG00000003680 | 225895 | 0.135874041806382 | 1.54845897275896 | 0.798676929951671 | 0.4384000468706 | 0.537417395126512 |
| Glrx2 | ENSMUSG00000018196 | 69367 | -0.0826083884812263 | 3.44281130225295 | -0.798460204576168 | 0.4385214922814 | 0.537505991492286 |
| Arl6ip4 | ENSMUSG00000029404 | 65105 | 0.0702992346535787 | 4.53247474751397 | 0.79814486598881 | 0.4386982356350 | 0.537662340366208 |
| Klhl22 | ENSMUSG00000022750 | 224023 | -0.0844675783018518 | 3.23344224424056 | -0.797635013375259 | 0.4389840982282 | 0.537952374185365 |
| Ciz1 | ENSMUSG00000039205 | 68379 | -0.0483898330594608 | 4.15962132859583 | -0.797489205537603 | 0.4390658712539 | 0.537992269776612 |
| Zfp729b | ENSMUSG00000058093 | 100416706 | -0.113307160132715 | 2.18636227106273 | -0.797036668013687 | 0.4393197288057 | 0.538242989865018 |
| Zbtb7b | ENSMUSG00000028042 | 22724 | -0.0796408501426527 | 3.74154895174107 | -0.796701782110233 | 0.4395076484520 | 0.538377319158872 |
| Ufsp1 | ENSMUSG00000051502 | 70240 | 0.119813525574916 | 1.77196141460189 | 0.79666574334561 | 0.4395278745013 | 0.538377319158872 |
| Dtnb | ENSMUSG00000071454 | 13528 | 0.154945648072836 | 1.53898408438395 | 0.796258193449394 | 0.4397566453473 | 0.53859718658729 |
| Bivm | ENSMUSG00000041684 | 246229 | 0.0992974760150442 | 1.8077005267526 | 0.796047244584978 | 0.4398750876438 | 0.538681893685269 |
| Zfp251 | ENSMUSG00000022526 | 71591 | 0.0873113784240354 | 3.20462697678203 | 0.795661868467974 | 0.4400915190038 | 0.538886568189894 |
| Spindoc | ENSMUSG00000024970 | 68229 | -0.10734014842795 | 2.07275823127145 | -0.795059021490677 | 0.4404302209340 | 0.539240898860874 |
| Usp33 | ENSMUSG00000025437 | 170822 | -0.0685656990851298 | 4.29081656738779 | -0.794616867788741 | 0.4406787453151 | 0.539484753616343 |
| Nbdy | ENSMUSG00000086316 | 70123 | -0.124846755595127 | 1.46278193761614 | -0.7942249034009 | 0.4408991343861 | 0.539694113994965 |
| Prrg3 | ENSMUSG00000033361 | 208748 | 0.140786767826705 | 2.75762016552178 | 0.79380292755119 | 0.4411364766640 | 0.539924177182264 |

| | | | | | | | |
|---|---|---|---|---|---|---|---|
| Ints5 | ENSMUSG00000071652 | 109077 | -0.0807293350941931 | 2.89763590961502 | -0.793358067013121 | 0.4413867789107 | 0.540170049291644 |
| Emsy | ENSMUSG00000035401 | 233545 | -0.0675784886715265 | 4.62640731391711 | -0.793064467246584 | 0.4415520234646 | 0.540311784226086 |
| Dmrt2 | ENSMUSG00000048138 | 226049 | -0.13198255806529 | 1.37494436357195 | -0.792340108612317 | 0.4419598777269 | 0.540750327225577 |
| Tnrc6b | ENSMUSG00000047888 | 213988 | -0.0684628987912266 | 6.44307741804226 | -0.792225990842333 | 0.4420241543066 | 0.5407684066997 |
| Axin2 | ENSMUSG00000000142 | 12006 | -0.0831537407090304 | 3.96943269958542 | -0.792138217683048 | 0.4420735964018 | 0.5407684066997 |
| Marf1 | ENSMUSG00000060657 | 223989 | -0.0522071146264489 | 5.26958573346373 | -0.790687847435865 | 0.4428910921487 | 0.541707791012471 |
| Smim15 | ENSMUSG00000071180 | 75616 | 0.0853911498228275 | 3.05061822452432 | 0.790558048588951 | 0.4429642997416 | 0.541736715202001 |
| Lzts2 | ENSMUSG00000035342 | 226154 | -0.0850700228518333 | 4.58589363968961 | -0.790091766637884 | 0.4432273501310 | 0.541997780991491 |
| Fbxo38 | ENSMUSG00000042211 | 107035 | -0.0580502337312856 | 4.48613387887553 | -0.789901111109899 | 0.4433349360623 | 0.542068700905729 |
| Klf4 | ENSMUSG00000003032 | 16600 | -0.260267670413067 | 4.53568870246873 | -0.78977236135054 | 0.4434075982991 | 0.542096908288549 |
| Supt5 | ENSMUSG00000003435 | 20924 | 0.0637893746755225 | 6.21097746870366 | 0.789594438447524 | 0.4435080247688 | 0.542159049289658 |
| Pacsin3 | ENSMUSG00000027257 | 80708 | 0.0543834913828638 | 6.60961671441553 | 0.789275042512581 | 0.4436883403595 | 0.542292618083704 |
| Rps25 | ENSMUSG00000009927 | 75617 | -0.0632904229066398 | 5.98880015667404 | -0.789225151774377 | 0.4437165104876 | 0.542292618083704 |
| Zfp74 | ENSMUSG00000059975 | 72723 | -0.105407108161589 | 2.45757495099465 | -0.788999762623708 | 0.4438437876084 | 0.542387528490575 |
| Pdxdc1 | ENSMUSG00000022680 | 94184 | -0.0509755070301985 | 6.22385689318259 | -0.788471445988629 | 0.4441422188013 | 0.542691548593496 |
| Dad1 | ENSMUSG00000022174 | 13135 | 0.06452711455661 | 3.47097689647485 | 0.787762255474404 | 0.4445430213076 | 0.543120572919835 |
| Eif2b3 | ENSMUSG00000028683 | 108067 | -0.0660124477170982 | 3.03450163469941 | -0.786260581573504 | 0.4453924588821 | 0.544097560129768 |
| Fbxo21 | ENSMUSG00000032898 | 231670 | -0.0651508050573124 | 3.45723037678856 | -0.786025260331265 | 0.4455256642510 | 0.544199467642039 |
| Pex11b | ENSMUSG00000028102 | 18632 | -0.101448191461933 | 2.34895450508113 | -0.785425758029072 | 0.4458651314440 | 0.544553268359152 |
| Smg8 | ENSMUSG00000020495 | 74133 | -0.0808582855398425 | 2.77626363815015 | -0.784847886097681 | 0.4461925059430 | 0.544892222373333 |
| Plekhg1 | ENSMUSG00000040624 | 213783 | 0.116579813138865 | 3.14821685704877 | 0.784655789082272 | 0.4463013660354 | 0.544964279728972 |
| Zfp975 | ENSMUSG00000069727 | 434179 | -0.166042190578298 | 1.52287538699075 | -0.784467359009485 | 0.4464081644759 | 0.545033803851947 |
| Lrrfip1 | ENSMUSG00000026305 | 16978 | 0.0510183788398688 | 6.70815418905805 | 0.784310927745944 | 0.4464968389468 | 0.545081186791141 |
| Colgalt2 | ENSMUSG00000032649 | 269132 | 0.161327761627771 | 2.23139647464511 | 0.78407347282791 | 0.4466314637695 | 0.54518464885143 |
| Sipa1 | ENSMUSG00000056917 | 20469 | -0.122441565988154 | 3.30347487743862 | -0.783879233459226 | 0.4467416067321 | 0.545252090645643 |
| Brms1l | ENSMUSG00000012076 | 52592 | -0.0592025214656831 | 3.41204000417414 | -0.783800113207859 | 0.4467864766181 | 0.545252090645643 |
| Nr2c2 | ENSMUSG00000005893 | 22026 | 0.068101028084258 | 4.77700312037477 | 0.783652166204782 | 0.4468703865154 | 0.54529361408795 |
| Pgrmc1 | ENSMUSG00000006373 | 53328 | 0.0698781885159949 | 3.05834138862895 | 0.783068048879398 | 0.4472017732274 | 0.545637078150336 |
| Ubap2l | ENSMUSG00000042520 | 74383 | -0.0500306148578478 | 6.10031509163895 | -0.782729709086447 | 0.4473937944709 | 0.545810442785894 |
| A330023F24Rik | ENSMUSG00000096929 | 320977 | -0.147990074924167 | 2.25490487182988 | -0.782615658824772 | 0.4474585342937 | 0.545828505564691 |
| Olfr1031 | ENSMUSG00000043267 | 257916 | -0.263715152910015 | 0.73566252175674 | -0.782203789082005 | 0.4476923786794 | 0.546052822064861 |
| Olfr767 | ENSMUSG00000059762 | 258315 | -0.45477023958686 | 0.5939271924729 | -0.781385519245518 | 0.4481571917188 | 0.546558770799771 |
| Olfr735 | ENSMUSG00000046210 | 257909 | -0.302980682284399 | 0.952814390360111 | -0.781001368055625 | 0.4483755117894 | 0.546764024918667 |
| Hsp90ab1 | ENSMUSG00000023944 | 15516 | 0.057482000168898 | 9.96852931163759 | 0.780681879475141 | 0.4485571342168 | 0.546918622535306 |
| Tmem9 | ENSMUSG00000026411 | 66241 | 0.154393625732135 | 1.20066871879649 | 0.780602340817307 | 0.4486023574834 | 0.546918622535306 |
| Fcf1 | ENSMUSG00000021243 | 73736 | -0.0835680706326425 | 2.33496397180307 | -0.779970600389919 | 0.4489616484062 | 0.547237999576268 |
| Meak7 | ENSMUSG00000034105 | 74347 | -0.120714873028617 | 1.44004239225772 | -0.779965679269731 | 0.4489644479187 | 0.547237999576268 |
| Pphln1 | ENSMUSG00000036167 | 223828 | 0.0707619826639441 | 3.490500881103 | 0.779618498856464 | 0.4491619787776 | 0.54741772661598 |
| Nubp2 | ENSMUSG00000039183 | 26426 | 0.094626022796608 | 2.07946613907471 | 0.779248524414096 | 0.4493725389689 | 0.547613291356657 |
| Hpf1 | ENSMUSG00000038005 | 72612 | -0.131499910272315 | 1.79233908783398 | -0.778951971119262 | 0.4495413587155 | 0.547757952527021 |
| Atp6v0a2 | ENSMUSG00000038023 | 21871 | -0.0486708470151198 | 5.26851747423726 | -0.778534986034234 | 0.4497788048409 | 0.547953917253975 |
| Atg3 | ENSMUSG00000022663 | 67841 | -0.0566388031762397 | 4.26324030450252 | -0.778493482877825 | 0.4498024425505 | 0.547953917253975 |
| Scyl3 | ENSMUSG00000026584 | 240880 | 0.10693310730209 | 2.80683110549288 | 0.77832372930421 | 0.4498991321822 | 0.548010632258128 |
| 2810025M15Rik | ENSMUSG00000049881 | 69953 | 0.130211910719191 | 1.36401729281092 | 0.778149507026867 | 0.4499983807934 | 0.548027845233342 |
| Wiz | ENSMUSG00000024050 | 22404 | 0.0757271842043186 | 3.71141128204676 | 0.778122909246831 | 0.4500135338774 | 0.548027845233342 |
| Pdcd2 | ENSMUSG00000014771 | 18567 | -0.119531297247399 | 2.13060527450068 | -0.777989591111168 | 0.4500894917258 | 0.548059288632652 |
| Kat5 | ENSMUSG00000024926 | 81601 | 0.0837680859570571 | 3.16935002636997 | 0.776872158167997 | 0.4507264662702 | 0.548773780663869 |
| Tsr2 | ENSMUSG00000025264 | 69499 | 0.0769448070148631 | 2.3998899270459 | 0.776570826141481 | 0.4508983331044 | 0.548921893213538 |
| Zfp39 | ENSMUSG00000037001 | 22698 | -0.119170026741577 | 2.42360523965359 | -0.77596236517905 | 0.45124549916 | 0.549267321680193 |
| Lrrc41 | ENSMUSG00000028703 | 230654 | 0.0580362279449753 | 3.40791982623159 | 0.775897403442739 | 0.4512825739648 | 0.549267321680193 |
| Snx8 | ENSMUSG00000029560 | 231834 | 0.091017850855406 | 2.23252002737786 | 0.775663204703334 | 0.4514162512557 | 0.549368853537826 |
| Cdkn2aip | ENSMUSG00000038069 | 70925 | -0.0743527248312676 | 2.68946071204387 | -0.774965414388223 | 0.4518146880505 | 0.549792536466763 |
| Samd1 | ENSMUSG00000079003 | 666704 | 0.0650217278059131 | 3.14158782071693 | 0.77478275574536 | 0.4519190223589 | 0.549858285107513 |
| Fam120b | ENSMUSG00000014763 | 67544 | -0.0662055320850006 | 3.93665983047122 | -0.774622065452633 | 0.4520108209407 | 0.54990876835873 |
| D630003M21Rik | ENSMUSG00000037813 | 228846 | 0.208224519219336 | 1.39262186253051 | 0.774365519267478 | 0.4521574040990 | 0.550025882952061 |
| Spen | ENSMUSG00000040761 | 56381 | 0.0608566308691998 | 6.09190938757947 | 0.77365134348847 | 0.4525656214880 | 0.550461200454616 |
| Zfp26 | ENSMUSG00000063108 | 22688 | 0.0642328489972193 | 3.18052575407453 | 0.773455188977366 | 0.4526777825278 | 0.550536364131285 |
| Gm33989 | ENSMUSG00000108621 | 102637087 | 0.0802473055879611 | 3.74874351520101 | 0.772620369360802 | 0.4531553276999 | 0.551055833472938 |
| Synrg | ENSMUSG00000034940 | 217030 | 0.0590847689248868 | 5.42355850150415 | 0.772497592373063 | 0.4532255870469 | 0.551079965740822 |
| Casd1 | ENSMUSG00000015189 | 213819 | -0.0547282225763586 | 3.94073200746124 | -0.772277014248635 | 0.4533518304891 | 0.551172156498378 |
| Polr3b | ENSMUSG00000034453 | 70428 | 0.083079940738767 | 3.49976395368057 | 0.771971461521751 | 0.4535267439866 | 0.551323491828046 |
| Thnsl1 | ENSMUSG00000048550 | 208967 | -0.151195479911041 | 1.45895680353734 | -0.771164384127353 | 0.4539889591683 | 0.551824008970192 |
| Pgap2 | ENSMUSG00000030990 | 233575 | 0.0855588504566724 | 3.14852006782072 | 0.770737835888494 | 0.4542333639855 | 0.552059695544367 |
| Trappc4 | ENSMUSG00000032112 | 60409 | -0.128832434179707 | 2.09807215191286 | -0.770612140990435 | 0.4543054007791 | 0.552085862803392 |
| Tbx2 | ENSMUSG00000000093 | 21385 | -0.169044706367537 | 1.8140507989429 | -0.770373214605005 | 0.4544423512780 | 0.552190900602557 |
| Pogk | ENSMUSG00000040596 | 71592 | 0.0655722627497779 | 3.15525677857626 | 0.769806157581964 | 0.4547674872106 | 0.552524552928756 |
| Sumf1 | ENSMUSG00000030101 | 58911 | 0.0916227930958118 | 3.05898908738785 | 0.76928442912709 | 0.4550667619501 | 0.55282671425616 |
| Fam193b | ENSMUSG00000021495 | 212483 | 0.106946652251608 | 3.21348079222449 | 0.769073511977112 | 0.4551877836925 | 0.552912286203235 |
| Dcun1d4 | ENSMUSG00000051674 | 100737 | -0.0741118376938532 | 2.24510797199294 | -0.768403984257154 | 0.4555720845685 | 0.553317606268789 |
| Ing4 | ENSMUSG00000030330 | 28019 | -0.107302905459498 | 3.06552896094175 | -0.767932724815661 | 0.4558427038942 | 0.553584779054314 |
| Lepr | ENSMUSG00000057722 | 16847 | 0.176611634365266 | 1.40564810870059 | 0.767741401636706 | 0.4559525994349 | 0.553656727885241 |
| Sema4c | ENSMUSG00000026121 | 20353 | 0.138411609645603 | 2.85626365373704 | 0.767448720255224 | 0.4561207470191 | 0.553799387500467 |
| Ift20 | ENSMUSG00000001105 | 55978 | 0.0693139288551654 | 3.02697213997951 | 0.766482975614981 | 0.4566758501413 | 0.554364118009813 |
| Bcar3 | ENSMUSG00000028121 | 29815 | 0.28660200969583 | 1.95777566870146 | 0.766463062373862 | 0.4566873005835 | 0.554364118009813 |
| Utp3 | ENSMUSG00000070697 | 65961 | -0.0542673362879809 | 4.56671376949997 | -0.765901280861287 | 0.4570104083518 | 0.554647493308362 |
| Yod1 | ENSMUSG00000046404 | 226418 | -0.10741312930565 | 2.16458399435112 | -0.765880735300035 | 0.4570222278134 | 0.554647493308362 |
| Sugp1 | ENSMUSG00000011306 | 70616 | -0.0741468173770601 | 3.6427020606603 | -0.765707344536244 | 0.4571219837726 | 0.554706972093593 |
| Map2 | ENSMUSG00000015222 | 17756 | 0.104793897408305 | 1.86730846331044 | 0.765454741696823 | 0.4572673367862 | 0.554821762505348 |
| Casp8 | ENSMUSG00000026029 | 12370 | 0.127886079820421 | 2.36502522159183 | 0.765291281455951 | 0.4573614106870 | 0.554874315229738 |
| Stxbp1 | ENSMUSG00000026797 | 20910 | 0.152924210257048 | 1.87052945776764 | 0.764540764247704 | 0.4577935004979 | 0.555336894234017 |
| Tmem115 | ENSMUSG00000010045 | 56395 | 0.0688316230432164 | 2.87740954852817 | 0.76427435026046 | 0.4579469425027 | 0.555412189713705 |
| Ddx17 | ENSMUSG00000055065 | 67040 | 0.0984016689343115 | 5.88502495739444 | 0.76425655651885 | 0.4579571920125 | 0.555412189713705 |
| Bach1 | ENSMUSG00000025612 | 12013 | -0.0801538943635576 | 4.80684007729729 | -0.764161135222281 | 0.4580121588152 | 0.555417229643866 |
| Glce | ENSMUSG00000032252 | 93683 | 0.0794985713287102 | 3.07979254619245 | 0.763689369661262 | 0.4582839768698 | 0.555685208115839 |
| Gtf2ird1 | ENSMUSG00000023079 | 57080 | -0.0788021911440064 | 3.43459788468994 | -0.763357294561595 | 0.4584753696975 | 0.555855619583441 |
| Tomm22 | ENSMUSG00000022427 | 223696 | -0.0777908167007615 | 3.9655218702812 | -0.762374420462464 | 0.4590421459676 | 0.556481057732294 |
| Srp9 | ENSMUSG00000026511 | 27058 | 0.0655443279205956 | 4.99729372464969 | 0.76108112310199 | 0.4597885950980 | 0.557262810894379 |
| Rnf216 | ENSMUSG00000045078 | 108086 | 0.0560669630427508 | 4.44570785757069 | 0.761027263127699 | 0.4598196977486 | 0.557262810894379 |
| Cpox | ENSMUSG00000022742 | 12892 | -0.0707505194146836 | 3.86036395950021 | -0.760992184208584 | 0.4598399555637 | 0.557262810894379 |
| Kdm5a | ENSMUSG00000030180 | 214899 | -0.064148850447812 | 5.0957507904905 | -0.760213040026509 | 0.4602900491197 | 0.557746427993327 |
| Zcchc2 | ENSMUSG00000038866 | 227449 | 0.0698392758988131 | 3.61042743340671 | 0.759795161325557 | 0.4605315611622 | 0.557977221798343 |
| Unc50 | ENSMUSG00000026111 | 67387 | -0.114031162792733 | 2.24347160314401 | -0.75938759761376 | 0.4607671877907 | 0.558200834505242 |
| Polr2l | ENSMUSG00000038489 | 66491 | 0.0514791408775901 | 4.58854830505261 | 0.758721189996424 | 0.4611526228856 | 0.558605864446286 |
| Ddi2 | ENSMUSG00000078515 | 68817 | 0.0487309664474989 | 5.63978788305318 | 0.758509221319282 | 0.4612752628807 | 0.55863275694094 |
| Rbm27 | ENSMUSG00000024491 | 225432 | 0.0483878194192524 | 4.69657003615715 | 0.758506159538243 | 0.4612770345026 | 0.55863275694094 |
| Pura | ENSMUSG00000043991 | 19290 | 0.049254780620462 | 5.19143102493327 | 0.75826579582348 | 0.4614161281023 | 0.558691382879145 |
| Rn7sk | ENSMUSG00000065037 | 19817 | -0.0803071653862837 | 11.688224566108 | -0.758245861498585 | 0.4614276648644 | 0.558691382879145 |
| Abhd17b | ENSMUSG00000047368 | 226016 | 0.0712059821560054 | 2.89619368110308 | 0.757634540571846 | 0.4617815470847 | 0.5590166305297 |
| Usp42 | ENSMUSG00000051306 | 76800 | 0.09336076527708 | 3.05007508425668 | 0.757524503101861 | 0.4618452636550 | 0.5590166305297 |
| Prpf40a | ENSMUSG00000061136 | 56194 | 0.0547791861724791 | 5.10634008265954 | 0.757516823062171 | 0.4618497109426 | 0.5590166305297 |
| Npm3 | ENSMUSG00000056209 | 18150 | 0.0977017985604376 | 2.19107029611653 | 0.757241717732062 | 0.4620090340192 | 0.55912248701726 |
| Psmd2 | ENSMUSG00000006998 | 21762 | 0.0464796291699789 | 7.11415917361592 | 0.757189173598433 | 0.4620394680474 | 0.55912248701726 |
| Rpl7 | ENSMUSG00000043716 | 19989 | -0.0546856723074605 | 7.30842477198593 | -0.756464567513247 | 0.4624592935645 | 0.559568578476216 |
| Tpm3 | ENSMUSG00000027940 | 59069 | 0.0965361236221334 | 11.8113675157377 | 0.755986515231978 | 0.4627363995244 | 0.559841901848513 |
| Wdr83 | ENSMUSG00000005150 | 67836 | -0.0856081917775446 | 2.92082825409301 | -0.75550610351007 | 0.4630149771500 | 0.560116945022897 |
| Zfp740 | ENSMUSG00000046897 | 68744 | 0.0523473741939586 | 4.26807745778045 | 0.754902208101566 | 0.4633653074550 | 0.560478718135555 |
| Cep44 | ENSMUSG00000038215 | 382010 | -0.103317826980269 | 1.60587912381888 | -0.754069868109591 | 0.4638484323355 | 0.561001019457799 |
| Heatr6 | ENSMUSG00000000976 | 217026 | 0.061292436032645 | 3.29282962256772 | 0.752889366330218 | 0.4645341810680 | 0.561748515220107 |
| Actr6 | ENSMUSG00000019948 | 67019 | 0.122835641499542 | 1.93720388546989 | 0.752757954029802 | 0.4646105568371 | 0.561748515220107 |
| Hmces | ENSMUSG00000030060 | 232210 | 0.15299327996789 | 1.56960495484306 | 0.752740588565936 | 0.4646206500873 | 0.561748515220107 |
| Kifc3 | ENSMUSG00000031788 | 16582 | -0.127791021297925 | 2.88543043243333 | -0.751672273622595 | 0.4652418435196 | 0.562383049312688 |
| Limd1 | ENSMUSG00000025239 | 29806 | -0.0500047706314595 | 5.69773613639022 | -0.751661056977903 | 0.4652483683958 | 0.562383049312688 |
| Rbm8a | ENSMUSG00000038374 | 60365 | 0.0591040179001405 | 3.40564679034394 | 0.751280132083171 | 0.4654699913074 | 0.562571305323558 |
| Twistnb | ENSMUSG00000020561 | 28071 | -0.0692383600064685 | 3.24493609298055 | -0.751216463472704 | 0.4655070402206 | 0.562571305323558 |
| Trappc6b | ENSMUSG00000020993 | 78232 | -0.0662817104783597 | 3.54681516024254 | -0.750811773951483 | 0.4657425727161 | 0.562744650115306 |
| Pacs2 | ENSMUSG00000021143 | 217893 | 0.0523110016110451 | 5.31409408086029 | 0.750793105435021 | 0.4657534397212 | 0.562744650115306 |

| | | | | | | | |
|---|---|---|---|---|---|---|---|
| Dgcr2 | ENSMUSG00000003166 | 13356 | -0.0510385933092055 | 4.26635828776255 | -0.749849090301623 | 0.4663031588112 | 0.563346577029756 |
| Plekha2 | ENSMUSG00000031557 | 83436 | 0.0760106612970102 | 3.37968613132892 | 0.749552215777806 | 0.4664761178338 | 0.563493252739758 |
| Bbs9 | ENSMUSG00000035919 | 319845 | -0.132294928222975 | 1.95840559296671 | -0.749055148899703 | 0.4667657977503 | 0.563780876818705 |
| Luzp1 | ENSMUSG00000001089 | 269593 | -0.0740427055146231 | 4.99312501313043 | -0.748628709905865 | 0.4670144059275 | 0.564018834514877 |
| Numb | ENSMUSG00000021224 | 18222 | -0.0845161213615606 | 3.66418655808674 | -0.748377369234063 | 0.4671609724985 | 0.564133516392132 |
| Nop16 | ENSMUSG00000025869 | 28126 | -0.0741262366848314 | 2.90126050013489 | -0.747767341096829 | 0.4675168220265 | 0.564500870603384 |
| Rbm26 | ENSMUSG00000022119 | 74213 | -0.0533387368460767 | 4.45600993396552 | -0.747190041438054 | 0.4678537342049 | 0.56484528038371 |
| Med17 | ENSMUSG00000031935 | 234959 | 0.0837722363935631 | 3.01703976477448 | 0.746489035933862 | 0.4682630427336 | 0.565277009400541 |
| Gpatch3 | ENSMUSG00000028850 | 242691 | 0.128330456119321 | 1.30692321946553 | 0.746278439648581 | 0.4683860505059 | 0.565363065159052 |
| Ints6 | ENSMUSG00000035161 | 18130 | 0.0667159469705436 | 3.31779857399439 | 0.744815343468892 | 0.4692411850183 | 0.566332714302234 |
| Hoxa7 | ENSMUSG00000038236 | 15404 | -0.0870800325395074 | 2.271343379958 | -0.743971523749741 | 0.4697348090785 | 0.566865886292494 |
| Map2k1 | ENSMUSG00000004936 | 26395 | -0.0658451751798603 | 4.45099469164712 | -0.743876496892931 | 0.4697904186538 | 0.566870412441284 |
| Zscan20 | ENSMUSG00000061894 | 269585 | 0.105213949845098 | 1.72844363623012 | 0.7436658485588 | 0.4699137042096 | 0.566956589482914 |
| 1700037H04Rik | ENSMUSG00000027327 | 67326 | 0.137575555632077 | 1.211869110815 | 0.743294416266127 | 0.4701311398986 | 0.567156328245496 |
| Rtn3 | ENSMUSG00000024758 | 20168 | 0.0636023625256774 | 4.33289517023093 | 0.743126690096278 | 0.4702293467571 | 0.567212203641883 |
| Ccnc | ENSMUSG00000028252 | 51813 | 0.0854638039659497 | 2.62079097995628 | 0.743011061586024 | 0.4702970568036 | 0.567231284113447 |
| Kmt5c | ENSMUSG00000059851 | 232811 | -0.111194011303844 | 1.97692526270469 | -0.742631514239565 | 0.4705193553333 | 0.56743679094756 |
| Atg4c | ENSMUSG00000028550 | 242557 | -0.0992035139092695 | 2.49831491732879 | -0.741652217401041 | 0.4710932221403 | 0.568066189874959 |
| Exoc4 | ENSMUSG00000029763 | 20336 | -0.041467064612589 | 5.54898070115287 | -0.74138688686334 | 0.4712487796383 | 0.568161586221026 |
| Dhrs1 | ENSMUSG00000002332 | 52585 | 0.0730027824207116 | 3.45303687083759 | 0.741339972435013 | 0.4712762878296 | 0.568161586221026 |
| Mtfmt | ENSMUSG00000059183 | 69606 | 0.100253272477227 | 2.17740138500421 | 0.74121503835468 | 0.4713495475214 | 0.56818724128333 |
| Atp2a2 | ENSMUSG00000029467 | 11938 | -0.0733764816943197 | 11.5698724940701 | -0.74070052513796 | 0.4716513250713 | 0.568488326646294 |
| Tstd2 | ENSMUSG00000035495 | 272027 | 0.0651889930906009 | 3.41378563091925 | 0.739345007367878 | 0.4724469456798 | 0.569334948481835 |
| 2610002M06Rik | ENSMUSG00000031242 | 67028 | -0.102615361906389 | 1.87918448551592 | -0.739326352032062 | 0.4724579011690 | 0.569334948481835 |
| Rcbtb2 | ENSMUSG00000022106 | 105670 | 0.0568183431250211 | 3.36526258882391 | 0.73857280377107 | 0.4729005585837 | 0.569805556203626 |
| Arid4a | ENSMUSG00000048118 | 238247 | -0.0543566899182846 | 4.12176284247876 | -0.738322508307883 | 0.4730476461355 | 0.569919962516044 |
| Pnkp | ENSMUSG00000002963 | 59047 | 0.0812156194852403 | 3.18379888231357 | 0.738091940134558 | 0.4731831656726 | 0.570020408195659 |
| Ccnt1 | ENSMUSG00000011960 | 12455 | 0.0569191909605964 | 4.19606173461083 | 0.737800873053081 | 0.4733542782658 | 0.570126000941156 |
| Fam32a | ENSMUSG00000003039 | 67922 | 0.073166823479669 | 3.01890006368966 | 0.737765402117564 | 0.4733751335231 | 0.570126000941156 |
| Orc4 | ENSMUSG00000026761 | 26428 | -0.0769704535004818 | 3.64501559878834 | -0.736706706803848 | 0.4739978564450 | 0.57081310662124 |
| Mpp3 | ENSMUSG00000052373 | 13384 | -0.162211677704944 | 1.91682823330708 | -0.736295315920954 | 0.4742399713160 | 0.571041763213895 |
| Cnot6l | ENSMUSG00000034724 | 231464 | 0.0642765943851824 | 4.64407748106947 | 0.735467464054022 | 0.4747274145484 | 0.571565741648779 |
| Casp8ap2 | ENSMUSG00000028282 | 26885 | -0.0734630744518721 | 3.67822897161153 | -0.733764458603462 | 0.4757311166107 | 0.572711105250244 |
| Ddx47 | ENSMUSG00000030204 | 67755 | -0.0642940405005992 | 3.47337579109573 | -0.733462257060224 | 0.4759093609299 | 0.57286259490425 |
| Bcl2l1 | ENSMUSG00000007659 | 12048 | -0.130132981170142 | 3.88322626451074 | -0.733226637114237 | 0.4760483624568 | 0.572966818984336 |
| Prpf8 | ENSMUSG00000020850 | 192159 | 0.0482910804884327 | 6.96405609001636 | 0.732846010708667 | 0.4762729614467 | 0.57316590763871 |
| Rnf20 | ENSMUSG00000028309 | 109331 | 0.0463015894687846 | 5.4408961221608 | 0.732728027629047 | 0.4763425937245 | 0.57316590763871 |
| Mphosph8 | ENSMUSG00000079184 | 75339 | -0.0701941609180285 | 3.75143418653978 | -0.73259960487422 | 0.4764183944502 | 0.57316590763871 |
| Slc9b2 | ENSMUSG00000037994 | 97086 | -0.185392191409957 | 1.84925093931601 | -0.732590930887343 | 0.4764235144822 | 0.57316590763871 |
| Rbak | ENSMUSG00000061898 | 57782 | 0.131956651579934 | 1.716395410616 | 0.732292879659041 | 0.4765994669017 | 0.573314490228106 |
| Iqcb1 | ENSMUSG00000022837 | 320299 | -0.097401963955128 | 1.59723763460789 | -0.732073538885483 | 0.4767289785141 | 0.573407181353256 |
| Asb3 | ENSMUSG00000020305 | 65257 | -0.090711970514235 | 1.78227365840355 | -0.731773897533199 | 0.4769059390128 | 0.573504847507029 |
| Cct2 | ENSMUSG00000034024 | 12461 | -0.0480666448022742 | 6.0444776233735 | -0.731758371007887 | 0.4769151096733 | 0.573504847507029 |
| Akap17b | ENSMUSG00000059708 | 338351 | 0.128849240898749 | 2.17660698309142 | 0.730507315128934 | 0.4776543930175 | 0.574330675401512 |
| Phrf1 | ENSMUSG00000038611 | 101471 | -0.0555857101032924 | 4.57013485360218 | -0.730066784519496 | 0.4779148810015 | 0.574580682383232 |
| Usp39 | ENSMUSG00000056305 | 28035 | -0.068798362628733 | 3.42478762770854 | -0.72981010414678 | 0.4780666973380 | 0.574658406857224 |
| Ngdn | ENSMUSG00000022204 | 68966 | 0.0707593997277298 | 3.12789473901859 | 0.729779716784279 | 0.4780846722129 | 0.574658406857224 |
| Mapk3 | ENSMUSG00000063065 | 26417 | 0.0658679417642096 | 3.80606004433544 | 0.729621305678204 | 0.4781783829634 | 0.574707850926118 |
| Vps9d1 | ENSMUSG00000001062 | 72325 | -0.0554199980449988 | 3.48157590441704 | -0.729437950127812 | 0.4782868639971 | 0.574775034119683 |
| Trim21 | ENSMUSG00000030966 | 20821 | -0.176225346864863 | 1.17532574611951 | -0.729183832742128 | 0.4784372355729 | 0.574892538424537 |
| Impact | ENSMUSG00000024423 | 16210 | 0.109908573378345 | 3.18545779254195 | 0.728896684586652 | 0.4786071874580 | 0.575033542108545 |
| Ap2s1 | ENSMUSG00000008036 | 232910 | 0.0782936025534622 | 3.23007424735264 | 0.728474621079708 | 0.4788570572645 | 0.575270523459613 |
| Pggt1b | ENSMUSG00000024477 | 225467 | -0.0621023103136755 | 4.70788693875799 | -0.727204811046326 | 0.4796092883767 | 0.57611089354351 |
| Bola1 | ENSMUSG00000015943 | 69168 | -0.0952995720756893 | 2.58290198336583 | -0.726714518968949 | 0.4798999280061 | 0.576396672127824 |
| Hnrnph2 | ENSMUSG00000045427 | 56258 | 0.0546793283820382 | 4.4923595739955 | 0.726280392623355 | 0.4801573626071 | 0.576642510509656 |
| Yrdc | ENSMUSG00000028889 | 230734 | 0.119776096122318 | 2.14055543668102 | 0.726179630306235 | 0.4802171261246 | 0.576650928888143 |
| Gypc | ENSMUSG00000090523 | 71683 | 0.0971889202752525 | 3.30459259298177 | 0.725638901752997 | 0.4805379169054 | 0.576972142272132 |
| St7 | ENSMUSG00000029534 | 64213 | -0.0801487042257243 | 2.31328953899619 | -0.725550812196071 | 0.4805901889477 | 0.576972142272132 |
| Trip12 | ENSMUSG00000026219 | 14897 | -0.0469545311158148 | 7.37043734281823 | -0.72538220912516 | 0.4806902470678 | 0.577028892016106 |
| Cpsf2 | ENSMUSG00000041781 | 51786 | -0.0552207145794403 | 4.33137968522909 | -0.725074249138504 | 0.4808730397332 | 0.577184934371845 |
| Atf7 | ENSMUSG00000099083 | 223922 | 0.0539286471014382 | 5.10078732567893 | 0.72437395850106 | 0.4812888612150 | 0.577519705271759 |
| Cops7b | ENSMUSG00000026240 | 26895 | -0.0737488971659408 | 2.60591006152303 | -0.724359020662408 | 0.4812977334439 | 0.577519705271759 |
| Sgms1 | ENSMUSG00000040451 | 208449 | -0.060735230874509 | 4.75538263482528 | -0.724337612710471 | 0.4813104487261 | 0.577519705271759 |
| Banp | ENSMUSG00000025316 | 53325 | -0.142953688913346 | 2.3926519043887 | -0.724069680683799 | 0.4814696045633 | 0.57761350324527 |
| Actn3 | ENSMUSG00000006457 | 11474 | 0.183929684937739 | 7.22909459382894 | 0.724028104359593 | 0.4814943044159 | 0.57761350324527 |
| Trim39 | ENSMUSG00000045409 | 79263 | -0.0914106409579034 | 2.62725966291581 | -0.723810301442147 | 0.4816237103330 | 0.577705341561609 |
| Entpd5 | ENSMUSG00000021236 | 12499 | -0.0801035879259708 | 3.65150737244441 | -0.723379133511239 | 0.4818799476471 | 0.577877156199874 |
| Fkbp1a | ENSMUSG00000032966 | 14225 | 0.04106713591045 | 5.3427039815046 | 0.72336514766815 | 0.4818882606303 | 0.577877156199874 |
| Fam120a | ENSMUSG00000038014 | 218236 | -0.0450704668809836 | 6.76927573681409 | -0.723302418264587 | 0.4819255471519 | 0.577877156199874 |
| Ubc | ENSMUSG00000008348 | 22190 | -0.121603937793953 | 9.76302698589916 | -0.722977364913784 | 0.4821187877379 | 0.578045461090652 |
| Lrrc75a | ENSMUSG00000046417 | 192976 | 0.165729690478145 | 0.885065663991601 | 0.722488851418091 | 0.4824092918992 | 0.578293590659632 |
| Flvcr1 | ENSMUSG00000066595 | 226844 | -0.157906643179141 | 3.20498932489239 | -0.722451432470737 | 0.4824315481863 | 0.578293590659632 |
| Erlin2 | ENSMUSG00000031483 | 244373 | 0.0848638482953628 | 3.25782666361724 | 0.721855766787834 | 0.4827859259275 | 0.578654929420445 |
| Ift27 | ENSMUSG00000016637 | 67042 | -0.110649356689063 | 2.16158677370325 | -0.721496893342695 | 0.4829995056221 | 0.578847450493996 |
| Hist1h4i | ENSMUSG00000060639 | 319158 | -0.124508150685719 | 1.52757259335698 | -0.721251288034055 | 0.4831457079729 | 0.578959190293006 |
| Dgat1 | ENSMUSG00000022555 | 13350 | 0.0999494785696609 | 1.6362786532053 | 0.721034872374313 | 0.4832745567034 | 0.579050112827537 |
| Otub1 | ENSMUSG00000024767 | 107260 | -0.06093570746423 | 4.45794208659281 | -0.720901362125763 | 0.4833540559008 | 0.579081892267931 |
| Ccar1 | ENSMUSG00000020074 | 67500 | 0.0390011107972548 | 5.65998168081708 | 0.72054272256045 | 0.4835676484172 | 0.579274297517662 |
| Tcf7l1 | ENSMUSG00000055799 | 21415 | -0.108129777761157 | 2.31514440467026 | -0.720277887046232 | 0.4837254113529 | 0.579399788680633 |
| Nfya | ENSMUSG00000023994 | 18044 | 0.140663285245382 | 3.07288914750118 | 0.720157757850396 | 0.4837969827708 | 0.579422024615812 |
| Lasp1 | ENSMUSG00000038366 | 16796 | 0.121874796168687 | 4.45387945862857 | 0.719902288139516 | 0.4839492096107 | 0.579540842490649 |
| Trp53rkb | ENSMUSG00000042854 | 76367 | -0.121463426827714 | 1.36567634736148 | -0.719743962832213 | 0.4840435655151 | 0.579590340086089 |
| Ythdc1 | ENSMUSG00000035851 | 231386 | -0.0420171474138717 | 5.4124749638226 | -0.719421040322409 | 0.4842360496213 | 0.579757311983689 |
| Thsd4 | ENSMUSG00000032289 | 207596 | 0.128392717298428 | 1.4367885574669 | 0.719213677009687 | 0.4843596768670 | 0.579841816650243 |
| Map10 | ENSMUSG00000050930 | 74393 | 0.117237106107373 | 1.53253849521594 | 0.718581137513743 | 0.4847369063414 | 0.580229864566151 |
| 1110059E24Rik | ENSMUSG00000035171 | 66206 | -0.126291655570623 | 1.52475815986179 | -0.717592274924747 | 0.4853269928328 | 0.580872589363413 |
| Taf2 | ENSMUSG00000037343 | 319944 | 0.053556844363543 | 4.22556305018116 | 0.717429263649583 | 0.4854243085910 | 0.580925456230373 |
| Whamm | ENSMUSG00000045795 | 434204 | -0.096952480078031 | 3.0869378197983 | -0.717268806827386 | 0.4855201108747 | 0.580976500489494 |
| Pik3c2a | ENSMUSG00000030660 | 18704 | -0.0842270121495456 | 3.66435109849263 | -0.717164775649597 | 0.4855822297842 | 0.580987232242896 |
| Prpf19 | ENSMUSG00000024735 | 28000 | -0.0633004412353939 | 6.63785196022321 | -0.716553558488708 | 0.4859472955407 | 0.581360390451553 |
| Ahr | ENSMUSG00000019256 | 11622 | -0.121383236826328 | 2.45123113341847 | -0.715966509438673 | 0.4862980820760 | 0.58171638598966 |
| Nsrp1 | ENSMUSG00000037958 | 237859 | 0.0790834494756778 | 4.25165791759689 | 0.714583845103705 | 0.4871248852758 | 0.582641658928809 |
| Plekha3 | ENSMUSG00000002733 | 83435 | 0.0762700715321607 | 2.98027449772968 | 0.714319934124371 | 0.4872827944783 | 0.582766764381068 |
| Gpatch1 | ENSMUSG00000063808 | 67471 | 0.0806210887112617 | 3.41041071450977 | 0.713957533045802 | 0.4874996847276 | 0.582962373236815 |
| Zkscan1 | ENSMUSG00000029729 | 74570 | 0.0580289705249433 | 4.95508304810815 | 0.713624049539678 | 0.4876993196859 | 0.583137307316419 |
| Zfp973 | ENSMUSG00000078879 | 545490 | -0.141110664718552 | 1.99979526454717 | -0.712910453578151 | 0.4881266685698 | 0.583584448664187 |
| Ston1 | ENSMUSG00000033855 | 77057 | -0.131478933333464 | 2.74624663963447 | -0.712572169602844 | 0.4883293342604 | 0.583762899475155 |
| Proser1 | ENSMUSG00000049504 | 212127 | 0.0858852827084622 | 2.89901776370696 | 0.712281175282125 | 0.4885037092270 | 0.583907495413998 |
| Daxx | ENSMUSG00000002307 | 13163 | -0.0658955267127039 | 3.48715094685717 | -0.711726484440425 | 0.4888362049327 | 0.584241040467977 |
| Tollip | ENSMUSG00000025139 | 54473 | -0.0428196024073895 | 4.73334290498407 | -0.711579320695978 | 0.4889244414200 | 0.58428261388028 |
| Sox9 | ENSMUSG00000000567 | 20682 | 0.16813008123704 | 4.25851870422163 | 0.711324410187441 | 0.4890773033892 | 0.584401399578966 |
| Zfp426 | ENSMUSG00000059475 | 235028 | 0.0913119367902135 | 2.46209969306722 | 0.711100076989787 | 0.4892118528347 | 0.584498279958116 |
| Tmem255b | ENSMUSG00000038457 | 272465 | -0.174885107138655 | 1.15967041465873 | -0.710523225585697 | 0.4895579359510 | 0.584847846762907 |
| Mospd2 | ENSMUSG00000061778 | 76763 | -0.0799176951081404 | 2.76854285662732 | -0.710209293488917 | 0.4897463418616 | 0.585008989497244 |
| Dst | ENSMUSG00000026131 | 13518 | 0.0714836548967916 | 10.3914546533923 | 0.709949030565712 | 0.4899025712797 | 0.585131665937363 |
| Zcchc7 | ENSMUSG00000035649 | 319885 | 0.0736047193700502 | 3.83142002343481 | 0.709694815167684 | 0.4900551993827 | 0.58525001468954 |
| Mknk2 | ENSMUSG00000020190 | 17347 | -0.0580937807295179 | 7.72641096339296 | -0.709518458738381 | 0.4901610985819 | 0.585312537535395 |
| Jrk | ENSMUSG00000046380 | 16469 | 0.148273893938973 | 1.51080803789311 | 0.708946875706582 | 0.4905044193204 | 0.585658526225243 |
| Pknox1 | ENSMUSG00000006705 | 18771 | 0.0601232955650834 | 3.19716983889029 | 0.708009589373451 | 0.4910677112819 | 0.58624569499741 |
| Arnt | ENSMUSG00000015522 | 11863 | -0.0474637013723225 | 5.63416956209803 | -0.707931885097236 | 0.4911144275223 | 0.58624569499741 |
| Per1 | ENSMUSG00000020893 | 18626 | 0.235520130663581 | 4.51845494892894 | 0.707860938976849 | 0.4911570830469 | 0.58624569499741 |
| Rel | ENSMUSG00000020275 | 19696 | -0.10656903376764 | 2.73011300947651 | -0.707525992608166 | 0.4913584955986 | 0.586422067407877 |
| Gpalpp1 | ENSMUSG00000022008 | 67467 | 0.0565390482470316 | 3.55001252011018 | 0.706780319170278 | 0.4918070673189 | 0.586893346382477 |
| Bmpr1a | ENSMUSG00000021796 | 12166 | -0.0522317226352671 | 5.90526952715219 | -0.706492785874138 | 0.4919801030005 | 0.587035750016278 |
| Snapc4 | ENSMUSG00000036281 | 227644 | -0.111254957583583 | 2.36451820437736 | -0.706253879850539 | 0.4921239028060 | 0.587143241821926 |
| Clcn4 | ENSMUSG00000000605 | 12727 | -0.0433087205753617 | 4.99106145259427 | -0.705857461790174 | 0.4923625660674 | 0.587355523625411 |

| | | | | | | | |
|---|---|---|---|---|---|---|---|
| Zfp965 | ENSMUSG00000078896 | 100503949 | -0.116674863082798 | 2.29557380643124 | -0.705779856546523 | 0.4924092963592 | 0.587355523625411 |
| Pomt1 | ENSMUSG00000039254 | 99011 | -0.077276575604087 | 2.40953242587007 | -0.704824930913298 | 0.4929845255003 | 0.587977506627917 |
| Fam120c | ENSMUSG00000025262 | 207375 | 0.0612696158204766 | 4.15494531927216 | 0.704140501530655 | 0.4933970597278 | 0.588405330556908 |
| Rad50 | ENSMUSG00000020380 | 19360 | 0.0685140362911278 | 3.48416901966507 | 0.703698856682572 | 0.4936633669538 | 0.588622055335422 |
| Jdp2 | ENSMUSG00000034271 | 81703 | 0.0940628928312202 | 2.14132109534633 | 0.70366051891582 | 0.4936864882732 | 0.588622055335422 |
| Rsl1d1 | ENSMUSG00000005846 | 66409 | -0.0519642331212804 | 4.9463927656921 | -0.703545623497807 | 0.4937557850086 | 0.588640471799445 |
| Plpp3 | ENSMUSG00000028517 | 67916 | -0.0499254418865958 | 6.37669079338277 | -0.703387585972083 | 0.4938511114757 | 0.588689912708978 |
| Zfp605 | ENSMUSG00000023284 | 675812 | 0.121175435304756 | 1.77415976211634 | 0.703026060712664 | 0.4940692207629 | 0.588885688819076 |
| Gtf3c3 | ENSMUSG00000041303 | 98488 | 0.0532038843794607 | 3.28762200169137 | 0.702224583192272 | 0.4945529594614 | 0.589397993880636 |
| Cops6 | ENSMUSG00000019494 | 26893 | -0.0615961833818428 | 4.80020126248761 | -0.701874073854773 | 0.4947646011305 | 0.589585942980239 |
| Tirap | ENSMUSG00000032041 | 117149 | 0.0916414924090296 | 2.168750094256 | 0.701368055850092 | 0.4950702359691 | 0.589832673309517 |
| Atxn7l2 | ENSMUSG00000048997 | 72522 | 0.178941909925782 | 1.3616077764775 | 0.70135260539559 | 0.4950795698119 | 0.589832673309517 |
| Eif2b2 | ENSMUSG00000004788 | 217715 | -0.0831339284634036 | 2.97709745444558 | -0.701012372292891 | 0.4952851361193 | 0.590013276255568 |
| Dot1l | ENSMUSG00000061589 | 208266 | 0.0738787270850416 | 5.7806700366415 | 0.700717554230745 | 0.4954633041031 | 0.590161204876487 |
| Ccar2 | ENSMUSG00000033712 | 219158 | 0.0575806608248044 | 3.74253289377333 | 0.700278397347371 | 0.4957287714061 | 0.590413074770152 |
| Taf1d | ENSMUSG00000031939 | 75316 | -0.070641973460071 | 4.30009161657032 | -0.700086712342167 | 0.4958446702062 | 0.59048677307168 |
| Plekha8 | ENSMUSG00000005225 | 231999 | 0.112203480662791 | 2.3940811622715 | 0.699934107721532 | 0.4959369513022 | 0.590532332754307 |
| Rps10 | ENSMUSG00000052146 | 67097 | 0.058362702149183 | 5.39681928289198 | 0.699735349971746 | 0.4960571568450 | 0.590611129667019 |
| Rnf213 | ENSMUSG00000070327 | 672511 | 0.116517005588976 | 5.90795839049983 | 0.698207225472037 | 0.4969819211398 | 0.591600710962129 |
| Atmin | ENSMUSG00000047388 | 234776 | -0.0677842413477094 | 4.26354006993583 | -0.69818305948343 | 0.4969965537247 | 0.591600710962129 |
| Zan | ENSMUSG00000079173 | 22635 | -0.204490221003666 | 1.90421123361403 | -0.697759924069365 | 0.4972528050777 | 0.591841290538419 |
| Fus | ENSMUSG00000030795 | 233908 | 0.0567616879437232 | 5.33345010395433 | 0.697566906853596 | 0.4973697226135 | 0.591915997592651 |
| Rttn | ENSMUSG00000023066 | 246102 | 0.123981739974151 | 1.68575753886661 | 0.697430054636448 | 0.4974526288611 | 0.591950216207396 |
| Utp15 | ENSMUSG00000041747 | 105372 | 0.0871176204460986 | 2.21123388589805 | 0.697096359307839 | 0.4976548187528 | 0.592126355043753 |
| Pramef8 | ENSMUSG00000046862 | 242736 | -0.108709099453156 | 2.20842020061809 | -0.696316577027275 | 0.4981274880926 | 0.592624246010146 |
| Neu1 | ENSMUSG00000007038 | 18010 | -0.0849446305693395 | 2.83577288769506 | -0.695978787587225 | 0.4983323236822 | 0.592803420412474 |
| Zfyve26 | ENSMUSG00000066440 | 211978 | 0.0693214890855263 | 4.10775748929091 | 0.695006161901291 | 0.4989224030460 | 0.593422422896251 |
| Eif2b4 | ENSMUSG00000029145 | 13667 | -0.0492411761084437 | 4.56238335242372 | -0.69494214472494 | 0.4989612559545 | 0.593422422896251 |
| Psmc3 | ENSMUSG00000002102 | 19182 | 0.052503085374819 | 6.63094358638454 | 0.693898608885071 | 0.4995948450338 | 0.594111319741598 |
| Reep1 | ENSMUSG00000052852 | 52250 | 0.0638806092090132 | 3.64007240402124 | 0.693630976933452 | 0.4997574161088 | 0.594239999508941 |
| Irak1bp1 | ENSMUSG00000032251 | 65099 | -0.143108715708574 | 1.04577651883 | -0.693199944119424 | 0.5000193096623 | 0.594486738516382 |
| Ptgis | ENSMUSG00000017969 | 19223 | 0.160712454546369 | 1.20013556841884 | 0.692511076560184 | 0.5004380309797 | 0.594919860428436 |
| Cep70 | ENSMUSG00000056267 | 68121 | -0.128579038320698 | 1.24309959830746 | -0.69233558219854 | 0.5005447366222 | 0.594982004786677 |
| Aoc3 | ENSMUSG00000019326 | 11754 | -0.105809944288923 | 2.83718417032674 | -0.691894201338382 | 0.5008131683968 | 0.59523635356593 |
| Klhl5 | ENSMUSG00000054920 | 71778 | 0.0981469346401065 | 2.51469926352049 | 0.690479681819189 | 0.5016740007209 | 0.596194662087488 |
| Cenpw | ENSMUSG00000075266 | 66311 | -0.0968803502006184 | 2.93894907344596 | -0.690220476210582 | 0.5018318398019 | 0.596317408509091 |
| Lrif1 | ENSMUSG00000056260 | 321000 | -0.110094246282459 | 2.56181361498076 | -0.689990514500744 | 0.5019718958155 | 0.596391311027669 |
| Ralb | ENSMUSG00000004451 | 64143 | -0.0505508867477715 | 3.86830255479149 | -0.689939199566965 | 0.5020031518402 | 0.596391311027669 |
| Clns1a | ENSMUSG00000025439 | 12729 | 0.0541826981242228 | 3.75301393207804 | 0.689208145728516 | 0.5024485628381 | 0.596855600998061 |
| Tnrc6a | ENSMUSG00000052707 | 233833 | 0.0519782730135443 | 5.09691505499675 | 0.688800309673728 | 0.5026971473731 | 0.597086006512692 |
| Zcchc14 | ENSMUSG00000061410 | 142682 | 0.0480738446659315 | 5.00454475313306 | 0.6883624804413 | 0.5029640940504 | 0.597338169498645 |
| Ube2q1 | ENSMUSG00000042572 | 70093 | -0.0602403147070998 | 3.49562733216596 | -0.688170110732432 | 0.5030814092362 | 0.597412589284276 |
| Parp10 | ENSMUSG00000063268 | 671535 | -0.143842008191897 | 1.99379450595335 | -0.687049269306364 | 0.5037652662843 | 0.59815969213062 |
| Rbm25 | ENSMUSG00000010608 | 67039 | 0.0407962539178577 | 6.01610254931144 | 0.686463029422847 | 0.5041231656599 | 0.598519639820728 |
| Nfe2l1 | ENSMUSG00000038615 | 18023 | -0.0542877603150949 | 9.26090727641207 | -0.685797719799054 | 0.5045295182653 | 0.598898771742436 |
| Susd6 | ENSMUSG00000021133 | 217684 | 0.0536900285177548 | 5.52819021223381 | 0.685760788971132 | 0.5045520802283 | 0.598898771742436 |
| Map3k6 | ENSMUSG00000028862 | 53608 | 0.203899675916317 | 2.03119683553442 | 0.684434706346098 | 0.5053626101158 | 0.599795731940942 |
| Lrsam1 | ENSMUSG00000026792 | 227738 | 0.0549120057408023 | 3.13208982306528 | 0.68425340249833 | 0.5054734862796 | 0.599835658521219 |
| Smpd2 | ENSMUSG00000019822 | 20598 | -0.103913776701906 | 1.82728910985013 | -0.684200240697413 | 0.5055060000272 | 0.599835658521219 |
| Plekhn1 | ENSMUSG00000078485 | 231002 | 0.0933317497635926 | 2.16167290923228 | 0.682764868833392 | 0.5063843372385 | 0.60077077021916 |
| C1galt1c1 | ENSMUSG00000048970 | 59048 | -0.0847254317582311 | 1.81838125465162 | -0.682714000036486 | 0.5064154814455 | 0.60077077021916 |
| Elavl1 | ENSMUSG00000040028 | 15568 | -0.052220046731092 | 4.79342873029719 | -0.682630833340134 | 0.5064664023249 | 0.60077077021916 |
| Uba1 | ENSMUSG00000001924 | 22201 | 0.0420669211994342 | 6.52847900754639 | 0.682553265394042 | 0.5065138979361 | 0.60077077021916 |
| Mpp7 | ENSMUSG00000057440 | 75739 | 0.0877940808195725 | 2.90339527681018 | 0.682158436444984 | 0.5067556960336 | 0.600992352538088 |
| Aftph | ENSMUSG00000049659 | 216549 | 0.0482728159503694 | 4.17207534451057 | 0.680853650331986 | 0.5075552441229 | 0.60187528460701 |
| Mettl9 | ENSMUSG00000030876 | 59052 | 0.0463729655075973 | 4.17834516624905 | 0.680457306620746 | 0.5077982621902 | 0.602037360588795 |
| Kctd6 | ENSMUSG00000021752 | 71393 | -0.106742029049963 | 1.8528848232734 | -0.680451091672454 | 0.5078020734268 | 0.602037360588795 |
| Wdr43 | ENSMUSG00000041057 | 72515 | -0.0653032476803181 | 4.85922302865744 | -0.679673247328355 | 0.5082792085464 | 0.602537688821259 |
| Usp49 | ENSMUSG00000090115 | 224836 | -0.116907179822537 | 2.12119867342015 | -0.679329603493653 | 0.5084900855461 | 0.602722308079104 |
| Rab40c | ENSMUSG00000025730 | 224624 | 0.088515486271753 | 1.91358322102419 | 0.679023257802517 | 0.5086781176658 | 0.602879811797168 |
| Cenpc1 | ENSMUSG00000029253 | 12617 | -0.0994794374358796 | 3.26414353671087 | -0.678454455091374 | 0.5090273512197 | 0.6032070075238 |
| Cdkl3 | ENSMUSG00000020389 | 213084 | -0.0905587265554564 | 2.20915986974656 | -0.678393875033204 | 0.5090645544238 | 0.6032070075238 |
| Pheta1 | ENSMUSG00000044134 | 231717 | -0.123506143872615 | 1.41339889670145 | -0.678217313479116 | 0.5091729928218 | 0.603235897271785 |
| Son | ENSMUSG00000022961 | 20658 | 0.0473588562410798 | 7.50162126658615 | 0.678174471626203 | 0.5091993069368 | 0.603235897271785 |
| Ptges3 | ENSMUSG00000071072 | 56351 | 0.0568352642636398 | 5.38256766544654 | 0.677649994256494 | 0.5095215132605 | 0.603513667373832 |
| Slmap | ENSMUSG00000021870 | 83997 | 0.0540198818512962 | 6.97659539428272 | 0.677596862691999 | 0.5095541606332 | 0.603513667373832 |
| Zfp867 | ENSMUSG00000054519 | 237775 | -0.0895877194329944 | 1.81412514433527 | -0.677457834789731 | 0.5096395938933 | 0.603513667373832 |
| Nagpa | ENSMUSG00000023143 | 27426 | 0.111127954001317 | 1.86090898985935 | 0.677433381966341 | 0.5096546211259 | 0.603513667373832 |
| Nup155 | ENSMUSG00000022142 | 170762 | 0.0555316186871408 | 3.62639254425034 | 0.677326376269876 | 0.5097203834250 | 0.603526160227403 |
| Spock2 | ENSMUSG00000058297 | 94214 | 0.121060431881428 | 3.09664229874486 | 0.676778288766455 | 0.5100572982217 | 0.60385966932329 |
| Bcar1 | ENSMUSG00000031955 | 12927 | 0.0769075524002103 | 3.66747837017594 | 0.676655242123106 | 0.5101329540620 | 0.603883833750509 |
| Naf1 | ENSMUSG00000014907 | 234344 | -0.114311505043434 | 1.97690568074782 | -0.676101848960551 | 0.5104732915103 | 0.604221283107668 |
| Thap3 | ENSMUSG00000039759 | 69876 | -0.121781566916166 | 2.18950351008748 | -0.675341236083607 | 0.5109412852845 | 0.60470974333536 |
| AI987944 | ENSMUSG00000056383 | 233168 | -0.0650895831676954 | 3.13174213667366 | -0.674953044809455 | 0.5111802299166 | 0.604927042678296 |
| Ankrd54 | ENSMUSG00000033055 | 223690 | 0.132709366058595 | 1.57618404120897 | 0.674745684112574 | 0.5113078939835 | 0.605012620603415 |
| Neb | ENSMUSG00000026950 | 17996 | 0.100319504747464 | 13.6276338588285 | 0.674382944164474 | 0.5115312637685 | 0.60521141295093 |
| Slc39a9 | ENSMUSG00000048833 | 328133 | 0.0679677145034154 | 3.40621199532087 | 0.674150781342286 | 0.5116742559407 | 0.605315074858011 |
| Cetn3 | ENSMUSG00000021537 | 12626 | -0.0608302246325261 | 3.61715895617497 | -0.674022720976937 | 0.5117531399792 | 0.605342882059679 |
| Nipbl | ENSMUSG00000022141 | 71175 | 0.0390231063926745 | 6.34532718905388 | 0.673888499979701 | 0.5118358265081 | 0.605375180649271 |
| Blvra | ENSMUSG00000001999 | 109778 | 0.0977908175015376 | 1.81799415864172 | 0.673683086640227 | 0.5119623859639 | 0.605459357456621 |
| Ebpl | ENSMUSG00000021928 | 68177 | -0.107721221427385 | 1.5425549065027 | -0.673197976961521 | 0.5122613444458 | 0.605747377340754 |
| Pebp1 | ENSMUSG00000032959 | 23980 | -0.0436504544562437 | 5.7151129234971 | -0.67282592898783 | 0.5124906951805 | 0.605953032884652 |
| Col11a2 | ENSMUSG00000024330 | 12815 | -0.177258859942058 | 3.7048257342772 | -0.672712146487194 | 0.5125608488606 | 0.605970434663235 |
| Lrrc28 | ENSMUSG00000030556 | 67867 | -0.0649921078958138 | 2.89653762230608 | -0.672489071425801 | 0.5126984041055 | 0.606067508951907 |
| Parp1 | ENSMUSG00000026496 | 11545 | -0.0496319926626594 | 4.99469506464815 | -0.671958988694599 | 0.5130253561864 | 0.606350085281108 |
| Rwdd2b | ENSMUSG00000041079 | 53858 | 0.110916751823103 | 1.39998037766552 | 0.671921653531889 | 0.5130483888724 | 0.606350085281108 |
| Mosmo | ENSMUSG00000046096 | 233812 | -0.0523656335730376 | 3.4033060053956 | -0.671416777158043 | 0.5133599145249 | 0.606601013378871 |
| Chml | ENSMUSG00000078185 | 12663 | -0.093038738272194 | 2.78839469716385 | -0.671329389940237 | 0.5134138465175 | 0.606601013378871 |
| Smurf2 | ENSMUSG00000018363 | 66313 | -0.0532006548062025 | 4.64800768455894 | -0.671307775667397 | 0.5134271865137 | 0.606601013378871 |
| Mettl6 | ENSMUSG00000021891 | 67011 | -0.0942746966432724 | 3.88855919983757 | -0.670677549118781 | 0.5138162410935 | 0.606995064454048 |
| Rac1 | ENSMUSG00000001847 | 19353 | -0.0395530087882241 | 6.47973539255177 | -0.670324059218426 | 0.5140345340979 | 0.607187323560092 |
| Fbxw11 | ENSMUSG00000020271 | 103583 | 0.0426475327935636 | 4.76778676527994 | 0.669854107392743 | 0.5143248299076 | 0.607464583006019 |
| Anapc4 | ENSMUSG00000029176 | 52206 | 0.0435524788808639 | 4.5315600246365 | 0.668823110794374 | 0.5149620238199 | 0.608151456609305 |
| Pgls | ENSMUSG00000031807 | 66171 | -0.0728847894012112 | 3.09755330470515 | -0.667997660021027 | 0.5154725123195 | 0.608574740605952 |
| Mtf2 | ENSMUSG00000029267 | 17765 | -0.063264143617614 | 3.30791393712682 | -0.667982797148495 | 0.5154817067396 | 0.608574740605952 |
| Zfyve21 | ENSMUSG00000021286 | 68520 | -0.0666093650658013 | 2.50906927627773 | -0.667973482529425 | 0.5154874689662 | 0.608574740605952 |
| Trp53bp2 | ENSMUSG00000026510 | 209456 | 0.0773777289402653 | 3.22541679580498 | 0.667870382931413 | 0.5155512511179 | 0.608584311659878 |
| Sec11a | ENSMUSG00000025724 | 56529 | 0.0712775995035484 | 3.17528647931484 | 0.667594174103796 | 0.5157221491076 | 0.608720312266088 |
| Hexdc | ENSMUSG00000039307 | 238023 | 0.117541065198927 | 3.63497923416374 | 0.667502317175062 | 0.5157789907761 | 0.608721674387157 |
| Tbca | ENSMUSG00000042043 | 21371 | 0.0684685390023905 | 3.40602349269636 | 0.667257589859284 | 0.5159304473330 | 0.608834688523928 |
| Zfp362 | ENSMUSG00000028799 | 230761 | -0.0716020028932212 | 3.0381014016131 | -0.666716681995493 | 0.5162652950225 | 0.60916406950464 |
| Wdr44 | ENSMUSG00000036769 | 72404 | 0.0575109994947232 | 3.74117288384606 | 0.666569657329372 | 0.5163563319979 | 0.609205727476457 |
| Ackr3 | ENSMUSG00000044337 | 12778 | -0.0782152715973314 | 3.33611531188463 | -0.665571565165534 | 0.5169745912204 | 0.609869334840321 |
| Rcl1 | ENSMUSG00000024785 | 59028 | 0.109472860281064 | 2.14442418094401 | 0.665131010867494 | 0.5172476245947 | 0.610125583732113 |
| St5 | ENSMUSG00000031024 | 76954 | -0.115405524887191 | 3.61041273890034 | -0.664976371701909 | 0.5173434819027 | 0.610172809740944 |
| Scfd1 | ENSMUSG00000020952 | 76983 | 0.0542295500826482 | 3.61015901445418 | 0.664527938721529 | 0.5176215133085 | 0.61043486481562 |
| Kdsr | ENSMUSG00000009905 | 70750 | 0.0858968247507139 | 2.41408716642782 | 0.66407185100623 | 0.5179043791272 | 0.610702563995654 |
| Bdnf | ENSMUSG00000048482 | 12064 | 0.173051660898417 | 1.27169291154129 | 0.663853352563176 | 0.5180399235512 | 0.610796505699307 |
| Zfp395 | ENSMUSG00000034522 | 380912 | -0.146986069494886 | 2.99084112407621 | -0.663729750813735 | 0.5181166083419 | 0.610821035999292 |
| Metap1 | ENSMUSG00000005813 | 75624 | -0.0576859184816669 | 4.61754159710348 | -0.663290424232188 | 0.5183892275805 | 0.611076528273851 |
| 5031425E22Rik | ENSMUSG00000073147 | 269630 | 0.0865033356002658 | 2.81402082972266 | 0.663127277852956 | 0.5184904873082 | 0.611129988868497 |
| Pnrc2 | ENSMUSG00000028675 | 52830 | 0.0569519913143042 | 4.9680230307405 | 0.662817013359615 | 0.5186830899842 | 0.611243556797616 |
| Gpatch8 | ENSMUSG00000034621 | 237943 | -0.0639424395670201 | 4.77923018367273 | -0.662766878937617 | 0.5187142157632 | 0.611243556797616 |
| Gnpda2 | ENSMUSG00000029209 | 67980 | 0.104179644868758 | 1.57798847630133 | 0.662701842741299 | 0.5187545948597 | 0.611243556797616 |
| Scrib | ENSMUSG00000022568 | 105782 | 0.0454080779926791 | 4.22815141356985 | 0.662472935885025 | 0.5188967309204 | 0.611345135340722 |
| Mill2 | ENSMUSG00000040987 | 243864 | -0.0897163078856942 | 1.73565991795431 | -0.662274069052227 | 0.5190202323154 | 0.61139905019694 |

| | | | | | | | |
|---|---|---|---|---|---|---|---|
| 1110038F14Rik | ENSMUSG00000063236 | 117171 | 0.0579640395817909 | 3.08043785811482 | 0.662219123547032 | 0.5190543578654 | 0.61139905019694 |
| Sil1 | ENSMUSG00000024357 | 81500 | -0.0894561564524121 | 2.4887042334694 | -0.661678250367089 | 0.5193903522905 | 0.611691458473083 |
| Polr3f | ENSMUSG00000027427 | 70408 | -0.0633294793764065 | 3.05582097378044 | -0.661634334091801 | 0.5194176389001 | 0.611691458473083 |
| Naa35 | ENSMUSG00000021555 | 78689 | -0.063815011102053 | 3.48411691087274 | -0.661549294225391 | 0.5194704792796 | 0.611691458473083 |
| Zfp553 | ENSMUSG00000045598 | 233887 | -0.0798690516239571 | 2.34764534973996 | -0.661433321053673 | 0.5195425453666 | 0.611710422598357 |
| Ripk2 | ENSMUSG00000041135 | 192656 | 0.117793242352462 | 1.42302798227737 | 0.660625764975884 | 0.5200445226836 | 0.612173493019148 |
| Borcs7 | ENSMUSG00000062376 | 66439 | -0.0888719018360917 | 2.70813867704557 | -0.660620412499956 | 0.5200478507159 | 0.612173493019148 |
| Ryr1 | ENSMUSG00000030592 | 20190 | -0.0823727949468651 | 11.4598211271237 | -0.659770800436976 | 0.5205762729357 | 0.612729540159457 |
| Slc37a3 | ENSMUSG00000029924 | 72144 | 0.0893047026495366 | 2.66215973220849 | 0.659650311006508 | 0.5206512371600 | 0.612751795154664 |
| Taf4 | ENSMUSG00000039117 | 228980 | -0.0606023918041623 | 3.25820043982111 | -0.659314328647399 | 0.5208603061509 | 0.612889347845494 |
| Ctnnd1 | ENSMUSG00000101645 | 12388 | 0.197026281032699 | 3.21968113704746 | 0.659282277895733 | 0.5208802526287 | 0.612889347845494 |
| Desi2 | ENSMUSG00000026502 | 78825 | 0.050300087781201 | 4.51036959181376 | 0.659041794131661 | 0.5210299293859 | 0.612999478863151 |
| Trappc1 | ENSMUSG00000049299 | 245828 | 0.0823790551000347 | 2.48395122977821 | 0.658917449353741 | 0.5211073310864 | 0.613024562645979 |
| Ttc23 | ENSMUSG00000030555 | 67009 | -0.0907124157759114 | 2.58753089548548 | -0.658387187262803 | 0.5214374808625 | 0.613346938911975 |
| Hoxc9 | ENSMUSG00000036139 | 15427 | 0.0808939274042229 | 2.60236713687899 | 0.657993284981345 | 0.5216828084936 | 0.613569483499457 |
| Rgs3 | ENSMUSG00000059810 | 50780 | -0.116183950162611 | 3.49046571573499 | -0.657816418913729 | 0.5217929845924 | 0.613633040837062 |
| Hnrnpu | ENSMUSG00000039630 | 51810 | -0.0385740884174828 | 7.52573214863491 | -0.657463010494527 | 0.5220131751076 | 0.613825948483439 |
| Zfp607a | ENSMUSG00000020420 | 545938 | 0.172961647982681 | 1.09767469963977 | 0.656701554275597 | 0.5224877801211 | 0.614296692613033 |
| Cdc42bpb | ENSMUSG00000021279 | 217866 | -0.0592728659185084 | 5.61739686613067 | -0.656640406810191 | 0.5225259032033 | 0.614296692613033 |
| Akap10 | ENSMUSG00000047804 | 56697 | -0.068499710682643 | 3.33434300704388 | -0.656445546267283 | 0.5226474018543 | 0.614373454099304 |
| Snx25 | ENSMUSG00000038291 | 102141 | -0.0831815621622076 | 2.22048293364483 | -0.656158773611892 | 0.5228262386010 | 0.614491889623768 |
| Cul4a | ENSMUSG00000031446 | 99375 | -0.0493019900978222 | 6.15238952872821 | -0.656103702491544 | 0.5228605859839 | 0.614491889623768 |
| Crem | ENSMUSG00000063889 | 12916 | 0.0769420196684565 | 3.12514506117055 | 0.655874956656943 | 0.5230032666182 | 0.614593496818381 |
| Zmynd8 | ENSMUSG00000039671 | 228880 | 0.0597201521681363 | 4.8736383239159 | 0.655454464716235 | 0.5232656073825 | 0.614835682500178 |
| Hirip3 | ENSMUSG00000042606 | 233876 | -0.118094281942507 | 1.34418888969575 | -0.655308143364886 | 0.5233569134876 | 0.614876872456294 |
| Sncg | ENSMUSG00000023064 | 20618 | -0.116793182502323 | 1.56832720120724 | -0.655075708708043 | 0.5235019739689 | 0.614935507785838 |
| Vsig10 | ENSMUSG00000066894 | 231668 | -0.1047069541663 | 1.60837706857485 | -0.655013091751559 | 0.5235410566088 | 0.614935507785838 |
| 1110065P20Rik | ENSMUSG00000078570 | 68920 | -0.103794912995346 | 1.66560321596116 | -0.654872677574677 | 0.5236287027830 | 0.614935507785838 |
| Tapbpl | ENSMUSG00000038213 | 213233 | -0.119061042683339 | 1.52927320764204 | -0.654867642527195 | 0.5236318458028 | 0.614935507785838 |
| Wdr45b | ENSMUSG00000025173 | 66840 | 0.053379075182109 | 3.48179485504396 | 0.654661599363818 | 0.5237604730589 | 0.61502048888247 |
| 1810009A15Rik | ENSMUSG00000071653 | 66276 | 0.0590072966010018 | 2.52131941769767 | 0.653174823631417 | 0.5246891629391 | 0.616044816335981 |
| Fance | ENSMUSG00000007570 | 72775 | 0.117248799700519 | 1.19255364969809 | 0.652941469783088 | 0.5248350089112 | 0.616149874600932 |
| Slc27a4 | ENSMUSG00000059316 | 26569 | -0.079260377848418 | 2.47304045980651 | -0.652188281800453 | 0.5253059087049 | 0.616636478528112 |
| Cuedc2 | ENSMUSG00000036748 | 67116 | -0.0584247268981297 | 4.4974328131803 | -0.651776633780027 | 0.5255633765473 | 0.616832155627525 |
| Zcchc10 | ENSMUSG00000018239 | 67966 | 0.0782224784875842 | 1.79753658463671 | 0.651741326380223 | 0.5255854631337 | 0.616832155627525 |
| Trim11 | ENSMUSG00000020455 | 94091 | 0.101602619355678 | 2.28673267364444 | 0.65164185048522 | 0.5256476932461 | 0.616838962412592 |
| Rftn1 | ENSMUSG00000039316 | 76438 | -0.0624475712586454 | 4.26828769296531 | -0.65080181297503 | 0.5261733712921 | 0.617389557921294 |
| Gtf2h4 | ENSMUSG00000001524 | 14885 | 0.083664797098022 | 2.39516367340329 | 0.650461089919494 | 0.5263866740842 | 0.617573546679764 |
| BC029722 | ENSMUSG00000074649 | 613262 | -0.0570437664123657 | 3.50213987992914 | -0.650285375427065 | 0.5264966958458 | 0.617636336369503 |
| 3110082I17Rik | ENSMUSG00000053553 | 73212 | -0.0735952914791485 | 2.024619056323 | -0.650134246022787 | 0.5265913343924 | 0.617681068381125 |
| Cnksr1 | ENSMUSG00000028841 | 194231 | 0.0703222029714121 | 4.83550972299282 | 0.649843825093242 | 0.5267732256967 | 0.617828125546277 |
| Gm6710 | ENSMUSG00000078887 | 626832 | 0.117537147119309 | 1.42010759727266 | 0.649165071258786 | 0.5271984702646 | 0.618260538403942 |
| Rbm28 | ENSMUSG00000029701 | 68272 | 0.0431081687162793 | 4.19239726025134 | 0.648981966127304 | 0.5273132205030 | 0.618304173221812 |
| Ost4 | ENSMUSG00000038803 | 67695 | 0.0500956330063413 | 3.84408351397863 | 0.648925186270894 | 0.5273488067831 | 0.618304173221812 |
| Arl13b | ENSMUSG00000022911 | 68146 | -0.111610550647064 | 2.62282551471876 | -0.647702231620074 | 0.5281156144458 | 0.61913682770356 |
| Ubl5 | ENSMUSG00000084786 | 66177 | -0.0580371774773241 | 4.12739962226966 | -0.647177719803546 | 0.5284446838765 | 0.61945617581283 |
| Vegfd | ENSMUSG00000031380 | 14205 | -0.122221710775056 | 2.04653502924324 | -0.647024406395039 | 0.5285408919857 | 0.619502518796506 |
| Senp7 | ENSMUSG00000052917 | 66315 | 0.0664445165640979 | 3.05871830355957 | 0.646684193853831 | 0.5287544196305 | 0.619686347269356 |
| Ubxn7 | ENSMUSG00000053774 | 224111 | 0.0584252798539255 | 4.38335115212042 | 0.646396488937955 | 0.5289350301170 | 0.619831562415236 |
| Ktn1 | ENSMUSG00000021843 | 16709 | -0.049727549211747 | 8.06367423938317 | -0.645997519382115 | 0.5291855463518 | 0.620058656573223 |
| Borcs8 | ENSMUSG00000002345 | 72368 | 0.0783907450472521 | 2.21692083622674 | 0.64572813722845 | 0.5293547316616 | 0.620190414983182 |
| Zc3hc1 | ENSMUSG00000039130 | 232679 | -0.0689818016751711 | 2.70809132965942 | -0.645607530830909 | 0.5294304883827 | 0.620212696228901 |
| Purg | ENSMUSG00000049184 | 75029 | 0.107425823306169 | 1.72747458508872 | 0.645406613712369 | 0.5295567044780 | 0.620241609597531 |
| Thoc2 | ENSMUSG00000037475 | 331401 | 0.0440226070561005 | 4.91424612375613 | 0.645387595381834 | 0.5295686526734 | 0.620241609597531 |
| Rab35 | ENSMUSG00000029518 | 77407 | 0.0445412418913538 | 4.39360784961528 | 0.64448721130618 | 0.5301344904903 | 0.62083780967961 |
| Tnpo2 | ENSMUSG00000031691 | 212999 | 0.0428069278505792 | 3.90713492286268 | 0.644016731776547 | 0.5304302950741 | 0.621117681355666 |
| Tmem117 | ENSMUSG00000063296 | 320709 | 0.0638156665053753 | 3.22301251258666 | 0.643154201691816 | 0.5309728362418 | 0.621677187812595 |
| Cdk7 | ENSMUSG00000069089 | 12572 | 0.0762398792375278 | 2.30359700610471 | 0.643076295596404 | 0.5310218555124 | 0.621677187812595 |
| Dhrs7b | ENSMUSG00000042569 | 216820 | -0.0768648333262454 | 2.66035940312915 | -0.642230482341952 | 0.5315542141591 | 0.622134258797664 |
| Prkrip1 | ENSMUSG00000039737 | 66801 | -0.0707161405917076 | 2.90638343646897 | -0.642185703865554 | 0.5315824063394 | 0.622134258797664 |
| Phf8 | ENSMUSG00000041229 | 320595 | 0.056248930039339 | 3.09104289898622 | 0.642184731284864 | 0.5315830186780 | 0.622134258797664 |
| Pja1 | ENSMUSG00000034403 | 18744 | 0.0505659001502647 | 3.26197132956162 | 0.641821129202505 | 0.5318119711616 | 0.622335580426951 |
| Polr3k | ENSMUSG00000038628 | 67005 | -0.0810263142698315 | 2.34384544961637 | -0.641183410628069 | 0.5322136634922 | 0.622738980478874 |
| Six4 | ENSMUSG00000034460 | 20474 | 0.0749315381629501 | 5.12211032746821 | 0.63907256298033 | 0.5335444859011 | 0.624229345540511 |
| Rtp4 | ENSMUSG00000033355 | 67775 | -0.148089147348042 | 1.58949115428396 | -0.638741956770406 | 0.5337530923905 | 0.624406576725239 |
| Lrrc57 | ENSMUSG00000027286 | 66606 | 0.0764677761038071 | 3.03880017477079 | 0.637781262809028 | 0.5343595334506 | 0.625049123611425 |
| Pck2 | ENSMUSG00000040618 | 74551 | 0.0907734271119893 | 2.18441495170358 | 0.637354532375459 | 0.5346290327261 | 0.625297448826166 |
| Lats2 | ENSMUSG00000021959 | 50523 | -0.0728343958964572 | 4.72340367527775 | -0.636929338666248 | 0.5348976375394 | 0.625531505668947 |
| Nabp2 | ENSMUSG00000025374 | 69917 | 0.0514340715877131 | 4.11466647348775 | 0.63685659005843 | 0.5349436021400 | 0.625531505668947 |
| Thoc3 | ENSMUSG00000025872 | 73666 | -0.0669670132177997 | 2.69947588162401 | -0.63670066816493 | 0.5350421254259 | 0.625579791745753 |
| Ttc7 | ENSMUSG00000036918 | 225049 | -0.0702117741218822 | 3.54404842432095 | -0.636045855350534 | 0.5354559972377 | 0.625996738588871 |
| Dnpep | ENSMUSG00000026209 | 13437 | -0.0563335210395781 | 4.21422205497081 | -0.635031686696303 | 0.5360973534058 | 0.626659831658962 |
| Acbd5 | ENSMUSG00000026781 | 74159 | -0.0442327140988441 | 4.72772340254988 | -0.63496768620449 | 0.5361378415217 | 0.626659831658962 |
| Zfp866 | ENSMUSG00000043090 | 330788 | -0.0649057598539491 | 3.37676534004799 | -0.634692940304018 | 0.5363116713091 | 0.626795988354609 |
| Rbm10 | ENSMUSG00000031060 | 236732 | -0.0511263232034684 | 3.68594976162754 | -0.634512525346243 | 0.5364258357794 | 0.626862391587065 |
| Arhgap23 | ENSMUSG00000049807 | 58996 | 0.0922838304382081 | 3.55815012401468 | 0.634207811560943 | 0.5366186860523 | 0.627020722312981 |
| A230057D06Rik | ENSMUSG00000109394 | 319893 | -0.245101436191079 | 0.524696653630263 | -0.633570176941748 | 0.5370223643665 | 0.627425338273873 |
| Copz1 | ENSMUSG00000060992 | 56447 | 0.0542723381246968 | 3.82212519970239 | 0.632881122847413 | 0.5374587870158 | 0.627868120216931 |
| Snrpd1 | ENSMUSG00000002477 | 20641 | -0.0795370502925321 | 2.41991351398652 | -0.631763003929686 | 0.5381673868580 | 0.6286287353863 |
| Tmco3 | ENSMUSG00000038497 | 234076 | 0.0698416376397373 | 3.08598438266657 | 0.630278905739345 | 0.5391087304207 | 0.629661024920363 |
| Psmd7 | ENSMUSG00000039067 | 17463 | 0.0420411263482245 | 6.47916445092078 | 0.629971551324803 | 0.5393037961618 | 0.629821559385188 |
| Ets2 | ENSMUSG00000022895 | 23872 | 0.0810268190984251 | 5.3055198821567 | 0.629595227835687 | 0.5395426875435 | 0.630033235502442 |
| Orai3 | ENSMUSG00000043964 | 269999 | -0.0762154774354804 | 2.18102199687195 | -0.629194913086972 | 0.5397968735086 | 0.630262724238694 |
| 4933421O10Rik | ENSMUSG00000085037 | 100504166 | 0.10929636246072 | 1.5102250931834 | 0.628110444477819 | 0.5404858091297 | 0.630990729493448 |
| Srsf3 | ENSMUSG00000071172 | 20383 | -0.0900368212801634 | 4.80709922988496 | -0.628031730780965 | 0.5405358330415 | 0.630990729493448 |
| Slc50a1 | ENSMUSG00000027953 | 19729 | 0.0832377138991891 | 1.92960038152681 | 0.627730901966753 | 0.5407270387251 | 0.631146530731952 |
| Inpp5j | ENSMUSG00000034570 | 170835 | -0.107944140154075 | 3.08876797693834 | -0.627206196998373 | 0.5410606294758 | 0.631468475421825 |
| Zfp758 | ENSMUSG00000044501 | 224598 | -0.0688117995906454 | 2.84373638589574 | -0.62709505519363 | 0.5411313046183 | 0.631483536968411 |
| Csrnp1 | ENSMUSG00000032515 | 215418 | -0.210466929103094 | 1.90146134530687 | -0.626681634179359 | 0.5413942444362 | 0.631722938293403 |
| Klhdc4 | ENSMUSG00000040263 | 234825 | 0.0736888372777456 | 2.8964200651164 | 0.62648192636657 | 0.5415212860440 | 0.631803733349097 |
| Adrm1 | ENSMUSG00000039041 | 56436 | 0.0469290216976139 | 5.4846480678347 | 0.626251276168146 | 0.5416680319119 | 0.63190749806082 |
| Slc16a9 | ENSMUSG00000037762 | 66859 | 0.137969722189228 | 1.41745091950888 | 0.62502406751291 | 0.5424491871550 | 0.63275126078237 |
| Abcf1 | ENSMUSG00000038762 | 224742 | 0.0452656729827838 | 6.62139824906451 | 0.624113493776222 | 0.5430291992710 | 0.633360240848518 |
| S100pbp | ENSMUSG00000040928 | 74648 | 0.0817081087697529 | 1.95282560543066 | 0.623835732737784 | 0.5432061944743 | 0.633499083729757 |
| Skil | ENSMUSG00000027660 | 20482 | 0.0686844524436658 | 4.72810436361027 | 0.622988880550411 | 0.5437460242474 | 0.63405939517998 |
| Arhgef18 | ENSMUSG00000004568 | 102098 | -0.0610496531182525 | 4.1337199653801 | -0.622900067560103 | 0.5438026557325 | 0.63405939517998 |
| Psma1 | ENSMUSG00000030751 | 26440 | -0.0524624471770645 | 4.91621537293465 | -0.622420117448667 | 0.5441087518950 | 0.634348631288953 |
| Lrg1 | ENSMUSG00000037095 | 76905 | -0.111024210929378 | 1.89182634394565 | -0.622097636986585 | 0.5443144728230 | 0.634520795822646 |
| Rnpc3 | ENSMUSG00000027981 | 67225 | 0.0631595535225196 | 2.74545668475312 | 0.621819130950485 | 0.5444921757065 | 0.634633736968979 |
| Stip1 | ENSMUSG00000024966 | 20867 | -0.0478792634606817 | 5.89031122101704 | -0.621763815354769 | 0.5445274740675 | 0.634633736968979 |
| Tug1 | ENSMUSG00000056579 | 544752 | -0.0603760957317583 | 4.47665507621885 | -0.621576182231611 | 0.5446472172268 | 0.634705621695853 |
| Synpo2 | ENSMUSG00000050315 | 118449 | 0.0730195327775114 | 9.4852379753407 | 0.621159150476147 | 0.5449134095753 | 0.634932791067357 |
| Mbd2 | ENSMUSG00000024513 | 17191 | -0.0411964633740105 | 5.38420152824335 | -0.621088795282761 | 0.5449583245626 | 0.634932791067357 |
| Rps15 | ENSMUSG00000063457 | 20054 | -0.0439207591426767 | 6.00113733389874 | -0.62029473542394 | 0.5454653971078 | 0.635455851623818 |
| Hypk | ENSMUSG00000027245 | 67693 | 0.0999819774295729 | 2.26637190669914 | 0.619752317515339 | 0.5458119255969 | 0.635791790142814 |
| Zfp281 | ENSMUSG00000041483 | 226442 | -0.0580057138446448 | 3.78694573458811 | -0.619407522935218 | 0.5460322639499 | 0.635980679513753 |
| Ttll5 | ENSMUSG00000012609 | 320244 | -0.0603037536672324 | 2.61931343495568 | -0.618104214027235 | 0.5468655776348 | 0.636883403912953 |
| Ftx | ENSMUSG00000086370 | 78878 | -0.0956876357875496 | 2.35813160474865 | -0.617843406815807 | 0.5470324176401 | 0.637009838843566 |
| Nfx1 | ENSMUSG00000028423 | 74164 | -0.0494293956644825 | 4.36424791531419 | -0.617189790586374 | 0.5474506634388 | 0.637428973375589 |
| Pik3r1 | ENSMUSG00000041417 | 18708 | 0.0886605847980926 | 5.74251022593179 | 0.61545630404005 | 0.5485607666508 | 0.638653503063176 |
| Ift46 | ENSMUSG00000002031 | 76568 | 0.0684473867743121 | 2.65112226830719 | 0.614921282954091 | 0.5489036376823 | 0.638984628701306 |
| Tm9sf2 | ENSMUSG00000025544 | 68059 | -0.0472328220026847 | 5.80117587650778 | -0.614511172341895 | 0.5491665390264 | 0.639222600159489 |
| Acd | ENSMUSG00000038000 | 497652 | -0.093331983261101 | 2.15895406991978 | -0.614371107044365 | 0.5492563437306 | 0.63925906019168 |
| Dexi | ENSMUSG00000038055 | 58239 | 0.0748600797530501 | 3.026693744612 | 0.613877216028442 | 0.5495730728889 | 0.639559593287481 |
| Samd4 | ENSMUSG00000021838 | 74480 | 0.0718033263681432 | 7.16490704570355 | 0.613284072035946 | 0.5499535850184 | 0.639934281226007 |
| Fam214a | ENSMUSG00000034858 | 235493 | 0.0954226681231694 | 4.87664424418318 | 0.612466678750715 | 0.5504781940512 | 0.640476544882836 |

| | | | | | | | |
|---|---|---|---|---|---|---|---|
| Vmn1r229 | ENSMUSG00000117148 | 171224 | -0.173763484431414 | 1.70520045987267 | -0.612268328042077 | 0.5506055384098 | 0.64052490894006 |
| Mcm3ap | ENSMUSG00000001150 | 54387 | 0.0433883625444063 | 4.61607597401425 | 0.61221939562316 | 0.5506369562994 | 0.64052490894006 |
| Rbm17 | ENSMUSG00000037197 | 76938 | 0.0526789079410417 | 4.66250384928191 | 0.612016382321401 | 0.5507673149423 | 0.640608376211413 |
| Agtpbp1 | ENSMUSG00000021557 | 67269 | -0.0510547965469188 | 5.63263192614856 | -0.6117393543741 | 0.5509452270930 | 0.640726066506725 |
| Med13l | ENSMUSG00000018076 | 76199 | 0.0469275081422929 | 5.59211987394347 | 0.611626544233544 | 0.5510176847535 | 0.640726066506725 |
| Synj2 | ENSMUSG00000023805 | 20975 | 0.0473003058408885 | 5.86932239307759 | 0.611585036362214 | 0.5510443464669 | 0.640726066506725 |
| Rnd3 | ENSMUSG00000017144 | 74194 | 0.115883735825148 | 3.72836715333221 | 0.611310980206162 | 0.5512203984669 | 0.640862601110619 |
| Camta1 | ENSMUSG00000014592 | 100072 | -0.0400838723986094 | 5.37425538397159 | -0.610516062149311 | 0.5517312233900 | 0.641388280641935 |
| Ash2l | ENSMUSG00000031575 | 23808 | -0.0436550794935737 | 4.53891130172138 | -0.609485016501939 | 0.5523941732360 | 0.64209067499398 |
| Bahcc1 | ENSMUSG00000039741 | 268515 | 0.068848849037809 | 3.72466284144166 | 0.609192876471852 | 0.5525820949756 | 0.64224081660595 |
| Chka | ENSMUSG00000024843 | 12660 | 0.0925034729411012 | 1.54762664536105 | 0.609081742054276 | 0.5526535923872 | 0.642255626024327 |
| Seh1l | ENSMUSG00000079614 | 72124 | 0.0567131124314821 | 3.51238180345003 | 0.607739260335268 | 0.5535176665941 | 0.643191412091036 |
| Pigc | ENSMUSG00000026698 | 67292 | -0.0717507183370674 | 2.50432255731238 | -0.606999034567852 | 0.5539944206290 | 0.643676978305279 |
| Slc4a7 | ENSMUSG00000021733 | 218756 | -0.0635238379661436 | 4.51799000518913 | -0.606768215541827 | 0.5541431290885 | 0.643722205868089 |
| Mthfd2l | ENSMUSG00000029376 | 665563 | -0.0921984467676051 | 1.94571469267515 | -0.606755804184175 | 0.5541511259005 | 0.643722205868089 |
| Sec22a | ENSMUSG00000034473 | 317717 | 0.085451648460656 | 1.84973930862494 | 0.606310977554542 | 0.5544377755169 | 0.643986752117272 |
| Gfpt1 | ENSMUSG00000029992 | 14583 | 0.0402201377193574 | 4.65433691508494 | 0.605829482498846 | 0.5547481456872 | 0.644220838194619 |
| Gstp2 | ENSMUSG00000038155 | 14869 | 0.0503381870036721 | 4.3322949696349 | 0.605769819354672 | 0.5547866109547 | 0.644220838194619 |
| Zc3h6 | ENSMUSG00000042851 | 78751 | 0.141729914724209 | 2.90105492777752 | 0.605724054902047 | 0.5548161166191 | 0.644220838194619 |
| Btbd10 | ENSMUSG00000038187 | 68815 | 0.0722007163849745 | 2.54027768850199 | 0.605320205472014 | 0.5550765270928 | 0.644386438693669 |
| Pxn | ENSMUSG00000029528 | 19303 | 0.0588707117069287 | 5.51804706882134 | 0.605320036793612 | 0.5550766358740 | 0.644386438693669 |
| Med13 | ENSMUSG00000034297 | 327987 | 0.0384808870009268 | 6.53886247029023 | 0.605124152025322 | 0.5552029704285 | 0.644464656446215 |
| Ecd | ENSMUSG00000021810 | 70601 | -0.0441898675952173 | 4.25011079638247 | -0.60492792953606 | 0.5553295384995 | 0.644543128619515 |
| Znhit6 | ENSMUSG00000074182 | 229937 | 0.0556102139133614 | 2.70030651081227 | 0.604739949589881 | 0.5554508046785 | 0.644555142321105 |
| Uba2 | ENSMUSG00000052997 | 50995 | -0.0375920737712034 | 4.7072072950168 | -0.604729074068167 | 0.5554578209372 | 0.644555142321105 |
| Tbx3 | ENSMUSG00000018604 | 21386 | -0.0516653324660883 | 4.41961069514705 | -0.604435133913627 | 0.5556474724338 | 0.644706774352458 |
| Dnajb4 | ENSMUSG00000028035 | 67035 | -0.0532182194963537 | 6.59096948030693 | -0.602450169168616 | 0.5569291028347 | 0.64612524125311 |
| Idi1 | ENSMUSG00000058258 | 319554 | -0.095595267548116 | 2.23107627598745 | -0.602088874138106 | 0.5571625525657 | 0.646327481916212 |
| Pex3 | ENSMUSG00000019809 | 56535 | 0.0710285126310262 | 2.56699227839205 | 0.60101609610109 | 0.5578560380465 | 0.647063280123817 |
| Rnf168 | ENSMUSG00000014074 | 70238 | -0.0581772916474468 | 3.21061488171471 | -0.600525435372234 | 0.5581733762572 | 0.647291365146261 |
| Rfx1 | ENSMUSG00000031706 | 19724 | 0.103493136956941 | 2.09279293501939 | 0.600480584811195 | 0.5582023885492 | 0.647291365146261 |
| Hddc3 | ENSMUSG00000030532 | 68695 | 0.0798603272389853 | 1.97579877649691 | 0.600437396248789 | 0.5582303265239 | 0.647291365146261 |
| Rcan1 | ENSMUSG00000022951 | 54720 | 0.107932125100018 | 7.2922460243054 | 0.600319008645729 | 0.5583069134159 | 0.647311505149502 |
| Faim | ENSMUSG00000032463 | 23873 | 0.109189006155459 | 1.18420614669956 | 0.599942362302409 | 0.5585506100297 | 0.647461759273423 |
| Ints6l | ENSMUSG00000035967 | 236790 | 0.080968181277584 | 2.4226584692603 | 0.599935622722186 | 0.5585549711781 | 0.647461759273423 |
| Syt11 | ENSMUSG00000068923 | 229521 | 0.0758546799955617 | 3.19178311462863 | 0.599647411952598 | 0.5587414881808 | 0.647609289291191 |
| Zfp646 | ENSMUSG00000049739 | 233905 | 0.0478176615071124 | 4.51563964550435 | 0.599304380873199 | 0.5589635263869 | 0.647797954509766 |
| Mtmr6 | ENSMUSG00000021987 | 219135 | 0.0368045196229465 | 5.0384305850832 | 0.599166472509785 | 0.5590528055999 | 0.647832738048689 |
| Msrb1 | ENSMUSG00000075705 | 27361 | -0.05205205801846 | 4.8605958301367 | -0.599040191322541 | 0.5591345643755 | 0.647858800422858 |
| Fig4 | ENSMUSG00000038417 | 103199 | -0.0519819432776241 | 2.98267319926739 | -0.598637993919643 | 0.5593950039169 | 0.648091869402825 |
| Hivep2 | ENSMUSG00000015501 | 15273 | 0.0464225933540474 | 6.21211973555208 | 0.598387374345157 | 0.5595573237069 | 0.648211223552383 |
| Cdc37 | ENSMUSG00000019471 | 12539 | -0.0435335293883199 | 6.40750420576451 | -0.598170699155764 | 0.5596976791376 | 0.648233036030635 |
| Rif1 | ENSMUSG00000036202 | 51869 | -0.0542852335788142 | 4.47437341872898 | -0.598094297924539 | 0.5597471740136 | 0.648233036030635 |
| Rassf1 | ENSMUSG00000010067 | 56289 | -0.078844320477137 | 2.57188820105795 | -0.598079598286005 | 0.5597566971263 | 0.648233036030635 |
| Ppp2r5b | ENSMUSG00000024777 | 225849 | 0.0506371566077171 | 4.38149662410522 | 0.597903164457079 | 0.5598710060423 | 0.648233036030635 |
| Bora | ENSMUSG00000022070 | 77744 | -0.189943920419466 | 0.605268101047696 | -0.597900605150558 | 0.5598726642727 | 0.648233036030635 |
| Npat | ENSMUSG00000033054 | 244879 | 0.0617622925856449 | 2.81538164158641 | 0.597438190917291 | 0.5601723161064 | 0.648511288642219 |
| Snx29 | ENSMUSG00000071669 | 74478 | 0.0637786578723207 | 2.53104210919907 | 0.59683472150419 | 0.5605635043012 | 0.648895442710726 |
| Pms2 | ENSMUSG00000079109 | 18861 | 0.0990067302422544 | 1.52164622205765 | 0.596703284632681 | 0.5606487254558 | 0.648925372507239 |
| Dtx3 | ENSMUSG00000040415 | 80904 | -0.0937800210093918 | 3.35798361397984 | -0.596023731819032 | 0.5610894460917 | 0.649301293047644 |
| Rrn3 | ENSMUSG00000022682 | 106298 | -0.0500291598610758 | 4.48872236737653 | -0.595905913296451 | 0.5611658757307 | 0.649301293047644 |
| Ncstn | ENSMUSG00000003458 | 59287 | -0.0525101472438738 | 4.45481498501746 | -0.595891509271963 | 0.5611752201001 | 0.649301293047644 |
| Srrm2 | ENSMUSG00000039218 | 75956 | -0.0455733451296174 | 7.90008849264046 | -0.595836191191559 | 0.5612111075577 | 0.649301293047644 |
| Fam160b1 | ENSMUSG00000033478 | 226252 | -0.0479206529287834 | 3.86725861055674 | -0.595309687433422 | 0.5615527374445 | 0.649627788443872 |
| Ddit4 | ENSMUSG00000020108 | 74747 | 0.216927065179225 | 1.23498108948566 | 0.594875208116681 | 0.5618347403280 | 0.649885243018654 |
| Coprs | ENSMUSG00000031458 | 66423 | -0.0818880266428588 | 1.87173497073218 | -0.594569527701121 | 0.5620331908128 | 0.650046006642156 |
| Cyb561a3 | ENSMUSG00000034445 | 225912 | -0.0862734831990437 | 1.50925953969038 | -0.594317268740924 | 0.5621969880500 | 0.650166660640561 |
| Katnb1 | ENSMUSG00000031787 | 74187 | -0.0650695729513462 | 2.61534790632849 | -0.593874682939376 | 0.5624844307793 | 0.650430266883483 |
| Smim14 | ENSMUSG00000037822 | 68552 | -0.0472641621453684 | 4.30209192890424 | -0.593716105171259 | 0.5625874402927 | 0.650480570112161 |
| Nudt21 | ENSMUSG00000031754 | 68219 | 0.0506146355120558 | 3.72107477324655 | 0.59358875401375 | 0.5626701728655 | 0.650507420369417 |
| Ccnl2 | ENSMUSG00000029068 | 56036 | 0.0686717796254017 | 4.50206088444431 | 0.592949779792397 | 0.5630853757591 | 0.650918595856998 |
| Hspb2 | ENSMUSG00000038086 | 69253 | -0.0575353570070902 | 4.60215868931783 | -0.592658649002847 | 0.5632746060235 | 0.651068490900244 |
| Gsto1 | ENSMUSG00000025068 | 14873 | 0.0629349694796693 | 5.2875467117409 | 0.592178060019335 | 0.5635870558296 | 0.651360764143997 |
| Golga7 | ENSMUSG00000015341 | 57437 | 0.0469169544469477 | 4.20856034937822 | 0.591858193372392 | 0.5637950653457 | 0.651532282407652 |
| Fut10 | ENSMUSG00000046152 | 171167 | 0.0984290802118259 | 1.80204229655459 | 0.591453665599035 | 0.5640581891155 | 0.651767448754937 |
| Ccdc32 | ENSMUSG00000039983 | 269336 | -0.116482852832391 | 1.76720843926376 | -0.591087664984613 | 0.5642963098392 | 0.651973677502715 |
| Zfyve16 | ENSMUSG00000021706 | 218441 | 0.0523897716494641 | 4.05989592364485 | 0.590567087380112 | 0.5646350916064 | 0.652296151590559 |
| Phf10 | ENSMUSG00000023883 | 72057 | 0.0424326471650689 | 5.41236770542934 | 0.590224352977009 | 0.5648581960458 | 0.652484935113282 |
| Fgf11 | ENSMUSG00000042826 | 14166 | 0.142808754315809 | 1.4346028214093 | 0.589707971629897 | 0.5651944259790 | 0.652796371085255 |
| Phactr4 | ENSMUSG00000066043 | 100169 | -0.0694558147775468 | 3.69902851168069 | -0.589557218930652 | 0.5652926053898 | 0.652796371085255 |
| Txnl1 | ENSMUSG00000024583 | 53382 | 0.0510604036491253 | 5.27867543401809 | 0.58952869067445 | 0.5653111857698 | 0.652796371085255 |
| Hs6st1 | ENSMUSG00000045216 | 50785 | 0.0410914596259619 | 4.49211801864017 | 0.5894434787347 | 0.5653666860364 | 0.652796371085255 |
| Stk35 | ENSMUSG00000037885 | 67333 | 0.0761001667747632 | 2.47904326427236 | 0.588951760370038 | 0.5656870092253 | 0.653097243118098 |
| Lrwd1 | ENSMUSG00000029703 | 71735 | -0.102924477282684 | 1.43252721126903 | -0.587663284224601 | 0.5665268302091 | 0.653965436184637 |
| Samd9l | ENSMUSG00000047735 | 209086 | -0.0805214697952086 | 3.70025314075738 | -0.587614474666305 | 0.5665586570916 | 0.653965436184637 |
| Gm14391 | ENSMUSG00000090093 | 665001 | -0.0922884213822985 | 1.97537180985116 | -0.587149315903595 | 0.5668620176530 | 0.654204047404585 |
| Sdf4 | ENSMUSG00000029076 | 20318 | -0.0406960824533211 | 5.61450848115155 | -0.587113968797395 | 0.5668850733708 | 0.654204047404585 |
| Taf1c | ENSMUSG00000031832 | 21341 | 0.0889597851601909 | 1.65397981116942 | 0.586739366470691 | 0.5671294445577 | 0.654416970174262 |
| Filip1l | ENSMUSG00000043336 | 78749 | -0.0371045432523482 | 6.96957890645053 | -0.586257392331976 | 0.5674439423478 | 0.654710759320724 |
| Mfsd4a | ENSMUSG00000059149 | 213006 | -0.105389845416498 | 1.36359982438722 | -0.586053261602088 | 0.5675771698166 | 0.654795360766787 |
| Glod4 | ENSMUSG00000017286 | 67201 | 0.0616940149728609 | 3.21179271779328 | 0.584376429210768 | 0.5686721989257 | 0.655989426599572 |
| Kdm4c | ENSMUSG00000028397 | 76804 | -0.0459164914515289 | 3.51473731769941 | -0.584276367589864 | 0.5687375781868 | 0.655995617511922 |
| Ptrhd1 | ENSMUSG00000096199 | 69709 | -0.110679961620801 | 1.36731670430688 | -0.583976707229803 | 0.5689333972237 | 0.656152243622323 |
| Mbtd1 | ENSMUSG00000059474 | 103537 | 0.061336555859977 | 2.74599985230043 | 0.583515127739372 | 0.5692350958051 | 0.65643093493473 |
| Wdr7 | ENSMUSG00000040560 | 104082 | 0.0793051831736307 | 4.40474619495198 | 0.582737256598896 | 0.5697437223043 | 0.656948167564186 |
| Ints12 | ENSMUSG00000028016 | 71793 | 0.0640698218796754 | 2.61310801127612 | 0.581780265200347 | 0.5703698017056 | 0.65760070693436 |
| Ift22 | ENSMUSG00000007987 | 67286 | 0.118378389557645 | 1.44712082621683 | 0.581060531632994 | 0.5708409039629 | 0.658074448557118 |
| Poc1b | ENSMUSG00000019952 | 382406 | 0.0708897199139482 | 1.72459943053735 | 0.580520698661108 | 0.5711943876981 | 0.65841251206671 |
| Lta4h | ENSMUSG00000015889 | 16993 | 0.0540272718438488 | 3.58737784449512 | 0.580269090442474 | 0.5713591809591 | 0.658533024785425 |
| Lap3 | ENSMUSG00000039682 | 66988 | -0.0468316347061126 | 4.86720768219539 | -0.580148078105136 | 0.5714384481551 | 0.658554947473257 |
| Atp11c | ENSMUSG00000062949 | 320940 | 0.0696957269486269 | 2.68158130966696 | 0.579946659743474 | 0.5715703969668 | 0.658637572132089 |
| Sec23a | ENSMUSG00000020986 | 20334 | 0.034140847251134 | 5.41717380607543 | 0.579622057474873 | 0.5717830773402 | 0.65877080957225 |
| Foxo6 | ENSMUSG00000052135 | 329934 | -0.140411968657812 | 2.17815416489524 | -0.57957792325976 | 0.5718119974496 | 0.65877080957225 |
| Zfx | ENSMUSG00000079509 | 22764 | -0.0385789458644279 | 4.53926481626311 | -0.578889759653819 | 0.5722630351145 | 0.65913973696946 |
| Ammecr1l | ENSMUSG00000041915 | 225339 | 0.0846526671751051 | 3.01817355880994 | 0.578840697192655 | 0.5722951989348 | 0.65913973696946 |
| Cops9 | ENSMUSG00000073616 | 66915 | -0.0649111120567438 | 2.83047424259251 | -0.578821659966111 | 0.5723076794050 | 0.65913973696946 |
| Rab18 | ENSMUSG00000073639 | 19330 | -0.0349146559216758 | 5.10062964513285 | -0.578356169181377 | 0.5726128918179 | 0.659421778388316 |
| Taf1a | ENSMUSG00000072258 | 21339 | -0.102346732453733 | 1.33087603913204 | -0.578256940948679 | 0.5726779648033 | 0.659427244629291 |
| Plxna2 | ENSMUSG00000026640 | 18845 | -0.0707335603335526 | 4.68159610405783 | -0.577799619014859 | 0.5729779230205 | 0.659703146886242 |
| Ythdf2 | ENSMUSG00000040025 | 213541 | -0.0368002054085883 | 4.42601142822681 | -0.57739587006641 | 0.5732428117099 | 0.659938617672653 |
| Lyve1 | ENSMUSG00000030787 | 114332 | -0.0867761126015636 | 2.50149686936648 | -0.57686959172974 | 0.5735881858059 | 0.660266686925495 |
| Trmt5 | ENSMUSG00000034442 | 76357 | -0.0793282276802421 | 2.17726383505274 | -0.575960775138346 | 0.5741848623667 | 0.660883934982774 |
| Ylpm1 | ENSMUSG00000021244 | 56531 | 0.0353438427403061 | 5.18405845633236 | 0.575837036988386 | 0.5742661270459 | 0.660907878999708 |
| Eif2s1 | ENSMUSG00000021116 | 13665 | -0.041626040168846 | 4.66258077983461 | -0.575662704639803 | 0.5743806296256 | 0.660970066579423 |
| Clcn6 | ENSMUSG00000029016 | 26372 | 0.0833319509703485 | 2.05075996022157 | 0.5751041726108 | 0.5747475581078 | 0.661322690281792 |
| Hira | ENSMUSG00000022702 | 15260 | -0.0545443637868783 | 3.3769741057379 | -0.57460676414105 | 0.5750744354480 | 0.661629160497074 |
| Ern1 | ENSMUSG00000020715 | 78943 | -0.0626219160705555 | 3.91517598783681 | -0.574046967230723 | 0.5754424292274 | 0.661982866121324 |
| Srbd1 | ENSMUSG00000024135 | 78586 | -0.0848913461687371 | 1.99126838408495 | -0.573656184119468 | 0.5756993919078 | 0.66219013174002 |
| Hddc2 | ENSMUSG00000000295 | 69692 | -0.0922392738902555 | 1.87659063306354 | -0.57358872338244 | 0.5757437573924 | 0.66219013174002 |
| Zfp35 | ENSMUSG00000063281 | 22694 | -0.0733610028007758 | 2.34178821702807 | -0.573484595401712 | 0.5758122405869 | 0.662199221657699 |
| Sra1 | ENSMUSG00000006050 | 24068 | -0.0421651519709916 | 3.91843549299259 | -0.573065985067057 | 0.5760875967347 | 0.662390704089691 |
| Il6st | ENSMUSG00000021756 | 16195 | 0.0297026437344545 | 7.37814826980352 | 0.573047225509357 | 0.5760999381374 | 0.662390704089691 |
| Cpsf1 | ENSMUSG00000034022 | 94230 | -0.0438357919548276 | 4.10175803640771 | -0.572733052398217 | 0.5763066447602 | 0.662521362300434 |
| Angel1 | ENSMUSG00000021257 | 68737 | -0.101324611730479 | 1.81742762722921 | -0.572690272870022 | 0.5763347940824 | 0.662521362300434 |
| Cenpj | ENSMUSG00000064128 | 219103 | 0.0925386262132697 | 2.00421465358611 | 0.572564612645567 | 0.5764174838464 | 0.662546741948035 |
| Cstf2 | ENSMUSG00000031256 | 108062 | 0.0712592085171848 | 3.25363493966503 | 0.572288840294431 | 0.5765989756401 | 0.662685669511397 |

| | | | | | | | |
|---|---|---|---|---|---|---|---|
| Obscn | ENSMUSG00000061462 | 380698 | -0.0601360102514068 | 11.5429155528744 | -0.571948129474937 | 0.5768232463492 | 0.662873728537022 |
| Vmn1r46 | ENSMUSG00000061653 | 113856 | -0.162661475339025 | 1.38568632472538 | -0.57148271190142 | 0.5771296785687 | 0.663156156463254 |
| Nelfcd | ENSMUSG00000016253 | 57314 | 0.0496581606802731 | 3.56974219040956 | 0.568421900859414 | 0.5791470534840 | 0.665404292793173 |
| Cecr2 | ENSMUSG00000071226 | 330409 | -0.119551493140424 | 3.86628280239912 | -0.567422672256706 | 0.5798064419579 | 0.66609187777637 |
| Sgce | ENSMUSG00000004631 | 20392 | 0.0930440811512785 | 2.11116094064242 | 0.566739151510558 | 0.5802577216706 | 0.666540264352898 |
| Hgsnat | ENSMUSG00000037260 | 52120 | -0.0495263429397279 | 3.41829940955921 | -0.566378211108895 | 0.5804960987658 | 0.666744021814612 |
| L3mbtl2 | ENSMUSG00000022394 | 214669 | -0.0654308914271708 | 2.72624137476676 | -0.564515499441949 | 0.5817271084085 | 0.668087730827216 |
| Sumf2 | ENSMUSG00000025538 | 67902 | -0.107133607856727 | 1.33973211881032 | -0.563754708239692 | 0.5822302834091 | 0.668583203539558 |
| Sbds | ENSMUSG00000025337 | 66711 | 0.0548212077467372 | 3.9762039408341 | 0.563678253971812 | 0.5822808615585 | 0.668583203539558 |
| Atp6v1g1 | ENSMUSG00000039105 | 66290 | 0.0395021595343454 | 5.23498029546411 | 0.563421395377647 | 0.5824508025409 | 0.66870808975684 |
| Mus81 | ENSMUSG00000024906 | 71711 | -0.0604731716165188 | 2.86956872776623 | -0.563261398802384 | 0.5825566714041 | 0.668759396672781 |
| Uggt2 | ENSMUSG00000042104 | 66435 | -0.114337866328769 | 2.79626156628037 | -0.563070469560312 | 0.5826830213523 | 0.668834202079431 |
| Rictor | ENSMUSG00000050310 | 78757 | 0.0405598546379173 | 3.86692908173015 | 0.562764333138683 | 0.5828856409358 | 0.668940964222707 |
| N4bp2l2 | ENSMUSG00000029655 | 381695 | -0.048337475230805 | 4.52594486002152 | -0.56274501957276 | 0.5828984250499 | 0.668940964222707 |
| Trim12a | ENSMUSG00000066258 | 76681 | 0.0747950515185205 | 2.45108492630283 | 0.56260649720824 | 0.5829901206120 | 0.668952693187901 |
| Kras | ENSMUSG00000030265 | 16653 | -0.0544985796012458 | 4.41204459426796 | -0.562544683543707 | 0.5830310408929 | 0.668952693187901 |
| Wdr48 | ENSMUSG00000032512 | 67561 | 0.0628631728741396 | 3.41764053982631 | 0.562063652041571 | 0.5833495319439 | 0.669247872972258 |
| Lipt1 | ENSMUSG00000037216 | 623661 | -0.104001796227626 | 1.30704438875207 | -0.5608005701112 | 0.5841862490164 | 0.670137463322324 |
| Tbl1xr1 | ENSMUSG00000027630 | 81004 | 0.0426314661748926 | 5.61569218969541 | 0.560602688823037 | 0.5843173900917 | 0.670193583541484 |
| Nol11 | ENSMUSG00000018433 | 68979 | -0.0470886150676355 | 3.58419446716274 | -0.560457552690797 | 0.5844135852917 | 0.670193583541484 |
| Sesn1 | ENSMUSG00000038332 | 140742 | 0.0717859617009489 | 3.96130462546536 | 0.560449224898163 | 0.5844191051429 | 0.670193583541484 |
| Mrpl17 | ENSMUSG00000030879 | 27397 | -0.0390622499558093 | 4.33343249594002 | -0.559661405260918 | 0.5849414122852 | 0.670722183750161 |
| Rrp7a | ENSMUSG00000018040 | 74778 | 0.0458591511047461 | 3.77790092572413 | 0.559376970413817 | 0.5851300457903 | 0.670868106831788 |
| Btaf1 | ENSMUSG00000040565 | 107182 | 0.041809889308051 | 4.0203762055415 | 0.558742111233341 | 0.5855511898432 | 0.671280551250839 |
| Ran | ENSMUSG00000029430 | 19384 | 0.0422845118445343 | 5.93688678796208 | 0.557747905157148 | 0.5862110276665 | 0.671966520912676 |
| Zfp653 | ENSMUSG00000038895 | 319601 | -0.100829588425465 | 1.2702940351643 | -0.557426392703942 | 0.5864244923366 | 0.672140728293178 |
| Ccdc61 | ENSMUSG00000074358 | 232933 | -0.0878759954615984 | 1.57547501071245 | -0.556962670051669 | 0.5867324466812 | 0.672423188789287 |
| Gramd1a | ENSMUSG00000001248 | 52857 | 0.0384376909920494 | 4.6859349736688 | 0.55673794839978 | 0.5868817125124 | 0.672523744572157 |
| Cnot11 | ENSMUSG00000003135 | 52846 | 0.0504349173082321 | 3.85798905706184 | 0.556284726805019 | 0.5871828134811 | 0.672798253056832 |
| Sdf2 | ENSMUSG00000002064 | 20316 | 0.0531703268340959 | 2.87252207393225 | 0.555838049940623 | 0.5874796444397 | 0.673067811903429 |
| Colq | ENSMUSG00000057606 | 382864 | 0.0990462886035779 | 3.38063290737428 | 0.555581871904543 | 0.5876499178150 | 0.673192334063753 |
| Letmd1 | ENSMUSG00000037353 | 68614 | 0.0873066207480298 | 1.8297548607786 | 0.554108298211998 | 0.5886298494441 | 0.674244250683638 |
| Fzd5 | ENSMUSG00000045005 | 14367 | 0.0917960537360345 | 1.72463294953296 | 0.55332531671985 | 0.5891508770730 | 0.674770351769222 |
| Med22 | ENSMUSG00000015776 | 20933 | -0.0597459753646565 | 2.51980647393107 | -0.552729898917429 | 0.5895472510094 | 0.675083765083785 |
| Vps11 | ENSMUSG00000032127 | 71732 | -0.0470764786035396 | 3.90046350201133 | -0.55272871444654 | 0.5895480396569 | 0.675083765083785 |
| Bbs1 | ENSMUSG00000006464 | 52028 | 0.0885625854469008 | 1.51733014550539 | 0.552625479990404 | 0.5896167775747 | 0.675091756119132 |
| Cdv3 | ENSMUSG00000032803 | 321022 | -0.0430395449435466 | 8.58656368574645 | -0.552248137853277 | 0.5898680631653 | 0.675308734652382 |
| Zswim1 | ENSMUSG00000017764 | 71971 | -0.0718094686591917 | 1.94490519683557 | -0.551848131798629 | 0.5901345015485 | 0.675543013554029 |
| Scarna13 | ENSMUSG00000088789 | 100306943 | 0.0652912245924671 | 3.84276976597763 | 0.551441861773943 | 0.5904051755379 | 0.675760787321657 |
| Eef2kmt | ENSMUSG00000022544 | 70511 | 0.0626326073719983 | 2.38182543782576 | 0.551377017378061 | 0.5904483834698 | 0.675760787321657 |
| Rgl2 | ENSMUSG00000041354 | 19732 | -0.0752784978168712 | 3.1458803711779 | -0.551053965909113 | 0.5906636673663 | 0.675927950067304 |
| Rara | ENSMUSG00000037992 | 19401 | 0.0433785349674842 | 4.27394755849625 | 0.550972273369729 | 0.5907181142615 | 0.675927950067304 |
| Dvl3 | ENSMUSG00000003233 | 13544 | 0.0605817183102255 | 5.45075718919414 | 0.550334414239866 | 0.5911433266753 | 0.676343699381262 |
| Arrdc4 | ENSMUSG00000042659 | 66412 | 0.0716051023810897 | 2.82265388527007 | 0.549850515933345 | 0.5914660097187 | 0.676642067214623 |
| Pik3r4 | ENSMUSG00000032571 | 75669 | 0.0505565161973318 | 3.85056702130072 | 0.549749661395616 | 0.5915332749860 | 0.676648203105136 |
| Gbf1 | ENSMUSG00000025224 | 107338 | 0.0298277450996638 | 5.7559307494883 | 0.54943107442612 | 0.5917457833620 | 0.676820462271736 |
| Psmb6 | ENSMUSG00000018286 | 19175 | -0.0412390428639613 | 5.50248697133612 | -0.548969576747546 | 0.5920536873878 | 0.677101784561315 |
| Dguok | ENSMUSG00000014554 | 27369 | -0.0699492985421317 | 2.34355558258309 | -0.548434253371695 | 0.5924109494781 | 0.677439490401258 |
| Mrgpre | ENSMUSG00000048965 | 244238 | -0.150787676668523 | 2.0927129009171 | -0.547690327275832 | 0.5929076111222 | 0.6779365164411 |
| B3gntl1 | ENSMUSG00000046605 | 210004 | 0.096875414529904 | 1.62325860086195 | 0.547543632432892 | 0.5930055729865 | 0.677977608860622 |
| Dnase1l1 | ENSMUSG00000019088 | 69537 | -0.0575696387571529 | 3.59986068435026 | -0.547096386161236 | 0.5933042921872 | 0.678200078912552 |
| Gpr180 | ENSMUSG00000022131 | 58245 | -0.0523139729113127 | 3.21810995676203 | -0.547046670536649 | 0.5933375023788 | 0.678200078912552 |
| Pced1a | ENSMUSG00000037773 | 319513 | -0.103356229574998 | 1.36350157593497 | -0.546947080545603 | 0.5934040316578 | 0.678200078912552 |
| Gnptg | ENSMUSG00000035521 | 214505 | 0.0835756068542897 | 2.07631293260611 | 0.54682886804134 | 0.5934830063117 | 0.678200078912552 |
| Rps11 | ENSMUSG00000003429 | 27207 | -0.0430521797991554 | 5.50022895573391 | -0.546787897074563 | 0.5935103791855 | 0.678200078912552 |
| Rbbp9 | ENSMUSG00000027428 | 26450 | -0.0720461736708335 | 2.3340412088817 | -0.546686661536853 | 0.5935780178346 | 0.678206471511531 |
| Ttc21b | ENSMUSG00000034848 | 73668 | -0.0886872004219457 | 1.81633231743642 | -0.544963432465957 | 0.5947299646073 | 0.679451634941778 |
| Isg15 | ENSMUSG00000035692 | 100038882 | -0.230329903684203 | 0.487004512105161 | -0.544240798418624 | 0.5952133708677 | 0.679913921306496 |
| Mcts2 | ENSMUSG00000042814 | 66405 | -0.0928904050871042 | 1.61892439784563 | -0.544172590421041 | 0.5952590089564 | 0.679913921306496 |
| Stk16 | ENSMUSG00000026201 | 20872 | 0.0561270943571324 | 2.79315774491911 | 0.544048354617224 | 0.5953421399231 | 0.679937825880202 |
| Phf5a | ENSMUSG00000061360 | 68479 | -0.0503837048279445 | 2.94162487084472 | -0.54383037657499 | 0.5954880117115 | 0.680033374009555 |
| Vmn2r52 | ENSMUSG00000091930 | 384534 | 0.274495406460242 | -0.465268970184555 | 0.543682776515566 | 0.5955867965934 | 0.680075135648522 |
| Dhx37 | ENSMUSG00000029480 | 208144 | -0.0524587034322748 | 3.23883045688891 | -0.542286130554834 | 0.5965219478016 | 0.681039103689788 |
| Ddx19b | ENSMUSG00000033658 | 234733 | 0.0443237495438895 | 3.82855117412753 | 0.542235870762782 | 0.5965556141093 | 0.681039103689788 |
| Tent4b | ENSMUSG00000036779 | 214627 | -0.0404421632406242 | 3.83825918643884 | -0.54176273778742 | 0.5968725875014 | 0.681227498064379 |
| Taf15 | ENSMUSG00000020680 | 70439 | -0.0444880412428899 | 3.3220581387902 | -0.541731232475986 | 0.5968936973837 | 0.681227498064379 |
| Ggt5 | ENSMUSG00000006344 | 23887 | -0.121935239193474 | 1.11436757850349 | -0.541710483794772 | 0.5969076000787 | 0.681227498064379 |
| Abca7 | ENSMUSG00000035722 | 27403 | -0.0538590605817371 | 3.26804023882117 | -0.541285136448467 | 0.5971926411190 | 0.681481653624811 |
| Zfp330 | ENSMUSG00000031711 | 30932 | 0.0487203961945708 | 3.63766090685336 | 0.541039143418882 | 0.5973575216803 | 0.681598650259665 |
| Dedd | ENSMUSG00000013973 | 21945 | -0.0474873031615326 | 3.7110618526224 | -0.540902195303961 | 0.5974493232337 | 0.681610678736148 |
| Paxx | ENSMUSG00000047617 | 227622 | 0.0787907318999828 | 1.93094345320408 | 0.540837378344722 | 0.5974927750114 | 0.681610678736148 |
| Nop2 | ENSMUSG00000038279 | 110109 | 0.0462808270287702 | 3.43090172402919 | 0.540478710165826 | 0.5977332467471 | 0.681802791688512 |
| Nktr | ENSMUSG00000032525 | 18087 | 0.0495845054346287 | 5.34610996181866 | 0.540400148033492 | 0.5977859258569 | 0.681802791688512 |
| Elk1 | ENSMUSG00000009406 | 13712 | 0.0618425180793825 | 2.8648921667518 | 0.539992395662285 | 0.5980593782114 | 0.682035493377331 |
| Faf2 | ENSMUSG00000025873 | 76577 | 0.0632052292332799 | 3.60497988964459 | 0.53990985340135 | 0.5981147415163 | 0.682035493377331 |
| Ncoa6 | ENSMUSG00000038369 | 56406 | 0.0419012142791582 | 4.82922810198567 | 0.53973291472125 | 0.5982334277706 | 0.682059448378801 |
| Dnlz | ENSMUSG00000075467 | 52838 | -0.0492418712353717 | 3.975302321563 | -0.539692494164844 | 0.5982605425904 | 0.682059448378801 |
| Rps6kb1 | ENSMUSG00000020516 | 72508 | -0.0468987807588759 | 4.05048312423235 | -0.538897068671594 | 0.5987942545356 | 0.682596725031753 |
| Prdm4 | ENSMUSG00000035529 | 72843 | 0.0383458802345982 | 3.81953335843551 | 0.538743340689782 | 0.5988974301844 | 0.682643150088254 |
| Slc35e2 | ENSMUSG00000042202 | 320541 | -0.0768013653581585 | 2.43203486057677 | -0.538507780736696 | 0.5990555453916 | 0.682752180865016 |
| Fam133b | ENSMUSG00000058503 | 68152 | -0.0676908824307878 | 3.11017199040853 | -0.538140140364865 | 0.5993023592641 | 0.682962269507591 |
| Fzd7 | ENSMUSG00000041075 | 14369 | -0.1215201522599 | 5.10673529877857 | -0.537435095492026 | 0.5997758319781 | 0.683420831428795 |
| Prpf31 | ENSMUSG00000008373 | 68988 | 0.0445804825004239 | 3.62228435410741 | 0.53735476179608 | 0.5998297920343 | 0.683420831428795 |
| Eif4h | ENSMUSG00000040731 | 22384 | 0.032353382893989 | 5.43231831700383 | 0.537113933466643 | 0.5999915708444 | 0.683533909421635 |
| Tmem120a | ENSMUSG00000039886 | 215210 | -0.0911527793940059 | 2.36275679078733 | -0.536940880082624 | 0.6001078347592 | 0.683595116897967 |
| Bloc1s5 | ENSMUSG00000038982 | 17828 | -0.065064170820921 | 2.62369542667013 | -0.536245688705016 | 0.6005750055563 | 0.684050990963629 |
| Snrpf | ENSMUSG00000020018 | 69878 | -0.0532220768820384 | 3.74874483270036 | -0.536159122666017 | 0.6006331910409 | 0.684050990963629 |
| Tpst2 | ENSMUSG00000029344 | 22022 | -0.0613767365159745 | 2.82921460053235 | -0.535219466763001 | 0.6012649648146 | 0.684699169745673 |
| Psmb1 | ENSMUSG00000014769 | 19170 | -0.0415597639389249 | 5.65085927386035 | -0.534826556556276 | 0.6015292355364 | 0.684900077799158 |
| Ankrd49 | ENSMUSG00000031931 | 56503 | -0.0663941291897867 | 2.31574512018133 | -0.534718874288387 | 0.6016016726459 | 0.684900077799158 |
| Hnrnpk | ENSMUSG00000021546 | 15387 | 0.0327771659112068 | 7.07050152567932 | 0.534657439291827 | 0.6016430015051 | 0.684900077799158 |
| Cep85l | ENSMUSG00000038594 | 100038725 | 0.145850540250357 | 6.06519288357225 | 0.534518720663618 | 0.6017363262506 | 0.684900077799158 |
| Cse1l | ENSMUSG00000002718 | 110750 | -0.0410636341185491 | 5.21332499577913 | -0.534400102739362 | 0.6018161337633 | 0.684900077799158 |
| Bcl6 | ENSMUSG00000022508 | 12053 | -0.0908860300828049 | 5.5763408598099 | -0.534336565184004 | 0.6018588847556 | 0.684900077799158 |
| Sf3a3 | ENSMUSG00000028902 | 75062 | -0.0446884257081059 | 3.46131986693635 | -0.534253720388109 | 0.6019146288419 | 0.684900077799158 |
| Trim26 | ENSMUSG00000024457 | 22670 | -0.0559759507369266 | 3.6279496209463 | -0.534212085959048 | 0.6019426445354 | 0.684900077799158 |
| Cnppd1 | ENSMUSG00000033159 | 69171 | 0.0665855754245332 | 2.82203148182671 | 0.533924440894955 | 0.6021362180234 | 0.685049021566858 |
| Nudcd3 | ENSMUSG00000053838 | 209586 | -0.0421547109749927 | 5.90033439027126 | -0.533645380269579 | 0.6023240444294 | 0.685191396571771 |
| Spryd7 | ENSMUSG00000021930 | 66674 | 0.0631172131394049 | 4.23572670304848 | 0.532450595627006 | 0.6031285468178 | 0.685969017630467 |
| Med19 | ENSMUSG00000027080 | 381379 | -0.064376561299353 | 2.38830511677798 | -0.53244379344879 | 0.6031331285743 | 0.685969017630467 |
| Tpr | ENSMUSG00000006005 | 108989 | -0.0427732788490118 | 7.38108777962866 | -0.531708686240018 | 0.6036283790664 | 0.686460862627526 |
| Thoc1 | ENSMUSG00000024287 | 225160 | -0.0513731174460048 | 3.43065147573734 | -0.53157801084292 | 0.6037164379751 | 0.686489585301814 |
| Tmem30a | ENSMUSG00000032328 | 69981 | -0.0386127352068515 | 5.80354919085718 | -0.531433836492526 | 0.6038136009632 | 0.686528653227499 |
| Nelfe | ENSMUSG00000024369 | 27632 | 0.0984614435351052 | 1.87987626826028 | 0.530932216905751 | 0.6041517169931 | 0.68684164518481 |
| Dnmt3a | ENSMUSG00000020661 | 13435 | 0.0530020792776069 | 6.30131513788577 | 0.530504385534699 | 0.6044401710723 | 0.687080264236952 |
| Mfsd14a | ENSMUSG00000089911 | 15247 | 0.057256099605608 | 3.23460029103082 | 0.530434463724597 | 0.6044873205714 | 0.687080264236952 |
| Tom1l2 | ENSMUSG00000000538 | 216810 | -0.0304624558059068 | 5.9389904899912 | -0.529860278504961 | 0.6048745733062 | 0.687448945811095 |
| Plvap | ENSMUSG00000034845 | 84094 | 0.0438219393684243 | 3.79969635854422 | 0.529509227056538 | 0.6051113970914 | 0.687646603764059 |
| Qsox2 | ENSMUSG00000036327 | 227638 | 0.0758642760185149 | 1.90640492068906 | 0.529370547746089 | 0.6052049646936 | 0.687680450014586 |
| Wdr34 | ENSMUSG00000039715 | 71820 | -0.0808922608615672 | 1.48369532439784 | -0.529278604872988 | 0.6052670029814 | 0.687680450014586 |
| Kdm5c | ENSMUSG00000025332 | 20591 | 0.0360769002421542 | 4.67137787263312 | 0.529075000375153 | 0.6054043960592 | 0.687703427074321 |
| Plekhg2 | ENSMUSG00000037552 | 101497 | 0.0795479637117271 | 3.35677882422956 | 0.529062172809884 | 0.6054130526700 | 0.687703427074321 |
| Fen1 | ENSMUSG00000024742 | 14156 | -0.0757499497797436 | 1.4030337855634 | -0.528565551991561 | 0.6057482419904 | 0.688012680091152 |
| Xrn1 | ENSMUSG00000032410 | 24127 | 0.0453035294652118 | 4.27541607333279 | 0.528272540762268 | 0.6059460504819 | 0.688117053975988 |
| Edrf1 | ENSMUSG00000039990 | 214764 | 0.0559569130966397 | 2.82806956252548 | 0.52822300019335 | 0.6059794979329 | 0.688117053975988 |
| Aox1 | ENSMUSG00000063558 | 11761 | -0.0814687915528207 | 5.08628543974717 | -0.528149699203198 | 0.6060289889879 | 0.688117053975988 |

| | | | | | | | |
|---|---|---|---|---|---|---|---|
| Eif5a | ENSMUSG00000078812 | 276770 | 0.0361929916061419 | 6.88801317601348 | 0.527796538652461 | 0.6062674627119 | 0.688316331003791 |
| Tmx4 | ENSMUSG00000034723 | 52837 | 0.0331028280152861 | 4.2803378294681 | 0.527473230395723 | 0.6064858195330 | 0.68849272959972 |
| Knop1 | ENSMUSG00000030980 | 66356 | -0.042354321347843 | 3.97396490135665 | -0.527120113124967 | 0.6067243536419 | 0.68862550601789 |
| AU022252 | ENSMUSG00000078584 | 230696 | -0.0468830694290565 | 4.80614755397113 | -0.527113567349914 | 0.6067287758172 | 0.68862550601789 |
| Cope | ENSMUSG00000055681 | 59042 | 0.0383045333026706 | 5.00117068980913 | 0.526550847019895 | 0.6071089967837 | 0.688985511196313 |
| Fdx2 | ENSMUSG00000079677 | 68165 | 0.0534146660542731 | 3.0703836035087 | 0.526028888596953 | 0.6074617816459 | 0.689314308644392 |
| Ss18l1 | ENSMUSG00000039086 | 269397 | 0.0866194230823691 | 1.62551998395312 | 0.525407990914517 | 0.6078815710838 | 0.689719063059727 |
| Ppp6r1 | ENSMUSG00000052296 | 243819 | -0.0324268066973712 | 6.31706553210483 | -0.524895138681371 | 0.6082284196656 | 0.690040981356043 |
| Ufl1 | ENSMUSG00000040359 | 67490 | 0.0371762647847369 | 4.89224652783192 | 0.523832518182218 | 0.6089473964789 | 0.690784972074628 |
| Sav1 | ENSMUSG00000021067 | 64010 | 0.0613527833822456 | 4.44740775874148 | 0.523671448688543 | 0.6090564140846 | 0.690836947427581 |
| Pus1 | ENSMUSG00000029507 | 56361 | 0.0602930327918725 | 2.52443039947714 | 0.523504834390706 | 0.6091691948020 | 0.69089317995237 |
| Cptp | ENSMUSG00000029073 | 79554 | 0.0889180717783078 | 1.74642304898713 | 0.522733228383229 | 0.6096916276340 | 0.69141396220228 |
| Trpm3 | ENSMUSG00000052387 | 226025 | 0.0968017317302576 | 1.3604492735805 | 0.522595901768447 | 0.6097846309150 | 0.691447697150718 |
| Nlk | ENSMUSG00000017376 | 18099 | -0.0614935868508937 | 2.74449347798126 | -0.522228518704623 | 0.6100334725224 | 0.691658115148117 |
| Sim2 | ENSMUSG00000062713 | 20465 | -0.0525314585229983 | 5.25550162644819 | -0.521851699221208 | 0.6102887579919 | 0.691830187000288 |
| Rcan3 | ENSMUSG00000059713 | 53902 | 0.0663297815104493 | 2.43192047207559 | 0.521817663412456 | 0.6103118189775 | 0.691830187000288 |
| C1qtnf12 | ENSMUSG00000023571 | 67389 | 0.0836570414793425 | 1.33128649480871 | 0.521090947032874 | 0.6108043090629 | 0.692316663455686 |
| Utp14a | ENSMUSG00000063785 | 72554 | -0.0433067717887496 | 3.7814054031364 | -0.520828753550842 | 0.6109820438032 | 0.692446316310336 |
| Acox3 | ENSMUSG00000029098 | 80911 | 0.0461469510758481 | 3.66094962383665 | 0.519812647610607 | 0.6116710793583 | 0.693155356465453 |
| Psmd1 | ENSMUSG00000026229 | 70247 | 0.036075883119036 | 6.69688573781197 | 0.519362102870876 | 0.6119767227472 | 0.6933494459476 |
| Inhba | ENSMUSG00000041324 | 16323 | 0.101951452131768 | 3.02346906970761 | 0.519183760088751 | 0.6120977288696 | 0.6933494459476 |
| Pwp2 | ENSMUSG00000032834 | 110816 | 0.0508255235241575 | 3.07785731586833 | 0.519148308763787 | 0.6121217841083 | 0.6933494459476 |
| Tmx2 | ENSMUSG00000050043 | 66958 | -0.0357825486555443 | 4.16877841316291 | -0.519134054237726 | 0.6121314565447 | 0.6933494459476 |
| Usp6nl | ENSMUSG00000039046 | 98910 | 0.0531187089810832 | 2.86994358420054 | 0.519092725357811 | 0.6121595007629 | 0.6933494459476 |
| Pex5 | ENSMUSG00000005069 | 19305 | 0.0419013927769534 | 4.01865098065366 | 0.517812008627647 | 0.6130288607672 | 0.69426217126467 |
| Sec62 | ENSMUSG00000027706 | 69276 | -0.0319914577536045 | 5.50302905531246 | -0.517479537523473 | 0.6132546439970 | 0.694445924948918 |
| Cyfip1 | ENSMUSG00000030447 | 20430 | -0.0458630909720865 | 4.89841420861007 | -0.516307896872443 | 0.6140506387201 | 0.695275277796788 |
| Rtcb | ENSMUSG00000001783 | 28088 | -0.0393022597986663 | 4.46063182384275 | -0.515790157436217 | 0.6144025446979 | 0.695529902953792 |
| AW146154 | ENSMUSG00000074166 | 101835 | 0.0907820855564487 | 1.51618061017663 | 0.51566401029007 | 0.6144883015505 | 0.695529902953792 |
| Nudt18 | ENSMUSG00000045211 | 213484 | -0.10107065280753 | 1.1926973226839 | -0.515653873985319 | 0.6144951926282 | 0.695529902953792 |
| Dvl1 | ENSMUSG00000029071 | 13542 | -0.038723636257509 | 5.97036801146821 | -0.515540620024585 | 0.6145721899184 | 0.695529902953792 |
| Rpl6 | ENSMUSG00000029614 | 19988 | 0.0395749072360152 | 7.39967282134452 | 0.51550903656896 | 0.6145936632175 | 0.695529902953792 |
| Gm10389 | ENSMUSG00000115667 | 100038608 | -0.132848101513423 | 1.37196876001712 | -0.515191693722162 | 0.6148094421258 | 0.695702071622932 |
| Umps | ENSMUSG00000022814 | 22247 | -0.0682971001507848 | 2.15837590324136 | -0.514083492573603 | 0.6155632607208 | 0.69642254347458 |
| Rragc | ENSMUSG00000028646 | 54170 | 0.0498961497540229 | 4.13264717852722 | 0.514068351916473 | 0.6155735628077 | 0.69642254347458 |
| Bbx | ENSMUSG00000022641 | 70508 | 0.0513748256539186 | 4.06128223421613 | 0.513881546780588 | 0.6157006767096 | 0.696494266492145 |
| Jun | ENSMUSG00000052684 | 16476 | 0.0429591336742696 | 6.99277898647703 | 0.513742070890578 | 0.6157955932171 | 0.696529556028244 |
| Ssu72 | ENSMUSG00000029038 | 68991 | -0.0506695670487076 | 4.46215916296303 | -0.513212154980747 | 0.6161562783541 | 0.696865419419449 |
| Gcfc2 | ENSMUSG00000035125 | 330361 | -0.112097900662776 | 1.21824432087183 | -0.513050754692601 | 0.6162661553570 | 0.696917581647835 |
| Itgb1bp1 | ENSMUSG00000062352 | 16413 | -0.0483446631467399 | 2.90946852346868 | -0.512504359539461 | 0.6166381977449 | 0.697266177671498 |
| Kat6b | ENSMUSG00000021767 | 54169 | 0.0341680423497177 | 5.11832178645426 | 0.512136392794473 | 0.6168888094869 | 0.697477407581874 |
| Cdk13 | ENSMUSG00000041297 | 69562 | 0.0274386709892482 | 5.16190928128897 | 0.512029846805595 | 0.6169613842414 | 0.697487319386013 |
| Aggf1 | ENSMUSG00000021681 | 66549 | 0.029097695185067 | 4.89230724682951 | 0.510780143001315 | 0.6178129425566 | 0.698377794734926 |
| Psmg3 | ENSMUSG00000029551 | 66506 | -0.0780359619902899 | 1.67658638803158 | -0.510472111092618 | 0.6180229260850 | 0.698542922659061 |
| Ift43 | ENSMUSG00000007867 | 76411 | 0.0895951334976004 | 1.81710210613733 | 0.509282337748998 | 0.6188343142679 | 0.699387705672324 |
| Pde6d | ENSMUSG00000026239 | 18582 | 0.0651644990897871 | 2.45708076272589 | 0.509115221659153 | 0.6189483237557 | 0.699444239323336 |
| Mzt2 | ENSMUSG00000022671 | 72083 | 0.0788739678120911 | 1.69066983220288 | 0.50889917239009 | 0.6190957314612 | 0.699471490392977 |
| Gpaa1 | ENSMUSG00000022561 | 14731 | -0.0605030096140182 | 2.68857754842421 | -0.508892303885456 | 0.6191004180360 | 0.699471490392977 |
| Chmp5 | ENSMUSG00000028419 | 76959 | 0.0494987021016144 | 3.88002357490016 | 0.508776902455039 | 0.6191791622755 | 0.69948815862274 |
| Gstp1 | ENSMUSG00000060803 | 14870 | -0.0403370649250556 | 5.06052300485027 | -0.508448658815419 | 0.6194031661788 | 0.699668906634429 |
| Zfp326 | ENSMUSG00000029290 | 54367 | 0.0684372099836268 | 2.67507238950246 | 0.50774316180053 | 0.6198847531708 | 0.700140549381182 |
| Pin4 | ENSMUSG00000079480 | 69713 | -0.0729961420075145 | 2.00892398599566 | -0.50750589902111 | 0.6200467545835 | 0.700251169992057 |
| Cpsf4 | ENSMUSG00000029625 | 54188 | 0.0881680230308521 | 1.40476195530587 | 0.507281204583094 | 0.6202001933811 | 0.700352098538159 |
| BC005561 | ENSMUSG00000079065 | 100042165 | 0.0652604983660554 | 3.12599239339878 | 0.507177548177477 | 0.6202709842345 | 0.700359686878173 |
| Lclat1 | ENSMUSG00000054469 | 225010 | -0.0358996382703602 | 3.93253960261211 | -0.506931551069338 | 0.6204390006064 | 0.700477041482053 |
| March7 | ENSMUSG00000026977 | 57438 | -0.0320714054872653 | 4.57432874791753 | -0.50678262117441 | 0.6205407306719 | 0.700519542184747 |
| Hnrnpa3 | ENSMUSG00000059005 | 229279 | -0.0479485039487964 | 6.25483241112 | -0.506303546077821 | 0.6208680290790 | 0.70081664868479 |
| Nufip2 | ENSMUSG00000037857 | 68564 | 0.0360664477138357 | 5.32173459308298 | 0.505634411102325 | 0.6213253142528 | 0.701260403727181 |
| Sprtn | ENSMUSG00000031986 | 244666 | 0.0629278726796425 | 3.02102933297896 | 0.505392999073195 | 0.6214903346947 | 0.701374235860865 |
| Strbp | ENSMUSG00000026915 | 20744 | -0.053829034566768 | 2.92325630139112 | -0.504680081915881 | 0.6219777827214 | 0.701851878076592 |
| Btf3 | ENSMUSG00000021660 | 218490 | 0.0310943116174629 | 5.64267982352856 | 0.503688059250313 | 0.6226563733398 | 0.702545088457629 |
| Casq1 | ENSMUSG00000007122 | 12372 | 0.044771259226275 | 9.09392923887462 | 0.503458760516686 | 0.6228132754759 | 0.70264959378965 |
| Anapc15 | ENSMUSG00000030649 | 75430 | -0.0583212413778562 | 2.94085900263858 | -0.503231227955753 | 0.6229689879582 | 0.702752735538811 |
| Fam192a | ENSMUSG00000031774 | 102122 | -0.0397622271529934 | 3.46464143440874 | -0.502913980708315 | 0.6231861283004 | 0.702925143788335 |
| Eif3i | ENSMUSG00000028798 | 54709 | -0.0366670657450658 | 5.26374022069979 | -0.502692625839928 | 0.6233376566083 | 0.703023516754578 |
| 2700081O15Rik | ENSMUSG00000053080 | 108899 | -0.0699932886467036 | 2.86764547600401 | -0.502419061518372 | 0.6235249494287 | 0.70316220181634 |
| Tsr3 | ENSMUSG00000015126 | 68327 | -0.0573547912263916 | 2.31756762959965 | -0.501992289925468 | 0.6238171881499 | 0.703386083793777 |
| Rnf125 | ENSMUSG00000033107 | 67664 | 0.13536318901464 | 1.29306057661651 | 0.50194120893389 | 0.6238521711079 | 0.703386083793777 |
| Crkl | ENSMUSG00000006134 | 12929 | 0.0302510132808278 | 4.48087576481383 | 0.501202091898638 | 0.6243584632994 | 0.703884319546872 |
| Pqlc2 | ENSMUSG00000028744 | 212555 | 0.0982862896865997 | 1.0911460923539 | 0.500981693404797 | 0.6245094736891 | 0.703981959048831 |
| Nt5m | ENSMUSG00000032615 | 103850 | -0.0575104683107606 | 2.2843965167959 | -0.500777677592103 | 0.6246492748377 | 0.704066944034967 |
| Tada1 | ENSMUSG00000026563 | 27878 | 0.0887986338839346 | 1.25861374118484 | 0.50026443165624 | 0.6250010415095 | 0.704390801602288 |
| Zfp384 | ENSMUSG00000038346 | 269800 | 0.0379969361305985 | 4.17949170046778 | 0.499845845503706 | 0.6252880011486 | 0.704641560882091 |
| Zfp773 | ENSMUSG00000063535 | 76373 | -0.0817963481465647 | 2.93809629722431 | -0.499227195381351 | 0.6257122296240 | 0.705046941760655 |
| Svil | ENSMUSG00000024236 | 225115 | 0.0433214225637002 | 8.90012163241827 | 0.498584798170062 | 0.6261528885580 | 0.70547075085837 |
| Wwox | ENSMUSG00000004637 | 80707 | -0.0886263920328412 | 1.68597835035325 | -0.498414977949091 | 0.6262694033692 | 0.705529305186998 |
| Lrp3 | ENSMUSG00000001802 | 435965 | 0.0553030108256105 | 2.59536251815215 | 0.497708090754941 | 0.6267545152655 | 0.706003050944772 |
| Tiprl | ENSMUSG00000040843 | 226591 | -0.0410969003989994 | 3.40206814867292 | -0.496625602342731 | 0.6274977381360 | 0.706767416338485 |
| Pi4ka | ENSMUSG00000041720 | 224020 | 0.0340666991485685 | 4.31892244000226 | 0.49589512344127 | 0.6279995140737 | 0.707259704135557 |
| Abhd17a | ENSMUSG00000003346 | 216169 | -0.0469639362426397 | 4.10232992593116 | -0.494925387223486 | 0.6286659354810 | 0.707937296872699 |
| Tmem9b | ENSMUSG00000031021 | 56786 | -0.0452129500389526 | 4.01091764104897 | -0.494187463086259 | 0.6291732773959 | 0.708435629915073 |
| Myl12b | ENSMUSG00000034868 | 67938 | -0.0836030211436296 | 3.41699005376065 | -0.493948846573222 | 0.6293373742113 | 0.708496645853375 |
| Zbtb34 | ENSMUSG00000068966 | 241311 | 0.06953125284282 | 1.95855477408911 | 0.493920168741217 | 0.6293570973594 | 0.708496645853375 |
| Frat1 | ENSMUSG00000067199 | 14296 | -0.151049237516302 | 1.18538686941112 | -0.493395663800626 | 0.6297178770663 | 0.708758791494338 |
| Ctnnbl1 | ENSMUSG00000027649 | 66642 | 0.0465648430629146 | 2.70835792978317 | 0.493393101428483 | 0.6297196398310 | 0.708758791494338 |
| Zbp1 | ENSMUSG00000027514 | 58203 | 0.284473406091221 | 0.457160243941647 | 0.492897616158344 | 0.6300605493960 | 0.709069480639133 |
| Rps19 | ENSMUSG00000040952 | 20085 | -0.0375710588642191 | 5.89344196387382 | -0.492435215680371 | 0.6303787748580 | 0.70934424276641 |
| Tmco4 | ENSMUSG00000041143 | 77056 | -0.0627245992271517 | 2.80189982253309 | -0.492354281896517 | 0.6304344816097 | 0.70934424276641 |
| Nkapd1 | ENSMUSG00000059820 | 270156 | 0.0552477025216005 | 3.06959123111947 | 0.492017846168539 | 0.6306660755498 | 0.709466367609213 |
| Adam17 | ENSMUSG00000052593 | 11491 | 0.0377703869905011 | 4.49993662981539 | 0.492008036302271 | 0.6306728290237 | 0.709466367609213 |
| Rnpep | ENSMUSG00000041926 | 215615 | -0.0468152339747836 | 3.28859305849059 | -0.491738736201422 | 0.6308582385840 | 0.709553141516867 |
| Zfp763 | ENSMUSG00000067430 | 73451 | -0.0806171368619328 | 1.39122985042245 | -0.49170743633629 | 0.6308797898090 | 0.709553141516867 |
| Cpsf6 | ENSMUSG00000055531 | 432508 | 0.0329126784368139 | 4.59982961723143 | 0.491307695587001 | 0.6311550582898 | 0.709751852045283 |
| Mturn | ENSMUSG00000038065 | 68235 | -0.0775426894885776 | 2.89061026703827 | -0.491262291112436 | 0.6311863282162 | 0.709751852045283 |
| Jrkl | ENSMUSG00000079083 | 77532 | 0.0702255684337532 | 2.42605764600417 | 0.491078769167292 | 0.6313127267354 | 0.709820964404924 |
| Lysmd3 | ENSMUSG00000035840 | 80289 | 0.0532807884371556 | 3.23249353846859 | 0.490856906414161 | 0.6314655480783 | 0.70985523456693 |
| Rnf19a | ENSMUSG00000022280 | 30945 | 0.0365026271272582 | 4.4353612246322 | 0.490845963964801 | 0.6314730858044 | 0.70985523456693 |
| Hps3 | ENSMUSG00000027615 | 12807 | 0.0669559886240401 | 2.2928693979956 | 0.490416319288498 | 0.6317690810986 | 0.710114943495025 |
| Met | ENSMUSG00000009376 | 17295 | 0.0369114878077039 | 3.6760534795648 | 0.490259533601851 | 0.6318771119307 | 0.710163346752567 |
| Abcb1b | ENSMUSG00000028970 | 18669 | -0.0414949008045515 | 3.34471405848017 | -0.490087220990465 | 0.6319958514698 | 0.710214333228333 |
| Tecr | ENSMUSG00000031708 | 106529 | -0.0371459670214271 | 6.90436253592086 | -0.489954792215943 | 0.6320871145017 | 0.710214333228333 |
| Myh7b | ENSMUSG00000074652 | 668940 | 0.049966896374209 | 5.22076401776589 | 0.489875796761665 | 0.6321415570419 | 0.710214333228333 |
| Zfp709 | ENSMUSG00000056019 | 236193 | -0.0947678277450719 | 1.25739004397118 | -0.489816582986019 | 0.6321823677899 | 0.710214333228333 |
| Mettl8 | ENSMUSG00000041975 | 228019 | 0.0717472528205727 | 1.43550569557015 | 0.48940804380169 | 0.6324639712779 | 0.710457678557067 |
| Dctn4 | ENSMUSG00000024603 | 67665 | 0.0327753107016182 | 4.83119033492438 | 0.489239113842981 | 0.6325804310103 | 0.710515484111586 |
| Nfib | ENSMUSG00000008575 | 18028 | 0.0383629830070982 | 7.37918595882699 | 0.489023653515506 | 0.6327289833647 | 0.710609320575397 |
| Pou6f1 | ENSMUSG00000009739 | 19009 | -0.0394715509990737 | 3.2475547815076 | -0.48794132024089 | 0.6334754641868 | 0.711374594105835 |
| Idh1 | ENSMUSG00000025950 | 15926 | -0.0570860894766989 | 4.14819546254384 | -0.486861121956582 | 0.6342208876443 | 0.712138523147496 |
| Dus3l | ENSMUSG00000007603 | 224907 | -0.0487895931773137 | 3.02794624620735 | -0.485784796781665 | 0.6349640501855 | 0.712899756838371 |
| Sun1 | ENSMUSG00000036817 | 77053 | -0.0388037933920433 | 4.6875384825111 | -0.485676309853143 | 0.6350389791598 | 0.71291065843655 |
| Smad5 | ENSMUSG00000021540 | 17129 | 0.0283318530898233 | 4.95943170609901 | 0.485494289734223 | 0.6351647048767 | 0.712978577634772 |
| Wwp2 | ENSMUSG00000031930 | 66894 | 0.0362489901297085 | 4.1120270206787 | 0.484137404697831 | 0.6361023078642 | 0.713957729465462 |
| Virma | ENSMUSG00000040720 | 66185 | 0.0378908432411897 | 4.46667622500582 | 0.483914842061614 | 0.6362561600521 | 0.714057092970197 |
| Aldh1a2 | ENSMUSG00000013584 | 19378 | 0.0724099955180826 | 2.95869997349908 | 0.483808868865512 | 0.6363294229201 | 0.714066001636444 |
| Thsd7a | ENSMUSG00000032625 | 330267 | -0.0770845769696902 | 1.86700681341177 | -0.483409181348918 | 0.6366057761625 | 0.714302785796889 |
| Pls3 | ENSMUSG00000016382 | 102866 | 0.0567688334808754 | 4.68054724174571 | 0.482253042268662 | 0.6374054750164 | 0.715126680427424 |

| | | | | | | | |
|---|---|---|---|---|---|---|---|
| Psmf1 | ENSMUSG00000032869 | 228769 | -0.0543073921423187 | 2.89351040078388 | -0.481606965029157 | 0.6378525704617 | 0.715554848903656 |
| Zgpat | ENSMUSG00000027582 | 229007 | 0.0778694498107475 | 3.47837256206015 | 0.480068615575007 | 0.6389177234351 | 0.716676206759353 |
| Cep135 | ENSMUSG00000036403 | 381644 | 0.0866935431949694 | 2.21075594974673 | 0.478460055122142 | 0.6400323795935 | 0.717852856693765 |
| Apeh | ENSMUSG00000032590 | 235606 | 0.0411038257067471 | 3.86408246556699 | 0.478300880490816 | 0.6401427294090 | 0.717902962467412 |
| Dbi | ENSMUSG00000026385 | 13167 | -0.0394982247869251 | 5.28490836215038 | -0.478119935828746 | 0.6402681823530 | 0.717969993978347 |
| Tyms | ENSMUSG00000025747 | 22171 | -0.0974207859895444 | 1.62086078819555 | -0.477703643144114 | 0.6405568508335 | 0.718220016048974 |
| Fbxo30 | ENSMUSG00000047648 | 71865 | -0.0390188601144139 | 4.46247110853082 | -0.477352667800308 | 0.6408002736565 | 0.718419260650202 |
| Desi1 | ENSMUSG00000022472 | 28075 | -0.0429032023481812 | 3.13120697497794 | -0.47696475985925 | 0.6410693616265 | 0.718580647372878 |
| Psmd3 | ENSMUSG00000017221 | 22123 | 0.0345045160564856 | 5.7232131483639 | 0.476955624057128 | 0.6410756996780 | 0.718580647372878 |
| Smchd1 | ENSMUSG00000024054 | 74355 | 0.0369650768810814 | 4.42400194941786 | 0.47584767292658 | 0.6418445677869 | 0.719279279187389 |
| Impdh2 | ENSMUSG00000062867 | 23918 | -0.030743535073622 | 4.86254942128002 | -0.475805577316558 | 0.6418737886953 | 0.719279279187389 |
| Tmcc2 | ENSMUSG00000042066 | 68875 | -0.0335131775603763 | 4.40081013951534 | -0.47572782414485 | 0.6419277631301 | 0.719279279187389 |
| Smpd4 | ENSMUSG00000005899 | 77626 | -0.0637309818677085 | 3.06115317134294 | -0.475678238652904 | 0.6419621853217 | 0.719279279187389 |
| Trim30a | ENSMUSG00000030921 | 20128 | 0.0769524860761841 | 2.47041398241338 | 0.475150012847693 | 0.6423289323108 | 0.719616435286409 |
| Vmn2r29 | ENSMUSG00000110105 | 76229 | -0.155702784423439 | 0.293979057470099 | -0.474927581457906 | 0.6424833954599 | 0.719715720001292 |
| Dhx30 | ENSMUSG00000032480 | 72831 | -0.0289281682059723 | 4.42673071710159 | -0.474470837282988 | 0.6428006266191 | 0.71998862578171 |
| Zfyve19 | ENSMUSG00000068580 | 72008 | -0.0617976750367326 | 1.96401012734585 | -0.474387164676881 | 0.6428587492113 | 0.71998862578171 |
| Bod1l | ENSMUSG00000061755 | 665775 | -0.0335644402639756 | 5.98078069709446 | -0.473968013565865 | 0.6431499463017 | 0.720240965374921 |
| Crls1 | ENSMUSG00000027357 | 66586 | 0.050008709050045 | 2.83022493404565 | 0.472991481744415 | 0.6438286099259 | 0.720927118940879 |
| Unc13d | ENSMUSG00000057948 | 70450 | 0.10270061488846 | 1.31751335537236 | 0.472478663550626 | 0.6441851376668 | 0.721252457219756 |
| Smc3 | ENSMUSG00000024974 | 13006 | -0.0296259055289969 | 5.38263712535678 | -0.4723404452656 | 0.6442812471012 | 0.721286185176547 |
| Gm15614 | ENSMUSG00000085234 | 102640951 | -0.169523693828937 | 0.325403974070003 | -0.47188301946266 | 0.6445993632702 | 0.721568421905507 |
| Gtf2ird2 | ENSMUSG00000015942 | 114674 | -0.0930546429002368 | 1.8226641690107 | -0.471661501953994 | 0.6447534434202 | 0.721666996726075 |
| Zfp160 | ENSMUSG00000067942 | 224585 | -0.0653329404816896 | 2.26578728172042 | -0.47150585966631 | 0.6448617131553 | 0.721714281406911 |
| Jagn1 | ENSMUSG00000051256 | 67767 | 0.0660575272730872 | 2.16623764177285 | 0.471182935960597 | 0.6450863759648 | 0.721857081704195 |
| Mief1 | ENSMUSG00000022412 | 239555 | 0.0404052321499027 | 4.36475882958887 | 0.47113262339907 | 0.6451213824140 | 0.721857081704195 |
| Syne2 | ENSMUSG00000063450 | 319565 | -0.0473620343423038 | 5.58457888872612 | -0.470870459249621 | 0.6453038050489 | 0.721958716320457 |
| Nif3l1 | ENSMUSG00000026036 | 65102 | -0.0555809381503574 | 2.49768119330319 | -0.47081225786843 | 0.6453443067575 | 0.721958716320457 |
| Cald1 | ENSMUSG00000029761 | 109624 | 0.0660584408829141 | 6.01205766514266 | 0.470170152978057 | 0.6457912184211 | 0.722384753229826 |
| Gxylt1 | ENSMUSG00000036197 | 223827 | -0.0553329808913912 | 3.66638389678888 | -0.469992724861199 | 0.6459147353929 | 0.722448989315451 |
| Ift81 | ENSMUSG00000029469 | 12589 | -0.0475379977937012 | 4.34911245750137 | -0.469860572445688 | 0.6460067406621 | 0.722473748230691 |
| 3110040N11Rik | ENSMUSG00000025102 | 67290 | 0.0727270852585007 | 2.10702989175166 | 0.469771065330343 | 0.6460690594597 | 0.722473748230691 |
| Smtnl1 | ENSMUSG00000027077 | 68678 | 0.0447482555319212 | 8.35301683904463 | 0.469516377203224 | 0.6462463997112 | 0.72259813781133 |
| Nfrkb | ENSMUSG00000042185 | 235134 | 0.0591361733472189 | 3.36157912593096 | 0.468913581586458 | 0.6466662177650 | 0.722993599917089 |
| Dact2 | ENSMUSG00000048826 | 240025 | 0.105794567466462 | 1.25166242909648 | 0.468418327645467 | 0.6470112321338 | 0.723305356765674 |
| Zfp346 | ENSMUSG00000021481 | 26919 | 0.0638657095170814 | 3.85642803480259 | 0.468065197538389 | 0.6472572888564 | 0.723506434654809 |
| Cnih4 | ENSMUSG00000062169 | 98417 | 0.0406059620223624 | 3.81913409650104 | 0.467565494209007 | 0.6476055494797 | 0.723821703615821 |
| Uvssa | ENSMUSG00000037355 | 71101 | 0.0449257331113007 | 2.81643001616801 | 0.467096833967618 | 0.6479322532914 | 0.724112816759943 |
| Unc45a | ENSMUSG00000030533 | 101869 | -0.0365687769887262 | 3.45222137724659 | -0.46678413398938 | 0.6481502790217 | 0.724282426905233 |
| Arid3a | ENSMUSG00000019564 | 13496 | -0.0763249426671735 | 1.57713969623322 | -0.466530016399766 | 0.6483274837803 | 0.724406391209576 |
| Usf1 | ENSMUSG00000026641 | 22278 | -0.0355957401629561 | 3.42580428515104 | -0.466099239555061 | 0.6486279297370 | 0.724668019210541 |
| Mecp2 | ENSMUSG00000031393 | 17257 | 0.037293998474861 | 5.13461352000607 | 0.465903215787942 | 0.6487646679310 | 0.724746712841518 |
| Heg1 | ENSMUSG00000075254 | 77446 | 0.0854668353791412 | 6.5031481737574 | 0.465777814341993 | 0.6488521497977 | 0.724758447359493 |
| Ube2n | ENSMUSG00000074781 | 93765 | -0.0298613010822466 | 4.50002095330154 | -0.465698077373422 | 0.6489077782734 | 0.724758447359493 |
| Gatad2b | ENSMUSG00000042390 | 229542 | 0.0319974575773427 | 5.63358924071078 | 0.46554336693612 | 0.6490157182012 | 0.724804946430087 |
| Chmp2a | ENSMUSG00000033916 | 68953 | -0.0418251133284731 | 4.84475971956507 | -0.462524327051724 | 0.6511237153240 | 0.727005594047699 |
| Zfp524 | ENSMUSG00000051184 | 66056 | -0.0602964601388947 | 1.60816340838921 | -0.462470166564142 | 0.6511615605270 | 0.727005594047699 |
| Nfil3 | ENSMUSG00000056749 | 18030 | -0.114498288211521 | 3.84798083520325 | -0.462435505362609 | 0.6511857809277 | 0.727005594047699 |
| Slc29a2 | ENSMUSG00000024891 | 13340 | -0.0968655474599416 | 1.22448381123758 | -0.461805389019159 | 0.6516261623949 | 0.72742295559023 |
| Ap5z1 | ENSMUSG00000039623 | 231855 | 0.080359717141181 | 1.70931967567922 | 0.460133331574642 | 0.6527954019926 | 0.728599463295559 |
| Per3 | ENSMUSG00000028957 | 18628 | -0.120534403480856 | 3.30818065983415 | -0.460067977050044 | 0.6528411225822 | 0.728599463295559 |
| Urb2 | ENSMUSG00000031976 | 382038 | 0.053060769688134 | 2.59452416454285 | 0.460012345476496 | 0.6528800423547 | 0.728599463295559 |
| Slx4 | ENSMUSG00000039738 | 52864 | 0.0522376789255325 | 2.83613721228655 | 0.459167174442668 | 0.6534714523974 | 0.729185019003315 |
| Alg13 | ENSMUSG00000041718 | 67574 | -0.0774354556964603 | 1.44394708633125 | -0.459012707388989 | 0.6535795672722 | 0.729231218725381 |
| Zup1 | ENSMUSG00000039531 | 72580 | -0.0653715419826986 | 2.81452485626361 | -0.458531009847358 | 0.6539167700057 | 0.729532987643708 |
| Snrnp40 | ENSMUSG00000074088 | 66585 | 0.0462107711197706 | 2.55492353917207 | 0.458302103475348 | 0.6540770389623 | 0.729579471882409 |
| Khnyn | ENSMUSG00000047153 | 219094 | 0.0515909079545143 | 3.7233120582025 | 0.458280844583821 | 0.6540919242930 | 0.729579471882409 |
| Exosc7 | ENSMUSG00000025785 | 66446 | -0.0644634162401148 | 1.92808481193162 | -0.457999147406154 | 0.6542891812185 | 0.729725032129413 |
| Orai1 | ENSMUSG00000049686 | 109305 | -0.0372796911317579 | 4.12813397452899 | -0.457702289039438 | 0.6544970838260 | 0.729882434533995 |
| Hoxc6 | ENSMUSG00000001661 | 15425 | 0.0977452808025312 | 1.18440373247452 | 0.457596437996531 | 0.6545712230674 | 0.729890649734768 |
| Zfr | ENSMUSG00000022201 | 22763 | -0.0300906464077242 | 5.99671546305455 | -0.457445902590799 | 0.6546766662750 | 0.729933765713207 |
| Firre | ENSMUSG00000085396 | 103012 | -0.0616959011647054 | 3.43390215083971 | -0.457223855901505 | 0.6548322139095 | 0.730032731284618 |
| Tcf3 | ENSMUSG00000020167 | 21423 | 0.0509112837362418 | 3.78471899700447 | 0.45673837102914 | 0.6551723628529 | 0.730337456490458 |
| Ccdc34 | ENSMUSG00000027160 | 68201 | -0.0740286219232817 | 1.6841989947921 | -0.456521928897796 | 0.6553240360716 | 0.730432042245244 |
| Ltn1 | ENSMUSG00000052299 | 78913 | -0.0313368725479695 | 4.92829676910585 | -0.456143073588126 | 0.6555895595432 | 0.730653494633703 |
| Zfp398 | ENSMUSG00000062519 | 272347 | 0.043642587659122 | 3.13299047451487 | 0.456038880832049 | 0.6556625923016 | 0.730660393154227 |
| Rbm15 | ENSMUSG00000048109 | 229700 | 0.0386161252276649 | 4.55583759474547 | 0.455774925818773 | 0.6558476250787 | 0.730792088658065 |
| Drg2 | ENSMUSG00000020537 | 13495 | -0.0415611418465627 | 3.18308895325316 | -0.453773503449602 | 0.6572513903041 | 0.732281617308565 |
| Picalm | ENSMUSG00000039361 | 233489 | -0.0450139662544196 | 6.36360235378406 | -0.453136195366003 | 0.6576986714464 | 0.732705276965088 |
| Zfp938 | ENSMUSG00000062931 | 237411 | -0.057503844378167 | 1.86387372001485 | -0.452917663300452 | 0.6578520750235 | 0.732801491091606 |
| 1110059G10Rik | ENSMUSG00000032551 | 66202 | -0.0820274428707317 | 1.64563130121466 | -0.452677565252443 | 0.6580206358627 | 0.732889502697956 |
| Npm1 | ENSMUSG00000057113 | 18148 | -0.0300476172773845 | 5.990626643132 | -0.452614122972539 | 0.6580651787558 | 0.732889502697956 |
| Exosc4 | ENSMUSG00000034259 | 109075 | -0.0459134625302253 | 2.7398429197547 | -0.45209487012192 | 0.6584297976135 | 0.733220875887739 |
| Fcsk | ENSMUSG00000033703 | 234730 | -0.073022327710522 | 1.41232810439819 | -0.451740576306113 | 0.6586786344029 | 0.733359868367607 |
| Srp14 | ENSMUSG00000009549 | 20813 | 0.040375101575604 | 5.48722928351492 | 0.45172387816202 | 0.6586903633155 | 0.733359868367607 |
| Dhx33 | ENSMUSG00000040620 | 216877 | -0.0423520400229833 | 3.01608483947274 | -0.451630603211273 | 0.6587558821243 | 0.733359868367607 |
| Sin3a | ENSMUSG00000042557 | 20466 | -0.0341853093601365 | 4.45931528278351 | -0.450933928206912 | 0.6592453375671 | 0.733830018833623 |
| Gm46430 | ENSMUSG00000113204 | 108168101 | -0.0397998384020801 | 2.87193970047171 | -0.45075161292519 | 0.6593734517309 | 0.733843566284201 |
| Tmem208 | ENSMUSG00000014856 | 66320 | 0.0774302803928808 | 1.45844543898608 | 0.45072553692728 | 0.6593917764196 | 0.733843566284201 |
| Plcd1 | ENSMUSG00000010660 | 18799 | -0.0449557284142892 | 2.97313986042298 | -0.449348790247955 | 0.6603595971307 | 0.734845847117609 |
| Dclre1a | ENSMUSG00000025077 | 55947 | -0.0815881577736195 | 1.60519045639065 | -0.44872654052054 | 0.6607972322715 | 0.735257995313572 |
| H3f3a | ENSMUSG00000060743 | 15078 | -0.030112196062146 | 6.7824683371047 | -0.44797211724136 | 0.6613280000583 | 0.735773676197186 |
| Dolpp1 | ENSMUSG00000026856 | 57170 | -0.0713423336363181 | 1.23324125145058 | -0.447766820223851 | 0.6614724678371 | 0.735859510067926 |
| Neurl2 | ENSMUSG00000039873 | 415115 | -0.0522134093770696 | 2.89380596886902 | -0.447595814928012 | 0.6615928152212 | 0.735918496304437 |
| Rbis | ENSMUSG00000078784 | 69126 | -0.0507113851512595 | 3.94906590723897 | -0.447231656927594 | 0.6618491290365 | 0.736128696530163 |
| Ptrh2 | ENSMUSG00000072582 | 217057 | -0.068347399477426 | 1.86484103855787 | -0.447128332108972 | 0.6619218626039 | 0.7361346912324 |
| Golph3l | ENSMUSG00000046519 | 229593 | 0.0510460923926264 | 2.8031166009949 | 0.446573166879762 | 0.6623127216508 | 0.736470964608647 |
| Exosc2 | ENSMUSG00000039356 | 227715 | -0.0735836713883169 | 1.72282681000012 | -0.446500587342066 | 0.6623638282033 | 0.736470964608647 |
| Irf9 | ENSMUSG00000002325 | 16391 | 0.0490441375269648 | 3.0934930059502 | 0.44641178772993 | 0.6624263584331 | 0.736470964608647 |
| Rbm5 | ENSMUSG00000032580 | 83486 | -0.0480069166148178 | 5.35175243443118 | -0.446121867978404 | 0.6626305301716 | 0.736623037252719 |
| 2210408I21Rik | ENSMUSG00000071252 | 72371 | -0.0971681252469287 | 1.30959714654026 | -0.446013215723733 | 0.6627070541350 | 0.736633191859943 |
| Commd8 | ENSMUSG00000029213 | 27784 | -0.0342577384206759 | 3.45734156971398 | -0.445743710673003 | 0.6628968839387 | 0.73676927690235 |
| Zfp59 | ENSMUSG00000078779 | 22717 | -0.0779341939612956 | 1.29131386018695 | -0.445125951133705 | 0.6633321029595 | 0.737178041627821 |
| Rhoq | ENSMUSG00000024143 | 104215 | -0.0278632744081859 | 6.59497520761528 | -0.443747041864056 | 0.6643040181313 | 0.738137504125485 |
| Slc35c2 | ENSMUSG00000017664 | 228875 | 0.0479538285795775 | 2.57081864581883 | 0.443709474007873 | 0.6643305064117 | 0.738137504125485 |
| Abcc10 | ENSMUSG00000032842 | 224814 | -0.100056296110905 | 1.21693297499888 | -0.443080949688068 | 0.6647737346066 | 0.738492732812529 |
| Tgfb1i1 | ENSMUSG00000030782 | 21804 | 0.0935599258010611 | 2.66101699635348 | 0.443064502900184 | 0.6647853344502 | 0.738492732812529 |
| Nme1 | ENSMUSG00000037601 | 18102 | 0.0511888103871954 | 3.25414719095498 | 0.442876467541878 | 0.6649179612826 | 0.738515330983431 |
| Smurf1 | ENSMUSG00000038780 | 75788 | -0.0320963101767378 | 4.48912614575709 | -0.442844088637761 | 0.6649408002505 | 0.738515330983431 |
| Snx15 | ENSMUSG00000024787 | 69024 | 0.060659536776658 | 1.80228384094495 | 0.44254606566968 | 0.6651510317110 | 0.738673770967591 |
| Decr2 | ENSMUSG00000036775 | 26378 | 0.0781171055476078 | 1.77266370850901 | 0.441669900891194 | 0.6657692661914 | 0.739258197135659 |
| Fmo2 | ENSMUSG00000040170 | 55990 | -0.0441953534310597 | 3.73909031784894 | -0.441608576643824 | 0.6658125469582 | 0.739258197135659 |
| Hist1h4d | ENSMUSG00000061482 | 319156 | 0.0468498693650714 | 3.66345659437635 | 0.441026883016372 | 0.6662231501431 | 0.73963896549003 |
| Stx3 | ENSMUSG00000041488 | 20908 | -0.083570411500983 | 1.20225765150932 | -0.440866169578573 | 0.6663366134745 | 0.739689806224263 |
| Atoh8 | ENSMUSG00000037621 | 71093 | 0.0631729892926169 | 2.48086635713099 | 0.440474288933608 | 0.6666133160364 | 0.739921827538011 |
| Ubald1 | ENSMUSG00000039568 | 207740 | -0.0346695160722872 | 4.43578966738823 | -0.439756369149419 | 0.6671203624362 | 0.740409450887432 |
| Cdc40 | ENSMUSG00000038446 | 71713 | 0.0395872805146236 | 3.34915076883003 | 0.438615890570297 | 0.6679261987746 | 0.741228556223901 |
| Trim56 | ENSMUSG00000043279 | 384309 | -0.0374887838844318 | 4.11920821366613 | -0.43804585004724 | 0.6683291366641 | 0.741528168139794 |
| Maz | ENSMUSG00000030678 | 17188 | 0.037865141226713 | 4.30298024641584 | 0.437999983973718 | 0.6683615621155 | 0.741528168139794 |
| Hps5 | ENSMUSG00000014418 | 246694 | -0.0534783631383564 | 3.0457751306492 | -0.437946049219238 | 0.6683996926796 | 0.741528168139794 |
| Efcab14 | ENSMUSG00000034210 | 230648 | 0.0249904800827423 | 5.63200686100817 | 0.437470703145184 | 0.6687357920860 | 0.741825750283379 |
| Mcm4 | ENSMUSG00000022673 | 17217 | -0.0738131036581002 | 2.26293905222909 | -0.436896986643919 | 0.6691415441627 | 0.742200529603508 |
| Snrpa | ENSMUSG00000061479 | 53607 | -0.0399556094561366 | 3.80875924425054 | -0.43667440141906 | 0.6692989931255 | 0.742299847185047 |
| Pfdn4 | ENSMUSG00000052033 | 109054 | -0.0644159949866261 | 1.92988308331502 | -0.435818276332656 | 0.6699047369867 | 0.742845033809324 |
| Thap1 | ENSMUSG00000037214 | 73754 | 0.0655481585210422 | 1.79493027796944 | 0.435787553487506 | 0.6699264791228 | 0.742845033809324 |
| Kcnj12 | ENSMUSG00000042529 | 16515 | 0.0780589192004966 | 4.45119778740068 | 0.435270741020401 | 0.6702922664400 | 0.743068606287332 |

| | | | | | | | |
|---|---|---|---|---|---|---|---|
| Ddx55 | ENSMUSG00000029389 | 67848 | -0.0577010471537023 | 1.91929587351364 | -0.435222418121324 | 0.6703264726739 | 0.743068606287332 |
| Ankrd46 | ENSMUSG00000048307 | 68839 | -0.0337953705186473 | 3.32841285377161 | -0.43521455364333 | 0.6703320397584 | 0.743068606287332 |
| Srgap2 | ENSMUSG00000026425 | 14270 | 0.0484708829401259 | 3.91312908358142 | 0.434965129019926 | 0.6705086122288 | 0.743188971838747 |
| Hoxa10 | ENSMUSG00000000938 | 15395 | -0.054850951566296 | 2.68560402378093 | -0.434531777541513 | 0.6708154383129 | 0.743442983257067 |
| Coil | ENSMUSG00000033983 | 12812 | 0.0767343204773641 | 1.33447511242288 | 0.434393315065178 | 0.6709134868856 | 0.743442983257067 |
| Hmgb3 | ENSMUSG00000015217 | 15354 | -0.0512045199976439 | 3.62156495804856 | -0.434353305774042 | 0.6709418195801 | 0.743442983257067 |
| Tspyl4 | ENSMUSG00000039485 | 72480 | -0.0643913442057142 | 2.29490874802773 | -0.433178287804919 | 0.6717741444517 | 0.744274918720814 |
| Yy1 | ENSMUSG00000021264 | 22632 | -0.0274187071499989 | 4.95703194178814 | -0.433101149800353 | 0.6718288009347 | 0.744274918720814 |
| Ppp3ca | ENSMUSG00000028161 | 19055 | 0.026686570328498 | 6.18246254163134 | 0.432736856400698 | 0.6720869488209 | 0.744478408306707 |
| Rab5c | ENSMUSG00000019173 | 19345 | -0.0487605885755382 | 4.0600496320724 | -0.432641951898718 | 0.6721542077308 | 0.744478408306707 |
| Ankrd2 | ENSMUSG00000025172 | 56642 | -0.0615526583038548 | 9.14570534961178 | -0.432553629538778 | 0.6722168044997 | 0.744478408306707 |
| Abt1 | ENSMUSG00000036376 | 30946 | -0.0443924877655908 | 2.3111343756815 | -0.432285235730592 | 0.6724070389859 | 0.744613650405791 |
| Cyld | ENSMUSG00000036712 | 74256 | 0.0451080144474343 | 5.20852663618746 | 0.431892157529262 | 0.6726856905713 | 0.744846766980868 |
| Tdg | ENSMUSG00000034674 | 21665 | 0.0283760316803576 | 4.05614089475037 | 0.429667476012904 | 0.6742637024331 | 0.74649544357376 |
| Ythdc2 | ENSMUSG00000034653 | 240255 | 0.0537679296787474 | 2.83039086175518 | 0.429600512486466 | 0.6743112258574 | 0.74649544357376 |
| Gtpbp4 | ENSMUSG00000021149 | 69237 | -0.0283319243603382 | 5.1849680239919 | -0.429208099565674 | 0.6745897469709 | 0.746728154343184 |
| Trnt1 | ENSMUSG00000013736 | 70047 | 0.0422371525955776 | 3.64036263650671 | 0.429035078641685 | 0.6747125670358 | 0.746788484231338 |
| Git2 | ENSMUSG00000041890 | 26431 | 0.0433449134782284 | 4.76447307345879 | 0.428658076490372 | 0.6749802180141 | 0.747009088094865 |
| Fam126a | ENSMUSG00000028995 | 84652 | 0.0324536002511087 | 3.75989564457706 | 0.428076810742541 | 0.6753929748555 | 0.747383283468766 |
| Apc | ENSMUSG00000005871 | 11789 | 0.0340691459191182 | 6.30819366784652 | 0.427989367153535 | 0.6754550779789 | 0.747383283468766 |
| Abraxas2 | ENSMUSG00000030965 | 109359 | -0.035581737055395 | 4.04078417972126 | -0.427248411128797 | 0.6759814096218 | 0.747889958357969 |
| Rsrc1 | ENSMUSG00000034544 | 66880 | 0.0341716016308391 | 3.27662314806131 | 0.426929005467482 | 0.6762083511540 | 0.748065326026202 |
| Afap1 | ENSMUSG00000029094 | 70292 | 0.0632095472561147 | 4.39067163826544 | 0.426752692790478 | 0.6763336374702 | 0.74812821202055 |
| Mrps22 | ENSMUSG00000032459 | 64655 | -0.0336111624920457 | 3.74678038495907 | -0.426458467307408 | 0.6765427338962 | 0.748283782679291 |
| Ikbkg | ENSMUSG00000004221 | 16151 | 0.0368652529758 | 4.39049837316813 | 0.426360729839474 | 0.6766121988733 | 0.748284899421779 |
| Slc7a6os | ENSMUSG00000033106 | 66432 | 0.0498321988578807 | 2.18488314734089 | 0.426134532379775 | 0.6767729760252 | 0.748386990483774 |
| Pts | ENSMUSG00000032067 | 19286 | -0.055876489777178 | 1.80202326118959 | -0.42583064179446 | 0.6769890019417 | 0.748550149729422 |
| Rexo4 | ENSMUSG00000052406 | 227656 | 0.0340192990338131 | 3.22123613585368 | 0.425475602746044 | 0.6772414251780 | 0.74875351659969 |
| Sept11 | ENSMUSG00000058013 | 52398 | 0.0350118768636598 | 5.74166246396671 | 0.425231401983372 | 0.6774150688383 | 0.748869752980615 |
| Xpot | ENSMUSG00000034667 | 73192 | -0.0313287308782417 | 4.7846776129809 | -0.423511155364431 | 0.6786388246106 | 0.7501467278119 |
| Cinp | ENSMUSG00000021276 | 67236 | -0.0509700340462618 | 2.55195357819902 | -0.423176267506532 | 0.6788771685213 | 0.750334310324291 |
| Cwc25 | ENSMUSG00000018541 | 67480 | 0.050182643154209 | 2.75342229204363 | 0.422619703532914 | 0.6792733613441 | 0.750696300965805 |
| Rfx2 | ENSMUSG00000024206 | 19725 | 0.0756872271189484 | 2.77988741036009 | 0.422501442284483 | 0.6793575589353 | 0.750713452964277 |
| Ccdc28a | ENSMUSG00000059554 | 215814 | -0.0826765249624881 | 1.30616135133084 | -0.422386247940901 | 0.6794395772944 | 0.750728193612175 |
| Cfdp1 | ENSMUSG00000031954 | 23837 | -0.0378842948590385 | 3.68616983590622 | -0.421806091034179 | 0.6798527132567 | 0.751108753649669 |
| Ctnnd1 | ENSMUSG00000034101 | 12388 | -0.0679112976683562 | 4.66291343243673 | -0.421606030099112 | 0.6799952036673 | 0.751190254804239 |
| Pik3cd | ENSMUSG00000039936 | 18707 | 0.0640118073279905 | 2.55121986494293 | 0.421486755030877 | 0.6800801616173 | 0.751208189838211 |
| Ubp1 | ENSMUSG00000009741 | 22221 | 0.0397681598281086 | 3.83436354595635 | 0.421257566852867 | 0.6802434218397 | 0.751312604236704 |
| Spout1 | ENSMUSG00000039660 | 227695 | 0.0671327327221376 | 1.53465950197777 | 0.419504441723449 | 0.6814927979152 | 0.752616465347685 |
| Odf2 | ENSMUSG00000026790 | 18286 | -0.0302690499418201 | 4.13405176950029 | -0.418932255893748 | 0.6819007810764 | 0.752974117019671 |
| Nhlrc1 | ENSMUSG00000044231 | 105193 | -0.0832429968392179 | 1.04237005586952 | -0.418857040352193 | 0.6819544194030 | 0.752974117019671 |
| Shisa2 | ENSMUSG00000044461 | 219134 | 0.0548064851434473 | 4.21832731398925 | 0.418452026141965 | 0.6822432772791 | 0.753216974440794 |
| Dbr1 | ENSMUSG00000032469 | 83703 | -0.0728560518065358 | 1.35700263861681 | -0.417743014523033 | 0.6827490724242 | 0.753699263852674 |
| Ube2v2 | ENSMUSG00000022674 | 70620 | -0.0407594533387394 | 3.48884189533178 | -0.41693892736762 | 0.6833228849421 | 0.754256533909151 |
| Plcb1 | ENSMUSG00000051177 | 18795 | 0.113100930298103 | 2.68253364675446 | 0.41662327888483 | 0.6835481937576 | 0.754429049471443 |
| Abhd14b | ENSMUSG00000042073 | 76491 | -0.0942169488594398 | 1.17412200577671 | -0.416251633291761 | 0.6838135134138 | 0.754645685525203 |
| Nrp2 | ENSMUSG00000025969 | 18187 | 0.0444180310367273 | 4.45949590072872 | 0.41530195140808 | 0.6844916938252 | 0.75531785838919 |
| Hook3 | ENSMUSG00000037234 | 320191 | 0.0338302238963877 | 7.13411280746254 | 0.414996335424677 | 0.6847099986397 | 0.755482486638901 |
| 2310033P09Rik | ENSMUSG00000020441 | 67862 | 0.0661801966843518 | 1.69388063540843 | 0.414264421842932 | 0.6852329318488 | 0.755983162902628 |
| Pitx3 | ENSMUSG00000025229 | 18742 | -0.0739418412610888 | 2.07760919961508 | -0.413499441022428 | 0.6857796705679 | 0.756501865980447 |
| Nup54 | ENSMUSG00000034826 | 269113 | -0.0447457829212476 | 2.75313012618976 | -0.413412940170718 | 0.6858415050650 | 0.756501865980447 |
| Trpm4 | ENSMUSG00000038260 | 68667 | -0.0400405063762004 | 3.64989387229944 | -0.412789203896529 | 0.6862874477830 | 0.756917373798485 |
| Tbc1d17 | ENSMUSG00000038520 | 233204 | -0.053550013314573 | 3.86281259471282 | -0.411896380433694 | 0.6869259874571 | 0.757474401231764 |
| Arntl | ENSMUSG00000055116 | 11865 | -0.1157584262457 | 4.23130676922957 | -0.411889248012107 | 0.6869310895078 | 0.757474401231764 |
| Sema6a | ENSMUSG00000019647 | 20358 | -0.0469038684637271 | 3.94228081132459 | -0.411663923630495 | 0.6870922794774 | 0.757542231898551 |
| Nutf2 | ENSMUSG00000071497 | 68051 | -0.116833000721772 | 0.900836290183153 | -0.411609510204411 | 0.6871312075083 | 0.757542231898551 |
| Large1 | ENSMUSG00000004383 | 16795 | -0.0394220067002387 | 5.21178610680263 | -0.411418236969335 | 0.6872680540634 | 0.757616690093555 |
| H2-T23 | ENSMUSG00000067212 | 15040 | -0.0660921184613539 | 3.18706797706141 | -0.410744259024218 | 0.6877503430582 | 0.758071896739929 |
| Pard3 | ENSMUSG00000025812 | 93742 | -0.0292736232078701 | 4.9276444940499 | -0.410420527986714 | 0.6879820507594 | 0.758250836544806 |
| Srsf1 | ENSMUSG00000018379 | 110809 | -0.0331248225721028 | 5.30168199623687 | -0.410096232817394 | 0.6882141949929 | 0.758430221339673 |
| 2210016F16Rik | ENSMUSG00000021550 | 70153 | -0.0493030487833539 | 3.02452724930811 | -0.409969942106363 | 0.6882852780473 | 0.758432094187079 |
| Ccdc82 | ENSMUSG00000079084 | 66396 | 0.037740733990272 | 3.78085156962454 | 0.409421895037024 | 0.6886970194574 | 0.758658446056609 |
| Qtrt2 | ENSMUSG00000022704 | 106248 | 0.0562802461015746 | 1.59458746885008 | 0.409390541306152 | 0.6887194721082 | 0.758658446056609 |
| Dhx16 | ENSMUSG00000024422 | 69192 | -0.0349279006058989 | 3.92105136859636 | -0.409387631148776 | 0.6887215561100 | 0.758658446056609 |
| Six1 | ENSMUSG00000051367 | 20471 | 0.0681425948887906 | 6.08087059619205 | 0.409322340883955 | 0.6887683120177 | 0.758658446056609 |
| Taf9b | ENSMUSG00000047242 | 407786 | -0.0568701805589398 | 2.29840054901296 | -0.40907344795088 | 0.6889465623147 | 0.758778324701483 |
| Zc3h12a | ENSMUSG00000042677 | 230738 | 0.11410637412262 | 1.63561056061243 | 0.408450265382527 | 0.6893929530662 | 0.75919346866487 |
| Bscl2 | ENSMUSG00000071657 | 14705 | -0.0398983939507187 | 2.72815879119809 | -0.408250033728098 | 0.6895364062421 | 0.759274952818848 |
| Tbc1d19 | ENSMUSG00000039178 | 67249 | 0.0556938873809241 | 2.05573347252681 | 0.408114907703752 | 0.6896332224320 | 0.759305071958554 |
| Dnal4 | ENSMUSG00000022420 | 54152 | -0.0474641699077636 | 1.73347876616065 | -0.405825778427527 | 0.6912742163367 | 0.761035195767695 |
| Lmf1 | ENSMUSG00000002279 | 76483 | 0.0393742390068001 | 2.77674824172327 | 0.40501191858964 | 0.691858033521 | 0.761601225016923 |
| Ankhd1 | ENSMUSG00000024483 | 108857 | -0.0243897099649315 | 5.5797971637099 | -0.404882542979018 | 0.6919508591243 | 0.761626708396811 |
| Psma4 | ENSMUSG00000032301 | 26441 | -0.0295756290848912 | 5.89943579202595 | -0.402987751860706 | 0.6933109420271 | 0.763046909715911 |
| Mkx | ENSMUSG00000061013 | 210719 | 0.0820138295307873 | 1.78192983005446 | 0.402880076162644 | 0.6933882649109 | 0.763055182094156 |
| Il6ra | ENSMUSG00000027947 | 16194 | -0.0817656719270419 | 2.61861505439159 | -0.402529755986624 | 0.6936398576079 | 0.763255212755456 |
| March8 | ENSMUSG00000025702 | 71779 | -0.0318805125460448 | 4.68462497703053 | -0.401016726930457 | 0.6947269161305 | 0.764301835521552 |
| Smarcc2 | ENSMUSG00000025369 | 68094 | -0.021550107610719 | 5.95002575978132 | -0.401007332804766 | 0.6947336676723 | 0.764301835521552 |
| Tmem56 | ENSMUSG00000028132 | 99887 | -0.0726333279377154 | 2.36695431877644 | -0.400913952747398 | 0.6948007812256 | 0.764301835521552 |
| Flywch1 | ENSMUSG00000040097 | 224613 | -0.0304754767121578 | 4.83216964508303 | -0.400564789002375 | 0.6950517537285 | 0.764500978064697 |
| Rsbn1 | ENSMUSG00000044098 | 229675 | -0.0339638088242276 | 3.43472315678903 | -0.400330287398469 | 0.6952203301115 | 0.76460946055433 |
| Nupr1 | ENSMUSG00000030717 | 56312 | -0.0523635235969859 | 3.1819024819455 | -0.400192462999392 | 0.6953194159056 | 0.764641502541702 |
| Swap70 | ENSMUSG00000031015 | 20947 | -0.0399822408933734 | 3.54419494550949 | -0.399979480874571 | 0.6954725460926 | 0.764732964625167 |
| Sctr | ENSMUSG00000026387 | 319229 | 0.0868459945109757 | 0.885722087681698 | 0.399847925727217 | 0.6955671387187 | 0.764760047609591 |
| Rps8 | ENSMUSG00000047675 | 20116 | 0.0268430221570126 | 7.0130570113856 | 0.399389945201603 | 0.6958964835043 | 0.765045203780099 |
| Zfp335 | ENSMUSG00000039834 | 329559 | 0.0514613228484572 | 3.44505426185604 | 0.398897722457116 | 0.6962505240241 | 0.765357449528178 |
| Kif1b | ENSMUSG00000063077 | 16561 | -0.0286014803220187 | 9.52416503760614 | -0.397237246868995 | 0.6974453967062 | 0.766593829200228 |
| Nudcd1 | ENSMUSG00000038736 | 67429 | -0.0686213403426921 | 2.75129532562162 | -0.397062166701069 | 0.6975714326601 | 0.766655271506976 |
| Rnf103 | ENSMUSG00000052656 | 22644 | -0.0363617084340722 | 3.83581720990076 | -0.396510059780049 | 0.6979689419317 | 0.767015030085099 |
| Tmem63a | ENSMUSG00000026519 | 208795 | -0.0394109084638665 | 3.46339211382189 | -0.394793280602172 | 0.6992055891131 | 0.768296772677488 |
| Clock | ENSMUSG00000029238 | 12753 | -0.0252430046996794 | 5.76791993558956 | -0.394374655765292 | 0.6995072722432 | 0.768551009437249 |
| Rptor | ENSMUSG00000025583 | 74370 | 0.0240768151102744 | 5.66085886680132 | 0.394256397525158 | 0.6995925049534 | 0.768567404185518 |
| A830082N09Rik | ENSMUSG00000112280 | 414093 | -0.0629226646729708 | 1.80733118492747 | -0.393856357881959 | 0.6998808583311 | 0.76880692015047 |
| Eipr1 | ENSMUSG00000036613 | 380752 | -0.0772645967998218 | 1.24042632558986 | -0.393501009157546 | 0.7001370384392 | 0.768938661016284 |
| Tcn2 | ENSMUSG00000020432 | 21452 | -0.0465224001431617 | 2.98007704115856 | -0.39349485185672 | 0.7001414777326 | 0.768938661016284 |
| Snrnp48 | ENSMUSG00000021431 | 67797 | 0.0381371200499765 | 2.79072218288533 | 0.393394273438816 | 0.7002139944198 | 0.76894104611242 |
| Hnrnpll | ENSMUSG00000024095 | 72692 | 0.0328954012187457 | 3.53610429929842 | 0.39252636762546 | 0.7008398780449 | 0.769551050417827 |
| Zdhhc4 | ENSMUSG00000001844 | 72881 | -0.0385027585681135 | 3.42894226519317 | -0.39191769100772 | 0.7012789557124 | 0.76995583215071 |
| Bud31 | ENSMUSG00000038722 | 231889 | 0.0283127919345327 | 3.70724977769988 | 0.391551545574546 | 0.7015431336275 | 0.770168524021605 |
| Atp11b | ENSMUSG00000037400 | 76295 | -0.0277217296221748 | 4.18981183464156 | -0.390864196302666 | 0.7020391722649 | 0.770635689097009 |
| Babam2 | ENSMUSG00000052139 | 107976 | 0.0351458077963036 | 3.52486021402746 | 0.390049710546375 | 0.7026271446683 | 0.771203666871137 |
| Vmn1r203 | ENSMUSG00000069289 | 171270 | -0.105854349651411 | 1.16145555060361 | -0.389929034414538 | 0.7027142769400 | 0.771221863577429 |
| Sf3a1 | ENSMUSG00000002129 | 67465 | -0.0331613355553069 | 4.97609759322187 | -0.389567458963199 | 0.7029753727917 | 0.771430960745596 |
| Tasp1 | ENSMUSG00000039033 | 75812 | -0.0793663710147441 | 1.73041652818033 | -0.389390748532089 | 0.7031029906974 | 0.771493554639003 |
| Mfge8 | ENSMUSG00000030605 | 17304 | -0.047305224563102 | 5.3950581508635 | -0.389036336698055 | 0.7033589702964 | 0.771659543431957 |
| Phf12 | ENSMUSG00000037791 | 268848 | -0.0287762076930695 | 4.11173676349188 | -0.388985830230127 | 0.7033954524522 | 0.771659543431957 |
| Kcp | ENSMUSG00000059022 | 333088 | 0.104332600665892 | 1.03138465001307 | 0.388385390797668 | 0.7038292240403 | 0.772043506762921 |
| Brd8 | ENSMUSG00000003778 | 78656 | -0.0268237320762728 | 4.04669534899159 | -0.388305833094583 | 0.7038867064677 | 0.772043506762921 |
| Fam234b | ENSMUSG00000030207 | 74525 | -0.0744817617203393 | 1.7209678206572 | -0.388031763575945 | 0.7040847429855 | 0.772127490515357 |
| En1 | ENSMUSG00000058665 | 13798 | 0.0553932121546303 | 3.48977613131137 | 0.387965383949675 | 0.7041327107894 | 0.772127490515357 |
| Tdrd3 | ENSMUSG00000022019 | 219249 | -0.0262294810002235 | 4.12695676591492 | -0.387906607354995 | 0.7041751855226 | 0.772127490515357 |
| Fbrs | ENSMUSG00000042423 | 14123 | 0.0306262701386135 | 4.47157132917128 | 0.387633794621045 | 0.7043723463135 | 0.772266210386479 |
| Intu | ENSMUSG00000060798 | 380614 | 0.0574463922703585 | 2.53776297687129 | 0.3874146646367 | 0.7045307268201 | 0.772362388413536 |
| Myoz3 | ENSMUSG00000049173 | 170947 | -0.0465134952014052 | 5.40715802406879 | -0.386093616486351 | 0.7054858428595 | 0.773331904161461 |
| Mad1l1 | ENSMUSG00000029554 | 17120 | -0.0484452579185954 | 2.96260857776417 | -0.385750040287919 | 0.7057343329081 | 0.773526721449837 |
| Erp44 | ENSMUSG00000028343 | 76299 | 0.0249336797549589 | 4.27040499314733 | 0.385539466683967 | 0.7058866466893 | 0.773603327206884 |
| Bche | ENSMUSG00000027792 | 12038 | 0.0434105592504674 | 2.6644424823517 | 0.385457736585893 | 0.7059457678976 | 0.773603327206884 |

| | | | | | | | |
|---|---|---|---|---|---|---|---|
| Ankrd50 | ENSMUSG00000044864 | 99696 | -0.0346336792429511 | 4.31412021519132 | -0.385062018548758 | 0.7062320469622 | 0.773839465251012 |
| Ociad1 | ENSMUSG00000029152 | 68095 | 0.0294061960649188 | 5.52179567198563 | 0.384309352053967 | 0.7067766854826 | 0.774358619726408 |
| Rmnd5b | ENSMUSG00000001054 | 66089 | -0.0435548151041654 | 2.72043082654874 | -0.383852673370896 | 0.7071072253170 | 0.774643122864369 |
| Nprl3 | ENSMUSG00000020289 | 17168 | 0.0491401369594624 | 2.70784296978999 | 0.382655837125582 | 0.7079737764731 | 0.775514715966283 |
| Tmem181a | ENSMUSG00000038141 | 77106 | 0.0421106603718449 | 3.14691890834732 | 0.382273968933984 | 0.7082503511998 | 0.775714163679723 |
| Papola | ENSMUSG00000021111 | 18789 | -0.0271543005344572 | 5.35537907352969 | -0.382208485333402 | 0.7082977831568 | 0.775714163679723 |
| Ppp1r13b | ENSMUSG00000021285 | 21981 | -0.034675008194716 | 4.10138861520072 | -0.38183002647779 | 0.7085719382516 | 0.775936671711919 |
| Hecw2 | ENSMUSG00000042807 | 329152 | 0.0530692284484644 | 2.98818551083891 | 0.381650392988049 | 0.7087020792561 | 0.775980160404912 |
| Tmem258 | ENSMUSG00000036372 | 69038 | 0.0486339007377383 | 2.44470335413709 | 0.381579242187503 | 0.7087536292638 | 0.775980160404912 |
| Atxn7l3b | ENSMUSG00000074748 | 382423 | -0.038202263222626 | 3.90775753364335 | -0.380381534521106 | 0.7096216126946 | 0.776852663832305 |
| Gadd45b | ENSMUSG00000015312 | 17873 | -0.0776745779555803 | 2.10810048086605 | -0.379954309585545 | 0.7099313260826 | 0.777113891989698 |
| Polr3a | ENSMUSG00000025280 | 218832 | 0.0359509184513023 | 2.65853194115523 | 0.37962326391393 | 0.7101713518229 | 0.77729879310871 |
| Cox18 | ENSMUSG00000035505 | 231430 | -0.067740448796223 | 1.44902047709477 | -0.377467485046254 | 0.7117351897258 | 0.778932454835166 |
| Uimc1 | ENSMUSG00000025878 | 20184 | 0.0414406460509973 | 3.34276187051964 | 0.377059493783069 | 0.7120313059179 | 0.77917851686742 |
| Cnksr3 | ENSMUSG00000015202 | 215748 | 0.0625431222511943 | 1.82756913328124 | 0.376092527485378 | 0.7127333142735 | 0.779868654486903 |
| Avl9 | ENSMUSG00000029787 | 78937 | 0.0477334781669025 | 2.60843780148556 | 0.373450849650052 | 0.7146525297858 | 0.781879521959231 |
| Zc3h11a | ENSMUSG00000116275 | 70579 | -0.0537134982095884 | 4.95303112330692 | -0.373366103302821 | 0.7147141325968 | 0.781879521959231 |
| Mthfsd | ENSMUSG00000031816 | 234814 | -0.0514960556499195 | 2.61077462164197 | -0.372970637950183 | 0.7150016269895 | 0.782115759493935 |
| Ubtf | ENSMUSG00000020923 | 21429 | 0.01985850349012 | 5.46361814453804 | 0.372259935351439 | 0.7155184050873 | 0.782602730239163 |
| Ppp4r2 | ENSMUSG00000052144 | 232314 | -0.0244485040983973 | 5.61973449301653 | -0.371906344690051 | 0.7157755681054 | 0.782805676334245 |
| Spc24 | ENSMUSG00000074476 | 67629 | -0.0873711821310936 | 1.3745852480911 | -0.370818589430286 | 0.7165669072508 | 0.783592723405312 |
| Fam160a1 | ENSMUSG00000051000 | 229488 | -0.0287068504011838 | 5.70568605888781 | -0.370510321053337 | 0.7167912334938 | 0.783753818413581 |
| Rnf144b | ENSMUSG00000038068 | 218215 | 0.0450343013237746 | 3.94564011352017 | 0.370419096062545 | 0.7168576229529 | 0.783753818413581 |
| Ahcy | ENSMUSG00000027597 | 269378 | 0.0609902928501431 | 3.66159740933716 | 0.369895236367276 | 0.7172389107408 | 0.784092262557098 |
| Gps1 | ENSMUSG00000025156 | 209318 | 0.0257797861009976 | 4.70493389596447 | 0.368823822804837 | 0.7180189766320 | 0.784863979446159 |
| Gm10516 | ENSMUSG00000097316 | 100038712 | -0.0896430138161739 | 1.13473309277306 | -0.368677371018422 | 0.7181256295312 | 0.784863979446159 |
| Pqbp1 | ENSMUSG00000031157 | 54633 | -0.0365236142660161 | 3.34635123551601 | -0.368629854685556 | 0.7181602344177 | 0.784863979446159 |
| Prdm10 | ENSMUSG00000042496 | 382066 | -0.0430018882264536 | 2.56609123107581 | -0.368504442171821 | 0.7182515721259 | 0.784885327892549 |
| Fam219b | ENSMUSG00000032305 | 78323 | -0.0491196277477361 | 2.2565529576325 | -0.368274994962907 | 0.7184186897383 | 0.784989473963409 |
| Pard3b | ENSMUSG00000052062 | 72823 | 0.0301500568508182 | 3.98397333411537 | 0.368099347990726 | 0.7185466321626 | 0.785050798217934 |
| Trappc11 | ENSMUSG00000038102 | 320714 | 0.0246841251042769 | 4.4182994140284 | 0.367910523369331 | 0.7186841830901 | 0.785060030666921 |
| Uri1 | ENSMUSG00000030421 | 19777 | -0.0448356928541509 | 4.59662956100157 | -0.367804464454204 | 0.7187614470826 | 0.785060030666921 |
| Prpf3 | ENSMUSG00000015748 | 70767 | -0.0355490081762255 | 3.22548792536698 | -0.367791981276981 | 0.7187705412967 | 0.785060030666921 |
| Inafm2 | ENSMUSG00000074918 | 100043272 | -0.0463562398481332 | 2.26296014231382 | -0.366902360018503 | 0.7194187603330 | 0.785689526346399 |
| Map3k20 | ENSMUSG00000004085 | 65964 | -0.0321499570901231 | 9.12636828266539 | -0.366394728774512 | 0.7197887447636 | 0.786010038777589 |
| Zfp212 | ENSMUSG00000052763 | 232784 | -0.041994542638363 | 2.40534642098109 | -0.366302392534877 | 0.7198560514319 | 0.786010038777589 |
| Zkscan8 | ENSMUSG00000063894 | 93681 | 0.0441600988340936 | 3.23390350950803 | 0.366020782679098 | 0.7200613403002 | 0.786155664285057 |
| Nbr1 | ENSMUSG00000017119 | 17966 | -0.022563400538143 | 6.1922502095943 | -0.3649290176344 | 0.7208574314457 | 0.786910373726926 |
| Mepce | ENSMUSG00000029726 | 231803 | -0.0390482564555383 | 3.18664966827127 | -0.364875345177556 | 0.7208965769372 | 0.786910373726926 |
| Vps45 | ENSMUSG00000015747 | 22365 | 0.0503919167353406 | 1.80122936251431 | 0.364574608722879 | 0.7211159312727 | 0.787071217647819 |
| Trmt2a | ENSMUSG00000022721 | 15547 | -0.0431754761775411 | 2.55227136050832 | -0.363616135920757 | 0.7218152029727 | 0.787714007504857 |
| Dcaf5 | ENSMUSG00000049106 | 320808 | -0.0313373072638949 | 5.51684043352136 | -0.363569846064756 | 0.7218489811706 | 0.787714007504857 |
| Alkbh3 | ENSMUSG00000040174 | 69113 | -0.0257743489157054 | 3.90653297029939 | -0.363084360933494 | 0.7222032812388 | 0.788021967181241 |
| Dmwd | ENSMUSG00000030410 | 13401 | -0.0343845298388636 | 4.61312321051586 | -0.36179945495406 | 0.7231413076879 | 0.788966726653045 |
| 1810030O07Rik | ENSMUSG00000044148 | 69155 | 0.0351807820500531 | 2.83220452277135 | 0.361374583608455 | 0.7234515808664 | 0.789226470104863 |
| Lysmd1 | ENSMUSG00000053769 | 217779 | -0.0716056840709285 | 1.83821873324833 | -0.36111952042641 | 0.7236378716446 | 0.78935092056158 |
| Tmem33 | ENSMUSG00000037720 | 67878 | 0.0413794873244604 | 3.32164770872158 | 0.36101641065139 | 0.7237131852290 | 0.789354303306648 |
| Mmp28 | ENSMUSG00000020682 | 118453 | -0.0819217209492302 | 1.34794224266764 | -0.360496651111771 | 0.7240928741239 | 0.789689634545475 |
| Kdm6a | ENSMUSG00000037369 | 22289 | -0.0259142352885209 | 3.54506740927701 | -0.359831801698912 | 0.7245786628104 | 0.790140598880787 |
| Rbm6 | ENSMUSG00000032582 | 19654 | 0.0282245074200453 | 4.51778518794124 | 0.359620609261158 | 0.7247330016933 | 0.790151039474023 |
| Ogfr | ENSMUSG00000049401 | 72075 | 0.0345167699064506 | 3.71477008317002 | 0.359507750663941 | 0.7248154835774 | 0.790151039474023 |
| Raly | ENSMUSG00000027593 | 19383 | -0.0231699492647959 | 5.16346684239367 | -0.359425300427395 | 0.7248757439842 | 0.790151039474023 |
| Pgam1 | ENSMUSG00000011752 | 18648 | -0.0619733545423857 | 3.94382816848678 | -0.359377401602372 | 0.7249107526727 | 0.790151039474023 |
| Rhobtb3 | ENSMUSG00000021589 | 73296 | 0.035272303386332 | 3.41752490484372 | 0.359285910762379 | 0.7249776240355 | 0.790151039474023 |
| Peli2 | ENSMUSG00000021846 | 93834 | 0.053138683431599 | 2.95334371298786 | 0.359225268630088 | 0.7250219491285 | 0.790151039474023 |
| Dis3l2 | ENSMUSG00000053333 | 208718 | -0.0346266574942464 | 3.24485211083289 | -0.359013749176081 | 0.7251765628744 | 0.790204864079715 |
| Mtx3 | ENSMUSG00000021704 | 382793 | 0.0593406193813082 | 2.42905864224259 | 0.35895991249796 | 0.7252159177063 | 0.790204864079715 |
| Psma7 | ENSMUSG00000027566 | 26444 | 0.0427543929252918 | 5.57397854491204 | 0.358717807282109 | 0.7253929076133 | 0.790318934827241 |
| Agap3 | ENSMUSG00000023353 | 213990 | -0.0222676447708094 | 4.74907545711719 | -0.35807328237548 | 0.7258641643202 | 0.79075355592833 |
| Timm29 | ENSMUSG00000048429 | 69773 | -0.0239848839826893 | 4.83669826536595 | -0.357758693518915 | 0.7260942240350 | 0.790925357541347 |
| Ankrd13c | ENSMUSG00000039988 | 433667 | -0.0272867702048876 | 4.46763404187062 | -0.356711345662881 | 0.7268603509835 | 0.791680998067104 |
| Wdr75 | ENSMUSG00000025995 | 73674 | 0.0345145119446877 | 3.2723370564963 | 0.356576218597612 | 0.7269592176099 | 0.791709794529654 |
| Hnrnpr | ENSMUSG00000066037 | 74326 | 0.0235274220070885 | 4.79406222849937 | 0.356332820286626 | 0.7271373144143 | 0.791824863903464 |
| St7l | ENSMUSG00000045576 | 229681 | 0.042000172402977 | 2.71363005592782 | 0.356098225906418 | 0.7273089848632 | 0.791841688149639 |
| Crhr2 | ENSMUSG00000003476 | 12922 | 0.0715804416361214 | 4.23229573401657 | 0.356021236723124 | 0.7273653269987 | 0.791841688149639 |
| Slc15a2 | ENSMUSG00000022899 | 57738 | 0.130418361198823 | 0.69853450231 | 0.35601473626836 | 0.7273700842293 | 0.791841688149639 |
| Fam160a2 | ENSMUSG00000044465 | 74349 | -0.0315191824653347 | 3.68450873419582 | -0.355804509694362 | 0.7275239407501 | 0.791930312322211 |
| Pip4k2c | ENSMUSG00000025417 | 117150 | 0.0324329117503561 | 3.18500427039963 | 0.355588930279189 | 0.7276817275274 | 0.792023196614775 |
| Ggps1 | ENSMUSG00000021302 | 14593 | 0.0298136881015201 | 3.44681233669003 | 0.355261675106972 | 0.7279212765771 | 0.792205045227509 |
| E2f3 | ENSMUSG00000016477 | 13557 | 0.0269082249441546 | 3.15808067514559 | 0.354842900283562 | 0.7282278607709 | 0.792456800525964 |
| Fam49b | ENSMUSG00000022378 | 223601 | 0.0329377650435045 | 4.68857782855351 | 0.3547476560488 | 0.7282975956530 | 0.792456800525964 |
| Tatdn3 | ENSMUSG00000026632 | 68972 | -0.0598539884687419 | 1.86986020028022 | -0.354611106938388 | 0.7283975770632 | 0.792460411238091 |
| Psme2 | ENSMUSG00000079197 | 19188 | -0.0472975556819657 | 3.49334702262589 | -0.354545102755223 | 0.7284459072473 | 0.792460411238091 |
| Rpl35a | ENSMUSG00000060636 | 57808 | -0.0294213887011114 | 5.5070339691263 | -0.354399321043605 | 0.7285526571261 | 0.792497670917152 |
| Orc2 | ENSMUSG00000026037 | 18393 | -0.0287886982767103 | 3.61792410517191 | -0.353549523710381 | 0.7291750451491 | 0.793095763037389 |
| Gm2026 | ENSMUSG00000078886 | 100039060 | -0.0565631998934286 | 2.21005399932967 | -0.353209505325837 | 0.7294241289930 | 0.793287747888075 |
| Smug1 | ENSMUSG00000036061 | 71726 | -0.0538347540263122 | 1.90677831866857 | -0.352324368214335 | 0.7300726942901 | 0.793914108762974 |
| Gar1 | ENSMUSG00000028010 | 68147 | 0.0656195331877365 | 1.29884731935938 | 0.352013692293666 | 0.7303003865205 | 0.79406273215357 |
| Rpusd4 | ENSMUSG00000032044 | 71989 | -0.0622632546288571 | 1.6827276979377 | -0.351840980447977 | 0.7304269773047 | 0.79406273215357 |
| Adnp2 | ENSMUSG00000053950 | 240442 | 0.0366131602920499 | 2.87578968599809 | 0.351840545752266 | 0.7304272959293 | 0.79406273215357 |
| Epdr1 | ENSMUSG00000002808 | 105298 | -0.0324585233145145 | 6.08222014494147 | -0.351244491782073 | 0.7308642422990 | 0.794425471015418 |
| Wfs1 | ENSMUSG00000039474 | 22393 | -0.0304501618804869 | 7.03547834148584 | -0.351187101856434 | 0.7309063179948 | 0.794425471015418 |
| Rilpl2 | ENSMUSG00000029401 | 80291 | -0.0541097659643155 | 1.34624118144522 | -0.350601718671945 | 0.7313355460069 | 0.794812970113589 |
| Plekhf2 | ENSMUSG00000049969 | 71801 | 0.03326124474304 | 3.51181464117275 | 0.350247986312565 | 0.7315949631985 | 0.795015860694212 |
| Dcaf12 | ENSMUSG00000028436 | 68970 | -0.0322821937226063 | 4.28172785692189 | -0.349801833072854 | 0.7319222080248 | 0.795257823295986 |
| Guk1 | ENSMUSG00000020444 | 14923 | 0.0333726226639617 | 3.47300800904497 | 0.349746048541504 | 0.7319631287330 | 0.795257823295986 |
| Rfwd3 | ENSMUSG00000033596 | 234736 | -0.0382846904885121 | 3.00102212923331 | -0.34958832036806 | 0.7320788347363 | 0.79530448643446 |
| Stard13 | ENSMUSG00000016128 | 243362 | -0.0367222103117086 | 3.62899953947684 | -0.349483677886545 | 0.7321556019644 | 0.795308842797737 |
| Rpsa | ENSMUSG00000032518 | 16785 | 0.0297433669240998 | 8.50013982035129 | 0.349159294358861 | 0.7323935933768 | 0.795423830521552 |
| Haus2 | ENSMUSG00000027285 | 66296 | -0.0348703112644646 | 3.07304002126799 | -0.349141030056579 | 0.7324069942548 | 0.795423830521552 |
| Cyth2 | ENSMUSG00000003269 | 19158 | -0.0324228307255132 | 3.654745084999 | -0.348838969709519 | 0.7326286350703 | 0.795585496717082 |
| Zfp944 | ENSMUSG00000033972 | 319615 | -0.0396554611349647 | 2.16579214340318 | -0.348315511645177 | 0.7330127883072 | 0.795923590840034 |
| Rpl22 | ENSMUSG00000028936 | 19934 | -0.0305649096459186 | 6.42554124240338 | -0.348170056024043 | 0.7331195479250 | 0.795960446798614 |
| Thoc5 | ENSMUSG00000034274 | 107829 | -0.0334644975044522 | 3.12916996642234 | -0.34764784473554 | 0.7335028811793 | 0.796185511223918 |
| Bbs7 | ENSMUSG00000037325 | 71492 | 0.0487014936779784 | 1.54596178682476 | 0.347600606149275 | 0.7335375606879 | 0.796185511223918 |
| Pdgfa | ENSMUSG00000025856 | 18590 | -0.0388805017539968 | 3.33270733328301 | -0.347589988263918 | 0.7335453557347 | 0.796185511223918 |
| Rhob | ENSMUSG00000054364 | 11852 | 0.0827729617272119 | 4.86692222878016 | 0.34712249483111 | 0.7338885931515 | 0.796478972571441 |
| Sf3b2 | ENSMUSG00000024853 | 319322 | 0.0228022019513423 | 6.51475953449871 | 0.346881228326074 | 0.7340657561627 | 0.796575412012654 |
| Rpl23 | ENSMUSG00000071415 | 65019 | 0.02495641747955 | 6.60167902625535 | 0.346731399284976 | 0.7341757842081 | 0.796575412012654 |
| Cdk17 | ENSMUSG00000020015 | 237459 | 0.0384708118768953 | 3.1964443092079 | 0.346703771974984 | 0.7341960731888 | 0.796575412012654 |
| Txnip | ENSMUSG00000038393 | 56338 | 0.0982628434166255 | 7.73096339037364 | 0.346593898381276 | 0.7342767643534 | 0.796577503005279 |
| Gna11 | ENSMUSG00000034781 | 14672 | 0.0226562535987203 | 5.08646829087452 | 0.346502694539641 | 0.7343437469384 | 0.796577503005279 |
| Map3k12 | ENSMUSG00000023050 | 26404 | 0.0955978156227242 | 1.21661498941195 | 0.346290419593498 | 0.7344996561692 | 0.796667567134941 |
| Pigv | ENSMUSG00000043257 | 230801 | -0.0469624420056252 | 2.1693490633581 | -0.346121836947376 | 0.7346234834852 | 0.796722819523513 |
| 1110051M20Rik | ENSMUSG00000040591 | 228356 | -0.0370310810060897 | 2.11666293575128 | -0.345690613930703 | 0.7349402604040 | 0.796987300245648 |
| Polr1c | ENSMUSG00000067148 | 20016 | 0.0435357479639205 | 2.05076510016447 | 0.345336797139942 | 0.7352002121904 | 0.797190112037894 |
| Rybp | ENSMUSG00000072872 | 56353 | -0.041810400023135 | 2.65047171762312 | -0.345010513884639 | 0.7354399648738 | 0.797370983538618 |
| Rnaseh2c | ENSMUSG00000024925 | 68209 | -0.0436835236889102 | 2.12953391803094 | -0.344699749511109 | 0.7356683410258 | 0.797539485842864 |
| Gm45871 | ENSMUSG00000110277 | 108168395 | -0.0888637449530081 | 1.0643993844834 | -0.344152303576141 | 0.7360707143397 | 0.797896566684588 |
| Nomo1 | ENSMUSG00000030835 | 211548 | -0.0259620252576012 | 6.05863773215903 | -0.343135402387886 | 0.7368183519880 | 0.798627804222279 |
| Setdb1 | ENSMUSG00000015697 | 84505 | -0.0343281542412313 | 3.7531124405644 | -0.342976688364691 | 0.7369350654883 | 0.798675114103997 |
| Caprin2 | ENSMUSG00000030309 | 232560 | -0.0555080990170096 | 1.66070549674179 | -0.34191913945694 | 0.7377129276885 | 0.799438882974502 |
| Adprhl1 | ENSMUSG00000031448 | 234072 | 0.0336739424794672 | 7.45664642634952 | 0.341480809572474 | 0.7380354217141 | 0.79970907957542 |
| Ints9 | ENSMUSG00000021975 | 210925 | 0.0347372040029361 | 2.63862346890608 | 0.34022935564505 | 0.7389564412528 | 0.800627699408748 |
| Mbnl2 | ENSMUSG00000022139 | 105559 | -0.0202218918266981 | 7.75135188125866 | -0.339081604973699 | 0.7398015068015 | 0.801458472813952 |
| Neu2 | ENSMUSG00000079434 | 23956 | 0.049130942936817 | 2.04391590158827 | 0.338988788704958 | 0.7398698608695 | 0.801458472813952 |

| | | | | | | | |
|---|---|---|---|---|---|---|---|
| Reep3 | ENSMUSG00000019873 | 28193 | 0.0439803967407499 | 5.07380968399117 | 0.338459478205203 | 0.7402597126688 | 0.801801319776426 |
| Zmym2 | ENSMUSG00000021945 | 76007 | -0.0270148461564865 | 4.59122858469568 | -0.337707015635038 | 0.7408140502074 | 0.802322241436408 |
| Dicer1 | ENSMUSG00000041415 | 192119 | -0.0189825717502359 | 5.44278289185747 | -0.337560196897155 | 0.7409222287638 | 0.802359905153375 |
| Hdac10 | ENSMUSG00000062906 | 170787 | -0.0551354710332626 | 1.70811333279434 | -0.337452604359668 | 0.7410015084125 | 0.802366269287572 |
| Rps2 | ENSMUSG00000044533 | 16898 | -0.0229256326647862 | 7.86118564429419 | -0.337284013510732 | 0.7411257408847 | 0.802421302853751 |
| Otud1 | ENSMUSG00000043415 | 71198 | 0.123494745976391 | 5.01835321904911 | 0.336897873194965 | 0.7414103111776 | 0.802649907049855 |
| Rps6 | ENSMUSG00000028495 | 20104 | 0.0268739968498162 | 7.171367899615 | 0.336489496096103 | 0.7417113120313 | 0.802896251912687 |
| Ncoa1 | ENSMUSG00000020647 | 17977 | -0.0239986447466443 | 5.11314149422845 | -0.335904060557993 | 0.7421428935076 | 0.803283886417656 |
| Col11a1 | ENSMUSG00000027966 | 12814 | 0.0737022298798635 | 4.30337712541503 | 0.335439457460052 | 0.7424854620201 | 0.803575107459631 |
| Nrf1 | ENSMUSG00000058440 | 18181 | 0.0280603448417498 | 3.14294762531439 | 0.334537234011817 | 0.7431508661660 | 0.804215633903637 |
| Shoc2 | ENSMUSG00000024976 | 56392 | -0.0238182070153066 | 4.54089111935063 | -0.334371324246899 | 0.7432732505735 | 0.804268451991607 |
| Pwp1 | ENSMUSG00000001785 | 103136 | 0.0312432626828891 | 3.55749919967445 | 0.333582519194545 | 0.7438552166321 | 0.804818506681752 |
| Ncl | ENSMUSG00000026234 | 17975 | 0.0237547517198102 | 7.81214715838296 | 0.333130775738971 | 0.7441885782866 | 0.805099500123844 |
| Ankrd39 | ENSMUSG00000079610 | 109346 | -0.070538357093209 | 1.18508683605081 | -0.332549675446861 | 0.7446174767457 | 0.805483784097762 |
| Nup160 | ENSMUSG00000051329 | 59015 | -0.0304608887204925 | 3.03828012643572 | -0.331598707482589 | 0.7453195576356 | 0.806095766358135 |
| Tut1 | ENSMUSG00000071645 | 70044 | -0.0341490269799007 | 2.46155396767797 | -0.331583614160038 | 0.7453307026421 | 0.806095766358135 |
| Stard8 | ENSMUSG00000031216 | 236920 | -0.028347485713816 | 4.2144178285238 | -0.331295714111253 | 0.7455433012715 | 0.806245926612492 |
| Epm2aip1 | ENSMUSG00000046785 | 77781 | -0.0206947265451287 | 4.75123246539139 | -0.330540962125715 | 0.7461007473947 | 0.806716933948879 |
| Mkrn1 | ENSMUSG00000029922 | 54484 | -0.0245870752805994 | 3.55928889796045 | -0.33050616984238 | 0.7461264479208 | 0.806716933948879 |
| Slc24a2 | ENSMUSG00000037996 | 76376 | -0.0735021559816056 | 1.42340539773184 | -0.330343913259786 | 0.7462463085298 | 0.80676672916027 |
| Rpl18 | ENSMUSG00000059070 | 19899 | 0.0261134081519332 | 6.28389111197122 | 0.32982840595984 | 0.7466271645108 | 0.807098648661803 |
| Fgf7 | ENSMUSG00000027208 | 14178 | -0.0516981882791656 | 2.08185833042635 | -0.329524204166229 | 0.7468519407105 | 0.807261798072455 |
| Ccp110 | ENSMUSG00000033904 | 101565 | 0.0580898057514474 | 1.7748926196996 | 0.328889606299007 | 0.7473209254881 | 0.807630089332415 |
| Parp3 | ENSMUSG00000023249 | 235587 | 0.0228374688721988 | 5.46813860010932 | 0.32877719986268 | 0.7474040077318 | 0.807630089332415 |
| Kdm7a | ENSMUSG00000042599 | 338523 | 0.0268092594246081 | 5.68677519171892 | 0.328755805454155 | 0.7474198212137 | 0.807630089332415 |
| Arl4a | ENSMUSG00000047446 | 11861 | 0.031827860994885 | 4.38531439636184 | 0.328663283807948 | 0.7474882091095 | 0.807630089332415 |
| R3hcc1 | ENSMUSG00000034194 | 71843 | -0.027796597292745 | 3.05982853389964 | -0.328319559568028 | 0.7477422942288 | 0.807824769540971 |
| Usp12 | ENSMUSG00000029640 | 22217 | 0.0231518697867141 | 4.77647410080403 | 0.327659848610218 | 0.7482300461709 | 0.808271828707845 |
| Thap11 | ENSMUSG00000036442 | 59016 | -0.0377529503422127 | 3.0647573202903 | -0.327524277274462 | 0.7483302936680 | 0.808300241115123 |
| Ttc14 | ENSMUSG00000027677 | 67120 | 0.0364610240344558 | 3.79444137097449 | 0.327359355774221 | 0.7484522503843 | 0.808352094551059 |
| Pdcd6 | ENSMUSG00000021576 | 18570 | 0.0376791117559563 | 2.88829124240794 | 0.326230274956302 | 0.7492873753086 | 0.809105859261926 |
| Rraga | ENSMUSG00000070934 | 68441 | 0.0287571487406124 | 4.21355811084344 | 0.32621564433887 | 0.7492981990036 | 0.809105859261926 |
| Cmtr1 | ENSMUSG00000024019 | 74157 | 0.0256953348050075 | 4.46865486536402 | 0.326095322334132 | 0.7493872150167 | 0.809122051298716 |
| Arpc1a | ENSMUSG00000029621 | 56443 | -0.0326158439919055 | 3.24810619512847 | -0.325748253943262 | 0.7496440022450 | 0.809319366769474 |
| Arl15 | ENSMUSG00000042348 | 218639 | -0.052166413224624 | 2.33938612308096 | -0.325226015768254 | 0.7500304515408 | 0.809656613252306 |
| Rpl7a | ENSMUSG00000062647 | 27176 | -0.0251338502432121 | 6.91521384553955 | -0.323594384840838 | 0.7512382871679 | 0.810880390740884 |
| Dennd2a | ENSMUSG00000038456 | 209773 | 0.0479805343311992 | 2.86010926216415 | 0.322977219793569 | 0.7516953289316 | 0.811293605899703 |
| Ppp2r5c | ENSMUSG00000017843 | 26931 | 0.0180018787553005 | 5.86737689539402 | 0.322333521117213 | 0.7521721236879 | 0.811728056474824 |
| Cln6 | ENSMUSG00000032245 | 76524 | 0.0333222628512027 | 2.63233695384273 | 0.32138815060974 | 0.7528725613210 | 0.812385371004726 |
| Kat7 | ENSMUSG00000038909 | 217127 | -0.0233305717616572 | 4.15375205646705 | -0.321283026973022 | 0.7529504628610 | 0.812385371004726 |
| Adcy1 | ENSMUSG00000020431 | 432530 | -0.0762866977312817 | 1.67479481303449 | -0.321210554891769 | 0.7530041697026 | 0.812385371004726 |
| Nom1 | ENSMUSG00000001569 | 433864 | -0.0221444989153285 | 4.59180453929809 | -0.320951512920343 | 0.7531961486782 | 0.812512296575772 |
| Dnaaf5 | ENSMUSG00000025857 | 433956 | -0.0279369195288721 | 3.44509119929495 | -0.320495585585216 | 0.7535340829918 | 0.812796631259483 |
| Pom121 | ENSMUSG00000053293 | 107939 | 0.038701682482507 | 3.94281498189925 | 0.320079644013192 | 0.7538424257127 | 0.813048994125871 |
| Cggbp1 | ENSMUSG00000054604 | 106143 | 0.0293477887231344 | 4.05299861810974 | 0.319710793672171 | 0.7541158957404 | 0.813263699323095 |
| Fto | ENSMUSG00000055932 | 26383 | 0.0206913120216718 | 5.86275803192644 | 0.318659194411354 | 0.7548957526318 | 0.814024412747221 |
| Irf1 | ENSMUSG00000018899 | 16362 | 0.0414935126503679 | 2.80706347513865 | 0.318378156283007 | 0.7551042153084 | 0.814074348628581 |
| Phlda3 | ENSMUSG00000041801 | 27280 | -0.0230051084085972 | 5.89103698739149 | -0.318367047014362 | 0.7551124561218 | 0.814074348628581 |
| Yipf4 | ENSMUSG00000024072 | 67864 | -0.0297385078074317 | 3.68181726932695 | -0.318295563536706 | 0.7551654830384 | 0.814074348628581 |
| Hltf | ENSMUSG00000002428 | 20585 | 0.0489042269546616 | 2.36287947342372 | 0.317846874543153 | 0.7554983528796 | 0.814352874009228 |
| Polr1a | ENSMUSG00000049553 | 20019 | 0.0273691748929057 | 5.77759021407814 | 0.315742467736 | 0.7570602281921 | 0.815955960793605 |
| Rsad2 | ENSMUSG00000020641 | 58185 | -0.0661157748287926 | 3.3213539791446 | -0.315536781894559 | 0.7572129462280 | 0.815973581846144 |
| Rell1 | ENSMUSG00000047881 | 100532 | 0.0244421614686066 | 3.43675315269616 | 0.315519372765626 | 0.7572258726783 | 0.815973581846144 |
| Lsg1 | ENSMUSG00000022538 | 224092 | -0.026318642250327 | 3.42249710572134 | -0.315220517798111 | 0.7574477872530 | 0.816132258498071 |
| Trappc2l | ENSMUSG00000015013 | 59005 | -0.0373712179379595 | 2.60341247786285 | -0.315033030061521 | 0.7575870175429 | 0.81620182227105 |
| Phax | ENSMUSG00000008301 | 56698 | -0.0320421620860563 | 3.14172742050171 | -0.314603186922768 | 0.7579062565098 | 0.81646528923909 |
| Nsun5 | ENSMUSG00000000916 | 100609 | -0.0426287481249121 | 1.92431104298252 | -0.314255510543298 | 0.7581645049753 | 0.816663007872015 |
| Plbd2 | ENSMUSG00000029598 | 71772 | 0.0228731793066546 | 4.80796963160896 | 0.313226614568756 | 0.7589289287189 | 0.817388770946596 |
| Ppp2r1b | ENSMUSG00000032058 | 73699 | 0.0618712711162894 | 1.06407769673517 | 0.313147342734635 | 0.7589878350661 | 0.817388770946596 |
| Lgals9 | ENSMUSG00000001123 | 16859 | 0.0540792070756866 | 3.42904749014906 | 0.313016854281117 | 0.7590848035232 | 0.817412667452712 |
| Slc16a6 | ENSMUSG00000041920 | 104681 | -0.0501390107509736 | 3.22973473223267 | -0.311625157530696 | 0.7601192624412 | 0.818445986775541 |
| Ddhd2 | ENSMUSG00000061313 | 72108 | -0.0268130278506806 | 3.37086462973053 | -0.310769112259257 | 0.7607558054159 | 0.818872283591134 |
| Cct4 | ENSMUSG00000007739 | 12464 | -0.0201881620386519 | 5.44492874727599 | -0.310757151547294 | 0.7607647005107 | 0.818872283591134 |
| Tle2 | ENSMUSG00000034771 | 21886 | 0.0514687843268626 | 1.68063927220342 | 0.310745640266249 | 0.7607732614002 | 0.818872283591134 |
| Ero1l | ENSMUSG00000021831 | 50527 | -0.0359890573174146 | 2.43178362595124 | -0.310689745584332 | 0.7608148304959 | 0.818872283591134 |
| Micu3 | ENSMUSG00000039478 | 78506 | 0.0457027988167926 | 1.96858885018918 | 0.309237145702938 | 0.7618954045064 | 0.819954579763721 |
| Snx11 | ENSMUSG00000020876 | 74479 | -0.032193771565681 | 2.37107021490099 | -0.309065097331572 | 0.7620234238620 | 0.820011621011665 |
| Naa16 | ENSMUSG00000022020 | 66897 | 0.0447733409858646 | 2.54278288764285 | 0.307959650007981 | 0.7628461485228 | 0.820816148194059 |
| Ppig | ENSMUSG00000042133 | 228005 | 0.0200626677949689 | 5.45771731599791 | 0.306752157282734 | 0.7637451616046 | 0.821646900217047 |
| Filip1 | ENSMUSG00000034898 | 70598 | -0.0241349019916042 | 7.04081300379087 | -0.306720733625229 | 0.7637685621723 | 0.821646900217047 |
| Eef1e1 | ENSMUSG00000001707 | 66143 | 0.0411291475121023 | 2.03145499402402 | 0.306309446920497 | 0.7640748613159 | 0.821895523424956 |
| Fn3krp | ENSMUSG00000039253 | 238024 | 0.0411126324227526 | 1.47633551128643 | 0.304715574022488 | 0.7652622610487 | 0.823091781513724 |
| Rsad1 | ENSMUSG00000039096 | 237926 | 0.0478389685906893 | 1.80040580547479 | 0.304341668522363 | 0.7655409015532 | 0.823310467815697 |
| Rnf25 | ENSMUSG00000026171 | 57751 | -0.0475288824455249 | 2.41712925113259 | -0.303834536954526 | 0.7659188783759 | 0.823635933057326 |
| Tomm34 | ENSMUSG00000018322 | 67145 | -0.0252405313044437 | 3.22568420130412 | -0.303562214005094 | 0.7661218725925 | 0.823739185484788 |
| Cd2bp2 | ENSMUSG00000042502 | 70233 | -0.0189365035026254 | 3.81921709652883 | -0.303503540129161 | 0.7661656114558 | 0.823739185484788 |
| Zfpl1 | ENSMUSG00000024792 | 81909 | 0.0435715248272124 | 1.83408629820804 | 0.303401471752203 | 0.7662417010431 | 0.82373997188267 |
| Zbtb24 | ENSMUSG00000019826 | 268294 | -0.046267328530877 | 1.86543395805396 | -0.30295933091672 | 0.7665713357256 | 0.824013302273263 |
| Efr3a | ENSMUSG00000015002 | 76740 | -0.0189134922576972 | 5.70388303687239 | -0.302521385471805 | 0.7668978890280 | 0.824283266958232 |
| Leo1 | ENSMUSG00000042487 | 235497 | -0.0267512445739415 | 4.41256361294496 | -0.301303793982471 | 0.7678060266128 | 0.825178220126292 |
| Taf4b | ENSMUSG00000054321 | 72504 | 0.0603226335647691 | 1.33464773847208 | 0.30074060019416 | 0.7682262036849 | 0.82549844405041 |
| H1f0 | ENSMUSG00000096210 | 14958 | 0.022085547803802 | 6.09590085235347 | 0.300701973482038 | 0.7682550243641 | 0.82549844405041 |
| Syde1 | ENSMUSG00000032714 | 71709 | -0.0498325808067404 | 2.41702403130641 | -0.300219766606538 | 0.7686148450696 | 0.82580389929787 |
| Rai2 | ENSMUSG00000043518 | 24004 | -0.027349202741084 | 3.93681196067873 | -0.29976609589854 | 0.7689534229788 | 0.826017232324992 |
| Ctbs | ENSMUSG00000028189 | 74245 | 0.0458789455819291 | 1.62811441193374 | 0.29967830795071 | 0.7690189454719 | 0.826017232324992 |
| Rft1 | ENSMUSG00000052395 | 328370 | -0.0558346586257321 | 1.29114452364095 | -0.299649960410175 | 0.7690401036841 | 0.826017232324992 |
| Stxbp6 | ENSMUSG00000046314 | 217517 | 0.0490717155172486 | 1.70643335548251 | 0.299498405303588 | 0.7691532255741 | 0.826057566196815 |
| Ddx50 | ENSMUSG00000020076 | 94213 | 0.0239374599323723 | 5.1951837988701 | 0.298631863644171 | 0.7698001240818 | 0.826590086002738 |
| Tubgcp5 | ENSMUSG00000033790 | 233276 | 0.0324097150838329 | 3.39257865389556 | 0.298631628493275 | 0.7698002996530 | 0.826590086002738 |
| Prrc2b | ENSMUSG00000039262 | 227723 | -0.017500071324613 | 7.46577102748625 | -0.297532373564625 | 0.7706211825426 | 0.827390251092608 |
| Dock4 | ENSMUSG00000035954 | 238130 | 0.0275017225446819 | 4.05373113816 | 0.296803345643584 | 0.7711657517686 | 0.827893619385489 |
| Tm2d2 | ENSMUSG00000031556 | 69742 | 0.029713939831626 | 3.27450544082338 | 0.296566165156458 | 0.7713429479696 | 0.828002530124332 |
| Rex1bd | ENSMUSG00000058833 | 66462 | -0.0244995040669154 | 3.94782135177071 | -0.296003116439 | 0.7717636517563 | 0.828372788427745 |
| Olfr784 | ENSMUSG00000094347 | 258724 | 0.0648123807998329 | 2.08817006381025 | 0.294767507509236 | 0.7726871473433 | 0.829264752236166 |
| Mertk | ENSMUSG00000014361 | 17289 | 0.0533074861076983 | 2.14617257967834 | 0.294688261852389 | 0.7727463879130 | 0.829264752236166 |
| Sptssa | ENSMUSG00000044408 | 104725 | -0.032310652096792 | 2.84707063443597 | -0.294105040790207 | 0.7731824237576 | 0.829651229419373 |
| Noc4l | ENSMUSG00000033294 | 100608 | 0.0435402435467198 | 2.2660928361822 | 0.292611962113703 | 0.7742990636125 | 0.830767870470017 |
| Tex10 | ENSMUSG00000028345 | 269536 | 0.0286158600098574 | 3.07344213105619 | 0.292118841006294 | 0.7746679723391 | 0.831082108709339 |
| Pcbp4 | ENSMUSG00000023495 | 59092 | 0.0248400364640489 | 5.30604808930736 | 0.291627742093701 | 0.7750354246423 | 0.83139472294071 |
| Stat2 | ENSMUSG00000040033 | 20847 | 0.0534108195387737 | 3.11985249614184 | 0.291321063694848 | 0.7752649174854 | 0.831502045201451 |
| Scarna6 | ENSMUSG00000089281 | 100217415 | 0.0808434371767777 | 2.67639219496505 | 0.291202694598384 | 0.7753535010210 | 0.831502045201451 |
| Ddx19a | ENSMUSG00000015023 | 13680 | 0.0234777458565301 | 4.26599317487384 | 0.291189098101046 | 0.7753636764009 | 0.831502045201451 |
| Fuz | ENSMUSG00000011658 | 70300 | -0.0369378357298147 | 2.38398569098855 | -0.289844523562132 | 0.7763701445658 | 0.832499710638596 |
| Tbc1d13 | ENSMUSG00000039678 | 70296 | 0.0264649033132973 | 2.81496612511993 | 0.289449897369787 | 0.7766656177597 | 0.832695889025801 |
| Rpl19 | ENSMUSG00000017404 | 19921 | -0.0233225297017929 | 7.23940440867712 | -0.289396700875465 | 0.7767054509657 | 0.832695889025801 |
| Rbbp7 | ENSMUSG00000031353 | 245688 | -0.0182740279739848 | 5.91381638375592 | -0.288990923416397 | 0.7770093161605 | 0.8329399661617 |
| Fbxo34 | ENSMUSG00000037536 | 78938 | -0.0300534878737526 | 3.06000777073383 | -0.288701948554016 | 0.7772257372963 | 0.833042098641192 |
| Ghdc | ENSMUSG00000017747 | 80860 | 0.0321748932910331 | 2.34566097878474 | 0.288660195515372 | 0.7772570088776 | 0.833042098641192 |
| Chmp3 | ENSMUSG00000053119 | 66700 | 0.0187381612591162 | 5.17291748313195 | 0.288038119382985 | 0.7777229700218 | 0.833459782873421 |
| Rassf8 | ENSMUSG00000030259 | 71323 | -0.0212412822298775 | 5.1618754935663 | -0.286881457026263 | 0.7785895956210 | 0.834306721863862 |
| Cxcl9 | ENSMUSG00000029417 | 17329 | 0.154970176449365 | 0.488973606761811 | 0.286768445182439 | 0.7786742858774 | 0.834315685054574 |
| Tmem18 | ENSMUSG00000043061 | 211986 | 0.0349048660083884 | 1.85229628126192 | 0.285684553295838 | 0.7794866953344 | 0.835033599940428 |
| Puf60 | ENSMUSG00000002524 | 67959 | -0.018830745349927 | 5.20257226936009 | -0.285670667484185 | 0.7794971049119 | 0.835033599940428 |
| Nupl1 | ENSMUSG00000114797 | 71844 | 0.0278118742328681 | 4.69044944088227 | 0.284221084778819 | 0.7805840350680 | 0.836116030115569 |
| Rrp1 | ENSMUSG00000061032 | 18114 | 0.0226797728391732 | 5.45855390240374 | 0.284014671621651 | 0.7807388473245 | 0.836199915746092 |
| Eps15 | ENSMUSG00000028552 | 13858 | -0.0153660924736743 | 5.60794073692625 | -0.283746245972504 | 0.7809401841678 | 0.836333609595225 |

| | | | | | | | |
|---|---|---|---|---|---|---|---|
| Tshz1 | ENSMUSG00000046982 | 110796 | -0.0294397352959095 | 4.25035854208744 | -0.283354077224886 | 0.7812343658966 | 0.836566698042314 |
| Gm32742 | ENSMUSG00000110266 | 102635385 | -0.0885980413224435 | 0.726055243049063 | -0.2827112253345 | 0.7817166707678 | 0.837001168396827 |
| Zfp809 | ENSMUSG00000057982 | 235047 | 0.0389089161743228 | 2.03763274287586 | 0.28155450609418 | 0.7825847445240 | 0.837848564387107 |
| Itpr1 | ENSMUSG00000030102 | 16438 | -0.0361152851012712 | 4.53566143369199 | -0.281377146204982 | 0.7827178731063 | 0.837862605325409 |
| Med1 | ENSMUSG00000018160 | 19014 | 0.0212495934382666 | 5.3189685897397 | 0.281332801918352 | 0.7827511595995 | 0.837862605325409 |
| Rexo2 | ENSMUSG00000032026 | 104444 | -0.0239123500827763 | 4.15833658748085 | -0.281196963164208 | 0.7828531280635 | 0.837889703599564 |
| Cnbp | ENSMUSG00000030057 | 12785 | -0.0235494095152973 | 7.95524406058979 | -0.280979271155476 | 0.7830165489792 | 0.837982562844399 |
| Dgkh | ENSMUSG00000034731 | 380921 | -0.0390382921598 | 3.35103986419113 | -0.2808099456726 | 0.7831436686442 | 0.838007445509383 |
| Zc3h15 | ENSMUSG00000027091 | 69082 | -0.0173236772617873 | 5.75425079298729 | -0.280744069618604 | 0.7831931262760 | 0.838007445509383 |
| Zfp276 | ENSMUSG00000001065 | 57247 | -0.0427976063524462 | 2.82646088326379 | -0.28018483514831 | 0.7836130209462 | 0.838374663009015 |
| Adam15 | ENSMUSG00000028041 | 11490 | -0.0357194773379694 | 4.38320945181913 | -0.279876035796997 | 0.7838449094650 | 0.838540683633086 |
| Kri1 | ENSMUSG00000035047 | 215194 | 0.0281712328234063 | 3.10100500044102 | 0.279083292543083 | 0.7844403068242 | 0.839095507769468 |
| Vps51 | ENSMUSG00000024797 | 68505 | 0.0327264631919726 | 2.95842586937867 | 0.278903606641844 | 0.7845752812094 | 0.839157768972602 |
| Rps3 | ENSMUSG00000030744 | 27050 | 0.0240927033736709 | 6.74852134684969 | 0.278448438862864 | 0.7849172212288 | 0.839381560433858 |
| Snap47 | ENSMUSG00000009894 | 67826 | 0.021732666032499 | 3.99945806627093 | 0.278420655778947 | 0.7849380944794 | 0.839381560433858 |
| Znfx1 | ENSMUSG00000039501 | 98999 | -0.0238313861429349 | 5.5515039194907 | -0.278206442036615 | 0.7850990376648 | 0.839471542669877 |
| Capn15 | ENSMUSG00000037326 | 50817 | 0.0409663701677467 | 2.5116118244255 | 0.277090819954461 | 0.7859373929598 | 0.840285763149838 |
| Srp68 | ENSMUSG00000020780 | 217337 | -0.0186548774908637 | 5.64906211040786 | -0.276736365655246 | 0.7862038123641 | 0.840488398332777 |
| Dcaf13 | ENSMUSG00000022300 | 223499 | 0.0267417362120601 | 3.04272503863405 | 0.27505068993933 | 0.7874712028001 | 0.841760973773593 |
| Taf11 | ENSMUSG00000024218 | 68776 | -0.0286443155894514 | 2.76251277660018 | -0.274724824360828 | 0.7877162803153 | 0.841940614073265 |
| Vps18 | ENSMUSG00000034216 | 228545 | -0.0252425831899346 | 2.97547240038201 | -0.272565393451151 | 0.7893409399226 | 0.8435946240022 |
| Setd3 | ENSMUSG00000056770 | 52690 | -0.0178336651142788 | 6.07929133783409 | -0.271913357154401 | 0.7898317042522 | 0.8440365978279 |
| Zfp444 | ENSMUSG00000044876 | 72667 | -0.0544773593709278 | 1.22008160838545 | -0.271221569178042 | 0.7903524899161 | 0.844495062583353 |
| Map7 | ENSMUSG00000019996 | 17761 | -0.0387294469003034 | 1.72590922229229 | -0.271138224575338 | 0.7904152397118 | 0.844495062583353 |
| Fbxw2 | ENSMUSG00000035949 | 30050 | -0.0205731699786371 | 4.22590270058042 | -0.269793034908613 | 0.7914282366023 | 0.845494727746285 |
| Nolc1 | ENSMUSG00000015176 | 70769 | -0.0181358805452678 | 4.70964758517604 | -0.26945172041535 | 0.7916853265214 | 0.845649627529203 |
| Dhx8 | ENSMUSG00000034931 | 217207 | -0.0221860356202761 | 4.16857038821386 | -0.269395128312304 | 0.7917279561004 | 0.845649627529203 |
| Ccni | ENSMUSG00000063015 | 12453 | 0.0230122370470749 | 5.75001754974187 | 0.269069751815745 | 0.7919730684895 | 0.845766987730062 |
| Slc5a3 | ENSMUSG00000089774 | 53881 | -0.0348520271590651 | 3.71321584322789 | -0.269043853083258 | 0.7919925794899 | 0.845766987730062 |
| Aimp1 | ENSMUSG00000028029 | 13722 | -0.0264808099892777 | 5.12585441198744 | -0.268539817982072 | 0.7923723269544 | 0.846041255148519 |
| Tef | ENSMUSG00000022389 | 21685 | 0.0447618806262495 | 6.0967110875461 | 0.268497509232562 | 0.7924042054899 | 0.846041255148519 |
| Prickle2 | ENSMUSG00000030020 | 243548 | 0.0430291852986806 | 2.60044728212174 | 0.268015120867045 | 0.7927676998505 | 0.846346686889988 |
| Trdn | ENSMUSG00000019787 | 76757 | 0.0304537942735319 | 9.41238057933004 | 0.26670068951954 | 0.7937584186938 | 0.847321608861542 |
| Vezt | ENSMUSG00000036099 | 215008 | -0.0218412059897677 | 4.8652200562372 | -0.266193257802802 | 0.7941409817405 | 0.847647209394245 |
| Ino80b | ENSMUSG00000030034 | 70020 | 0.0264000032473248 | 2.90941339116989 | 0.265691437456887 | 0.7945193685902 | 0.847865211928262 |
| Zfp213 | ENSMUSG00000071256 | 449521 | -0.0466589135666675 | 1.27781491999213 | -0.265640287943892 | 0.7945579398119 | 0.847865211928262 |
| Zbtb33 | ENSMUSG00000048047 | 56805 | -0.0272503925309504 | 2.87232892565598 | -0.265613793916457 | 0.7945779188540 | 0.847865211928262 |
| Ccn1 | ENSMUSG00000028195 | 16007 | -0.0678911087676903 | 5.38955113799075 | -0.264300164476186 | 0.7955687098443 | 0.84883958685301 |
| Nasp | ENSMUSG00000028693 | 50927 | -0.0205250166963769 | 3.29473525941745 | -0.263831054684106 | 0.7959226201140 | 0.849134311972132 |
| D2hgdh | ENSMUSG00000073609 | 98314 | -0.028506156155983 | 2.89558629827244 | -0.263018261489291 | 0.7965359261003 | 0.84970569027062 |
| Cd24a | ENSMUSG00000047139 | 12484 | 0.0225757827531323 | 5.05426704801343 | 0.260749802001158 | 0.7982483685084 | 0.851449347791389 |
| Trappc3 | ENSMUSG00000028847 | 27096 | -0.0296558513122815 | 3.02281592407347 | -0.259952492412228 | 0.7988505090501 | 0.852008480283611 |
| Odr4 | ENSMUSG00000006010 | 226499 | 0.0234813656032982 | 4.18836475102213 | 0.259113719245636 | 0.7994841075641 | 0.852455507844815 |
| Slc25a16 | ENSMUSG00000071253 | 73132 | -0.020965634232752 | 3.58392203943727 | -0.259078813782168 | 0.7995104778991 | 0.852455507844815 |
| Washc3 | ENSMUSG00000020056 | 67282 | 0.0235099073286333 | 2.71344534864487 | 0.259071947365809 | 0.7995156653605 | 0.852455507844815 |
| Cep162 | ENSMUSG00000056919 | 382090 | -0.0315702417459769 | 2.83754551257061 | -0.258984691579757 | 0.7995815864909 | 0.852455507844815 |
| Rnf166 | ENSMUSG00000014470 | 68718 | -0.026505496520822 | 4.24653461166349 | -0.258574513901141 | 0.7998914941669 | 0.852702742611594 |
| Sh3bp5l | ENSMUSG00000013646 | 79566 | 0.0239417223296101 | 3.62397814597266 | 0.257905504856788 | 0.8003970359672 | 0.85315845930352 |
| 5930430L01Rik | ENSMUSG00000106951 | 319982 | -0.0503594938767042 | 1.59825756099086 | -0.257252103530612 | 0.8008908737183 | 0.853601612774459 |
| Vamp2 | ENSMUSG00000020894 | 22318 | 0.0183968282847848 | 4.76610069004061 | 0.256998126526983 | 0.8010828523528 | 0.853656464532301 |
| Zfp868 | ENSMUSG00000060427 | 234362 | 0.0427987957465974 | 1.49809562639479 | 0.256977388868737 | 0.8010985283297 | 0.853656464532301 |
| Natd1 | ENSMUSG00000018931 | 24083 | -0.0256648987040426 | 3.44344157615381 | -0.256663084496488 | 0.8013361277130 | 0.85382641700275 |
| Gtf3c1 | ENSMUSG00000032777 | 233863 | 0.0168342369806744 | 5.47127679551681 | 0.256250701183203 | 0.8016479012675 | 0.854075361477182 |
| Rplp1 | ENSMUSG00000007892 | 56040 | 0.0207974546999106 | 7.60241862364695 | 0.255134325544261 | 0.8024920897595 | 0.854891436815236 |
| Por | ENSMUSG00000005514 | 18984 | 0.0187380940116162 | 4.23962714852111 | 0.254570856836743 | 0.8029182750679 | 0.855207302404829 |
| Tpgs1 | ENSMUSG00000020308 | 110012 | -0.0381306679887419 | 2.65268630919268 | -0.254459535350431 | 0.8030024819493 | 0.855207302404829 |
| Cct5 | ENSMUSG00000022234 | 12465 | 0.0193602988561148 | 6.06290833565043 | 0.254432007204213 | 0.8030233054508 | 0.855207302404829 |
| Nsdhl | ENSMUSG00000031349 | 18194 | 0.0508099547652636 | 1.24950015095871 | 0.25368962192699 | 0.8035849371249 | 0.855586198381974 |
| Vstm4 | ENSMUSG00000050666 | 320736 | -0.0571648476591173 | 1.28759639633427 | -0.253689141926497 | 0.8035853002931 | 0.855586198381974 |
| Trp53rka | ENSMUSG00000039725 | 381406 | -0.0312984060048414 | 2.63184393576647 | -0.253651346889732 | 0.8036138961474 | 0.855586198381974 |
| Memo1 | ENSMUSG00000058704 | 76890 | -0.0190648720529012 | 4.09243626475675 | -0.252994010119015 | 0.8041112862579 | 0.856032379244752 |
| Znrf2 | ENSMUSG00000058446 | 387524 | -0.0143112076446199 | 4.90153518315273 | -0.252816436138538 | 0.8042456671853 | 0.85609206232379 |
| Ccdc115 | ENSMUSG00000042111 | 69668 | -0.0276863295989322 | 2.96127874808337 | -0.252560273033322 | 0.8044395326039 | 0.856193425579399 |
| Nabp1 | ENSMUSG00000026107 | 109019 | 0.0417682340174617 | 2.65473866403234 | 0.252483620912732 | 0.8044975458902 | 0.856193425579399 |
| Ccdc127 | ENSMUSG00000021578 | 67433 | -0.0156378731459217 | 4.35940010799555 | -0.252096541394838 | 0.8047905209599 | 0.85642184429651 |
| Sobp | ENSMUSG00000038248 | 109205 | -0.0255781012503147 | 3.99897973913085 | -0.251983145251309 | 0.8048763547070 | 0.856429809530148 |
| Mr1 | ENSMUSG00000026471 | 15064 | -0.0311438907632189 | 3.66488394538365 | -0.251444118460592 | 0.8052843999306 | 0.856780589414244 |
| Nipa2 | ENSMUSG00000030452 | 93790 | -0.0204765871300641 | 3.43230946655998 | -0.251160717299907 | 0.8054989593653 | 0.856925462269809 |
| Mia3 | ENSMUSG00000056050 | 338366 | 0.0181666663264712 | 6.73226465419583 | 0.250478924116525 | 0.8060152032499 | 0.85739122097363 |
| Czib | ENSMUSG00000028608 | 74098 | -0.0204879882414306 | 3.29220308264434 | -0.250373932848747 | 0.8060947094965 | 0.857392358616976 |
| Sart3 | ENSMUSG00000018974 | 53890 | 0.0245270918530602 | 3.37345321909221 | 0.250078997581463 | 0.8063180657053 | 0.857546485330283 |
| Krit1 | ENSMUSG00000000600 | 79264 | 0.0177795079864405 | 4.49904462623854 | 0.247676385961399 | 0.8081382323486 | 0.859398678646035 |
| Triap1 | ENSMUSG00000029535 | 69076 | -0.0324551802856252 | 1.88311302814863 | -0.247484641447732 | 0.8082835439247 | 0.85946959325303 |
| Slc25a27 | ENSMUSG00000023912 | 74011 | 0.0595184303429785 | 1.07899825719189 | 0.246964287003849 | 0.8086779262593 | 0.859712859395306 |
| Kcne4 | ENSMUSG00000047330 | 57814 | 0.0851956035472697 | 0.693947340598056 | 0.246801844671493 | 0.8088010541928 | 0.859712859395306 |
| Bloc1s4 | ENSMUSG00000060708 | 117197 | -0.0321331440110968 | 2.29837654377977 | -0.24669528440035 | 0.8088818275463 | 0.859712859395306 |
| Snx12 | ENSMUSG00000046032 | 55988 | -0.0202334315758037 | 3.38541282709844 | -0.246678065453339 | 0.8088948798293 | 0.859712859395306 |
| Esr1 | ENSMUSG00000019768 | 13982 | 0.0234371789421717 | 3.90994723589422 | 0.246663965964233 | 0.8089055675492 | 0.859712859395306 |
| Lyrm2 | ENSMUSG00000045854 | 108755 | 0.0331188284173241 | 2.51490755933974 | 0.245733100145773 | 0.8096112719870 | 0.860379235280047 |
| Ing2 | ENSMUSG00000063049 | 69260 | 0.0321884426757699 | 2.39141503584681 | 0.244643582900064 | 0.8104374719691 | 0.8611735205691 |
| L1cam | ENSMUSG00000031391 | 16728 | -0.0574346996557622 | 1.5882920239465 | -0.244418994558392 | 0.8106078105602 | 0.861270798720284 |
| Tmem248 | ENSMUSG00000053094 | 71667 | 0.0196102754762482 | 4.66404413571586 | 0.244194260438491 | 0.8107782697301 | 0.861368186064757 |
| Ring1 | ENSMUSG00000024325 | 19763 | 0.0278295490351126 | 2.38106312518192 | 0.244023940952398 | 0.8109074624693 | 0.86142171742009 |
| Ppp1r9a | ENSMUSG00000032827 | 243725 | -0.0386879267130609 | 2.32411561225955 | -0.243819032064003 | 0.8110628999815 | 0.86150311531419 |
| Spred2 | ENSMUSG00000045671 | 114716 | 0.0413987760233652 | 3.43561912002257 | 0.243571762167011 | 0.8112504822997 | 0.861535003611596 |
| Snx17 | ENSMUSG00000029146 | 266781 | -0.0159883463577284 | 3.92975141764855 | -0.243571669295135 | 0.8112505527558 | 0.861535003611596 |
| Ado | ENSMUSG00000057134 | 211488 | 0.0158745478580009 | 4.59063643168421 | 0.243169861163765 | 0.8115553956693 | 0.861775017491934 |
| Ccne2 | ENSMUSG00000028212 | 12448 | 0.0478167094943178 | 1.16666037267869 | 0.243060667196856 | 0.8116382442182 | 0.861779276109711 |
| Dph6 | ENSMUSG00000057147 | 66632 | 0.0394938205136255 | 1.5109105294494 | 0.24247626712135 | 0.8120816850362 | 0.86210171054254 |
| Rbm33 | ENSMUSG00000048271 | 381626 | -0.0148998527524222 | 5.32494176073109 | -0.242452588654472 | 0.8120996535958 | 0.86210171054254 |
| Odf2l | ENSMUSG00000028256 | 52184 | -0.032080644020029 | 2.3858725511112 | -0.242007033315361 | 0.8124377867479 | 0.862376912695889 |
| Trp53i11 | ENSMUSG00000068735 | 277414 | 0.0582587141697823 | 2.78625740672018 | 0.241503340156733 | 0.8128200878246 | 0.862698939703966 |
| Arhgef3 | ENSMUSG00000021895 | 71704 | -0.0228656235356191 | 3.58245808781672 | -0.241265332319292 | 0.8130007521313 | 0.862806914713414 |
| Stk10 | ENSMUSG00000020272 | 20868 | 0.0394455846782162 | 3.00039304144212 | 0.240287611162466 | 0.8137430260368 | 0.863510825901237 |
| Ppp1r12b | ENSMUSG00000073557 | 329251 | 0.019585309522931 | 9.19189780285257 | 0.239695037476893 | 0.8141929915788 | 0.863904444860028 |
| Mta2 | ENSMUSG00000071646 | 23942 | 0.0205089038525221 | 4.22664857466792 | 0.239523217029482 | 0.8143234747000 | 0.863959031539 |
| Champ1 | ENSMUSG00000047710 | 101994 | 0.0202944541668013 | 3.06783005941962 | 0.239336713805921 | 0.8144651146535 | 0.864025443349955 |
| Pelo | ENSMUSG00000042275 | 105083 | 0.031396934711275 | 2.58720151512689 | 0.238846790637181 | 0.8148372192304 | 0.864336306977948 |
| Tenm3 | ENSMUSG00000031561 | 23965 | 0.0237651153066096 | 4.4681397259563 | 0.237577989955365 | 0.8158011102612 | 0.865274785199423 |
| Pogz | ENSMUSG00000038902 | 229584 | -0.0190907901038431 | 3.97622576958308 | -0.236775322198193 | 0.8164110469857 | 0.865837698127582 |
| Lrrc32 | ENSMUSG00000090958 | 434215 | -0.0511862491711352 | 2.69437415424183 | -0.236425661828257 | 0.8166767881413 | 0.865992334924426 |
| Rpl32 | ENSMUSG00000057841 | 19951 | -0.0191780295305435 | 5.91798424974695 | -0.236374986373297 | 0.8167153033332 | 0.865992334924426 |
| Ufc1 | ENSMUSG00000062963 | 66155 | 0.0262193213775346 | 4.29238343754939 | 0.235955595140116 | 0.8170340748910 | 0.866246311603692 |
| Zdhhc21 | ENSMUSG00000028403 | 68268 | 0.0238244134789412 | 4.07452098884675 | 0.235541897399159 | 0.8173485520058 | 0.86649568635201 |
| Gas6 | ENSMUSG00000031451 | 14456 | 0.0173106303504147 | 5.80913996685914 | 0.235414574542439 | 0.8174453445479 | 0.866514261176693 |
| Ccdc137 | ENSMUSG00000049957 | 67291 | -0.0329590514359646 | 2.01372173278324 | -0.2352255680564 | 0.8175890355369 | 0.866582541205742 |
| Drap1 | ENSMUSG00000024914 | 66556 | -0.0234026876026467 | 5.25047766837374 | -0.234722597045464 | 0.8179714493672 | 0.866839323123546 |
| Zfp280d | ENSMUSG00000038535 | 235469 | -0.0241421206085777 | 3.87513251775232 | -0.234698328858857 | 0.8179899019320 | 0.866839323123546 |
| Calm2 | ENSMUSG00000036438 | 12314 | -0.0179486701744 72 | 5.8823831281777 | -0.23326813788668 | 0.8190775605885 | 0.86790779515252 |
| Mrps34 | ENSMUSG00000038880 | 79044 | 0.0209630952728137 | 4.37813353193813 | 0.23312771329793 | 0.8191843743106 | 0.867936841677764 |
| Arhgef10 | ENSMUSG00000071176 | 234094 | 0.0341718313163666 | 3.48114799025059 | 0.232892230833723 | 0.8193635020413 | 0.868042492810021 |
| Rpl30 | ENSMUSG00000058600 | 19946 | -0.0167912364956235 | 5.67668992081956 | -0.232337311014079 | 0.8197856622059 | 0.86840556968438 |
| Hebp1 | ENSMUSG00000042770 | 15199 | 0.0312581730586963 | 1.69635795848937 | 0.231913340783987 | 0.8201082405768 | 0.86864791720379 |
| Rpf2 | ENSMUSG00000038510 | 67239 | 0.0227900311132196 | 2.50152846105092 | 0.231827739963907 | 0.8201733742073 | 0.86864791720379 |
| Trim37 | ENSMUSG00000018548 | 68729 | 0.0253162911254465 | 2.38287897505398 | 0.231069920318908 | 0.8207500591611 | 0.869174471680879 |
| Zfp592 | ENSMUSG00000005621 | 233410 | 0.0190400414548542 | 4.64932776347842 | 0.230895329435161 | 0.8208829345171 | 0.869230975221014 |

| | | | | | | | |
|---|---|---|---|---|---|---|---|
| Edc3 | ENSMUSG00000038957 | 353190 | -0.0293665722615701 | 2.13364574865756 | -0.230268171314361 | 0.8213602911115 | 0.869577347386383 |
| Cds2 | ENSMUSG00000058793 | 110911 | -0.0187072858258404 | 5.22752535354813 | -0.230256541331704 | 0.8213691438811 | 0.869577347386383 |
| Spice1 | ENSMUSG00000043065 | 212514 | -0.0306030805839734 | 2.29876942838783 | -0.230027297746273 | 0.8215436498047 | 0.869586839363701 |
| Kansl1 | ENSMUSG00000018412 | 76719 | -0.0176618460911184 | 4.68469853035423 | -0.229982955772097 | 0.8215774051564 | 0.869586839363701 |
| Rasl11b | ENSMUSG00000049907 | 68939 | 0.0470023718949613 | 1.75914232525627 | 0.229927247489277 | 0.8216198136389 | 0.869586839363701 |
| Miip | ENSMUSG00000029022 | 28010 | -0.0364927357865645 | 1.89156362804851 | -0.229826750980283 | 0.8216963190730 | 0.869586839363701 |
| Usp36 | ENSMUSG00000033909 | 72344 | -0.0251574611869136 | 4.07410741174034 | -0.229484489261764 | 0.8219568883921 | 0.869778387900779 |
| Tcerg1 | ENSMUSG00000024498 | 56070 | -0.0194661823131179 | 5.09023903167548 | -0.229059128234075 | 0.8222807530727 | 0.869872579292858 |
| Rab6b | ENSMUSG00000032549 | 270192 | 0.0444085887068746 | 2.43574532989602 | 0.229052097892655 | 0.8222861061726 | 0.869872579292858 |
| Ufd1 | ENSMUSG00000005262 | 22230 | -0.0162181866052151 | 4.07355351504881 | -0.228865630503416 | 0.8224280910582 | 0.869872579292858 |
| Med23 | ENSMUSG00000019984 | 70208 | -0.0208733073225043 | 3.08387725726294 | -0.228864212113228 | 0.8224291711109 | 0.869872579292858 |
| Cep152 | ENSMUSG00000068394 | 99100 | -0.0359104234543925 | 1.47183852808881 | -0.2288450091312 | 0.8224437935222 | 0.869872579292858 |
| Kmt2c | ENSMUSG00000038056 | 231051 | 0.0176149633728295 | 6.40415121568686 | 0.228653739101587 | 0.8225894428344 | 0.86991674771709 |
| Nsmce1 | ENSMUSG00000030750 | 67711 | -0.0280104331647877 | 2.54954680478554 | -0.228581151781734 | 0.8226447187953 | 0.86991674771709 |
| Cfl2 | ENSMUSG00000062929 | 12632 | 0.0173698372739191 | 7.79087250154821 | 0.228392924461476 | 0.8227880603312 | 0.869984164004691 |
| Aamp | ENSMUSG00000006299 | 227290 | -0.016157204513628 | 5.28496259107656 | -0.22816904388472 | 0.8229585616286 | 0.870080282151393 |
| Gdi1 | ENSMUSG00000015291 | 14567 | 0.0218551673148744 | 4.25925661176515 | 0.227764685133313 | 0.8232665337674 | 0.870321709923762 |
| Nemf | ENSMUSG00000020982 | 66244 | 0.013446039524843 | 4.87596903030262 | 0.227163817049903 | 0.8237242295631 | 0.870721357059738 |
| Hist2h2ac | ENSMUSG00000068855 | 319176 | 0.0337208082160414 | 2.31031416390146 | 0.225358176677594 | 0.8251000314430 | 0.872091321185891 |
| Patj | ENSMUSG00000061859 | 12695 | 0.0205545699624978 | 2.90364163945168 | 0.224716576857052 | 0.8255890409450 | 0.872523813842238 |
| Arhgap6 | ENSMUSG00000031355 | 11856 | -0.0245132155795584 | 2.15260953059211 | -0.224323256799894 | 0.8258888558651 | 0.872756291904726 |
| Actr3b | ENSMUSG00000056367 | 242894 | -0.0479888133374495 | 1.29156395867292 | -0.223912567096135 | 0.8262019413189 | 0.873002747274765 |
| Lyrm1 | ENSMUSG00000030922 | 73919 | 0.0324234375566215 | 1.42219450831178 | 0.223172491054042 | 0.8267662091535 | 0.873481719003484 |
| Gdpd5 | ENSMUSG00000035314 | 233552 | 0.0390297037848812 | 1.57627379950422 | 0.223108434890583 | 0.8268150531999 | 0.873481719003484 |
| Sfpq | ENSMUSG00000028820 | 71514 | -0.0157333443332925 | 5.7682004277914 | -0.222466431344067 | 0.8273046343978 | 0.873854788657579 |
| Zscan25 | ENSMUSG00000070420 | 666311 | -0.0315547052331726 | 1.77483770431618 | -0.222435692996407 | 0.8273280768289 | 0.873854788657579 |
| Pdcd11 | ENSMUSG00000025047 | 18572 | -0.0177917660765718 | 4.00927585097324 | -0.222044994093268 | 0.8276260561346 | 0.874085064696443 |
| Rps18 | ENSMUSG00000008668 | 20084 | 0.0187815977200918 | 6.22338602989411 | 0.221646802934693 | 0.8279297781336 | 0.874321360716732 |
| Srsf10 | ENSMUSG00000028676 | 14105 | -0.0214138758920415 | 4.25906841252415 | -0.221187865846401 | 0.8282798699492 | 0.874606574421905 |
| Frg1 | ENSMUSG00000031590 | 14300 | -0.024934872280733 | 3.30087478613284 | -0.219385319136401 | 0.8296552774783 | 0.875974291327688 |
| Cd99l2 | ENSMUSG00000035776 | 171486 | -0.016700853333588 | 5.37496971100438 | -0.21858939534181 | 0.8302627814252 | 0.876531047301449 |
| Pigo | ENSMUSG00000028454 | 56703 | 0.0232804053779847 | 2.32017283490706 | 0.218411514715054 | 0.8303985676790 | 0.876589738609296 |
| Zfyve27 | ENSMUSG00000018820 | 319740 | 0.0285076022916906 | 3.20773498657177 | 0.217938060357228 | 0.8307600094464 | 0.876886603250228 |
| Tcta | ENSMUSG00000039461 | 102791 | -0.0158964697465365 | 4.31950039374421 | -0.217795010399719 | 0.8308692236717 | 0.876917204205423 |
| Mfap1b | ENSMUSG00000048222 | 100034361 | -0.0155432704308112 | 5.43124857798783 | -0.217017151938466 | 0.8314631583030 | 0.877459334177517 |
| Ermard | ENSMUSG00000036552 | 381062 | 0.0306740194796822 | 2.49037352278458 | 0.216657124930849 | 0.8317380936680 | 0.877664745813798 |
| Pagr1a | ENSMUSG00000107068 | 67278 | -0.128997303784286 | 2.57200651344178 | -0.216224984915289 | 0.8320681286034 | 0.877928254224381 |
| Itpk1 | ENSMUSG00000057963 | 217837 | -0.0260789463359514 | 1.99639004960163 | -0.215899639887851 | 0.8323166236275 | 0.878105686021837 |
| Mtss1 | ENSMUSG00000022353 | 211401 | -0.0183672756891653 | 4.2513365183913 | -0.215599816838639 | 0.8325456417909 | 0.878262537195203 |
| Camk2n1 | ENSMUSG00000046447 | 66259 | -0.0314181681700951 | 2.50524822105105 | -0.214553486176611 | 0.8333449997012 | 0.878951655936142 |
| Prnd | ENSMUSG00000027338 | 26434 | 0.111017725936403 | 0.592810487065287 | 0.214534236626164 | 0.8333597074487 | 0.878951655936142 |
| Cryab | ENSMUSG00000032060 | 12955 | -0.021431547265772 | 9.70529408181861 | -0.214291176246579 | 0.8335454249483 | 0.879062714916553 |
| Nol8 | ENSMUSG00000021392 | 70930 | 0.0164307719559549 | 4.23879689699859 | 0.213959728054391 | 0.8337986944737 | 0.879188567108202 |
| Steap3 | ENSMUSG00000026389 | 68428 | -0.0263234142094251 | 3.56842212920002 | -0.213924489945862 | 0.8338256221032 | 0.879188567108202 |
| Tgfbrap1 | ENSMUSG00000070939 | 73122 | 0.0145663538561447 | 4.27290127470177 | 0.213667005548392 | 0.8340223885352 | 0.879311220011489 |
| Masp1 | ENSMUSG00000022887 | 17174 | -0.0344096801692459 | 2.98873011876767 | -0.213252026560939 | 0.8343395348036 | 0.87956075368297 |
| Heyl | ENSMUSG00000032744 | 56198 | 0.0314582255314146 | 3.70393740019665 | 0.212993448742919 | 0.8345371672185 | 0.879600205347107 |
| Haus6 | ENSMUSG00000038047 | 230376 | -0.0361060617743338 | 2.42058435054806 | -0.212992496628847 | 0.8345378949460 | 0.879600205347107 |
| Exosc9 | ENSMUSG00000027714 | 50911 | -0.0195541118964924 | 3.91066076845016 | -0.212536610807304 | 0.8348863593892 | 0.879882645052437 |
| Leng8 | ENSMUSG00000035545 | 232798 | -0.0359101602707419 | 4.86579584100454 | -0.212417103270021 | 0.8349777130881 | 0.879894088669324 |
| Slc35a3 | ENSMUSG00000027957 | 229782 | -0.0211707717247518 | 3.02830693307733 | -0.210961230350592 | 0.8360908074222 | 0.880982129933762 |
| Cnnm2 | ENSMUSG00000064105 | 94219 | 0.030736250686235 | 2.37167324960483 | 0.209899710667869 | 0.8369026281670 | 0.881752542505442 |
| Per2 | ENSMUSG00000055866 | 18627 | 0.0831901148785756 | 2.16854379048418 | 0.209716644368304 | 0.8370426517992 | 0.881815075830529 |
| Cfap20 | ENSMUSG00000031796 | 14894 | 0.0285237417814002 | 2.52158520715265 | 0.208814031824731 | 0.8377331255881 | 0.882457434307001 |
| Pcid2 | ENSMUSG00000038542 | 234069 | 0.0322998320339325 | 2.4423427597666 | 0.208440626727368 | 0.8380188110232 | 0.88267331116737 |
| Hmbox1 | ENSMUSG00000021972 | 219150 | -0.0155067439701266 | 4.02245143609208 | -0.208249975141128 | 0.8381646842650 | 0.882699511539151 |
| Imp3 | ENSMUSG00000032288 | 102462 | 0.0269033038003967 | 2.73562377895904 | 0.208197037153291 | 0.8382051898066 | 0.882699511539151 |
| Tmem245 | ENSMUSG00000055296 | 242474 | 0.016936183261981 | 5.19313012908795 | 0.207631803497318 | 0.8386377085551 | 0.883069915443226 |
| Runx1t1 | ENSMUSG00000006586 | 12395 | 0.0441017514780588 | 2.02177633938046 | 0.207374890233768 | 0.8388343174558 | 0.883165226515591 |
| Tnfsfm13 | ENSMUSG00000018752 | 619441 | -0.0568703967872748 | 0.942079519785226 | -0.207302375494134 | 0.8388898130922 | 0.883165226515591 |
| Lin54 | ENSMUSG00000035310 | 231506 | 0.0261091327415097 | 2.63883040422124 | 0.207110403344808 | 0.8390367339725 | 0.883234836195516 |
| Ccdc51 | ENSMUSG00000025645 | 66658 | -0.0292696812761697 | 1.68434856295561 | -0.206654074234413 | 0.8393859986673 | 0.883517414678178 |
| Gm10705 | ENSMUSG00000074506 | 100042355 | -0.0271062768718719 | 2.46100585997542 | -0.206440708517529 | 0.8395493164053 | 0.883604234318009 |
| Ldhd | ENSMUSG00000031958 | 52815 | -0.022669538882929 | 4.23012200098112 | -0.206093891419395 | 0.8398147990351 | 0.883798552830802 |
| Il34 | ENSMUSG00000031750 | 76527 | 0.0614635245646258 | 0.95213346538045 | 0.205665828352619 | 0.8401425021067 | 0.884058306750906 |
| Tdp1 | ENSMUSG00000021177 | 104884 | -0.0367732259537774 | 1.29437125510432 | -0.205401331836285 | 0.8403450024905 | 0.884186276082791 |
| Bcap31 | ENSMUSG00000002015 | 27061 | -0.0162352961060672 | 3.75260692353276 | -0.203471536998744 | 0.8418228226886 | 0.885655945602471 |
| Ap1s3 | ENSMUSG00000054702 | 252903 | -0.0558667409358482 | 1.29002410919514 | -0.202803927792667 | 0.8423342170883 | 0.886068857277366 |
| Akap9 | ENSMUSG00000040407 | 100986 | -0.0176491956586595 | 7.06717545199017 | -0.202662155190547 | 0.8424428256793 | 0.886068857277366 |
| Tia1 | ENSMUSG00000071337 | 21841 | 0.0237765847655251 | 3.83552721442889 | 0.202641722465483 | 0.8424584789757 | 0.886068857277366 |
| Adar | ENSMUSG00000027951 | 56417 | 0.0296935376878644 | 3.15511752085687 | 0.202257179762482 | 0.8427530860508 | 0.886293436946476 |
| Ncoa5 | ENSMUSG00000039804 | 228869 | -0.025417434451712 | 3.84572373000109 | -0.201666142409989 | 0.8432059413083 | 0.88668438137963 |
| Mars2 | ENSMUSG00000046994 | 212679 | -0.0248305760836995 | 2.96651962767648 | -0.201188121644945 | 0.8435722450992 | 0.886984245015343 |
| Ppp1r26 | ENSMUSG00000035829 | 241289 | -0.0376787521624053 | 1.66723369398464 | -0.200946542148355 | 0.8437573800506 | 0.887052820064408 |
| Abca8b | ENSMUSG00000020620 | 27404 | 0.019806606224722 | 4.88949170598199 | 0.200891237483156 | 0.8437997642511 | 0.887052820064408 |
| Xndc1 | ENSMUSG00000099481 | 102443350 | -0.0536156423575544 | 1.05441107237249 | -0.200224679183 | 0.8443106385709 | 0.887504528341109 |
| Nlrc5 | ENSMUSG00000074151 | 434341 | 0.0413202480483932 | 2.00270155492825 | 0.199752006237649 | 0.8446729564469 | 0.887763187550682 |
| Mcl1 | ENSMUSG00000038612 | 17210 | 0.014171987668341 | 7.07519611236718 | 0.199691760170431 | 0.8447191394853 | 0.887763187550682 |
| Usp8 | ENSMUSG00000027363 | 84092 | -0.0127024408659162 | 4.85127182646086 | -0.199157318612949 | 0.8451288542626 | 0.888108393188301 |
| E2f4 | ENSMUSG00000014859 | 104394 | -0.0132470506462585 | 4.53486768185846 | -0.197354126258709 | 0.8465115656949 | 0.889475913160735 |
| Ears2 | ENSMUSG00000030871 | 67417 | -0.0269773663878143 | 1.81030250459301 | -0.196895462461456 | 0.8468633593899 | 0.889760032823073 |
| Ciao2b | ENSMUSG00000031879 | 68523 | -0.0210936490232765 | 3.67043107490451 | -0.19669897186096 | 0.8470140774850 | 0.889772404146852 |
| Snhg8 | ENSMUSG00000104960 | 69895 | 0.0320295181636971 | 1.52929343633283 | 0.196638996543817 | 0.8470600827895 | 0.889772404146852 |
| Zfp738 | ENSMUSG00000048280 | 408068 | -0.0226645393724847 | 3.15176984784743 | -0.196561757751133 | 0.8471193312557 | 0.889772404146852 |
| Utp20 | ENSMUSG00000004356 | 70683 | -0.0159466956638967 | 4.34809061858514 | -0.196448255084881 | 0.8472063988252 | 0.889778357567125 |
| Selenoi | ENSMUSG00000075703 | 28042 | -0.0209350740377703 | 2.8082965835588 | -0.195557808027898 | 0.8478895302921 | 0.890343608696363 |
| Ppp4r3a | ENSMUSG00000041846 | 68734 | -0.0116597975052786 | 4.81829912535827 | -0.195436978000126 | 0.8479822382764 | 0.890343608696363 |
| Chuk | ENSMUSG00000025199 | 12675 | -0.0177677042559452 | 3.69255610434665 | -0.195428218813331 | 0.8479889589367 | 0.890343608696363 |
| Acot8 | ENSMUSG00000017307 | 170789 | 0.0422665330600911 | 1.10966866742251 | 0.195027489767846 | 0.8482964394431 | 0.89050532518822 |
| Rplp2 | ENSMUSG00000025508 | 67186 | -0.0145176979296311 | 7.37778017052982 | -0.19496933510667 | 0.8483410638166 | 0.89050532518822 |
| Bcl2 | ENSMUSG00000057329 | 12043 | -0.037052396355997 | 1.5235853833622 | -0.194908975736268 | 0.8483873805242 | 0.89050532518822 |
| Rabggtb | ENSMUSG00000038975 | 19352 | -0.0148580246698913 | 4.65416163364149 | -0.194634668457606 | 0.8485978773168 | 0.890640748627006 |
| H2afz | ENSMUSG00000037894 | 51788 | 0.0260128117088541 | 3.41538116447105 | 0.192523604359888 | 0.8502182587878 | 0.892217531866268 |
| Rab33b | ENSMUSG00000027739 | 19338 | -0.0210863686743861 | 3.34529211380491 | -0.192464710921144 | 0.8502634735569 | 0.892217531866268 |
| Fam3a | ENSMUSG00000031399 | 66294 | 0.0200563893956134 | 2.86210939548365 | 0.192260460377578 | 0.8504202888478 | 0.892296427087307 |
| Zfp930 | ENSMUSG00000059897 | 234358 | 0.0313437103019454 | 2.02179945955941 | 0.191540400244432 | 0.8509731744373 | 0.892790840750087 |
| Lypla2 | ENSMUSG00000028670 | 26394 | 0.0225389686033086 | 3.03204400100324 | 0.191103554526403 | 0.8513086386369 | 0.893057075714026 |
| Twsg1 | ENSMUSG00000024098 | 65960 | 0.0207251241373827 | 5.00362649206831 | 0.190886046471493 | 0.8514756794335 | 0.893146593598316 |
| Flot2 | ENSMUSG00000061981 | 14252 | 0.0129314310474697 | 5.63521558007102 | 0.190481263564109 | 0.8517865624574 | 0.893386961641033 |
| Mapkbp1 | ENSMUSG00000033902 | 26390 | -0.0180760016517523 | 3.08142973973711 | -0.189968106170087 | 0.8521807166189 | 0.893714613197915 |
| P4ha1 | ENSMUSG00000019916 | 18451 | -0.0244858923911959 | 4.62894853115458 | -0.189701830456532 | 0.8523852581749 | 0.893843366952231 |
| Trmt12 | ENSMUSG00000037085 | 68260 | 0.0267104706539552 | 1.58212945114969 | 0.189280957753406 | 0.8527085770695 | 0.894096638460184 |
| Mre11a | ENSMUSG00000031928 | 17535 | -0.0278666965691432 | 2.87750940253131 | -0.188492112629396 | 0.8533146506123 | 0.894613277288427 |
| Zbtb8os | ENSMUSG00000057572 | 67106 | 0.0261962108933429 | 2.686538527687 | 0.188343462734189 | 0.8534288698769 | 0.894613277288427 |
| Tyw1 | ENSMUSG00000056310 | 100929 | -0.020751629133609 | 2.76050831248999 | -0.188236760968598 | 0.8535108592473 | 0.894613277288427 |
| 4933431E20Rik | ENSMUSG00000086968 | 329735 | -0.0486752331794487 | 1.20356263396258 | -0.188213584915059 | 0.8535286679023 | 0.894613277288427 |
| Ddx49 | ENSMUSG00000057788 | 234374 | -0.0171405186918707 | 2.94894230506068 | -0.187678712432321 | 0.8539396910819 | 0.894874660858956 |
| Mad2l2 | ENSMUSG00000029003 | 71890 | -0.0297662759003099 | 1.32166388873679 | -0.187567816641787 | 0.8540249145784 | 0.894874660858956 |
| Gng11 | ENSMUSG00000032766 | 66066 | -0.0273975316294912 | 3.12886815166542 | -0.187436221733947 | 0.8541260478266 | 0.894874660858956 |
| Tbce | ENSMUSG00000039233 | 70430 | 0.0153784370704115 | 3.14733789624767 | 0.187358939780354 | 0.8541854417585 | 0.894874660858956 |
| Gpr146 | ENSMUSG00000044197 | 80290 | 0.0127026887260193 | 4.08780397752926 | 0.187356422494708 | 0.8541873763976 | 0.894874660858956 |
| Myo5b | ENSMUSG00000025885 | 17919 | 0.0269601598926812 | 2.76974635354746 | 0.186684787086363 | 0.8547035911485 | 0.895329655471517 |
| Ik | ENSMUSG00000024474 | 24010 | 0.012711669084271 | 5.74909969302634 | 0.186513870858072 | 0.8548349673702 | 0.895381470709481 |
| Sgcg | ENSMUSG00000035296 | 24053 | -0.0185655149198105 | 6.31949092382274 | -0.186018983540302 | 0.8552153920445 | 0.895676200189739 |
| Cep350 | ENSMUSG00000033671 | 74081 | -0.0159460597493855 | 6.44049663417845 | -0.185934643904357 | 0.8552802284860 | 0.895676200189739 |
| Setd6 | ENSMUSG00000031671 | 66083 | -0.0291820938828828 | 2.08439372904835 | -0.185606960988836 | 0.8555321464111 | 0.895839795456586 |

| | | | | | | | |
|---|---|---|---|---|---|---|---|
| Gtf2f2 | ENSMUSG00000067995 | 68705 | -0.0240270050026119 | 2.80184994848355 | -0.185518244057173 | 0.8556003535419 | 0.895839795456586 |
| 1300002E11Rik | ENSMUSG00000097290 | 100043489 | 0.0307016564975453 | 1.69125015925426 | 0.185228244642065 | 0.8558233185632 | 0.89598742411789 |
| Rnaseh2a | ENSMUSG00000052926 | 69724 | -0.0217376823979953 | 2.1007326131982 | -0.184511019640065 | 0.8563748095581 | 0.896478935384036 |
| Mga | ENSMUSG00000033943 | 29808 | 0.0199207891538755 | 6.05465953326328 | 0.182622630193518 | 0.8578272095465 | 0.897913360868856 |
| Prr5 | ENSMUSG00000036106 | 109270 | -0.0372641663367607 | 0.995980864719047 | -0.182228837010567 | 0.8581301520005 | 0.89814445533493 |
| Plk2 | ENSMUSG00000021701 | 20620 | -0.0238299021388704 | 3.58832581289976 | -0.181934694802064 | 0.8583564488355 | 0.898244142512838 |
| Prkd1 | ENSMUSG00000002688 | 18760 | 0.0254366316282875 | 2.25470032990707 | 0.181891416439081 | 0.8583897459234 | 0.898244142512838 |
| Ctnna1 | ENSMUSG00000037815 | 12385 | -0.0136082430694353 | 5.85950375296126 | -0.181226444816242 | 0.8589013906386 | 0.898693510201093 |
| Klhl20 | ENSMUSG00000026705 | 226541 | -0.0189218025308282 | 3.67682071856119 | -0.180478016907444 | 0.8594773276249 | 0.899164866121507 |
| Rpl31 | ENSMUSG00000073702 | 114641 | 0.0145156994083937 | 5.68144676902588 | 0.180427256218643 | 0.8595163924712 | 0.899164866121507 |
| Hsdl1 | ENSMUSG00000034189 | 72552 | -0.0217598857140984 | 3.16217330997871 | -0.18021457329095 | 0.8596800749921 | 0.899250038252584 |
| Prelid3b | ENSMUSG00000016257 | 66390 | 0.0165915225996631 | 4.68278012028078 | 0.179717444373809 | 0.8600626956929 | 0.899485357097161 |
| Gramd3 | ENSMUSG00000001700 | 107022 | -0.018188574835512 | 3.02446961844849 | -0.179708455832358 | 0.8600696141596 | 0.899485357097161 |
| Dnmbp | ENSMUSG00000025195 | 71972 | 0.0172000830728516 | 3.75845958810032 | 0.179223099987876 | 0.8604432095897 | 0.899789986035195 |
| Fam160b2 | ENSMUSG00000022095 | 239170 | 0.0150400610579005 | 3.87580774955497 | 0.177463285276658 | 0.8617980922583 | 0.901120618564749 |
| Sbf2 | ENSMUSG00000038371 | 319934 | 0.0173526465083391 | 5.0044379945609 | 0.177216867675388 | 0.8619878456668 | 0.901232820754155 |
| Commd4 | ENSMUSG00000032299 | 66199 | 0.0248129132344506 | 2.33352312426714 | 0.176936551769738 | 0.8622037132012 | 0.901372301932182 |
| Gtf2b | ENSMUSG00000028271 | 229906 | 0.0185041086339455 | 2.97517720839467 | 0.176486750967806 | 0.8625501227703 | 0.901569532476421 |
| Sptlc1 | ENSMUSG00000021468 | 268656 | 0.0184993982844097 | 2.55745827694417 | 0.176402566500979 | 0.8626149598694 | 0.901569532476421 |
| Ubqln1 | ENSMUSG00000005312 | 56085 | 0.011942372004051 | 5.78921252617021 | 0.176289867120273 | 0.8627017601690 | 0.901569532476421 |
| Paf1 | ENSMUSG00000003437 | 54624 | -0.0124272721912844 | 4.75558053652963 | -0.176263217064954 | 0.8627222861314 | 0.901569532476421 |
| Rpl7l1 | ENSMUSG00000063888 | 66229 | -0.0109490793332814 | 4.34075874890038 | -0.174955853219448 | 0.8637293487980 | 0.902446292721846 |
| Rad23b | ENSMUSG00000028426 | 19359 | 0.0107468890097921 | 6.4551198100789 | 0.174898873172939 | 0.8637732461708 | 0.902446292721846 |
| Dip2a | ENSMUSG00000020231 | 64451 | -0.0296171701620334 | 2.80218110186056 | -0.174852537543803 | 0.8638089434405 | 0.902446292721846 |
| Zfp292 | ENSMUSG00000039967 | 30046 | 0.0151990907895421 | 4.89920518769019 | 0.17426156408015 | 0.8642642604071 | 0.902776300814351 |
| Coasy | ENSMUSG00000001755 | 71743 | -0.0207441201790992 | 2.72563212316478 | -0.174208554444458 | 0.8643051042716 | 0.902776300814351 |
| Gnb5 | ENSMUSG00000032192 | 14697 | 0.017740982228498 | 2.4062467611557 | 0.174120970552241 | 0.8643725884478 | 0.902776300814351 |
| Psmg2 | ENSMUSG00000024537 | 107047 | -0.0159056290129014 | 2.6341086301713 | -0.17268176580245 | 0.8654816662906 | 0.903742960242573 |
| Zfp235 | ENSMUSG00000047603 | 56525 | -0.0275113191467848 | 1.81681922391863 | -0.172631467239546 | 0.8655204326544 | 0.903742960242573 |
| Cidea | ENSMUSG00000024526 | 12683 | -0.0302632461866141 | 1.32788007853249 | -0.172598089664329 | 0.8655461577878 | 0.903742960242573 |
| Ostm1 | ENSMUSG00000038280 | 14628 | -0.0168751422147131 | 3.65507979402106 | -0.172320563408307 | 0.8657600620401 | 0.903879965440405 |
| Clk2 | ENSMUSG00000068917 | 12748 | -0.0186399695718507 | 3.5142822429129 | -0.171720066355291 | 0.8662229345915 | 0.904276850159468 |
| Pstpip2 | ENSMUSG00000025429 | 19201 | 0.0101957609289753 | 5.48304927064433 | 0.171459385542226 | 0.8664238873945 | 0.904400259082277 |
| Meis1 | ENSMUSG00000020160 | 17268 | -0.0268208757275525 | 2.13682223176497 | -0.171230347469543 | 0.8666004554904 | 0.904489089475149 |
| Slc48a1 | ENSMUSG00000081534 | 67739 | 0.0152194104439811 | 4.63913128161179 | 0.171134330385391 | 0.8666744783791 | 0.904489089475149 |
| Rad54l2 | ENSMUSG00000040661 | 81000 | -0.0146041562868158 | 3.93286617842404 | -0.170622344025088 | 0.8670692083610 | 0.904814655522204 |
| Mbd1 | ENSMUSG00000024561 | 17190 | -0.012367871303923 | 3.90997428668299 | -0.170312130538771 | 0.8673083939860 | 0.904977859360569 |
| Bmp4 | ENSMUSG00000021835 | 12159 | -0.0382372907087838 | 1.64161367000048 | -0.169859026859305 | 0.8676577773733 | 0.905256005007938 |
| Vangl1 | ENSMUSG00000027860 | 229658 | 0.0213621842014047 | 2.05199526805209 | 0.169139510032802 | 0.8682126483015 | 0.905748469038428 |
| Btf3l4 | ENSMUSG00000028568 | 70533 | 0.0150460928710974 | 4.18969226131243 | 0.168953848600479 | 0.8683558368873 | 0.905751065665007 |
| Arhgap21 | ENSMUSG00000036591 | 71435 | 0.0134434715234466 | 5.19980273723364 | 0.168921405211547 | 0.8683808588573 | 0.905751065665007 |
| Gabarap | ENSMUSG00000018567 | 56486 | -0.0122180719707288 | 5.75977111181782 | -0.168151893195047 | 0.8689743881947 | 0.906283659703903 |
| Ccdc122 | ENSMUSG00000034795 | 108811 | 0.030489148202657 | 1.32537886761826 | 0.167717722449574 | 0.8693093031206 | 0.906546459875209 |
| Actl6a | ENSMUSG00000027671 | 56456 | 0.0138293958476581 | 3.33639000929418 | 0.167508633990428 | 0.8694706011795 | 0.906573450591227 |
| Tnks1bp1 | ENSMUSG00000033955 | 228140 | 0.0211287050585947 | 4.89481505366709 | 0.167469155181155 | 0.8695010571766 | 0.906573450591227 |
| Yju2 | ENSMUSG00000003208 | 72886 | 0.0227366068656831 | 1.84508914928786 | 0.167268078779138 | 0.8696561812717 | 0.906643271657278 |
| Cdk5rap2 | ENSMUSG00000039298 | 214444 | 0.0122577132619948 | 3.98023120969612 | 0.167167330049505 | 0.8697339078399 | 0.906643271657278 |
| Htatsf1 | ENSMUSG00000067873 | 72459 | -0.0156996688313793 | 6.88952723859731 | -0.166931933463842 | 0.8699155192602 | 0.906746118149477 |
| Hnrnpm | ENSMUSG00000059208 | 76936 | 0.00899627933758218 | 5.97740065185825 | 0.166542774501551 | 0.8702157770633 | 0.906972602887064 |
| Ska2 | ENSMUSG00000020492 | 66140 | 0.0270850346855724 | 1.40584481964577 | 0.166195116133809 | 0.8704840324987 | 0.907165693511665 |
| Snx14 | ENSMUSG00000032422 | 244962 | 0.0145147488873962 | 3.40423989943248 | 0.16572388026057 | 0.8708476676617 | 0.907382944334049 |
| Dnttip1 | ENSMUSG00000017299 | 76233 | 0.0159953266181958 | 3.18001805715355 | 0.16570981884493 | 0.8708585188005 | 0.907382944334049 |
| Ddx42 | ENSMUSG00000020705 | 72047 | -0.0112481559952498 | 4.80324125037518 | -0.165338029530327 | 0.8711454369733 | 0.907595384250415 |
| Irf2bp2 | ENSMUSG00000051495 | 270110 | 0.0227552689438957 | 5.33880359816723 | 0.164886294682299 | 0.8714940763607 | 0.907872081263632 |
| Rufy2 | ENSMUSG00000020070 | 70432 | 0.0254280728098477 | 2.03093833692238 | 0.164290169699764 | 0.8719541956093 | 0.908264847742141 |
| Zc3h10 | ENSMUSG00000039810 | 103284 | 0.0262156064548495 | 1.91666922226291 | 0.163930788922465 | 0.8722316071328 | 0.908467241311984 |
| Gars | ENSMUSG00000029777 | 353172 | 0.0106020268310569 | 5.61840586246282 | 0.163297001909009 | 0.8727208797203 | 0.908890237826152 |
| Uty | ENSMUSG00000068457 | 22290 | 0.013926081017829 | 3.85293584312963 | 0.162380937456971 | 0.8734281615233 | 0.909540176584938 |
| Reep5 | ENSMUSG00000005873 | 13476 | -0.0107936919829585 | 5.43647142286689 | -0.162054963170022 | 0.8736798694104 | 0.909668648040278 |
| Rtca | ENSMUSG00000000339 | 66368 | -0.0167260645425338 | 3.48602369168681 | -0.162005621937719 | 0.8737179705218 | 0.909668648040278 |
| Bhlhb9 | ENSMUSG00000072964 | 70237 | 0.025353520561141 | 2.02654557067192 | 0.161661263019073 | 0.8739838922555 | 0.909858850118575 |
| Crip2 | ENSMUSG00000006356 | 68337 | 0.0253214891172258 | 4.38384329020837 | 0.160497843612177 | 0.8748824282402 | 0.910707534815214 |
| 1110012L19Rik | ENSMUSG00000045237 | 68618 | 0.0270864390919445 | 1.5493226428677 | 0.160148701479562 | 0.8751521140309 | 0.910901519565032 |
| Zfp790 | ENSMUSG00000011427 | 233056 | 0.0164110482049407 | 3.147106897938 | 0.159753676534575 | 0.8754572603106 | 0.911132372889251 |
| Iws1 | ENSMUSG00000024384 | 73473 | 0.00943006324456697 | 5.08523381793006 | 0.158129342305947 | 0.8767122319115 | 0.912281319473978 |
| Il1rap | ENSMUSG00000022514 | 16180 | 0.0219591001158457 | 1.35640932452141 | 0.15810875945028 | 0.8767281365908 | 0.912281319473978 |
| Crppa | ENSMUSG00000043153 | 75847 | -0.0243584200216962 | 1.61894886703332 | -0.157695605164965 | 0.8770473987956 | 0.912526662501006 |
| Acvr1 | ENSMUSG00000026836 | 11477 | 0.0146307564869224 | 3.9572089496179 | 0.15713209414006 | 0.8774828841213 | 0.912892872021598 |
| Ercc3 | ENSMUSG00000024382 | 13872 | 0.0133184964510202 | 3.42240773274987 | 0.156973437849487 | 0.877605502299 | 0.912933550212255 |
| Psmd6 | ENSMUSG00000021737 | 66413 | 0.0106275792929491 | 6.0112486580737 | 0.156234731288595 | 0.8781764578329 | 0.913440561478004 |
| Sltm | ENSMUSG00000032212 | 66660 | 0.00855524619382475 | 5.00356164308136 | 0.156110483821278 | 0.8782724972067 | 0.913453536343148 |
| Tbc1d22a | ENSMUSG00000051864 | 223754 | 0.014146101909482 | 3.54586505514542 | 0.155836027591194 | 0.8784846502964 | 0.913587262143525 |
| Nek1 | ENSMUSG00000031644 | 18004 | -0.0193954651149202 | 3.52651268773491 | -0.155663056362942 | 0.8786183610637 | 0.913639393530071 |
| Zdhhc1 | ENSMUSG00000039199 | 70796 | 0.0187727739704573 | 3.14276096287719 | 0.15550438209037 | 0.8787410233494 | 0.913659180143233 |
| Tarbp2 | ENSMUSG00000023051 | 21357 | -0.0187132010346254 | 3.01545206690372 | -0.155422196588385 | 0.8788045576823 | 0.913659180143233 |
| Rabggta | ENSMUSG00000040472 | 56187 | -0.0222468115605103 | 2.20426897662579 | -0.154640097840724 | 0.8794092105198 | 0.914200863546641 |
| Mmp12 | ENSMUSG00000049723 | 17381 | 0.0713308991423725 | 1.47663724385857 | 0.154333454058474 | 0.8796463030791 | 0.914299645342696 |
| Adrb2 | ENSMUSG00000045730 | 11555 | 0.025767061121396 | 3.27734756162979 | 0.154300842438233 | 0.8796715186231 | 0.914299645342696 |
| Cdc25a | ENSMUSG00000032477 | 12530 | 0.032126310223243 | 1.33967863456189 | 0.153776132024659 | 0.8800772472565 | 0.914601907457799 |
| Exosc5 | ENSMUSG00000061286 | 27998 | -0.0219876758083999 | 1.90846178930775 | -0.153657640474247 | 0.8801688748787 | 0.914601907457799 |
| Alg1 | ENSMUSG00000039427 | 208211 | -0.0197645045076928 | 2.14382684770517 | -0.153600134089588 | 0.8802133442920 | 0.914601907457799 |
| Tspan18 | ENSMUSG00000027217 | 241556 | 0.0256822153422744 | 2.26227844035451 | 0.153172900059701 | 0.8805437355153 | 0.914858242839857 |
| Fam241a | ENSMUSG00000050549 | 70617 | -0.0210079258892688 | 2.11661297380171 | -0.152797158525127 | 0.8808343254199 | 0.91507318106492 |
| Plscr4 | ENSMUSG00000032377 | 235527 | 0.024266167659746 | 2.25563950082783 | 0.152024285130141 | 0.8814321042385 | 0.915464124181156 |
| Psmg4 | ENSMUSG00000071451 | 69666 | 0.0165005631562622 | 2.09236277966231 | 0.151862735440741 | 0.8815570643566 | 0.915464124181156 |
| Nlrx1 | ENSMUSG00000032109 | 270151 | -0.0211532420426805 | 3.16733150646079 | -0.151825928487072 | 0.8815855353220 | 0.915464124181156 |
| Eif5 | ENSMUSG00000021282 | 217869 | -0.0146552501944317 | 6.56505032417089 | -0.151778670440462 | 0.8816220906814 | 0.915464124181156 |
| Scaf8 | ENSMUSG00000046201 | 106583 | -0.00995675179414324 | 4.72742298545773 | -0.151769237242852 | 0.8816293875450 | 0.915464124181156 |
| Zmym6 | ENSMUSG00000042408 | 100177 | 0.0228012921001084 | 2.45740878470213 | 0.151412997804802 | 0.8819049577197 | 0.91556156634869 |
| Zfp623 | ENSMUSG00000050846 | 78834 | -0.0189004921580409 | 2.60808996456831 | -0.151333312532387 | 0.8819666007048 | 0.91556156634869 |
| Fam168b | ENSMUSG00000037503 | 214469 | 0.0112656549454639 | 4.59030042273308 | 0.15132309722805 | 0.8819745031242 | 0.91556156634869 |
| Rbm24 | ENSMUSG00000038132 | 666794 | 0.0102104759706325 | 6.63954255920519 | 0.151132335711083 | 0.8821220760265 | 0.915627804866177 |
| Cdadc1 | ENSMUSG00000021982 | 71891 | 0.0114433849772659 | 4.28268509840267 | 0.150866583719769 | 0.8823276690852 | 0.915733288434078 |
| Errfi1 | ENSMUSG00000028967 | 74155 | 0.0329719983008676 | 4.40235065691874 | 0.150784403755066 | 0.8823912475597 | 0.915733288434078 |
| Stk25 | ENSMUSG00000026277 | 59041 | -0.00920262952618778 | 4.91888841441673 | -0.150390374159447 | 0.8826961000253 | 0.915962698820718 |
| Jak3 | ENSMUSG00000031805 | 16453 | 0.0353852424984793 | 2.31236574511462 | 0.150049721468878 | 0.8829596715168 | 0.916149232971163 |
| Zfp407 | ENSMUSG00000048410 | 240476 | -0.0177841795645892 | 3.85620745474009 | -0.149714892155379 | 0.8832187514014 | 0.916331071713087 |
| Rpl37a | ENSMUSG00000046330 | 19981 | -0.0113973417139333 | 5.5796304650159 | -0.148680193606279 | 0.8840194549513 | 0.917074752023686 |
| Zbtb37 | ENSMUSG00000043467 | 240869 | 0.0173598855370192 | 3.82079447913447 | 0.148092653212998 | 0.8844741831521 | 0.917459413174842 |
| Ogfod2 | ENSMUSG00000023707 | 66627 | -0.0186174936651764 | 1.70218892453558 | -0.147558081509227 | 0.8848879530038 | 0.917776349985071 |
| Hspa14 | ENSMUSG00000109865 | 50497 | -0.0134729111004184 | 2.99229835453897 | -0.147358564778985 | 0.8850423921706 | 0.917776349985071 |
| Zfp846 | ENSMUSG00000058192 | 244721 | -0.0329536543372595 | 1.5011571094905 | -0.147329243557808 | 0.8850650891490 | 0.917776349985071 |
| Rasal2 | ENSMUSG00000070565 | 226525 | -0.018678118862189 | 4.31692541659228 | -0.147264031738877 | 0.8851155687075 | 0.917776349985071 |
| Ptar1 | ENSMUSG00000074925 | 72351 | -0.0129543106317183 | 4.96387723315769 | -0.147089630007138 | 0.8852505732080 | 0.917829272098739 |
| Ap1m1 | ENSMUSG00000003033 | 11767 | 0.0132749757404937 | 4.1506148622164 | 0.146380641444687 | 0.8857994399520 | 0.9182890969284 |
| Mettl15 | ENSMUSG00000057234 | 76894 | -0.0212245790198713 | 1.93934823317234 | -0.146299716900319 | 0.8858620919504 | 0.9182890969284 |
| Ice2 | ENSMUSG00000032235 | 93697 | 0.021275819496726 | 1.98725500857987 | 0.145859571199465 | 0.8862028678516 | 0.918485314874978 |
| Zmat1 | ENSMUSG00000052676 | 215693 | 0.029426874866227 | 1.20045628304856 | 0.145838177174303 | 0.8862194324386 | 0.918485314874978 |
| Zfp273 | ENSMUSG00000030446 | 212569 | 0.05159565603978 | 0.53847301431394 | 0.145052789694914 | 0.8868275666664 | 0.918959138117485 |
| Abl2 | ENSMUSG00000026596 | 11352 | -0.00821187335089135 | 5.11205737323023 | -0.145030601004955 | 0.8868447486964 | 0.918959138117485 |
| Znrd1 | ENSMUSG00000036315 | 66136 | -0.0257726702183747 | 1.72686669837152 | -0.144844015724038 | 0.8869892351880 | 0.918994621941891 |
| Gpr89 | ENSMUSG00000028096 | 67549 | 0.0134325052496109 | 3.21718442853326 | 0.144769244491585 | 0.8870471371428 | 0.918994621941891 |
| Sertad2 | ENSMUSG00000049800 | 58172 | 0.0119697666767542 | 3.84761244191029 | 0.14443927388742 | 0.8873026704892 | 0.919172241387207 |
| Tut7 | ENSMUSG00000035248 | 214290 | 0.00918718097812743 | 5.91149511299003 | 0.1437457632621 | 0.8878397763152 | 0.919417134234707 |
| Naa80 | ENSMUSG00000079334 | 56441 | -0.0154895522288827 | 3.01692710556671 | -0.143733502335414 | 0.8878492725997 | 0.919417134234707 |
| Arhgef11 | ENSMUSG00000041977 | 213498 | -0.0111537041530338 | 4.85934222619129 | -0.143700160772902 | 0.8878750962655 | 0.919417134234707 |

| | | | | | | | |
|---|---|---|---|---|---|---|---|
| Sel1l3 | ENSMUSG00000029189 | 231238 | 0.0139425052602888 | 5.36746812528763 | 0.143699618421488 | 0.8878755163280 | 0.919417134234707 |
| Zfp984 | ENSMUSG00000078495 | 100041677 | -0.0243813146816587 | 1.61675110842175 | -0.143486430096681 | 0.8880406378836 | 0.919501014700771 |
| D16Ertd472e | ENSMUSG00000022864 | 67102 | 0.0265856365857422 | 2.537543954459 | 0.143135911329717 | 0.8883121381157 | 0.919655259002842 |
| Slc30a6 | ENSMUSG00000024069 | 210148 | 0.0218290596698314 | 1.37052505169383 | 0.143012506528202 | 0.8884077269942 | 0.919655259002842 |
| Rbm4b | ENSMUSG00000033760 | 66704 | 0.0225177270577757 | 1.47854445720841 | 0.142968256789168 | 0.8884420030025 | 0.919655259002842 |
| Zfp383 | ENSMUSG00000099689 | 73729 | 0.0214691361781831 | 2.03315862446512 | 0.14233485940724 | 0.8889326614695 | 0.92007602713039 |
| Btg2 | ENSMUSG00000020423 | 12227 | 0.0276171470962225 | 4.7336779613426 | 0.141688674003201 | 0.8894332750091 | 0.92050701847427 |
| Socs4 | ENSMUSG00000048379 | 67296 | 0.0143131488652242 | 3.8063551883453 | 0.140989465147671 | 0.8899750223366 | 0.920980495045127 |
| Ccl27a | ENSMUSG00000073888 | 20301 | -0.0249736410067087 | 1.38053083882498 | -0.140685397146526 | 0.8902106322532 | 0.921137109159425 |
| Tcea1 | ENSMUSG00000033813 | 21399 | 0.00850338910338788 | 4.79671295663498 | 0.140151719303349 | 0.8906241836246 | 0.921449204192321 |
| Fbxl5 | ENSMUSG00000039753 | 242960 | -0.0106948975890866 | 4.22573017646732 | -0.140078605510763 | 0.8906808426947 | 0.921449204192321 |
| Insr | ENSMUSG00000005534 | 16337 | -0.00774649055297871 | 6.4657936594913 | -0.139231670575635 | 0.8913372148543 | 0.922040985669825 |
| Mybbp1a | ENSMUSG00000040463 | 18432 | 0.00806138929523925 | 5.91216834045421 | 0.138611626268786 | 0.8918178001145 | 0.922393254286903 |
| D6Wsu163e | ENSMUSG00000030347 | 28040 | -0.0212982997217321 | 2.34847765148599 | -0.138574574748307 | 0.8918465194911 | 0.922393254286903 |
| Zbtb43 | ENSMUSG00000026788 | 71834 | -0.0198956866173212 | 3.75951044175099 | -0.138150707210553 | 0.8921750789701 | 0.92264576993594 |
| Cramp1l | ENSMUSG00000038002 | 57354 | 0.0144278348627322 | 3.8895393477427 | 0.13753018645754 | 0.8926561107386 | 0.923055902609436 |
| Fuca2 | ENSMUSG00000019810 | 66848 | 0.0183006587301211 | 2.34280365055352 | 0.137146473592005 | 0.8929535895313 | 0.923276171695719 |
| Lamtor5 | ENSMUSG00000087260 | 68576 | -0.01473357046551 | 2.85439409663822 | -0.137019076326222 | 0.8930523597694 | 0.92329096232285 |
| Mier2 | ENSMUSG00000042570 | 70427 | -0.0136552145302913 | 2.8358966664329 | -0.13663803775357 | 0.8933477874321 | 0.923509047136411 |
| Bves | ENSMUSG00000071317 | 23828 | -0.0105397136518723 | 4.87229736692601 | -0.135460991683915 | 0.8942604814239 | 0.924357948993355 |
| Med21 | ENSMUSG00000030291 | 108098 | -0.014459398427066 | 3.24852396079055 | -0.1353609131286 | 0.8943380906187 | 0.924357948993355 |
| Pik3c3 | ENSMUSG00000033628 | 225326 | -0.0136231147654065 | 3.59774231885618 | -0.135164203719452 | 0.8944906386627 | 0.924423866767685 |
| Map2k3 | ENSMUSG00000018932 | 26397 | 0.0107702177944837 | 6.26820902496159 | 0.135055058384521 | 0.8945752826908 | 0.924423866767685 |
| Arhgap18 | ENSMUSG00000039031 | 73910 | -0.0265346416341954 | 3.08598575116748 | -0.134951526472572 | 0.8946555746533 | 0.924423866767685 |
| Tmem129 | ENSMUSG00000019295 | 68366 | -0.0228556536884184 | 2.64217662229774 | -0.134774805560341 | 0.8947926295620 | 0.924478093926567 |
| Cys1 | ENSMUSG00000062563 | 12879 | 0.0400206790141986 | 1.02772634114511 | 0.13452204802618 | 0.8949886603321 | 0.92459323750977 |
| Foxn2 | ENSMUSG00000034998 | 14236 | -0.00970719110678697 | 4.3971050814587 | -0.13350694118061 | 0.8957760170967 | 0.925316176795745 |
| Sf3b1 | ENSMUSG00000025982 | 81898 | -0.00949568967316594 | 6.21303533199227 | -0.133401570787511 | 0.8958577530902 | 0.925316176795745 |
| Rps21 | ENSMUSG00000039001 | 66481 | -0.0121474219105098 | 5.81735445150483 | -0.133130153852271 | 0.8960682973455 | 0.925446197872678 |
| Zfp518a | ENSMUSG00000049164 | 72672 | -0.0167604867111462 | 3.5011607271958 | -0.132423960122785 | 0.8966161460110 | 0.925924524520217 |
| Stx4a | ENSMUSG00000030805 | 20909 | 0.00797046315306016 | 4.58989767712841 | 0.132273505591744 | 0.8967328722508 | 0.925957588000562 |
| Nsf | ENSMUSG00000034187 | 18195 | -0.0124674241769931 | 3.65358555490974 | -0.132124693512462 | 0.8968483266895 | 0.925989332109481 |
| Cables2 | ENSMUSG00000038990 | 252966 | -0.0148903657625055 | 2.9896067399473 | -0.132011669001157 | 0.8969360173113 | 0.925992406991953 |
| Mtr | ENSMUSG00000021311 | 238505 | 0.0145318318077923 | 4.1024849215138 | 0.131704719880124 | 0.8971741723751 | 0.926150805386055 |
| Sde2 | ENSMUSG00000038806 | 208768 | 0.0114006198097094 | 3.31721115468768 | 0.131570850571858 | 0.8972780418608 | 0.926151404517571 |
| Helq | ENSMUSG00000035266 | 191578 | 0.0186991978763396 | 1.66432259196578 | 0.131485576958854 | 0.8973442068653 | 0.926151404517571 |
| Ints14 | ENSMUSG00000034263 | 69882 | -0.0114577686235529 | 3.13987572366172 | -0.131143245605012 | 0.8976098346218 | 0.926338095001048 |
| Tmem19 | ENSMUSG00000069520 | 67226 | -0.0112043534379818 | 3.46478104021209 | -0.131022361710388 | 0.8977036360357 | 0.926347441282668 |
| Prdm2 | ENSMUSG00000057637 | 110593 | -0.00788723269472325 | 5.44500543847915 | -0.130613200403001 | 0.8980211418422 | 0.926553982602846 |
| Thap6 | ENSMUSG00000102644 | 381650 | -0.0196653241561846 | 1.73429891851084 | -0.130451522055685 | 0.8981466079318 | 0.926553982602846 |
| Gfra2 | ENSMUSG00000022103 | 14586 | -0.022933105685903 | 1.65490565039394 | -0.130436735876285 | 0.8981580824864 | 0.926553982602846 |
| Imp4 | ENSMUSG00000026127 | 27993 | -0.0137467620169565 | 2.91758433864726 | -0.13032015768348 | 0.8982485517758 | 0.926559867836343 |
| Psen1 | ENSMUSG00000019969 | 19164 | 0.0100332401826284 | 4.09634503693698 | 0.1299916384992 | 0.8985035035902 | 0.926735402693413 |
| Hdac7 | ENSMUSG00000022475 | 56233 | 0.0196233502800176 | 4.25647779736029 | 0.129495985896707 | 0.8988881836896 | 0.926981713632051 |
| Setx | ENSMUSG00000043535 | 269254 | -0.0104355769430473 | 6.21744197879515 | -0.129465407148108 | 0.8989119169792 | 0.926981713632051 |
| Ell2 | ENSMUSG00000001542 | 192657 | 0.0208245770773359 | 3.23691084263763 | 0.129300737429436 | 0.8990397249271 | 0.926998875438897 |
| Rin3 | ENSMUSG00000044456 | 217835 | 0.0188215971962702 | 3.58602385172119 | 0.129225439379736 | 0.8990981682740 | 0.926998875438897 |
| Pithd1 | ENSMUSG00000028669 | 66193 | -0.0136371555283112 | 3.73942659804729 | -0.128881201599206 | 0.8993653596955 | 0.927186904350848 |
| Pi15 | ENSMUSG00000067780 | 94227 | -0.0321538255797451 | 2.10088755068027 | -0.128322787343639 | 0.8997988182438 | 0.927546292174934 |
| Zscan12 | ENSMUSG00000036721 | 22758 | -0.0159368419381671 | 2.08728756474759 | -0.127787951446057 | 0.9002140057937 | 0.927886779569248 |
| Gdap2 | ENSMUSG00000027865 | 14547 | 0.01619127181488 | 2.7387191848423 | 0.127400702550473 | 0.9005146421046 | 0.928109141320597 |
| Gipc3 | ENSMUSG00000034872 | 209047 | 0.0222828653307388 | 1.48765249652891 | 0.126644906541316 | 0.9011014416214 | 0.928517391164429 |
| Pold3 | ENSMUSG00000030726 | 67967 | -0.0166849576377248 | 2.24402979625977 | -0.126582856962636 | 0.9011496195556 | 0.928517391164429 |
| 9930111J21Rik1 | ENSMUSG00000069893 | 667214 | -0.023830394715294 | 2.1137574061249 | -0.126562295576691 | 0.9011655843805 | 0.928517391164429 |
| 9330159M07Rik | ENSMUSG00000097177 | 319673 | -0.0161460057192986 | 1.73717693908142 | -0.126191796079879 | 0.9014532652097 | 0.928726262206201 |
| Rrnad1 | ENSMUSG00000004896 | 229503 | -0.0282138050022263 | 1.64917042422759 | -0.125984596937908 | 0.9016141549128 | 0.928804479064418 |
| Zmat2 | ENSMUSG00000001383 | 66492 | -0.0108108986338902 | 4.66204884714981 | -0.12576155786842 | 0.9017873493245 | 0.928895355773342 |
| Lonrf1 | ENSMUSG00000039633 | 244421 | -0.0217936856129777 | 2.00123859691815 | -0.125581247389593 | 0.9019273679213 | 0.928952045486532 |
| Psmb3 | ENSMUSG00000069744 | 26446 | 0.00982707872010469 | 4.60437644507339 | 0.125418254606747 | 0.9020539415346 | 0.928994877983325 |
| Rnaseh1 | ENSMUSG00000020630 | 19819 | 0.022222629276395 | 1.67859139638108 | 0.124791124625782 | 0.9025409712220 | 0.929408888971072 |
| Rpf1 | ENSMUSG00000028187 | 70285 | -0.010477018652495 | 3.24787714826191 | -0.124359327890547 | 0.9028763287359 | 0.929666649341839 |
| Atf3 | ENSMUSG00000026628 | 11910 | -0.0366698215670782 | 4.08360441040735 | -0.124131757876042 | 0.9030530801295 | 0.929761064226857 |
| Parp2 | ENSMUSG00000036023 | 11546 | -0.0138882482501148 | 2.50264036536595 | -0.123248345350853 | 0.9037392686515 | 0.930379915768467 |
| Gls | ENSMUSG00000026103 | 14660 | 0.00663296439957328 | 5.05707082086579 | 0.122389592319861 | 0.9044063798487 | 0.930979012913295 |
| Psme4 | ENSMUSG00000040850 | 103554 | -0.00863021060421065 | 7.33948305772436 | -0.121916771114712 | 0.9047737170550 | 0.931269444550712 |
| Ccs | ENSMUSG00000034108 | 12460 | 0.0189627500283109 | 1.74754942165997 | 0.121431902554545 | 0.9051504375185 | 0.931569478628676 |
| Cars2 | ENSMUSG00000056228 | 71941 | -0.0174653939600375 | 2.59985034528184 | -0.121256853111534 | 0.9052864487005 | 0.931577868196432 |
| Tor1aip2 | ENSMUSG00000050565 | 240832 | 0.00725129290627269 | 5.52621954611797 | 0.12120204296468 | 0.9053290361221 | 0.931577868196432 |
| Phf1 | ENSMUSG00000024193 | 21652 | 0.0134965338635258 | 3.07803552860208 | 0.120669494804086 | 0.9057428412245 | 0.931909208541351 |
| Clec1a | ENSMUSG00000033082 | 243653 | -0.017747381230897 | 1.88630042060336 | -0.120568207013381 | 0.9058215479601 | 0.931909208541351 |
| Plxnb1 | ENSMUSG00000053646 | 235611 | -0.0307241120280599 | 1.03573134431484 | -0.120265095559657 | 0.9060570900411 | 0.932063810581621 |
| Pdap1 | ENSMUSG00000029623 | 231887 | -0.00952960234434208 | 5.53502456019985 | -0.119832465265653 | 0.9063932946828 | 0.932321925677773 |
| Asap1 | ENSMUSG00000022377 | 13196 | 0.0142483472066433 | 4.40863500686777 | 0.119473323976943 | 0.9066724039366 | 0.932443686619681 |
| Abce1 | ENSMUSG00000058355 | 24015 | -0.00718835219875569 | 5.54380191161679 | -0.119460623125823 | 0.9066822747306 | 0.932443686619681 |
| Eefsec | ENSMUSG00000033216 | 65967 | 0.0206054244945753 | 1.47945714379289 | 0.119290768274334 | 0.9068142833538 | 0.932476537348818 |
| Trim41 | ENSMUSG00000040365 | 211007 | -0.0108444153099665 | 3.73166409112171 | -0.119151273267105 | 0.9069226989124 | 0.932476537348818 |
| Klhl25 | ENSMUSG00000055652 | 207952 | -0.0210092856793156 | 1.29446758355808 | -0.119090239546335 | 0.9069701349458 | 0.932476537348818 |
| Sri | ENSMUSG00000003161 | 109552 | 0.0151240777607868 | 3.06640711885031 | 0.118910475446701 | 0.9071098515962 | 0.932532473224692 |
| Dennd1a | ENSMUSG00000035392 | 227801 | -0.00795268664852333 | 3.81318706308167 | -0.117467202719283 | 0.9082317110073 | 0.933597971884664 |
| H2-D1 | ENSMUSG00000073411 | 14964 | -0.0140598044122973 | 6.20083316191151 | -0.11663156263065 | 0.9088813495836 | 0.934177906186995 |
| Slc12a6 | ENSMUSG00000027130 | 107723 | 0.00764968071657179 | 4.72005247093027 | 0.116323861438029 | 0.9091205780470 | 0.934335938194065 |
| Rarb | ENSMUSG00000017491 | 218772 | 0.0182499058351593 | 2.3982343089203 | 0.115622884381282 | 0.9096656010042 | 0.934808186949102 |
| Rsph3b | ENSMUSG00000023806 | 100037282 | -0.00872191402543177 | 5.17610303869206 | -0.115202715794415 | 0.9099923128003 | 0.93505602286343 |
| Thada | ENSMUSG00000024251 | 240174 | 0.0111645952444932 | 3.06321539919847 | 0.114422644461875 | 0.9105989202687 | 0.935576230945646 |
| Trim34a | ENSMUSG00000056144 | 94094 | -0.0270604482931873 | 1.31434343035702 | -0.114261043500869 | 0.9107245934292 | 0.935576230945646 |
| Ifit1 | ENSMUSG00000034459 | 15957 | -0.042319450415734 | 2.61362150239517 | -0.114139407322486 | 0.9108191885880 | 0.935576230945646 |
| Gigyf2 | ENSMUSG00000048000 | 227331 | 0.00620604004951986 | 5.4539182439544 | 0.114111445992482 | 0.9108409340155 | 0.935576230945646 |
| Krtcap2 | ENSMUSG00000042747 | 66059 | 0.0122602289022757 | 3.03155901428013 | 0.113222263898872 | 0.9115324865380 | 0.93619859159521 |
| Fbxl7 | ENSMUSG00000043556 | 448987 | -0.0288693326407036 | 0.886747572755643 | -0.112637991528953 | 0.9119869392892 | 0.936577342481305 |
| Ift80 | ENSMUSG00000027778 | 68259 | 0.0157168747736267 | 2.20618684805587 | 0.112204868089675 | 0.9123238476606 | 0.936835319753739 |
| Hnrnpul1 | ENSMUSG00000040725 | 232989 | -0.00627180736631683 | 6.17357341239818 | -0.111821747249662 | 0.9126218757825 | 0.937053327463791 |
| Zfp422 | ENSMUSG00000059878 | 67255 | -0.0126003875406036 | 2.2903264805269 | -0.110839772720151 | 0.9133858122046 | 0.937749630244146 |
| 1700030K09Rik | ENSMUSG00000052794 | 72254 | -0.0165295189294528 | 1.31317714585475 | -0.110495788763788 | 0.9136534389478 | 0.937936301759874 |
| Gnpda1 | ENSMUSG00000052102 | 26384 | -0.0230777317211426 | 2.72545472826529 | -0.109807573699496 | 0.9141889174732 | 0.93839788307827 |
| Gstm4 | ENSMUSG00000027890 | 14865 | 0.0172762273486322 | 1.91333450081148 | 0.109362796386894 | 0.9145350078604 | 0.938614030892033 |
| Lztfl1 | ENSMUSG00000025245 | 93730 | 0.0108540230209293 | 3.29408039383756 | 0.109316255403313 | 0.9145712234041 | 0.938614030892033 |
| Usp34 | ENSMUSG00000056342 | 17847 | -0.00778681124536718 | 6.60273736598588 | -0.109165031021777 | 0.9146888989734 | 0.93864667242575 |
| Tspan14 | ENSMUSG00000037824 | 52588 | 0.016583537204736 | 2.03490979490805 | 0.108003349704793 | 0.9155929330634 | 0.939486187113216 |
| Senp1 | ENSMUSG00000033075 | 223870 | 0.0113529762049667 | 3.22107583533917 | 0.107656562393603 | 0.9158628306990 | 0.939623698496045 |
| C87436 | ENSMUSG00000046679 | 232196 | -0.0126044890624954 | 1.93399384695769 | -0.10761026186416 | 0.9158988662954 | 0.939623698496045 |
| Tbcel | ENSMUSG00000037287 | 272589 | 0.00770295806564913 | 4.23737034098929 | 0.10718141487665 | 0.9162326460780 | 0.939877914252889 |
| Nup205 | ENSMUSG00000038759 | 70699 | 0.00895072037437306 | 3.79112501205469 | 0.106552075094421 | 0.9167225030300 | 0.940292172339413 |
| Dhx36 | ENSMUSG00000027770 | 72162 | -0.00748829498946968 | 4.27375964161407 | -0.106384510943556 | 0.9168529352920 | 0.940337721493433 |
| Ifi203 | ENSMUSG00000039997 | 15950 | 0.0195454013896703 | 3.87512566347291 | 0.105763829083964 | 0.9173360969826 | 0.940744992599393 |
| Yipf6 | ENSMUSG00000047694 | 77929 | -0.00897130586208394 | 3.3554425257416 | -0.10485427047262 | 0.9180441924747 | 0.941382839394117 |
| Trim33 | ENSMUSG00000033014 | 94093 | -0.0070282474491945 | 4.26068088081527 | -0.104677045317379 | 0.9181821715200 | 0.941436011339019 |
| Rnpepl1 | ENSMUSG00000026269 | 108657 | 0.00607632877930175 | 5.13919890713264 | 0.104410255838286 | 0.9183898862535 | 0.941560668414701 |
| Cox6a1 | ENSMUSG00000041697 | 12861 | -0.0152925891245386 | 3.20093368063577 | -0.104083008952547 | 0.9186446798942 | 0.941733564280614 |
| Rbm41 | ENSMUSG00000031433 | 237073 | -0.011150801175611 | 2.52168692887644 | -0.103468280574371 | 0.9191233311064 | 0.942135890127993 |
| Mbd3 | ENSMUSG00000035478 | 17192 | -0.00893893856752204 | 5.78745705244612 | -0.10319164029937 | 0.9193387445661 | 0.942268337257639 |
| Dbp | ENSMUSG00000059824 | 13170 | -0.0387248300816738 | 4.90784796624614 | -0.102528225187676 | 0.9198553576226 | 0.942709442543888 |
| Gne | ENSMUSG00000028479 | 50798 | 0.011399827268475 | 2.61910193232818 | 0.102302081509985 | 0.9200314683312 | 0.942801535607829 |
| BC031181 | ENSMUSG00000036299 | 407819 | 0.0130247118968421 | 2.5809540353018 | 0.101697943269098 | 0.9205019659406 | 0.943117139382946 |
| Arl14ep | ENSMUSG00000027122 | 212772 | -0.013891886726646 | 2.49419635763659 | -0.101685049214104 | 0.9205120080575 | 0.943117139382946 |
| Pcdh17 | ENSMUSG00000035566 | 219228 | -0.0211089820777288 | 1.9512082221242 | -0.10104691187513 | 0.9210090182729 | 0.943537917407864 |
| E130308A19Rik | ENSMUSG00000045071 | 230259 | 0.0129872757461273 | 1.71598384957543 | 0.100710168268549 | 0.9212713033061 | 0.943718172283743 |

| | | | | | | | |
|---|---|---|---|---|---|---|---|
| Sptan1 | ENSMUSG00000057738 | 20740 | -0.0107720904734681 | 7.18923757238906 | -0.0998942381120278 | 0.9219068597992 | 0.94425428544983 |
| Gapvd1 | ENSMUSG00000026867 | 66691 | -0.00623989585962903 | 5.28823292350944 | -0.0998164802400083 | 0.9219674310315 | 0.94425428544983 |
| Commd10 | ENSMUSG00000042705 | 69456 | -0.013573757112336 | 2.33416929068432 | -0.0995276507042408 | 0.9221924256709 | 0.944396234308525 |
| Tfb1m | ENSMUSG00000036983 | 224481 | -0.0157319895438379 | 1.87440183508579 | -0.0992582455906381 | 0.9224022951862 | 0.944508427985123 |
| Utp11 | ENSMUSG00000028907 | 67205 | 0.00853991127112685 | 4.34204571752895 | 0.099165181711914 | 0.9224747943618 | 0.944508427985123 |
| Zfp229 | ENSMUSG00000061544 | 381067 | 0.0132036616809841 | 2.50093929138311 | 0.0985070100746649 | 0.9229875476096 | 0.944944917385116 |
| Grasp | ENSMUSG00000000531 | 56149 | -0.0159586557949867 | 1.4456486142779 | -0.0982109171182351 | 0.9232182325686 | 0.945008813026342 |
| Aagab | ENSMUSG00000037257 | 66939 | 0.0107813211224013 | 3.20365649245019 | 0.0982049740119275 | 0.9232228628952 | 0.945008813026342 |
| Agk | ENSMUSG00000029916 | 69923 | -0.0112411437325211 | 2.22989959811701 | -0.0976742240252443 | 0.9236363865773 | 0.945343571318081 |
| G3bp1 | ENSMUSG00000018583 | 27041 | 0.00692007342248502 | 6.42508742117962 | 0.0972873684445793 | 0.9239378122022 | 0.945563545097114 |
| Zfp449 | ENSMUSG00000073176 | 78619 | 0.0140283354444952 | 1.56118958763026 | 0.0969897957047418 | 0.9241696797551 | 0.945712298202959 |
| Tomm6 | ENSMUSG00000033475 | 66119 | 0.00715206979937566 | 4.39076275291343 | 0.0965926549665534 | 0.9244791415074 | 0.945940418966388 |
| Chmp4c | ENSMUSG00000027536 | 66371 | 0.0162255981548456 | 2.51062977918587 | 0.095592474673782 | 0.9252585627581 | 0.94664932136926 |
| Dnajc24 | ENSMUSG00000027166 | 99349 | -0.0205144266767814 | 1.24581452993856 | -0.0954421412869614 | 0.9253757216115 | 0.946680581463377 |
| Cbx3 | ENSMUSG00000029836 | 12417 | -0.00571984570017925 | 4.71469380356639 | -0.0952409588882862 | 0.9255325112920 | 0.946752375157529 |
| Crtc3 | ENSMUSG00000030527 | 70461 | 0.00900098591275178 | 3.74175706862609 | 0.0946832960652201 | 0.9259671376077 | 0.947108335472134 |
| Aqr | ENSMUSG00000040383 | 11834 | -0.00784309953594597 | 4.07688172302671 | -0.094395181018843 | 0.9261916958274 | 0.947249385019657 |
| Plekhm3 | ENSMUSG00000051344 | 241075 | -0.0098842400234167 | 3.42642847744828 | -0.0942046022532594 | 0.9263402374172 | 0.947312670521807 |
| Aup1 | ENSMUSG00000068328 | 11993 | -0.00700699182453235 | 4.7238230950588 | -0.0938038852639029 | 0.9266525750349 | 0.947543432900607 |
| Slc20a1 | ENSMUSG00000027397 | 20515 | 0.0274492925897714 | 3.47300280662067 | 0.0936122348256439 | 0.9268019608365 | 0.947607542222867 |
| Snapc1 | ENSMUSG00000021113 | 75627 | -0.00929158135929553 | 3.4238141995273 | -0.0930449191998827 | 0.9272441832971 | 0.947971022037189 |
| E130311K13Rik | ENSMUSG00000048581 | 329659 | -0.0136737314408084 | 1.87017633176539 | -0.0916173874950093 | 0.9283570546531 | 0.949020009764631 |
| Fyttd1 | ENSMUSG00000022800 | 69823 | -0.00668577290755223 | 5.49317826949073 | -0.0910547010791025 | 0.9287957553126 | 0.949379689669197 |
| Ube3c | ENSMUSG00000039000 | 100763 | -0.00732744608151492 | 5.27924666086299 | -0.0907672726677894 | 0.9290198593189 | 0.949519970286664 |
| Fzr1 | ENSMUSG00000020235 | 56371 | -0.00675680450652916 | 3.52294068296607 | -0.0905292626604612 | 0.9292054372241 | 0.94962085212199 |
| Zfp81 | ENSMUSG00000003929 | 224694 | -0.0162821960294952 | 2.04433195048601 | -0.0898344361750267 | 0.9297472223472 | 0.950085714450598 |
| Srsf5 | ENSMUSG00000021134 | 20384 | 0.00849630171306792 | 5.60239492362605 | 0.0896897290829942 | 0.9298600610673 | 0.950112201114872 |
| Sypl | ENSMUSG00000020570 | 19027 | -0.00596190223221615 | 5.52483446414208 | -0.0894341132464688 | 0.9300593873013 | 0.950227045760495 |
| 1810026B05Rik | ENSMUSG00000101970 | 69170 | -0.0153411340020515 | 2.41585758803806 | -0.0887937492803536 | 0.9305587570746 | 0.950627434349325 |
| Lsm2 | ENSMUSG00000007050 | 27756 | -0.0135679154376591 | 1.86323133160355 | -0.0887085361832064 | 0.9306252104082 | 0.950627434349325 |
| Tgoln1 | ENSMUSG00000056429 | 22134 | 0.00670771682906413 | 5.37363716426009 | 0.0883135131661462 | 0.9309332756824 | 0.950853264481872 |
| Zmiz2 | ENSMUSG00000041164 | 52915 | -0.00593543001174179 | 5.26502405301492 | -0.0880213784743707 | 0.9311611092110 | 0.950882834685987 |
| Adck2 | ENSMUSG00000046947 | 57869 | 0.0218376315262787 | 1.48321805839821 | 0.0880120414521421 | 0.9311683911845 | 0.950882834685987 |
| Rarres2 | ENSMUSG00000009281 | 71660 | 0.0118624132221288 | 2.42913814689997 | 0.0879417716030307 | 0.9312231950703 | 0.950882834685987 |
| Klf7 | ENSMUSG00000025959 | 93691 | -0.0135218695711008 | 3.88218072007957 | -0.0872613140010335 | 0.9317539068936 | 0.95133588233271 |
| Rpap1 | ENSMUSG00000034032 | 68925 | -0.00588301733079177 | 3.76280694563094 | -0.08709955523047 | 0.9318800730008 | 0.951375836179382 |
| Ngly1 | ENSMUSG00000021785 | 59007 | 0.00593684049632248 | 4.52687073655668 | 0.0868693985077503 | 0.9320595903761 | 0.951470244901193 |
| Prmt5 | ENSMUSG00000023110 | 27374 | -0.00736075550998283 | 4.20331873390462 | -0.0864169599872869 | 0.9324124940793 | 0.951712689046959 |
| Wrap73 | ENSMUSG00000029029 | 59002 | 0.0110822558932511 | 1.9657183678127 | 0.0863097510842889 | 0.9324961195910 | 0.951712689046959 |
| Hist2h4 | ENSMUSG00000091405 | 97122 | 0.0164419712289463 | 1.38461259575552 | 0.0862300550911391 | 0.9325582849128 | 0.951712689046959 |
| Glul | ENSMUSG00000026473 | 14645 | 0.0113214171925158 | 6.38915949022671 | 0.0858350328783145 | 0.9328664211333 | 0.95193827944439 |
| Lig4 | ENSMUSG00000049717 | 319583 | -0.0130372964888656 | 2.35591937795619 | -0.0853719061274902 | 0.9332276964373 | 0.952172507745668 |
| Hinfp | ENSMUSG00000032119 | 102423 | 0.0107549983940598 | 2.2704355080618 | 0.0853174573159595 | 0.9332701718037 | 0.952172507745668 |
| Gm12185 | ENSMUSG00000048852 | 620913 | 0.0476116498247763 | 1.49702361167687 | 0.0848992810053012 | 0.9335963970346 | 0.952416445728951 |
| Plk3 | ENSMUSG00000028680 | 12795 | 0.0336739623893411 | 0.550387567690069 | 0.0844537342062245 | 0.9339439880550 | 0.952682132645574 |
| Myl12a | ENSMUSG00000024048 | 67268 | -0.00818279955740486 | 8.31010674442073 | -0.0841993719651088 | 0.9341424337397 | 0.952795646469132 |
| Gng12 | ENSMUSG00000036402 | 14701 | 0.00724618354840345 | 4.09384654850007 | 0.0832373471147021 | 0.9348930174366 | 0.953465910648213 |
| Olfr624 | ENSMUSG00000045780 | 258189 | 0.0157435878027448 | 2.08273868958794 | 0.0830407806184399 | 0.9350463890173 | 0.953465910648213 |
| Tecpr2 | ENSMUSG00000021275 | 104859 | 0.00889964152301514 | 2.6291813040587 | 0.0830217299333874 | 0.9350612535128 | 0.953465910648213 |
| Fam78a | ENSMUSG00000050592 | 241303 | -0.0169245750856282 | 1.47891584167209 | -0.0828025648126293 | 0.9352322611962 | 0.953496265008182 |
| Hist1h1c | ENSMUSG00000036181 | 50708 | -0.00925808091261082 | 6.74502750597978 | -0.0827599927566885 | 0.9352654792258 | 0.953496265008182 |
| Vgll4 | ENSMUSG00000030315 | 232334 | -0.0125853194382524 | 2.47247847176528 | -0.0826263282956923 | 0.9353697754521 | 0.953513663663851 |
| Celf1 | ENSMUSG00000005506 | 13046 | -0.00469392991160156 | 5.66404771465049 | -0.0806181181376034 | 0.9369368973068 | 0.955022120893463 |
| Dchs2 | ENSMUSG00000102692 | 100534287 | -0.0199370258676093 | 1.63792653175866 | -0.0798211012247522 | 0.9375589313513 | 0.955556275705147 |
| Tmem50b | ENSMUSG00000022964 | 77975 | -0.00951232791895285 | 3.20257407242456 | -0.0797226497452842 | 0.9376357710377 | 0.955556275705147 |
| Nt5dc3 | ENSMUSG00000054027 | 103466 | -0.00474927093432509 | 4.74084514398893 | -0.0790908930741618 | 0.9381288616348 | 0.955969664074814 |
| Ogfod3 | ENSMUSG00000025169 | 66179 | -0.0122873571117079 | 1.85735798454886 | -0.0785614492104739 | 0.9385421167456 | 0.956301629208347 |
| Vopp1 | ENSMUSG00000037788 | 232023 | -0.0125622820185918 | 1.70594166313423 | -0.0783110040810914 | 0.9387376070449 | 0.956344836579023 |
| Kat2b | ENSMUSG00000000708 | 18519 | -0.00618891420444001 | 6.24687900999259 | -0.0782829568308654 | 0.9387595001823 | 0.956344836579023 |
| Taf8 | ENSMUSG00000023980 | 63856 | 0.00722881577665824 | 3.11116915418192 | 0.0772407057500909 | 0.9395730975319 | 0.957084477599671 |
| Chid1 | ENSMUSG00000025512 | 68038 | 0.00985566312741915 | 3.07686722464477 | 0.0762394140538801 | 0.9403547876256 | 0.95779148187996 |
| Ypel5 | ENSMUSG00000039770 | 383295 | 0.00586373482445435 | 4.15806327367973 | 0.0758823419865126 | 0.9406335627990 | 0.957884197806933 |
| Snn | ENSMUSG00000037972 | 20621 | -0.014390143258423 | 1.70530156532302 | -0.0758595586393724 | 0.9406513506088 | 0.957884197806933 |
| Spopl | ENSMUSG00000026771 | 76857 | 0.00587616334354024 | 3.48808146693846 | 0.0757860960339043 | 0.9407087058327 | 0.957884197806933 |
| Bin3 | ENSMUSG00000022089 | 57784 | -0.0101565594973967 | 1.83853403957751 | -0.0756603960142336 | 0.9408068457115 | 0.95789489851652 |
| Zfp157 | ENSMUSG00000036898 | 72154 | -0.0144234412527102 | 1.41192277599898 | -0.0750586790626439 | 0.9412766481092 | 0.958283975084456 |
| Sept7 | ENSMUSG00000001833 | 235072 | 0.00481188440732172 | 6.34505099906583 | 0.0738624686804816 | 0.9422106805139 | 0.959135532372376 |
| Zfp780b | ENSMUSG00000063047 | 338354 | -0.00867844511679072 | 2.61748978472784 | -0.0737627067134799 | 0.9422885812960 | 0.959135532372376 |
| Ss18 | ENSMUSG00000037013 | 268996 | 0.00824716697444742 | 3.74320637945158 | 0.0731345108656977 | 0.9427791325277 | 0.959545502575509 |
| Josd2 | ENSMUSG00000038695 | 66124 | -0.00936968394593274 | 2.74420962439574 | -0.0730149593414771 | 0.9428724917471 | 0.95955117840873 |
| Unc5a | ENSMUSG00000025876 | 107448 | -0.00879490741697264 | 2.50315034185991 | -0.0724134329619989 | 0.9433422441667 | 0.959939868831383 |
| Heatr5b | ENSMUSG00000039414 | 320473 | -0.00648684841983416 | 3.47162769582456 | -0.0719627622947093 | 0.9436942026398 | 0.960208631579265 |
| Utp25 | ENSMUSG00000016181 | 215193 | 0.0131857942958473 | 2.17040774463986 | 0.0714286338658136 | 0.9441113546597 | 0.960438906620646 |
| Gmfb | ENSMUSG00000062014 | 63985 | 0.00642826499855786 | 3.81679254281151 | 0.0713625001855391 | 0.9441630059733 | 0.960438906620646 |
| Yme1l1 | ENSMUSG00000026775 | 27377 | -0.00502613123729478 | 5.41336155825186 | -0.0713354808149907 | 0.9441841085486 | 0.960438906620646 |
| Ilf2 | ENSMUSG00000001016 | 67781 | -0.00725347551407873 | 3.84095033669334 | -0.0711272108636083 | 0.9443467722904 | 0.960441602660067 |
| Dmpk | ENSMUSG00000030409 | 13400 | -0.00544531190563902 | 7.60335498051809 | -0.0711070903804124 | 0.9443624869993 | 0.960441602660067 |
| Ccdc57 | ENSMUSG00000048445 | 71276 | 0.0117299862040583 | 1.5386636199208 | 0.0702386374000382 | 0.9450407981006 | 0.961042047077727 |
| Stx12 | ENSMUSG00000028879 | 100226 | 0.0045124018769771 | 4.66586976211973 | 0.0692585256221936 | 0.9458063744238 | 0.961674884888717 |
| Samhd1 | ENSMUSG00000027639 | 56045 | 0.00881735910216186 | 4.4719933858175 | 0.0692166913174804 | 0.9458390529172 | 0.961674884888717 |
| Alg11 | ENSMUSG00000063362 | 207958 | -0.00888682335304969 | 2.30868132655569 | -0.0673318345637129 | 0.9473114964616 | 0.963082400281055 |
| Ppm1h | ENSMUSG00000034613 | 319468 | -0.0118256499927655 | 1.67595815220747 | -0.0663885832031894 | 0.9480484368660 | 0.963741975577325 |
| Mpv17l2 | ENSMUSG00000035559 | 234384 | 0.0107723671243685 | 1.67303886440432 | 0.0659727251131695 | 0.9483733529319 | 0.963913033297252 |
| Fer | ENSMUSG00000000127 | 14158 | -0.00517826725140758 | 3.86940126014851 | -0.0659474868832904 | 0.9483930722817 | 0.963913033297252 |
| Mettl2 | ENSMUSG00000020691 | 52686 | -0.00799355018491221 | 1.82452272453297 | -0.0651391015140361 | 0.9490247041362 | 0.964465325484722 |
| Kdm2b | ENSMUSG00000029475 | 30841 | 0.00633719739214555 | 3.50583592824563 | 0.0649405373913423 | 0.9491798576932 | 0.964526479156914 |
| Mvd | ENSMUSG00000006517 | 192156 | -0.0124571339975785 | 1.94514141014982 | -0.0646217262686751 | 0.9494289740549 | 0.964526479156914 |
| Commd2 | ENSMUSG00000036513 | 52245 | -0.00925576017562202 | 1.95628361523594 | -0.0644923960645695 | 0.9495300331750 | 0.964526479156914 |
| Kctd18 | ENSMUSG00000054770 | 51960 | 0.0103293638894717 | 1.64820820908563 | 0.064426881526755 | 0.9495812268332 | 0.964526479156914 |
| Foxo6os | ENSMUSG00000084929 | 402730 | 0.0119449489770505 | 3.83284530676613 | 0.0644033798547231 | 0.9495995913109 | 0.964526479156914 |
| Dkc1 | ENSMUSG00000031403 | 245474 | -0.00740335811333726 | 3.60929219115681 | -0.0643845501680518 | 0.9496143050669 | 0.964526479156914 |
| Clk3 | ENSMUSG00000032316 | 102414 | -0.00549581476034255 | 4.05256137069591 | -0.0639133577869998 | 0.9499825069951 | 0.964810813338622 |
| Rala | ENSMUSG00000008859 | 56044 | -0.00480312247190273 | 4.59051418013548 | -0.0631274872461633 | 0.9505966330447 | 0.965343644210475 |
| Btbd3 | ENSMUSG00000062098 | 228662 | -0.0061810515634475 | 4.0014473544325 | -0.0630159806596615 | 0.9506837736644 | 0.965343644210475 |
| Noc2l | ENSMUSG00000095567 | 57741 | 0.00503471899742625 | 4.83834431097123 | 0.0622768957810354 | 0.9512613727522 | 0.965840429644655 |
| Maoa | ENSMUSG00000025037 | 17161 | 0.00945016563409151 | 2.16328154602856 | 0.0620778911341588 | 0.9514169009429 | 0.965908623034079 |
| Zfp622 | ENSMUSG00000052253 | 52521 | 0.0067571176801513 | 3.49629965218752 | 0.0611513486487998 | 0.9521410491781 | 0.966535311764342 |
| Exoc2 | ENSMUSG00000021357 | 66482 | -0.00604514041994664 | 3.49895246344587 | -0.0609929625083796 | 0.9522648418271 | 0.966535311764342 |
| Rbck1 | ENSMUSG00000027466 | 24105 | -0.00406536541026625 | 4.43835564382045 | -0.0609486811657312 | 0.9522994518069 | 0.966535311764342 |
| Vps16 | ENSMUSG00000027411 | 80743 | 0.00523348540649859 | 2.96994190244442 | 0.0607985422163488 | 0.9524168000795 | 0.966564668245227 |
| Naif1 | ENSMUSG00000039164 | 71254 | 0.0098534726692141 | 1.31619490150908 | 0.0601500415521335 | 0.9529236799563 | 0.966989300557235 |
| Mon1b | ENSMUSG00000078908 | 270096 | -0.00661948273266377 | 2.42692712269252 | -0.0597412870015745 | 0.9532431808523 | 0.967223726900304 |
| Ptprd | ENSMUSG00000028399 | 19266 | 0.00542053989939273 | 3.74364753650979 | 0.0591318694101046 | 0.9537195445310 | 0.967502143782407 |
| Cyp26b1 | ENSMUSG00000063415 | 232174 | 0.0274784887227972 | 2.32722413785756 | 0.0590060527113194 | 0.9538178940204 | 0.967502143782407 |
| Setdb2 | ENSMUSG00000071350 | 239122 | -0.00736232304204054 | 1.77366880505774 | -0.0588055481842459 | 0.9539746277603 | 0.967502143782407 |
| Hps4 | ENSMUSG00000042328 | 192232 | 0.00859220821949136 | 1.82294498201133 | 0.0587853605755083 | 0.9539904084588 | 0.967502143782407 |
| Arhgef10l | ENSMUSG00000040964 | 72754 | 0.00323820261650454 | 5.50664377544425 | 0.0587759292405475 | 0.9539977809612 | 0.967502143782407 |
| Rtel1 | ENSMUSG00000038685 | 269400 | -0.00761797212724602 | 2.13917779846235 | -0.05858220640739 | 0.9541492156416 | 0.967502143782407 |
| Rnf170 | ENSMUSG00000013878 | 77733 | -0.00482198622274055 | 4.02553150646873 | -0.0584963377850321 | 0.9542163404180 | 0.967502143782407 |
| Gm21992 | ENSMUSG00000096370 | 102902673 | -0.00890563133256588 | 2.36021408117294 | -0.0584844240914816 | 0.9542256529245 | 0.967502143782407 |
| Clk1 | ENSMUSG00000026034 | 12747 | -0.00856482071438069 | 4.58370321513682 | -0.0576030686224745 | 0.9549146435640 | 0.968030910776386 |
| Rack1 | ENSMUSG00000020372 | 14694 | 0.00451719723857469 | 6.97810381733371 | 0.0575140987823803 | 0.9549841968621 | 0.968030910776386 |
| Agpat5 | ENSMUSG00000031467 | 52123 | 0.00531482259902916 | 3.74772199031373 | 0.0574774611813721 | 0.9550128388813 | 0.968030910776386 |
| Ubald2 | ENSMUSG00000050628 | 319370 | -0.00584141504334168 | 3.6411060916427 | -0.0573120931344415 | 0.9551421187516 | 0.968072183595229 |
| Zfp207 | ENSMUSG00000017421 | 22680 | -0.00399925139421686 | 5.17372172524799 | -0.0559954783316847 | 0.9561714560884 | 0.969025606017308 |
| Maco1 | ENSMUSG00000028826 | 66146 | -0.00479582824673219 | 4.28087233092088 | -0.055692432477355 | 0.9564083907643 | 0.969175870904326 |

| | | | | | | | |
|---|---|---|---|---|---|---|---|
| Casz1 | ENSMUSG00000028977 | 69743 | 0.0039559787526402 | 5.80052957206632 | 0.0554779619318862 | 0.9565760759367 | 0.969255940495471 |
| Psmc1 | ENSMUSG00000021178 | 19179 | 0.00451578141524145 | 6.08165659075543 | 0.0543046972734706 | 0.9574934381599 | 0.969984651138246 |
| Plrg1 | ENSMUSG00000027998 | 53317 | -0.00450190580912713 | 3.48096706829713 | -0.0542470724449278 | 0.9575384959835 | 0.969984651138246 |
| Eif3k | ENSMUSG00000053565 | 73830 | -0.00423397056215613 | 4.73915996338259 | -0.0542176976589734 | 0.9575614646814 | 0.969984651138246 |
| Raver1 | ENSMUSG00000010205 | 71766 | 0.00347363416676938 | 4.50777988626273 | 0.0539554941820519 | 0.9577664882527 | 0.970061474401265 |
| Col24a1 | ENSMUSG00000028197 | 71355 | -0.0128357627035756 | 1.31515022240562 | -0.0538052750021068 | 0.9578839498575 | 0.970061474401265 |
| Smim27 | ENSMUSG00000028407 | 100504309 | -0.00912916881985137 | 1.37271085603998 | -0.0537802265102028 | 0.9579035362443 | 0.970061474401265 |
| Epc2 | ENSMUSG00000069495 | 227867 | -0.00355971682652712 | 4.03764619662274 | -0.0533370926686884 | 0.9582500444483 | 0.970322485953189 |
| Rps28 | ENSMUSG00000067288 | 54127 | -0.00351657407895356 | 5.79898800113248 | -0.0531773159288328 | 0.9583749839227 | 0.9703591097869 |
| Tjp2 | ENSMUSG00000024812 | 21873 | 0.00565119339169406 | 3.26455579093951 | 0.052862485132614 | 0.9586211732430 | 0.970518481496648 |
| Tceal1 | ENSMUSG00000049536 | 237052 | -0.00668834430599285 | 1.94088417230086 | -0.0518111570446128 | 0.9594433156294 | 0.971198168620963 |
| Rbm12b2 | ENSMUSG00000052137 | 77604 | -0.00742475452890279 | 2.03234313581185 | -0.0516312519443751 | 0.9595840069199 | 0.971198168620963 |
| Arf1 | ENSMUSG00000048076 | 11840 | -0.00318655259933465 | 6.81082873794162 | -0.0515975976170435 | 0.9596103257892 | 0.971198168620963 |
| Fan1 | ENSMUSG00000033458 | 330554 | 0.00665246715153822 | 2.26083345230118 | 0.0515495247794809 | 0.9596479205264 | 0.971198168620963 |
| Fdps | ENSMUSG00000059743 | 110196 | 0.00627580724250487 | 3.00997532950891 | 0.051194587103629 | 0.9599254980103 | 0.971389151893266 |
| Atp6v1h | ENSMUSG00000033793 | 108664 | 0.00634030715808054 | 3.84647150576101 | 0.0504580533809896 | 0.9605015182897 | 0.971882078721205 |
| Ercc4 | ENSMUSG00000022545 | 50505 | 0.00686278751663361 | 2.72419383862073 | 0.0499418628483641 | 0.9609052286929 | 0.97211998075623 |
| Farsa | ENSMUSG00000003808 | 66590 | -0.00569541092818593 | 3.25168303051029 | -0.0499300094453601 | 0.9609144993203 | 0.97211998075623 |
| Hist3h2a | ENSMUSG00000078851 | 319162 | -0.0070506915546753 | 2.04720886229999 | -0.0495076473787684 | 0.9612448354146 | 0.972364176931092 |
| Fbxw8 | ENSMUSG00000032867 | 231672 | 0.00287656876910702 | 4.85712109360251 | 0.0493217843642038 | 0.9613902041915 | 0.972421238273188 |
| Jarid2 | ENSMUSG00000038518 | 16468 | -0.00389642329229822 | 4.47467896778702 | -0.0481289680031151 | 0.9623231738149 | 0.973274853161705 |
| 6820431F20Rik | ENSMUSG00000071796 | 547150 | 0.00411251064747109 | 5.21683509005982 | 0.0480085722790705 | 0.9624173453900 | 0.973280044634938 |
| Myl2 | ENSMUSG00000013936 | 17906 | 0.00581314636220576 | 10.4752460018144 | 0.0478474392101796 | 0.9625433819679 | 0.973286935332617 |
| Pknox2 | ENSMUSG00000035934 | 208076 | -0.00370815256772783 | 4.03349829067449 | -0.0477721950897597 | 0.9626022374788 | 0.973286935332617 |
| Dnajb1 | ENSMUSG00000005483 | 81489 | -0.0084245677595236 | 2.53514669514743 | -0.0473377687241223 | 0.9629420476747 | 0.973494262966595 |
| Pcna | ENSMUSG00000027342 | 18538 | 0.00463428988434567 | 3.28503579495729 | 0.0472823393537195 | 0.9629854053112 | 0.973494262966595 |
| Brap | ENSMUSG00000029458 | 72399 | 0.00277439904657228 | 4.88278445350083 | 0.0466701430315033 | 0.9634642820118 | 0.97382199578455 |
| Ctsf | ENSMUSG00000083282 | 56464 | 0.00482792029048573 | 3.27364749600412 | 0.0466401082062821 | 0.9634877764531 | 0.97382199578455 |
| Hspa4 | ENSMUSG00000020361 | 15525 | 0.00274820130025155 | 7.08416962375149 | 0.0457828212650857 | 0.9641583954007 | 0.974352617396108 |
| Safb | ENSMUSG00000071054 | 224903 | 0.00304744844361426 | 5.29507161740614 | 0.0457410904275123 | 0.9641910403781 | 0.974352617396108 |
| Mettl7a1 | ENSMUSG00000054619 | 70152 | -0.00531122653400917 | 3.815642939493 | -0.0455288614449905 | 0.9643570627714 | 0.974430306263163 |
| Ano10 | ENSMUSG00000037949 | 102566 | 0.0030651923695908 | 4.07792483365686 | 0.0453643321280802 | 0.9644857718782 | 0.974470281209053 |
| Slc40a1 | ENSMUSG00000025993 | 53945 | 0.00713575062286353 | 1.9240295612571 | 0.0441771924435101 | 0.9654144859017 | 0.975318460757151 |
| Epn2 | ENSMUSG00000001036 | 13855 | -0.00345121844947016 | 3.82792412491195 | -0.0438645703336759 | 0.9656590627984 | 0.97533179729132 |
| Mphosph6 | ENSMUSG00000031843 | 68533 | 0.00594246775419593 | 1.38764669517679 | 0.0438520679848425 | 0.9656688439657 | 0.97533179729132 |
| Itsn2 | ENSMUSG00000020640 | 20403 | -0.0030681281341165 | 5.23532785484524 | -0.0438181678211386 | 0.9656953656649 | 0.97533179729132 |
| Inf2 | ENSMUSG00000037679 | 70435 | 0.00725923804299988 | 3.69561558224695 | 0.0434828748811003 | 0.9659576834428 | 0.97550659993664 |
| Rasl12 | ENSMUSG00000041696 | 70784 | 0.00566626830192085 | 1.80404588453654 | 0.0432037090256202 | 0.9661760931071 | 0.97563304467271 |
| Glmp | ENSMUSG00000001418 | 56700 | 0.00532905426826691 | 2.48586315378084 | 0.0430890008861344 | 0.9662658375908 | 0.97563304467271 |
| Ppox | ENSMUSG00000062729 | 19044 | -0.00646721084509982 | 2.04234541807257 | -0.0429805950224242 | 0.9663506517858 | 0.97563304467271 |
| Cebpb | ENSMUSG00000056501 | 12608 | 0.00582005899112747 | 6.10099709365215 | 0.0426872060800235 | 0.9665801945122 | 0.975774668102487 |
| Zfp827 | ENSMUSG00000071064 | 622675 | 0.00319077492469244 | 4.17736050969609 | 0.0419192831176744 | 0.9671810194901 | 0.976291044791498 |
| Chst15 | ENSMUSG00000030930 | 77590 | 0.0101319521687588 | 2.35152867219306 | 0.041667108385283 | 0.9673783262259 | 0.976400044688423 |
| Inip | ENSMUSG00000038544 | 66209 | -0.00655169494637486 | 1.68173272023801 | -0.0414579060389412 | 0.9675420121675 | 0.976471866033861 |
| Ebf3 | ENSMUSG00000010476 | 13593 | 0.00672739216226015 | 3.02093829921905 | 0.041347822912633 | 0.9676281449893 | 0.976471866033861 |
| Bcl9l | ENSMUSG00000063382 | 80288 | 0.00410410817145834 | 5.70347336599467 | 0.0407418428695123 | 0.9681022921157 | 0.976860163861997 |
| Spata6 | ENSMUSG00000034401 | 67946 | 0.00767309662555538 | 1.74903807978584 | 0.0395649629825657 | 0.9690231737470 | 0.977699124259642 |
| Atr | ENSMUSG00000032409 | 245000 | 0.00469053592204594 | 3.37620484609602 | 0.0389101798278493 | 0.9695355466459 | 0.978125801604679 |
| Bc1 | ENSMUSG00000115783 | 100568459 | -0.0131063530583373 | 0.973397650878177 | -0.0384782104926164 | 0.9698735736960 | 0.978376525846367 |
| Sec13 | ENSMUSG00000030298 | 110379 | -0.00274530014227459 | 4.47628779006691 | -0.0364676364767581 | 0.9714469776301 | 0.979873296343577 |
| Ankra2 | ENSMUSG00000021661 | 68558 | -0.00369641426541678 | 2.67149144585241 | -0.0360592129578455 | 0.9717666107640 | 0.980105261326982 |
| Bclaf3 | ENSMUSG00000044150 | 382252 | 0.00577171416976752 | 1.73480121236467 | 0.0355064384963285 | 0.9721992213043 | 0.980451119852136 |
| Necap1 | ENSMUSG00000030327 | 67602 | -0.00468449945831351 | 2.90831305953706 | -0.035277115638658 | 0.9723786958944 | 0.98047518637572 |
| Tbl3 | ENSMUSG00000040688 | 213773 | -0.00370375402166623 | 2.97473955340702 | -0.0352065073087648 | 0.9724339562960 | 0.98047518637572 |
| Lpcat1 | ENSMUSG00000021608 | 210992 | -0.0049041628533617 | 1.91038921968222 | -0.035081567899721 | 0.9725317383489 | 0.98047518637572 |
| Cnn2 | ENSMUSG00000004665 | 12798 | -0.00663056723989186 | 3.03922495857028 | -0.0350175103250079 | 0.9725818722781 | 0.98047518637572 |
| Nucks1 | ENSMUSG00000026434 | 98415 | 0.00234224465833144 | 5.94660078263189 | 0.0348415057362538 | 0.9727196208805 | 0.980523623740788 |
| Plekhg3 | ENSMUSG00000052609 | 263406 | -0.0041343798075828 | 3.7509870136754 | -0.0345117197732392 | 0.9729777277020 | 0.980693364823498 |
| Psmg1 | ENSMUSG00000022913 | 56088 | 0.00512279787788872 | 1.57859803600607 | 0.0337169741818632 | 0.9735997478586 | 0.981229839926548 |
| Hmgcr | ENSMUSG00000021670 | 15357 | 0.004860625636113 | 2.80510021377099 | 0.0327425816516097 | 0.9743623959190 | 0.981907933049757 |
| Acsl4 | ENSMUSG00000031278 | 50790 | 0.00366831322133443 | 4.35089425969461 | 0.0322432855166606 | 0.9747532005858 | 0.982211212721646 |
| Pdgfb | ENSMUSG00000000489 | 18591 | 0.00443676048362285 | 3.96721379513389 | 0.031186185968032 | 0.9755806264005 | 0.982905607127512 |
| Lrrc40 | ENSMUSG00000063052 | 67144 | 0.00409696427920969 | 1.88530330930121 | 0.0311331238814695 | 0.9756221605830 | 0.982905607127512 |
| Plac9b | ENSMUSG00000072674 | 100039246 | -0.00629174890838469 | 2.11914569031198 | -0.0306351513706565 | 0.9760119505864 | 0.98320768886375 |
| Mthfr | ENSMUSG00000029009 | 17769 | -0.00527589231797387 | 2.8417740832403 | -0.0301764433320885 | 0.9763710118210 | 0.983478762460016 |
| Atp6v1f | ENSMUSG00000004285 | 66144 | 0.00254665144993327 | 3.59887957699073 | 0.0299808413663765 | 0.9765241240942 | 0.983542356995691 |
| Gm20939 | ENSMUSG00000095193 | 100044193 | -0.006803211653438 | 1.68230800067368 | -0.0298439974115288 | 0.9766312426561 | 0.983559619815208 |
| Abhd13 | ENSMUSG00000040396 | 68904 | -0.00283873399981838 | 2.80506896784329 | -0.0291373852708868 | 0.9771843711706 | 0.984026012092751 |
| Pxk | ENSMUSG00000033885 | 218699 | 0.00276488062572877 | 3.1176457197414 | 0.0284130903071045 | 0.9777513543100 | 0.984506268788101 |
| Ccdc86 | ENSMUSG00000024732 | 108673 | -0.00268725623880162 | 3.38256264981288 | -0.0281916888859546 | 0.9779246714005 | 0.984590087784785 |
| Smap1 | ENSMUSG00000026155 | 98366 | -0.00235795095978508 | 3.84390157634907 | -0.0275893145959584 | 0.9783962268926 | 0.984899672287763 |
| Lars2 | ENSMUSG00000035202 | 102436 | 0.00430347428235045 | 11.4180587412482 | 0.0275687020398214 | 0.9784123631298 | 0.984899672287763 |
| Pea15a | ENSMUSG00000013698 | 18611 | 0.00397824572987648 | 4.79989574521606 | 0.0274065037785763 | 0.9785393380028 | 0.98493678671351 |
| Capzb | ENSMUSG00000028745 | 12345 | 0.00176434223673905 | 6.56429151790393 | 0.0272342909545706 | 0.9786741533142 | 0.984981785275577 |
| Oasl2 | ENSMUSG00000029561 | 23962 | -0.00868280125359977 | 2.6766428928841 | -0.0271011415757637 | 0.9787783886133 | 0.984996001282661 |
| Nol4l | ENSMUSG00000061411 | 329540 | 0.00704566140671803 | 1.20302987482011 | 0.0268254056011428 | 0.9789942483706 | 0.985122537875321 |
| Hipk1 | ENSMUSG00000008730 | 15257 | -0.00188969229481373 | 6.47688174059316 | -0.0264503276447438 | 0.9792878806950 | 0.985327303376067 |
| Eif3g | ENSMUSG00000070319 | 53356 | -0.00206565547633009 | 5.27653681781897 | -0.0263023469191618 | 0.9794037292742 | 0.985353167482431 |
| Dhx38 | ENSMUSG00000037993 | 64340 | 0.00187010485927708 | 4.25566510624497 | 0.0260081104919167 | 0.9796340774126 | 0.985494211319509 |
| Bms1 | ENSMUSG00000030138 | 213895 | -0.00169159783935147 | 5.22243755638291 | -0.0258822465692061 | 0.9797326127677 | 0.985502640118228 |
| Ptges3l | ENSMUSG00000097487 | 73635 | -0.0029196893616259 | 3.1340175064841 | -0.0253978290055131 | 0.9801118529730 | 0.985793399415576 |
| Dpp7 | ENSMUSG00000026958 | 83768 | -0.00699015833687699 | 1.16957604559373 | -0.0247101098718897 | 0.9806502622409 | 0.98624418222065 |
| Nudt4 | ENSMUSG00000020029 | 71207 | 0.00192980752645511 | 8.10217395522229 | 0.0243287374032301 | 0.9809488396515 | 0.986440155033287 |
| Nit2 | ENSMUSG00000022751 | 52633 | 0.00371228912099575 | 2.00684031307749 | 0.0242306814217064 | 0.9810256083951 | 0.986440155033287 |
| Dtd2 | ENSMUSG00000020956 | 328092 | 0.00335738459722745 | 1.63951605673656 | 0.0239188501631643 | 0.9812697446620 | 0.986594884004865 |
| Srsf11 | ENSMUSG00000055436 | 69207 | 0.00197192858408357 | 4.83096208496401 | 0.0237321465835659 | 0.9814159179728 | 0.986651098993991 |
| Taf10 | ENSMUSG00000043866 | 24075 | -0.00154609107925108 | 4.26009819177502 | -0.022938546040211 | 0.9820372486233 | 0.987184951692263 |
| Gspt1 | ENSMUSG00000062203 | 14852 | 0.00141111392659621 | 5.90758816340903 | 0.0225921519077529 | 0.9823084534970 | 0.987366777487496 |
| Dnah7b | ENSMUSG00000041144 | 227058 | 0.00793310691423627 | 0.315131876102291 | 0.0221786714022103 | 0.9826321857750 | 0.987546873182799 |
| Colgalt1 | ENSMUSG00000034807 | 234407 | 0.00147774481147511 | 4.81668367854361 | 0.0221325276963451 | 0.9826683139337 | 0.987546873182799 |
| Saysd1 | ENSMUSG00000045107 | 67509 | 0.00366106571996072 | 1.40683646970748 | 0.0218628176637192 | 0.9828794838674 | 0.987668288118702 |
| Mypn | ENSMUSG00000020067 | 68802 | -0.00151807438914098 | 8.22690119470258 | -0.021600295491507 | 0.9830850273387 | 0.987784027377474 |
| Itpa | ENSMUSG00000074797 | 16434 | 0.00206149056429794 | 2.93230276653018 | 0.0209275138721161 | 0.9836117918395 | 0.988086813290409 |
| Morc2a | ENSMUSG00000034543 | 74522 | -0.00176262982072206 | 4.04395547797707 | -0.0209041449467407 | 0.9836300890362 | 0.988086813290409 |
| Slc35a2 | ENSMUSG00000031156 | 22232 | 0.00333543966019745 | 1.38454566563775 | 0.0208217057764782 | 0.9836946366132 | 0.988086813290409 |
| Scaper | ENSMUSG00000034007 | 244891 | -0.00162491324397029 | 3.91489199634726 | -0.0207385406998746 | 0.9837597526753 | 0.988086813290409 |
| Zfp994 | ENSMUSG00000096433 | 240038 | -0.00290694817148474 | 2.09880691565621 | -0.0206381727548239 | 0.9838383382904 | 0.988086813290409 |
| Jsrp1 | ENSMUSG00000020216 | 71912 | 0.00169418887020178 | 7.29075013639724 | 0.0205030073405454 | 0.9839441697383 | 0.988102316895749 |
| Mier3 | ENSMUSG00000032727 | 218613 | -0.00215629591807964 | 3.36720625301365 | -0.0199487273330471 | 0.9843781616626 | 0.988427972422291 |
| Riok2 | ENSMUSG00000116564 | 67045 | -0.00153415498608034 | 4.02841737778076 | -0.0198578680780035 | 0.9844493034270 | 0.988427972422291 |
| Hspb3 | ENSMUSG00000051456 | 56534 | -0.00204997968130982 | 3.029554808194 | -0.0189225951795611 | 0.9851816193843 | 0.989038939719473 |
| Cntrl | ENSMUSG00000057110 | 26920 | -0.00222178702295484 | 3.99446172791438 | -0.0188496040700144 | 0.9852387718054 | 0.989038939719473 |
| Rps9 | ENSMUSG00000006333 | 76846 | -0.00119689124407962 | 6.77481451153134 | -0.0157207389762325 | 0.9876887647146 | 0.991407335821436 |
| Syap1 | ENSMUSG00000031357 | 67043 | -0.000974331858879296 | 4.49421163308623 | -0.0153635835491757 | 0.9879684365819 | 0.99159700489187 |
| Xrn2 | ENSMUSG00000027433 | 24128 | -0.00119369865350105 | 4.40549657739429 | -0.0148441438270654 | 0.9883751886947 | 0.991914174405292 |
| Mark2 | ENSMUSG00000024969 | 13728 | -0.000881894320832188 | 4.87407396969571 | -0.0142881059222209 | 0.9888106030871 | 0.992260047952452 |
| Pla2g4e | ENSMUSG00000050211 | 329502 | -0.00305088680784582 | 3.48575708262024 | -0.014017559199918 | 0.9890224603871 | 0.992381541627696 |
| Zfp668 | ENSMUSG00000049728 | 244219 | -0.00148884317657627 | 2.12687782124754 | -0.013135379898398 | 0.9897132753500 | 0.992983553267653 |
| Stx18 | ENSMUSG00000029125 | 71116 | -0.00139419855166567 | 2.90653120733903 | -0.0126460088712074 | 0.9900964946484 | 0.99327431657392 |
| Kdm2a | ENSMUSG00000054611 | 225876 | -0.00076511151047054 | 5.45616500432402 | -0.0124632230014966 | 0.9902396322502 | 0.99327431657392 |
| Rps27l | ENSMUSG00000036781 | 67941 | 0.00154073417515654 | 3.61869055917418 | 0.012417185175906 | 0.9902756840199 | 0.99327431657392 |
| Clp1 | ENSMUSG00000027079 | 98985 | 0.00302997827295837 | 1.42325862719326 | 0.0119657896160414 | 0.9906291686164 | 0.99353770457344 |
| Atg101 | ENSMUSG00000037204 | 68118 | -0.00149601950795564 | 2.60146750986974 | -0.0113425193586202 | 0.9911172504007 | 0.993936024232208 |
| Tmc7 | ENSMUSG00000042246 | 209760 | -0.00102230317190477 | 3.82013317314988 | -0.0110570339200156 | 0.9913408147695 | 0.994069025433047 |
| Tmem184b | ENSMUSG00000009035 | 223693 | 0.00129950282592706 | 4.53253128816414 | 0.010341665761962 | 0.9919010246704 | 0.994505497977483 |
| Dimt1 | ENSMUSG00000021692 | 66254 | -0.00160901984520323 | 1.6326422765452 | -0.0102688467777511 | 0.9919580499907 | 0.994505497977483 |

| | | | | | | | |
|---|---|---|---|---|---|---|---|
| R3hdm1 | ENSMUSG00000056211 | 226412 | 0.000645738741283708 | 6.02264994536932 | 0.00979536575401204 | 0.9923288391141 | 0.994785999665894 |
| Abhd8 | ENSMUSG00000007950 | 64296 | 0.00128350655333742 | 2.12170594960632 | 0.0094242745382809 | 0.9926194467487 | 0.994986077249977 |
| Ntan1 | ENSMUSG00000022681 | 18203 | -0.00100622682078511 | 3.06404509103544 | -0.0088108211370707 | 0.9930998545716 | 0.995376353413029 |
| Ppp1r16a | ENSMUSG00000033819 | 73062 | 0.000691765986574833 | 3.29987848907073 | 0.00842257125881911 | 0.9934039023939 | 0.995589809944866 |
| Hyal2 | ENSMUSG00000010047 | 15587 | 0.00106675143479107 | 2.32724014390398 | 0.00823716539423077 | 0.9935490985680 | 0.995644040745073 |
| Hmg20b | ENSMUSG00000020232 | 15353 | 0.00076403579183637 | 3.32772270696158 | 0.00760276239325101 | 0.9940459178941 | 0.996050593867495 |
| Uhrf1bp1 | ENSMUSG00000039512 | 224648 | 0.000688656991187031 | 4.8706441809576 | 0.00737258705899363 | 0.9942261754989 | 0.996139901409654 |
| Snx3 | ENSMUSG00000019804 | 54198 | -0.00044542650492474 | 5.55345665214075 | -0.00706438828890793 | 0.9944675362440 | 0.996290407724686 |
| Map7d1 | ENSMUSG00000028849 | 245877 | 0.000367918643751963 | 7.94495879156927 | 0.00608072653623286 | 0.9952378782560 | 0.996970788784521 |
| Pole3 | ENSMUSG00000028394 | 59001 | 0.000597552092836295 | 3.50340007239642 | 0.00564784152644245 | 0.9955768881833 | 0.997128888886438 |
| Cwf19l2 | ENSMUSG00000025898 | 244672 | 0.000438706990131266 | 5.21013954350599 | 0.00564623781924383 | 0.9955781441136 | 0.997128888886438 |
| Magoh | ENSMUSG00000028609 | 17149 | -0.000487804171095174 | 3.54610211241445 | -0.00532503201922977 | 0.9958296940734 | 0.997248242836573 |
| Ensa | ENSMUSG00000038619 | 56205 | 0.000366157199565897 | 6.47654050653647 | 0.00526108362693089 | 0.9958797748425 | 0.997248242836573 |
| Alg9 | ENSMUSG00000032059 | 102580 | -0.000655165142034342 | 2.06882356646609 | -0.00455396179144114 | 0.9964335538849 | 0.997680973599571 |
| Sco1 | ENSMUSG00000069844 | 52892 | 0.000447477090042447 | 2.84569707836458 | 0.00437879481252417 | 0.9965707353641 | 0.997680973599571 |
| Hint1 | ENSMUSG00000020267 | 15254 | 0.000359172299013721 | 5.07894713399833 | 0.0043596556405575 | 0.9965857241545 | 0.997680973599571 |
| Jup | ENSMUSG00000001552 | 16480 | 0.000213365686536626 | 6.07638094669403 | 0.00330119833141122 | 0.9974146540410 | 0.998419375762177 |
| Rrp15 | ENSMUSG00000001305 | 67223 | -0.000334036950222494 | 2.39182679093783 | -0.00289973357981173 | 0.9977290616671 | 0.998642649307194 |
| Polr2g | ENSMUSG00000071662 | 67710 | 0.00017130138937782 | 2.65221227957247 | 0.00174840261498872 | 0.9986307299564 | 0.999453626547675 |
| Ipp | ENSMUSG00000028696 | 16351 | -0.000145025144619559 | 3.10264345483717 | -0.00147668672755785 | 0.9988435253986 | 0.999575077921165 |
| Acta1 | ENSMUSG00000031972 | 11459 | -0.000111125414310465 | 13.9467435939853 | -0.00106155810009215 | 0.9991686353040 | 0.999707269727581 |
| Rps14 | ENSMUSG00000024608 | 20044 | -0.000100413515391473 | 6.24897851005559 | -0.00103378988733794 | 0.9991903821145 | 0.999707269727581 |
| Zfp266 | ENSMUSG00000060510 | 77519 | 8.40534640369484e-05 | 4.08386415345728 | 0.000957678736726968 | 0.9992499889345 | 0.999707269727581 |
| Magi1 | ENSMUSG00000045095 | 14924 | -7.72877518571136e-05 | 2.6556439069963 | -0.000603928846629441 | 0.9995270299473 | 0.999892922518027 |
| Lmbrd2 | ENSMUSG00000039704 | 320506 | 2.72707021906626e-05 | 4.26177259611839 | 0.00041507333551198 | 0.9996749331229 | 0.99989315699222 |
| Nfkbia | ENSMUSG00000021025 | 18035 | 4.96781381191802e-05 | 3.92701308185021 | 0.000370027935904386 | 0.9997102106639 | 0.99989315699222 |
| Acsl6 | ENSMUSG00000020333 | 216739 | 1.96536496974709e-05 | 3.36710685605145 | 0.000123746144041372 | 0.9999030875503 | 0.999994569992055 |
| Aff1 | ENSMUSG00000029313 | 17355 | 1.96408163510853e-07 | 5.65952770317174 | 2.85002954060124e-06 | 0.9999977679842 | 0.99999776798423 |
